# No evidence that shrinking and shapeshifting meaningfully affect how birds respond to warming and cooling

**DOI:** 10.1101/2024.03.22.586255

**Authors:** Joshua K.R. Tabh, Elin Persson, Maria Correia, Ciarán Ó Cuív, Elisa Thoral, Andreas Nord

## Abstract

Across the globe, birds and mammals are becoming smaller and longer-limbed. Although the cause of these changes is unclear, many argue that each provide thermoregulatory benefits in a warmer world by easing heat dissipation. Here, we show that neither body size nor limb length in a model species (the Japanese quail) influenced metabolic costs of warming during a cold challenge. In the heat, larger body sizes increased metabolic costs of thermoregulation, however, this effect was moderate and almost always negated by cooling from the limbs (>97% of cases). Rearing in the warmth (30°C) relative to the cold (10°C) reduced body sizes and increased limb lengths at adulthood but thermoregulatory benefits of these changes in later heat exposures were absent. Our findings demonstrate that shrinking and shape-shifting are unlikely to ease thermoregulation in contemporary birds or reflect selection for such. Alternative contributors, including neutral or non-adaptive plasticity, should be further investigated.

**Teaser:** Using experimental data, we show for the first time that shrinking and shape-shifting – which has been described as the third general response to climate change in animals – does not inherently provide thermoregulatory benefits to birds in a warming world. Further research evaluating the drivers of shape-shifts (including neutral plasticity and temporal reductions in resource abundance) is needed before we can determine why animals shrink under climate change.

## Main

In 2022, “Dippy”, the cast replica of a *Diplodocus carnegii* skeleton unearthed in the late 1800s, returned for display at the United Kingdom’s Natural History Museum after several years of absence. Within 6 months of return, over 1 million attendants sought to view the cast, rendering Dippy the museums “most popular exhibition” of the year (1). That Dippy stands an impressive 6 metres tall and 26 metres long is undoubtedly one of the keys to its allure; we are fascinated with large-bodied animals (2,3). Yet opposing this fascination, amassing evidence now indicates that terrestrial mammals and birds are shrinking and changing shape (4–7). How or why these “shape-shifts” have occurred is not yet known, however, given a concurrence with rising global temperatures, many have interpreted them as beneficial for thermoregulation (6,8,9) and potentially driven by selection. By decreasing body size and adjusting shape, surface area to volume ratios may increase, thus easing heat dissipation in a warmer world.

The logic behind thermoregulation as a driver for species’ shape-shifts is not novel. Over a century ago, Bergmann (10) and Allen (11) applied the same logic to explain their observations that body size generally decreases, and appendage length increases, with increasing environmental temperatures across the globe (Bergmann’s and Allen’s rules respectively). Despite this, whether body size and shape do have meaningful impacts on costs of thermoregulation in endotherms with evolved heat retention and dissipation mechanisms is tenuous. Already 70 years ago, for example, Scholander (12) argued that low surface area to volume ratios should be irrelevant for endotherm survival in the cold since heat loss is so readily mitigated via insulation, heat exchangers, and peripheral vascular constriction (supported by ref. 13). In the heat, predictions from long-established physiological allometries have raised similar doubts about the importance of morphology on thermoregulatory costs (14). Yet even with such longstanding theoretical discussion, remarkably little is empirically known about whether body size and shape do directly alter costs of thermoregulatory within a species. One probable hinderance is that adult morphometry and thermal physiology are often part of an interlaced phenotype that emerges from early temperature experience (15–17). Thus, teasing out any direct effects of morphology on thermoregulation is empirically challenging, and probably explains mixed conclusions of the few striving to do so (18–22). Still, if we wish to understand whether shape-shifting in contemporary bird species is driven by selection on improved thermoregulatory efficiency in the heat, direct effects of morphology on thermoregulation must first be clarified.

In this study, we used the Japanese quail (*Coturnix japonica*) to test whether body size (here, body mass, a strong predictor of skeletal size in this species; see Methods) and limb length (here, tarsus length) have a direct influence on costs of thermoregulation in the heat and cold. The Japanese quail was chosen as our model for its precociality, allowing us to control for early thermal experience by rearing individuals at fixed ambient temperatures without bias from parental brooding. To measure thermoregulatory costs, we quantified the rate at which individuals increased their resting metabolism across moderate cooling (from 30°C to 10°C) and heating (from 30°C to 40°C) events (here, defined as “metabolic slopes”). To better understand mechanisms linking morphology to thermoregulation in the heat, these measurements were supplemented with those of evaporative cooling efficiency (i.e. the ratio of evaporative heat loss to metabolic heat production) at temperatures limiting sensible heat loss (40°C). Effects of body mass and tarsus length on all measures of thermoregulatory costs were evaluated with simple Bayesian regressions. Last, we varied rearing conditions of quail (10°C [cold] and 30°C [warm]) then compared metabolic slopes and growth curves across rearing treatments to: (1) compare the importance of morphological contributions to thermoregulation against those of physiological acclimation, and (2) evaluate neutral phenotypic plasticity as an alternative driver of modern avian shape-shifts. All analyses were carried out on juveniles (3 week old) and adults (8 week old, the age of reproductive maturity; 23). Together, our findings cast doubt on whether body size and limb length alone (24–25) provides any meaningful influence on thermoregulation, including at temperatures matching heat-waves in a warming world. We therefore question whether selection on improved heat dissipation is driving contemporary shape-shifts in birds.

## Results and Discussion

Bayes Factors (BF) are given for test statistics (e.g. model coefficients, or βs) in place of credible intervals and represent relative support for the alternative hypothesis over the null. A BF of ≥ 3 represents moderate support for an alternative hypothesis, with corresponding 50% credible intervals not crossing 0. Credible intervals (50% and 95%) are provided in the supplement information.

### Body size and limb length weakly influence thermoregulatory costs in the cold, but only in juveniles

Large bodies and short appendages are widely assumed to reduce costs of thermoregulation in the cold by decreasing surface area to volume ratios and lowering rates of sensible heat loss. Our findings did not support this assumption. At maturity, neither body mass nor tarsus length influenced metabolic slopes below thermoneutrality (Figs. 1A-1B; n=56; mass: β≈0, BF=2.272, partial R^2^=0.020 [95%: 0, 0.174]; tarsus length: β≈0, BF=1.780; partial R^2^=0.016 [95%: 0, 0.164]; Supp. 3 Tab. 17). Even when dramatically misaligned with assumed optima (i.e., a body mass 2 standard deviations [SDs] below the mean, or tarsus length 2 SDs above the mean), predicted metabolic responses to cooling were virtually identical to those of average-sized individuals (metabolic slope at small mass - metabolic slope at average mass ≈ 0, BF=1.18; metabolic slope at long tarsus length - metabolic slope at average tarsus length ≈ 0, BF = 1.12). Control over life-time temperature experience in our study indicates that previous failures to link morphology with metabolic responses to cold (i.e. as in wild birds; 18,22) are probably not due to masking effects of thermal acclimation.

**Figure 1.**
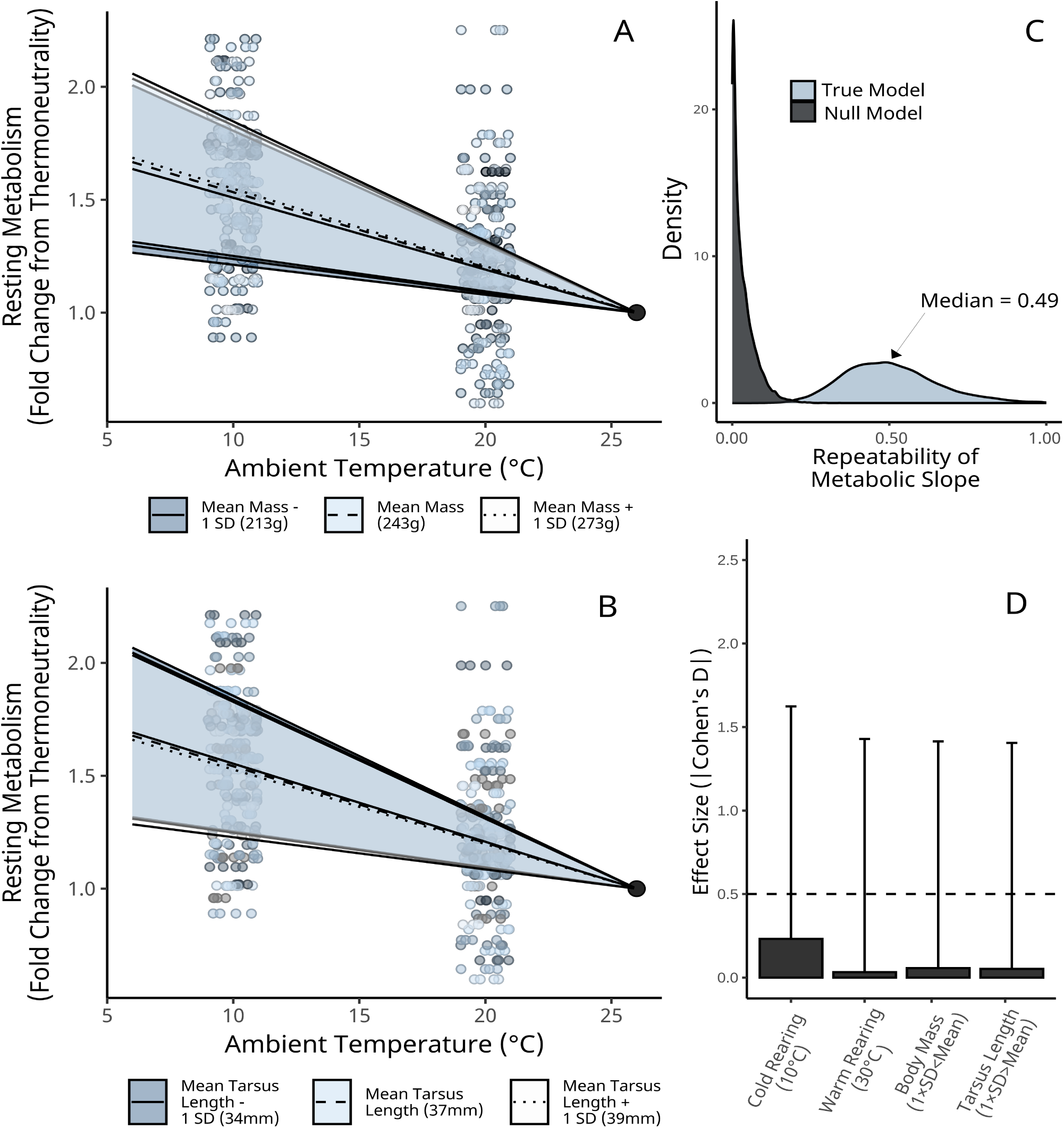
Contributions of morphology, individual identity, and prior temperature experience on metabolic responses to cold in adult Japanese quail (n = 53). Panels A and B show effects of body mass and tarsus length respectively on resting metabolic rate. Resting metabolism is relativised by individual to represent their fold change from thermoneutrality (here, 30°C). Lines and ribbons represent predicted effects ± one standard error respectively, holding other variables constant. Small dots represent raw values, with colour scaled by relative morphological size. Panel C displays conditional repeatabilities of metabolic slopes below thermoneutrality (calculated following Schielzeth and Nakagawa, 2022). “True model” indicates repeatability estimates derived from a model where individual identities were known and correctly labelled; “Null” model indicates repeatability estimates derived from a model where individual identities were randomly scrambled. Panel D shows absolute effect sizes (here, Cohen’s D) of given predictors on metabolic slopes of adult quail. Bars display means and errorbars indicate standard deviations.

During development (i.e. 3 weeks of age), effects of body mass on metabolic slopes were slightly clearer (β=2.0×10^-4^, BF=29.651; partial R^2^=0.092 [0, 0.292]) but still absent for tarsus length (β=- 2.0×10^-4^, BF=1.976; partial R^2^=0.020 [0, 0.244]; Supp. 3 Tab. 10). At this stage, individuals with smaller body masses (and thus, higher surface area to volume ratios) increased their resting metabolism slightly more across a cooling event relative to those with larger body masses (Supp. 3, Fig. 30), as predicted by compensation for increased rates of heat loss. In Japanese quail, plumage development remains incomplete until near sexual maturation (three weeks after our juvenile measurements; 26). Early juveniles may therefore be more dependant on body size scaling for heat loss and heat retention relative to better-insulated adults. Moreover, larger individuals may also be better insulated owing to advanced developmental stages, thus reducing demands for compensatory heat production. Still, effects of mass on metabolic slopes in juveniles were relatively small (mean Cohen’s D between mean mass and mean mass + 1 SD = 0.32; partial R^2^=0.078 [0, 0.284]), with atypically small body mass (2 SDs below the mean) predicted to increase metabolic slopes by ∼9% more than average sized birds at 10°C.

One possible explanation for why we did not detect effects of adult morphometry on metabolic responses to cold is that metabolic slopes were either highly variable within individuals (i.e., ‘noisy’) or highly similar among individuals. In these cases, thermoregulatory phenotypes would be too indistinguishable to detect any causative influence of morphometry. However, similar to other thermal physiological traits (e.g. cold-induced maximal metabolism and peripheral vasomotor actions; 27,28), metabolic slopes were indeed distinct among quail (Fig. 1C; conditional repeatability = 0.488 [95%: 0.250, 0.840]; probability of exceeding repeatability in null model ≈ 100%; Supp. 3 Fig. 22), yet neither body size nor appendage length were able to explain their distinctions. Alternatively, larger body sizes and shortened appendage lengths may have benefited thermoregulation by either: (1) reducing risk of hypothermia, or (2) reducing lower critical temperatures (proposed in ref. 29) rather than flattening metabolic responses below such. Again, however, our data provided little evidence to support this (similar to others; 22). Neither body mass nor tarsus length meaningfully influenced body temperature responses to cooling (body mass: β=-7.0×10^-4^, BF=1.474; tarsus length: β=-9.1×10^-3^, BF=1.559; n=55; Supp. 6 Tab. 3) or lower critical temperature in our adult quail (body mass: β=- 1.0×10^-3^, BF=4.615; tarsus length: β=0.014, BF=2.350; Supp. 5 Tab. 3) as estimated from remote, intraperitoneal temperature readings and break-point regressions (metabolism by ambient temperature) respectively.

Unlike morphology, we found that thermal history did, at least weakly, influence metabolic responses to the cold among both adults and juveniles. Here, cold-rearing (10°C), but not warm-rearing (30°C) had a direct effect on metabolic slopes below 30°C (adults; cold rearing: β=-4.4×10^-3^, BF = 2.916; warm rearing: β≈0, BF = 1.040; Supp. 3 Tab. 17; juveniles; cold rearing: 3.8×10^-3^, BF = 6.240; warm rearing: β=-2.5×10^-3^, BF = 2.509; Supp. 3 Tab. 10), with overall effects on such being even larger than those of relatively substantial changes in body mass and tarsus length (1 SD; Fig. 1D) at adulthood. While we cannot rule out possible effects of cold rearing on plumage density (i.e. by stunting or promoting feather growth), these outcomes strongly indicate that acclimation of thermogenic performance is arguably more important for shifting cold tolerance than acclimation of insulation via morphological plasticity (Fig. 1D; see ref. 30). Supporting this, cold tolerance and thermogenic capacity are widely known to vary across seasons in temperate birds (31-33; reviewed in ref. 34), with some even showing their necessity for survival through harsh winters (35). That selection pressures from low ambient temperatures necessarily drives changes in morphology therefore appears unlikely. This is particularly true given that physiological acclimation is predicted to buffer any heat-loss costs of high surface area to volume ratios, argued nearly a century ago by Scholander (12).

### Body size and tarsus length weakly influence thermoregulation to the heat, but in opposite directions

Ecogeographical rules describing temperature-size relationships across endotherms primarily focussed on an influence of low ambient temperatures on morphology, rather than high (10,11). However, by physical principles, warm environments should also favour smaller and longer-limbed animals owing to their larger surface area to volume ratios and thus higher sensible heat dissipation rates. We tested this assumption by evaluating whether and how body mass and tarsus length influenced the extent to which metabolism of quail (n=84) increased during a heat exposure (40°; near normal body temperature, 42°C, in quail; 36) relative to thermoneutrality (30°C).

As predicted by size-related heat dissipation capacities, the rate at which adult quail increased metabolism in the heat increased with body mass (Fig. 2A; β=3.0×10^-4^, BF=59.606; Supp. 3 Tab. 33), although with limited explanatory power (partial R^2^=0.043 [95%: 0, 0.158]). Trends in evaporative cooling efficiency suggest that elevated metabolic responses in large birds may represent an inability to counteract metabolic heat production with evaporative cooling (Fig. 2B; effect of mass: β=-1.4×10^-3^, BF=27.369; n=64; Supp. 4 Tab. 9), the dominant form of heat-dissipation at ambient temperatures near body temperature (as here; 37; but see no effect of mass on body temperature response to heating; β ≈0; BF=1.107; n=84; Supp. 6 Tab. 6). Among atypically heavy birds (i.e., mean mass + 2 SDs), heat exposure (40°C) also bore apparent metabolic costs, with relative metabolism increasing by ∼42% from thermoneutrality compared with a modest 10% among birds of average mass (holding tarsus length constant). Superficially, these findings imply that a large body size may well impede thermoregulatory performance in a warmer world, as assumed by many (*sensu* 4,8,9). However, metabolic responses to heat also marginally *decreased* with tarsus length (Fig 2C; β=-2.6×10^-3^, BF=11.719, partial R^2^ = 0.026 [0, 0.152]; Supp. 3 Tab. 33), a trait with clear and positive body size allometry (β=0.028, BF>1000). Combining these effects showed that thermoregulatory costs accrued by large individuals were at least partly offset by benefits of their elongated limbs (presumably by increasing maximal sensible heat loss, given that tarsus length did not influence evaporative cooling efficiency: β=3.7×10^-3^, BF=1.679; Supp. 4 Tab. 9). Phenotypes lending to moderate (mean Cohen’s D ≥ 0.5) or large (mean Cohen’s D ≥ 0.8) metabolic costs in the heat therefore required extreme deviations from allometry (i.e. a large mass and short tarsi), which was rare in our population (Fig. 2E ∼2.5% and 0% for moderate and large effects respectively). Likewise, allometric deviations leading to moderate (Cohen’s D ≤ -0.5) or large (Cohen’s D ≤ -0.8) thermoregulatory *benefits* (i.e. by having a relatively small body size and long tarsi) was equally rare (Fig. 2D; 1% and 0% for moderate and large effects respectively).

**Figure 2.**
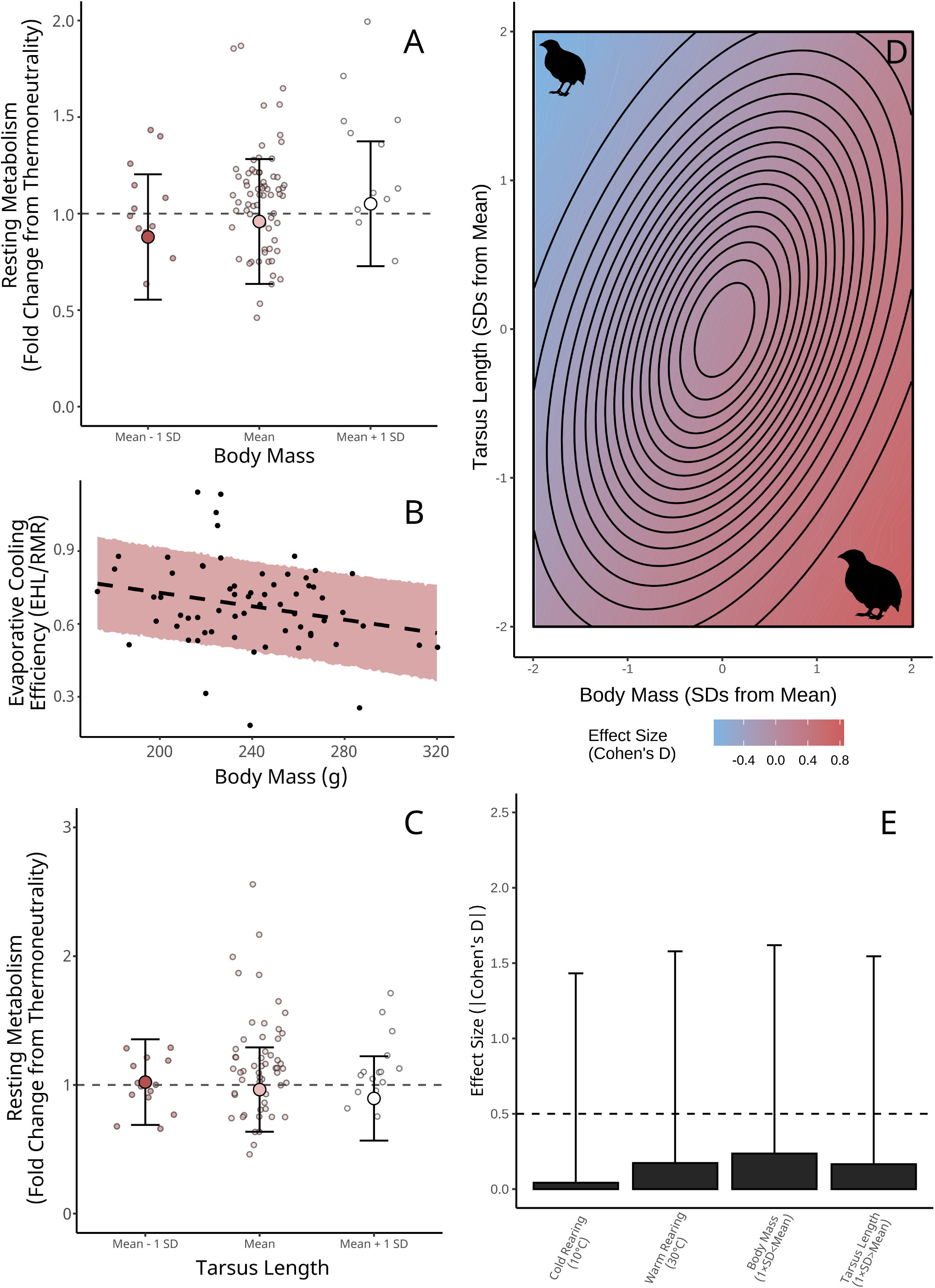
Contributions of morphology and prior temperature experience on metabolic responses to heat in adult Japanese quail (n = 84). Panels A and C display effects of body mass and tarsus length respectively on resting metabolic rate, with resting metabolism relativised to represent an individual’s fold change from thermoneutrality (here, 30°C). Large dots indicate predicted effects, holding other variables constant. Errorbars indicate ± 1 standard deviation around predictions. Small dots display raw values, with individuals coloured according to whether their given morphometric measured was below, above or within one standard deviation of the mean. Panel B displays the effect of body mass on evaporative cooling efficiency. Evaporative cooling efficiency is calculated as the ratio of evaporative heat loss (in watts) by metabolic heat production (also in watts; assuming 20 J/mL of O_2_), with evaporative heat loss estimated from measured evaporative water loss and assuming 2406 J required to evaporating 1 mL of water. The dashed line and ribbon indicate the predicted relationship ± one standard deviation, respectively. Dots represent raw values. Panel D shows the combined effects of body mass and tarsus length on metabolic slopes in the heat relative to the mean. Effects are shown as mean Cohen’s D values. Contours show the multivariate, normal distribution of phenotypes among quail. The centre circle indicates the modal phenotype for our population (i.e. the peak of the distribution). Outer circles represent the tails of the distribution, where phenotypes within are rare. Panel E displays absolute effect sizes (here, Cohen’s D) of given predictors on metabolic slopes. Bars display means and errorbars indicate standard deviations.

When we analysed heat-induced metabolic responses in juveniles, moderate effects of body mass and tarsus length again emerged. Consistent with adults and predictions from physical principles, larger juveniles with shorter tarsi tended to increase their metabolism slightly more in the heat than smaller juveniles with longer tarsi (mass: β=3.0×10^-4^, BF=6.055; partial R^2^=6.0×10^-3^ [0, 0.147]; tarsus length: β=-1.5×10^-3^, BF=5.768; partial R^2^=0.007 [0, 0.146]; Supp. 3 Tab. 26). Again, elevated metabolic responses among large juveniles appeared to be a consequence of their reduced capacity to dissipate heat evaporatively (effect of body mass on evaporative cooling capacity: β=-2.60×10^-3^, BF=194.12; Supp. 4 Tab. 3). Nevertheless, given that large individuals generally had longer tarsi (β=0.065, BF>1000) that offset these costs (i.e. by increasing dry heat loss), phenotypes displaying moderate (Cohen’s D ≥ 0.5) or high (Cohen’s D ≥ 0.8) increases in metabolism relative to average remained rare at this life stage (1% and 0% for moderate and large effects respectively).

The above findings provide tentative evidence that shifts in body size and appendage length could benefit thermoregulation in a warming world (at least among endotherms). Exactly how and to what degree these benefits might shape selection in nature, however, is not obvious (see ref. 38). That effects of body mass and limb length on metabolic responses to heat evidently counteract each other demonstrates that assuming benefits of independent size reductions or limb elongations is too simplistic. Rather, for thermoregulatory benefits of each to emerge, dissolving or dramatically modifying allometries between these traits (i.e. by increasing tarsus lengths relative to body sizes, or decreasing body sizes while keeping tarsus lengths constant) is first required. Supporting this, allometry between body size and appendage length appears to explain thermal niche across species better then each trait individually (25). In our study, however, deviations from allometry that led to at least moderate thermoregulatory costs were uncommon (∼2.5%), and removal of these individuals (simulating selective disappearance) had little effect on the relationship between tarsus length and body mass across our population (change in slope = 6.61×10^-3^ ± 0.011; BF = 2.962; holding all other effects constant). As such, consequences of selection on these extremes on population-level phenotypes could be limited (but see ref. 39). Even then, evidence from desert birds has shown that, in extreme heat waves (∼48°C), size reductions can carry thermoregulatory *costs,* not benefits, by lowering maximal heat tolerance and time to dehydration (21), which may help explain continued positive selection on body size in our warming world (40). Evidently, if climate warming is creating new selective pressures on thermoregulation, whether and how that may change body size and shape in avian populations is likely too complex to explain unidirectional shape-shifts reported across species (see ref. 14).

Finally, similar to metabolic responses in the cold, our results again revealed a tentative, but weak effect of prior temperature exposures on metabolic responses to heat, particularly when those exposures were to warmth (adults; warm rearing; β=0.066, BF=3.065; cold rearing [10°C]: β=1.40×10^-3^, BF=1.289; Supp. 3 Tab. 33). These effects were more apparent in juveniles than adults (juveniles; warm rearing: β=-0.018, BF=29.303; cold rearing [10°C]: β=-6.3×10^-3^, BF=3.553; Supp. 3 Tab. 26) and could reflect changes in physiology facilitating evaporative and sensible cooling seen in several bird species (e.g. 41-43). Effects of prior experience of warmth on metabolic responses to heat were also comparable to those of relatively large changes in body mass and tarsus length (+1 SD; Fig. 2E). Thus, any thermoregulatory costs imposed by severely mismatching morphology with expected optima (discussed above) could, at least partly, be compensated by physiological acclimation.

### Plasticity recapitulates observations of shape-shifting, but with no thermoregulatory benefit

Beyond selection on thermoregulatory response to heat, warming climates may directly alter avian size and shape through neutral, or even non-adaptive phenotypic plasticity (16,44). To test this, we raised quail in cold (10°C), mild (20°C) and warm (30°C) temperatures, then compared body mass and tarsus length at maturity.

Supporting a plastic origin of size declines and limb elongations (6,7), quail raised at warm temperatures (n=47) has smaller asymptotic masses (Fig. 3A; Δ Gompertz a=-13.25, BF=113.286; Supp. 2 Tab. 5) but longer tarsus lengths (Fig. 3B; Δ Gompertz a=0.641, BF=7.252; Supp. 2 Tab. 14) than cold reared quail (10°C; n = 46), despite growing more quickly (mass Δ Gompertz c=0.061, BF>1000; tarsus length Δ Gompertz c=0.088, BF=53.983; Supp. 2 Tab. 5; Supp 2. Tab. 14). These findings largely agree with those of others (43,45), revealing temperature-dependant developmental plasticity as a possible contributor to negative temperature scaling of mass and positive temperature scaling of tarsus length.

**Figure 3.**
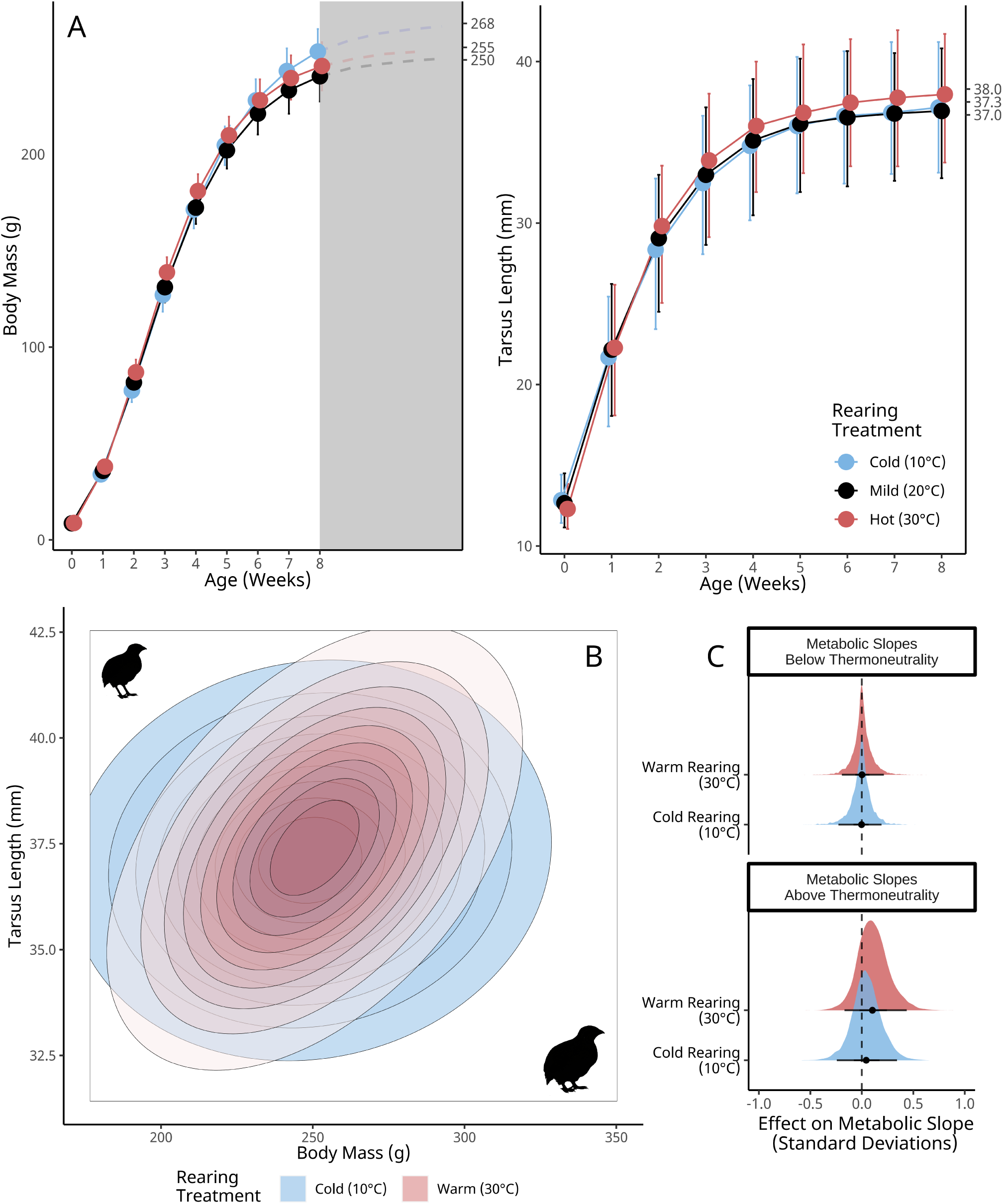
Effects of rearing temperature on growth, tarsus elongation, and thermoregulatory responses to heat and cold at adulthood. Rearing treatments were maintained until at least three weeks of age. Panel A displays mass gain and tarsus elongation across weeks. Large dots display predicted means from a Bayesian Gompertz model, by age, and errorbars represent 95% quantile intervals. Asymptotes per treatment are displayed on the right-hand y-axes. In panel A, the grey rectangle indicates the period of estimated further growth after sexual maturation. Panel B displays distributions of combined mass and tarsus lengths among adult quail reared in two temperature treatments. Contours show the distribution of phenotypes assuming multivariate normality. Panel C displays the direct effects of plastic differences in mass and tarsus length observed between temperature treatments on subsequent metabolic responses to cold (30°C - 10°C) and heat (30°C – 40°C) exposures. Densities represent posterior densities from Bayesian path analyses and the dashed line indicates zero.

Despite evidence that thermal conditions can alter phenotypes to reduce surface area to volume ratios in the heat, or increase them in the cold (Fig. 3C), we find no evidence that emergent phenotypes benefit thermoregulation in response to heat or cold exposures. In all cases, plastic changes in morphology led to remarkably weak and uncertain effects on metabolic responses to both heat and cold challenges (Fig. 3D). Thus, temperature-dependant plasticity of size and limb length does not evidently occur to reduce thermoregulatory costs (i.e. via adaptive plasticity). Instead, heat-induced reductions in mass and increases in limb length may better reflect the emergent consequences of balancing thermoregulation and growth, and direct effects of temperature on peripheral tissue proliferation. For example, in the warmth, energetic demands of dissipating heat (including that produced *from* growth) may compete with those *of* growth, thus shrinking tentative asymptotic masses (44). Further, stimulating effects of heat on chondrocyte proliferation in developing limbs may also directly increase asymptotic limb lengths (46,47), but without a subsequent thermoregulatory value.

## Conclusions

In birds and mammals, body size and limb length are widely assumed to influence costs of thermoregulation by affecting surface area to volume ratios, and thus, rates of sensible heat loss. Here, we show that this assumption is tenuous and context specific. In the cold, body size and limb length had remarkably limited effects on costs of thermoregulation, and only during early development. In the heat, each variable independently influenced cost of thermoregulation at all ages, however, phenotypes expected to suffer large costs required strong allometric decoupling (i.e. a large body and short limbs) that was rare in our population (∼2.5%). More critically still, prior exposure to cold and warmth (i.e., acclimation, or physiological plasticity) had subsequent effects on thermoregulatory costs that were comparable to, or even larger than, dramatic changes in body mass or limb length (± 1 SD). If changing climates alter selection pressures on thermoregulatory performance, our results indicate that adapting physiology is arguably more effective and probable than adapting morphology. Even if pressures to adapt morphology occur to match warming temperatures, effects on population-wide phenotypes may be minimal and unlikely to drive shape-shifts observed across avian species.

## Methods

All animal handling, measurements, and euthanasia for this study were approved by the Malmö/Lund Animal Ethics Committee (permit no. 9246-19).

### Animal husbandry

Japanese quail eggs (n_batch 1_ = 92, n_batch 2_ = 60 and n_batch 3_ = 85) were acquired from a commercial supplier (Sigvard Månsgård, Åstorp, Sweden) and held at room temperature, manually or mechanically turned, for a maximum of 6 days until incubation in Brinsea OvaEasy 190 incubators (Brinsea, Weston-super- Mare, United Kingdom; calibrated prior to use). Incubation was completed in three batches between 2021 and 2022, with temperatures fixed at 37.5°C and relative humidity (RH) fixed at 50% per batch (23). In all batches, eggs were shifted to hatching trays within incubators at day 15 of incubation and monitored twice daily until hatching. Once hatched (n = 47, 45, and 55 for batches one to three; average hatching success ≈ 63.6%), chicks were collected within 12 hours of their earliest possible hatch time, colour banded, measured (see below), and placed into one of three possible animal housing rooms (14L:10D), each containing identical open pens (310 × 120 × 60 cm) lined with wood shavings. Ground food (turkey starter pellets, [Kalkonfoder Start, Lantmännen, Stockholm, Sweden]), water and crushed seashells were provided *ad libitum* and supplemented with mealworms and mixed shredded vegetables (e.g. lettuce and carrots) haphazardly but equally between pens.

At least 24 hours before use, housing rooms were set to one of three ambient temperature treatments (10°C [cold], 20°C [mild], or 30°C [warm]) and monitored daily for temperature deviations. Relative humidity was left at ambient. To aid survivorship before development of endothermy, all pens were equipped with a hanging, infrared heat lamp until chicks reached two weeks (experimental batch three) or three weeks (experimental batches one and two; 43,48). In batch three, heating from lamps was further restricted by allowing for 6 cooling bouts per day (10 mins/2 hours in week one, and 30 mins/2 hours in week two). Food and water were placed to require regular departure from lamps, thus ensuring consistent exposure to treatment temperatures (see ref. 49). Surface temperatures under lamps averaged approximately 37.5°C across pens (determined by infrared thermography) and duration of lamp placement did not differ between treatment groups.

At three weeks of age, ground feed was altered to lower relative protein density (turkey grower [Kalkonfoder Tillväxt, Lantmännen, Stockholm, Sweden]; 22.5% protein relative to 25.5% in starter feed) and maintained *ad libitum* until study completion. At this time, warm- and cold-reared birds from experimental batches one and two were also shifted to mild conditions (20°C) until adulthood, as part of another study. All birds reared in warm- or cold- conditions until at least this time are nonetheless considered “warm-” and “cold-” reared respectively, recognising that study outcomes are conservative.

Quail were euthanised upon study completion.

### Morphometry

Quail were weighed to the nearest 0.1 g on a digital scale within 12 hours of hatching, then weekly until 8 weeks of age (the age of reproductive maturity for this species; 23). Tarsus lengths were measured weekly until 3 weeks of age, then again at 8 weeks of age. To measure tarsus length, individuals were flat-lay photographed on 1 mm × 1 mm grid paper (for calibration), with their left tarsus exposed and digits angled roughly perpendicularly to the tarsometatarsus (Supp. 1, Fig. 5; Digital measurements were used to minimise animal handling time and reduce risk of measurement error reported for analogue measurements (50; but see 51). Blurred images were removed from analysis and means taken when multiple images were available (61.5% of all images). From remaining images, length measurements were calculated in FIJI (52) as the calibrated straight-line distance between the ankle and the distal end of the tarsometatarsus (matching traditional analogue measurements; 50). A correlation between analogue and digital tarsus length measurements was confirmed with a subsample (n=43; β=0.728 [95% CI: 0.498, 0.975]; R^2^=0.563; BF>1000; prior for correlation between digital and analogue measures = skew-normal[ξ=0.25, ω=1, α=3]). In some cases, calibration paper was laid above the imaged tarsus (n=49 images; 10.0% of total) or replaced with a ruler (n=14 images; 2.9% of total). In these cases, tarsus lengths were adjusted by modelling the effect of calibration type on tarsus length (controlling for categorical weekly age) and subtracting mean effects from estimated tarsus lengths (grid over: β=3.260 [95% CI: 2.502, 4.021]; ruler under: β=2.406 [95% CI:0.801, 4.004]).

To confirm that body mass predicted structural size in quail, we measured the maximum external body height (the maximal straight-line distance between the base of the keel and spinal dorsum in the transverse plane) and synsacrum width (the maximal distance between the fossa renalii) of a subsample of adults (n = 20), then tested whether body mass predicted these measurements across individuals. Measurements were obtained after euthanasia (approximately 9 weeks of age) and collected using analogue and digital calipers, to the nearest 0.1 mm. To reduce observer biases, all skeletal measurements were collected by two independent researchers, and precision of each measurement subsequently confirmed (mean CVs = 1.78% and 2.24% for maximum body depth and synsacrum width respectively). For both skeletal measurements, strong correlations with body mass were detected via Bayesian linear models, indicating that body mass was a clear predictor of structural size (Supp. 1; Fig. 2; maximum body height: β=0.032 [95% CI: 0.009, 0.054], BF=194; synsacrum width: β=0.029 [95% CI: 0.010, 0.046], BF=499; priors for main effects normal with mean = 0 and s.d. = 2).

### Respirometry

Resting metabolic rate of quail was measured at 3-4 weeks and 8-9 weeks using flow-through respirometry methods described previously (49). Briefly, quail were placed in glass chambers (3.3 L at 3 weeks; 8.0 L at 8 weeks) ventilated with dried (via drierite; Sigma-Aldrich, Stockholm, Sweden) atmospheric air at least 30 minutes before measurement. To capture faeces and negate its effects on water loss estimates (i.e. via its evaporation), chambers were fitted with mineral oil reservoirs, over which a metallic mesh grid was placed for standing (53). All chambers were secured within a sealed climate chamber (Weiss Umwelttechnik C180, Reiskirchen, Germany) before bird placement, and the climate chamber set to 10°C (batches 1 and 2) or 30°C (batch 3) for acclimation. Throughout the experiment, air temperature was measured in chambers using thermocouples (36-gauge type T, copper- constantan; thermocouple box: TC-2000, Sable Systems) secured out of reach and influence from the birds. In total, 1-4 quail were held in our climate chamber at a time for any given set of measurements.

Following acclimation, chamber temperature was sequentially increased by 10°C increments until 40°C, with initial baseline (7-15 mins), measurement (10 mins per bird; totalling 30-40 mins), and terminal baseline periods (at least 5 mins) collected at each temperature. Air flow rates across temperatures averaged 2.2 L/min (± pooled sem = 0.014; 1.9-2.6 L/min) for 3 week old quail and 4.2 L/min (± sem=0.016; 3.7-4.8 L/min) for 8 week old quail (measured using a FB8 mass flow meter; Sable Systems, Las Vegas, NV, USA) in batches 1 and 2, from which we subsampled at, on average, 351 mL/min (308-388 mL/min) for analysis. In batch 3, mean flow rates were increased to 10.1 L/min (± sem= 0.006; 9.5-10.6 L/min) between 30°C and 40°C with subsampling averaging 403 mL/min (390-419 mL/min). Subsampling was achieved using a Sable Systems SS-4 sub-sampler and 99% equilibration times prior to subsampling ranged from 5.8-8.1 min and 7.8-10.1 min for 3- and 8-week- old quail respectively. To measure oxygen from subsampled air, we used a FC-10 analyser (Sable Systems) and stripped water vapour and carbon dioxide prior to measurement with drierite and ascarite (II; Acros Organics, Geel, Belgium). A RH-300 water vapour meter (Sable Systems) was used to measure water vapour. Calibration of oxygen and water vapour analysers is described elsewhere (49), and all birds showing signs of distress for >5 min were removed from study (n = 5 adults from batch 1 and n = 2 juveniles from batch 2).

To calculate oxygen consumption per bird, we extracted oxygen readings from the most stable 2 mins of 10 min recordings (batch 1 and 2) or from the most stable 2 mins after gas concentrations had stabilized (experiment 3) using the software ExpeData (version 1.9.27; Sable Systems), then converted these to mL O_2_/min following equations described in ref. 54. To calculate evaporative cooling efficiency, we first collected water vapour pressure readings from the same 2 min period then converted these to estimates of evaporative heat loss (W) following ref. 54 while assuming 2406 J consumed for every 1 mL of water evaporated (55). Evaporative cooling efficiency then represented the ratio of this evaporative heat loss value to metabolic heat production (estimated from O_2_ consumption and assuming 20 J = 1 mL O_2_; 56).

### Data organisation and statistical analyses

Data organisation and statistical analyses were conducted using R statistical software (version 4.2.3; 57) and package `brms` (58). Plots were produced using the package `ggplot2` (59). Model diagnostics (i.e. Hamiltonian Monte Carlo [HMC] chain diagnostics, prior predictive checks, posterior predictive checks, posterior density checks, residual visualisation etc.) were achieved visually, guided by ref. 60. For all models, Gelman-Rubin statistics 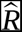 ; 61) exceeded 0.9 and ratios of effective sample sizes to sample sizes (N_eff_/N) exceeded 0.75, indicating strong chain convergence and little autocorrelation within HMC chains. To minimise bias from skewed posterior distributions, posterior estimates and credible intervals were calculated as medians and quantile intervals respectively, unless otherwise stated. Detailed description of analyses, prior derivations, and model validations (including R code) are provided in the supplemental material.

### Calculation of metabolic slopes

Metabolic slopes were calculated as the rate at which an individual increased their resting metabolism from thermoneutrality (30°C; 49,62) to our lowest (10°C) and highest (40°C) temperature exposures. Because we were interested in the efficiency with which an individual expended their own energy toward warming, or as consequence of heating, resting metabolism values (mL O_2_/min) at all temperatures were first relativised as an individual’s fold change from that observed at 30°C. Doing so allowed us to restrict among-individual variation in metabolism to that explained by differences in their response to a given temperature (i.e. by fixing resting metabolism at 30°C [thermoneutrality] for all individuals at 1) rather than also differences in their metabolism at thermoneutrality.

To estimate metabolic slopes in the cold, we used Bayesian linear mixed effects models with relative resting metabolism as our Gaussian-distributed response variable, ambient temperature (°C; 10°C, 20°C, and 30°C, encoded continuously) as the sole population-level predictor, and individual identity as a group-level slope; metabolic slopes per individual then represented their group-level slope of the metabolism by temperature relationship. Separate models were constructed for juveniles (3 weeks of age) and adults (8 weeks of age). Since, by definition, metabolism values were invariable at 30°C (equaling 1; see above), we set 30°C as our x-intercept and fixed our corresponding y-intercepts at 1. Equations for our model were therefore:

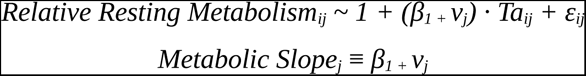

with *β_1_* representing the population-level effect of ambient temperature on relative metabolism, *v_ij_* representing an individuals’ deviation from that population-level effect (here, normally-distributed with a mean of 0 and variance *v_v_*), *i* representing an individual observation, *j* representing an individual, and *ε* representing a normally-distributed error term. To capture heterogeneity in relative metabolism observed across ambient temperature (Supp. 3 Fig. 7 & Supp. 3 Fig. 16), *ε* was allowed to vary between measurement temperatures.

In our adults, thermoneutrality extended below 30°C, ending at approximately 26°C (Supp. 5). Thus, to more accurately capture the linear relationship between ambient temperature and relative metabolism at this stage, we adjusted our x-intercept to 26°C (again, fixing our corresponding y-intercept to 1; linearity confirmed in Supp. 3 Fig. 16).

Priors for the population-level effect of ambient temperature on relative resting metabolism were informed by refs. 49 and 63, and set as skew-normal for all ages (*ξ*=-0.018, *ω*=0.02, *α*=-2.5; defining the mean near that reported by ref. 63). For the variance around individual slopes ( *v_v_*), we used a weak exponential prior (*λ*=2.5), and for that of our error term (*ε*) at 10°C (here, natural-log transformed to fix values above 0), we used a weak skew-normal prior (*ξ*=-0.2-, *ω*=1.0, *α*=-5). Variance is metabolism decreased at increases temperatures. As such, priors for the change in error between 10°C and 20°C were normal with a mean above 0 (0.25, SD = 0.25).

In response to heat, metabolic slopes were calculated manually as the change in relative metabolism observed among individual between 40°C and 30°C, divided by 10. Manual calculation was done as resting metabolism was only measured at these two temperatures.

### Repeatability of metabolic slopes

To calculated repeatability of metabolic slopes, we followed methods described by others (64). Repeatability values represented conditional repeatabilities (i.e. conditional on ambient temperature; 64) and were only calculated for responses to cold since we were unable to estimate within-individual variation explained by ambient temperature in the heat (owing to manual calculation). To statistically test our repeatability values against null expectations (which can be greater than 0), we again followed methods described by others (65). Briefly, models predicting relative metabolism in response to temperature we re-run but while randomly scrambling individual identities among measurements. Repeatabilities calculated from these new models (or “null models”) were then compared against those from our initial models using one-way hypothesis tests (here, using the Savage-Dickey density ratio method: 66). Priors for hypothesis tests are described in the supplemental material (Suppl. 3).

### Effects of morphology and rearing temperature on metabolic slopes

To partition direct effects of morphology and rearing temperature on metabolic slopes in the cold and heat, we used Bayesian path analyses composed of the following mixed effects models:

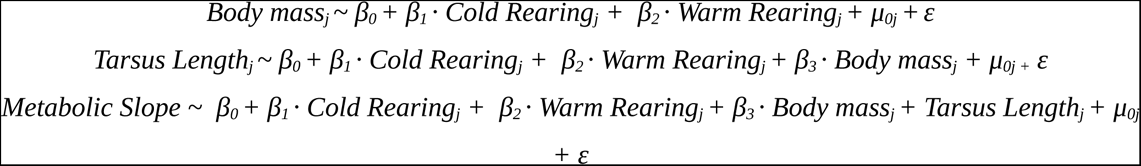

with residual variance between models assumed to be uncorrelated. *Cold* and *warm* rearing were binomial variables encoding whether an individual was reared at 10°C or 30°C respectively (0 equaling “no” and 1 equaling “yes”). *μ_0_* represents a group-level intercept of the egg batch that an individual was derived from. For the majority of models, response variables (and thus ε) were assumed to be normally distributed, and both continuous predictors and response variables were mean-centred to ease interpretation of model intercepts. Among juveniles in the heat, however, variance in metabolic slopes differed widely between egg batches (Supp. 3 Fig. 53), and was thus corrected by allowing error (ε) to vary by batch. Further, among adults in the heat, some metabolic slopes fell outside of expectations from a normally-distributed error but with no evidence of measurement error. To thus balance the influence of these individuals, a student’s-T error was assumed, centred on zero and with degrees of freedom calculated from our data. In all, four path analyses were constructed: two ultimately predicted metabolic slopes in the cold (one for juveniles and one for adults), and two ultimately predicting metabolic slopes in the heat (again, one for juveniles and one for adults).

For path analyses pertaining to juveniles, priors for the effects of cold and warming rearing on both body mass and tarsus length were normal and conservative (assuming no effect; means = 0, SDs = 15 for body mass, and SDs = 2.5 for tarsus length). Priors for body mass and tarsus length intercepts were also normal and conservative (means = 0, SDs = 5 and 2.5 for body mass and tarsus length respectively) and those for egg batch effects (*μ_0_*) on, and error (ε) around, body mass and tarsus length were exponential and weak (*μ_0_*: *λ* = 2.5 and 2; ε: *λ* = 0.15 and 1 for body mass and tarsus length respectively; derivation described in Suppl. 3). For path analyses pertaining to adults, priors on predictors of body mass and tarsus length were similar to those for juveniles but broadened to account for increased variance in each variable with age (body mass: intercept = *N*[0, 10], cold rearing = *N*[0, 25], warm- rearing = *N*[0, 25], batch effects = exponential[2.5], error = exponential[0.15]; tarsus length: *N*[0, 2.5], cold rearing = *N*[0, 2.5], warm- rearing = *N*[0, 2.5], batch effects = exponential[2], error = exponential[1]). At all ages, priors for the effect of body mass on tarsus length with skew-normal, following visually-confirmed, positive allometry (*ξ*=0, *ω*=0.25, *α*=5).

For models predicting metabolic slopes in the cold (as part of our path analyses), priors for juveniles and adults were uninformed and as follows: intercepts = *N*(0, 0.01), cold rearing = *N*(0, 0.025), warm rearing = *N*(0, 0.025), mass = *N*(0, 1.0×10^-3^), tarsus length = *N*(0, 2.5×10^-3^), egg batch effects = exponential(50), and error = exponential(10). These priors assumed no previous evidence that each parameter predicted metabolic slopes, and that effects of body mass and tarsus length greater than the range of metabolic slopes divided by their own ranges were unlikely. For models predicting metabolic slopes in the warmth, uninformed priors were also used and were as follows for juveniles and adults respectively: intercepts = *N*(0, 0.125) and *N*(0, 0.1), cold rearing = *N*(0, 0.125) and *N*(0, 0.1), warm rearing = *N*(0, 0.125) and *N*(0, 0.1), mass = *N*(0, 4.0×10^-3^) and *N*(0, 2.5×10^-3^), tarsus length = *N*(0, 0.015) and *N*(0, 0.03), egg batch effects = exponential(50). For juveniles, among which our error term varied by egg batch, error term priors were as follows (again, natural-log transform to fix values above 0): egg batch A = *N*(-3, 1.5), change from egg batch 1 and egg batch 2 = *N*(0, 0.5), and change from egg batch 1 and egg batch 3 = *N*(1, 1.5). For adults, our prior for the centrality of our error term was exponential (10), and that for our degrees of freedom (ν) was conservatively gamma-distributed (ɑ = 10, β = 1).

### Effects of morphology and rearing temperature on evaporative cooling efficiency

Effects of morphology and rearing condition on evaporative cooling efficiency (here, at 40°C) were again partitioned using Bayesian path analyses. Path analyses followed the same structure as those ultimately predicting metabolic slopes and were as follows:

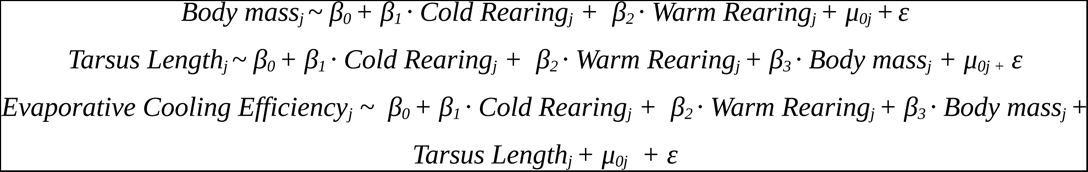

Priors for all predictors of body mass and tarsus length at each life stage remained identical to those described above. For predictors of evaporative cooling efficiency, priors for juveniles and adults were as follows: intercepts = *N*(0.5, 0.2) and *N*(0.75, 0.2), cold rearing = *N*(0, 0.25), warm rearing = *N*(0, 0.25), mass = *N*(0, 0.1), tarsus length = *N*(0, 0.1), egg batch effects = exponential(15). Intercept priors were informed from ref. 49 and again, those for body mass and tarsus length assumed that effects greater than the range of evaporative cooling efficiency divided by their own ranges were unlikely. Given that error in evaporative cooling efficiency again varied by egg batch among juveniles (Supp. 4 Fig. 7), priors for our error term at this age class was set as: egg batch 1 = *N*(-2, 1), change from egg batch 1 to egg batch 2 = *N*(0, 0.5), change from egg batch 1 to egg batch 3 = *N*(1, 1). For adults, the prior for our error term was exponential (*λ*=5).

### Effects of rearing temperature on mass gain and tarsus elongation

To evaluate whether and how environmental temperature shaped mass gain and tarsus elongation after hatching, we modelled body mass (g) and tarsus length (mm) as Gompertz functions of developmental age, in weeks, from hatching until maturity (8 weeks). Gompertz parameters (*a*, the asymptote, *b*, the x-axis displacement, and *c*, the growth rate) were then each modelled as functions of cold rearing (binomial; 0 = “no”, 1 = “yes”), warm rearing (binomial; 0 = “no”, 1 = “yes”), and egg batch. Here, cold rearing and warm rearing were treated as population-level predictors, and egg batch a group-level intercept. To account for repeated measurements of individuals across all ages, individual identity was also included as a group-level predictor (here, intercept) of body mass and tarsus length in addition to Gompertz parameters. Further, to evaluate and adjust for effect of body size scaling on tarsus length measurements, body mass, mean-centred by week of measurement, was also included as a population-level predictor of tarsus length in alongside Gompertz parameters. Our models for body mass and tarsus length were therefore as follows:

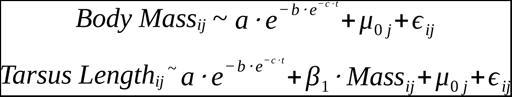

where “*t*” represents age in weeks, *μ_0j_* represents a group-level intercept of individual identity, “*Mass*” represents the relative mass of an individual *j* at a given week (*t*) of measurement relative to the mean at that week, and:

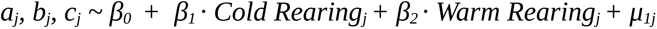

where *μ*_1j_ represents the group-level intercepts of egg batch on each Gompertz parameter. Last, because variance in body mass and tarsus length increased natural-logarithmically with age, error terms (*ε*) was modelled the following:

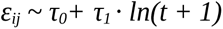

where τ_0_ indicates the error intercept, and τ_1_ indicates the rate at which error increased across the natural-log of time + 1.

Priors for our model asymptotes (Gompertz *a*) were informed by data from ref. 43 while those for other Gompertz parameters were informed by refs. 49 and 67. For our model predicting body mass, priors were as follows: Gompertz *a* intercept = *N*(250, 25), effect of cold rearing on *a* = *N*(7.5, 25), effect of warm rearing on *a* = *N*(-7.5, 25), effect of egg batch on *a* = exponential(2.5); Gompertz *b* intercept *= N*(3, 1), effects of cold and warm rearing on *b* = *N*(0, 0.5), effect of egg batch on *b =* exponential(10); Gompertz *c* intercept = skew-normal(0.5, 0.1, 2.5; assuming no negative growth), effects of cold and warm rearing on *c* = *N*(0, 0.2), effect of egg batch on *c* = exponential(25); effect of individual identity (*μ_0j_*) on mass = exponential(0.5); error term intercept (τ_0_; natural log-transformed to fix above 0) = skew-normal(1, 0.5, -10); effect of age on error term τ_1_ = skew-normal(1, 0.5, 10). For our model predicting tarsus length, priors remained similar and were: Gompertz *a* intercept = *N*(37.5, 2.5), effect of cold rearing on *a* = *N*(-0.35, 2), effect of warm rearing on *a* = *N*(0.35, 2), effect of egg batch on *a* = exponential(1); Gompertz *b* intercept *=* skew-normal(0.6, 0.5, 2.5), effects of cold and warm rearing on *b* = *N*(0, 0.5), effect of egg batch on *b =* exponential(10); Gompertz *c* intercept = skew-normal(0.5, 0.1, 2.5; assuming no regression), effects of cold and warm rearing on *c* = *N*(0, 0.25), effect of egg batch on *c* = exponential(25); effect of scaled body mass on tarsus length = skew-normal(0.5, 0.15, 5); effect of individual identity (*μ_0j_*) on tarsus length = exponential(1.5); error term intercept (τ_0_; again, natural log-transformed) = skew-normal(1, 0.5, 10); effect of age on error term τ_1_ = skew-normal(1, 0.5, 10).

### Effect size calculation

Where reported, Cohen’s D values (68) were calculated as the difference in raw posterior predictions (n = 1000 samples) between comparison states (i.e. average tarsus length and a tarsus length 1 SD above average), divided by the standard deviation of posterior predictions from our original path analyses.

## Acknowledgements

We thank Camilla Björklund for her invaluable assistance with animal care and Fredrik Andreasson for his helpful comments on an earlier version of this manuscript.

## Conflict of Interests

The authors have no conflicts of interest to report.

## Data Availability

All data and statistical code required to reproduce this study are provided in the supplement.

## Funding Statement

Funding for this research was provided by the Wenner-Gren Foundation (UP2021-0038), the Craaford Foundation (20211007, 20221018), the Royal Physiographic Society of Lund (2021-41891, 20221103), Stiftelsen Lunds Djurskyddsfond, the Lars Hierta Minne (20221121), and the Swedish Research Council (2020-04686).

## Notes

### Competing Interest Statement

The authors have declared no competing interest.

