## Supplementary material for "No evidence that shrinking and shapeshifting meaningfully affect how birds respond to warming and cooling": Supp. 1

### Size Measurement Validations

#### Validating body mass as a proxy for structural size in Japanese quail

In our study, we use body mass as a proxy for body size (or structural size) in quail. However, tarsus length is also often assumed as a proxy of structural size in other avian species (e.g. Weeks et al, 2020). If the latter is indeed true but not the former, effects of body mass on thermal physiology, conditional upon tarsus length, may be more suitably interpreted as effects of body *condition* rather than body *size* on such. Conversely, effects of tarsus length on thermal physiology, conditional upon body mass, may be interpreted as effects of *inverse* body condition, rather than mere appendage length, on such. For example, a positive effect of tarsus length on metabolic slopes in the cold, when body mass is set at its average (commonly our model intercepts) may indicate a negative effect of body condition on thermal resistance, rather than a positive effect of appendage length.

To assess whether body mass or tarsus length are indeed predictors of structural size, we tested how well each predicted other measures of structural size in a sub-sample of mature and sacrificed Japanese quail (n = 20). To do so, we began by measuring tarsus length (here, digital, as described in Tabh et al, 2024) and five other possible indicators of structural size, including: (1) maximum keel depth (mm), (2) maximum skull height (mm), (3) maximum body width (mm), (4) synsacrum width, and (5) maximum body height (from back to deepest point of keel); measurements were collected by two independent measurers (J.T. and E.P.). Following measurement, we then quantified precision of each structural size metric by calculating the mean coefficient of variation (CV) per metric across all birds measured. Metrics with the lowest mean CV were considered to be the most precise and those with the highest mean CV the least precise. Finally, the two size metrics with the lowest mean CV were regressed against tarsus length and body mass using simple Bayesian linear regressions.

We begin below by loading in packages and functions necessary for our analysis.

```
def.chunk.hook <- knitr::knit_hooks$get("chunk")
knitr::knit_hooks$set(chunk = function(x, options) {
  x <- def.chunk.hook(x, options)
  ifelse(options$size != "normalsize",
    paste0("\n \\", options$size, "\n\n", x, "\n\n \\normalsize"), x
  )
})

knitr::opts_chunk$set(fig.pos = "H", out.extra = "")
knitr::opts_chunk$set(size = "footnotesize")

# First loading in packages

library("tidyverse")
library("easypackages")
```

```
packageList <- c("bayesplot", "brms", "brmsMethods",
                 "doParallel", "foreach", "ggpubr",
                 "kableExtra", "latex2exp", "patchwork",
                 "priorsense", "showtext", "tidybayes",
                 "wesanderson")

libraries(packageList)

caption <- paste0("R packages and their respective versions used for",
                  " data organisation and analysis in this study."
)

sapply(packageList, function(x) {
  y <- as.character(packageVersion(x))
  return(y)
}, simplify = FALSE) %>%
  enframe(., name = "Package", value = "Version") %>%
  as.data.frame(.) %>%
  kbl(.,
    longtable = T, booktabs = T,
    caption = caption
  ) %>%
  kable_styling(latex_options = "striped")
```

***Table 1:*** *R packages and their respective versions used for data organisation and analysis in this study.*

| Package | Version |
|---|---|
| bayesplot | 1.11.1 |
| brms | 2.20.4 |
| brmsMethods | 0.0.0.9000 |
| doParallel | 1.0.17 |
| foreach | 1.5.2 |
| ggpubr | 0.6.0 |
| kableExtra | 1.3.4 |
| latex2exp | 0.9.6 |
| patchwork | 1.2.0 |
| priorsense | 0.0.0.9000 |
| showtext | 0.9.6 |
| tidybayes | 3.0.4 |
| wesanderson | 0.3.6.9000 |

```
# Loading additional functions

pp_check2 <- function(model, resp = NA, ndraws = 500,
                      xlab = "label", colour = "lightblue") {
  require(brms)
  require(ggplot2)
  stopifnot("Model must be a brmsfit object" = is.brmsfit(model))

  if (is.na(resp)) {
    resp <- model$formula$resp
```

```
  }

  p1 <- brms::pp_check(model, ndraws = ndraws, resp = resp) +
    scale_colour_manual(
      values = c("black", colour),
      labels = c("y", "yhat"),
      name = NULL
    ) +
    xlab(xlab) +
    ylab("Density") +
    theme_classic()
  return(p1)
}

chainCheck <- function(model, rDig = 3) {
  require(brms)
  stopifnot("Model must be a brmsfit object" = is.brmsfit(model))

  Rhat <- paste0(
    "Rhat range: ",
    round(min(rhat(model)), digits = rDig),
    " - ",
    round(max(rhat(model)), digits = rDig)
  )
  Neff <- paste0(
    "Neff/N range: ",
    round(min(neff_ratio(model)), digits = rDig),
    " - ",
    round(max(neff_ratio(model)), digits = rDig)
  )
  cat(paste0(Rhat, "\n", Neff))
}

quantileCIs <- function(x, rnd = 3, cis = c(50, 95), sci_note = FALSE) {
  require(tidyverse)

  if (class(x)[1] != "brmsfit") {
    return("x must be a brmsfit object.")
  }
  if (length(cis) != 2) {
    return("cis must be a vector of integers with length 2")
  }

  prbs = c()
  nColNames = c()
  for (i in 1:length(cis)){
    prbs = c(prbs, c(0.5 - (cis[i]/100)/2, 0.5 + (cis[i]/100)/2))
    nColNames = c(nColNames,
                  paste0("Low_CI_", cis[i]),
                  paste0("High_CI_", cis[i])
    )
  }
```

```
  modelFrame = as.data.frame(x)

  Results <- apply(modelFrame, MARGIN = 2, FUN = quantile,
        probs = prbs, type = 8) %>%
    t() %>%
    as.data.frame() %>%
    rownames_to_column(var = "par") %>%
    `colnames<-`(c("Parameter", nColNames))

  if (sci_note == FALSE) {
    Results <- apply(modelFrame, MARGIN = 2, FUN = quantile,
        probs = prbs, type = 8) %>%
    t() %>%
    as.data.frame() %>%
    rownames_to_column(var = "par") %>%
    `colnames<-`(c("Parameter", nColNames))

  } else if (sci_note == TRUE) {
    Results <- apply(modelFrame, MARGIN = 2, FUN = quantile,
        probs = prbs, type = 8) %>%
    t() %>%
    as.data.frame() %>%
    rownames_to_column(var = "par") %>%
    `colnames<-`(c("Parameter", nColNames)) %>%
    mutate_at(.vars = vars(-Parameter),
              .funs = function(x){
                return(format(x, scientific = TRUE))
              }
    )
  }

  return(Results)
}

# Installing font

font_add_google(name = "Noto Sans", family = "Noto Sans")

# Setting working directory

setwd(paste0("/Users/joshuatabh/Documents/",
  "researchProjects/lund/functionOfAllen/functionData")
)
```

Next, we load in our validation data and plot CVs by size metric.

```
# Loading in data.

structuralData <- read.csv("structuralSizeData.csv") %>%
  select(-c("measurer", "date", "headPhoto", "legPhoto", "notes")) %>%
  group_by(ring) %>%
  summarise_all(.funs = list("Mean" = mean, "CV" = function(x) {
    sd(x) / mean(x)
  })) %>%
  merge(., read.csv("digitalDataForStructuralSizeAnalyses.csv"),
```

```
    by = "ring"
  ) %>%
  select(-wingLength_CV)

# Renaming columns

colnames(structuralData) <- gsub(
  "_", "",
  colnames(structuralData)
)

# Plotting coefficients of variation for structural
# size measures

plotOrder <- structuralData %>%
  select(
    bodyWidthCV, bodyHeightCV, keelDepthCV,
    synsacrumWidthCV, skullHeightCV
  ) %>%
  pivot_longer(everything(),
    names_to = "metric",
    values_to = "CV"
  ) %>%
  merge(., tribble(
    ~Metric, ~metric,
    "Body Width (mm)", "bodyWidthCV",
    "Body Height (mm)", "bodyHeightCV",
    "Keel Depth (mm)", "keelDepthCV",
    "Synsacrum Width\n(mm)", "synsacrumWidthCV",
    "Skull Height (mm)", "skullHeightCV"
  ), by = "metric") %>%
  group_by(Metric) %>%
  summarise("Mean" = mean(CV)) %>%
  arrange(desc(Mean)) %>%
  pull(Metric)

showtext_auto()

cvPlot <- structuralData %>%
  select(
    bodyWidthCV, bodyHeightCV, keelDepthCV,
    synsacrumWidthCV, skullHeightCV
  ) %>%
  pivot_longer(everything(),
    names_to = "metric",
    values_to = "CV"
  ) %>%
  merge(., tribble(
    ~Metric, ~metric,
    "Body Width (mm)", "bodyWidthCV",
    "Body Height (mm)", "bodyHeightCV",
    "Keel Depth (mm)", "keelDepthCV",
    "Synsacrum Width\n(mm)", "synsacrumWidthCV",
    "Skull Height (mm)", "skullHeightCV"
  ), by = "metric") %>%
  select(Metric, CV) %>%
  mutate(
    "CV" = CV * 100,
    Metric = factor(Metric, levels = plotOrder)
  ) %>%
  ggplot(aes(x = Metric, y = CV, fill = Metric)) +
  geom_point(
    pch = 21, size = 2.5, colour = "black", alpha = 0.5,
    position = position_jitter(width = 0.3)
  ) +
  stat_summary(
    geom = "errorbar", fun.data = "mean_se",
```

```
    colour = "black", width = 0.3
  ) +
  stat_summary(
    geom = "point", fun = "mean",
    pch = 21, size = 4, colour = "black"
  ) +
  geom_segment(aes(x = 4.5, y = 7.5, xend = 4.05, yend = 3),
    size = 0.4,
    arrow = arrow(length = unit(0.2, "cm"))
  ) +
  geom_segment(aes(x = 4.5, y = 7.5, xend = 4.95, yend = 2.7),
    size = 0.4,
    arrow = arrow(length = unit(0.2, "cm"))
  ) +
  ylab("Coefficient of Variation (%)") +
  scale_fill_manual(
    values =
      c(
        "#ff9690", "#ffada9",
        "#fec4c1", "#fee3e2",
        "#fff3f2"
      )
  ) +
  theme_classic() +
  theme(
    axis.title.x = element_blank(),
    legend.position = "none",
    axis.title.y = element_text(family = "Noto Sans"),
    axis.text = element_text(family = "Noto Sans")
  )

cvPlot
```

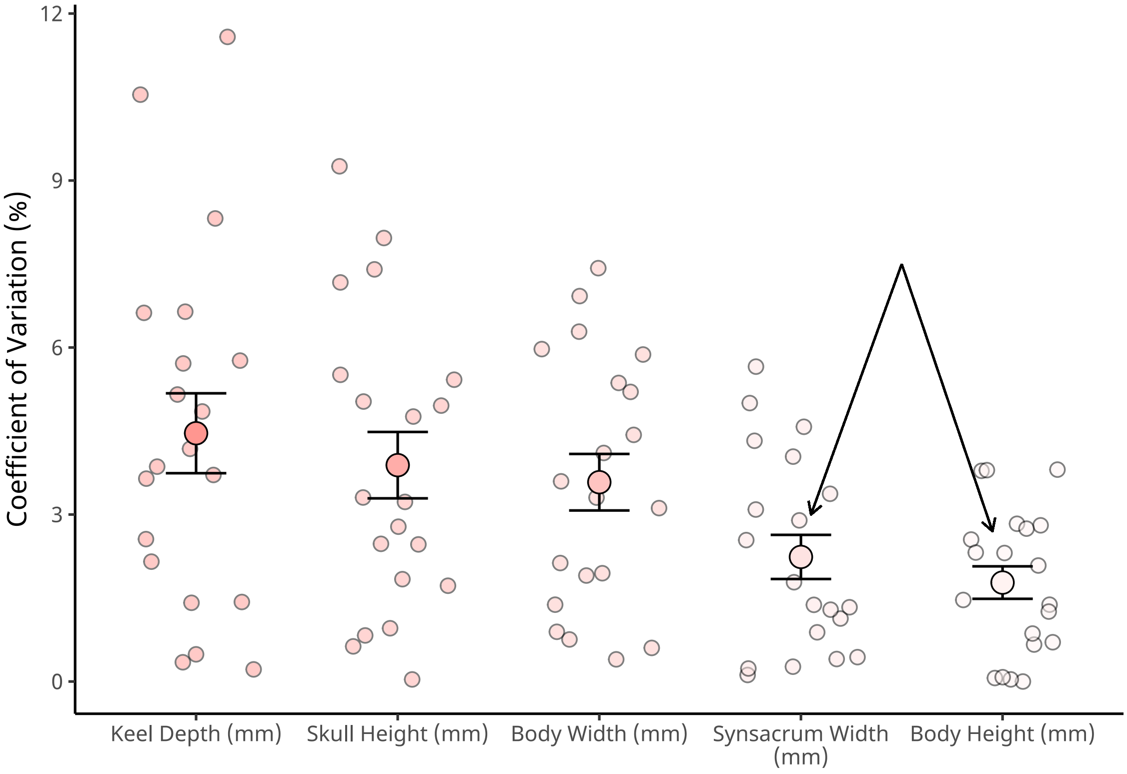


***Figure 1:*** *Precision of skeletal size metrics derived from mature and deceased Japanese quail (n = 20) by two observers. Small dots indicate raw coefficients of variation (CVs) per bird, and large dots indicate mean CVs per size metric. Errorbars indicate +/- one standard error around means. The black errors identify the two skeletal size metrics with highest precision (i.e. lowest mean CV).*

```
ggsave("../plots/structuralSizeCVs.pdf",
  cvPlot,
  dpi = 800, height = 7, width = 8
)

showtext_auto(enable = FALSE)
```

Synsacrum width and maximum body height, being the most precise metrics of skeletal size, are selected for use in further analyses. Next, tentative correlations between tarsus length, body mass, and each selected size metric are visualised.

```
# Plotting tarsus and body mass by two structural size
# metrics with the lowest coefficients of variation
# (body height and synsacrum width)

showtext_auto()

p1 <- structuralData %>%
  select(ring,
    tarsusLengthMean,
    "Body Height (mm)" = bodyHeightMean,
    "Synsacrum Width\n(mm)" = synsacrumWidthMean
  ) %>%
  pivot_longer(c("Body Height (mm)", "Synsacrum Width\n(mm)"),
    names_to = c("par"), values_to = c("measure")
  ) %>%
  ggplot(aes(x = tarsusLengthMean, y = measure, fill = par)) +
  facet_wrap(~par,
```

```
    scales = "free",
    strip.position = "left"
  ) +
  geom_point(
    pch = 21, size = 2.5,
    colour = "black", alpha = 0.7
  ) +
  geom_smooth(
    method = "lm", se = FALSE,
    colour = "black", linetype = "dashed"
  ) +
  scale_fill_manual(values = c("#fff3f2", "#fee3e2")) +
  xlab("Tarsus Length (mm)") +
  theme_classic() +
  theme(
    legend.position = "none",
    strip.background = element_blank(),
    strip.placement = "outside",
    axis.title.y = element_blank(),
    strip.text = element_text(
      size = 12,
      colour = "black",
      family = "Noto Sans"
    ),
    axis.title.x = element_text(
      size = 12,
      colour = "black",
      family = "Noto Sans"
    )
  )

p2 <- structuralData %>%
  select(ring,
    massMean,
    "Body Height (mm)" = bodyHeightMean,
    "Synsacrum Width\n(mm)" = synsacrumWidthMean
  ) %>%
  pivot_longer(c("Body Height (mm)", "Synsacrum Width\n(mm)"),
    names_to = c("par"), values_to = c("measure")
  ) %>%
  ggplot(aes(x = massMean, y = measure, fill = par)) +
  facet_wrap(~par, scales = "free", strip.position = "left", ) +
  geom_point(
    pch = 21, size = 2.5,
    colour = "black", alpha = 0.7
  ) +
  geom_smooth(
    method = "lm", se = FALSE,
    colour = "black", linetype = "dashed"
  ) +
  scale_fill_manual(values = c("#fff3f2", "#fee3e2")) +
  xlab("Body Mass (g)") +
  theme_classic() +
  theme(
    legend.position = "none",
    strip.background = element_blank(),
    strip.placement = "outside",
    axis.title.y = element_blank(),
    strip.text = element_text(
      size = 12,
      colour = "black",
      family = "Noto Sans"
    ),
    axis.title.x = element_text(
      size = 12,
      colour = "black",
      family = "Noto Sans"
```

```
    )
  )

structuralSizePlot <- p1 / p2
structuralSizePlot
```

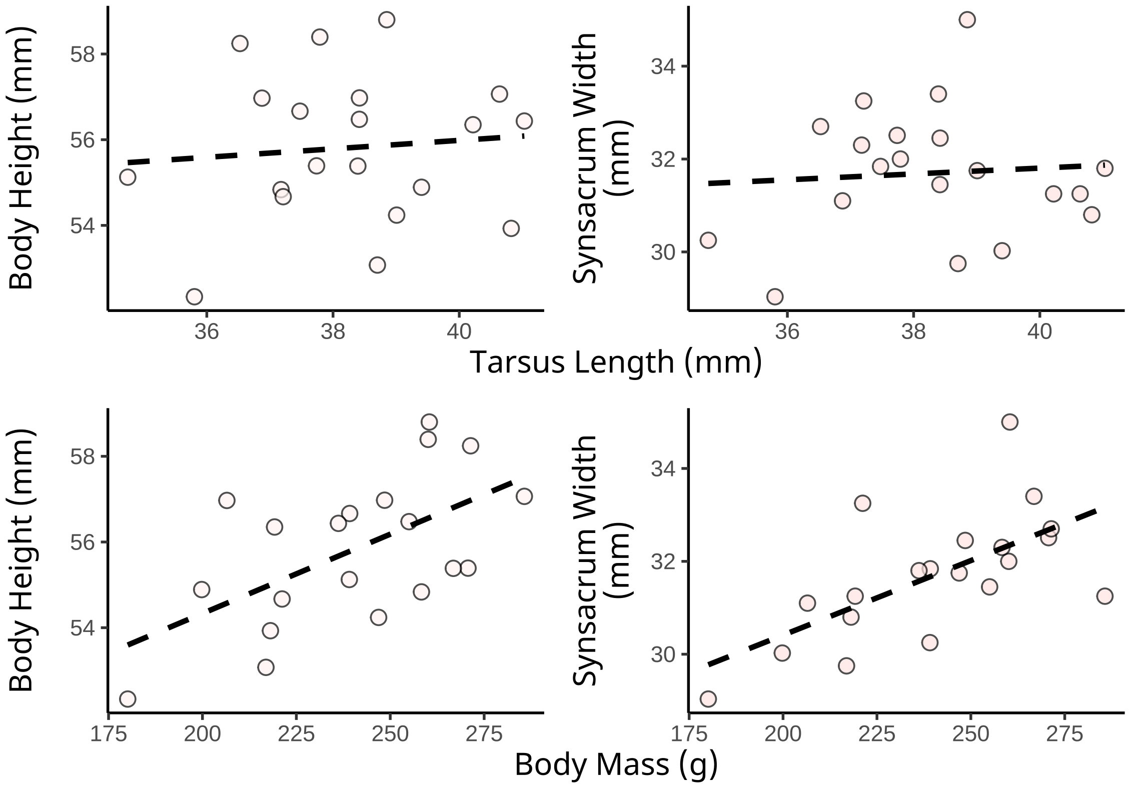


***Figure 2:*** *Relationship between tarsus length (mm) or body mass (g) and both maximum body height (mm) and synsacrum width (mm) in twenty mature and deceased Japanese quail. Dots indicate raw measurements and dashed lines indicate predicted linear relationships estimated by the R package ggplot2 (Wickham, 2011). All measurements were collected using digital and analogue calipers.*

```
ggsave("../plots/tarsusMassByStructuralSizeMeasures.pdf",
  structuralSizePlot,
  dpi = 800, height = 7, width = 8
)

showtext_auto(enable = FALSE)
```

Last, linear regressions between tarsus length and both synsacrum width and body height are constructed, followed by regressions between body mass and both synsacrum width and body height. In these models, all size metrics are mean-centred prior to analysis to simplify interpretation of model intercepts. Priors for slope coefficients were normal, centred at zero, and wide, with 2.5 percentiles equaling the range of our observed response variable divided by the range of our observed dependent variable. All intercept priors were also normal, centred at zero, and with standard deviations of 2, and error ($\epsilon$) priors were exponential with scaling factors of 1.

```
# Running basic linear models to compare predictive
# capacity of tarsus length and body mass on
# structural size metrics. Note that only coefficients,
# 95% hdis and r2 values are pulled from each model.
# Priors for each slope are weak and wide. Results
```

```
# from body mass as predictor are added for comparison
# purposes.

upperLimit <- with(
  structuralData,
  diff(range(bodyHeightMean, na.rm = T)) /
    diff(range(tarsusLengthMean, na.rm = T))
)

tarsusHeightModel <- brm(
  data = structuralData %>%
    select(
      "bodyHeight" = bodyHeightMean,
      "tarsus" = tarsusLengthMean
    ) %>%
    mutate(
      "bodyHeight" = bodyHeight -
        mean(bodyHeight, na.rm = T),
      "tarsus" = tarsus - mean(tarsus, na.rm = T)
    ),
  family = "gaussian",
  bodyHeight ~ tarsus,
  prior = c(
    set_prior("normal(0, 2)", class = "Intercept"),
    set_prior(paste0("normal(0, ", upperLimit / 2, ")"),
      class = "b"
    ),
    set_prior("exponential(1)", class = "sigma")
  ),
  iter = 50000, warmup = 10000, cores = 4, chains = 4, thin = 20,
  silent = TRUE, refresh = 0,
  file = "./models/tarsusBodyHeight.Rds"
)

p1 <- pp_check2(tarsusHeightModel,
  xlab = "Max. Body Height (mm;\nMean-Centred)"
) +
  theme(
    legend.position = "none"
  )

p2 <- tarsusHeightModel$data %>%
  mutate(
    "Residuals" =
      residuals(tarsusHeightModel,
        type = "ordinary",
        robust = TRUE
      )[, "Estimate"]
  ) %>%
  ggplot(aes(x = Residuals)) +
  geom_density(
    colour = "black",
    fill = "lightblue", alpha = 0.5
  ) +
  xlab("Maximum Body Height\nResiduals (mm)") +
  ylab("Density") +
  theme_classic()

p3 <- tarsusHeightModel$data %>%
  mutate(
    "Residuals" =
      residuals(tarsusHeightModel,
        type = "ordinary",
        robust = TRUE
      )[, "Estimate"]
  ) %>%
  ggplot(aes(x = tarsus, y = Residuals)) +
```

```
  geom_point(
    size = 2, pch = 21, colour = "black",
    fill = "lightblue", alpha = 0.5
  ) +
  xlab("Tarsus Length (mm)") +
  ylab("Max. Body Height\nResiduals (mm)") +
  theme_classic()

upperLimit <- with(
  structuralData,
  diff(range(synsacrumWidthMean, na.rm = T)) /
    diff(range(tarsusLengthMean, na.rm = T))
)

tarsusSynsacrumModel <- brm(
  data = structuralData %>%
    select(
      "synsacrumWidth" = synsacrumWidthMean,
      "tarsus" = tarsusLengthMean
    ) %>%
    mutate(
      "synsacrumWidth" = synsacrumWidth -
        mean(synsacrumWidth, na.rm = T),
      "tarsus" = tarsus - mean(tarsus, na.rm = T)
    ),
  family = "gaussian",
  synsacrumWidth ~ tarsus,
  prior = c(
    set_prior("normal(0, 2)", class = "Intercept"),
    set_prior(paste0("normal(0, ", upperLimit / 2, ")"),
      class = "b"
    ),
    set_prior("exponential(1)", class = "sigma")
  ),
  iter = 50000, warmup = 10000, cores = 4, chains = 4, thin = 20,
  silent = TRUE, refresh = 0,
  file = "./models/tarsusSynsacrumWidth.Rds"
)

p4 <- pp_check2(tarsusSynsacrumModel,
  xlab = "Max. Synsacrum Width\n(mm; Mean-Centred)"
) +
  theme(legend.position = "none")

p5 <- tarsusSynsacrumModel$data %>%
  mutate(
    "Residuals" =
      residuals(tarsusSynsacrumModel,
        type = "ordinary",
        robust = TRUE
      )[, "Estimate"]
  ) %>%
  ggplot(aes(x = Residuals)) +
  geom_density(
    colour = "black", fill = "lightblue",
    alpha = 0.5
  ) +
  xlab("Max. Synsacrum Width\nResiduals (mm)") +
  ylab("Density") +
  theme_classic()

p6 <- tarsusSynsacrumModel$data %>%
  mutate(
    "Residuals" =
      residuals(tarsusSynsacrumModel,
        type = "ordinary",
        robust = TRUE
```

```
      )[, "Estimate"]
  ) %>%
  ggplot(aes(x = tarsus, y = Residuals)) +
  geom_point(
    size = 2, pch = 21, colour = "black",
    fill = "lightblue", alpha = 0.5
  ) +
  xlab("Tarsus Length (mm)") +
  ylab("Max. Synsacrum Width\nResiduals (mm)") +
  theme_classic()

(p1 + p2 + p3)/
  (p4 + p5 + p6) +
  plot_annotation(tag_levels = "A")
```

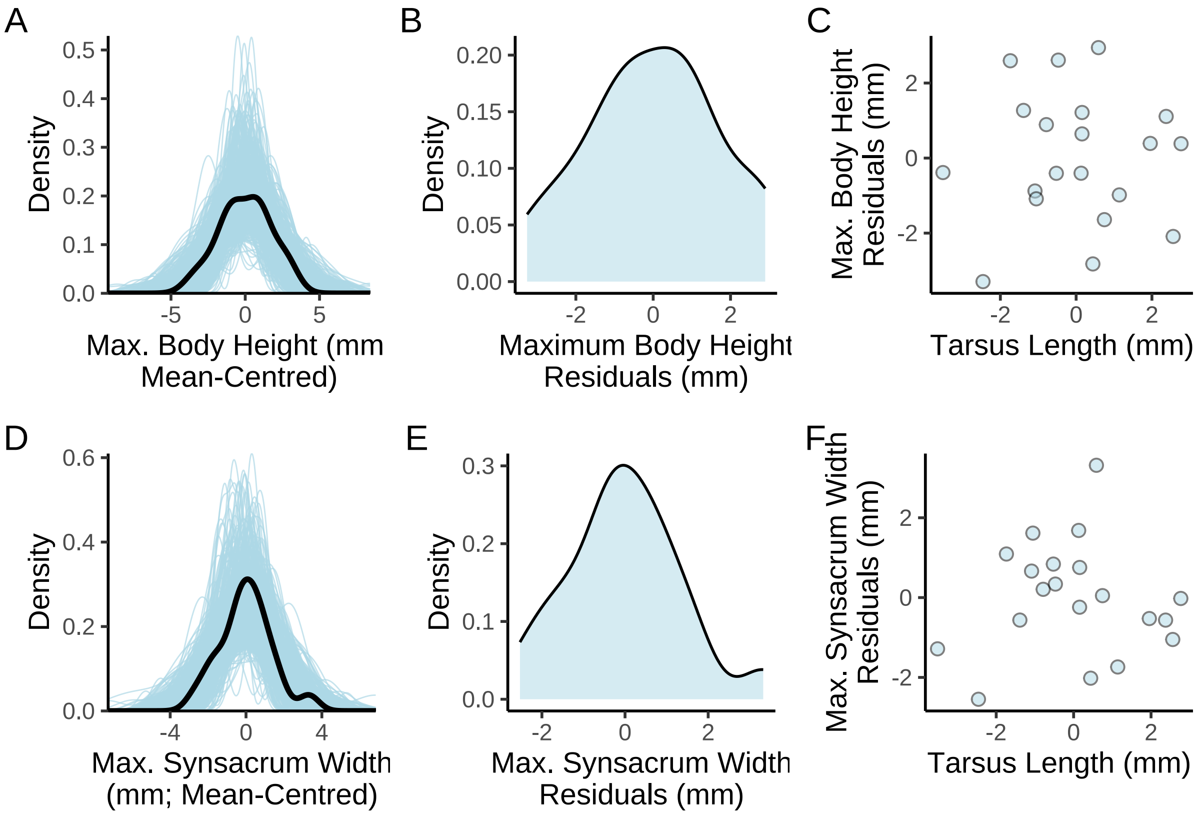


***Figure 3:*** *Validations for linear models with tarsus length (mm) predicting maximum body height (mm) and maximum synsacrum width (mm) in twenty mature and deceased Japanese quail. Maximum body height represents the maximum vertial distance between the keel and back of feathered quail. Maximum synsacrum width represents that measured after dissection. Panels A and D represent posterior predictive checks, with black lines indicating true density of response variable values and light blue lines indicating estimated densities, as drawn from model posteriors. All other panels display spread (density in panels B and E) of model residuals.*

```
# Collating results

tarsus <- data.frame(
  "tarsusHeight" = as.data.frame(tarsusHeightModel)$b_tarsus,
  "tarsusSynsacrum" = as.data.frame(tarsusSynsacrumModel)$b_tarsus
) %>%
  summarise_all(., .funs = median) %>%
  pivot_longer(everything(),
    names_to = "Parameter",
    values_to = "Estimate"
```

```
  ) %>%
  merge(.,
    bind_rows(
      quantileCIs(tarsusHeightModel, cis = c(50, 95)) %>%
        mutate(Parameter = ifelse(Parameter == "b_tarsus", "tarsusHeight", Parameter)),
      quantileCIs(tarsusSynsacrumModel, cis = c(50, 95)) %>%
        mutate(Parameter = ifelse(Parameter == "b_tarsus", "tarsusSynsacrum", Parameter))
    ),
    by = "Parameter", all.x = TRUE
  ) %>%
  rowwise() %>%
  mutate("BF" = ifelse(Parameter == "tarsusHeight",
    ifelse(Estimate < 0,
      (2 * mean(as.data.frame(
        tarsusHeightModel
      )[, "b_tarsus"] <= 0)) /
        (2 * mean(as.data.frame(
          tarsusHeightModel
        )[, "b_tarsus"] >= 0)),
      (2 * mean(as.data.frame(
        tarsusHeightModel
      )[, "b_tarsus"] >= 0)) /
        (2 * mean(as.data.frame(
          tarsusHeightModel
        )[, "b_tarsus"] <= 0))
    ),
    ifelse(Estimate < 0,
      (2 * mean(as.data.frame(
        tarsusSynsacrumModel
      )[, "b_tarsus"] <= 0)) /
        (2 * mean(as.data.frame(
          tarsusSynsacrumModel
        )[, "b_tarsus"] >= 0)),
      (2 * mean(as.data.frame(
        tarsusSynsacrumModel
      )[, "b_tarsus"] >= 0)) /
        (2 * mean(as.data.frame(
          tarsusSynsacrumModel
        )[, "b_tarsus"] <= 0))
    )
  )) %>%
  ungroup() %>%
  mutate(
    "Response" = c("Body Height (mm)", "Synsacrum Width (mm)"),
    "Predictor" = "Tarsus Length (mm)",
    "Estimate" = round(Estimate, digits = 4),
    "BF" = round(BF, digits = 4),
    `50\\% CIs` = paste0(
      "[", round(Low_CI_50, digits = 4),
      ", ", round(High_CI_50, digits = 4),
      "]"
    ),
    `95\\% CIs` = paste0(
      "[", round(Low_CI_95, digits = 4),
      ", ", round(High_CI_95, digits = 4),
      "]"
    ),
    "N" = 20
  ) %>%
  select(
    Predictor, Response, N, Estimate,
    `50\\% CIs`, `95\\% CIs`, BF
  )
```

```
# Body mass models

upperLimit <- with(
  structuralData,
```

```
  diff(range(bodyHeightMean, na.rm = T)) /
    diff(range(massMean, na.rm = T))
)

massHeightModel <- brm(
  data = structuralData %>%
    select(
      "bodyHeight" = bodyHeightMean,
      "mass" = massMean
    ) %>%
    mutate(
      "bodyHeight" = bodyHeight -
        mean(bodyHeight, na.rm = T),
      "mass" = mass - mean(mass, na.rm = T)
    ),
  family = "gaussian",
  bodyHeight ~ mass,
  prior = c(
    set_prior("normal(0, 2)", class = "Intercept"),
    set_prior(paste0("normal(0, ", upperLimit / 2, ")"),
      class = "b"
    ),
    set_prior("exponential(1)", class = "sigma")
  ),
  iter = 50000, warmup = 10000, cores = 4,
  chains = 4, thin = 20,
  silent = TRUE, refresh = 0,
  file = "./models/massBodyHeightModel.Rds"
)

p1 <- pp_check2(massHeightModel,
  xlab = "Max. Body Height (mm;\nMean-Centred)"
) +
  theme(legend.position = "none")

p2 <- massHeightModel$data %>%
  mutate("Residuals" = residuals(massHeightModel,
    type = "ordinary", robust = TRUE
  )[, "Estimate"]) %>%
  ggplot(aes(x = Residuals)) +
  geom_density(colour = "black", fill = "lightblue", alpha = 0.5) +
  xlab("Max. Body Height\nResiduals (mm)") +
  ylab("Density") +
  theme_classic()

p3 <- massHeightModel$data %>%
  mutate("Residuals" = residuals(massHeightModel,
    type = "ordinary", robust = TRUE
  )[, "Estimate"]) %>%
  ggplot(aes(x = mass, y = Residuals)) +
  geom_point(
    size = 2, pch = 21, colour = "black",
    fill = "lightblue", alpha = 0.5
  ) +
  xlab("Body Mass (g)") +
  ylab("Max. Body Height\nResiduals (mm)") +
  theme_classic()

upperLimit <- with(
  structuralData,
  diff(range(synsacrumWidthMean, na.rm = T)) /
    diff(range(massMean, na.rm = T))
)

massSynsacrumModel <- brm(
  data = structuralData %>%
    select(
```

```
      "synsacrumWidth" = synsacrumWidthMean,
      "mass" = massMean
    ) %>%
    mutate(
      "synsacrumWidth" = synsacrumWidth -
        mean(synsacrumWidth, na.rm = T),
      "mass" = mass - mean(mass, na.rm = T)
    ),
  family = "gaussian",
  synsacrumWidth ~ mass,
  prior = c(
    set_prior("normal(0, 2)", class = "Intercept"),
    set_prior(paste0(
      "normal(0, ",
      upperLimit / 2, ")"
    ), class = "b"),
    set_prior("exponential(1)", class = "sigma")
  ),
  iter = 50000, warmup = 10000, cores = 4,
  chains = 4, thin = 20,
  silent = TRUE, refresh = 0,
  file = "./models/massSynsacrumModel.Rds"
)

p4 <- pp_check2(massSynsacrumModel,
  xlab = "Max. Synsacrum Width\n(mm; Mean-Centred)"
) +
  theme(legend.position = "none")

p5 <- massSynsacrumModel$data %>%
  mutate(
    "Residuals" =
      residuals(massSynsacrumModel, type = "ordinary",
                robust = TRUE)[, "Estimate"]
  ) %>%
  ggplot(aes(x = Residuals)) +
  geom_density(colour = "black", fill = "lightblue", alpha = 0.5) +
  xlab("Max. Synsacrum Width\nResiduals (mm)") +
  ylab("Density") +
  theme_classic()

p6 <- massSynsacrumModel$data %>%
  mutate(
    "Residuals" =
      residuals(massSynsacrumModel, type = "ordinary",
                robust = TRUE)[, "Estimate"]
  ) %>%
  ggplot(aes(x = mass, y = Residuals)) +
  geom_point(
    size = 2, pch = 21, colour = "black",
    fill = "lightblue", alpha = 0.5
  ) +
  xlab("Body Mass (g)") +
  ylab("Max. Synsacrum Width\nResiduals (mm)") +
  theme_classic()

(p1 + p2 + p3)/
  (p4 + p5 + p6) +
  plot_annotation(tag_levels = "A")
```

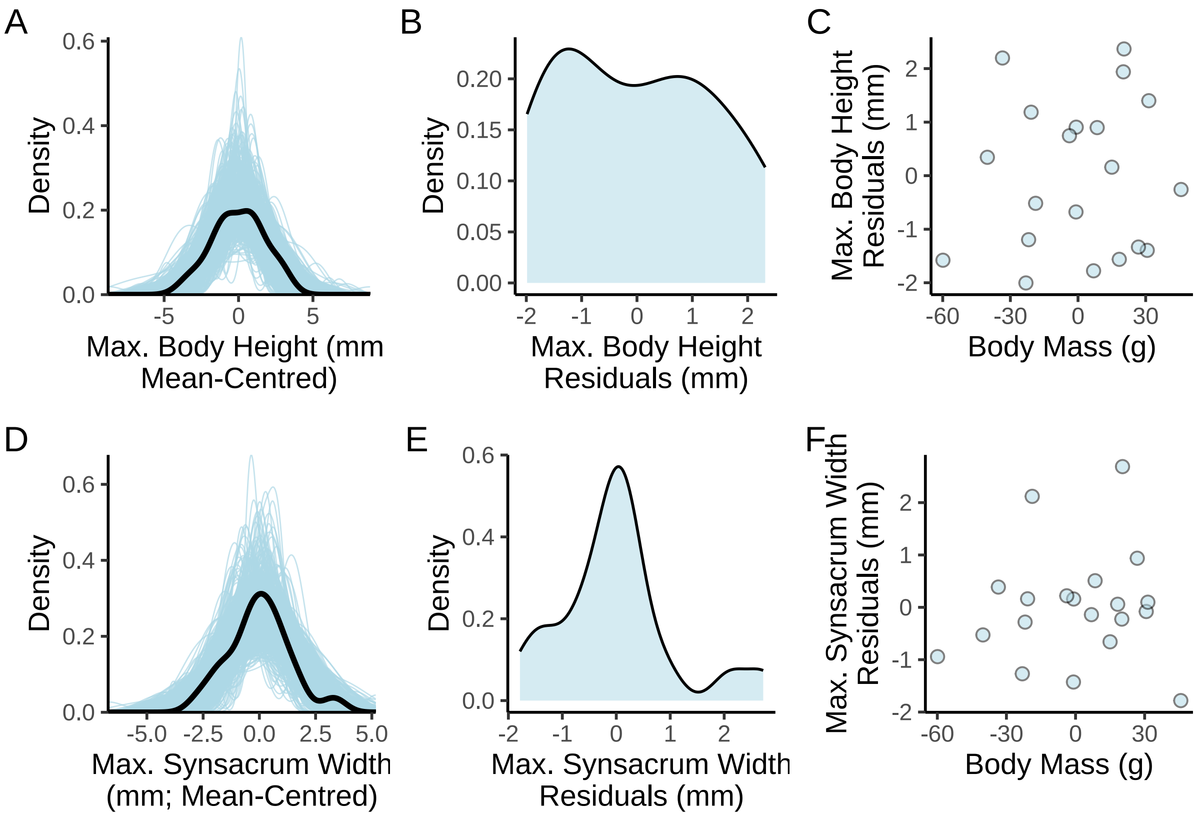


***Figure 4:*** *Validations for linear models with body mass (g) predicting maximum body height (mm) and maximum synsacrum width (mm) in twenty mature and deceased Japanese quail. Maximum body height represents the maximum vertial distance between the keel and back of feathered quail. Maximum synsacrum width represents that measured after dissection. Panels A and D represent posterior predictive checks, with black lines indicating true density of response variable values and light blue lines indicating estimated densities, as drawn from model posteriors. All other panels display spread (density in panels B and E) of model residuals.*

```
# Again, collating results

mass <- data.frame(
  "massHeight" = as.data.frame(massHeightModel)$b_mass,
  "massSynsacrum" = as.data.frame(massSynsacrumModel)$b_mass
) %>%
  summarise_all(., .funs = median) %>%
  pivot_longer(everything(),
    names_to = "Parameter",
    values_to = "Estimate"
  ) %>%
  merge(.,
    bind_rows(
      quantileCIs(massHeightModel, cis = c(50, 95)) %>%
        mutate(Parameter = ifelse(Parameter == "b_mass", "massHeight", Parameter)),
      quantileCIs(massSynsacrumModel, cis = c(50, 95)) %>%
        mutate(Parameter = ifelse(Parameter == "b_mass", "massSynsacrum", Parameter))
    ),
    by = "Parameter", all.x = TRUE
  ) %>%
  rowwise() %>%
  mutate("BF" = ifelse(Parameter == "massHeight",
    ifelse(Estimate < 0,
      (2 * mean(as.data.frame(
        massHeightModel
      )[, "b_mass"] <= 0)) /
```

```
        (2 * mean(as.data.frame(
          massHeightModel
        )[, "b_mass"] >= 0)),
      (2 * mean(as.data.frame(
        massHeightModel
      )[, "b_mass"] >= 0)) /
        (2 * mean(as.data.frame(
          massHeightModel
        )[, "b_mass"] <= 0))
    ),
    ifelse(Estimate < 0,
      (2 * mean(as.data.frame(
        massSynsacrumModel
      )[, "b_mass"] <= 0)) /
        (2 * mean(as.data.frame(
          massSynsacrumModel
        )[, "b_mass"] >= 0)),
      (2 * mean(as.data.frame(
        massSynsacrumModel
      )[, "b_mass"] >= 0)) /
        (2 * mean(as.data.frame(
          massSynsacrumModel
        )[, "b_mass"] <= 0))
    )
  )) %>%
  ungroup() %>%
  mutate(
    "Response" = c("Body Height (mm)", "Synsacrum Width (mm)"),
    "Predictor" = "Body Mass (g)",
    "Estimate" = round(Estimate, digits = 4),
    "BF" = round(BF, digits = 4),
    `50\\% CIs` = paste0(
      "[", round(Low_CI_50, digits = 4),
      ", ", round(High_CI_50, digits = 4),
      "]"
    ),
    `95\\% CIs` = paste0(
      "[", round(Low_CI_95, digits = 4),
      ", ", round(High_CI_95, digits = 4),
      "]"
    ),
    "N" = 20
  ) %>%
  select(
    Predictor, Response, N, Estimate,
    `50\\% CIs`, `95\\% CIs`, BF
  )

caption <- paste0(
  "Correlations between tarsus length (mm) ",
  "or body mass (g) and two measures of skeletal size ",
  "(maximum body height [mm] and maximum synsacrum width [mm]) ",
  "in twenty mature and deceased Japanese quail. ",
  "Results are derived from Bayesian linear models. ",
  "'CIs' indicate highest posterior density ",
  "credible intervals, and 'BFs' indicate Bayes factors."
)

structuralSizeResults <- rbind(tarsus, mass) %>%
  kbl(format = "latex", escape = FALSE, caption = caption) %>%
  column_spec(column = c(1:2), width = "2.1cm") %>%
  column_spec(column = c(3:10), width = "1.9cm") %>%
  kable_styling(latex_options = "striped")

structuralSizeResults
```

```
# save_kable(structuralSizeResults,
# "../tables/structuralSizePredictionResults.html")
```

***Table 2:*** *Correlations between tarsus length (mm) or body mass (g) and two measures of skeletal size (maximum body height [mm] and maximum synsacrum width [mm]) in twenty mature and deceased Japanese quail. Results are derived from Bayesian linear models. 'CIs' indicate highest posterior density credible intervals, and 'BFs' indicate Bayes factors.*

| Predictor | Response | N | Estimate | 50% CIs | 95% CIs | BF |
|---|---|---|---|---|---|---|
| Tarsus Length (mm) | Body Height (mm) | 20 | 0.0804 | [-0.0668, 0.2257] | [-0.3627, 0.5242] | 1.8369 |
| Tarsus Length (mm) | Synsacrum Width (mm) | 20 | 0.0522 | [-0.0732, 0.1735] | [-0.3193, 0.4153] | 1.5690 |
| Body Mass (g) | Body Height (mm) | 20 | 0.0320 | [0.0241, 0.0396] | [0.0087, 0.054] | 194.1220 |
| Body Mass (g) | Synsacrum Width (mm) | 20 | 0.0291 | [0.0229, 0.0348] | [0.01, 0.0464] | 499.0000 |

```
# Body mass, but not tarsus length, is
# a good predictor of structural size in our quail.
```

#### Validating accuracy of digital tarsus length measurements

Next, we confirm precision of our digital tarsus length measurements (mm) by comparing them against analogue measures of tarsus length (mm) in a subset of mature Japanese quail (n = 43). Comparisons are first done visually, then statistically by a simple Bayesian, measurement error model with analogue tarsus length as the Gaussian-distributed response and digital tarsus length as the sole population-level predictor with known measurement error as follows:

$$Analogue\ Tarsus\ Length_i \sim \beta_0 + \beta_1 * Digital\ Tarsus\ Length_{i*} + \epsilon_i$$

$$Digital\ Tarsus\ Length_i \sim Digital\ Tarsus\ Length_i + \eta_i$$

where $Digital\ Tarsus\ Length_{i*}$ represents the true but unknown digital tarsus length measurement, $Digital\ Tarsus\ Length_i$ mean, digital tarsus length measurement for an individual $i$, and $\eta_i$ represents the standard deviation of digital tarsus length measurements for individual $i$.

Priors for this regression were:

$$\beta_0 \sim \mathcal{N}(0, 5)$$

$$\beta_1 \sim S\mathcal{N}(0.25, 1, 3)$$

$$\epsilon i \sim exponential(1)$$

with $\xi = 0.25$ being chosen for our digital tarsus length prior to place its distributional mean at approximately 1.

Below, we begin by displaying a representative image used for tarsus length calculation. Data are then subsequently loaded in and plotted.

```
plotJpeg <- function(path, add=FALSE) {
  image = jpeg::readJPEG(path, native=T)
  res = dim(image)[2:1]
  if (!add)
    plot(1, 1, xlim=c(1,res[1]),
    ylim=c(1,res[2]), asp=1,type='n',
    xaxs='i', yaxs='i',
    xaxt='n',yaxt='n',
    xlab='',ylab='',
```

```
    bty='n'
    )
  rasterImage(image, 1, 1, res[1], res[2])
}

plotJpeg("tarsusImage.jpg")
```

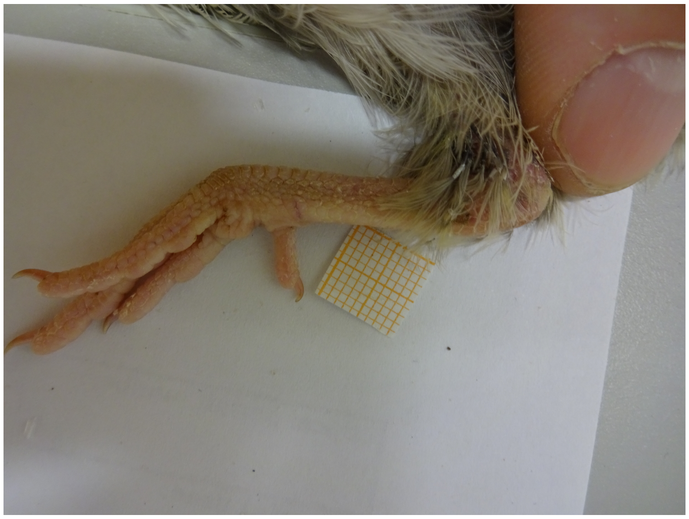

***Figure 5:*** *Photo of a mature (8 weeks) Japaense quail tarsus with 1 mm x 1 mm grid paper used for digital tarsus length measurement.*

```
# Loading in data and correcting typo in header

comparisonData <-
  merge(read.csv("quailAnalogueMeasures20Weeks.csv"),
    read.csv("quailDigitalMeasurements20Weeks.csv"),
    by = "ring", all = TRUE
  ) %>%
  rename("tarsusLengthMean" = tarsusLengthhMean) %>%
  select(ring, tarsusLength, tarsusLengthMean,
         tarsusLengthSD, tarsusLengthN) %>%
  mutate(tarsusLengthSD = ifelse(is.na(tarsusLengthSD),
                                 mean(tarsusLengthSD, na.rm = T),
                                 tarsusLengthSD)
  ) %>%
  drop_na()

# Plotting digital measures of tarsus and bill length against analogue measures

comparisonData %>%
  ggplot(aes(x = tarsusLengthMean, y = tarsusLength)) +
  geom_errorbarh(
    aes(
      xmin = tarsusLengthMean - tarsusLengthSD /
        sqrt(tarsusLengthN),
      xmax = tarsusLengthMean + tarsusLengthSD /
        sqrt(tarsusLengthN),
      y = tarsusLength
    ),
    colour = "black", height = 0.2
  ) +
  geom_point(size = 2.5, colour = "black", pch = 21,
             fill = "#90B1DB", alpha = 0.7) +
  theme_classic() +
  xlab("Digital Tarsus Length\nMeasurement (mm)") +
  ylab("Analogue Tarsus Length\nMeasurement (mm)")
```

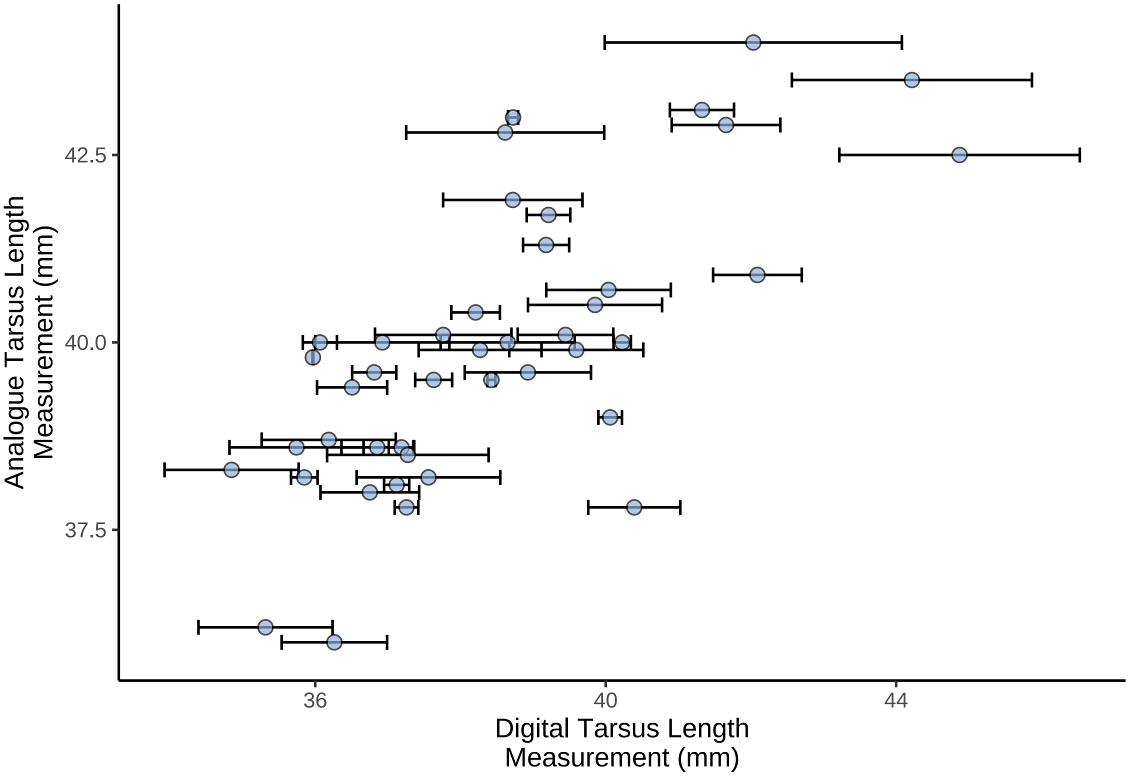


***Figure 6:*** *Analogue tarsus length measurements as a function of digital tarsus length measurements from 43 mature Japanese quail. Digital tarsus length was measured more than once per individual (range of measurements = 2-4) and dots represent means per individual. Horizontal errorbars represent +/- one standard error around means.*

```
# Modelling

tarsusModel <- brm(
  data = comparisonData %>%
    mutate(
      tarsusLength = tarsusLength -
        mean(tarsusLength, na.rm = T),
      tarsusLengthMean = tarsusLengthMean -
        mean(tarsusLengthMean, na.rm = T)
    ),
  family = "gaussian",
  tarsusLength ~ me(tarsusLengthMean, tarsusLengthSD),
  prior = c(
    set_prior("normal(0, 5)", class = "Intercept"),
    set_prior("skew_normal(0.25, 1, 3)", class = "b"),
    set_prior("exponential(1)", class = "sigma")
  ),
  iter = 50000, warmup = 10000, cores = 4, chains = 4, thin = 20,
  silent = TRUE, refresh = 0,
  file = "./models/digitalTarsusValidation.Rds"
)
```

To ensure that our priors are not restrictive, we check the sensitivity of our model likelihood to power-scaling of our prior (Kallioninen et al, 2024).

```
powerscale_sensitivity(tarsusModel)
```

```
## Sensitivity based on cjs_dist:
## # A tibble: 6 x 4
##   variable                              prior likelihood diagnosis
##   <chr>                                 <dbl>      <dbl> <chr>
## 1 b_Intercept                         0.00378     0.193  -
## 2 bsp_metarsusLengthMeantarsusLengthSD 0.00819     0.968  -
## 3 sigma                               0.0322      0.546  -
## 4 Intercept                           0.00378     0.193  -
## 5 meanme_metarsusLengthMean           0.00118     0.0547 -
## 6 sdme_metarsusLengthMean             0.00373     0.228  -
```

We find no evidence that our priors are restrictive and therefore proceed to simple model validations.

```
# Checking fit and residuals.

p1 <- pp_check2(tarsusModel,
  xlab = "Analogue Tarsus Length (mm)"
) + theme(legend.position = "none")

p2 <- tarsusModel$data %>%
  mutate(
    "Fit" = fitted(tarsusModel)[, "Estimate"],
    "SE" = fitted(tarsusModel)[, "Est.Error"]
  ) %>%
  ggplot(aes(x = Fit, y = tarsusLength)) +
  geom_errorbarh(aes(xmin = Fit - SE, xmax = Fit + SE)) +
  geom_point(
    pch = 21, colour = "black",
    fill = "lightblue", size = 2, alpha = 0.8
  ) +
  xlab("Fitted Tarsus\nLength(mm)") +
  ylab("Tarsus Length\n(mm)") +
  theme_classic()

p3 <- ggplot(
  data =
    data.frame("X" = c(brms::bayes_R2(tarsusModel, summary = FALSE))),
  aes(x = X)
) +
  geom_density(
    colour = "black",
    fill = "lightblue", alpha = 0.7
  ) +
  xlab(
    TeX("$R^{2}$")
  ) +
  ylab("Density") +
  theme_classic()

p4 <- tarsusModel$data %>%
  mutate("Residuals" = residuals(tarsusModel,
    robust = TRUE
  )[, "Estimate"]) %>%
  ggplot(aes(sample = Residuals)) +
  stat_qq(colour = "lightblue") +
  stat_qq_line() +
  xlab("Theoretical Residual\nQuantiles") +
  ylab("Sample Residual\nQuantiles") +
  theme_classic()

p5 <- tarsusModel$data %>%
  mutate("Residuals" = residuals(tarsusModel,
    robust = TRUE
  )[, "Estimate"]) %>%
  ggplot(aes(x = tarsusLengthMean, y = Residuals)) +
```

```
  geom_point(
    size = 2, pch = 21,
    colour = "black", fill = "lightblue", alpha = 0.5
  ) +
  xlab("Digital Tarsus Length\nMeasurement (mm)") +
  ylab("Ordinary Residuals") +
  theme_classic()

(p1 + p2) / (p3 + p4 + p5) +
  plot_annotation(tag_levels = "A")
```

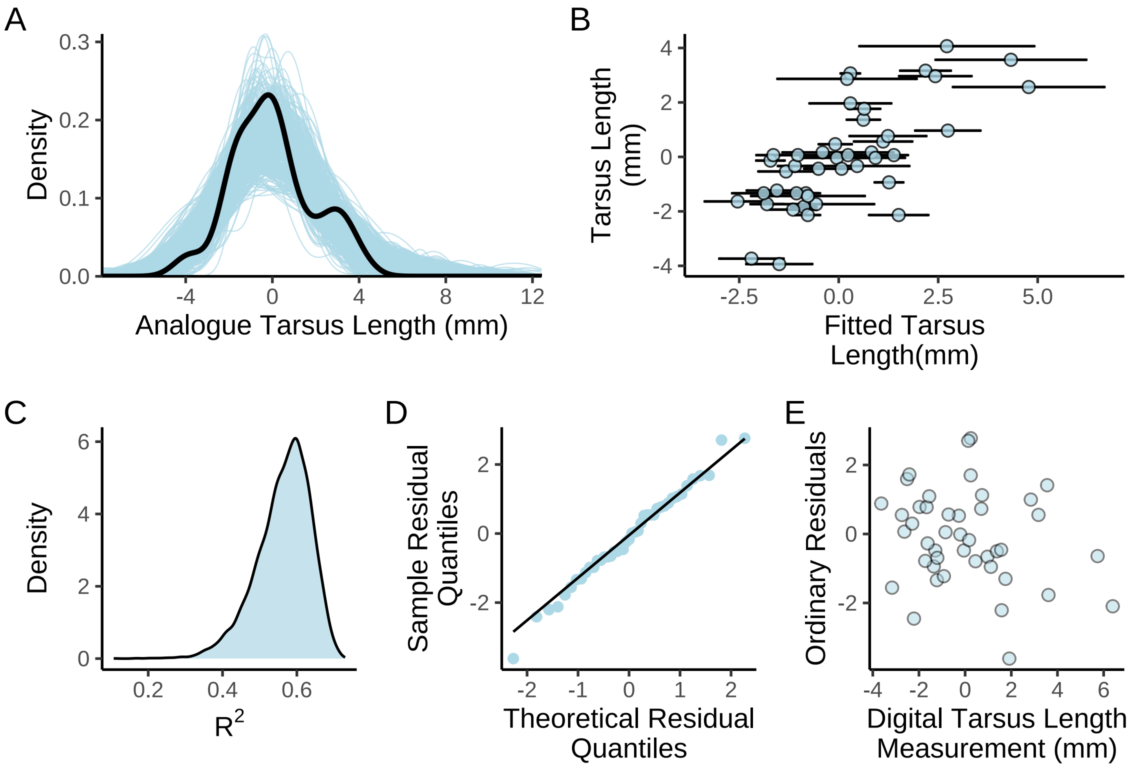


***Figure 7:*** *Spread of fitted and residual values for a simple Bayesian regression predicting anoalogue tarsus length measurements as a function of digital tarsus length measurements. Panel A displays a posterior predictive check, with the density of true analogue tarsus length measurements indicated by a black line, and estimated densities (drawn from model posteriors) indicated by blue lines. Panel B displays raw analogue tarsus lengths measurements as a function of their fitted values (blue dots) +/- one standard error around fitted values. Panel C displays the esitmated* $R^2$ *density. Panels D and E display the distribution oresiduals from regression.*

```
# Summarising model.

caption <- paste0(
  "Results from a Bayesian linear model ",
  "predicting analogue tarsus length measurements ",
  "in mature Japanese quail. 'BF' indicates Bayes ",
  "Factor and R\\textsuperscript{2} indicates ",
  "a Bayesian R\\textsuperscript{2}."
)

digitalAnalogueTable <- as.data.frame(tarsusModel) %>%
  select(bsp_metarsusLengthMeantarsusLengthSD) %>%
  summarise_all(., .funs = mean) %>%
  pivot_longer(everything(),
```

***Table 3:*** *Results from a Bayesian linear model predicting analogue tarsus length measurements in mature Japanese quail. 'BF' indicates Bayes Factor and $R^2$ indicates a Bayesian $R^2$.*

| Predictor | N | Estimate | 50% CIs | 95% CIs | BF | $R^2$ |
|---|---|---|---|---|---|---|
| Digital Tarsus Length (mm) | 43 | 0.7279 | [0.6479, 0.8064] | [0.4981, 0.9745] | Inf | 0.5626 |

```
    names_to = "Parameter",
    values_to = "Mean"
  ) %>%
  merge(., quantileCIs(tarsusModel, cis = c(50, 95)),
    by = "Parameter", all.x = TRUE
  ) %>%
  mutate("BF" = ifelse(Mean < 0,
    (2 * mean(as.data.frame(
      tarsusModel
    )[, Parameter] <= 0)) /
      (2 * mean(as.data.frame(
        tarsusModel
      )[, Parameter] >= 0)),
    (2 * mean(as.data.frame(
      tarsusModel
    )[, Parameter] >= 0)) /
      (2 * mean(as.data.frame(
        tarsusModel
      )[, Parameter] <= 0))
  )) %>%
   mutate(
    "Predictor" = "Digital Tarsus\nLength (mm)",
    "Estimate" = round(Mean, digits = 4),
    "BF" = round(BF, digits = 4),
    `50\\% CIs` = paste0(
      "[", round(Low_CI_50, digits = 4),
      ", ", round(High_CI_50, digits = 4),
      "]"
    ),
    `95\\% CIs` = paste0(
      "[", round(Low_CI_95, digits = 4),
      ", ", round(High_CI_95, digits = 4),
      "]"
    ),
     `R\\textsuperscript{2}` =
      round(brms::bayes_R2(tarsusModel)[, "Estimate"], digits = 4),
    "N" = nrow(tarsusModel$data)
  ) %>%
  select(Predictor, N, Estimate,
         `50\\% CIs`, `95\\% CIs`,
         BF, `R\\textsuperscript{2}`
  ) %>%
  kbl(., format = "latex", escape = FALSE, caption = caption) %>%
  column_spec(column = c(1:10), width = "2cm") %>%
  kable_styling(latex_options = "striped")

digitalAnalogueTable
```
