## Supplementary material for "No evidence that shrinking and shapeshifting meaningfully affect how birds respond to warming and cooling": Supp. 2

### Data Organisation and Analysis of Growth

#### Overview

In this document, we compile morphological data from Japanese quail (n = 108) reared across three distinct thermal treatments: (1) "cold" (10°C for 3-8 weeks of age, followed by a constant 20°C), (2) "mild" (20°C until 8 weeks of age), and (3) "warm" (30°C for 3-8 weeks of life, followed by a constant 20°C). Morphological measurements taken include body mass (g) and tarsus length (mm), with mass measured weekly from 0-8 weeks of age, and tarsus length measured weekly between 0-3 weeks of age, then again at 8 weeks of age. A full description of measurement methodology is provided in the 'Materials and Methods' of Tabh et al (2024). Once compiled, we proceed by visualising these data for oddities or erroneous values. Then, to test for an effect of rearing temperature treatment on patterns of growth and adulthood phenotypes (here, 8 weeks of age), we follow by building analytical models, scrutinising these models, and visualising outcomes.

##### Data compilation

Below, we begin by importing all packages and functions required for data compilation, visualisation, and analysis. Versions of R packages loaded into our R session are printed to facilitate reproduction. We recommend that those seeking to reproduce our models and results use the versions reported before proceeding.

```
def.chunk.hook <- knitr::knit_hooks$get("chunk")
knitr::knit_hooks$set(chunk = function(x, options) {
  x <- def.chunk.hook(x, options)
  ifelse(options$size != "normalsize",
         paste0("\n \\", options$size, "\n\n", x, "\n\n \\normalsize"), x)
})

knitr::opts_chunk$set(fig.pos = "H", out.extra = "")
knitr::opts_chunk$set(size = "footnotesize")

# Much of the below data organisation relies on the tidyverse
# language set. Tidyverse is imported first to avoid
# interference between its functions and those of 'brms'.

library("tidyverse")

# 'easypackages' is used to load all other packages in with one simple execution.

library("easypackages")

packageList <- c("bayesplot", "brms", "doParallel",
                 "foreach", "ggpubr", "kableExtra",
                 "latex2exp", "patchwork", "priorsense",
                 "showtext", "tidybayes", "wesanderson")
```

```
libraries(packageList)

# R methods from the 'brms' packages that allow for the
# addition of priors in hypothesis tests are loaded from
# a local source (available on GitHub). This first
# requires installation of the dependent package "mgsub".

#install.packages("mgsub")
#install.packages("/Users/joshuatabh/rPackageDevelopment/brmsMethods",
  #repos = NULL, type = "source")

library("brmsMethods")

# Printing package version numbers

caption <- paste0("R packages and their respective versions used for",
                  " data organisation and analysis in this study."
)

sapply(packageList, function(x) {
  y <- as.character(packageVersion(x))
  return(y)
}, simplify = FALSE) %>%
  enframe(., name = "Package", value = "Version") %>%
  as.data.frame(.) %>%
  kbl(.,
    longtable = T, booktabs = T,
    caption = caption
  ) %>%
  kable_styling(latex_options = "striped")
```

***Table 1:*** *R packages and their respective versions used for data organisation and analysis in this study.*

| Package | Version |
|---|---|
| bayesplot | 1.11.1 |
| brms | 2.20.4 |
| doParallel | 1.0.17 |
| foreach | 1.5.2 |
| ggpubr | 0.6.0 |
| kableExtra | 1.3.4 |
| latex2exp | 0.9.6 |
| patchwork | 1.2.0 |
| priorsense | 0.0.0.9000 |
| showtext | 0.9.6 |
| tidybayes | 3.0.4 |
| wesanderson | 0.3.6.9000 |

```
# Adding custom functions

## A function to calculate the mode of a vector
```

```
md <- function(x) {
  all_values <- unique(x)
  all_values[which.max(tabulate(match(x, all_values)))]
}

# A function to cleanly view autocorrelation between posterior
# draws of specified coefficients/variables

clean_ac <- function(x, prs = NA, names = NA) {
  require("rstan")

  if (class(x)[1] != "brmsfit") {
    return("x must be a brmsfit object.")
  }

  if (is.na(prs[1])) {
    return(stan_ac(x$fit))
  }

  if (!is.na(prs[1]) & is.na(names[1])) {
    return(stan_ac(x$fit, pars = prs))
  }

  if (length(prs) != length(names)) {
    return("Length of pars and names must be equal.")
  }

  Base_plot <- stan_ac(x$fit, pars = prs, fill = nice_pink)
  Base_plot$data$parameters <- as.character(Base_plot$data$parameters)

  for (i in 1:length(prs)) {
    Base_plot$data$parameters[c(which(Base_plot$data$parameters == prs[i]))] <-
      names[i]
  }
  Base_plot$data$parameters <- as.factor(Base_plot$data$parameters)

  return(Base_plot)
}

# A function to simplify the output of bayestestR's hdi function.

simple_hdi <- function(x, rnd = 3, cis = c(50, 95), sci_note = FALSE) {
  if (class(x)[1] != "brmsfit") {
    return("x must be a brmsfit object.")
  }
  if (length(cis) != 2) {
    return("cis must be a vector of integers with length 2")
  }

  HDI_low <- bayestestR::hdi(x, effects = "all", ci = min(cis) / 100)
  HDI_high <- bayestestR::hdi(x, effects = "all", ci = max(cis) / 100)

  if (sci_note == FALSE) {
```

```
    Results <- data.frame(
      "Parameter" = HDI_low$Parameter,
      "1" = round(HDI_low$CI_low, rnd),
      "2" = round(HDI_low$CI_high, rnd),
      "3" = round(HDI_high$CI_low, rnd),
      "4" = round(HDI_high$CI_high, rnd)
    )
    colnames(Results)[c(2:5)] <-
      c(paste0("Low_HDI_", min(cis)), paste0("High_HDI_", min(cis)),
        paste0("Low_HDI_", max(cis)), paste0("High_HDI_", max(cis)))
  } else if (sci_note == TRUE) {
    Results <- data.frame(
      "Parameter" = HDI_low$Parameter,
      "1" = format(round(HDI_low$CI_low, rnd), scientific = TRUE),
      "2" = format(round(HDI_low$CI_high, rnd), scientific = TRUE),
      "3" = format(round(HDI_high$CI_low, rnd), scientific = TRUE),
      "4" = format(round(HDI_high$CI_high, rnd), scientific = TRUE)
    )
    colnames(Results)[c(2:5)] <- c(paste0("Low_HDI_", min(cis)),
                                   paste0("High_HDI_", min(cis)),
                                   paste0("Low_HDI_", max(cis)),
                                   paste0("High_HDI_", max(cis)))
  }

  return(Results)
}

modeHDI <- function(x, cis = c(50, 95), rnd = 4, collapse = FALSE){
  stopifnot("x must be a 'brmsfit' object" = class(x) == "brmsfit",
            "collapse must be logical TRUE/FALSE" = is.logical(collapse))

  out <- lapply(X = as.data.frame(x), MARGIN = 2, FUN = ggdist::mode_hdi,
                .width = c(cis/100))
  hold <- names(out)
  out <- out %>%
    map2(hold, ~mutate(.x, name = .y)) %>%
    bind_rows() %>%
    mutate(y = round(y, digits = rnd),
           ymin = round(ymin, digits = rnd),
           ymax = round(ymax, digits = rnd)) %>%
    select("par" = name, "mode" = y, "lcl" = ymin,
           "ucl"= ymax, "confidenceLevel" = .width)

  if (collapse == FALSE){
    return(out)
  } else if (collapse == TRUE){
    out <- out %>%
      mutate("cis" = paste0("[", lcl, ", ", ucl, "]")) %>%
      select(-c(lcl, ucl))

    return(out)
  }
}
```

```r
# A function to calculate quantile intervals from a brmsfit object

quantileCIs <- function(x, rnd = 3, cis = c(50, 95), sci_note = FALSE) {
  require(tidyverse)

  if (class(x)[1] != "brmsfit") {
    return("x must be a brmsfit object.")
  }
  if (length(cis) != 2) {
    return("cis must be a vector of integers with length 2")
  }

  prbs = c()
  nColNames = c()
  for (i in 1:length(cis)){
    prbs = c(prbs, c(0.5 - (cis[i]/100)/2, 0.5 + (cis[i]/100)/2))
    nColNames = c(nColNames,
                  paste0("Low_CI_", cis[i]),
                  paste0("High_CI_", cis[i])
    )
  }

  modelFrame = as.data.frame(x)

  Results <- apply(modelFrame, MARGIN = 2, FUN = quantile,
        probs = prbs, type = 8) %>%
    t() %>%
    as.data.frame() %>%
    rownames_to_column(var = "par") %>%
    `colnames<-`(c("Parameter", nColNames))

  if (sci_note == FALSE) {
    Results <- apply(modelFrame, MARGIN = 2, FUN = quantile,
        probs = prbs, type = 8) %>%
    t() %>%
    as.data.frame() %>%
    rownames_to_column(var = "par") %>%
    `colnames<-`(c("Parameter", nColNames))

  } else if (sci_note == TRUE) {
    Results <- apply(modelFrame, MARGIN = 2, FUN = quantile,
        probs = prbs, type = 8) %>%
    t() %>%
    as.data.frame() %>%
    rownames_to_column(var = "par") %>%
    `colnames<-`(c("Parameter", nColNames)) %>%
    mutate_at(.vars = vars(-Parameter),
              .funs = function(x){
                return(format(x, scientific = TRUE))
              }
    )
  }
```

```
  return(Results)
}

# A function that allows users to assign multiple objects
# to different variable names at once. The below is reported by "ellbur" at
# https://strugglingthroughproblems.wordpress.com/author/ellbur/page/3/.

{
  "%=%" <- function(l, r, ...) UseMethod("%=%")

  "%=%.lbunch" <- function(l, r, ..., List = NA) {
    Envir <- as.environment(-1)

    if (!is.na(List)) {
      l <- List[[1]]
      r <- List[[2]]
    }

    if (length(r) > length(l)) {
      warning("RHS has more args than LHS. Only first",
              length(l), "used.")
    }

    if (length(l) > length(r)) {
      warning("LHS has more args than RHS. RHS will be repeated.")
      r <- extendToMatch(r, l)
    }

    for (II in 1:length(l)) {
      do.call("<-", list(l[[II]], r[[II]]), envir = Envir)
    }
  }

  extendToMatch <- function(source, destin) {
    s <- length(source)
    d <- length(destin)

    if (d == 1 && s > 1 && !is.null(as.numeric(destin))) {
      d <- destin
    }

    dif <- d - s
    if (dif > 0) {
      source <- rep(source, ceiling(d / s))[1:d]
    }
    return(source)
  }

  g <- function(...) {
    List <- as.list(substitute(list(...)))[-1L]
    class(List) <- "lbunch"
    return(List)
  }
```

```r
}

# A function to simplify output of bayes_R2 from brms

simpleR2 <- function(x, ndraws = 1000, roundDigits = 5,
                     robust = TRUE) {
  stopifnot("x must be a 'brmsfit' object" = class(x) == "brmsfit",
            "robust must be logical (TRUE/FALSE)" = is.logical(robust))
  grab <- brms::bayes_R2(x, ndraws = ndraws, robust = robust)
  toPrint <- paste0(
    "R2 = ", round(grab[, "Estimate"], digits = roundDigits),
    " [",
    round(grab[, "Q2.5"], digits = roundDigits),
    ",",
    round(grab[, "Q97.5"], digits = roundDigits),
    "]"
  )
  cat(toPrint)
}

# A function to calculate the position of a skew-normal
# distribution given its mean, omega, and alpha values

skewxi <- function(mean, omega, alpha) {
  delta <- alpha / (sqrt(1 + alpha^2))
  xi <- mean - omega * delta * sqrt(2 / pi)
  return(xi)
}

# A function to produce clean posterior or prior
# predictive checks, based upon "pp_check" from the R-package 'brms'.

pp_check2 <- function(model, resp = NA, ndraws = 500,
                      xlab = "label", colour = "lightblue") {
  require(brms)
  require(ggplot2)
  stopifnot("Model must be a brmsfit object" = is.brmsfit(model))

  if (is.na(resp)) {
    resp <- model$formula$resp
  }

  p1 <- brms::pp_check(model, ndraws = ndraws, resp = resp) +
    scale_colour_manual(
      values = c("black", colour),
      labels = c("y", "yhat"),
      name = NULL
    ) +
    xlab(xlab) +
    ylab("Density") +
    theme_classic()
  return(p1)
}
```

```
# A function to summarise and print Gelman-Rubin statistics and effective
# sample sizes to sample sizes for brmsfit objects.

neffBase <- function(x){
  stopifnot("Model must be a brmsfit object" = is.brmsfit(x))
  out <- as.data.frame(brms::neff_ratio(x)) %>%
    rownames_to_column(var = "var") %>%
    filter(!(var %in% c("lprior", "lp__"))) %>%
    pull(.)
  return(out)
}

chainCheck <- function(model, rDig = 3) {
  require(brms)
  stopifnot("Model must be a brmsfit object" = is.brmsfit(model))

  Rhat <- paste0(
    "Rhat range: ",
    round(min(rhat(model)), digits = rDig),
    " - ",
    round(max(rhat(model)), digits = rDig)
  )
  Neff <- paste0(
    "Neff/N range: ",
    round(min(neffBase(model)), digits = rDig),
    " - ",
    round(max(neffBase(model)), digits = rDig)
  )
  cat(paste0(Rhat, "\n", Neff))
}

# Setting working directory. Note that this Rmarkdown
# file should be placed with the below defined folder
# for scripts to correctly identify other folder and
# file paths referenced in the remaining code.

setwd(
  paste0("/Users/joshuatabh/Documents/researchProjects/lund/",
  "functionOfAllen/functionData"
  )
)
```

Next, data are imported and bound.

```
data <- rbind(
  read.csv("exp1Morphology.csv") %>%
    mutate("exp" = "A"),
  read.csv("exp2Morphology.csv") %>%
    mutate("exp" = "B"),
  read.csv("exp3Morphology.csv") %>%
    mutate("exp" = "C")
) %>%
  select(-c(billCalibration, billLengthMean,
            billLengthSD, billDepthMean, billDepthSD)) %>%
  mutate(
```

```
    sex = tolower(sex),
    treatment = tolower(treatment),
    pretreatment = tolower(pretreatment),
    posttreatment = tolower(posttreatment)
  )
```

In this study, tarsus length measurements were derived from digital photographs and calibrated to standardised grids within view (see Materials and Methods of Tabh et al, 2024, for complete details). In some cases, calibration methods varied (e.g. calibration grid layed above or below the subject). To ensure that these variations did not bias our estimates of true appendage length, we first modelled appendage length (± its standard deviation when multiple photos were captured per individual) as a function of age (here, categorical) and calibration method, then used mean differences in length measures per calibration method (derived from our model) to correct raw measurement values accordingly. Before modelling, however, raw tarsus length measurements are plotted by age to check for erroneous values that may bias model convergence.

```
# Checking for aberrant, raw tarsus length measurements

data %>%
  filter(week < 10) %>%
  ggplot(aes(x = week, y = tarsusLengthMean)) +
  geom_point(size = 2, pch = 21, fill = "slateblue", colour = "black",
             position = position_jitter(width = 0.5), alpha = 0.3) +
  stat_summary(geom = "errorbar", fun.data = "mean_cl_boot",
               colour = "black", width = 0.3) +
  stat_summary(geom = "point", fun = "mean", size = 3, pch = 21,
               colour = "black", fill = "black") +
  theme_classic() +
  xlab("Age (weeks)") +
  ylab("Tarsus Length (mm)")
```

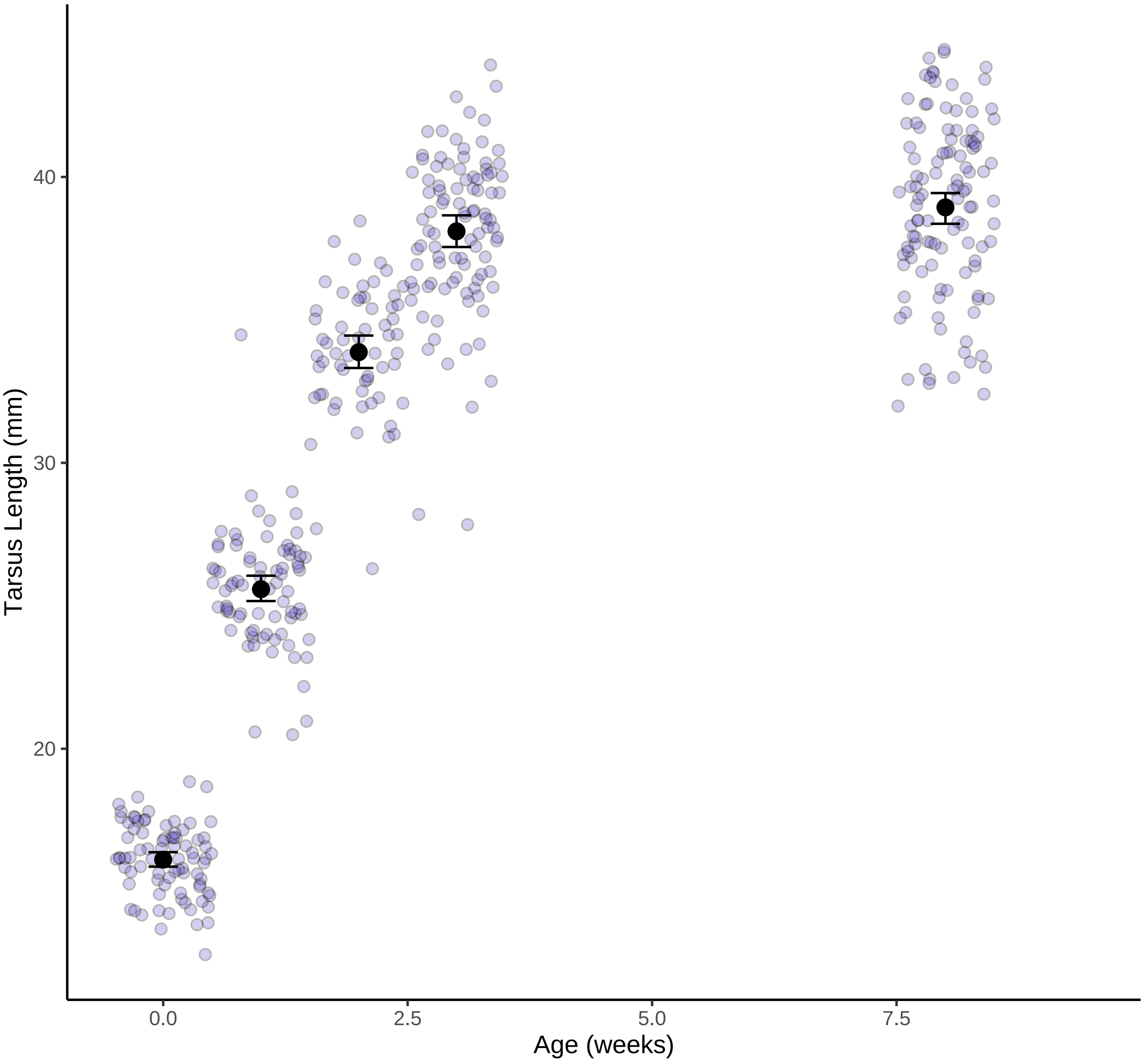


***Figure 1:*** *Effect of post-hatch age on tarsus length (mm) in Japanese quail. Black dots represent means and errorbars indicate standard errors; purple dots represent raw values.*

Our model is then constructed using uninformative, flat priors for population level slopes. For our intercept, a student-t distributed prior with three degrees of freedom, $\mu$ of 35.8, and and $\sigma$ of 6.9 is used.

```
# Adjusting for calibration method in tarsus length
# measurements then replotting. Note that this is achieved
# by modelling mean appendage length (mm; as derived from
# multiple digital photographs per individual) as a function
# of age (weeks; factorial) and calibration method (categorical).
# Uncertainty around mean appendage length is also accounted
# for within these models. All missing values for tarsus
# length measurements are first removed.

tarsusCorrection <- brm(tarsusLengthMean |
                          se(tarsusLengthSD, sigma = TRUE) ~
                          week + tarsusCalibration,
```

```r
  data = data %>%
    filter(!(is.na(tarsusLengthMean))) %>%
    mutate(
      week = factor(week),
      tarsusLengthSD = ifelse(is.na(tarsusLengthSD),
                              0.001,
                              tarsusLengthSD)
    ),
  iter = 50000, warmup = 5000, cores = 4, chains = 4,
  inits = 0,
  control = list(adapt_delta = .95),
  silent = TRUE, refresh = 0,
  file = "./models/tarsusCorrectionModel.Rds"
)

## Checking distribution of calibration coefficients,
# distribution of posterior predictions and leave-one-out "r2".

tarsusCorrection %>%
  as.data.frame() %>%
  mutate(b_Intercept = b_Intercept - mean(b_Intercept)) %>%
  select(
    "Grid over" = b_Intercept,
    "Grid under" = b_tarsusCalibrationgridUnder,
    "Ruler under" = b_tarsusCalibrationrulerUnder
  ) %>%
  pivot_longer(everything(),
    names_to = "method",
    values_to = "effect"
  ) %>%
  ggplot(aes(x = effect)) +
  facet_wrap(~method, scales = "free") +
  geom_density(colour = "black", fill = "grey50", alpha = 0.5) +
  theme_classic() +
  xlab("Mean Effect of Tarsus Length Estimate (mm)") +
  ylab("Density")
```

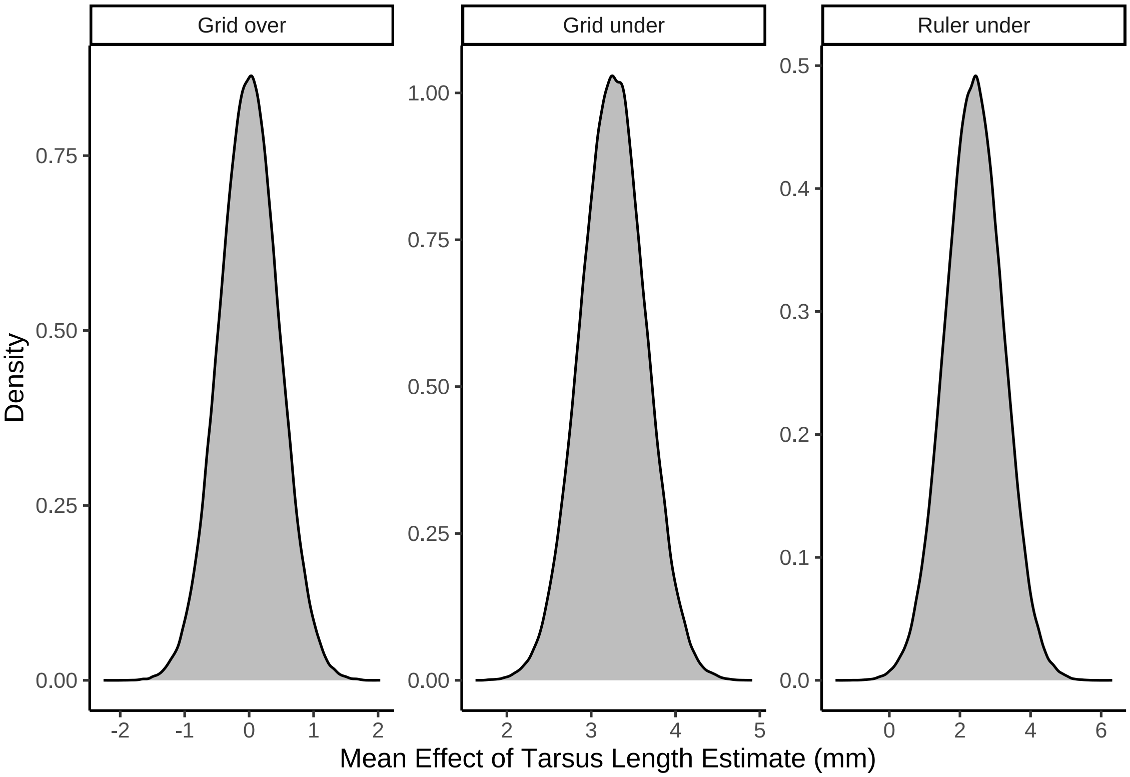


***Figure 2:*** *Effects (model coefficients) of measurement calibration method on tarsus length estimate in Japanese quail. Distributions represent posterior densities from a Bayesian, linear model with tarsus length (mm) as the response variable and age (weeks) and calibration method as predictors.*

Posteriors of our model are briefly visualised, a Bayesian $R^2$ estimated (here, using posterior median as a measure of centrality) and effects of calibration method displayed.

```
# Clear effects of calibration method.

pp_check2(tarsusCorrection, xlab = "Tarsus Length (mm)")
```

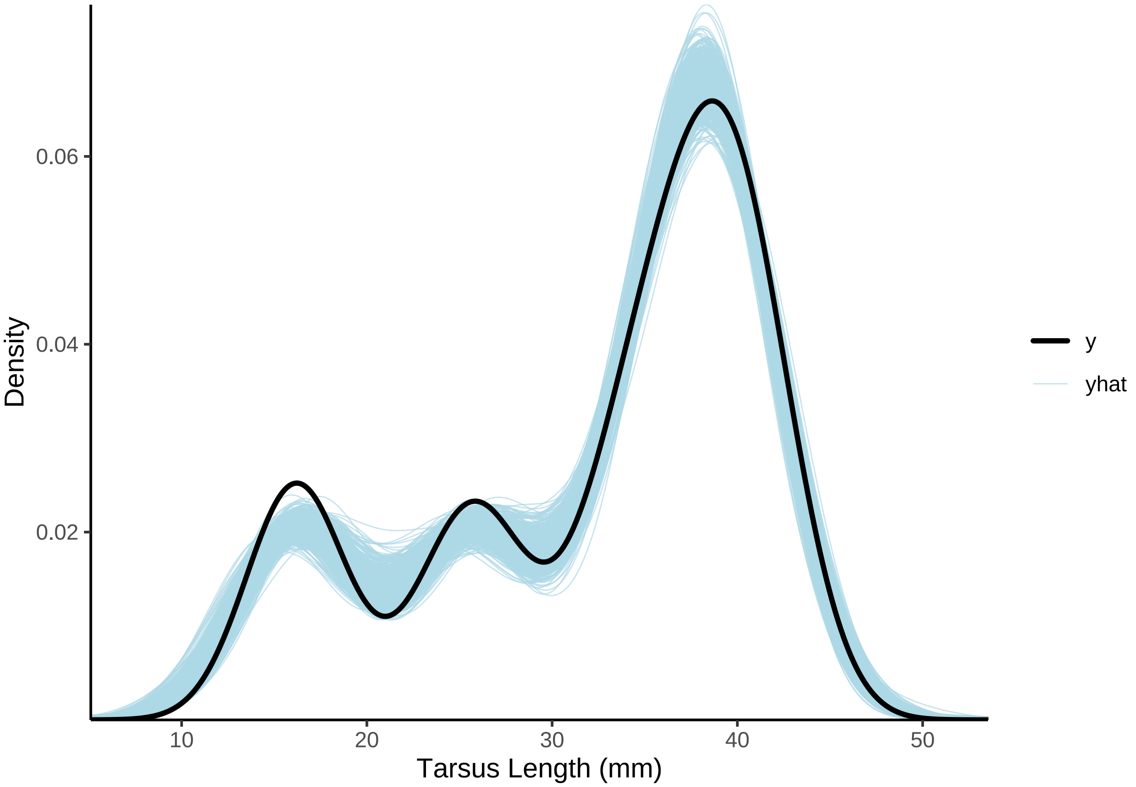


***Figure 3:*** *Density of posterior draws from a Bayesian linear mixed effects model predicting tarsus length in Japanese quail (mm; draws in grey). Density is overlayed with that of true tarsus length measurements (mm; line in black).*

```
simpleR2(tarsusCorrection, robust = TRUE)
```

```
## R2 = 0.93768 [0.9342,0.94011]
```

```
## Normal and remarkably high R2. Proceeding to calculate
# means, and errors around means, by quantiles

tarsusDeltas <- as.data.frame(tarsusCorrection) %>%
  summarise(
    "gridUnder" = mean(b_tarsusCalibrationgridUnder),
    "gridUnderLCL" = quantile(b_tarsusCalibrationgridUnder,
                              0.025, type = 8),
    "gridUnderUCL" = quantile(b_tarsusCalibrationgridUnder,
                              0.975, type = 8),
    "rulerUnder" = mean(b_tarsusCalibrationrulerUnder),
    "rulerUnderLCL" = quantile(b_tarsusCalibrationrulerUnder,
                               0.025, type = 8),
    "rulerUnderUCL" = quantile(b_tarsusCalibrationrulerUnder,
                               0.975, type = 8)
  )

caption <- paste0("Estimated effect of measurement calibration ",
                  "method on tarsus length estimates (mm; posterior means) ",
                  "across ages in Japanese quail. Credible invervals ",
                  "(2.5 and 97.5\\%) are in braces."
)

tarsusDeltas %>%
  mutate(
    "Grid Under" = paste0(
      round(gridUnder, digits = 5),
```

***Table 2:*** *Estimated effect of measurement calibration method on tarsus length estimates (mm; posterior means) across ages in Japanese quail. Credible invervals (2.5 and 97.5%) are in braces.*

| Calibration Method | Effect on tarsus length estimate (mm) |
|---|---|
| Grid Under | 3.2597 [2.50213,4.02127] |
| Ruler Under | 2.40562 [0.80056,4.00386] |

```
      " [",
      round(gridUnderLCL, digits = 5),
      ",",
      round(gridUnderUCL, digits = 5),
      "]"
    ),
    "Ruler Under" = paste0(
      round(rulerUnder, digits = 5),
      " [",
      round(rulerUnderLCL, digits = 5),
      ",",
      round(rulerUnderUCL, digits = 5),
      "]"
    )
  ) %>%
  select(`Grid Under`, `Ruler Under`) %>%
  pivot_longer(everything(), names_to = "Calibration Method",
               values_to = "Effect on tarsus length estimate (mm)") %>%
  kbl(., format = "latex", caption = caption, escape = FALSE) %>%
  kable_styling()
```

```
rm(tarsusCorrection)

## Correcting with these means

data <- data %>%
  mutate("tarsusLengthMeanAdjusted" =
           ifelse(tarsusCalibration == "gridUnder",
                  tarsusLengthMean - tarsusDeltas$gridUnder,
                  ifelse(tarsusCalibration == "rulerUnder",
                         tarsusLengthMean - tarsusDeltas$rulerUnder,
                         tarsusLengthMean)
                  )
         )
```

We now check for measurement oddities in light of corrections.

```
ggplot(data %>%
         filter(week %in% c(0:3, 8)) %>%
         group_by(week) %>%
         mutate("ID" = 1:n()) %>%
         ungroup() %>%
         mutate("week" = ifelse(week == 1,
                                paste0("Age = ", week, " week"),
                                paste0("Age = ", week, " weeks"))
                ),
       aes(x = ID, y = tarsusLengthMeanAdjusted)) +
  facet_wrap(~week, scales = "free") +
  geom_point(size = 2, colour = "black",
             fill = "slateblue", alpha = 0.7) +
  theme_classic() +
  xlab("Sample Number") +
  ylab("Tarsus Length (mm)")
```

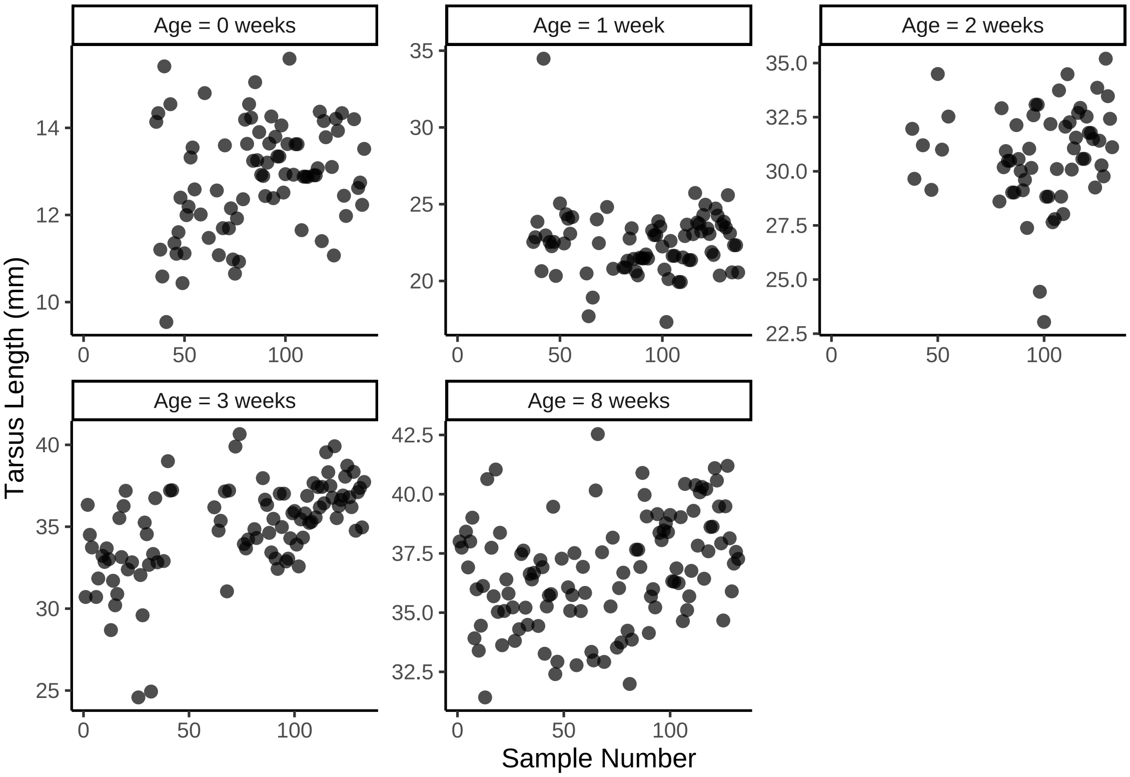


***Figure 4:*** *Cleveland dotplot of adjusted tarsus length estimates (mm) from Japanese quail at 0, 1, 2, 3, and 8 weeks of age.*

```
# One particularly high tarsus length measurement at 1
# week of age, although its value falls within the
# distribution of 2 weeks measurements. Retaining for
# this reason and reproducing tarsus timeline plot.
```

```
# And across ages

data %>%
  filter(week < 10) %>%
  ggplot(aes(x = week, y = tarsusLengthMeanAdjusted)) +
  geom_point(size = 2, pch = 21, fill = "slateblue", colour = "black",
             position = position_jitter(width = 0.5), alpha = 0.3) +
  stat_summary(geom = "errorbar", fun.data = "mean_cl_boot",
               colour = "black", width = 0.3) +
  stat_summary(geom = "point", fun = "mean", size = 3, pch = 21,
               colour = "black", fill = "black") +
  theme_classic() +
  xlab("Age (weeks)") +
  ylab("Tarsus Length (mm)")
```

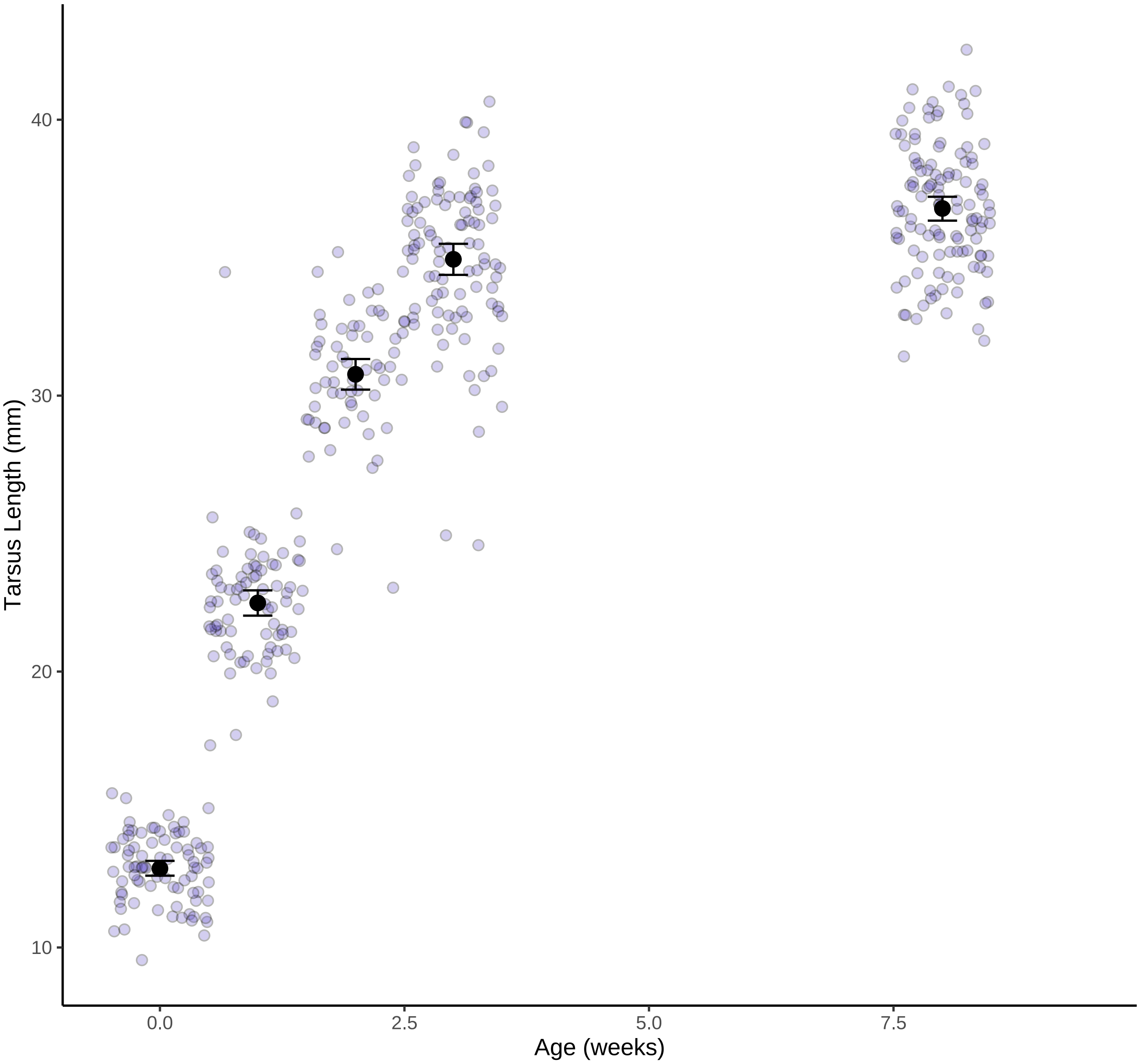


***Figure 5:*** *Raw tarsus length (mm) measurements during growth in Japanese quail (n = 133). Measurements are jittered by age to simplify visualisation.*

```
# No obvious oddities are detected.
```

With tarsus length measurements calibrated, we visualise trends of elongation across ages by treatment type. Here, effects of body mass are not corrected.

```
showtext_auto()

tarsusPlotA <- data %>%
  filter(!is.na(pretreatment) & !is.na(treatment) & week <= 8) %>%
  mutate(pretreatment = ifelse(pretreatment == "cold", "Cold (10°C)",
    ifelse(pretreatment == "neutral", "Mild (20°C)",
      "Warm (30°C)"
    )
  )) %>%
  ggplot(aes(x = week, y = tarsusLengthMean, fill = pretreatment)) +
  stat_summary(
```

```
    geom = "line", fun.data = "mean_se",
    position = position_dodge(width = 0.8), colour = "black",
    aes(linetype = pretreatment)
  ) +
  stat_summary(
    geom = "errorbar", fun.data = "mean_se", width = 0.3,
    position = position_dodge(width = 0.8)
  ) +
  stat_summary(
    geom = "point", fun = "mean", size = 3, colour = "black",
    pch = 21, position = position_dodge(width = 0.8)
  ) +
  scale_fill_manual(values = c("#7BB4E3", "black", "#CD5C5C"),
                    name = "Rearing\nConditions") +
  scale_linetype_manual(values = c("dashed", "solid", "dotted"),
                        name = "Rearing\nConditions") +
  xlab("Age (weeks)") +
  ylab("Tarsus Length (mm)") +
  ylim(c(15, 45)) +
  theme_classic() +
  theme(
    axis.title = element_text(size = 12, family = "Noto Sans", colour = "black"),
    legend.title = element_text(size = 12, family = "Noto Sans", colour = "black"),
    legend.text = element_text(size = 11, family = "Noto Sans", colour = "black"),
    legend.position = "bottom"
  )

p2 <- ggtexttable(
    data %>%
      filter(!is.na(pretreatment) & !is.na(treatment) &
               !is.na(tarsusLengthMean) & week <= 8 &
        pretreatment == "cold") %>%
      group_by(week) %>%
      count() %>%
      rename("Age (weeks)" = week),
    rows = NULL, theme = ttheme("light")
  ) %>%
    tab_add_title(text = "Cold\nRearing", face = "bold")

p3 <- ggtexttable(
      data %>%
        filter(!is.na(pretreatment) & !is.na(treatment) &
                 !is.na(tarsusLengthMean) & week <= 8 &
          pretreatment == "neutral") %>%
        group_by(week) %>%
        rename("Age (weeks)" = week) %>%
        count(),
      rows = NULL, theme = ttheme("light")
    ) %>%
      tab_add_title(text = "Mild\nRearing", face = "bold")

p4 <- ggtexttable(
        data %>%
          filter(!is.na(pretreatment) & !is.na(treatment) &
                   !is.na(tarsusLengthMean) & week <= 8 &
            pretreatment == "warm") %>%
          group_by(week) %>%
          rename("Age (weeks)" = week) %>%
          count(),
        rows = NULL, theme = ttheme("light")
      ) %>%
      tab_add_title(text = "Warm\nRearing", face = "bold")

tarsusPlotA/(p2 + p3 + p4)
```

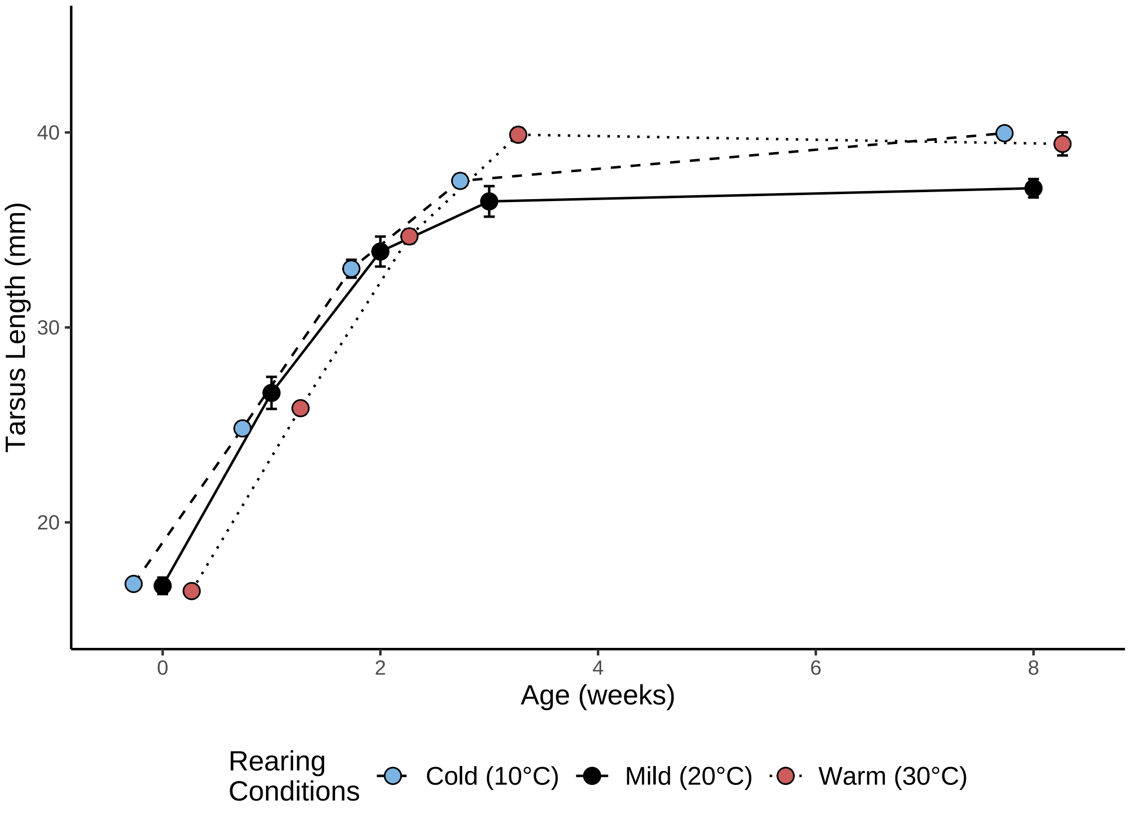


**Cold Rearing**

| Age (weeks) | n |
|---|---|
| 0 | 26 |
| 1 | 28 |
| 2 | 27 |
| 3 | 41 |
| 8 | 40 |

**Mild Rearing**

| Age (weeks) | n |
|---|---|
| 0 | 15 |
| 1 | 12 |
| 2 | 4 |
| 3 | 20 |
| 8 | 33 |

**Warm Rearing**

| Age (weeks) | n |
|---|---|
| 0 | 34 |
| 1 | 37 |
| 2 | 29 |
| 3 | 34 |
| 8 | 39 |

***Figure 6:*** *Effects of age and thermal environment during post-hatch development on tarsus length (mm) in Japanese quail. Dots represent means and errorbars represent standard errors.*

The above plot is reproduced below but only including individuals that were reared at 10°C, 20°C, or 30°C fully until maturity.

```
tarsusPlotB <- data %>%
  filter(!is.na(pretreatment) & !is.na(treatment) & week <= 8) %>%
  filter(!(exp == "A" & pretreatment == "cold") &
           !(exp == "B" & pretreatment == "warm")) %>%
  mutate(pretreatment = ifelse(pretreatment == "cold", "Cold (10°C)",
```

```
    ifelse(pretreatment == "neutral", "Mild (20°C)",
      "Warm (30°C)"
    )
  )) %>%
  ggplot(aes(x = week, y = tarsusLengthMean, fill = pretreatment)) +
  stat_summary(
    geom = "line", fun.data = "mean_se",
    position = position_dodge(width = 0.8), colour = "black",
    aes(linetype = pretreatment)
  ) +
  stat_summary(
    geom = "errorbar", fun.data = "mean_se", width = 0.3,
    position = position_dodge(width = 0.8)
  ) +
  stat_summary(
    geom = "point", fun = "mean", size = 3, colour = "black",
    pch = 21, position = position_dodge(width = 0.8)
  ) +
  scale_fill_manual(values = c("#7BB4E3", "black", "#CD5C5C"),
                    name = "Rearing\nConditions") +
  scale_linetype_manual(values = c("dashed", "solid", "dotted"),
                        name = "Rearing\nConditions") +
  xlab("Age (weeks)") +
  ylab("Tarsus Length (mm)") +
  ylim(c(15, 45)) +
  theme_classic() +
  theme(
    axis.title = element_text(size = 12, family = "Noto Sans", colour = "black"),
    legend.title = element_text(size = 12, family = "Noto Sans", colour = "black"),
    legend.text = element_text(size = 11, family = "Noto Sans", colour = "black"),
    legend.position = "bottom"
  )

p2 <- ggtexttable(
    data %>%
      filter(!is.na(pretreatment) & !is.na(treatment) &
               !is.na(tarsusLengthMean) & week <= 8) %>%
      filter(!(exp == "A" & pretreatment == "cold")) %>%
      filter(pretreatment == "cold") %>%
      group_by(week) %>%
      count() %>%
      rename("Age (weeks)" = week),
    rows = NULL, theme = ttheme("light")
  ) %>%
    tab_add_title(text = "Cold\nRearing", face = "bold")

p3 <- ggtexttable(
      data %>%
        filter(!is.na(pretreatment) & !is.na(treatment) &
                 !is.na(tarsusLengthMean) & week <= 8 &
          pretreatment == "neutral") %>%
        group_by(week) %>%
        rename("Age (weeks)" = week) %>%
        count(),
      rows = NULL, theme = ttheme("light")
    ) %>%
      tab_add_title(text = "Mild\nRearing", face = "bold")

p4 <- ggtexttable(
        data %>%
          filter(!is.na(pretreatment) & !is.na(treatment) &
                   !is.na(tarsusLengthMean) & week <= 8) %>%
          filter(!(exp == "B" & pretreatment == "warm")) %>%
          filter(pretreatment == "warm") %>%
          group_by(week) %>%
          rename("Age (weeks)" = week) %>%
          count(),
```

```
    rows = NULL, theme = ttheme("light")
  ) %>%
  tab_add_title(text = "Warm\nRearing", face = "bold")

tarsusPlotB/(p2 + p3 + p4)
```

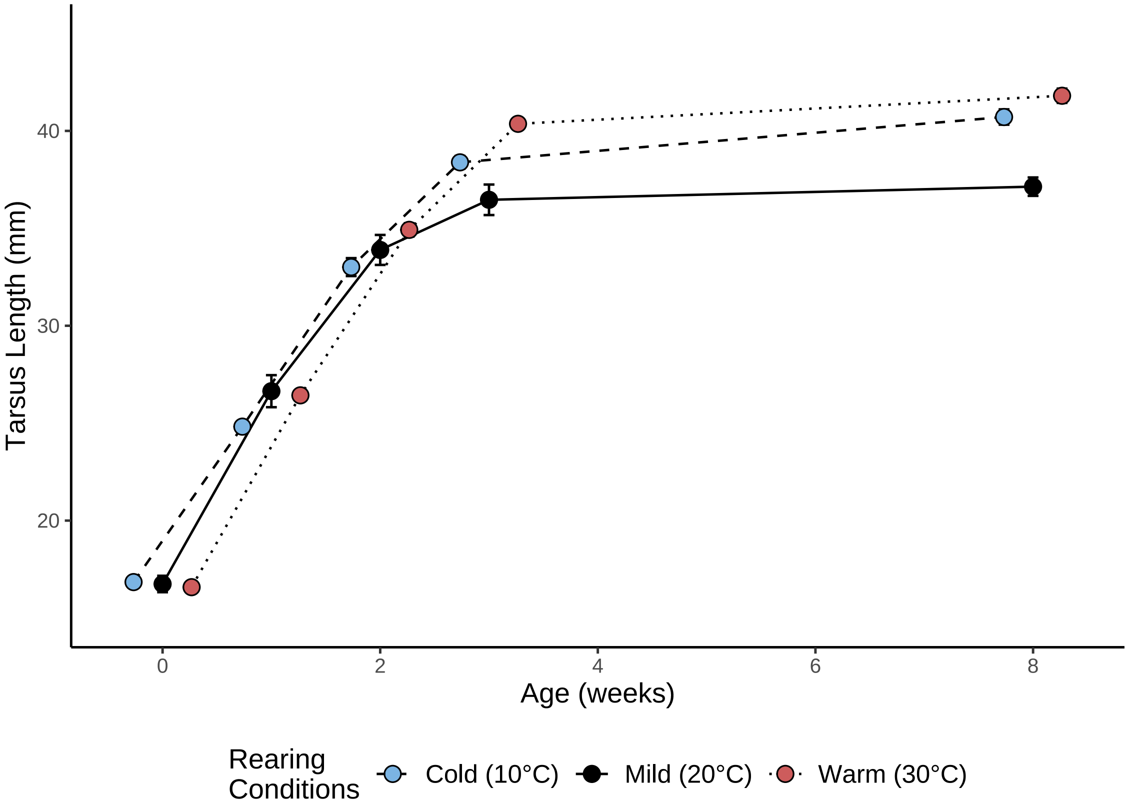


**Cold Rearing**

| Age (weeks) | n |
|---|---|
| 0 | 26 |
| 1 | 28 |
| 2 | 27 |
| 3 | 25 |
| 8 | 24 |

**Mild Rearing**

| Age (weeks) | n |
|---|---|
| 0 | 15 |
| 1 | 12 |
| 2 | 4 |
| 3 | 20 |
| 8 | 33 |

**Warm Rearing**

| Age (weeks) | n |
|---|---|
| 0 | 18 |
| 1 | 26 |
| 2 | 26 |
| 3 | 24 |
| 8 | 23 |

***Figure 7:*** *Effects of age and thermal environment during post-hatch development on tarsus length (mm) in Japanese quail, where only individuals reared in selected thermal environments for 8 weeks of are included. Dots represent group means and errorbars represent group standard errors.*

```
showtext_auto(enable = FALSE)

# Errorbars consistently small and warm treatment lengths
# generally greater than cold treatment lengths.
```

Data inspection is next continued, with particular focus on the spread of body mass data. A final data-frame containing all collated data is then saved in a single .csv file for future use.

```
# Checking for aberrant mass values

ggplot(
  data %>%
    filter(week <= 8) %>%
    group_by(week) %>%
    mutate("ID" = 1:n()) %>%
    ungroup() %>%
    mutate("week" = ifelse(week == 1,
                           paste0("Age = ", week, " week"),
                           paste0("Age = ", week, " weeks"))
           ),
  aes(x = ID, y = mass)
) +
  facet_wrap(~week, scales = "free") +
  geom_point(size = 2, colour = "black",
             fill = "slateblue", alpha = 0.7) +
  theme_classic() +
  xlab("Sample Number") +
  ylab("Body Mass (g)")
```

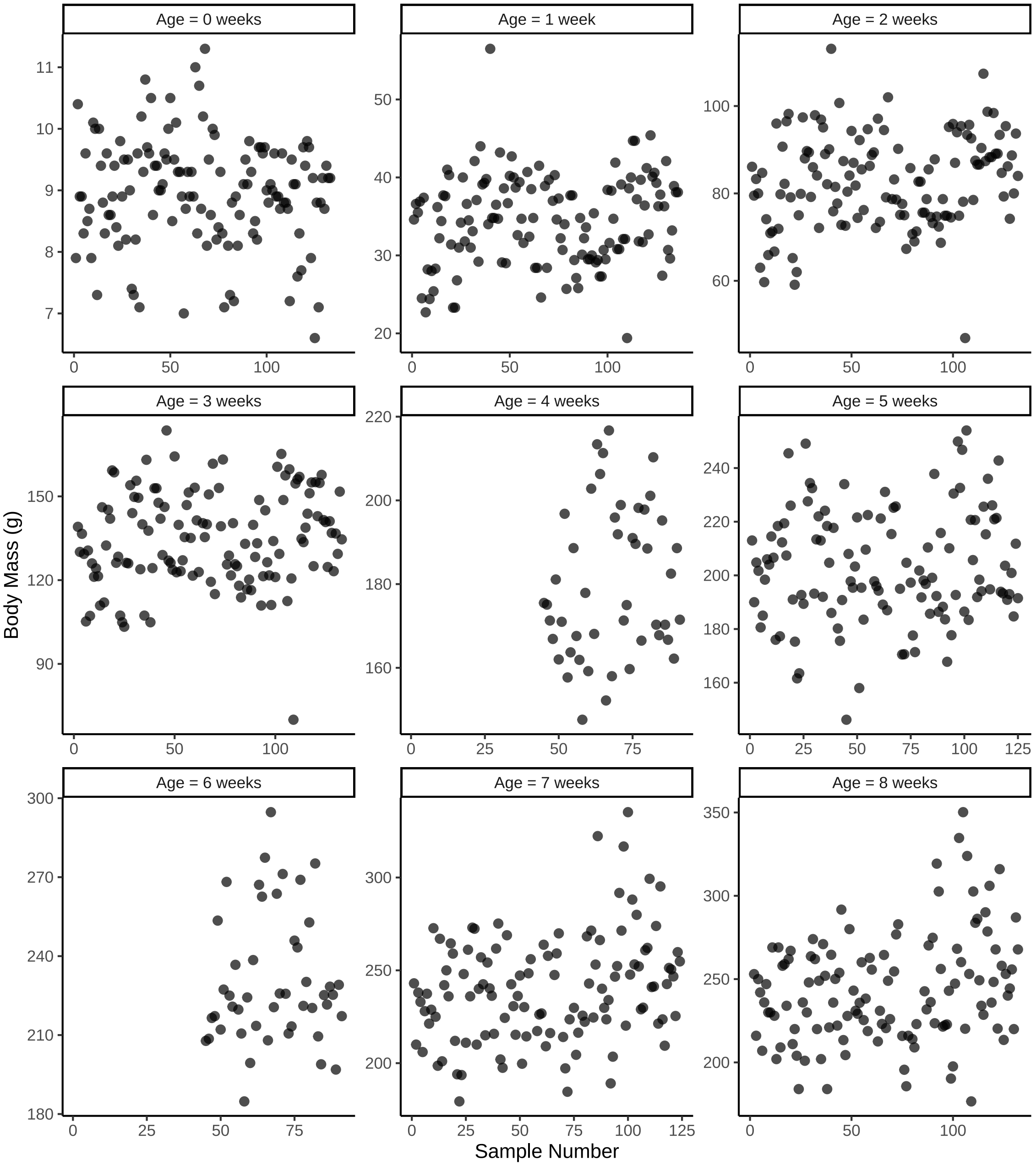


***Figure 8:*** *Cleveland dotplots displaying body mass (g) by sample number of Japanese quail.*

```
# One peculiarly low value (week 3) and one peculiarly high (week 1).
# Each, however, appear to fall within the distributions of
# surrounding weeks. Retaining for this reason.

# Removing raw, unadjusted tarsus measurements,
# then renaming adjusted values for simplicity.

data <- data %>%
  select(-c(tarsusLengthMean)) %>%
  rename("tarsusLengthMean" = tarsusLengthMeanAdjusted)
```

```
## Visualising change in mass across age

showtext_auto()
```

```
massPlot <- data %>%
  filter(!is.na(pretreatment) & !is.na(treatment) &
    week <= 8) %>%
  mutate(pretreatment = ifelse(pretreatment == "cold", "Cold (10°C)",
    ifelse(pretreatment == "neutral", "Mild (20°C)",
      "Warm (30°C)"
    )
  )) %>%
  ggplot(aes(x = week, y = mass, fill = pretreatment)) +
  stat_summary(
    geom = "line", fun.data = "mean_cl_boot",
    position = position_dodge(width = 0.5), colour = "black",
    aes(linetype = pretreatment)
  ) +
  stat_summary(
    geom = "errorbar", fun.data = "mean_cl_boot", width = 0.3,
    position = position_dodge(width = 0.5)
  ) +
  stat_summary(
    geom = "point", fun = "mean", size = 3, colour = "black",
    pch = 21, position = position_dodge(width = 0.5)
  ) +
  scale_fill_manual(values = c("#7BB4E3", "black", "#CD5C5C"),
                    name = "Rearing\nConditions") +
  scale_linetype_manual(values = c("dashed", "solid", "dotted"),
                        name = "Rearing\nConditions") +
  xlab("Age (weeks)") +
  ylab("Mass (g)") +
  ylim(c(0, 280)) +
  theme_classic() +
  theme(
    axis.title = element_text(size = 12, family = "Noto Sans", colour = "black"),
    legend.title = element_text(size = 12, family = "Noto Sans", colour = "black"),
    legend.text = element_text(size = 11, family = "Noto Sans", colour = "black"),
    legend.position = "bottom"
  )

p2 <- ggtexttable(
    data %>%
      filter(!is.na(pretreatment) & !is.na(treatment) &
               !is.na(mass) & week <= 8 &
        pretreatment == "cold") %>%
      group_by(week) %>%
      count() %>%
      rename("Age (weeks)" = week),
    rows = NULL, theme = ttheme("light")
  ) %>%
    tab_add_title(text = "Cold\nRearing", face = "bold")

p3 <- ggtexttable(
      data %>%
        filter(!is.na(pretreatment) & !is.na(treatment) &
                 !is.na(mass) & week <= 8 &
          pretreatment == "neutral") %>%
        group_by(week) %>%
        rename("Age (weeks)" = week) %>%
        count(),
      rows = NULL, theme = ttheme("light")
    ) %>%
    tab_add_title(text = "Mild\nRearing", face = "bold")

p4 <- ggtexttable(
      data %>%
        filter(!is.na(pretreatment) & !is.na(treatment) &
                 !is.na(mass) & week <= 8 &
          pretreatment == "warm") %>%
        group_by(week) %>%
```

```
        rename("Age (weeks)" = week) %>%
        count(),
      rows = NULL, theme = ttheme("light")
    ) %>%
    tab_add_title(text = "Warm\nRearing", face = "bold")

massPlot/(p2 + p3 + p4)
```

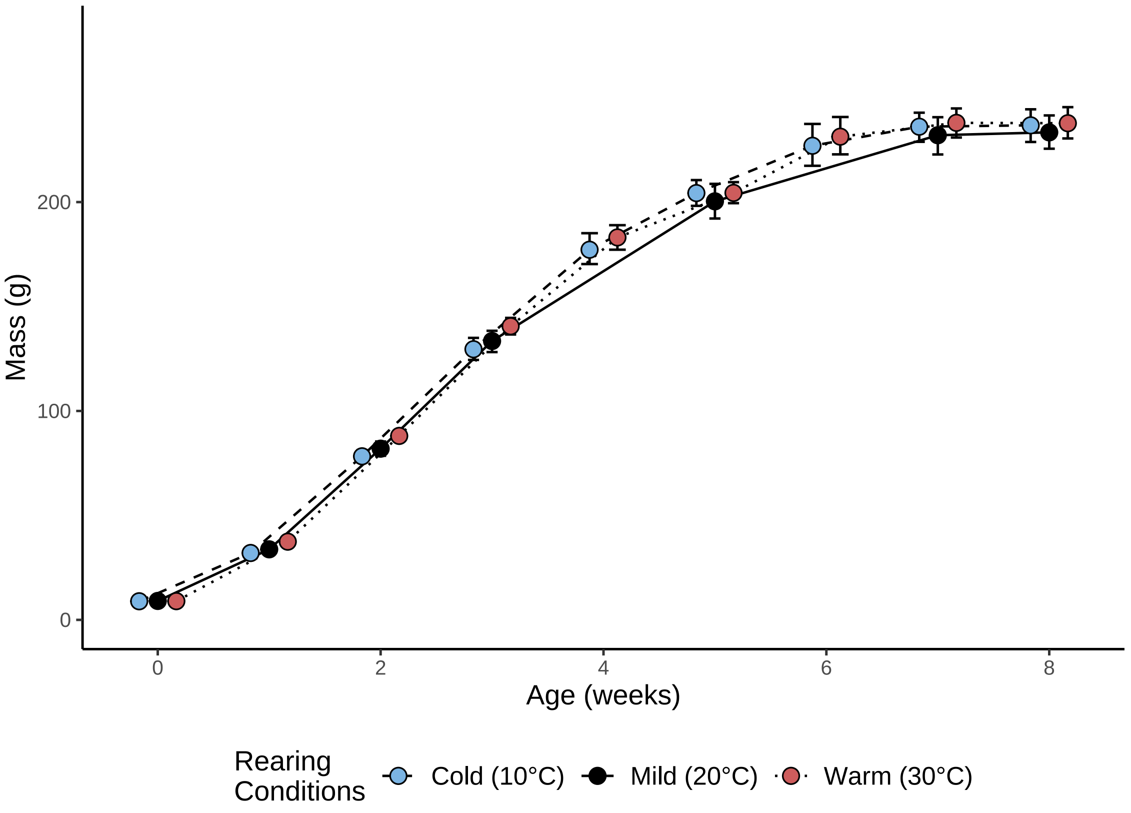


**Cold Rearing**

| Age (weeks) | n |
|---|---|
| 0 | 44 |
| 1 | 48 |
| 2 | 46 |
| 3 | 43 |
| 4 | 24 |
| 5 | 42 |
| 6 | 24 |
| 7 | 41 |
| 8 | 42 |

**Mild Rearing**

| Age (weeks) | n |
|---|---|
| 0 | 36 |
| 1 | 35 |
| 2 | 34 |
| 3 | 38 |
| 5 | 34 |
| 7 | 33 |
| 8 | 36 |

**Warm Rearing**

| Age (weeks) | n |
|---|---|
| 0 | 51 |
| 1 | 49 |
| 2 | 48 |
| 3 | 47 |
| 4 | 23 |
| 5 | 42 |
| 6 | 23 |
| 7 | 43 |
| 8 | 43 |

***Figure 9:*** *Effect of age and rearing conditions on body mass of Japanese quail. Dots represent group means and errorbars represent 2.5% and 97.5% quantiles.*

```
# Combining tarsus and mass plots.

earlyPanel <- ggarrange(tarsusPlotB, massPlot,
    common.legend = TRUE,
    legend = "bottom", ncol = 2
  )
```

```
print(earlyPanel)
```

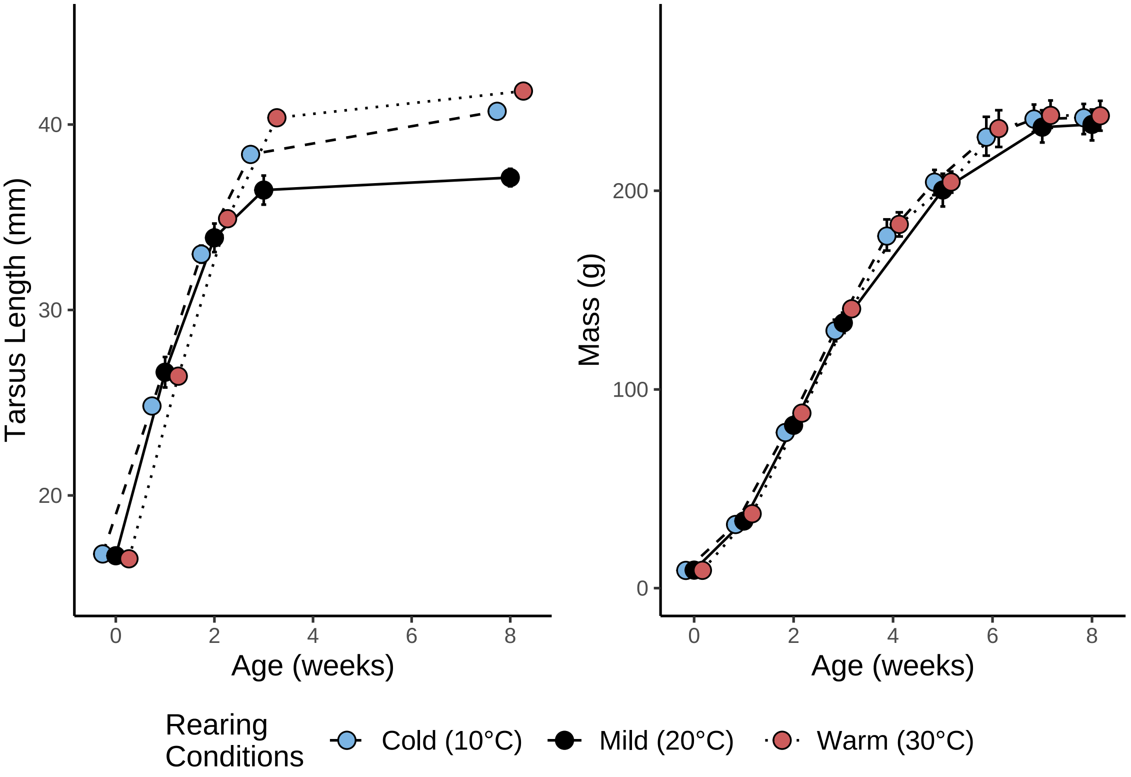


***Figure 10:*** *Effect of age and rearing conditions on morphometry of Japanese quail. Dots represent group means in both plots, while errorbars around tarsus length (mm) means represent standard errors and those around body mass (g) means represent 2.5% and 97.5% quantiles.*

Last, we visualise trends in relative tarsus length by age and treatment type. Tarsus length is relativised by estimating tarsus length residuals from a basic linear model with tarsus length (mm) as the Gaussian-distributed response variable and body mass (g) as the sole population-level predictor. Priors for this model are set broadly as follow:

$$Intercept\ (\beta_0) \sim exponential(0.075)$$

$$Body\ Mass\ (\beta_1) \sim \mathcal{N}(1, 2)$$

```
# Exploring residual tarsus and bill length trends

residualTarsus <- data %>%
  drop_na(tarsusLengthMean, mass) %>%
  mutate(
    "residualTarsus" =
      residuals(brm(tarsusLengthMean ~ mass,
        prior = c(
          set_prior("normal(1, 2)", class = "b"),
          set_prior("exponential(0.075)", class = "Intercept")
        ),
        data = .,
        cores = 1, chains = 4,
        seed = 100, family = "gaussian",
```

```
        iter = 50000, warmup = 5000, thin = 10,
        control = list(adapt_delta = 0.97, max_treedepth = 14),
        silent = TRUE, refresh = 0,
        file = "./models/residualModelPlotA-Median.Rds"
      ), robust = TRUE)[, "Estimate"]
  ) %>%
  filter(!is.na(pretreatment) & !is.na(treatment) & week <= 8) %>%
  mutate(pretreatment = ifelse(pretreatment == "cold", "Cold (10°C)",
    ifelse(pretreatment == "neutral", "Mild (20°C)",
      "Warm (30°C)"
    )
  )) %>%
  ggplot(aes(x = week, y = residualTarsus, fill = pretreatment)) +
  stat_summary(
    geom = "line", fun.data = "mean_cl_boot",
    position = position_dodge(width = 0.4), colour = "black",
    aes(linetype = pretreatment)
  ) +
  stat_summary(
    geom = "errorbar", fun.data = "mean_cl_boot", width = 0.3,
    position = position_dodge(width = 0.4)
  ) +
  stat_summary(
    geom = "point", fun = "mean", size = 3, colour = "black",
    pch = 21, position = position_dodge(width = 0.4)
  ) +
  scale_fill_manual(values = c("#7BB4E3", "black", "#CD5C5C"),
                    name = "Rearing\nConditions") +
  scale_linetype_manual(values = c("dashed", "solid", "dotted"),
                        name = "Rearing\nConditions") +
  xlab("Age (weeks)") +
  ylab("Residual\nTarsus Length (mm)") +
  theme_classic() +
  theme(
    axis.title = element_text(size = 12, family = "Noto Sans", colour = "black"),
    legend.title = element_text(size = 12, family = "Noto Sans", colour = "black"),
    legend.text = element_text(size = 11, family = "Noto Sans", colour = "black"),
    legend.position = "bottom"
  )

p2 <- ggtexttable(
    data %>%
      filter(!is.na(pretreatment) & !is.na(treatment) &
               !is.na(mass) &
        !is.na(tarsusLengthMean) & week <= 8 &
          pretreatment == "cold") %>%
      group_by(week) %>%
      count() %>%
      rename("Age (weeks)" = week),
    rows = NULL, theme = ttheme("light")
  ) %>%
    tab_add_title(text = "Cold\nRearing", face = "bold")

p3 <- ggtexttable(
      data %>%
        filter(!is.na(pretreatment) & !is.na(treatment) &
                 !is.na(mass) &
          !is.na(tarsusLengthMean) & week <= 8 &
            pretreatment == "neutral") %>%
        group_by(week) %>%
        rename("Age (weeks)" = week) %>%
        count(),
      rows = NULL, theme = ttheme("light")
    ) %>%
      tab_add_title(text = "Mild\nRearing", face = "bold")

p4 <- ggtexttable(
```

```
        data %>%
          filter(!is.na(pretreatment) & !is.na(treatment) &
                   !is.na(mass) &
            !is.na(tarsusLengthMean) & week <= 8 &
              pretreatment == "warm") %>%
          group_by(week) %>%
          rename("Age (weeks)" = week) %>%
          count(),
        rows = NULL, theme = ttheme("light")
      ) %>%
      tab_add_title(text = "Warm\nRearing", face = "bold")

residualTarsus/(p2 + p3 + p4)
```

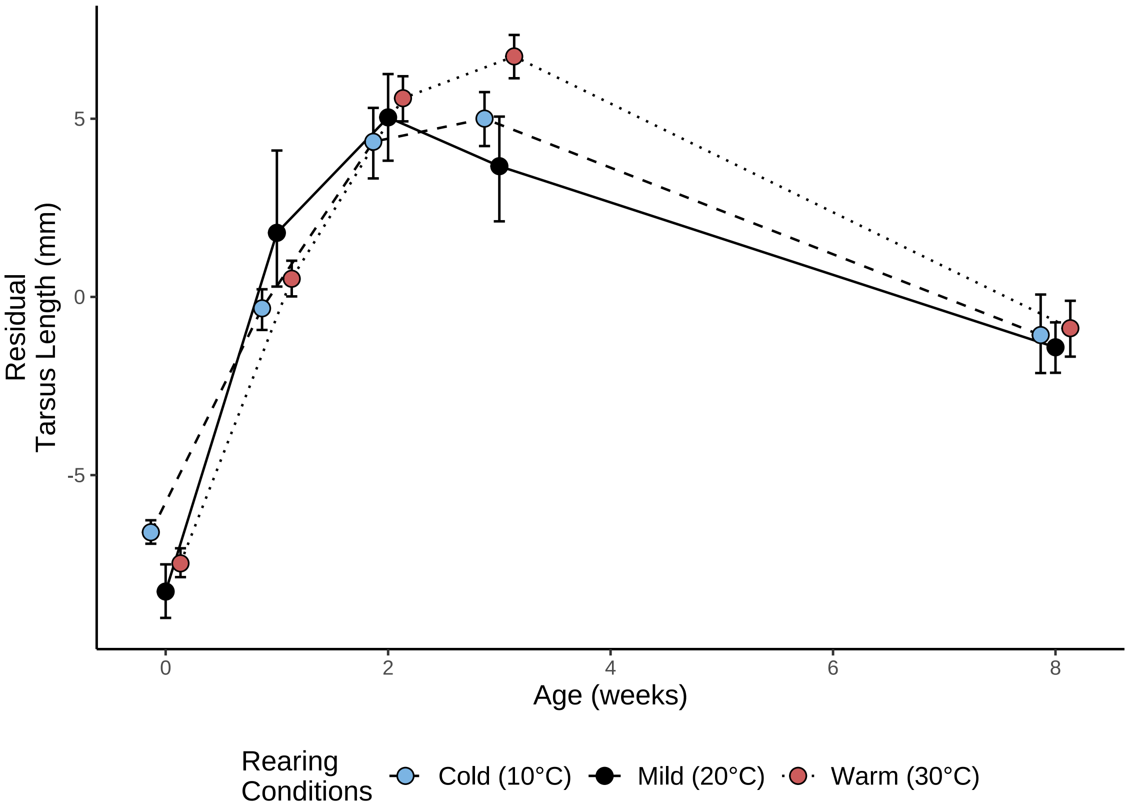


**Cold Rearing**

| Age (weeks) | n |
|---|---|
| 0 | 25 |
| 1 | 28 |
| 2 | 27 |
| 3 | 41 |
| 8 | 40 |

**Mild Rearing**

| Age (weeks) | n |
|---|---|
| 0 | 15 |
| 1 | 12 |
| 2 | 4 |
| 3 | 20 |
| 8 | 33 |

**Warm Rearing**

| Age (weeks) | n |
|---|---|
| 0 | 34 |
| 1 | 37 |
| 2 | 29 |
| 3 | 34 |
| 8 | 39 |

***Figure 11:*** *Effect of age and rearing conditions on relative tarsus length (mm) of Japanese quail. Relative tarsus length represents median residuals from a Bayesian linear model with tarsus length (mm) as as the response variable and body mass (g) as the sole predictor. Dots represent group means and errorbars represent 2.5% and 97.5% quantiles.*

Variance in morphological traits is calculated for reference, then our data-set saved.

```
caption = paste0('Average and variance of size metrics among ',
                 'captive-reared Japanese quail. "SD" indicates ',
                 'standard deviation and "CV" indicates ',
                 'coefficient of variation.')
```

```
data %>%
  filter(week <= 8) %>%
  select(week, "tarsus" = tarsusLengthMean, mass) %>%
  pivot_longer(!week, names_to = "Metric", values_to = "Size") %>%
  group_by(week, Metric) %>%
  summarise("Mean" = mean(Size, na.rm = T),
            "SD" = sd(Size, na.rm = T)) %>%
  mutate("CV" = (SD/Mean)*100,
         Metric = ifelse(Metric == "tarsus",
                         "Tarsus Length\n(mm)",
                         "Body Mass\n(g)")
         ) %>%
  select(Metric, "Age (Weeks)" = week, Mean, SD, "CV (\\%)" = CV) %>%
  kbl(., longtable = T, booktabs = T,
      format = "latex", caption = caption,
      escape = FALSE) %>%
  kable_styling()
```

***Table 3:*** *Average and variance of size metrics among captive-reared Japanese quail. "SD" indicates standard deviation and "CV" indicates coefficient of variation.*

| Metric | Age (Weeks) | Mean | SD | CV (%) |
|---|---|---|---|---|
| Body Mass (g) | 0 | 8.970992 | 0.8800777 | 9.810260 |
| Tarsus Length (mm) | 0 | 12.861453 | 1.2539701 | 9.749833 |
| Body Mass (g) | 1 | 34.494697 | 5.7158720 | 16.570292 |
| Tarsus Length (mm) | 1 | 22.487715 | 2.1651457 | 9.628127 |
| Body Mass (g) | 2 | 82.934375 | 10.6426229 | 12.832584 |
| Tarsus Length (mm) | 2 | 30.774077 | 2.2388453 | 7.275101 |
| Body Mass (g) | 3 | 134.767188 | 16.8622913 | 12.512164 |
| Tarsus Length (mm) | 3 | 34.940364 | 2.8407332 | 8.130234 |
| Body Mass (g) | 4 | 180.074468 | 17.9338697 | 9.959141 |
| Tarsus Length (mm) | 4 | NaN | NA | NA |
| Body Mass (g) | 5 | 203.204237 | 21.2254137 | 10.445360 |
| Tarsus Length (mm) | 5 | NaN | NA | NA |
| Body Mass (g) | 6 | 230.534043 | 24.6884173 | 10.709228 |
| Tarsus Length (mm) | 6 | NaN | NA | NA |
| Body Mass (g) | 7 | 239.803419 | 28.6671232 | 11.954426 |
| Tarsus Length (mm) | 7 | NaN | NA | NA |
| Body Mass (g) | 8 | 243.977686 | 32.4864600 | 13.315341 |
| Tarsus Length (mm) | 8 | 36.782459 | 2.3369011 | 6.353303 |

```
# Saving compiled data-frame

write.csv(data, "compiledData.csv", row.names = F)
data <- read.csv("compiledData.csv")
```

#### Modelling effects of the thermal environment during development on morphology

In this subsection, we evaluate the effect of the post-hatch thermal environment on body size (here, body mass in g) and appendage length (here, tarsus length in mm) of captive-reared Japanese quail. As described above, this is achieved by modelling each variable as a Gompertz function of age in weeks, and testing for an effect of rearing condition on any, and all, parameters of the Gompertz curve (i.e. a, b, and c). For body mass, our model was therefore as follows:

$$Mass_{ij} \sim a \cdot e^{-b \cdot e^{-c \cdot Age_{ij}}} + \mu_{0j} + \epsilon_{ij}$$

$$a_{ij} \sim \beta_{a0} + \beta_{a1} \cdot Cold\ Reared_j + \beta_{a2} \cdot Warm\ Reared_j + \mu_{0aj}$$

$$b_{ij} \sim \beta_{b0} + \beta_{b1} \cdot Cold\ Reared_j + \beta_{b2} \cdot Warm\ Reared_j + \mu_{0bj}$$

$$c_{ij} \sim \beta_{c0} + \beta_{c1} \cdot Cold\ Reared_j + \beta_{c2} \cdot Warm\ Reared_j + \mu_{0cj}$$

where $a$, $b$, and $c$ represent growth curve parameters implicit in the Gompertz function, *Cold Reared* and *Warmed Reared* represent logical, true/false variables per individual (with true equaling 1 and false equaling 0), $\beta_x$ values represent model coefficients for growth curve parameters $x$ ($a$ - $c$), $i$ represents an observation, $j$ represents an individual, $\mu_0$ represents a group-level intercept for individual $j$, $\mu_{0a}$ - $\mu_{0c}$ represent group-level intercepts corresponding to the batch of eggs from which an individual was derived from, and $\epsilon$ represents the model error structure. Because we expected the standard deviation of body mass to also increase with age, $\epsilon$ was therefore modeled as:

$$ln(\epsilon_{ij}) \sim \tau_0 + \tau_1 * ln(Age_{ij} + 1)$$

where $\tau_0$ represents the intercept for $\epsilon$, and $\tau_1$ represents the rate at which $\epsilon$, or its natural logarithm, increases with the natural logarithm of an individual's age in weeks plus 1 (i.e. to limit week 0 being equal to negative infinity).

Priors for all model parameters were moderately informative and selected according to findings by Narinc et al (2010), Burness et al (2013), Haqani et al (2021), and Persson et al (2024). Specifically, our prior for the asymptote of our growth curve ($\beta_{a0}$) was normally distributed with a mean of 250 and standard deviation of 25, while that for our x-axis displacement ($\beta_{b0}$) was normally distributed with mean of 3 and standard deviation of 1 (i.e. in line with expectations from Narinc et al. 2010). For our growth rate ($\beta_{c0}$), we used a skew-normal prior with $\xi$ set to 0.5, $\omega$ set to 0.1, and $\alpha$ set to 2.5 (thus assuming that growth rate could not be $< 0$, and should lay near 0.5). Priors for the effect of rearing condition on growth parameters $a$, $b$, and $c$ (i.e. $\beta_{a1-2}$, $\beta_{b1-2}$, and $\beta_{c1-2}$ respectively) were all normally distributed. Here, for effects of cold-rearing and warm-rearing on $a$, we assumed means of 7.5 and -7.5 respectively, given that quail reared at 15°C were approximately 15 g heavier that those reared at 30°C by 66 days in Burness et al (2013); standard deviations were, however, set broadly to 25. Others treatment-specific effects were assigned means of 0 and standard deviations 0.5 ($\beta_{b1-2}$), and 0.2 ($\beta_{c1-2}$) respectively. For our group-level effects of egg batch, we used exponential priors with lambda values determined from preliminary plots ($\mu_{0a}$: $\lambda = 2.5$; $\mu_{0b}$: $\lambda = 10$; $\mu_{0c}$: $\lambda = 25$). Similarly, for our group-level effect of bird identity, we also used an exponential prior, however, with lambda set broadly to 0.5. Finally, skew-normal priors were used for our error structure parameters $\tau_0$ and $\tau_1$ with $\xi$ values of 1, $\omega$ values of 0.5, and $\alpha$ values of 10 and -10 respectively.

For this model, 4 Hamiltonian Monte Carlo (HMC) chains were used, with each run for 50000 iterations and 10000 warm-up iterations, then sampled every 10 iterations. Suitability of priors is evaluated by a prior predictive check below.

First, we check that the standard deviation of body mass does indeed increase with age as anticipated.

```
data %>%
  filter(week <= 8) %>%
  mutate(pretreatment = factor(pretreatment,
    levels = c("neutral", "cold", "warm")
  )) %>%
  select(ring, pretreatment, week, mass) %>%
  distinct() %>%
  mutate(weekB = log(week + 1)) %>%
  group_by(week, weekB) %>%
  summarise(SD = log(sd(mass, na.rm = T))) %>%
  ungroup() %>%
  ggplot(aes(x = weekB, y = SD)) +
  geom_point(size = 3, pch = 21, colour = "black", fill = "grey50") +
  geom_smooth(method = "lm", colour = "black",
              linetype = "dashed", se = FALSE) +
  xlab("Age (Weeks + 1; Natural-log Transformed )") +
  ylab("Standard Deviation of Body Mass (Natural-log Transformed)") +
  theme_classic()
```

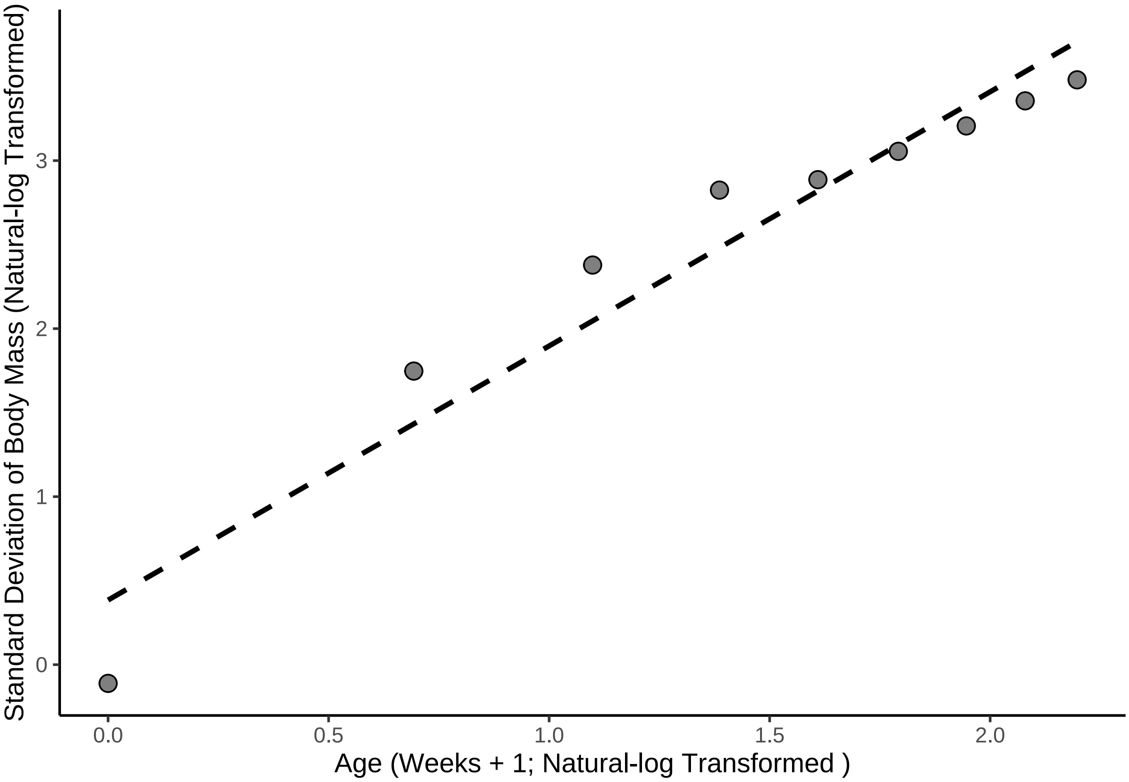


***Figure 12:*** *Effect of natural log-transformed age (in weeks + 1) on the natural log-transformed standard deviation of body mass (g). Dotted line represents a line of body best fit estimated from a linear relationship by internal functions of the R package ggplot (Wickham, 2011).*

```
# Reasonably linear.
```

Our prior predictive check is then produced.

```
growthModel_priorCheck <- brm(
  data = data %>%
    filter(week <= 8) %>%
    mutate(pretreatment = factor(pretreatment,
                                 levels = c("neutral", "cold", "warm")
    )) %>%
    select(ring, pretreatment, week, mass, "batch" = exp) %>%
    distinct() %>%
    mutate(weekB = week + 1),
  formula = bf(mass ~ A * exp(-B * exp(-C * week)) + D,
               A ~ 1 + pretreatment + (1|batch),
               B ~ 1 + pretreatment + (1|batch),
               C ~ 1 + pretreatment + (1|batch),
               D ~ 0 + (1 | ring),
               sigma ~ 0 + intercept + log(weekB),
               nl = TRUE
  ),
  prior = c(
    set_prior("normal(250, 25)",
              class = "b", coef = "Intercept",
              nlpar = "A"
    ),
    set_prior("normal(7.5, 25)",
              class = "b", coef = "pretreatmentcold",
              nlpar = "A"
    ),
```

```
    set_prior("normal(-7.5, 25)",
              class = "b", coef = "pretreatmentwarm",
              nlpar = "A"
    ),
    set_prior("exponential(2.5)", class = "sd",
              coef = "Intercept",
              group = "batch", nlpar = "A"
    ),
    set_prior("normal(3, 1)",
              class = "b", coef = "Intercept",
              nlpar = "B"
    ),
    set_prior("normal(0, 0.5)",
              class = "b", coef = "pretreatmentcold",
              nlpar = "B"
    ),
    set_prior("normal(0, 0.5)",
              class = "b", coef = "pretreatmentwarm",
              nlpar = "B"
    ),
    set_prior("exponential(10)", class = "sd",
              coef = "Intercept", group = "batch",
              nlpar = "B"
    ),
    set_prior("skew_normal(0.5, 0.1, 2.5)",
              class = "b", coef = "Intercept",
              nlpar = "C"
    ),
    set_prior("normal(0, 0.2)",
              class = "b", coef = "pretreatmentcold",
              nlpar = "C"
    ),
    set_prior("normal(0, 0.2)",
              class = "b", coef = "pretreatmentwarm",
              nlpar = "C"
    ),
    set_prior("exponential(25)", class = "sd",
              coef = "Intercept", group = "batch",
              nlpar = "C"
    ),
    set_prior("exponential(0.5)", class = "sd",
              coef = "Intercept", group = "ring", nlpar = "D"),
    set_prior("skew_normal(1, 0.5, -10)", class = "b",
              coef = "intercept", dpar = "sigma"),
    set_prior("skew_normal(1, 0.5, 10)", class = "b",
              coef = "logweekB", dpar = "sigma")
  ),
  family = "gaussian",
  seed = 100,
  cores = 4, chains = 4,
  iter = 50000, warmup = 10000, thin = 10,
  control = list(adapt_delta = 0.98, max_treedepth = 16),
  sample_prior = "only",
  silent = TRUE, refresh = 0,
  file = "./models/ppCheckMass.Rds"
)

pp_check2(growthModel_priorCheck, xlab = "Body Mass (g)") +
  xlim(c(0, 1000))
```

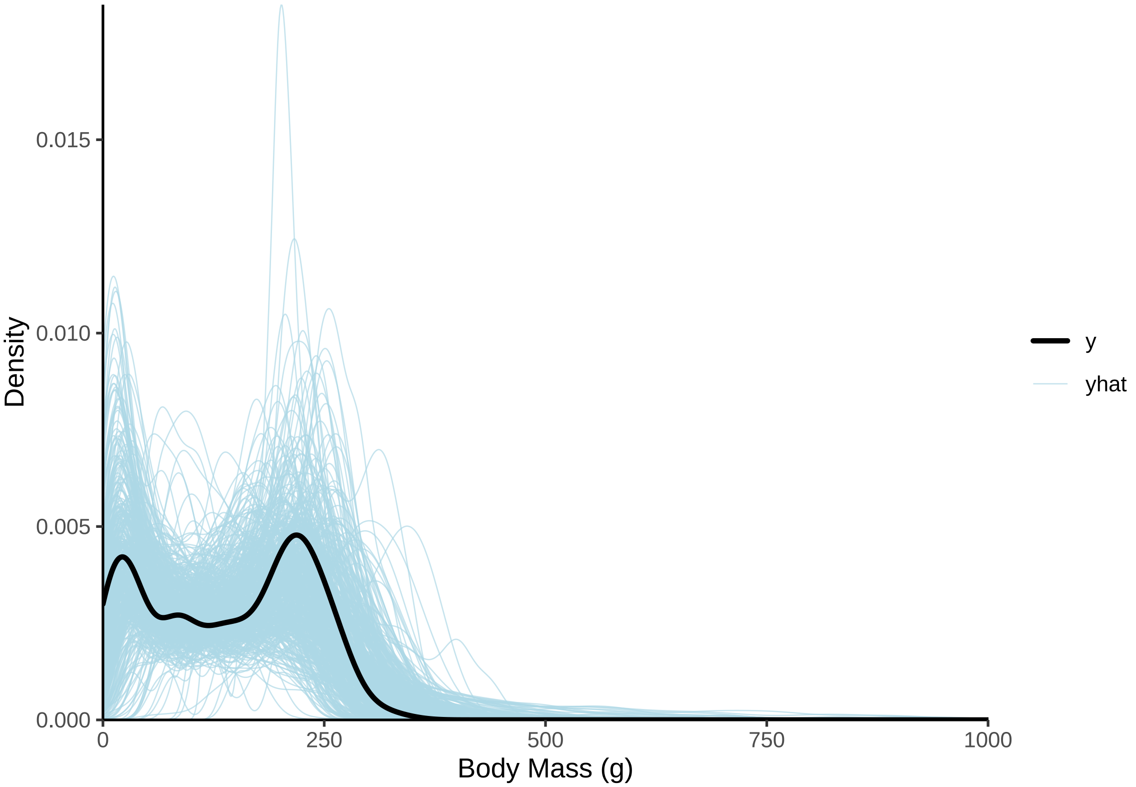


***Figure 13:*** *Overlay of predicted (blue) and true (black) body mass densities, where predicted densities are derived from priors in a Bayesian non-linear model. Clear overlap betwen the black and blue lines suggests that model priors are reasonable with respect to the data.*

```
# Some extreme values permitted, although reasonable overlay.
```

To guide our HMC chains for our full model, initial values were selected from within prior distributions.

```
# Acceptable prior selection. Creating function from
# which to draw initial HMC chain values.

initFunction <- function(chain_id = 1) {
  list(
    "b_A" = c(rnorm(1, 250, 25), 7.5, -7.5),
    "b_B" = c(3, 0, 0),
    "b_C" = c(0.5, 0, 0),
    "b_sigma" = c(rskew_normal(1, xi = 1, omega = 0.5, alpha = -5), 1),
    "sd_1" = rexp(1, 2.5),
    "sd_2" = rexp(1, 10),
    "sd_3" = rexp(1, 25),
    "sd_4" = rexp(1, 0.5)
  )
}

initList <- lapply(1:4, initFunction)

growthModel <- brm(
  data = data %>%
    filter(week <= 8) %>%
    mutate(pretreatment = factor(pretreatment,
                                 levels = c("neutral", "cold", "warm")
    )) %>%
    select(ring, pretreatment, week, mass, "batch" = exp) %>%
    distinct() %>%
```

```
  mutate(weekB = week + 1),
formula = bf(mass ~ A * exp(-B * exp(-C * week)) + D,
             A ~ 1 + pretreatment + (1|batch),
             B ~ 1 + pretreatment + (1|batch),
             C ~ 1 + pretreatment + (1|batch),
             D ~ 0 + (1 | ring),
             sigma ~ 0 + intercept + log(weekB),
             nl = TRUE
),
prior = c(
  set_prior("normal(250, 25)",
            class = "b", coef = "Intercept",
            nlpar = "A"
  ),
  set_prior("normal(7.5, 25)",
            class = "b", coef = "pretreatmentcold",
            nlpar = "A"
  ),
  set_prior("normal(-7.5, 25)",
            class = "b", coef = "pretreatmentwarm",
            nlpar = "A"
  ),
  set_prior("exponential(2.5)", class = "sd",
            coef = "Intercept",
            group = "batch", nlpar = "A"
  ),
  set_prior("normal(3, 1)",
            class = "b", coef = "Intercept",
            nlpar = "B"
  ),
  set_prior("normal(0, 0.5)",
            class = "b", coef = "pretreatmentcold",
            nlpar = "B"
  ),
  set_prior("normal(0, 0.5)",
            class = "b", coef = "pretreatmentwarm",
            nlpar = "B"
  ),
  set_prior("exponential(10)", class = "sd",
            coef = "Intercept", group = "batch",
            nlpar = "B"
  ),
  set_prior("skew_normal(0.5, 0.1, 2.5)",
            class = "b", coef = "Intercept",
            nlpar = "C"
  ),
  set_prior("normal(0, 0.2)",
            class = "b", coef = "pretreatmentcold",
            nlpar = "C"
  ),
  set_prior("normal(0, 0.2)",
            class = "b", coef = "pretreatmentwarm",
            nlpar = "C"
  ),
  set_prior("exponential(25)", class = "sd",
            coef = "Intercept", group = "batch",
            nlpar = "C"
  ),
  set_prior("exponential(0.5)", class = "sd",
            coef = "Intercept", group = "ring", nlpar = "D"),
  set_prior("skew_normal(1, 0.5, -10)", class = "b",
            coef = "intercept", dpar = "sigma"),
  set_prior("skew_normal(1, 0.5, 10)", class = "b",
            coef = "logweekB", dpar = "sigma")
),
family = "gaussian",
init = initList,
```

```
  seed = 100,
  cores = 4, chains = 4,
  iter = 50000, warmup = 10000, thin = 10,
  control = list(adapt_delta = 0.98, max_treedepth = 16),
  silent = TRUE, refresh = 0,
  file = "./models/growthModelMass.Rds"
)

# Very few divergences (~0.01%). Checking chain mixing and chain autocorrelation

ggarrange(
  ggplot(data = data.frame("Rhat" = brms::rhat(growthModel)),
         aes(x = Rhat)) +
    geom_density() +
    theme_classic() +
    xlab(
      TeX('$\\hat{R}$')
    ) +
    ylab("Density"),
  ggplot(data = data.frame("Neff" = neffBase(growthModel)),
         aes(x = Neff)) +
    geom_density() +
    theme_classic() +
    xlab(
      TeX('$N_{eff}/N~Ratio$')
    ) +
    ylab("Density")
)
```

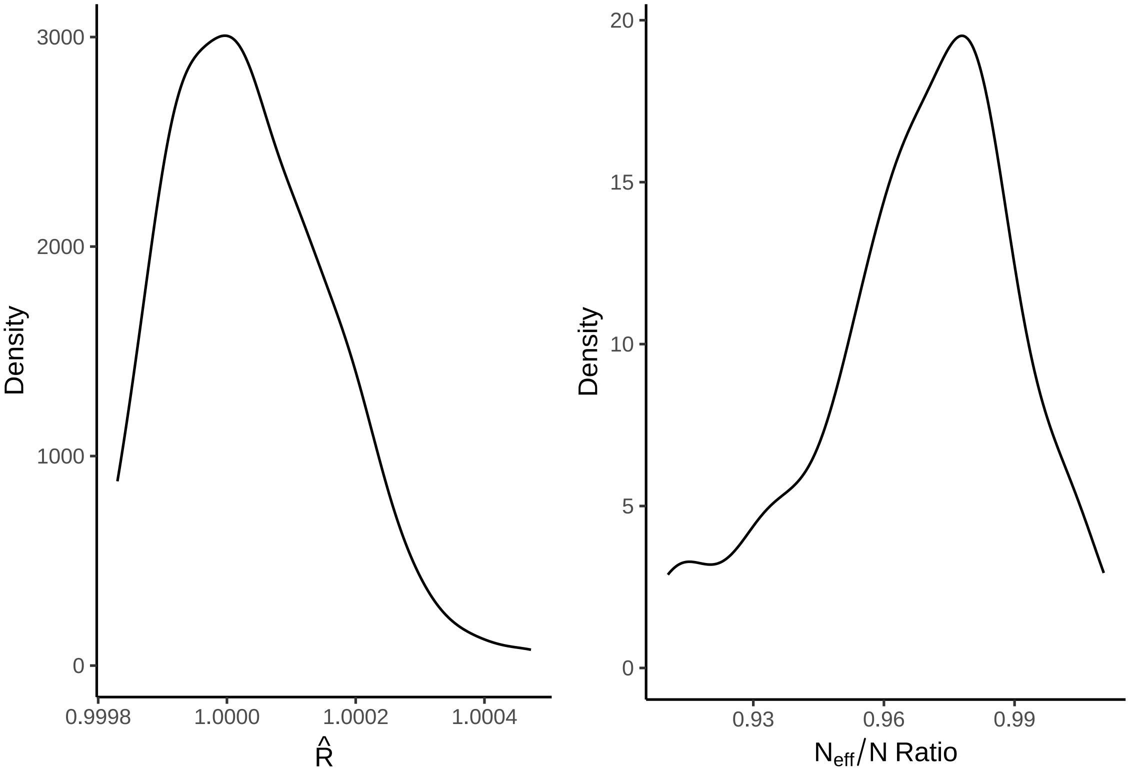


***Figure 14:*** *Gelman-Rubin statistics ($\hat{R}$) and ratio of effective samples sizes by samples sizes per parameter from a Bayesian non-linear model estimating growth of mass (g) among Japanese quail.*

No obvious autocorrelation is detected in our HMC chains, and all appear well converged. We next visualise our posterior predictions.

```r
pp_check2(growthModel, xlab = "Body Mass (g)")
```

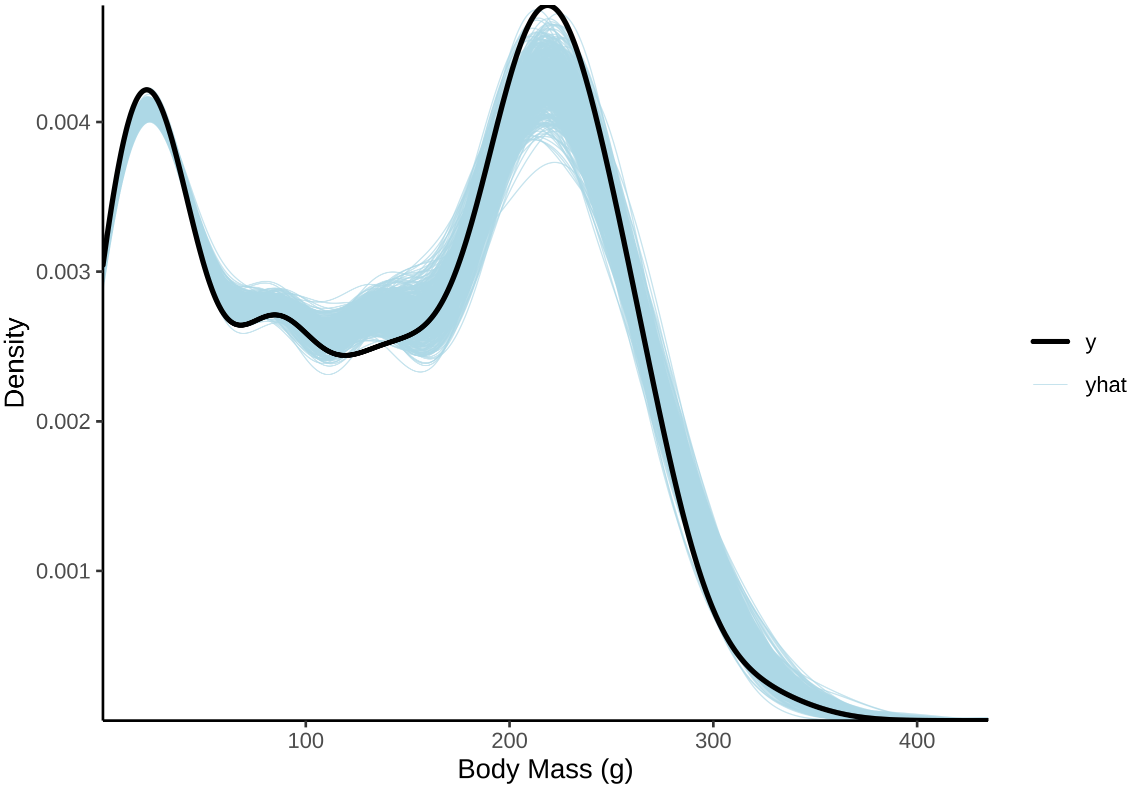


***Figure 15:*** *Posterior predictions of a Bayesian non-linear model predicting Japanese quail mass (g) across time, overlayed with true distributions of quail mass. Blue lines represent predictions from posterior draws, while the black line represents true mass distributions.*

```r
# Clear overlay. Checking fitted values using a scatter-plot.
```

```r
growthModel$data %>%
  mutate("Fit" = predict(growthModel)[,"Estimate"]) %>%
  ggplot(aes(x = mass, y = Fit, fill = week)) +
  geom_point(pch = 21, colour = "black", size = 2, alpha = 0.5) +
  geom_smooth(method = "lm", colour = "black",
              linetype = "dashed", se = FALSE) +
  scale_fill_gradient2(name = "Age (Weeks)") +
  theme_classic() +
  xlab("Body Mass (g)") +
  ylab("Expected Body Mass (g)")
```

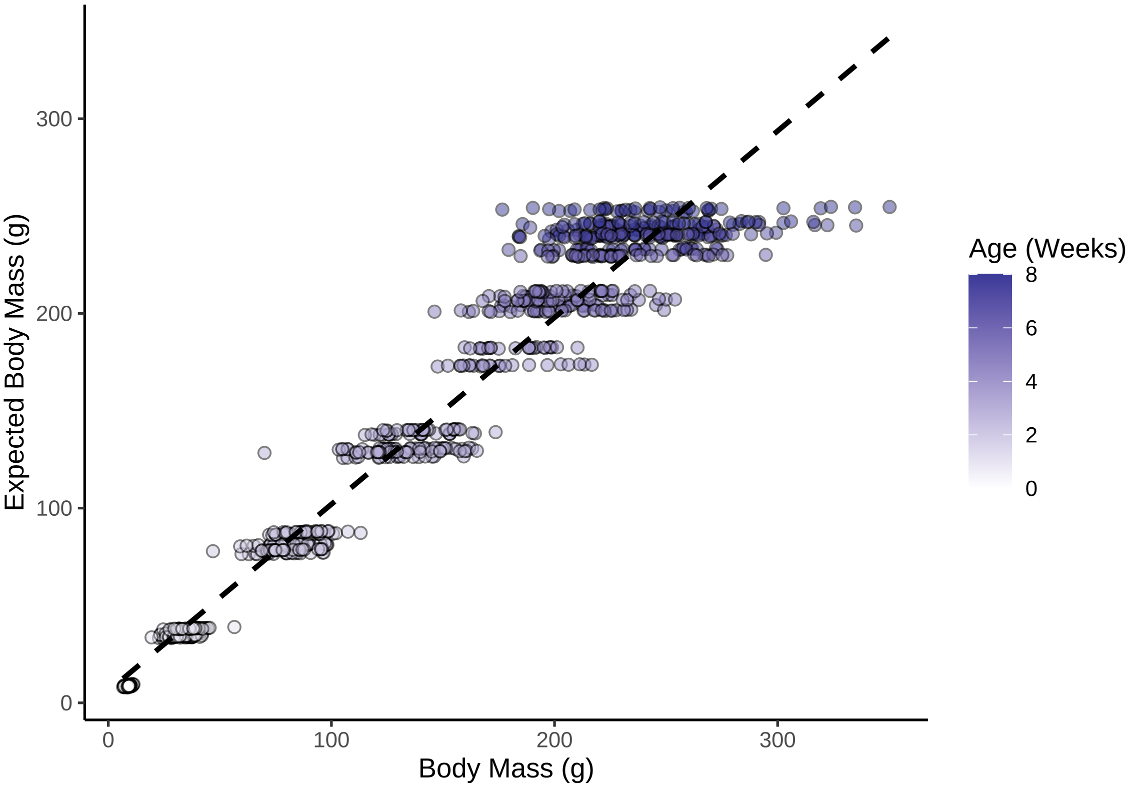


***Figure 16:*** *Scatterplot of Japanese quail body mass (g) against predicted body masses (g) from a Bayesian non-linear model.*

```
# Reasonable, but spread appears large by week.
# Plotting mean responses by rearing treatment.
```

```
with(
  growthModel$data,
  expand.grid(
    "week" = seq(0, 8, by = 0.1),
    "pretreatment" = c("cold", "neutral", "warm")
  )
) %>%
  mutate("weekB" = week + 1) %>%
  mutate(
    "Fit" = predict(growthModel, re_form = NA, robust = TRUE,
                    newdata = .)[, "Estimate"],
    "SE" = predict(growthModel, re_form = NA, robust = TRUE,
                   newdata = .)[, "Est.Error"]
  ) %>%
  mutate(
    "LCL" = Fit - SE,
    "UCL" = Fit + SE
  ) %>%
  mutate(pretreatment = str_to_title(pretreatment)) %>%
  mutate(pretreatment = factor(pretreatment,
                               levels = c("Cold", "Neutral", "Warm"))) %>%
  ggplot(aes(x = week, y = Fit, fill = pretreatment,
             linetype = pretreatment)) +
  # facet_wrap(~sex) +
  geom_ribbon(aes(x = week, ymin = LCL, ymax = UCL),
    colour = NA, size = 0.25, alpha = 0.3
  ) +
  geom_line(colour = "black", alpha = 0.7) +
  stat_summary(
```

```
  data = growthModel$data %>%
    mutate(pretreatment = str_to_title(pretreatment)) %>%
    mutate(pretreatment = factor(pretreatment,
                                 levels = c("Cold", "Neutral", "Warm"))),
  aes(x = week, y = mass),
  geom = "errorbar", fun.data = "mean_se",
  colour = "black", alpha = 0.7, width = 0.25,
  position = position_dodge(width = 0.15)
) +
stat_summary(
  data = growthModel$data %>%
    mutate(pretreatment = str_to_title(pretreatment)) %>%
    mutate(pretreatment = factor(pretreatment,
                                 levels = c("Cold", "Neutral", "Warm"))),
  aes(x = week, y = mass),
  geom = "point", fun = "mean", pch = 21, size = 3,
  colour = "black", alpha = 0.7,
  position = position_dodge(width = 0.15)
) +
theme_classic() +
scale_fill_manual(values = c("#7BB4E3", "black", "#CD5C5C"),
                  name = "Rearing\nConditions",
                  labels = c("Cold (10°C)", "Mild (20°C)", "Warm (30°C)")) +
scale_linetype_manual(values = c("dotted", "solid", "dashed"),
                      name = "Rearing\nConditions",
                  labels = c("Cold (10°C)", "Mild (20°C)", "Warm (30°C)")) +
xlab("Age (weeks)") +
ylab("Body Mass (g)")
```

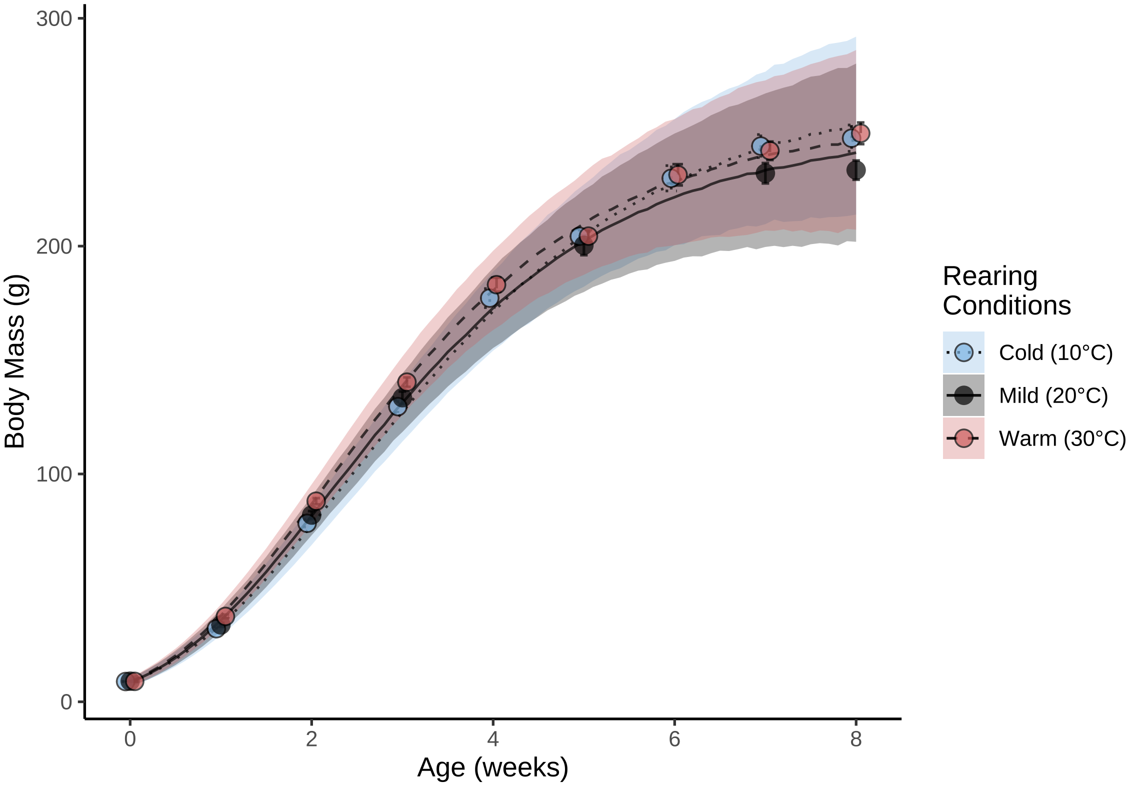


***Figure 17:*** *Body mass (g) growth curves of Japanese quail reared in the cold (10°C), mild conditions (20°C), or the warm (30°C) until at least 3 weeks of age. Dots represent mean values and errorbars represent standard errors around means. Lines represent estimated trends in growth from a Bayesian non-linear model, while ribbons represent confidence around trends (± one standard error).*

```
# No obvious issues. Proceeding with further model checking,
# beginning with assessment of residual distributions.
```

Below, we proceed by checking residual distributions.

```
p1 <- growthModel$data %>%
  mutate("residuals" =
           residuals(growthModel,
                     type = "pearson",
                     robust = TRUE)[, "Estimate"]) %>%
  ggplot(aes(x = residuals)) +
  geom_density(colour = "black", fill = "white") +
  xlab("Pearson Residuals") +
  ylab("Density") +
  theme_classic()

p2 <- growthModel$data %>%
  mutate(
    "residuals" = residuals(growthModel,
      type = "pearson",
      robust = TRUE
    )[, "Estimate"],
    "fitted" = fitted(growthModel,
                      robust = TRUE)[, "Estimate"]
  ) %>%
  ggplot(aes(x = fitted, y = residuals)) +
  geom_point(colour = "black", pch = 21,
             size = 2, fill = "grey75", alpha = 0.5) +
  ylab("Pearson Residuals") +
  xlab("Fitted Values (g)") +
  theme_classic()

p3 <- growthModel$data %>%
  mutate("residuals" = residuals(growthModel,
    type = "pearson",
    robust = TRUE
  )[, "Estimate"]) %>%
  ggplot(aes(x = week, y = residuals)) +
  geom_point(
    colour = "black", pch = 21, size = 2,
    fill = "grey75", alpha = 0.5,
    position = position_jitter(width = 0.25)
  ) +
  stat_summary(geom = "errorbar", fun.data = "mean_se",
               colour = "black", width = 0.25) +
  stat_summary(geom = "point", fun = "mean", pch = 21,
               colour = "black", fill = "white", size = 4) +
  ylab("Pearson Residuals") +
  xlab("Age (weeks)") +
  theme_classic()

p4 <- growthModel$data %>%
  mutate("residuals" = residuals(growthModel,
    type = "pearson",
    robust = TRUE
  )[, "Estimate"]) %>%
  mutate(pretreatment = str_to_title(pretreatment)) %>%
  mutate(pretreatment = ifelse(pretreatment == "Cold", "Cold\n(10°C)",
    ifelse(pretreatment == "Neutral", "Mild\n(20°C)",
      "Warm\n(30°C)"
    )
  )) %>%
  mutate(pretreatment = factor(pretreatment,
                               levels = c("Cold\n(10°C)",
                                          "Mild\n(20°C)",
                                          "Warm\n(30°C)"))
```

```
        ) %>%
  ggplot(aes(x = pretreatment, y = residuals)) +
  geom_point(
    colour = "black", pch = 21, size = 2, fill = "grey75", alpha = 0.5,
    position = position_jitter(width = 0.25)
  ) +
  stat_summary(geom = "errorbar", fun.data = "mean_se",
               colour = "black", width = 0.25) +
  stat_summary(geom = "point", fun = "mean", pch = 21,
               colour = "black", fill = "white", size = 4) +
  ylab("Pearson Residuals") +
  xlab("Rearing Conditions)") +
  theme_classic()

(p1 + p2) / (p3 + p4)
```

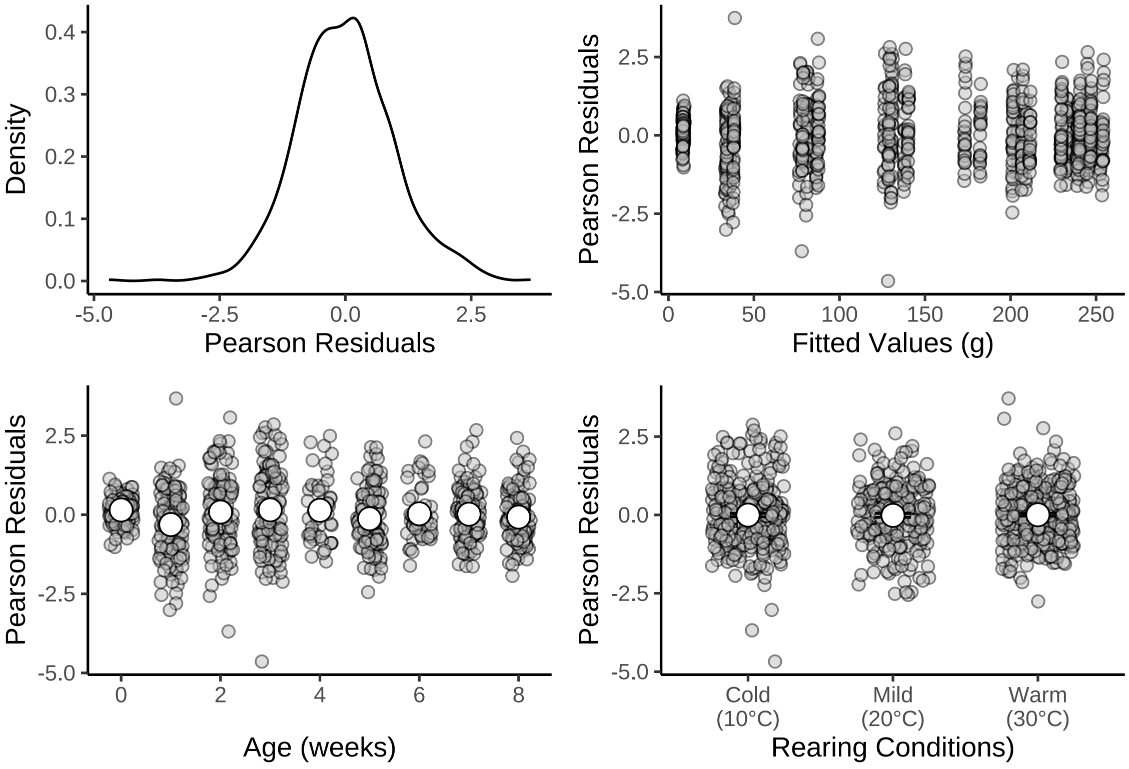


***Figure 18:*** *Density and distributions of Pearson residuals across fitted values and model predictors (including age and rearing conditions). All residuals pertain to those drawn from a Bayesian non-linear model predicting body mass (g) during growth in Japanese quail. Pearson residuals are shown rather than ordinary residuals to correct for the age-dependence of model error.*

```
# No heteroskedasticity and residuals appear normal, albeit large. Plotting model outcomes.
```

And our model outcomes are visualised.

```
as.data.frame(growthModel) %>%
  pivot_longer(everything(), names_to = "Parameter", values_to = "Values") %>%
  filter(grepl("b_|sd_", Parameter)) %>%
  arrange(Parameter) %>%
  merge(., tribble(
    ~Parameter, ~Par,
    "b_A_Intercept", "Beta a0",
    "b_A_pretreatmentcold", "Beta a1\n(Cold-reared)",
```

```
    "b_A_pretreatmentwarm", "Beta a2\n(Warm-reared)",
    "sd_batch__A_Intercept", "Mu 0a",
    "b_B_Intercept", "Beta b0",
    "b_B_pretreatmentcold", "Beta b1\n(Cold-reared)",
    "b_B_pretreatmentwarm", "Beta b2\n(Warm-reared)",
    "sd_batch__B_Intercept", "Mu 0b",
    "b_C_Intercept", "Beta c0",
    "b_C_pretreatmentcold", "Beta c1\n(Cold-reared)",
    "b_C_pretreatmentwarm", "Beta c2\n(Warm-reared)",
    "sd_batch__C_Intercept", "Mu 0c",
    "sd_ring__D_Intercept", "Mu 0\n(Individual Intercept)",
    "b_sigma_intercept", "Tau 0",
    "b_sigma_logweekB", "Tau 1"
  ),
  by = "Parameter", all.x = TRUE
  ) %>%
  ggplot(aes(x = Values)) +
  facet_wrap(~Par, scales = "free") +
  geom_density() +
  geom_vline(xintercept = 0, linetype = "dashed",
             colour = "firebrick4") +
  scale_x_continuous(n.breaks = 3) +
  ylab("Density") +
  theme_classic() +
  theme(axis.title.x = element_blank())
```

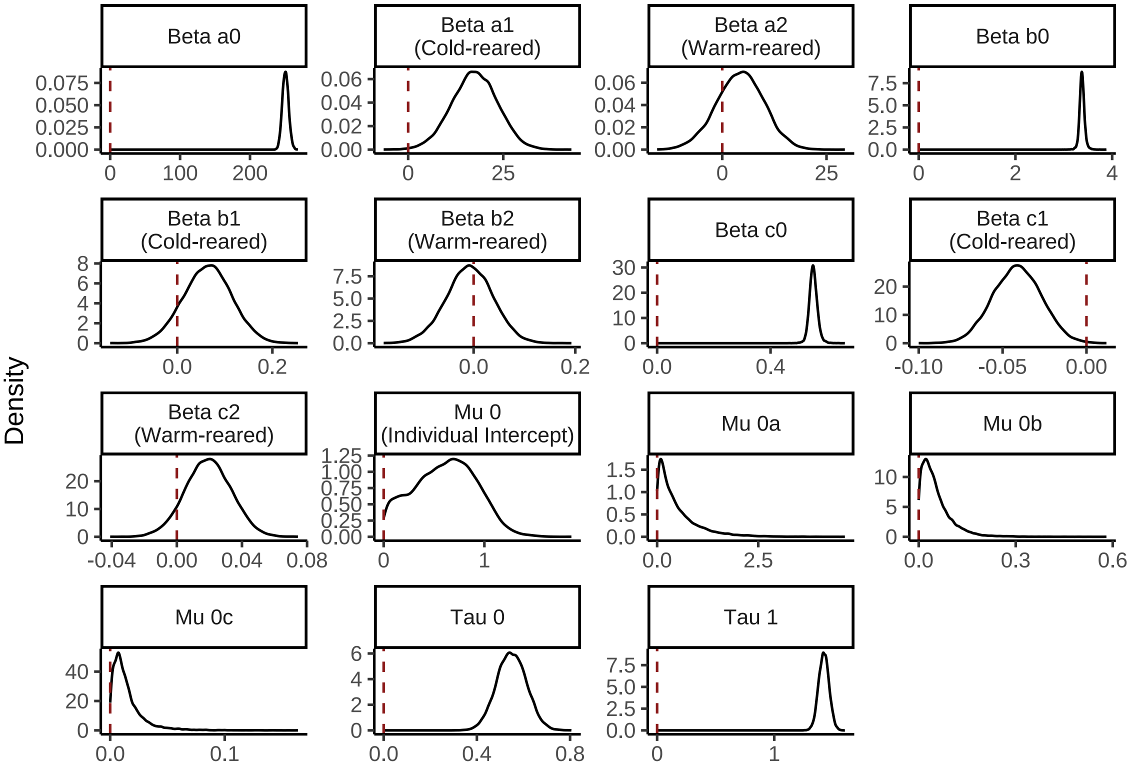


***Figure 19:*** *Density of model coefficients from a Bayesian non-linear model predicting body mass (g) of Japanese quail during growth. Dashed red lines indicate zero values for each coefficient.*

```
# Posterior densities are generally normal, however,
# given that some stray, summarising model coefficients
# by their medians and quantile intervals.

caption <- paste0("Coefficients from a Bayesian non-linear ",
  "effects model predicting body mass of Japanese ",
```

```
  "quail as a Gompertz function of age in weeks. ",
  "Coefficients represent posterior medians and are ",
  "estimated from body mass data collected weekly ",
  "between 0 and 8 weeks of age. Credible intervals ",
  "(CIs) represent quantile intervals."
)

growthModelMassTable <-
  merge(
    as.data.frame(growthModel) %>%
      summarise_all(., .funs = median) %>%
      mutate_all(., .funs = round, 3) %>%
      pivot_longer(everything(), names_to = "Parameter",
                   values_to = "Values") %>%
      filter(grepl("b_|sd_", Parameter)) %>%
      arrange(Parameter),
    quantileCIs(growthModel, cis = c(50, 95)) %>%
    mutate(
    `50\\% CIs` = paste0("[", round(Low_CI_50, digits = 3),
                         ", ", round(High_CI_50, digits = 3), "]"),
    `95\\% CIs` = paste0("[", round(Low_CI_95, digits = 3),
                         ", ", round(High_CI_95, digits = 3), "]")
    ) %>%
      select(Parameter, `50\\% CIs`, `95\\% CIs`),
      by = "Parameter"
  ) %>%
  merge(., tribble(
    ~Parameter, ~Par,
    "b_A_Intercept", "Beta a0",
    "b_A_pretreatmentcold", "Beta a1 Cold-reared)",
    "b_A_pretreatmentwarm", "Beta a2 (Warm-reared)",
    "sd_batch__A_Intercept", "Mu 0a",
    "b_B_Intercept", "Beta b0",
    "b_B_pretreatmentcold", "Beta b1 (Cold-reared)",
    "b_B_pretreatmentwarm", "Beta b2 (Warm-reared)",
    "sd_batch__B_Intercept", "Mu 0b",
    "b_C_Intercept", "Beta c0",
    "b_C_pretreatmentcold", "Beta c1 (Cold-reared)",
    "b_C_pretreatmentwarm", "Beta c2 (Warm-reared)",
    "sd_batch__C_Intercept", "Mu 0c",
    "sd_ring__D_Intercept", "Mu (Individual Intercept)",
    "b_sigma_intercept", "Tau 0",
    "b_sigma_logweekB", "Tau 1"
  ),
  by = "Parameter", all.x = TRUE
  ) %>%
  select(-Parameter) %>%
  select("Parameter" = Par, "Value" = Values,
         `50\\% CIs`, `95\\% CIs`) %>%
  kbl(., longtable = T, booktabs = T, format = "latex",
      caption = caption, escape = FALSE) %>%
  column_spec(column = c(1, 4), width = "2.5cm") %>%
  kable_styling(latex_options = "striped")

growthModelMassTable
```

***Table 4:*** *Coefficients from a Bayesian non-linear effects model predicting body mass of Japanese quail as a Gompertz function of age in weeks. Coefficients represent posterior medians and are estimated from body mass data collected weekly between 0 and 8 weeks of age. Credible intervals (CIs) represent quantile intervals.*

| Parameter | Value | 50% CIs | 95% CIs |
|---|---|---|---|
| Beta a0 | 250.641 | [247.628, 253.685] | [242.164, 259.563] |
| Beta a1 Cold-reared) | 17.521 | [13.457, 21.511] | [5.524, 29.054] |
| Beta a2 (Warm-reared) | 4.399 | [0.493, 8.164] | [-7.023, 15.284] |

| | | | |
|---|---|---|---|
| Beta b0 | 3.375 | [3.345, 3.407] | [3.276, 3.488] |
| Beta b1 (Cold-reared) | 0.064 | [0.03, 0.098] | [-0.037, 0.162] |
| Beta b2 (Warm-reared) | -0.008 | [-0.038, 0.023] | [-0.1, 0.081] |
| Beta c0 | 0.550 | [0.542, 0.56] | [0.524, 0.581] |
| Beta c1 (Cold-reared) | -0.041 | [-0.051, -0.031] | [-0.069, -0.013] |
| Beta c2 (Warm-reared) | 0.019 | [0.01, 0.029] | [-0.009, 0.047] |
| Tau 0 | 0.548 | [0.505, 0.592] | [0.425, 0.674] |
| Tau 1 | 1.423 | [1.393, 1.453] | [1.337, 1.509] |
| Mu 0a | 0.324 | [0.128, 0.675] | [0.011, 1.947] |
| Mu 0b | 0.041 | [0.02, 0.071] | [0.002, 0.176] |
| Mu 0c | 0.011 | [0.006, 0.018] | [0.001, 0.053] |
| Mu (Individual Intercept) | 0.623 | [0.38, 0.835] | [0.043, 1.184] |

```
#save_kable(growthModelMassTable, "../tables/growthModelMassTable.html")
```

From our model, it is evident that cold-reared individuals (i.e. those raised at 10°C for at least their first three weeks of life) have, on average, a higher asymptote (value for *a*) than those of both warm-reared (30°C for at least 3 weeks of life) and mild-reared (constant 20°C) individuals. By contrast, the growth rate of cold-reared individuals is lower than warm-reared and mild-reared individuals alike. To test these distinctions formally, we calculated the ratio of probabilities that one asymptote or growth rate (i.e. from one experimental treatment group) was indeed larger than another, as indicated from our model.

```
caption <- paste0("Coefficients from a Bayesian non-linear ",
                  "effects model predicting body mass of ",
                  "Japanese quail as a Gompertz function of ",
                  "age in weeks. Coefficients represent ",
                  "posterior medians and are estimated from ",
                  "body mass data collected weekly between ",
                  "0 and 8 weeks of age. Credible ",
                  "intervals (CIs) represent quantile ",
                  "intervals around medians."
)

pairwiseTableGrowthMass <- rbind(
  hypothesis(growthModel,
    hypothesis = paste0(
      "(A_Intercept + A_pretreatmentcold) - ",
      "A_Intercept > 0"
    ),
    class = "b",
    robust = TRUE
  )$hypothesis,
  hypothesis(growthModel,
    hypothesis = paste0(
      "A_Intercept - (A_Intercept + A_pretreatmentwarm) > 0 "
    ),
    class = "b",
    robust = TRUE
  )$hypothesis,
  hypothesis(growthModel,
    hypothesis = paste0(
      "(A_Intercept + A_pretreatmentcold) - ",
      "(A_Intercept + A_pretreatmentwarm) > 0"
    ),
    class = "b",
    robust = TRUE
  )$hypothesis,
  hypothesis(growthModel,
    hypothesis = paste0(
```

```
      "(B_Intercept + B_pretreatmentcold) - ",
      "B_Intercept > 0"
    ),
    class = "b",
    robust = TRUE
  )$hypothesis,
  hypothesis(growthModel,
    hypothesis = paste0(
      "(B_Intercept + B_pretreatmentwarm) - ",
      "B_Intercept > 0"
    ),
    class = "b",
    robust = TRUE
  )$hypothesis,
  hypothesis(growthModel,
    hypothesis = paste0(
      "(B_Intercept + B_pretreatmentcold) - ",
      "(B_Intercept + B_pretreatmentwarm) > 0"
    ),
    class = "b",
    robust = TRUE
  )$hypothesis,
  hypothesis(growthModel,
    hypothesis = paste0(
      "C_Intercept - (C_Intercept + C_pretreatmentcold) > 0"
    ),
    class = "b",
    robust = TRUE
  )$hypothesis,
  hypothesis(growthModel,
    hypothesis = paste0(
      "(C_Intercept + C_pretreatmentwarm) - C_Intercept > 0"
    ),
    class = "b",
    robust = TRUE
  )$hypothesis,
  hypothesis(growthModel,
    hypothesis = paste0(
      "(C_Intercept + C_pretreatmentwarm) - ",
      "(C_Intercept + C_pretreatmentcold) > 0"
    ),
    class = "b",
    robust = TRUE
  )$hypothesis
) %>%
  merge(.,
    rbind(
      hypothesis(growthModel,
        hypothesis = paste0(
          "(A_Intercept + A_pretreatmentcold) - ",
          "A_Intercept > 0"
        ),
        class = "b", alpha = 0.20,
        robust = TRUE
      )$hypothesis,
      hypothesis(growthModel,
        hypothesis = paste0(
          "A_Intercept - (A_Intercept + A_pretreatmentwarm) > 0 "
        ),
        class = "b", alpha = 0.20,
        robust = TRUE
      )$hypothesis,
      hypothesis(growthModel,
        hypothesis = paste0(
          "(A_Intercept + A_pretreatmentcold) - ",
          "(A_Intercept + A_pretreatmentwarm) > 0"
        ),
```

```
        class = "b", alpha = 0.20,
        robust = TRUE
      )$hypothesis,
      hypothesis(growthModel,
        hypothesis = paste0(
          "(B_Intercept + B_pretreatmentcold) - ",
          "B_Intercept > 0"
        ),
        class = "b", alpha = 0.20,
        robust = TRUE
      )$hypothesis,
      hypothesis(growthModel,
        hypothesis = paste0(
          "(B_Intercept + B_pretreatmentwarm) - ",
          "B_Intercept > 0"
        ),
        class = "b", alpha = 0.20,
        robust = TRUE
      )$hypothesis,
      hypothesis(growthModel,
        hypothesis = paste0(
          "(B_Intercept + B_pretreatmentcold) - ",
          "(B_Intercept + B_pretreatmentwarm) > 0"
        ),
        class = "b", alpha = 0.20,
        robust = TRUE
      )$hypothesis,
      hypothesis(growthModel,
        hypothesis = paste0(
          "C_Intercept - (C_Intercept + C_pretreatmentcold) > 0"
        ),
        class = "b", alpha = 0.20,
        robust = TRUE
      )$hypothesis,
      hypothesis(growthModel,
        hypothesis = paste0(
          "(C_Intercept + C_pretreatmentwarm) - C_Intercept > 0"
        ),
        class = "b", alpha = 0.20,
        robust = TRUE
      )$hypothesis,
      hypothesis(growthModel,
        hypothesis = paste0(
          "(C_Intercept + C_pretreatmentwarm) - ",
          "(C_Intercept + C_pretreatmentcold) > 0"
        ),
        class = "b", alpha = 0.20,
        robust = TRUE
      )$hypothesis
    ) %>% mutate(`50\\% CIs` = paste0(
      "[",
      round(CI.Lower, digits = 3),
      ", ",
      round(CI.Upper, digits = 3),
      "]"
    )) %>%
      select(Hypothesis, `50\\% CIs`),
    by = "Hypothesis"
  ) %>%
  merge(.,
    tribble(
      ~hypothesis, ~Hypothesis,
      "Cold-reared Asymptote (a) > Mild-reared Asymptote (a)",
      "((A_Intercept+A_pretreatmentcold)-A_Intercept) > 0",
      "Warm-reared Asymptote (a) < Mild-reared Asymptote (a)",
      "(A_Intercept-(A_Intercept+A_pretreatmentwarm)) > 0",
      "Warm-reared Asymptote (a) < Cold-reared Asymptote (a)",
```

```
      paste0(
        "((A_Intercept+A_pretreatmentcold)",
        "-(A_Intercept+A_pretreatmentwarm)) > 0"
      ),
      "Cold-reared Displacement (b) > Mild-reared Displacement (b)",
      "((B_Intercept+B_pretreatmentcold)-B_Intercept) > 0",
      "Warm-reared Displacement (b) > Mild-reared Displacement (b)",
      "((B_Intercept+B_pretreatmentwarm)-B_Intercept) > 0",
      "Warm-reared Displacement (b) > Cold-reared Displacement (b)",
      paste0(
        "((B_Intercept+B_pretreatmentcold)",
        "-(B_Intercept+B_pretreatmentwarm)) > 0"
      ),
      "Cold-reared Growth Rate (c) < Mild-reared Growth Rate (c)",
      "(C_Intercept-(C_Intercept+C_pretreatmentcold)) > 0",
      "Warm-reared Growth Rate (c) > Mild-reared Growth Rate (c)",
      "((C_Intercept+C_pretreatmentwarm)-C_Intercept) > 0",
      "Warm-reared Growth Rate (c) > Cold-reared Growth Rate (c)",
      paste0(
        "((C_Intercept+C_pretreatmentwarm)",
        "-(C_Intercept+C_pretreatmentcold)) > 0"
      )
    ),
    by = "Hypothesis"
  ) %>%
  mutate(`95\\% CIs` = paste0(
    "[", round(CI.Lower, digits = 3), ", ",
    round(CI.Upper, digits = 3), "]"
  )) %>%
  select(
    "Hypothesis" = hypothesis, "Delta" = Estimate,
    `50\\% CIs`, `95\\% CIs`, "Evidence Ratio" = Evid.Ratio
  ) %>%
  kbl(., longtable = T, booktabs = T, format = "latex",
      caption = caption, escape = FALSE
  ) %>%
  column_spec(column = c(1:10), width = "2.5cm") %>%
  kable_styling(latex_options = "striped")

pairwiseTableGrowthMass
```

***Table 5:*** *Coefficients from a Bayesian non-linear effects model predicting body mass of Japanese quail as a Gompertz function of age in weeks. Coefficients represent posterior medians and are estimated from body mass data collected weekly between 0 and 8 weeks of age. Credible intervals (CIs) represent quantile intervals around medians.*

| Hypothesis | Delta | 50% CIs | 95% CIs | Evidence Ratio |
|---|---|---|---|---|
| Warm-reared Asymptote (a) < Cold-reared Asymptote (a) | 13.2504503 | [8.51, 17.781] | [4.071, 22.297] | 113.2857143 |
| Cold-reared Asymptote (a) > Mild-reared Asymptote (a) | 17.5206766 | [12.414, 22.527] | [7.579, 27.25] | 469.5882353 |
| Warm-reared Displacement (b) > Cold-reared Displacement (b) | 0.0719893 | [0.033, 0.11] | [-0.003, 0.145] | 16.3535792 |
| Cold-reared Displacement (b) > Mild-reared Displacement (b) | 0.0642949 | [0.021, 0.106] | [-0.021, 0.147] | 8.3731693 |

| | | | | |
|---|---|---|---|---|
| Warm-reared Displacement (b) > Mild-reared Displacement (b) | -0.0077255 | [-0.046, 0.031] | [-0.085, 0.067] | 0.7654198 |
| Warm-reared Growth Rate (c) > Cold-reared Growth Rate (c) | 0.0606325 | [0.049, 0.072] | [0.039, 0.082] | Inf |
| Warm-reared Growth Rate (c) > Mild-reared Growth Rate (c) | 0.0193074 | [0.007, 0.031] | [-0.004, 0.043] | 10.5025162 |
| Warm-reared Asymptote (a) < Mild-reared Asymptote (a) | -4.3991811 | [-9.059, 0.472] | [-13.447, 5.26] | 0.2875191 |
| Cold-reared Growth Rate (c) < Mild-reared Growth Rate (c) | 0.0412728 | [0.029, 0.054] | [0.018, 0.065] | 515.1290323 |

```
#save_kable(pairwiseTableGrowthMass,
# "../tables/pairwiseTableGrowthMass.html")
```

Distinctions in body mass among treatments at 8 weeks of age were evaluated using a Bayesian one-way ANOVA with priors on body mass for our mild treatment being normally distributed with a mean of 250 and standard deviation of 25. Priors for our warm- and cold-reared treatments were assigned means of 242.5 and 257.5 respectively (again, as per Burness et al, 2013), and standard deviations held liberally at 25. Posterior probabilities for differences *between* treatments were calculated using the Savage-Dickey density ratio method.

```
growthAnova <- brm(
  data = data %>%
    filter(week == 8) %>%
    select(pretreatment, mass, "batch" = exp) %>%
    drop_na() %>%
    mutate(pretreatment =
             factor(pretreatment,
                    levels = c("neutral", "cold", "warm"))),
  formula = mass ~ 0 + pretreatment + (1|batch),
  prior = c(
    set_prior("normal(250, 25)", class = "b",
              coef = "pretreatmentneutral"),
    set_prior("normal(257.5, 25)", class = "b",
              coef = "pretreatmentcold"),
    set_prior("normal(242.5, 25)", class = "b",
              coef = "pretreatmentwarm"),
    set_prior("exponential(2.5)", class = "sd",
              group = "batch")
  ),
  family = "gaussian",
  seed = 200,
  cores = 4, chains = 4,
  iter = 50000, warmup = 10000, thin = 10,
  control = list(adapt_delta = 0.95, max_treedepth = 13),
  silent = TRUE, refresh = 0,
  file = "./models/massGrowthAnova.Rds"
)

hypotheses <- c(
  "pretreatmentcold - pretreatmentneutral > 0",
  "pretreatmentwarm - pretreatmentneutral > 0",
  "pretreatmentcold - pretreatmentwarm > 0"
)
```

```
caption <- paste0("Results from a Bayesian, one-way ANOVA comparing ",
                  "body mass at 8 weeks of Japanese quail reared in the ",
                  "cold (10°C until at least 3 weeks of age; n = ",
                  nrow(subset(growthAnova$data, pretreatment == "cold")),
                  "), mild temperature (constant 20°C; n = ",
                  nrow(subset(growthAnova$data, pretreatment == "neutral")),
                  "), or warmth (30°C until at least 3 weeks of age; n = ",
                  nrow(subset(growthAnova$data, pretreatment == "warm")), ")."
                  )

growthMassAnovaTable <- bind_rows(
  lapply(hypotheses, FUN = function(x) {
    hold <- hypothesis(growthAnova,
                       hypothesis = x, class = "b",
                       alpha = 0.5, robust = TRUE)
    return(data.frame(
      "hyp" = x,
      "deltaMass" = hold$hypothesis$Estimate,
      "pProb" = hold$hypothesis$Post.Prob
    ))
  })
) %>%
  mutate("Hypothesis" = c(
    "Cold-Reared Mass > Mild-Reared Mass",
    "Warm-Reared Mass > Mild-Reared Mass",
    "Cold-Reared Mass > Warm-Reared Mass"
  )) %>%
  select(Hypothesis, "Delta Mass (g)" = deltaMass,
         "Posterior Probability" = pProb) %>%
  kbl(.,
    longtable = T, booktabs = T,
    caption = caption
  ) %>%
  kable_styling(latex_options = "striped")

growthMassAnovaTable
```

***Table 6:*** *Results from a Bayesian, one-way ANOVA comparing body mass at 8 weeks of Japanese quail reared in the cold (10°C until at least 3 weeks of age; n = 42), mild temperature (constant 20°C; n = 36), or warmth (30°C until at least 3 weeks of age; n = 43).*

| Hypothesis | Delta Mass (g) | Posterior Probability |
|---|---|---|
| Cold-Reared Mass > Mild-Reared Mass | 13.556696 | 0.9673125 |
| Warm-Reared Mass > Mild-Reared Mass | 15.072196 | 0.9818750 |
| Cold-Reared Mass > Warm-Reared Mass | -1.589709 | 0.4123750 |

```
#save_kable(growthMassAnovaTable, "../tables/growthMassAnovaTable.html")

# Convincing evidence that cold and warm reared birds are
# heavier at 8 weeks of age than mild-reared birds, but
# not that cold- and warm-reared birds differ in mass
# (~61% probability that warm-reared birds are largest).
```

Thermal rearing conditions in our study varied in duration, with some warmth (30°C) and cold (10°C) exposures lasting between hatch to three weeks of age and some lasting between hatch and maturity (eight weeks of age). To test whether longer exposure to thermal environments during development differentially shaped growth patterns relative to more transient exposures (3 weeks), we repeated our above analysis but while only including individuals who experience continuous cold (10°C), or warmth (30°C) treatments until maturity. Priors for these analyses remained identical to those used previously, however, effect of egg batch was removed from our model given that all individuals in this analysis were derived from the same batch.

```
initFunction <- function(chain_id = 1) {
  list(
```

```
    "b_A" = c(rnorm(1, 250, 25), -14),
    "b_B" = c(rskew_normal(1, xi = 2.5, omega = 2, alpha = 4), 0),
    "b_C" = c(rbeta(1, 5, 5), 0),
    "b_sigma" = c(rskew_normal(1, xi = 1, omega = 0.5, alpha = -5), 1),
    "sd_1" = rexp(2, 0.5)
  )
}

# Loading initial values into a list.

initList <- lapply(1:4, initFunction)

growthModelStrict <- brm(
  data = data %>%
    filter(week <= 8 & exp == "C") %>%
    mutate(pretreatment = factor(pretreatment,
      levels = c("cold", "warm")
    )) %>%
    select(ring, pretreatment, week, mass) %>%
    distinct() %>%
    mutate(weekB = week + 1),
  formula = bf(mass ~ A * exp(-B * exp(-C * week)) + D,
    A ~ 1 + pretreatment,
    B ~ 1 + pretreatment,
    C ~ 1 + pretreatment,
    D ~ 0 + (1 | ring),
    sigma ~ 0 + intercept + log(weekB),
    nl = TRUE
  ),
  prior = c(
    set_prior("normal(250, 25)",
      class = "b", coef = "Intercept",
      nlpar = "A"
    ),
    set_prior("normal(-14, 25)",
      class = "b", coef = "pretreatmentwarm",
      nlpar = "A"
    ),
    set_prior("normal(3, 1)",
      class = "b", coef = "Intercept",
      nlpar = "B"
    ),
    set_prior("normal(0, 0.5)",
      class = "b", coef = "pretreatmentwarm",
      nlpar = "B"
    ),
    set_prior("skew_normal(0.5, 0.1, 2.5)",
      class = "b", coef = "Intercept",
      nlpar = "C"
    ),
    set_prior("normal(0, 0.2)",
      class = "b", coef = "pretreatmentwarm",
      nlpar = "C"
    ),
    set_prior("exponential(0.5)", class = "sd",
              coef = "Intercept", group = "ring", nlpar = "D"),
    set_prior("skew_normal(1, 0.5, -10)", class = "b",
              coef = "intercept", dpar = "sigma"),
    set_prior("skew_normal(1, 0.5, 10)", class = "b",
              coef = "logweekB", dpar = "sigma")
  ),
  family = "gaussian",
  init = initList,
  seed = 200,
  cores = 4, chains = 4,
  iter = 50000, warmup = 10000, thin = 10,
  control = list(adapt_delta = 0.95, max_treedepth = 13),
```

```
  silent = TRUE, refresh = 0,
  file = "./models/growthModelMassStrict.Rds"
)

ggarrange(
  ggplot(data = data.frame("Rhat" = brms::rhat(growthModelStrict)),
         aes(x = Rhat)) +
    geom_density() +
    theme_classic() +
    xlab(
      TeX('$\\hat{R}$')
    ) +
    ylab("Density"),
  ggplot(data = data.frame("Neff" = brms::neff_ratio(growthModelStrict)),
         aes(x = Neff)) +
    geom_density() +
    theme_classic() +
    xlab(
      TeX('$N_{eff}/N~Ratio$')
      ) +
    ylab("Density")
)
```

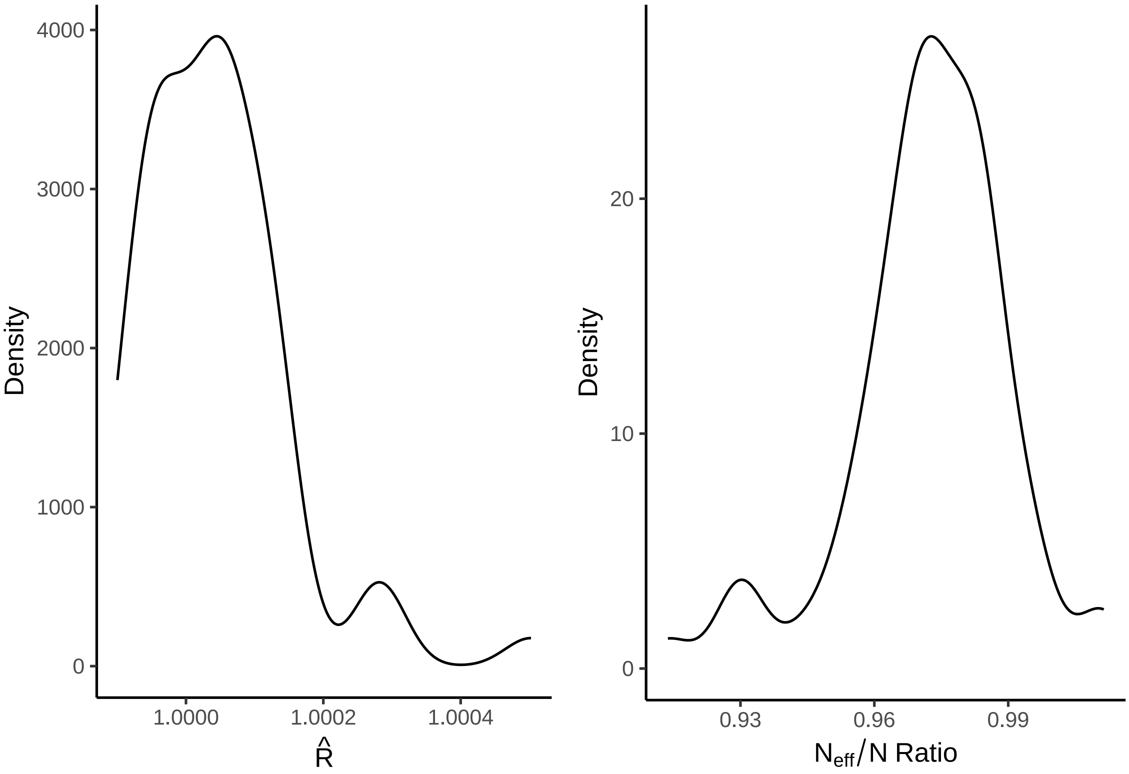


***Figure 20:*** *Gelman-Rubin statistics ($\hat{R}$) and ratio of effective samples sizes by samples sizes per parameter from a Bayesian non-linear model estimating growth of mass (g) among Japanese quail. Here, our sample population is restricted to include only quail reared in cold (10°C), mild (20°C) or warm (30°C) conditions until maturity (8 weeks).*

```
# Good. Quickly checking residuals.
```

```
p1 <- growthModelStrict$data %>%
  mutate("residuals" =
           residuals(growthModelStrict,
                     type = "pearson",
```

```
                      robust = TRUE)[, "Estimate"]) %>%
  ggplot(aes(x = residuals)) +
  geom_density(colour = "black", fill = "white") +
  xlab("Pearson Residuals") +
  ylab("Density") +
  theme_classic()

p2 <- growthModelStrict$data %>%
  mutate(
    "residuals" = residuals(growthModelStrict,
      type = "pearson",
      robust = TRUE
    )[, "Estimate"],
    "fitted" = fitted(growthModelStrict,
                       robust = TRUE)[, "Estimate"]
  ) %>%
  ggplot(aes(x = fitted, y = residuals)) +
  geom_point(colour = "black", pch = 21,
             size = 2, fill = "grey75", alpha = 0.5) +
  ylab("Pearson Residuals") +
  xlab("Fitted Values (g)") +
  theme_classic()

p3 <- growthModelStrict$data %>%
  mutate("residuals" = residuals(growthModelStrict,
    type = "pearson",
    robust = TRUE
  )[, "Estimate"]) %>%
  ggplot(aes(x = week, y = residuals)) +
  geom_point(
    colour = "black", pch = 21, size = 2,
    fill = "grey75", alpha = 0.5,
    position = position_jitter(width = 0.25)
  ) +
  stat_summary(geom = "errorbar", fun.data = "mean_se",
               colour = "black", width = 0.25) +
  stat_summary(geom = "point", fun = "mean", pch = 21,
               colour = "black", fill = "white", size = 4) +
  ylab("Pearson Residuals") +
  xlab("Age (weeks)") +
  theme_classic()

p4 <- growthModelStrict$data %>%
  mutate("residuals" = residuals(growthModelStrict,
    type = "pearson",
    robust = TRUE
  )[, "Estimate"]) %>%
  mutate(pretreatment = str_to_title(pretreatment)) %>%
  mutate(pretreatment = ifelse(pretreatment == "Cold", "Cold\n(10°C)",
      "Warm\n(30°C)"
    )
  ) %>%
  mutate(pretreatment = factor(pretreatment,
                               levels = c("Cold\n(10°C)",
                                          "Warm\n(30°C)"))
         ) %>%
  ggplot(aes(x = pretreatment, y = residuals)) +
  geom_point(
    colour = "black", pch = 21, size = 2, fill = "grey75", alpha = 0.5,
    position = position_jitter(width = 0.25)
  ) +
  stat_summary(geom = "errorbar", fun.data = "mean_se",
               colour = "black", width = 0.25) +
  stat_summary(geom = "point", fun = "mean", pch = 21,
               colour = "black", fill = "white", size = 4) +
  ylab("Pearson Residuals") +
  xlab("Rearing Conditions)") +
```

```
  theme_classic()

(p1 + p2) / (p3 + p4)
```

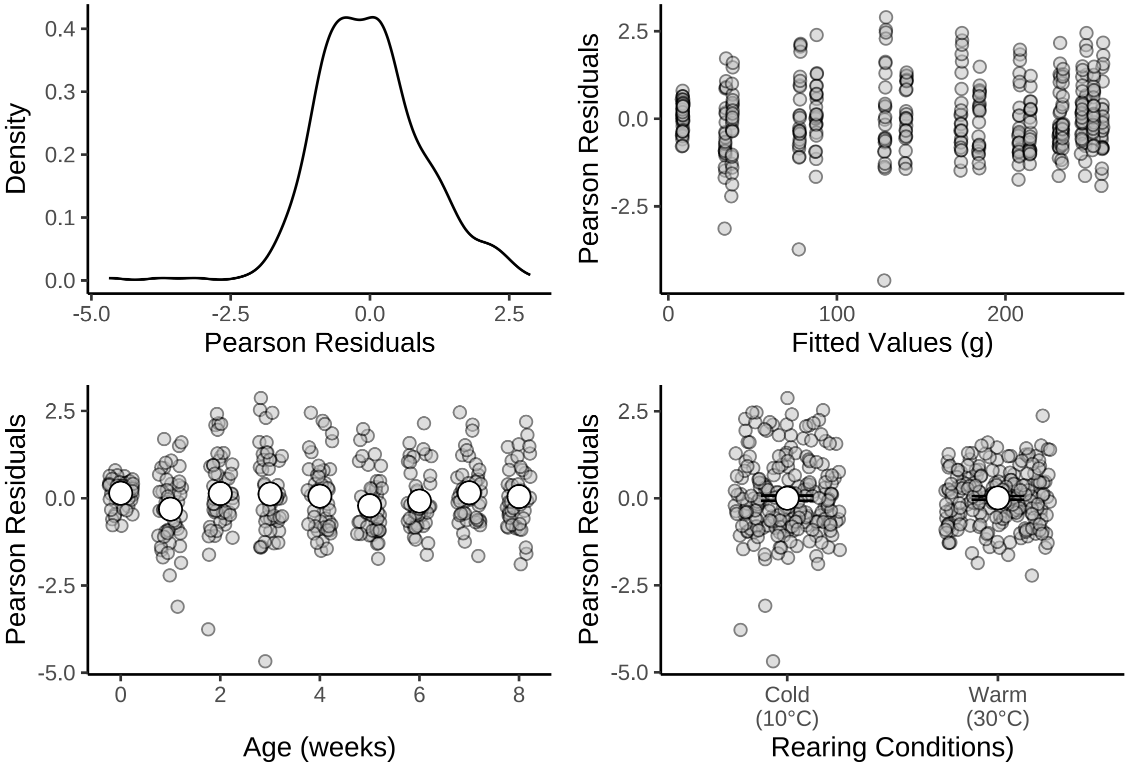


***Figure 21:*** *Density and distributions of Pearson residuals across fitted values and model predictors (including age and rearing conditions). All residuals pertain to those drawn from a Bayesian non-linear model predicting body mass (g) during growth in Japanese quail. Pearson residuals are shown rather than ordinary residuals to correct for the age-dependence of model error. Here, our sample population is restricted to include only quail reared in cold (10°C) or warm (30°C) conditions until maturity (8 weeks).*

```
# Homoskedastic.
# Summarising sample sizes

caption <- paste0("Number of body mass measurements (samples) ",
                  "drawn from Japanese quail between 0 and ",
                  "8 weeks of age (maturity) across three distict ",
                  "thermal rearing conditions.")

growthModelStrict$data %>%
  group_by(week, pretreatment) %>%
  count(name = "Samples (n)") %>%
  mutate(pretreatment = ifelse(pretreatment == "cold",
                               "Cold (10°C)", "Warm (30°C)"
                               )
         ) %>%
  arrange(pretreatment, week) %>%
  rename("Age (Weeks)" = week,
         "Rearing Conditions" = pretreatment) %>%
  kbl(.,
    longtable = T, booktabs = T,
    caption = caption
  ) %>%
  kable_styling(latex_options = "striped")
```

***Table 7:*** *Number of body mass measurements (samples) drawn from Japanese quail between 0 and 8 weeks of age (maturity) across three distict thermal rearing conditions.*

| Age (Weeks) | Rearing Conditions | Samples (n) |
|---|---|---|
| 0 | Cold (10°C) | 24 |
| 1 | Cold (10°C) | 25 |
| 2 | Cold (10°C) | 25 |
| 3 | Cold (10°C) | 25 |
| 4 | Cold (10°C) | 24 |
| 5 | Cold (10°C) | 24 |
| 6 | Cold (10°C) | 24 |
| 7 | Cold (10°C) | 23 |
| 8 | Cold (10°C) | 24 |
| 0 | Warm (30°C) | 24 |
| 1 | Warm (30°C) | 24 |
| 2 | Warm (30°C) | 24 |
| 3 | Warm (30°C) | 24 |
| 4 | Warm (30°C) | 23 |
| 5 | Warm (30°C) | 23 |
| 6 | Warm (30°C) | 23 |
| 7 | Warm (30°C) | 23 |
| 8 | Warm (30°C) | 23 |

Our predicted growth curve is then replotted.

```
showtext_auto()

growthCurveStrict <- with(
  growthModelStrict$data,
  expand.grid(
    "week" = seq(0, 8, by = 0.1),
    "pretreatment" = c("cold", "warm")
  )
) %>%
  mutate("weekB" = week + 1) %>%
  mutate(
    "Fit" = predict(growthModelStrict, re_form = NA,
                    newdata = .,
                    robust = TRUE)[, "Estimate"],
    "SE" = predict(growthModelStrict, re_form = NA,
                   newdata = .,
                   robust = TRUE)[, "Est.Error"]
  ) %>%
  mutate(
    "LCL" = Fit - SE,
    "UCL" = Fit + SE
  ) %>%
  mutate(pretreatment = str_to_title(pretreatment)) %>%
  mutate(pretreatment = factor(pretreatment,
                               levels = c("Cold", "Warm"))) %>%
  ggplot(aes(x = week, y = Fit, fill = pretreatment,
             linetype = pretreatment)) +
  # facet_wrap(~sex) +
  geom_ribbon(aes(x = week, ymin = LCL, ymax = UCL),
    colour = NA, size = 0.25, alpha = 0.3
  ) +
  geom_line(colour = "black", alpha = 0.7) +
  stat_summary(
    data = growthModelStrict$data %>%
      mutate(pretreatment = str_to_title(pretreatment)) %>%
      mutate(pretreatment = factor(pretreatment,
                                   levels = c("Cold", "Warm"))),
    aes(x = week, y = mass),
```

```
    geom = "errorbar", fun.data = "mean_se",
    colour = "black", alpha = 0.7, width = 0.25,
    position = position_dodge(width = 0.15)
  ) +
  stat_summary(
    data = growthModelStrict$data %>%
      mutate(pretreatment = str_to_title(pretreatment)) %>%
      mutate(pretreatment = factor(pretreatment,
                                   levels = c("Cold", "Warm"))),
    aes(x = week, y = mass),
    geom = "point", fun = "mean", pch = 21, size = 3,
    colour = "black", alpha = 0.7,
    position = position_dodge(width = 0.15)
  ) +
  theme_classic() +
  scale_fill_manual(values = c("#7BB4E3", "#CD5C5C"),
                    name = "Rearing\nConditions",
                    labels = c("Cold (10°C)",
                               "Warm (30°C)")) +
  scale_linetype_manual(values = c("solid", "dashed"),
                        name = "Rearing\nConditions",
                    labels = c("Cold (10°C)",
                               "Warm (30°C)")) +
  xlab("Age (weeks)") +
  ylab("Body Mass (g)")

growthCurveStrict
```

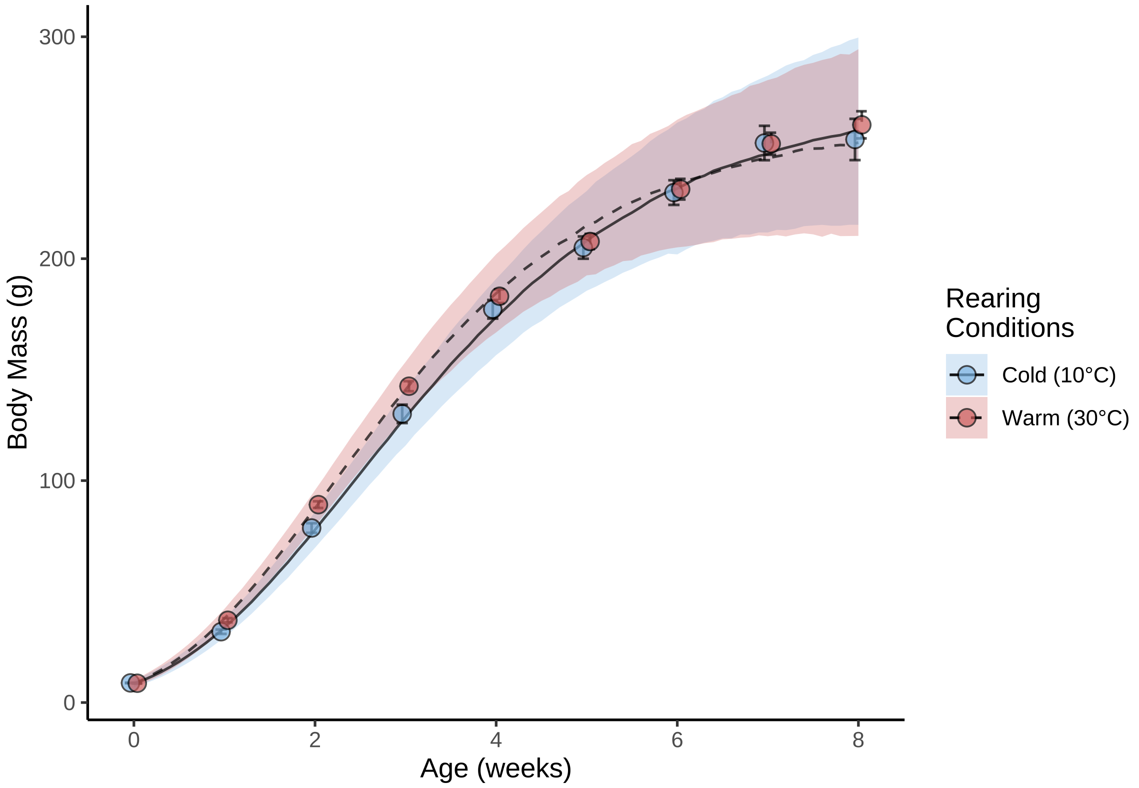


***Figure 22:*** *Body mass (g) growth curves of Japanese quail reared in the cold (10°C), or the warmth (30°C) until at eight weeks of age. Dots represent mean values and errorbars represent standard errors around means. Lines represent estimated trends in growth from a Bayesian non-linear model, while ribbons represent confidence around trends (± one standard error).*

```
ggsave("../plots/strictGrowthCurve.jpg", dpi = 800,
       width = 7, height = 6.5,
       growthCurveStrict)
showtext_auto(enable = "FALSE")

# Summarising growth curve parameters

caption <- paste0("Coefficients from a Bayesian non-linear effects model ",
                  "predicting body mass of Japanese quail as a Gompertz ",
                  "function of age in weeks. Coefficients represent posterior ",
                  "medians and are estimated from body mass data collected ",
                  "weekly between 0 and 8 weeks of age. Credible intervals ",
                  "(CIs) represent quantile intervals around medians."
)

growthModelStrictMassTable <-
  merge(
    as.data.frame(growthModelStrict) %>%
      summarise_all(., .funs = median) %>%
      mutate_all(., .funs = round, 3) %>%
      pivot_longer(everything(), names_to = "Parameter",
                   values_to = "Values") %>%
      filter(grepl("b_|sd_", Parameter)) %>%
      arrange(Parameter),
    quantileCIs(growthModelStrict, cis = c(50, 95)) %>%
    mutate(
    `50\\% CIs` = paste0("[", round(Low_CI_50, digits = 3),
                         ", ", round(High_CI_50, digits = 3), "]"),
    `95\\% CIs` = paste0("[", round(Low_CI_95, digits = 3),
                         ", ", round(High_CI_95, digits = 3), "]")
    ) %>%
      select(Parameter, `50\\% CIs`, `95\\% CIs`),
    by = "Parameter"
  ) %>%
  merge(., tribble(
    ~Parameter, ~Par,
    "b_A_Intercept", "Beta a0 (Cold-reared)",
    "b_A_pretreatmentwarm", "Beta a1 (Warm-reared)",
    "b_B_Intercept", "Beta b0 (Cold-reared)",
    "b_B_pretreatmentwarm", "Beta b1 (Warm-reared)",
    "b_C_Intercept", "Beta c0 (Cold-reared)",
    "b_C_pretreatmentwarm", "Beta c1 (Warm-reared)",
    "sd_ring__D_Intercept", "Mu (Individual Intercept)",
    "b_sigma_intercept", "Tau 0",
    "b_sigma_logweekB", "Tau 1"
  ),
  by = "Parameter", all.x = TRUE
  ) %>%
  select(-Parameter) %>%
  select("Parameter" = Par, "Value" = Values, `50\\% CIs`,`95\\% CIs`) %>%
  kbl(., longtable = T, booktabs = T, format = "latex",
      caption = caption, escape = FALSE) %>%
  column_spec(column = c(1, 4), width = "2.5cm") %>%
  kable_styling(latex_options = "striped")

growthModelStrictMassTable
```

***Table 8:*** *Coefficients from a Bayesian non-linear effects model predicting body mass of Japanese quail as a Gompertz function of age in weeks. Coefficients represent posterior medians and are estimated from body mass data collected weekly between 0 and 8 weeks of age. Credible intervals (CIs) represent quantile intervals around medians.*

| Parameter | Value | 50% CIs | 95% CIs |
|---|---|---|---|
| Beta a0 (Cold-reared) | 273.063 | [269.478, 276.567] | [263.04, 283.521] |

| | | | |
|---|---|---|---|
| Beta a1 (Warm-reared) | -11.627 | [-16.325, -6.913] | [-25.128, 1.589] |
| Beta b0 (Cold-reared) | 3.479 | [3.455, 3.504] | [3.41, 3.556] |
| Beta b1 (Warm-reared) | -0.063 | [-0.096, -0.029] | [-0.166, 0.035] |
| Beta c0 (Cold-reared) | 0.510 | [0.503, 0.517] | [0.49, 0.531] |
| Beta c1 (Warm-reared) | 0.060 | [0.05, 0.071] | [0.03, 0.09] |
| Tau 0 | 0.420 | [0.354, 0.489] | [0.227, 0.619] |
| Tau 1 | 1.509 | [1.465, 1.553] | [1.38, 1.638] |
| Mu (Individual Intercept) | 0.617 | [0.338, 0.878] | [0.036, 1.355] |

And again, growth curve parameters are formally compared between rearing conditions groups.

```
caption <- paste0("Coefficients from a Bayesian non-linear ",
                  "effects model predicting body mass of Japanese quail as a ",
                  "Gompertz function of age in weeks. Coefficients represent ",
                  "posterior medians and are estimated from body mass data ",
                  "collected weekly between 0 and 8 weeks of age. Credible ",
                  "intervals (CIs) represent quantile intervals. ",
                  "Rearing conditions persisted from ",
                  "hatch to maturity (week 8)."
)

pairwiseTableGrowthMassStrict <- rbind(
  hypothesis(growthModelStrict,
             hypothesis = "A_Intercept - (A_Intercept + A_pretreatmentwarm) > 0",
             class = "b",
             robust = TRUE)$hypothesis,
  hypothesis(growthModelStrict,
             hypothesis = "B_Intercept - (B_Intercept + B_pretreatmentwarm) > 0",
             class = "b",
             robust = TRUE)$hypothesis,
  hypothesis(growthModelStrict,
             hypothesis = "(C_Intercept + C_pretreatmentwarm) - C_Intercept > 0",
             class = "b",
             robust = TRUE)$hypothesis
) %>%
  cbind(., data.frame("hypothesis" = c(
    "Cold-reared Asymptote (a) > Warm-reared Asymptote (a)",
    "Cold-reared Displacement (b) > Warm-reared Displacement (b)",
    "Cold-reared Growth Rate (c) < Warm-reared Growth Rate (c)"
  ))) %>%
  mutate(`95% CIs` = paste0("[", round(CI.Lower, digits = 3), ", ",
                            round(CI.Upper, digits = 3), "]"),
         Estimate = round(Estimate, digits = 3),
         Evid.Ratio = round(Evid.Ratio, digits = 3)) %>%
  select("Hypothesis" = hypothesis, "Delta" = Estimate,
         `95% CIs`, "Evidence Ratio" = Evid.Ratio) %>%
  kbl(., longtable = T, booktabs = T, format = "latex", caption = caption) %>%
  column_spec(column = c(1:10), width = "2.5cm")  %>%
  kable_styling(latex_options = "striped")

pairwiseTableGrowthMassStrict
```

***Table 9:*** *Coefficients from a Bayesian non-linear effects model predicting body mass of Japanese quail as a Gompertz function of age in weeks. Coefficients represent posterior medians and are estimated from body mass data collected weekly between 0 and 8 weeks of age. Credible intervals (CIs) represent quantile intervals. Rearing conditions persisted from hatch to maturity (week 8).*

| Hypothesis | Delta | 95% CIs | Evidence Ratio |
|---|---|---|---|

| | | | |
|---|---|---|---|
| Cold-reared Asymptote (a) > Warm-reared Asymptote (a) | 11.627 | [0.451, 22.91] | 22.222 |
| Cold-reared Displacement (b) > Warm-reared Displacement (b) | 0.063 | [-0.02, 0.149] | 8.434 |
| Cold-reared Growth Rate (c) < Warm-reared Growth Rate (c) | 0.060 | [0.035, 0.086] | 7999.000 |

```
#save_kable(pairwiseTableGrowthMassStrict,
# "../tables/pairwiseTableGrowthMassStrict")
```

```
growthAnovaStrict <- brm(
  data = data %>%
    filter(week == 8 & exp == "C") %>%
    mutate(pretreatment = factor(pretreatment,
      levels = c("cold", "warm")
    )) %>%
    select(pretreatment, mass) %>%
    drop_na(),
  formula = mass ~ 0 + pretreatment,
  prior = c(
    set_prior("normal(257.5, 25)", class = "b",
              coef = "pretreatmentcold"),
    set_prior("normal(242.5, 25)", class = "b",
              coef = "pretreatmentwarm")
  ),
  family = "gaussian",
  seed = 200,
  cores = 4, chains = 4,
  iter = 50000, warmup = 10000, thin = 10,
  control = list(adapt_delta = 0.95, max_treedepth = 13),
  silent = TRUE, refresh = 0,
  file = "./models/massGrowthAnovaStrict.Rds"
)

caption <- paste0("Results from a Bayesian, one-way ANOVA comparing ",
                  "body mass at 8 weeks of Japanese quail reared in the ",
                  "cold (10°C until 8 weeks of age; n = ",
                  nrow(subset(growthAnovaStrict$data, pretreatment == "cold")),
                  "), or warmth (30°C until at least 3 weeks of age; n = ",
                  nrow(subset(growthAnovaStrict$data, pretreatment == "warm")), ")."
                  )

growthMassStrictAnovaTable <-
  data.frame(
    hypothesis(growthAnovaStrict,
               hypothesis = "pretreatmentcold - pretreatmentwarm > 0",
               class = "b", robust = TRUE, alpha = 0.5)$hypothesis
    ) %>%
  select("deltaMass" = Estimate,
         "pProb" = Post.Prob) %>%
  mutate("Hypothesis" = "Cold-Reared Mass > Warm-Reared Mass") %>%
  mutate(deltaMass = round(deltaMass, digits = 3),
         pProb = round(pProb, digits = 3)) %>%
  select(Hypothesis, "Delta Mass (g)" = deltaMass,
         "Posterior Probability" = pProb) %>%
  kbl(.,
    longtable = T, booktabs = T,
    caption = caption
  ) %>%
  kable_styling(latex_options = "striped")
```

```
growthMassStrictAnovaTable
```

***Table 10:*** *Results from a Bayesian, one-way ANOVA comparing body mass at 8 weeks of Japanese quail reared in the cold (10°C until 8 weeks of age; n = 24), or warmth (30°C until at least 3 weeks of age; n = 23).*

| Hypothesis | Delta Mass (g) | Posterior Probability |
|---|---|---|
| Cold-Reared Mass > Warm-Reared Mass | -4.704 | 0.335 |

To evaluate an effect of rearing condition on extremity (tarsus) elongation during development, we again used a non-linear model with tarsus length being a Gompertz function of age in weeks with a log-log, age-dependent error term. Because body mass may also explain some variance in tarsus length within ages, individual body mass, mean-centred and scaled by standard deviation per week of age, was included as an additional linear predictor of tarsus length. Our model predicting tarsus length was therefore as follows:

$$Tarsus\ Length_{ij} \sim a \cdot e^{-b \cdot e^{-c \cdot Age_{ij}}} + \beta_1 \cdot Scaled\ Mass + \mu_{0j} + \epsilon_{ij}$$
$$a_{ij} \sim \beta_{a0} + \beta_{a1} \cdot Cold\ Reared_j + \beta_{a2} \cdot Warm\ Reared_j + \mu_{0aj}$$
$$b_{ij} \sim \beta_{b0} + \beta_{b1} \cdot Cold\ Reared_j + \beta_{b2} \cdot Warm\ Reared_j + \mu_{0bj}$$
$$c_{ij} \sim \beta_{c0} + \beta_{c1} \cdot Cold\ Reared_j + \beta_{c2} \cdot Warm\ Reared_j + \mu_{0cj}$$
$$ln(\epsilon_{ij}) \sim \tau_0 + \tau_1 * ln(Age_{ij} + 1)$$

where $\beta_1$ represents the slope of the predicted relationship between body mass (g) and tarsus length (mm), and all other variables remain as previously described.

Priors for model parameters were again moderately informative and derived from findings of Burness et al (2013) and Persson et al (2024), while assuming a y-intercept (tarsus length at hatching) of approximately 10 mm. For the asymptote ($\beta_{a0}$), x-axis displacement ($\beta_{b0}$), and growth rate ($\beta_{c0}$) of our population growth curve, priors were normal ($\beta_{a0}$: mean = 37.5, standard deviation = 2.5; $\beta_{b0}$: mean = 1.35, standard deviation = 0.5) and skew-normal ($\beta_{c0}$: $\xi = 0.5$, $\omega = 0.1$, $\alpha = 2.5$) while those for the effect of rearing condition on growth parameters $a$, $b$, and $c$ (i.e. $\beta_{a1-2}$, $\beta_{b1-2}$, and $\beta_{c1-2}$ respectively) were all normal-distributed. For effects of cold- and warming-rearing on asymptotes, means were derived from Burness et al (2013) and represented the calculated difference in tarsus length from mean values among birds raised at 15°C and 30°C until 66 days of age (-0.35 for cold-rearing and 0.35 for warm-rearing respectively). Standard deviations were set broadly at 2. For similar effects of rearing treatment on x-axis displacement and growth rate, we assumed means of 0 and standard deviations of 0.5 and 0.25 respectively. Since we expected the correlation between body mass and tarsus length to be positive and low ($< 1$) but above 0, we used a skew normal distribution for our prior on $\beta_1$, with $\xi$ equaling 0.5, $\omega$ equaling 0.15, and $\alpha$ equaling 5. For our priors on our batch and individual-level effects ($\mu_{0a}$ - $\mu_{0c}$, $\mu_{0j}$), we assumed exponential distributions with lambda values of 1 ($\mu_{0a}$), 10 ($\mu_{0b}$), 25 ($\mu_{0c}$), and 1.5 ($\mu_{0j}$). Finally, for our and error structure terms ($\tau_0$ and $\tau_1$), we assumed both exponential ($\lambda = 1.5$) and skew normal distributions ($\tau_0$: $\xi = 0$, $\omega = 0.5$, $\alpha = 10$; $\tau_1$: $\xi = 0$, $\omega = 0.25$, $\alpha = 10$) respectively.

As previous, 4 Hamiltonian Monte Carlo (HMC) chains were used, with each run for 50000 iterations and 10000 warm-up iterations, then sampled every 10 iterations. Prior suitability is evaluated by a prior predictive check below, after graphically evaluating those of $\tau_0$, $\tau_1$, and $\beta_1$.

```
# Checking how the standard deviation of tarsus length scales with age

data %>%
  filter(week <= 8) %>%
  mutate(pretreatment = factor(pretreatment,
    levels = c("neutral", "cold", "warm")
```

```
)) %>%
  select(ring, pretreatment, week, tarsusLengthMean) %>%
  distinct() %>%
  mutate(weekB = log(week + 1)) %>%
  group_by(week, weekB) %>%
  summarise(SD = log(sd(tarsusLengthMean, na.rm = T))) %>%
  ungroup() %>%
  ggplot(aes(x = weekB, y = SD)) +
  geom_point(size = 3, pch = 21, colour = "black", fill = "grey50") +
  geom_smooth(method = "lm", colour = "black",
              linetype = "dashed", se = FALSE) +
  xlab("Age (Weeks + 1; Natural-log Transformed )") +
  ylab("Standard Deviation of Tarsus Length\n(Natural-log Transformed)") +
  theme_classic()
```

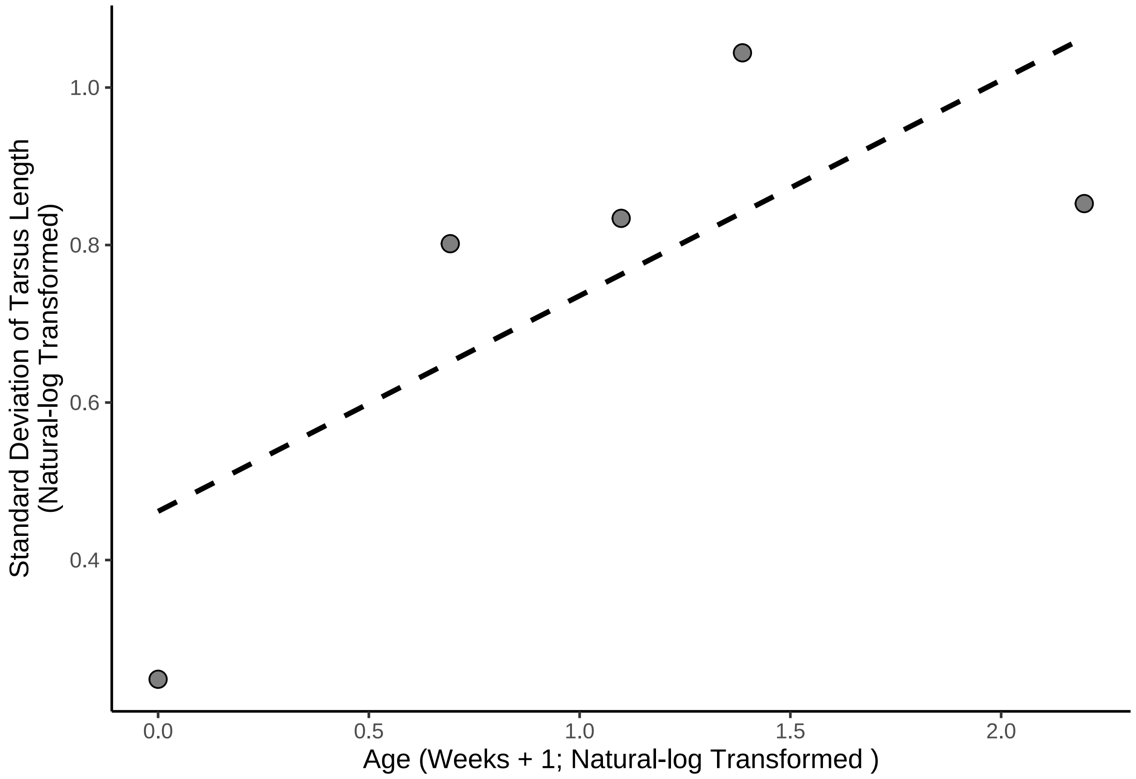


***Figure 23:*** *Effect of natural log-transformed age (in weeks + 1) on the natural log-transformed standard deviation of tarsus length (mm). Dotted line represents a line of body best fit estimated from a linear relationship by internal functions of the R package ggplot (Wickham, 2011).*

```
# Weakly linear, but clearly increasing between 0
# and 3 weeks of age. For this reason, retaining linear expectations.
```

```
# Confirming prior for age-independent effect of natural-log
# transformed body mass (g) on tarsus length (mm).

ggplot(data %>%
  filter(week <= 8) %>%
  mutate(pretreatment = factor(pretreatment,
    levels = c("neutral", "cold", "warm")
  )) %>%
  select(ring, pretreatment, week, mass, tarsusLengthMean) %>%
  distinct() %>%
  drop_na() %>%
  group_by(week) %>%
```

```
  mutate(
    "adjustedTarsus" = tarsusLengthMean - mean(tarsusLengthMean),
    "centredMass" = (mass - mean(mass, na.rm = T)) / sd(mass, na.rm = T)
  ) %>%
  ungroup(), aes(x = centredMass, y = adjustedTarsus)) +
  stat_summary_bin(geom = "errorbar", fun.data = "mean_se",
                   binwidth = 0.5, colour = "black") +
  stat_summary_bin(geom = "point", fun = "mean", binwidth = 0.5,
                   pch = 21, size = 5, colour = "black",
                   fill = "lightblue2", alpha = 0.5) +
  geom_smooth(method = "lm", colour = "black", se = FALSE,
              linetype = "dashed") +
  xlab("Age-Corrected Body Mass\n(g; Mean-centred Per Week of Age)") +
  ylab("Age-Corrected Tarsus Length\n(mm; Mean-centred Per Week of Age)") +
  theme_classic()
```

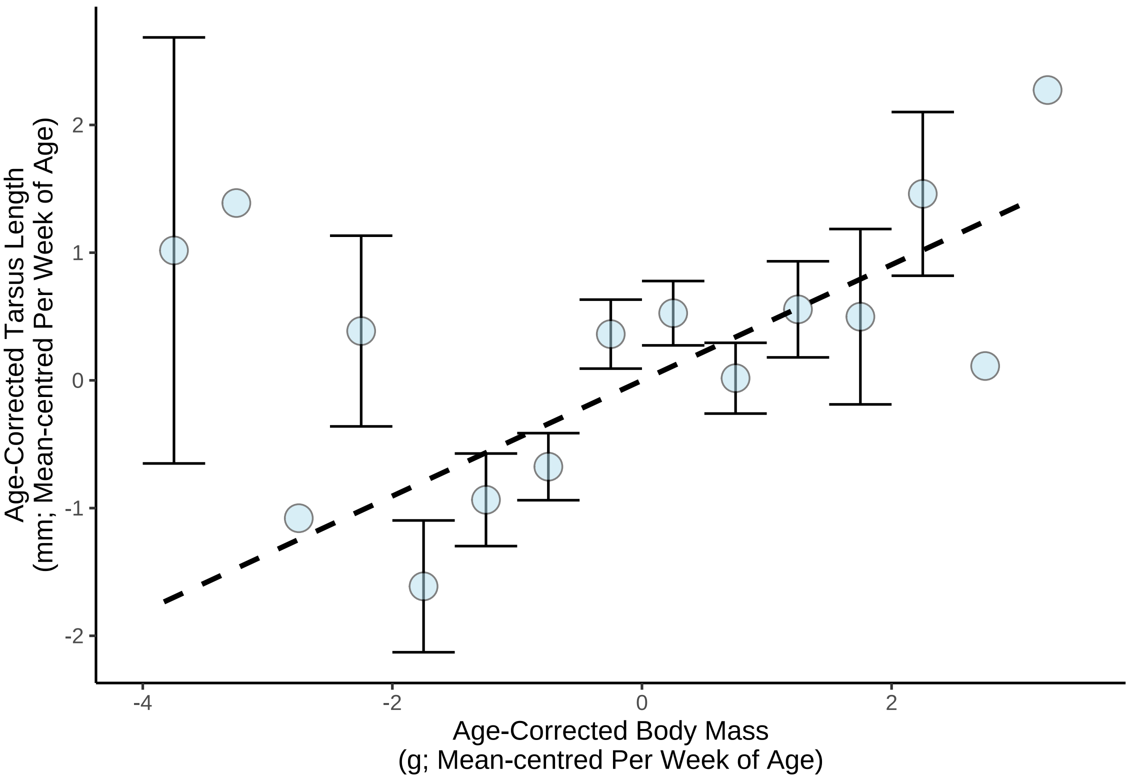


***Figure 24:*** *Effect of natural log-transformed body mass (g) on the tarsus length (mm), mean-centred by age in weeks. Dots represent means binned at intervals of 0.5 and errorbars represent standard errors around means. Dotted line represents a line of body best fit estimated from a linear relationship by internal functions of the R package ggplot (Wickham, 2011).*

```
data %>%
  filter(week == 0) %>%
  summarise(min(tarsusLengthMean, na.rm = T))
```

```
## # A tibble: 1 x 1
##   `min(tarsusLengthMean, na.rm = T)`
##                                 <dbl>
## 1                                9.54
```

```
bAtY <- function(a, yint) {
  b <- -log(yint / a)
  return(b)
}
```

```
cat(paste0("Estimated b value = ",
           round(bAtY(37.5, 10), digits = 3)))
```

```
## Estimated b value = 1.322
```

```
bXi <- round(skewxi(round(bAtY(37.5, 10), digits = 3),
                    omega = 1, alpha = 5), digits = 3)
cat(paste("xi value for skew-normal distribution on b = ", bXi))
```

```
## xi value for skew-normal distribution on b =  0.54
```

```
growthModelTarsus_ppCheck <- brm(
  data = data %>%
    filter(week <= 8) %>%
    mutate(pretreatment = factor(pretreatment,
      levels = c("neutral", "cold", "warm")
    )) %>%
    select(ring, pretreatment, week, mass,
           "tarsus" = tarsusLengthMean, "batch" = exp) %>%
    distinct() %>%
    mutate(
      weekB = week + 1
    ) %>%
    group_by(week) %>%
    mutate(cScaledMass = (mass - mean(mass, na.rm = T)) /
             sd(mass, na.rm = T)) %>%
    ungroup(),
  formula = bf(tarsus ~ A * exp(-B * exp(-C * week)) + D,
    A ~ 1 + pretreatment + (1|batch),
    B ~ 1 + pretreatment + (1|batch),
    C ~ 1 + pretreatment + (1|batch),
    D ~ 1 + cScaledMass + (1 | ring),
    sigma ~ 0 + intercept + log(weekB),
    nl = TRUE
  ),
  prior = c(
    set_prior("normal(37.5, 2.5)",
              class = "b", coef = "Intercept",
              nlpar = "A"
    ),
    set_prior("normal(-0.35, 2)",
              class = "b", coef = "pretreatmentcold",
              nlpar = "A"
    ),
    set_prior("normal(0.35, 2)",
              class = "b", coef = "pretreatmentwarm",
              nlpar = "A"
    ),
    set_prior("exponential(1)", class = "sd",
              coef = "Intercept",
              group = "batch", nlpar = "A"
    ),
    set_prior("skew_normal(0.6, 0.5, 2.5)",
              class = "b", coef = "Intercept",
              nlpar = "B"
    ),
    set_prior("normal(0, 0.5)",
              class = "b", coef = "pretreatmentcold",
              nlpar = "B"
    ),
    set_prior("normal(0, 0.5)",
              class = "b", coef = "pretreatmentwarm",
              nlpar = "B"
    ),
    set_prior("exponential(10)", class = "sd",
              coef = "Intercept",
              group = "batch", nlpar = "B"
    ),
```

```
    set_prior("skew_normal(0.5, 0.1, 2.5)",
              class = "b", coef = "Intercept",
              nlpar = "C"
    ),
    set_prior("normal(0, 0.25)",
              class = "b", coef = "pretreatmentcold",
              nlpar = "C"
    ),
    set_prior("normal(0, 0.25)",
              class = "b", coef = "pretreatmentwarm",
              nlpar = "C"
    ),
    set_prior("exponential(25)", class = "sd",
              coef = "Intercept",
              group = "batch", nlpar = "C"
    ),
    set_prior("normal(0, 1.5)", class = "b",
              coef = "Intercept", nlpar = "D"),
    set_prior("skew_normal(0.5, 0.15, 5)", class = "b",
              coef = "cScaledMass", nlpar = "D"),
    set_prior("exponential(1.5)", class = "sd",
              coef = "Intercept", group = "ring", nlpar = "D"),
    set_prior("skew_normal(0, 0.25, 10)", class = "b",
              coef = "intercept", dpar = "sigma"),
    set_prior("skew_normal(0, 0.25, 10)", class = "b",
              coef = "logweekB", dpar = "sigma")
  ),
  family = "gaussian",
  seed = 103,
  cores = 4, chains = 4,
  iter = 50000, warmup = 10000, thin = 10,
  control = list(adapt_delta = 0.95, max_treedepth = 13),
  silent = TRUE, refresh = 0,
  sample_prior = "only",
  file = "./models/ppCheckTarsus.Rds"
)

# Checking how posterior predictions overlay with true tarsus length values

pp_check2(growthModelTarsus_ppCheck, xlab = "Tarsus Length (mm)")
```

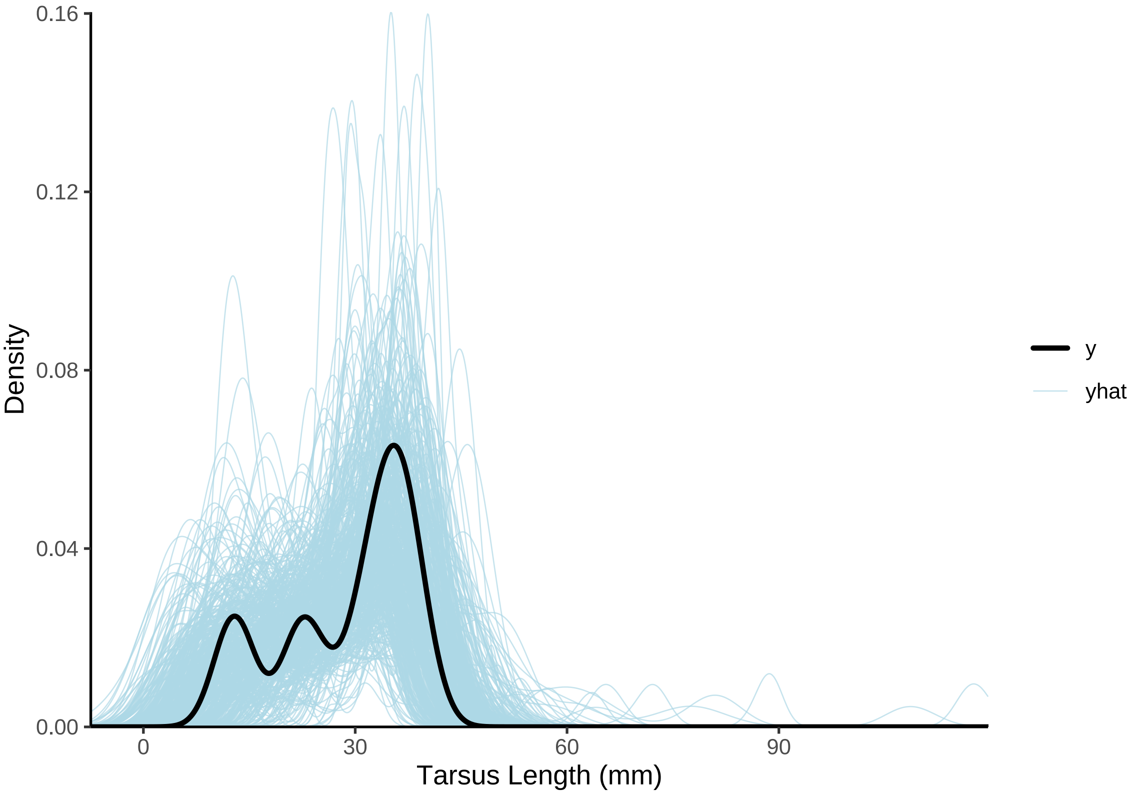


***Figure 25:*** *Overlay of predicted (blue) and true (black) tarsus length densities, where predicted densities are derived from priors in a Bayesian non-linear model. Clear overlap betwen the black and blue lines suggests that model priors are reasonable with respect to the data.*

```
# Reasonable overlap, despite some individuals having
# unrealistically high tarsus length. Proceeding to full model,
# given our relative uncertainty around true parameter values.
```

Similar to our model estimating body mass during growth, initial values for our HMC chains are drawn from within prior distributions to facilitate chain convergence.

```
initFunction <- function(chain_id = 1) {
  list(
    "b_A" = c(rnorm(1, 37.5, 2.5), -0.35, 0.35),
    "b_B" = c(1.35, 0, 0),
    "b_C" = c(0.5, 0, 0),
    "b_D" = c(rnorm(1, 0, 1.5),
              rskew_normal(1, xi = 0.5, omega = 0.15, alpha = 5)),
    "b_sigma" = rskew_normal(2, xi = 0, omega = 0.25, alpha = 10),
    "sd_1" = rexp(1, 1),
    "sd_2" = rexp(1, 10),
    "sd_3" = rexp(1, 25),
    "sd_4" = rexp(1, 0.5)
  )
}

# Loading initial values into a list.

initList <- lapply(1:4, initFunction)

growthModelTarsus <- brm(
  data = data %>%
    filter(week <= 8) %>%
```

```
    mutate(pretreatment = factor(pretreatment,
                                  levels = c("neutral", "cold", "warm")
    )) %>%
    select(ring, pretreatment, week, mass,
           "tarsus" = tarsusLengthMean, "batch" = exp) %>%
    distinct() %>%
    mutate(
      weekB = week + 1
    ) %>%
    group_by(week) %>%
    mutate(cScaledMass = (mass - mean(mass, na.rm = T)) /
             sd(mass, na.rm = T)) %>%
    ungroup(),
  formula = bf(tarsus ~ A * exp(-B * exp(-C * week)) + D,
               A ~ 1 + pretreatment + (1|batch),
               B ~ 1 + pretreatment + (1|batch),
               C ~ 1 + pretreatment + (1|batch),
               # D ~ 1 + mass + (1|ring),
               D ~ 1 + cScaledMass + (1 | ring),
               sigma ~ 0 + intercept + log(weekB),
               nl = TRUE
  ),
  prior = c(
    set_prior("normal(37.5, 2.5)",
              class = "b", coef = "Intercept",
              nlpar = "A"
    ),
    set_prior("normal(-0.35, 2)",
              class = "b", coef = "pretreatmentcold",
              nlpar = "A"
    ),
    set_prior("normal(0.35, 2)",
              class = "b", coef = "pretreatmentwarm",
              nlpar = "A"
    ),
    set_prior("exponential(1)", class = "sd",
              coef = "Intercept",
              group = "batch", nlpar = "A"
    ),
    set_prior("skew_normal(0.6, 0.5, 2.5)",
              class = "b", coef = "Intercept",
              nlpar = "B"
    ),
    set_prior("normal(0, 0.5)",
              class = "b", coef = "pretreatmentcold",
              nlpar = "B"
    ),
    set_prior("normal(0, 0.5)",
              class = "b", coef = "pretreatmentwarm",
              nlpar = "B"
    ),
    set_prior("exponential(10)", class = "sd",
              coef = "Intercept",
              group = "batch", nlpar = "B"
    ),
    set_prior("skew_normal(0.5, 0.1, 2.5)",
              class = "b", coef = "Intercept",
              nlpar = "C"
    ),
    set_prior("normal(0, 0.25)",
              class = "b", coef = "pretreatmentcold",
              nlpar = "C"
    ),
    set_prior("normal(0, 0.25)",
              class = "b", coef = "pretreatmentwarm",
              nlpar = "C"
    ),
```

```
    set_prior("exponential(25)", class = "sd",
              coef = "Intercept",
              group = "batch", nlpar = "C"
    ),
    set_prior("normal(0, 1.5)", class = "b",
              coef = "Intercept", nlpar = "D"),
    set_prior("skew_normal(0.5, 0.15, 5)", class = "b",
              coef = "cScaledMass", nlpar = "D"),
    set_prior("exponential(1.5)", class = "sd",
              coef = "Intercept", group = "ring", nlpar = "D"),
    set_prior("skew_normal(0, 0.25, 10)", class = "b",
              coef = "intercept", dpar = "sigma"),
    set_prior("skew_normal(0, 0.25, 10)", class = "b",
              coef = "logweekB", dpar = "sigma")
  ),
  family = "gaussian",
  seed = 100,
  cores = 4, chains = 4,
  init = initList,
  iter = 50000, warmup = 10000, thin = 10,
  control = list(adapt_delta = 0.95, max_treedepth = 14),
  silent = TRUE, refresh = 0,
  file = "./models/growthModelTarsus.Rds"
)

# Taking a look at effective sample size to sample size
# ratios and Gelman-Rubin statistics.

ggarrange(
  ggplot(data = data.frame("Rhat" = brms::rhat(growthModelTarsus)),
         aes(x = Rhat)) +
    geom_density() +
    theme_classic() +
    xlab(
      TeX('$\\hat{R}$')
    ) +
    ylab("Density"),
  ggplot(data = data.frame("Neff" = neffBase(growthModelTarsus)),
         aes(x = Neff)) +
    geom_density() +
    theme_classic() +
    xlab(
      TeX('$N_{eff}/N~Ratio$')
    ) +
    ylab("Density")
)
```

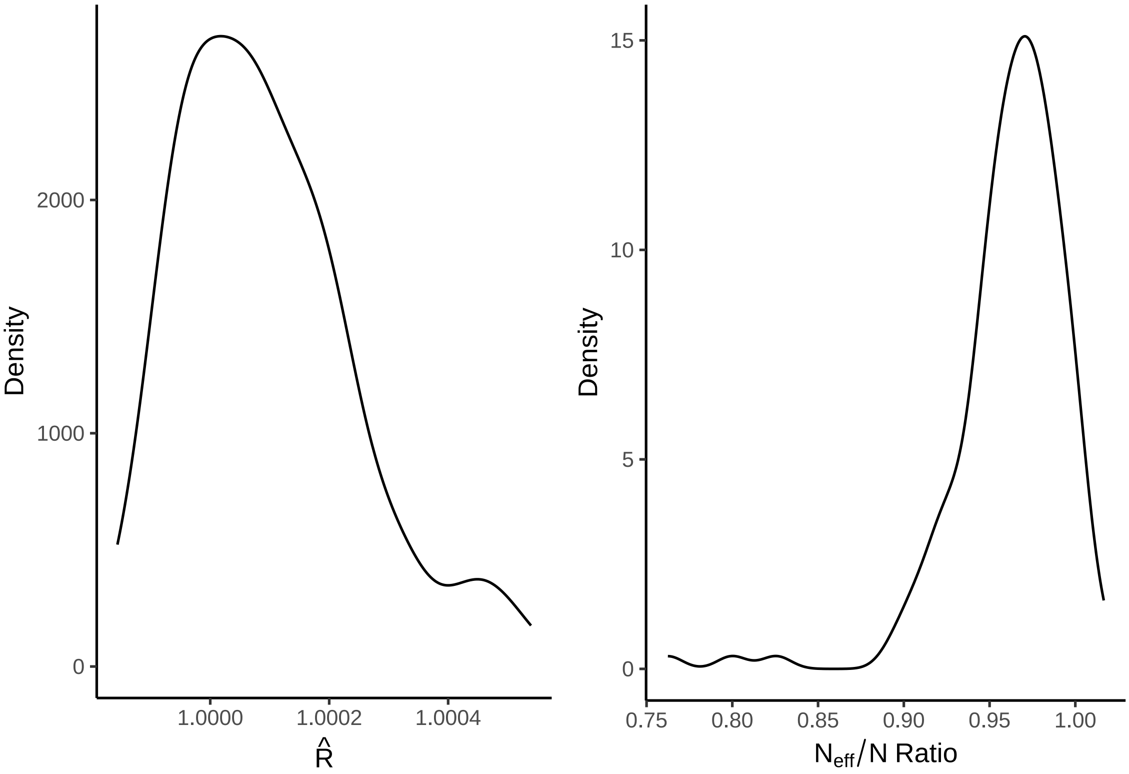


***Figure 26:*** *Gelman-Rubin statistics ($\hat{R}$) and ratio of effective samples sizes by samples sizes per parameter from a Bayesian non-linear model estimating tarsus length (mm) among Japanese quail.*

```
# Good, although some loss in effective sample sizes.
```

```
pp_check2(growthModelTarsus, xlab = "Tarsus Length (mm)")
```

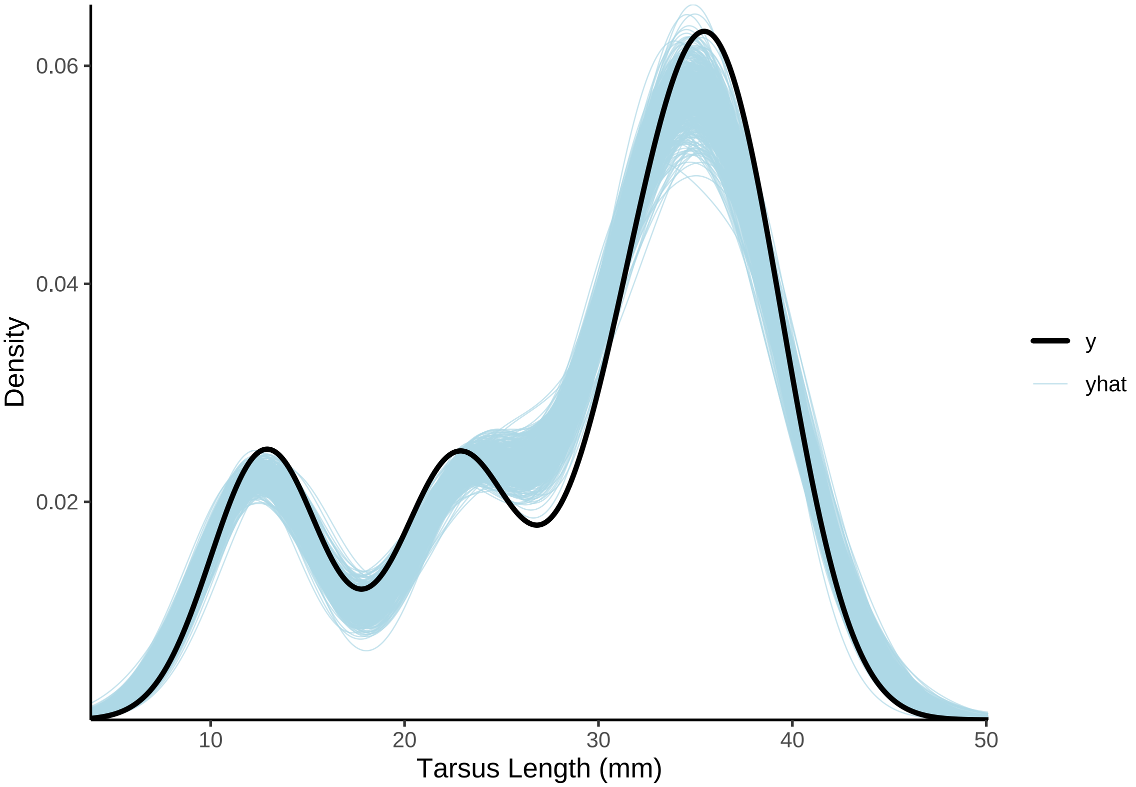


***Figure 27:*** *Posterior predictions of a Bayesian non-linear model predicting Japanese tarsus length (mm) across time, overlayed with true distributions of quail mass. Blue lines represent predictions from posterior draws, while the black line represents true mass distributions.*

```
# Distinct overlay. Checking fit with a scatterplot.
```

```
growthModelTarsus$data %>%
  mutate("Fit" = predict(growthModelTarsus,
                         robust = TRUE)[, "Estimate"]) %>%
  ggplot(aes(x = tarsus, y = Fit, fill = week)) +
  geom_point(pch = 21, colour = "black", size = 2, alpha = 0.5) +
  geom_smooth(method = "lm", colour = "black",
              linetype = "dashed", se = FALSE) +
  scale_fill_gradient2() +
  theme_classic() +
  xlab("Tarsus Length (mm)") +
  ylab("Predicted Tarsus Length (mm)")
```

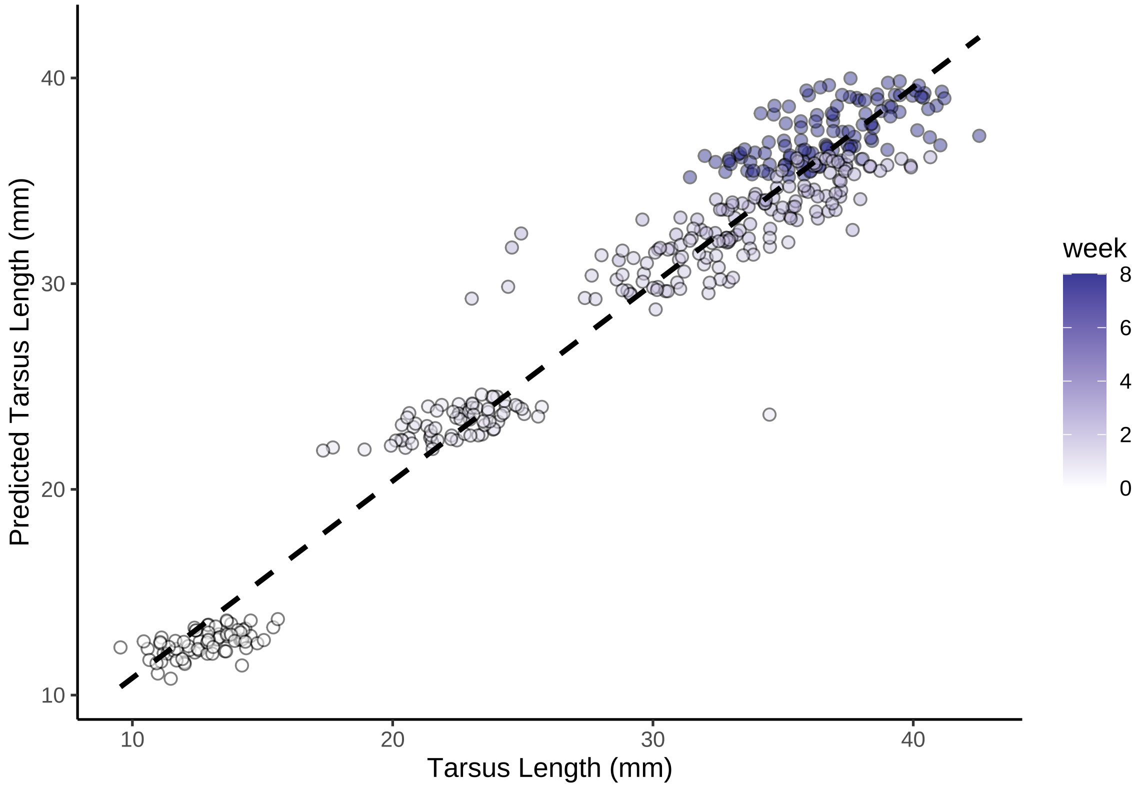


***Figure 28:*** *Scatterplot of Japanese quail tarsus length (mm) against predicted tarsus length (mm) from a Bayesian non-linear model. Dashed line indicates line of best fit, as estimated by the R package ggplot2 (Wickham, 2011).*

```
# Good. Slightly over-fitting at 0 weeks of age (suggesting that
# 'b' may be slightly over-estimated), but otherwise little worry.
# Plotting mean responses
```

Posterior predictions appear to overlay with raw data well. Below, tarsus elongation curves are predicted from our model for each rearing treatment. These curves are then plotted and overlaid with raw tarsus length means, per treatment, across our observed period of elongation.

```
with(
  growthModelTarsus$data,
  expand.grid(
    "week" = seq(0, 8, by = 0.1),
    "pretreatment" = c("cold", "neutral", "warm"),
    "cScaledMass" = 0
  )
) %>%
  mutate("weekB" = week + 1) %>%
  mutate(
    "Fit" = predict(growthModelTarsus, re_form = NA,
                    newdata = .,
                    robust = TRUE)[, "Estimate"],
    "SE" = predict(growthModelTarsus, re_form = NA,
                   newdata = .,
                    robust = TRUE)[, "Est.Error"]
  ) %>%
  mutate(
    "LCL" = Fit - SE,
    "UCL" = Fit + SE
  ) %>%
```

```
mutate(pretreatment = ifelse(pretreatment == "cold",
                             "Cold (10°C)",
                             ifelse(pretreatment == "neutral",
                                    "Mild (20°C)",
                                    "Warm (30°C)"))) %>%
mutate(pretreatment = factor(pretreatment,
                             levels =
                               c("Cold (10°C)",
                                 "Mild (20°C)",
                                "Warm (30°C)")
)
) %>%
ggplot(aes(x = week, y = Fit,
 fill = pretreatment, linetype = pretreatment)) +
geom_ribbon(aes(x = week, ymin = LCL, ymax = UCL),
  colour = NA, size = 0.25, alpha = 0.3
) +
geom_line(colour = "black", alpha = 0.7) +
stat_summary(
  data = growthModelTarsus$data %>%
  mutate(pretreatment = ifelse(pretreatment == "cold",
                             "Cold (10°C)",
                             ifelse(pretreatment == "neutral",
                                    "Mild (20°C)",
                                    "Warm (30°C)"))) %>%
  mutate(pretreatment = factor(pretreatment,
                             levels =
                               c("Cold (10°C)",
                                "Mild (20°C)",
                                "Warm (30°C)")
)
),
  aes(x = week, y = tarsus),
  geom = "errorbar", fun.data = "mean_cl_boot",
  colour = "black", alpha = 0.7, width = 0.25,
  position = position_dodge(width = 0.15)
) +
stat_summary(
  data = growthModelTarsus$data %>%
  mutate(pretreatment = ifelse(pretreatment == "cold",
                             "Cold (10°C)",
                             ifelse(pretreatment == "neutral",
                                    "Mild (20°C)",
                                    "Warm (30°C)"))) %>%
  mutate(pretreatment = factor(pretreatment,
                             levels =
                               c("Cold (10°C)",
                                "Mild (20°C)",
                                "Warm (30°C)")
)
),
  aes(x = week, y = tarsus),
  geom = "point", fun = "mean", pch = 21, size = 3,
  colour = "black", alpha = 0.7,
  position = position_dodge(width = 0.15)
) +
theme_classic() +
scale_fill_manual(values = c("#7BB4E3", "black", "#CD5C5C"),
                  name = "Rearing\nConditions") +
scale_linetype_manual(values = c("dotted", "solid", "dashed"),
                      name = "Rearing\nConditions") +
xlab("Age (weeks)") +
ylab("Tarsus Length (mm)")
```

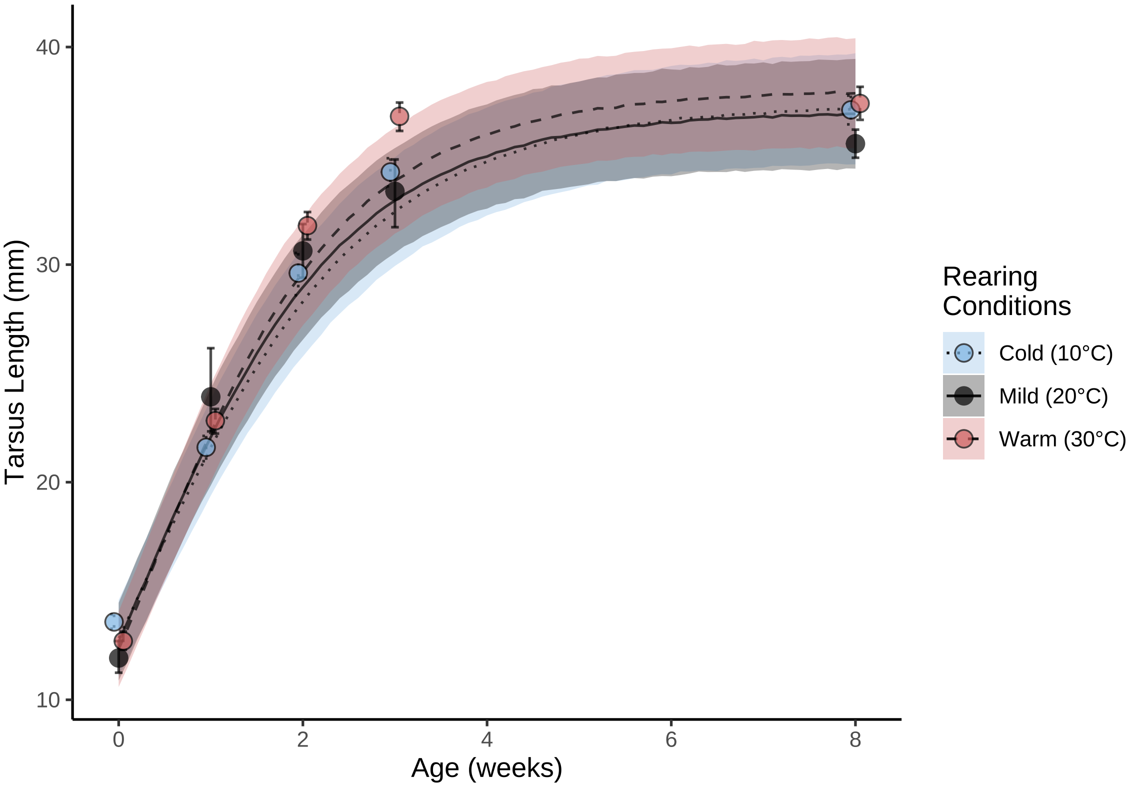


***Figure 29:*** *Tarsus length (mm) elongation curves of Japanese quail reared in the cold (10°C), mild conditions (20°C), or the warm (30°C) until at least 3 weeks of age. Dots represent mean values and errorbars represent quantile-based 95% credible intervals. Lines represent estimated trends in growth from a Bayesian non-linear model, while ribbons represent confidence around trends (± one standard error).*

```
# Good. Very rapid growth proceeded by some weak convergence
# among groups at asymptotes. Checking residuals next.
```

Next, residuals are again visualised to check for heteroskedasticity.

```
p1 <- growthModelTarsus$data %>%
  mutate("residuals" = residuals(growthModelTarsus,
                                 method = "posterior_predict",
                                 type = "pearson",
                                 robust = TRUE)[, "Estimate"]) %>%
  ggplot(aes(x = residuals)) +
  geom_density(colour = "black", fill = "white") +
  xlab("Pearson Residuals") +
  ylab("Density") +
  theme_classic()

p2 <- growthModelTarsus$data %>%
  mutate(
    "residuals" = residuals(growthModelTarsus,
      method = "posterior_predict",
      type = "pearson",
      robust = TRUE
    )[, "Estimate"],
    "fitted" = fitted(growthModelTarsus,
                                 robust = TRUE)[, "Estimate"]
  ) %>%
  ggplot(aes(x = fitted, y = residuals)) +
  geom_point(colour = "black", pch = 21, size = 2,
```

```
            fill = "grey75", alpha = 0.5) +
  ylab("Pearson Residuals") +
  xlab("Fitted Values (mm)") +
  theme_classic()

p3 <- growthModelTarsus$data %>%
  mutate("residuals" =
           residuals(growthModelTarsus,
                     method = "posterior_predict",
                     type = "pearson",
                     robust = TRUE
  )[, "Estimate"]) %>%
  ggplot(aes(x = week, y = residuals)) +
  geom_point(
    colour = "black", pch = 21, size = 2, fill = "grey75", alpha = 0.5,
    position = position_jitter(width = 0.25)
  ) +
  stat_summary(geom = "errorbar", fun.data = "mean_se",
               colour = "black", width = 0.25) +
  stat_summary(geom = "point", fun = "mean", pch = 21,
               colour = "black", fill = "white", size = 4) +
  ylab("Pearson Residuals") +
  xlab("Age (weeks)") +
  theme_classic()

p4 <- growthModelTarsus$data %>%
  mutate("residuals" =
           residuals(growthModelTarsus,
                     method = "posterior_predict",
                     type = "pearson",
                     robust = TRUE
  )[, "Estimate"]) %>%
  ggplot(aes(x = cScaledMass, y = residuals)) +
  geom_point(
    colour = "black", pch = 21, size = 2, fill = "grey75", alpha = 0.5,
    position = position_jitter(width = 0.25)
  ) +
  stat_summary(geom = "errorbar", fun.data = "mean_se",
               colour = "black", width = 0.25) +
  stat_summary(geom = "point", fun = "mean", pch = 21,
               colour = "black", fill = "white", size = 4) +
  ylab("Pearson Residuals") +
  xlab("Centred and Scaled Body Mass") +
  theme_classic()

p5 <- growthModelTarsus$data %>%
  mutate("residuals" =
           residuals(growthModelTarsus,
                     method = "posterior_predict",
                     type = "pearson",
                     robust = TRUE
  )[, "Estimate"]) %>%
  mutate(pretreatment = str_to_title(pretreatment)) %>%
  mutate(pretreatment = ifelse(pretreatment == "Cold",
                               "Cold\n(10°C)",
    ifelse(pretreatment == "Neutral", "Mild\n(20°C)",
      "Warm\n(30°C)"
    )
  )) %>%
  mutate(pretreatment = factor(pretreatment,
                               levels = c("Cold\n(10°C)",
                                          "Mild\n(20°C)",
                                          "Warm\n(30°C)"))) %>%
  ggplot(aes(x = pretreatment, y = residuals)) +
  geom_point(
    colour = "black", pch = 21, size = 2, fill = "grey75", alpha = 0.5,
    position = position_jitter(width = 0.25)
```

```
  ) +
  stat_summary(geom = "errorbar", fun.data = "mean_se",
               colour = "black", width = 0.25) +
  stat_summary(geom = "point", fun = "mean", pch = 21,
               colour = "black", fill = "white", size = 4) +
  ylab("Pearson Residuals") +
  xlab("Rearing Conditions)") +
  theme_classic()

((p1 + p2) / (p3 + p4)) + p5
```

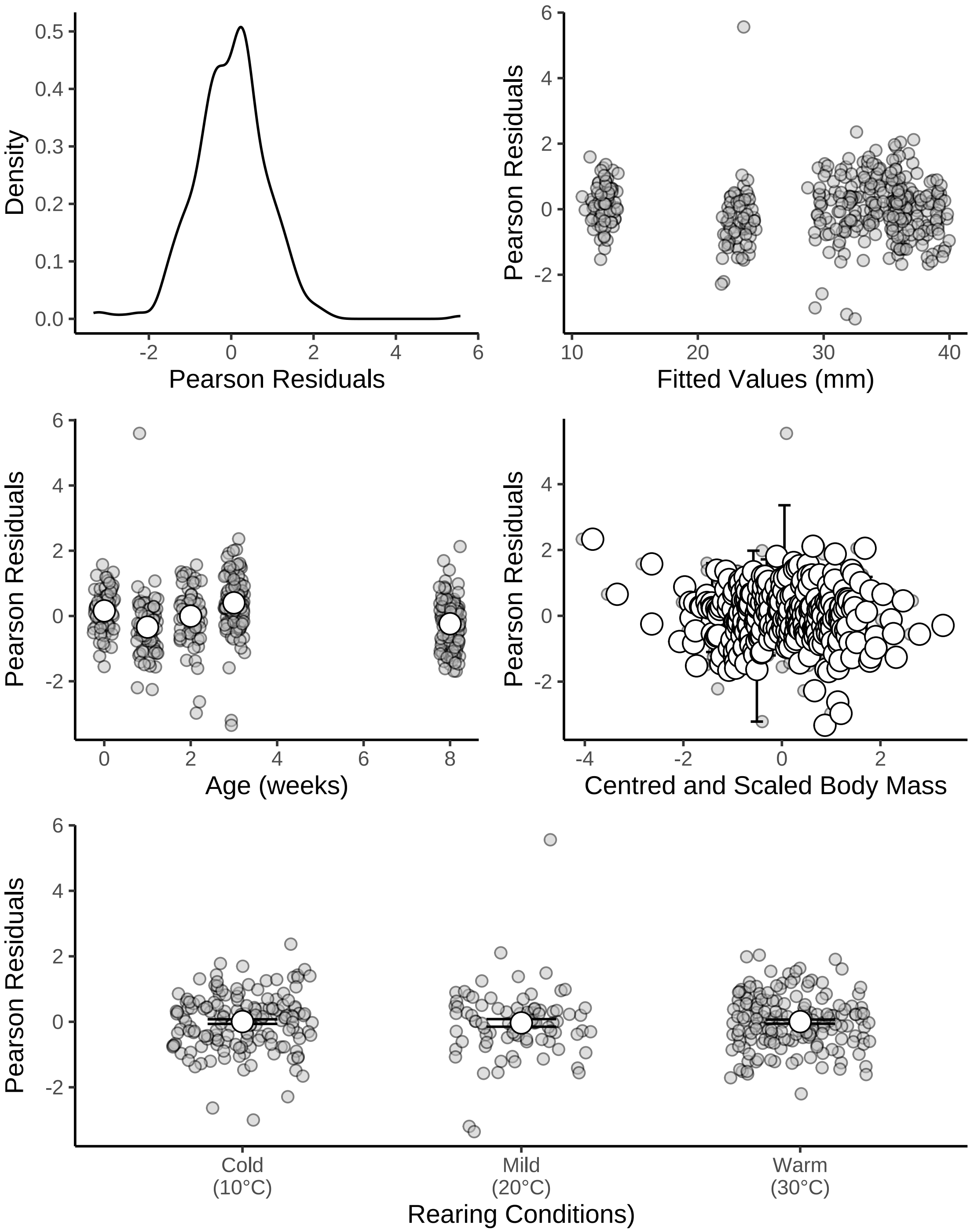


***Figure 30:*** *Density and distributions of Pearson residuals across fitted values and model predictors (including age and rearing conditions). All residuals pertain to those drawn from a Bayesian non-linear model predicting tarsus length (mm) during growth in Japanese quail. Pearson residuals are shown rather than ordinary residuals to correct for the age-dependence of model error.*

Residuals appear homogenous, but reveal one potential extreme value. Below, the value is identified and scrutinised.

```r
caption <- paste0("Possible outlier identified from Bayesian ",
                  "non-linear model predicting tarsus length (mm) ",
                  "of Japanese quail across growth."
)

growthModelTarsus$data %>%
  mutate("Residuals" = residuals(growthModelTarsus,
                                 type = "pearson",
                                 robust = TRUE)[, "Estimate"]) %>%
  filter(Residuals > 4) %>%
  select("Bird Identity" = ring, "Age (weeks)" = week,
         "Centred & Scaled Mass (g)" = cScaledMass,
         "Tarsus Length (mm)" = tarsus) %>%
  kbl(., longtable = T, booktabs = T, format = "latex",
      caption = caption) %>%
  column_spec(column = c(1:10), width = "2.5cm")  %>%
  kable_styling(latex_options = "striped")
```

***Table 11:*** *Possible outlier identified from Bayesian non-linear model predicting tarsus length (mm) of Japanese quail across growth.*

| Bird Identity | Age (weeks) | Centred & Scaled Mass (g) | Tarsus Length (mm) |
|---|---|---|---|
| R3 | 1 | 0.0516127 | 34.477 |

```r
# Clearly a large tarsus length for its age. Plotting trends
# in this individuals' tarsus length across time.

growthModelTarsus$data %>%
  filter(ring == "R3") %>%
  ggplot(aes(x = week, y = tarsus)) +
  geom_line(colour = "black", linetype = "dashed", size = 0.5) +
  geom_point(size = 3, pch = 21, colour = "black", fill = "lightblue2") +
  xlab("Age (weeks)") +
  ylab("Tarsus Length (mm)") +
  theme_classic()
```

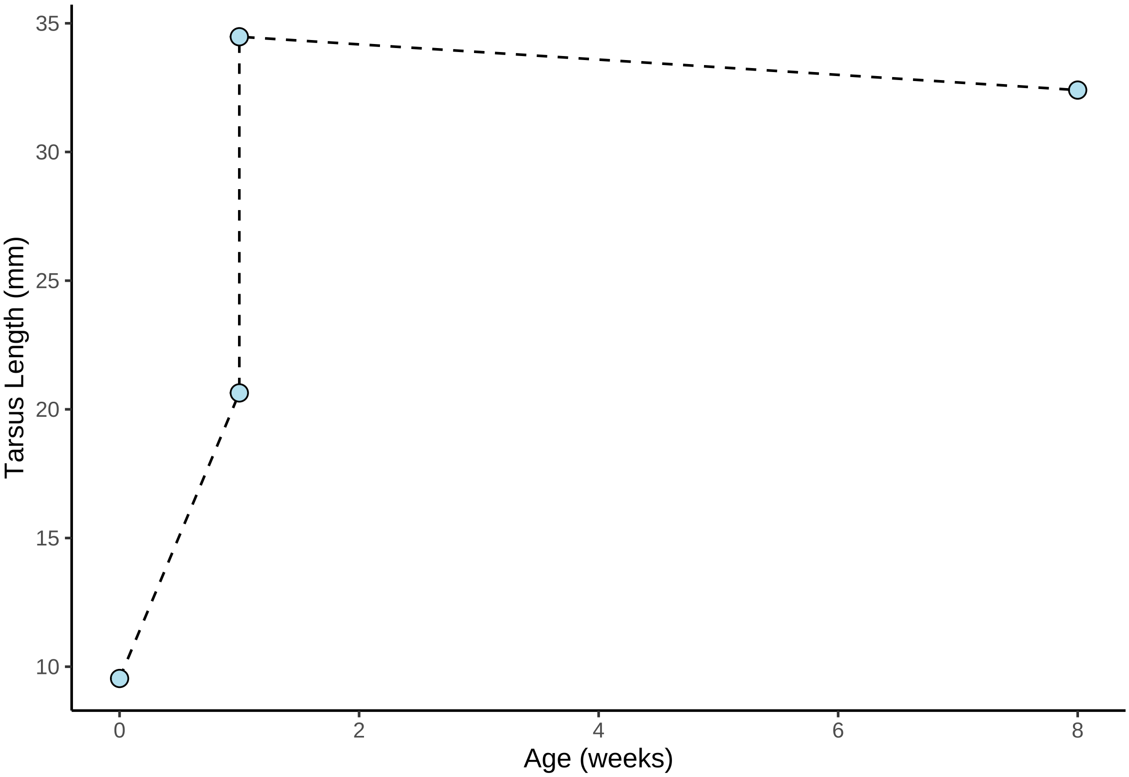


***Figure 31:*** *Tarsus elongation trends of possible outlier identified from Bayesian non-linear model predicting tarsus length (mm) of across growth in Japanese quail.*

```
# Two measurements labelled as being drawn from one. Assessing why.

caption <- paste0("Further information regarding possible tarsus ",
                  "length outlier identified from Bayesian non-linear ",
                  "model predicting tarsus length (mm) of ",
                  "Japanese quail across growth."
)

data %>%
  filter(ring == "R3" & week == 1) %>%
  mutate(
    sex = str_to_title(sex),
    tarsusCalibration = ifelse(tarsusCalibration == "gridUnder", "Grid Under",
      ifelse(tarsusCalibration == "gridOver", "Grid Over",
        tarsusCalibration
      )
    )
  ) %>%
  select(
    "Bird Identity" = ring, "Age (weeks)" = week,
    "Sex" = sex, "Body Mass (g)" = mass,
    "Tarsus Length (mm)" = tarsusLengthMean,
    "Calibration Method" = tarsusCalibration
  ) %>%
  kbl(., longtable = T, booktabs = T,
      caption = caption, format = "latex") %>%
  column_spec(column = c(1:10), width = "2cm")  %>%
  kable_styling(latex_options = "striped")
```

***Table 12:*** *Further information regarding possible tarsus length outlier identified from Bayesian non-linear model predicting tarsus length (mm) of Japanese quail across growth.*

| Bird Identity | Age (weeks) | Sex | Body Mass (g) | Tarsus Length (mm) | Calibration Method |
|---|---|---|---|---|---|
| R3 | 1 | Male | 34.8 | 20.6383 | Grid Under |
| R3 | 1 | Male | 34.8 | 34.4770 | Grid Over |

```
# Two different calibration methods used, so duplicates
# are not true duplications. Given that both tarsus length
# measurements are within the realistic range of
# Japanese quail morphometrics, both values should be retained.
# Nevertheless, our model is re-run without this data-point to
# dispel concerns about our bias entering our model coefficients.
```

To ensure that potential outliers are not biasing our model outcomes, our model is re-run without their inclusion and coefficients are visually compared between model iterations. Importantly, our revised model is ran with two HMC chains rather than four to improve computational efficiency.

```
initList <- lapply(1:2, initFunction)

growthModelTarsusB <- brm(
  data = data %>%
    filter(!(ring == "R3" & week == 1 &
               tarsusLengthMean > 34)) %>%
    filter(week <= 8) %>%
    mutate(pretreatment = factor(pretreatment,
      levels = c("neutral", "cold", "warm")
    )) %>%
    select(ring, pretreatment, week, mass, "batch" = exp,
           "tarsus" = tarsusLengthMean) %>%
    distinct() %>%
    mutate(weekB = week + 1) %>%
    group_by(week) %>%
    mutate(cScaledMass = (mass - mean(mass, na.rm = T)) /
             sd(mass, na.rm = T)) %>%
    ungroup(),
  formula = bf(tarsus ~ A * exp(-B * exp(-C * week)) + D,
               A ~ 1 + pretreatment + (1|batch),
               B ~ 1 + pretreatment + (1|batch),
               C ~ 1 + pretreatment + (1|batch),
               D ~ 1 + cScaledMass + (1 | ring),
               sigma ~ 0 + intercept + log(weekB),
               nl = TRUE
  ),
  prior = c(
    set_prior("normal(37.5, 2.5)",
              class = "b", coef = "Intercept",
              nlpar = "A"
    ),
    set_prior("normal(-0.35, 2)",
              class = "b", coef = "pretreatmentcold",
              nlpar = "A"
    ),
    set_prior("normal(0.35, 2)",
              class = "b", coef = "pretreatmentwarm",
              nlpar = "A"
    ),
    set_prior("exponential(1)", class = "sd",
              coef = "Intercept",
              group = "batch", nlpar = "A"
    ),
    set_prior("skew_normal(0.6, 0.5, 2.5)",
              class = "b", coef = "Intercept",
```

```
              nlpar = "B"
    ),
    set_prior("normal(0, 0.5)",
              class = "b", coef = "pretreatmentcold",
              nlpar = "B"
    ),
    set_prior("normal(0, 0.5)",
              class = "b", coef = "pretreatmentwarm",
              nlpar = "B"
    ),
    set_prior("exponential(10)", class = "sd",
              coef = "Intercept",
              group = "batch", nlpar = "B"
    ),
    set_prior("skew_normal(0.5, 0.1, 2.5)",
              class = "b", coef = "Intercept",
              nlpar = "C"
    ),
    set_prior("normal(0, 0.25)",
              class = "b", coef = "pretreatmentcold",
              nlpar = "C"
    ),
    set_prior("normal(0, 0.25)",
              class = "b", coef = "pretreatmentwarm",
              nlpar = "C"
    ),
    set_prior("exponential(25)", class = "sd",
              coef = "Intercept",
              group = "batch", nlpar = "C"
    ),
    set_prior("normal(0, 1.5)", class = "b",
              coef = "Intercept", nlpar = "D"),
    set_prior("skew_normal(0.5, 0.15, 5)", class = "b",
              coef = "cScaledMass", nlpar = "D"),
    set_prior("exponential(1.5)", class = "sd",
              coef = "Intercept", group = "ring", nlpar = "D"),
    set_prior("skew_normal(0, 0.25, 10)", class = "b",
              coef = "intercept", dpar = "sigma"),
    set_prior("skew_normal(0, 0.25, 10)", class = "b",
              coef = "logweekB", dpar = "sigma")
  ),
  family = "gaussian",
  seed = 100,
  cores = 2, chains = 2,
  init = initList,
  iter = 50000, warmup = 10000, thin = 10,
  control = list(adapt_delta = 0.95, max_treedepth = 13),
  silent = TRUE, refresh = 0,
  file = "./models/growthModelTarsusB.Rds"
)

# Comparing model coefficients visually.

as.data.frame(growthModelTarsus) %>%
  pivot_longer(everything(), names_to = "Parameter",
               values_to = "Values") %>%
  filter(grepl("b_|sd_", Parameter)) %>%
  arrange(Parameter) %>%
  mutate("model" = "Model A") %>%
  rbind(
    .,
    as.data.frame(growthModelTarsusB) %>%
      pivot_longer(everything(), names_to = "Parameter",
                   values_to = "Values") %>%
      filter(grepl("b_|sd_", Parameter)) %>%
      arrange(Parameter) %>%
      mutate("model" = "Model B")
```

```
  ) %>%
  merge(., tribble(
    ~Parameter, ~Par,
    "b_A_Intercept", "Beta a0",
    "b_A_pretreatmentcold", "Beta a1\n(Cold-reared)",
    "b_A_pretreatmentwarm", "Beta a2\n(Warm-reared)",
    "sd_batch__A_Intercept", "Mu 0a",
    "b_B_Intercept", "Beta b0",
    "b_B_pretreatmentcold", "Beta b1\n(Cold-reared)",
    "b_B_pretreatmentwarm", "Beta b2\n(Warm-reared)",
    "sd_batch__B_Intercept", "Mu 0b",
    "b_C_Intercept", "Beta c0",
    "b_C_pretreatmentcold", "Beta c1\n(Cold-reared)",
    "b_C_pretreatmentwarm", "Beta c2\n(Warm-reared)",
    "sd_batch__C_Intercept", "Mu 0c",
    "b_D_Intercept", "Beta 0",
    "b_D_cScaledMass", "Beta 1",
    "sd_ring__D_Intercept", "Mu 0 (Individual\nIntercept)",
    "b_sigma_intercept", "Tau 0",
    "b_sigma_logweekB", "Tau 1"
  ),
  by = "Parameter", all.x = TRUE
  ) %>%
  ggplot(aes(x = Values, fill = model)) +
  facet_wrap(~Par, scales = "free") +
  geom_density(colour = "black", alpha = 0.5) +
  geom_vline(xintercept = 0, linetype = "dashed", colour = "firebrick4") +
  ylab("Density") +
  scale_fill_manual(values = c("lightblue2", "grey40"), name = NULL) +
  theme_classic() +
  theme(axis.title.x = element_blank(),
        legend.position = "bottom")
```

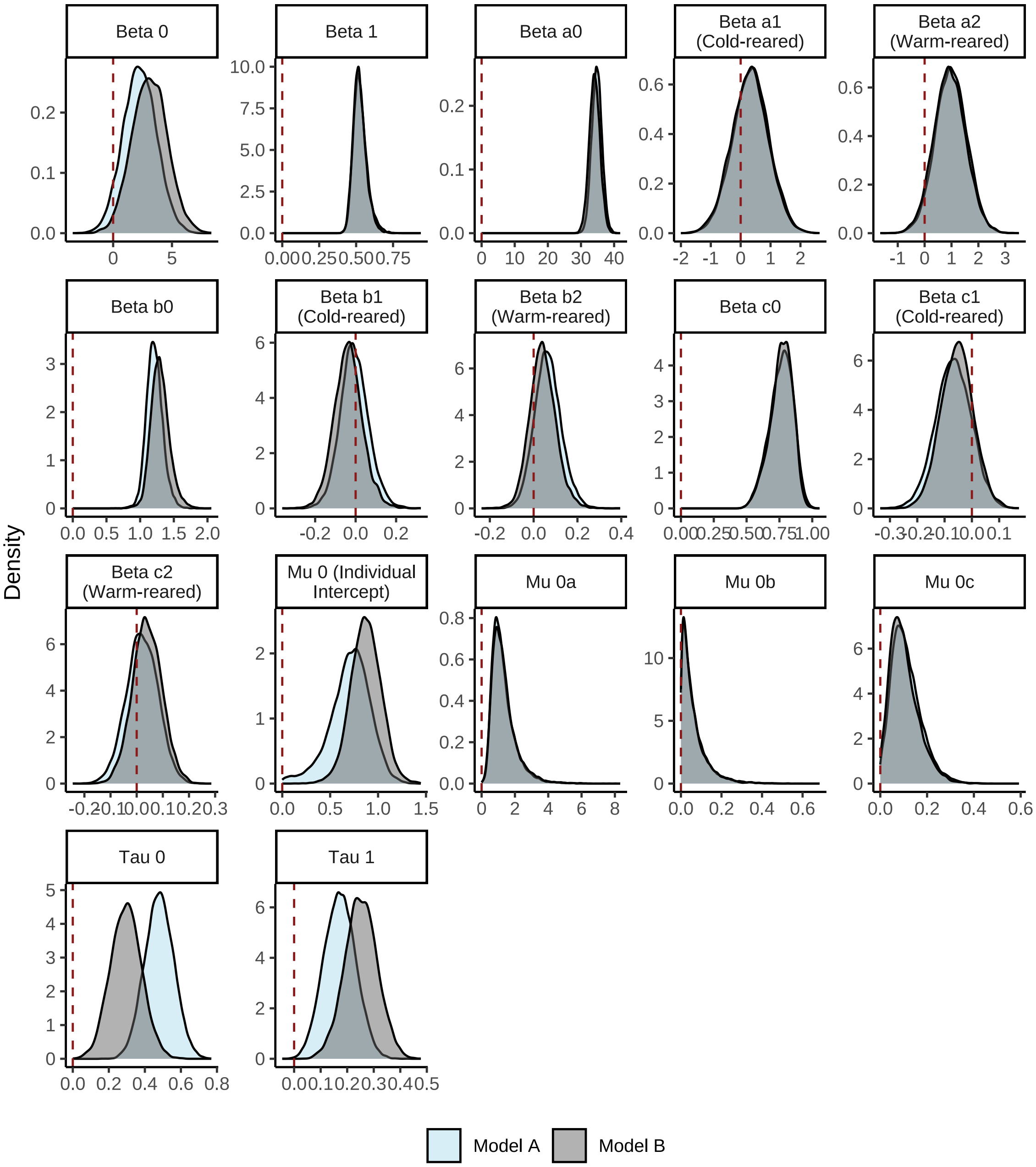


***Figure 32:*** *Comparions of model coefficients derived from two, Bayesian, non-linear models predicting tarsus length (mm) of Japanese quail between 0 and 8 weeks of age. 'Model A' includes one possible outlier (as determined by visually inspecting model residuals), and 'Model B' excludes this outlier. Densities represent posterior densities for each coefficient.*

Some distinctions in variance and error structure emerge after the potential outlier is removed, but all population-level coefficients remain essentially unchanged. As such, our initial model is summarised going forward.

```
as.data.frame(growthModelTarsus) %>%
  pivot_longer(everything(), names_to = "Parameter", values_to = "Values") %>%
  filter(grepl("b_|sd_", Parameter)) %>%
  arrange(Parameter) %>%
```

```
merge(., tribble(
  ~Parameter, ~Par,
  "b_A_Intercept", "Beta a0",
  "b_A_pretreatmentcold", "Beta a1\n(Cold-reared)",
  "b_A_pretreatmentwarm", "Beta a2\n(Warm-reared)",
  "sd_batch__A_Intercept", "Mu 0a",
  "b_B_Intercept", "Beta b0",
  "b_B_pretreatmentcold", "Beta b1\n(Cold-reared)",
  "b_B_pretreatmentwarm", "Beta b2\n(Warm-reared)",
  "sd_batch__B_Intercept", "Mu 0b",
  "b_C_Intercept", "Beta c0",
  "b_C_pretreatmentcold", "Beta c1\n(Cold-reared)",
  "b_C_pretreatmentwarm", "Beta c2\n(Warm-reared)",
  "sd_batch__C_Intercept", "Mu 0c",
  "b_D_Intercept", "Beta 0",
  "b_D_centredMass", "Beta 1",
  "sd_ring__D_Intercept", "Mu 0\n(Individual Intercept)",
  "b_sigma_intercept", "Tau 0",
  "b_sigma_logweekB", "Tau 1"
),
by = "Parameter", all.x = TRUE
) %>%
filter(!is.na(Par)) %>%
ggplot(aes(x = Values)) +
facet_wrap(~Par, scales = "free") +
geom_density() +
geom_vline(
  xintercept = 0, linetype = "dashed",
  colour = "firebrick4"
) +
ylab("Density") +
theme_classic() +
theme(axis.title.x = element_blank())
```

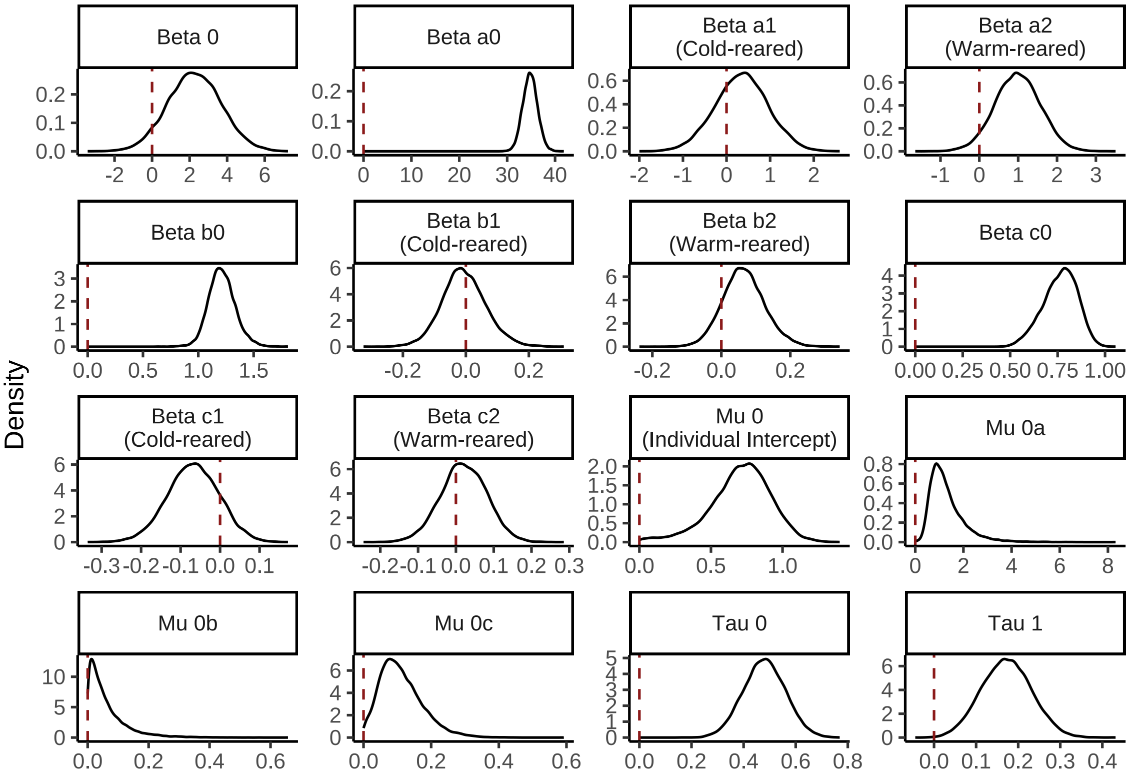


***Figure 33:*** *Density of model coefficients from a Bayesian non-linear model predicting tarsus length (mm) of Japanese quail during growth. Dashed red lines indicate zero values for each coefficient.*

```
# Growth rate (parameter 'c') slowed among cold-reared
# birds, and asymptote evidently raised in warm-reared birds.
# Summarising model outcomes.

caption <- paste0(
  "Coefficients from a Bayesian non-linear effects ",
  "model predicting tarsus length (mm) of Japanese quail as a ",
  "Gompertz function of age in weeks. Coefficients represent ",
  "posterior medians and are estimated from body mass data ",
  "collected weekly between 0 and 8 weeks of age. Credible ",
  "intervals (CIs) represent quantile ",
  "intervals around medians"
)

tarsusGrowthModelTable <-
  merge(
    as.data.frame(growthModelTarsus) %>%
      summarise_all(., .funs = median) %>%
      mutate_all(., .funs = round, 3) %>%
      pivot_longer(everything(),
        names_to = "Parameter",
        values_to = "Values"
      ) %>%
      filter(grepl("b_|sd_", Parameter)) %>%
      arrange(Parameter),
    quantileCIs(growthModelTarsus, cis = c(50, 95)) %>%
      mutate(
        `50\\% CIs` = paste0(
          "[", round(Low_CI_50, digits = 3),
          ", ", round(High_CI_50, digits = 3), "]"
        ),
        `95\\% CIs` = paste0(
```

```
        "[", round(Low_CI_95, digits = 3),
        ", ", round(High_CI_95, digits = 3), "]"
      )
    ) %>%
    select(Parameter, `50\\% CIs`, `95\\% CIs`),
  by = "Parameter"
) %>%
merge(., tribble(
  ~Parameter, ~Par,
  "b_A_Intercept", "Beta a0",
  "b_A_pretreatmentcold", "Beta a1 (Cold-reared)",
  "b_A_pretreatmentwarm", "Beta a2 (Warm-reared)",
  "sd_batch__A_Intercept", "Mu 0a",
  "b_B_Intercept", "Beta b0",
  "b_B_pretreatmentcold", "Beta b1 (Cold-reared)",
  "b_B_pretreatmentwarm", "Beta b2 (Warm-reared)",
  "sd_batch__B_Intercept", "Mu 0b",
  "b_C_Intercept", "Beta c0",
  "b_C_pretreatmentcold", "Beta c1 (Cold-reared)",
  "b_C_pretreatmentwarm", "Beta c2 (Warm-reared)",
  "sd_batch__C_Intercept", "Mu 0c",
  "b_D_Intercept", "Beta 0",
  "b_D_cScaledMass", "Beta 1",
  "sd_ring__D_Intercept", "Mu 0 (Individual Intercept)",
  "b_sigma_intercept", "Tau 0",
  "b_sigma_logweekB", "Tau 1"
),
by = "Parameter", all.x = TRUE
) %>%
select(-Parameter) %>%
select("Parameter" = Par, "Value" = Values, `50\\% CIs`, `95\\% CIs`) %>%
kbl(.,
  longtable = T, booktabs = T, format = "latex",
  escape = FALSE, caption = caption
) %>%
column_spec(column = c(1:10), width = "2.5cm") %>%
kable_styling(latex_options = "striped")

tarsusGrowthModelTable
```

***Table 13:*** *Coefficients from a Bayesian non-linear effects model predicting tarsus length (mm) of Japanese quail as a Gompertz function of age in weeks. Coefficients represent posterior medians and are estimated from body mass data collected weekly between 0 and 8 weeks of age. Credible intervals (CIs) represent quantile intervals around medians*

| Parameter | Value | 50% CIs | 95% CIs |
|---|---|---|---|
| Beta a0 | 34.786 | [33.759, 35.798] | [31.819, 37.843] |
| Beta a1 (Cold-reared) | 0.352 | [-0.056, 0.751] | [-0.846, 1.527] |
| Beta a2 (Warm-reared) | 0.984 | [0.593, 1.379] | [-0.157, 2.126] |
| Beta b0 | 1.208 | [1.134, 1.288] | [0.994, 1.467] |
| Beta b1 (Cold-reared) | -0.009 | [-0.052, 0.039] | [-0.14, 0.14] |
| Beta b2 (Warm-reared) | 0.063 | [0.024, 0.104] | [-0.05, 0.192] |
| Beta c0 | 0.770 | [0.704, 0.828] | [0.572, 0.92] |
| Beta c1 (Cold-reared) | -0.068 | [-0.112, -0.024] | [-0.198, 0.059] |
| Beta c2 (Warm-reared) | 0.020 | [-0.02, 0.062] | [-0.1, 0.138] |
| Beta 1 | 0.523 | [0.496, 0.555] | [0.452, 0.641] |
| Beta 0 | 2.255 | [1.299, 3.228] | [-0.491, 5.068] |
| Tau 0 | 0.480 | [0.425, 0.534] | [0.322, 0.638] |
| Tau 1 | 0.172 | [0.131, 0.213] | [0.058, 0.289] |

| | | | |
|---|---|---|---|
| Mu 0a | 1.136 | [0.817, 1.603] | [0.417, 3.154] |
| Mu 0b | 0.042 | [0.018, 0.083] | [0.002, 0.238] |
| Mu 0c | 0.102 | [0.066, 0.147] | [0.014, 0.261] |
| Mu 0 (Individual Intercept) | 0.724 | [0.585, 0.849] | [0.21, 1.075] |

```
# save_kable(tarsusGrowthModelTable,
# "../tables/tarsusGrowthModelTable.html")
```

Our coefficients indicate that warm-reared individuals (i.e. those raised at 30°C for at least their first three weeks of life) had a higher asymptote (value for $a$) than both those who were reared in mild conditions (20°C) and cold conditions (10°C for at least 3 weeks of life). Similarly, the growth rate of warm-reared individuals appeared higher (or more rapid) than those from our other experimental treatments. Evidence ratios for these differences were next calculated formally, again using the Savage-Dickey density ratio method.

```
caption <- paste0(
  "Pairwise comparisons of Gompertz function ",
  "variables between cold-reared (10°C until at least 3 ",
  "weeks of age), mild-reared (constant 20°C), ",
  "and warm-reared (30°C until at least 3 weeks of age) ",
  "Japanese quail. Gompertz function variables being ",
  "compared are indicated in parenthesis, per hypothesis."
)

pairwiseTarsusGrowthTable <- rbind(
  hypothesis(growthModelTarsus,
    hypothesis = "A_Intercept - (A_Intercept + A_pretreatmentcold) > 0",
    class = "b",
    robust = TRUE
  )$hypothesis,
  hypothesis(growthModelTarsus,
    hypothesis = "(A_Intercept + A_pretreatmentwarm) - A_Intercept > 0 ",
    class = "b",
    robust = TRUE
  )$hypothesis,
  hypothesis(growthModelTarsus,
    hypothesis = paste0(
      "(A_Intercept + A_pretreatmentwarm) - ",
      "(A_Intercept + A_pretreatmentcold) > 0"
    ),
    class = "b",
    robust = TRUE
  )$hypothesis,
  hypothesis(growthModelTarsus,
    hypothesis = "B_Intercept - (B_Intercept + B_pretreatmentcold) > 0",
    class = "b",
    robust = TRUE
  )$hypothesis,
  hypothesis(growthModelTarsus,
    hypothesis = "(B_Intercept + B_pretreatmentwarm) - B_Intercept > 0",
    class = "b",
    robust = TRUE
  )$hypothesis,
  hypothesis(growthModelTarsus,
    hypothesis = paste0(
      "(B_Intercept + B_pretreatmentwarm) - ",
      "(B_Intercept + B_pretreatmentcold) > 0"
    ),
    class = "b",
    robust = TRUE
  )$hypothesis,
  hypothesis(growthModelTarsus,
    hypothesis = "C_Intercept - (C_Intercept + C_pretreatmentcold) > 0",
    class = "b",
```

```
    robust = TRUE
  )$hypothesis,
  hypothesis(growthModelTarsus,
    hypothesis = "(C_Intercept + C_pretreatmentwarm) - C_Intercept > 0",
    class = "b",
    robust = TRUE
  )$hypothesis,
  hypothesis(growthModelTarsus,
    hypothesis = paste0(
      "(C_Intercept + C_pretreatmentwarm) - ",
      "(C_Intercept + C_pretreatmentcold) > 0"
    ),
    class = "b",
    robust = TRUE
  )$hypothesis
) %>%
  mutate(
    `95\\% CIs` = paste0(
      "[", round(CI.Lower, digits = 3), ", ",
      round(CI.Upper, digits = 3), "]"
    ),
    Estimate = round(Estimate, digits = 3),
    Evid.Ratio = round(Evid.Ratio, digits = 3)
  ) %>%
  select(Hypothesis, Estimate, `95\\% CIs`, Evid.Ratio) %>%
  merge(.,
    rbind(
      hypothesis(growthModelTarsus,
        hypothesis = "A_Intercept - (A_Intercept + A_pretreatmentcold) > 0",
        class = "b", alpha = 0.2,
        robust = TRUE
      )$hypothesis,
      hypothesis(growthModelTarsus,
        hypothesis = "(A_Intercept + A_pretreatmentwarm) - A_Intercept > 0 ",
        class = "b", alpha = 0.2,
        robust = TRUE
      )$hypothesis,
      hypothesis(growthModelTarsus,
        hypothesis = paste0(
          "(A_Intercept + A_pretreatmentwarm) - ",
          "(A_Intercept + A_pretreatmentcold) > 0"
        ),
        class = "b", alpha = 0.2,
        robust = TRUE
      )$hypothesis,
      hypothesis(growthModelTarsus,
        hypothesis = "B_Intercept - (B_Intercept + B_pretreatmentcold) > 0",
        class = "b", alpha = 0.2,
        robust = TRUE
      )$hypothesis,
      hypothesis(growthModelTarsus,
        hypothesis = "(B_Intercept + B_pretreatmentwarm) - B_Intercept > 0",
        class = "b", alpha = 0.2,
        robust = TRUE
      )$hypothesis,
      hypothesis(growthModelTarsus,
        hypothesis = paste0(
          "(B_Intercept + B_pretreatmentwarm) - ",
          "(B_Intercept + B_pretreatmentcold) > 0"
        ),
        class = "b", alpha = 0.2,
        robust = TRUE
      )$hypothesis,
      hypothesis(growthModelTarsus,
        hypothesis = "C_Intercept - (C_Intercept + C_pretreatmentcold) > 0",
        class = "b", alpha = 0.2,
        robust = TRUE
```

```
    )$hypothesis,
    hypothesis(growthModelTarsus,
      hypothesis = "(C_Intercept + C_pretreatmentwarm) - C_Intercept > 0",
      class = "b", alpha = 0.2,
      robust = TRUE
    )$hypothesis,
    hypothesis(growthModelTarsus,
      hypothesis = paste0(
        "(C_Intercept + C_pretreatmentwarm) - ",
        "(C_Intercept + C_pretreatmentcold) > 0"
      ),
      class = "b", alpha = 0.2,
      robust = TRUE
    )$hypothesis
  ) %>%
    mutate(
      `50\\% CIs` = paste0(
        "[", round(CI.Lower, digits = 3), ", ",
        round(CI.Upper, digits = 3), "]"
      ),
      Estimate = round(Estimate, digits = 3),
      Evid.Ratio = round(Evid.Ratio, digits = 3)
    ) %>%
    select(Hypothesis, `50\\% CIs`),
  by = "Hypothesis"
) %>%
merge(.,
  tribble(
    ~hypothesis, ~Hypothesis,
    "Cold-reared Asymptote (a) < Mild-reared Asymptote (a)",
    "(A_Intercept-(A_Intercept+A_pretreatmentcold)) > 0",
    "Warm-reared Asymptote (a) > Mild-reared Asymptote (a)",
    "((A_Intercept+A_pretreatmentwarm)-A_Intercept) > 0",
    "Warm-reared Asymptote (a) > Cold-reared Asymptote (a)",
    paste0(
      "((A_Intercept+A_pretreatmentwarm)-",
      "(A_Intercept+A_pretreatmentcold)) > 0"
    ),
    "Cold-reared Displacement (b) < Mild-reared Displacement (b)",
    "(B_Intercept-(B_Intercept+B_pretreatmentcold))",
    "Warm-reared Displacement (b) > Mild-reared Displacement (b)",
    "((B_Intercept+B_pretreatmentwarm)-B_Intercept) > 0",
    "Warm-reared Displacement (b) > Cold-reared Displacement (b)",
    paste0(
      "((B_Intercept+B_pretreatmentwarm)-",
      "(B_Intercept+B_pretreatmentcold)) > 0"
    ),
    "Cold-reared Growth Rate (c) < Mild-reared Growth Rate (c)",
    "(C_Intercept-(C_Intercept+C_pretreatmentcold)) > 0",
    "Warm-reared Growth Rate (c) > Mild-reared Growth Rate (c)",
    "((C_Intercept+C_pretreatmentwarm)-C_Intercept) > 0",
    "Warm-reared Growth Rate (c) > Cold-reared Growth Rate (c)",
    paste0(
      "((C_Intercept+C_pretreatmentwarm)-",
      "(C_Intercept+C_pretreatmentcold)) > 0"
    )
  ),
  by = "Hypothesis"
) %>%
select(
  "Hypothesis" = hypothesis, "Delta" = Estimate,
  `50\\% CIs`, `95\\% CIs`, "Evidence Ratio" = Evid.Ratio
) %>%
kbl(.,
  longtable = T, booktabs = T, escape = FALSE,
  format = "latex", caption = caption
) %>%
```

```
  column_spec(column = c(1:10), width = "2.5cm") %>%
  kable_styling(latex_options = "striped")

pairwiseTarsusGrowthTable
```

***Table 14:*** *Pairwise comparisons of Gompertz function variables between cold-reared (10°C until at least 3 weeks of age), mild-reared (constant 20°C), and warm-reared (30°C until at least 3 weeks of age) Japanese quail. Gompertz function variables being compared are indicated in parenthesis, per hypothesis.*

| Hypothesis | Delta | 50% CIs | 95% CIs | Evidence Ratio |
|---|---|---|---|---|
| Warm-reared Asymptote (a) > Cold-reared Asymptote (a) | 0.641 | [0.178, 1.1] | [-0.259, 1.539] | 7.252 |
| Warm-reared Asymptote (a) > Mild-reared Asymptote (a) | 0.984 | [0.501, 1.474] | [0.029, 1.939] | 21.346 |
| Warm-reared Displacement (b) > Cold-reared Displacement (b) | 0.070 | [0.03, 0.111] | [-0.008, 0.152] | 12.901 |
| Warm-reared Displacement (b) > Mild-reared Displacement (b) | 0.063 | [0.015, 0.114] | [-0.032, 0.169] | 6.306 |
| Warm-reared Growth Rate (c) > Cold-reared Growth Rate (c) | 0.088 | [0.054, 0.124] | [0.019, 0.158] | 53.983 |
| Warm-reared Growth Rate (c) > Mild-reared Growth Rate (c) | 0.020 | [-0.03, 0.072] | [-0.08, 0.119] | 1.713 |
| Cold-reared Asymptote (a) < Mild-reared Asymptote (a) | -0.352 | [-0.85, 0.159] | [-1.35, 0.661] | 0.392 |
| Cold-reared Growth Rate (c) < Mild-reared Growth Rate (c) | 0.068 | [0.012, 0.122] | [-0.037, 0.176] | 5.636 |

```
#save_kable(pairwiseTarsusGrowthTable, "../tables/pairwiseTarsusGrowthTable.html")
```

Whether tarsus lengths at maturity (8 weeks of age; the end of our growth curves) differed among rearing treatments was next analysed using a Bayesian one-way ANOVA. Here, priors on tarsus length (mm) for cold-, mild-, and warm-reared birds were normally-distributed with means of 37.1, 37.5, and 37.9 respectively (following findings of Burness et al, 2013), and standard deviations of 2. Our prior for batch effects (included as a group-level intercept) was again exponential with a lambda of 2.5, and that for our error term was half-student t distributed with three degrees of freedom and location and scale parameters of 0 and 2.5 respectively.

```
growthAnovaTarsus <- brm(
  data = data %>%
    filter(week == 8) %>%
    select(pretreatment, mass, "tarsus" = tarsusLengthMean,
           "batch" = exp) %>%
    drop_na() %>%
    mutate(pretreatment = factor(pretreatment, levels = c("neutral", "cold", "warm"))),
  formula = tarsus ~ 0 + pretreatment + (1|batch),
  prior = c(
```

```
    set_prior("normal(37.5, 2)", class = "b",
              coef = "pretreatmentneutral"),
    set_prior("normal(37.1, 2)", class = "b",
              coef = "pretreatmentcold"),
    set_prior("normal(37.9, 2)", class = "b",
              coef = "pretreatmentwarm"),
    set_prior("exponential(2.5)", class = "sd",
              group = "batch")
  ),
  family = "gaussian",
  seed = 200,
  cores = 4, chains = 4,
  iter = 50000, warmup = 10000, thin = 10,
  control = list(adapt_delta = 0.95, max_treedepth = 13),
  silent = TRUE, refresh = 0,
  file = "./models/tarsusAtMaturityANOVA.Rds"
)

hypotheses <- c(
  "pretreatmentcold - pretreatmentneutral > 0",
  "pretreatmentwarm - pretreatmentneutral > 0",
  "pretreatmentwarm - pretreatmentcold > 0"
)

caption <- paste0("Results from a Bayesian, one-way ANOVA ",
                  "comparing tarsus length (mm) at 8 weeks of Japanese ",
                  "quail reared in the cold (10°C until at least ",
                  "3 weeks of age; n = ",
                  nrow(subset(growthAnovaTarsus$data, pretreatment == "cold")),
                  "), mild temperature (constant 20°C; n = ",
                  nrow(subset(growthAnovaTarsus$data, pretreatment == "neutral")),
                  "), or warmth (30°C until at least 3 weeks of age; n = ",
                  nrow(subset(growthAnovaTarsus$data, pretreatment == "warm")),
                  ")."
                  )

tarsusGrowthAnovaTable <- bind_rows(
  lapply(hypotheses, FUN = function(x) {
    hold <- hypothesis(growthAnovaTarsus,
                       hypothesis = x,
                       class = "b",
                       alpha = 0.5,
                       robust = TRUE)
    return(data.frame(
      "hyp" = x,
      "deltaMass" = hold$hypothesis$Estimate,
      "pProb" = hold$hypothesis$Post.Prob
    ))
  })
) %>%
  mutate("Hypothesis" = c(
    "Cold-Reared Tarsus Length > Mild-Reared Tarsus Length",
    "Warm-Reared Tarsus Length > Mild-Reared Tarsus Length",
    "Cold-Reared Tarsus Length < Warm-Reared Tarsus Length"
  )) %>%
  mutate(deltaMass = round(deltaMass, digits = 3),
         pProb = round(pProb, digits = 3)) %>%
  select(Hypothesis, "Delta Tarsus Length (mm)" = deltaMass,
         "Posterior Probability" = pProb) %>%
  kbl(.,
    longtable = T, booktabs = T, format = "latex",
    caption = caption
  ) %>%
  column_spec(column = c(1:10), width = "2.5cm")  %>%
  kable_styling(latex_options = "striped")

tarsusGrowthAnovaTable
```

***Table 15:*** *Results from a Bayesian, one-way ANOVA comparing tarsus length (mm) at 8 weeks of Japanese quail reared in the cold (10°C until at least 3 weeks of age; n = 40), mild temperature (constant 20°C; n = 33), or warmth (30°C until at least 3 weeks of age; n = 39).*

| Hypothesis | Delta Tarsus Length (mm) | Posterior Probability |
|---|---|---|
| Cold-Reared Tarsus Length > Mild-Reared Tarsus Length | 0.561 | 0.830 |
| Warm-Reared Tarsus Length > Mild-Reared Tarsus Length | 1.188 | 0.981 |
| Cold-Reared Tarsus Length < Warm-Reared Tarsus Length | 0.623 | 0.885 |

```
#save_kable(tarsusGrowthAnovaTable, "../tables/tarsusGrowthAnovaTable.html")
```

The above-described ANOVA compares absolute tarsus length measurements among treatment groups at maturity. As such, it does not: (1) adjust for potential differences in body mass that may occur among treatment groups, and (2) consider how *allometry* may vary by treatment groups. To address such limitations, we then construct: (1) an otherwise identical ANOVA, but with our response variable being residual tarsus length at maturity (derived from a linear model regressing tarsus length [mm] by body mass [g]), and (2) a simple linear model with absolute tarsus length (mm) at maturity as the Gaussian-distributed response, rearing condition (categorical), body mass (g) and the interaction between rearing condition and body mass as population level predictors, and egg batch as a group-level predictor. Priors for rearing condition in our ANOVA are broad (mild-rearing: $\mathcal{N}$[0, 2]; cold-rearing: $\mathcal{N}$[-0.4, 2]; warm-rearing: $\mathcal{N}$[0.4, 2]; egg batch: exponential[$\lambda$ = 2.5]). To simplify prior construction for our linear model, body tarsus length and body mass are first mean-centred (placing our intercept near or at 0); thus, our prior for our model intercept was centred at zero, and normally-distributed with a standard deviation of 2.5. For the effect of body mass on tarsus length, we assumed a skew-normal distribution with $\xi$ equaling 0, $\omega$ equaling 0.25, and $\alpha$ equaling 5. Next, for our effect of cold- and warm-rearing, we assumed normal distributions with means of -0.4 and 0.4 respectively, and standard deviations of 2. Given we had no *a prior* information about how rearing conditions and body mass might interact to shape tarsus length, we assumed broad and normal priors with means of 0 and standard deviations of 0.05 respectively. Finally, for our prior on egg batch and $\epsilon$ (our model error), we used exponential distributions with lambda values of 2.5 and 1 respectively.

```
growthAnovaResidualTarsus <- brm(
  data = data %>%
    filter(week == 8) %>%
    select(pretreatment, mass, "tarsus" = tarsusLengthMean,
           "batch" = exp) %>%
    drop_na() %>%
    mutate(pretreatment = factor(pretreatment,
                                 levels = c("neutral", "cold", "warm"))) %>%
    mutate("residualTarsus" =
          residuals(
            brm(tarsus ~ mass,
              data = .,
              prior = c(
                set_prior("skew_normal(0, 0.25, 5)",
                          class = "b"),
                set_prior("normal(35, 2.5)",
                          class = "Intercept")
              ),
              family = "gaussian",
              iter = 50000, warmup = 10000, cores = 1,
```

```
              chains = 4, thin = 20,
              silent = TRUE, refresh = 0,
              file = "./models/residualTarsusForAnova.Rds"
            ), robust = TRUE
          )[, "Estimate"]
      ),
  formula = residualTarsus ~ 0 + pretreatment + (1|batch),
  prior = c(
    set_prior("normal(0, 2)", class = "b",
              coef = "pretreatmentneutral"),
    set_prior("normal(-0.4, 2)", class = "b",
              coef = "pretreatmentcold"),
    set_prior("normal(0.4, 2)", class = "b",
              coef = "pretreatmentwarm"),
    set_prior("exponential(2.5)", class = "sd",
              group = "batch")
  ),
  family = "gaussian",
  seed = 200,
  cores = 4, chains = 4,
  iter = 50000, warmup = 10000, thin = 10,
  control = list(adapt_delta = 0.95, max_treedepth = 13),
  silent = TRUE, refresh = 0,
  file = "./models/residualTarsusAtMaturityANOVA.Rds"
)

hypotheses <- c(
  "pretreatmentcold - pretreatmentneutral > 0",
  "pretreatmentwarm - pretreatmentneutral > 0",
  "pretreatmentwarm - pretreatmentcold > 0"
)

caption <- paste0("Results from a Bayesian, one-way ANOVA ",
                  "comparing residual tarsus length (mm) at 8 weeks of Japanese ",
                  "quail reared in the cold (10°C until at least ",
                  "3 weeks of age; n = ",
                  nrow(subset(growthAnovaResidualTarsus$data,
                              pretreatment == "cold")),
                  "), mild temperature (constant 20°C; n = ",
                  nrow(subset(growthAnovaResidualTarsus$data,
                              pretreatment == "neutral")),
                  "), or warmth (30°C until at least 3 weeks of age; n = ",
                  nrow(subset(growthAnovaResidualTarsus$data,
                              pretreatment == "warm")),
                  ")."
                  )

residualTarsusGrowthAnovaTable <- bind_rows(
  lapply(hypotheses, FUN = function(x) {
    hold <- hypothesis(growthAnovaResidualTarsus,
                       hypothesis = x,
                       class = "b",
                       alpha = 0.5,
                       robust = TRUE)
    return(data.frame(
      "hyp" = x,
      "deltaMass" = hold$hypothesis$Estimate,
      "pProb" = hold$hypothesis$Post.Prob
    ))
  })
) %>%
  mutate("Hypothesis" = c(
    "Cold-Reared Tarsus Length > Mild-Reared Tarsus Length",
    "Warm-Reared Tarsus Length > Mild-Reared Tarsus Length",
    "Warm-Reared Tarsus Length > Cold-Reared Tarsus Length"
  )) %>%
  mutate(deltaMass = round(deltaMass, digits = 3),
```

```
        pProb = round(pProb, digits = 3)) %>%
  select(Hypothesis, "Delta Tarsus Length (mm)" = deltaMass,
         "Posterior Probability" = pProb) %>%
  kbl(.,
    longtable = T, booktabs = T, format = "latex",
    caption = caption
  ) %>%
  column_spec(column = c(1:10), width = "2.5cm")  %>%
  kable_styling(latex_options = "striped")

residualTarsusGrowthAnovaTable
```

***Table 16:*** *Results from a Bayesian, one-way ANOVA comparing residual tarsus length (mm) at 8 weeks of Japanese quail reared in the cold (10°C until at least 3 weeks of age; n = 40), mild temperature (constant 20°C; n = 33), or warmth (30°C until at least 3 weeks of age; n = 39).*

| Hypothesis | Delta Tarsus Length (mm) | Posterior Probability |
|---|---|---|
| Cold-Reared Tarsus Length > Mild-Reared Tarsus Length | 0.471 | 0.788 |
| Warm-Reared Tarsus Length > Mild-Reared Tarsus Length | 1.007 | 0.961 |
| Warm-Reared Tarsus Length > Cold-Reared Tarsus Length | 0.526 | 0.850 |

```
#save_kable(tarsusGrowthAnovaTable, "../tables/tarsusGrowthAnovaTable.html")
```

```
tarsusAllometryModel <- brm(
  data = data %>%
    filter(week == 8) %>%
    select(pretreatment, mass,
           "tarsus" = tarsusLengthMean,
           "batch" = exp) %>%
    drop_na() %>%
    mutate(pretreatment = factor(pretreatment,
                                 levels = c("neutral", "cold", "warm")
                                 )
           ) %>%
    group_by(pretreatment) %>%
    mutate(mass = mass - mean(mass, na.rm = T),
           tarsus = tarsus - mean(tarsus, na.rm = T)
           ) %>% ungroup(),
  formula = tarsus ~ mass*pretreatment + (1|batch),
  prior = c(
    set_prior("normal(0, 2.5)", class = "Intercept"),
    set_prior("skew_normal(0, 0.25, 5)", class = "b",
              coef = "mass"),
    set_prior("normal(0, 2)", class = "b",
              coef = "pretreatmentcold"),
    set_prior("normal(0, 2)", class = "b",
              coef = "mass:pretreatmentwarm"),
    set_prior("normal(0, 0.05)", class = "b",
              coef = "mass:pretreatmentcold"),
    set_prior("normal(0, 0.05)", class = "b",
              coef = "pretreatmentwarm"),
    set_prior("exponential(2.5)", class = "sd",
              group = "batch"),
    set_prior("exponential(1)", class = "sigma")
  ),
```

```
  family = "gaussian",
  seed = 200,
  cores = 4, chains = 4,
  iter = 50000, warmup = 10000, thin = 10,
  control = list(adapt_delta = 0.95, max_treedepth = 13),
  silent = TRUE, refresh = 0,
  file = "./models/tarsusAllometryModel.Rds"
)

caption <- paste0("Results from a Bayesian linear model ",
                  "predicting tarsus length (mm) of mature Japanese quail ",
                  "as a function of body mass (g), rearing conditions ",
                  "and interactions between body mass and rearing conditions. ",
                  "week old Japanese quail. Physiological measurements ",
                  "Cold rearing indicates ",
                  "post-hatch rearing at 10°C, ",
                  "relative to 20°C (intercept), ",
                  "or 30°C ('warm rearing'). ",
                  "CI indicates quantile ",
                  "intervals and BF indicates Bayes Factors."
)

allometryModelResults <-
  as.data.frame(tarsusAllometryModel) %>%
  summarise_all(., .funs = median) %>%
  pivot_longer(everything(),
    names_to = "Parameter",
    values_to = "Estimate"
  ) %>%
  merge(., quantileCIs(tarsusAllometryModel, cis = c(50, 95)),
    by = "Parameter", all.x = TRUE
  ) %>%
  filter(grepl("b_|sd_", Parameter)) %>%
  rowwise() %>%
  mutate("BF" = ifelse(Estimate < 0,
    (2 * mean(as.data.frame(
      tarsusAllometryModel
    )[, Parameter] <= 0)) /
      (2 * mean(as.data.frame(
        tarsusAllometryModel
      )[, Parameter] >= 0)),
    (2 * mean(as.data.frame(
      tarsusAllometryModel
    )[, Parameter] >= 0)) /
      (2 * mean(as.data.frame(
        tarsusAllometryModel
      )[, Parameter] <= 0))
  )) %>%
  ungroup() %>%
  mutate(
    "Estimate" = round(Estimate, digits = 4),
    "BF" = round(BF, digits = 4),
    "N" = nrow(tarsusAllometryModel$data)
  ) %>%
  mutate("Parameter" = ifelse(grepl("b_", Parameter),
    gsub("b_", "", Parameter),
    gsub(
      "Intercept", "batch",
      gsub(".*__", "", Parameter)
    )
  )) %>%
  merge(., tribble(
    ~Parameter, ~parameter, ~level,
    "Intercept", "Intercept", "A",
    "mass", "Body Mass (g)", "B",
    "mass:pretreatmentcold", "Body Mass:Cold Rearing", "C",
    "mass:pretreatmentwarm", "Body Mass:Warm Rearing", "D",
```

```
    "pretreatmentcold", "Cold Rearing", "E",
    "pretreatmentwarm", "Warm Rearing", "F",
    "batch", "Egg Batch [mu]", "G"
  ),
  by = "Parameter"
  ) %>%
  mutate(
    `50\\% HDI` = paste0("(", paste(
      round(Low_CI_50, digits = 4),
      round(High_CI_50, digits = 4),
      sep = ", "
    ), ")"),
    `95\\% HDI` = paste0("(", paste(
      round(Low_CI_95, digits = 4),
      round(High_CI_95, digits = 4),
      sep = ", "
    ), ")")
  ) %>%
  select(-c(Low_CI_50, High_CI_50, Low_CI_95, High_CI_95)) %>%
  select(
    "Parameter" = "parameter", N,
    Estimate, `50\\% HDI`, `95\\% HDI`, BF, level
  ) %>%
  arrange(level) %>%
  select(-c(level)) %>%
  kbl(.,
    longtable = T, booktabs = T, format = "latex", escape = FALSE,
    caption = caption
  ) %>%
  column_spec(column = c(1:2), width = "2.2cm") %>%
  column_spec(column = c(3:10), width = "1.9cm") %>%
  kable_styling(latex_options = "striped")

allometryModelResults
```

***Table 17:*** *Results from a Bayesian linear model predicting tarsus length (mm) of mature Japanese quail as a function of body mass (g), rearing conditions and interactions between body mass and rearing conditions. week old Japanese quail. Physiological measurements Cold rearing indicates post-hatch rearing at 10°C, relative to 20°C (intercept), or 30°C ('warm rearing'). CI indicates quantile intervals and BF indicates Bayes Factors.*

| Parameter | N | Estimate | 50% HDI | 95% HDI | BF |
|---|---|---|---|---|---|
| Intercept | 112 | 0.0617 | (-0.2014, 0.3343) | (-0.8236, 1.0356) | 1.2844 |
| Body Mass (g) | 112 | 0.0332 | (0.0245, 0.042) | (0.0074, 0.059) | 169.2128 |
| Body Mass:Cold Rearing | 112 | -0.0277 | (-0.0381, -0.0179) | (-0.0583, 0.0022) | 27.8288 |
| Body Mass:Warm Rearing | 112 | -0.0029 | (-0.0145, 0.0089) | (-0.0363, 0.031) | 1.3061 |
| Cold Rearing | 112 | -0.3775 | (-0.6805, -0.0635) | (-1.2488, 0.5173) | 3.7747 |
| Warm Rearing | 112 | -0.0033 | (-0.0366, 0.0307) | (-0.1006, 0.0952) | 1.1097 |
| Egg Batch [mu] | 112 | 0.5445 | (0.3433, 0.7866) | (0.0487, 1.4942) | Inf |

```
allometryPlot <- expand.grid(
  "mass" = seq(min(tarsusAllometryModel$data$mass),
    max(tarsusAllometryModel$data$mass),
    by = 1
  ),
  "pretreatment" = unique(tarsusAllometryModel$data$pretreatment)
) %>%
```

```
  mutate(
    "tarsus" = predict(tarsusAllometryModel,
      newdata = .,
      re_form = NA
    )[, "Estimate"],
    "tarsusSE" = predict(tarsusAllometryModel,
      newdata = .,
      re_form = NA
    )[, "Est.Error"]
  ) %>%
  merge(., subset(data, week == 8) %>%
          group_by(pretreatment) %>%
          summarise("deltaMass" = mean(mass, na.rm = T),
                    "deltaTarsus" = mean(tarsusLengthMean, na.rm = T)
                    ) %>%
          drop_na(),
        by = "pretreatment", all.x = TRUE
        ) %>%
  mutate(
    mass = mass + deltaMass,
    tarsus = tarsus + deltaTarsus,
    pretreatment = factor(pretreatment,
      levels = c("cold", "neutral", "warm")
    )
  ) %>%
  ggplot(aes(
    x = mass, y = tarsus, fill = pretreatment,
    linetype = pretreatment
  )) +
  geom_ribbon(aes(ymin = tarsus - tarsusSE, ymax = tarsus + tarsusSE),
    alpha = 0.5
  ) +
  geom_point(data = subset(data, week == "8") %>%
    select(pretreatment, mass,
      "tarsus" = tarsusLengthMean
    ) %>%
    drop_na() %>%
    mutate(pretreatment = factor(pretreatment,
      levels = c("cold", "neutral", "warm")
    )), pch = 21, colour = "black", size = 2, alpha = 0.5) +
  geom_line(colour = "black") +
  xlab("Body Mass (g)") +
  ylab("Tarsus Length (mm)") +
  scale_fill_manual(
    values = c("#7BB4E3", "black", "#CD5C5C"),
    name = "Rearing\nConditions",
    labels = c("Cold (10°C)", "Mild (20°C)", "Warm (30°C)")
  ) +
  scale_linetype_manual(
    values = c("solid", "dashed", "dotted"),
    name = "Rearing\nConditions",
    labels = c("Cold (10°C)", "Mild (20°C)", "Warm (30°C)")
  ) +
  theme_classic() +
  theme(axis.title = element_text(family = "Noto Sans"),
        axis.text = element_text(family = "Noto Sans"),
        legend.title = element_text(family = "Noto Sans"),
        legend.text = element_text(family = "Noto Sans")
        )

showtext_auto(enable = TRUE)
allometryPlot
```

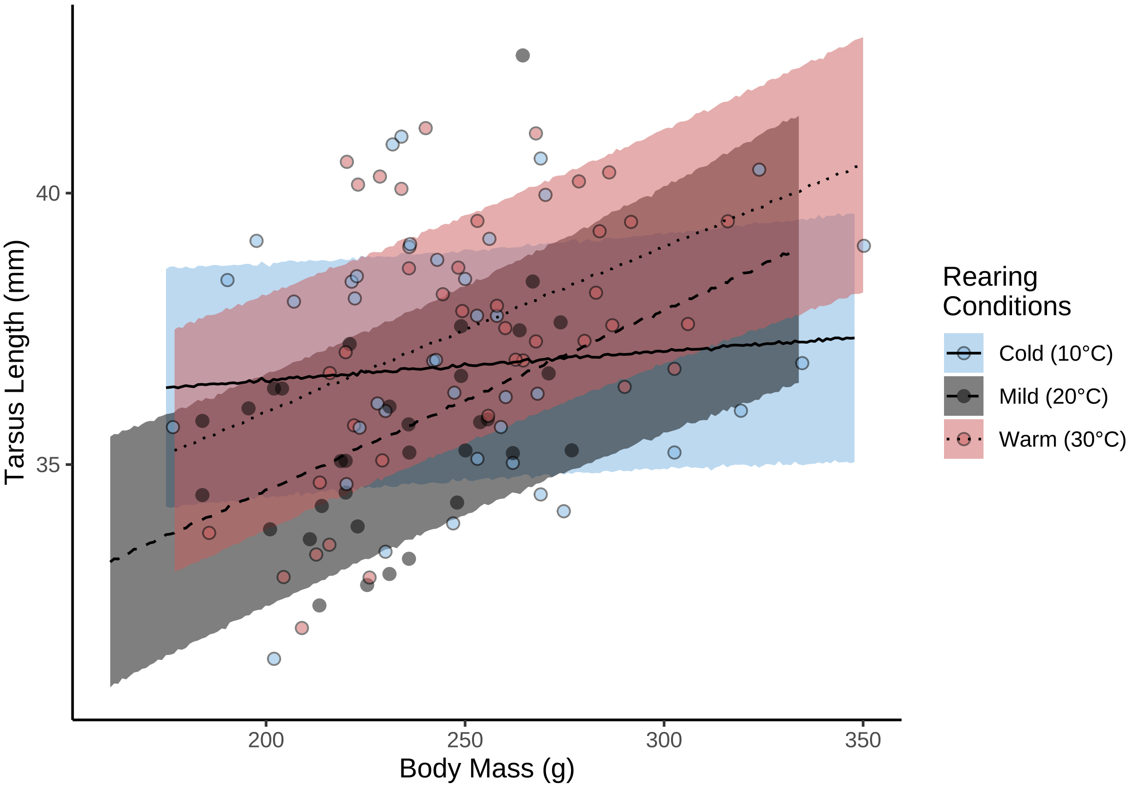


***Figure 34:*** *Effect of body mass (g) and rearing temperature on tarsus length (mm) in eight week old Japanese quail. Rearing temperatures were applied from hatching to at least three weeks of age, after which, temperatures to switched to 20°C in approximately two-thirds half of individuals. Lines represent predicted relationships from a Bayesian mixed effects model and ribbons represent +/- one standard deviation around predicted relationships.*

```
ggsave("../plots/allometryPlot.pdf",
  dpi = 800, width = 8, height = 8,
  allometryPlot
)
showtext_auto(enable = FALSE)
```

Similar to our analyses regarding mass gain during growth, we again tested whether continuous exposure to cold (10°C) or warm (30°C) conditions until maturity (8 weeks of age) differentially shifted appendage elongation trajectories relative to more transient exposure (at least 3 weeks). This is acheived by repeating our above analysis but while only including individuals exposed to their given thermal treatment from hatch until 8 weeks of age. All priors remain the same as above.

```
initFunction <- function(chain_id = 1) {
  list(
    "b_A" = c(rnorm(1, 37.5, 2.5), -0.35, 0.35),
    "b_B" = c(1.35, 0, 0),
    "b_C" = c(0.5, 0, 0),
    "b_D" = c(rnorm(1, 0, 1.5),
              rskew_normal(1, xi = 0.5, omega = 0.15, alpha = 5)),
    "b_sigma" = rskew_normal(2, xi = 0, omega = 0.25, alpha = 10),
    "sd_1" = rexp(1, 0.5)
  )
}

initList <- lapply(1:4, initFunction)
```

```
# Executing model

growthModelTarsusStrict <- brm(
  data = data %>%
    rename("batch" = exp) %>%
    filter(week <= 8 & batch == "C") %>%
    mutate(pretreatment = factor(pretreatment,
                                 levels = c("cold", "warm")
    )) %>%
    select(ring, pretreatment, week, mass,
           "tarsus" = tarsusLengthMean) %>%
    distinct() %>%
    mutate(
      weekB = week + 1
    ) %>%
    group_by(week) %>%
    mutate(cScaledMass = (mass - mean(mass, na.rm = T)) /
             sd(mass, na.rm = T)) %>%
    ungroup(),
  formula = bf(tarsus ~ A * exp(-B * exp(-C * week)) + D,
               A ~ 1 + pretreatment,
               B ~ 1 + pretreatment,
               C ~ 1 + pretreatment,
               D ~ 1 + cScaledMass + (1 | ring),
               sigma ~ 0 + intercept + log(weekB),
               nl = TRUE
  ),
  prior = c(
    set_prior("normal(37.1, 2.5)",
              class = "b", coef = "Intercept",
              nlpar = "A"
    ),
    set_prior("normal(0.8, 2)",
              class = "b", coef = "pretreatmentwarm",
              nlpar = "A"
    ),
    set_prior("skew_normal(0.6, 0.5, 2.5)",
              class = "b", coef = "Intercept",
              nlpar = "B"
    ),
    set_prior("normal(0, 0.5)",
              class = "b", coef = "pretreatmentwarm",
              nlpar = "B"
    ),
    set_prior("skew_normal(0.5, 0.1, 2.5)",
              class = "b", coef = "Intercept",
              nlpar = "C"
    ),
    set_prior("normal(0, 0.25)",
              class = "b", coef = "pretreatmentwarm",
              nlpar = "C"
    ),
    set_prior("normal(0, 1.5)", class = "b",
              coef = "Intercept", nlpar = "D"),
    set_prior("skew_normal(0.5, 0.15, 5)", class = "b",
              coef = "cScaledMass", nlpar = "D"),
    set_prior("exponential(1.5)", class = "sd",
              coef = "Intercept", group = "ring", nlpar = "D"),
    set_prior("skew_normal(0, 0.25, 10)", class = "b",
              coef = "intercept", dpar = "sigma"),
    set_prior("skew_normal(0, 0.25, 10)", class = "b",
              coef = "logweekB", dpar = "sigma")
  ),
  family = "gaussian",
  seed = 100,
  cores = 4, chains = 4,
  init = initList,
```

```
  iter = 50000, warmup = 10000, thin = 10,
  control = list(adapt_delta = 0.95, max_treedepth = 14),
  silent = TRUE, refresh = 0,
  file = "./models/growthModelTarsusStrict.Rds"
  )

ggarrange(
  ggplot(data = data.frame("Rhat" = brms::rhat(growthModelTarsusStrict)),
         aes(x = Rhat)) +
    geom_density() +
    theme_classic() +
    xlab(
      TeX('$\\hat{R}$')
    ) +
    ylab("Density"),
  ggplot(data = data.frame("Neff" = brms::neff_ratio(growthModelTarsusStrict)),
         aes(x = Neff)) +
    geom_density() +
    theme_classic() +
    xlab(
      TeX('$N_{eff}/N~Ratio$')
    ) +
    ylab("Density")
)
```

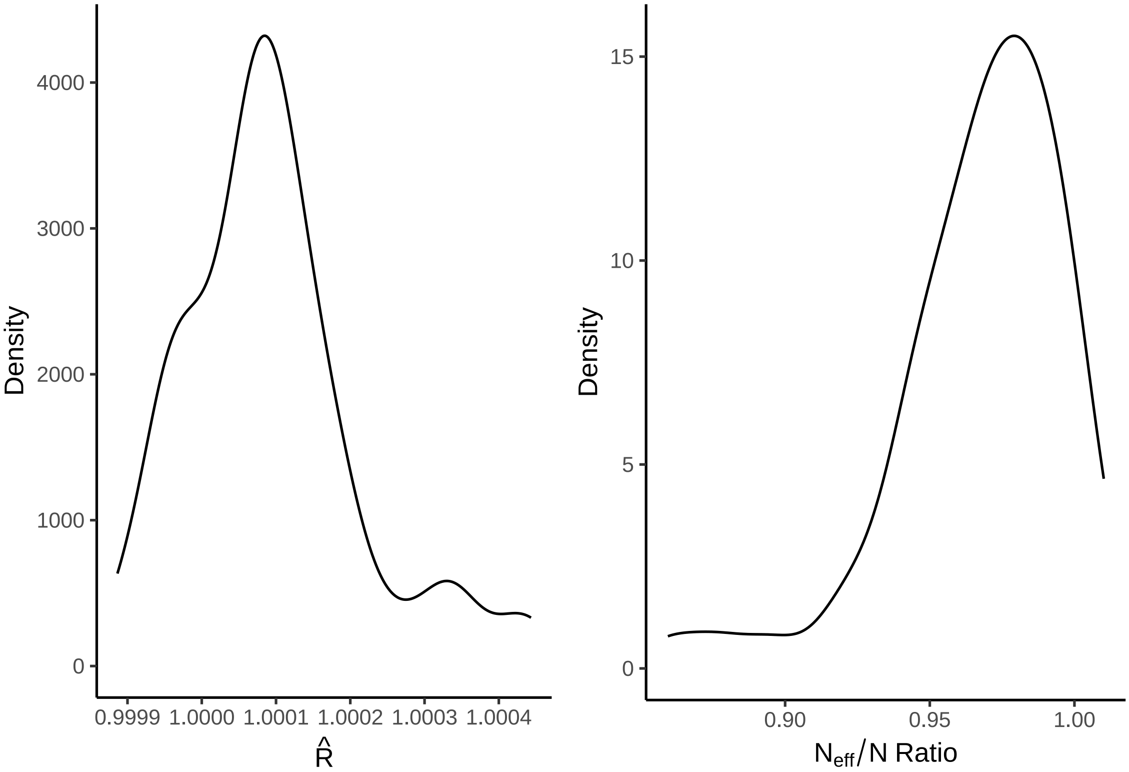


***Figure 35:*** *Gelman-Rubin statistics ($\hat{R}$) and ratio of effective samples sizes by samples sizes per parameter from a Bayesian non-linear model estimating tarsus length (mm) among Japanese quail. Here, only quail exposed to their assigned rearing conditions until 8 weeks of age are included.*

```
# No obvious concerns.
```

Residuals of this new model are visualised and evaluated.

```
p1 <- growthModelTarsusStrict$data %>%
  mutate("residuals" = residuals(growthModelTarsusStrict,
                                method = "posterior_predict",
                                type = "pearson",
                                robust = TRUE)[, "Estimate"]) %>%
  ggplot(aes(x = residuals)) +
  geom_density(colour = "black", fill = "white") +
  xlab("Pearson Residuals") +
  ylab("Density") +
  theme_classic()

p2 <- growthModelTarsusStrict$data %>%
  mutate(
    "residuals" =
      residuals(growthModelTarsusStrict,
      method = "posterior_predict",
      type = "pearson",
      robust = TRUE
    )[, "Estimate"],
    "fitted" = fitted(growthModelTarsusStrict,
                      robust = TRUE)[, "Estimate"]
  ) %>%
  ggplot(aes(x = fitted, y = residuals)) +
  geom_point(colour = "black", pch = 21, size = 2,
             fill = "grey75", alpha = 0.5) +
  ylab("Pearson Residuals") +
  xlab("Fitted Values (mm)") +
  theme_classic()

p3 <- growthModelTarsusStrict$data %>%
  mutate("residuals" =
           residuals(growthModelTarsusStrict,
                     method = "posterior_predict",
                     type = "pearson",
                     robust = TRUE
  )[, "Estimate"]) %>%
  ggplot(aes(x = week, y = residuals)) +
  geom_point(
    colour = "black", pch = 21, size = 2, fill = "grey75", alpha = 0.5,
    position = position_jitter(width = 0.25)
  ) +
  stat_summary(geom = "errorbar", fun.data = "mean_se",
               colour = "black", width = 0.25) +
  stat_summary(geom = "point", fun = "mean", pch = 21,
               colour = "black", fill = "white", size = 4) +
  ylab("Pearson Residuals") +
  xlab("Age (weeks)") +
  theme_classic()

p4 <- growthModelTarsusStrict$data %>%
  mutate("residuals" =
           residuals(growthModelTarsusStrict,
                     method = "posterior_predict",
                     type = "pearson",
                     robust = TRUE
  )[, "Estimate"]) %>%
  ggplot(aes(x = cScaledMass, y = residuals)) +
  geom_point(
    colour = "black", pch = 21, size = 2, fill = "grey75", alpha = 0.5,
    position = position_jitter(width = 0.25)
  ) +
  stat_summary(geom = "errorbar", fun.data = "mean_se",
               colour = "black", width = 0.25) +
  stat_summary(geom = "point", fun = "mean", pch = 21,
               colour = "black", fill = "white", size = 4) +
  ylab("Pearson Residuals") +
  xlab("Centred and Scaled Body Mass") +
```

```
  theme_classic()

p5 <- growthModelTarsusStrict$data %>%
  mutate("residuals" =
           residuals(growthModelTarsusStrict,
                     method = "posterior_predict",
                     type = "pearson",
                     robust = TRUE
  )[, "Estimate"]) %>%
  mutate(pretreatment = str_to_title(pretreatment)) %>%
  mutate(pretreatment = ifelse(pretreatment == "Cold",
                               "Cold\n(10°C)",
    ifelse(pretreatment == "Neutral", "Mild\n(20°C)",
      "Warm\n(30°C)"
    )
  )) %>%
  mutate(pretreatment = factor(pretreatment,
                               levels = c("Cold\n(10°C)",
                                          "Mild\n(20°C)",
                                          "Warm\n(30°C)"))) %>%
  ggplot(aes(x = pretreatment, y = residuals)) +
  geom_point(
    colour = "black", pch = 21, size = 2, fill = "grey75", alpha = 0.5,
    position = position_jitter(width = 0.25)
  ) +
  stat_summary(geom = "errorbar", fun.data = "mean_se",
               colour = "black", width = 0.25) +
  stat_summary(geom = "point", fun = "mean", pch = 21,
               colour = "black", fill = "white", size = 4) +
  ylab("Pearson Residuals") +
  xlab("Rearing Conditions)") +
  theme_classic()

((p1 + p2) / (p3 + p4)) + p5
```

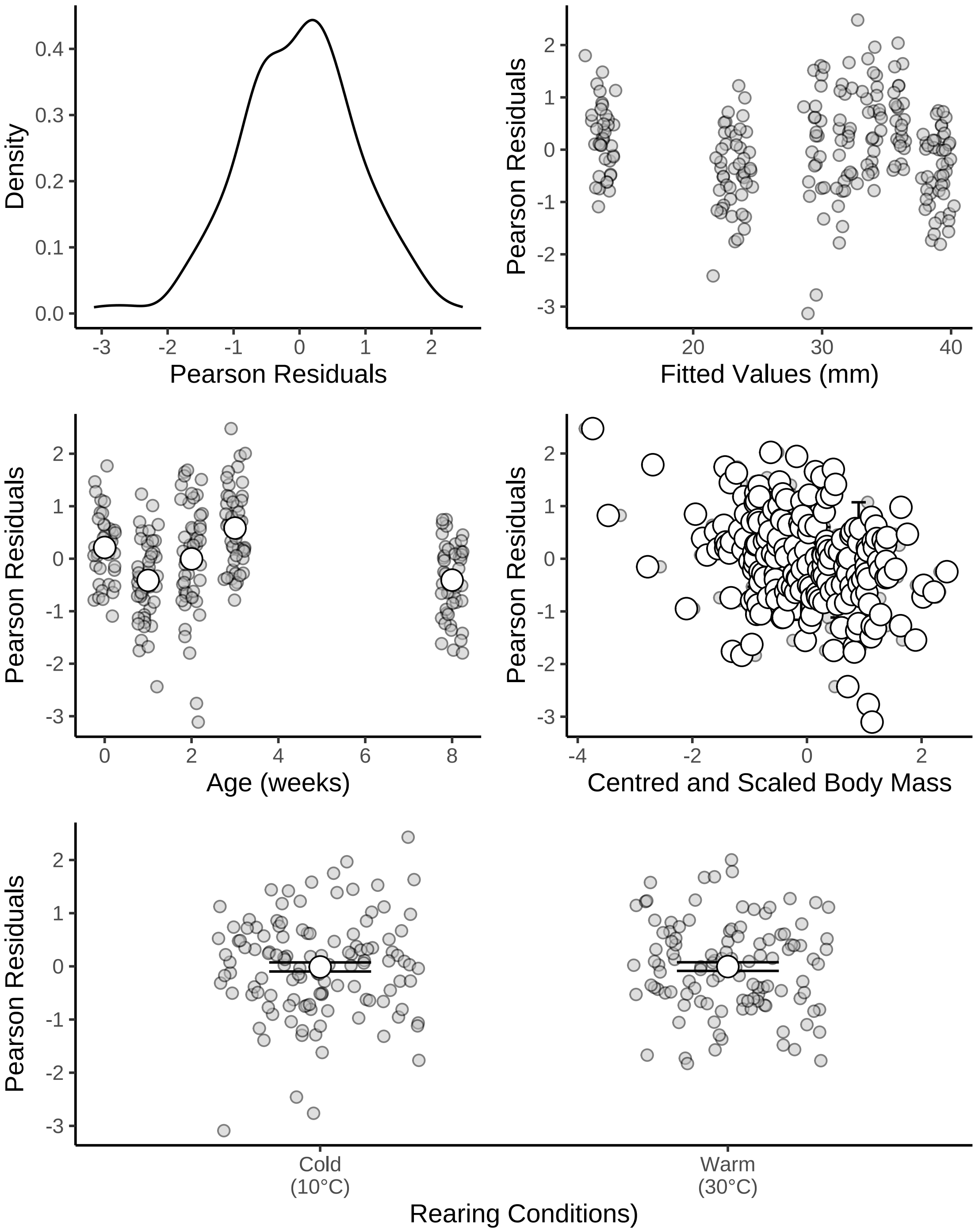


***Figure 36:*** *Density and distributions of Pearson residuals across fitted values and model predictors (including age and rearing conditions). All residuals pertain to those drawn from a Bayesian non-linear model predicting tarsus length (mm) during growth in Japanese quail. Pearson residuals are shown rather than ordinary residuals to correct for the age-dependence of model error. Here, only quail exposed to their assigned rearing conditions until 8 weeks of age were included in our model.*

```
# Again, little concern.
# Summarising sample sizes

caption <- paste0("Number of tarsus length measurements (samples) ",
```

```
                  "drawn from Japanese quail between 0 and ",
                  "8 weeks of age (maturity) across three distict ",
                  "thermal rearing conditions.")

growthModelTarsusStrict$data %>%
  group_by(week, pretreatment) %>%
  count(name = "Samples (n)") %>%
  mutate(pretreatment = ifelse(pretreatment == "cold",
                               "Cold (10°C)",
                               ifelse(pretreatment == "neutral",
                                      "Mild (20°C)", "Warm (30°C)")
                               )
         ) %>%
  arrange(pretreatment, week) %>%
  rename("Age (Weeks)" = week,
         "Rearing Conditions" = pretreatment) %>%
  kbl(.,
    longtable = T, booktabs = T,
    caption = caption
  ) %>%
  kable_styling(latex_options = "striped")
```

***Table 18:*** *Number of tarsus length measurements (samples) drawn from Japanese quail between 0 and 8 weeks of age (maturity) across three distict thermal rearing conditions.*

| Age (Weeks) | Rearing Conditions | Samples (n) |
|---|---|---|
| 0 | Cold (10°C) | 23 |
| 1 | Cold (10°C) | 23 |
| 2 | Cold (10°C) | 24 |
| 3 | Cold (10°C) | 25 |
| 8 | Cold (10°C) | 24 |
| 0 | Warm (30°C) | 16 |
| 1 | Warm (30°C) | 24 |
| 2 | Warm (30°C) | 24 |
| 3 | Warm (30°C) | 24 |
| 8 | Warm (30°C) | 23 |

Our tarsus elongation curves are replotted.

```
growthCurveTarsusStrict <- with(
  growthModelTarsusStrict$data,
  expand.grid(
    "week" = seq(0, 8, by = 0.1),
    "pretreatment" = c("cold", "warm"),
    "cScaledMass" = 0
  )
) %>%
  mutate("weekB" = week + 1) %>%
  mutate(
    "Fit" = predict(growthModelTarsusStrict, re_form = NA,
                    newdata = .,
                    robust = TRUE)[, "Estimate"],
    "SE" = predict(growthModelTarsusStrict, re_form = NA,
                   newdata = .,
                   robust = TRUE)[, "Est.Error"]
  ) %>%
  mutate(
    "LCL" = Fit - SE,
    "UCL" = Fit + SE
  ) %>%
  mutate(pretreatment = ifelse(pretreatment == "cold",
                               "Cold (10°C)",
                               "Warm (30°C)")
```

```
        ) %>%
  mutate(pretreatment = factor(pretreatment,
                               levels =
                                 c("Cold (10°C)",
                                  "Warm (30°C)")
  )
  ) %>%
  ggplot(aes(x = week, y = Fit,
   fill = pretreatment, linetype = pretreatment)) +
  geom_ribbon(aes(x = week, ymin = LCL, ymax = UCL),
    colour = NA, size = 0.25, alpha = 0.3
  ) +
  geom_line(colour = "black", alpha = 0.7) +
  stat_summary(
    data = growthModelTarsusStrict$data %>%
    mutate(pretreatment = ifelse(pretreatment == "cold",
                               "Cold (10°C)",
                               "Warm (30°C)")
          ) %>%
    mutate(pretreatment = factor(pretreatment,
                               levels =
                                 c("Cold (10°C)",
                                  "Warm (30°C)")
  )
  ),
    aes(x = week, y = tarsus),
    geom = "errorbar", fun.data = "mean_cl_boot",
    colour = "black", alpha = 0.7, width = 0.25,
    position = position_dodge(width = 0.15)
  ) +
  stat_summary(
    data = growthModelTarsusStrict$data %>%
    mutate(pretreatment = ifelse(pretreatment == "cold",
                               "Cold (10°C)",
                               "Warm (30°C)")
          ) %>%
    mutate(pretreatment = factor(pretreatment,
                               levels =
                                 c("Cold (10°C)",
                                  "Warm (30°C)")
  )
  ),
    aes(x = week, y = tarsus),
    geom = "point", fun = "mean", pch = 21, size = 3,
    colour = "black", alpha = 0.7,
    position = position_dodge(width = 0.15)
  ) +
  theme_classic() +
  scale_fill_manual(values = c("#7BB4E3", "#CD5C5C"),
                    name = "Rearing\nConditions") +
  scale_linetype_manual(values = c("dotted", "solid", "dashed"),
                        name = "Rearing\nConditions") +
  xlab("Age (weeks)") +
  ylab("Tarsus Length (mm)")

showtext_auto()
growthCurveTarsusStrict
```

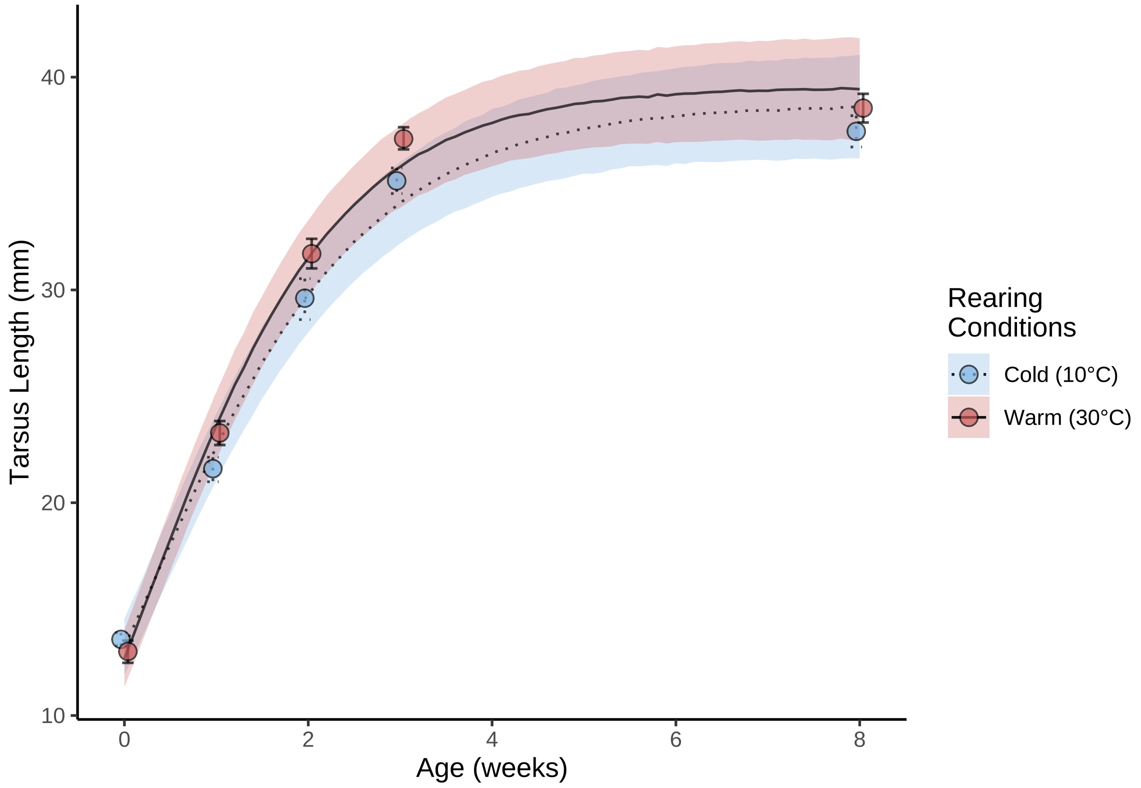


***Figure 37:*** *Tarsus length (mm) elongation curves of Japanese quail reared in the cold (10°C), mild conditions (20°C), or the warm (30°C) until at least 8 weeks of age. Dots represent mean values and errorbars represent quantile-based 95% credible intervals. Lines represent estimated trends in growth from a Bayesian non-linear model, while ribbons represent confidence around trends (± one standard error).*

```
ggsave("../plots/growthCurveTarsusStrict.jpg", dpi = 800,
       width = 7, height = 6.5,
       growthCurveTarsusStrict)
showtext_auto(enable = "FALSE")

# Summarising growth curve parameters

caption <- paste0('Coefficients from a Bayesian non-linear effects ',
                  "model predicting tarsus length (mm) of Japanese quail as a ",
                  "Gompertz function of age in weeks. Coefficients represent ",
                  "posterior medians and are estimated from body mass data ",
                  "collected weekly between 0 and 8 weeks of age. Credible ",
                  "intervals (CIs) represent quantile ",
                  'intervals around medians Here, only individuals ',
                  'who experienced continuous exposure to thermal treatment ',
                  'until maturity (8 weeks of age) were included in our ',
                  'analysis.'
)

tarsusGrowthModelStrictTable <-
  merge(
    as.data.frame(growthModelTarsusStrict) %>%
      summarise_all(., .funs = median) %>%
      mutate_all(., .funs = round, 3) %>%
      pivot_longer(everything(), names_to = "Parameter",
                   values_to = "Values") %>%
      filter(grepl("b_|sd_", Parameter)) %>%
      arrange(Parameter),
    quantileCIs(growthModelTarsusStrict, cis = c(50, 95)),
```

```
    by = "Parameter"
  ) %>%
    mutate(`50\\% CI` = paste0("[",
                               round(Low_CI_50, digits = 3),
                               ", ",
                               round(High_CI_50, digits = 3),
                               "]"),
           `95\\% CI` = paste0("[",
                               round(Low_CI_95, digits = 3),
                               ", ",
                               round(High_CI_95, digits = 3),
                               "]"),
           ) %>%
    select(Parameter, Values, `50\\% CI`, `95\\% CI`) %>%
  merge(., tribble(
    ~Parameter, ~Par,
    "b_A_Intercept", "Beta a0",
    "b_A_pretreatmentwarm", "Beta a2 (Warm-reared)",
    "b_B_Intercept", "Beta b0",
    "b_B_pretreatmentwarm", "Beta b2 (Warm-reared)",
    "b_C_Intercept", "Beta c0",
    "b_C_pretreatmentwarm", "Beta c2 (Warm-reared)",
    "b_D_Intercept", "Beta 0",
    "b_D_cScaledMass", "Beta 1",
    "sd_ring__D_Intercept", "Mu 0 (Individual Intercept)",
    "b_sigma_intercept", "Tau 0",
    "b_sigma_logweekB", "Tau 1"
  ),
  by = "Parameter", all.x = TRUE
  ) %>%
  select(-Parameter) %>%
  select("Parameter" = Par, "Value" = Values, `50\\% CI`, `95\\% CI`) %>%
  kbl(., longtable = T, booktabs = T, format = "latex",
      escape = FALSE, caption = caption) %>%
  column_spec(column = c(1:10), width = "2.5cm")  %>%
  kable_styling(latex_options = "striped")

tarsusGrowthModelStrictTable
```

***Table 19:*** *Coefficients from a Bayesian non-linear effects model predicting tarsus length (mm) of Japanese quail as a Gompertz function of age in weeks. Coefficients represent posterior medians and are estimated from body mass data collected weekly between 0 and 8 weeks of age. Credible intervals (CIs) represent quantile intervals around medians Here, only individuals who experienced continuous exposure to thermal treatment until maturity (8 weeks of age) were included in our analysis.*

| Parameter | Value | 50% CI | 95% CI |
|---|---|---|---|
| Beta a0 | 36.244 | [35.284, 37.18] | [33.49, 39.036] |
| Beta a2 (Warm-reared) | 0.798 | [0.386, 1.218] | [-0.421, 1.995] |
| Beta b0 | 1.216 | [1.159, 1.281] | [1.061, 1.421] |
| Beta b2 (Warm-reared) | 0.074 | [0.045, 0.103] | [-0.009, 0.165] |
| Beta c0 | 0.729 | [0.708, 0.749] | [0.671, 0.789] |
| Beta c2 (Warm-reared) | 0.105 | [0.079, 0.131] | [0.029, 0.184] |
| Beta 1 | 0.487 | [0.464, 0.511] | [0.423, 0.567] |
| Beta 0 | 2.487 | [1.585, 3.401] | [-0.15, 5.097] |
| Tau 0 | 0.253 | [0.182, 0.324] | [0.061, 0.465] |
| Tau 1 | 0.278 | [0.224, 0.332] | [0.117, 0.433] |
| Mu 0 (Individual Intercept) | 0.688 | [0.568, 0.806] | [0.256, 1.044] |

And we proceed by comparing: (1) growth curve parameters in a pairwise manner (as above), and (2) mean tarsus lengths at maturity using a Bayesian permutation test. Our prior for the difference between tarsus lengths of warm-reared and cold-reared quail in our permutation test is set as normal with a mean of 0.8 and standard deviation of 1 (according to results of Burness et al, 2013).

```
caption <- paste0('Pairwise comparisons of Gompertz function ',
                  "variables between cold-reared (10°C until 8 ",
                  "weeks of age), mild-reared (constant 20°C), ",
                  "and warm-reared (30°C until 8 weeks of age) ",
                  "Japanese quail. Gompertz function variables being ",
                  "compared are indicated in parenthesis, per hypothesis."
)

pairwiseTarsusGrowthTableStrict <- rbind(
  hypothesis(growthModelTarsusStrict,
             hypothesis = "(A_Intercept + A_pretreatmentwarm) - A_Intercept > 0",
             class = "b",
             robust = TRUE)$hypothesis,
  hypothesis(growthModelTarsusStrict,
             hypothesis = "(B_Intercept + B_pretreatmentwarm) - B_Intercept > 0",
             class = "b",
             robust = TRUE)$hypothesis,
  hypothesis(growthModelTarsusStrict,
             hypothesis = "(C_Intercept + C_pretreatmentwarm) - C_Intercept > 0",
             class = "b",
             robust = TRUE)$hypothesis
  ) %>%
  cbind(., data.frame("hypothesis" = c(
    "Warm-reared Asymptote (a) > Cold-reared Asymptote (a)",
    "Warm-reared Displacement (b) > Cold-reared Displacement (b)",
    "Warm-reared Growth Rate (c) > Cold-reared Growth Rate (c)"))
    ) %>%
  mutate(`95\\% CIs` = paste0("[", round(CI.Lower, digits = 3), ", ",
                              round(CI.Upper, digits = 3), "]"),
         Estimate = round(Estimate, digits = 3),
         Evid.Ratio = round(Evid.Ratio, digits = 3)) %>%
  select("Hypothesis" = hypothesis, "Delta" = Estimate,
         `95\\% CIs`, "Evidence Ratio" = Evid.Ratio) %>%
  kbl(., longtable = T, booktabs = T,
      format = "latex", caption = caption,
      escape = FALSE) %>%
  column_spec(column = c(1:10), width = "2.5cm")  %>%
  kable_styling(latex_options = "striped")

pairwiseTarsusGrowthTableStrict
```

***Table 20:*** *Pairwise comparisons of Gompertz function variables between cold-reared (10°C until 8 weeks of age), mild-reared (constant 20°C), and warm-reared (30°C until 8 weeks of age) Japanese quail. Gompertz function variables being compared are indicated in parenthesis, per hypothesis.*

| Hypothesis | Delta | 95% CIs | Evidence Ratio |
|---|---|---|---|
| Warm-reared Asymptote (a) > Cold-reared Asymptote (a) | 0.798 | [-0.226, 1.813] | 8.919 |
| Warm-reared Displacement (b) > Cold-reared Displacement (b) | 0.074 | [0.003, 0.148] | 22.358 |
| Warm-reared Growth Rate (c) > Cold-reared Growth Rate (c) | 0.105 | [0.04, 0.17] | 443.444 |

```r
# Setting up permutations

permutationData <- data %>%
  rename("batch" = exp) %>%
  filter(week == 8 & batch == "C") %>%
  mutate(pretreatment = factor(pretreatment,
    levels = c("cold", "warm")
  )) %>%
  select(pretreatment, "tarsus" = tarsusLengthMean)

trueDeltaT <- mean(subset(permutationData, pretreatment == "warm")$tarsus) -
  mean(subset(permutationData, pretreatment == "cold")$tarsus)

# Permuting differences between means

drawDeltaT <- sapply(
  c(1:10000),
  function(x) {
    data <- permutationData %>%
      mutate("pretreatment" = sample(pretreatment,
        size = nrow(permutationData),
        replace = FALSE
      ))
    return(mean(subset(data, pretreatment == "warm")$tarsus) -
      mean(subset(data, pretreatment == "cold")$tarsus))
  }
)

require(loo)

# Examining whether observed difference in means is evidently larger than
# that calculated after iterations. Note that

pTest <- hypothesis_df(paste0(trueDeltaT, " - delta > 0"),
  alpha = 0.05,
  x = data.frame(
    "b_delta" = drawDeltaT,
    "prior_b_delta" = rnorm(10000, mean = 0.8, sd = 1)
  )
)

cat(paste0("Evidence Ratio = ", round(pTest$hypothesis$Evid.Ratio, digits = 3)))
```

```
## Evidence Ratio = 37.023
```

```r
cat(paste0(
  "Posterior Probability = ",
  round(pTest$hypothesis$Post.Prob, digits = 3)
))
```

```
## Posterior Probability = 0.974
```

Overall, our findings strongly indicate that the thermal environment during post-hatch development shapes both the magnitude and rate of body mass gain and extremity elongation. Under the hypothesis that Allen's rule emerges through phenotypic plasticity, one should expect that rates of body mass gain and extremity length elongation should also differ across developmental thermal environments, with extremities elongating more quickly than the rates of mass gain in the warmth, and elongating at more similar rates to those of mass gain in the cold. To qualitatively test this prediction, we next compared body mass growth rates and tarsus elongation rates among rearing conditions, as estimated from our previous non-linear models.

```r
# Checking differences in growth rate and inflection
# points for tarsus length and body mass, per rearing condition

caption <- paste0("Growth rates and extremity elongation ",
                  "rates at inflection points of growth and ",
                  "elongation curves repectively for Japanese quail."
```

```
)

growthInflectionTable <- growthModel %>%
  as.data.frame() %>%
  mutate(
    "coldGrowthRate" = b_C_Intercept + b_C_pretreatmentcold,
    "neutralGrowthRate" = b_C_Intercept,
    "warmGrowthRate" = b_C_Intercept + b_C_pretreatmentwarm
  ) %>%
  select(coldGrowthRate, neutralGrowthRate, warmGrowthRate) %>%
  pivot_longer(everything()) %>%
  arrange(name) %>%
  mutate("pretreatment" = rep(c("cold", "neutral", "warm"),
    each = nrow(as.data.frame(growthModel))
  )) %>%
  group_by(pretreatment) %>%
  summarise(
    "Body Mass Growth Rate" =
      paste0(round(median(value), digits = 4),
             " [",
             round(quantile(value, 0.025, type = 8), digits = 4),
             ", ",
             round(quantile(value, 0.975, type = 8), digits = 4),
      "]")
  ) %>%
  cbind(
    .,
    growthModelTarsus %>%
      as.data.frame() %>%
      mutate(
        "coldGrowthRate" = b_C_Intercept + b_C_pretreatmentcold,
        "neutralGrowthRate" = b_C_Intercept,
        "warmGrowthRate" = b_C_Intercept + b_C_pretreatmentwarm
      ) %>%
      select(coldGrowthRate, neutralGrowthRate, warmGrowthRate) %>%
      pivot_longer(everything()) %>%
      arrange(name) %>%
      mutate("pretreatment" = rep(c("cold", "neutral", "warm"),
        each = nrow(as.data.frame(growthModelTarsus))
      )) %>%
      group_by(pretreatment) %>%
      summarise(
        "Tarsus Elongation Rate" = paste0(round(median(value), digits = 4),
             " [",
             round(quantile(value, 0.025, type = 8), digits = 4),
             ", ",
             round(quantile(value, 0.975, type = 8), digits = 4),
      "]")
      ) %>%
      select(-pretreatment)
  ) %>%
  mutate(
    pretreatment = str_to_title(pretreatment),
    "Delta Growth Rate" =
      as.numeric(gsub("[[:space:]].*", "", `Tarsus Elongation Rate`)) -
      as.numeric(gsub("[[:space:]].*", "", `Body Mass Growth Rate`))
  ) %>%
  mutate(pretreatment = ifelse(pretreatment == "Cold", "Cold (10°C)",
    ifelse(pretreatment == "Neutral", "Mild (20°C)",
      "Warm (30°C)"
    )
  )) %>%
  rename("Rearing Conditions" = pretreatment) %>%
  kbl(.,
    longtable = T, booktabs = T,
    caption = caption, escape = FALSE
  ) %>%
```

```
  column_spec(column = c(1:10), width = "2cm") %>%
  kable_styling(latex_options = "striped")

growthInflectionTable
```

***Table 21:*** *Growth rates and extremity elongation rates at inflection points of growth and elongation curves repectively for Japanese quail.*

| Rearing Conditions | Body Mass Growth Rate | Tarsus Elongation Rate | Delta Growth Rate |
|---|---|---|---|
| Cold (10°C) | 0.5098 [0.4809, 0.5369] | 0.711 [0.4756, 0.8413] | 0.2012 |
| Mild (20°C) | 0.5504 [0.524, 0.5814] | 0.7703 [0.5722, 0.9196] | 0.2199 |
| Warm (30°C) | 0.5702 [0.5425, 0.5984] | 0.7983 [0.561, 0.9332] | 0.2281 |

```
#save_kable(growthInflectionTable, "../tables/growthInflectionTable.html")

# Visually comparing mass and tarsus growth curves

expand.grid(
  "pretreatment" = na.omit(unique(data$pretreatment)),
  "week" = seq(0, 8, by = 0.1)
) %>%
  mutate("weekB" = week + 1) %>%
  mutate("massFit" = predict(growthModel, newdata = .,
                             re_form = NA,
                             robust = TRUE)[, "Estimate"]) %>%
  group_by(week) %>%
  mutate("cScaledMass" = (massFit - mean(massFit)) / sd(massFit)) %>%
  ungroup() %>%
  mutate("tarsusFit" = predict(growthModelTarsus,
                               newdata = .,
                               robust = TRUE,
                               re_form = NA)[, "Estimate"]) %>%
  select(-c(cScaledMass, weekB)) %>%
  group_by(pretreatment) %>%
  mutate(
    "Body Mass" = ((massFit - min(massFit)) /
                   (max(massFit) - min(massFit))) * 100,
    "Tarsus Length" = ((tarsusFit - min(tarsusFit)) /
                       (max(tarsusFit) - min(tarsusFit))) * 100
  ) %>%
  ungroup() %>%
  select(-c(massFit, tarsusFit)) %>%
  pivot_longer(!c("pretreatment", "week"),
               names_to = "Parameter", values_to = "size") %>%
  mutate(pretreatment = ifelse(pretreatment == "cold", "10°C",
    ifelse(pretreatment == "neutral", "20°C", "30°C")
  )) %>%
  ggplot(aes(x = week, y = size, linetype = Parameter)) +
  geom_line() +
  facet_wrap(~pretreatment, scales = "free") +
  xlab("Age (Weeks)") +
  ylab("% of Growth Completed") +
  theme_classic()
```

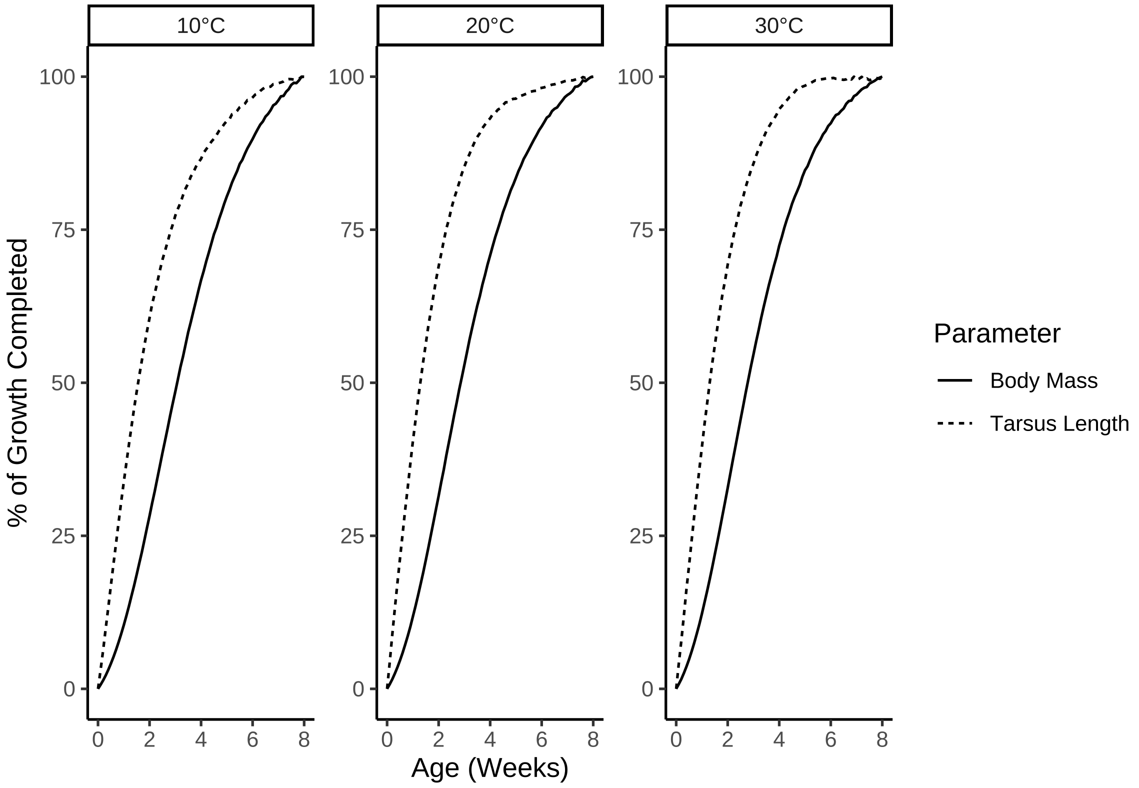


***Figure 38:*** *Comparison of mass and tarsus growth curves for Japanese quail reared in one of three different thermal conditions: cold (10°C until at least 3 weeks of age), mild (20°C until 8 weeks of age), or warm (30°C until at least 3 weeks of age). Growth curves are estimated from two, Bayesian, non-linear mixed effects models.*

```
# Tarsus clearly elongating more rapidly in the wamrth relative
# to mass when compared with other treatments.
```

```
expand.grid(
  "pretreatment" = na.omit(unique(data$pretreatment)),
  "week" = seq(0, 8, by = 0.1)
) %>%
  mutate("weekB" = week + 1) %>%
  mutate("massFit" = predict(growthModel, newdata = .,
                             re_form = NA,
                             robust = TRUE)[, "Estimate"]) %>%
  group_by(week) %>%
  mutate("cScaledMass" = (massFit - mean(massFit)) / sd(massFit)) %>%
  ungroup() %>%
  mutate("tarsusFit" = predict(growthModelTarsus,
                               newdata = .,
                               re_form = NA,
                               robust = TRUE)[, "Estimate"]) %>%
  select(-c(cScaledMass, weekB)) %>%
  group_by(pretreatment) %>%
  mutate(
    massFit = (massFit / max(massFit)) * 100,
    tarsusFit = (tarsusFit / max(tarsusFit)) * 100
  ) %>%
  ungroup() %>%
  mutate(
    "delta" = tarsusFit - massFit,
    pretreatment = ifelse(pretreatment == "cold", "10°C",
      ifelse(pretreatment == "neutral", "20°C", "30°C")
    )
```

```
  ) %>%
  mutate(pretreatment = factor(pretreatment)) %>%
  ggplot(aes(x = week, y = delta, linetype = pretreatment)) +
  geom_line() +
  scale_linetype_manual(values = c("solid", "dashed", "dotted"),
                        name = "Rearing\nConditions") +
  xlab("Age (Weeks)") +
  ylab(paste0("Difference in Relative Growth Completion\n",
              "Between Tarsus Length and Body Mass (%)")) +
  theme_classic() +
  theme(
    axis.title = element_text(family = "Noto Sans"),
    text = element_text(family = "Noto Sans")
  )
```

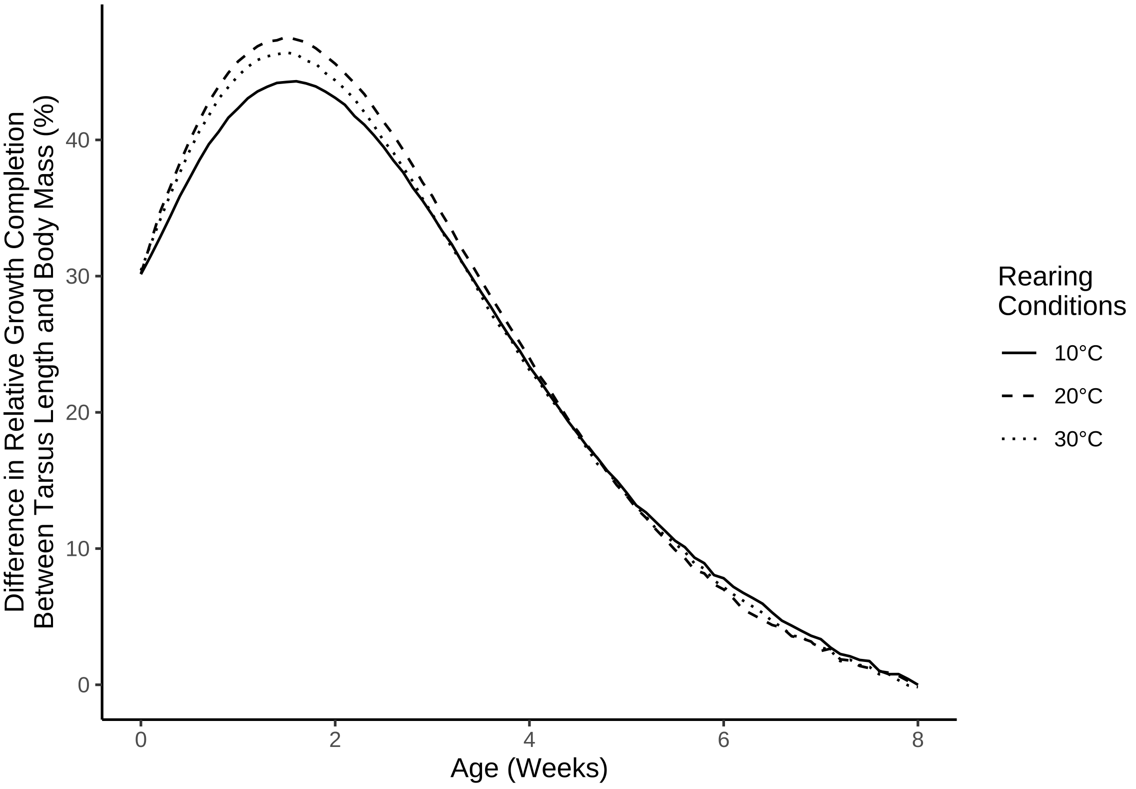


***Figure 39:*** *Differences between mass and tarsus growth curves (percent growth completed) across ages for Japanese quail reared in one of three different thermal conditions: cold (10°C until at least 3 weeks of age), mild (20°C until 8 weeks of age), or warm (30°C until at least 3 weeks of age). Growth curves are estimated from two, Bayesian, non-linear mixed effects models.*

```
# Very telling plot. Cold-reared individuals lag in growth
# considerably at around 2 weeks of age. Checking relative completion of
# tarsus growth when mass growth is 80% completed.

caption <- paste0('Total tarsus elongation of Japanese ',
                  "quail at ages where mass gain is 50\\% complete. ",
                  "Elongation rates are shown for quail reared in the ",
                  "cold (10°C until at least 3 weeks of age), mild ",
                  "conditions (20°C until 8 weeks of age), or the ",
                  "warmth (30°C until at least 3 weeks of age)."
)

elongationAt80 <- expand.grid(
```

```
  "pretreatment" = na.omit(unique(data$pretreatment)),
  "week" = seq(0, 8, by = 0.1)
) %>%
  mutate("weekB" = week + 1) %>%
  mutate("massFit" = predict(growthModel,
                             newdata = .,
                             re_form = NA,
                             robust = TRUE)[, "Estimate"]) %>%
  group_by(week) %>%
  mutate("cScaledMass" = (massFit - mean(massFit)) / sd(massFit)) %>%
  ungroup() %>%
  mutate("tarsusFit" = predict(growthModelTarsus,
                               newdata = .,
                               re_form = NA,
                               robust = TRUE)[, "Estimate"]) %>%
  select(-c(cScaledMass, weekB)) %>%
  group_by(pretreatment) %>%
  mutate(
    massFit = ((massFit - min(massFit)) /
                 (max(massFit) - min(massFit)) * 100),
    tarsusFit = ((tarsusFit - min(tarsusFit)) /
                   (max(tarsusFit) - min(tarsusFit)) * 100)
  ) %>%
  mutate("grab" = abs(massFit - 80)) %>%
  filter(grab == min(grab)) %>%
  select(-c(grab, massFit)) %>%
  mutate(pretreatment = ifelse(pretreatment == "cold", "Cold (10°C)",
    ifelse(pretreatment == "neutral", "Mild (20°C)",
      "Warm (30°C)"
    )
  )) %>%
  rename("Age (Weeks)" = week,
         "Tarsus Elongation Completed (%)" = tarsusFit,
         "Rearing Conditions" = pretreatment) %>%
  kbl(.,
    longtable = T, booktabs = T, format = "latex",
    caption = caption
  ) %>%
  column_spec(column = c(1:10), width = "2.5cm")  %>%
  kable_styling(latex_options = "striped")

elongationAt80
```

***Table 22:*** *Total tarsus elongation of Japanese quail at ages where mass gain is 50% complete. Elongation rates are shown for quail reared in the cold (10°C until at least 3 weeks of age), mild conditions (20°C until 8 weeks of age), or the warmth (30°C until at least 3 weeks of age).*

| Rearing Conditions | Age (Weeks) | Tarsus Elongation Completed (%) |
|---|---|---|
| Warm (30°C) | 4.6 | 97.40176 |
| Mild (20°C) | 4.7 | 95.74330 |
| Cold (10°C) | 5.0 | 92.87750 |

```
#save_kable(elongationAt80, "../tables/tarsusElongationAt80PercentGrowth.html")

# When mass is growth is 80% complete, tarsus growth is already
# 98.9% completed in warm-reared birds relative to 94.7% completed
# in cold-reared and 97.4% completed in mild-reared birds.
```

```
# And finally, evaluating ages of most rapid mass gain and tarsus elongation per treatment.

caption <- paste0('Ages of most rapid mass gain and tarsus ',
                  "elogation among Japanese quail, between 0 and 8 weeks ",
                  "of age. Quail were reared in the cold ",
                  "(10°C until at least ",
```

```
                  "3 weeks of age), mild conditions ",
                  "(20°C until 8 weeks of age), ",
                  "or the warmth (30°C ",
                  "until at least 3 weeks of age)."
)

ageOfRapidGrowth <- with(
  as.data.frame(growthModel),
  data.frame(
    "pretreatment" = c("neutral", "cold", "warm"),
    "weeks" = c(
      mean(log(b_B_Intercept) / b_C_Intercept),
      mean(log(b_B_Intercept + b_B_pretreatmentcold) /
             (b_C_Intercept + b_C_pretreatmentcold)),
      mean(log(b_B_Intercept + b_B_pretreatmentwarm) /
             (b_C_Intercept + b_C_pretreatmentwarm))
    )
  )
) %>%
  mutate("days" = weeks * 7) %>%
  select(-weeks) %>%
  rename("Age of Most Rapid Mass Gain (Days)" = days) %>%
  merge(.,
    with(
      as.data.frame(growthModelTarsus),
      data.frame(
        "pretreatment" = c("neutral", "cold", "warm"),
        "weeks" = c(
          mean(log(b_B_Intercept) / b_C_Intercept),
          mean(log(b_B_Intercept + b_B_pretreatmentcold) /
                 (b_C_Intercept + b_C_pretreatmentcold)),
          mean(log(b_B_Intercept + b_B_pretreatmentwarm) /
                 (b_C_Intercept + b_C_pretreatmentwarm))
        )
      )
    ) %>%
      mutate("days" = weeks * 7) %>%
      select(-weeks) %>%
      rename("Age of Most Rapid Tarsus Elongation (Days)" = days),
    by = "pretreatment"
  ) %>%
  mutate(pretreatment = ifelse(pretreatment == "cold", "Cold (10°C)",
    ifelse(pretreatment == "neutral", "Mild (20°C)",
      "Warm (30°C)"
    )
  )) %>%
  rename("Rearing Conditions" = pretreatment) %>%
  kbl(.,
    longtable = T, booktabs = T, format = "latex",
    caption = caption
  ) %>%
  column_spec(column = c(1:10), width = "2.5cm")  %>%
  kable_styling(latex_options = "striped")

ageOfRapidGrowth
```

***Table 23:*** *Ages of most rapid mass gain and tarsus elogation among Japanese quail, between 0 and 8 weeks of age. Quail were reared in the cold (10°C until at least 3 weeks of age), mild conditions (20°C until 8 weeks of age), or the warmth (30°C until at least 3 weeks of age).*

| Rearing Conditions | Age of Most Rapid Mass Gain (Days) | Age of Most Rapid Tarsus Elongation (Days) |
|---|---|---|
| Cold (10°C) | 16.97926 | 1.886318 |
| Mild (20°C) | 15.46763 | 1.755900 |

| | | |
|---|---|---|
| Warm (30°C) | 14.91451 | 2.185562 |

```
# Cold-reared birds reach peak mass gain rates at ~2 days
# after warm-reared birds, however, tarsus elongation rates
# reach peaks at ~1.5 days for all treatment groups.
```
