## Supplementary material for "No evidence that shrinking and shapeshifting meaningfully affect how birds respond to warming and cooling": Supp. 3

### Metabolic Slope and Repeatability Analyses

#### Overview

In a previous document (entitled: "Estimating Lower Critical Temperature in a Population of Japanese Quail"), we showed that patterns of growth and limb elongation in Japanese quail (n = 108) depend on the thermal environment experienced during post-hatch development. Quail reared in cold environmental (10°C) grew more slowly than those reared in warm environments (30°C) but reached higher asymptotic masses and displayed relatively shorter tarsi by adulthood. In this document we test whether morphology during development (here, 3 weeks of age) or at maturity (8 weeks of age) influenced thermoregulatory costs in these quail, while also accounting for direct effects of rearing temperature (and thus, physiological acclimation) on such costs.

To approximate thermoregulatory costs, we measured resting energy expenditure at thermoneutrality (30°C), throughout a cold challenge (10°C - 30°C) and throughout a heat challenge (30°C - 40°C). Details of these measurements are described in the Materials and Methods of Tabh et al (2024). We then quantified the rate at which metabolism increased across cold and heat challenges (using simple Bayesian linear mixed effects models) assuming that higher rates (or steeper slopes) indicate lower thermoregulatory efficiency and thus, higher thermoregulatory costs relative to lower rates (or shallower slopes). Next, we tested whether morphology and rearing conditions influenced these rates. Throughout this document, we described the steps taken: (1) to quantify metabolic responses to cold and heat challenges, (2) estimate how consistently variable these responses are among individuals, and (3) analyse their potential dependence on morphology and thermal history (i.e. rearing conditions).

##### Data compilation

We again begin by importing packages and functions required to execute the below analyses. Versions of R packages used are displayed for code reproduction. We recommend that packages versions used here and installed for the those wishing to reproduce our results.

```
def.chunk.hook <- knitr::knit_hooks$get("chunk")
knitr::knit_hooks$set(chunk = function(x, options) {
  x <- def.chunk.hook(x, options)
  ifelse(options$size != "normalsize",
         paste0("\n \\", options$size, "\n\n", x, "\n\n \\normalsize"), x)
})

knitr::opts_chunk$set(fig.pos = "H", out.extra = "")
knitr::opts_chunk$set(size = "footnotesize")

library("tidyverse")
library("easypackages")

packageList <- c("bayesplot", "brms", "doParallel",
                 "foreach", "ggpubr", "kableExtra",
```

```
                 "latex2exp", "patchwork", "priorsense",
                 "showtext", "tidybayes", "wesanderson")

libraries(packageList)
library("brmsMethods")

# Printing package version numbers

caption <- paste0("R packages and their respective versions used for",
                  " data organisation and analysis in this study."
)

sapply(packageList, function(x) {
  y <- as.character(packageVersion(x))
  return(y)
}, simplify = FALSE) %>%
  enframe(., name = "Package", value = "Version") %>%
  as.data.frame(.) %>%
  kbl(.,
    longtable = T, booktabs = T,
    caption = caption
  ) %>%
  kable_styling(latex_options = "striped")
```

***Table 1:*** *R packages and their respective versions used for data organisation and analysis in this study.*

| Package | Version |
|---|---|
| bayesplot | 1.11.1 |
| brms | 2.20.4 |
| doParallel | 1.0.17 |
| foreach | 1.5.2 |
| ggpubr | 0.6.0 |
| kableExtra | 1.3.4 |
| latex2exp | 0.9.6 |
| patchwork | 1.2.0 |
| priorsense | 0.0.0.9000 |
| showtext | 0.9.6 |
| tidybayes | 3.0.4 |
| wesanderson | 0.3.6.9000 |

```
# Adding custom functions

## A function to calculate the mode of a vector

md <- function(x) {
  all_values <- unique(x)
  all_values[which.max(tabulate(match(x, all_values)))]
}

# A function to cleanly view autocorrelation between posterior
# draws of specified coefficients/variables
```

```
clean_ac <- function(x, prs = NA, names = NA) {
  require("rstan")

  if (class(x)[1] != "brmsfit") {
    return("x must be a brmsfit object.")
  }

  if (is.na(prs[1])) {
    return(stan_ac(x$fit))
  }

  if (!is.na(prs[1]) & is.na(names[1])) {
    return(stan_ac(x$fit, pars = prs))
  }

  if (length(prs) != length(names)) {
    return("Length of pars and names must be equal.")
  }

  Base_plot <- stan_ac(x$fit, pars = prs, fill = nice_pink)
  Base_plot$data$parameters <- as.character(Base_plot$data$parameters)

  for (i in 1:length(prs)) {
    Base_plot$data$parameters[c(which(Base_plot$data$parameters == prs[i]))] <-
      names[i]
  }
  Base_plot$data$parameters <- as.factor(Base_plot$data$parameters)

  return(Base_plot)
}

# A function to simplify the output of bayestestR's hdi function.

simple_hdi <- function(x, rnd = 3, cis = c(50, 95), sci_note = FALSE) {
  if (class(x)[1] != "brmsfit") {
    return("x must be a brmsfit object.")
  }
  if (length(cis) != 2) {
    return("cis must be a vector of integers with length 2")
  }

  HDI_low <- bayestestR::hdi(x, effects = "all", ci = min(cis) / 100)
  HDI_high <- bayestestR::hdi(x, effects = "all", ci = max(cis) / 100)

  if (sci_note == FALSE) {
    Results <- data.frame(
      "Parameter" = HDI_low$Parameter,
      "1" = round(HDI_low$CI_low, rnd),
      "2" = round(HDI_low$CI_high, rnd),
      "3" = round(HDI_high$CI_low, rnd),
      "4" = round(HDI_high$CI_high, rnd)
    )
    colnames(Results)[c(2:5)] <-
```

```
      c(paste0("Low_HDI_", min(cis)), paste0("High_HDI_", min(cis)),
        paste0("Low_HDI_", max(cis)), paste0("High_HDI_", max(cis)))
  } else if (sci_note == TRUE) {
    Results <- data.frame(
      "Parameter" = HDI_low$Parameter,
      "1" = format(round(HDI_low$CI_low, rnd), scientific = TRUE),
      "2" = format(round(HDI_low$CI_high, rnd), scientific = TRUE),
      "3" = format(round(HDI_high$CI_low, rnd), scientific = TRUE),
      "4" = format(round(HDI_high$CI_high, rnd), scientific = TRUE)
    )
    colnames(Results)[c(2:5)] <- c(paste0("Low_HDI_", min(cis)),
                                   paste0("High_HDI_", min(cis)),
                                   paste0("Low_HDI_", max(cis)),
                                   paste0("High_HDI_", max(cis)))
  }

  return(Results)
}

modeHDI <- function(x, cis = c(50, 95), rnd = 4, collapse = FALSE){
  stopifnot("x must be a 'brmsfit' object" = class(x) == "brmsfit",
            "collapse must be logical TRUE/FALSE" = is.logical(collapse))

  out <- lapply(X = as.data.frame(x), MARGIN = 2, FUN = ggdist::mode_hdi,
                .width = c(cis/100))
  hold <- names(out)
  out <- out %>%
    map2(hold, ~mutate(.x, name = .y)) %>%
    bind_rows() %>%
    mutate(y = round(y, digits = rnd),
           ymin = round(ymin, digits = rnd),
           ymax = round(ymax, digits = rnd)) %>%
    select("par" = name, "mode" = y, "lcl" = ymin,
           "ucl"= ymax, "confidenceLevel" = .width)

  if (collapse == FALSE){
    return(out)
  } else if (collapse == TRUE){
    out <- out %>%
      mutate("cis" = paste0("[", lcl, ", ", ucl, "]")) %>%
      select(-c(lcl, ucl))

    return(out)
  }
}


# A function to calculate quantile intervals from a brmsfit object

quantileCIs <- function(x, rnd = 3, cis = c(50, 95), sci_note = FALSE) {
  require(tidyverse)

  if (class(x)[1] != "brmsfit") {
```

```
    return("x must be a brmsfit object.")
  }
  if (length(cis) != 2) {
    return("cis must be a vector of integers with length 2")
  }

  prbs = c()
  nColNames = c()
  for (i in 1:length(cis)){
    prbs = c(prbs, c(0.5 - (cis[i]/100)/2, 0.5 + (cis[i]/100)/2))
    nColNames = c(nColNames,
                  paste0("Low_CI_", cis[i]),
                  paste0("High_CI_", cis[i])
    )
  }

  modelFrame = as.data.frame(x)

  Results <- apply(modelFrame, MARGIN = 2, FUN = quantile,
        probs = prbs, type = 8) %>%
    t() %>%
    as.data.frame() %>%
    rownames_to_column(var = "par") %>%
    `colnames<-`(c("Parameter", nColNames))

  if (sci_note == FALSE) {
    Results <- apply(modelFrame, MARGIN = 2, FUN = quantile,
        probs = prbs, type = 8) %>%
    t() %>%
    as.data.frame() %>%
    rownames_to_column(var = "par") %>%
    `colnames<-`(c("Parameter", nColNames))

  } else if (sci_note == TRUE) {
    Results <- apply(modelFrame, MARGIN = 2, FUN = quantile,
        probs = prbs, type = 8) %>%
    t() %>%
    as.data.frame() %>%
    rownames_to_column(var = "par") %>%
    `colnames<-`(c("Parameter", nColNames)) %>%
    mutate_at(.vars = vars(-Parameter),
              .funs = function(x){
                return(format(x, scientific = TRUE))
              }
    )
  }

  return(Results)
}

# A function that allows users to assign multiple objects
# to different variable names at once. The below is reported by "ellbur" at
# https://strugglingthroughproblems.wordpress.com/author/ellbur/page/3/.
```

```
{
  "%=%" <- function(l, r, ...) UseMethod("%=%")

  "%=%.lbunch" <- function(l, r, ..., List = NA) {
    Envir <- as.environment(-1)

    if (!is.na(List)) {
      l <- List[[1]]
      r <- List[[2]]
    }

    if (length(r) > length(l)) {
      warning("RHS has more args than LHS. Only first",
              length(l), "used.")
    }

    if (length(l) > length(r)) {
      warning("LHS has more args than RHS. RHS will be repeated.")
      r <- extendToMatch(r, l)
    }

    for (II in 1:length(l)) {
      do.call("<-", list(l[[II]], r[[II]]), envir = Envir)
    }
  }

  extendToMatch <- function(source, destin) {
    s <- length(source)
    d <- length(destin)

    if (d == 1 && s > 1 && !is.null(as.numeric(destin))) {
      d <- destin
    }

    dif <- d - s
    if (dif > 0) {
      source <- rep(source, ceiling(d / s))[1:d]
    }
    return(source)
  }

  g <- function(...) {
    List <- as.list(substitute(list(...)))[-1L]
    class(List) <- "lbunch"
    return(List)
  }
}

# A function to simplify output of bayes_R2 from brms

simpleR2 <- function(x, ndraws = 1000, roundDigits = 5,
                     robust = TRUE) {
  stopifnot("x must be a 'brmsfit' object" = class(x) == "brmsfit",
```

```
            "robust must be logical (TRUE/FALSE)" = is.logical(robust))
  grab <- brms::bayes_R2(x, ndraws = ndraws, robust = robust)
  toPrint <- paste0(
    "R2 = ", round(grab[, "Estimate"], digits = roundDigits),
    " [",
    round(grab[, "Q2.5"], digits = roundDigits),
    ",",
    round(grab[, "Q97.5"], digits = roundDigits),
    "]"
  )
  cat(toPrint)
}

# A function to calculate the position of a skew-normal
# distribution given its mean, omega, and alpha values

skewxi <- function(mean, omega, alpha) {
  delta <- alpha / (sqrt(1 + alpha^2))
  xi <- mean - omega * delta * sqrt(2 / pi)
  return(xi)
}

# A function to produce clean posterior or prior
# predictive checks, based upon "pp_check" from the R-package 'brms'.

pp_check2 <- function(model, resp = NA, ndraws = 500,
                      xlab = "label", colour = "lightblue") {
  require(brms)
  require(ggplot2)
  stopifnot("Model must be a brmsfit object" = is.brmsfit(model))

  if (is.na(resp)) {
    resp <- model$formula$resp
  }

  p1 <- brms::pp_check(model, ndraws = ndraws, resp = resp) +
    scale_colour_manual(
      values = c("black", colour),
      labels = c("y", "yhat"),
      name = NULL
    ) +
    xlab(xlab) +
    ylab("Density") +
    theme_classic()
  return(p1)
}

# A function to summarise and print Gelman-Rubin statistics and effective
# sample sizes to sample sizes for brmsfit objects.

neffBase <- function(x){
  stopifnot("Model must be a brmsfit object" = is.brmsfit(x))
  out <- as.data.frame(brms::neff_ratio(x)) %>%
```

```
    rownames_to_column(var = "var") %>%
    filter(!(var %in% c("lprior", "lp__"))) %>%
    pull(.)
  return(out)
}

chainCheck <- function(model, rDig = 3) {
  require(brms)
  stopifnot("Model must be a brmsfit object" = is.brmsfit(model))

  Rhat <- paste0(
    "Rhat range: ",
    round(min(rhat(model)), digits = rDig),
    " - ",
    round(max(rhat(model)), digits = rDig)
  )
  Neff <- paste0(
    "Neff/N range: ",
    round(min(neffBase(model)), digits = rDig),
    " - ",
    round(max(neffBase(model)), digits = rDig)
  )
  cat(paste0(Rhat, "\n", Neff))
}

# Setting working directory.

setwd(
  paste0(
  "/Users/joshuatabh/Documents/researchProjects/lund/",
  "functionOfAllen/functionData"
  )
)
```

#### Import, collation, and visualisation of resting energy expenditure data

Data pertaining to both morphometry and resting energy expenditure of our quail are first loaded, organised, and quality-checked. Visualisations of data and identification of erroneous data points is conducted below as above. Following quality checks, energy expenditure data is bound to morphometric data that was previously compiled above.

Compilation of morphometric data is described fully in the supplemental document entitled “Data Organisation and Analysis of Growth”.

```
# Loading in morphological data

data <- read.csv("compiledData.csv")

# Loading in raw metabolic rate data

vo2Data <- bind_rows(
  read.csv("exp1VO2.csv"),
  read.csv("exp2VO2.csv"),
  read.csv("exp3VO2.csv")
) %>%
  select(-c(startTime, endTime, initialTb, fileName)) %>%
```

```
  mutate(pretreatment = ifelse(pretreatment == "control",
                              "neutral", pretreatment))

# Joining metabolic data with morphology data

all <- merge(
  data %>%
    select(-Ta),
  vo2Data %>%
    distinct() %>%
    select(-c(pretreatment, posttreatment, birdID)),
  by = c("ring", "week", "exp"),
  all.x = TRUE
) %>%
  distinct()

# Re-ordering columns more logically

all <- all %>%
  select(ring, birdID, sex, exp, week, pretreatment,
         posttreatment, treatment, Ta, mass, wingLength,
         meanTb, tarsusLengthMean, tarsusLengthSD,
         tarsusHeightMean, tarsusHeightSD,
         tarsusCalibration, VO2, RMR)

# Ambient temperature measurements from experiment C are binned and
# only metabolic data obtained at 30°C and 40°C are retained
# for consistency with experiments A and B.

all <- all %>%
  mutate(Ta = ifelse(exp == "C" & Ta > 28 & Ta < 31, 30, Ta)) %>%
  mutate(Ta = ifelse(exp == "C" & Ta > 38 & Ta < 41, 40, Ta)) %>%
  filter(Ta %in% c(10, 20, 30, 40))

# Checking for spurious VO2 measurements with Cleveland dotplot

ggplot(
  all %>%
    filter(week %in% c(3, 8)) %>%
    group_by(week) %>%
    mutate("ID" = 1:n()) %>%
    ungroup() %>%
    mutate(
      "week" = paste0("Age = ", week, " weeks"),
      "Ta" = paste0(Ta, "°C")
    ),
  aes(x = ID, y = VO2)
) +
  facet_grid(week ~ Ta, scales = "free") +
  geom_point(size = 2, colour = "black",
             fill = "slateblue", alpha = 0.7) +
  geom_rect(
    data = all %>%
      filter(week %in% c(3, 8)) %>%
      group_by(week, Ta) %>%
      summarise(
        "Mean" = mean(VO2, na.rm = T),
        "LCL" = Mean - 3 * sd(VO2, na.rm = T),
        "UCL" = Mean + 3 * sd(VO2, na.rm = T),
        "ID" = 1, "VO2" = 1
      ) %>%
      ungroup() %>%
      mutate(
        "week" = paste0("Age = ", week, " weeks"),
        "Ta" = paste0(Ta, "°C")
      ) %>%
      as.data.frame(),
```

```
    aes(xmin = -Inf, xmax = Inf, ymin = LCL, ymax = UCL),
    colour = "black", fill = "grey70", alpha = 0.5
  ) +
  theme_classic() +
  xlab("Sample Number") +
  ylab(TeX('$Resting~Metabolism~(mL~O_{2}/min)$'))
```

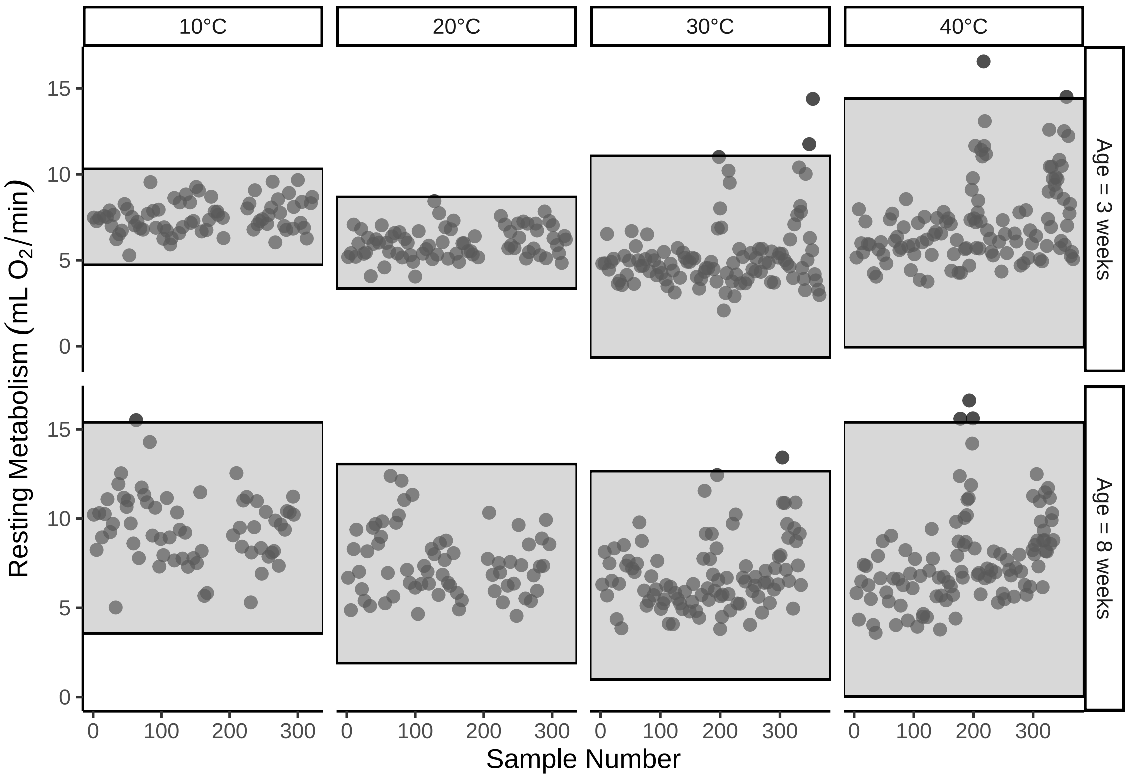


***Figure 1:*** *Cleveland dotplot of resting metabolism measures drawn from 3 and 8 week old Japanese quail. Measurements are divided by ambient temperature during collection. Grey boxes represent means ± three standard deviations. Dots outside of grey rectangles represent tentative outliers.*

A few data points observed at 30°C and 40°C appear high relative to mean values, but are not particularly abnormal from general trends. We retaining these values given that we have no reason to expect that they are erroneous.

Next, we check for duplications in our data and further visualise resting metabolism as a function of morphology (here, body mass, tarsus length, and relative [or residual] tarsus length) across rearing treatments and ambient temperature of measurement.

```
# Checking for, and removing, duplicates

all %>%
  filter(week %in% c(3, 8)) %>%
  group_by(ring, week, Ta, VO2) %>%
  count() %>%
  filter(n > 1)
```

```
## # A tibble: 4 x 5
##   ring   week    Ta   VO2     n
##   <chr> <int> <dbl> <dbl> <int>
## 1 B14       3    10  7.83     2
## 2 B14       3    20  5.53     2
```

```
## 3 B14        3    30  4.55     2
## 4 B14        3    40  4.28     2
```

```
# Duplicate measurements for one individual at each temperature treatment

all %>%
  mutate("Row" = 1:n()) %>%
  filter((ring == "B14" & week == "3")) %>%
  select(-c(
    birdID, sex, exp, posttreatment, treatment, meanTb,
    tarsusLengthSD, tarsusHeightMean, tarsusHeightSD,
    tarsusCalibration, Row
  )) %>%
  arrange(Ta)
```

```
## # A tibble: 8 x 9
##   ring   week pretreatment    Ta  mass wingLength tarsusLengthMean   VO2   RMR
##   <chr> <int> <chr>        <dbl> <dbl>      <dbl>            <dbl> <dbl> <dbl>
## 1 B14       3 warm            10  153.        103             39.0  7.83  2.61
## 2 B14       3 warm            10  153.        103             37.2  7.83  2.61
## 3 B14       3 warm            20  153.        103             39.0  5.53  1.84
## 4 B14       3 warm            20  153.        103             37.2  5.53  1.84
## 5 B14       3 warm            30  153.        103             39.0  4.55  1.52
## 6 B14       3 warm            30  153.        103             37.2  4.55  1.52
## 7 B14       3 warm            40  153.        103             39.0  4.28  1.43
## 8 B14       3 warm            40  153.        103             37.2  4.28  1.43
```

```
# Tarsus measurements uniquely doubled. Averaging values and proceeding.

all <- rbind(
  all %>%
    filter(ring == "B14" & week == "3") %>%
    group_by(Ta) %>%
    mutate(
      "tarsusLengthMean" = mean(tarsusLengthMean, na.rm = T),
      "tarsusHeightMean" = mean(tarsusHeightMean, na.rm = T)
    ) %>%
    mutate("n" = 1:2) %>%
    ungroup() %>%
    arrange(n, Ta) %>%
    slice(1:4) %>%
    select(-n),
  all %>%
    filter(!(ring == "B14" & week == "3"))
) %>%
  arrange(exp, week, ring, Ta)

# Plotting VO2 as kJ/day across tarsus length and mass

mass3Weeks <- ggplot(all %>%
  mutate(
    Ta = paste0(Ta, "°C"),
    pretreatment = ifelse(pretreatment == "cold", "Cold (10°C)",
      ifelse(pretreatment == "neutral", "Mild (20°C)",
        "Warm (30°C)"
      )
    )
  ) %>%
  filter(week == 3) %>%
  drop_na(pretreatment), aes(x = mass, y = VO2)) +
  facet_wrap(~Ta, scale = "free") +
  geom_point(pch = 21, colour = "black", size = 2, aes(fill = pretreatment)) +
  theme_classic() +
  scale_fill_manual(values = c("#7BB4E3", "black", "#CD5C5C"),
                    name = "Rearing\nConditions") +
  xlab("Body Mass at 3 Weeks (g)")  +
  ylab(TeX('$Resting~Metabolism~(mL~O_{2}/min)$'))

mass8Weeks <- ggplot(all %>%
```

```
  mutate(
    Ta = paste0(Ta, "°C"),
    pretreatment = ifelse(pretreatment == "cold", "Cold (10°C)",
      ifelse(pretreatment == "neutral", "Mild (20°C)",
        "Warm (30°C)"
      )
    )
  ) %>%
  filter(week == 8) %>%
  drop_na(pretreatment), aes(x = mass, y = VO2)) +
  facet_wrap(~Ta, scales = "free") +
  geom_point(pch = 21, colour = "black", size = 2, aes(fill = pretreatment)) +
  theme_classic() +
  scale_fill_manual(values = c("#7BB4E3", "black", "#CD5C5C"),
                    name = "Rearing\nConditions") +
  xlab("Body Mass at Maturity (g)")  +
  ylab(TeX('$Resting~Metabolism~(mL~O_{2}/min)$'))

showtext_auto()
ggarrange(mass3Weeks, mass8Weeks, ncol = 1,
  common.legend = T
)
```

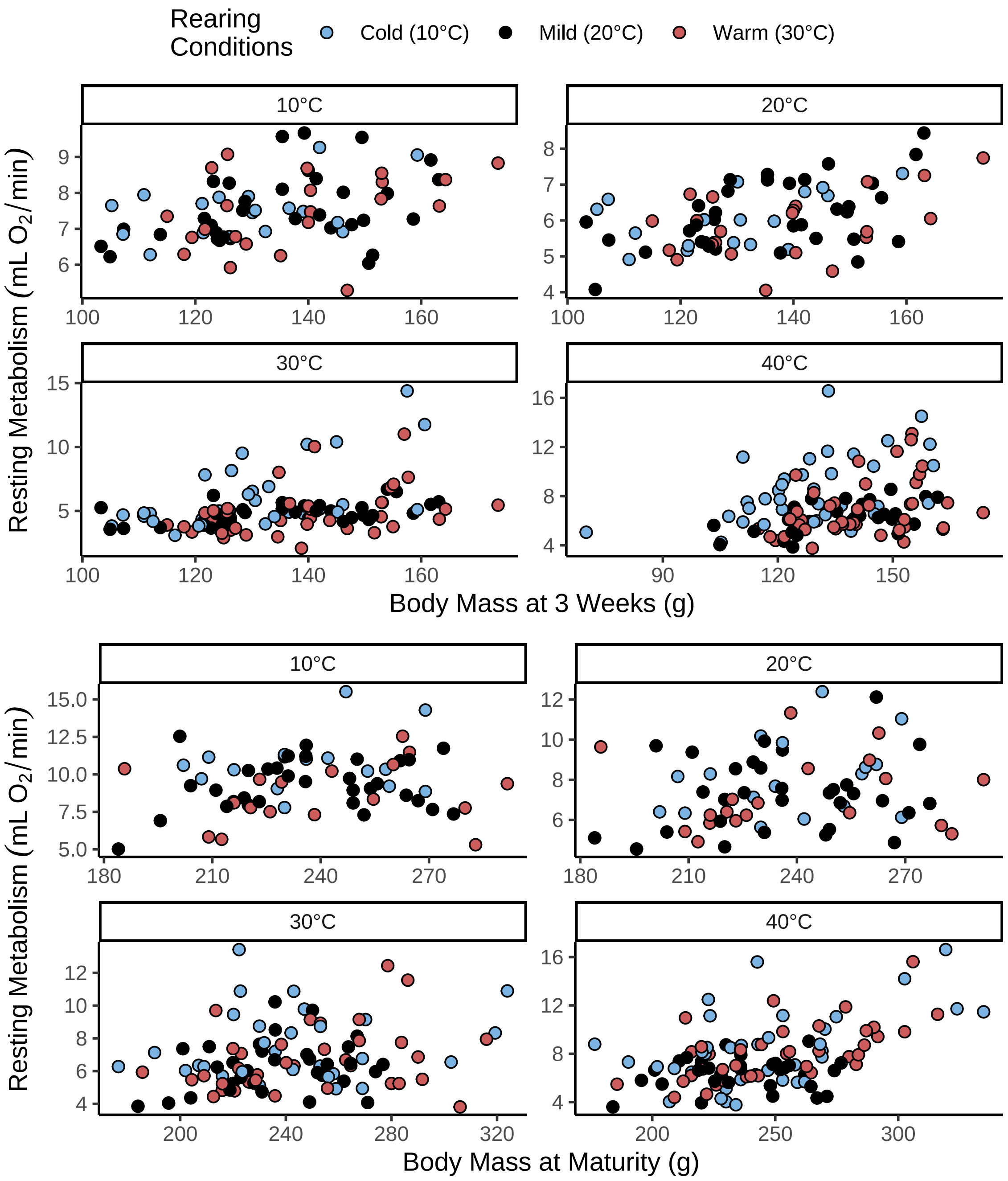


***Figure 2:*** *Resting metabolic rate (mL $O_2$/min) as a function of body mass (g) in 3 and 8 week old Japanese quail. Resting metabolic rate was measured at four different thermal environments (10°C, 20°C, 30°C and 40°C), with individuals measured in at least two environments. Measured quail were reared in one of three difference rearing conditions: (1) cold (10°C) until at least three weeks of age, (2) mild conditions (20°C) for eight weeks of age, or (3) warmth (30°C) until at least three weeks of age.*

```
showtext_auto(enable = FALSE)
```

```
tarsus3Weeks <- ggplot(all %>%
  mutate(
    Ta = paste0(Ta, "°C"),
    pretreatment = ifelse(pretreatment == "cold", "Cold (10°C)",
      ifelse(pretreatment == "neutral", "Mild (20°C)",
```

```
        "Warm (30°C)"
      )
    )
  ) %>%
  filter(week == 3) %>%
  drop_na(pretreatment), aes(x = tarsusLengthMean, y = VO2)) +
  facet_wrap(~Ta, scale = "free") +
  geom_point(pch = 21, colour = "black", size = 2, aes(fill = pretreatment)) +
  theme_classic() +
  scale_fill_manual(values = c("#7BB4E3", "black", "#CD5C5C"),
                    name = "Rearing\nConditions") +
  xlab("Tarsus Length at Week 3 (mm)")  +
  ylab(TeX('$Resting~Metabolism~(mL~O_{2}/min)$')) +
  theme(legend.position = "none")

tarsus8Weeks <- ggplot(all %>%
  mutate(
    Ta = paste0(Ta, "°C"),
    pretreatment = ifelse(pretreatment == "cold", "Cold (10°C)",
      ifelse(pretreatment == "neutral", "Mild (20°C)",
        "Warm (30°C)"
      )
    )
  ) %>%
  filter(week == 8) %>%
  drop_na(pretreatment), aes(x = tarsusLengthMean, y = VO2)) +
  facet_wrap(~Ta, scales = "free") +
  geom_point(pch = 21, colour = "black", size = 2, aes(fill = pretreatment)) +
  theme_classic() +
  scale_fill_manual(values = c("#7BB4E3", "black", "#CD5C5C"),
                    name = "Rearing\nConditions") +
  xlab("Tarsus Length at Maturity (mm)")  +
  ylab(TeX('$Resting~Metabolism~(mL~O_{2}/min)$')) +
  theme(legend.position = "bottom")

showtext_auto()
tarsus3Weeks/tarsus8Weeks
```

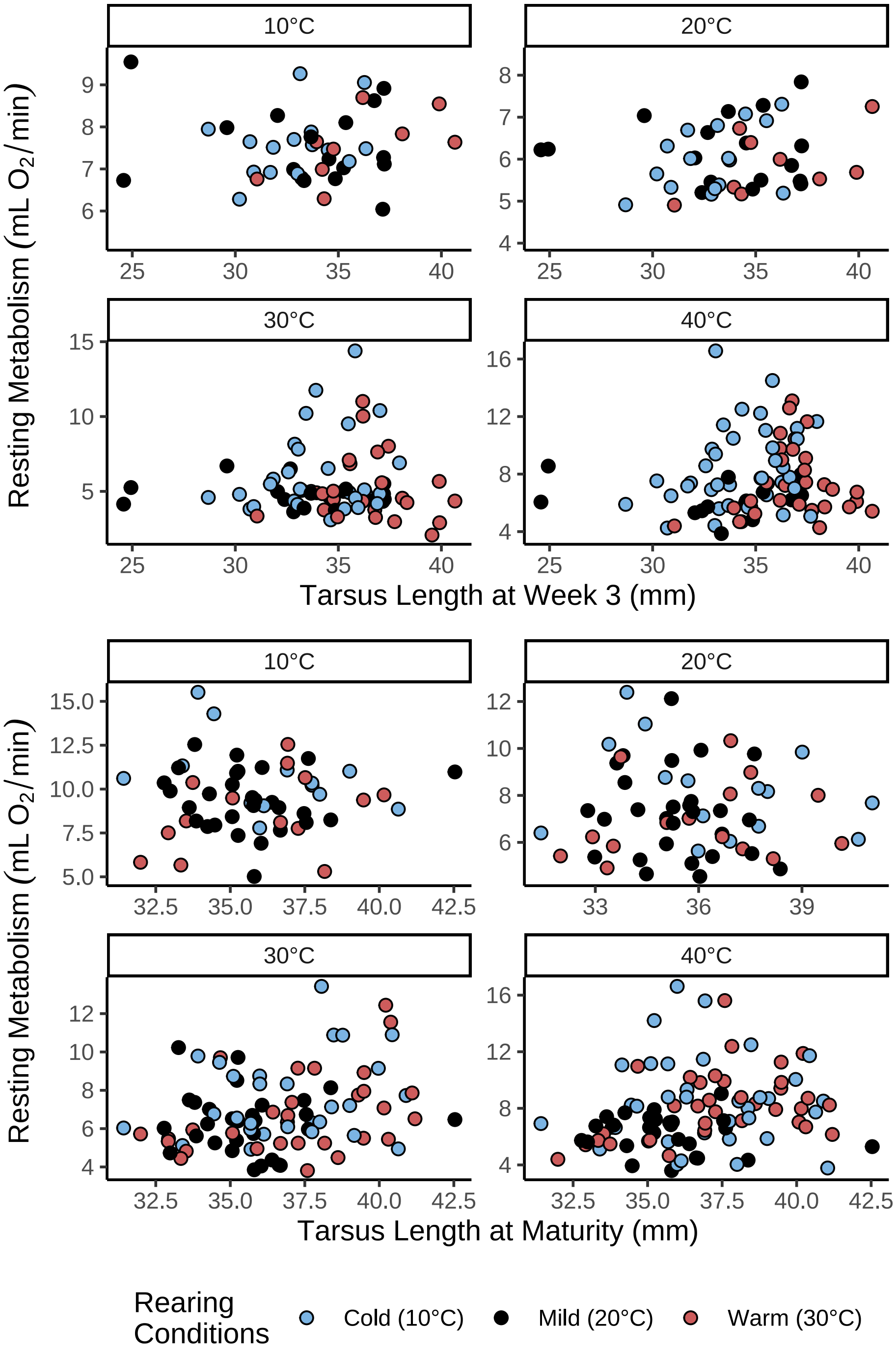


***Figure 3:*** *Resting metabolic rate (mL $O_2$/min) as a function of tarsus length (mm) in 3 and 8 week old Japanese quail. Resting metabolic rate was measured at four different thermal environments (10°C, 20°C, 30°C and 40°C), with individuals measured in at least two environments. Measured quail were reared in one of three difference rearing conditions: (1) cold (10°C) until at least three weeks of age, (2) mild conditions (20°C) for eight weeks of age, or (3) warmth (30°C) until at least three weeks of age.*

```
showtext_auto(enable = FALSE)
```

```
# Repeating exploratory plots with mass adjusted energy expenditure in place
# of un-adjusted energy expenditure.
```

```
residualVO2Tarsus3Weeks <- ggplot(all %>%
  filter(week == 3) %>%
  mutate(Ta = paste0(Ta, "°C")) %>%
  mutate(
    "VO2" = ((VO2 * 20 / 60) / 1000) * (3600 * 24),
    pretreatment = ifelse(pretreatment == "cold", "Cold (10°C)",
      ifelse(pretreatment == "neutral", "Mild (20°C)",
        "Warm (30°C)"
      )
    )
  ) %>%
  drop_na(mass, tarsusLengthMean, VO2, Ta) %>%
  mutate(
    "residualTarsus" =
      residuals(brm(tarsusLengthMean ~ mass,
        prior = c(
          set_prior("normal(1, 2)", class = "b"),
          set_prior("exponential(0.05)", class = "Intercept")
        ),
        data = .,
        cores = 1, chains = 4,
        seed = 100, family = "gaussian",
        iter = 50000, warmup = 5000, thin = 10,
        control = list(adapt_delta = 0.99, max_treedepth = 16),
        silent = TRUE, refresh = 0,
        file = "./models/vo2ResidualPlotTarsus3Weeks-Median.Rds"
      ),
      robust = TRUE)[, "Estimate"]
  ) %>%
  drop_na(pretreatment), aes(x = residualTarsus, y = VO2)) +
  facet_wrap(~Ta, scales = "free") +
  geom_point(pch = 21, colour = "black", size = 2, aes(fill = pretreatment)) +
  theme_classic() +
  scale_fill_manual(values = c("#7BB4E3", "black", "#CD5C5C"),
                    name = "Rearing\nConditions") +
  xlab("Residual Tarsus Length at Week 3 (mm)")  +
  ylab(TeX('$Resting~Metabolism~(mL~O_{2}/min)$'))

residualVO2Tarsus8Weeks <- ggplot(all %>%
  filter(week == 8) %>%
  mutate(Ta = paste0(Ta, "°C")) %>%
  mutate(
    "VO2" = ((VO2 * 20 / 60) / 1000) * (3600 * 24),
    pretreatment = ifelse(pretreatment == "cold", "Cold (10°C)",
      ifelse(pretreatment == "neutral", "Mild (20°C)",
        "Warm (30°C)"
      )
    )
  ) %>%
  drop_na(mass, tarsusLengthMean, VO2, Ta) %>%
  mutate(
    "residualTarsus" =
      residuals(brm(tarsusLengthMean ~ mass,
        prior = c(
          set_prior("normal(1, 2)", class = "b"),
          set_prior("exponential(0.05)", class = "Intercept")
        ),
        data = .,
        cores = 1, chains = 4,
        seed = 100, family = "gaussian",
        iter = 50000, warmup = 5000, thin = 10,
        control = list(adapt_delta = 0.99, max_treedepth = 16),
        silent = TRUE, refresh = 0,
        file = "./models/vo2ResidualPlotTarsus8Weeks-Median.Rds"
      ),
      robust = TRUE)[, "Estimate"]
  ) %>%
```

```
  drop_na(pretreatment), aes(x = residualTarsus, y = VO2)) +
  facet_wrap(~Ta, scales = "free") +
  geom_point(pch = 21, colour = "black", size = 2, aes(fill = pretreatment)) +
  theme_classic() +
  scale_fill_manual(values = c("#7BB4E3", "black", "#CD5C5C"),
                    name = "Rearing\nConditions") +
  xlab("Residual Tarsus Length at Maturity (mm)")  +
  ylab(TeX('$Resting~Metabolism~(mL~O_{2}/min)$'))

showtext_auto()
ggarrange(residualVO2Tarsus3Weeks,
  residualVO2Tarsus8Weeks,
  ncol = 1,
  common.legend = TRUE
)
```

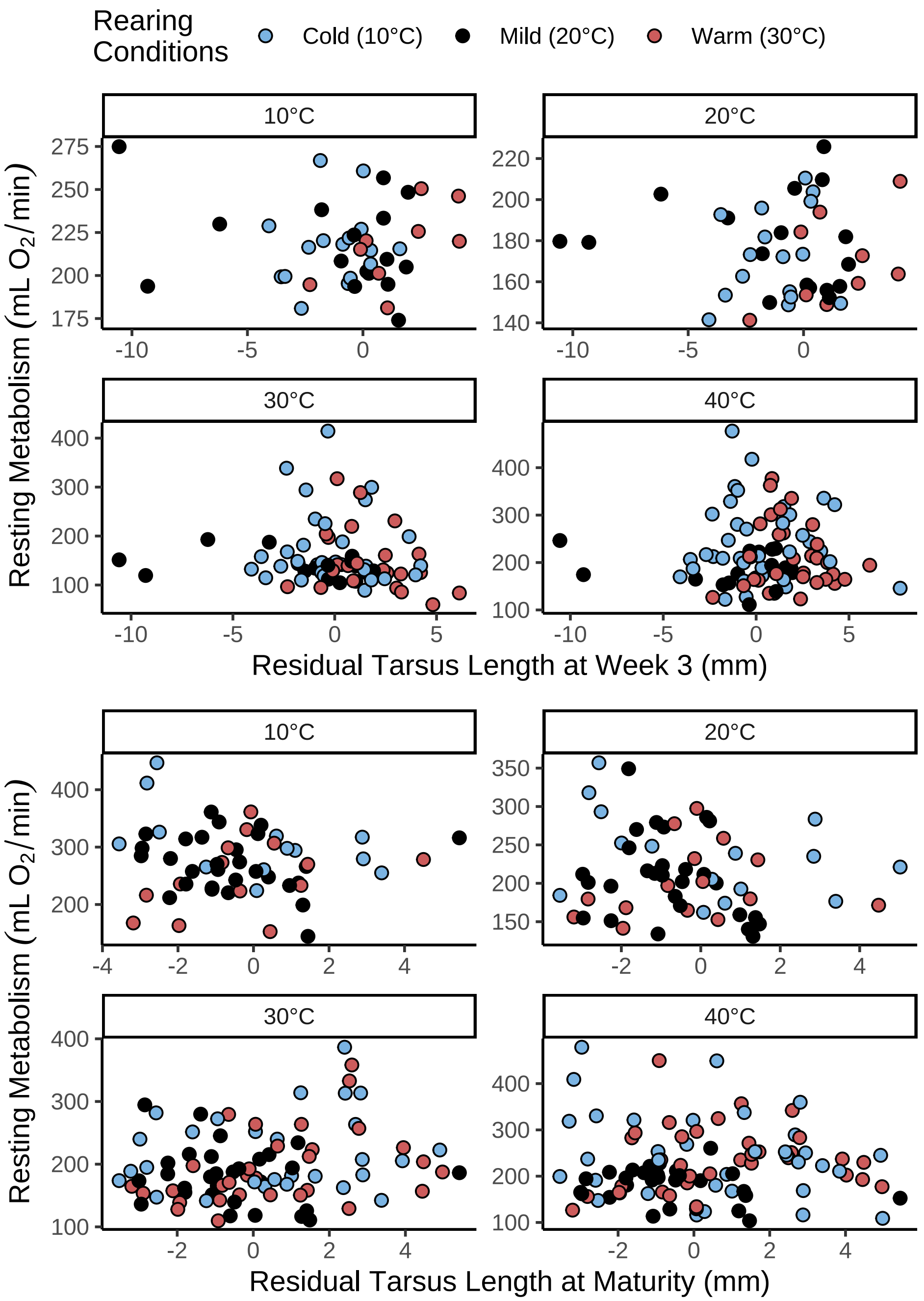


***Figure 4:*** *Resting metabolic rate (mL $O_2$/min) as a function of relative tarsus length (mm; residuals from a tarsus length [mm] by body mass [g] regression) in 3 and 8 week old Japanese quail. Resting metabolic rate was measured at four different thermal environments (10°C, 20°C, 30°C and 40°C), with individuals measured in at least two environments. Measured quail were reared in one of three difference rearing conditions: (1) cold (10°C) until at least three weeks of age, (2) mild conditions (20°C) for eight weeks of age, or (3) warmth (30°C) until at least three weeks of age.*

```
showtext_auto(enable = FALSE)
```

#### Visualising and quantifying costs of thermoregulation (metabolic slopes) in the cold

With data collated, we proceed by visualising the rates at which resting metabolism increases in cold (i.e. metabolic slopes), depending on individual morphology (here, body mass and tarsus length) and rearing conditions. Visualisation are first conducted using raw resting metabolism data obtained from quail during development (three weeks of age) and at adulthood (eight weeks of age).

```
## Visualising metabolic slopes by tarsus size at
# temperatures below thermoneutrality

rSlopesMass <- all %>%
  filter(week %in% c(3, 8) & Ta < 35) %>%
  mutate("Age" = paste0("Age = ", week, " weeks")) %>%
  drop_na(Ta, week, mass, VO2, pretreatment) %>%
  group_by(Age) %>%
  mutate(
    massScaledA = (mass - min(mass)) /
      (max(mass) - min(mass)),
    massScaledB = (mass - mean(mass)) / sd(mass)
  ) %>%
  mutate(massScaledA = exp(massScaledA)) %>%
  ungroup() %>%
  ggplot(aes(
    x = Ta, y = VO2,
    colour = massScaledB,
    group = ring
  )) +
  facet_wrap(~Age, scales = "free") +
  geom_smooth(
    method = "lm", linetype = "solid", alpha = 0.5, se = FALSE,
    aes(size = massScaledA)
  ) +
  scale_colour_gradient(
    low = "black", high = "grey99", name = "Relative\nBody Mass",
    breaks = c(-2, -1, 0, 1, 2),
    labels = c("Mean - 2 SDs", "Mean - 1 SD",
               "Mean", "Mean + 1 SD", "Mean + 2 SDs")
  ) +
  scale_size_continuous(range = c(0.1, 2), guide = "none") +
  xlab("Ambient Temperature (°C)") +
  ylab(
   TeX('$\\overset{Resting~Energy~Expenditure}{(mL~O_{2}/min)}$')
   ) +
  theme_classic() +
  theme(
    axis.title = element_text(family = "Noto Sans"),
    legend.title = element_text(family = "Noto Sans")
  )

rSlopesTarsus <- all %>%
  filter(week %in% c(3, 8) & Ta < 35) %>%
  mutate("Age" = paste0("Age = ", week, " weeks")) %>%
  drop_na(Ta, week, tarsusLengthMean, VO2, pretreatment) %>%
  group_by(Age) %>%
  mutate(
    tarsusScaledA = (tarsusLengthMean - min(tarsusLengthMean)) /
      (max(tarsusLengthMean) - min(tarsusLengthMean)),
    tarsusScaledB = (tarsusLengthMean - mean(tarsusLengthMean)) /
      sd(tarsusLengthMean)
  ) %>%
  mutate(tarsusScaledA = exp(tarsusScaledA)) %>%
  ungroup() %>%
  ggplot(aes(
    x = Ta, y = VO2,
    colour = tarsusScaledB,
```

```
    group = ring
  )) +
  facet_wrap(~Age, scales = "free") +
  geom_smooth(
    method = "lm", linetype = "solid", alpha = 0.5, se = FALSE,
    aes(size = tarsusScaledA)
  ) +
  scale_colour_gradient(
    low = "black", high = "grey99",
    name = "Relative\nTarsus Length",
    breaks = c(-2, -1, 0, 1, 2),
    labels = c("Mean - 2 SDs", "Mean - 1 SD", "Mean", "Mean + 1 SD", "Mean + 2 SDs")
  ) +
  scale_size_continuous(range = c(0.1, 2), guide = "none") +
  xlab("Ambient Temperature (°C)") +
  ylab(
   TeX('$\\overset{Resting~Energy~Expenditure}{(mL~O_{2}/min)}$')
   ) +
  theme_classic() +
  theme(
    axis.title = element_text(family = "Noto Sans"),
    legend.title = element_text(family = "Noto Sans")
  )

rSlopesPlot <- rSlopesMass / rSlopesTarsus

showtext_auto()
print(rSlopesPlot)
```

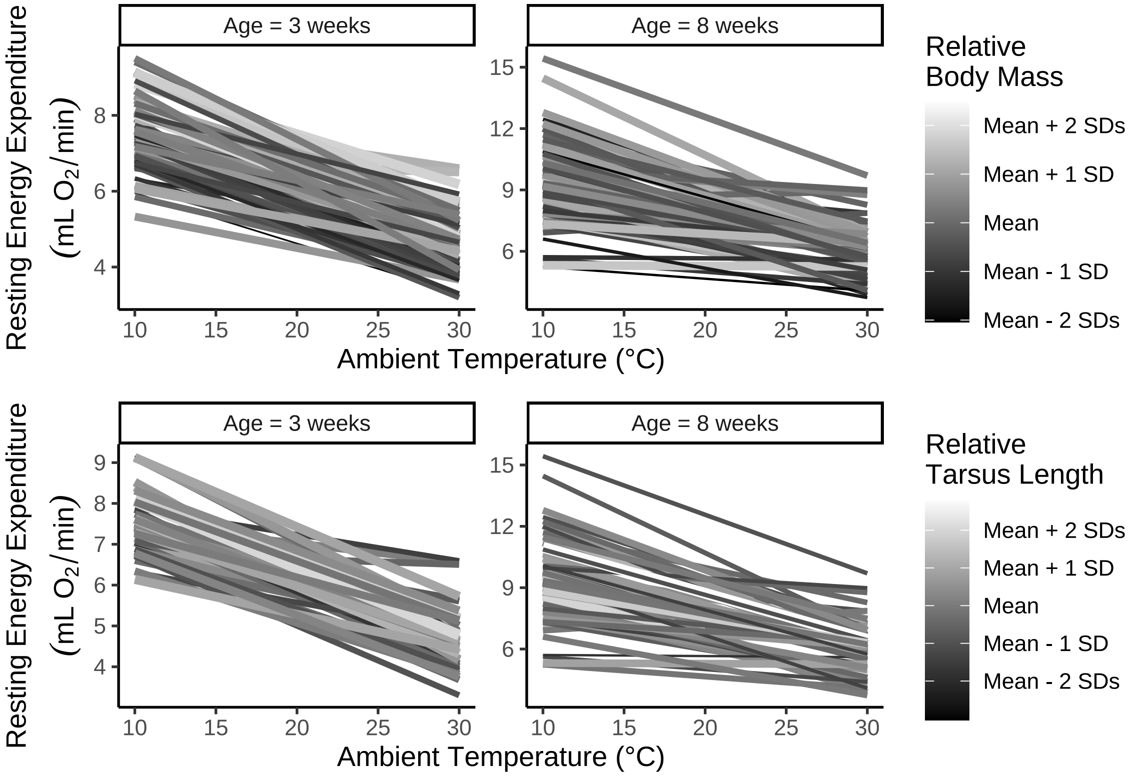


***Figure 5:*** *Resting energy expenditure of Japanese quail at various ambient temperatures (°C) and according to body mass (g) and tarsus lengths (mm). Each line represents energy expenditure for one individual. Line size is scaled by relative body mass and tarsus length for each plot respectively, with thicker lines representing relatively large phenotype and thinner lines representing relative small phenotype. Lines are scaled per age of measurement.*

```r
showtext_auto(enable = FALSE)

ggsave("./plots/resistanceSlopes.jpg",
  rSlopesPlot,
  dpi = 800, height = 7, width = 9
)
```

Tentative correlations between tarsus length and metabolic slopes are further investigated by relativising tarsus length measurements with respect to body mass (here, as tarsus length residuals from a simple Bayesian linear model with broad priors).

```r
rSlopesPlotResiduals <- all %>%
  filter(week %in% c(3, 8) & Ta < 35) %>%
  drop_na(tarsusLengthMean, mass, week) %>%
  mutate(
    "residualTarsus" =
      residuals(brm(tarsusLengthMean ~ mass + week,
        prior = c(
          set_prior("normal(1, 2)", class = "b"),
          set_prior("exponential(0.05)", class = "Intercept")
        ),
        data = .,
        cores = 4, chains = 4,
        seed = 100, family = "gaussian",
        iter = 50000, warmup = 5000, thin = 10,
        control = list(adapt_delta = 0.99, max_treedepth = 16),
        silent = TRUE, refresh = 0,
        file = "./models/residualTarsusPlotModel-Median.Rds"
      ),
      robust = TRUE)[, "Estimate"]
  ) %>%
  mutate("tarsusBin" = ifelse(residualTarsus < 0,
                              "Below Average", "Above Average")) %>%
  mutate(
    "Age" = paste0("Age = ", week, " weeks"),
    "tarsusBin" = factor(tarsusBin, levels = c("Below Average",
                                               "Above Average"))
  ) %>%
  ggplot(aes(
    x = Ta, y = VO2,
    colour = tarsusBin,
    linetype = tarsusBin,
    group = ring
  )) +
  facet_wrap(~Age, scales = "free") +
  geom_smooth(method = "lm", linetype = "solid", alpha = 0.5, se = FALSE) +
  scale_colour_manual(values = c("black", "grey80"),
                      name = "Relative\nTarsus Length") +
  scale_linetype_manual(values = c("solid", "dashed"),
                        name = "Relative\nTarsus Length") +
  xlab("Ambient Temperature (°C)") +
  ylab(TeX('$\\overset{Resting~Energy~Expenditure}{(mL~O_{2}/min)}$')) +
  theme_classic() +
  theme(
    axis.title = element_text(family = "Noto Sans", size = 14),
    legend.title = element_text(family = "Noto Sans", size = 14)
  )

showtext_auto()
print(rSlopesPlotResiduals)
```

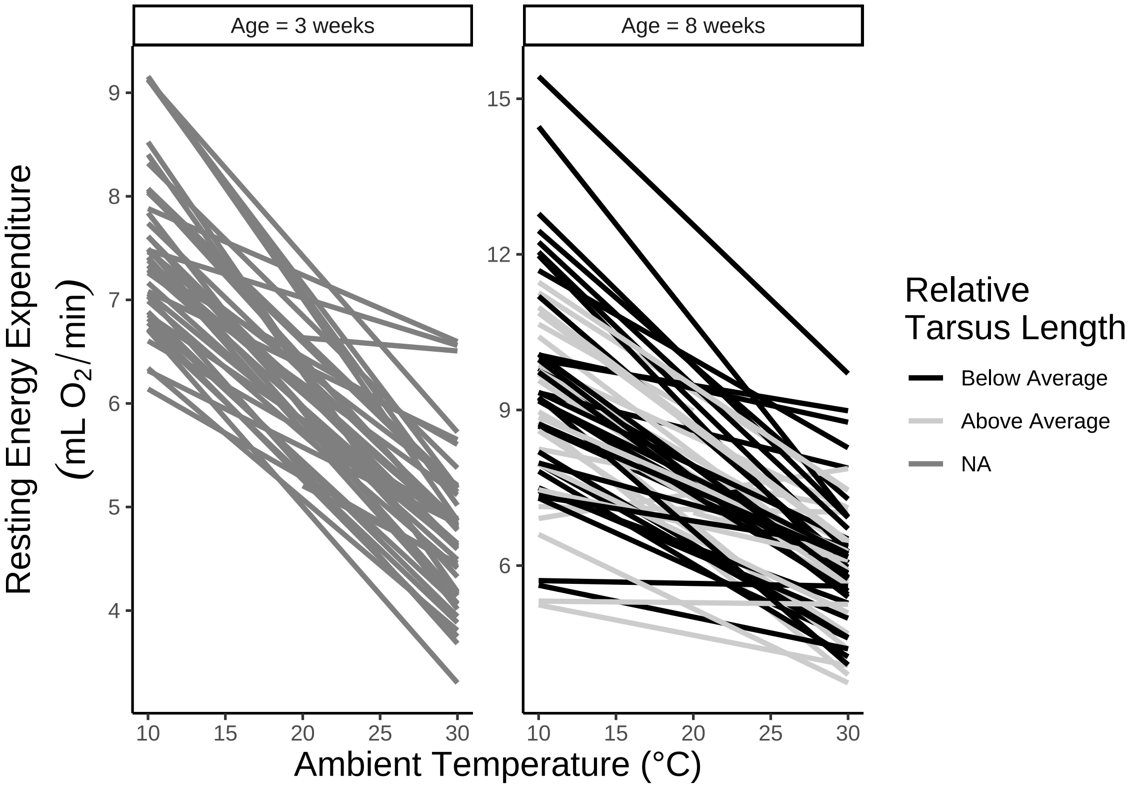


***Figure 6:*** *Resting energy expenditure of Japanese quail at various ambient temperatures (°C) and according to relative tarsus lengths (mm; residuals from a regression against body mass [g]). Each line represents energy expenditure for one individual and lines are now coloured according to whether an individual's relative tarsus length fell above or below the average for its age.*

```
showtext_auto(enable = FALSE)

ggsave("./plots/resistanceSlopesResiduals.jpg",
  rSlopesPlotResiduals,
  dpi = 800, height = 7, width = 9
)
```

Next, we formally quantify metabolic slopes in response to our cold challenge per individual and during development (3 weeks of age). To do so, we model resting metabolism as a function of air temperatures between 10°C - 30°C while allowing slopes to vary by individual. However, given that: (1) morphology can influence both intercepts and slopes of temperature-metabolism relationships, and (2) intercepts and slopes may be inter-related (thus complicating analyses), we first relativised resting metabolism measurements as fold-changes from those observed at thermoneutrality (30°C; Persson et al, 2024) per individual; thus, resting metabolism at thermoneutrality for all individuals was rooted at 1. To accommodate this rooting, we then set thermoneutrality (30°C) as our model intercept and proceeded to estimating slopes of temperature-metabolism relationships alone. Last, since variance in metabolism measurements can vary greatly by measurement temperature, we allow our error structure to vary by categorical ambient temperature while assuming an increase in error at lower ambient temperature. Formulae for our model were therefore:

$$Relative\ VO_{2ij} \sim 1 + (\beta_1 + v_j) \cdot Ta_{ij} + \epsilon_{ij}$$

where:

$$Relative\ VO_{2ij} = VO_{2ij}/Thermoneutral\ VO_{2j}$$
$$v_j \sim \mathcal{N}(0, V_v)$$
$$\epsilon_{ij} \sim S\mathcal{N}(0, Ve)$$
$$Ve \sim e^{(\tau_0 + \tau_1 \cdot Ta_{10°C;ij})}$$

and where $i$ represents an observation, $j$ represents an individual, $v$ represents an individual's deviance from the population metabolic slope ($\beta_1$), which is normally-distributed around 0 with a standard deviation of $V_v$, $\epsilon$ represents the model error term with standard deviation $Ve$, $\tau_0$ indicates the natural-log transformed standard deviation of $\epsilon$ for measures at 20°C, $Ta_{10°C;\ ij}$ indicates whether an observation was made at 10°C (true = 1, false = 0) and $\tau_1$ indicates the natural-log transformed adjustment to $\epsilon$ made for these observations. Metabolic slopes are then calculated per individual as:

$$Metabolic\ Slope_j = \beta_1 + v_j$$

Priors for $\beta_1$ were informed by Saarela and Heldmaier (1987) and set as skew-normal with an $\xi$ value of -0.018, $\omega$ value of 0.02, and $\alpha$ value of -5 (thus setting the mean for our prior at approximately that reported by Saarela and Heldmaier, 1987). Priors for other variables were weak, with that for $V_v$ set as exponential (*Exp*) with a $\lambda$ of 10, and those for $\tau_0$, $\tau_1$, and $\tau_2$ set as normal with means of -1.5, 0, and -5 respectively, and standard deviations of 1, 0.25, and 1 respectively.

Only individuals for whom resting metabolism was measured at the lower critical temperature (~30°C) and at least one ambient temperature below (10°C or 20°) were used in this model and subsequent analyses. Further, a linear relationship between ambient temperature and resting metabolism is first visually confirmed.

```r
birdVector <- merge(
  all %>%
    drop_na(VO2) %>%
    filter(week == 3 & Ta == 30) %>%
    select(ring) %>%
    distinct(),
  all %>%
    drop_na(VO2) %>%
    filter(week == 3 & Ta %in% c(10, 20)) %>%
    select(ring) %>%
    distinct(),
  by = "ring", all = FALSE
) %>% pull(ring)

modData <- subset(all, week == 3 & Ta < 35 & ring %in% birdVector)

# Checking correlation between ambient temperature
# and metabolic rate visually.

modData %>%
  ggplot(aes(x = Ta, y = VO2)) +
  geom_point(
    pch = 21, size = 1.5, alpha = 0.5, colour = "black", fill = "grey50",
    position = position_jitter(width = 0.2)
  ) +
  stat_summary(geom = "errorbar", fun.data = "mean_se",
               width = 0.3, colour = "black") +
  stat_summary(
    geom = "point", fun = "mean", size = 4, pch = 21,
    colour = "black",
    fill = "grey50"
  ) +
  geom_smooth(method = "lm", colour = "black", linetype = "dashed", se = FALSE) +
  xlab("Ambient Temperature (°C)") +
```

```
  ylab(TeX('$\\overset{Resting~Energy~Expenditure}{(mL~O_{2}/min)}$')) +
  theme_classic()
```

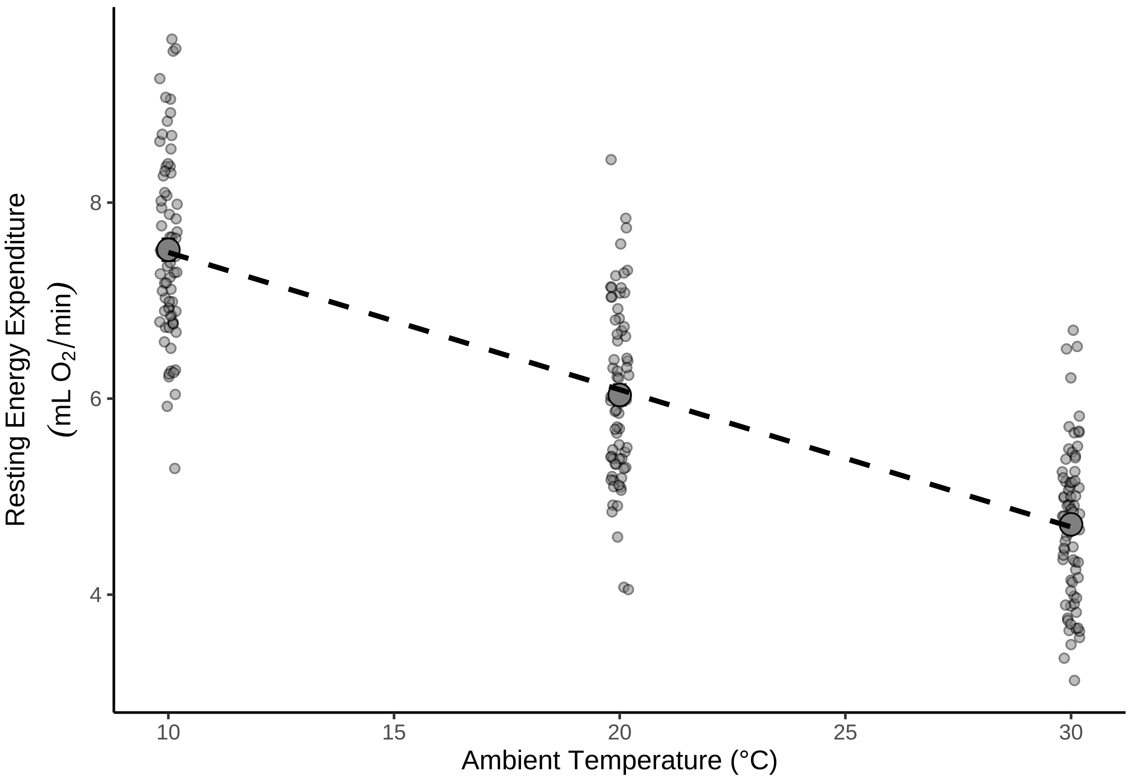


***Figure 7:*** *Relationship between resting metabolism and ambient temperature in three week old Japanese quail. Large dots represent means across quail and small dots represent individual measurements. Errorbars around large dots represent standard errors and the dotted line represents the linear, line of best fit as estimated by the R package ggplot2 (Wickham, 2011).*

To confirm suitability of our priors, we visualise how well predictions from our priors alone capture distributions of metabolism measurements in our data.

```
# Setting 30°C as 0°C and checking suitability of priors.

slopes3Weeks_ppCheck <- brm(
  data = modData %>%
    select(ring, Ta, VO2) %>%
    pivot_wider(id_cols = "ring", names_from = Ta, values_from = VO2) %>%
    mutate(`10` = `10`/`30`,
           `20` = `20`/`30`,
           `30` = 1) %>%
    pivot_longer(-c("ring"), names_to = "Ta", values_to = "VO2") %>%
    mutate(Ta = as.integer(Ta)) %>%
    mutate(Ta = Ta - 30) %>%
    filter(Ta < 0) %>%
    mutate("discreteTa" = ifelse(Ta == -20, "A", "B")),
  family = "gaussian",
  bf(VO2 ~ 1 + B,
     B ~ 0 + Ta + (0 + Ta | ring),
     sigma ~ discreteTa,
     nl = TRUE),
  prior = c(
    set_prior("skew_normal(-0.018, 0.02, -2.5)", nlpar = "B",
```

```
              class = "b", coef = "Ta"),
    set_prior("exponential(2.5)", nlpar = "B", class = "sd",
              coef = "Ta", group = "ring"),
    set_prior("skew_normal(-0.2, 1, -5)", dpar = "sigma",
              class = "Intercept"),
    set_prior("normal(0.25, 0.25)", dpar = "sigma",
              class = "b")
  ),
  iter = 50000, warmup = 10000, thin = 10,
  chains = 4, cores = 4,
  control = list(adapt_delta = .96),
  silent = TRUE, refresh = 0,
  sample_prior = "only",
  file = "./models/threeWeekSlopesPPCheck.Rds"
)

pp_check2(slopes3Weeks_ppCheck,
          xlab = "Fold Metabolism at 30°C") +
  xlim(c(-10, 10))
```

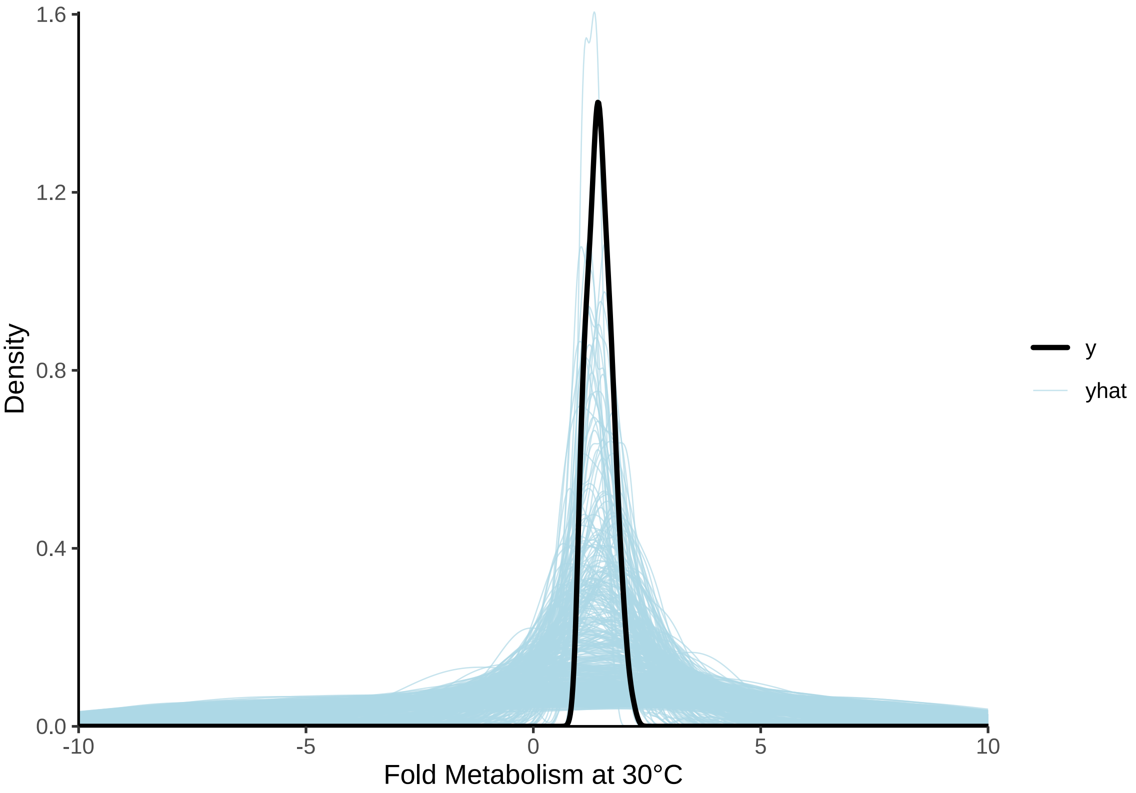


***Figure 8:*** *Prior predictive check for a Bayesian linear, mixed effects model predicting relative resting metabolism (fold metabolism at thermoneutrality) of three week old Japanese quail by ambient temperature (raw ambient temperatures ranging from 10°C - 30°C). Light blue lines represent densities of resting metabolism values as predicted by model priors alone. The dark blue line represent the true density of resting metabolism values. Clear overlap between the dark blue and light blue lines indicates that priors are suitable.*

Priors are wide but capture our peak density well. We proceed to constructing our full model accordingly and evaluating chain mixing and intra-train autocorrelation.

```
# Priors are slightly low and wide, but acceptable. Proceeding to full model.

slopes3Weeks <- brm(
```

```
  data = modData %>%
    select(ring, Ta, VO2) %>%
    pivot_wider(id_cols = "ring", names_from = Ta, values_from = VO2) %>%
    mutate(`10` = `10`/`30`,
           `20` = `20`/`30`,
           `30` = 1) %>%
    pivot_longer(-c("ring"), names_to = "Ta", values_to = "VO2") %>%
    mutate(Ta = as.integer(Ta)) %>%
    mutate(Ta = Ta - 30) %>%
    filter(Ta < 0) %>%
    mutate("discreteTa" = ifelse(Ta == -20, "A", "B")),
  family = "gaussian",
  bf(VO2 ~ 1 + B,
     B ~ 0 + Ta + (0 + Ta | ring),
     sigma ~ discreteTa,
     nl = TRUE),
  prior = c(
    set_prior("skew_normal(-0.018, 0.02, -2.5)", nlpar = "B",
              class = "b", coef = "Ta"),
    set_prior("exponential(2.5)", nlpar = "B", class = "sd",
              coef = "Ta", group = "ring"),
    set_prior("skew_normal(-0.2, 1, -5)", dpar = "sigma",
              class = "Intercept"),
    set_prior("normal(0.25, 0.25)", dpar = "sigma",
              class = "b")
  ),
  iter = 100000, warmup = 50000, thin = 10,
  chains = 4, cores = 4,
  control = list(adapt_delta = .98, max_treedepth = 14),
  silent = TRUE, refresh = 0,
  file = "./models/threeWeekSlopes.Rds"
)

# Checking Gelman-Rubin statistic and ratio of effective sample sizes to sample sizes.

ggarrange(
  mcmc_rhat(rhat(slopes3Weeks)) +
    theme(legend.position = "none") +
    xlab(
      TeX('$\\hat{R}$')
    ),
  mcmc_neff(neff_ratio(slopes3Weeks), size = 2) +
    theme(legend.position = "none") +
    xlab(
      TeX('$N_{eff}/N~Ratio$')
    )
)
```

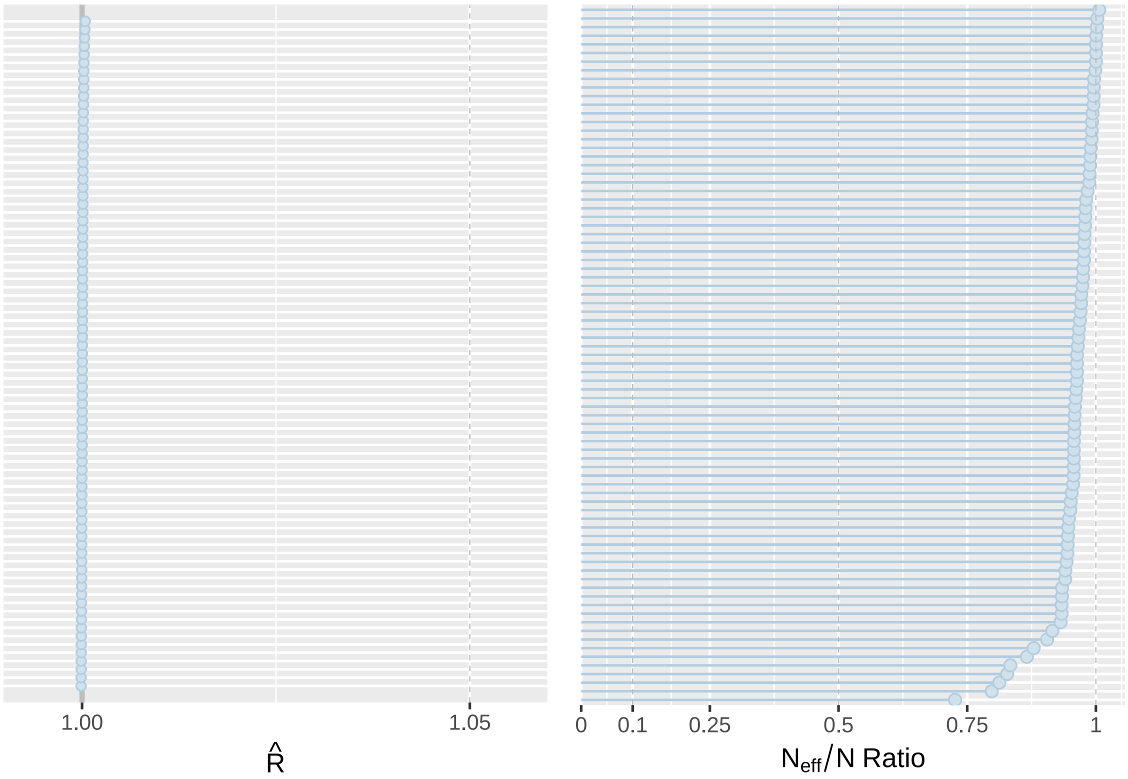


***Figure 9:*** *Gelman-Rubin statistics ($\hat{R}$s) and ratio of effective sample sizes to sample sizes for coefficients from a Bayesian, linear-mixed effects model predicting relative resting metabolism fold metabolism at thermoneutrality) of three week old Japanese quail across 10°C - 30°C thermal environemnts.*

Chains appear well mixed (Gelman-Rubin statistics [$\hat{R}$] laying near 1) and high effective sample size to sample size ratios indicate minimal chain autocorrelation. We proceed by visualising posterior densities and predictions, alongside estimating fit (here, using a Bayesian "$R^2$").

```
pp_check2(slopes3Weeks,
          xlab = paste0("Relative Resting Energy Expenditure\n",
          "(Fold Metabolism at 30°C)")
)
```

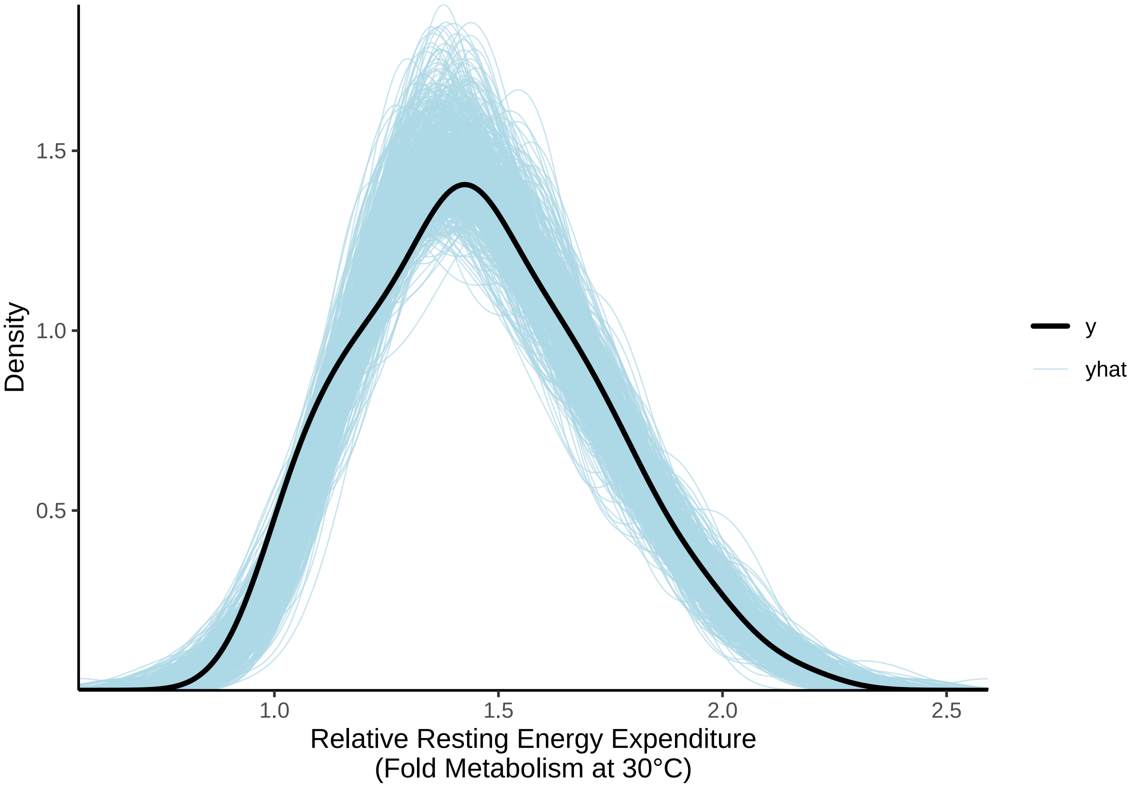


***Figure 10:*** *Posterior predictive check for a Bayesian linear, mixed effects model predicting relative resting metabolism (fold resting metabolism at thermoneutrality) of three week old Japanese quail by ambient temperature (raw ambient temperatures ranging from 10°C - 30°C). Light blue lines represent densities of resting metabolism values as drawn from model posteriors. The dark blue line represent the true density of resting metabolism values. Clear overlap between the dark blue and light blue lines indicates strong model fit.*

```
# Good. Checking distributions of metabolic slope estimates per bird
```

```
as.data.frame(slopes3Weeks) %>%
  select(starts_with("r_")) %>%
  pivot_longer(everything(), names_to = "ring",
               values_to = "slope") %>%
  mutate("ring" =
           gsub("\\,.*", "",
                gsub(".*\\[", "", ring)
                )
  ) %>%
  mutate(slope = fixef(slopes3Weeks)[1,"Estimate"] +
           slope) %>%
  ggplot(aes(x = slope)) +
  facet_wrap(~ring) +
  geom_density() +
  xlab("Metabolic Slope") +
  ylab("Density") +
  theme_classic() +
  theme(axis.text.x = element_blank(),
        axis.ticks.x = element_blank(),
        axis.text.y = element_blank(),
        axis.ticks.y = element_blank()
        )
```

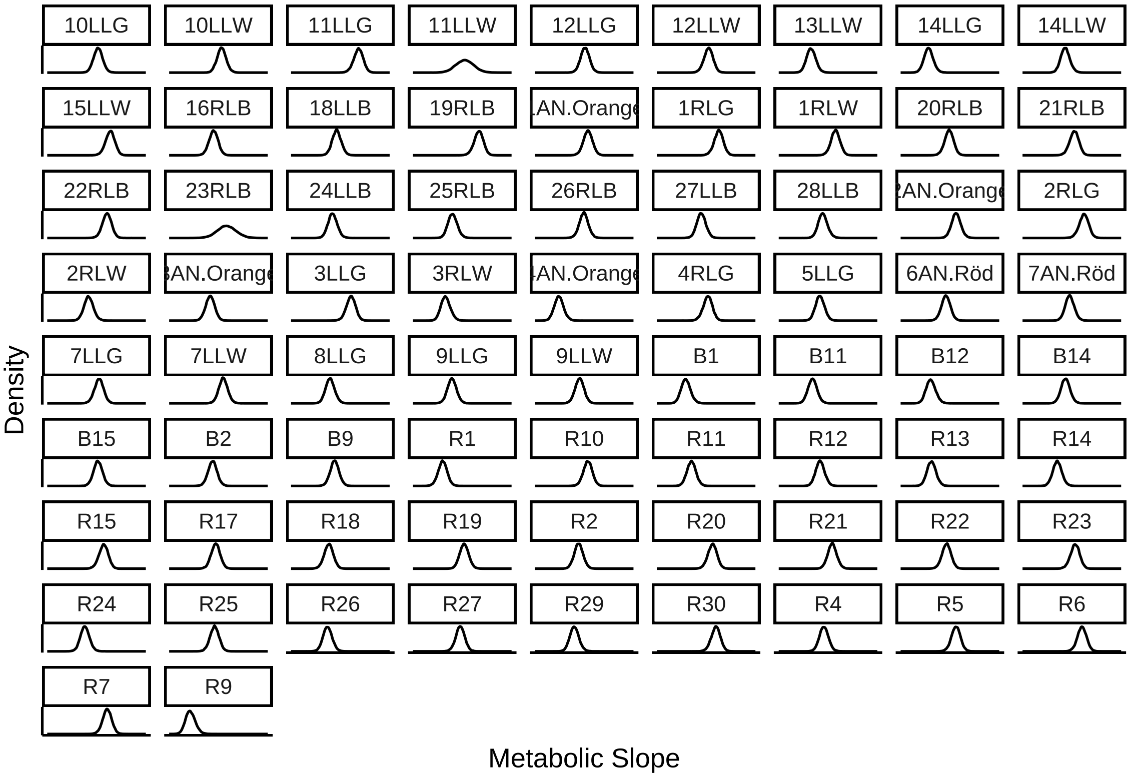


***Figure 11:*** *Posterior densities of metabolic slopes, per individual, from a Bayesian linear, mixed effects model predicting relative resting metabolism (fold resting metabolism at thermoneutrality) of three week old Japanese quail by ambient temperature (raw ambient temperatures ranging from 10°C - 30°C).*

```
# Reasonably normal and no multi-modality. Using medians for measure of
# central tendancy.

caption <- paste0("R\\textsuperscript{2} for Bayesian linear, mixed effects",
                  " model predicting relative resting metabolism (fold ",
                  "resting metabolism at thermoneutrality) ",
                  "of three week old Japanese quail by mean-centred ambient ",
                  "temperature (raw ambient temperatures ranging from ",
                  "10°C - ",
                  "30°C).")
)

brms::bayes_R2(slopes3Weeks,
               robust = TRUE, ndraws = 1000) %>%
  as.data.frame() %>%
  remove_rownames() %>%
  mutate(Estimate = round(Estimate, digits = 4),
         Est.Error = round(Est.Error, digits = 4),
         Q2.5 = round(Q2.5, digits = 4),
         Q97.5 = round(Q97.5, digits = 4)
         ) %>%
  rename(
    "R\\textsuperscript{2}" = Estimate, "Standard Error" = Est.Error,
    `2.5\\% CI` = `Q2.5`, `97.5\\% CI` = `Q97.5`
  ) %>%
  kbl(.,
    longtable = T, booktabs = T, format = "latex",
    caption = caption, escape = FALSE,
  ) %>%
  kable_styling(latex_options = "striped")
```

***Table 2:*** *$R^2$ for Bayesian linear, mixed effects model predicting relative resting metabolism (fold resting metabolism at thermoneutrality) of three week old Japanese quail by mean-centred ambient temperature (raw ambient temperatures ranging from 10°C - 30°C).*

| $R^2$ | Standard Error | 2.5% CI | 97.5% CI |
|---|---|---|---|
| 0.788 | 0.028 | 0.7119 | 0.8281 |

```
# Good and R2 reasonably high. Checking residuals.
```

```
p1 <- slopes3Weeks$data %>%
  mutate("Residuals" = residuals(slopes3Weeks,
    method = "posterior_predict",
    type = "pearson",
    robust = TRUE
  )[, "Estimate"]) %>%
  ggplot(aes(x = Residuals)) +
  geom_density(colour = "black", fill = "grey50", adjust = 1) +
  theme_classic() +
  xlab("Pearson Residuals") +
  ylab("Density")

p2 <- as.data.frame(slopes3Weeks) %>%
  select("Ta" = b_B_Ta, "ring" = sd_ring__B_Ta) %>%
  pivot_longer(everything(),
    names_to = "var",
    values_to = "value"
  ) %>%
  ggplot(aes(sample = value)) +
  facet_wrap(~var, scales = "free") +
  stat_qq(colour = "grey50") +
  stat_qq_line() +
  xlab("Theoretical") +
  ylab("Sample") +
  theme_classic()

p3 <- slopes3Weeks$data %>%
  mutate(
    "Residuals" = residuals(slopes3Weeks,
      type = "pearson",
      robust = TRUE
    )[, "Estimate"],
    "Fitted" = fitted(slopes3Weeks,
      robust = TRUE
    )[, "Estimate"]
  ) %>%
  ggplot(aes(x = Fitted, y = Residuals)) +
  geom_point(size = 2, pch = 21, colour = "black", fill = "grey50") +
  xlab("Fitted Values") +
  ylab("Pearson Residuals") +
  theme_classic()

(p1 + p2) / p3
```

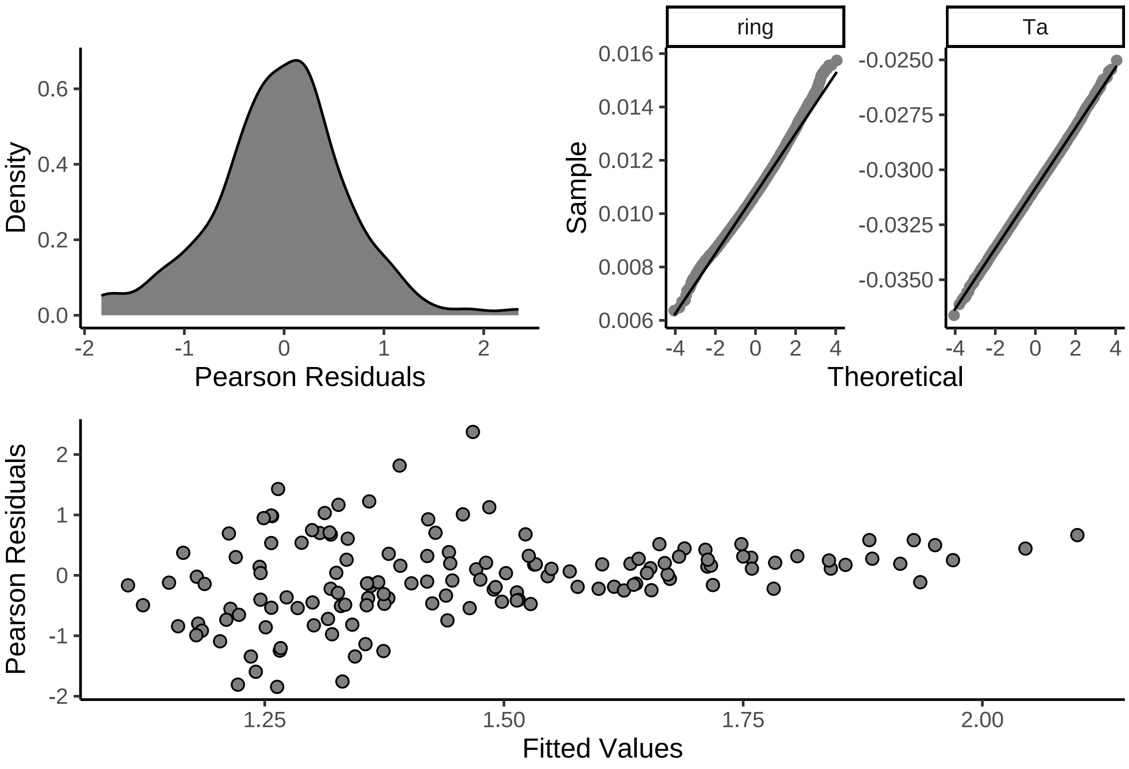


***Figure 12:*** *Distributions of Pearson residuals and posterior densities drawn from Bayesian linear, mixed effects model predicting relative resting metabolism (fold resting metabolism at thermoneutrality) of three week old Japanese quail by ambient temperature (raw ambient temperatures ranging from 10°C - 30°C).*

```
# Good. Some straying from normality among group-level effect of ring, although minor.
```

For metabolic slopes to vary by morphology or thermal history, we first expect that consistent differences in slopes among individual are detectable. To test this expectation, we quantify conditional repeatability of our metabolic slopes according to Schielzeth and Nakagawa (2022), then visualise and formally test its deviance from null expectations.

```
# Visualising repeatability of slopes and intercepts.
# Note that intercept repeatabilities are calculated classically,
# and slope repeatabilities calculated as per Schielzeth and Nakagawa (2022).

as.data.frame(slopes3Weeks) %>%
  mutate(
    "Vs" = sd_ring__B_Ta^2 * var(slopes3Weeks$data$Ta) +
      mean(slopes3Weeks$data$Ta)^2 * sd_ring__B_Ta^2,
    "Vf" = b_B_Ta^2 * var(slopes3Weeks$data$Ta)
  ) %>%
  mutate("Slope" = Vs / (Vf + exp(b_sigma_Intercept)^2 +
    exp(b_sigma_Intercept + b_sigma_discreteTaB)^2)) %>%
  ggplot(aes(x = Slope)) +
  geom_density(alpha = 0.5, colour = "black", fill = "grey80") +
  theme_classic() +
  xlab("Metabolic Slope Repeatability") +
  ylab("Density") +
  theme(legend.position = "none")
```

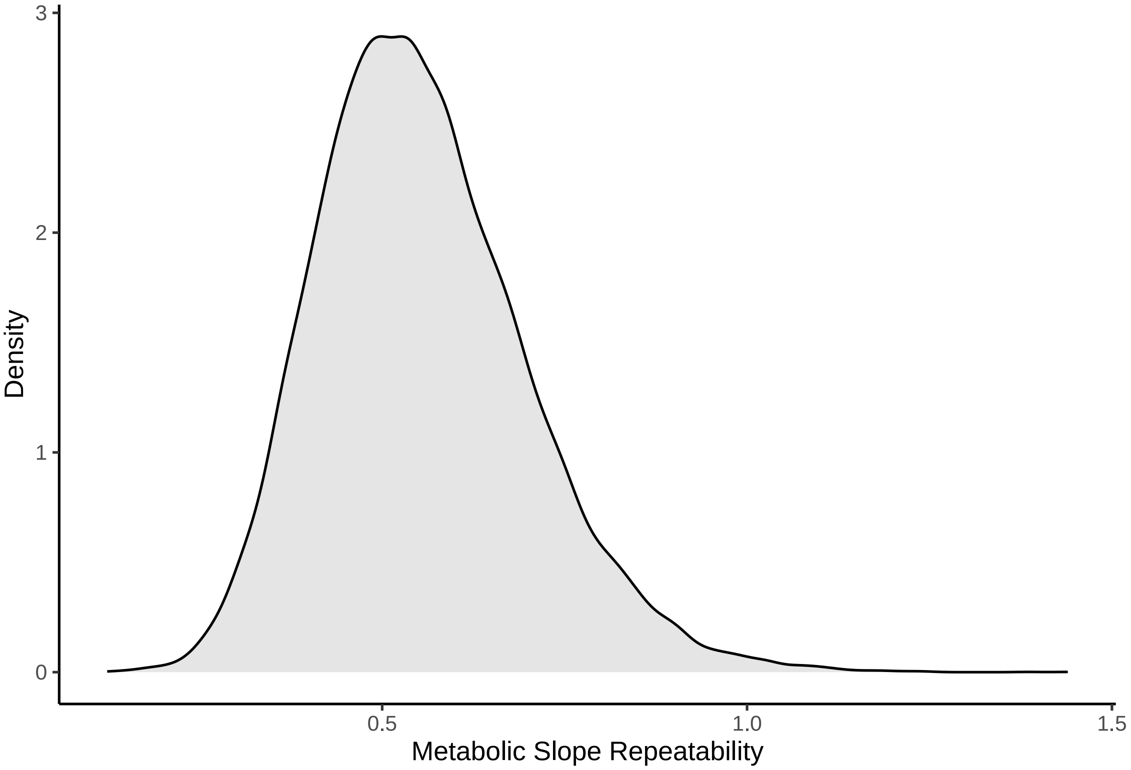


***Figure 13:*** *Conditional repeatability of group-level (here, individual-level) slopes derived from a Bayesian, linear mixed effects model predicting relative resting metabolism (fold metabolism at thermoneutrality) of three week old Japanese quail across ambient temperature (10°C - 30°C). Trends are estimated per individual and repeatabilities calculated from variability within and among trend parameters, as per Schielzeth and Nakagawa (2022).*

```
# Quantifying median and 95% quantile intervals.

repeatabilityFrame <-
  as.data.frame(slopes3Weeks) %>%
  mutate(
    "Vs" = sd_ring__B_Ta^2 * var(slopes3Weeks$data$Ta) +
      mean(slopes3Weeks$data$Ta)^2 * sd_ring__B_Ta^2,
    "Vf" = b_B_Ta^2 * var(slopes3Weeks$data$Ta)
  ) %>%
  mutate("Slope" = Vs / (Vf + exp(b_sigma_Intercept)^2 +
                           exp(b_sigma_Intercept + b_sigma_discreteTaB)^2)
        ) %>%
  select(Slope)

caption <- paste0('Conditional repeatability of individual-level ',
                  "slopes for the relationship between ",
                  "relative resting metabolism ",
                  "(fold metabolism at thermoneutrality) ",
                  "across ambient temperature (10°C ",
                  "- 30°C) in three week old ",
                  "Japanese quail. Repeatability is calculated following ",
                  "Schielzeth and Nakagawa (2022) and represents ",
                  "the posterior median. 95\\% credible intervals indicate ",
                  "quantile intervals."
)

repeatability3Weeks <-
    repeatabilityFrame %>%
      summarise("Median" = round(median(Slope), digits = 4),
```

```
                "95% CIs" =
                  paste0("[",
                         round(quantile(Slope, 0.025, type = 8),
                               digits = 4),
                         ", ",
                         round(quantile(Slope, 0.975, type = 8),
                               digits = 4),
                         "]")
                ) %>%
  kbl(.,
    longtable = T, booktabs = T, format = "latex",
    caption = caption
  ) %>%
  kable_styling(latex_options = "striped")

repeatability3Weeks
```

***Table 3:*** *Conditional repeatability of individual-level slopes for the relationship between relative resting metabolism (fold metabolism at thermoneutrality) across ambient temperature (10°C - 30°C) in three week old Japanese quail. Repeatability is calculated following Schielzeth and Nakagawa (2022) and represents the posterior median. 95% credible intervals indicate quantile intervals.*

| Median | 95% CIs |
|---|---|
| 0.5402 | [0.3141, 0.8706] |

```
#save_kable(repeatability3Weeks, "../tables/VO2Repeatability3Weeks.html")
```

To formally test an increase an repeatability from null expectations, we follow the approach described by Tabh et al (2022). Specifically, we randomly scramble individual identities, re-execute our model described above, recalculate conditional repeatability (here, a null repeatability) then test for a relative increase in true conditional repeatability from null repeatabilities.

```
slopes3WeeksScrambled <- brm(
  data = modData %>%
    select(ring, Ta, VO2) %>%
    pivot_wider(id_cols = "ring", names_from = Ta, values_from = VO2) %>%
    mutate(`10` = `10`/`30`,
           `20` = `20`/`30`,
           `30` = 1) %>%
    pivot_longer(-c("ring"), names_to = "Ta", values_to = "VO2") %>%
    mutate(Ta = as.integer(Ta)) %>%
    mutate(Ta = Ta - 30) %>%
    filter(Ta < 0) %>%
    mutate("ringScrambled" = sample(ring, nrow(.), replace = FALSE)) %>%
    mutate("discreteTa" = ifelse(Ta == -20, "A", "B")),
  family = "gaussian",
  bf(VO2 ~ 1 + B,
     B ~ 0 + Ta + (0 + Ta | ringScrambled),
     sigma ~ discreteTa,
     nl = TRUE),
  prior = c(
    set_prior("skew_normal(-0.018, 0.02, -2.5)", nlpar = "B",
              class = "b", coef = "Ta"),
    set_prior("exponential(2.5)", nlpar = "B", class = "sd",
              coef = "Ta", group = "ringScrambled"),
    set_prior("skew_normal(-0.2, 1, -5)", dpar = "sigma",
              class = "Intercept"),
    set_prior("normal(0.25, 0.25)", dpar = "sigma",
              class = "b")
  ),
  iter = 100000, warmup = 50000, thin = 10,
  chains = 4, cores = 4,
  control = list(adapt_delta = .98, max_treedepth = 14),
```

```
  silent = TRUE, refresh = 0,
  file = "./models/threeWeekSlopesScrambled.Rds"
)

# Visually comparing first

repeatabilityFrameScrambled <-
  as.data.frame(slopes3WeeksScrambled) %>%
  mutate(
    "Vs" = sd_ringScrambled__B_Ta^2 *
      var(slopes3WeeksScrambled$data$Ta) +
      mean(slopes3WeeksScrambled$data$Ta)^2 *
        sd_ringScrambled__B_Ta^2,
    "Vf" = b_B_Ta^2 * var(slopes3WeeksScrambled$data$Ta)
  ) %>%
  mutate("Slope" = Vs / (Vf + exp(b_sigma_Intercept)^2 +
    exp(b_sigma_Intercept + b_sigma_discreteTaB)^2)) %>%
  select(Slope)

repeatPlot <- rbind(
  repeatabilityFrame %>%
    mutate("Type" = "True Model"),
  repeatabilityFrameScrambled %>%
    mutate("Type" = "Null Model")
) %>%
  ggplot(aes(x = Slope, fill = Type)) +
  geom_density(colour = "black", adjust = 1, alpha = 0.8) +
  scale_fill_manual(values = c("grey20", "grey80")) +
  xlab("Conditional Repeatability at 3 Weeks") +
  ylab("Density") +
  theme_classic() +
  theme(
    axis.text = element_text(family = "Noto Sans"),
    axis.title = element_text(family = "Noto Sans"),
    strip.text.x = element_text(family = "Noto Sans", colour = "black")
  )

showtext_auto()
repeatPlot
```

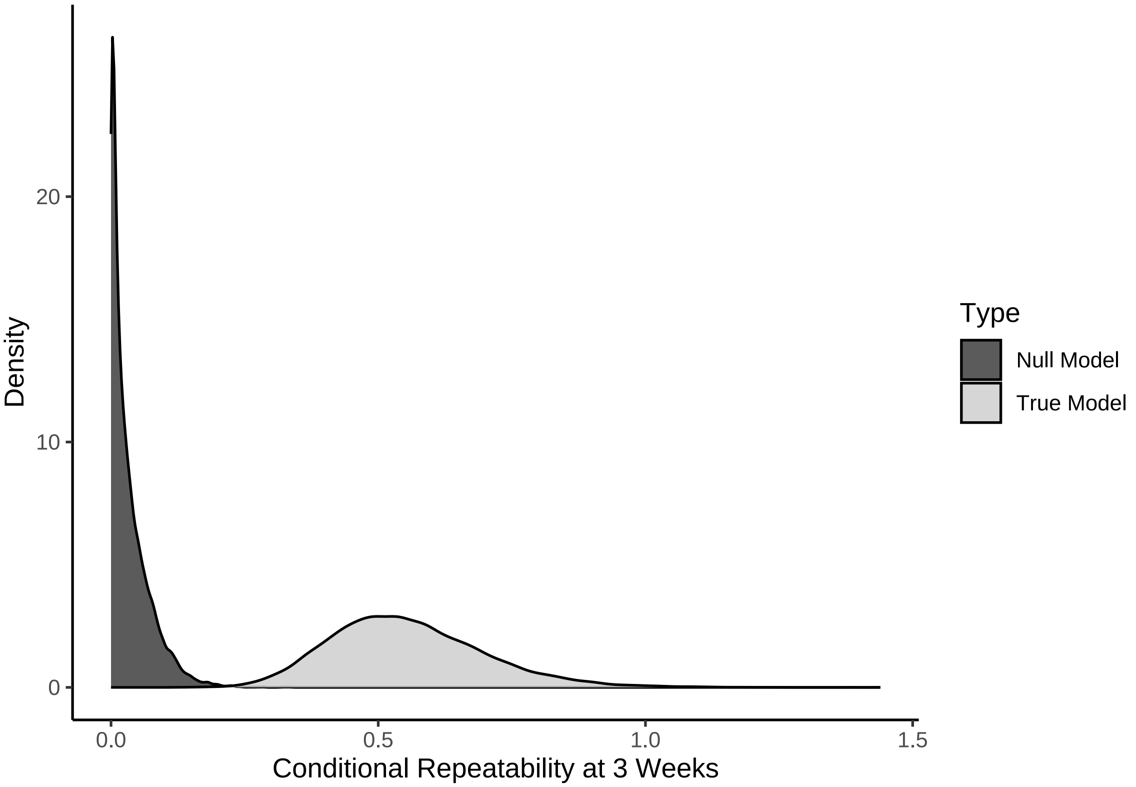


***Figure 14:*** *Density of slope repeatabilites calculated from coefficients of a Bayesian, linear mixed effects model predicting relative resting metabolism (fold resting metabolism at thermoneutrality) across ambient temperature (10°C - 30°C) in three week old Japanese quail. Light grey densities represent true repeatability estimates and dark grey densities represent those calculated from models with scrambled individual identities (null model). All repeatabilities are calculated as per Schielzeth and Nakagawa (2022).*

```
ggsave("../plots/traitRepeatability3Weeks_3.pdf",
  repeatPlot,
  dpi = 800, height = 5, width = 7.5
)
showtext_auto(enable = FALSE)
```

Comparisons between true and null conditional repeatabilities are achieved using a one-way hypothesis test via the Savage-Dickey density comparison method. Priors for our null and true repeatabilies are assigned generous beta distributions with $\alpha = 1$ and $\beta = 4$.

```
# Clear effect of identity on metabolic slopes.

require(loo)

u1RepeatabilityTestFrame <- build_hdf(
  vars = list(
    repeatabilityFrame$Slope,
    repeatabilityFrameScrambled$Slope
  ),
  priors = list(
    rbeta(nrow(repeatabilityFrame), 1, 4),
    rbeta(nrow(repeatabilityFrameScrambled), 1, 4)
  ),
  names = c("True", "Null")
)

u1RepeatabilityTest <- hypothesis_df("True > Null",
```

```
                                    u1RepeatabilityTestFrame,
                                    class = "b", alpha = 0.05
)

# Relabelling for clearer interpretation of plots

u1RepeatabilityTest$hypothesis$Hypothesis <- paste0(
  "Slope: ",
  u1RepeatabilityTest$hypothesis$Hypothesis
)

# Viewing outcome and summarising results

repeatTestPlot <-
    data.frame(
      "values" = c(
        u1RepeatabilityTest$samples$H1,
        u1RepeatabilityTest$prior_samples$H1
      ),
      "Type" = c(
        rep("Posterior", length(u1RepeatabilityTest$samples$H1)),
        rep("Prior", length(u1RepeatabilityTest$prior_samples$H1))
      ),
      "Hypothesis" = "Slope Repeatability > Null"
    ) %>%
  ggplot(aes(x = values, fill = Type)) +
  facet_wrap(~Hypothesis) +
  geom_density(colour = "black", alpha = 0.4) +
  geom_vline(xintercept = 0, colour = "black", linetype = "longdash") +
  scale_fill_manual(values = c("black", "grey80")) +
  xlab("True - Null Repeatability") +
  ylab("Density") +
  theme_classic()

repeatTestPlot
```

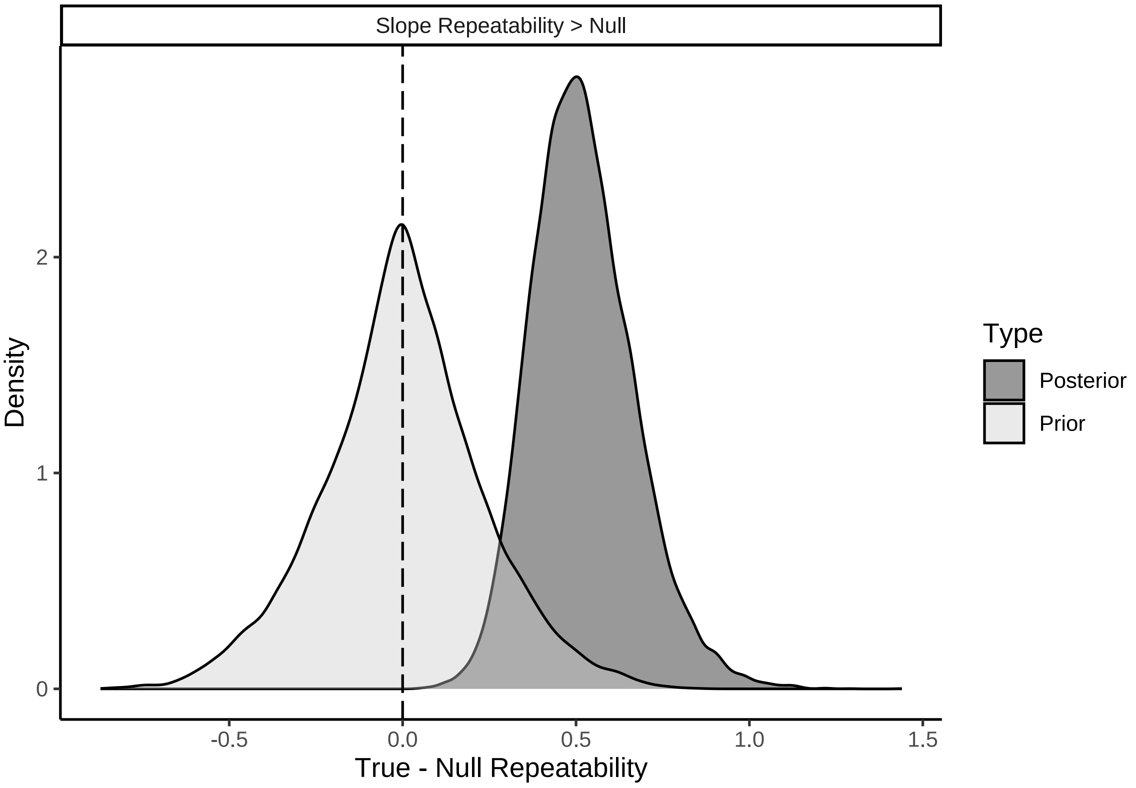


***Figure 15:*** *Differences between true and null repeatability estimates for individual-level slopes of relative resting metabolism (fold metabolism at thermoneutrality) by ambient temperature (°C; between 10-30°C) curves in three week old Japanese quail.*

```
caption <- paste0("Results of a non-linear hypothesis ",
                  "tests comparing true and null repeatabilities ",
                  "of individual-level slopes from relative ",
                  "resting metabolism (fold metabolism at ",
                  "thermoneutrality) by ambient temperature ",
                  "(10-30°C) curves. ",
                  "Curves are derived from three week old ",
                  "Japanese quail. Posterior probabilities are ",
                  "calculated using the Savage-Dickey ",
                  "density ratio method."
)

repeatabilityTest3Weeks <- u1RepeatabilityTest$hypothesis %>%
  mutate(Hypothesis = c(
    "True Slope Repeatability > Null"
  )) %>%
  select(-c(Evid.Ratio, Star)) %>%
  mutate(Estimate = round(Estimate, digits = 4),
         "Est.Error" = round(Est.Error, digits = 4),
         "CI.Lower" = round(CI.Lower, digits = 4),
         "CI.Upper" = round(CI.Upper, digits = 4)) %>%
  rename(
    "Difference Between True and Null Repeatabilities" = Estimate,
    "Standard Error" = Est.Error,
    `2.5 % CI` = "CI.Lower",
    `97.5 % CI` = "CI.Upper",
    "Posterior Probability" = Post.Prob
  ) %>%
  kbl(.,
    longtable = T, booktabs = T, format = "latex",
    caption = caption
```

```
  ) %>%
  column_spec(column = c(1:2), width = "2.5cm")  %>%
  column_spec(column = c(3:10), width = "2cm")  %>%
  kable_styling(latex_options = "striped")

repeatabilityTest3Weeks
```

***Table 4:*** *Results of a non-linear hypothesis tests comparing true and null repeatabilities of individual-level slopes from relative resting metabolism (fold metabolism at thermoneutrality) by ambient temperature (10-30°C) curves. Curves are derived from three week old Japanese quail. Posterior probabilities are calculated using the Savage-Dickey density ratio method.*

| Hypothesis | Difference Between True and Null Repeatabilities | Standard Error | 2.5 % CI | 97.5 % CI | Posterior Probability |
|---|---|---|---|---|---|
| True Slope Repeatability > Null | 0.5206 | 0.1475 | 0.3003 | 0.7798 | 1 |

```
#save_kable(repeatabilityTest3Weeks, "../tables/repeatabilityTest3Weeks.html")

# Clear repeatability.
```

Models including only eight week old individuals are next constructed. Again, only individuals for whom resting metabolism was measured at thermoneutrality (30°C) and at least one ambient temperature below were used here.

Using data derived from other adult Japanese quail in our sample population, we have estimated the lower critical temperature of our quail strain to be 26°C (see supplemental document entitled "Estimating Lower Critical Temperature in a Population of Japanese Quail"). For this reason, slopes of resting metabolism by ambient temperature are not likely to be linear at and below 30°C, potentially biasing estimation of metabolic slopes. To therefore better estimate metabolic slopes in our sample population, we reassigned our metabolism measurements at 30°C to 26°C, since: (1) metabolism at these temperatures is expect to be constant given that both temperatures fall within thermoneutrality, and (2) doing so will linearise metabolism by ambient temperature measurements as expected below thermoneutrality. Below, we visually test whether linearisation does occur with ambient temperature reassignment.

```
## Repeating model for 8 week measurements while retaining
# equivalent priors

birdVector <- merge(
  all %>%
    drop_na(VO2) %>%
    filter(week == 8 & Ta == 30) %>%
    select(ring) %>%
    distinct(),
  all %>%
    drop_na(VO2) %>%
    filter(week == 8 & Ta %in% c(10, 20)) %>%
    select(ring) %>%
    distinct(),
  by = "ring", all = FALSE
) %>% pull(ring)

modData <- all %>%
  filter(week == 8 & Ta < 35 & ring %in% birdVector)

vo2ByTaAdjusted <- modData %>%
  mutate(Ta = ifelse(Ta == 30, 26, Ta)) %>%
  ggplot(aes(x = Ta, y = VO2)) +
  geom_point(
```

```
    pch = 21, size = 1.5, alpha = 0.5,
    colour = "black", fill = "grey50",
    position = position_jitter(width = 0.2)
  ) +
  stat_summary(geom = "errorbar", fun.data = "mean_se",
               width = 0.5, colour = "black", size = 1) +
  stat_summary(
    geom = "point", fun = "mean", size = 4, pch = 21, colour = "black",
    fill = "grey50"
  ) +
  geom_smooth(method = "lm", colour = "black",
              linetype = "dashed", se = FALSE) +
  xlab("Ambient Temperature (°C)") +
  ylab(bquote(Metabolism ~ at ~ 8 ~ weeks ~ (mL ~ O[2] / min))) +
  theme_classic() +
  theme(axis.title = element_text(family = "Noto Sans", size = 12))

showtext_auto()
vo2ByTaAdjusted
```

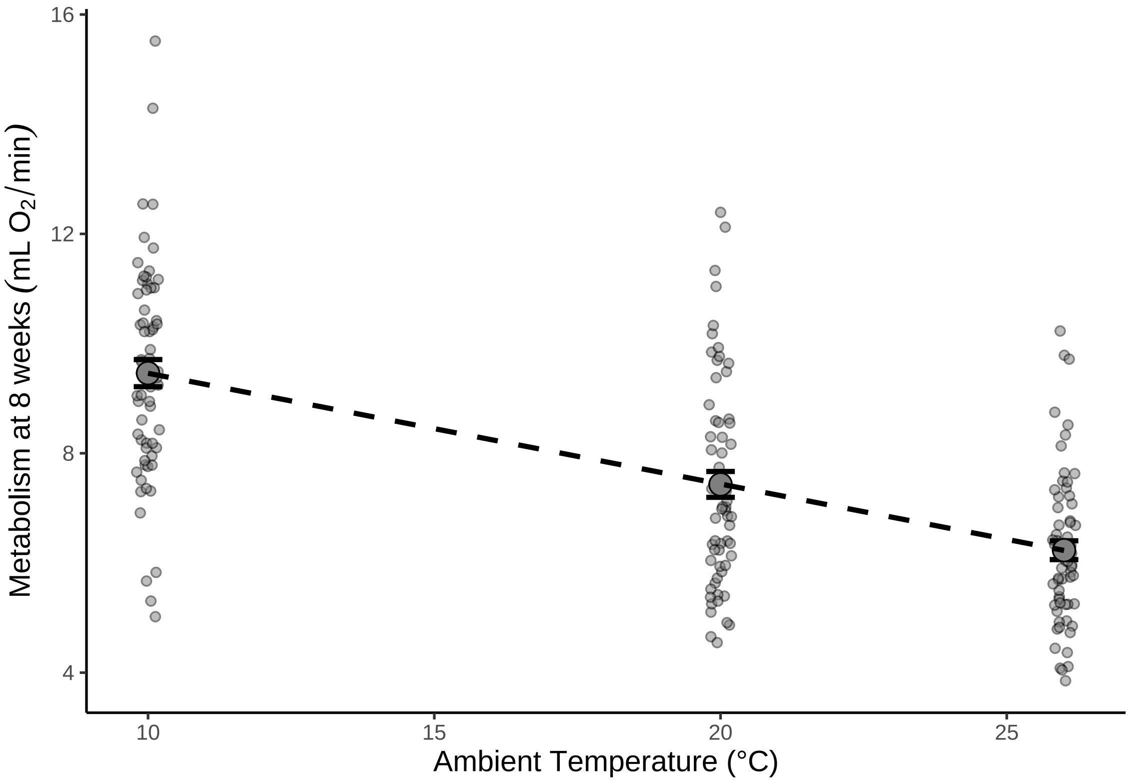


***Figure 16:*** *Relationship between resting metabolism and ambient temperature in eight week old Japanese quail. Large dots represent means across quail and small dots represent individual measurements. Errorbars around large dots represent standard errors and the dotted line represents the linear, line of best fit as estimated by the R package ggplot2 (Wickham, 2011). Metabolism measurements at 26°C were collected at 30°C and expected to be constant at 26°C.*

```
showtext_auto(enable = FALSE)
```

Clearly, the correlation between resting metabolism and ambient temperature is linearised after reassigning measurements at 30°C to 26°C. We proceed by constructing a model of fold-metabolism by ambient temperature using these data as previously described. Priors remain the same as those used for our model addressing metabolic trends in three week old Japanese quail. However, despite this constancy of priors, we still begin by checking their suitability against data obtained from mature quail.

```
# Running prior predictive check.

slopes8WeeksPPCheck <- brm(
  data = modData %>%
    select(ring, Ta, VO2) %>%
    pivot_wider(id_cols = "ring", names_from = Ta, values_from = VO2) %>%
    mutate(`10` = `10`/`30`,
           `20` = `20`/`30`,
           `30` = 1) %>%
    rename(`26` = `30`) %>%
    pivot_longer(-c("ring"), names_to = "Ta", values_to = "VO2") %>%
    mutate(Ta = as.integer(Ta)) %>%
    mutate(Ta = Ta - 26) %>%
    filter(Ta < 0) %>%
    mutate("discreteTa" = ifelse(Ta == -16, "A", "B")),
  family = "gaussian",
  bf(VO2 ~ 1 + B,
     B ~ 0 + Ta + (0 + Ta | ring),
     sigma ~ discreteTa,
     nl = TRUE),
  prior = c(
    set_prior("skew_normal(-0.018, 0.02, -2.5)", nlpar = "B",
              class = "b", coef = "Ta"),
    set_prior("exponential(2.5)", nlpar = "B", class = "sd",
              coef = "Ta", group = "ring"),
    set_prior("skew_normal(-0.2, 1, -5)", dpar = "sigma",
              class = "Intercept"),
    set_prior("normal(0.25, 0.25)", dpar = "sigma",
              class = "b")
  ),
  iter = 50000, warmup = 10000, thin = 10,
  chains = 4, cores = 4,
  control = list(adapt_delta = .96),
  silent = TRUE, refresh = 0,
  sample_prior = "only",
  file = "./models/eightWeekSlopesPPCheck.Rds"
)

pp_check2(slopes8WeeksPPCheck,
          xlab = TeX('$Relative~Resting~Metabolism$')) +
  xlim(c(-10, 10))
```

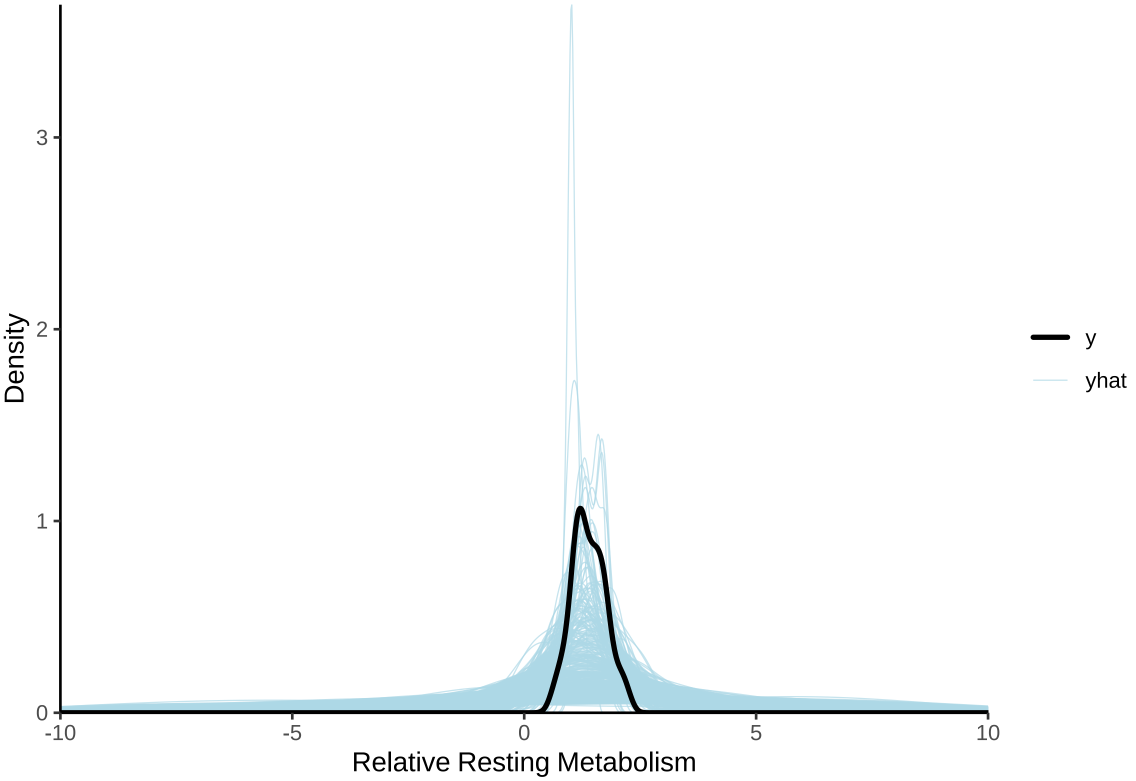


***Figure 17:*** *Prior predictive check for a Bayesian linear, mixed effects model predicting relative resting metabolism (fold metabolism at thermoneutrality) of eight week old Japanese quail by mean-centred ambient temperature (raw ambient temperatures ranging from 10°C - 26°C). Light blue lines represent densities of resting metabolism values as predicted by model priors alone. The dark blue line represent the true density of resting metabolism values. Clear overlap between the dark blue and light blue lines indicates that priors are suitable.*

Again, centrality of relative metabolism measures is well predicted. We proceed with the selected priors.

```
slopes8Weeks <- brm(
  data = modData %>%
    select(ring, Ta, VO2) %>%
    pivot_wider(id_cols = "ring", names_from = Ta, values_from = VO2) %>%
    mutate(`10` = `10`/`30`,
           `20` = `20`/`30`,
           `30` = 1) %>%
    rename(`26` = `30`) %>%
    pivot_longer(-c("ring"), names_to = "Ta", values_to = "VO2") %>%
    mutate(Ta = as.integer(Ta)) %>%
    mutate(Ta = Ta - 26) %>%
    filter(Ta < 0) %>%
    mutate("discreteTa" = ifelse(Ta == -16, "A", "B")),
  family = "gaussian",
  bf(VO2 ~ 1 + B,
     B ~ 0 + Ta + (0 + Ta | ring),
     sigma ~ discreteTa,
     nl = TRUE),
  prior = c(
    set_prior("skew_normal(-0.018, 0.02, -2.5)", nlpar = "B",
              class = "b", coef = "Ta"),
    set_prior("exponential(2.5)", nlpar = "B", class = "sd",
              coef = "Ta", group = "ring"),
    set_prior("skew_normal(-0.2, 1, -5)", dpar = "sigma",
```

```
              class = "Intercept"),
    set_prior("normal(0.25, 0.25)", dpar = "sigma",
              class = "b")
  ),
  iter = 50000, warmup = 10000, thin = 10, cores = 4, chains = 4,
  control = list(adapt_delta = .96),
  silent = TRUE, refresh = 0,
  file = "./models/eightWeekSlopes.Rds"
)

# Again, assessing Rubin-Gelman statistic, ratio of effective
# sample sizes to sample sizes, and distribution of posterior predictions.

p1 <- mcmc_neff(neff_ratio(slopes8Weeks), size = 2) +
  theme(legend.position = "none") +
  xlab(
    TeX('$N_{eff}/N~Ratio$')
  )
p2 <- mcmc_rhat(rhat(slopes8Weeks)) +
  theme(legend.position = "none") +
  xlab(
    TeX('$\\hat{R}$')
  )

p3 <- pp_check2(slopes8Weeks,
                xlab = TeX('$Relative Resting~Metabolism$'))
p4 <- ppc_scatter_avg_grouped(
  slopes8Weeks$data %>%
    pull(VO2),
  posterior_predict(slopes8Weeks,
                    ndraws = 100, robust = TRUE),
  group = slopes8Weeks$data %>%
    merge(., modData %>%
            select(ring, pretreatment) %>%
            distinct(),
          by = "ring", all.x = TRUE) %>%
    mutate(pretreatment = ifelse(pretreatment == "cold", "Cold\n(10°C)",
      ifelse(pretreatment == "neutral", "Mild\n(20°C)",
        "Warm\n(30°C)"
      )
    )) %>%
    pull(pretreatment)
) +
  geom_smooth(method = "lm", colour = "black") +
  xlab("Predicted Metabolism\n(Fold from Thermoneutrality)") +
  ylab("True Metabolism\n(Fold from Thermoneutrality") +
  scale_fill_manual(values = "lightblue", guide = "none") +
  theme_classic()

(p1 + p2 + p3) / (p4) + plot_annotation(tag_levels = "A")
```

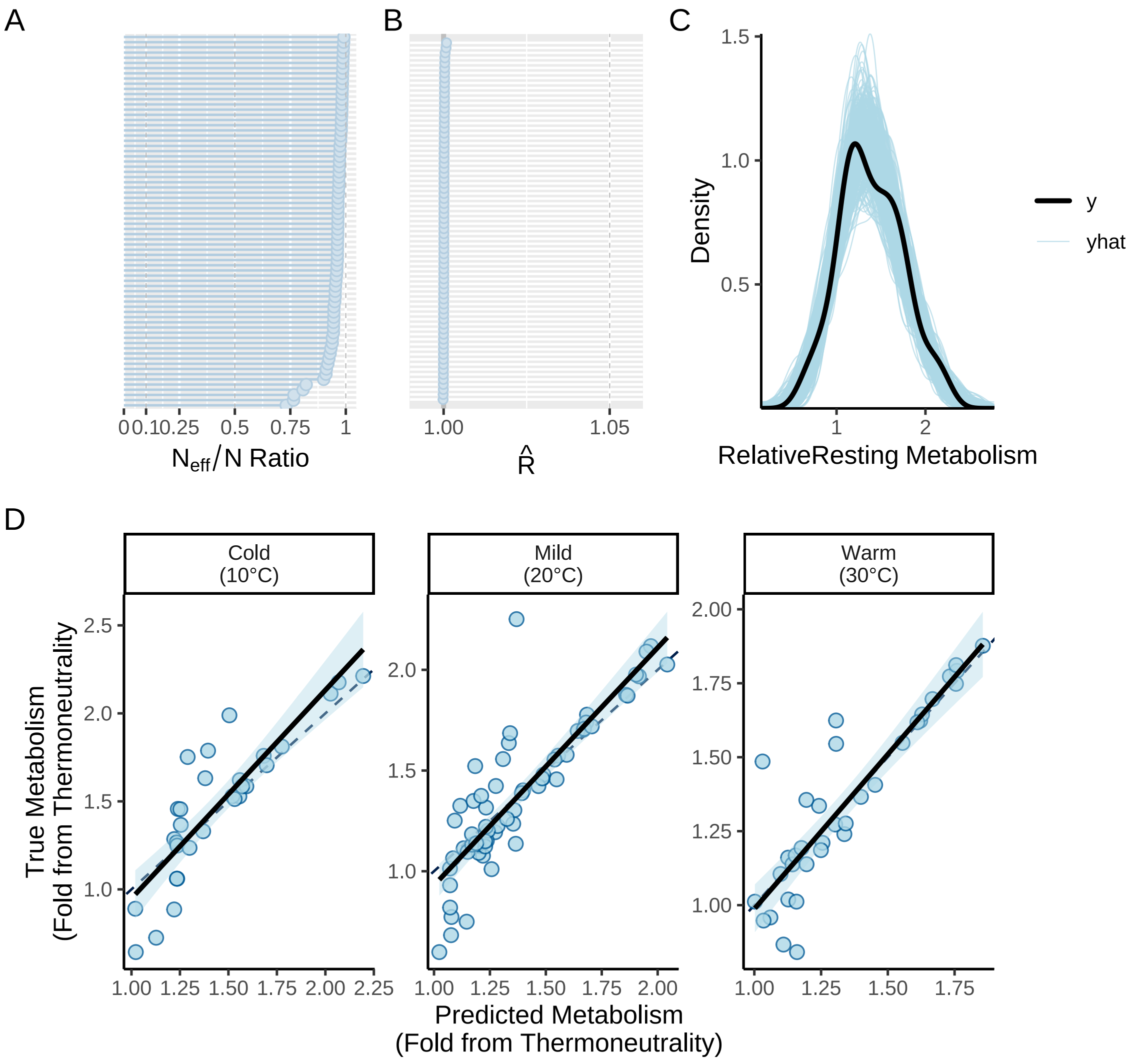


***Figure 18:*** *Validations for a Bayesian mixed effects model predicting relative resting metabolism (fold metabolism at thermoneutrality) of eight week old Japanese quail across 10°C - 26°C. Quail were reared across three different thermal environments: (1) cold (10°C until at least 3 weeks of age), mild (20°C until 8 weeks of age), or warm (30°C until at least 3 weeks of age). Panel A displays the ratio of effective sample sizes to sample sizes for all model parameters. Panel B displays Gelman-Rubin statistics, again, for each model parameter. Panel C displays densities of true (y) and predicted (yhat) values of resting metabolism (mL $O_2$/min). Panel D displays true and predicted resting metabolism (mL $O_2$/min) values across each rearing treatment.*

```
rm(p1, p2, p3, p4)
```

Residuals from this model are next visualised and relative fit (Bayesian $R^2$) evaluated.

```
# No obvious bias by treatment groups. Checking residuals.

g(p1, p2, p3) %=% list(
  ggplot(
    slopes8Weeks$data %>%
      mutate(
        "residuals" =
          residuals(slopes8Weeks,
            type = "pearson",
```

```
          robust = TRUE
        )[, "Estimate"]
      ),
    aes(x = residuals)
  ) +
    geom_density(colour = "black") +
    theme_classic() +
    xlab("Pearson Residuals") +
    ylab("Denstiy"),
  ggplot(
    slopes8Weeks$data %>%
      mutate("residuals" = residuals(slopes8Weeks,
        robust = TRUE
      )[, "Estimate"]),
    aes(x = ring, y = residuals)
  ) +
    geom_boxplot(fill = "lightblue", colour = "black", width = 0.3) +
    geom_point(
      pch = 21, size = 3, fill = "lightblue",
      alpha = 0.5, position = position_jitter(width = 0.2)
    ) +
    xlab("Bird Identity") +
    ylab("Ordinary Residuals") +
    theme_classic() +
    theme(
      axis.text.x = element_blank(),
      axis.ticks.x = element_blank()
    ),
  ggplot(
    slopes8Weeks$data %>%
      mutate(
        "residuals" = residuals(slopes8Weeks,
          type = "pearson",
          robust = TRUE
        )[, "Estimate"],
        "residualSE" = residuals(slopes8Weeks,
          type = "pearson",
          robust = TRUE
        )[, "Est.Error"],
        "fitted" = fitted(slopes8Weeks,
          robust = TRUE
        )[, "Estimate"],
        "fittedSE" = fitted(slopes8Weeks,
          robust = TRUE
        )[, "Est.Error"]
      ),
    aes(x = fitted, y = residuals)
  ) +
    geom_errorbar(
      aes(
        ymin = residuals - residualSE,
        ymax = residuals + residualSE
      ),
      width = 0.1, alpha = 0.5
    ) +
    geom_errorbar(
      aes(
        xmin = fitted - fittedSE,
        xmax = fitted + fittedSE
      ),
      height = 0.5, alpha = 0.5
    ) +
    geom_point(pch = 21, size = 3, fill = "lightblue", alpha = 0.5) +
    theme_classic() +
    xlab("Fitted Values") +
    ylab("Pearson Residuals")
)
```

```
(p1 + p3) / p2 + plot_annotation(tag_levels = "A")
```

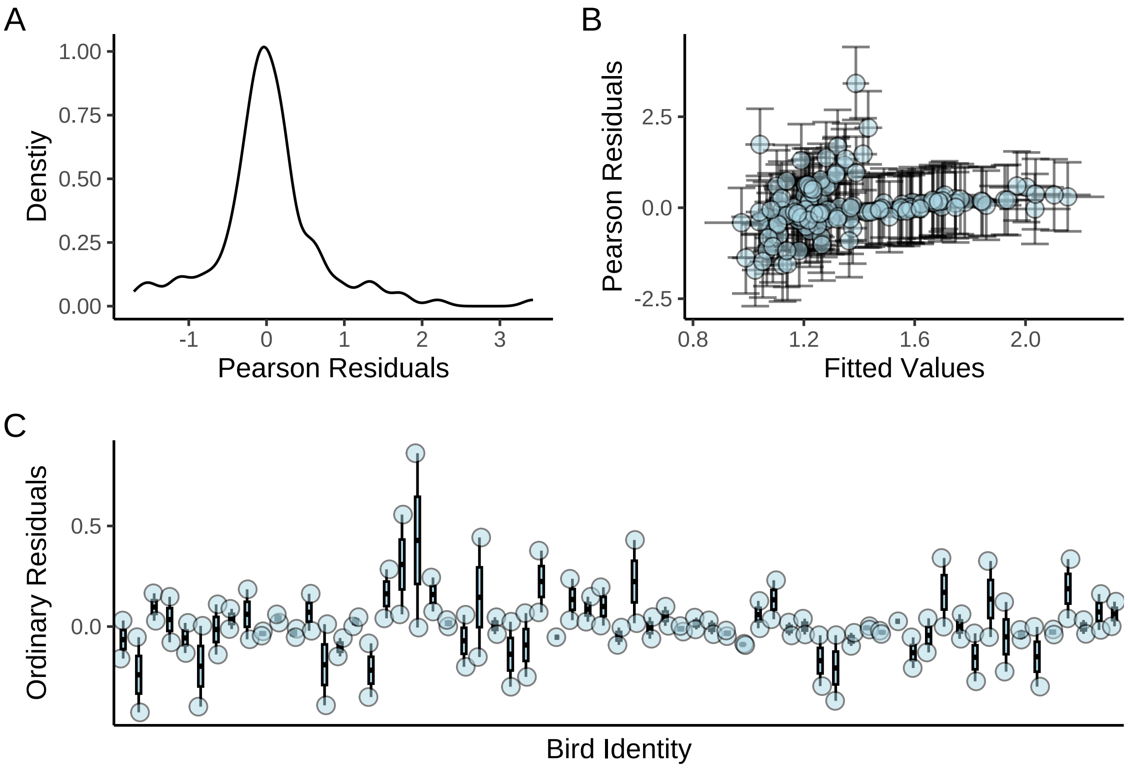


***Figure 19:*** *Ordinary residuals from a Bayesian mixed effects model predicting relative resting metabolism (fold metabolism at thermoneutrality) of eight week old Japanese quail across 10°C - 26°C. Panel A displays residuals against mean-centred ambient temperature values, while panel B displays residuals against fitted values. Panel C displays residuals across individual bird identities.*

```
# Checking R2 values

caption <- paste0("R\\textsuperscript{2} for Bayesian linear, mixed effects ",
                  "model predicting relative resting metabolism (fold ",
                  "metabolism at thermoneutrality) of eight week old Japanese ",
                  "quail by mean-centred ambient temperature ",
                  "(raw ambient temperatures ranging from ",
                  "10°C - ",
                  "26°C)."
)

brms::bayes_R2(slopes8Weeks,
               robust = TRUE, ndraws = 1000) %>%
  as.data.frame() %>%
  remove_rownames() %>%
  mutate(Estimate = round(Estimate, digits = 4),
         Est.Error = round(Est.Error, digits = 4),
         Q2.5 = round(Q2.5, digits = 4),
         Q97.5 = round(Q97.5, digits = 4)
         ) %>%
  rename(
    "R\\textsuperscript{2}" = Estimate,
    "Standard Error" = Est.Error,
    `2.5\\%CI` = "Q2.5", `97.5\\% CI` = "Q97.5"
  ) %>%
  kbl(.,
    longtable = T, booktabs = T, format = "latex",
```

```
    caption = caption, escape = FALSE
  ) %>%
  kable_styling(latex_options = "striped")
```

***Table 5:*** *$R^2$ for Bayesian linear, mixed effects model predicting relative resting metabolism (fold metabolism at thermoneutrality) of eight week old Japanese quail by mean-centred ambient temperature (raw ambient temperatures ranging from 10°C - 26°C).*

| $R^2$ | Standard Error | 2.5%CI | 97.5% CI |
|---|---|---|---|
| 0.6737 | 0.0378 | 0.5545 | 0.7277 |

```
# Again, reasonable
```

Posterior densities per individual are plotted for visual evaluation of spread and skeweness.

```
# Checking densities of metabolic slope estimates

as.data.frame(slopes8Weeks) %>%
  select(starts_with("r_")) %>%
  pivot_longer(everything(), names_to = "ring",
               values_to = "slope") %>%
  mutate("ring" =
           gsub("\\,.*", "",
                gsub(".*\\[", "", ring)
                )
  ) %>%
  mutate(slope = fixef(slopes8Weeks)[1,"Estimate"] +
           slope) %>%
  ggplot(aes(x = slope)) +
  facet_wrap(~ring) +
  geom_density() +
  xlab("Metabolic Slope") +
  ylab("Density") +
  theme_classic() +
  theme(axis.text.x = element_blank(),
        axis.ticks.x = element_blank(),
        axis.text.y = element_blank(),
        axis.ticks.y = element_blank())
```

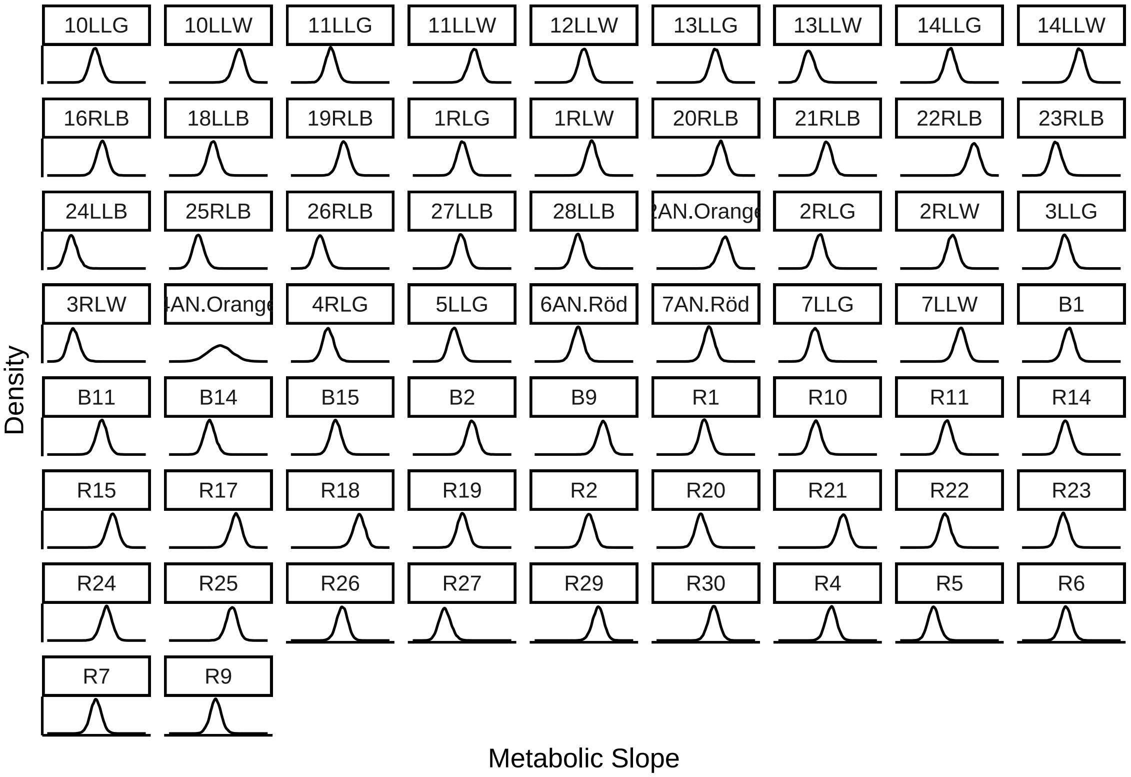


***Figure 20:*** *Posterior densities of metabolic slopes, per individual, from a Bayesian linear, mixed effects model predicting relative resting metabolism (fold resting metabolism at thermoneutrality) of eight week old Japanese quail by ambient temperature (raw ambient temperatures ranging from 10°C - 26°C).*

```
# Again, means appear to be good estimates of central tendency.
```

As done for metabolic slopes during development, we next test whether individuals consistently vary in their metabolic responses to decreasing temperature. This is acheived by quantifying condition repeatability of metabolic slopes, then testing whether this repeatability differs from null expectations (as described above).

```
as.data.frame(slopes8Weeks) %>%
  mutate(
    "Vs" = sd_ring__B_Ta^2 * var(slopes8Weeks$data$Ta) +
      mean(slopes8Weeks$data$Ta)^2 * sd_ring__B_Ta^2,
    "Vf" = b_B_Ta^2 * var(slopes8Weeks$data$Ta)
  ) %>%
  mutate("Slope" = Vs / (Vf + exp(b_sigma_Intercept)^2 +
                          exp(b_sigma_Intercept + b_sigma_discreteTaB)^2)
        ) %>%
  ggplot(aes(x = Slope)) +
  geom_density(alpha = 0.5, colour = "black", fill = "grey80") +
  theme_classic() +
  xlab("Metabolic Slope Repeatability") +
  ylab("Density") +
  theme(legend.position = "none")
```

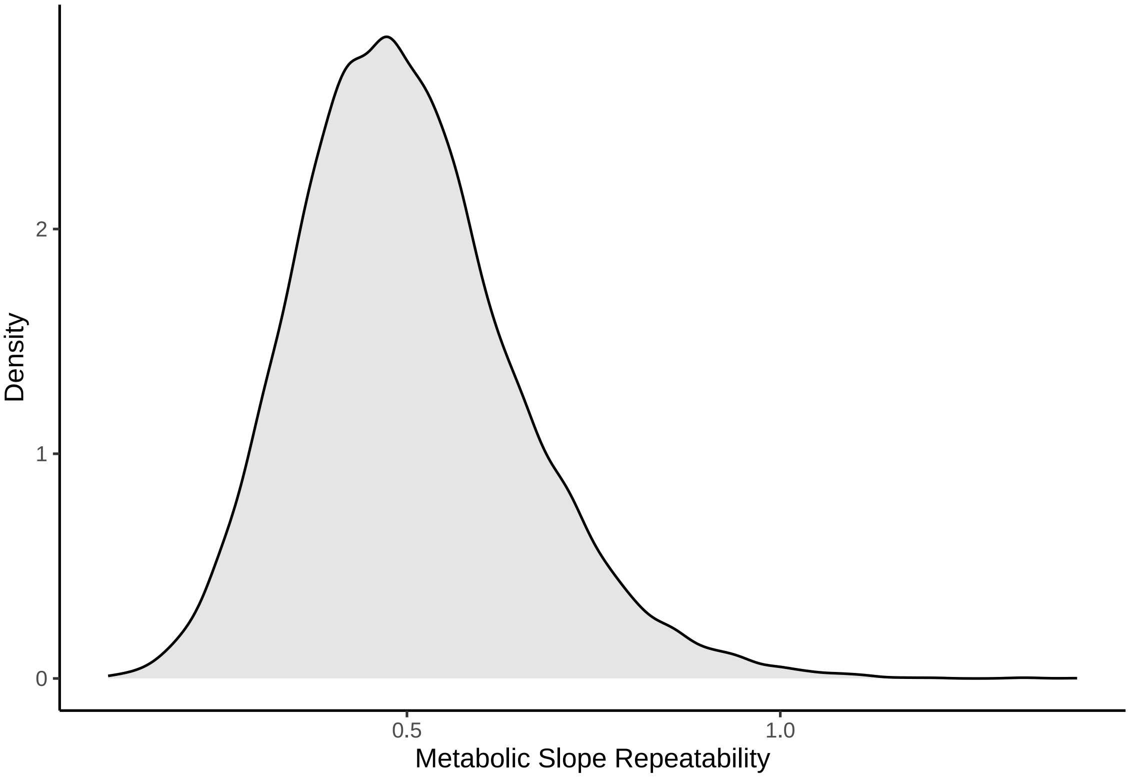


***Figure 21:*** *Repeatability of group-level (here, individual-level) slopes derived from a Bayesian, linear mixed effects model predicting relative resting metabolism (fold metabolism at thermoneutrality) of eight week old Japanese quail across ambient temperature (10°C - 26°C). Trends are estimated per individual and repeatabilities calculated from variability within and among trend parameters, as per Schielzeth and Nakagawa (2022).*

Again, repeatability values appear quite high but are formally tested for biological significance following Tabh et al (2022).

```
repeatabilityFrame = as.data.frame(slopes8Weeks) %>%
  mutate(
    "Vs" = sd_ring__B_Ta^2 * var(slopes8Weeks$data$Ta) +
      mean(slopes8Weeks$data$Ta)^2 * sd_ring__B_Ta^2,
    "Vf" = b_B_Ta^2 * var(slopes8Weeks$data$Ta)
  ) %>%
  mutate("Slope" = Vs / (Vf + exp(b_sigma_Intercept)^2 +
         exp(b_sigma_Intercept + b_sigma_discreteTaB)^2)
         ) %>%
  select(Slope)

caption <- paste("Repeatability of individual-level ",
                 "slopes for the relationship ",
                 "between relative resting metabolism (fold ",
                 "metabolism at thermoneutrality) across ambient ",
                 "temperature (10°C ",
                 "- 26°C) in eight week ",
                 "old Japanese quail. Repeatabilities are ",
                 "calculated using group-level coefficients ",
                 "derived from a Bayesian linear-mixed effects ",
                 "model. Calculations follow Schielzeth and ",
                 "Nakagawa (2022)."
)
```

```r
repeatability8Weeks <-
    repeatabilityFrame %>%
      summarise("Median" = median(Slope),
                "95% CIs" =
                  paste0("[",
                         round(quantile(Slope, 0.025, type = 8),
                               digits = 4),
                         ", ",
                         round(quantile(Slope, 0.975, type = 8),
                               digits = 4),
                         "]")
                ) %>%
  kbl(.,
    longtable = T, booktabs = T, format = "latex",
    caption = caption
  ) %>%
  kable_styling(latex_options = "striped")

repeatability8Weeks
```

***Table 6:*** *Repeatability of individual-level slopes for the relationship between relative resting metabolism (fold metabolism at thermoneutrality) across ambient temperature (10°C - 26°C) in eight week old Japanese quail. Repeatabilities are calculated using group-level coefficients derived from a Bayesian linear-mixed effects model. Calculations follow Schielzeth and Nakagawa (2022).*

| Median | 95% CIs |
|---|---|
| 0.487585 | [0.2503, 0.8395] |

```r
#save_kable(repeatability8Weeks,
# "../tables/VO2Repeatability8Weeks.html")

# And testing formally with scrambled model.

slopes8WeeksScrambled <- brm(
data = modData %>%
    select(ring, Ta, VO2) %>%
    pivot_wider(id_cols = "ring", names_from = Ta, values_from = VO2) %>%
    mutate(`10` = `10`/`30`,
           `20` = `20`/`30`,
           `30` = 1) %>%
    rename(`26` = `30`) %>%
    pivot_longer(-c("ring"), names_to = "Ta", values_to = "VO2") %>%
    mutate(Ta = as.integer(Ta)) %>%
    mutate(Ta = Ta - 26) %>%
    filter(Ta < 0) %>%
    mutate("discreteTa" = ifelse(Ta == -16, "A", "B"),
           "ringScrambled" = sample(ring, nrow(.), replace = FALSE)),
  family = "gaussian",
  bf(VO2 ~ 1 + B,
     B ~ 0 + Ta + (0 + Ta | ringScrambled),
     sigma ~ discreteTa,
     nl = TRUE),
  prior = c(
    set_prior("skew_normal(-0.018, 0.02, -2.5)", nlpar = "B",
              class = "b", coef = "Ta"),
    set_prior("exponential(2.5)", nlpar = "B", class = "sd",
              coef = "Ta", group = "ringScrambled"),
    set_prior("skew_normal(-0.2, 1, -5)", dpar = "sigma",
              class = "Intercept"),
    set_prior("normal(0.25, 0.25)", dpar = "sigma",
              class = "b")
  ),
  iter = 50000, warmup = 10000, thin = 10, cores = 4, chains = 4,
  control = list(adapt_delta = .96),
```

```
  silent = TRUE, refresh = 0,
  file = "./models/eightWeekSlopesScrambled.Rds"
)

repeatabilityFrameScrambled =
  as.data.frame(slopes8WeeksScrambled) %>%
  mutate(
    "Vs" = sd_ringScrambled__B_Ta^2 *
      var(slopes8WeeksScrambled$data$Ta) +
      mean(slopes8WeeksScrambled$data$Ta)^2 *
      sd_ringScrambled__B_Ta^2,
    "Vf" = b_B_Ta^2 * var(slopes8WeeksScrambled$data$Ta)
  ) %>%
  mutate("Slope" = Vs / (Vf + exp(b_sigma_Intercept)^2 +
         exp(b_sigma_Intercept + b_sigma_discreteTaB)^2)
         ) %>%
  select(Slope)

repeatPlot8Weeks <- rbind(
  repeatabilityFrame %>%
    mutate("Type" = "True"),
  repeatabilityFrameScrambled %>%
    mutate("Type" = "Null")
) %>%
  pivot_longer(!c("Type"), names_to = "Par",
               values_to = "Repeatability") %>%
  mutate(Par = "Individual Slope") %>%
  ggplot(aes(x = Repeatability, fill = Type)) +
  #facet_wrap(~Par, scales = "free") +
  geom_density(colour = "black", adjust = 1, alpha = 0.7) +
  scale_fill_manual(
    values = c("grey20", "grey80"),
    labels = c("Null Model", "True Model"),
    name = NULL
  ) +
  ylab("Density") +
  theme_classic()

showtext_auto()
repeatPlot8Weeks
```

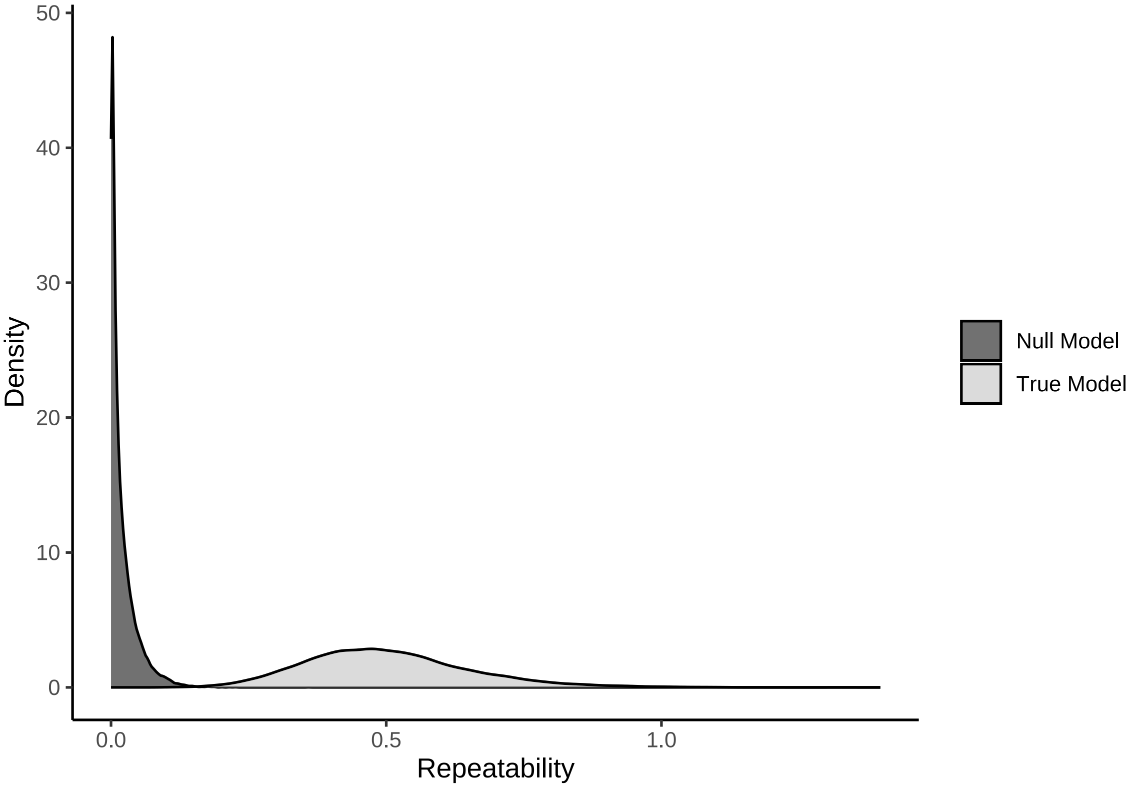


***Figure 22:*** *Density conditional repeatabilities calculated from coefficients of a Bayesian, linear mixed effects model predicting relative resting metabolism (fold metabolism at thermoneutrality) across ambient temperature (10°C - 26°C) in eight week old Japanese quail. Light grey densities represent true repeatability estimates and dark grey densities represent those calculated from models with scrambled individual identities (null model). All repeatabilities are calculated as per Schielzeth and Nakagawa (2022).*

```
ggsave("../plots/traitRepeatability8Weeks_2.pdf",
  repeatPlot8Weeks,
  dpi = 800, height = 5, width = 7.5
)
showtext_auto(enable = FALSE)

# Formally testing possible distinctions.
```

```
u1RepeatabilityTestFrame <- build_hdf(
  vars = list(
    repeatabilityFrame$Slope,
    repeatabilityFrameScrambled$Slope
  ),
  priors = list(
    rbeta(nrow(repeatabilityFrame), 1, 4),
    rbeta(nrow(repeatabilityFrameScrambled), 1, 4)
  ),
  names = c("True", "Null")
)

u1RepeatabilityTest <- hypothesis_df("True > Null",
                                     u1RepeatabilityTestFrame,
  class = "b", alpha = 0.05
)

u1RepeatabilityTest$hypothesis$Hypothesis <- paste0(
  "Slope: ",
  u1RepeatabilityTest$hypothesis$Hypothesis
```

```
)

# Visualising and summarising outcomes

repeatTestPlot <-
    data.frame(
      "values" = c(
        u1RepeatabilityTest$samples$H1,
        u1RepeatabilityTest$prior_samples$H1
      ),
      "Type" = c(
        rep("Posterior", length(u1RepeatabilityTest$samples$H1)),
        rep("Prior", length(u1RepeatabilityTest$prior_samples$H1))
      ),
      "Hypothesis" = "Slope Repeatability > Null"
    ) %>%
  ggplot(aes(x = values, fill = Type)) +
  facet_wrap(~Hypothesis) +
  geom_density(colour = "black", alpha = 0.4) +
  geom_vline(xintercept = 0, colour = "black", linetype = "longdash") +
  scale_fill_manual(values = c("black", "grey80")) +
  xlab("True - Null Repeatability") +
  ylab("Density") +
  theme_classic()

repeatTestPlot
```

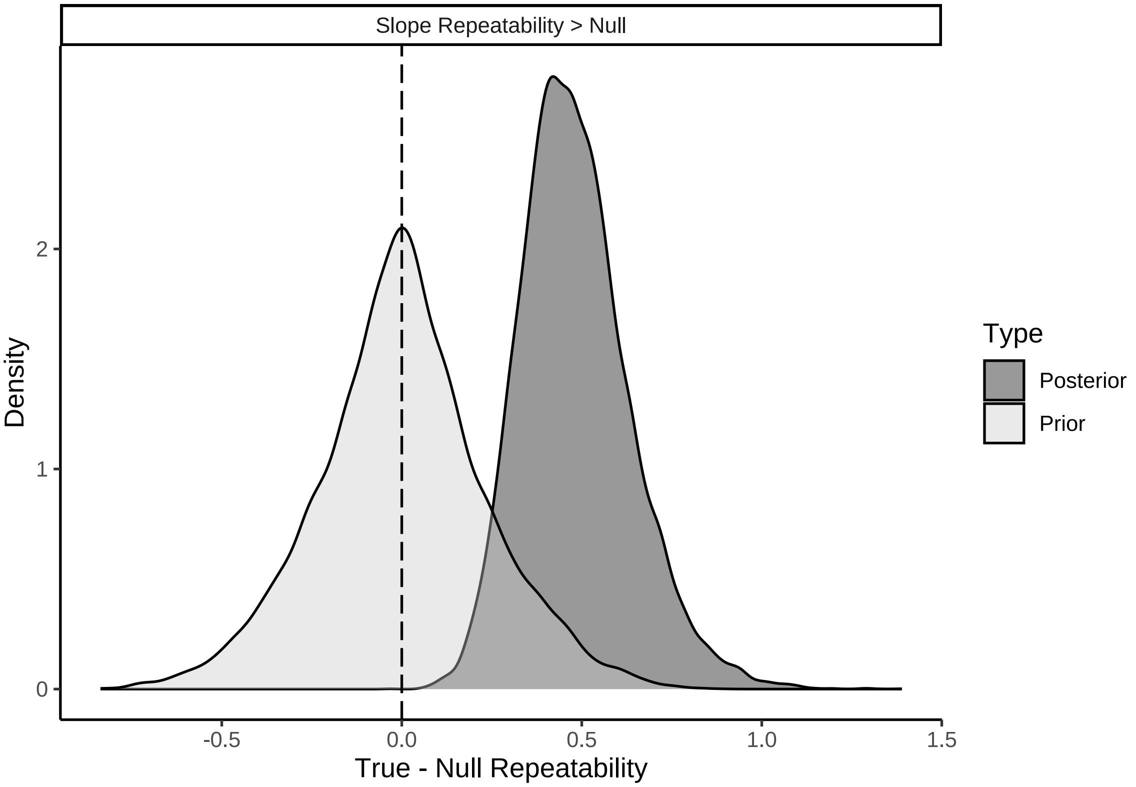


***Figure 23:*** *Differences between true and null repeatability estimates for individual-level slopes of relative resting metabolism (fold metabolism at thermoneutrality) by ambient temperature (°C; between 10-26°C) curves in eight week old Japanese quail.*

```
caption <- paste0("Results of two, non-linear hypothesis ",
                  "tests comparing true and null repeatabilities of ",
                  "individual-level slopes from relative ",
```

```
                  "resting metabolism (fold metabolism at thermoneutrality) ",
                  "by ambient temperature (°C; ",
                  "10-26°C) curves. ",
                  "Curves are derived from eight week old ",
                  "Japanese quail. Posterior probabilities are ",
                  "calculated using the Savage-Dickey density ratio method."
)

repeatabilityTest8Weeks <-
  u1RepeatabilityTest$hypothesis %>%
  mutate(Hypothesis =
    "True Slope Repeatability > Null"
    ) %>%
  select(-c(Evid.Ratio, Star)) %>%
  mutate(Estimate = round(Estimate, digits = 4),
         "Est.Error" = round(Est.Error, digits = 4),
         "CI.Lower" = round(CI.Lower, digits = 4),
         "CI.Upper" = round(CI.Upper, digits = 4)) %>%
  rename(
    "Difference Between True and Null Repeatabilities" = Estimate,
    "Standard Error" = Est.Error,
    `2.5\\% CI` = CI.Lower,
    `97.5\\% CI` = CI.Upper,
    "Posterior Probability" = Post.Prob
  ) %>%
  kbl(.,
    longtable = T, booktabs = T, format = "latex",
    caption = caption, escape = FALSE
  ) %>%
  column_spec(column = c(1:10), width = "2.1cm")  %>%
  kable_styling(latex_options = "striped")

repeatabilityTest8Weeks
```

***Table 7:*** *Results of two, non-linear hypothesis tests comparing true and null repeatabilities of individual-level slopes from relative resting metabolism (fold metabolism at thermoneutrality) by ambient temperature (°C; 10-26°C) curves. Curves are derived from eight week old Japanese quail. Posterior probabilities are calculated using the Savage-Dickey density ratio method.*

| Hypothesis | Difference Between True and Null Repeatabilities | Standard Error | 2.5% CI | 97.5% CI | Posterior Probability |
|---|---|---|---|---|---|
| True Slope Repeatability > Null | 0.483 | 0.1509 | 0.2614 | 0.7506 | 0.9999375 |

```
#save_kable(repeatabilityTest8Weeks,
# "../tables/VO2RepeatabilityTest8Weeks.html")
```

Finally, all metabolic slope values are extracted from models and appended to a data frame for use in subsequent analyses.

```
u1sTrue <- rbind(
  as.data.frame(slopes3Weeks) %>%
    select(c(contains("r_ring") |
             contains("b_B_Ta"))
           ) %>%
    mutate_at(vars(contains("r_ring")), ~ b_B_Ta + .) %>%
    select(-b_B_Ta) %>%
    summarise_all(.funs = median) %>%
    pivot_longer(everything(), names_to = "ring",
                 values_to = "mslope") %>%
```

```
    mutate(
      "ring" = gsub(
        "\\,.*", "",
        gsub(".*\\[", "", ring)
      ),
      "week" = "3"
    ),
  as.data.frame(slopes8Weeks) %>%
    select(c(contains("r_ring") |
             contains("b_B_Ta"))
           ) %>%
    mutate_at(vars(contains("r_ring")), ~ b_B_Ta + .) %>%
    select(-b_B_Ta) %>%
    summarise_all(.funs = median) %>%
    pivot_longer(everything(), names_to = "ring",
                 values_to = "mslope") %>%
    mutate(
      "ring" = gsub(
        "\\,.*", "",
        gsub(".*\\[", "", ring)
      ),
      "week" = "8"
    )
)

u1sRaw <-  rbind(
  as.data.frame(slopes3Weeks) %>%
    select(contains("r_ring")) %>%
    summarise_all(.funs = median) %>%
    pivot_longer(everything(), names_to = "ring",
                 values_to = "rawSlope") %>%
    mutate(
      "ring" = gsub(
        "\\,.*", "",
        gsub(".*\\[", "", ring)
      ),
      "week" = "3"
    ),
    as.data.frame(slopes8Weeks) %>%
    select(contains("r_ring")) %>%
    summarise_all(.funs = median) %>%
    pivot_longer(everything(), names_to = "ring",
                 values_to = "rawSlope") %>%
    mutate(
      "ring" = gsub(
        "\\,.*", "",
        gsub(".*\\[", "", ring)
      ),
      "week" = "8"
    )
)

# Note that ring identities are corrected to bind properly

slopeData <-
  merge(u1sTrue, u1sRaw, by = c("ring", "week")) %>%
  mutate(ring = gsub("\\.", " ", ring)) %>%
  merge(
    all %>%
      filter(Ta < 35 & week %in% c(3, 8)) %>%
      distinct(ring, week, mass, tarsusLengthMean, pretreatment),
    .,
    by = c("ring", "week"), all = TRUE
  )

# Summarising slope values
```

```
caption = paste0("Summary of metabolic slope values estimated for ",
                 "captive reared Japanese quail.")

slopeData %>%
  group_by(week) %>%
  summarise("Mean" = mean(mslope, na.rm = T),
    "SD" = sd(rawSlope, na.rm = T)
  ) %>%
  mutate("CV" = abs((SD/Mean))*100,
    "Parameter" = "Metabolic Slope (RMR/°C)"
  ) %>%
  select(Parameter, "Age (weeks)" = week, Mean, "S.D." = SD, "CV (\\%)" = CV) %>%
  kbl(.,
    longtable = T, booktabs = T, format = "latex",
    caption = caption, escape = FALSE
  ) %>%
  column_spec(column = c(1:10), width = "2.1cm")  %>%
  kable_styling(latex_options = "striped")
```

***Table 8:*** *Summary of metabolic slope values estimated for captive reared Japanese quail.*

| Parameter | Age (weeks) | Mean | S.D. | CV (%) |
|---|---|---|---|---|
| Metabolic Slope (RMR/°C) | 3 | -0.0308816 | 0.0097171 | 31.46575 |
| Metabolic Slope (RMR/°C) | 8 | -0.0346739 | 0.0171759 | 49.53539 |

#### Analysing effects of morphometry on metabolic slopes in the cold

In this subsection, we test whether and how body mass and appendage length affect metabolic slopes of Japanese quail in the cold. These potential effects are contrasted against those of thermal history (here, rearing temperature treatment). Our conceptual model for this subsection is therefore as shown below.

```
data.frame(x = c(-5:5), y = c(-5:5)) %>%
  ggplot(aes(x = x, y = y)) +
  annotate("text", x = 0, y = 5, label = paste0("Developmental, ",
           "Thermal\nEnvironment"), colour = "black") +
  annotate("text", x = -2.5, y = 2.5, label = "Body Mass (g)",
           colour = "black") +
  annotate("text", x = 2.5, y = 2.5, label = "Appendage Length\n(mm)",
           colour = "black") +
  annotate("text", x = 0, y = -0.1,
           label = "Metabolic Slope (fold RMR/°C)",
           colour = "black") +
  geom_segment(
    lineend = "round", linejoin = "round",
    size = 0.3, linetype = "solid", colour = "grey10",
    aes(x = -0.25, y = 4.6, xend = -2.5, yend = 2.75),
    arrow = arrow(length = unit(0.2, "cm"))
  ) +
  geom_segment(
    lineend = "round", linejoin = "round",
    size = 0.3, linetype = "solid", colour = "grey10",
    aes(x = 0.25, y = 4.6, xend = 2.5, yend = 2.75),
    arrow = arrow(length = unit(0.2, "cm"))
  ) +
  geom_segment(
    lineend = "round", linejoin = "round",
    size = 0.3, linetype = "solid", colour = "grey10",
    aes(x = -1.25, y = 2.5, xend = 1, yend = 2.5),
    arrow = arrow(length = unit(0.2, "cm"))
  ) +
  geom_segment(
```

```
    lineend = "round", linejoin = "round",
    size = 0.3, linetype = "dotted", colour = "grey10",
    aes(x = -2.5, y = 2.25, xend = -0.25, yend = 0.25),
    arrow = arrow(length = unit(0.2, "cm"))
  ) +
  geom_segment(
    lineend = "round", linejoin = "round",
    size = 0.3, linetype = "longdash", colour = "grey10",
    aes(x = 2.5, y = 2.25, xend = 0.25, yend = 0.25),
    arrow = arrow(length = unit(0.2, "cm"))
  ) +
  geom_curve(
    lineend = "round",
    size = 0.3, linetype = "solid", colour = "grey10",
    aes(x = -1.75, y = 4.75, xend = -1.75, yend = 0.25),
    arrow = arrow(length = unit(0.2, "cm"))
  ) +
  xlim(c(-6, 6)) +
  ylim(c(-0.5, 6)) +
  theme_void()
```

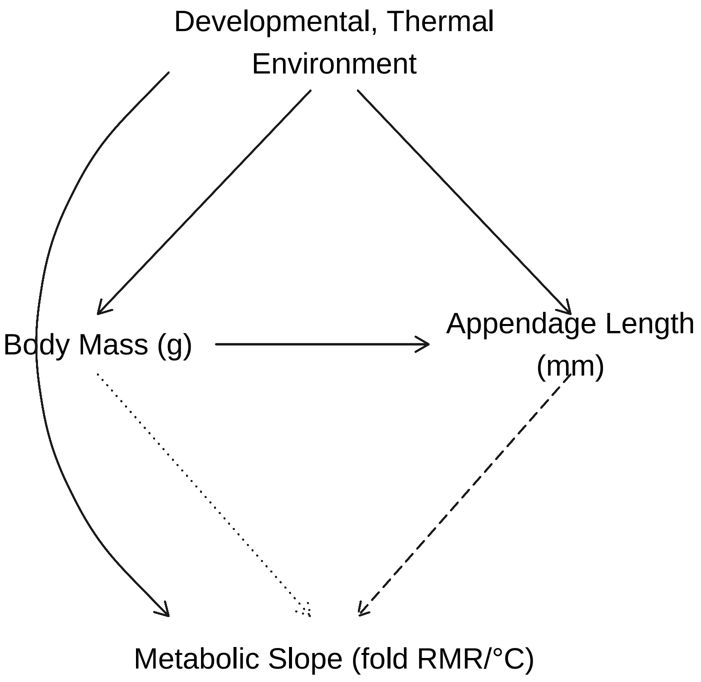


***Figure 24:*** *Flow-chart describing expected effects of developmental, thermal environment on morphology, and subsequently, thermal physiology of Japanese quail. Dashed and dotted lines represent effects predicted under a adaptive hypotheses of Allen's rule and Bergmann's rule respectively.*

Adaptive hypotheses of Bergmann's and Allen's rules predict direct effects of body size (here, body mass) and appendage length on metabolic slopes. However, appendage length is likely to be correlated with body mass, and each are clearly influenced by the thermal environment during rearing (shown in Tabh et al 2023). For these reasons, rather than analysing effects of body mass or appendage length on metabolic slopes alone, we chose to analyse each as part of a comprehensive path analysis, terminating in metabolic slope. This path analysis was composed of three separate models as follows:

$$Body\ Mass_j \sim \beta_{a0} + \beta_{a1} \cdot Cold\ Rearing_j + \beta_{a2} \cdot Warm\ Rearing_j + \mu_{0a} + \epsilon_a$$

$$Tarsus\ Length_j \sim \beta_{b0} + \beta_{b1} \cdot Cold\ Rearing_j + \beta_{b2} \cdot Warm\ Rearing_j + \beta_{b3} \cdot Body\ Mass_j + \mu_{0b} + \epsilon_b$$

and:

$$Metabolic\ Slope_j \sim \beta_{c0} + \beta_{c1} \cdot Cold\ Rearing_j + \beta_{c2} \cdot Warm\ Rearing_j + \beta_{c3} \cdot Body\ Mass_j + \beta_{c4} \cdot Tarsus\ Length_j + \mu_{0c} + \epsilon_c$$

with $j$ representing individual (and thus, observation) identity, “warm rearing” and “cold rearing” representing binomial terms with “0” corresponding to “false” and “1” corresponding to “true”, $\mu_{0a}$ - $\mu_{0c}$ representing group-level intercepts, per response variable, of egg batch, and body mass and tarsus length measures being mean-centred to simplify interpretation of model intercepts ($\beta_0$ terms). Error terms ($\epsilon$) were normally distributed with a mean of zero for each model, and residuals for all models were assumed to be uncorrelated with each response values.

Since we were interested in testing age effects on the functional significance of Bergmann’s and Allen’s rules, we ran our path analysis for data drawn from both three week and eight week old Japanese quail, then compare outcomes thereafter. This approach was chosen to avoid over-parameterising a single model.

**Modelling effects during development (3 weeks of age)**

Below, we begin by constructing and running our path analysis for data from three week old quail. Priors for each model parameter here were moderately informative and as follows:

$$\beta_{a0} \sim \mathcal{N}(0, 5)$$
$$\beta_{a1} \sim \mathcal{N}(0, 15)$$
$$\beta_{a2} \sim \mathcal{N}(0, 15)$$
$$\mu 0a \sim \exp(2.5)$$
$$\epsilon_a \sim \exp(0.15)$$

$$\beta_{b0} \sim \mathcal{N}(0, 2.5)$$
$$\beta_{b1} \sim \mathcal{N}(0, 2.5)$$
$$\beta_{b2} \sim \mathcal{N}(0, 2.5)$$
$$\beta_{b3} \sim \Gamma(1, 5)$$
$$\mu 0b \sim \exp(2)$$
$$\epsilon_b \sim \exp(1)$$

$$\beta_{c0} \sim \mathcal{N}(0, 0.01)$$
$$\beta_{c1} \sim \mathcal{N}(0, 0.025)$$
$$\beta_{c2} \sim \mathcal{N}(0, 0.025)$$
$$\beta_{c3} \sim \mathcal{N}(0, 0.001)$$

$$\beta_{c4} \sim \mathcal{N}(0, 0.0025)$$

$$\mu 0c \sim \exp(25)$$

$$\epsilon_c \sim \exp(10)$$

with distributions for group-level effects of egg batch on body mass and tarsus length derived from growth models described in the supplemental document entitled: "Data Organisation and Analysis of Growth".

Suitability of priors is examined using prior predictive checks (as described above).

Below, we begin with constructing and evaluating models pertaining to data obtained from quail during development.

```
# Testing how appendage length and body mass predict metabolic slopes
# Beginning with prior predictive check

functionModel3WeeksPPCheck <-
  brm(
    data = slopeData %>%
      filter(week == "3") %>%
      mutate(
        mass = mass - mean(mass, na.rm = T),
        tarsus = tarsusLengthMean - mean(tarsusLengthMean, na.rm = T),
        pretreatment = ifelse(pretreatment == "cold", "A",
          ifelse(pretreatment == "neutral", "B", "C")
        )
      ) %>%
      mutate(pretreatment = factor(pretreatment, levels = c("B", "A", "C"))) %>%
      drop_na() %>%
      merge(., data %>%
              select(ring, "batch" = exp) %>%
              distinct(),
            by = "ring", all.x = TRUE),
    family = "gaussian",
    bf(mass ~ pretreatment + (1|batch)) +
      bf(tarsus ~ mass + pretreatment + (1|batch)) +
      bf(rawSlope ~ tarsus + mass + pretreatment + (1|batch)) +
      set_rescor(FALSE),
    prior = c(
      set_prior("normal(0, 5)", class = "Intercept", resp = "mass"),
      set_prior("normal(0, 15)", class = "b",
                coef = "pretreatmentA", resp = "mass"),
      set_prior("normal(0, 15)", class = "b",
                coef = "pretreatmentC", resp = "mass"),
      set_prior("exponential(2.5)", class = "sd",
                group = "batch", resp = "mass"),
      set_prior("exponential(0.15)", class = "sigma", resp = "mass"),
      set_prior("normal(0, 2.5)", class = "Intercept", resp = "tarsus"),
      set_prior("normal(0, 2.5)", class = "b",
                coef = "pretreatmentA", resp = "tarsus"),
      set_prior("normal(0, 2.5)", class = "b",
                coef = "pretreatmentC", resp = "tarsus"),
      set_prior("gamma(1, 5)", class = "b",
                coef = "mass", resp = "tarsus"),
      set_prior("exponential(2)", class = "sd",
                group = "batch", resp = "tarsus"),
      set_prior("exponential(1)", class = "sigma", resp = "tarsus"),
      set_prior("normal(0, 0.01)", class = "Intercept",
                resp = "rawSlope"),
      set_prior("normal(0, 0.025)", class = "b",
                coef = "pretreatmentA", resp = "rawSlope"),
      set_prior("normal(0, 0.025)", class = "b",
                coef = "pretreatmentC", resp = "rawSlope"),
      set_prior("normal(0, 0.001)", class = "b",
                coef = "mass", resp = "rawSlope"),
      set_prior("normal(0, 0.0025)", class = "b",
```

```
                coef = "tarsus", resp = "rawSlope"),
      set_prior("exponential(50)", class = "sd",
                group = "batch", resp = "rawSlope"),
      set_prior("exponential(10)", class = "sigma",
                resp = "rawSlope")
    ),
    iter = 50000, warmup = 10000, cores = 4, chains = 4, thin = 20,
    control = list(adapt_delta = .97, max_treedepth = 14),
    silent = TRUE, refresh = 0,
    sample_prior = "only",
    file = "./models/threeWeekFunctionModelPPCheck.Rds"
  )

(pp_check2(functionModel3WeeksPPCheck,
  resp = "mass",
  xlab = "Relative Body Mass (g)"
) +
  pp_check2(functionModel3WeeksPPCheck,
    resp = "tarsus",
    xlab = "Relative Tarsus Length (mm)"
  )
) /
  pp_check2(functionModel3WeeksPPCheck,
    resp = "rawSlope",
    xlab = "Metabolic Slope (fold RMR/°C)"
  )
```

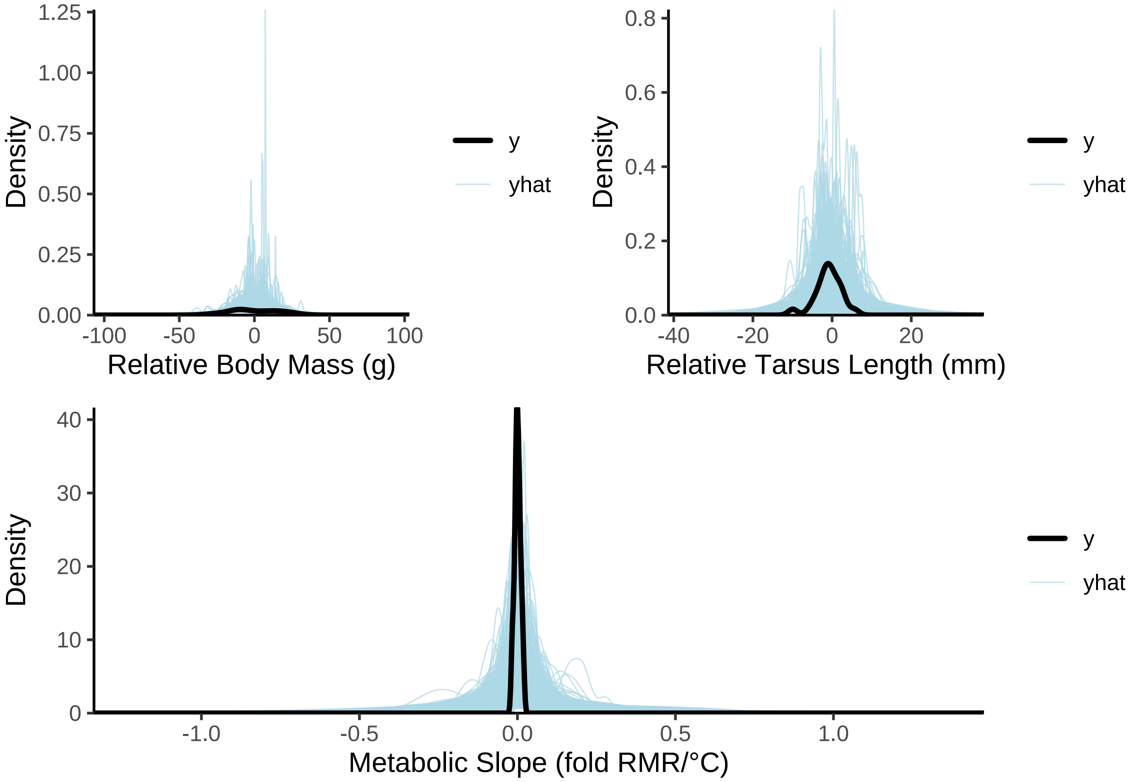


***Figure 25:*** *Prior predictive check for a Bayesian path analysis predicting metabolic slope in the cold (fold metabolism at thermoneutrality/°C) as a direct and indirect function of body mass (g) and tarsus length (mm) in three week old Japanese quail. Light blue lines represent densities of resting metabolism values as predicted by model priors alone. The dark blue line represent the true density of resting metabolism values. Clear overlap between the dark blue and light blue lines indicates that priors are suitable.*

Priors are clearly suitable and broad. We therefore continue to full model construction.

```
functionModel3Weeks <-
  brm(
    data = slopeData %>%
      filter(week == "3") %>%
      mutate(
        mass = mass - mean(mass, na.rm = T),
        tarsus = tarsusLengthMean - mean(tarsusLengthMean, na.rm = T),
        pretreatment = ifelse(pretreatment == "cold", "A",
          ifelse(pretreatment == "neutral", "B", "C")
        )
      ) %>%
      mutate(pretreatment = factor(pretreatment, levels = c("B", "A", "C"))) %>%
      drop_na() %>%
      merge(., data %>%
              select(ring, "batch" = exp) %>%
              distinct(),
            by = "ring", all.x = TRUE),
    family = "gaussian",
    bf(mass ~ pretreatment + (1|batch)) +
      bf(tarsus ~ mass + pretreatment + (1|batch)) +
      bf(rawSlope ~ tarsus + mass + pretreatment + (1|batch)) +
      set_rescor(FALSE),
    prior = c(
      set_prior("normal(0, 5)", class = "Intercept", resp = "mass"),
      set_prior("normal(0, 15)", class = "b",
                coef = "pretreatmentA", resp = "mass"),
      set_prior("normal(0, 15)", class = "b",
                coef = "pretreatmentC", resp = "mass"),
      set_prior("exponential(2.5)", class = "sd",
                group = "batch", resp = "mass"),
      set_prior("exponential(0.15)", class = "sigma", resp = "mass"),
      set_prior("normal(0, 2.5)", class = "Intercept", resp = "tarsus"),
      set_prior("normal(0, 2.5)", class = "b",
                coef = "pretreatmentA", resp = "tarsus"),
      set_prior("normal(0, 2.5)", class = "b",
                coef = "pretreatmentC", resp = "tarsus"),
      set_prior("gamma(1, 5)", class = "b",
                coef = "mass", resp = "tarsus"),
      set_prior("exponential(2)", class = "sd",
                group = "batch", resp = "tarsus"),
      set_prior("exponential(1)", class = "sigma", resp = "tarsus"),
      set_prior("normal(0, 0.01)", class = "Intercept",
                resp = "rawSlope"),
      set_prior("normal(0, 0.025)", class = "b",
                coef = "pretreatmentA", resp = "rawSlope"),
      set_prior("normal(0, 0.025)", class = "b",
                coef = "pretreatmentC", resp = "rawSlope"),
      set_prior("normal(0, 0.001)", class = "b",
                coef = "mass", resp = "rawSlope"),
      set_prior("normal(0, 0.0025)", class = "b",
                coef = "tarsus", resp = "rawSlope"),
      set_prior("exponential(50)", class = "sd",
                group = "batch", resp = "rawSlope"),
      set_prior("exponential(10)", class = "sigma",
                resp = "rawSlope")
    ),
    iter = 50000, warmup = 10000, cores = 4, chains = 4, thin = 20,
    control = list(adapt_delta = .97, max_treedepth = 14),
    silent = TRUE, refresh = 0,
    file = "./models/threeWeekFunctionModel.Rds"
  )

# Some divergent transitions. Checking where in chains they emerge

grab <- paste0(grep("b_|sd_|sigma_",
                    get_variables(functionModel3Weeks), value = TRUE),
```

```
  collapse = "|"
)

posteriorDraws <- as.array(functionModel3Weeks)
divergences <- nuts_params(functionModel3Weeks) %>%
  filter(Parameter == "divergent__" & Value == "1") %>%
  select(".chain" = Chain, ".iteration" = Iteration) %>%
  mutate("divergent" = "TRUE")

spread_draws(functionModel3Weeks, !!sym(grab), regex = TRUE) %>%
  select(-c(.draw)) %>%
  pivot_longer(-c(".chain", ".iteration"),
               names_to = "par", values_to = "values") %>%
  mutate("chain" = paste0("Chain ", .chain)) %>%
  merge(.,
    tribble(
      ~par, ~Par,
      "b_mass_Intercept", " a0",
      "b_mass_pretreatmentA", " a1",
      "b_mass_pretreatmentC", " a2",
      "sd_batch__mass_Intercept", " 0a",
      "sigma_mass", " a",
      "b_tarsus_Intercept", " b0",
      "b_tarsus_pretreatmentA", " b1",
      "b_tarsus_pretreatmentC", " b2",
      "b_tarsus_mass", " b3",
      "sd_batch__tarsus_Intercept", " 0b",
      "sigma_tarsus", " b",
      "b_rawSlope_Intercept", " c0",
      "b_rawSlope_pretreatmentA", " c1",
      "b_rawSlope_pretreatmentC", " c2",
      "b_rawSlope_mass", " c3",
      "b_rawSlope_tarsus", " c4",
      "sd_batch__rawSlope_Intercept", " 0c",
      "sigma_rawSlope", " c"
    ),
    by = "par", all.x = TRUE
  ) %>%
  merge(., divergences, by = c(".chain", ".iteration"), all.x = TRUE) %>%
  mutate(divergent = ifelse(is.na(divergent), "FALSE", divergent)) %>%
  ggplot(aes(x = .iteration, y = values,
             fill = divergent, alpha = divergent)) +
  facet_grid(Par ~ chain, scales = "free") +
  geom_point(pch = 21, colour = "black") +
  scale_fill_manual(values = c("grey90", "darkred"),
                    name = "Divergent") +
  scale_alpha_manual(values = c(0.2, 1), name = "Divergent") +
  xlab("Iteration") +
  ylab("Values") +
  theme_classic() +
  theme(
    axis.text.y = element_blank(),
    axis.ticks.y = element_blank()
  )
```

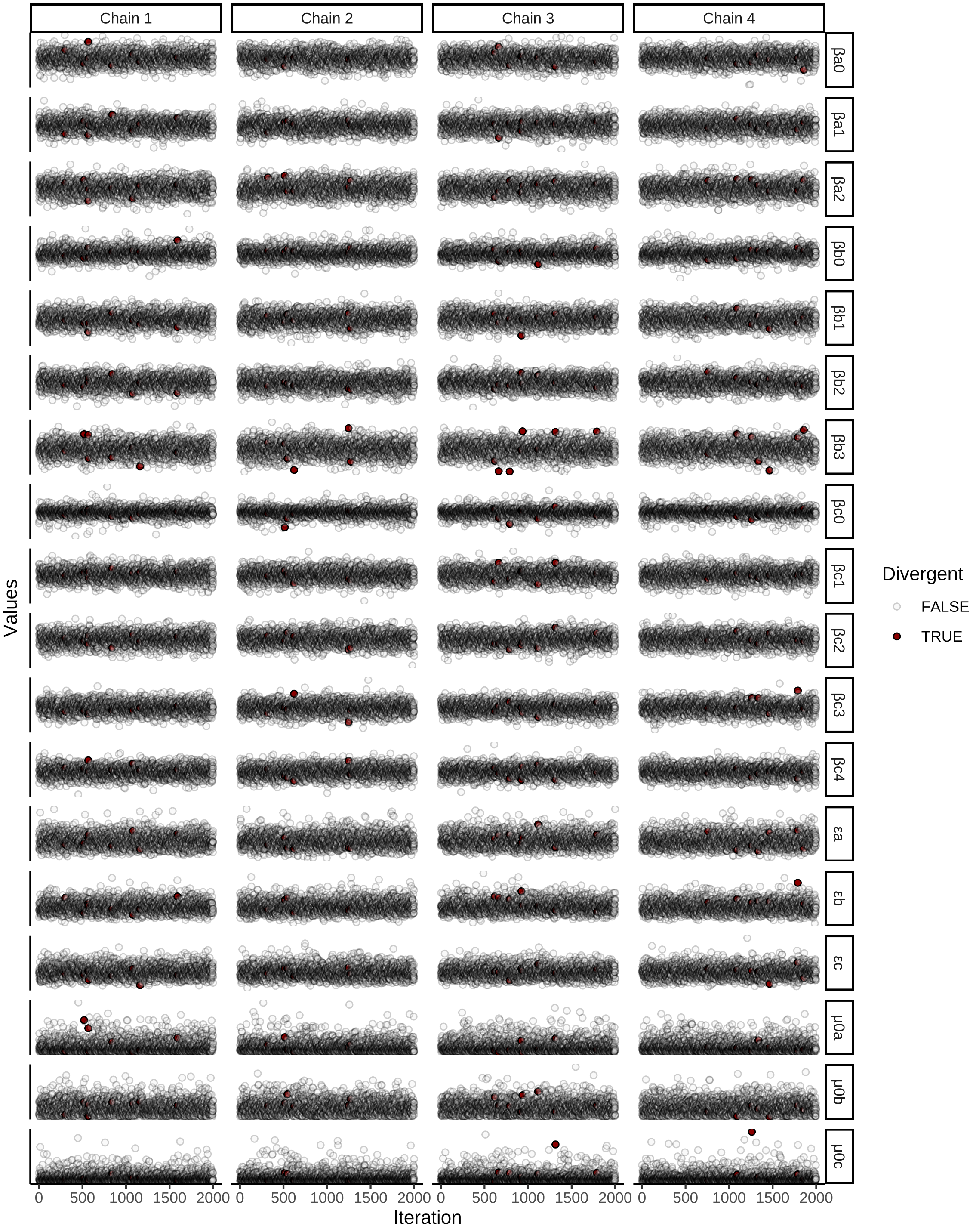


***Figure 26:*** *Posterior draws from Bayesian path analysis predicting metabolic slope in the cold (fold metabolism at thermoneutrality/°C) as a direct and indirect function of body mass (g) and tarsus length (mm) in three week old Japanese quail. Draws are seperated by model parameter and Hamiltonian Monte Carlo (HMC) chains.*

Restricting our prior for the effect of body mass on tarsus length ($\beta_{b3}$) to values above zero is limiting adequate exploration of posterior space. For this reason, we adjust our prior on this parameter as follows:

$$\beta_{b3} \sim S\mathcal{N}(0, 0.25, 5)$$

with $\mathcal{SN}$ representing a skew-normal distribution. Our model is then re-run below and inspected.

```
functionModel3Weeks <-
  brm(
    data = slopeData %>%
      filter(week == "3") %>%
      mutate(
        mass = mass - mean(mass, na.rm = T),
        tarsus = tarsusLengthMean - mean(tarsusLengthMean, na.rm = T),
        pretreatment = ifelse(pretreatment == "cold", "A",
          ifelse(pretreatment == "neutral", "B", "C")
        )
      ) %>%
      mutate(pretreatment = factor(pretreatment, levels = c("B", "A", "C"))) %>%
      drop_na() %>%
      merge(., data %>%
              select(ring, "batch" = exp) %>%
              distinct(),
            by = "ring", all.x = TRUE),
    family = "gaussian",
    bf(mass ~ pretreatment + (1|batch)) +
      bf(tarsus ~ mass + pretreatment + (1|batch)) +
      bf(rawSlope ~ tarsus + mass + pretreatment + (1|batch)) +
      set_rescor(FALSE),
    prior = c(
      set_prior("normal(0, 5)", class = "Intercept", resp = "mass"),
      set_prior("normal(0, 15)", class = "b",
                coef = "pretreatmentA", resp = "mass"),
      set_prior("normal(0, 15)", class = "b",
                coef = "pretreatmentC", resp = "mass"),
      set_prior("exponential(2.5)", class = "sd",
                group = "batch", resp = "mass"),
      set_prior("exponential(0.15)", class = "sigma", resp = "mass"),
      set_prior("normal(0, 2.5)", class = "Intercept", resp = "tarsus"),
      set_prior("normal(0, 2.5)", class = "b",
                coef = "pretreatmentA", resp = "tarsus"),
      set_prior("normal(0, 2.5)", class = "b",
                coef = "pretreatmentC", resp = "tarsus"),
      set_prior("skew_normal(0, 0.25, 5)", class = "b",
                coef = "mass", resp = "tarsus"),
      set_prior("exponential(2)", class = "sd",
                group = "batch", resp = "tarsus"),
      set_prior("exponential(1)", class = "sigma", resp = "tarsus"),
      set_prior("normal(0, 0.01)", class = "Intercept",
                resp = "rawSlope"),
      set_prior("normal(0, 0.025)", class = "b",
                coef = "pretreatmentA", resp = "rawSlope"),
      set_prior("normal(0, 0.025)", class = "b",
                coef = "pretreatmentC", resp = "rawSlope"),
      set_prior("normal(0, 0.001)", class = "b",
                coef = "mass", resp = "rawSlope"),
      set_prior("normal(0, 0.0025)", class = "b",
                coef = "tarsus", resp = "rawSlope"),
      set_prior("exponential(50)", class = "sd",
                group = "batch", resp = "rawSlope"),
      set_prior("exponential(10)", class = "sigma",
                resp = "rawSlope")
    ),
    iter = 50000, warmup = 10000, cores = 4, chains = 4, thin = 20,
    control = list(adapt_delta = .97, max_treedepth = 14),
    silent = TRUE, refresh = 0,
    file = "./models/threeWeekFunctionModelRevised.Rds"
  )

# Divergences dramatically reduced.
# Checking Rhat values and Neff/N ratios.
```

```
chainCheck(functionModel3Weeks)
```

```
## Rhat range: 1 - 1.001
## Neff/N range: 0.895 - 1.007
```

```
# Good. And R2 values?

caption <- paste0("R\\textsuperscript{2} for Bayesian ",
                  "path analysis predicting metabolic slopes ",
                  "(fold metabolism at thermoneutrality/°C) ",
                  "of three week old Japanese quail as a function ",
                  "of body mass (g) and tarsus length (mm). ",
                  "Metabolic slopes are measured below thermoneutrality ",
                  "(<30°C)."
)

brms::bayes_R2(functionModel3Weeks, ndraws = 1000,
               robust = TRUE) %>%
  as.data.frame() %>%
  rownames_to_column("var") %>%
  merge(., tribble(
    ~var, ~Var,
    "R2mass", "Body Mass (g)",
    "R2tarsus", "Tarsus Length (mm)",
    "R2rawSlope",
    "Metabolic Slope (fold RMR/°C)"
  ), by = c("var")) %>%
  mutate(Estimate = round(Estimate, digits = 4),
         Est.Error = round(Est.Error, digits = 4),
         Q2.5 = round(Q2.5, digits = 4),
         Q97.5 = round(Q97.5, digits = 4)
         ) %>%
  select(
    "Response" = Var, "R\\textsuperscript{2}" = Estimate,
    "Standard Error" = Est.Error,
    `2.5\\% CI` = Q2.5, `97.5\\% CI` = Q97.5
  ) %>%
  kbl(.,
    longtable = T, booktabs = T, format = "latex",
    caption = caption, escape = FALSE
  ) %>%
  column_spec(column = c(1:10), width = "2.5cm")  %>%
  kable_styling(latex_options = "striped")
```

***Table 9:*** *$R^2$ for Bayesian path analysis predicting metabolic slopes (fold metabolism at thermoneutrality/°C) of three week old Japanese quail as a function of body mass (g) and tarsus length (mm). Metabolic slopes are measured below thermoneutrality (<30°C).*

| Response | $R^2$ | Standard Error | 2.5% CI | 97.5% CI |
|---|---|---|---|---|
| Body Mass (g) | 0.0731 | 0.0594 | 0.0060 | 0.2247 |
| Metabolic Slope (fold RMR/°C) | 0.2114 | 0.0853 | 0.0668 | 0.3785 |
| Tarsus Length (mm) | 0.3730 | 0.0986 | 0.1752 | 0.5267 |

```
pp_check2(functionModel3Weeks, resp = "rawSlope",
          xlab = "Metabolic Slope (fold RMR/°C)"
          )
```

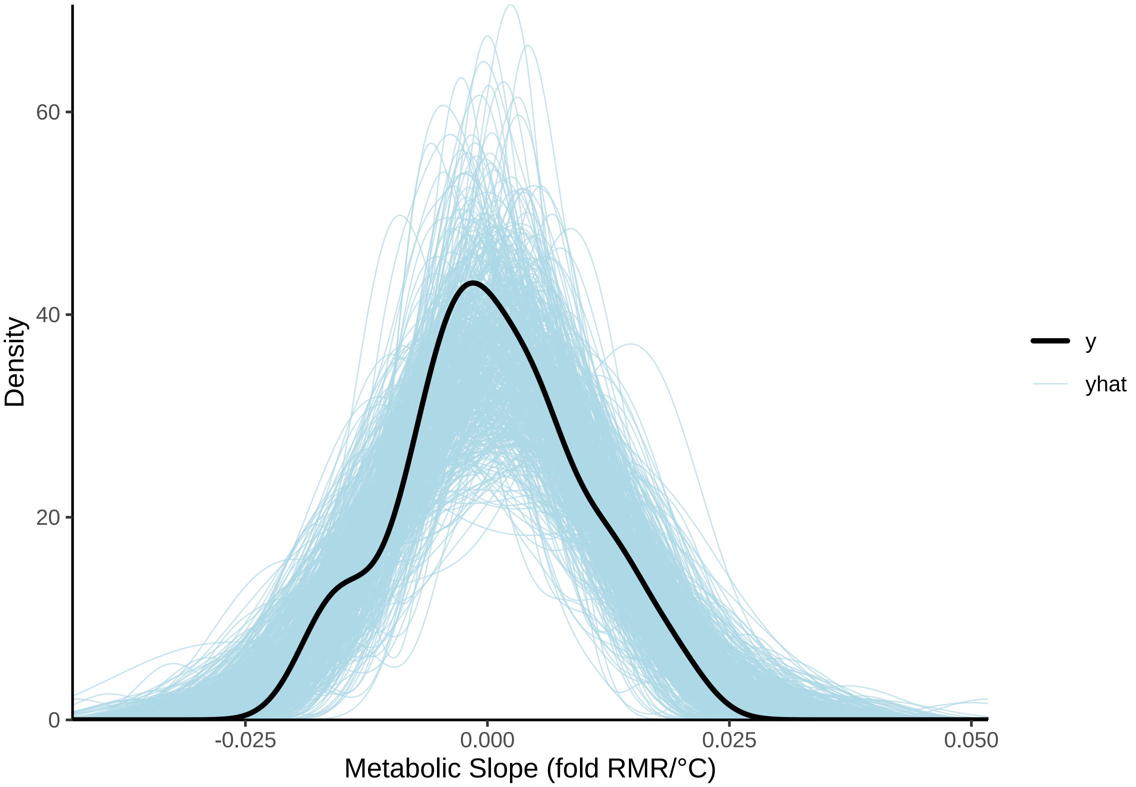


***Figure 27:*** *Densities of posterior predictions from a Bayesian path analysis predicting metabolic slopes (fold metabolism at thermoneutrality/°C) as a direct and indirect function of body mass (g) and tarsus length (mm) in three week old Japanese quail. Prediction are overlayed with true distributions of quail metabolic slopes. Blue lines represent predictions from posterior draws, while the black line represents true, mean-centred resitance density.*

```
# Reasonable overlay. Checking residuals next.
```

Residuals of this model are visualised as before.

```
p1 <- functionModel3Weeks$data %>%
  mutate("Residuals" = residuals(functionModel3Weeks,
                                 resp = "rawSlope",
                                 robust = TRUE)[, "Estimate"]) %>%
  ggplot(aes(x = mass, y = Residuals)) +
  geom_point(size = 2, pch = 21, colour = "black", fill = "grey50") +
  xlab("Relative Body Mass (g)") +
  ylab("Ordinary Residuals") +
  theme_classic()

p2 <- functionModel3Weeks$data %>%
  mutate("Residuals" = residuals(functionModel3Weeks,
                                 resp = "rawSlope",
                                 robust = TRUE)[, "Estimate"]) %>%
  ggplot(aes(x = tarsus, y = Residuals)) +
  geom_point(size = 2, pch = 21, colour = "black", fill = "grey50") +
  xlab("Relative Tarsus Length (mm)") +
  ylab("Ordinary Residuals") +
  theme_classic()

p3 <- functionModel3Weeks$data %>%
  mutate("Residuals" = residuals(functionModel3Weeks,
                                 resp = "rawSlope",
```

```
                                 robust = TRUE)[, "Estimate"]) %>%
  ggplot(aes(x = factor(pretreatment), y = Residuals)) +
  geom_boxplot(width = 0.5) +
  geom_point(size = 2, pch = 21,
             position = position_jitter(width = 0.25),
             colour = "black", fill = "grey50") +
  xlab("Rearing Conditions") +
  ylab("Ordinary Residuals") +
  theme_classic()

p4 <- functionModel3Weeks$data %>%
  mutate(
    "Residuals" = residuals(functionModel3Weeks,
                            resp = "rawSlope",
                                 robust = TRUE)[, "Estimate"],
    "Fitted" = fitted(functionModel3Weeks,
                      resp = "rawSlope",
                                 robust = TRUE)[, "Estimate"]
  ) %>%
  ggplot(aes(x = Fitted, y = Residuals)) +
  geom_point(size = 2, pch = 21, colour = "black", fill = "grey50") +
  xlab("Fitted Values") +
  ylab("Ordinary Residuals") +
  theme_classic()

p5 <- functionModel3Weeks$data %>%
  mutate("Residuals" = residuals(functionModel3Weeks,
                                 resp = "rawSlope",
                                 robust = TRUE)[, "Estimate"]) %>%
  ggplot(aes(sample = Residuals)) +
  stat_qq(colour = "grey50") +
  stat_qq_line() +
  xlab("Theoretical") +
  ylab("Sample") +
  theme_classic()

((p1 + p2) / (p3 + p4)) + p5 + plot_annotation(tag_levels = "A")
```

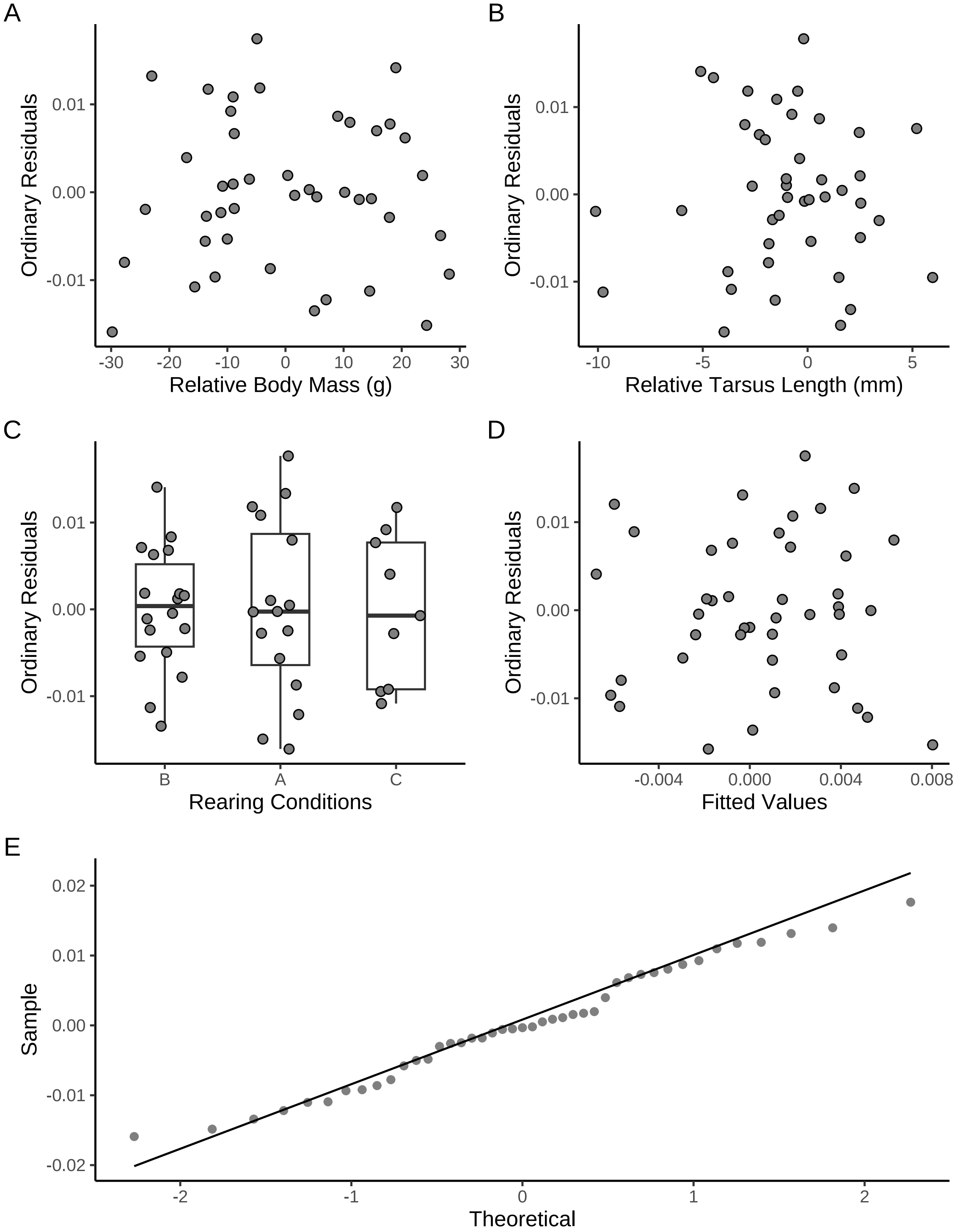


***Figure 28:*** *Ordinary residuals from a Bayesian path analysis predicting metabolic slopes below thermoneutrality (fold metabolism at thermoneutrality/°C) as a direct and indirect function of body mass (g) and tarsus length (mm) in three week old Japanese quail. Panel A displays residuals against mean-centred body mass values, panel B displays residuals against mean-centred tarsus length values, panel C displays residuals across post-hatch rearing conditions, panel D displays residuals against fitted, metabolic slope values, and panel E displays theoretical against sample residuals (qq-plot).*

Next, predicted effects of thermal history (here, rearing treatment) on morphology and metabolic slopes are visualised.

```
p1 <- data.frame("pretreatment" = c("A", "B", "C")) %>%
  mutate(
    "mass" = predict(functionModel3Weeks,
      newdata = ., robust = "TRUE",
      resp = "mass", re_form = NA
    )[, "Estimate"],
    "SE" = predict(functionModel3Weeks,
      newdata = ., robust = "TRUE",
      resp = "mass", re_form = NA
    )[, "Est.Error"]
  ) %>%
  mutate("Mass" = mass +
           mean(subset(slopeData, week == "3")$mass,
                na.rm = T)) %>%
  ggplot(aes(x = pretreatment, y = Mass)) +
  geom_errorbar(aes(x = pretreatment, ymin = Mass - SE,
                    ymax = Mass + SE), colour = "black", width = 0.3) +
  geom_point(size = 5, pch = 21, colour = "black",
             aes(x = pretreatment, fill = factor(pretreatment))) +
  geom_line(linetype = "dashed", colour = "black") +
  geom_point(
    size = 2, pch = 21, colour = "black",
    data = subset(slopeData, week == "3") %>%
      mutate("pretreatment" = ifelse(pretreatment == "cold", "A",
        ifelse(pretreatment == "neutral", "B", "C")
      )),
    aes(x = pretreatment, y = mass, fill = factor(pretreatment)),
    position = position_jitter(width = 0.3)
  ) +
  scale_fill_manual(values = c("#7BB4E3", "black", "#CD5C5C")) +
  scale_x_discrete(breaks = c("A", "B", "C"),
                   labels = c("Cold\n(10°C)",
                              "Mild\n(20°C)",
                              "Warm\n(30°C)")) +
  theme_classic() +
  theme(legend.position = "none",
        axis.text = element_text(family = "Noto Sans"),
        axis.title = element_text(family = "Noto Sans")) +
  xlab("Rearing Conditions") +
  ylab("Mass (g)")

p2 <- data.frame(
  "pretreatment" = c("A", "B", "C"),
  "mass" = 0
) %>%
  mutate(
    "tarsus" = predict(functionModel3Weeks,
      newdata = ., robust = TRUE,
      resp = "tarsus", re_form = NA
    )[, "Estimate"],
    "SE" = predict(functionModel3Weeks,
      newdata = ., robust = TRUE,
      resp = "tarsus", re_form = NA
    )[, "Est.Error"]
  ) %>%
  mutate("Tarsus" = tarsus +
           mean(subset(slopeData, week == "3")$tarsusLengthMean,
                na.rm = T)) %>%
  ggplot(aes(x = pretreatment, y = Tarsus)) +
  geom_errorbar(aes(x = pretreatment, ymin = Tarsus - SE,
                    ymax = Tarsus + SE), colour = "black", width = 0.3) +
  geom_point(size = 5, pch = 21, colour = "black",
             aes(x = pretreatment, fill = factor(pretreatment))) +
  geom_line(linetype = "dashed", colour = "black") +
```

```r
  geom_point(
    size = 2, pch = 21, colour = "black",
    data = subset(slopeData, week == "3") %>%
      mutate("pretreatment" = ifelse(pretreatment == "cold", "A",
        ifelse(pretreatment == "neutral", "B", "C")
      )),
    aes(x = pretreatment, y = tarsusLengthMean,
        fill = factor(pretreatment)),
    position = position_jitter(width = 0.3)
  ) +
  scale_fill_manual(values = c("#7BB4E3", "black", "#CD5C5C")) +
  scale_x_discrete(breaks = c("A", "B", "C"),
                   labels = c("Cold\n(10°C)",
                              "Mild\n(20°C)",
                              "Warm\n(30°C)")
                   ) +
  theme_classic() +
  theme(legend.position = "none",
        axis.text = element_text(family = "Noto Sans"),
        axis.title = element_text(family = "Noto Sans")) +
  xlab("Rearing Conditions") +
  ylab("Tarsus Length (mm)")

p3 <- data.frame(
  "pretreatment" = c("A", "B", "C"),
  "mass" = 0,
  "tarsus" = 0
) %>%
  mutate(
    "slope" = predict(functionModel3Weeks,
      newdata = ., robust = TRUE,
      resp = "rawSlope", re_form = NA
    )[, "Estimate"],
    "SE" = predict(functionModel3Weeks,
      newdata = ., robust = TRUE,
      resp = "rawSlope", re_form = NA
    )[, "Est.Error"]
  ) %>%
  mutate(slope = slope +
           mean(subset(slopeData, week == "3")$mslope,
                na.rm = T)) %>%
  ggplot(aes(x = pretreatment, y = slope)) +
  geom_errorbar(aes(x = pretreatment,
                    ymin = slope - SE, ymax = slope + SE),
                colour = "black", width = 0.3) +
  geom_point(size = 5, pch = 21, colour = "black",
             aes(x = pretreatment, fill = factor(pretreatment))) +
  geom_line(linetype = "dashed", colour = "black") +
  geom_point(
    size = 2, pch = 21, colour = "black",
    data = subset(slopeData, week == "3") %>%
      mutate("pretreatment" = ifelse(pretreatment == "cold", "A",
        ifelse(pretreatment == "neutral", "B", "C")
      )),
    aes(x = pretreatment, y = mslope,
        fill = factor(pretreatment)),
    position = position_jitter(width = 0.3)
  ) +
  scale_fill_manual(values = c("#7BB4E3", "black", "#CD5C5C")) +
  scale_x_discrete(breaks = c("A", "B", "C"),
                   labels = c("Cold\n(10°C)",
                              "Mild\n(20°C)",
                              "Warm\n(30°C)")) +
  theme_classic() +
  theme(legend.position = "none",
        axis.text = element_text(family = "Noto Sans"),
        axis.title = element_text(family = "Noto Sans")) +
```

```
  xlab("Rearing Conditions") +
  ylab("Metabolic Slope\n(fold Metabolism at Thermoneutrality/°C)")

pretreatmentPanel <- ((p1/p2)| p3) +
  plot_annotation(tag_levels = "A")

showtext_auto()
pretreatmentPanel
```

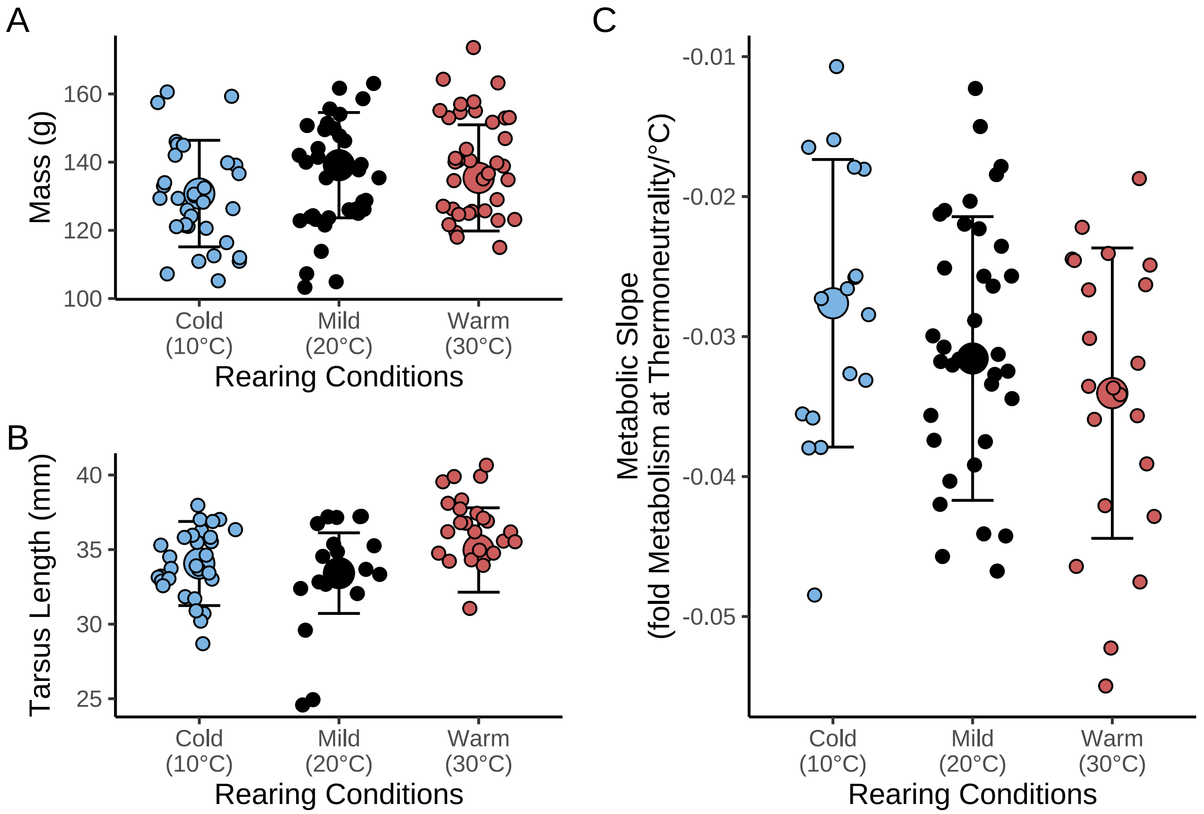


***Figure 29:*** *Effect of post-hatch thermal environment on morphology (body mass, g, and tarsus length, mm) and thermal physiology (metabolic slope, fold RMR at thermoneutrality/°C) in three week old Japanese quail. Metabolic slope is measured at ambient temperatures at and below 30°C (10°C - 30°C).*

```
ggsave("../plots/rearingConditionEffects3Weeks_3fold.pdf",
  pretreatmentPanel,
  dpi = 800, height = 7, width = 9
)
showtext_auto(enable = "FALSE")
```

And predicted effects of morphology on metabolic slopes are also visualised.

```
# And morphology effects on thermal physiology

p1 <- data.frame(
  "pretreatment" = "B",
  "mass" = seq(min(functionModel3Weeks$data$mass, na.rm = T),
    max(functionModel3Weeks$data$mass, na.rm = T),
    by = 0.1
  ),
  "tarsus" = 0
) %>%
  mutate(
```

```
    "slope" = predict(functionModel3Weeks,
      newdata = ., robust = TRUE,
      resp = "rawSlope", re_form = NA
    )[, "Estimate"],
    "SE" = predict(functionModel3Weeks,
      newdata = ., robust = TRUE,
      resp = "rawSlope", re_form = NA
    )[, "Est.Error"]
  ) %>%
  mutate("Mass" = mass +
           mean(subset(slopeData, week == "3")$mass,
                na.rm = T)) %>%
  mutate("slope" = slope +
           mean(subset(slopeData, week == "3")$mslope,
                na.rm = T)) %>%
  ggplot(aes(x = Mass, y = slope)) +
  geom_ribbon(aes(x = Mass, ymin = slope - SE,
                  ymax = slope + SE),
    fill = "grey50", alpha = 0.5
  ) +
  geom_smooth(method = "lm", linetype = "dashed",
              se = FALSE, colour = "black") +
  geom_point(
    size = 2.5, pch = 21, colour = "black", alpha = 0.5,
    fill = "grey50",
    data = subset(slopeData, week == "3"),
    aes(x = mass, y = mslope)
  ) +
  theme_classic() +
  theme(legend.position = "none",
        axis.text = element_text(family = "Noto Sans"),
        axis.title = element_text(family = "Noto Sans")) +
  xlab("Body Mass (g)") +
  ylab("Metabolic Slope (Fold Metabolism\nat Thermoneutrality/°C)")

p2 <- data.frame(
  "pretreatment" = "B",
  "tarsus" = seq(min(functionModel3Weeks$data$tarsus, na.rm = T),
    max(functionModel3Weeks$data$tarsus, na.rm = T),
    by = 0.1
  ),
  "mass" = 0
) %>%
  mutate(
    "slope" = predict(functionModel3Weeks,
      newdata = ., robust = TRUE,
      resp = "rawSlope", re_form = NA
    )[, "Estimate"],
    "SE" = predict(functionModel3Weeks,
      newdata = ., robust = TRUE,
      resp = "rawSlope", re_form = NA
    )[, "Est.Error"]
  ) %>%
  mutate("Tarsus" = tarsus +
           mean(subset(slopeData, week == "3")$tarsusLengthMean,
                na.rm = T)) %>%
  mutate("slope" = slope +
           mean(subset(slopeData, week == "3")$mslope,
                na.rm = T)) %>%
  ggplot(aes(x = Tarsus, y = slope)) +
  geom_ribbon(aes(x = Tarsus, ymin = slope - SE, ymax = slope + SE),
    fill = "grey50", alpha = 0.5
  ) +
  geom_smooth(method = "lm", linetype = "dashed", se = FALSE, colour = "black") +
  geom_point(
    size = 2.5, pch = 21, colour = "black", alpha = 0.5, fill = "grey50",
    data = subset(slopeData, week == "3"),
```

```
    aes(x = tarsusLengthMean, y = mslope)
  ) +
  theme_classic() +
  theme(legend.position = "none",
        axis.text = element_text(family = "Noto Sans"),
        axis.title = element_text(family = "Noto Sans")) +
  xlab("Tarsus Length (mm)") +
  ylab("Metabolic Slope (Fold Metabolism\nat Thermoneutrality/°C)")

massPred <- data.frame(
  "pretreatment" = "B",
  "mass" = c(
    mean(functionModel3Weeks$data$mass, na.rm = T) -
      sd(functionModel3Weeks$data$mass, na.rm = T),
    mean(functionModel3Weeks$data$mass, na.rm = T),
    mean(functionModel3Weeks$data$mass, na.rm = T) +
      sd(functionModel3Weeks$data$mass, na.rm = T)
  ),
  "tarsus" = 0
) %>%
  predict(functionModel3Weeks, re_form = NA,
    resp = "rawSlope", newdata = .,
    summary = TRUE, robust = TRUE
  ) %>%
  as.data.frame() %>%
  mutate("slope" = Estimate +
           mean(
             subset(slopeData,
                    week == 3 & pretreatment == "neutral"
                    )$mslope,
             na.rm = TRUE
           )
  ) %>%
  mutate("Size" = c("Small", "Average", "Large")) %>%
  select(Size, slope, "slopeSE" = Est.Error) %>%
  slice(rep(1:n(), each = length(seq(-20, -0.1, by = 0.1)))) %>%
  mutate("Ta" = rep(seq(-20, -0.1, by = 0.1), 3)) %>%
  mutate("VO2" = 1 + Ta*slope,
         "LL" = 1 + Ta*(slope - slopeSE),
         "UL" = 1 + Ta*(slope + slopeSE)
  ) %>%
  mutate("Ta" = Ta + 30,
         Size = factor(Size, levels = c("Small", "Average", "Large"))
         )

massPlot <- massPred %>%
  ggplot(aes(x = Ta, y = VO2)) +
  geom_point(
    data = slopes3Weeks$data %>%
      merge(., subset(all, week == "3") %>%
              select(ring, mass),
            by = "ring", all.x = TRUE) %>%
      mutate(Ta = Ta + 30),
    aes(x = Ta, y = VO2, colour = mass),
    position = position_jitter(width = 1), alpha = 0.5
  ) +
  geom_point(
    data = data.frame("Ta" = 30, "VO2" = 1),
    pch = 21, colour = "black", fill = "grey15",
    size = 3
  ) +
  geom_ribbon(aes(x = Ta, ymin = LL, ymax = UL, fill = Size),
              alpha = 0.4, colour = "black") +
  geom_line(
    aes(linetype = Size)
  ) +
  theme_classic() +
```

```
  xlab("Ambient Temperature (°C)") +
  ylab("Relative Resting Metabolism\n(Fold Metabolism at Thermoneutrality)") +
  scale_fill_manual(values = c("#DECCC1", "#855E46", "#231709"),
                    labels = c(
    paste0(
      "Mean Mass -\n1 SD (",
      round(
        mean(subset(slopeData, week == "3")$mass, na.rm = T) -
          sd(subset(slopeData, week == "3")$mass, na.rm = T),
            digits = 0.1
        ),
      "g)"
    ),
    paste0(
      "Mean Mass\n(",
      round(mean(subset(slopeData, week == "3")$mass, na.rm = T),
            digits = 0.1),
      "g)"
    ),
    paste0(
      "Mean Mass +\n1 SD (",
      round(
        mean(subset(slopeData, week == "3")$mass, na.rm = T) +
          sd(subset(slopeData, week == "3")$mass, na.rm = T),
            digits = 0.1
        ),
      "g)"
    )
  ), name = NULL) +
  scale_linetype_manual(
    values = c("solid", "dashed", "dotted"),
    labels = c(
    paste0(
      "Mean Mass -\n1 SD (",
      round(
        mean(subset(slopeData, week == "3")$mass, na.rm = T) -
          sd(subset(slopeData, week == "3")$mass, na.rm = T),
            digits = 0.1
        ),
      "g)"
    ),
    paste0(
      "Mean Mass\n(",
      round(mean(subset(slopeData, week == "3")$mass, na.rm = T),
            digits = 0.1),
      "g)"
    ),
    paste0(
      "Mean Mass +\n1 SD (",
      round(
        mean(subset(slopeData, week == "3")$mass, na.rm = T) +
          sd(subset(slopeData, week == "3")$mass, na.rm = T),
            digits = 0.1
        ),
      "g)"
    )
    ), name = NULL
  ) +
  scale_colour_gradient(low = "#DECCC1", high = "#231709",
                        guide = NULL) +
  theme(
    axis.text = element_text(family = "Noto Sans"),
    axis.title = element_text(family = "Noto Sans"),
    legend.position = "bottom"
  )

tarsusPred <- data.frame(
```

```
  "pretreatment" = "B",
  "tarsus" = c(
    mean(functionModel3Weeks$data$tarsus, na.rm = T) -
      sd(functionModel3Weeks$data$tarsus, na.rm = T),
    mean(functionModel3Weeks$data$tarsus, na.rm = T),
    mean(functionModel3Weeks$data$tarsus, na.rm = T) +
      sd(functionModel3Weeks$data$tarsus, na.rm = T)
  ),
  "mass" = 0
) %>%
  predict(functionModel3Weeks, re_form = NA,
    resp = "rawSlope", newdata = .,
    summary = TRUE, robust = TRUE
  ) %>%
  as.data.frame() %>%
  mutate("slope" = Estimate +
           mean(
             subset(slopeData,
                    week == 3 & pretreatment == "neutral"
                    )$mslope,
             na.rm = TRUE
           )
  ) %>%
  mutate("Size" = c("Small", "Average", "Large")) %>%
  select(Size, slope, "slopeSE" = Est.Error) %>%
  slice(rep(1:n(), each = length(seq(-20, -0.1, by = 0.1)))) %>%
  mutate("Ta" = rep(seq(-20, -0.1, by = 0.1), 3)) %>%
  mutate("VO2" = 1 + Ta*slope,
         "LL" = 1 + Ta*(slope - slopeSE),
         "UL" = 1 + Ta*(slope + slopeSE)
  ) %>%
  mutate("Ta" = Ta + 30,
         Size = factor(Size, levels = c("Small", "Average", "Large"))
         )

tarsusPlot <- tarsusPred %>%
  ggplot(aes(x = Ta, y = VO2)) +
  geom_point(
    data = slopes3Weeks$data %>%
      merge(., subset(all, week == "3") %>%
              select(ring, "tarsus" = tarsusLengthMean),
            by = "ring", all.x = TRUE) %>%
      mutate(Ta = Ta + 30),
    aes(x = Ta, y = VO2, colour = tarsus),
    position = position_jitter(width = 1), alpha = 0.5
  ) +
  geom_point(
    data = data.frame("Ta" = 30, "VO2" = 1),
    pch = 21, colour = "black", fill = "grey15",
    size = 3
  ) +
  geom_ribbon(aes(x = Ta, ymin = LL, ymax = UL, fill = Size),
              alpha = 0.4, colour = "black") +
  geom_line(
    aes(linetype = Size)
  ) +
  theme_classic() +
  xlab("Ambient Temperature (°C)") +
  ylab("Relative Resting Metabolism\n(Fold Metabolism at Thermoneutrality)") +
  scale_fill_manual(values = c("#DECCC1", "#855E46", "#231709"),
                    labels = c(
    paste0(
      "Mean Tarsus\nLength -\n1 SD (",
      round(
        mean(subset(slopeData, week == "3")$tarsusLengthMean,
             na.rm = T) -
          sd(subset(slopeData, week == "3")$tarsusLengthMean,
```

```
          na.rm = T),
         digits = 0.1
      ),
    "g)"
  ),
  paste0(
    "Mean Tarsus\nLength (",
    round(
      mean(subset(slopeData, week == "3")$tarsusLengthMean,
           na.rm = T),
         digits = 0.1),
    "g)"
  ),
  paste0(
    "Mean Tarsus\nLength +\n1 SD (",
    round(
      mean(subset(slopeData, week == "3")$tarsusLengthMean,
           na.rm = T) +
        sd(subset(slopeData, week == "3")$tarsusLengthMean,
          na.rm = T),
         digits = 0.1
      ),
    "g)"
  )
), name = NULL) +
scale_linetype_manual(
  values = c("solid", "dashed", "dotted"),
  labels = c(
  paste0(
    "Mean Tarsus\nLength -\n1 SD (",
    round(
      mean(subset(slopeData, week == "3")$tarsusLengthMean,
           na.rm = T) -
        sd(subset(slopeData, week == "3")$tarsusLengthMean,
          na.rm = T),
         digits = 0.1
      ),
    "g)"
  ),
  paste0(
    "Mean Tarsus\nLength (",
    round(
      mean(subset(slopeData, week == "3")$tarsusLengthMean,
           na.rm = T),
         digits = 0.1),
    "g)"
  ),
  paste0(
    "Mean Tarsus\nLength +\n1 SD (",
    round(
      mean(subset(slopeData, week == "3")$tarsusLengthMean,
           na.rm = T) +
        sd(subset(slopeData, week == "3")$tarsusLengthMean,
          na.rm = T),
         digits = 0.1
      ),
    "g)"
  )
  ), name = NULL
) +
scale_colour_gradient(low = "#DECCC1", high = "#231709",
                      guide = NULL) +
theme(
  axis.text = element_text(family = "Noto Sans"),
  axis.title = element_text(family = "Noto Sans"),
  legend.position = "bottom"
)
```

```
showtext.auto()

allMorphology <- (p1 + massPlot) / (p2 + tarsusPlot) +
  plot_annotation(tag_levels = "A")

allMorphology
```

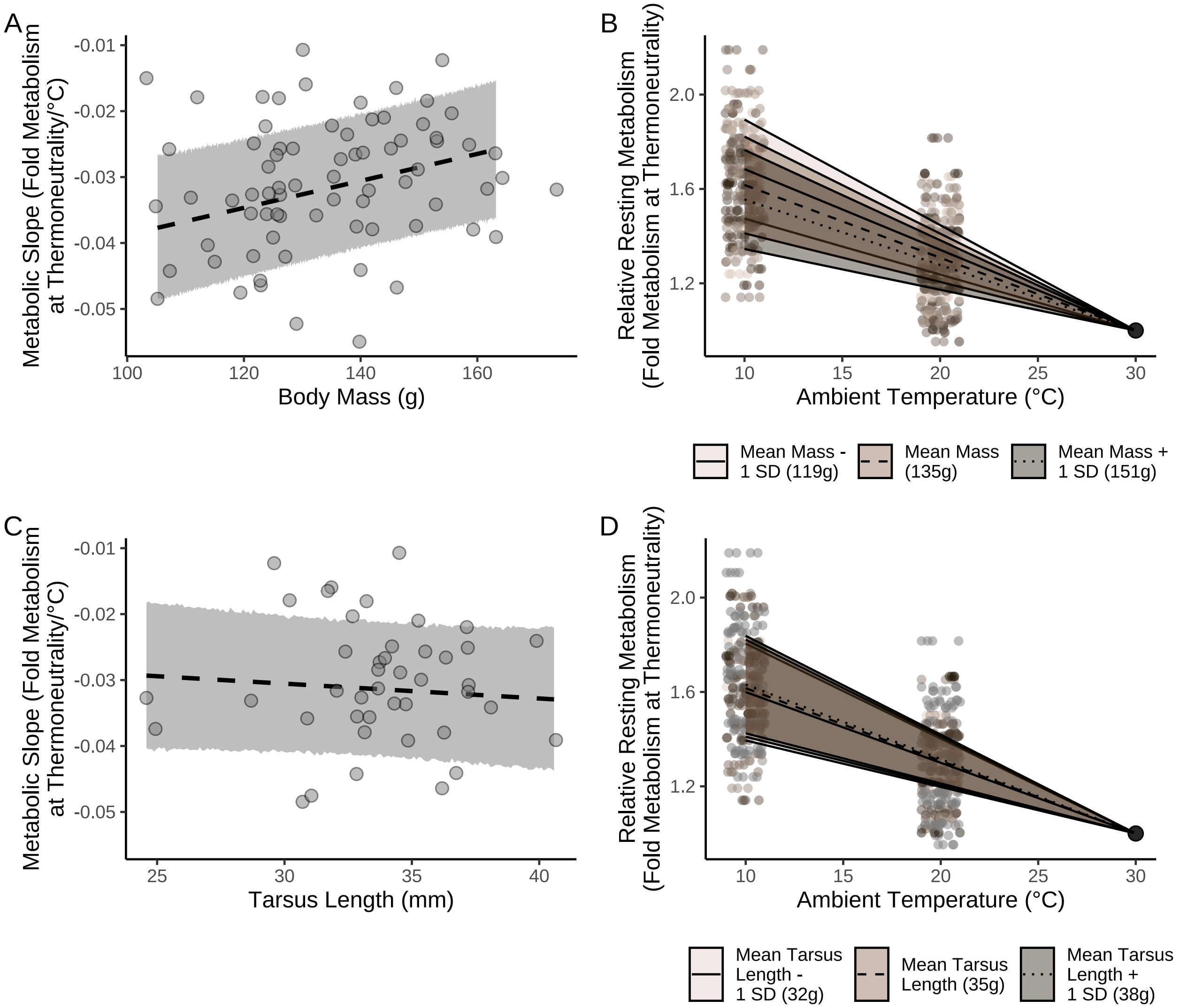


***Figure 30:*** *Effect of morphology (body mass, g, and tarsus length, mm) on metabolic slopes (fold RMR at thermoneutrality/°C) at ambient temperatures below thermoneutrality (<30°C) in three week old Japanese quail. Dots represent raw data points and lines and ribbons represent predicted relationships +/- 1 standard error from a Bayesian path analysis.*

```
ggsave("../plots/morphologyEffectsResistance3Weeks_3Fold.pdf",
  allMorphology,
  dpi = 800, height = 7, width = 9
)

showtext.auto(enable = "FALSE")
```

```
# As fold increase at 10°C

massPlot10 <- massPred %>%
```

```
  filter(Ta == 10) %>%
  ggplot(aes(x = Size, y = VO2)) +
  geom_point(
    data = slopes3Weeks$data %>%
      filter(Ta == -20) %>%
      merge(., subset(all, week == "3") %>%
              select(ring, mass) %>%
              distinct() %>%
              mutate(Size = ifelse(mass < (mean(mass, na.rm = T) -
                                            sd(mass, na.rm = T)),
                                  "Small",
                                  ifelse((mass > (mean(mass, na.rm = T) -
                                            sd(mass, na.rm = T))) &
                                           (mass < (mean(mass, na.rm = T) +
                                            sd(mass, na.rm = T))),
                                         "Average", "Large"
                                  )
                                  )
                     ),
            by = "ring", all.x = TRUE) %>%
      mutate(Size = factor(Size, levels = c("Small", "Average", "Large"))
             ),
    aes(x = Size, y = VO2, colour = Size),
    position = position_jitter(width = 0.25), alpha = 0.5
  ) +
  geom_errorbar(aes(ymin = LL, ymax = UL),
                colour = "black", width = 0.25) +
  geom_point(size = 3, pch = 21, colour = "black",
             aes(fill = Size)
  ) +
  xlab("Body Mass") +
  ylab("Fold Increase in Metabolism at 10°C\n(From Thermoneutrality)") +
  scale_x_discrete(labels = c("Mean - 1 SD", "Mean", "Mean + 1 SD")) +
  scale_fill_manual(values = c("#DECCC1", "#855E46", "#231709")) +
  scale_colour_manual(values = c("#DECCC1", "#855E46", "#231709")) +
  theme_classic() +
  theme(
    legend.position = "none",
    axis.text = element_text(family = "Noto Sans"),
    axis.title = element_text(family = "Noto Sans")
  )

tarsusPlot10 <- tarsusPred %>%
  filter(Ta == 10) %>%
  ggplot(aes(x = Size, y = VO2)) +
  geom_point(
    data = slopes3Weeks$data %>%
      filter(Ta == -20) %>%
      merge(., subset(all, week == "3") %>%
              select(ring, "tarsus" = tarsusLengthMean) %>%
              distinct() %>%
              mutate(Size = ifelse(tarsus < (mean(tarsus, na.rm = T) -
                                            sd(tarsus, na.rm = T)),
                                  "Small",
                                  ifelse((tarsus > (mean(tarsus, na.rm = T) -
                                            sd(tarsus, na.rm = T))) &
                                           (tarsus < (mean(tarsus, na.rm = T) +
                                            sd(tarsus, na.rm = T))),
                                         "Average", "Large"
                                  )
                                  )
                     ),
            by = "ring", all.x = TRUE) %>%
      drop_na(tarsus) %>%
      mutate(Size = factor(Size, levels = c("Small", "Average", "Large"))
             ),
    aes(x = Size, y = VO2, colour = Size),
```

```
    position = position_jitter(width = 0.25), alpha = 0.5
  ) +
  geom_errorbar(aes(ymin = LL, ymax = UL),
                colour = "black", width = 0.25) +
  geom_point(size = 3, pch = 21, colour = "black",
             aes(fill = Size)
  ) +
  xlab("Tarsus Length") +
  ylab("Fold Increase in Metabolism at 10°C\n(From Thermoneutrality)") +
  scale_x_discrete(labels = c("Mean - 1 SD",
                              "Mean",
                              "Mean + 1 SD")) +
  scale_fill_manual(values = c("#DECCC1", "#855E46", "#231709")) +
  scale_colour_manual(values = c("#DECCC1", "#855E46", "#231709")) +
  theme_classic() +
  theme(
    legend.position = "none",
    axis.text = element_text(family = "Noto Sans"),
    axis.title = element_text(family = "Noto Sans")
  )

allMorphology10 <- (p1 + massPlot10) / (p2 + tarsusPlot10) +
  plot_annotation(tag_levels = "A")

ggsave("../plots/morphologyEffectsResistance3Weeks_3Fold10.pdf",
  allMorphology10,
  dpi = 800, height = 7, width = 9
)
```

Finally, we summarise model outcomes and estimate partial $R^2$ values for population-level model predictors.

```
caption <- paste0("Results from a Bayesian path analysis ",
                  "predicting metabolic slopes (fold resting ",
                  "metabolism at thermoneutrality/°C) ",
                  "as a function of morphometry in three ",
                  "week old Japanese quail. ",
                  "Cold rearing indicates post-hatch rearing at ",
                  "10°C, relative to ",
                  "20°C (intercept), or ",
                  "30°C ('warm rearing'). ",
                  "Coefficients represent medians and credible intervals ",
                  "(CIs) represent quantile intervals. ",
                  "BF indicates Bayes Factors."
)

week3Results <-
  as.data.frame(functionModel3Weeks) %>%
  summarise_all(., .funs = median) %>%
  pivot_longer(everything(), names_to = "Parameter",
               values_to = "Estimate") %>%
  merge(., quantileCIs(functionModel3Weeks, cis = c(50, 95)),
        by = "Parameter", all.x = TRUE) %>%
  filter(grepl("b_|sd_", Parameter)) %>%
  rowwise() %>%
  mutate("BF" = ifelse(Estimate < 0,
                       (2 * mean(as.data.frame(
                         functionModel3Weeks)[, Parameter] <= 0)) /
    (2 * mean(as.data.frame(
      functionModel3Weeks)[, Parameter] >= 0)),
  (2 * mean(as.data.frame(
    functionModel3Weeks)[, Parameter] >= 0)) /
    (2 * mean(as.data.frame(
      functionModel3Weeks)[, Parameter] <= 0))
  )) %>%
  ungroup() %>%
  mutate(
    "Estimate" = round(Estimate, digits = 4),
```

```
    "BF" = round(BF, digits = 4),
    "N" = nrow(functionModel3Weeks$data)
  ) %>%
  mutate("Parameter" = ifelse(grepl("b_", Parameter),
                              gsub("b_", "",
                            Parameter),
                            gsub("Intercept", "batch",
                                 gsub("sd_batch__", "", Parameter)
                                 )
  )
  ) %>%
  mutate(
    "Response" = gsub("_.*", "", Parameter),
    "Parameter" = gsub(".*_", "", Parameter)
  ) %>%
  merge(., tribble(
    ~Response, ~response, ~level,
    "mass", "Body Mass (g)", "A",
    "rawSlope",
    "Metabolic Slope", "D",
    "tarsus", "Tarsus Length (mm)", "B"
  ),
  by = "Response"
  ) %>%
  merge(., tribble(
    ~Parameter, ~parameter, ~number,
    "Intercept", "Intercept", "1",
    "mass", "Body Mass (g)", "4",
    "tarsus", "Tarsus Length (mm)", "5",
    "pretreatmentA", "Cold Rearing", "2",
    "pretreatmentC", "Warm Rearing", "3",
    "batch", "Egg Batch [mu]", "6"
  ),
  by = "Parameter"
  ) %>%
  mutate(
    `50\\% HDI` = paste0("(", paste(
      round(Low_CI_50, digits = 4),
      round(High_CI_50, digits = 4),
      sep = ", "
    ), ")"),
    `95\\% HDI` = paste0("(", paste(
      round(Low_CI_95, digits = 4),
      round(High_CI_95, digits = 4),
      sep = ", "
    ), ")")
  ) %>%
  select(-c(Low_CI_50, High_CI_50, Low_CI_95, High_CI_95)) %>%
  select(
    "Response" = "response", "Parameter" = "parameter", N,
    Estimate, `50\\% HDI`, `95\\% HDI`, BF, level, number
  ) %>%
  arrange(level, number) %>%
  select(-c(level, number)) %>%
  kbl(.,
    longtable = T, booktabs = T, format = "latex", escape = FALSE,
    caption = caption
  ) %>%
  column_spec(column = c(1:2), width = "2.2cm")  %>%
  column_spec(column = c(3:10), width = "1.9cm") %>%
  kable_styling(latex_options = "striped")

week3Results
```

***Table 10:*** *Results from a Bayesian path analysis predicting metabolic slopes (fold resting metabolism at thermoneutrality/°C) as a function of morphometry in three week old Japanese quail. Cold rearing indicates post-hatch rearing at 10°C, relative to 20°C (intercept), or 30°C ('warm rearing'). Coefficients represent medians and credible intervals (CIs) represent quantile intervals. BF indicates Bayes Factors.*

| Response | Parameter | N | Estimate | 50% HDI | 95% HDI | BF |
|---|---|---|---|---|---|---|
| Body Mass (g) | Intercept | 43 | 3.7121 | (1.5743, 5.9732) | (-2.7568, 10.3314) | 6.7146 |
| Body Mass (g) | Cold Rearing | 43 | -8.0843 | (-11.3903, -4.7943) | (-17.6359, 2.0016) | 18.0024 |
| Body Mass (g) | Warm Rearing | 43 | -3.0488 | (-6.9634, 0.8108) | (-14.3291, 8.2615) | 2.3884 |
| Body Mass (g) | Egg Batch [mu] | 43 | 0.2777 | (0.1167, 0.5548) | (0.0103, 1.4707) | Inf |
| Tarsus Length (mm) | Intercept | 43 | -1.2945 | (-1.8426, -0.7214) | (-2.9712, 0.738) | 10.5108 |
| Tarsus Length (mm) | Cold Rearing | 43 | 0.7069 | (0.0564, 1.3746) | (-1.1967, 2.6091) | 3.2965 |
| Tarsus Length (mm) | Warm Rearing | 43 | 1.5660 | (0.7349, 2.4057) | (-0.8813, 3.9412) | 8.5352 |
| Tarsus Length (mm) | Body Mass (g) | 43 | 0.0967 | (0.0797, 0.1144) | (0.0465, 0.1505) | 1599.0000 |
| Tarsus Length (mm) | Egg Batch [mu] | 43 | 0.8230 | (0.4089, 1.2746) | (0.038, 2.4098) | Inf |
| Metabolic Slope | Intercept | 43 | -0.0006 | (-0.0031, 0.0017) | (-0.0112, 0.0092) | 1.3269 |
| Metabolic Slope | Cold Rearing | 43 | 0.0038 | (0.0015, 0.0061) | (-0.003, 0.0106) | 6.2398 |
| Metabolic Slope | Warm Rearing | 43 | -0.0025 | (-0.0054, 5e-04) | (-0.0112, 0.0063) | 2.5088 |
| Metabolic Slope | Body Mass (g) | 43 | 0.0002 | (1e-04, 3e-04) | (0, 4e-04) | 29.6513 |
| Metabolic Slope | Tarsus Length (mm) | 43 | -0.0002 | (-6e-04, 1e-04) | (-0.0013, 9e-04) | 1.9762 |
| Metabolic Slope | Egg Batch [mu] | 43 | 0.0039 | (0.0017, 0.0082) | (2e-04, 0.0267) | Inf |

```
#save_kable(week3Results, "../tables/threeWeekResults_3.html")

# And checking partial R2 values of morphometry on metabolic slopes

functionModel3WeeksR2Tarsus <-
  brm(
    data = slopeData %>%
      filter(week == "3") %>%
      mutate(
        mass = mass - mean(mass, na.rm = T),
        tarsus = tarsusLengthMean - mean(tarsusLengthMean, na.rm = T),
        pretreatment = ifelse(pretreatment == "cold", "A",
          ifelse(pretreatment == "neutral", "B", "C")
        )
      ) %>%
      mutate(pretreatment = factor(pretreatment, levels = c("B", "A", "C"))) %>%
      drop_na() %>%
      merge(., data %>%
              select(ring, "batch" = exp) %>%
              distinct(),
            by = "ring", all.x = TRUE),
    family = "gaussian",
    bf(mass ~ pretreatment + (1|batch)) +
      bf(tarsus ~ mass + pretreatment + (1|batch)) +
      bf(rawSlope ~ mass + pretreatment + (1|batch)) +
      set_rescor(FALSE),
    prior = c(
      set_prior("normal(0, 5)", class = "Intercept", resp = "mass"),
```

```
      set_prior("normal(0, 15)", class = "b",
                coef = "pretreatmentA", resp = "mass"),
      set_prior("normal(0, 15)", class = "b",
                coef = "pretreatmentC", resp = "mass"),
      set_prior("exponential(2.5)", class = "sd",
                group = "batch", resp = "mass"),
      set_prior("exponential(0.15)", class = "sigma", resp = "mass"),
      set_prior("normal(0, 2.5)", class = "Intercept", resp = "tarsus"),
      set_prior("normal(0, 2.5)", class = "b",
                coef = "pretreatmentA", resp = "tarsus"),
      set_prior("normal(0, 2.5)", class = "b",
                coef = "pretreatmentC", resp = "tarsus"),
      set_prior("gamma(1, 5)", class = "b",
                coef = "mass", resp = "tarsus"),
      set_prior("exponential(2)", class = "sd",
                group = "batch", resp = "tarsus"),
      set_prior("exponential(1)", class = "sigma", resp = "tarsus"),
      set_prior("normal(0, 0.01)", class = "Intercept",
                resp = "rawSlope"),
      set_prior("normal(0, 0.025)", class = "b",
                coef = "pretreatmentA", resp = "rawSlope"),
      set_prior("normal(0, 0.025)", class = "b",
                coef = "pretreatmentC", resp = "rawSlope"),
      set_prior("normal(0, 0.001)", class = "b",
                coef = "mass", resp = "rawSlope"),
      set_prior("exponential(50)", class = "sd",
                group = "batch", resp = "rawSlope"),
      set_prior("exponential(10)", class = "sigma",
                resp = "rawSlope")
    ),
    iter = 50000, warmup = 10000, cores = 4, chains = 4, thin = 20,
    control = list(adapt_delta = .97, max_treedepth = 14),
    silent = TRUE, refresh = 0,
    file = "./models/threeWeekFunctionModelTarsusR2.Rds"
  )

functionModel3R2Mass <-
  brm(
    data = slopeData %>%
      filter(week == "3") %>%
      mutate(
        mass = mass - mean(mass, na.rm = T),
        tarsus = tarsusLengthMean - mean(tarsusLengthMean, na.rm = T),
        pretreatment = ifelse(pretreatment == "cold", "A",
          ifelse(pretreatment == "neutral", "B", "C")
        )
      ) %>%
      mutate(pretreatment = factor(pretreatment, levels = c("B", "A", "C"))) %>%
      drop_na() %>%
      merge(., data %>%
              select(ring, "batch" = exp) %>%
              distinct(),
            by = "ring", all.x = TRUE),
    family = "gaussian",
    bf(mass ~ pretreatment + (1|batch)) +
      bf(tarsus ~ mass + pretreatment + (1|batch)) +
      bf(rawSlope ~ tarsus + pretreatment + (1|batch)) +
      set_rescor(FALSE),
    prior = c(
      set_prior("normal(0, 5)", class = "Intercept", resp = "mass"),
      set_prior("normal(0, 15)", class = "b",
                coef = "pretreatmentA", resp = "mass"),
      set_prior("normal(0, 15)", class = "b",
                coef = "pretreatmentC", resp = "mass"),
      set_prior("exponential(2.5)", class = "sd",
                group = "batch", resp = "mass"),
      set_prior("exponential(0.15)", class = "sigma", resp = "mass"),
```

```
      set_prior("normal(0, 2.5)", class = "Intercept", resp = "tarsus"),
      set_prior("normal(0, 2.5)", class = "b",
                coef = "pretreatmentA", resp = "tarsus"),
      set_prior("normal(0, 2.5)", class = "b",
                coef = "pretreatmentC", resp = "tarsus"),
      set_prior("gamma(1, 5)", class = "b",
                coef = "mass", resp = "tarsus"),
      set_prior("exponential(2)", class = "sd",
                group = "batch", resp = "tarsus"),
      set_prior("exponential(1)", class = "sigma", resp = "tarsus"),
      set_prior("normal(0, 0.01)", class = "Intercept",
                resp = "rawSlope"),
      set_prior("normal(0, 0.025)", class = "b",
                coef = "pretreatmentA", resp = "rawSlope"),
      set_prior("normal(0, 0.025)", class = "b",
                coef = "pretreatmentC", resp = "rawSlope"),
      set_prior("normal(0, 0.0025)", class = "b",
                coef = "tarsus", resp = "rawSlope"),
      set_prior("exponential(50)", class = "sd",
                group = "batch", resp = "rawSlope"),
      set_prior("exponential(10)", class = "sigma",
                resp = "rawSlope")
    ),
    iter = 50000, warmup = 10000, cores = 4, chains = 4, thin = 20,
    control = list(adapt_delta = .97, max_treedepth = 14),
    silent = TRUE, refresh = 0,
    file = "./models/threeWeekFunctionModelMassR2.Rds"
  )

massPR2 = brms::bayes_R2(functionModel3Weeks, ndraws = 1000,
               resp = "rawSlope", summary = FALSE,
               robust = TRUE) %>%
  as.data.frame() %>%
  select("fullR2" = R2rawSlope) %>%
  cbind(.,
        brms::bayes_R2(functionModel3R2Mass, ndraws = 1000,
               resp = "rawSlope", summary = FALSE,
               robust = TRUE)
        ) %>%
  mutate("pR2" = round(fullR2 - R2rawSlope, digits = 3)) %>%
  pull(pR2)

tarsusPR2 = brms::bayes_R2(functionModel3Weeks, ndraws = 1000,
               resp = "rawSlope", summary = FALSE,
               robust = TRUE) %>%
  as.data.frame() %>%
  select("fullR2" = R2rawSlope) %>%
  cbind(.,
        brms::bayes_R2(functionModel3WeeksR2Tarsus, ndraws = 1000,
               resp = "rawSlope", summary = FALSE,
               robust = TRUE)
        ) %>%
  mutate("pR2" = round(fullR2 - R2rawSlope, digits = 3)) %>%
  pull(pR2)

caption = paste0('Variance in metabolic slope ',
  '(fold resting metabolsim at thermoneutrality/°C) explained by ',
  'morphometry in three week old Japanese quail. Metabolic ',
  'slopes are measured below thermoneutrality (<30°C).'
  )

quantile(massPR2, probs = c(0.025, 0.975), type = 8) %>%
  t() %>%
  as.data.frame() %>%
  mutate(`2.5%` = ifelse(`2.5%` < 0, 0, `2.5%`)) %>%
  mutate("95\\% CI" = paste0("[",
                           round(`2.5%`, digits = 3),
```

```
                          ",",
                          round(`97.5%`, digits = 3),
                          "]")
  ) %>%
  mutate("massR2" =
           round(median(massPR2), digits = 3)) %>%
  select("Mass Partial R\\textsuperscript{2}" = massR2,
         "95\\% CI") %>%
  cbind(.,
    quantile(tarsusPR2, probs = c(0.025, 0.975), type = 8) %>%
      t() %>%
      as.data.frame() %>%
      mutate(`2.5%` = ifelse(`2.5%` < 0, 0, `2.5%`)) %>%
      mutate("95\\% CI" = paste0("[",
                          round(`2.5%`, digits = 3),
                          ",",
                          round(`97.5%`, digits = 3),
                          "]")
           ) %>%
      mutate("tarsusR2" = round(median(tarsusPR2), digits = 3)) %>%
      select("Tarsus Partial R\\textsuperscript{2}" = tarsusR2,
             "95\\% CI")
  ) %>%
  kbl(.,
    longtable = T, booktabs = T, format = "latex",
    caption = caption, escape = FALSE
  ) %>%
  column_spec(column = c(1:10), width = "2.5cm")  %>%
  kable_styling(latex_options = "striped")
```

***Table 11:*** *Variance in metabolic slope (fold resting metabolsim at thermoneutrality/°C) explained by morphometry in three week old Japanese quail. Metabolic slopes are measured below thermoneutrality (<30°C).*

| Mass Partial $R^2$ | 95% CI | Tarsus Partial $R^2$ | 95% CI |
|---|---|---|---|
| 0.092 | [0,0.292] | 0.02 | [0,0.244] |

To better understand the potential of selection on thermoregulatory efficiency to influence morphology in quail, we estimate the metabolic costs of misalignment with Bergmann's and Allen's rules (i.e. by falling 2 standard deviations below average mass or above average tarsus length respectively).

```
data.frame(
  "Size" = c("Average", "Small (2x s.d. < mean)"),
  "pretreatment" = "B",
  "mass" = c(mean(functionModel3Weeks$data$mass, na.rm = T),
             mean(functionModel3Weeks$data$mass, na.rm = T) -
               2*sd(functionModel3Weeks$data$mass, na.rm = T)),
  "tarsus" = 0
) %>%
  mutate("slope" = predict(functionModel3Weeks, newdata = .,
                           robust = TRUE, re_form = NA,
                           resp = "rawSlope")[, "Estimate"]) %>%
  mutate(slope = slope +
           mean(subset(slopeData,
                       week == 3 & pretreatment == "neutral")$mslope,
                na.rm = T)
  ) %>%
  mutate(cost = 1 + slope*-30) %>%
  select("Body Size" = Size, "Expenditure Rate at 0°C\n(fold RMR)" = cost)
```

```
## # A tibble: 2 x 2
##   `Body Size`            `Expenditure Rate at 0°C\n(fold RMR)`
##   <chr>                                                  <dbl>
## 1 Average                                                 1.93
```

```
## 2 Small (2x s.d. < mean)                                    2.11
```

```
data.frame(
  "Size" = c("Average", "Long (2x s.d. > mean)"),
  "pretreatment" = "B",
  "tarsus" = c(mean(functionModel3Weeks$data$tarsus, na.rm = T),
             mean(functionModel3Weeks$data$tarsus, na.rm = T) +
               2*sd(functionModel3Weeks$data$tarsus, na.rm = T)),
  "mass" = 0
) %>%
  mutate("slope" = predict(functionModel3Weeks, newdata = .,
                             robust = TRUE, re_form = NA,
                           resp = "rawSlope")[, "Estimate"]) %>%
  mutate(slope = slope +
           mean(subset(slopeData, week == 3)$mslope, na.rm = T)
  ) %>%
  mutate(cost = 1 + slope*-20) %>%
  select("Appendage Length" = Size,
         "Expenditure Rate at 0°C\n(fold RMR)" = cost)
```

```
## # A tibble: 2 x 2
##   `Appendage Length`    `Expenditure Rate at 0°C\n(fold RMR)`
##   <chr>                                                 <dbl>
## 1 Average                                                1.62
## 2 Long (2x s.d. > mean)                                  1.66
```

```
# Checking whether these differences are statistically clear

mismatchDFMass <- data.frame(
  "Size" = c("Average", "Small (2x s.d. < mean)"),
  "pretreatment" = "B",
  "mass" = c(mean(functionModel3Weeks$data$mass, na.rm = T),
             mean(functionModel3Weeks$data$mass, na.rm = T) -
               2*sd(functionModel3Weeks$data$mass, na.rm = T)),
  "tarsus" = 0
) %>% predict(functionModel3Weeks, newdata = .,
              robust = TRUE, re_form = NA,
              resp = "rawSlope", summary = FALSE) %>%
  as.data.frame() %>%
  rename("Average" = "V1", "Small" = "V2") %>%
  mutate_all(.funs = function(x)
    {x + mean(subset(slopeData, week == 3)$mslope, na.rm = T)}
    )

mismatchDFMassTest <- build_hdf(
  vars = list(
    mismatchDFMass$Average,
    mismatchDFMass$Small
    ),
  priors = list(
    rnorm(nrow(mismatchDFMass), 0, 0.01),
    rnorm(nrow(mismatchDFMass), 0, 0.01)
    ),
  names = c("Average", "Small")
)

mismatchTestMass <- hypothesis_df("Small > Average",
                                  mismatchDFMassTest,
                                  class = "b", alpha = 0.05
                                  )

mismatchTestMass$hypothesis$Hypothesis =
  "Small Metabolic Slope > Average Slope"

# Summarising

caption <- paste0(
  "Results of a non-linear hypothesis ",
  "tests comparing predicted metabolic slopes in the cold (30°C - 10°C) ",
```

```
  "among three week old Japanese quail of average or atypically small ",
  "body size (mean - 2 standard deviations). Posterior probabilities are ",
  "calculated using the Savage-Dickey ",
  "density ratio method."
)

mismatchTestMassTable <- mismatchTestMass$hypothesis %>%
  mutate(Hypothesis = c(
    "Small Metabolic Slope > Average Slope"
  )) %>%
  select(-c(Evid.Ratio, Star)) %>%
  mutate(
    Estimate = round(Estimate, digits = 4),
    "Est.Error" = round(Est.Error, digits = 4),
    "CI.Lower" = round(CI.Lower, digits = 4),
    "CI.Upper" = round(CI.Upper, digits = 4)
  ) %>%
  rename(
    "Difference Between Metabolic Slopes" = Estimate,
    "Standard Error" = Est.Error,
    `2.5 % CI` = "CI.Lower",
    `97.5 % CI` = "CI.Upper",
    "Posterior Probability" = Post.Prob
  ) %>%
  kbl(.,
    longtable = T, booktabs = T, format = "latex",
    caption = caption
  ) %>%
  column_spec(column = c(1:2), width = "2.5cm") %>%
  column_spec(column = c(3:10), width = "2cm") %>%
  kable_styling(latex_options = "striped")

mismatchTestMassTable
```

***Table 12:*** *Results of a non-linear hypothesis tests comparing predicted metabolic slopes in the cold (30°C - 10°C) among three week old Japanese quail of average or atypically small body size (mean - 2 standard deviations). Posterior probabilities are calculated using the Savage-Dickey density ratio method.*

| Hypothesis | Difference Between Metabolic Slopes | Standard Error | 2.5 % CI | 97.5 % CI | Posterior Probability |
|---|---|---|---|---|---|
| Small Metabolic Slope > Average Slope | -0.0063 | 0.0137 | -0.029 | 0.0162 | 0.3185 |

```
# For tarsus length

mismatchDFTarsus <- data.frame(
  "Size" = c("Average", "Long (2x s.d. > mean)"),
  "pretreatment" = "B",
  "tarsus" = c(mean(functionModel3Weeks$data$tarsus, na.rm = T),
            mean(functionModel3Weeks$data$tarsus, na.rm = T) +
              2*sd(functionModel3Weeks$data$tarsus, na.rm = T)),
  "mass" = 0
) %>% predict(functionModel3Weeks, newdata = .,
              robust = TRUE, re_form = NA,
              resp = "rawSlope", summary = FALSE) %>%
  as.data.frame() %>%
  rename("Average" = "V1", "Long" = "V2") %>%
  mutate_all(.funs = function(x)
    {x + mean(subset(slopeData, week == 3)$mslope, na.rm = T)}
    )

mismatchDFTarsusTest <- build_hdf(
  vars = list(
    mismatchDFTarsus$Average,
```

```r
    mismatchDFTarsus$Long
    ),
  priors = list(
    rnorm(nrow(mismatchDFTarsus), 0, 0.01),
    rnorm(nrow(mismatchDFTarsus), 0, 0.01)
    ),
  names = c("Average", "Long")
)

mismatchTestTarsus <- hypothesis_df("Long > Average",
                                     mismatchDFTarsusTest,
                                     class = "b", alpha = 0.05
                                     )

mismatchTestTarsus$hypothesis$Hypothesis =
  "Long Tarsus Metabolic Slope > Average Slope"

caption <- paste0(
  "Results of a non-linear hypothesis ",
  "tests comparing predicted metabolic slopes in the cold (30°C - 10°C) ",
  "among three week old Japanese quail of average or atypically long ",
  "tarsus length (mean + 2 standard deviations). Posterior probabilities are ",
  "calculated using the Savage-Dickey ",
  "density ratio method."
)

mismatchTestTarsusTable <- mismatchTestTarsus$hypothesis %>%
  mutate(Hypothesis = c(
    "Small Metabolic Slope > Average Slope"
  )) %>%
  select(-c(Evid.Ratio, Star)) %>%
  mutate(
    Estimate = round(Estimate, digits = 4),
    "Est.Error" = round(Est.Error, digits = 4),
    "CI.Lower" = round(CI.Lower, digits = 4),
    "CI.Upper" = round(CI.Upper, digits = 4)
  ) %>%
  rename(
    "Difference Between Metabolic Slopes" = Estimate,
    "Standard Error" = Est.Error,
    `2.5 % CI` = "CI.Lower",
    `97.5 % CI` = "CI.Upper",
    "Posterior Probability" = Post.Prob
  ) %>%
  kbl(.,
    longtable = T, booktabs = T, format = "latex",
    caption = caption
  ) %>%
  column_spec(column = c(1:2), width = "2.5cm") %>%
  column_spec(column = c(3:10), width = "2cm") %>%
  kable_styling(latex_options = "striped")

mismatchTestTarsusTable
```

***Table 13:*** *Results of a non-linear hypothesis tests comparing predicted metabolic slopes in the cold (30°C - 10°C) among three week old Japanese quail of average or atypically long tarsus length (mean + 2 standard deviations). Posterior probabilities are calculated using the Savage-Dickey density ratio method.*

| Hypothesis | Difference Between Metabolic Slopes | Standard Error | 2.5 % CI | 97.5 % CI | Posterior Probability |
|---|---|---|---|---|---|
| Small Metabolic Slope > Average Slope | -0.0017 | 0.0136 | -0.0239 | 0.0205 | 0.4535 |

Any effects of prior thermal acclimation or thermal history (i.e. via our distinct rearing treatments) on metabolic slopes may be direct, indirect through their influence on morphology, or both. Using outcomes from our path analysis, we visualise possible contributions of each pathway below. To both ease interpretation and comparison of effects, each are first scaled to represent the effect that a change in one standard deviation (or one categorical level) of a variable is predicted to have on another.

```
# Visualising indirect and direct effects

scaledBetas <- as.data.frame(functionModel3Weeks) %>%
  mutate(b_mass_pretreatmentA = b_mass_pretreatmentA/
           sd(functionModel3Weeks$data$mass),
         b_mass_pretreatmentC = b_mass_pretreatmentC/
           sd(functionModel3Weeks$data$mass),
         b_tarsus_pretreatmentA = b_tarsus_pretreatmentA/
           sd(functionModel3Weeks$data$tarsus),
         b_tarsus_pretreatmentC = b_tarsus_pretreatmentC/
           sd(functionModel3Weeks$data$tarsus),
         b_tarsus_mass =
           (b_tarsus_mass*sd(functionModel3Weeks$data$mass))/
           sd(functionModel3Weeks$data$tarsus),
         b_rawSlope_mass =
           (b_rawSlope_mass*sd(functionModel3Weeks$data$mass))/
           sd(functionModel3Weeks$data$rawSlope),
         b_rawSlope_tarsus =
           (b_rawSlope_tarsus*sd(functionModel3Weeks$data$tarsus))/
           sd(functionModel3Weeks$data$rawSlope),
         b_rawSlope_pretreatmentA = b_rawSlope_pretreatmentA/
           sd(functionModel3Weeks$data$rawSlope),
         b_rawSlope_pretreatmentC = b_rawSlope_pretreatmentC/
           sd(functionModel3Weeks$data$rawSlope),
  )

fullEffectThreeWeeksBetas <- scaledBetas %>%
  mutate("Effects" = "Direct Effects") %>%
  mutate(
    "Body Mass" = b_rawSlope_mass,
    "Tarsus Length" = b_rawSlope_tarsus,
    "Cold Rearing\n(10°C)" = b_rawSlope_pretreatmentA,
    "Warm Rearing\n(30°C)" = b_rawSlope_pretreatmentC
  ) %>%
  select(Effects, `Body Mass`, `Tarsus Length`,
         `Cold Rearing\n(10°C)`,
         `Warm Rearing\n(30°C)`) %>%
  rbind(
    .,
    scaledBetas %>%
      mutate("Effects" = "Indirect Effects") %>%
      mutate(
        "Body Mass" = b_tarsus_mass *
          b_rawSlope_tarsus,
        "Tarsus Length" = NA,
        "Cold Rearing\n(10°C)" =
          b_tarsus_pretreatmentA * b_rawSlope_tarsus +
          b_mass_pretreatmentA * b_rawSlope_mass +
          b_mass_pretreatmentA * b_tarsus_mass *
          b_rawSlope_tarsus,
        "Warm Rearing\n(30°C)" =
          b_tarsus_pretreatmentC * b_rawSlope_tarsus +
          b_mass_pretreatmentC * b_rawSlope_mass +
          b_mass_pretreatmentC * b_tarsus_mass *
          b_rawSlope_tarsus,
      ) %>%
      select(Effects, `Body Mass`, `Tarsus Length`,
             `Cold Rearing\n(10°C)`, `Warm Rearing\n(30°C)`)
  ) %>%
  rbind(., scaledBetas %>%
```

```
    mutate("Effects" = "Total Effects") %>%
    mutate(
      "Body Mass" =
        b_rawSlope_mass +
          b_tarsus_mass * b_rawSlope_tarsus,
      "Tarsus Length" =
        b_rawSlope_tarsus,
      "Cold Rearing\n(10°C)" =
        b_rawSlope_pretreatmentA +
          b_mass_pretreatmentA *
        b_rawSlope_mass +
          b_tarsus_pretreatmentA *
        b_rawSlope_tarsus +
          b_mass_pretreatmentA *
        b_tarsus_mass * b_rawSlope_tarsus,
      "Warm Rearing\n(30°C)" =
         b_rawSlope_pretreatmentC +
          b_mass_pretreatmentC *
        b_rawSlope_mass +
          b_tarsus_pretreatmentC *
        b_rawSlope_tarsus +
          b_mass_pretreatmentC *
        b_tarsus_mass * b_rawSlope_tarsus,
    ) %>%
    select(Effects, `Body Mass`, `Tarsus Length`,
             `Cold Rearing\n(10°C)`, `Warm Rearing\n(30°C)`)
    ) %>%
  pivot_longer(c(-Effects), names_to = "var", values_to = "values") %>%
  mutate(var = factor(var,
                      levels = c("Tarsus Length",
                                 "Body Mass",
                                 "Warm Rearing\n(30°C)",
                                 "Cold Rearing\n(10°C)")))

fullEffectThreeWeeks <- fullEffectThreeWeeksBetas %>%
  ggplot(aes(x = values, y = var, fill = var)) +
  facet_wrap(~Effects) +
  stat_halfeye(normalize = "xy", colour = "black", alpha = 0.7) +
  geom_vline(xintercept = 0, linetype = "dashed",
             colour = "black") +
  xlab(".                   Effect on Metabolic Slope (standard deviations)") +
  scale_fill_manual(values = c("black", "grey50", "#CD5C5C", "#7BB4E3")) +
  theme_classic() +
  theme(
    legend.position = "none", axis.title.y = element_blank(),
    axis.text.y = element_text(size = 11, colour = "black",
                               family = "Noto Sans"),
    axis.title.x = element_text(family = "Noto Sans", hjust = -0.005),
    axis.text.x = element_text(family = "Noto Sans")
  )

showtext_auto(enable = TRUE)

fullEffectThreeWeeks
```

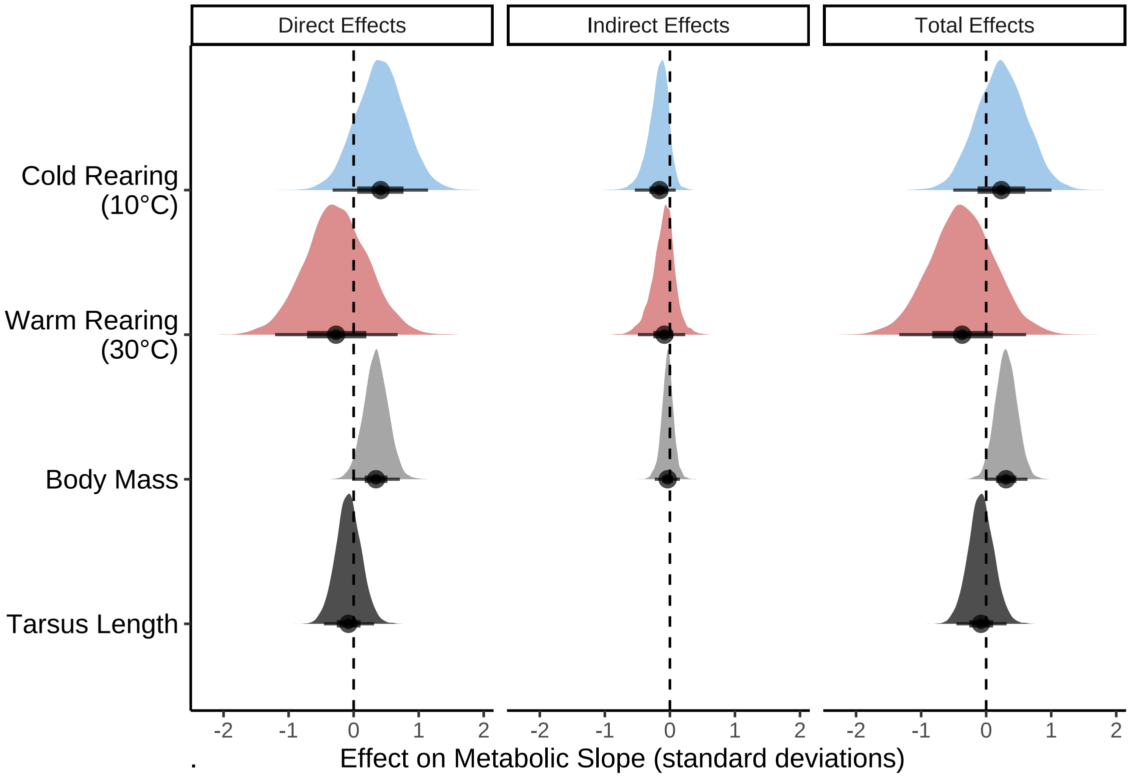


***Figure 31:*** *Effects of morphology (body mass, g, and tarsus length, mm) on metabolic slope (in standard deviations) at ambient temperatures below thermoneutrality (<30°C) in three week old Japanese quail. Distributions represent those derived from posteriors of a Bayesian path analysis.*

```
ggsave("../plots/indirectEffectsOnResistance3Weeks_3ScaledFold.pdf",
       fullEffectThreeWeeks,
  dpi = 800, width = 8, height = 6
)
showtext_auto(enable = FALSE)

# Summarising results

caption <- paste0("Direct, indirect, and total effects of morphology and ",
                  "rearing temperature on metabolic slope ",
                  "(fold metabolism at thermoneutrality/ ",
                  "°C) ",
                  "of three week old Japanese quail. Effects are derived from ",
                  "a Bayesian path analysis and represent those predicted for a ",
                  "change in one standard deviation (or categorical level) of ",
                  "a given predictor on the standard deviation of metabolic ",
                  "slopes. Estimates indicate posterior medians and credible ",
                  "intervels (CIs) indicate quantile intervals."
)

week3ResultsScaled <-
  fullEffectThreeWeeksBetas %>%
  filter(!is.na(values) & !is.nan(values)) %>%
  group_by(Effects, var) %>%
  summarise(
    "Estimate" = median(values),
    "50\\% CIs" = paste0(
      "[",
      round(
        quantile(values, probs = 0.1, type = 8),
        digits = 4
```

```
      ),
      ", ",
      round(
        quantile(values, probs = 0.9, type = 8),
        digits = 4
      ),
      "]"
    ),
    "95\\% CIs" = paste0(
      "[",
      round(
        quantile(values, probs = 0.025, type = 8),
        digits = 4
      ),
      ", ",
      round(
        quantile(values, probs = 0.975, type = 8),
        digits = 4
      ),
      "]"
    )
  ) %>%
  mutate("Effects" = gsub("[[:space:]].*", "", Effects)) %>%
  select(
    "Predictor" = "var", "Effect Level" = "Effects",
    Estimate, `50\\% CIs`, `95\\% CIs`
  ) %>%
  arrange(Predictor, `Effect Level`) %>%
  kbl(.,
    longtable = T, booktabs = T, format = "latex", escape = FALSE,
    caption = caption
  ) %>%
  column_spec(column = c(1:2), width = "2.2cm") %>%
  column_spec(column = c(3:10), width = "1.9cm") %>%
  kable_styling(latex_options = "striped")

week3ResultsScaled
```

***Table 14:*** *Direct, indirect, and total effects of morphology and rearing temperature on metabolic slope (fold metabolism at thermoneutrality/ °C) of three week old Japanese quail. Effects are derived from a Bayesian path analysis and represent those predicted for a change in one standard deviation (or categorical level) of a given predictor on the standard deviation of metabolic slopes. Estimates indicate posterior medians and credible intervels (CIs) indicate quantile intervals.*

| Predictor | Effect Level | Estimate | 50% CIs | 95% CIs |
|---|---|---|---|---|
| Tarsus Length | Direct | -0.0801793 | [-0.3233, 0.17] | [-0.4569, 0.3147] |
| Tarsus Length | Total | -0.0801793 | [-0.3233, 0.17] | [-0.4569, 0.3147] |
| Body Mass | Direct | 0.3438260 | [0.1069, 0.5813] | [-0.0241, 0.711] |
| Body Mass | Indirect | -0.0339407 | [-0.1507, 0.0764] | [-0.2332, 0.1531] |
| Body Mass | Total | 0.3059695 | [0.0986, 0.5177] | [-0.0115, 0.6359] |
| Warm Rearing (30°C) | Direct | -0.2703813 | [-0.8753, 0.3458] | [-1.2094, 0.6784] |
| Warm Rearing (30°C) | Indirect | -0.0843165 | [-0.3247, 0.0997] | [-0.4929, 0.2358] |
| Warm Rearing (30°C) | Total | -0.3687310 | [-0.9937, 0.2615] | [-1.3379, 0.6143] |
| Cold Rearing (10°C) | Direct | 0.4147416 | [-0.0657, 0.8889] | [-0.3237, 1.1447] |
| Cold Rearing (10°C) | Indirect | -0.1590820 | [-0.3804, -0.0022] | [-0.5402, 0.089] |

| | | | | |
|---|---|---|---|---|
| Cold Rearing (10°C) | Total | 0.2330359 | [-0.2582, 0.7322] | [-0.5066, 1.0048] |

```
#save_kable(week3ResultsScaled, "../tables/threeWeekResults_3Scaled.html")
```

To alleviate concerns over possible collinearity between body mass and tarsus length in our previous model, our path analysis is re-run but while relativising tarsus length by body mass prior to use in our models. This was achieved by first regressing tarsus length against mean-centred body mass (priors: intercept = $\mathcal{N}$[30, 2.5]; body mass = $\mathcal{SN}$[0, 0.25, 5]), then extracting ordinary residuals from the regression and replacing true tarsus length values with these values in our path analysis; body mass is then removed as a predictor of tarsus length in our analysis. Priors for all model parameters remained the same as described for our previous path analysis (with variables unscaled).

```
# Note that our prior for the intercept of tarsus length (here, the mean,
# since body mass is mean-centred) is informed from Gebhardt-Henrich
# & Mark (1993: Gen Res Camb) and van der Ziel & Visser
# (2001; Physiol Biochem Zool).

functionModel3WeeksResidual <- brm(
  data = slopeData %>%
    filter(week == "3") %>%
    mutate(
      massCentred = mass - mean(mass, na.rm = T),
      pretreatment = ifelse(pretreatment == "cold", "A",
        ifelse(pretreatment == "neutral", "B", "C")
      )
    ) %>%
    mutate(pretreatment = factor(pretreatment, levels = c("B", "A", "C"))) %>%
    select(-mass) %>%
    rename("mass" = massCentred) %>%
    drop_na() %>%
    merge(., data %>%
      select(ring, "batch" = exp) %>%
      distinct(),
    by = "ring", all.x = TRUE
    ) %>%
    mutate(
      "residualTarsus" =
        residuals(
          brm(tarsusLengthMean ~ mass,
            data = .,
            prior = c(
              set_prior("skew_normal(0, 0.25, 5)",
                class = "b"
              ),
              set_prior("normal(30, 2.5)",
                class = "Intercept"
              )
            ),
            family = "gaussian",
            iter = 50000, warmup = 10000, cores = 1,
            chains = 4, thin = 20,
            silent = TRUE, refresh = 0
          )
        )[, "Estimate"]
    ),
  family = "gaussian",
  bf(mass ~ pretreatment + (1 | batch)) +
    bf(residualTarsus ~ pretreatment + (1 | batch)) +
    bf(rawSlope ~ residualTarsus + mass + pretreatment + (1 | batch)) +
    set_rescor(rescor = FALSE),
  prior = c(
    set_prior("normal(0, 5)",
      class = "Intercept",
```

```
      resp = "mass"
    ),
    set_prior("normal(0, 15)",
      class = "b",
      coef = "pretreatmentA", resp = "mass"
    ),
    set_prior("normal(0, 15)",
      class = "b",
      coef = "pretreatmentC", resp = "mass"
    ),
    set_prior("exponential(2.5)",
      class = "sd",
      group = "batch", resp = "mass"
    ),
    set_prior("exponential(0.15)",
      class = "sigma",
      resp = "mass"
    ),
    set_prior("normal(0, 2.5)",
      class = "Intercept",
      resp = "residualTarsus"
    ),
    set_prior("normal(0, 2.5)",
      class = "b",
      coef = "pretreatmentA", resp = "residualTarsus"
    ),
    set_prior("normal(0, 2.5)",
      class = "b",
      coef = "pretreatmentC", resp = "residualTarsus"
    ),
    set_prior("exponential(2)",
      class = "sd",
      group = "batch", resp = "residualTarsus"
    ),
    set_prior("exponential(1)",
      class = "sigma",
      resp = "residualTarsus"
    ),
    set_prior("normal(0, 0.01)",
      class = "Intercept",
      resp = "rawSlope"
    ),
    set_prior("normal(0, 0.025)",
      class = "b",
      coef = "pretreatmentA", resp = "rawSlope"
    ),
    set_prior("normal(0, 0.025)",
      class = "b",
      coef = "pretreatmentC", resp = "rawSlope"
    ),
    set_prior("normal(0, 0.001)",
      class = "b",
      coef = "mass", resp = "rawSlope"
    ),
    set_prior("normal(0, 0.0025)",
      class = "b",
      coef = "residualTarsus", resp = "rawSlope"
    ),
    set_prior("exponential(50)",
      class = "sd",
      group = "batch", resp = "rawSlope"
    ),
    set_prior("exponential(10)",
      class = "sigma",
      resp = "rawSlope"
    )
  ),
```

```
  iter = 50000, warmup = 10000, cores = 4, chains = 4, thin = 20,
  control = list(adapt_delta = .96),
  silent = TRUE, refresh = 0,
  file = "./models/threeWeekFunctionModelResidual.Rds"
)

# Running quick diagnostics. Note that these are not fulsome,
# given the supplemental nature of the analysis.

chainCheck(functionModel3WeeksResidual)
```

```
## Rhat range: 1 - 1
## Neff/N range: 0.906 - 1.008
```

```
pp_check2(functionModel3WeeksResidual,
  resp = "rawSlope",
  xlab = "Metabolic Slope\n(fold Metabolism at Thermoneutrality)"
  )
```

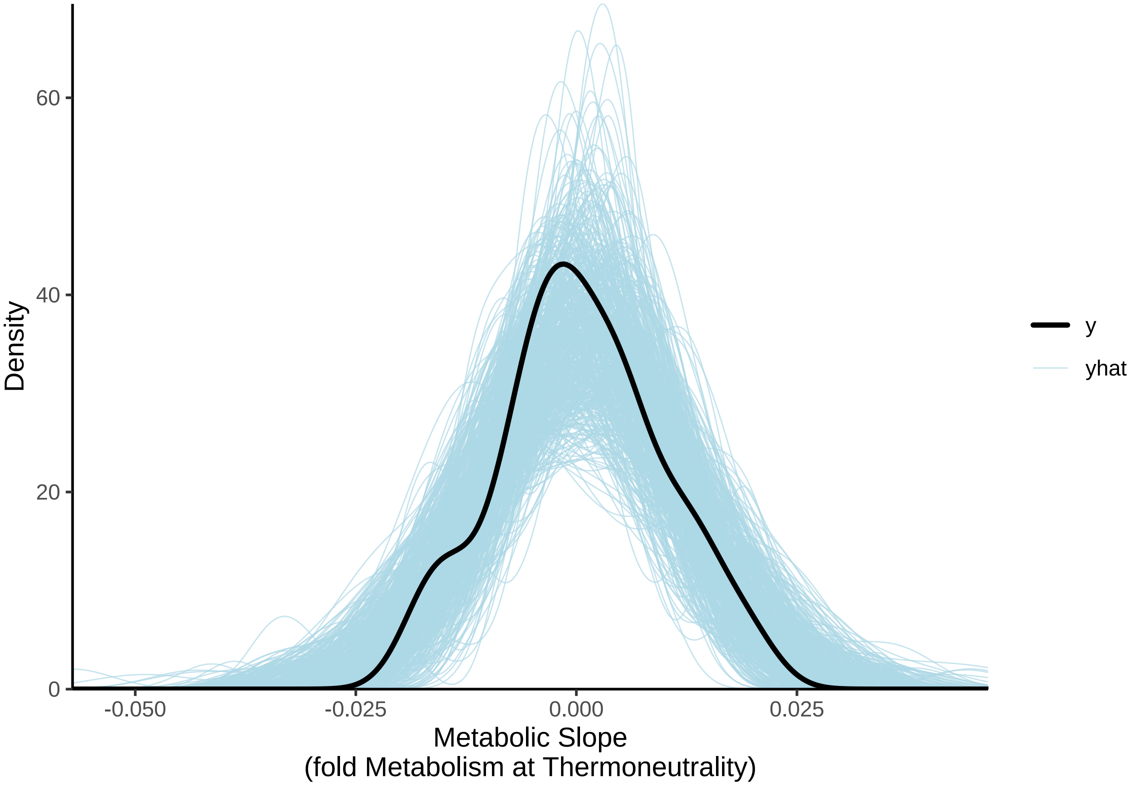


***Figure 32:*** *Posterior predictive check for a Bayesian path analysis predicting metabolic slope (fold metabolism at thermoneutrality/°C) of three week old Japanese quail by morphometry. Light blue lines represent densities of resting metabolism values as drawn from model posteriors. The dark blue line represent the true density of resting metabolism values. Clear overlap between the dark blue and light blue lines indicates strong model fit. Morphometry measures include body mass (g) and tarsus length residuals.*

Density of posterior predictions overlays clearly with true density of metabolic slope values. Next, we visualise the spread of residuals, autocorrelation factors in chains (per predictor) and posterior densities (again, per predictor).

```
lbs <- c("Mass ~ Cold Rearing")
names(lbs) <- c("b_mass_pretreatmentA")
p1 <- mcmc_acf(functionModel3WeeksResidual,
```

```
  pars = "b_mass_pretreatmentA", lags = 10,
  facet_args = list(labeller = labeller(Parameter = lbs))
) +
  theme_classic()

lbs <- c("Mass ~ Warm Rearing")
names(lbs) <- c("b_mass_pretreatmentC")
p2 <- mcmc_acf(functionModel3WeeksResidual,
  pars = "b_mass_pretreatmentC", lags = 10,
  facet_args = list(labeller = labeller(Parameter = lbs))
) +
  theme_classic()

lbs <- c("Tarsus ~ Cold Rearing")
names(lbs) <- c("b_residualTarsus_pretreatmentA")
p3 <- mcmc_acf(functionModel3WeeksResidual,
  pars = "b_residualTarsus_pretreatmentA", lags = 10,
  facet_args = list(labeller = labeller(Parameter = lbs))
) +
  theme_classic()

lbs <- c("Tarsus ~ Warm Rearing")
names(lbs) <- c("b_residualTarsus_pretreatmentC")
p4 <- mcmc_acf(functionModel3WeeksResidual,
  pars = "b_residualTarsus_pretreatmentC", lags = 10,
  facet_args = list(labeller = labeller(Parameter = lbs))
) +
  theme_classic()

lbs <- c("Slope ~ Tarsus")
names(lbs) <- c("b_rawSlope_residualTarsus")
p5 <- mcmc_acf(functionModel3WeeksResidual,
  pars = "b_rawSlope_residualTarsus", lags = 10,
  facet_args = list(labeller = labeller(Parameter = lbs))
) +
  theme_classic()

lbs <- c("Slope ~ Mass")
names(lbs) <- c("b_rawSlope_mass")
p6 <- mcmc_acf(functionModel3WeeksResidual,
  pars = "b_rawSlope_mass", lags = 10,
  facet_args = list(labeller = labeller(Parameter = lbs))
) +
  theme_classic()

lbs <- c("Slope ~\nCold Rearing")
names(lbs) <- c("b_rawSlope_pretreatmentA")
p7 <- mcmc_acf(functionModel3WeeksResidual,
  pars = "b_rawSlope_pretreatmentA", lags = 10,
  facet_args = list(labeller = labeller(Parameter = lbs))
) +
  theme_classic()

lbs <- c("Slope ~\nWarm Rearing")
names(lbs) <- c("b_rawSlope_pretreatmentC")
p8 <- mcmc_acf(functionModel3WeeksResidual,
  pars = "b_rawSlope_pretreatmentC", lags = 10,
  facet_args = list(labeller = labeller(Parameter = lbs))
) +
  theme_classic()

p9 <- functionModel3WeeksResidual$data %>%
  mutate("Res" =
           residuals(functionModel3WeeksResidual,
                     resp = "rawSlope",
                     robust = TRUE)[, "Estimate"]) %>%
  ggplot(aes(sample = Res)) +
```

```
  stat_qq(colour = "grey50") +
  stat_qq_line() +
  xlab("Theoretical") +
  ylab("Sample") +
  theme_classic()

p10 <- functionModel3WeeksResidual$data %>%
  mutate("Fit" =
           fitted(functionModel3WeeksResidual,
                  resp = "rawSlope",
                     robust = TRUE)[, "Estimate"]) %>%
  ggplot(aes(x = Fit, y = rawSlope)) +
  geom_point(pch = 21, colour = "black", size = 2, fill = "grey90") +
  geom_smooth(method = "lm", linetype = "dashed",
              colour = "black", se = FALSE) +
  xlab("Fitted Values") +
  ylab("True Metabolic Slope") +
  theme_classic()

(p1 + p2 + p3) /
  (p4 + p5 + p6) /
  (p7 + p8 + p9) /
  (p10)
```

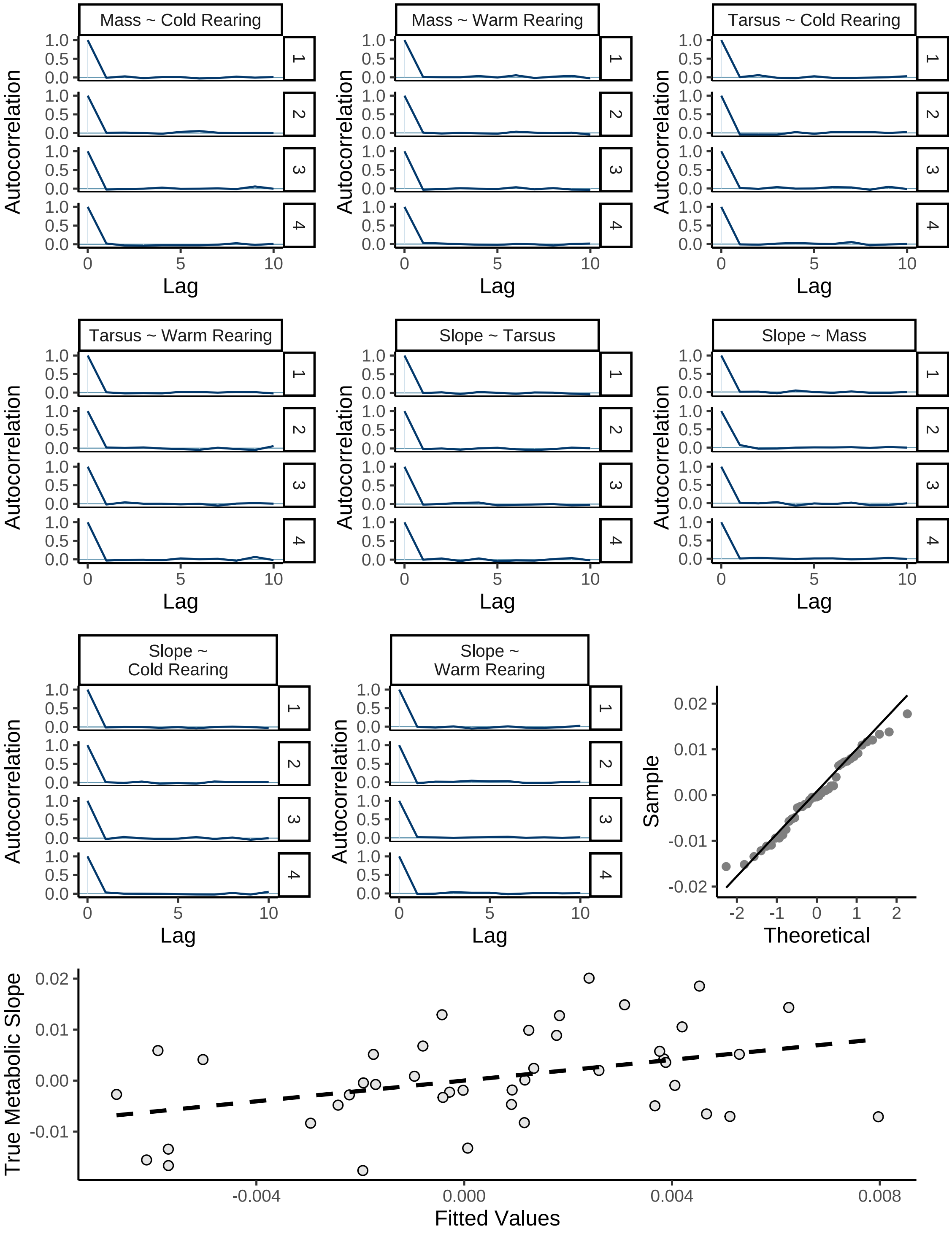


***Figure 33:*** *Diagnostics for a Bayesian path analysis predicting metabolic slope in the cold (fold metabolism at thermoneutrality/°C) of three week old Japanese quail by morphometry (here, body mass [g] and residual tarsus length [mm]). Autocorrelation plots display autocorrelation between adjacent Hamiltonian Monte Carlo chain draws, divided by chain. QQ-plot displays sample ordinary residuals by theoretical residual quantiles, and fitted values are displayed against true, mean-centred metabolic slope values.*

```r
# Plotting coefficients

as.data.frame(functionModel3WeeksResidual) %>%
    select(
      "Intercept" = b_rawSlope_Intercept,
      "Residual Tarsus (mm)" =
        b_rawSlope_residualTarsus,
      "Body Mass (g)" = b_rawSlope_mass,
      "Cold Rearing\n(10°C)" =
        b_rawSlope_pretreatmentA,
      "Warm Rearing\n(30°C)" =
        b_rawSlope_pretreatmentC,
      "Sigma" = sigma_rawSlope
    ) %>%
    pivot_longer(everything(), names_to = "Par",
                 values_to = "Coefs") %>%
    ggplot(aes(x = Coefs)) +
    facet_wrap(~Par, scales = "free", ncol = 2) +
    geom_density(colour = "black", alpha = 0.5,
                 fill = "grey70") +
    geom_vline(xintercept = 0, linetype = "dashed",
               colour = "black") +
    ylab("Density") +
    theme_classic() +
    theme(axis.title.x = element_blank())
```

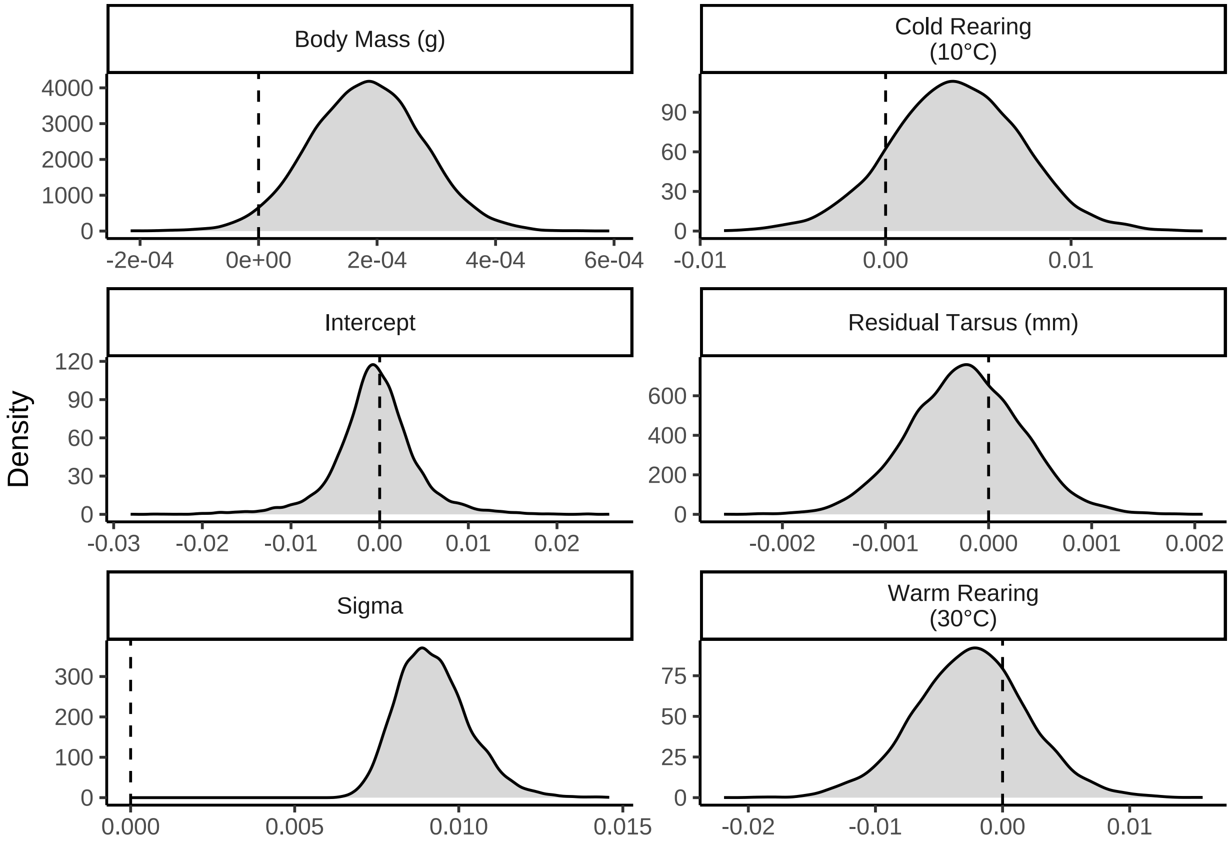


***Figure 34:*** *Posterior densities for model coefficients derived from a Bayesian path analysis predicting metabolic slope (fold metabolism at thermoneutrality/°C) of three week old Japanese quail in the cold (10°C - 30°C). Vertical dashed lines indicate 0.*

```
ggarrange(
  as.data.frame(functionModel3WeeksResidual) %>%
    select(
      "Intercept" = b_residualTarsus_Intercept,
      "Cold Rearing" = b_mass_pretreatmentA,
      "Warm Rearing" = b_mass_pretreatmentC,
      "Sigma" = sigma_mass
    ) %>%
    pivot_longer(everything(), names_to = "Par", values_to = "Coefs") %>%
    ggplot(aes(x = Coefs)) +
    facet_wrap(~Par, scales = "free") +
    geom_density(colour = "black", alpha = 0.5, fill = "grey70") +
    geom_vline(xintercept = 0, linetype = "dashed", colour = "black") +
    ylab("Density") +
    theme_classic() +
    theme(axis.title.x = element_blank()) +
    ggtitle("Body Mass (g)"),

  as.data.frame(functionModel3WeeksResidual) %>%
    select(
      "Intercept" = b_residualTarsus_Intercept,
      "Cold Rearing" = b_residualTarsus_pretreatmentA,
      "Warm Rearing" = b_residualTarsus_pretreatmentC,
      "Sigma" = sigma_residualTarsus
    ) %>%
    pivot_longer(everything(), names_to = "Par",
                 values_to = "Coefs") %>%
    ggplot(aes(x = Coefs)) +
    facet_wrap(~Par, scales = "free") +
    geom_density(colour = "black", alpha = 0.5, fill = "grey70") +
    geom_vline(xintercept = 0, linetype = "dashed", colour = "black") +
    ylab("Density") +
    theme_classic() +
    theme(axis.title.x = element_blank()) +
    ggtitle("Residual Tarsus Length (mm)"),
  ncol = 1
)
```

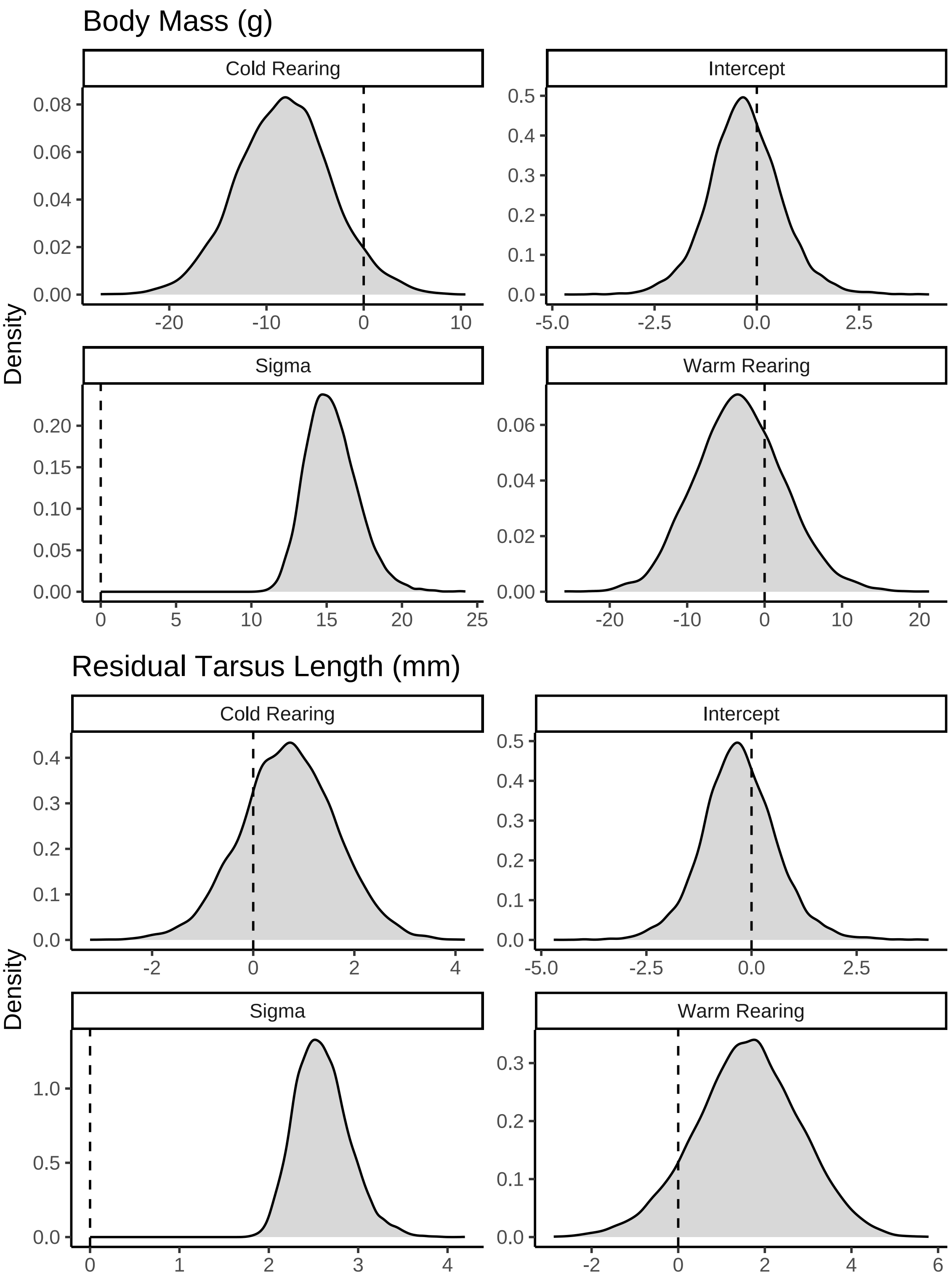


***Figure 35:*** *Continued. Posterior densities for model coefficients derived from a Bayesian path analysis predicting metabolic slopes (fold metabolism at thermoneutrality/°C) of three week old Japanese quail in the cold (10°C - 30°C). Densities are split by their respective response values (indicated with titles). Vertical dashed lines indicate 0.*

Some skewing in predictors is evident. As such, we proceed by using medians as our metric of central tendency for posterior summaries.

```
caption <- paste0("Results of a Bayesian path analysis ",
                  "predicting metabolic slopes (fold metabolism ",
                  "at thermoneutrality) of three week old Japanese quail by ",
                  "morphometry and thermal history. Tarsus length is ",
                  "relativised by body mass (tarsus ",
                  "length residuals) as estimated from a Bayesian ",
                  "linear model. Metabolic slopes represent those in response ",
                  "to cold (10°C - 30°C). Estimates are posterior medians, ",
                  "credible intervals are quantile intervals, and BF indicates ",
                  "Bayes Factors for each model parameter."
)

functionModel3WeeksResidualTable <-
  as.data.frame(functionModel3WeeksResidual) %>%
  summarise_all(., .funs = median) %>%
  pivot_longer(everything(),
    names_to = "Parameter",
    values_to = "Estimate"
  ) %>%
  merge(., quantileCIs(functionModel3WeeksResidual, cis = c(50, 95)),
    by = "Parameter", all.x = TRUE
  ) %>%
  filter(grepl("b_|sd_", Parameter)) %>%
  rowwise() %>%
  mutate("BF" = ifelse(Estimate < 0,
    (2 * mean(as.data.frame(
      functionModel3WeeksResidual
    )[, Parameter] <= 0)) /
      (2 * mean(as.data.frame(
        functionModel3WeeksResidual
      )[, Parameter] >= 0)),
    (2 * mean(as.data.frame(
      functionModel3WeeksResidual
    )[, Parameter] >= 0)) /
      (2 * mean(as.data.frame(
        functionModel3WeeksResidual
      )[, Parameter] <= 0))
  )) %>%
  ungroup() %>%
  mutate(
    "Estimate" = round(Estimate, digits = 4),
    "BF" = round(BF, digits = 4),
    "N" = nrow(functionModel3WeeksResidual$data)
  ) %>%
  mutate("Parameter" = ifelse(grepl("b_", Parameter),
                              gsub("b_", "", Parameter),
                              gsub("Intercept", "batch",
                                   gsub(".*__", "", Parameter)
                                   )
                              )
         ) %>%
  mutate(
    "Response" = gsub("_.*", "", Parameter),
    "Parameter" = gsub(".*_", "", Parameter)
  ) %>%
  merge(., tribble(
    ~Response, ~response, ~level,
    "mass", "Body Mass (g)", "A",
    "rawSlope",
    "Metabolic Slope", "D",
    "residualTarsus", "Residuial Tarsus Length (mm)",
    "B"
  ),
  by = "Response"
```

```
  ) %>%
  merge(., tribble(
    ~Parameter, ~parameter, ~number,
    "Intercept", "Intercept", "1",
    "mass", "Body Mass (g)", "4",
    "residualTarsus", "Residual Tarsus Length (mm)", "5",
    "pretreatmentA", "Cold Rearing", "2",
    "pretreatmentC", "Warm Rearing", "3",
    "batch", "Egg Batch [mu]", "6"
  ),
  by = "Parameter"
  ) %>%
  mutate(
    `50\\% CI` = paste0("(", paste(
      round(Low_CI_50, digits = 4),
      round(High_CI_50, digits = 4),
      sep = ", "
    ), ")"),
    `95\\% CI` = paste0("(", paste(
      round(Low_CI_95, digits = 4),
      round(High_CI_95, digits = 4),
      sep = ", "
    ), ")")
  ) %>%
  select(-c(Low_CI_50, High_CI_50, Low_CI_95, High_CI_95)) %>%
  select(
    "Response" = "response", "Parameter" = "parameter", N,
    Estimate, `50\\% CI`, `95\\% CI`, BF, level, number
  ) %>%
  arrange(level, number) %>%
  select(-c(level, number)) %>%
  kbl(.,
    longtable = T, booktabs = T, format = "latex", escape = FALSE,
    caption = caption
  ) %>%
  column_spec(column = c(1:2), width = "2.2cm") %>%
  column_spec(column = c(3:10), width = "1.9cm") %>%
  kable_styling(latex_options = "striped")

functionModel3WeeksResidualTable
```

***Table 15:*** *Results of a Bayesian path analysis predicting metabolic slopes (fold metabolism at thermoneutrality) of three week old Japanese quail by morphometry and thermal history. Tarsus length is relativised by body mass (tarsus length residuals) as estimated from a Bayesian linear model. Metabolic slopes represent those in response to cold (10°C - 30°C). Estimates are posterior medians, credible intervals are quantile intervals, and BF indicates Bayes Factors for each model parameter.*

| Response | Parameter | N | Estimate | 50% CI | 95% CI | BF |
|---|---|---|---|---|---|---|
| Body Mass (g) | Intercept | 43 | 3.8112 | (1.6088, 5.9647) | (-2.8562, 10.3557) | 6.8585 |
| Body Mass (g) | Cold Rearing | 43 | -8.1479 | (-11.4139, -4.9861) | (-17.7537, 1.7421) | 19.0501 |
| Body Mass (g) | Warm Rearing | 43 | -3.3032 | (-7.0709, 0.6078) | (-14.0957, 8.5473) | 2.5119 |
| Body Mass (g) | Egg Batch [mu] | 43 | 0.2748 | (0.1125, 0.5541) | (0.0095, 1.4623) | Inf |
| Residuial Tarsus Length (mm) | Intercept | 43 | -0.3131 | (-0.8562, 0.2608) | (-2.0417, 1.5684) | 1.8551 |
| Residuial Tarsus Length (mm) | Cold Rearing | 43 | 0.6990 | (0.0872, 1.322) | (-1.1491, 2.5361) | 3.5506 |
| Residuial Tarsus Length (mm) | Warm Rearing | 43 | 1.6149 | (0.8048, 2.4072) | (-0.8943, 3.9088) | 9.4031 |
| Residuial Tarsus Length (mm) | Egg Batch [mu] | 43 | 0.7941 | (0.3953, 1.2557) | (0.0346, 2.3603) | Inf |

| | | | | | | |
|---|---|---|---|---|---|---|
| Metabolic Slope | Intercept | 43 | -0.0005 | (-0.0028, 0.0019) | (-0.0105, 0.0092) | 1.2644 |
| Metabolic Slope | Cold Rearing | 43 | 0.0038 | (0.0015, 0.0062) | (-0.0032, 0.0107) | 6.3801 |
| Metabolic Slope | Warm Rearing | 43 | -0.0024 | (-0.0054, 4e-04) | (-0.0115, 0.0066) | 2.5398 |
| Metabolic Slope | Body Mass (g) | 43 | 0.0002 | (1e-04, 2e-04) | (0, 4e-04) | 38.0244 |
| Metabolic Slope | Residual Tarsus Length (mm) | 43 | -0.0002 | (-6e-04, 1e-04) | (-0.0013, 9e-04) | 1.9795 |
| Metabolic Slope | Egg Batch [mu] | 43 | 0.0039 | (0.0016, 0.0083) | (1e-04, 0.0275) | Inf |

```
#save_kable(resistanceResidual3WeeksTable,
# "../residualTarsusAnalysisResults3Weeks.html")
```

#### Modelling effects at maturity (8 weeks of age)

Here, we repeating the above described path analyses but while only using data obtained from mature Japanese quail (8 weeks of age). Priors used in our previous analyses are retained, except for a broadening of our body mass priors (intercept: $\mathcal{N}[0, 10]$; cold rearing: $\mathcal{N}[0, 25]$; warm rearing: $\mathcal{N}[0, 25]$). Again, priors are tested for suitability via a prior predictive check.

```
functionModel8WeeksPPCheck <-
  brm(
    data = slopeData %>%
      filter(week == "8") %>%
      mutate(
        mass = mass - mean(mass, na.rm = T),
        tarsus = tarsusLengthMean - mean(tarsusLengthMean, na.rm = T),
        pretreatment = ifelse(pretreatment == "cold", "A",
          ifelse(pretreatment == "neutral", "B", "C")
        )
      ) %>%
      mutate(pretreatment = factor(pretreatment, levels = c("B", "A", "C"))) %>%
      drop_na() %>%
      merge(., data %>%
        select(ring, "batch" = exp) %>%
        distinct(),
      by = "ring", all.x = TRUE
      ),
    family = "gaussian",
    bf(mass ~ pretreatment + (1 | batch)) +
      bf(tarsus ~ mass + pretreatment + (1 | batch)) +
      bf(rawSlope ~ tarsus + mass + pretreatment + (1 | batch)) +
      set_rescor(FALSE),
    prior = c(
      set_prior("normal(0, 10)",
        class = "Intercept",
        resp = "mass"
      ),
      set_prior("normal(0, 25)",
        class = "b",
        coef = "pretreatmentA", resp = "mass"
      ),
      set_prior("normal(0, 25)",
        class = "b",
        coef = "pretreatmentC", resp = "mass"
      ),
      set_prior("exponential(2.5)",
        class = "sd",
        group = "batch", resp = "mass"
      ),
      set_prior("exponential(0.15)",
        class = "sigma",
```

```r
        resp = "mass"
      ),
      set_prior("normal(0, 2.5)",
        class = "Intercept",
        resp = "tarsus"
      ),
      set_prior("normal(0, 2.5)",
        class = "b",
        coef = "pretreatmentA", resp = "tarsus"
      ),
      set_prior("normal(0, 2.5)",
        class = "b",
        coef = "pretreatmentC", resp = "tarsus"
      ),
      set_prior("skew_normal(0, 0.25, 5)",
        class = "b",
        coef = "mass", resp = "tarsus"
      ),
      set_prior("exponential(2)",
        class = "sd",
        group = "batch", resp = "tarsus"
      ),
      set_prior("exponential(1)",
        class = "sigma",
        resp = "tarsus"
      ),
      set_prior("normal(0, 0.01)",
        class = "Intercept",
        resp = "rawSlope"
      ),
      set_prior("normal(0, 0.025)",
        class = "b",
        coef = "pretreatmentA", resp = "rawSlope"
      ),
      set_prior("normal(0, 0.025)",
        class = "b",
        coef = "pretreatmentC", resp = "rawSlope"
      ),
      set_prior("normal(0, 0.001)",
        class = "b",
        coef = "mass", resp = "rawSlope"
      ),
      set_prior("normal(0, 0.0025)",
        class = "b",
        coef = "tarsus", resp = "rawSlope"
      ),
      set_prior("exponential(50)",
        class = "sd",
        group = "batch", resp = "rawSlope"
      ),
      set_prior("exponential(10)",
        class = "sigma",
        resp = "rawSlope"
      )
    ),
    iter = 50000, warmup = 10000, cores = 4, chains = 4, thin = 20,
    control = list(adapt_delta = .97, max_treedepth = 14),
    silent = TRUE, refresh = 0,
    sample_prior = "only",
    file = "./models/eightWeekFunctionModelPPCheck.Rds"
  )

# Plotting

p1 <- pp_check2(functionModel8WeeksPPCheck, resp = "mass",
            xlab = "Body Mass (g; Mean-Centred)")
p2 <- pp_check2(functionModel8WeeksPPCheck, resp = "tarsus",
```

```
           xlab = "Tarsus Length (mm; Mean-Centred)")
p3 <- pp_check2(functionModel8WeeksPPCheck,
    resp = "rawSlope",
    xlab =
    "Metabolic Slope (fold RMR at\nThermoneutrality/°C)"
  )

(p1/p2) | p3
```

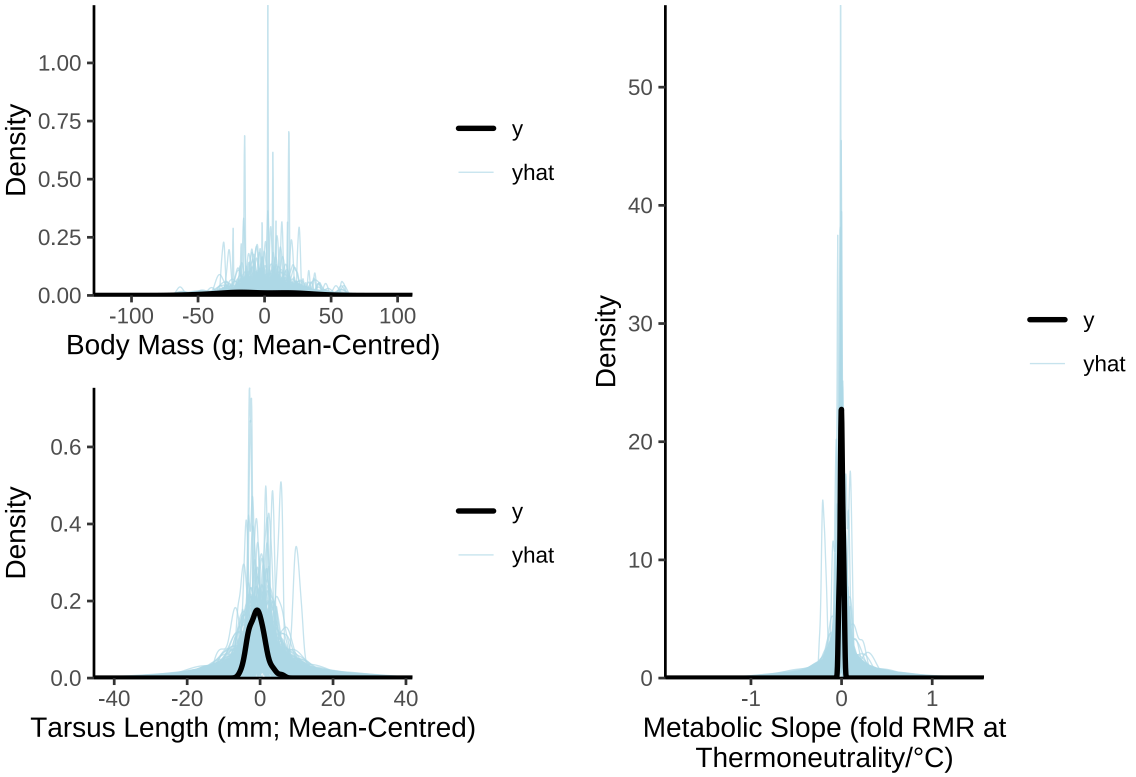


***Figure 36:*** *Prior predictive check for a Bayesian path analysis predicting metabolic slopes (fold metabolism at thermoneutrality/°C) as a direct and indirect function of body mass (g) and tarsus length (mm) in eight week old Japanese quail. Metabolic slopes pertain to those estimated in response to cold (10°C - 30°C). Light blue lines represent densities of resting metabolism values as predicted by model priors alone. The dark blue line represent the true density of resting metabolism values. Clear overlap between the dark blue and light blue lines indicates that priors are suitable.*

```
# Good. Proceeding.
```

```
functionModel8Weeks <-
  brm(
    data = slopeData %>%
      filter(week == "8") %>%
      mutate(
        mass = mass - mean(mass, na.rm = T),
        tarsus = tarsusLengthMean - mean(tarsusLengthMean, na.rm = T),
        pretreatment = ifelse(pretreatment == "cold", "A",
          ifelse(pretreatment == "neutral", "B", "C")
        )
      ) %>%
      mutate(pretreatment = factor(pretreatment, levels = c("B", "A", "C"))) %>%
      drop_na() %>%
      merge(., data %>%
        select(ring, "batch" = exp) %>%
```

```
      distinct(),
    by = "ring", all.x = TRUE
    ),
  family = "gaussian",
  bf(mass ~ pretreatment + (1 | batch)) +
    bf(tarsus ~ mass + pretreatment + (1 | batch)) +
    bf(rawSlope ~ tarsus + mass + pretreatment + (1 | batch)) +
    set_rescor(FALSE),
  prior = c(
    set_prior("normal(0, 10)",
      class = "Intercept",
      resp = "mass"
    ),
    set_prior("normal(0, 25)",
      class = "b",
      coef = "pretreatmentA", resp = "mass"
    ),
    set_prior("normal(0, 25)",
      class = "b",
      coef = "pretreatmentC", resp = "mass"
    ),
    set_prior("exponential(2.5)",
      class = "sd",
      group = "batch", resp = "mass"
    ),
    set_prior("exponential(0.15)",
      class = "sigma",
      resp = "mass"
    ),
    set_prior("normal(0, 2.5)",
      class = "Intercept",
      resp = "tarsus"
    ),
    set_prior("normal(0, 2.5)",
      class = "b",
      coef = "pretreatmentA", resp = "tarsus"
    ),
    set_prior("normal(0, 2.5)",
      class = "b",
      coef = "pretreatmentC", resp = "tarsus"
    ),
    set_prior("skew_normal(0, 0.25, 5)",
      class = "b",
      coef = "mass", resp = "tarsus"
    ),
    set_prior("exponential(2)",
      class = "sd",
      group = "batch", resp = "tarsus"
    ),
    set_prior("exponential(1)",
      class = "sigma",
      resp = "tarsus"
    ),
    set_prior("normal(0, 0.01)",
      class = "Intercept",
      resp = "rawSlope"
    ),
    set_prior("normal(0, 0.025)",
      class = "b",
      coef = "pretreatmentA", resp = "rawSlope"
    ),
    set_prior("normal(0, 0.025)",
      class = "b",
      coef = "pretreatmentC", resp = "rawSlope"
    ),
    set_prior("normal(0, 0.001)",
      class = "b",
```

```
        coef = "mass", resp = "rawSlope"
      ),
      set_prior("normal(0, 0.0025)",
        class = "b",
        coef = "tarsus", resp = "rawSlope"
      ),
      set_prior("exponential(50)",
        class = "sd",
        group = "batch", resp = "rawSlope"
      ),
      set_prior("exponential(10)",
        class = "sigma",
        resp = "rawSlope"
      )
    ),
    iter = 50000, warmup = 10000, cores = 4, chains = 4, thin = 20,
    control = list(adapt_delta = .97, max_treedepth = 14),
    silent = TRUE, refresh = 0,
    file = "./models/eightWeekFunctionModel.Rds"
  )

# Negligible divergences. Checking Rhat values, Neff/N ratios,
# and posterior distribution

chainCheck(functionModel8Weeks)
```

```
## Rhat range: 1 - 1.001
## Neff/N range: 0.855 - 1.009
```

```
pp_check2(functionModel8Weeks, resp = "rawSlope",
     xlab = "Metabolic Slope (fold RMR at Thermoneutrality/°C)"
    )
```

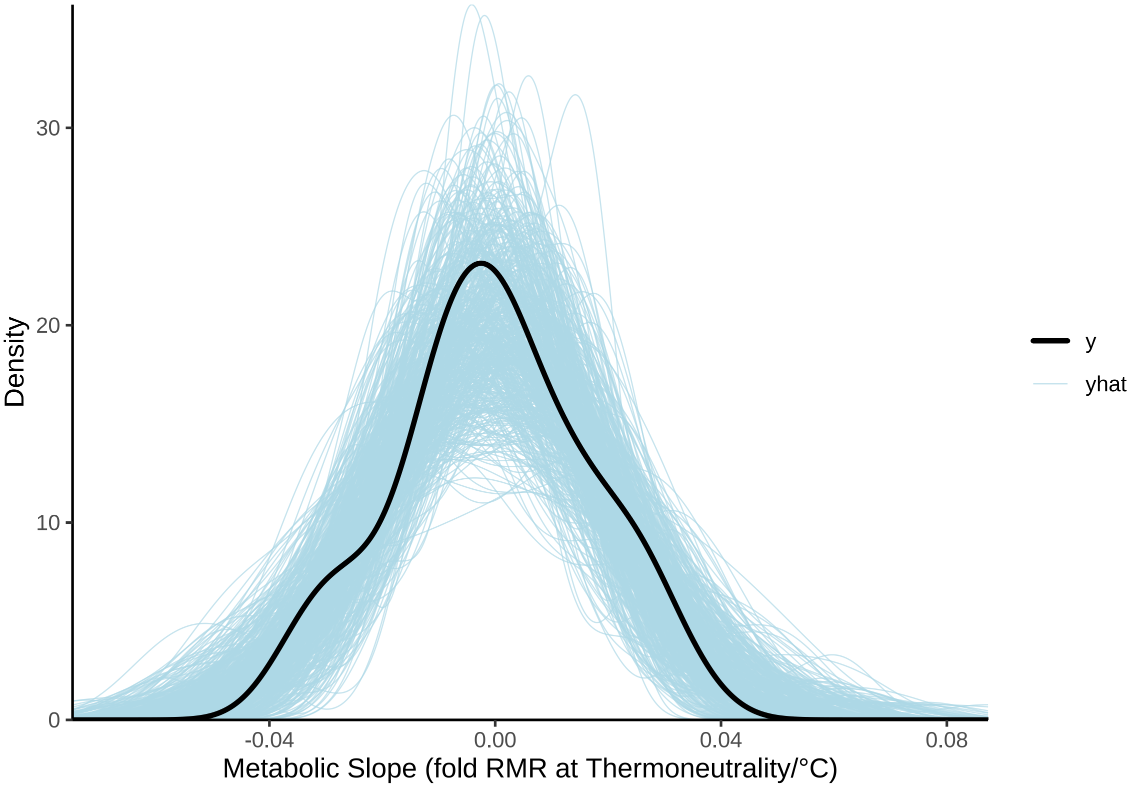


***Figure 37:*** *Posterior predictive check for a Bayesian path analysis predicting metabolic slopes of eight week old Japanese quail, in response to cold (10°C - 30°C), by mean-centred morphometry (body mass and tarsus length). Light blue lines represent densities of resting metabolism values as drawn from model posteriors. The dark blue line represent the true density of resting metabolism values. Clear overlap between the dark blue and light blue lines indicates strong model fit.*

Bayesian $R^2$ values for each element of our path analysis are next estimated.

```
caption = paste0("Estimates of fit for each element of a Bayesian path ",
                 "analysis predicting metabolic slopes (fold metabolism at ",
                 "thermoneutrality/°C) of mature Japanese quail by morphometry ",
                 "and thermal history. Metabolic slopes are measured in ",
                 "response to cold (10°C - 30°C). Credible intervals are ",
                 "quantile intervals. ")

brms::bayes_R2(functionModel8Weeks, ndraws = 1000,
               robust = TRUE) %>%
  as.data.frame() %>%
  rownames_to_column("var") %>%
  merge(., tribble(
    ~var, ~Var,
    "R2mass", "Body Mass (g)",
    "R2tarsus", "Tarsus Length (mm)",
    "R2rawSlope",
      "Metabolic Slope"
  ), by = c("var")) %>%
  mutate(Estimate = round(Estimate, digits = 4),
         Est.Error = round(Est.Error, digits = 4),
         "95\\% CI" = paste0("[", round(Q2.5, digits = 4),
                             ", ", round(Q97.5, digits = 4),
                             "]"
         )
         ) %>%
```

```
  select(
    "Response" = Var, "R\\textsuperscript{2}" = Estimate,
    "Standard Error" = Est.Error,
    `95\\% CI`
  ) %>%
  kbl(.,
    longtable = T, booktabs = T, format = "latex",
    caption = caption, escape = FALSE
  ) %>%
  column_spec(column = c(1:10), width = "2.5cm")  %>%
  kable_styling(latex_options = "striped")
```

***Table 16:*** *Estimates of fit for each element of a Bayesian path analysis predicting metabolic slopes (fold metabolism at thermoneutrality/°C) of mature Japanese quail by morphometry and thermal history. Metabolic slopes are measured in response to cold (10°C - 30°C). Credible intervals are quantile intervals.*

| Response | $R^2$ | Standard Error | 95% CI |
|---|---|---|---|
| Body Mass (g) | 0.0232 | 0.0235 | [9e-04, 0.1087] |
| Metabolic Slope | 0.0964 | 0.0550 | [0.0174, 0.2279] |
| Tarsus Length (mm) | 0.2953 | 0.0847 | [0.1347, 0.4311] |

Residuals and posterior densities are again visualised as before.

```
p1 <- functionModel8Weeks$data %>%
    mutate("Residuals" =
             residuals(functionModel8Weeks,
                       resp = "rawSlope",
                       robust = TRUE)[, "Estimate"]) %>%
    ggplot(aes(x = mass, y = Residuals)) +
    geom_point(size = 2, pch = 21, colour = "black", fill = "grey50") +
    xlab("Relative Body Mass (g)") +
    ylab("Ordinary Residuals") +
    theme_classic()

p2 <- functionModel8Weeks$data %>%
    mutate("Residuals" =
             residuals(functionModel8Weeks,
                       resp = "rawSlope",
                       robust = TRUE)[, "Estimate"]) %>%
    ggplot(aes(x = tarsus, y = Residuals)) +
    geom_point(size = 2, pch = 21, colour = "black", fill = "grey50") +
    xlab("Relative Tarsus\nLength (mm)") +
    ylab("Ordinary Residuals") +
    theme_classic()

p3 <- functionModel8Weeks$data %>%
    mutate("Residuals" =
             residuals(functionModel8Weeks,
                       resp = "rawSlope",
                       robust = TRUE)[, "Estimate"]) %>%
    ggplot(aes(x = factor(pretreatment), y = Residuals)) +
    geom_boxplot(width = 0.5) +
    geom_point(size = 2, pch = 21,
               position = position_jitter(width = 0.25),
               colour = "black", fill = "grey50") +
    xlab("Rearing Conditions") +
    ylab("Ordinary Residuals") +
    theme_classic()

p4 <- functionModel8Weeks$data %>%
    mutate("Residuals" =
             residuals(functionModel8Weeks,
```

```
                           resp = "rawSlope",
                           robust = TRUE)[, "Estimate"]) %>%
    ggplot(aes(sample = Residuals)) +
    stat_qq(colour = "grey50") +
    stat_qq_line() +
    xlab("Theoretical") +
    ylab("Sample") +
    theme_classic()

p5 <- functionModel8Weeks$data %>%
    mutate(
      "Residuals" =
        residuals(functionModel8Weeks,
                  resp = "rawSlope",
                       robust = TRUE)[, "Estimate"],
      "Fitted" =
        fitted(functionModel8Weeks,
               resp = "rawSlope",
                       robust = TRUE)[, "Estimate"]
    ) %>%
    ggplot(aes(x = Fitted, y = Residuals)) +
    geom_point(size = 2, pch = 21, colour = "black", fill = "grey50") +
    xlab("Fitted Values") +
    ylab("Ordinary Residuals") +
    theme_classic()

(p1 + p2 + p3)/(p4 + p5)
```

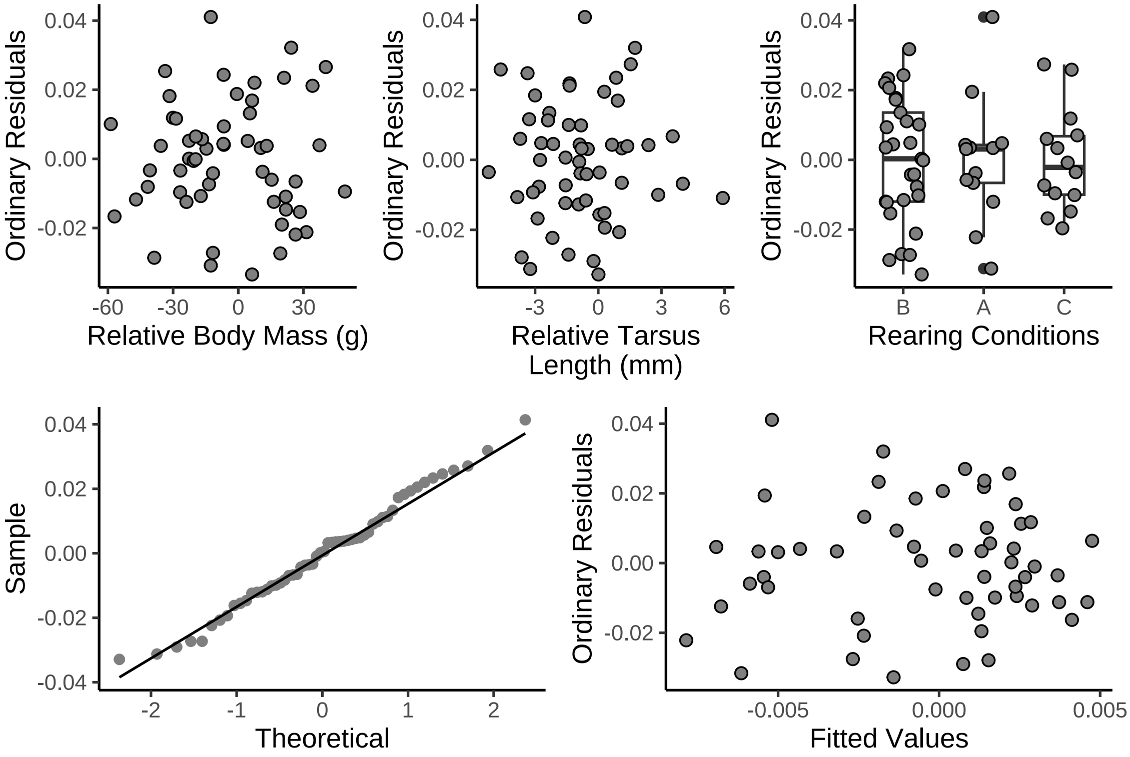


***Figure 38:*** *Ordinary residuals from a Bayesian path analysis predicting metabolic slopes (fold metabolism at thermoneutrality/°C) of mature Japanese quail as a function of body mass, tarsus length, and thermal history (here, rearing temperature). Metabolic slopes are measures in response to cold (10°C - 30°C). Residuals are displayed against model predictors, as a function of theoretical quantiles, and against model fitted values.*

```
# Evenly dispersed. Plotting model coefficients
```

```r
ggarrange(
  as.data.frame(functionModel8Weeks) %>%
    select(
      "Tarsus Length\n(mm)" = b_rawSlope_tarsus,
      "Body Mass (g)" = b_rawSlope_mass,
      "Cold Rearing\n(10°C)" = b_rawSlope_pretreatmentA,
      "Warm Rearing\n(10°C)" = b_rawSlope_pretreatmentC
    ) %>%
    pivot_longer(everything(), names_to = "Par",
                 values_to = "Coefs") %>%
    ggplot(aes(x = Coefs)) +
    facet_wrap(~Par, scales = "free", ncol = 2) +
    geom_density(colour = "black", alpha = 0.5, fill = "grey70") +
    geom_vline(xintercept = 0, linetype = "dashed",
               colour = "black") +
    ylab("Density") +
    scale_x_continuous(n.breaks = 3) +
    theme_classic() +
    theme(axis.title.x = element_blank()) +
    ggtitle("Metabolic Slope"),

  as.data.frame(functionModel8Weeks) %>%
    select(
      "Intercept" = b_mass_Intercept,
      "Cold Rearing\n(10°C)" = b_mass_pretreatmentA,
      "Warm Rearing\n(30°C)" = b_mass_pretreatmentC,
      "Sigma" = sigma_tarsus
    ) %>%
    pivot_longer(everything(), names_to = "Par",
                 values_to = "Coefs") %>%
    ggplot(aes(x = Coefs)) +
    facet_wrap(~Par, scales = "free") +
    geom_density(colour = "black",
                 alpha = 0.5, fill = "grey70") +
    geom_vline(xintercept = 0, linetype = "dashed",
               colour = "black") +
    ylab("Density") +
    theme_classic() +
    theme(axis.title.x = element_blank()) +
    ggtitle("Body Mass (g)")
)
```

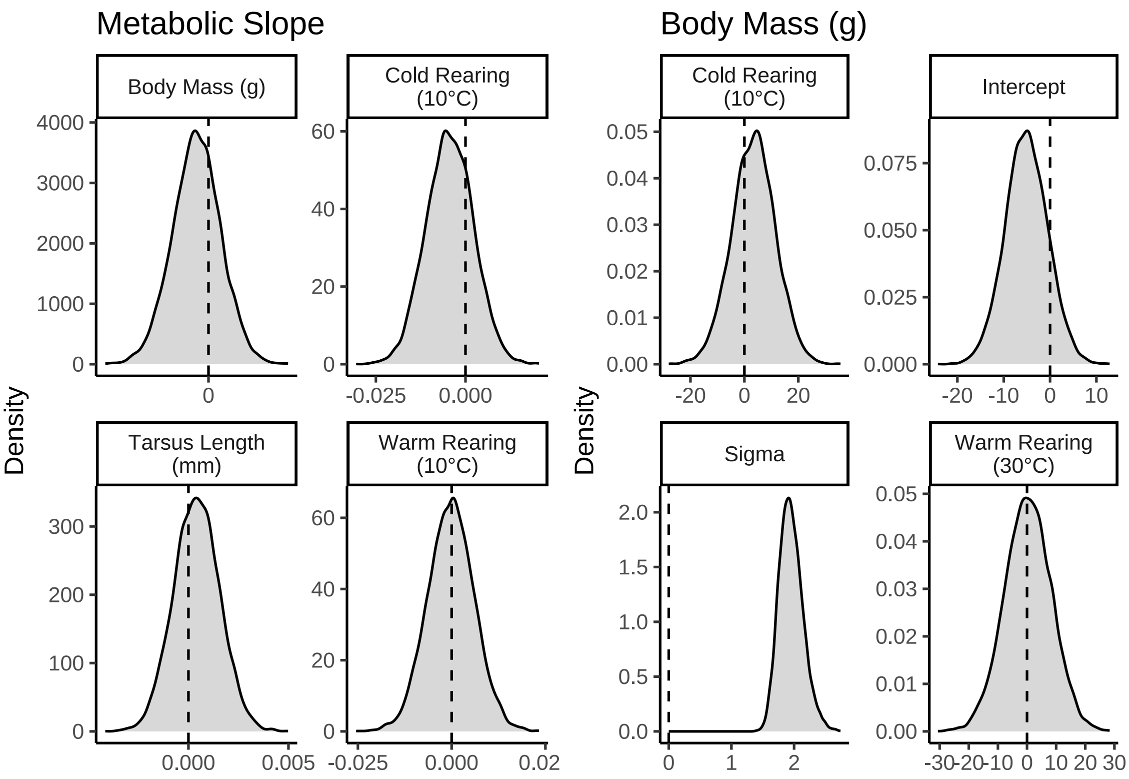


***Figure 39:*** *Coefficient densities (posteriors) from a Bayesian path analysis predicting metabolic slopes (fold metabolism at thermoneutrality/°C) as a function of body mass and tarsus length in eight week old Japanese quail.*

```
as.data.frame(functionModel8Weeks) %>%
    select(
      "Intercept" = b_tarsus_Intercept,
      "Cold Rearing\n(10°C)" = b_tarsus_pretreatmentA,
      "Warm Rearing\n(30°C)" = b_tarsus_pretreatmentC,
      "Sigma" = sigma_tarsus
    ) %>%
    pivot_longer(everything(), names_to = "Par", values_to = "Coefs") %>%
    ggplot(aes(x = Coefs)) +
    facet_wrap(~Par, scales = "free") +
    geom_density(colour = "black", alpha = 0.5, fill = "grey70") +
    geom_vline(xintercept = 0, linetype = "dashed", colour = "black") +
    ylab("Density") +
    theme_classic() +
    theme(axis.title.x = element_blank()) +
    ggtitle("Tarsus Length (mm)")
```

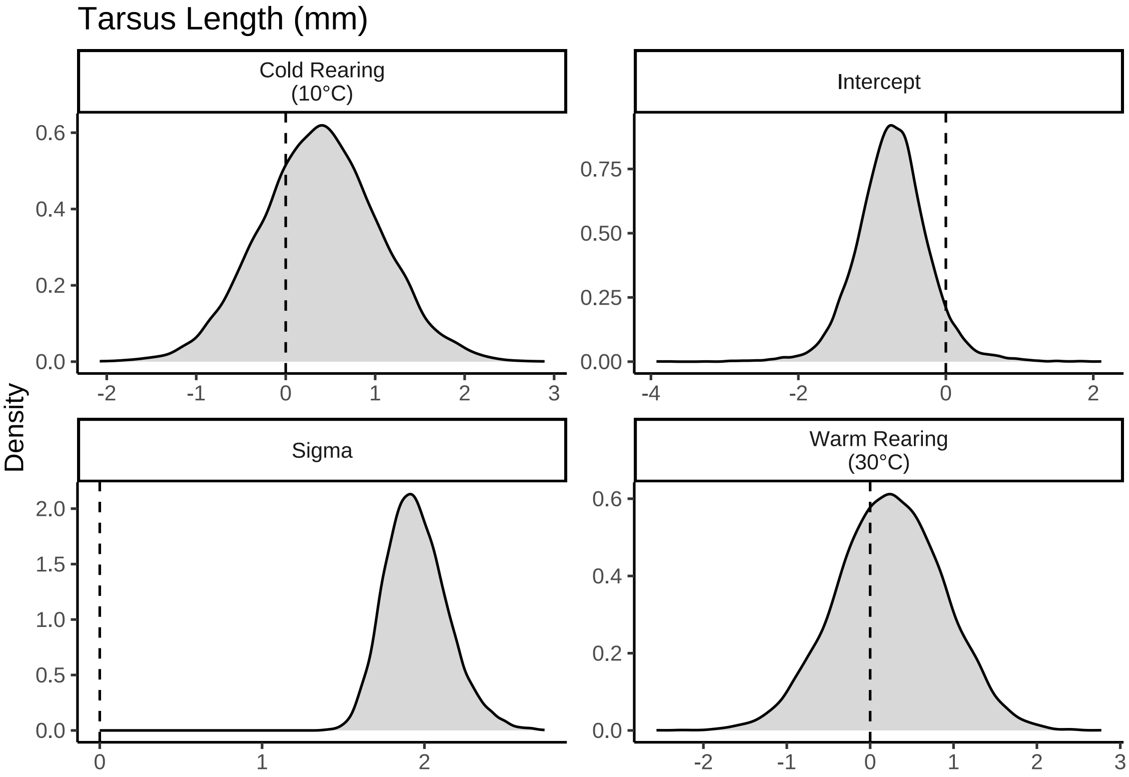


***Figure 40:*** *Coefficient densities (posteriors) from a Bayesian path analysis predicting metabolic slopes (fold metabolism at thermoneutrality/°C) as a function of body mass and tarsus length in eight week old Japanese quail. Continued from previous figure.*

```
# Very little evidence of biological effects. Summarising results

caption <- paste0("Results from a Bayesian path analysis ",
                  "predicting metabolic slope (fold  ",
                  "metabolism at thermoneutrality/{\\circ}C) as a function of ",
                  "morphometry and rearing temperature in ",
                  "eight week old Japanese quail. Metabolic slopes are ",
                  "measured in response to cold (10°C - 30°C). Cold rearing ",
                  "indicates post-hatch rearing at ",
                  "10°C, relative to ",
                  "20°C (intercept), or ",
                  "30°C ('warm rearing). Estimates indicate medians, ",
                  "CIs indicate quantile intervals, ",
                  "and BF indicates Bayes Factors."
)

functionModel8WeeksTable <-
  as.data.frame(functionModel8Weeks) %>%
  summarise_all(., .funs = median) %>%
  pivot_longer(everything(),
    names_to = "Parameter",
    values_to = "Estimate"
  ) %>%
  merge(., quantileCIs(functionModel8Weeks, cis = c(50, 95)),
    by = "Parameter", all.x = TRUE
  ) %>%
  filter(grepl("b_|sd_", Parameter)) %>%
  rowwise() %>%
  mutate("BF" = ifelse(Estimate < 0,
    (2 * mean(as.data.frame(
      functionModel8Weeks
```

```
    )[, Parameter] <= 0)) /
      (2 * mean(as.data.frame(
        functionModel8Weeks
      )[, Parameter] >= 0)),
    (2 * mean(as.data.frame(
      functionModel8Weeks
    )[, Parameter] >= 0)) /
      (2 * mean(as.data.frame(
        functionModel8Weeks
      )[, Parameter] <= 0))
  )) %>%
  ungroup() %>%
  mutate(
    "Estimate" = round(Estimate, digits = 4),
    "BF" = round(BF, digits = 4),
    "N" = nrow(functionModel8Weeks$data)
  ) %>%
  mutate("Parameter" = ifelse(grepl("b_", Parameter),
    gsub("b_", "", Parameter),
    gsub(
      "Intercept", "batch",
      gsub(".*__", "", Parameter)
    )
  )) %>%
  mutate(
    "Response" = gsub("_.*", "", Parameter),
    "Parameter" = gsub(".*_", "", Parameter)
  ) %>%
  merge(., tribble(
    ~Response, ~response, ~level,
    "mass", "Body Mass (g)", "A",
    "rawSlope",
    "Metabolic Slope", "D",
    "tarsus", "Tarsus Length (mm)", "B"
  ),
  by = "Response"
  ) %>%
  merge(., tribble(
    ~Parameter, ~parameter, ~number,
    "Intercept", "Intercept", "1",
    "mass", "Body Mass (g)", "4",
    "tarsus", "Tarsus Length (mm)", "5",
    "pretreatmentA", "Cold Rearing", "2",
    "pretreatmentC", "Warm Rearing", "3",
    "batch", "Egg Batch [mu]", "6"
  ),
  by = "Parameter"
  ) %>%
  mutate(
    `50\\% CI` = paste0("(", paste(
      round(Low_CI_50, digits = 4),
      round(High_CI_50, digits = 4),
      sep = ", "
    ), ")"),
    `95\\% CI` = paste0("(", paste(
      round(Low_CI_95, digits = 4),
      round(High_CI_95, digits = 4),
      sep = ", "
    ), ")")
  ) %>%
  select(-c(Low_CI_50, High_CI_50, Low_CI_95, High_CI_95)) %>%
  select(
    "Response" = "response", "Parameter" = "parameter", N,
    Estimate, `50\\% CI`, `95\\% CI`, BF, level, number
  ) %>%
  arrange(level, number) %>%
  select(-c(level, number)) %>%
```

```
  kbl(.,
    longtable = T, booktabs = T, format = "latex", escape = FALSE,
    caption = caption
  ) %>%
  column_spec(column = 1, width = "2.2cm") %>%
  column_spec(column = 2, width = "2cm") %>%
  column_spec(column = c(3:10), width = "1.9cm") %>%
  kable_styling(latex_options = "striped")

functionModel8WeeksTable
```

***Table 17:*** *Results from a Bayesian path analysis predicting metabolic slope (fold metabolism at thermoneutrality/ C) as a function of morphometry and rearing temperature in eight week old Japanese quail. Metabolic slopes are measured in response to cold (10°C - 30°C). Cold rearing indicates post-hatch rearing at 10°C, relative to 20°C (intercept), or 30°C ('warm rearing). Estimates indicate medians, CIs indicate quantile intervals, and BF indicates Bayes Factors.*

| Response | Parameter | N | Estimate | 50% CI | 95% CI | BF |
|---|---|---|---|---|---|---|
| Body Mass (g) | Intercept | 56 | -5.1243 | (-8.1664, -2.0232) | (-14.1508, 4.0641) | 6.4836 |
| Body Mass (g) | Cold Rearing | 56 | 3.5391 | (-1.9013, 8.8432) | (-12.6338, 19.1795) | 1.9773 |
| Body Mass (g) | Warm Rearing | 56 | 0.3330 | (-5.0674, 5.7114) | (-15.6609, 15.927) | 1.0656 |
| Body Mass (g) | Egg Batch [mu] | 56 | 0.2771 | (0.1196, 0.5628) | (0.01, 1.4359) | Inf |
| Tarsus Length (mm) | Intercept | 56 | -0.7247 | (-1.0174, -0.4386) | (-1.6626, 0.2637) | 15.6320 |
| Tarsus Length (mm) | Cold Rearing | 56 | 0.3803 | (-0.0562, 0.8179) | (-0.8838, 1.6924) | 2.5778 |
| Tarsus Length (mm) | Warm Rearing | 56 | 0.2445 | (-0.1885, 0.6764) | (-1.0234, 1.4799) | 1.8460 |
| Tarsus Length (mm) | Body Mass (g) | 56 | 0.0437 | (0.0371, 0.0504) | (0.0245, 0.0634) | Inf |
| Tarsus Length (mm) | Egg Batch [mu] | 56 | 0.2505 | (0.1045, 0.4851) | (0.0088, 1.3218) | Inf |
| Metabolic Slope | Intercept | 56 | 0.0007 | (-0.0026, 0.0041) | (-0.011, 0.0121) | 1.2605 |
| Metabolic Slope | Cold Rearing | 56 | -0.0044 | (-0.0088, 1e-04) | (-0.017, 0.0084) | 2.9158 |
| Metabolic Slope | Warm Rearing | 56 | -0.0001 | (-0.0043, 0.004) | (-0.0124, 0.0121) | 1.0398 |
| Metabolic Slope | Body Mass (g) | 56 | -0.0001 | (-1e-04, 0) | (-3e-04, 2e-04) | 2.2720 |
| Metabolic Slope | Tarsus Length (mm) | 56 | 0.0004 | (-4e-04, 0.0012) | (-0.0018, 0.0027) | 1.7797 |
| Metabolic Slope | Egg Batch [mu] | 56 | 0.0055 | (0.0024, 0.0106) | (2e-04, 0.031) | Inf |

```
#save_kable(resistance8WeeksTable,
# "../tables/pathAnalysisResults8Weeks.html")

# And calculating partial R2 of morphometry on metabolic slopes

functionModel8WeeksMassR2 <-
  brm(
    data = slopeData %>%
      filter(week == "8") %>%
      mutate(
        mass = mass - mean(mass, na.rm = T),
        tarsus = tarsusLengthMean - mean(tarsusLengthMean, na.rm = T),
        pretreatment = ifelse(pretreatment == "cold", "A",
          ifelse(pretreatment == "neutral", "B", "C")
        )
```

```
    ) %>%
    mutate(pretreatment = factor(pretreatment, levels = c("B", "A", "C"))) %>%
    drop_na() %>%
    merge(., data %>%
      select(ring, "batch" = exp) %>%
      distinct(),
    by = "ring", all.x = TRUE
    ),
  family = "gaussian",
  bf(mass ~ pretreatment + (1 | batch)) +
    bf(tarsus ~ mass + pretreatment + (1 | batch)) +
    bf(rawSlope ~ tarsus + pretreatment + (1 | batch)) +
    set_rescor(FALSE),
  prior = c(
    set_prior("normal(0, 10)",
      class = "Intercept",
      resp = "mass"
    ),
    set_prior("normal(0, 25)",
      class = "b",
      coef = "pretreatmentA", resp = "mass"
    ),
    set_prior("normal(0, 25)",
      class = "b",
      coef = "pretreatmentC", resp = "mass"
    ),
    set_prior("exponential(2.5)",
      class = "sd",
      group = "batch", resp = "mass"
    ),
    set_prior("exponential(0.15)",
      class = "sigma",
      resp = "mass"
    ),
    set_prior("normal(0, 2.5)",
      class = "Intercept",
      resp = "tarsus"
    ),
    set_prior("normal(0, 2.5)",
      class = "b",
      coef = "pretreatmentA", resp = "tarsus"
    ),
    set_prior("normal(0, 2.5)",
      class = "b",
      coef = "pretreatmentC", resp = "tarsus"
    ),
    set_prior("skew_normal(0, 0.25, 5)",
      class = "b",
      coef = "mass", resp = "tarsus"
    ),
    set_prior("exponential(2)",
      class = "sd",
      group = "batch", resp = "tarsus"
    ),
    set_prior("exponential(1)",
      class = "sigma",
      resp = "tarsus"
    ),
    set_prior("normal(0, 0.01)",
      class = "Intercept",
      resp = "rawSlope"
    ),
    set_prior("normal(0, 0.025)",
      class = "b",
      coef = "pretreatmentA", resp = "rawSlope"
    ),
    set_prior("normal(0, 0.025)",
```

```
        class = "b",
        coef = "pretreatmentC", resp = "rawSlope"
      ),
      set_prior("normal(0, 0.0025)",
        class = "b",
        coef = "tarsus", resp = "rawSlope"
      ),
      set_prior("exponential(50)",
        class = "sd",
        group = "batch", resp = "rawSlope"
      ),
      set_prior("exponential(10)",
        class = "sigma",
        resp = "rawSlope"
      )
    ),
    iter = 50000, warmup = 10000, cores = 4, chains = 4, thin = 20,
    control = list(adapt_delta = .97, max_treedepth = 14),
    silent = TRUE, refresh = 0,
    file = "./models/eightWeekFunctionModelMassR2.Rds"
  )

functionModel8WeeksTarsusR2 <-
  brm(
    data = slopeData %>%
      filter(week == "8") %>%
      mutate(
        mass = mass - mean(mass, na.rm = T),
        tarsus = tarsusLengthMean - mean(tarsusLengthMean, na.rm = T),
        pretreatment = ifelse(pretreatment == "cold", "A",
          ifelse(pretreatment == "neutral", "B", "C")
        )
      ) %>%
      mutate(pretreatment = factor(pretreatment, levels = c("B", "A", "C"))) %>%
      drop_na() %>%
      merge(., data %>%
        select(ring, "batch" = exp) %>%
        distinct(),
      by = "ring", all.x = TRUE
      ),
    family = "gaussian",
    bf(mass ~ pretreatment + (1 | batch)) +
      bf(tarsus ~ mass + pretreatment + (1 | batch)) +
      bf(rawSlope ~ mass + pretreatment + (1 | batch)) +
      set_rescor(FALSE),
    prior = c(
      set_prior("normal(0, 10)",
        class = "Intercept",
        resp = "mass"
      ),
      set_prior("normal(0, 25)",
        class = "b",
        coef = "pretreatmentA", resp = "mass"
      ),
      set_prior("normal(0, 25)",
        class = "b",
        coef = "pretreatmentC", resp = "mass"
      ),
      set_prior("exponential(2.5)",
        class = "sd",
        group = "batch", resp = "mass"
      ),
      set_prior("exponential(0.15)",
        class = "sigma",
        resp = "mass"
      ),
      set_prior("normal(0, 2.5)",
```

```
        class = "Intercept",
        resp = "tarsus"
      ),
      set_prior("normal(0, 2.5)",
        class = "b",
        coef = "pretreatmentA", resp = "tarsus"
      ),
      set_prior("normal(0, 2.5)",
        class = "b",
        coef = "pretreatmentC", resp = "tarsus"
      ),
      set_prior("skew_normal(0, 0.25, 5)",
        class = "b",
        coef = "mass", resp = "tarsus"
      ),
      set_prior("exponential(2)",
        class = "sd",
        group = "batch", resp = "tarsus"
      ),
      set_prior("exponential(1)",
        class = "sigma",
        resp = "tarsus"
      ),
      set_prior("normal(0, 0.01)",
        class = "Intercept",
        resp = "rawSlope"
      ),
      set_prior("normal(0, 0.025)",
        class = "b",
        coef = "pretreatmentA", resp = "rawSlope"
      ),
      set_prior("normal(0, 0.025)",
        class = "b",
        coef = "pretreatmentC", resp = "rawSlope"
      ),
      set_prior("normal(0, 0.001)",
        class = "b",
        coef = "mass", resp = "rawSlope"
      ),
      set_prior("exponential(50)",
        class = "sd",
        group = "batch", resp = "rawSlope"
      ),
      set_prior("exponential(10)",
        class = "sigma",
        resp = "rawSlope"
      )
    ),
    iter = 50000, warmup = 10000, cores = 4, chains = 4, thin = 20,
    control = list(adapt_delta = .97, max_treedepth = 14),
    silent = TRUE, refresh = 0,
    file = "./models/eightWeekFunctionModelTarsusR2.Rds"
  )

massPR2 = brms::bayes_R2(functionModel8Weeks, ndraws = 1000,
                         robust = TRUE, resp = "rawSlope",
                         summary = FALSE) %>%
  as.data.frame() %>%
  select("fullR2" = R2rawSlope) %>%
  cbind(.,
        brms::bayes_R2(functionModel8WeeksMassR2, ndraws = 1000,
                         robust = TRUE, resp = "rawSlope",
                         summary = FALSE)
        ) %>%
  mutate("pR2" = round(fullR2 - R2rawSlope, digits = 3)) %>%
  pull(pR2)
```

```
tarsusPR2 = brms::bayes_R2(functionModel8Weeks, ndraws = 1000,
                           robust = TRUE, resp = "rawSlope",
                           summary = FALSE) %>%
  as.data.frame() %>%
  select("fullR2" = R2rawSlope) %>%
  cbind(.,
        brms::bayes_R2(functionModel8WeeksTarsusR2, ndraws = 1000,
                           robust = TRUE, resp = "rawSlope",
                           summary = FALSE)
        ) %>%
  mutate("pR2" = round(fullR2 - R2rawSlope, digits = 3)) %>%
  pull(pR2)

caption = paste0('Variance in metabolic slopes ',
  '(fold metabolism at thermoneutrality/°C) explained by ',
  'morphometry in eight week old Japanese quail. Metabolic ',
  'slopes are measured below thermoneutrality (<30°C).'
  )

quantile(massPR2, probs = c(0.025, 0.975), type = 8) %>%
  t() %>%
  as.data.frame() %>%
  mutate(`2.5%` = ifelse(`2.5%` < 0, 0, `2.5%`)) %>%
  mutate("95\\% CI" = paste0("[",
                             round(`2.5%`, digits = 3),
                             ",",
                             round(`97.5%`, digits = 3),
                             "]")
  ) %>%
  mutate("massR2" =
           round(median(massPR2), digits = 3)) %>%
  mutate(massR2 = ifelse(massR2 < 0, 0, massR2)) %>%
  select("Mass Partial R\\textsuperscript{2}" = massR2,
         "95\\% CI") %>%
  cbind(.,
    quantile(tarsusPR2, probs = c(0.025, 0.975), type = 8) %>%
      t() %>%
      as.data.frame() %>%
      mutate(`2.5%` = ifelse(`2.5%` < 0, 0, `2.5%`)) %>%
      mutate("95\\% CI" = paste0("[",
                             round(`2.5%`, digits = 3),
                             ",",
                             round(`97.5%`, digits = 3),
                             "]")
           ) %>%
      mutate("tarsusR2" = round(median(tarsusPR2), digits = 3)) %>%
      select("Tarsus Partial R\\textsuperscript{2}" = tarsusR2,
           "95\\% CI")
  ) %>%
  kbl(.,
    longtable = T, booktabs = T, format = "latex",
    caption = caption, escape = FALSE
  ) %>%
  column_spec(column = c(1:10), width = "2.5cm")  %>%
  kable_styling(latex_options = "striped")
```

***Table 18:*** *Variance in metabolic slopes (fold metabolism at thermoneutrality/°C) explained by morphometry in eight week old Japanese quail. Metabolic slopes are measured below thermoneutrality (<30°C).*

| Mass Partial $R^2$ | 95% CI | Tarsus Partial $R^2$ | 95% CI |
|---|---|---|---|
| 0.02 | [0,0.174] | 0.016 | [0,0.164] |

Similar to above, we estimate the costs of misaligning with Bergmann's and Allen's rule at 0°C (i.e. by being two standard deviations below average mass, or two standard deviations above average tarsus length).

```
data.frame(
  "Size" = c("Average", "Small (2x s.d. < mean)"),
  "pretreatment" = "B",
  "mass" = c(mean(functionModel8Weeks$data$mass, na.rm = T),
             mean(functionModel8Weeks$data$mass, na.rm = T) -
               2*sd(functionModel8Weeks$data$mass, na.rm = T)),
  "tarsus" = 0
) %>%
  mutate("slope" = predict(functionModel8Weeks, newdata = .,
                                robust = TRUE, re_form = NA,
                               resp = "rawSlope")[, "Estimate"]) %>%
  mutate(slope = slope +
           mean(subset(slopeData,
                       week == 8 & pretreatment == "neutral")$mslope,
                na.rm = T)
  ) %>%
  mutate(cost = 1 + slope*-30) %>%
  select("Body Size" = Size, "Fold Increase in RMR at 0°C" = cost)
```

```
## # A tibble: 2 x 2
##   `Body Size`            `Fold Increase in RMR at 0°C`
##   <chr>                                          <dbl>
## 1 Average                                         1.99
## 2 Small (2x s.d. < mean)                          1.90
```

```
data.frame(
  "Size" = c("Average", "Long (2x s.d. > mean)"),
  "pretreatment" = "B",
  "tarsus" = c(mean(functionModel8Weeks$data$tarsus, na.rm = T),
             mean(functionModel8Weeks$data$tarsus, na.rm = T) +
               2*sd(functionModel8Weeks$data$tarsus, na.rm = T)),
  "mass" = 0
) %>%
  mutate("slope" = predict(functionModel8Weeks, newdata = .,
                                robust = TRUE, re_form = NA,
                               resp = "rawSlope")[, "Estimate"]) %>%
  mutate(slope = slope +
           mean(subset(slopeData, week == 8 &
                         pretreatment == "neutral")$mslope,
                na.rm = T)
  ) %>%
  mutate(cost = 1 + slope*-30) %>%
  select("Appendage Length" = Size,
         "Fold Increase in RMR at 0°C" = cost)
```

```
## # A tibble: 2 x 2
##   `Appendage Length`    `Fold Increase in RMR at 0°C`
##   <chr>                                         <dbl>
## 1 Average                                        2.01
## 2 Long (2x s.d. > mean)                          1.96
```

```
# Again, formally testing distinctions

mismatchDFMass <- data.frame(
  "Size" = c("Average", "Small (2x s.d. < mean)"),
  "pretreatment" = "B",
  "mass" = c(mean(functionModel8Weeks$data$mass, na.rm = T),
             mean(functionModel8Weeks$data$mass, na.rm = T) -
               2*sd(functionModel8Weeks$data$mass, na.rm = T)),
  "tarsus" = 0
) %>% predict(functionModel8Weeks, newdata = .,
              robust = TRUE, re_form = NA,
              resp = "rawSlope", summary = FALSE) %>%
  as.data.frame() %>%
  rename("Average" = "V1", "Small" = "V2") %>%
  mutate_all(.funs = function(x)
    {x + mean(subset(slopeData, week == 8)$mslope, na.rm = T)}
    )
```

```r
mismatchDFMassTest <- build_hdf(
  vars = list(
    mismatchDFMass$Average,
    mismatchDFMass$Small
    ),
  priors = list(
    rnorm(nrow(mismatchDFMass), 0, 0.01),
    rnorm(nrow(mismatchDFMass), 0, 0.01)
    ),
  names = c("Average", "Small")
)

mismatchTestMass <- hypothesis_df("Small > Average",
                                  mismatchDFMassTest,
                                  class = "b", alpha = 0.05
                                  )

mismatchTestMass$hypothesis$Hypothesis =
  "Small Metabolic Slope > Average Slope"

# Summarising

caption <- paste0(
  "Results of a non-linear hypothesis ",
  "tests comparing predicted metabolic slopes in the cold (30°C - 10°C) ",
  "among eight week old Japanese quail of average or atypically small ",
  "body size (mean - 2 standard deviations). Posterior probabilities are ",
  "calculated using the Savage-Dickey ",
  "density ratio method."
)

mismatchTestMassTable <- mismatchTestMass$hypothesis %>%
  mutate(Hypothesis = c(
    "Small Metabolic Slope > Average Slope"
  )) %>%
  select(-c(Evid.Ratio, Star)) %>%
  mutate(
    Estimate = round(Estimate, digits = 4),
    "Est.Error" = round(Est.Error, digits = 4),
    "CI.Lower" = round(CI.Lower, digits = 4),
    "CI.Upper" = round(CI.Upper, digits = 4)
  ) %>%
  rename(
    "Difference Between Metabolic Slopes" = Estimate,
    "Standard Error" = Est.Error,
    `2.5 % CI` = "CI.Lower",
    `97.5 % CI` = "CI.Upper",
    "Posterior Probability" = Post.Prob
  ) %>%
  kbl(.,
    longtable = T, booktabs = T, format = "latex",
    caption = caption
  ) %>%
  column_spec(column = c(1:2), width = "2.5cm") %>%
  column_spec(column = c(3:10), width = "2cm") %>%
  kable_styling(latex_options = "striped")

mismatchTestMassTable
```

***Table 19:*** *Results of a non-linear hypothesis tests comparing predicted metabolic slopes in the cold (30°C - 10°C) among eight week old Japanese quail of average or atypically small body size (mean - 2 standard deviations). Posterior probabilities are calculated using the Savage-Dickey density ratio method.*

| Hypothesis | Difference Between Metabolic Slopes | Standard Error | 2.5 % CI | 97.5 % CI | Posterior Probability |
|---|---|---|---|---|---|

| | | | | | |
|---|---|---|---|---|---|
| Small Metabolic Slope > Average Slope | 0.0026 | 0.0263 | -0.0406 | 0.0462 | 0.538375 |

```
# Tarsus length comparison

mismatchDFTarsus <- data.frame(
  "Size" = c("Average", "Long (2x s.d. > mean)"),
  "pretreatment" = "B",
  "tarsus" = c(mean(functionModel8Weeks$data$tarsus, na.rm = T),
             mean(functionModel8Weeks$data$tarsus, na.rm = T) +
               2*sd(functionModel8Weeks$data$tarsus, na.rm = T)),
  "mass" = 0
) %>% predict(functionModel8Weeks, newdata = .,
              robust = TRUE, re_form = NA,
              resp = "rawSlope", summary = FALSE) %>%
  as.data.frame() %>%
  rename("Average" = "V1", "Long" = "V2") %>%
  mutate_all(.funs = function(x)
    {x + mean(subset(slopeData, week == 8)$mslope, na.rm = T)}
    )

mismatchDFTarsusTest <- build_hdf(
  vars = list(
    mismatchDFTarsus$Average,
    mismatchDFTarsus$Long
    ),
  priors = list(
    rnorm(nrow(mismatchDFTarsus), 0, 0.01),
    rnorm(nrow(mismatchDFTarsus), 0, 0.01)
    ),
  names = c("Average", "Long")
)

mismatchTestTarsus <- hypothesis_df("Long > Average",
                                    mismatchDFTarsusTest,
                                    class = "b", alpha = 0.05
                                    )

mismatchTestTarsus$hypothesis$Hypothesis =
  "Long Tarsus Metabolic Slope > Average Slope"

caption <- paste0(
  "Results of a non-linear hypothesis ",
  "tests comparing predicted metabolic slopes in the cold (30°C - 10°C) ",
  "among eight week old Japanese quail of average or atypically long ",
  "tarsus length (mean + 2 standard deviations). Posterior probabilities are ",
  "calculated using the Savage-Dickey ",
  "density ratio method."
)

mismatchTestTarsusTable <- mismatchTestTarsus$hypothesis %>%
  mutate(Hypothesis = c(
    "Small Metabolic Slope > Average Slope"
  )) %>%
  select(-c(Evid.Ratio, Star)) %>%
  mutate(
    Estimate = round(Estimate, digits = 4),
    "Est.Error" = round(Est.Error, digits = 4),
    "CI.Lower" = round(CI.Lower, digits = 4),
    "CI.Upper" = round(CI.Upper, digits = 4)
  ) %>%
  rename(
    "Difference Between Metabolic Slopes" = Estimate,
    "Standard Error" = Est.Error,
    `2.5 % CI` = "CI.Lower",
    `97.5 % CI` = "CI.Upper",
```

```
    "Posterior Probability" = Post.Prob
  ) %>%
  kbl(.,
    longtable = T, booktabs = T, format = "latex",
    caption = caption
  ) %>%
  column_spec(column = c(1:2), width = "2.5cm") %>%
  column_spec(column = c(3:10), width = "2cm") %>%
  kable_styling(latex_options = "striped")

mismatchTestTarsusTable
```

***Table 20:*** *Results of a non-linear hypothesis tests comparing predicted metabolic slopes in the cold (30°C - 10°C) among eight week old Japanese quail of average or atypically long tarsus length (mean + 2 standard deviations). Posterior probabilities are calculated using the Savage-Dickey density ratio method.*

| Hypothesis | Difference Between Metabolic Slopes | Standard Error | 2.5 % CI | 97.5 % CI | Posterior Probability |
|---|---|---|---|---|---|
| Small Metabolic Slope > Average Slope | 0.0023 | 0.026 | -0.04 | 0.0456 | 0.531875 |

We then visualise effects of the thermal environment and morphometry on metabolic slopes in eight week old Japanese quail.

```
# Plotting outcomes

p1_8 <- data.frame("pretreatment" = c("B", "A", "C")) %>%
  mutate(
    "mass" = predict(functionModel8Weeks,
      newdata = ., robust = TRUE,
      resp = "mass", re_form = NA
    )[, "Estimate"],
    "SE" = predict(functionModel8Weeks,
      newdata = ., robust = TRUE,
      resp = "mass", re_form = NA
    )[, "Est.Error"]
  ) %>%
  mutate("Mass" = mass +
           mean(subset(slopeData, week == "8")$mass,
                na.rm = T)) %>%
  mutate("pretreatment" = ifelse(pretreatment == "A",
                                 "Cold\n(10°C)",
    ifelse(pretreatment == "B", "Mild\n(20°C)",
      "Warm\n(30°C)"
    )
  )) %>%
  ggplot(aes(x = pretreatment, y = Mass)) +
  geom_errorbar(aes(x = pretreatment, ymin = Mass - SE,
                    ymax = Mass + SE), colour = "black", width = 0.3) +
  geom_point(size = 5, pch = 21, colour = "black",
             aes(x = pretreatment, fill = factor(pretreatment))) +
  geom_line(linetype = "dashed", colour = "black") +
  geom_point(
    size = 2, pch = 21, colour = "black",
    data = subset(slopeData, week == "8") %>%
      mutate("pretreatment" = ifelse(pretreatment == "cold",
                                     "Cold\n(10°C)",
        ifelse(pretreatment == "neutral", "Mild\n(20°C)",
               "Warm\n(30°C)")
      )),
    aes(x = pretreatment, y = mass, fill = factor(pretreatment)),
    position = position_jitter(width = 0.3)
```

```
  ) +
  scale_fill_manual(values = c("#7BB4E3", "black", "#CD5C5C")) +
  theme_classic() +
  theme(legend.position = "none",
        axis.text = element_text(family = "Noto Sans"),
        axis.title = element_text(family = "Noto Sans")) +
  xlab("Rearing Conditions") +
  ylab("Mass (g)")

p2_8 <- data.frame(
  "pretreatment" = c("A", "B", "C"),
  "mass" = 0
) %>%
  mutate(
    "tarsus" = predict(functionModel8Weeks,
      newdata = ., robust = TRUE,
      resp = "tarsus", re_form = NA
    )[, "Estimate"],
    "SE" = predict(functionModel8Weeks,
      newdata = ., robust = TRUE,
      resp = "tarsus", re_form = NA
    )[, "Est.Error"]
  ) %>%
  mutate("Tarsus" = tarsus +
           mean(subset(slopeData, week == "8")$tarsusLengthMean,
                na.rm = T)) %>%
  mutate("Mass" = mass +
           mean(subset(slopeData, week == "8")$mass,
                na.rm = T)) %>%
  mutate("pretreatment" = ifelse(pretreatment == "A",
                                 "Cold\n(10°C)",
    ifelse(pretreatment == "B", "Mild\n(20°C)",
      "Warm\n(30°C)"
    )
  )) %>%
  ggplot(aes(x = pretreatment, y = Tarsus)) +
  geom_errorbar(aes(x = pretreatment, ymin = Tarsus - SE,
                    ymax = Tarsus + SE), colour = "black", width = 0.3) +
  geom_point(size = 5, pch = 21, colour = "black",
             aes(x = pretreatment, fill = factor(pretreatment))) +
  geom_line(linetype = "dashed", colour = "black") +
  geom_point(
    size = 2, pch = 21, colour = "black",
    data = subset(slopeData, week == "8") %>%
      mutate("pretreatment" = ifelse(pretreatment == "cold", "Cold\n(10°C)",
        ifelse(pretreatment == "neutral", "Mild\n(20°C)", "Warm\n(30°C)")
      )),
    aes(x = pretreatment, y = tarsusLengthMean,
        fill = factor(pretreatment)),
    position = position_jitter(width = 0.3)
  ) +
  scale_fill_manual(values = c("#7BB4E3", "black", "#CD5C5C")) +
  theme_classic() +
  theme(legend.position = "none",
        axis.text = element_text(family = "Noto Sans"),
        axis.title = element_text(family = "Noto Sans")) +
  xlab("Rearing Conditions") +
  ylab("Tarsus Length (mm)")

p3_8 <- data.frame(
  "pretreatment" = c("A", "B", "C"),
  "mass" = 0,
  "tarsus" = 0
) %>%
  mutate(
    "slope" = predict(functionModel8Weeks,
      newdata = ., robust = TRUE, re_form = NA,
```

```
      resp = "rawSlope"
    )[, "Estimate"],
    "SE" = predict(functionModel8Weeks,
      newdata = ., robust = TRUE, re_form = NA,
      resp = "rawSlope"
    )[, "Est.Error"]
  ) %>%
  mutate("pretreatment" = ifelse(pretreatment == "A",
                                 "Cold\n(10°C)",
                                 ifelse(pretreatment == "B",
                                        "Mild\n(20°C)",
                                        "Warm\n(30°C)"
                                        )
                                 )
         ) %>%
  mutate("slope" = slope +
           mean(subset(slopeData, week == "8")$mslope,
                na.rm = T)) %>%
  ggplot(aes(x = pretreatment, y = slope)) +
  geom_errorbar(aes(x = pretreatment,
                    ymin = slope - SE, ymax = slope + SE),
                colour = "black", width = 0.3) +
  geom_point(size = 5, pch = 21, colour = "black",
             aes(x = pretreatment, fill = factor(pretreatment))) +
  geom_line(linetype = "dashed", colour = "black") +
  geom_point(
    size = 2, pch = 21, colour = "black",
    data = subset(slopeData, week == "8") %>%
      mutate("pretreatment" = ifelse(pretreatment == "cold",
                                     "Cold\n(10°C)",
                                     ifelse(pretreatment == "neutral",
                                            "Mild\n(20°C)",
                                            "Warm\n(30°C)"
                                            )
                                     )
             ),
    aes(x = pretreatment, y = mslope,
        fill = factor(pretreatment)),
    position = position_jitter(width = 0.3)
  ) +
  scale_fill_manual(values = c("#7BB4E3", "black", "#CD5C5C")) +
  theme_classic() +
  theme(legend.position = "none",
        axis.text = element_text(family = "Noto Sans"),
        axis.title = element_text(family = "Noto Sans")) +
  xlab("Rearing Conditions") +
  ylab("Metabolic Slope\n(Fold Metabolism at Thermoneutrality)")

pretreatmentPanel8 <- ((p1_8/p2_8) | (p3_8)) +
  plot_annotation(tag_levels = "A")

showtext_auto()
pretreatmentPanel8
```

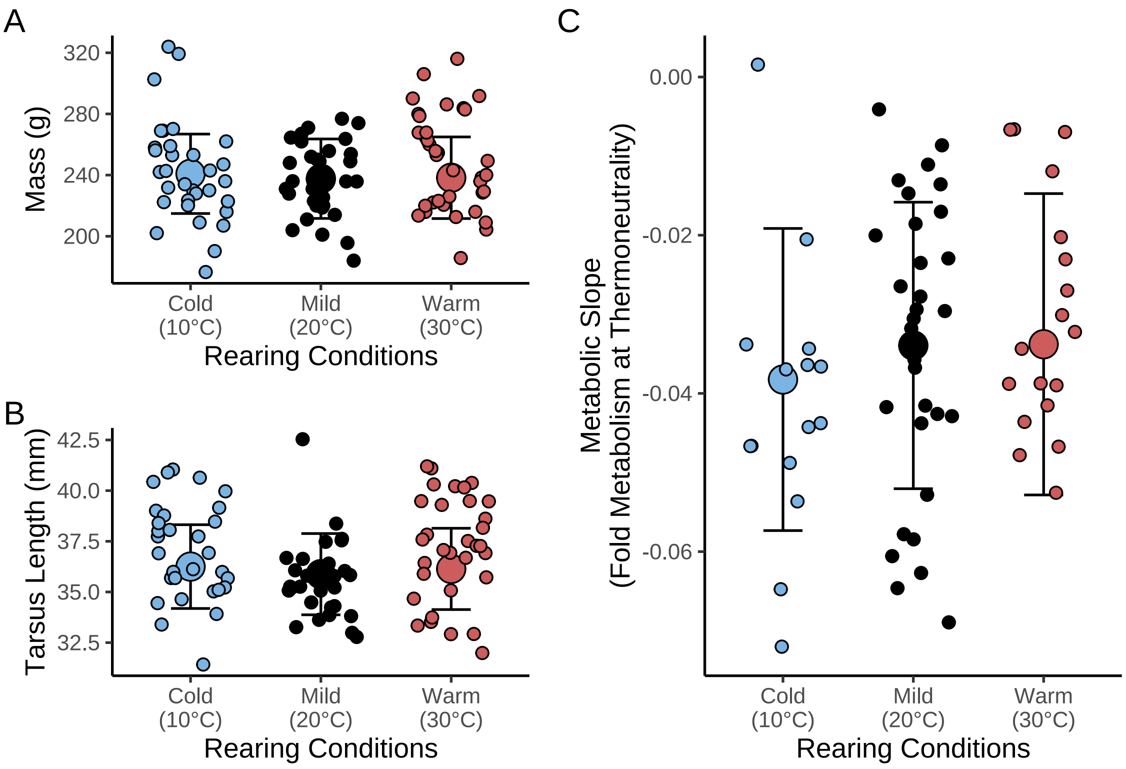


***Figure 41:*** *Effects of post-hatch thermal conditions on morphometry and thermal physiology in eight week old Japanese quail. Small dots represent raw data points while large dots represent estimated effectgs per grouping from a Bayesian path analysis. Errorbars represent +/- one standard error. Panels A and B display effects of rearing conditions on morphology, while panels C and D represent effects of rearing conditions on thermal physiology. Cold and warm rearing treatments persisted for at least 3 weeks of life, followed by holding at 20°C.*

```
ggsave("../plots/rearingConditionEffects8Weeks_2Fold.pdf",
  pretreatmentPanel8,
  dpi = 800, height = 7, width = 9
)
showtext_auto(enable = "FALSE")
```

```
p1 <- data.frame(
  "pretreatment" = "B",
  "mass" = seq(min(functionModel8Weeks$data$mass, na.rm = T),
    max(functionModel8Weeks$data$mass, na.rm = T),
    by = 0.1
  ),
  "tarsus" = 0
) %>%
  mutate(
    "slope" = predict(functionModel8Weeks,
      newdata = ., robust = TRUE, re_form = NA,
      resp = "rawSlope"
    )[, "Estimate"],
    "SE" = predict(functionModel8Weeks,
      newdata = ., robust = TRUE, re_form = NA,
      resp = "rawSlope"
    )[, "Est.Error"]
  ) %>%
  mutate(slope = slope +
           mean(subset(slopeData, week == 8)$mslope,
                na.rm = T)) %>%
  mutate("Mass" = mass +
```

```
          mean(subset(slopeData, week == "8")$mass,
               na.rm = T)) %>%
  ggplot(aes(x = Mass, y = slope)) +
  geom_ribbon(aes(x = Mass, ymin = slope - SE,
                  ymax = slope + SE),
              fill = "grey50", alpha = 0.5
  ) +
  geom_smooth(method = "lm", linetype = "dashed",
              se = FALSE, colour = "black") +
  geom_point(
    size = 2.5, pch = 21, colour = "black", alpha = 0.5, fill = "grey50",
    data = subset(slopeData, week == "8"),
    aes(x = mass, y = mslope)
  ) +
  xlim(c(180, 300)) +
  theme_classic() +
  theme(axis.text = element_text(family = "Noto Sans"),
        axis.title = element_text(family = "Noto Sans"),
        #axis.text = element_text(family = "Noto Sans", size = 16),
        #axis.title = element_text(family = "Noto Sans", size = 16),
        legend.position = "none"
        ) +
  xlab("Body Mass (g)") +
  ylab("Metabolic Slope < 30°C\n(Fold Metabolism/°C)")

p2 <- data.frame(
  "pretreatment" = "B",
  "tarsus" = seq(min(functionModel8Weeks$data$tarsus,
                     na.rm = T),
                 max(functionModel8Weeks$data$tarsus,
                     na.rm = T),
    by = 0.1
  ),
  "mass" = 0
) %>%
  mutate(
    "slope" = predict(functionModel8Weeks,
      newdata = .,  robust = TRUE, re_form = NA,
      resp = "rawSlope"
    )[, "Estimate"],
    "SE" = predict(functionModel8Weeks,
      newdata = .,  robust = TRUE, re_form = NA,
      resp = "rawSlope"
    )[, "Est.Error"]
  ) %>%
  mutate("Tarsus" = tarsus +
          mean(subset(slopeData, week == "8")$tarsusLengthMean,
               na.rm = T)) %>%
  mutate(slope = slope +
          mean(subset(slopeData, week == "8")$mslope,
               na.rm = T)) %>%
  ggplot(aes(x = Tarsus, y = slope)) +
  geom_ribbon(aes(x = Tarsus, ymin = slope - SE,
                  ymax = slope + SE),
    fill = "grey50", alpha = 0.5
  ) +
  geom_smooth(method = "lm", linetype = "dashed",
              se = FALSE, colour = "black") +
  geom_point(
    size = 2.5, pch = 21, colour = "black",
    alpha = 0.5, fill = "grey50",
    data = subset(slopeData, week == "8"),
    aes(x = tarsusLengthMean, y = mslope)
  ) +
  xlim(c(30, 42)) +
  theme_classic() +
  theme(legend.position = "none",
```

```
        axis.text = element_text(family = "Noto Sans"),
        axis.title = element_text(family = "Noto Sans")) +
  xlab("Tarsus Length (mm)") +
  ylab("Metabolic Slope (Fold\nMetabolism at Thermoneutrality/°C)")

massPred <- data.frame(
  "pretreatment" = "B",
  "mass" = c(
    mean(functionModel8Weeks$data$mass, na.rm = T) -
      sd(functionModel8Weeks$data$mass, na.rm = T),
    mean(functionModel8Weeks$data$mass, na.rm = T),
    mean(functionModel8Weeks$data$mass, na.rm = T) +
      sd(functionModel8Weeks$data$mass, na.rm = T)
  ),
  "tarsus" = 0
) %>%
  predict(functionModel8Weeks, re_form = NA,
    resp = "rawSlope", newdata = .,
    summary = TRUE, robust = TRUE
  ) %>%
  as.data.frame() %>%
  mutate("slope" = Estimate +
           mean(
             subset(slopeData,
                    week == 8 & pretreatment == "neutral"
                    )$mslope,
             na.rm = TRUE
           )
  ) %>%
  mutate("Size" = c("Small", "Average", "Large")) %>%
  select(Size, slope, "slopeSE" = Est.Error) %>%
  slice(rep(1:n(), each = length(seq(-20, -0.1, by = 0.1)))) %>%
  mutate("Ta" = rep(seq(-20, -0.1, by = 0.1), 3)) %>%
  mutate("VO2" = 1 + Ta*slope,
         "LL" = 1 + Ta*(slope - slopeSE),
         "UL" = 1 + Ta*(slope + slopeSE)
  ) %>%
  mutate("Ta" = Ta + 26,
         Size = factor(Size, levels = c("Small", "Average", "Large"))
         )

jitterer <- position_jitter(width = 1, seed = 100)

massPlot <- massPred %>%
  ggplot(aes(x = Ta, y = VO2)) +
  geom_point(
    data = data.frame("Ta" = 26, "VO2" = 1),
    pch = 21, colour = "black", fill = "grey15",
    size = 3
  ) +
    geom_point(
    data = slopes8Weeks$data %>%
      merge(., subset(all, week == "8") %>%
              select(ring, mass),
            by = "ring", all.x = TRUE) %>%
      mutate(Ta = Ta + 26),
    aes(x = Ta, y = VO2),
    position = jitterer,
    pch = 21, colour = "black", fill = "transparent"
  ) +
    geom_point(
    data = slopes8Weeks$data %>%
      merge(., subset(all, week == "8") %>%
              select(ring, mass),
            by = "ring", all.x = TRUE) %>%
      mutate(Ta = Ta + 26),
    aes(x = Ta, y = VO2, colour = mass),
```

```
  position = jitterer, alpha = 0.5
) +
scale_colour_gradient2(
  low = "#19242E",
  mid = "#BBD8F0",
  high = "grey98",
  midpoint = mean(subset(slopeData, week == "8")$mass, na.rm = T),
  guide = NULL
  ) +
geom_ribbon(aes(x = Ta, ymin = LL, ymax = UL, fill = Size),
            alpha = 0.4, colour = "black") +
scale_fill_manual(values = c("#194A7A", "#BBD8F0", "grey98"),
                  labels = c(
  paste0(
    "Mean Mass -\n1 SD (",
    round(
      mean(subset(slopeData, week == "8")$mass, na.rm = T) -
        sd(subset(slopeData, week == "8")$mass, na.rm = T),
        digits = 0.1
      ),
    "g)"
  ),
  paste0(
    "Mean Mass\n(",
    round(mean(subset(slopeData, week == "8")$mass, na.rm = T),
          digits = 0.1),
    "g)"
  ),
  paste0(
    "Mean Mass +\n1 SD (",
    round(
      mean(subset(slopeData, week == "8")$mass, na.rm = T) +
        sd(subset(slopeData, week == "8")$mass, na.rm = T),
        digits = 0.1
      ),
    "g)"
  )
), name = NULL) +
scale_linetype_manual(
  values = c("solid", "dashed", "dotted"),
  labels = c(
  paste0(
    "Mean Mass -\n1 SD (",
    round(
      mean(subset(slopeData, week == "8")$mass, na.rm = T) -
        sd(subset(slopeData, week == "8")$mass, na.rm = T),
        digits = 0.1
      ),
    "g)"
  ),
  paste0(
    "Mean Mass\n(",
    round(mean(subset(slopeData, week == "8")$mass, na.rm = T),
          digits = 0.1),
    "g)"
  ),
  paste0(
    "Mean Mass +\n1 SD (",
    round(
      mean(subset(slopeData, week == "8")$mass, na.rm = T) +
        sd(subset(slopeData, week == "8")$mass, na.rm = T),
        digits = 0.1
      ),
    "g)"
  )
  ), name = NULL
) +
```

```
  geom_line(
    aes(linetype = Size)
  ) +
  theme_classic() +
  xlab("Ambient Temperature (°C)") +
  ylab("Resting Metabolism\n(Fold Change from Thermoneutrality)") +
  scale_linetype_manual(
    values = c("solid", "dashed", "dotted"),
    labels = c(
    paste0(
      "Mean Mass -\n1 SD (",
      round(
        mean(subset(slopeData, week == "8")$mass, na.rm = T) -
          sd(subset(slopeData, week == "8")$mass, na.rm = T),
            digits = 0.1
        ),
      "g)"
    ),
    paste0(
      "Mean Mass\n(",
      round(mean(subset(slopeData, week == "8")$mass, na.rm = T),
            digits = 0.1),
      "g)"
    ),
    paste0(
      "Mean Mass +\n1 SD (",
      round(
        mean(subset(slopeData, week == "8")$mass, na.rm = T) +
          sd(subset(slopeData, week == "8")$mass, na.rm = T),
            digits = 0.1
        ),
      "g)"
    )
    ), name = NULL
  ) +
  guides(fill = guide_legend(override.aes = list(size = 3)),
         linetype = guide_legend(override.aes = list(size = 4))
         ) +
  theme(
    #axis.text = element_text(family = "Noto Sans", size = 16.5),
    #axis.title = element_text(family = "Noto Sans", size = 16.5),
    #legend.text = element_text(family = "Noto Sans", size = 15.5),
    #legend.title = element_text(family = "Noto Sans", size = 16),
    axis.text = element_text(family = "Noto Sans"),
    axis.title = element_text(family = "Noto Sans"),
    legend.text = element_text(family = "Noto Sans", size = 7),
    legend.title = element_text(family = "Noto Sans"),
    legend.position = "bottom"
  )

showtext_auto()
ggsave("../plots/massPlot8Weeks.pdf",
  massPlot,
  dpi = 800, height = 7, width = 7
)
showtext_auto(enable = FALSE)

tarsusPred <- data.frame(
  "pretreatment" = "B",
  "tarsus" = c(
    mean(functionModel8Weeks$data$tarsus, na.rm = T) -
      sd(functionModel8Weeks$data$tarsus, na.rm = T),
    mean(functionModel8Weeks$data$tarsus, na.rm = T),
    mean(functionModel8Weeks$data$tarsus, na.rm = T) +
      sd(functionModel8Weeks$data$tarsus, na.rm = T)
  ),
  "mass" = 0
```

```
) %>%
  predict(functionModel8Weeks, re_form = NA,
    resp = "rawSlope", newdata = .,
    summary = TRUE, robust = TRUE
  ) %>%
  as.data.frame() %>%
  mutate("slope" = Estimate +
           mean(
             subset(slopeData,
                    week == 8 & pretreatment == "neutral"
                    )$mslope,
             na.rm = TRUE
           )
  ) %>%
  mutate("Size" = c("Small", "Average", "Large")) %>%
  select(Size, slope, "slopeSE" = Est.Error) %>%
  slice(rep(1:n(), each = length(seq(-20, -0.1, by = 0.1)))) %>%
  mutate("Ta" = rep(seq(-20, -0.1, by = 0.1), 3)) %>%
  mutate("VO2" = 1 + Ta*slope,
         "LL" = 1 + Ta*(slope - slopeSE),
         "UL" = 1 + Ta*(slope + slopeSE)
  ) %>%
  mutate("Ta" = Ta + 26,
         Size = factor(Size, levels = c("Small", "Average", "Large"))
         )

tarsusPlot <- tarsusPred %>%
  ggplot(aes(x = Ta, y = VO2)) +
  geom_point(
    data = slopes8Weeks$data %>%
      merge(., subset(all, week == "8") %>%
              select(ring, "tarsus" = tarsusLengthMean),
            by = "ring", all.x = TRUE) %>%
      mutate(Ta = Ta + 26),
    aes(x = Ta, y = VO2),
    position = jitterer, alpha = 0.5,
    pch = 21, colour = "black", fill = "transparent"
  ) +
  geom_point(
    data = slopes8Weeks$data %>%
      merge(., subset(all, week == "8") %>%
              select(ring, "tarsus" = tarsusLengthMean),
            by = "ring", all.x = TRUE) %>%
      mutate(Ta = Ta + 26),
    aes(x = Ta, y = VO2, colour = tarsus),
    position = jitterer, alpha = 0.5
  ) +
  geom_point(
    data = data.frame("Ta" = 26, "VO2" = 1),
    pch = 21, colour = "black", fill = "grey15",
    size = 3
  ) +
  geom_ribbon(aes(x = Ta, ymin = LL, ymax = UL, fill = Size),
              alpha = 0.4, colour = "black") +
  geom_line(
    aes(linetype = Size)
  ) +
  theme_classic() +
  xlab("Ambient Temperature (°C)") +
  ylab("Resting Metabolism\n(Fold Change from Thermoneutrality)") +
  scale_fill_manual(values = c("#194A7A", "#BBD8F0", "grey98"),
                    labels = c(
    paste0(
      "Mean Tarsus\nLength -\n1 SD (",
      round(
        mean(subset(slopeData, week == "8")$tarsusLengthMean,
             na.rm = T) -
```

```
        sd(subset(slopeData, week == "8")$tarsusLengthMean,
           na.rm = T),
          digits = 0.1
      ),
    "mm)"
  ),
  paste0(
    "Mean Tarsus\nLength (",
    round(
      mean(subset(slopeData, week == "8")$tarsusLengthMean,
           na.rm = T),
          digits = 0.1),
    "mm)"
  ),
  paste0(
    "Mean Tarsus\nLength +\n1 SD (",
    round(
      mean(subset(slopeData, week == "8")$tarsusLengthMean,
           na.rm = T) +
        sd(subset(slopeData, week == "8")$tarsusLengthMean,
           na.rm = T),
          digits = 0.1
      ),
    "mm)"
  )
), name = NULL) +
scale_linetype_manual(
  values = c("solid", "dashed", "dotted"),
  labels = c(
  paste0(
    "Mean Tarsus\nLength -\n1 SD (",
    round(
      mean(subset(slopeData, week == "8")$tarsusLengthMean,
           na.rm = T) -
        sd(subset(slopeData, week == "8")$tarsusLengthMean,
           na.rm = T),
          digits = 0.1
      ),
    "mm)"
  ),
  paste0(
    "Mean Tarsus\nLength (",
    round(
      mean(subset(slopeData, week == "8")$tarsusLengthMean,
           na.rm = T),
          digits = 0.1),
    "mm)"
  ),
  paste0(
    "Mean Tarsus\nLength +\n1 SD (",
    round(
      mean(subset(slopeData, week == "8")$tarsusLengthMean,
           na.rm = T) +
        sd(subset(slopeData, week == "8")$tarsusLengthMean,
           na.rm = T),
          digits = 0.1
      ),
    "mm)"
  )
  ), name = NULL
) +
scale_colour_gradient2(
  low = "#19242E",
  mid = "#BBD8F0",
  high = "grey98",
  midpoint = mean(subset(slopeData, week == "8")$tarsus, na.rm = T),
  guide = NULL
```

```
    ) +
  guides(fill = guide_legend(override.aes = list(size = 3)),
         linetype = guide_legend(override.aes = list(size = 4))
         ) +
  theme(
    # axis.text = element_text(family = "Noto Sans", size = 16.5),
    # axis.title = element_text(family = "Noto Sans", size = 16.5),
    # legend.text = element_text(family = "Noto Sans", size = 15.5),
    # legend.title = element_text(family = "Noto Sans", size = 16),
    axis.text = element_text(family = "Noto Sans"),
    axis.title = element_text(family = "Noto Sans"),
    legend.text = element_text(family = "Noto Sans", size = 7),
    legend.title = element_text(family = "Noto Sans"),
    legend.position = "bottom"
  )

showtext_auto()
ggsave("../plots/tarsusPlot8Weeks.pdf",
  tarsusPlot,
  dpi = 800, height = 7, width = 7
)

allMorphology <- (p1 + massPlot) / (p2 + tarsusPlot) +
  plot_annotation(tag_levels = "A")

allMorphology
```

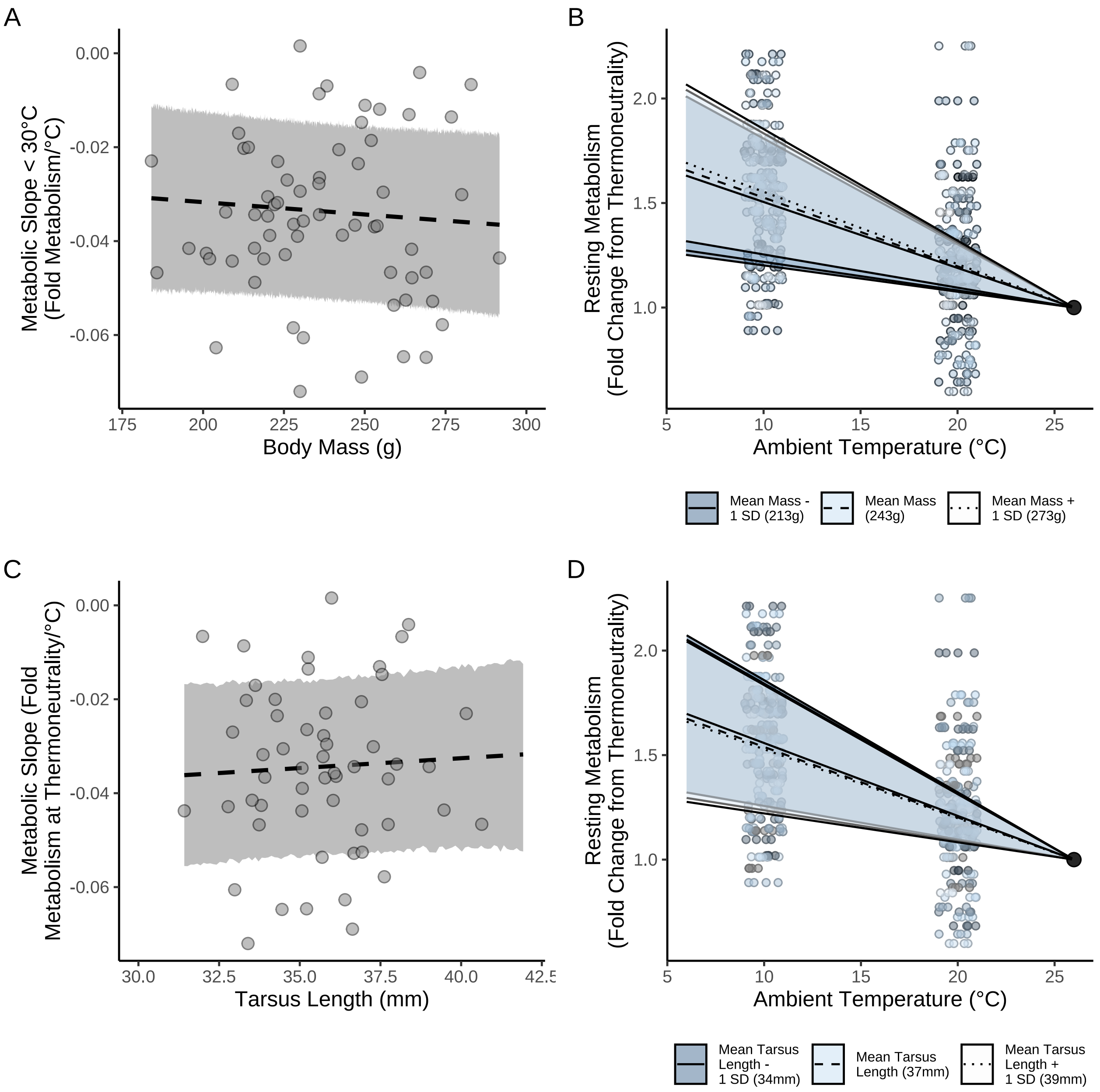


***Figure 42:*** *Effect of morphology on metabolic slopes (fold metabolism at thermoneutrality/°C) of eight week old Japanese quail in response to cold (10°C - 30°C). Dots represent raw values, trend lines are marginalised predicted from a Bayesian path analysis, and ribbons represent one standard error around predictions. Predictions in panels B and D assume rearing at mild conditions (20°C).*

```r
ggsave("../plots/morphologyEffectsResistance8Weeks_3Fold.pdf",
  allMorphology,
  dpi = 800, height = 7, width = 9
)

showtext_auto(enable = FALSE)
```

```r
# As fold increase at 10°C

massPlot10 <- massPred %>%
  filter(Ta == 10) %>%
  ggplot(aes(x = Size, y = VO2)) +
```

```
  geom_point(
    data = slopes8Weeks$data %>%
      filter(Ta == -16) %>%
      merge(., subset(all, week == "8") %>%
              select(ring, mass) %>%
              distinct() %>%
              mutate(Size = ifelse(mass < (mean(mass, na.rm = T) -
                                             sd(mass, na.rm = T)),
                                   "Small",
                                   ifelse((mass > (mean(mass, na.rm = T) -
                                             sd(mass, na.rm = T))) &
                                            (mass < (mean(mass, na.rm = T) +
                                             sd(mass, na.rm = T))),
                                          "Average", "Large"
                                   )
                                   )
                     ),
            by = "ring", all.x = TRUE) %>%
      mutate(Size = factor(Size, levels = c("Small", "Average", "Large"))
             ),
    aes(x = Size, y = VO2, colour = Size),
    position = position_jitter(width = 0.25), alpha = 0.5
  ) +
  geom_errorbar(aes(ymin = LL, ymax = UL),
                colour = "black", width = 0.25) +
  geom_point(size = 3, pch = 21, colour = "black",
             aes(fill = Size)
  ) +
  xlab("Body Mass") +
  ylab("Fold Increase in Metabolism at 10°C\n(From Thermoneutrality)") +
  scale_x_discrete(labels = c("Mean - 1 SD", "Mean", "Mean + 1 SD")) +
  scale_fill_manual(values = c("#DECCC1", "#855E46", "#231709")) +
  scale_colour_manual(values = c("#DECCC1", "#855E46", "#231709")) +
  theme_classic() +
  theme(
    legend.position = "none",
    axis.text = element_text(family = "Noto Sans"),
    axis.title = element_text(family = "Noto Sans")
  )

tarsusPlot10 <- tarsusPred %>%
  filter(Ta == 10) %>%
  ggplot(aes(x = Size, y = VO2)) +
  geom_point(
    data = slopes8Weeks$data %>%
      filter(Ta == -16) %>%
      merge(., subset(all, week == "8") %>%
              select(ring, "tarsus" = tarsusLengthMean) %>%
              distinct() %>%
              mutate(Size = ifelse(tarsus < (mean(tarsus, na.rm = T) -
                                             sd(tarsus, na.rm = T)),
                                   "Small",
                                   ifelse((tarsus > (mean(tarsus, na.rm = T) -
                                             sd(tarsus, na.rm = T))) &
                                            (tarsus < (mean(tarsus, na.rm = T) +
                                             sd(tarsus, na.rm = T))),
                                          "Average", "Large"
                                   )
                                   )
                     ),
            by = "ring", all.x = TRUE) %>%
      drop_na(tarsus) %>%
      mutate(Size = factor(Size, levels = c("Small", "Average", "Large"))
             ),
    aes(x = Size, y = VO2, colour = Size),
    position = position_jitter(width = 0.25), alpha = 0.5
  ) +
```

```
  geom_errorbar(aes(ymin = LL, ymax = UL),
                colour = "black", width = 0.25) +
  geom_point(size = 3, pch = 21, colour = "black",
             aes(fill = Size)
  ) +
  xlab("Tarsus Length") +
  ylab("Fold Increase in Metabolism at 10°C\n(From Thermoneutrality)") +
  scale_x_discrete(labels = c("Mean - 1 SD",
                              "Mean",
                              "Mean + 1 SD")) +
  scale_fill_manual(values = c("#DECCC1", "#855E46", "#231709")) +
  scale_colour_manual(values = c("#DECCC1", "#855E46", "#231709")) +
  theme_classic() +
  theme(
    legend.position = "none",
    axis.text = element_text(family = "Noto Sans"),
    axis.title = element_text(family = "Noto Sans")
  )

allMorphology10 <- (p1 + massPlot10) / (p2 + tarsusPlot10) +
  plot_annotation(tag_levels = "A")

allMorphology10
```

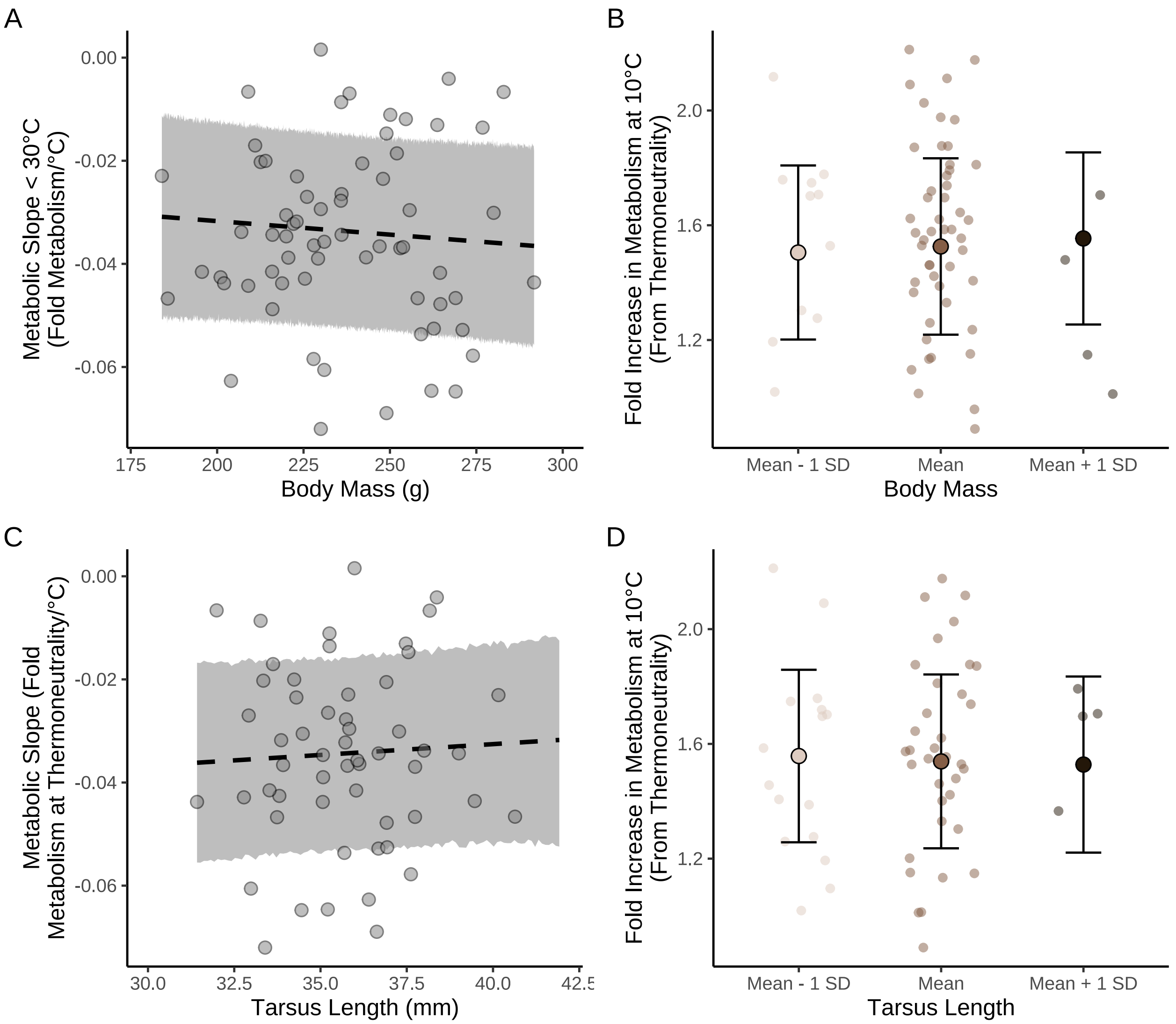


***Figure 43:*** *Effect of morphology and thermal environment on fold resting metabolism at 10°C in eight week old Japanese quail. Dots represent raw values, trend lines are marginalised predicted from a Bayesian path analysis, and ribbons represent one standard error around predictions.*

```
ggsave("../plots/morphologyEffectsResistance8Weeks_Fold10.pdf",
  allMorphology10,
  dpi = 800, height = 7, width = 9
)
```

Again, we visualise and summarise direct and indirect effects of thermal history (here, rearing conditions) and morphometry on metabolic slopes in our quail. To simplify comparisons, model coefficients are scaled to represent the effect that a change in one standard deviation (or categorical level) of a predictor elicits on metabolic slopes (in standard deviations).

```
scaledBetas <- as.data.frame(functionModel8Weeks) %>%
  mutate(b_mass_pretreatmentA = b_mass_pretreatmentA/
           sd(functionModel8Weeks$data$mass),
         b_mass_pretreatmentC = b_mass_pretreatmentC/
           sd(functionModel8Weeks$data$mass),
         b_tarsus_pretreatmentA = b_tarsus_pretreatmentA/
           sd(functionModel8Weeks$data$tarsus),
```

```
        b_tarsus_pretreatmentC = b_tarsus_pretreatmentC/
          sd(functionModel8Weeks$data$tarsus),
        b_tarsus_mass =
          (b_tarsus_mass*sd(functionModel8Weeks$data$mass))/
          sd(functionModel8Weeks$data$tarsus),
        b_rawSlope_mass =
          (b_rawSlope_mass*sd(functionModel8Weeks$data$mass))/
          sd(functionModel8Weeks$data$rawSlope),
        b_rawSlope_tarsus =
          (b_rawSlope_tarsus*sd(functionModel8Weeks$data$tarsus))/
          sd(functionModel8Weeks$data$rawSlope),
        b_rawSlope_pretreatmentA = b_rawSlope_pretreatmentA/
          sd(functionModel8Weeks$data$rawSlope),
        b_rawSlope_pretreatmentC = b_rawSlope_pretreatmentC/
          sd(functionModel8Weeks$data$rawSlope),
  )

fullEffectEightWeeksBetas <- scaledBetas %>%
  mutate("Effects" = "Direct Effects") %>%
  mutate(
    "Body Mass" = b_rawSlope_mass,
    "Tarsus Length" = b_rawSlope_tarsus,
    "Cold Rearing\n(10°C)" = b_rawSlope_pretreatmentA,
    "Warm Rearing\n(30°C)" = b_rawSlope_pretreatmentC
  ) %>%
  select(Effects, `Body Mass`, `Tarsus Length`,
         `Cold Rearing\n(10°C)`,
         `Warm Rearing\n(30°C)`) %>%
  rbind(
    .,
    scaledBetas %>%
      mutate("Effects" = "Indirect Effects") %>%
      mutate(
        "Body Mass" = b_tarsus_mass *
          b_rawSlope_tarsus,
        "Tarsus Length" = NA,
        "Cold Rearing\n(10°C)" =
          b_tarsus_pretreatmentA * b_rawSlope_tarsus +
          b_mass_pretreatmentA * b_rawSlope_mass +
          b_mass_pretreatmentA * b_tarsus_mass *
          b_rawSlope_tarsus,
        "Warm Rearing\n(30°C)" =
          b_tarsus_pretreatmentC * b_rawSlope_tarsus +
          b_mass_pretreatmentC * b_rawSlope_mass +
          b_mass_pretreatmentC * b_tarsus_mass *
          b_rawSlope_tarsus,
      ) %>%
      select(Effects, `Body Mass`, `Tarsus Length`,
             `Cold Rearing\n(10°C)`, `Warm Rearing\n(30°C)`)
  ) %>%
  rbind(., scaledBetas %>%
    mutate("Effects" = "Total Effects") %>%
    mutate(
      "Body Mass" =
        b_rawSlope_mass +
          b_tarsus_mass * b_rawSlope_tarsus,
      "Tarsus Length" =
        b_rawSlope_tarsus,
      "Cold Rearing\n(10°C)" =
        b_rawSlope_pretreatmentA +
          b_mass_pretreatmentA *
        b_rawSlope_mass +
          b_tarsus_pretreatmentA *
        b_rawSlope_tarsus +
          b_mass_pretreatmentA *
        b_tarsus_mass * b_rawSlope_tarsus,
      "Warm Rearing\n(30°C)" =
```

```
         b_rawSlope_pretreatmentC +
          b_mass_pretreatmentC *
        b_rawSlope_mass +
          b_tarsus_pretreatmentC *
        b_rawSlope_tarsus +
          b_mass_pretreatmentC *
        b_tarsus_mass * b_rawSlope_tarsus,
    ) %>%
    select(Effects, `Body Mass`, `Tarsus Length`,
             `Cold Rearing\n(10°C)`, `Warm Rearing\n(30°C)`)
    ) %>%
  pivot_longer(c(-Effects), names_to = "var", values_to = "values") %>%
  mutate(var = factor(var,
                      levels = c("Tarsus Length",
                                 "Body Mass",
                                 "Warm Rearing\n(30°C)",
                                 "Cold Rearing\n(10°C)")))

fullEffectEightWeeks <- fullEffectEightWeeksBetas %>%
  ggplot(aes(x = values, y = var, fill = var)) +
  facet_wrap(~Effects) +
  stat_halfeye(normalize = "xy", colour = "black", alpha = 0.7) +
  geom_vline(xintercept = 0, linetype = "dashed",
             colour = "black") +
  xlab(".                   Effect on Metabolic Slope (standard deviations)") +
  scale_fill_manual(values = c("black", "grey50", "#CD5C5C", "#7BB4E3")) +
  theme_classic() +
  theme(
    legend.position = "none", axis.title.y = element_blank(),
    axis.text.y = element_text(size = 11, colour = "black",
                               family = "Noto Sans"),
    axis.title.x = element_text(family = "Noto Sans", hjust = -0.005),
    axis.text.x = element_text(family = "Noto Sans")
  )

showtext_auto(enable = TRUE)

fullEffectEightWeeks
```

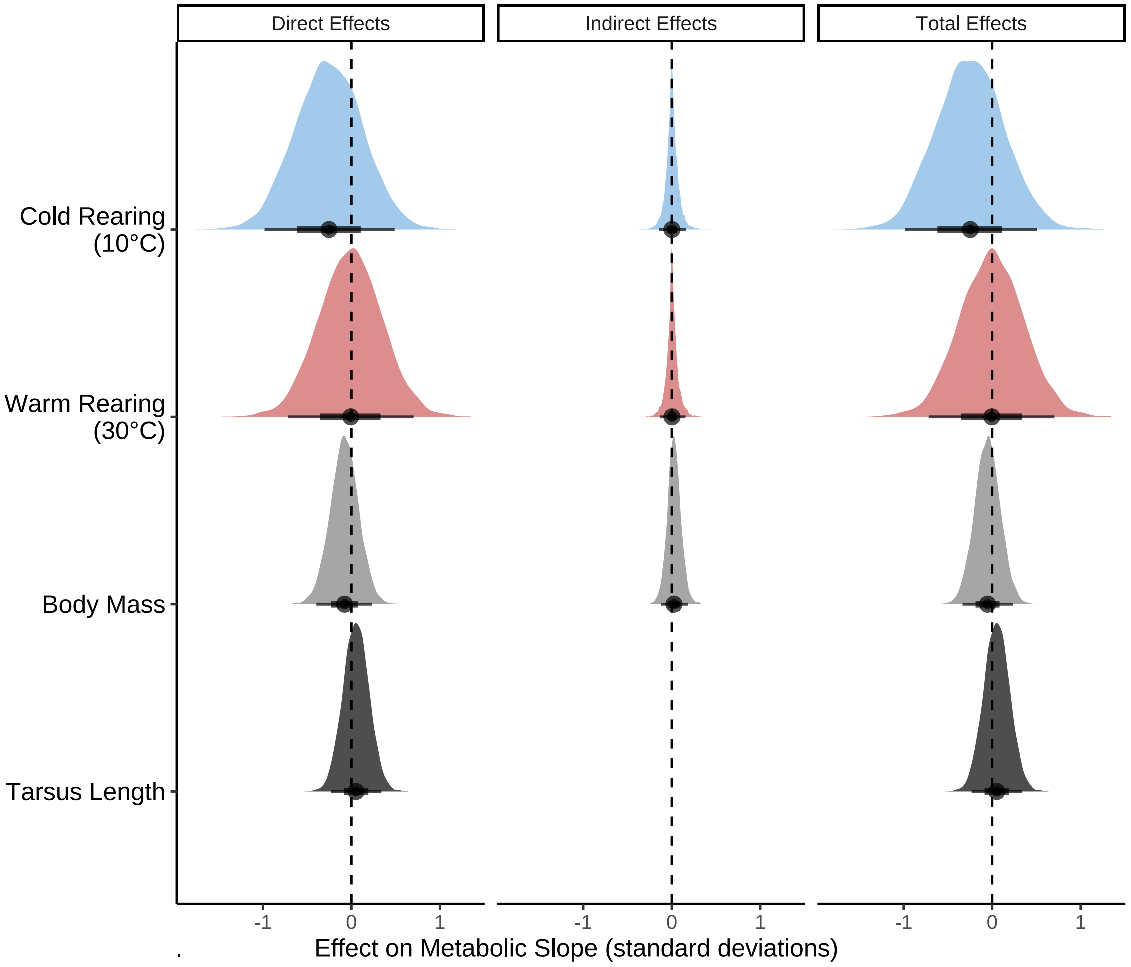


***Figure 44:*** *Effects of morphology (body mass, g, and tarsus length, mm) on metabolic slopes (in standard deviations) at ambient temperatures below thermoneutrality (<30°C) in eight week old Japanese quail. Distributions represent those derived from posteriors of a Bayesian path analysis.*

```
ggsave("../plots/indirectEffectsOnResistance8Weeks_3ScaledFold.pdf",
       fullEffectEightWeeks,
  dpi = 800, width = 8, height = 6
)
showtext_auto(enable = FALSE)

# Summarising results

caption <- paste0("Direct, indirect, and total effects of morphology and ",
                  "rearing temperature on metabolic slope ",
                  "(fold metabolism at thermoneutrality/ ",
                  "°C) ",
                  "of mature Japanese quail. Effects are derived from ",
                  "a Bayesian path analysis and represent those predicted for a ",
                  "change in one standard deviation (or categorical level) of ",
                  "a given predictor on the standard deviation of metabolic ",
                  "slopes. Estimates indicate posterior medians and credible ",
                  "intervels (CIs) indicate quantile intervals."
)

week8ResultsScaled <-
  fullEffectEightWeeksBetas %>%
  filter(!is.na(values) & !is.nan(values)) %>%
```

```
  group_by(Effects, var) %>%
  summarise(
    "Estimate" = median(values),
    "50\\% CIs" = paste0(
      "[",
      round(
        quantile(values, probs = 0.1, type = 8),
        digits = 4
      ),
      ", ",
      round(
        quantile(values, probs = 0.9, type = 8),
        digits = 4
      ),
      "]"
    ),
    "95\\% CIs" = paste0(
      "[",
      round(
        quantile(values, probs = 0.025, type = 8),
        digits = 4
      ),
      ", ",
      round(
        quantile(values, probs = 0.975, type = 8),
        digits = 4
      ),
      "]"
    )
  ) %>%
  mutate("Effects" = gsub("[[:space:]].*", "", Effects)) %>%
  select(
    "Predictor" = "var", "Effect Level" = "Effects",
    Estimate, `50\\% CIs`, `95\\% CIs`
  ) %>%
  arrange(Predictor, `Effect Level`) %>%
  kbl(.,
    longtable = T, booktabs = T, format = "latex", escape = FALSE,
    caption = caption
  ) %>%
  column_spec(column = c(1:2), width = "2.2cm") %>%
  column_spec(column = c(3:10), width = "1.9cm") %>%
  kable_styling(latex_options = "striped")

week8ResultsScaled
```

***Table 21:*** *Direct, indirect, and total effects of morphology and rearing temperature on metabolic slope (fold metabolism at thermoneutrality/ °C) of mature Japanese quail. Effects are derived from a Bayesian path analysis and represent those predicted for a change in one standard deviation (or categorical level) of a given predictor on the standard deviation of metabolic slopes. Estimates indicate posterior medians and credible intervels (CIs) indicate quantile intervals.*

| Predictor | Effect Level | Estimate | 50% CIs | 95% CIs |
|---|---|---|---|---|
| Tarsus Length | Direct | 0.0518263 | [-0.1345, 0.2381] | [-0.2326, 0.3388] |
| Tarsus Length | Total | 0.0518263 | [-0.1345, 0.2381] | [-0.2326, 0.3388] |
| Body Mass | Direct | -0.0776849 | [-0.2827, 0.1226] | [-0.3962, 0.2338] |
| Body Mass | Indirect | 0.0250892 | [-0.0693, 0.1244] | [-0.125, 0.1827] |
| Body Mass | Total | -0.0508449 | [-0.2356, 0.1328] | [-0.3348, 0.2351] |
| Warm Rearing (30°C) | Direct | -0.0078442 | [-0.4697, 0.4416] | [-0.7177, 0.7028] |

| | | | | |
|---|---|---|---|---|
| Warm Rearing (30°C) | Indirect | 0.0027137 | [-0.0715, 0.0824] | [-0.1366, 0.155] |
| Warm Rearing (30°C) | Total | -0.0032679 | [-0.4724, 0.4572] | [-0.7179, 0.7052] |
| Cold Rearing (10°C) | Direct | -0.2534004 | [-0.7424, 0.2304] | [-0.9824, 0.4884] |
| Cold Rearing (10°C) | Indirect | 0.0014471 | [-0.0772, 0.0876] | [-0.1472, 0.1604] |
| Cold Rearing (10°C) | Total | -0.2461172 | [-0.7491, 0.2438] | [-0.9866, 0.5104] |

```
#save_kable(week3ResultsScaled, "../tables/threeWeekResults_3Scaled.html")
```

Similar to our analysis of data from three week old quail, we now repeat our model using tarsus length residuals in place of tarsus length as our predictor of thermal physiology in these quail. Priors from our initial analysis of metabolic slopes across eight week old individuals are retained.

```
functionModel8WeeksResidual <-
  brm(
    data = slopeData %>%
      filter(week == "8") %>%
      mutate(
        mass = mass - mean(mass, na.rm = T),
        pretreatment = ifelse(pretreatment == "cold", "A",
          ifelse(pretreatment == "neutral", "B", "C")
        )
      ) %>%
      mutate(pretreatment = factor(pretreatment, levels = c("B", "A", "C"))) %>%
      drop_na() %>%
      merge(., data %>%
        select(ring, "batch" = exp) %>%
        distinct(),
      by = "ring", all.x = TRUE
      ) %>%
    mutate(
      "residualTarsus" =
        residuals(
          brm(tarsusLengthMean ~ mass,
            data = .,
            prior = c(
              set_prior("skew_normal(0, 0.25, 5)", class = "b"),
              set_prior("normal(30, 2.5)", class = "Intercept")
            ),
            family = "gaussian",
            iter = 50000, warmup = 10000,
            cores = 1, chains = 4, thin = 20,
            silent = TRUE, refresh = 0
          ), robust = TRUE
        )[, "Estimate"]
    ),
    family = "gaussian",
      bf(mass ~ pretreatment + (1 | batch)) +
      bf(residualTarsus ~ pretreatment + (1 | batch)) +
      bf(rawSlope ~ residualTarsus + mass + pretreatment + (1 | batch)) +
      set_rescor(FALSE),
    prior = c(
      set_prior("normal(0, 10)",
        class = "Intercept",
        resp = "mass"
      ),
      set_prior("normal(0, 25)",
        class = "b",
        coef = "pretreatmentA", resp = "mass"
      ),
      set_prior("normal(0, 25)",
```

```
        class = "b",
        coef = "pretreatmentC", resp = "mass"
      ),
      set_prior("exponential(2.5)",
        class = "sd",
        group = "batch", resp = "mass"
      ),
      set_prior("exponential(0.15)",
        class = "sigma",
        resp = "mass"
      ),
      set_prior("normal(0, 2.5)",
        class = "Intercept",
        resp = "residualTarsus"
      ),
      set_prior("normal(0, 2.5)",
        class = "b",
        coef = "pretreatmentA", resp = "residualTarsus"
      ),
      set_prior("normal(0, 2.5)",
        class = "b",
        coef = "pretreatmentC", resp = "residualTarsus"
      ),
      set_prior("exponential(2)",
        class = "sd",
        group = "batch", resp = "residualTarsus"
      ),
      set_prior("exponential(1)",
        class = "sigma",
        resp = "residualTarsus"
      ),
      set_prior("normal(0, 0.01)",
        class = "Intercept",
        resp = "rawSlope"
      ),
      set_prior("normal(0, 0.025)",
        class = "b",
        coef = "pretreatmentA", resp = "rawSlope"
      ),
      set_prior("normal(0, 0.025)",
        class = "b",
        coef = "pretreatmentC", resp = "rawSlope"
      ),
      set_prior("normal(0, 0.001)",
        class = "b",
        coef = "mass", resp = "rawSlope"
      ),
      set_prior("normal(0, 0.0025)",
        class = "b",
        coef = "residualTarsus", resp = "rawSlope"
      ),
      set_prior("exponential(50)",
        class = "sd",
        group = "batch", resp = "rawSlope"
      ),
      set_prior("exponential(10)",
        class = "sigma",
        resp = "rawSlope"
      )
    ),
    iter = 50000, warmup = 10000, cores = 4, chains = 4, thin = 20,
    control = list(adapt_delta = .97, max_treedepth = 14),
    silent = TRUE, refresh = 0,
    file = "./models/eightWeekFunctionModelResidual.Rds"
  )

# Quick diagnostics
```

```
chainCheck(functionModel8WeeksResidual)

## Rhat range: 1 - 1.001
## Neff/N range: 0.878 - 1.024

pp_check2(functionModel8WeeksResidual,
  resp = "rawSlope",
  xlab = "Metabolic Slope\n(fold metabolism at thermoneutrality/°C)"
)
```

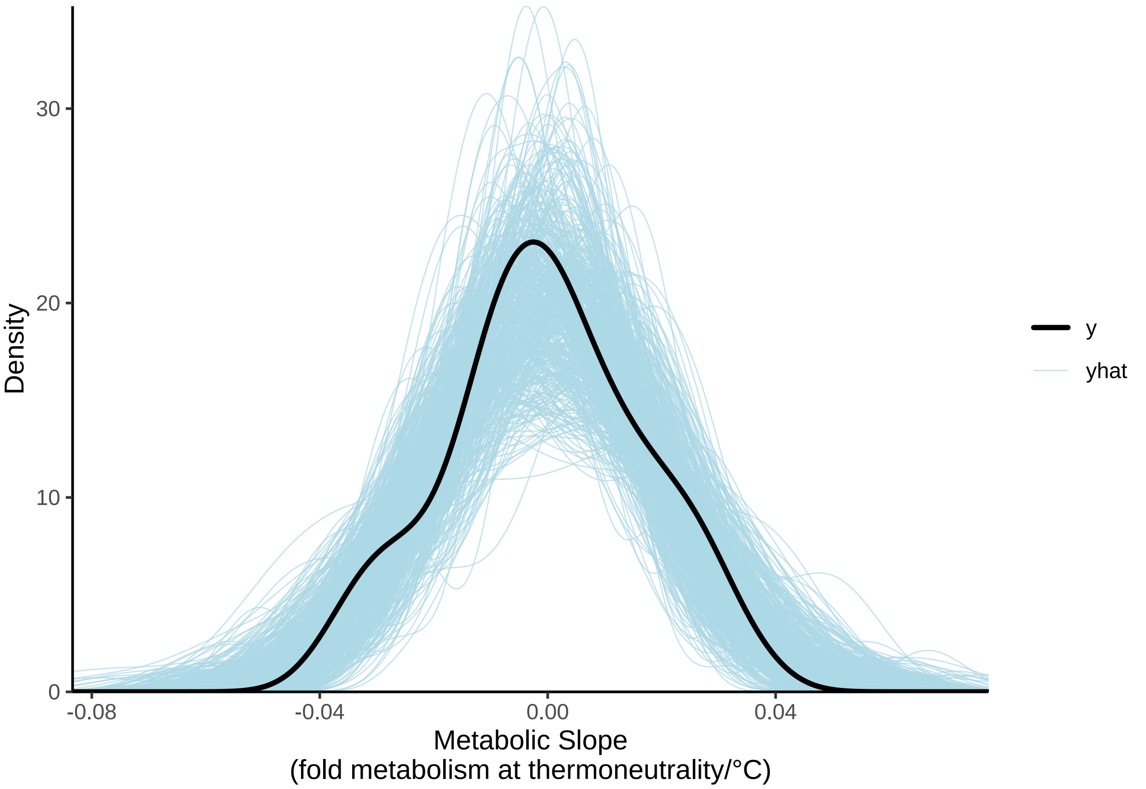


***Figure 45:*** *Posterior predictive check for a Bayesian path analysis predicting metabolic slopes (fold metabolism at thermoneutrality/°C) of eight week old Japanese quail by morphometry and thermal history. Metabolic slopes are measured in the cold (10°C - 30°C). Light blue lines represent densities of resting metabolism values as drawn from model posteriors. The dark blue line represent the true density of resting metabolism values. Clear overlap between the dark blue and light blue lines indicates strong model fit. Morphometry measures include body mass (g) and tarsus length residuals.*

```
# Clean overlap

lbs <- c("Mass ~ Cold Rearing")
names(lbs) <- c("b_mass_pretreatmentA")
p1 <- mcmc_acf(functionModel8WeeksResidual,
  pars = "b_mass_pretreatmentA", lags = 10,
  facet_args = list(labeller = labeller(Parameter = lbs))
) +
  theme_classic()

lbs <- c("Mass ~ Warm Rearing")
names(lbs) <- c("b_mass_pretreatmentC")
p2 <- mcmc_acf(functionModel8WeeksResidual,
  pars = "b_mass_pretreatmentC", lags = 10,
  facet_args = list(labeller = labeller(Parameter = lbs))
) +
```

```
  theme_classic()

lbs <- c("Tarsus ~ Cold Rearing")
names(lbs) <- c("b_residualTarsus_pretreatmentA")
p3 <- mcmc_acf(functionModel8WeeksResidual,
  pars = "b_residualTarsus_pretreatmentA", lags = 10,
  facet_args = list(labeller = labeller(Parameter = lbs))
) +
  theme_classic()

lbs <- c("Tarsus ~ Warm Rearing")
names(lbs) <- c("b_residualTarsus_pretreatmentC")
p4 <- mcmc_acf(functionModel8WeeksResidual,
  pars = "b_residualTarsus_pretreatmentC", lags = 10,
  facet_args = list(labeller = labeller(Parameter = lbs))
) +
  theme_classic()

lbs <- c("Slope ~ Tarsus")
names(lbs) <- c("b_rawSlope_residualTarsus")
p5 <- mcmc_acf(functionModel8WeeksResidual,
  pars = "b_rawSlope_residualTarsus", lags = 10,
  facet_args = list(labeller = labeller(Parameter = lbs))
) +
  theme_classic()

lbs <- c("Slope ~ Mass")
names(lbs) <- c("b_rawSlope_mass")
p6 <- mcmc_acf(functionModel8WeeksResidual,
  pars = "b_rawSlope_mass", lags = 10,
  facet_args = list(labeller = labeller(Parameter = lbs))
) +
  theme_classic()

lbs <- c("Slope ~\nCold Rearing")
names(lbs) <- c("b_rawSlope_pretreatmentA")
p7 <- mcmc_acf(functionModel8WeeksResidual,
  pars = "b_rawSlope_pretreatmentA", lags = 10,
  facet_args = list(labeller = labeller(Parameter = lbs))
) +
  theme_classic()

lbs <- c("Slope ~\nWarm Rearing")
names(lbs) <- c("b_rawSlope_pretreatmentC")
p8 <- mcmc_acf(functionModel8WeeksResidual,
  pars = "b_rawSlope_pretreatmentC", lags = 10,
  facet_args = list(labeller = labeller(Parameter = lbs))
) +
  theme_classic()

p9 <- functionModel8WeeksResidual$data %>%
  mutate("Res" = residuals(functionModel8WeeksResidual,
                           resp = "rawSlope",
                           robust = TRUE)[, "Estimate"]) %>%
  ggplot(aes(sample = Res)) +
  stat_qq(colour = "grey50") +
  stat_qq_line() +
  xlab("Theoretical") +
  ylab("Sample") +
  theme_classic()

p10 <- functionModel8WeeksResidual$data %>%
  mutate("Fit" = fitted(functionModel8WeeksResidual,
                        resp = "rawSlope",
                        robust = TRUE)[, "Estimate"]) %>%
  ggplot(aes(x = Fit, y = rawSlope)) +
  geom_point(pch = 21, colour = "black", size = 2, fill = "grey90") +
```

```
  geom_smooth(method = "lm", linetype = "dashed",
              colour = "black", se = FALSE) +
  xlab("Fitted Values") +
  ylab("Metabolic Slope (Fold\nMetabolism at Themoneutrality/°C)") +
  theme_classic()

(p1 + p2 + p3) /
  (p4 + p5 + p6) /
  (p7 + p8 + p9) /
  (p10)
```

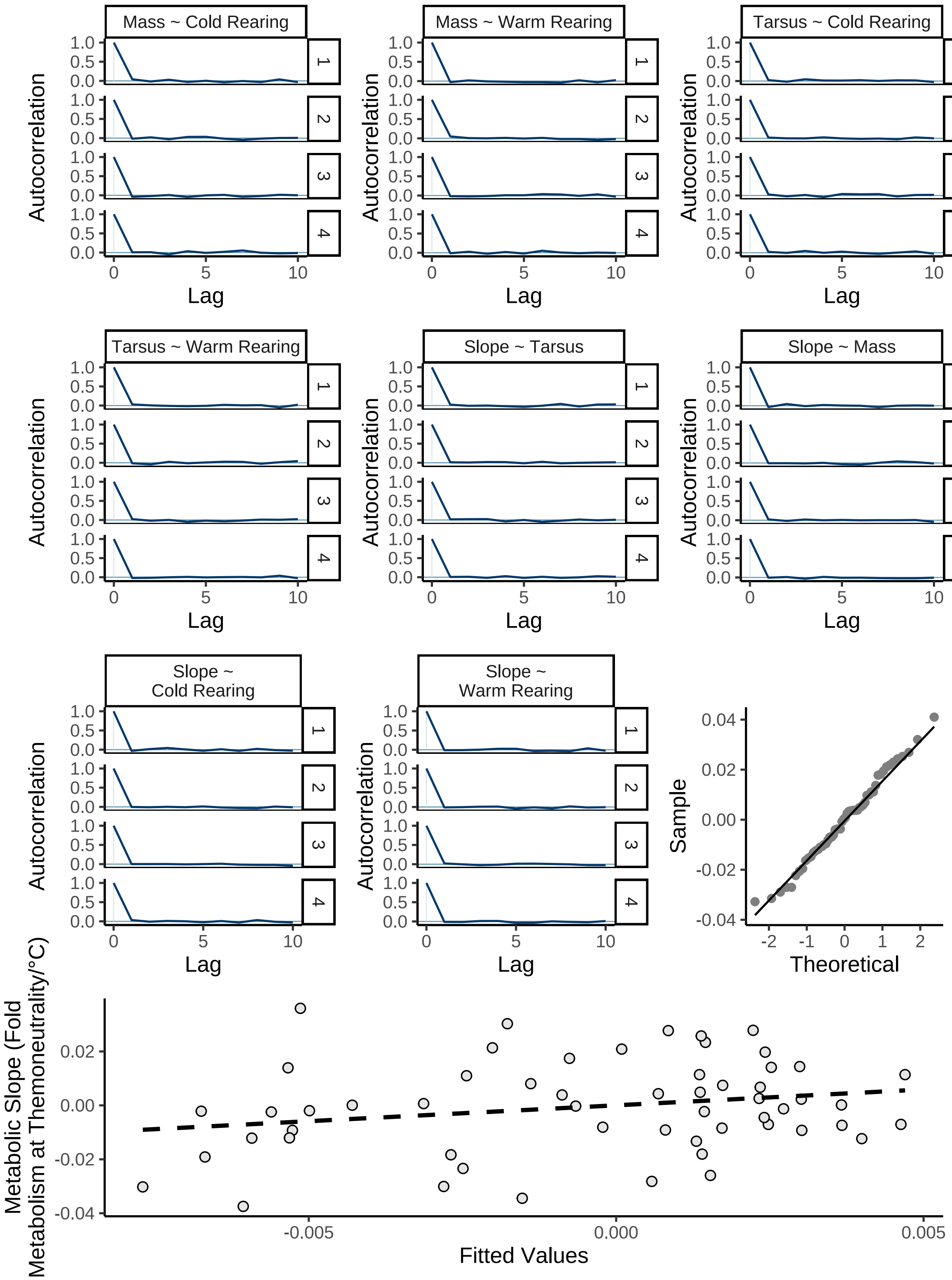


***Figure 46:*** *Diagnostics for a Bayesian path analysis predicting metabolic slopes (fold metabolism at thermoneutrality/°C) of eight week old Japanese quail by morphometry (here, body mass [g] and residual tarsus length [mm]) and thermal history. Metabolic slopes are measured in the cold (10°C - 30°C). Autocorrelation plots display autocorrelation between adjacent Hamiltonian Monte Carlo chain draws, divided by chain. QQ-plot displays sample ordinary residuals by theoretical residual quantiles, and fitted values are displayed against true, mean-centred metabolic slope values.*

```
# Heterogenous and little evidence of autocorrelation.
```

Below, we plot posterior densities per predictor.

```
# Plotting coefficients

  as.data.frame(functionModel8WeeksResidual) %>%
    select(
      "Intercept" = b_rawSlope_Intercept,
      "Residual Tarsus\n(mm)" = b_rawSlope_residualTarsus,
      "Body Mass (g)" = b_rawSlope_mass,
      "Cold Rearing\n(10°C)" = b_rawSlope_pretreatmentA,
      "Warm Rearing\n(30°C)" = b_rawSlope_pretreatmentC,
      "Sigma" = sigma_rawSlope
    ) %>%
    pivot_longer(everything(), names_to = "Par",
                 values_to = "Coefs") %>%
    ggplot(aes(x = Coefs)) +
    facet_wrap(~Par, scales = "free", ncol = 2) +
    geom_density(colour = "black", alpha = 0.5, fill = "grey70") +
    geom_vline(xintercept = 0, linetype = "dashed", colour = "black") +
    ylab("Density") +
    theme_classic() +
    theme(axis.title.x = element_blank())
```

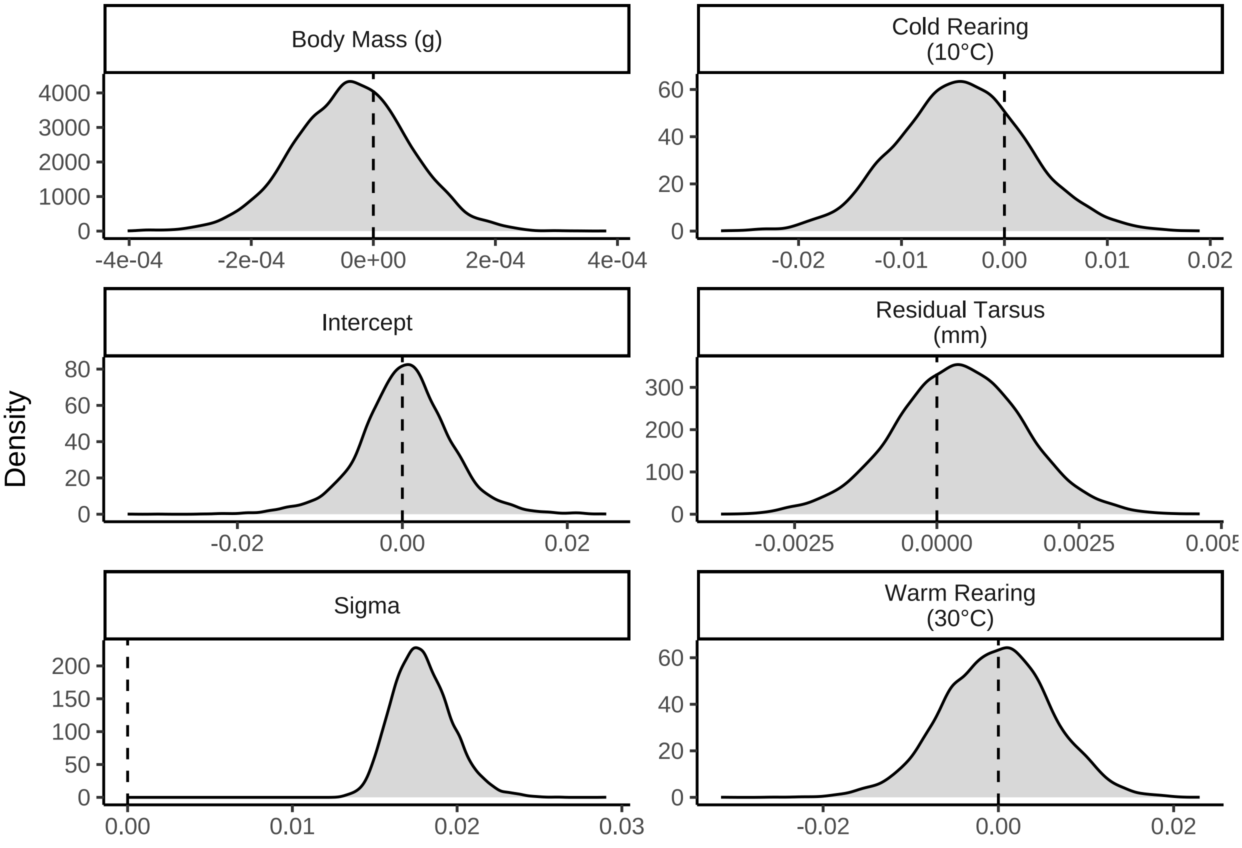


***Figure 47:*** *Posterior densities for model coefficients derived from a Bayesian path analysis predicting metabolic slopes (fold metabolism at thermoneutrality/°C) of eight week old Japanese quail. Metabolic slopes represent those displayed in the cold (10°C - 30°C). Densities are split by their respective response values (indicated with titles). Vertical dashed lines indicate 0.*

```
ggarrange(
  as.data.frame(functionModel8WeeksResidual) %>%
    select(
      "Intercept" = b_residualTarsus_Intercept,
      "Cold Rearing" = b_mass_pretreatmentA,
      "Warm Rearing" = b_mass_pretreatmentC,
      "Sigma" = sigma_mass
    ) %>%
    pivot_longer(everything(), names_to = "Par",
                 values_to = "Coefs") %>%
    ggplot(aes(x = Coefs)) +
    facet_wrap(~Par, scales = "free") +
    geom_density(colour = "black", alpha = 0.5, fill = "grey70") +
    geom_vline(xintercept = 0, linetype = "dashed",
               colour = "black") +
    ylab("Density") +
    theme_classic() +
    theme(axis.title.x = element_blank()) +
    ggtitle("Body Mass (g)"),

  as.data.frame(functionModel8WeeksResidual) %>%
    select(
      "Intercept" = b_residualTarsus_Intercept,
      "Cold Rearing" = b_residualTarsus_pretreatmentA,
      "Warm Rearing" = b_residualTarsus_pretreatmentC,
      "Sigma" = sigma_residualTarsus
    ) %>%
    pivot_longer(everything(), names_to = "Par",
                 values_to = "Coefs") %>%
    ggplot(aes(x = Coefs)) +
    facet_wrap(~Par, scales = "free") +
    geom_density(colour = "black", alpha = 0.5, fill = "grey70") +
    geom_vline(xintercept = 0, linetype = "dashed",
               colour = "black") +
    ylab("Density") +
    theme_classic() +
    theme(axis.title.x = element_blank()) +
    ggtitle("Residual Tarsus Length (mm)"),
  ncol = 1
)
```

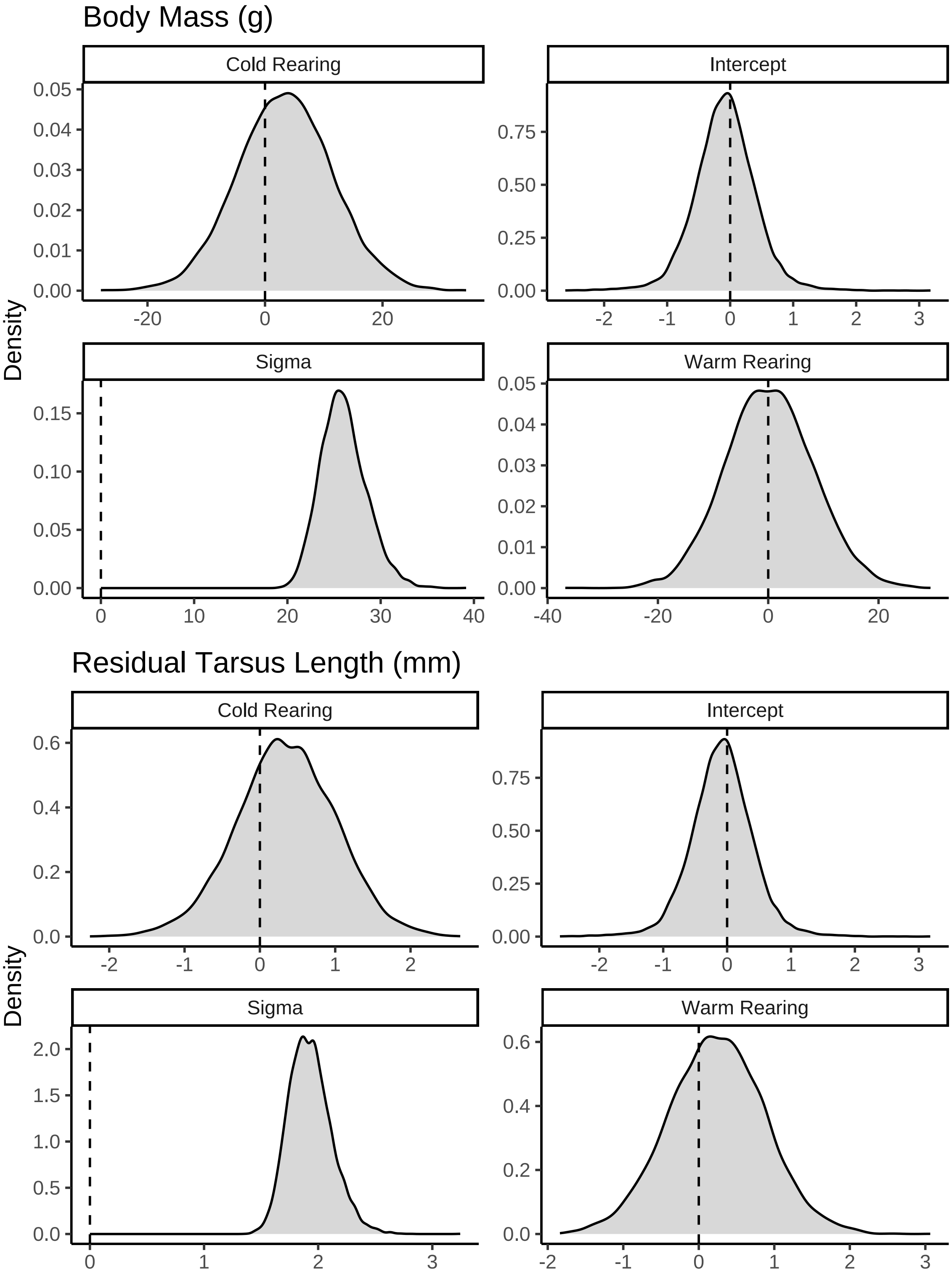


***Figure 48:*** *Continued. Posterior densities for model coefficients derived from a Bayesian path analysis predicting metabolic slopes (fold metabolism at thermoneutrality/°C) of eight week old Japanese quail. Densities are split by their respective response values (indicated with titles). Vertical dashed lines indicate 0.*

```
# Summarising model
```

```
caption <- paste0("Results of a Bayesian path analysis ",
                  "predicting metabolic slopes (fold metabolism  ",
                  "at thermoneutrality/°C) ",
                  "of eight week old Japanese quail by morphology and ",
                  "thermal history (here, rearing temperature). ",
                  "Tarsus length is relativised by body mass (tarsus ",
                  "length residuals) as estimated from a Bayesian ",
                  "linear model. Metabolic slopes are measured in the cold ",
                  "(10°C - 30°C). Credible intervals are quantile ",
                  "intervals. BF indicates ",
                  "Bayes Factors for each model parameter."
)

functionModel8WeeksResidualTable <-
  as.data.frame(functionModel8WeeksResidual) %>%
  summarise_all(., .funs = median) %>%
  pivot_longer(everything(),
    names_to = "Parameter",
    values_to = "Estimate"
  ) %>%
  merge(., quantileCIs(functionModel8WeeksResidual, cis = c(50, 95)),
    by = "Parameter", all.x = TRUE
  ) %>%
  filter(grepl("b_|sd_", Parameter)) %>%
  rowwise() %>%
  mutate("BF" = ifelse(Estimate < 0,
    (2 * mean(as.data.frame(
      functionModel8WeeksResidual
    )[, Parameter] <= 0)) /
      (2 * mean(as.data.frame(
        functionModel8WeeksResidual
      )[, Parameter] >= 0)),
    (2 * mean(as.data.frame(
      functionModel8WeeksResidual
    )[, Parameter] >= 0)) /
      (2 * mean(as.data.frame(
        functionModel8WeeksResidual
      )[, Parameter] <= 0))
  )) %>%
  ungroup() %>%
  mutate(
    "Estimate" = round(Estimate, digits = 4),
    "BF" = round(BF, digits = 4),
    "N" = nrow(functionModel8WeeksResidual$data)
  ) %>%
  mutate("Parameter" = ifelse(grepl("b_", Parameter),
    gsub("b_", "", Parameter),
    gsub(
      "Intercept", "batch",
      gsub(".*__", "", Parameter)
    )
  )) %>%
  mutate(
    "Response" = gsub("_.*", "", Parameter),
    "Parameter" = gsub(".*_", "", Parameter)
  ) %>%
  merge(., tribble(
    ~Response, ~response, ~level,
    "mass", "Body Mass (g)", "A",
    "rawSlope",
    "Metabolic Slope", "D",
    "residualTarsus", "Residiual Tarsus Length (mm)",
    "B"
  ),
  by = "Response"
  ) %>%
  merge(., tribble(
```

```
    ~Parameter, ~parameter, ~number,
    "Intercept", "Intercept", "1",
    "mass", "Body Mass (g)", "4",
    "residualTarsus", "Residual Tarsus Length (mm)", "5",
    "pretreatmentA", "Cold Rearing", "2",
    "pretreatmentC", "Warm Rearing", "3",
    "batch", "Egg Batch [mu]", "6"
  ),
  by = "Parameter"
  ) %>%
  mutate(
    `50\\% CI` = paste0("(", paste(
      round(Low_CI_50, digits = 4),
      round(High_CI_50, digits = 4),
      sep = ", "
    ), ")"),
    `95\\% CI` = paste0("(", paste(
      round(Low_CI_95, digits = 4),
      round(High_CI_95, digits = 4),
      sep = ", "
    ), ")")
  ) %>%
  select(-c(Low_CI_50, High_CI_50, Low_CI_95, High_CI_95)) %>%
  select(
    "Response" = "response", "Parameter" = "parameter", N,
    Estimate, `50\\% CI`, `95\\% CI`, BF, level, number
  ) %>%
  arrange(level, number) %>%
  select(-c(level, number)) %>%
  kbl(.,
    longtable = T, booktabs = T, format = "latex", escape = FALSE,
    caption = caption
  ) %>%
  column_spec(column = c(1:2), width = "2.2cm") %>%
  column_spec(column = c(3:10), width = "1.9cm") %>%
  kable_styling(latex_options = "striped")

functionModel8WeeksResidualTable
```

***Table 22:*** *Results of a Bayesian path analysis predicting metabolic slopes (fold metabolism at thermoneutrality/°C) of eight week old Japanese quail by morphology and thermal history (here, rearing temperature). Tarsus length is relativised by body mass (tarsus length residuals) as estimated from a Bayesian linear model. Metabolic slopes are measured in the cold (10°C - 30°C). Credible intervals are quantile intervals. BF indicates Bayes Factors for each model parameter.*

| Response | Parameter | N | Estimate | 50% CI | 95% CI | BF |
|---|---|---|---|---|---|---|
| Body Mass (g) | Intercept | 56 | -5.0857 | (-8.1612, -1.9698) | (-13.8419, 3.7883) | 6.5758 |
| Body Mass (g) | Cold Rearing | 56 | 3.3954 | (-2.0485, 8.8305) | (-12.3678, 19.6113) | 1.9685 |
| Body Mass (g) | Warm Rearing | 56 | 0.1320 | (-5.1477, 5.5695) | (-15.2129, 16.2373) | 1.0222 |
| Body Mass (g) | Egg Batch [mu] | 56 | 0.2826 | (0.1185, 0.5554) | (0.0111, 1.4649) | Inf |
| Residuial Tarsus Length (mm) | Intercept | 56 | -0.0800 | (-0.3752, 0.2067) | (-1.034, 0.8734) | 1.3592 |
| Residuial Tarsus Length (mm) | Cold Rearing | 56 | 0.3424 | (-0.0811, 0.7942) | (-0.9836, 1.6243) | 2.4072 |
| Residuial Tarsus Length (mm) | Warm Rearing | 56 | 0.2376 | (-0.1971, 0.663) | (-1.0182, 1.4988) | 1.8319 |
| Residuial Tarsus Length (mm) | Egg Batch [mu] | 56 | 0.2575 | (0.1135, 0.4938) | (0.0107, 1.3615) | Inf |
| Metabolic Slope | Intercept | 56 | 0.0004 | (-0.0029, 0.0037) | (-0.0113, 0.0116) | 1.1350 |

| | | | | | | |
|---|---|---|---|---|---|---|
| Metabolic Slope | Cold Rearing | 56 | -0.0042 | (-0.0084, 0) | (-0.0165, 0.0085) | 2.9880 |
| Metabolic Slope | Warm Rearing | 56 | -0.0001 | (-0.0044, 0.004) | (-0.0125, 0.0118) | 1.0207 |
| Metabolic Slope | Body Mass (g) | 56 | 0.0000 | (-1e-04, 0) | (-2e-04, 1e-04) | 1.8031 |
| Metabolic Slope | Residual Tarsus Length (mm) | 56 | 0.0004 | (-4e-04, 0.0012) | (-0.0019, 0.0026) | 1.7501 |
| Metabolic Slope | Egg Batch [mu] | 56 | 0.0056 | (0.0025, 0.011) | (3e-04, 0.0315) | Inf |

```
#save_kable(resistanceResidual8WeeksTable,
# "../tables/residualTarsusAnalysisResults8Weeks.html")
```

#### Testing effects of morphometry on metabolic slopes in the heat

Here, we evaluate the degree to which body size and appendage length (i.e. body mass and tarsus length respectively) influences metabolic responses to heat in our captive reared Japanese quail. To achieve this end, we first calculate metabolic slopes displayed in response to a heat challenge (30°C - 40°C) on an individual basis. These slopes are then regressed against morphometry, while controlling for thermal history (i.e. our experimental rearing treatment. Contrasting our approach used to quantify metabolic slopes in the cold, slopes are here calculated manually as fold increase in resting metabolism between 30°C and 40°C divided by 10°C. After calculation, data are visually inspected for erroneous values before proceeding to any further analysis.

```
# Beginning with metabolic data from birds three weeks of age
# Subsetting data to include only metabolism measurements
# at and above 30°C.

birdVector <- merge(
  all %>%
    drop_na(VO2) %>%
    filter(week == 3 & Ta == 30) %>%
    select(ring) %>%
    distinct(),
  all %>%
    drop_na(VO2) %>%
    filter(week == 3 & Ta == 40) %>%
    select(ring) %>%
    distinct(),
  by = "ring", all = FALSE
) %>% pull(ring)

modDataHot <- subset(all, week == 3 & Ta > 25 &
                       ring %in% birdVector)

# Calculating metabolic slopes manually.

slopeDataHot3 <- modDataHot %>%
  select(ring, Ta, VO2) %>%
  pivot_wider(id_cols = ring, names_from = Ta,
              values_from = VO2) %>%
  mutate(
    slope = (`40`/`30`)/10
  ) %>%
  select(ring, slope) %>%
  merge(modDataHot %>%
    select(
      ring, sex, exp, week, pretreatment,
      posttreatment, treatment, mass, wingLength,
      tarsusLengthMean,
    ) %>%
    distinct(), ., by = "ring")
```

```
# Checking for peculiar values.

slopeDataHot3 %>%
  ggplot(aes(x = 1:nrow(.), y = slope)) +
  geom_rect(
    colour = "black", fill = "grey80", alpha = 0.5,
    aes(
      xmin = -Inf, xmax = Inf,
      ymin = mean(slope, na.rm = T) -
        3.5 * sd(slope, na.rm = T),
      ymax = mean(slope, na.rm = T) +
        3.5 * sd(slope, na.rm = T)
    )
  ) +
  geom_point() +
  xlab("Sample Number") +
  ylab("Metabolic Slope\n(Fold Metabolsim at Thermoneutrality/°C)") +
  theme_classic()
```

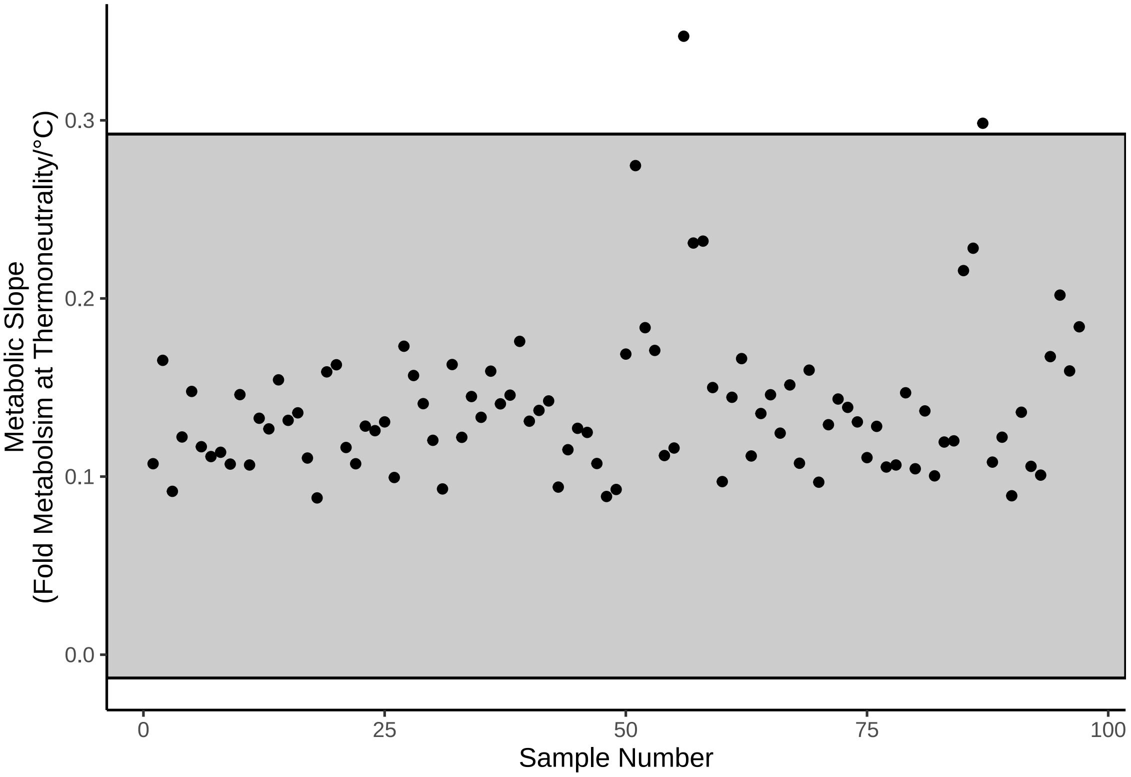


***Figure 49:*** *Cleveland dotplot displaying metabolic slope values of three week old Japanese quail at 30°C - 40°C. Dots represent raw values and the grey rectangle represents the range covered by mean metabolic slope +/- 3.5x the standard deviation.*

Two values appears particularly high and are investigated further below.

```
caption <- paste0("Potential metabolic slope outliers ",
                  "among three week old Japanese quail with ",
                  "metabolism measured at ambient temperatures ",
                  "between 30°C and 40°C."
)

slopeDataHot3 %>%
  filter(slope > 0.29) %>%
```

```
  merge(., modDataHot %>%
    select(ring, Ta, VO2),
  by = "ring", all.x = TRUE
  ) %>%
  select(
    "Bird Identity" = ring,
    "Rearing Conditions" = pretreatment,
    "Mass (g)" = mass,
    "Tarsus Length (mm)" = tarsusLengthMean,
    "Ambient Temperature (°C)" = Ta,
    "Resting Metabolism (mL O\\textsubscript{2}/min)" =
      VO2,
    "Metabolic Slope\n(Fold RMR at thermoneutrality/°C)" =
      slope
  ) %>%
    kbl(.,
    longtable = T, booktabs = T, format = "latex", escape = FALSE,
    caption = caption
  ) %>%
  column_spec(colum = c(1:10), width = "1.9cm") %>%
  kable_styling(latex_options = "striped")
```

***Table 23:*** *Potential metabolic slope outliers among three week old Japanese quail with metabolism measured at ambient temperatures between 30°C and 40°C.*

| Bird Identity | Rearing Conditions | Mass (g) | Tarsus Length (mm) | Ambient Temperature (°C) | Resting Metabolism (mL $O_2$/min) | Metabolic Slope (Fold RMR at thermoneutrality/°C) |
|---|---|---|---|---|---|---|
| LLW26 | warm | 155.0 | 36.7683 | 40 | 13.091570 | 0.3472285 |
| LLW26 | warm | 155.0 | 36.7683 | 30 | 3.770304 | 0.3472285 |
| RLG28 | warm | 124.7 | 36.8043 | 40 | 9.718958 | 0.2983607 |
| RLG28 | warm | 124.7 | 36.8043 | 30 | 3.257453 | 0.2983607 |

Resting metabolism values for these individuals are quite high at 40°C and quite low and 30°C. We next plot how these values fit within the distribution of others.

```
modDataHot %>%
  ggplot(aes(x = VO2, y = Ta)) +
  geom_line(aes(group = ring),
            colour = "grey70", alpha = 0.5) +
  geom_line(
    data = slopeDataHot3 %>%
      filter(ring == "LLW26") %>%
      merge(., modDataHot %>%
        select(ring, Ta, VO2),
      by = "ring", all.x = TRUE
      ),
    colour = "black", alpha = 0.5, size = 1.5
  ) +
  geom_line(
    data = slopeDataHot3 %>%
      filter(ring == "RLG28") %>%
      merge(., modDataHot %>%
        select(ring, Ta, VO2),
      by = "ring", all.x = TRUE
      ),
    colour = "black", alpha = 0.5, size = 1.5
  ) +
  geom_point(pch = 21, colour = "black",
             fill = "lightblue", alpha = 0.5) +
  geom_errorbarh(
```

```
    data = modDataHot %>%
      group_by(Ta) %>%
      summarise(
        "vo2" = mean(VO2, na.rm = T),
        "SE" = sd(VO2, na.rm = T) / sqrt(n())
      ) %>%
      rename("VO2" = vo2),
    aes(xmin = VO2 - 1.96 * SE, xmax = VO2 + 1.96 * SE, y = Ta),
    colour = "black", height = 1
  ) +
  geom_point(
    data = modDataHot %>%
      group_by(Ta) %>%
      summarise("mean" = mean(VO2, na.rm = T)),
    aes(x = mean, y = Ta), pch = 21,
    colour = "black", fill = "lightblue", size = 5
  ) +
  scale_y_continuous(limits = c(28, 43), breaks = c(30, 40)) +
  xlab(TeX('$\\overset{Resting~Metabolism}{(mL~O_{2}/min)}$')) +
  ylab("Ambient Temperature (°C)") +
  theme_classic()
```

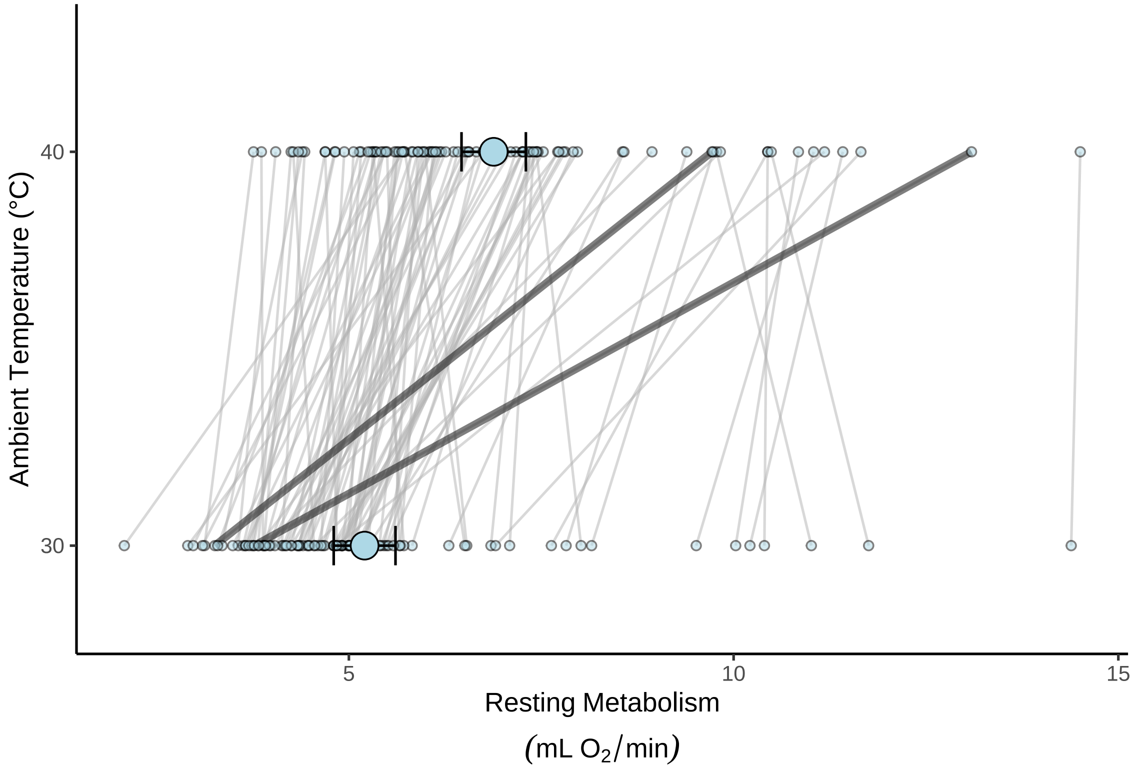


***Figure 50:*** *Spread of resting metabolism values among three week old Japanese quail at 30°C and 40°C. Small dots represent raw values and lines connect dots for individual birds. The black line indicates possible metabolic slope (fold metabolism at thermoneutrality/°C) outliers. Large dots represent mean values at given ambient temperatures and errorbars indicate means +/- 1.96x the standard error.*

```
# Quite extreme. Checking how many standard deviations from
# the mean slope values lies.

cat(paste0(
  "Slope values for LLW26 lies ",
  (subset(slopeDataHot3, ring == "LLW26")$slope -
    mean(slopeDataHot3$slope, na.rm = T)) /
```

```
    sd(slopeDataHot3$slope, na.rm = T),
  " standard deviations above the mean"
))
```

```
## Slope values for LLW26 lies 4.76155057760552 standard deviations above the mean
```

```
cat(paste0(
  "Slope values for RLG28 lies ",
  (subset(slopeDataHot3, ring == "RLG28")$slope -
    mean(slopeDataHot3$slope, na.rm = T)) /
    sd(slopeDataHot3$slope, na.rm = T),
  " standard deviations above the mean"
))
```

```
## Slope values for RLG28 lies 3.64096241661706 standard deviations above the mean
```

Although these metabolic slopes are relatively large with respect to mean values, we retain them in the absence of other reasons for removal.

Below, we summarise metabolic slope values at their means, medians, and standard deviations.

```
caption = paste0("Summary of metabolic parameters estimated for ",
                 "captive-reared Japanese quail above ",
                 "thermoneutrality (30°C).")

slopeDataHot3 %>%
  group_by(week) %>%
  summarise(
    "Mean" = mean(slope, na.rm = T),
    "Median" = median(slope, na.rm = T),
    "SD" = sd(slope, na.rm = T)
  ) %>%
  mutate(
    "CV" = (SD / Mean) * 100,
    "Parameter" = "Metabolic Slope (fold metabolism at thermoneutrality/°C)"
  ) %>%
  select(Parameter,
    "Age (weeks)" = week,
    Mean, Median, "S.D." = SD, "CV (\\%)" = CV
  ) %>%
  kbl(.,
    longtable = T, booktabs = T, format = "latex",
    caption = caption, escape = FALSE
  ) %>%
  column_spec(column = c(1:10), width = "2.1cm") %>%
  kable_styling(latex_options = "striped")
```

***Table 24:*** *Summary of metabolic parameters estimated for captive-reared Japanese quail above thermoneutrality (30°C).*

| Parameter | Age (weeks) | Mean | Median | S.D. | CV (%) |
|---|---|---|---|---|---|
| Metabolic Slope (fold metabolism at thermoneutrality/°C) | 3 | 0.1395815 | 0.1307003 | 0.0436091 | 31.24275 |

Next, distributions of, and trends in, our data are broadly explored to better inform model construction.

```
# Visualising data

p1 <- ggplot(slopeDataHot3, aes(x = slope)) +
  geom_density(colour = "black", fill = "grey50", alpha = 0.5) +
```

```
  theme_classic() +
  xlab("Metabolic Slope\n(Fold Metabolism at Thermoneutrality/°C)") +
  ylab("Density")

p2 <- ggplot(
  modDataHot %>%
    mutate(
      "tarsusGroup" =
        ifelse(tarsusLengthMean <
          mean(tarsusLengthMean, na.rm = T) -
            sd(tarsusLengthMean, na.rm = T),
        "Small",
        ifelse(tarsusLengthMean >
          mean(tarsusLengthMean, na.rm = T) +
            sd(tarsusLengthMean, na.rm = T),
        "Large", "Average"
        )
        )
    ) %>%
    mutate(
      tarsusGroup =
        factor(tarsusGroup,
          levels = c("Small", "Average", "Large")
        )
    ) %>%
    mutate(
      pretreatment =
        ifelse(pretreatment == "cold", "Cold\n(10°C)",
          ifelse(pretreatment == "neutral", "Mild\n(20°C)",
            "Warm\n(30°C)"
          )
        )
    ),
  aes(x = Ta, y = VO2, group = ring, colour = tarsusGroup)
) +
  facet_wrap(~pretreatment) +
  geom_line(aes(size = tarsusGroup)) +
  scale_size_manual(
    values = c("Small" = 0.5, "Average" = 1, "Large" = 2),
    labels = c("< Mean - 1SD", "Mean", "> Mean + 1SD"),
    name = "Relative Tarsus\nLength"
  ) +
  scale_colour_manual(
    values = c(
      "Small" = "grey80",
      "Average" = "grey30",
      "Large" = "black"
    ),
    name = "Relative Tarsus\nLength",
    labels = c("< Mean - 1SD", "Mean", "> Mean + 1SD"),
  ) +
  scale_x_continuous(breaks = c(30, 40)) +
  theme_classic() +
  theme(legend.position = "bottom") +
  xlab("Ambient Temperature (°C)") +
  ylab(TeX(
    "$\\overset{Resting~Metabolism}{(mL~O_{2}/min)}$"
  ))

p3 <- ggplot(
  modDataHot %>%
    mutate(
      "massGroup" =
        ifelse(mass <=
          mean(mass, na.rm = T) -
            sd(mass, na.rm = T),
        "Small",
```

```
        ifelse(mass >=
          mean(mass, na.rm = T) +
            sd(mass, na.rm = T),
        "Large", "Average"
        )
        )
    ) %>%
    mutate(
      massGroup =
        factor(massGroup,
          levels = c("Small", "Average", "Large")
        )
    ) %>%
    mutate(
      pretreatment =
        ifelse(pretreatment == "cold", "Cold\n(10°C)",
          ifelse(pretreatment == "neutral", "Mild\n(20°C)",
            "Warm\n(30°C)"
          )
        )
    ),
  aes(x = Ta, y = VO2, group = ring, colour = massGroup)
) +
  facet_wrap(~pretreatment) +
  geom_line(aes(size = massGroup)) +
  scale_size_manual(
    values = c("Small" = 0.5, "Average" = 1, "Large" = 2),
    name = "Relative Body\nMass",
    labels = c("< Mean - 1SD", "Mean", "> Mean + 1SD"),
  ) +
  scale_colour_manual(
    values = c(
      "Small" = "grey80",
      "Average" = "grey30",
      "Large" = "black"
    ),
    name = "Relative Body\nMass",
    labels = c(
      "< Mean - 1SD", "Mean",
      "> Mean + 1SD"
    ),
  ) +
  scale_x_continuous(breaks = c(30, 40)) +
  theme_classic() +
  theme(legend.position = "bottom") +
  xlab("Ambient Temperature (°C)") +
  ylab(TeX(
    "$\\overset{Resting~Metabolism}{(mL~O_{2}/min)}$"
  ))

(p1 / p2 / p3) +
  plot_annotation(tag_levels = "A")
```

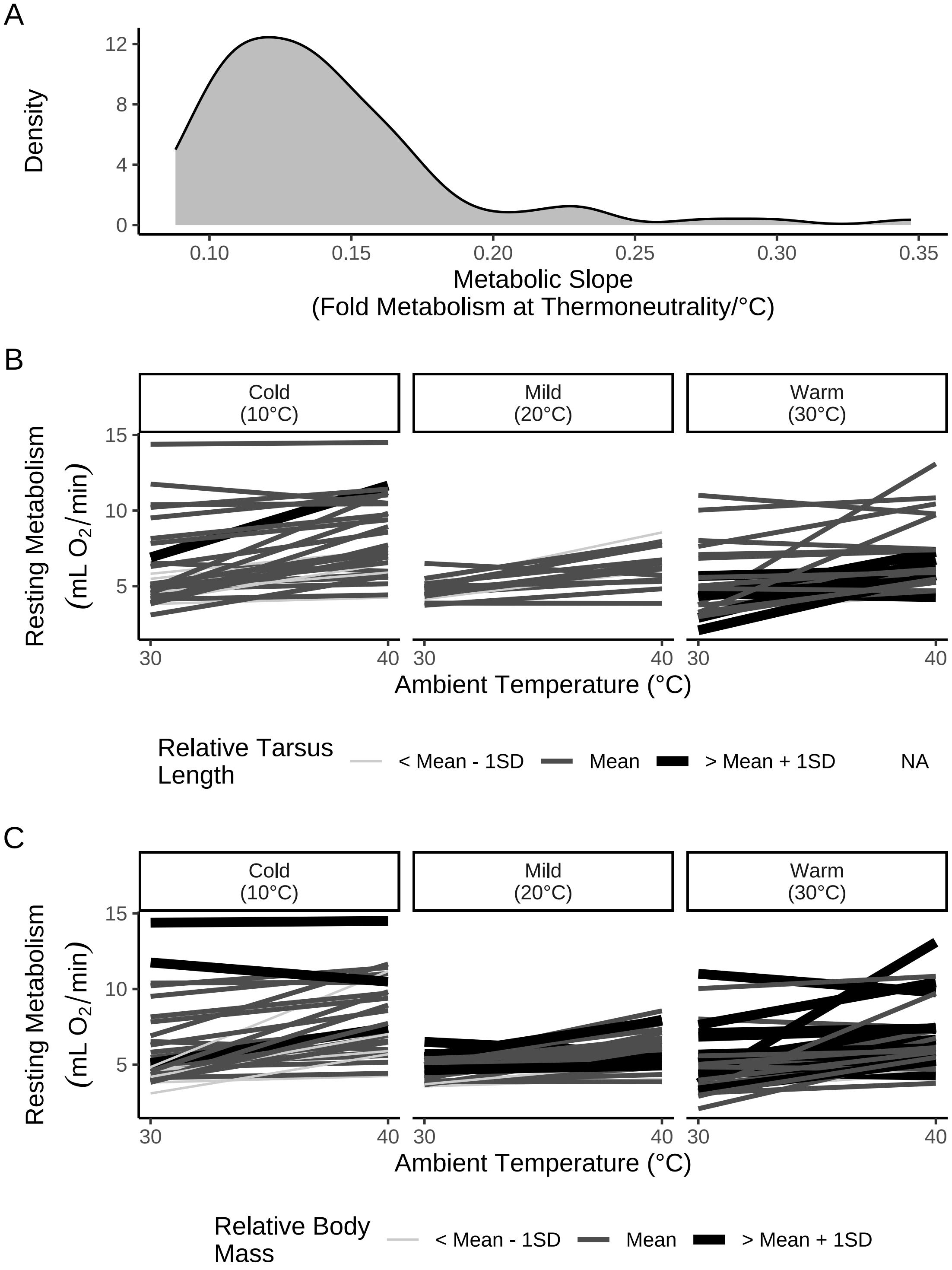


***Figure 51:*** *Thermal physiology of three week old Japanese quail at ambient temperature within and above thermoneutrality (≥ 30°C). Panel A displays density of metabolic slopes at this temperature range. Panels B and C display correlations between ambient temperature and resting metabolism by morphology. In panels B and C, each line represents one individual with line colour and line size scaled by relative size (tarsus length [mm] or body mass [g]); data is divided by rearing conditions up to three months of age.*

```
p1 <- ggplot(slopeDataHot3,
             aes(x = tarsusLengthMean, y = slope)) +
  geom_point(pch = 21, size = 1.5,
             colour = "black", fill = "grey50") +
  geom_smooth(method = "lm", se = FALSE,
              colour = "black", linetype = "dashed") +
  theme_classic() +
  xlab("Tarsus Length (mm)") +
  ylab("Metabolic Slope\n(Fold Metabolism at Thermoneutraltiy/°C)")

p2 <- ggplot(slopeDataHot3,
  aes(x = mass, y = slope)) +
  geom_point(pch = 21, size = 1.5,
             colour = "black", fill = "grey50") +
  geom_smooth(method = "lm", se = FALSE,
              colour = "black", linetype = "dashed") +
  xlab("Body Mass (g)") +
  theme_classic() +
  theme(axis.title.y = element_blank())

p3 <- ggplot(
  slopeDataHot3 %>%
    drop_na(tarsusLengthMean, mass) %>%
    mutate("tarsusResiduals" = residuals(
      brm(tarsusLengthMean ~ mass,
        data = .,
        prior = c(set_prior("gamma(1, 1)",
                            class = "b", coef = "mass")),
        iter = 50000, warmup = 5000, cores = 4, chains = 4,
        silent = TRUE, refresh = 0,
        file = "./models/residualPlotModel3.Rds"
      ), robust = TRUE
    )[, "Estimate"]),
  aes(x = tarsusResiduals, y = slope)
) +
  geom_point(pch = 21, size = 1.5,
             colour = "black", fill = "grey50") +
  geom_smooth(method = "lm", se = FALSE,
              colour = "black", linetype = "dashed") +
  xlab("Residual Tarsus\nLength (mm)") +
  theme_classic() +
  theme(axis.title.y = element_blank())

(p1 + p2)|p3
```

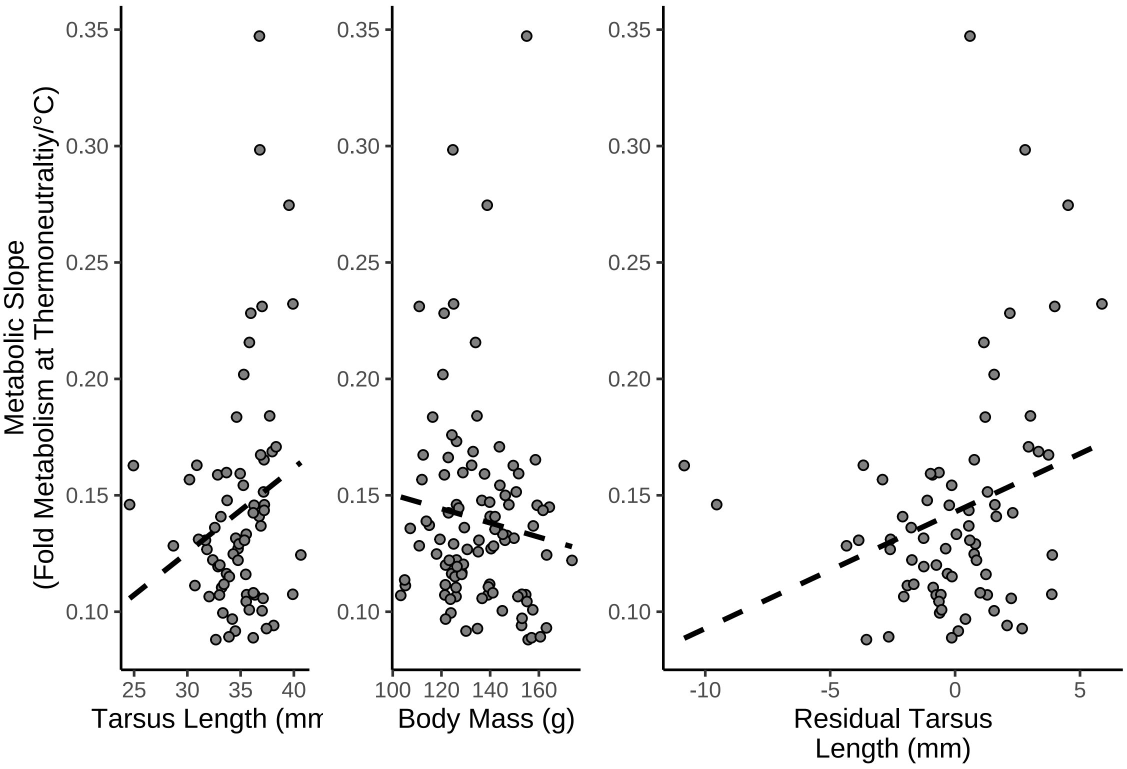


***Figure 52:*** *Raw correlations between morphology and metabolic slopes above thermoneutrality in 3 week old Japanese quail. Metabolic slopes are measured as the fold change in resting metabolism between 30°C and 40°C. Relative tarsus length indicates ordinary residuals from a Bayesian regression of tarsus length (mm) against body mass (g). Dots indicate raw values per individual and dashed lines indicate estimated relationships as predicted by the R package ggplot2 (Wickham, 2011).*

Because flow-rates differed notably between experimental batch 3 and batches 1 and 2 we visualise the spread of metabolic slope values, excluding the extreme value noted above.

```
slopeDataHot3 %>%
  rename("batch" = exp) %>%
  filter(slope < 0.3) %>%
  ggplot(aes(x = slope, fill = batch)) +
  geom_density(colour = "black", alpha = 0.5) +
  scale_fill_manual(values = c("black", "firebrick", "beige"),
                    labels = c("1", "2", "3"),
                    name = "Experimental\nBatch") +
  theme_classic() +
  xlab("Metabolic Slope\n(Fold Metabolism at Thermoneutrality") +
  ylab("Density")
```

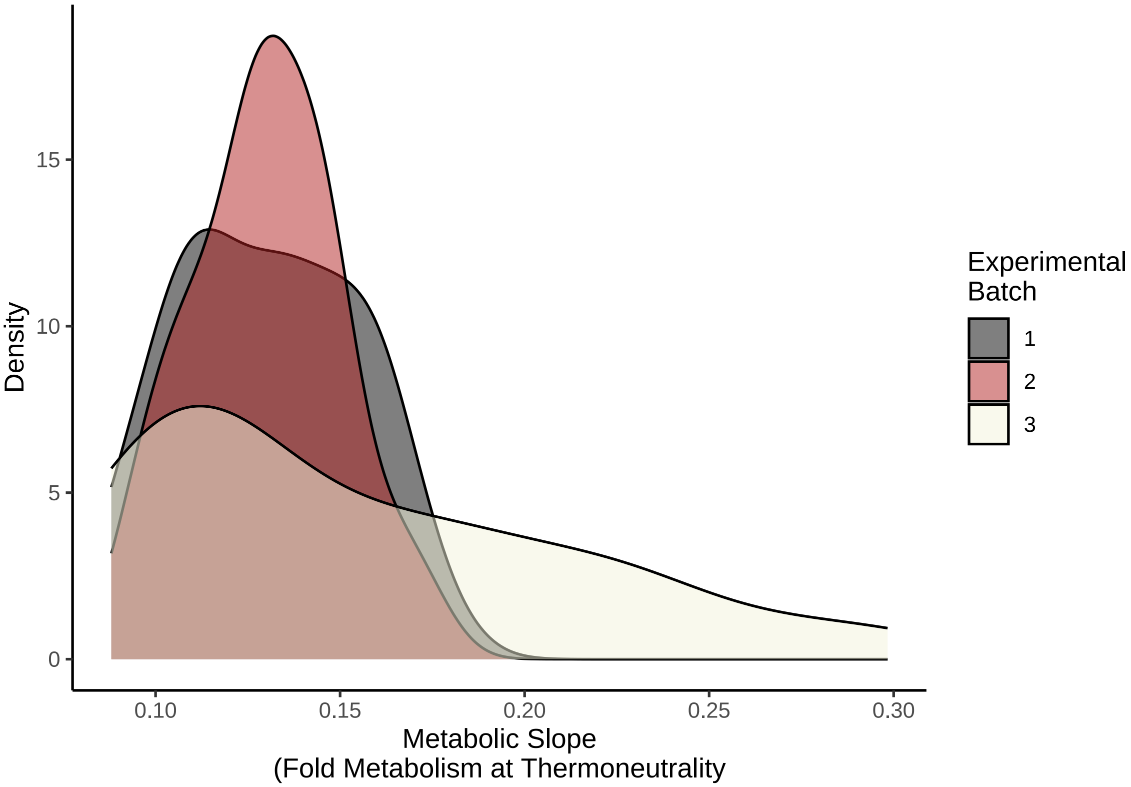


***Figure 53:*** *Densities of metabolic slope values below 30°C from three week old Japanese quail. Densities are divided by experimental batches.*

Clearly, variance in slopes among experimental batch 3 is considerably larger. For this reason, error ($\epsilon$) around metabolic slopes must be permitted to vary by batch.

To test for an effect of morphometry on metabolic slopes of Japanese quail in response to heat, we construct a new path analysis similar to that described for responses to the cold. Models encompassed within this analysis were identical to those previously and are as follows:

$$Body\ Mass_j \sim \beta_{a0} + \beta_{a1} \cdot Cold\ Rearing_j + \beta_{a2} \cdot Warm\ Rearing_j + \mu_{0a} + \epsilon_a$$

$$Tarsus\ Length_j \sim \beta_{b0} + \beta_{b1} \cdot Cold\ Rearing_j + \beta_{b2} \cdot Warm\ Rearing_j + \beta_{b3} \cdot Body\ Mass_j + \mu_{0b} + \epsilon_b$$

and:

$$Metabolic\ Slope_j \sim \beta_{d0} + \beta_{d1} \cdot Cold\ Rearing_j + \beta_{d2} \cdot Warm\ Rearing_j + \beta_{d3} \cdot Body\ Mass_j + \beta_{d4} \cdot Tarsus\ Length_j + \mu_{0c} + \epsilon_{cj}$$

where:

$$\epsilon_{cj} \sim e^{(\tau_0 + \tau_1 \cdot Batch_{2j} + \tau_2 \cdot Batch_{3j})}$$

Variable names largely remain as described above. Further, $\tau_0$ indicates the natural log-transformed error term for experimental batch 1, $\tau_1$ indicates the change in error associated with measures derived from experimental batch 2, and $\tau_2$ indicates the change in error associated with measures derived from experimental batch 3. $Batch_{2j}$ and $Batch_{3j}$ indicate binomial variables pertaining to whether a slope from individual $j$ was derived from experimental batch 2 or 3 respectively (0 = no; 1 = yes).

As before, model coefficients are mean-centred to ease interpretation of model intercepts. Priors for this path analysis were also weakly informed by Persson et al (2022) and Saarela and Heldmaier (1987), but were slightly more broad owing towider uncertainty around how quail at three weeks of age respond to heat exposure. Further, standard deviations for effects of morphometry on metabolic slopes were set to approximately the metabolic slope range divided by the morphometric measurement range. Priors were therefore as follows:

$$\beta_{a0} \sim \mathcal{N}(0, 5)$$
$$\beta_{a1} \sim \mathcal{N}(0, 15)$$
$$\beta_{a2} \sim \mathcal{N}(0, 15)$$
$$\mu_{0a} \sim \exp(2.5)$$
$$\epsilon_a \sim \exp(0.15)$$

$$\beta_{b0} \sim \mathcal{N}(0, 1.5)$$
$$\beta_{b1} \sim \mathcal{N}(0, 2.5)$$
$$\beta_{b2} \sim \mathcal{N}(0, 2.5)$$
$$\beta_{b3} \sim S\mathcal{N}(0, 0.25, 5)$$
$$\mu_{0b} \sim \exp(2)$$
$$\epsilon_b \sim \exp(1)$$

$$\beta_{c0} \sim \mathcal{N}(0, 0.125)$$
$$\beta_{c1} \sim \mathcal{N}(0, 0.125)$$
$$\beta_{c2} \sim \mathcal{N}(0, 0.125)$$
$$\beta_{c3} \sim \mathcal{N}(0, 0.004)$$
$$\beta_{c4} \sim \mathcal{N}(0, 0.015)$$
$$\mu_{0c} \sim \exp(50)$$

$$\tau_0 \sim \mathcal{N}(-3, 1.5)$$
$$\tau_1 \sim \mathcal{N}(0, 0.5)$$
$$\tau_2 \sim \mathcal{N}(1, 1.5)$$

Below, we test the suitability of our priors using a prior predictive check. In the event that priors are suitable, we continue with executing our full path analysis and: (1) assessing chain convergence (here, by calculating and summarising Gelman-Rubin statistics), (2) checking for evidence of autocorrelation in chains, and (3) summarising the general fit of each model within our path analysis.

```
# Running prior predictive check

functionModel3WeeksHotPPCheck <-
  brm(
    data = slopeDataHot3 %>%
      mutate(
        mass = mass - mean(mass, na.rm = T),
        tarsus = tarsusLengthMean -
          mean(tarsusLengthMean, na.rm = T),
        pretreatment = ifelse(pretreatment == "cold", "A",
          ifelse(pretreatment == "neutral", "B", "C")
        ),
        slope = slope -
          mean(slope, na.rm = T)
      ) %>%
      mutate(pretreatment = factor(pretreatment,
        levels = c("B", "A", "C")
      )) %>%
      drop_na() %>%
      merge(., data %>%
        select(ring, "batch" = exp) %>%
        distinct(),
      by = "ring", all.x = TRUE
      ),
    family = "gaussian",
    bf(mass ~ pretreatment + (1 | batch)) +
      bf(tarsus ~ mass + pretreatment + (1 | batch)) +
      bf(slope ~ tarsus + mass + pretreatment + (1 | batch),
         sigma ~ batch) +
      set_rescor(FALSE),
    prior = c(
      set_prior("normal(0, 5)",
        class = "Intercept",
        resp = "mass"
      ),
      set_prior("normal(0, 15)",
        class = "b",
        coef = "pretreatmentA", resp = "mass"
      ),
      set_prior("normal(0, 15)",
        class = "b",
        coef = "pretreatmentC", resp = "mass"
      ),
      set_prior("exponential(2.5)",
        class = "sd",
        group = "batch", resp = "mass"
      ),
      set_prior("exponential(0.15)",
        class = "sigma",
        resp = "mass"
      ),
      set_prior("normal(0, 2.5)",
        class = "Intercept",
        resp = "tarsus"
      ),
      set_prior("normal(0, 2.5)",
        class = "b",
        coef = "pretreatmentA",
        resp = "tarsus"
      ),
      set_prior("normal(0, 2.5)",
        class = "b",
        coef = "pretreatmentC",
        resp = "tarsus"
      ),
      set_prior("skew_normal(0, 0.25, 5)",
        class = "b",
```

```
        coef = "mass", resp = "tarsus"
      ),
      set_prior("exponential(2)",
        class = "sd",
        group = "batch", resp = "tarsus"
      ),
      set_prior("exponential(1)",
        class = "sigma",
        resp = "tarsus"
      ),
      set_prior("normal(0, 0.125)",
        class = "Intercept",
        resp = "slope"
      ),
      set_prior("normal(0, 0.125)",
        class = "b",
        coef = "pretreatmentA",
        resp = "slope"
      ),
      set_prior("normal(0, 0.125)",
        class = "b",
        coef = "pretreatmentC",
        resp = "slope"
      ),
      set_prior("normal(0, 0.004)",
        class = "b",
        coef = "mass", resp = "slope"
      ),
      set_prior("normal(0, 0.015)",
        class = "b",
        coef = "tarsus", resp = "slope"
      ),
      set_prior("exponential(50)",
        class = "sd",
        group = "batch", resp = "slope"
      ),
      set_prior("normal(-3, 1.5)",
        dpar = "sigma",
        class = "Intercept",
        resp = "slope"
      ),
      set_prior("normal(0, 0.5)",
        dpar = "sigma",
        class = "b",
        coef = "batchB",
        resp = "slope"
      ),
      set_prior("normal(1, 1.5)",
        dpar = "sigma",
        class = "b",
        coef = "batchC",
        resp = "slope"
      )
    ),
    iter = 50000, warmup = 10000, cores = 4, chains = 4, thin = 20,
    control = list(adapt_delta = .97, max_treedepth = 14),
    sample_prior = "only",
    silent = TRUE, refresh = 0,
    file = "./models/threeWeekFunctionModelHotPPCheck.Rds"
  )

pp_check2(functionModel3WeeksHotPPCheck, resp = "slope",
   xlab = "Metabolic Slope\n(Fold Metabolism at Thermoneutrality/°C)"
   ) + xlim(c(-5, 5))
```

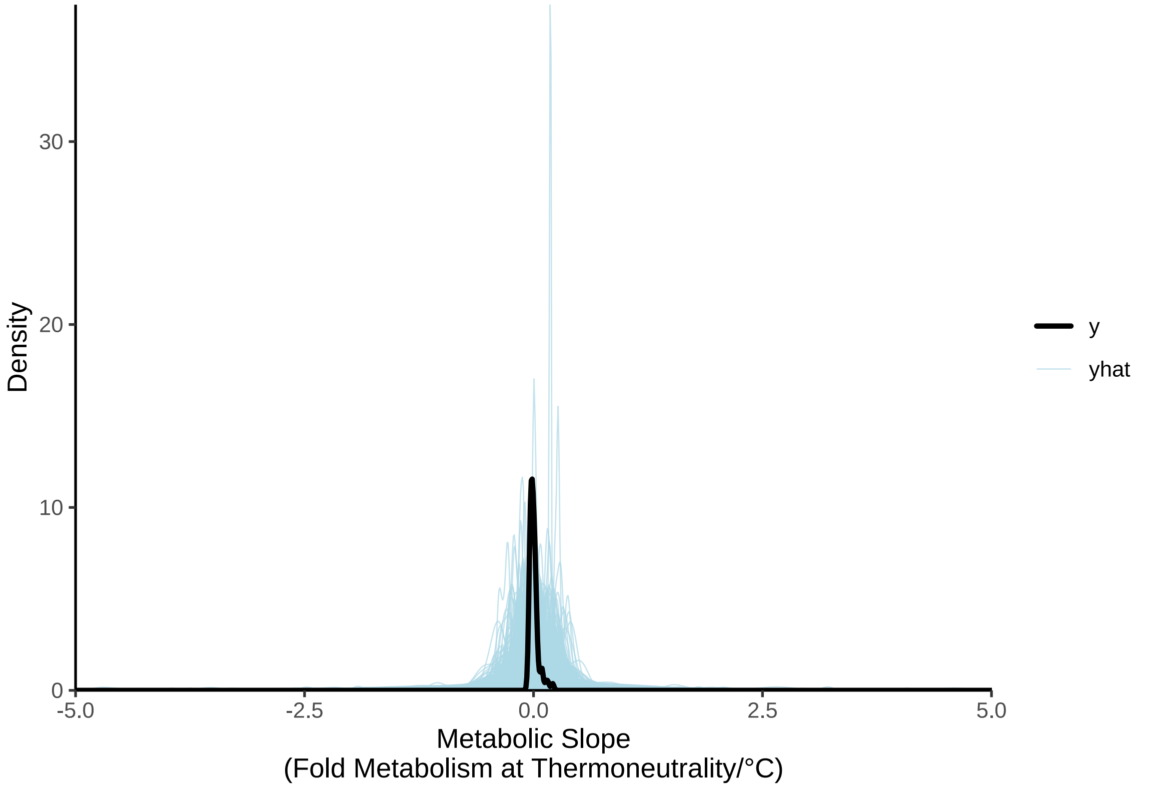


***Figure 54:*** *Prior predictive check for a Bayesian path analysis predicting physiological responses to heat in three week old Japanese quail. Black lines represent true densities of metabolic slopes (fold metabolism at thermoneutrality/°C) while blue lines represent densities estimated from model priors alone.*

```
# Broad, but acceptable. Running full model.

functionModel3WeeksHot <-
  brm(
    data = slopeDataHot3 %>%
      mutate(
        mass = mass - mean(mass, na.rm = T),
        tarsus = tarsusLengthMean -
          mean(tarsusLengthMean, na.rm = T),
        pretreatment = ifelse(pretreatment == "cold", "A",
          ifelse(pretreatment == "neutral", "B", "C")
        ),
        slope = slope -
          mean(slope, na.rm = T)
      ) %>%
      mutate(pretreatment = factor(pretreatment,
        levels = c("B", "A", "C")
      )) %>%
      drop_na() %>%
      merge(., data %>%
        select(ring, "batch" = exp) %>%
        distinct(),
      by = "ring", all.x = TRUE
      ),
    family = "gaussian",
    bf(mass ~ pretreatment + (1 | batch)) +
      bf(tarsus ~ mass + pretreatment + (1 | batch)) +
      bf(
        slope ~ tarsus + mass + pretreatment + (1 | batch),
        sigma ~ batch
      ) +
```

```
      set_rescor(FALSE),
    prior = c(
      set_prior("normal(0, 5)",
        class = "Intercept",
        resp = "mass"
      ),
      set_prior("normal(0, 15)",
        class = "b",
        coef = "pretreatmentA", resp = "mass"
      ),
      set_prior("normal(0, 15)",
        class = "b",
        coef = "pretreatmentC", resp = "mass"
      ),
      set_prior("exponential(2.5)",
        class = "sd",
        group = "batch", resp = "mass"
      ),
      set_prior("exponential(0.15)",
        class = "sigma",
        resp = "mass"
      ),
      set_prior("normal(0, 2.5)",
        class = "Intercept",
        resp = "tarsus"
      ),
      set_prior("normal(0, 2.5)",
        class = "b",
        coef = "pretreatmentA",
        resp = "tarsus"
      ),
      set_prior("normal(0, 2.5)",
        class = "b",
        coef = "pretreatmentC",
        resp = "tarsus"
      ),
      set_prior("skew_normal(0, 0.25, 5)",
        class = "b",
        coef = "mass", resp = "tarsus"
      ),
      set_prior("exponential(2)",
        class = "sd",
        group = "batch", resp = "tarsus"
      ),
      set_prior("exponential(1)",
        class = "sigma",
        resp = "tarsus"
      ),
      set_prior("normal(0, 0.125)",
        class = "Intercept",
        resp = "slope"
      ),
      set_prior("normal(0, 0.125)",
        class = "b",
        coef = "pretreatmentA",
        resp = "slope"
      ),
      set_prior("normal(0, 0.125)",
        class = "b",
        coef = "pretreatmentC",
        resp = "slope"
      ),
      set_prior("normal(0, 0.004)",
        class = "b",
        coef = "mass", resp = "slope"
      ),
      set_prior("normal(0, 0.015)",
```

```
        class = "b",
        coef = "tarsus", resp = "slope"
      ),
      set_prior("exponential(50)",
        class = "sd",
        group = "batch", resp = "slope"
      ),
      set_prior("normal(-3, 1.5)",
        dpar = "sigma",
        class = "Intercept",
        resp = "slope"
      ),
      set_prior("normal(0, 0.5)",
        dpar = "sigma",
        class = "b",
        coef = "batchB",
        resp = "slope"
      ),
      set_prior("normal(1, 1.5)",
        dpar = "sigma",
        class = "b",
        coef = "batchC",
        resp = "slope"
      )
    ),
    iter = 50000, warmup = 10000, cores = 4, chains = 4, thin = 20,
    control = list(adapt_delta = .97, max_treedepth = 14),
    silent = TRUE, refresh = 0,
    file = "./models/threeWeekFunctionModelHot.Rds"
  )

chainCheck(functionModel3WeeksHot)
```

```
## Rhat range: 1 - 1.001
## Neff/N range: 0.902 - 1.007
```

```
# Good. Checking r2 and posterior density/fit

caption <- paste0("R\\textsuperscript{2} for a Bayesian ",
                  "path analysis predicting metabolic slopes ",
                  "(fold metabolism at thermoneutrality; 30°C) ",
                  "of three week old Japanese quail as a ",
                  "function of body mass (g), tarsus length (mm) and rearing ",
                  "temperature. Metabolic slopes are measured above ",
                  "thermoneutrality (30°C - 40°C)."
)

brms::bayes_R2(functionModel3WeeksHot,
               ndraws = 1000,
               robust = TRUE) %>%
  as.data.frame() %>%
  rownames_to_column("var") %>%
  merge(., tribble(
    ~var, ~Var,
    "R2mass", "Body Mass (g)",
    "R2tarsus", "Tarsus Length (mm)",
    "R2slope", "Metabolic Slope"
  ), by = c("var")) %>%
  mutate(Estimate = round(Estimate, digits = 4),
         Est.Error = round(Est.Error, digits = 4),
         `95\\% CI` = paste0("[", round(Q2.5, digits = 4),
                             ", ",
                             round(Q97.5, digits = 4),
                             "]"
         )
         ) %>%
  select(
    "Response" = Var, "R\\textsuperscript{2}" = Estimate,
```

```
    "Standard Error" = Est.Error, `95\\% CI`
  ) %>%
  kbl(.,
    longtable = T, booktabs = T, format = "latex",
    caption = caption, escape = FALSE
  ) %>%
  column_spec(column = c(1:10), width = "2.5cm")  %>%
  kable_styling(latex_options = "striped")
```

***Table 25:*** *$R^2$ for a Bayesian path analysis predicting metabolic slopes (fold metabolism at thermoneutrality; 30°C) of three week old Japanese quail as a function of body mass (g), tarsus length (mm) and rearing temperature. Metabolic slopes are measured above thermoneutrality (30°C - 40°C).*

| Response | $R^2$ | Standard Error | 95% CI |
|---|---|---|---|
| Body Mass (g) | 0.1011 | 0.0571 | [0.0111, 0.232] |
| Metabolic Slope | 0.0604 | 0.0352 | [0.0146, 0.1953] |
| Tarsus Length (mm) | 0.3641 | 0.0754 | [0.1978, 0.4888] |

The fit of our path analysis is next broadly assessed.

```
ggarrange(
  pp_check2(functionModel3WeeksHot,
            resp = "slope",
            xlab = paste0("Metabolic Slope\n(Fold Metabolism ",
                          "at\nThermoneutrality/°C)")
            ),

  functionModel3WeeksHot$data %>%
    mutate("Fit" = fitted(functionModel3WeeksHot,
                          resp = "slope",
                          robust = TRUE)[, "Estimate"]) %>%
    ggplot(aes(x = Fit, y = slope)) +
    geom_point() +
    geom_line(data = data.frame("Fit" = c(-0.05, 0.05),
                                "slope" = c(-0.05, 0.05))) +
    xlab(
      "Fitted Metabolic Slope\n(Fold Metabolism at\nThermoneutrality/°C)"
    ) +
    ylab(
      "True Metabolic Slope\n(Fold Metabolism at\nThermoneutrality/°C)"
    ) +
    theme_classic(),
  labels = c("A", "B")
)
```

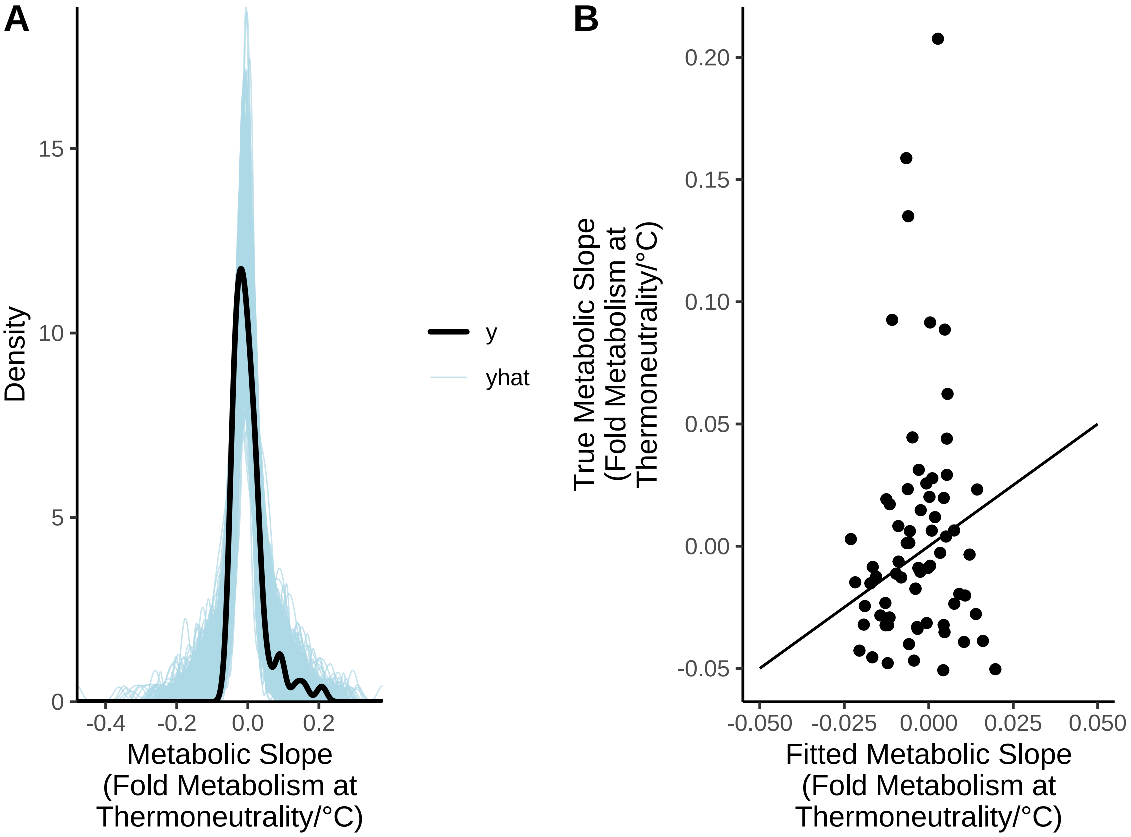


***Figure 55:*** *Characteristics of Bayesian path analysis predicting physiological responses to heat in three week old Japanese quail. Panels A displays a posterior predictive check from path analysis with the black line indicating the true density of metabolic slopes and blue lines indicating densities estimated from posterior draws. Panel B displays model fitted values by true values with dots indicating individual data points and the black line indicating a 1:1 relationship.*

Residuals of this path analysis are diagnosed for pathologies as above.

```
p1 <- functionModel3WeeksHot$data %>%
  mutate("Residuals" = residuals(functionModel3WeeksHot,
                                  resp = "slope",
                                  robust = TRUE)[, "Estimate"]) %>%
  ggplot(aes(x = Residuals)) +
  geom_density(colour = "black", fill = "grey50", alpha = 0.5) +
  xlab("Metabolic Slope\nResiduals") +
  ylab("Density") +
  theme_classic()

p2 <- functionModel3WeeksHot$data %>%
  mutate("Residuals" = residuals(functionModel3WeeksHot,
                                  resp = "slope",
                                  robust = TRUE)[, "Estimate"]) %>%
  ggplot(aes(sample = Residuals)) +
  stat_qq(colour = "grey50") +
  stat_qq_line() +
  xlab("Theoretical Quantiles") +
  ylab(TeX(
  '$\\overset{Sample~Quantiles}{(Slope)}$'
    )) +
  theme_classic()
```

```
p3 <- functionModel3WeeksHot$data %>%
  mutate("Residuals" = residuals(functionModel3WeeksHot,
                                 resp = "slope",
                                 robust = TRUE)[, "Estimate"]) %>%
  ggplot(aes(x = mass, y = Residuals)) +
  geom_point(pch = 21, colour = "black", size = 2,
             fill = "lightblue", alpha = 0.5) +
  xlab("Body Mass (g; Mean-Centred)") +
  ylab("Metabolic Slope\nResiduals") +
  theme_classic()

p4 <- functionModel3WeeksHot$data %>%
  mutate("Residuals" = residuals(functionModel3WeeksHot,
                                 resp = "slope",
                                 robust = TRUE)[, "Estimate"]) %>%
  ggplot(aes(x = tarsus, y = Residuals)) +
  geom_point(pch = 21, colour = "black", size = 2,
             fill = "lightblue", alpha = 0.5) +
  xlab("Tarsus Length (mm; Mean-Centred)") +
  ylab("Metabolic Slope\nResiduals") +
  theme_classic()

p5 <- functionModel3WeeksHot$data %>%
  mutate("Residuals" =
           residuals(functionModel3WeeksHot,
                                 resp = "slope",
                                 robust = TRUE)[, "Estimate"]) %>%
  mutate("pretreatment" = ifelse(pretreatment == "A", "Cold (10°C)",
    ifelse(pretreatment == "B", "Mild (20°C)",
      "Warm (30°C)"
    )
  )) %>%
  ggplot(aes(x = pretreatment, y = Residuals)) +
  geom_point(pch = 21, colour = "black", size = 2,
             fill = "lightblue", alpha = 0.5) +
  stat_summary(geom = "errorbar", fun.data = "mean_se",
               colour = "black", width = 0.25) +
  stat_summary(geom = "point", fun = "mean", size = 4,
               pch = 21, colour = "black", fill = "lightblue") +
  xlab("Rearing Conditions") +
  ylab("Metabolic Slope\nResiduals") +
  theme_classic()

p6 <- functionModel3WeeksHot$data %>%
  mutate(
    "Residuals" = residuals(functionModel3WeeksHot,
                            resp = "slope",
                                 robust = TRUE)[, "Estimate"],
    "Fit" = fitted(functionModel3WeeksHot,
                   resp = "slope",
                                 robust = TRUE)[, "Estimate"]
  ) %>%
  ggplot(aes(x = Fit, y = Residuals)) +
  geom_point(pch = 21, colour = "black", size = 2,
             fill = "lightblue", alpha = 0.5) +
  xlab("Fitted Metabolic Slope\n(Fold Metabolism at Thermoneutrality/°C)") +
  ylab("Metabolic Slope\nResiduals") +
  theme_classic()

(p1 + p2) / (p3 + p4) / (p5 + p6) +
  plot_annotation(tag_levels = "A")
```

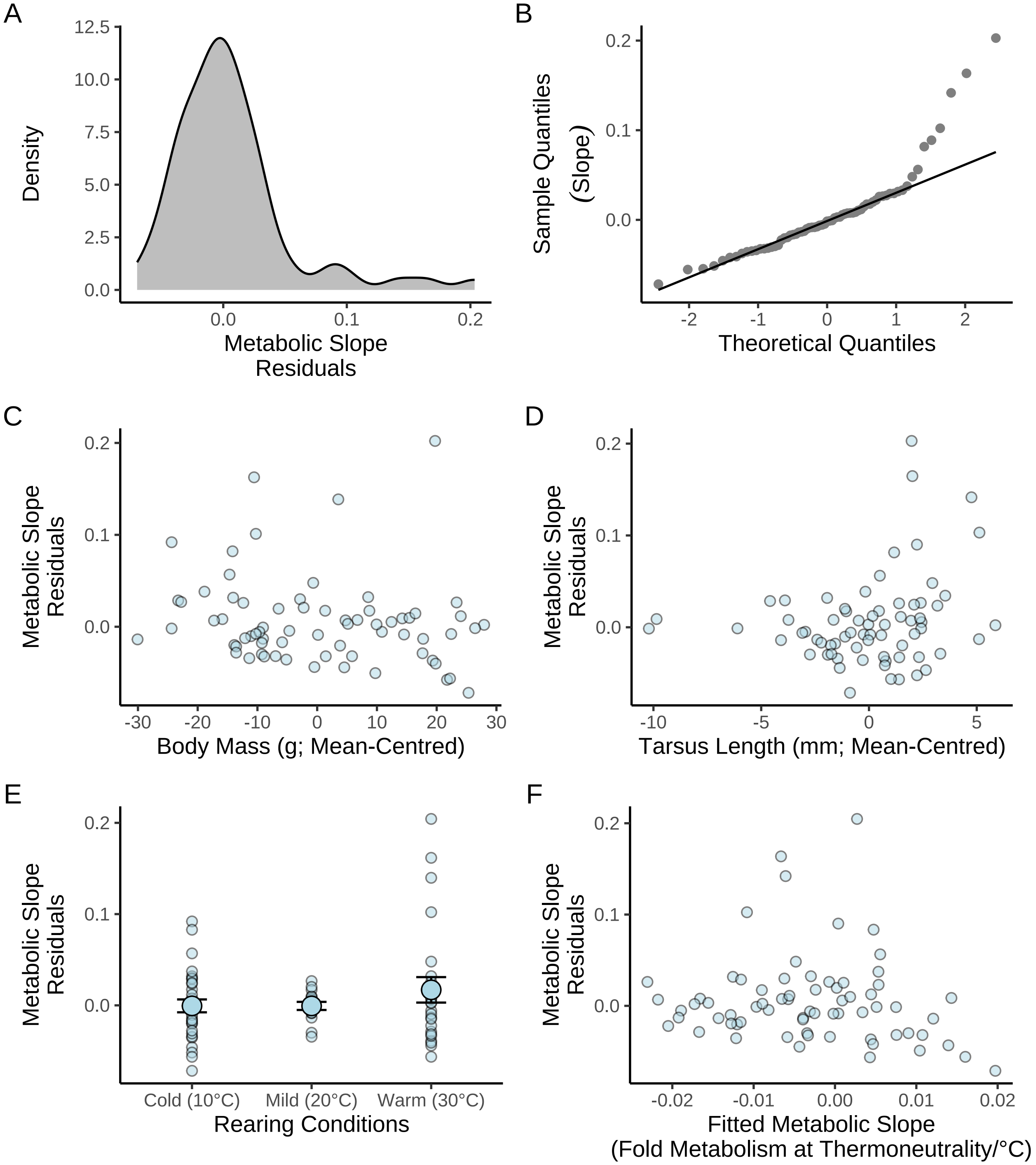


***Figure 56:*** *Spread of ordinary residuals from Bayesian path analysis predicting physiological responses to heat in three week old Japanese quail. Quail were reared in one of three thermal conditions: (1) cold (10°C), (2) mild (20°C), or warm (30°C). Panels A and B display distributions of ordinary residuals of metabolic slopes (fold metabolism at thermoneutrality/°C). Panels C to F display ordinary residuals against predictors and fitted values; dots in these panels represent values per individual.*

Evidently, there is some right skewing in our median residuals. This is arguably expected given that our slope values are manually calculated and therefore not balanced by a population-averaged response. Below, we check whether these values have undue influence on our model by first using leave-one-out ("LOO") cross-validation via Pareto-smoothed importance sampling, then evaluating importance weights (here, Pareto K) per data point (Vehtari et al, 2017).

```
loo(functionModel3WeeksHot)$diagnostics$pareto_k %>%
  as_tibble() %>%
  ggplot(aes(x = 1:nrow(.), y = value)) +
  geom_point(pch = 21, colour = "black", fill = "grey50", size = 2) +
  geom_hline(yintercept = 0.5, colour = "black", linetype = "dashed") +
  geom_hline(yintercept = 0.7, colour = "red4", linetype = "dashed") +
  xlab("Sample") +
  ylab("Pareto K") +
  theme_classic()
```

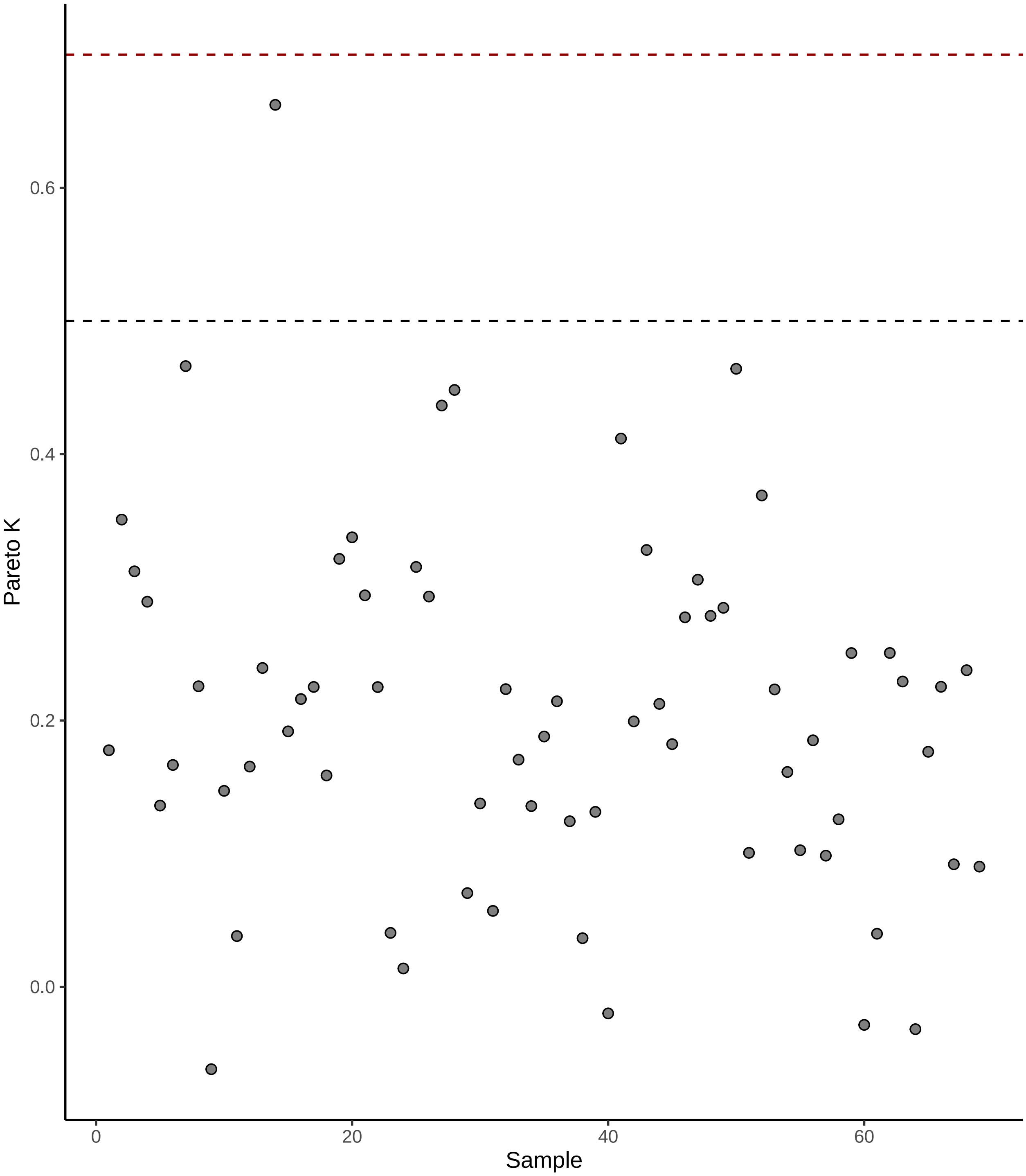


***Figure 57:*** *Relative importance of individual metabolic slope values (dots) to outcomes of our Bayesian path analysis, as estimated using leave-one-out cross validations. Importance is measured here as Pareto K (Vehtari et al, 2017). The black, dashed horizontal line indicated a Pareto K of 0.5 and the red horizontal dashed line indicates as Pareto K of 0.7, above which, samples hold significant importance on analysis outcomes.*

A few datum points are identified as potentially problematic but not severely so (see Vehtari et al 2020). We therefore proceed with our model as currently structured.

Coefficient estimates are next drawn from posteriors and visualised for normality. After-hand, these coefficients are summarised then used to plot conditional effects.

```
as.data.frame(functionModel3WeeksHot) %>%
    select(
      "Intercept" = b_slope_Intercept,
      "Tarsus Length\n(mm; Mean-Centred)" = b_slope_tarsus,
      "Body Mass\n(g; Mean-Centred)" = b_slope_mass,
      "Cold Rearing\n(10°C)" = b_slope_pretreatmentA,
      "Warm Rearing\n(30°C)" = b_slope_pretreatmentC,
      #"Sigma" = sigma_slope
    ) %>%
    pivot_longer(everything(), names_to = "Par",
                 values_to = "Coefs") %>%
    ggplot(aes(x = Coefs)) +
    facet_wrap(~Par, scales = "free", ncol = 2) +
    geom_density(colour = "black",
                 alpha = 0.5, fill = "grey70") +
    geom_vline(xintercept = 0, linetype = "dashed",
               colour = "black") +
    ylab("Density") +
    scale_x_continuous(n.breaks = 4) +
    theme_classic() +
    theme(axis.title.x = element_blank()) +
    ggtitle(paste0("Metabolic Slope\n(Fold Metabolism ",
                   "at Thermoneutrality/°C)")
      )
```

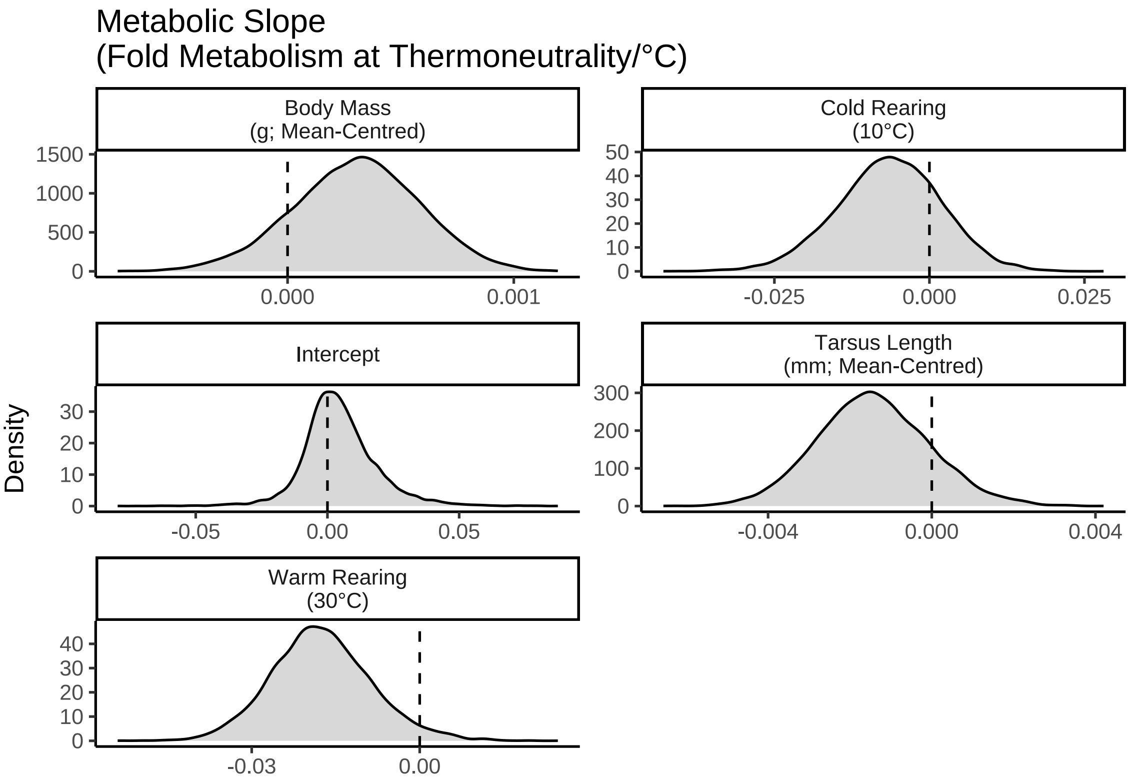


***Figure 58:*** *Posterior densities of coefficients for a Bayesian path analysis predicting physiological responses to heat in three week old Japanese quail. Quail were reared in one of three thermal conditions: (1) cold (10°C), (2) mild (20°C), or warm (30°C).*

```
caption <- paste0("Results from a Bayesian path analysis ",
                  "predicting metabolic responses to heat (metabolic slope ",
                  "[fold metabolism at thermoneutrality/°C]) as ",
                  "a function of morphometry in three week old Japanese quail. ",
                  "Physiological measurements are made at ambient temperatures ",
                  "between 30°C and 40°C. Cold rearing indicates ",
                  "post-hatch rearing at 10°C, relative to 20°C (intercept), ",
                  "or 30°C ('warm rearing'). CIs indicates quantile intervals ",
                  "and BF indicates Bayes Factors."
)

week3ResultsHeat <-
  as.data.frame(functionModel3WeeksHot) %>%
  summarise_all(., .funs = median) %>%
  pivot_longer(everything(),
    names_to = "Parameter",
    values_to = "Estimate"
  ) %>%
  merge(., quantileCIs(functionModel3WeeksHot, cis = c(50, 95)),
    by = "Parameter", all.x = TRUE
  ) %>%
  filter(grepl("b_|sd_", Parameter)) %>%
  rowwise() %>%
  mutate("BF" = ifelse(Estimate < 0,
    (2 * mean(as.data.frame(
      functionModel3WeeksHot
    )[, Parameter] <= 0)) /
      (2 * mean(as.data.frame(
        functionModel3WeeksHot
      )[, Parameter] >= 0)),
    (2 * mean(as.data.frame(
      functionModel3WeeksHot
    )[, Parameter] >= 0)) /
      (2 * mean(as.data.frame(
        functionModel3WeeksHot
      )[, Parameter] <= 0))
  )) %>%
  ungroup() %>%
  mutate(
    "Estimate" = round(Estimate, digits = 4),
    "BF" = round(BF, digits = 4),
    "N" = nrow(functionModel3WeeksHot$data)
  ) %>%
  mutate("Parameter" = ifelse(grepl("b_", Parameter),
    gsub("b_", "", Parameter),
    gsub(
      "Intercept", "batch",
      gsub(".*__", "", Parameter)
    )
  )) %>%
  mutate(
    "Response" = gsub("_.*", "", Parameter),
    "Parameter" = gsub(".*_", "", Parameter)
  ) %>%
  merge(., tribble(
    ~Response, ~response, ~level,
    "mass", "Body Mass (g)", "A",
    "slope",
    "Metabolic Slope", "D",
    "tarsus", "Tarsus Length (mm)", "B"
  ),
  by = "Response"
  ) %>%
  merge(., tribble(
    ~Parameter, ~parameter, ~number,
    "Intercept", "Intercept", "1",
    "mass", "Body Mass (g)", "4",
```

```
    "tarsus", "Tarsus Length (mm)", "5",
    "pretreatmentA", "Cold Rearing", "2",
    "pretreatmentC", "Warm Rearing", "3",
    "batch", "Egg Batch [mu]", "6"
  ),
  by = "Parameter"
  ) %>%
  mutate(
    `50\\% HDI` = paste0("(", paste(
      round(Low_CI_50, digits = 4),
      round(High_CI_50, digits = 4),
      sep = ", "
    ), ")"),
    `95\\% HDI` = paste0("(", paste(
      round(Low_CI_95, digits = 4),
      round(High_CI_95, digits = 4),
      sep = ", "
    ), ")")
  ) %>%
  select(-c(Low_CI_50, High_CI_50, Low_CI_95, High_CI_95)) %>%
  select(
    "Response" = "response", "Parameter" = "parameter", N,
    Estimate, `50\\% HDI`, `95\\% HDI`, BF, level, number
  ) %>%
  arrange(level, number) %>%
  select(-c(level, number)) %>%
  kbl(.,
    longtable = T, booktabs = T, format = "latex", escape = FALSE,
    caption = caption
  ) %>%
  column_spec(column = c(1:2), width = "2.2cm") %>%
  column_spec(column = c(3:10), width = "1.9cm") %>%
  kable_styling(latex_options = "striped")

week3ResultsHeat
```

***Table 26:*** *Results from a Bayesian path analysis predicting metabolic responses to heat (metabolic slope [fold metabolism at thermoneutrality/°C]) as a function of morphometry in three week old Japanese quail. Physiological measurements are made at ambient temperatures between 30°C and 40°C. Cold rearing indicates post-hatch rearing at 10°C, relative to 20°C (intercept), or 30°C ('warm rearing'). CIs indicates quantile intervals and BF indicates Bayes Factors.*

| Response | Parameter | N | Estimate | 50% HDI | 95% HDI | BF |
|---|---|---|---|---|---|---|
| Body Mass (g) | Intercept | 69 | 3.7324 | (1.4811, 6.0599) | (-3.0409, 10.5202) | 6.4766 |
| Body Mass (g) | Cold Rearing | 69 | -8.6607 | (-11.4814, -5.809) | (-17.0343, -0.0632) | 41.7807 |
| Body Mass (g) | Warm Rearing | 69 | 0.5418 | (-2.4201, 3.4241) | (-8.3102, 9.2061) | 1.1996 |
| Body Mass (g) | Egg Batch [mu] | 69 | 0.2665 | (0.1107, 0.5192) | (0.0106, 1.3774) | Inf |
| Tarsus Length (mm) | Intercept | 69 | -0.7154 | (-1.2593, -0.1298) | (-2.3545, 1.1171) | 3.8251 |
| Tarsus Length (mm) | Cold Rearing | 69 | 0.4519 | (-0.0802, 0.9886) | (-1.1125, 1.9555) | 2.5180 |
| Tarsus Length (mm) | Warm Rearing | 69 | 1.4727 | (0.8819, 2.071) | (-0.2022, 3.1667) | 21.2841 |
| Tarsus Length (mm) | Body Mass (g) | 69 | 0.0655 | (0.052, 0.0793) | (0.0275, 0.1055) | 1599.0000 |
| Tarsus Length (mm) | Egg Batch [mu] | 69 | 0.9446 | (0.6617, 1.2749) | (0.1661, 2.176) | Inf |
| Metabolic Slope | Intercept | 69 | 0.0033 | (-0.0036, 0.0117) | (-0.0209, 0.0363) | 1.6290 |
| Metabolic Slope | Cold Rearing | 69 | -0.0063 | (-0.012, -8e-04) | (-0.0235, 0.0095) | 3.5532 |

| | | | | | | |
|---|---|---|---|---|---|---|
| Metabolic Slope | Warm Rearing | 69 | -0.0179 | (-0.0235, -0.0119) | (-0.0344, 0.0015) | 29.3030 |
| Metabolic Slope | Body Mass (g) | 69 | 0.0003 | (1e-04, 5e-04) | (-3e-04, 8e-04) | 6.0547 |
| Metabolic Slope | Tarsus Length (mm) | 69 | -0.0015 | (-0.0023, -5e-04) | (-0.004, 0.0014) | 5.7682 |
| Metabolic Slope | Egg Batch [mu] | 69 | 0.0145 | (0.0079, 0.0233) | (9e-04, 0.0516) | Inf |

```
#save_kable(week3ResultsHeat,
# "../tables/pathAnalysisInHeat3Weeks.html")

functionModel3WeeksHotMassR2 <-
  brm(
    data = slopeDataHot3 %>%
      mutate(
        mass = mass - mean(mass, na.rm = T),
        tarsus = tarsusLengthMean -
          mean(tarsusLengthMean, na.rm = T),
        pretreatment = ifelse(pretreatment == "cold", "A",
          ifelse(pretreatment == "neutral", "B", "C")
        ),
        slope = slope -
          mean(slope, na.rm = T)
      ) %>%
      mutate(pretreatment = factor(pretreatment,
        levels = c("B", "A", "C")
      )) %>%
      drop_na() %>%
      merge(., data %>%
        select(ring, "batch" = exp) %>%
        distinct(),
      by = "ring", all.x = TRUE
      ),
    family = "gaussian",
    bf(mass ~ pretreatment + (1 | batch)) +
      bf(tarsus ~ mass + pretreatment + (1 | batch)) +
      bf(slope ~ tarsus + pretreatment + (1 | batch),
         sigma ~ batch) +
      set_rescor(FALSE),
    prior = c(
      set_prior("normal(0, 5)",
        class = "Intercept",
        resp = "mass"
      ),
      set_prior("normal(0, 15)",
        class = "b",
        coef = "pretreatmentA", resp = "mass"
      ),
      set_prior("normal(0, 15)",
        class = "b",
        coef = "pretreatmentC", resp = "mass"
      ),
      set_prior("exponential(2.5)",
        class = "sd",
        group = "batch", resp = "mass"
      ),
      set_prior("exponential(0.15)",
        class = "sigma",
        resp = "mass"
      ),
      set_prior("normal(0, 2.5)",
        class = "Intercept",
        resp = "tarsus"
      ),
      set_prior("normal(0, 2.5)",
        class = "b",
        coef = "pretreatmentA",
```

```
        resp = "tarsus"
      ),
      set_prior("normal(0, 2.5)",
        class = "b",
        coef = "pretreatmentC",
        resp = "tarsus"
      ),
      set_prior("skew_normal(0, 0.25, 5)",
        class = "b",
        coef = "mass", resp = "tarsus"
      ),
      set_prior("exponential(2)",
        class = "sd",
        group = "batch", resp = "tarsus"
      ),
      set_prior("exponential(1)",
        class = "sigma",
        resp = "tarsus"
      ),
      set_prior("normal(0, 0.125)",
        class = "Intercept",
        resp = "slope"
      ),
      set_prior("normal(0, 0.125)",
        class = "b",
        coef = "pretreatmentA",
        resp = "slope"
      ),
      set_prior("normal(0, 0.125)",
        class = "b",
        coef = "pretreatmentC",
        resp = "slope"
      ),
      set_prior("normal(0, 0.015)",
        class = "b",
        coef = "tarsus", resp = "slope"
      ),
      set_prior("exponential(50)",
        class = "sd",
        group = "batch", resp = "slope"
      ),
      set_prior("normal(-3, 1.5)",
        dpar = "sigma",
        class = "Intercept",
        resp = "slope"
      ),
      set_prior("normal(0, 0.5)",
        dpar = "sigma",
        class = "b",
        coef = "batchB",
        resp = "slope"
      ),
      set_prior("normal(1, 1.5)",
        dpar = "sigma",
        class = "b",
        coef = "batchC",
        resp = "slope"
      )
    ),
    iter = 50000, warmup = 10000, cores = 4, chains = 4, thin = 20,
    control = list(adapt_delta = .97, max_treedepth = 14),
    silent = TRUE, refresh = 0,
    file = "./models/threeWeekFunctionModelHotMassR2.Rds"
  )

functionModel3WeeksHotTarsusR2 <-
  brm(
```

```
  data = slopeDataHot3 %>%
    mutate(
      mass = mass - mean(mass, na.rm = T),
      tarsus = tarsusLengthMean -
        mean(tarsusLengthMean, na.rm = T),
      pretreatment = ifelse(pretreatment == "cold", "A",
        ifelse(pretreatment == "neutral", "B", "C")
      ),
      slope = slope -
        mean(slope, na.rm = T)
    ) %>%
    mutate(pretreatment = factor(pretreatment,
      levels = c("B", "A", "C")
    )) %>%
    drop_na() %>%
    merge(., data %>%
      select(ring, "batch" = exp) %>%
      distinct(),
    by = "ring", all.x = TRUE
    ),
  family = "gaussian",
  bf(mass ~ pretreatment + (1 | batch)) +
    bf(tarsus ~ mass + pretreatment + (1 | batch)) +
    bf(slope ~ mass + pretreatment + (1 | batch),
       sigma ~ batch) +
    set_rescor(FALSE),
  prior = c(
    set_prior("normal(0, 5)",
      class = "Intercept",
      resp = "mass"
    ),
    set_prior("normal(0, 15)",
      class = "b",
      coef = "pretreatmentA", resp = "mass"
    ),
    set_prior("normal(0, 15)",
      class = "b",
      coef = "pretreatmentC", resp = "mass"
    ),
    set_prior("exponential(2.5)",
      class = "sd",
      group = "batch", resp = "mass"
    ),
    set_prior("exponential(0.15)",
      class = "sigma",
      resp = "mass"
    ),
    set_prior("normal(0, 2.5)",
      class = "Intercept",
      resp = "tarsus"
    ),
    set_prior("normal(0, 2.5)",
      class = "b",
      coef = "pretreatmentA",
      resp = "tarsus"
    ),
    set_prior("normal(0, 2.5)",
      class = "b",
      coef = "pretreatmentC",
      resp = "tarsus"
    ),
    set_prior("skew_normal(0, 0.25, 5)",
      class = "b",
      coef = "mass", resp = "tarsus"
    ),
    set_prior("exponential(2)",
      class = "sd",
```

```
        group = "batch", resp = "tarsus"
      ),
      set_prior("exponential(1)",
        class = "sigma",
        resp = "tarsus"
      ),
      set_prior("normal(0, 0.125)",
        class = "Intercept",
        resp = "slope"
      ),
      set_prior("normal(0, 0.125)",
        class = "b",
        coef = "pretreatmentA",
        resp = "slope"
      ),
      set_prior("normal(0, 0.125)",
        class = "b",
        coef = "pretreatmentC",
        resp = "slope"
      ),
      set_prior("normal(0, 0.004)",
        class = "b",
        coef = "mass", resp = "slope"
      ),
      set_prior("exponential(50)",
        class = "sd",
        group = "batch", resp = "slope"
      ),
      set_prior("normal(-3, 1.5)",
        dpar = "sigma",
        class = "Intercept",
        resp = "slope"
      ),
      set_prior("normal(0, 0.5)",
        dpar = "sigma",
        class = "b",
        coef = "batchB",
        resp = "slope"
      ),
      set_prior("normal(1, 1.5)",
        dpar = "sigma",
        class = "b",
        coef = "batchC",
        resp = "slope"
      )
    ),
    iter = 50000, warmup = 10000, cores = 4, chains = 4, thin = 20,
    control = list(adapt_delta = .97, max_treedepth = 14),
    silent = TRUE, refresh = 0,
    file = "./models/threeWeekFunctionModelHotTarsusR2.Rds"
  )

massPR2 <- brms::bayes_R2(functionModel3WeeksHot,
  ndraws = 1000,
  robust = TRUE, resp = "slope",
  summary = FALSE
) %>%
  as.data.frame() %>%
  select("fullR2" = R2slope) %>%
  cbind(
    .,
    brms::bayes_R2(functionModel3WeeksHotMassR2,
      ndraws = 1000,
      robust = TRUE, resp = "slope",
      summary = FALSE
    )
  ) %>%
```

```
  mutate("pR2" = round(fullR2 - R2slope, digits = 3)) %>%
  pull(pR2)

tarsusPR2 <- brms::bayes_R2(functionModel3WeeksHot,
  ndraws = 1000,
  robust = TRUE, resp = "slope",
  summary = FALSE
) %>%
  as.data.frame() %>%
  select("fullR2" = R2slope) %>%
  cbind(
    .,
    brms::bayes_R2(functionModel3WeeksHotTarsusR2,
      ndraws = 1000,
      robust = TRUE, resp = "slope",
      summary = FALSE
    )
  ) %>%
  mutate("pR2" = round(fullR2 - R2slope, digits = 3)) %>%
  pull(pR2)

caption <- paste0(
  "Variance in metabolic slope ",
  "(fold metabolism at thermoneutrality/°C) above thermnoeutrality ",
  "explained by morphometry in three week old Japanese ",
  "quail."
)

quantile(massPR2, probs = c(0.025, 0.975), type = 8) %>%
  t() %>%
  as.data.frame() %>%
  mutate(`2.5%` = ifelse(`2.5%` < 0, 0, `2.5%`)) %>%
  mutate("95\\% CI" = paste0(
    "[",
    round(`2.5%`, digits = 3),
    ",",
    round(`97.5%`, digits = 3),
    "]"
  )) %>%
  mutate(
    "massR2" =
      round(median(massPR2), digits = 3)
  ) %>%
  mutate(massR2 = ifelse(massR2 < 0, 0, massR2)) %>%
  select(
    "Mass Partial R\\textsuperscript{2}" = massR2,
    "95\\% CI"
  ) %>%
  cbind(
    .,
    quantile(tarsusPR2, probs = c(0.025, 0.975), type = 8) %>%
      t() %>%
      as.data.frame() %>%
      mutate(`2.5%` = ifelse(`2.5%` < 0, 0, `2.5%`)) %>%
      mutate("95\\% CI" = paste0(
        "[",
        round(`2.5%`, digits = 3),
        ",",
        round(`97.5%`, digits = 3),
        "]"
      )) %>%
      mutate("tarsusR2" = round(median(tarsusPR2), digits = 3)) %>%
      select(
        "Tarsus Partial R\\textsuperscript{2}" = tarsusR2,
        "95\\% CI"
      )
  ) %>%
```

```
  kbl(.,
    longtable = T, booktabs = T, format = "latex",
    caption = caption, escape = FALSE
  ) %>%
  column_spec(column = c(1:10), width = "2.5cm") %>%
  kable_styling(latex_options = "striped")
```

***Table 27:*** *Variance in metabolic slope (fold metabolism at thermoneutrality/°C) above thermnoeutrality explained by morphometry in three week old Japanese quail.*

| Mass Partial $R^2$ | 95% CI | Tarsus Partial $R^2$ | 95% CI |
|---|---|---|---|
| 0.006 | [0,0.147] | 0.007 | [0,0.146] |

Similar to analysis of metabolic responses to the cold, we estimate costs of misaligning with Bergmann's and Allen's rule in the heat. This is achieved by calculating the relative increase in fold metabolism among large- (2 standard deviations above mean body mass) and short-limbed (2 standard deviations below mean tarsus length) quail at 40°C compared with average quail.

```
data.frame(
  "Size" = c("Average", "Large (2x s.d. > mean)"),
  "pretreatment" = "B",
  "batch" = "A",
  "mass" = c(mean(functionModel3WeeksHot$data$mass, na.rm = T),
             mean(functionModel3WeeksHot$data$mass, na.rm = T) +
               2*sd(functionModel3WeeksHot$data$mass, na.rm = T)),
  "tarsus" = mean(functionModel3WeeksHot$data$tarsus, na.rm = T)
) %>%
  mutate("slope" = predict(functionModel3WeeksHot, newdata = .,
                           resp = "slope", re_form = NA,
                           robust = TRUE)[, "Estimate"]) %>%
  mutate(slope = slope +
          mean(slopeDataHot3$slope, na.rm = T)
  ) %>%
  select("Body Mass" = Size,
         "Metabolic Slope" = slope)
```

```
## # A tibble: 2 x 2
##   `Body Mass`             `Metabolic Slope`
##   <chr>                               <dbl>
## 1 Average                             0.144
## 2 Large (2x s.d. > mean)              0.153
```

```
data.frame(
  "Size" = c("Average", "Small (2x s.d. < mean)"),
  "pretreatment" = "B",
  "batch" = "A",
  "tarsus" = c(mean(functionModel3WeeksHot$data$tarsus, na.rm = T),
             mean(functionModel3WeeksHot$data$tarsus, na.rm = T) -
               2*sd(functionModel3WeeksHot$data$tarsus, na.rm = T)),
  "mass" = mean(functionModel3WeeksHot$data$mass, na.rm = T)
) %>%
  mutate("slope" = predict(functionModel3WeeksHot, newdata = .,
                           resp = "slope", re_form = NA,
                           robust = TRUE)[, "Estimate"]) %>%
  mutate(slope = slope +
          mean(slopeDataHot3$slope, na.rm = T)
  ) %>%
  select("Tarsus Length" = Size,
         "Metabolic Slope" = slope)
```

```
## # A tibble: 2 x 2
##   `Tarsus Length`         `Metabolic Slope`
##   <chr>                               <dbl>
## 1 Average                             0.144
```

```
## 2 Small (2x s.d. < mean)              0.153
```

```
# Conducting formal comparisons

mismatchDFMass <- data.frame(
  "Size" = c("Average", "Large (2x s.d. > mean)"),
  "pretreatment" = "B",
  "batch" = "A",
  "mass" = c(mean(functionModel3WeeksHot$data$mass, na.rm = T),
             mean(functionModel3WeeksHot$data$mass, na.rm = T) +
               2*sd(functionModel3WeeksHot$data$mass, na.rm = T)),
  "tarsus" = 0
) %>% predict(functionModel3WeeksHot, newdata = .,
              robust = TRUE, re_form = NA,
              resp = "slope", summary = FALSE) %>%
  as.data.frame() %>%
  rename("Average" = "V1", "Large" = "V2") %>%
  mutate_all(.funs = function(x)
    {x + mean(slopeDataHot3$slope, na.rm = T)}
    )

mismatchDFMassTest <- build_hdf(
  vars = list(
    mismatchDFMass$Average,
    mismatchDFMass$Large
    ),
  priors = list(
    rnorm(nrow(mismatchDFMass), 0, 0.01),
    rnorm(nrow(mismatchDFMass), 0, 0.01)
    ),
  names = c("Average", "Large")
)

mismatchTestMass <- hypothesis_df("Large > Average",
                                  mismatchDFMassTest,
                                  class = "b", alpha = 0.05
                                  )

mismatchTestMass$hypothesis$Hypothesis =
  "Large Metabolic Slope > Average Slope"

# Summarising

caption <- paste0(
  "Results of a non-linear hypothesis ",
  "tests comparing predicted metabolic slopes in the heat (30°C - 40°C) ",
  "among three week old Japanese quail of average or atypically large ",
  "body size (mean + 2 standard deviations). Posterior probabilities are ",
  "calculated using the Savage-Dickey ",
  "density ratio method."
)

mismatchTestMassTable <- mismatchTestMass$hypothesis %>%
  mutate(Hypothesis = c(
    "Large Metabolic Slope > Average Slope"
  )) %>%
  select(-c(Evid.Ratio, Star)) %>%
  mutate(
    Estimate = round(Estimate, digits = 4),
    "Est.Error" = round(Est.Error, digits = 4),
    "CI.Lower" = round(CI.Lower, digits = 4),
    "CI.Upper" = round(CI.Upper, digits = 4)
  ) %>%
  rename(
    "Difference Between Metabolic Slopes" = Estimate,
    "Standard Error" = Est.Error,
    `2.5 % CI` = "CI.Lower",
    `97.5 % CI` = "CI.Upper",
```

```
    "Posterior Probability" = Post.Prob
  ) %>%
  kbl(.,
    longtable = T, booktabs = T, format = "latex",
    caption = caption
  ) %>%
  column_spec(column = c(1:2), width = "2.5cm") %>%
  column_spec(column = c(3:10), width = "2cm") %>%
  kable_styling(latex_options = "striped")

mismatchTestMassTable
```

***Table 28:*** *Results of a non-linear hypothesis tests comparing predicted metabolic slopes in the heat (30°C - 40°C) among three week old Japanese quail of average or atypically large body size (mean + 2 standard deviations). Posterior probabilities are calculated using the Savage-Dickey density ratio method.*

| Hypothesis | Difference Between Metabolic Slopes | Standard Error | 2.5 % CI | 97.5 % CI | Posterior Probability |
|---|---|---|---|---|---|
| Large Metabolic Slope > Average Slope | 0.0093 | 0.031 | -0.0417 | 0.0586 | 0.6225 |

```
# Tarsus length comparison

mismatchDFTarsus <- data.frame(
  "Size" = c("Average", "Short (2x s.d. < mean)"),
  "pretreatment" = "B",
  "batch" = "A",
  "tarsus" = c(mean(functionModel3WeeksHot$data$tarsus, na.rm = T),
             mean(functionModel3WeeksHot$data$tarsus, na.rm = T) -
               2*sd(functionModel3WeeksHot$data$tarsus, na.rm = T)),
  "mass" = 0
) %>% predict(functionModel3WeeksHot, newdata = .,
              robust = TRUE, re_form = NA,
              resp = "slope", summary = FALSE) %>%
  as.data.frame() %>%
  rename("Average" = "V1", "Long" = "V2") %>%
  mutate_all(.funs = function(x)
    {x + mean(slopeDataHot3$slope, na.rm = T)}
    )

mismatchDFTarsusTest <- build_hdf(
  vars = list(
    mismatchDFTarsus$Average,
    mismatchDFTarsus$Long
    ),
  priors = list(
    rnorm(nrow(mismatchDFTarsus), 0, 0.01),
    rnorm(nrow(mismatchDFTarsus), 0, 0.01)
    ),
  names = c("Average", "Short")
)

mismatchTestTarsus <- hypothesis_df("Short > Average",
                                    mismatchDFTarsusTest,
                                    class = "b", alpha = 0.05
                                    )

mismatchTestTarsus$hypothesis$Hypothesis =
  "Short Tarsus Metabolic Slope > Average Slope"

caption <- paste0(
  "Results of a non-linear hypothesis ",
  "tests comparing predicted metabolic slopes in the cold (30°C - 10°C) ",
  "among three week old Japanese quail of average or atypically long ",
```

```
  "tarsus length (mean + 2 standard deviations). Posterior probabilities are ",
  "calculated using the Savage-Dickey ",
  "density ratio method."
)

mismatchTestTarsusTable <- mismatchTestTarsus$hypothesis %>%
  mutate(Hypothesis = c(
    "Long Metabolic Slope > Average Slope"
  )) %>%
  select(-c(Evid.Ratio, Star)) %>%
  mutate(
    Estimate = round(Estimate, digits = 4),
    "Est.Error" = round(Est.Error, digits = 4),
    "CI.Lower" = round(CI.Lower, digits = 4),
    "CI.Upper" = round(CI.Upper, digits = 4)
  ) %>%
  rename(
    "Difference Between Metabolic Slopes" = Estimate,
    "Standard Error" = Est.Error,
    `2.5 % CI` = "CI.Lower",
    `97.5 % CI` = "CI.Upper",
    "Posterior Probability" = Post.Prob
  ) %>%
  kbl(.,
    longtable = T, booktabs = T, format = "latex",
    caption = caption
  ) %>%
  column_spec(column = c(1:2), width = "2.5cm") %>%
  column_spec(column = c(3:10), width = "2cm") %>%
  kable_styling(latex_options = "striped")

mismatchTestTarsusTable
```

***Table 29:*** *Results of a non-linear hypothesis tests comparing predicted metabolic slopes in the cold (30°C - 10°C) among three week old Japanese quail of average or atypically long tarsus length (mean + 2 standard deviations). Posterior probabilities are calculated using the Savage-Dickey density ratio method.*

| Hypothesis | Difference Between Metabolic Slopes | Standard Error | 2.5 % CI | 97.5 % CI | Posterior Probability |
|---|---|---|---|---|---|
| Long Metabolic Slope > Average Slope | 0.0086 | 0.0311 | -0.0427 | 0.0588 | 0.6135 |

Predicted effects of morphology and rearing conditions on metabolic slopes in the heat are next visualised.

```
showtext.auto(enable = TRUE)

p1 <- data.frame(
  "pretreatment" = c("A", "B", "C"),
  "mass" = 0,
  "tarsus" = 0,
  "batch" = "A"
) %>%
  mutate(
    "slope" = predict(functionModel3WeeksHot,
      newdata = .,
      resp = "slope",
      robust = TRUE,
      re_form = NA
    )[, "Estimate"],
    "SE" = predict(functionModel3WeeksHot,
      newdata = .,
      resp = "slope",
      robust = TRUE,
      re_form = NA
```

```
    )[, "Est.Error"]
  ) %>%
  mutate(slope = slope +
           mean(slopeDataHot3$slope,
                na.rm = T)) %>%
  ggplot(aes(x = pretreatment, y = slope)) +
  geom_errorbar(aes(x = pretreatment, ymin = slope - SE,
                ymax = slope + SE),
                colour = "black", width = 0.3) +
  geom_point(size = 5, pch = 21, colour = "black",
             aes(x = pretreatment, fill = factor(pretreatment))) +
  geom_line(linetype = "dashed", colour = "black") +
  geom_point(
    size = 2, pch = 21, colour = "black",
    data = slopeDataHot3 %>%
      mutate("pretreatment" =
               ifelse(pretreatment == "cold", "A",
        ifelse(pretreatment == "neutral", "B", "C")
      )),
    aes(x = pretreatment, y = slope,
        fill = factor(pretreatment)),
    position = position_jitter(width = 0.3)
  ) +
  scale_fill_manual(values = c("#7BB4E3", "black", "#CD5C5C")) +
  scale_x_discrete(breaks = c("A", "B", "C"),
                   labels = c("Cold\n(10°C)",
                              "Mild\n(20°C)",
                              "Warm\n(30°C)")) +
  theme_classic() +
  theme(legend.position = "none",
        axis.text = element_text(family = "Noto Sans"),
        axis.title = element_text(family = "Noto Sans")) +
  xlab("Rearing Conditions") +
  ylab("Metabolic Slope\n(Fold Metabolism at Thermoneutrality/°C)")

p2 <- data.frame(
  "pretreatment" = "B",
  "mass" = seq(min(functionModel3WeeksHot$data$mass, na.rm = T),
    max(functionModel3WeeksHot$data$mass, na.rm = T),
    by = 0.1
  ),
  "tarsus" = 0,
  "batch" = "A"
) %>%
  mutate(
    "slope" = predict(functionModel3WeeksHot,
      newdata = .,
      resp = "slope",
      robust = TRUE,
      re_form = NA

    )[, "Estimate"],
    "SE" = predict(functionModel3WeeksHot,
      newdata = .,
      resp = "slope",
      robust = TRUE,
      re_form = NA
    )[, "Est.Error"]
  ) %>%
  mutate("Mass" = mass +
           mean(slopeDataHot3$mass, na.rm = T)) %>%
  mutate("slope" = slope +
           mean(slopeDataHot3$slope, na.rm = T)) %>%
  ggplot(aes(x = Mass, y = slope)) +
  geom_ribbon(aes(x = Mass, ymin = slope - SE,
                  ymax = slope + SE),
    fill = "grey50", alpha = 0.5
```

```
  ) +
  geom_smooth(method = "lm", linetype = "dashed",
              se = FALSE, colour = "black") +
  geom_point(
    size = 2.5, pch = 21, colour = "black", alpha = 0.5, fill = "grey50",
    data = slopeDataHot3,
    aes(x = mass, y = slope)
  ) +
  theme_classic() +
  theme(legend.position = "none",
        axis.text = element_text(family = "Noto Sans"),
        axis.title = element_text(family = "Noto Sans")) +
  xlab("Body Mass (g)") +
  ylab("Metabolic Slope\n(Fold Metabolism at\nThermoneutrality/°C)")

p3 <- data.frame(
  "pretreatment" = "B",
  "tarsus" = seq(min(functionModel3WeeksHot$data$tarsus, na.rm = T),
    max(functionModel3WeeksHot$data$tarsus, na.rm = T),
    by = 0.1
  ),
  "mass" = 0,
  "batch" = "A"
) %>%
  mutate(
    "slope" = predict(functionModel3WeeksHot,
      newdata = .,
      resp = "slope",
      robust = TRUE,
      re_form = NA
    )[, "Estimate"],
    "SE" = predict(functionModel3WeeksHot,
      newdata = .,
      resp = "slope",
      robust = TRUE,
      re_form = NA
    )[, "Est.Error"]
  ) %>%
  mutate("Tarsus" = tarsus +
           mean(slopeDataHot3$tarsusLengthMean, na.rm = T)) %>%
  mutate("slope" = slope +
           mean(slopeDataHot3$slope, na.rm = T)) %>%
  ggplot(aes(x = Tarsus, y = slope)) +
  geom_ribbon(aes(x = Tarsus, ymin = slope - SE,
                  ymax = slope + SE),
    fill = "grey50", alpha = 0.5
  ) +
  geom_smooth(method = "lm", linetype = "dashed",
              se = FALSE, colour = "black") +
  geom_point(
    size = 2.5, pch = 21, colour = "black",
    alpha = 0.5, fill = "grey50",
    data = slopeDataHot3,
    aes(x = tarsusLengthMean, y = slope)
  ) +
  theme_classic() +
  theme(legend.position = "none",
        axis.text = element_text(family = "Noto Sans"),
        axis.title = element_text(family = "Noto Sans")) +
  xlab("Tarsus Length (mm)") +
  ylab("Metabolic Slope\n(Fold Metabolism at\nThermoneutrality/°C)")

massPred <- data.frame(
  "pretreatment" = "B",
  "mass" = c(
    mean(functionModel3WeeksHot$data$mass, na.rm = T) -
      sd(functionModel3WeeksHot$data$mass, na.rm = T),
```

```
    mean(functionModel3WeeksHot$data$mass, na.rm = T),
    mean(functionModel3WeeksHot$data$mass, na.rm = T) +
      sd(functionModel3WeeksHot$data$mass, na.rm = T)
  ),
  "tarsus" = 0,
  "batch" = "A"
) %>%
  predict(functionModel3WeeksHot, re_form = NA,
    resp = "slope", newdata = .,
    summary = TRUE, robust = TRUE
  ) %>%
  as.data.frame() %>%
  mutate("slope" = Estimate +
           mean(
             subset(slopeDataHot3,
                    week == 3 & pretreatment == "neutral"
                    )$slope,
             na.rm = TRUE
           )
  ) %>%
  mutate("Size" = c("Small", "Average", "Large")) %>%
  select(Size, slope, "slopeSE" = Est.Error) %>%
  slice(rep(1:n(), each = 2)) %>%
  mutate("Ta" = rep(c(30, 40), 3)) %>%
  mutate("foldRMR" = ifelse(Ta == 30, 1, 10*slope),
         "LL" = ifelse(Ta == 30, 1, 10*(slope - slopeSE)),
         "UL" = ifelse(Ta == 30, 1, 10*(slope + slopeSE))
  ) %>%
  mutate(Size = factor(Size, levels = c("Small", "Average", "Large")))

massPlot <- massPred %>%
  filter(Ta == 40) %>%
  ggplot(aes(x = Size, y = foldRMR)) +
  geom_point(
    data = modDataHot %>%
      select(ring, mass, Ta, VO2) %>%
      pivot_wider(id_cols = c("ring", "mass"),
                  names_from = "Ta",
                  values_from = "VO2") %>%
      mutate("foldRMR" = `40`/`30`,
             "Ta" = 40,
             "Size" = ifelse(mass < mean(mass, na.rm = T) -
                               sd(mass, na.rm = T),
                             "Small",
                             ifelse(mass > mean(mass, na.rm = T) +
                                      sd(mass, na.rm = T),
                                    "Large", "Average")
                             )
             ) %>%
      mutate(Size = factor(Size, levels = c("Small", "Average", "Large"))
             ),
    aes(x = Size, y = foldRMR, colour = mass),
    position = position_jitter(width = 0.25), alpha = 0.5
  ) +
  geom_hline(yintercept = 1, colour = "grey30", linetype = "dashed") +
  geom_errorbar(aes(ymin = LL, ymax = UL),
                colour = "grey80",
                width = 0.25
  ) +
  geom_errorbar(aes(ymin = LL, ymax = UL),
                colour = "black",
                width = 0.25
  ) +
  geom_point(aes(x = Size, y = foldRMR, fill = Size),
             size = 4, colour = "black", pch = 21) +
  ylim(c(0, 4.5)) +
  theme_classic() +
```

```
  ylab("Fold Resting Metabolism") +
  scale_fill_manual(values = c("#DECCC1", "#855E46", "#231709"),
                    guide = NULL) +
  scale_colour_gradient(low = "#DECCC1", high = "#231709",
                        name = "Body Mass (g)") +
  scale_x_discrete(labels = c("Mean - 1 SD", "Mean", "Mean + 1 SD"),
                   name = "Body Mass") +
  theme(
    axis.text = element_text(family = "Noto Sans"),
    axis.title = element_text(family = "Noto Sans"),
    legend.text = element_text(family = "Noto Sans", size = 8),
    legend.title = element_text(family = "Noto Sans", size = 8),
    legend.position = "right"
  )

rm(massPred)

tarsusPred <- data.frame(
  "pretreatment" = "B",
  "tarsus" = c(
    mean(functionModel3WeeksHot$data$tarsus, na.rm = T) -
      sd(functionModel3WeeksHot$data$tarsus, na.rm = T),
    mean(functionModel3WeeksHot$data$tarsus, na.rm = T),
    mean(functionModel3WeeksHot$data$tarsus, na.rm = T) +
      sd(functionModel3WeeksHot$data$tarsus, na.rm = T)
  ),
  "mass" = 0,
  "batch" = "A"
) %>%
  predict(functionModel3WeeksHot, re_form = NA,
    resp = "slope", newdata = .,
    summary = TRUE, robust = TRUE
  ) %>%
  as.data.frame() %>%
  mutate("slope" = Estimate +
           mean(
             subset(slopeDataHot3,
                    pretreatment == "neutral"
                    )$slope,
             na.rm = TRUE
           )
  ) %>%
  mutate("Size" = c("Small", "Average", "Large")) %>%
  select(Size, slope, "slopeSE" = Est.Error) %>%
  slice(rep(1:n(), each = 2)) %>%
  mutate("Ta" = rep(c(30, 40), 3)) %>%
  mutate("foldRMR" = ifelse(Ta == 30, 1, 10*slope),
         "LL" = ifelse(Ta == 30, 1, 10*(slope - slopeSE)),
         "UL" = ifelse(Ta == 30, 1, 10*(slope + slopeSE))
  ) %>%
  mutate(Size = factor(Size, levels = c("Small", "Average", "Large")))

tarsusPlot <- tarsusPred %>%
  filter(Ta == 40) %>%
  ggplot(aes(x = Size, y = foldRMR)) +
  geom_point(
    data = modDataHot %>%
      select(ring, "tarsus" = tarsusLengthMean, Ta, VO2) %>%
      pivot_wider(id_cols = c("ring", "tarsus"),
                  names_from = "Ta",
                  values_from = "VO2") %>%
      mutate("foldRMR" = `40`/`30`,
             "Ta" = 40,
             "Size" = ifelse(tarsus < mean(tarsus, na.rm = T) -
                               sd(tarsus, na.rm = T),
                             "Small",
                             ifelse(tarsus > mean(tarsus, na.rm = T) +
```

```
                                      sd(tarsus, na.rm = T),
                                    "Large", "Average")
             )
      ) %>%
      drop_na() %>%
      mutate(Size = factor(Size, levels = c("Small", "Average", "Large"))
      ),
    aes(x = Size, y = foldRMR, colour = tarsus),
    position = position_jitter(width = 0.25), alpha = 0.5
  ) +
  geom_hline(yintercept = 1, colour = "grey30", linetype = "dashed") +
  geom_errorbar(aes(ymin = LL, ymax = UL),
                colour = "grey80",
                width = 0.25
  ) +
  geom_errorbar(aes(ymin = LL, ymax = UL),
                colour = "black",
                width = 0.25
  ) +
  geom_point(aes(x = Size, y = foldRMR, fill = Size),
             size = 4, colour = "black", pch = 21) +
  theme_classic() +
  xlab("Ambient Temperature (°C)") +
  ylab("Fold Resting Metabolism") +
  ylim(c(0, 4.5)) +
  scale_fill_manual(values = c("#DECCC1", "#855E46", "#231709"),
                    guide = NULL) +
  scale_colour_gradient(low = "#DECCC1", high = "#231709",
                        name = "Tarsus Length\n(mm)") +
  theme(
    axis.text = element_text(family = "Noto Sans"),
    axis.title = element_text(family = "Noto Sans"),
    legend.text = element_text(family = "Noto Sans", size = 8),
    legend.title = element_text(family = "Noto Sans", size = 8),
    legend.position = "right"
  )

allMorphology <- (p1 / (p2 + massPlot) / (p3 + tarsusPlot)) +
  plot_annotation(tag_levels = "A")

allMorphology
```

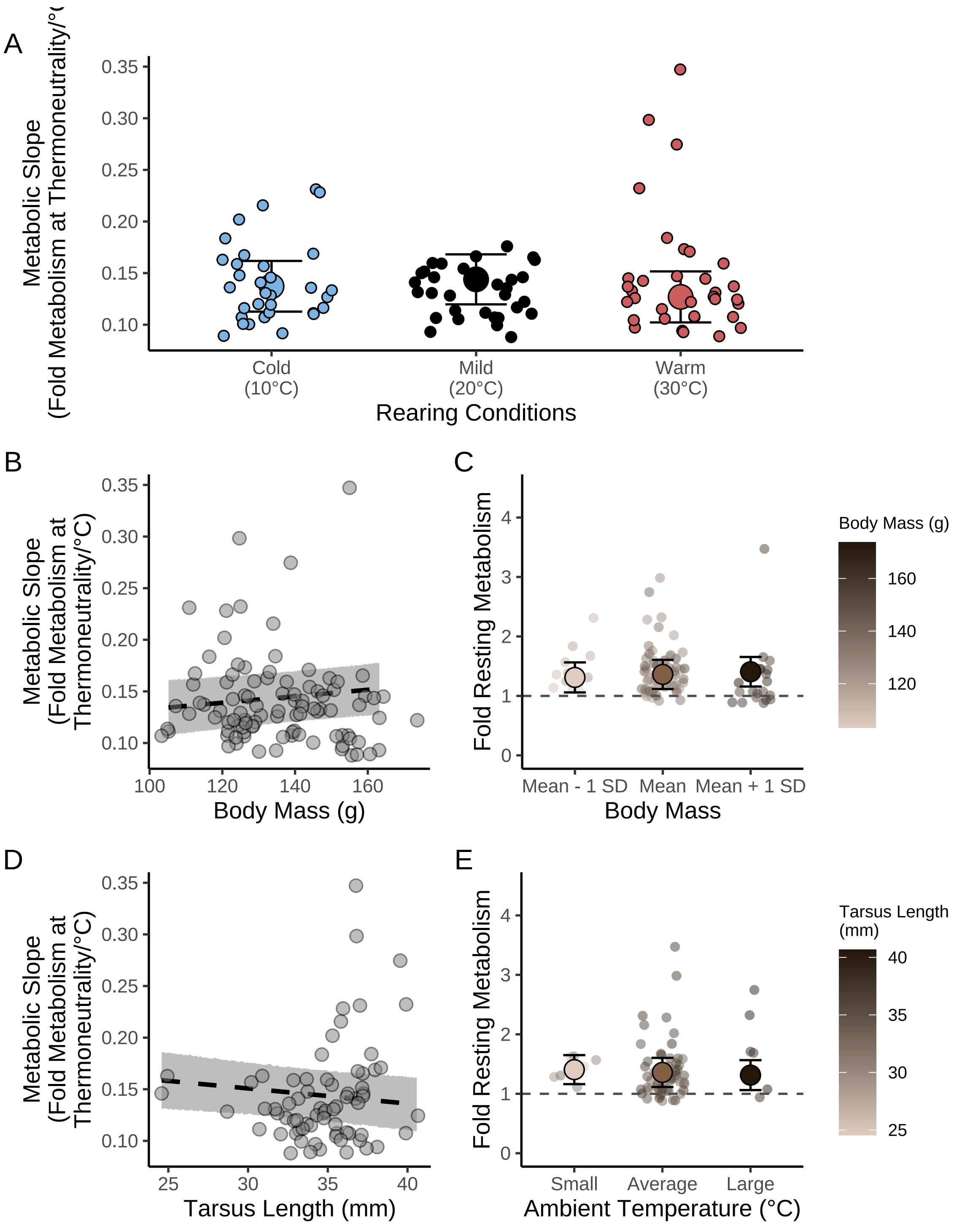


***Figure 59:*** *Conditional effects of post-hatch rearing environments (10°C, 20°C or 30°C) and morphometry (tarsus length [mm] and body mass [g]) on physiological responses to heat in three week old Japanese quail. Dots represent raw data points per individual. Large dots in panel A and B represent estimated effects at average morphometric values and errorbars indicate +/- one standard error around means. Lines in panels C to F indicate estimate effects across given parameters and grouping (assuming mild [20°C] rearing conditions) and ribbons indicate standard errors around effects. All estimated effects are drawn from a Bayesian path analysis.*

```
ggsave("../plots/morphologyEffectsResistanceHeat3Weeks_3.pdf",
  allMorphology,
  dpi = 800, height = 7, width = 9
)

showtext.auto(enable = "FALSE")
```

Next, the above analysis is repeated for data derived from eight week old Japanese quail. Model parameterisation and priors remain as described but with slight modifications described below within this subsection. We begin by calculating metabolic slopes manually and visualising resulting values for oddities. Then, raw trends in the data are inspected with plots, and our model constructed.

```
birdVector <- merge(
  all %>%
    drop_na(VO2) %>%
    filter(week == 8 & Ta == 30) %>%
    select(ring) %>%
    distinct(),
  all %>%
    drop_na(VO2) %>%
    filter(week == 8 & Ta == 40) %>%
    select(ring) %>%
    distinct(),
  by = "ring", all = FALSE
) %>% pull(ring)

modData8 <- subset(all, week == 8 & Ta > 25 &
                     ring %in% birdVector)

# Calculating metabolic slopes for each
# bird. Again, 30°C is set as 0 to root slopes.

slopeDataHot8 <- modData8 %>%
  select(ring, Ta, VO2) %>%
  pivot_wider(id_cols = ring, names_from = Ta,
              values_from = VO2) %>%
  mutate(
    slope = (`40`/`30`)/10
  ) %>%
  select(ring, slope) %>%
  merge(modData8 %>%
    select(
      ring, sex, exp, week, pretreatment,
      posttreatment, treatment, mass, wingLength,
      tarsusLengthMean,
    ) %>%
    distinct(), ., by = "ring")

# Building dotplot

slopeDataHot8 %>%
  ggplot(aes(x = 1:nrow(.), y = slope)) +
  geom_rect(
    colour = "black", fill = "grey80", alpha = 0.5,
    aes(
      xmin = -Inf, xmax = Inf,
      ymin = mean(slope, na.rm = T) -
        3.5 * sd(slope, na.rm = T),
      ymax = mean(slope, na.rm = T) +
        3.5 * sd(slope, na.rm = T)
    )
  ) +
  geom_point() +
  xlab("Sample Number") +
  ylab("Metabolic Slope\n(Fold Resting Metabolism/°C)") +
  theme_classic()
```

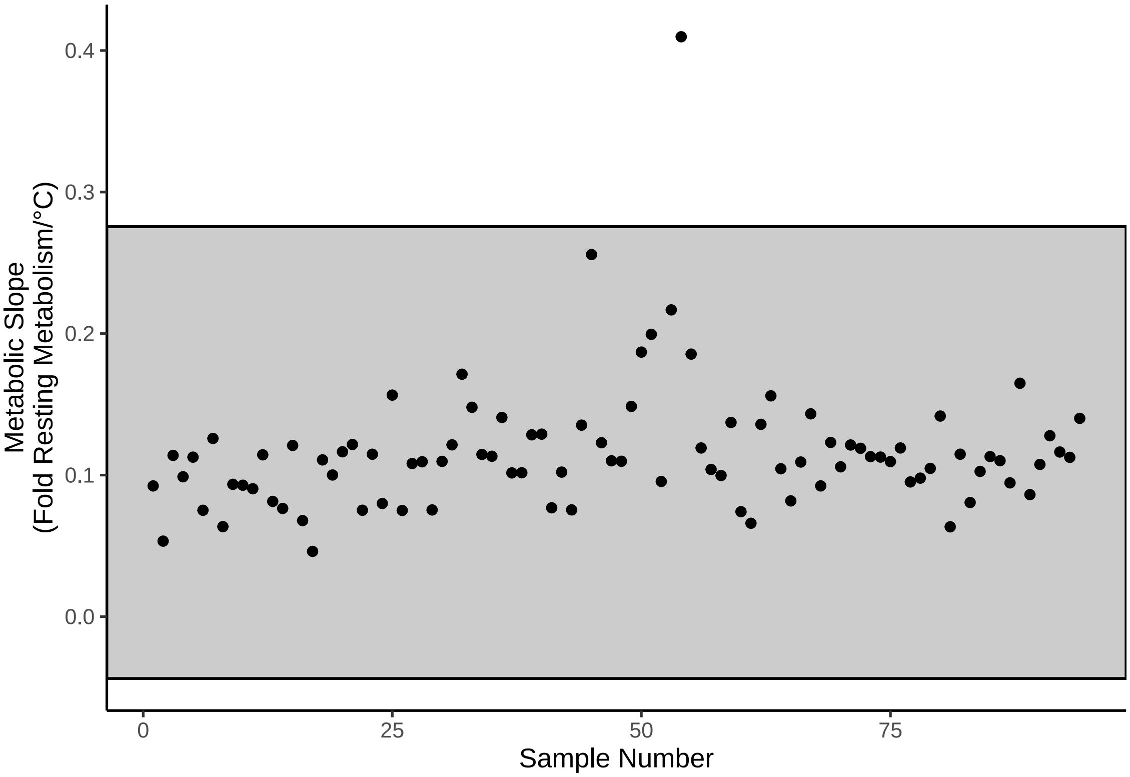


***Figure 60:*** *Cleveland dotplot displaying metabolic slope values of eight week old Japanese quail at 30°C - 40°C. Dots represent raw values and the grey rectangle represents the range covered by mean metabolic slope +/- 3.5x the standard deviation.*

Again, a potential outlier evident in our dotplot is investigated.

```
# One high value, as above. Identifying.

caption <- paste0("Potential metabolic slope outlier ",
                  "among eight week old Japanese quail with metabolism ",
                  "measured at ambient temperatures between ",
                  "30°C and 40°C."
)

slopeDataHot8 %>%
  filter(slope > 0.4) %>%
  merge(., modData8 %>%
    select(ring, Ta, VO2),
  by = "ring", all.x = TRUE
  ) %>%
  mutate(pretreatment = ifelse(pretreatment == "cold",
                               "Cold (10°C)",
                               ifelse(pretreatment == "warm",
                                      "Warm (30°C)", "Mild (20°C)")
                               )
         ) %>%
  select(
    "Bird Identity" = ring,
    "Rearing Conditions" = pretreatment,
    "Mass (g)" = mass,
    "Tarsus Length (mm)" = tarsusLengthMean,
    "Ambient Temperature (°C)" = Ta,
    "Metabolic Slope\n(Fold Resting Metabolism at Thermoneutrality/°C)" =
      slope
```

```
  ) %>%
  kbl(., longtable =T,booktabs =T,
      format = "latex", escape = FALSE,
      caption = caption) %>%
  column_spec(column = c(2,4:7), width = "2.1cm") %>%
  kable_styling(latex_options = "striped")
```

***Table 30:*** *Potential metabolic slope outlier among eight week old Japanese quail with metabolism measured at ambient temperatures between 30°C and 40°C.*

| Bird Identity | Rearing Conditions | Mass (g) | Tarsus Length (mm) | Ambient Temperature (°C) | Metabolic Slope (Fold Resting Metabolism at Thermoneutrality/°C) |
|---|---|---|---|---|---|
| LLW26 | Warm (30°C) | 306 | 37.5893 | 30 | 0.4097247 |
| LLW26 | Warm (30°C) | 306 | 37.5893 | 40 | 0.4097247 |

The individual from which this extreme point came was also flagged as extreme at three weeks of age as well. At this time, however, its response now lays more than six standard deviations above the mean. Below, we retain the value but check its relative importance on our analysis outcomes, again, using Pareto K as our metric of importance.

```
# Visualising data

p1 <- ggplot(slopeDataHot8, aes(x = slope)) +
  geom_density(colour = "black", fill = "grey50", alpha = 0.5) +
  theme_classic() +
  xlab(
    paste0("Metabolic Slope\n(Fold Resting ",
    "Metabolism at\nThermoneutrality/°C)"
    )
  ) +
  ylab("Density")

p2 <- ggplot(
  modData8 %>%
    mutate(
      "tarsusGroup" =
        ifelse(tarsusLengthMean <
          mean(tarsusLengthMean, na.rm = T) -
            sd(tarsusLengthMean, na.rm = T),
        "Small",
        ifelse(tarsusLengthMean >
          mean(tarsusLengthMean, na.rm = T) +
            sd(tarsusLengthMean, na.rm = T),
        "Large", "Average"
        )
        )
    ) %>%
    mutate(
      tarsusGroup =
        factor(tarsusGroup,
          levels = c("Small", "Average", "Large")
        )
    ) %>%
    mutate(pretreatment = ifelse(pretreatment == "cold",
                                 "Cold\n(10°C)",
      ifelse(pretreatment == "neutral", "Mild\n(20°C)",
        "Warm\n(30°C)"
      )
    )),
  aes(x = Ta, y = VO2, group = ring, colour = tarsusGroup)
```

```
) +
  facet_wrap(~pretreatment) +
  geom_line(aes(size = tarsusGroup)) +
  scale_size_manual(
    values = c("Small" = 0.5, "Average" = 1, "Large" = 2),
    labels = c("< Mean - 1SD", "Mean", "> Mean + 1SD"),
    name = "Relative Tarsus\nLength"
  ) +
  scale_colour_manual(
    values = c(
      "Small" = "grey80",
      "Average" = "grey30",
      "Large" = "black"
    ),
    name = "Relative Tarsus\nLength",
    labels = c("< Mean - 1SD", "Mean", "> Mean + 1SD"),
  ) +
  scale_x_continuous(breaks = c(30, 40)) +
  theme_classic() +
  theme(legend.position = "bottom") +
  xlab("Ambient Temperature (°C)") +
  ylab(
   TeX(
    '$\\overset{Resting~Metabolism}{(mL~O_{2}/min)}$'
    )
   )

p3 <- ggplot(
  modData8 %>%
    mutate(
      "massGroup" =
        ifelse(mass <=
          mean(mass, na.rm = T) -
            sd(mass, na.rm = T),
        "Small",
        ifelse(mass >=
          mean(mass, na.rm = T) +
            sd(mass, na.rm = T),
        "Large", "Average"
        )
        )
    ) %>%
    mutate(
      massGroup =
        factor(massGroup,
          levels = c("Small", "Average", "Large")
        )
    ) %>%
    mutate(pretreatment =
             ifelse(pretreatment == "cold", "Cold\n(10°C)",
      ifelse(pretreatment == "neutral", "Mild\n(20°C)",
        "Warm\n(30°C)"
      )
    )),
  aes(x = Ta, y = VO2, group = ring, colour = massGroup)
) +
  facet_wrap(~pretreatment) +
  geom_line(aes(size = massGroup)) +
  scale_size_manual(
    values = c("Small" = 0.5, "Average" = 1, "Large" = 2),
    name = "Relative Body\nMass",
    labels = c("< Mean - 1SD", "Mean", "> Mean + 1SD"),
  ) +
  scale_colour_manual(
    values = c(
      "Small" = "grey80",
      "Average" = "grey30",
```

```
      "Large" = "black"
    ),
    name = "Relative Body\nMass",
    labels = c("< Mean - 1SD", "Mean", "> Mean + 1SD"),
  ) +
  scale_x_continuous(breaks = c(30, 40)) +
  theme_classic() +
  theme(legend.position = "bottom") +
  xlab("Ambient Temperature (°C)") +
  ylab(
   TeX(
    '$\\overset{Resting~Metabolism}{(mL~O_{2}/min)}$'
    )
   )

(p1/p2/p3) +
  plot_annotation(tag_levels = "A")
```

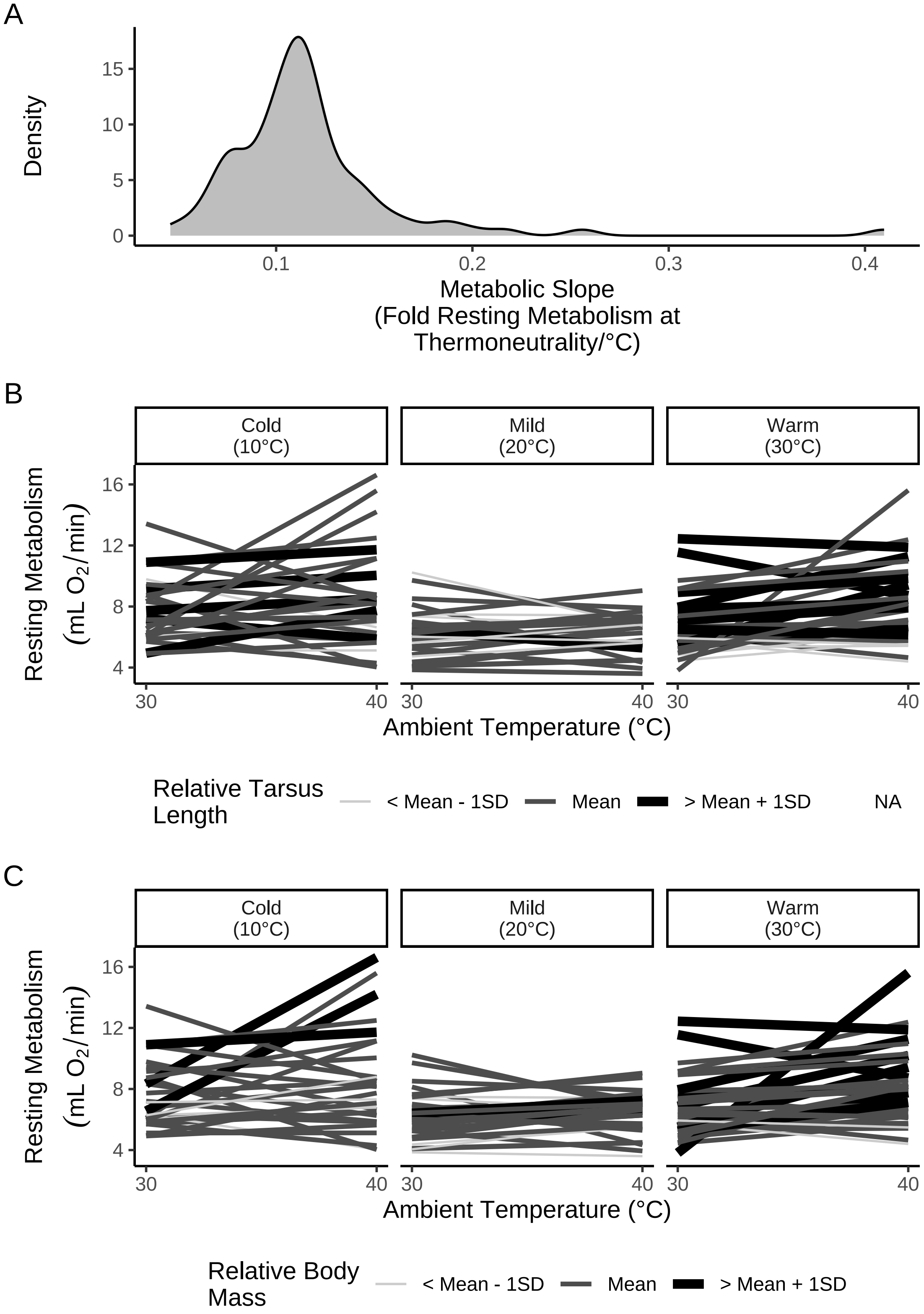


***Figure 61:*** *Thermal physiology of eight week old Japanese quail at ambient temperature within and about thermoneutral ( 30°C). Panel A displays the density of metabolic slopes at this temperature range. Panels B and C display correlations between ambient temperature and resting metabolism. Each line represents one individual with line colour and line size scaled by relative size (tarsus length [mm] or body mass [g]); data is divided by rearing conditions up to at least three months of age.*

```
p1 <- ggplot(slopeDataHot8,
             aes(x = tarsusLengthMean, y = slope)) +
  geom_point(pch = 21, size = 1.5,
             colour = "black", fill = "grey50") +
  geom_smooth(method = "lm", se = FALSE,
              colour = "black", linetype = "dashed") +
  theme_classic() +
  scale_x_continuous(n.breaks = 4) +
  xlab("Tarsus Length\n(mm)") +
  ylab("Metabolic Slope\n(Fold Metabolism at Thermoneutrality/°C)")

p2 <- ggplot(slopeDataHot8,
             aes(x = mass, y = slope)) +
  geom_point(pch = 21, size = 1.5,
             colour = "black", fill = "grey50") +
  geom_smooth(method = "lm", se = FALSE,
              colour = "black", linetype = "dashed") +
  xlab("Body Mass (g)") +
  theme_classic() +
  theme(axis.title.y = element_blank(),
        axis.text.y = element_blank(),
        axis.ticks.y = element_blank(),
        axis.line.y = element_blank())

p3 <- ggplot(
  slopeDataHot8 %>%
    drop_na(tarsusLengthMean, mass) %>%
    mutate("tarsusResiduals" = residuals(
      brm(tarsusLengthMean ~ mass,
        data = .,
        prior = c(set_prior("gamma(1, 1)",
                            class = "b", coef = "mass")),
        iter = 50000, warmup = 5000,
        cores = 4, chains = 4,
        silent = TRUE, refresh = 0,
        file = "./models/resPlotModelStudent.Rds"
      )
    )[, "Estimate"]),
  aes(x = tarsusResiduals, y = slope)
) +
  geom_point(pch = 21, size = 1.5,
             colour = "black", fill = "grey50") +
  geom_smooth(method = "lm", se = FALSE,
              colour = "black", linetype = "dashed") +
  xlab("Residual Tarsus\nLength (mm)") +
  theme_classic() +
  theme(axis.title.y = element_blank(),
        axis.text.y = element_blank(),
        axis.ticks.y = element_blank(),
        axis.line.y = element_blank())

(p1 + p2 + p3)
```

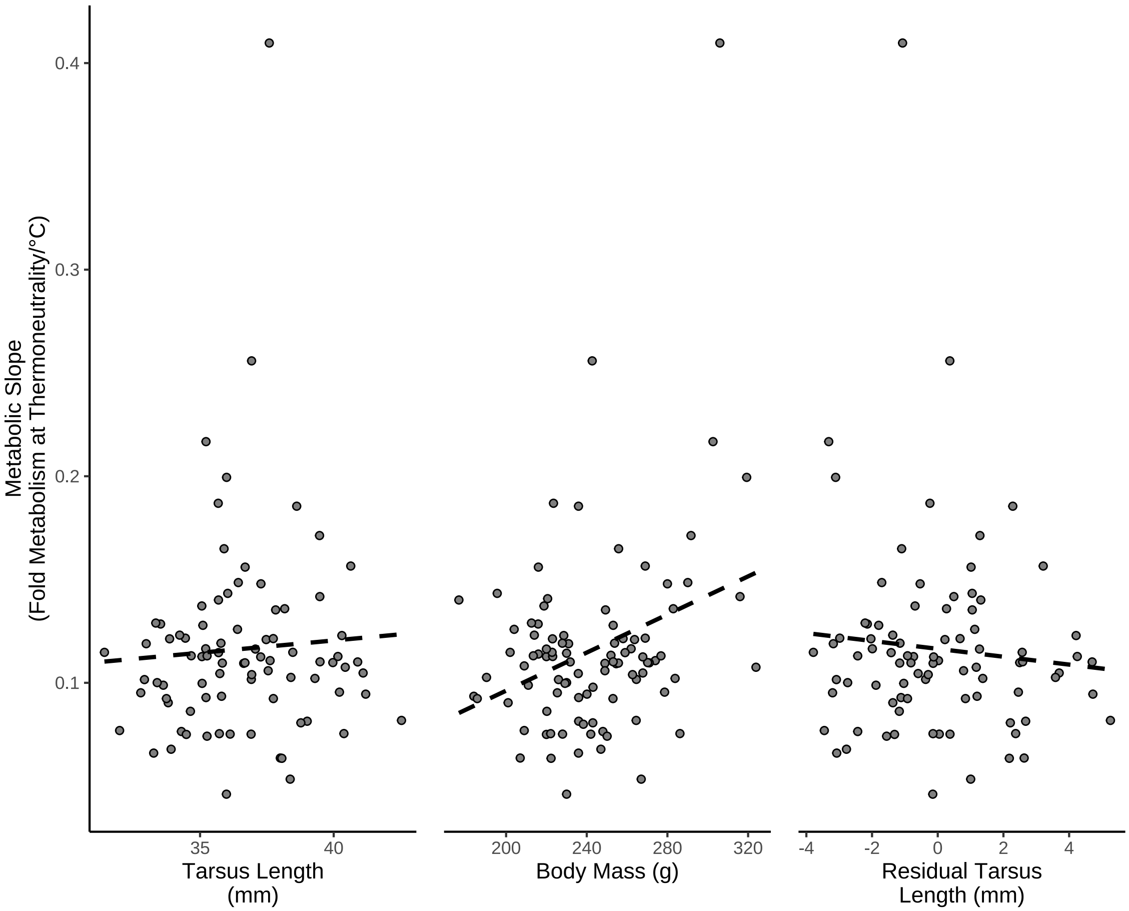


***Figure 62:*** *Raw correlations between morphology and metabolic slopes above thermoneutrality in 8 week old Japanese quail. Metabolic slopes are measured as the fold change in resting metabolism between 30°C and 40°C, divided by 10°C. Relative tarsus length indicates ordinary residuals from a Bayesian regression of tarsus length (mm) against body mass (g). Dots indicate raw values per individual and dashed lines indicate estimated relationships as predicted by the R package ggplot2 (Wickham, 2011).*

And again, we check whether variance in metabolic slopes differs between experimental batches owing to differences in flow-rates.

```
slopeDataHot8 %>%
  rename("batch" = exp) %>%
  filter(slope < 0.4) %>%
  ggplot(aes(x = slope, fill = batch)) +
  geom_density(colour = "black", alpha = 0.5) +
  scale_fill_manual(values = c("black", "firebrick", "beige"),
                    labels = c("1", "2", "3"),
                    name = "Experimental\nBatch") +
  theme_classic() +
  xlab("Metabolic Slope\n(Fold Metabolism at Thermoneutrality") +
  ylab("Density")
```

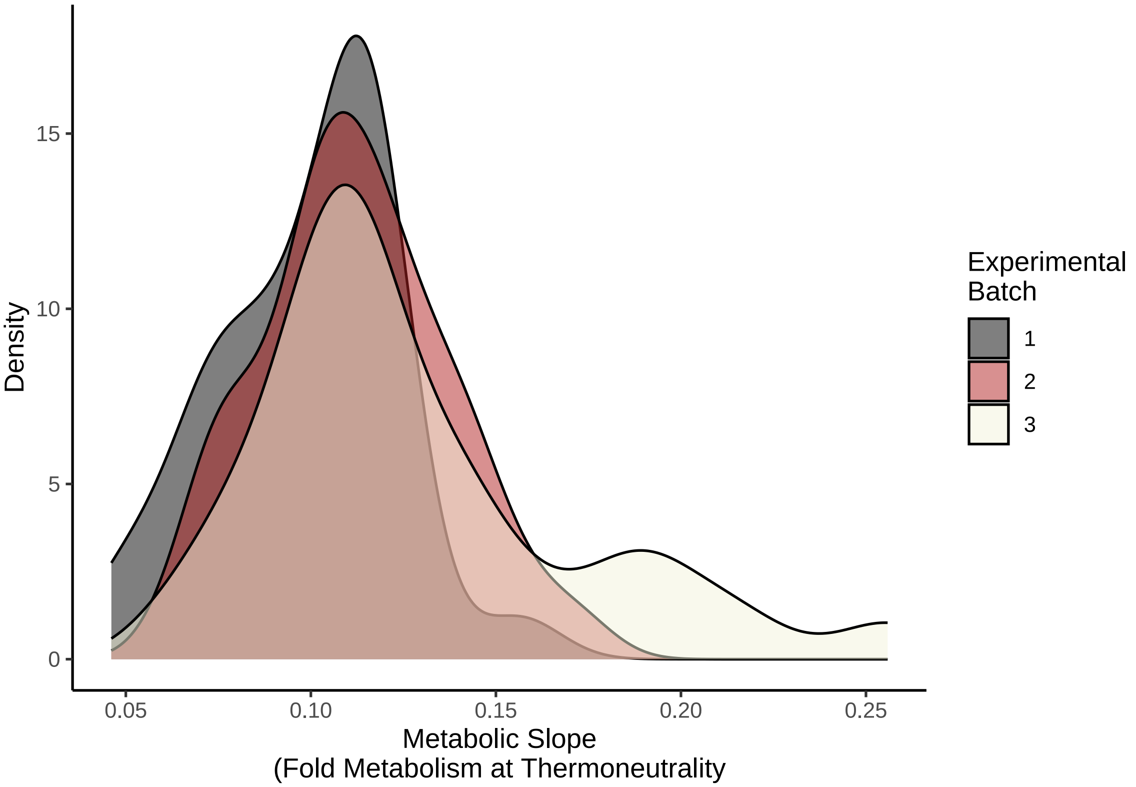


***Figure 63:*** *Densities of metabolic slope values below 30°C from eight week old Japanese quail. Densities are divided by experimental batches.*

Here, variance appears largely similar between batches. As such, no adjustments to the error term of our model predicting metabolic slope at this age are made.

For our path analysis of data from mature quail, we used models and priors largely equivalent to those applied to our data from three week old quail. Minor adjustments are made to priors to accommodate: (1) an increased variance in body size and tarsus length, and (2) a decrease in metabolic slope at this age (see Persson et al, 2024). These adjustments are as follows:

$$\beta_{a0} \sim \mathcal{N}(0, 10)$$
$$\beta_{a1} \sim \mathcal{N}(0, 25)$$
$$\beta_{a2} \sim \mathcal{N}(0, 25)$$
$$\mu_{0a} \sim exp(2.5)$$
$$\epsilon_a \sim exp(0.05)$$

$$\beta_{b0} \sim \mathcal{N}(0, 3)$$
$$\beta_{b1} \sim \mathcal{N}(0, 3)$$
$$\beta_{b2} \sim \mathcal{N}(0, 3)$$
$$\beta_{b3} \sim S\mathcal{N}(0, 0.25, 5)$$
$$\mu_{0b} \sim exp(2)$$

$$\epsilon_b \sim exp(0.75)$$

and:

$$\beta_{c0} \sim \mathcal{N}(0, 0.1)$$
$$\beta_{c1} \sim \mathcal{N}(0, 0.1)$$
$$\beta_{c2} \sim \mathcal{N}(0, 0.1)$$
$$\beta_{c3} \sim \mathcal{N}(0, 0.0025)$$
$$\beta_{c4} \sim \mathcal{N}(0, 0.03)$$
$$\mu_{0c} \sim exp(50)$$
$$\epsilon_c \sim exp(0.75)$$

with all other priors remaining the same as those for data from three week old quail.

```
# Running prior predictive check

functionModel8WeeksHotPPCheck <-
  brm(
    data = slopeDataHot8 %>%
      mutate(
        mass = mass - mean(mass, na.rm = T),
        tarsus = tarsusLengthMean -
          mean(tarsusLengthMean, na.rm = T),
        pretreatment = ifelse(pretreatment == "cold", "A",
          ifelse(pretreatment == "neutral", "B", "C")
        ),
        slope = slope -
          mean(slope, na.rm = T)
      ) %>%
      mutate(pretreatment = factor(pretreatment,
        levels = c("B", "A", "C")
      )) %>%
      drop_na() %>%
      merge(., data %>%
        select(ring, "batch" = exp) %>%
        distinct(),
      by = "ring", all.x = TRUE
      ),
    family = "gaussian",
    bf(mass ~ pretreatment + (1 | batch)) +
      bf(tarsus ~ mass + pretreatment + (1 | batch)) +
      bf(slope ~ mass + tarsus + pretreatment + (1 | batch),
       sigma ~ batch) +
      set_rescor(FALSE),
    prior = c(
      set_prior("normal(0, 10)",
        class = "Intercept",
        resp = "mass"
      ),
      set_prior("normal(0, 25)",
        class = "b",
        coef = "pretreatmentA",
        resp = "mass"
      ),
      set_prior("normal(0, 25)",
        class = "b",
        coef = "pretreatmentC",
        resp = "mass"
      ),
```

```
    set_prior("exponential(2.5)",
      class = "sd",
      group = "batch",
      resp = "mass"
    ),
    set_prior("exponential(0.05)",
      class = "sigma",
      resp = "mass"
    ),
    set_prior("normal(0, 3)",
      class = "Intercept",
      resp = "tarsus"
    ),
    set_prior("normal(0, 3)",
      class = "b",
      coef = "pretreatmentA",
      resp = "tarsus"
    ),
    set_prior("normal(0, 3)",
      class = "b",
      coef = "pretreatmentC",
      resp = "tarsus"
    ),
    set_prior("skew_normal(0, 0.25, 5)",
      class = "b",
      coef = "mass",
      resp = "tarsus"
    ),
    set_prior("exponential(2)",
      class = "sd",
      group = "batch",
      resp = "tarsus"
    ),
    set_prior("exponential(0.75)",
      class = "sigma",
      resp = "tarsus"
    ),
    set_prior("normal(0, 0.1)",
      class = "Intercept",
      resp = "slope"
    ),
    set_prior("normal(0, 0.1)",
      class = "b",
      coef = "pretreatmentA",
      resp = "slope"
    ),
    set_prior("normal(0, 0.1)",
      class = "b",
      coef = "pretreatmentC",
      resp = "slope"
    ),
    set_prior("normal(0, 0.0025)",
      class = "b",
      coef = "mass",
      resp = "slope"
    ),
    set_prior("normal(0, 0.03)",
      class = "b",
      coef = "tarsus",
      resp = "slope"
    ),
    set_prior("exponential(50)",
      class = "sd",
      group = "batch",
      resp = "slope"
    ),
    set_prior("normal(-3, 1.5)",
```

```
        dpar = "sigma",
        class = "Intercept",
        resp = "slope"
      ),
      set_prior("normal(0, 0.5)",
        dpar = "sigma",
        class = "b",
        coef = "batchB",
        resp = "slope"
      ),
      set_prior("normal(1, 1.5)",
        dpar = "sigma",
        class = "b",
        coef = "batchC",
        resp = "slope"
      )
      ),
    iter = 50000, warmup = 10000, cores = 4, chains = 4, thin = 20,
    control = list(adapt_delta = .97, max_treedepth = 14),
    silent = TRUE, refresh = 0,
    sample_prior = "only",
    file = "./models/eightWeekFunctionModelHotPPCheckNew2.Rds"
  )

pp_check2(functionModel8WeeksHotPPCheck,
          resp = "slope",
          xlab = paste0("Metabolic Slope\n",
                        "(Fold Metabolism at Thermoneutrality/°C)"
                        )
          )
```

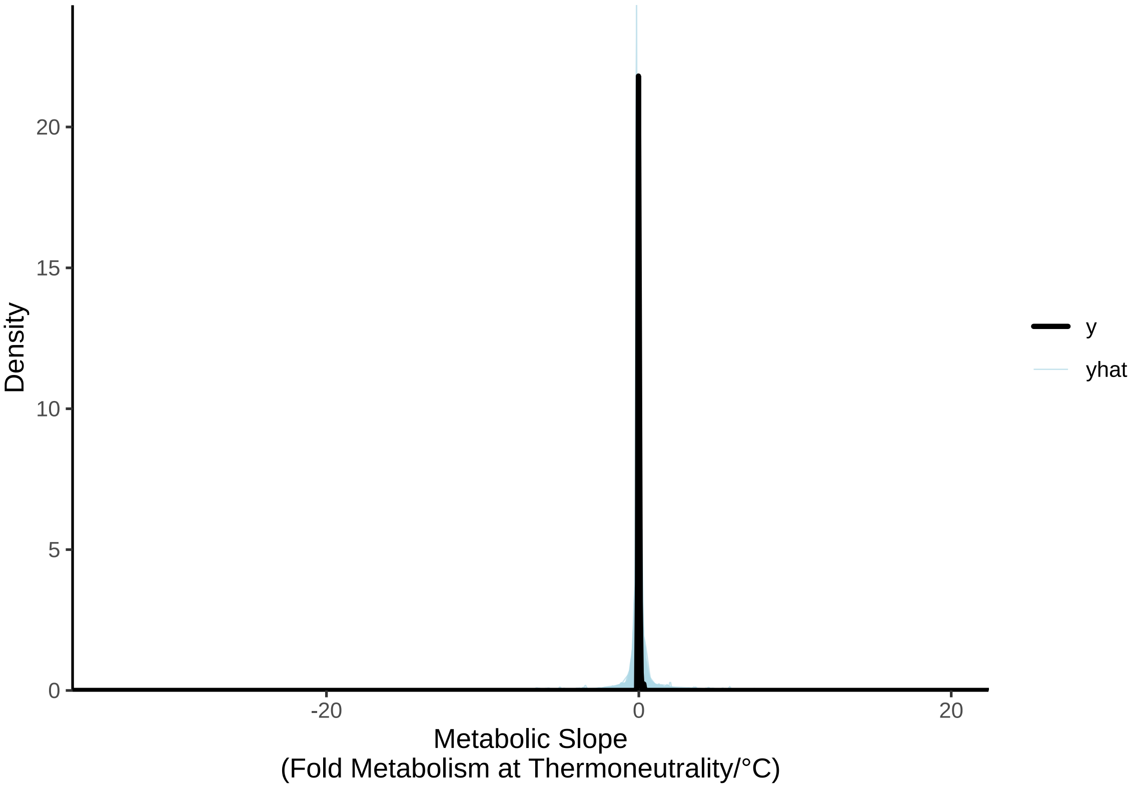


***Figure 64:*** *Prior predictive check for a Bayesian path analysis predicting physiological responses to heat in eight week old Japanese quail. Black lines represent true density of metabolic slopes (fold metabolism at thermoneutrality/°C) while blue lines represent densities estimated from model priors alone.*

```
# Priors clearly not constraining. Constructing full model.

functionModel8WeeksHot <-
  brm(
    data = slopeDataHot8 %>%
      mutate(
        mass = mass - mean(mass, na.rm = T),
        tarsus = tarsusLengthMean -
          mean(tarsusLengthMean, na.rm = T),
        pretreatment = ifelse(pretreatment == "cold", "A",
          ifelse(pretreatment == "neutral", "B", "C")
        ),
        slope = slope -
          mean(slope, na.rm = T)
      ) %>%
      mutate(pretreatment = factor(pretreatment,
        levels = c("B", "A", "C")
      )) %>%
      drop_na() %>%
      merge(., data %>%
        select(ring, "batch" = exp) %>%
        distinct(),
      by = "ring", all.x = TRUE
      ),
      family = "gaussian",
      bf(mass ~ pretreatment + (1 | batch)) +
      bf(tarsus ~ mass + pretreatment + (1 | batch)) +
      bf(slope ~ mass + tarsus + pretreatment + (1 | batch)) +
      set_rescor(FALSE),
    prior = c(
      set_prior("normal(0, 10)",
        class = "Intercept",
        resp = "mass"
      ),
      set_prior("normal(0, 25)",
        class = "b",
        coef = "pretreatmentA",
        resp = "mass"
      ),
      set_prior("normal(0, 25)",
        class = "b",
        coef = "pretreatmentC",
        resp = "mass"
      ),
      set_prior("exponential(2.5)",
        class = "sd",
        group = "batch",
        resp = "mass"
      ),
      set_prior("exponential(0.05)",
        class = "sigma",
        resp = "mass"
      ),
      set_prior("normal(0, 3)",
        class = "Intercept",
        resp = "tarsus"
      ),
      set_prior("normal(0, 3)",
        class = "b",
        coef = "pretreatmentA",
        resp = "tarsus"
      ),
      set_prior("normal(0, 3)",
        class = "b",
        coef = "pretreatmentC",
        resp = "tarsus"
      ),
```

```
      set_prior("skew_normal(0, 0.25, 5)",
        class = "b",
        coef = "mass",
        resp = "tarsus"
      ),
      set_prior("exponential(2)",
        class = "sd",
        group = "batch",
        resp = "tarsus"
      ),
      set_prior("exponential(0.75)",
        class = "sigma",
        resp = "tarsus"
      ),
      set_prior("normal(0, 0.1)",
        class = "Intercept",
        resp = "slope"
      ),
      set_prior("normal(0, 0.1)",
        class = "b",
        coef = "pretreatmentA",
        resp = "slope"
      ),
      set_prior("normal(0, 0.1)",
        class = "b",
        coef = "pretreatmentC",
        resp = "slope"
      ),
      set_prior("normal(0, 0.0025)",
        class = "b",
        coef = "mass",
        resp = "slope"
      ),
      set_prior("normal(0, 0.03)",
        class = "b",
        coef = "tarsus",
        resp = "slope"
      ),
      set_prior("exponential(50)",
        class = "sd",
        group = "batch",
        resp = "slope"
      ),
      set_prior("exponential(10)",
        class = "sigma",
        resp = "slope"
      )
    ),
    iter = 50000, warmup = 10000, cores = 4, chains = 4, thin = 20,
    control = list(adapt_delta = .97, max_treedepth = 14),
    silent = TRUE, refresh = 0,
    file = "./models/eightWeekFunctionModelHot.Rds"
  )

chainCheck(functionModel8WeeksHot)
```

```
## Rhat range: 1 - 1.001
## Neff/N range: 0.92 - 1.007
```

```
# Great. Very little attrition in effective sample sizes.

caption <- paste0("R\\textsuperscript{2} for a Bayesian ",
                  "path analysis predicting metabolic slopes ",
                  "(fold metabolism at thermoneutrality/°C) of ",
                  "eight week old Japanese quail as a function of ",
                  "body mass (g) and tarsus length (mm). Metabolic slopes ",
                  "are measured above thermoneutrality (30°C - 40°C)."
)
```

```
brms::bayes_R2(functionModel8WeeksHot,
               ndraws = 1000,
               robust = TRUE) %>%
  as.data.frame() %>%
  rownames_to_column("var") %>%
  merge(., tribble(
    ~var, ~Var,
    "R2mass", "Body Mass (g)",
    "R2tarsus", "Tarsus Length (mm)",
    "R2slope",
      "Metabolic Slope"
  ), by = c("var")) %>%
  mutate(Estimate = round(Estimate, digits = 4),
         Est.Error = round(Est.Error, digits = 4),
         `95\\% CI` = paste0("[",
                             round(Q2.5, digits = 4),
                             ", ",
                             round(Q97.5, digits = 4),
                             "]"
         )
         ) %>%
  select(
    "Response" = Var, "R\\textsuperscript{2}" = Estimate,
    "Standard Error" = Est.Error, `95\\% CI`
  ) %>%
  kbl(.,
    longtable = T, booktabs = T, format = "latex",
    caption = caption, escape = FALSE
  ) %>%
  column_spec(column = c(1:10), width = "2.5cm")  %>%
  kable_styling(latex_options = "striped")
```

***Table 31:*** *$R^2$ for a Bayesian path analysis predicting metabolic slopes (fold metabolism at thermoneutrality/°C) of eight week old Japanese quail as a function of body mass (g) and tarsus length (mm). Metabolic slopes are measured above thermoneutrality (30°C - 40°C).*

| Response | $R^2$ | Standard Error | 95% CI |
|---|---|---|---|
| Body Mass (g) | 0.0391 | 0.0338 | [0.0017, 0.1275] |
| Metabolic Slope | 0.2188 | 0.0695 | [0.0951, 0.3347] |
| Tarsus Length (mm) | 0.2954 | 0.0736 | [0.144, 0.411] |

```
# R2 for metabolic slope slightly increased relative
# to three week old data.
```

```
ggarrange(
  pp_check2(functionModel8WeeksHot, resp = "slope",
    xlab = "Metabolic Slope\n(Fold Metabolism at\nThermoneutrality/°C)"
  ),

  functionModel8WeeksHot$data %>%
    mutate(
      "Fit" = fitted(functionModel8WeeksHot,
                     resp = "slope",
                     robust = TRUE)[, "Estimate"],
      "SE" = fitted(functionModel8WeeksHot,
                    resp = "slope",
                     robust = TRUE)[, "Est.Error"]
    ) %>%
    ggplot(aes(x = Fit, y = slope)) +
    geom_errorbarh(aes(y = slope, xmin = Fit - SE,
                       xmax = Fit + SE),
      colour = "black", height = 0.01
    ) +
    geom_point() +
```

```
    geom_line(
      data = data.frame("Fit" = c(-0.1, 0.1),
                        "slope" = c(-0.1, 0.1)),
      colour = "grey50"
    ) +
    xlab("Fitted Metabolic Slope\n(Fold Metabolism at\nThermoneutrality/°C)") +
    ylab("Metabolic Slope\n(Fold Metabolism at\nThermoneutrality/°C)") +
    theme_classic(),
  labels = c("A", "B")
)
```

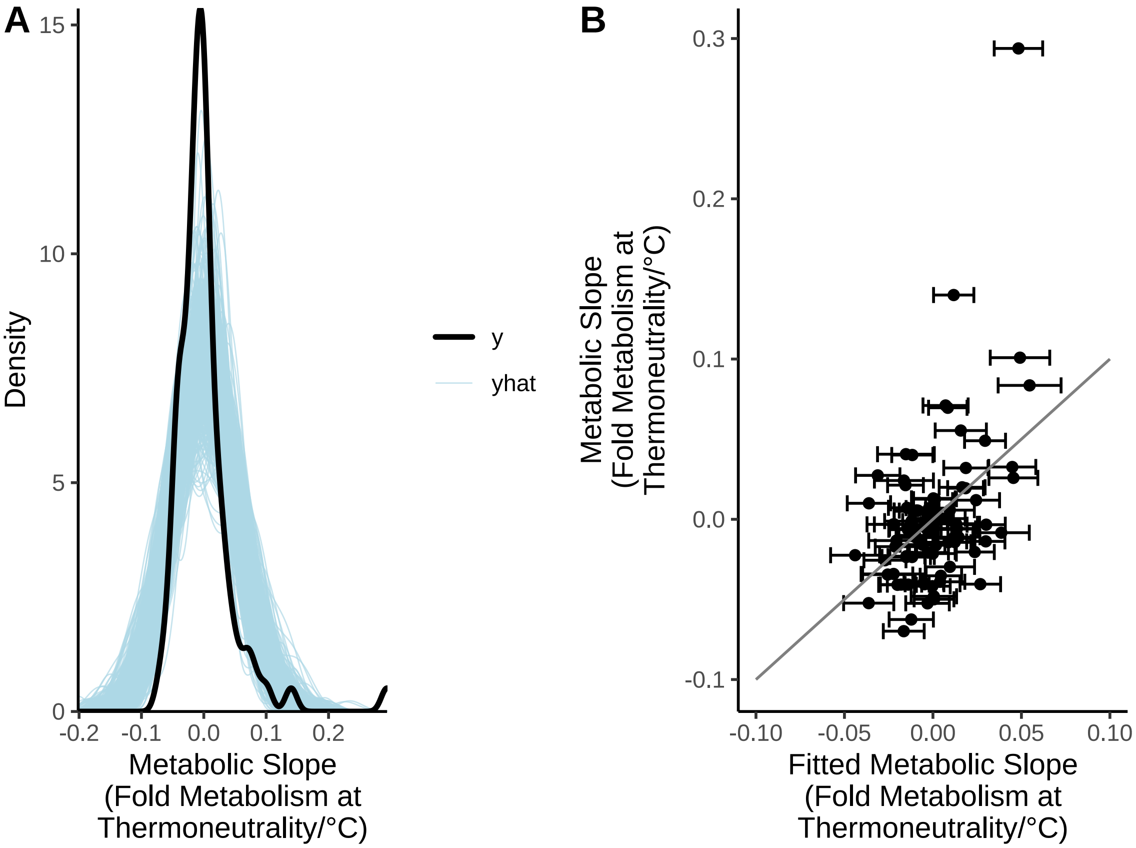


***Figure 65:*** *Characteristics of Bayesian path analysis predicting physiological responses to heat in three week old Japanese quail. Panels A and B display posterior predictive checks from path analysis with black lines indicating true densities of a given physiological variable and blue lines indicating densities estimated from posterior draws. Panels C and D disply model fitted values by true values with dots indicating individual data points, errorbars indicating standard errors around fitted values, and grey lines indicating 1:1 relationships.*

Model residuals are next visualised.

```
p1 <- functionModel8WeeksHot$data %>%
  mutate("Residuals" = residuals(functionModel8WeeksHot,
    resp = "slope",
    robust = TRUE
  )[, "Estimate"]) %>%
  ggplot(aes(x = Residuals)) +
  geom_density(colour = "black", fill = "grey50", alpha = 0.5) +
  xlab("Metabolic Slope\n(Fold Metabolism at\nThermoneutrality/°C)") +
  ylab("Density") +
  theme_classic() +
  theme(axis.title = element_text(size = 8))
```

```r
p2 <- functionModel8WeeksHot$data %>%
  mutate(
    "Residuals" = residuals(functionModel8WeeksHot,
      resp = "slope",
      robust = TRUE
    )[, "Estimate"],
    "SE" = residuals(functionModel8WeeksHot,
      resp = "slope",
      robust = TRUE
    )[, "Est.Error"]
  ) %>%
  ggplot(aes(x = mass, y = Residuals)) +
  geom_errorbar(aes(
    x = mass, ymin = Residuals - SE,
    ymax = Residuals + SE
  ), width = 1) +
  geom_point(
    pch = 21, colour = "black", size = 2,
    fill = "lightblue", alpha = 0.5
  ) +
  xlab("Body Mass\n(g; Mean-Centred)") +
  ylab(
    "Metabolic Slope Residuals\n(Fold Metabolism at\nThermoneutrality/C°)"
  ) +
  theme_classic() +
  theme(axis.title = element_text(size = 8))

p3 <- functionModel8WeeksHot$data %>%
  mutate(
    "Residuals" = residuals(functionModel8WeeksHot,
      resp = "slope",
      robust = TRUE
    )[, "Estimate"],
    "SE" = residuals(functionModel8WeeksHot,
      resp = "slope",
      robust = TRUE
    )[, "Est.Error"]
  ) %>%
  ggplot(aes(x = tarsus, y = Residuals)) +
  geom_errorbar(
    aes(
      x = tarsus, ymin = Residuals - SE,
      ymax = Residuals + SE
    ),
    width = 0.1
  ) +
  geom_point(
    pch = 21, colour = "black", size = 2,
    fill = "lightblue", alpha = 0.5
  ) +
  xlab("Tarsus Length\n(mm; Mean-Centred)") +
  ylab(
    "Metabolic Slope\n(Fold Metabolism at\nThermoneutrality/C°)"
  ) +
  theme_classic() +
  theme(axis.title = element_text(size = 8))

p4 <- functionModel8WeeksHot$data %>%
  mutate("Residuals" = residuals(functionModel8WeeksHot,
    resp = "slope",
    robust = TRUE
  )[, "Estimate"]) %>%
  mutate(
    "pretreatment" =
      ifelse(pretreatment == "A", "Cold (10°C)",
        ifelse(pretreatment == "B", "Mild (20°C)",
          "Warm (30°C)"
```

```
        )
      )
  ) %>%
  ggplot(aes(x = pretreatment, y = Residuals)) +
  geom_point(
    pch = 21, colour = "black", size = 2,
    fill = "lightblue", alpha = 0.5
  ) +
  stat_summary(
    geom = "errorbar", fun.data = "mean_se",
    colour = "black", width = 0.25
  ) +
  stat_summary(
    geom = "point", fun = "mean", size = 4,
    pch = 21, colour = "black", fill = "lightblue"
  ) +
  xlab("Rearing Conditions") +
  ylab(
    "Metabolic Slope\n(Fold Metabolism at\nThermoneutrality/C°)"
  ) +
  theme_classic() +
  theme(axis.title = element_text(size = 8))

p5 <- functionModel8WeeksHot$data %>%
  mutate(
    "Residuals" = residuals(functionModel8WeeksHot,
      resp = "slope",
      robust = TRUE
    )[, "Estimate"],
    "rSE" = residuals(functionModel8WeeksHot,
      resp = "slope",
      robust = TRUE
    )[, "Est.Error"],
    "Fit" = fitted(functionModel8WeeksHot,
      resp = "slope",
      robust = TRUE
    )[, "Estimate"],
    "fSE" = fitted(functionModel8WeeksHot,
      resp = "slope",
      robust = TRUE
    )[, "Est.Error"]
  ) %>%
  ggplot(aes(x = Fit, y = Residuals)) +
  geom_errorbar(
    aes(
      x = Fit, ymin = Residuals - rSE,
      ymax = Residuals + rSE
    ),
    colour = "black", width = 0.001
  ) +
  geom_errorbarh(
    aes(
      y = Residuals, xmin = Fit - fSE,
      xmax = Fit + fSE
    ),
    colour = "black", height = 0.01
  ) +
  geom_point(
    pch = 21, colour = "black", size = 2,
    fill = "lightblue", alpha = 0.5
  ) +
  xlab(
    "Fitted Metabolic Slope\n(Fold Metabolism at\nThermoneutrality/C°)"
  ) +
  ylab(
    "Metabolic Slope Residuals\n(Fold Metabolism at\nThermoneutrality/C°)"
  ) +
```

```
  theme_classic() +
  theme(axis.title = element_text(size = 8))

((p1 + p2)/(p3 + p4)/p5) +
  plot_annotation(tag_levels = "A")
```

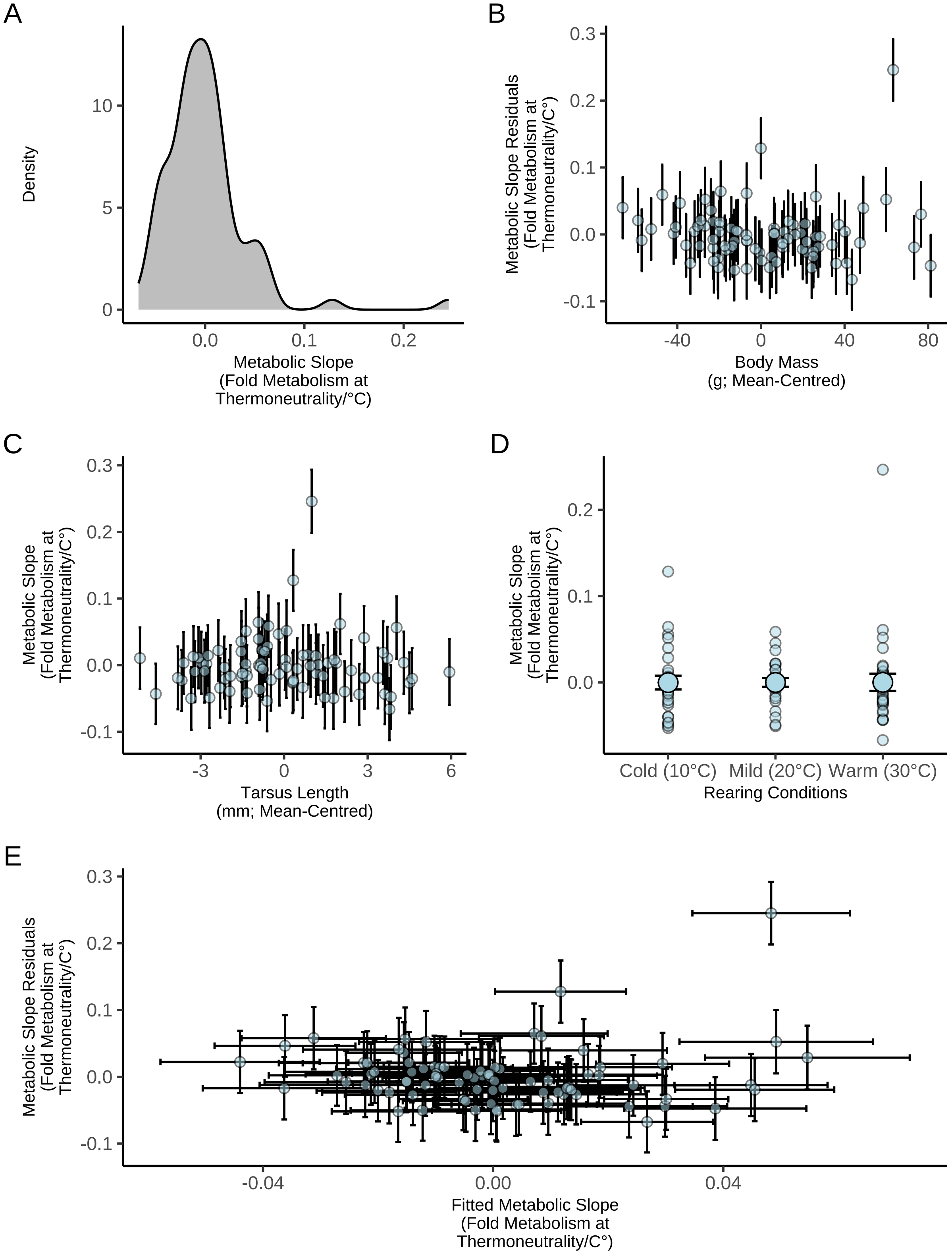


***Figure 66:*** *Spread of ordinary residuals from Bayesian path analysis predicting physiological responses to heat in eight week old Japanese quail. Quail were reared in one of three thermal conditions: (1) cold (10°C), (2) mild (20°C), or warm (30°C). Panel displays distribution of ordinary residuals for metabolic slopes (fold metabolism at thermoneutrality/°C). Panels B to E display ordinary residuals against predictors and fitted values; dots in these panels represent values per individual and errorbars represent +/- one standard error around values.*

Again, skewing in our median residuals is observed, as expected from our data-generating process (discussed above) and from our observation of a tentative outlying individual. We therefore visualise importance of individual data points to our analysis outcome as described for metabolic responses to heat in three-week old individuals.

```
loo(functionModel8WeeksHot)$diagnostics$pareto_k %>%
  as_tibble() %>%
  ggplot(aes(x = 1:nrow(.), y = value)) +
  geom_point(pch = 21, colour = "black", fill = "grey50", size = 2) +
  geom_hline(yintercept = 0.5, colour = "black", linetype = "dashed") +
  geom_hline(yintercept = 0.7, colour = "red4", linetype = "dashed") +
  xlab("Sample") +
  ylab("Pareto K") +
  theme_classic()
```

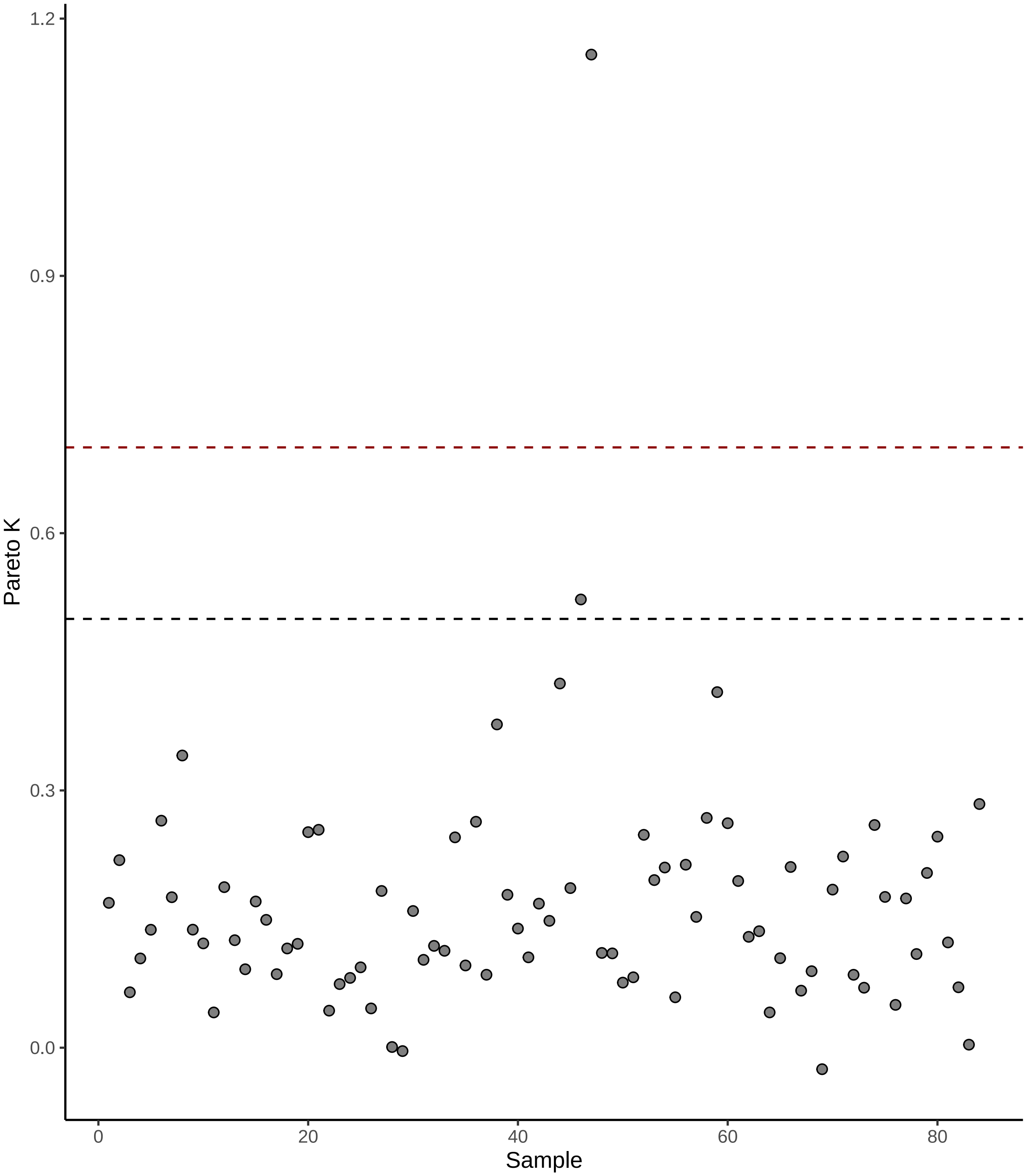


***Figure 67:*** *Relative importance of individual metabolic slope values (dots; derived from mature quail) to outcomes of our Bayesian path analysis, as estimated using leave-one-out cross validations. Importance is measured here as Pareto K (Vehtari et al, 2017). The black, dashed horizontal line indicated a Pareto K of 0.5 and the red horizontal dashed line indicates as Pareto K of 0.7, above which, samples hold significant importance on analysis outcomes.*

One sample clearly has dramatic influence on our model outcome (Pareto K $> 1$), and one moderately strong (Pareto K $> 0.5$). We have no *a priori* reason to exclude these samples. However, to balance their effects on our model estimates and acknowledge that our data-generating process is permissive of extreme values, our analysis is re-run but while assuming a Students-T distributed error around our metabolic slope values, rather than a Gaussian distributed error. Acknowledging the fact that our data is unlikely to contain numerous extreme values, we estimate the degrees of freedom ($\nu$) of our distribution from our data but while assuming that $\nu$ is probably high (prior on $\nu = \gamma(10, 1)$). Coefficients from this updated model are then plotted against those of our first model for reader interest.

```
functionModel8WeeksHotUpdated <-
  brm(
    data = slopeDataHot8 %>%
      mutate(
        mass = mass - mean(mass, na.rm = T),
        tarsus = tarsusLengthMean -
          mean(tarsusLengthMean, na.rm = T),
        pretreatment = ifelse(pretreatment == "cold", "A",
          ifelse(pretreatment == "neutral", "B", "C")
        ),
        slope = slope -
          mean(slope, na.rm = T)
      ) %>%
      mutate(pretreatment = factor(pretreatment,
        levels = c("B", "A", "C")
      )) %>%
      drop_na() %>%
      merge(., data %>%
        select(ring, "batch" = exp) %>%
        distinct(),
      by = "ring", all.x = TRUE
      ),
      bf(mass ~ pretreatment + (1 | batch),
         family = "gaussian") +
      bf(tarsus ~ mass + pretreatment + (1 | batch),
         family = "gaussian") +
      bf(slope ~ mass + tarsus + pretreatment + (1 | batch),
         family = student) +
      set_rescor(FALSE),
    prior = c(
      set_prior("normal(0, 10)",
        class = "Intercept",
        resp = "mass"
      ),
      set_prior("normal(0, 25)",
        class = "b",
        coef = "pretreatmentA",
        resp = "mass"
      ),
      set_prior("normal(0, 25)",
        class = "b",
        coef = "pretreatmentC",
        resp = "mass"
      ),
      set_prior("exponential(2.5)",
        class = "sd",
        group = "batch",
        resp = "mass"
      ),
      set_prior("exponential(0.05)",
        class = "sigma",
        resp = "mass"
      ),
      set_prior("normal(0, 3)",
        class = "Intercept",
        resp = "tarsus"
      ),
      set_prior("normal(0, 3)",
        class = "b",
```

```
        coef = "pretreatmentA",
        resp = "tarsus"
      ),
      set_prior("normal(0, 3)",
        class = "b",
        coef = "pretreatmentC",
        resp = "tarsus"
      ),
      set_prior("skew_normal(0, 0.25, 5)",
        class = "b",
        coef = "mass",
        resp = "tarsus"
      ),
      set_prior("exponential(2)",
        class = "sd",
        group = "batch",
        resp = "tarsus"
      ),
      set_prior("exponential(0.75)",
        class = "sigma",
        resp = "tarsus"
      ),
      set_prior("normal(0, 0.1)",
        class = "Intercept",
        resp = "slope"
      ),
      set_prior("normal(0, 0.1)",
        class = "b",
        coef = "pretreatmentA",
        resp = "slope"
      ),
      set_prior("normal(0, 0.1)",
        class = "b",
        coef = "pretreatmentC",
        resp = "slope"
      ),
      set_prior("normal(0, 0.0025)",
        class = "b",
        coef = "mass",
        resp = "slope"
      ),
      set_prior("normal(0, 0.03)",
        class = "b",
        coef = "tarsus",
        resp = "slope"
      ),
      set_prior("exponential(50)",
        class = "sd",
        group = "batch",
        resp = "slope"
      ),
      set_prior("exponential(10)",
        class = "sigma",
        resp = "slope"
      ),
      set_prior("gamma(10, 1)",
                class = "nu",
                resp = "slope")
    ),
    iter = 50000, warmup = 10000, cores = 4, chains = 4, thin = 20,
    control = list(adapt_delta = .97, max_treedepth = 14),
    silent = TRUE, refresh = 0,
    file = "./models/eightWeekFunctionModelHotST.Rds"
  )

bind_rows(
  as.data.frame(functionModel8WeeksHot) %>%
```

```
    select(
      "Intercept" = b_slope_Intercept,
      "Tarsus Length\n(mm; Mean-Centred)" =
        b_slope_tarsus,
      "Body Mass\n(g; Mean-Centred)" =
        b_slope_mass,
      "Cold Rearing\n(10°C)" =
        b_slope_pretreatmentA,
      "Warm Rearing\n(30°C)" =
        b_slope_pretreatmentC
    ) %>%
    pivot_longer(everything(),
      names_to = "Par",
      values_to = "Coefs"
    ) %>%
    mutate("Error" = "Gaussian"),
  as.data.frame(functionModel8WeeksHotUpdated) %>%
    select(
      "Intercept" = b_slope_Intercept,
      "Tarsus Length\n(mm; Mean-Centred)" =
        b_slope_tarsus,
      "Body Mass\n(g; Mean-Centred)" =
        b_slope_mass,
      "Cold Rearing\n(10°C)" =
        b_slope_pretreatmentA,
      "Warm Rearing\n(30°C)" =
        b_slope_pretreatmentC
    ) %>%
    pivot_longer(everything(),
      names_to = "Par",
      values_to = "Coefs"
    ) %>%
    mutate("Error" = "Student's T")
) %>% ggplot(aes(x = Coefs, fill = Error)) +
  facet_wrap(~Par, scales = "free", ncol = 2) +
  geom_density(colour = "black", alpha = 0.5) +
  geom_vline(
    xintercept = 0, linetype = "dashed",
    colour = "black"
  ) +
  scale_x_continuous(n.breaks = 4) +
  scale_fill_manual(values = c("grey80", "grey20")) +
  ylab("Density") +
  theme_classic() +
  theme(axis.title.x = element_blank()) +
  ggtitle(
    "Metabolic Slope\n(Fold Metabolism at Thermoneutrality/C°)"
  )
```

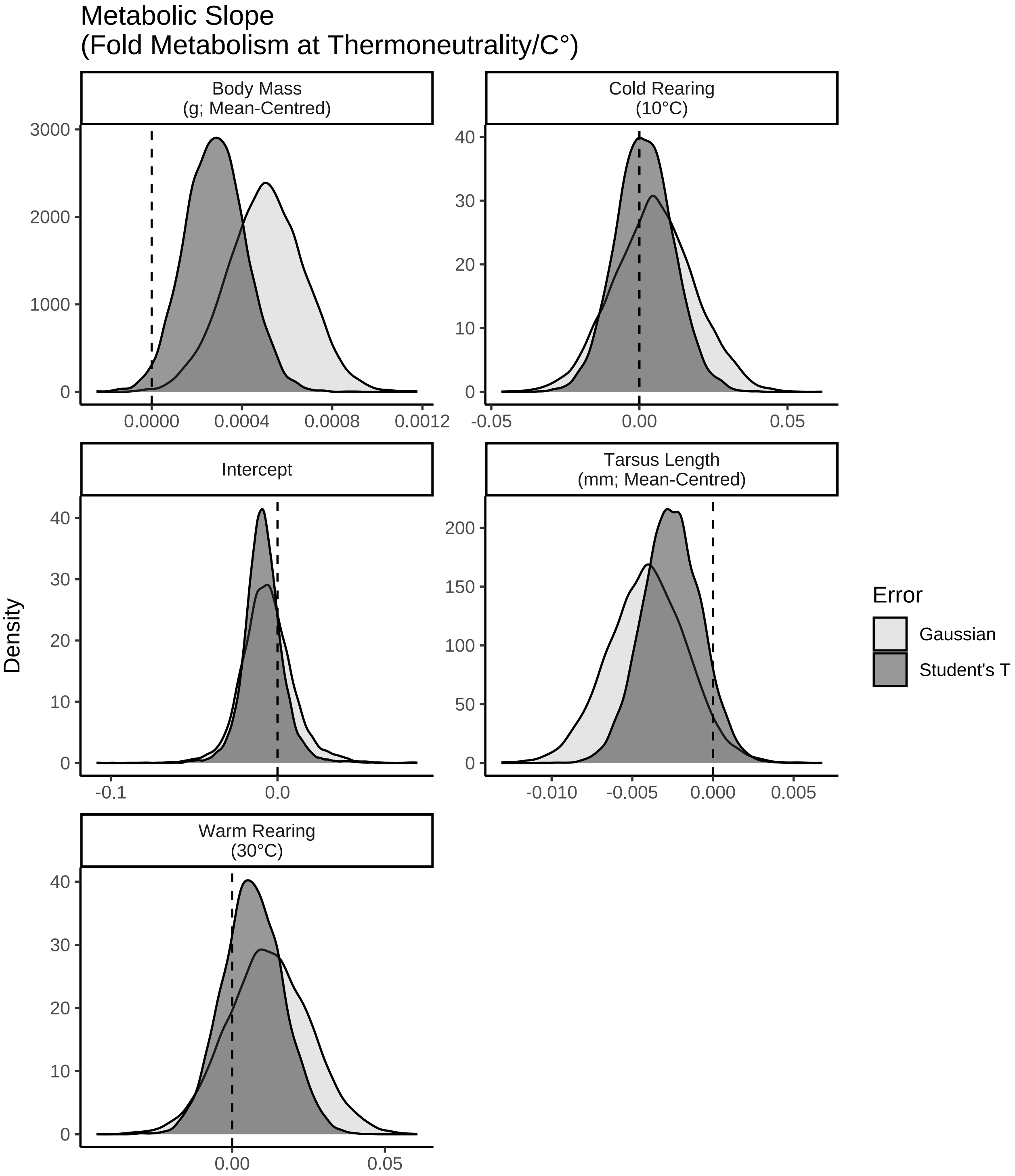


***Figure 68:*** *Posterior densities of coefficients from two Bayesian path analysis predicting metabolic responses to heating (30°C to 40°C) in eight week old Japanese quail. Quail were reared in one of three thermal conditions: (1) cold (10°C), (2) mild (20°C), or warm (30°C). Light grey densities represent those derived from posteriors of a model assuming Gaussian distributed error around metabolic responses, while dark grey densities represent those derived from a model assuming a Student's T distributed error.*

Last, we re-check the influence of our individual datum points on our model estimates.

```
loo(functionModel8WeeksHotUpdated)$diagnostics$pareto_k %>%
  as_tibble() %>%
  ggplot(aes(x = 1:nrow(.), y = value)) +
  geom_point(pch = 21, colour = "black", fill = "grey50", size = 2) +
  geom_hline(yintercept = 0.5, colour = "black", linetype = "dashed") +
  geom_hline(yintercept = 0.7, colour = "red4", linetype = "dashed") +
  xlab("Sample") +
  ylab("Pareto K") +
  theme_classic()
```

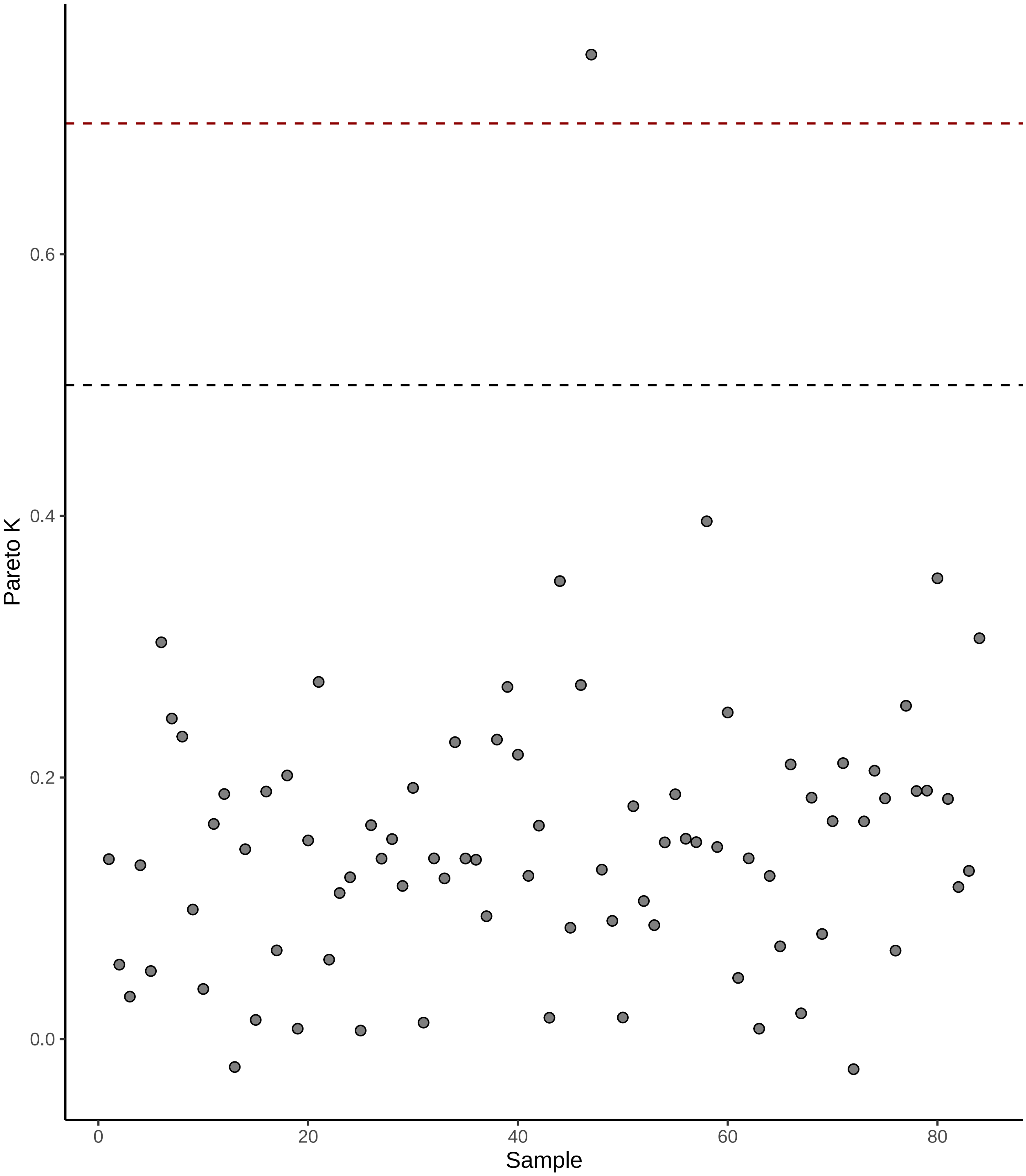


***Figure 69:*** *Relative importance of individual metabolic slope values (dots; derived from mature quail) to outcomes of our Bayesian path analysis, as estimated using leave-one-out cross validations. Importance is measured here as Pareto K (Vehtari et al, 2017). The black, dashed horizontal line indicated a Pareto K of 0.5 and the red horizontal dashed line indicates as Pareto K of 0.7, above which, samples hold significant importance on analysis outcomes.*

Clearly, the influence of our extreme values is lessened and balanced with those of our other data points. Thus, we proceed to visualising and evaluating predictions drawn from our updated model. We begin by plotting posterior densities for our model coefficients. In the process, variance explained by morphology alone on metabolic responses to heat (as partial $R^2$ values) is calculated.

```
functionModel8WeeksHot <- functionModel8WeeksHotUpdated

caption <- paste0("R\\textsuperscript{2} for a Bayesian ",
                  "path analysis predicting metabolic slopes ",
                  "(fold metabolism at thermoneutrality/°C) of ",
                  "eight week old Japanese quail as a function of ",
                  "body mass (g) and tarsus length (mm). Metabolic slopes ",
                  "are measured above thermoneutrality (30°C - 40°C)."
)

brms::bayes_R2(functionModel8WeeksHot,
               ndraws = 1000,
               robust = TRUE) %>%
  as.data.frame() %>%
  rownames_to_column("var") %>%
  merge(., tribble(
    ~var, ~Var,
    "R2mass", "Body Mass (g)",
    "R2tarsus", "Tarsus Length (mm)",
    "R2slope",
      "Metabolic Slope"
  ), by = c("var")) %>%
  mutate(Estimate = round(Estimate, digits = 4),
         Est.Error = round(Est.Error, digits = 4),
         `95\\% CI` = paste0("[",
                             round(Q2.5, digits = 4),
                             ", ",
                             round(Q97.5, digits = 4),
                             "]"
         )
         ) %>%
  select(
    "Response" = Var, "R\\textsuperscript{2}" = Estimate,
    "Standard Error" = Est.Error, `95\\% CI`
  ) %>%
  kbl(.,
    longtable = T, booktabs = T, format = "latex",
    caption = caption, escape = FALSE
  ) %>%
  column_spec(column = c(1:10), width = "2.5cm")  %>%
  kable_styling(latex_options = "striped")
```

***Table 32:*** *$R^2$ for a Bayesian path analysis predicting metabolic slopes (fold metabolism at thermoneutrality/°C) of eight week old Japanese quail as a function of body mass (g) and tarsus length (mm). Metabolic slopes are measured above thermoneutrality (30°C - 40°C).*

| Response | $R^2$ | Standard Error | 95% CI |
|---|---|---|---|
| Body Mass (g) | 0.0397 | 0.0320 | [0.0029, 0.1335] |
| Metabolic Slope | 0.0939 | 0.0472 | [0.0278, 0.201] |
| Tarsus Length (mm) | 0.2982 | 0.0703 | [0.1513, 0.4183] |

```
as.data.frame(functionModel8WeeksHot) %>%
    select(
      "Intercept" = b_slope_Intercept,
      "Tarsus Length\n(mm; Mean-Centred)" =
        b_slope_tarsus,
      "Body Mass\n(g; Mean-Centred)" =
        b_slope_mass,
      "Cold Rearing\n(10°C)" =
        b_slope_pretreatmentA,
```

```
      "Warm Rearing\n(30°C)" =
        b_slope_pretreatmentC,
      "Sigma" = sigma_slope
    ) %>%
    pivot_longer(everything(), names_to = "Par",
                 values_to = "Coefs") %>%
    ggplot(aes(x = Coefs)) +
    facet_wrap(~Par, scales = "free", ncol = 2) +
    geom_density(colour = "black", alpha = 0.5, fill = "grey70") +
    geom_vline(xintercept = 0, linetype = "dashed",
               colour = "black") +
    scale_x_continuous(n.breaks = 4) +
    ylab("Density") +
    theme_classic() +
    theme(axis.title.x = element_blank()) +
    ggtitle(
     "Metabolic Slope\n(Fold Metabolism at Thermoneutrality/C°)"
     )
```

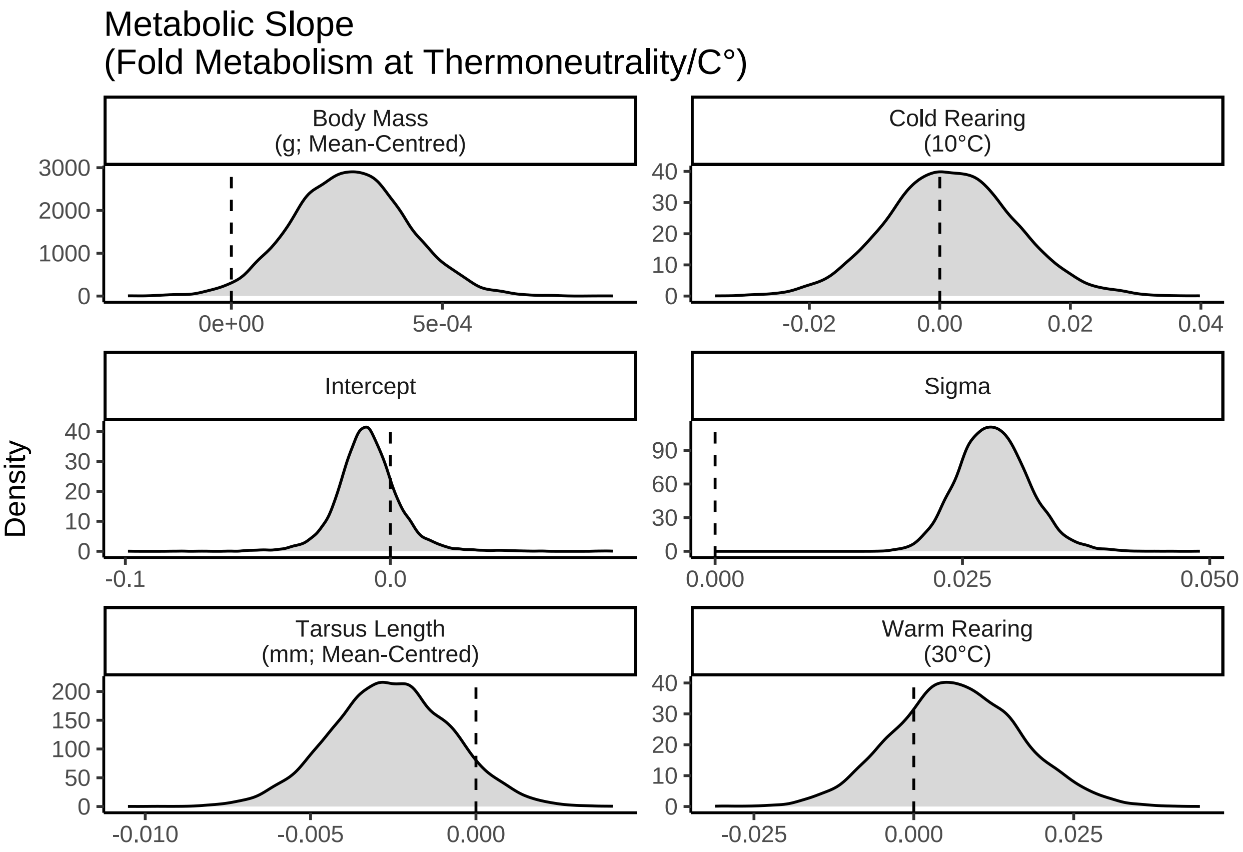


***Figure 70:*** *Posterior densities of coefficients for a Bayesian path analysis predicting physiological responses to heat in eight week old Japanese quail. Quail were reared in one of three thermal conditions: (1) cold (10°C), (2) mild (20°C), or warm (30°C).*

```
caption <- paste0("Results from a Bayesian path analysis ",
                  "predicting metabolic responses to heat (thermal ",
                  "resistace [fold metabolism at thermoneutrality",
                  "/°C] as a ",
                  "function of morphometry in eight week old ",
                  "Japanese quail. Physiological measurements are ",
                  "made at ambient temperatures between ",
                  "30°C and ",
```

```
                  "40°C. Cold rearing indicates ",
                  "post-hatch rearing at 10°C, ",
                  "relative to 20°C (intercept), ",
                  "or 30°C ('warm rearing'). Estimates ",
                  "indicate posterior medians, credible intervals ('CIs') ",
                  "indicate quantile intervals and BF indicates Bayes Factors."
)

week8ResultsHeat <-
  as.data.frame(functionModel8WeeksHot) %>%
  summarise_all(., .funs = median) %>%
  pivot_longer(everything(),
    names_to = "Parameter",
    values_to = "Estimate"
  ) %>%
  merge(., quantileCIs(functionModel8WeeksHot, cis = c(50, 95)),
    by = "Parameter", all.x = TRUE
  ) %>%
  filter(grepl("b_|sd_", Parameter)) %>%
  rowwise() %>%
  mutate("BF" = ifelse(Estimate < 0,
    (2 * mean(as.data.frame(
      functionModel8WeeksHot
    )[, Parameter] <= 0)) /
      (2 * mean(as.data.frame(
        functionModel8WeeksHot
      )[, Parameter] >= 0)),
    (2 * mean(as.data.frame(
      functionModel8WeeksHot
    )[, Parameter] >= 0)) /
      (2 * mean(as.data.frame(
        functionModel8WeeksHot
      )[, Parameter] <= 0))
  )) %>%
  ungroup() %>%
  mutate(
    "Estimate" = round(Estimate, digits = 4),
    "BF" = round(BF, digits = 4),
    "N" = nrow(functionModel8WeeksHot$data)
  ) %>%
  mutate("Parameter" = ifelse(grepl("b_", Parameter),
    gsub("b_|sd_", "", Parameter),
    gsub(
      "Intercept", "batch",
      gsub(".*__", "", Parameter)
    )
  )) %>%
  mutate(
    "Response" = gsub("_.*", "", Parameter),
    "Parameter" = gsub(".*_", "", Parameter)
  ) %>%
  merge(., tribble(
    ~Response, ~response, ~level,
    "mass", "Body Mass (g)", "A",
    "slope",
    "Metabolic Slope", "D",
    "tarsus", "Tarsus Length (mm)", "B"
  ),
  by = "Response"
  ) %>%
  merge(., tribble(
    ~Parameter, ~parameter, ~number,
    "Intercept", "Intercept", "1",
    "mass", "Body Mass (g)", "4",
    "tarsus", "Tarsus Length (mm)", "5",
    "pretreatmentA", "Cold Rearing", "2",
    "pretreatmentC", "Warm Rearing", "3",
```

```
    "batch", "Egg Batch [mu]", "6"
  ),
  by = "Parameter"
  ) %>%
  mutate(
    `50\\% HDI` = paste0("(", paste(
      round(Low_CI_50, digits = 4),
      round(High_CI_50, digits = 4),
      sep = ", "
    ), ")"),
    `95\\% HDI` = paste0("(", paste(
      round(Low_CI_95, digits = 4),
      round(High_CI_95, digits = 4),
      sep = ", "
    ), ")")
  ) %>%
  select(-c(Low_CI_50, High_CI_50, Low_CI_95, High_CI_95)) %>%
  select(
    "Response" = "response", "Parameter" = "parameter", N,
    Estimate, `50\\% HDI`, `95\\% HDI`, BF, level, number
  ) %>%
  arrange(level, number) %>%
  select(-c(level, number)) %>%
  kbl(.,
    longtable = T, booktabs = T, format = "latex", escape = FALSE,
    caption = caption
  ) %>%
  column_spec(column = c(1:2), width = "2.2cm") %>%
  column_spec(column = c(3:10), width = "1.9cm") %>%
  kable_styling(latex_options = "striped")

week8ResultsHeat
```

***Table 33:*** *Results from a Bayesian path analysis predicting metabolic responses to heat (thermal resistace [fold metabolism at thermoneutrality/°C] as a function of morphometry in eight week old Japanese quail. Physiological measurements are made at ambient temperatures between 30°C and 40°C. Cold rearing indicates post-hatch rearing at 10°C, relative to 20°C (intercept), or 30°C ('warm rearing'). Estimates indicate posterior medians, credible intervals ('CIs') indicate quantile intervals and BF indicates Bayes Factors.*

| Response | Parameter | N | Estimate | 50% HDI | 95% HDI | BF |
|---|---|---|---|---|---|---|
| Body Mass (g) | Intercept | 84 | -5.7954 | (-9.5272, -2.0963) | (-16.8274, 5.0508) | 5.6390 |
| Body Mass (g) | Cold Rearing | 84 | 5.9285 | (0.6606, 11.4371) | (-10.0298, 21.9769) | 3.4029 |
| Body Mass (g) | Warm Rearing | 84 | 12.7116 | (7.5107, 17.9427) | (-2.4341, 28.0855) | 19.4082 |
| Body Mass (g) | Egg Batch [mu] | 84 | 0.2760 | (0.1142, 0.564) | (0.0099, 1.4847) | Inf |
| Tarsus Length (mm) | Intercept | 84 | -0.5579 | (-0.9356, -0.1767) | (-1.6872, 0.7977) | 4.9128 |
| Tarsus Length (mm) | Cold Rearing | 84 | 0.5116 | (0.0883, 0.9645) | (-0.7688, 1.7911) | 3.8077 |
| Tarsus Length (mm) | Warm Rearing | 84 | 0.8816 | (0.4651, 1.3084) | (-0.3661, 2.1074) | 11.5000 |
| Tarsus Length (mm) | Body Mass (g) | 84 | 0.0277 | (0.0229, 0.0329) | (0.0135, 0.043) | 3999.0000 |
| Tarsus Length (mm) | Egg Batch [mu] | 84 | 0.6122 | (0.3697, 0.9029) | (0.0432, 1.7018) | Inf |
| Metabolic Slope | Intercept | 84 | -0.0089 | (-0.0154, -0.0021) | (-0.0311, 0.0165) | 4.2083 |
| Metabolic Slope | Cold Rearing | 84 | 0.0016 | (-0.0049, 0.0082) | (-0.0171, 0.0209) | 1.2890 |
| Metabolic Slope | Warm Rearing | 84 | 0.0066 | (1e-04, 0.0135) | (-0.0124, 0.0266) | 3.0650 |
| Metabolic Slope | Body Mass (g) | 84 | 0.0003 | (2e-04, 4e-04) | (0, 5e-04) | 59.6061 |

| | | | | | | |
|---|---|---|---|---|---|---|
| Metabolic Slope | Tarsus Length (mm) | 84 | -0.0026 | (-0.0038, -0.0013) | (-0.0061, 0.001) | 11.7186 |
| Metabolic Slope | Egg Batch [mu] | 84 | 0.0117 | (0.0066, 0.0186) | (8e-04, 0.0438) | Inf |

```
#save_kable(week8ResultsHeat,
# "../tables/pathAnalysisInHeat8Weeks.html")

functionModel8WeeksHotMassR2 <-
  brm(
    data = slopeDataHot8 %>%
      mutate(
        mass = mass - mean(mass, na.rm = T),
        tarsus = tarsusLengthMean -
          mean(tarsusLengthMean, na.rm = T),
        pretreatment = ifelse(pretreatment == "cold", "A",
          ifelse(pretreatment == "neutral", "B", "C")
        ),
        slope = slope -
          mean(slope, na.rm = T)
      ) %>%
      mutate(pretreatment = factor(pretreatment,
        levels = c("B", "A", "C")
      )) %>%
      drop_na() %>%
      merge(., data %>%
        select(ring, "batch" = exp) %>%
        distinct(),
      by = "ring", all.x = TRUE
      ),
      bf(mass ~ pretreatment + (1 | batch),
         family = "gaussian") +
      bf(tarsus ~ mass + pretreatment + (1 | batch),
         family = "gaussian") +
      bf(slope ~ tarsus + pretreatment + (1 | batch),
         family = student) +
      set_rescor(FALSE),
    prior = c(
      set_prior("normal(0, 10)",
        class = "Intercept",
        resp = "mass"
      ),
      set_prior("normal(0, 25)",
        class = "b",
        coef = "pretreatmentA",
        resp = "mass"
      ),
      set_prior("normal(0, 25)",
        class = "b",
        coef = "pretreatmentC",
        resp = "mass"
      ),
      set_prior("exponential(2.5)",
        class = "sd",
        group = "batch",
        resp = "mass"
      ),
      set_prior("exponential(0.05)",
        class = "sigma",
        resp = "mass"
      ),
      set_prior("normal(0, 3)",
        class = "Intercept",
        resp = "tarsus"
      ),
      set_prior("normal(0, 3)",
        class = "b",
```

```
        coef = "pretreatmentA",
        resp = "tarsus"
      ),
      set_prior("normal(0, 3)",
        class = "b",
        coef = "pretreatmentC",
        resp = "tarsus"
      ),
      set_prior("skew_normal(0, 0.25, 5)",
        class = "b",
        coef = "mass",
        resp = "tarsus"
      ),
      set_prior("exponential(2)",
        class = "sd",
        group = "batch",
        resp = "tarsus"
      ),
      set_prior("exponential(0.75)",
        class = "sigma",
        resp = "tarsus"
      ),
      set_prior("normal(0, 0.1)",
        class = "Intercept",
        resp = "slope"
      ),
      set_prior("normal(0, 0.1)",
        class = "b",
        coef = "pretreatmentA",
        resp = "slope"
      ),
      set_prior("normal(0, 0.1)",
        class = "b",
        coef = "pretreatmentC",
        resp = "slope"
      ),
      set_prior("normal(0, 0.03)",
        class = "b",
        coef = "tarsus",
        resp = "slope"
      ),
      set_prior("exponential(50)",
        class = "sd",
        group = "batch",
        resp = "slope"
      ),
      set_prior("exponential(10)",
        class = "sigma",
        resp = "slope"
      ),
      set_prior("gamma(10, 1)",
                class = "nu",
                resp = "slope")
    ),
    iter = 50000, warmup = 10000, cores = 4, chains = 4, thin = 20,
    control = list(adapt_delta = .97, max_treedepth = 14),
    silent = TRUE, refresh = 0,
    file = "./models/eightWeekFunctionModelHotMassR2Student.Rds"
  )


functionModel8WeeksHotTarsusR2 <-
  brm(
    data = slopeDataHot8 %>%
      mutate(
        mass = mass - mean(mass, na.rm = T),
        tarsus = tarsusLengthMean -
```

```
        mean(tarsusLengthMean, na.rm = T),
      pretreatment = ifelse(pretreatment == "cold", "A",
        ifelse(pretreatment == "neutral", "B", "C")
      ),
      slope = slope -
        mean(slope, na.rm = T)
    ) %>%
    mutate(pretreatment = factor(pretreatment,
      levels = c("B", "A", "C")
    )) %>%
    drop_na() %>%
    merge(., data %>%
      select(ring, "batch" = exp) %>%
      distinct(),
    by = "ring", all.x = TRUE
    ),
    bf(mass ~ pretreatment + (1 | batch),
       family = "gaussian") +
    bf(tarsus ~ mass + pretreatment + (1 | batch),
       family = "gaussian") +
    bf(slope ~ mass + pretreatment + (1 | batch),
       family = student) +
    set_rescor(FALSE),
  prior = c(
    set_prior("normal(0, 10)",
      class = "Intercept",
      resp = "mass"
    ),
    set_prior("normal(0, 25)",
      class = "b",
      coef = "pretreatmentA",
      resp = "mass"
    ),
    set_prior("normal(0, 25)",
      class = "b",
      coef = "pretreatmentC",
      resp = "mass"
    ),
    set_prior("exponential(2.5)",
      class = "sd",
      group = "batch",
      resp = "mass"
    ),
    set_prior("exponential(0.05)",
      class = "sigma",
      resp = "mass"
    ),
    set_prior("normal(0, 3)",
      class = "Intercept",
      resp = "tarsus"
    ),
    set_prior("normal(0, 3)",
      class = "b",
      coef = "pretreatmentA",
      resp = "tarsus"
    ),
    set_prior("normal(0, 3)",
      class = "b",
      coef = "pretreatmentC",
      resp = "tarsus"
    ),
    set_prior("skew_normal(0, 0.25, 5)",
      class = "b",
      coef = "mass",
      resp = "tarsus"
    ),
    set_prior("exponential(2)",
```

```
        class = "sd",
        group = "batch",
        resp = "tarsus"
      ),
      set_prior("exponential(0.75)",
        class = "sigma",
        resp = "tarsus"
      ),
      set_prior("normal(0, 0.1)",
        class = "Intercept",
        resp = "slope"
      ),
      set_prior("normal(0, 0.1)",
        class = "b",
        coef = "pretreatmentA",
        resp = "slope"
      ),
      set_prior("normal(0, 0.1)",
        class = "b",
        coef = "pretreatmentC",
        resp = "slope"
      ),
      set_prior("normal(0, 0.0025)",
        class = "b",
        coef = "mass",
        resp = "slope"
      ),
      set_prior("exponential(50)",
        class = "sd",
        group = "batch",
        resp = "slope"
      ),
      set_prior("exponential(10)",
        class = "sigma",
        resp = "slope"
      ),
      set_prior("gamma(10, 1)",
                class = "nu",
                resp = "slope")
    ),
    iter = 50000, warmup = 10000, cores = 4, chains = 4, thin = 20,
    control = list(adapt_delta = .97, max_treedepth = 14),
    silent = TRUE, refresh = 0,
    file = "./models/eightWeekFunctionModelHotTarsusR2Student.Rds"
  )

massPR2 <- brms::bayes_R2(functionModel8WeeksHot,
  ndraws = 1000,
  robust = TRUE, resp = "slope",
  summary = FALSE
) %>%
  as.data.frame() %>%
  select("fullR2" = R2slope) %>%
  cbind(
    .,
    brms::bayes_R2(functionModel8WeeksHotMassR2,
      ndraws = 1000,
      robust = TRUE, resp = "slope",
      summary = FALSE
    )
  ) %>%
  mutate("pR2" = round(fullR2 - R2slope, digits = 3)) %>%
  pull(pR2)

tarsusPR2 <- brms::bayes_R2(functionModel8WeeksHot,
  ndraws = 1000,
  robust = TRUE, resp = "slope",
```

```
  summary = FALSE
) %>%
  as.data.frame() %>%
  select("fullR2" = R2slope) %>%
  cbind(
    .,
    brms::bayes_R2(functionModel8WeeksHotTarsusR2,
      ndraws = 1000,
      robust = TRUE, resp = "slope",
      summary = FALSE
    )
  ) %>%
  mutate("pR2" = round(fullR2 - R2slope, digits = 3)) %>%
  pull(pR2)

caption = paste0('Variance in metabolic slope ',
  '(fold metabolism at thermoneutrality/°C) above thermnoeutrality ',
  'explained by morphometry in eight week old Japanese ',
  'quail.')

quantile(massPR2, probs = c(0.025, 0.975), type = 8) %>%
  t() %>%
  as.data.frame() %>%
  mutate(`2.5%` = ifelse(`2.5%` < 0, 0, `2.5%`)) %>%
  mutate("95\\% CI" = paste0(
    "[",
    round(`2.5%`, digits = 3),
    ",",
    round(`97.5%`, digits = 3),
    "]"
  )) %>%
  mutate(
    "massR2" =
      round(median(massPR2), digits = 3)
  ) %>%
  mutate(massR2 = ifelse(massR2 < 0, 0, massR2)) %>%
  select(
    "Mass Partial R\\textsuperscript{2}" = massR2,
    "95\\% CI"
  ) %>%
  cbind(
    .,
    quantile(tarsusPR2, probs = c(0.025, 0.975), type = 8) %>%
      t() %>%
      as.data.frame() %>%
      mutate(`2.5%` = ifelse(`2.5%` < 0, 0, `2.5%`)) %>%
      mutate("95\\% CI" = paste0(
        "[",
        round(`2.5%`, digits = 3),
        ",",
        round(`97.5%`, digits = 3),
        "]"
      )) %>%
      mutate("tarsusR2" = round(median(tarsusPR2), digits = 3)) %>%
      select(
        "Tarsus Partial R\\textsuperscript{2}" = tarsusR2,
        "95\\% CI"
      )
  ) %>%
  kbl(.,
    longtable = T, booktabs = T, format = "latex",
    caption = caption, escape = FALSE
  ) %>%
  column_spec(column = c(1:10), width = "2.5cm") %>%
  kable_styling(latex_options = "striped")
```

***Table 34:*** *Variance in metabolic slope (fold metabolism at thermoneutrality/°C) above thermnoeutrality explained by morphometry in eight week old Japanese quail.*

| Mass Partial $R^2$ | 95% CI | Tarsus Partial $R^2$ | 95% CI |
|---|---|---|---|
| 0.043 | [0,0.158] | 0.026 | [0,0.152] |

```
showtext.auto(enable = TRUE)

p1 <- data.frame(
  "pretreatment" = c("A", "B", "C"),
  "mass" = 0,
  "tarsus" = 0
) %>%
  mutate(
    "slope" = predict(functionModel8WeeksHot,
      newdata = .,
      resp = "slope",
      robust = TRUE,
      re_form = NA
    )[, "Estimate"],
    "SE" = predict(functionModel8WeeksHot,
      newdata = .,
      resp = "slope",
      robust = TRUE,
      re_form = NA
    )[, "Est.Error"]
  ) %>%
  mutate(slope = slope +
           mean(slopeDataHot8$slope,
                na.rm = T)) %>%
  ggplot(aes(x = pretreatment, y = slope)) +
  geom_errorbar(aes(x = pretreatment, ymin = slope - SE,
                ymax = slope + SE),
                colour = "black", width = 0.3) +
  geom_point(size = 5, pch = 21, colour = "black",
             aes(x = pretreatment, fill = factor(pretreatment))) +
  geom_line(linetype = "dashed", colour = "black") +
  geom_point(
    size = 2, pch = 21, colour = "black",
    data = slopeDataHot8 %>%
      mutate("pretreatment" =
               ifelse(pretreatment == "cold", "A",
        ifelse(pretreatment == "neutral", "B", "C")
      )),
    aes(x = pretreatment, y = slope,
        fill = factor(pretreatment)),
    position = position_jitter(width = 0.3)
  ) +
  scale_fill_manual(values = c("#7BB4E3", "black", "#CD5C5C")) +
  scale_x_discrete(breaks = c("A", "B", "C"),
                   labels = c("Cold\n(10°C)",
                              "Mild\n(20°C)",
                              "Warm\n(30°C)")) +
  theme_classic() +
  theme(legend.position = "none",
        axis.text = element_text(family = "Noto Sans"),
        axis.title = element_text(family = "Noto Sans")) +
  xlab("Rearing Conditions") +
  ylab("Metabolic Slope\n(Fold Metabolism at Thermoneutrality/°C)")

p2 <- data.frame(
  "pretreatment" = "B",
  "mass" = seq(min(functionModel8WeeksHot$data$mass, na.rm = T),
    max(functionModel8WeeksHot$data$mass, na.rm = T),
    by = 0.1
  ),
```

```
  "tarsus" = 0
) %>%
  mutate(
    "slope" = predict(functionModel8WeeksHot,
      newdata = .,
      resp = "slope",
      robust = TRUE,
      re_form = NA
    )[, "Estimate"],
    "SE" = predict(functionModel8WeeksHot,
      newdata = .,
      resp = "slope",
      robust = TRUE,
      re_form = NA
    )[, "Est.Error"]
  ) %>%
  mutate("Mass" = mass +
           mean(slopeDataHot8$mass, na.rm = T)) %>%
  mutate("slope" = slope +
           mean(slopeDataHot8$slope, na.rm = T)) %>%
  ggplot(aes(x = Mass, y = slope)) +
  geom_ribbon(aes(x = Mass, ymin = slope - SE,
                  ymax = slope + SE),
    fill = "grey50", alpha = 0.5
  ) +
  geom_smooth(method = "lm", linetype = "dashed",
              se = FALSE, colour = "black") +
  geom_point(
    size = 2.5, pch = 21, colour = "black", alpha = 0.5, fill = "grey50",
    data = slopeDataHot8,
    aes(x = mass, y = slope)
  ) +
  theme_classic() +
  theme(legend.position = "none",
        axis.text = element_text(family = "Noto Sans"),
        axis.title = element_text(family = "Noto Sans")) +
  xlab("Body Mass (g)") +
  ylab("Metabolic Slope\n(Fold Metabolism at\nThermoneutrality/°C)")

p3 <- data.frame(
  "pretreatment" = "B",
  "tarsus" = seq(min(functionModel8WeeksHot$data$tarsus, na.rm = T),
    max(functionModel8WeeksHot$data$tarsus, na.rm = T),
    by = 0.1
  ),
  "mass" = 0
) %>%
  mutate(
    "slope" = predict(functionModel8WeeksHot,
      newdata = .,
      resp = "slope",
      robust = TRUE,
      re_form = NA
    )[, "Estimate"],
    "SE" = predict(functionModel8WeeksHot,
      newdata = .,
      resp = "slope",
      robust = TRUE,
      re_form = NA
    )[, "Est.Error"]
  ) %>%
  mutate("Tarsus" = tarsus +
           mean(slopeDataHot8$tarsusLengthMean, na.rm = T)) %>%
  mutate("slope" = slope +
           mean(slopeDataHot8$slope, na.rm = T)) %>%
  ggplot(aes(x = Tarsus, y = slope)) +
  geom_ribbon(aes(x = Tarsus, ymin = slope - SE,
```

```
                  ymax = slope + SE),
    fill = "grey50", alpha = 0.5
  ) +
  geom_smooth(method = "lm", linetype = "dashed",
              se = FALSE, colour = "black") +
  geom_point(
    size = 2.5, pch = 21, colour = "black",
    alpha = 0.5, fill = "grey50",
    data = slopeDataHot8,
    aes(x = tarsusLengthMean, y = slope)
  ) +
  theme_classic() +
  theme(legend.position = "none",
        axis.text = element_text(family = "Noto Sans"),
        axis.title = element_text(family = "Noto Sans")) +
  xlab("Tarsus Length (mm)") +
  ylab("Metabolic Slope\n(Fold Metabolism at\nThermoneutrality/°C)")

massPred <- data.frame(
  "pretreatment" = "B",
  "mass" = c(
    mean(functionModel8WeeksHot$data$mass, na.rm = T) -
      sd(functionModel8WeeksHot$data$mass, na.rm = T),
    mean(functionModel8WeeksHot$data$mass, na.rm = T),
    mean(functionModel8WeeksHot$data$mass, na.rm = T) +
      sd(functionModel8WeeksHot$data$mass, na.rm = T)
  ),
  "tarsus" = 0,
  "batch" = "A"
) %>%
  predict(functionModel8WeeksHot,
    re_form = NA,
    resp = "slope", newdata = .,
    summary = TRUE, robust = TRUE
  ) %>%
  as.data.frame() %>%
  mutate("slope" = Estimate +
    mean(
      subset(
        slopeDataHot8,
        pretreatment == "neutral"
      )$slope,
      na.rm = TRUE
    )) %>%
  mutate("Size" = c("Small", "Average", "Large")) %>%
  select(Size, slope, "slopeSE" = Est.Error) %>%
  slice(rep(1:n(), each = 2)) %>%
  mutate("Ta" = rep(c(30, 40), 3)) %>%
  mutate(
    "foldRMR" = ifelse(Ta == 30, 1, 10 * slope),
    "LL" = ifelse(Ta == 30, 1, 10 * (slope - slopeSE)),
    "UL" = ifelse(Ta == 30, 1, 10 * (slope + slopeSE))
  ) %>%
  mutate(Size = factor(Size, levels = c("Small", "Average", "Large")))

massPlot <- massPred %>%
  filter(Ta == 40) %>%
  ggplot(aes(x = Size, y = foldRMR)) +
  geom_point(
    data = modData8 %>%
      select(ring, mass, Ta, VO2) %>%
      pivot_wider(id_cols = c("ring", "mass"),
                  names_from = "Ta",
                  values_from = "VO2") %>%
      mutate("foldRMR" = `40`/`30`,
             "Ta" = 40,
             "Size" = ifelse(mass < mean(mass, na.rm = T) -
```

```
                              sd(mass, na.rm = T),
                            "Small",
                            ifelse(mass > mean(mass, na.rm = T) +
                                     sd(mass, na.rm = T),
                                   "Large", "Average")
                            )
            ) %>%
      mutate(Size = factor(Size, levels = c("Small", "Average", "Large"))
            ),
    aes(x = Size, y = foldRMR, fill = Size),
    position = position_jitter(width = 0.25), alpha = 0.5,
    colour = "black", pch = 21
  ) +
  geom_hline(yintercept = 1, colour = "grey30", linetype = "dashed") +
  geom_errorbar(aes(ymin = LL, ymax = UL),
                colour = "grey80",
                width = 0.25
  ) +
  geom_errorbar(aes(ymin = LL, ymax = UL),
                colour = "black",
                width = 0.25
  ) +
  geom_point(aes(x = Size, y = foldRMR, fill = Size),
             size = 4, colour = "black", pch = 21) +
  ylim(c(0, 2)) +
  theme_classic() +
  ylab("Resting Metabolism\n(Fold Change from Thermoneutrality)") +
  scale_fill_manual(values = c("#B35050", "#EABCBC", "white"),
                    guide = NULL) +
  scale_x_discrete(labels = c("Mean - 1 SD", "Mean", "Mean + 1 SD"),
                   name = "Body Mass") +
  theme(
    axis.text = element_text(family = "Noto Sans"),
    axis.title = element_text(family = "Noto Sans"),
    legend.text = element_text(family = "Noto Sans", size = 8),
    legend.title = element_text(family = "Noto Sans", size = 8),
    legend.position = "right"
  )

ggsave("../plots/massPlotHot8Weeks.pdf", dpi = 800,
       width = 7, height = 7,
       massPlot)

rm(massPred)

tarsusPred <- data.frame(
  "pretreatment" = "B",
  "tarsus" = c(
    mean(functionModel8WeeksHot$data$tarsus, na.rm = T) -
      sd(functionModel8WeeksHot$data$tarsus, na.rm = T),
    mean(functionModel8WeeksHot$data$tarsus, na.rm = T),
    mean(functionModel8WeeksHot$data$tarsus, na.rm = T) +
      sd(functionModel8WeeksHot$data$tarsus, na.rm = T)
  ),
  "mass" = 0,
  "batch" = "A"
) %>%
  predict(functionModel8WeeksHot, re_form = NA,
    resp = "slope", newdata = .,
    summary = TRUE, robust = TRUE
  ) %>%
  as.data.frame() %>%
  mutate("slope" = Estimate +
           mean(
             subset(slopeDataHot8,
                    pretreatment == "neutral"
                    )$slope,
```

```
          na.rm = TRUE
        )
  ) %>%
  mutate("Size" = c("Small", "Average", "Large")) %>%
  select(Size, slope, "slopeSE" = Est.Error) %>%
  slice(rep(1:n(), each = 2)) %>%
  mutate("Ta" = rep(c(30, 40), 3)) %>%
  mutate("foldRMR" = ifelse(Ta == 30, 1, 10*slope),
         "LL" = ifelse(Ta == 30, 1, 10*(slope - slopeSE)),
         "UL" = ifelse(Ta == 30, 1, 10*(slope + slopeSE))
  ) %>%
  mutate(Size = factor(Size, levels = c("Small", "Average", "Large")))

tarsusPlot <- tarsusPred %>%
  filter(Ta == 40) %>%
  ggplot(aes(x = Size, y = foldRMR)) +
  geom_point(
    data = modData8 %>%
      select(ring, "tarsus" = tarsusLengthMean, Ta, VO2) %>%
      pivot_wider(id_cols = c("ring", "tarsus"),
                  names_from = "Ta",
                  values_from = "VO2") %>%
      mutate("foldRMR" = `40`/`30`,
             "Ta" = 40,
             "Size" = ifelse(tarsus < mean(tarsus, na.rm = T) -
                               sd(tarsus, na.rm = T),
                             "Small",
                             ifelse(tarsus > mean(tarsus, na.rm = T) +
                                      sd(tarsus, na.rm = T),
                                    "Large", "Average")
             )
      ) %>%
      drop_na() %>%
      mutate(Size = factor(Size, levels = c("Small", "Average", "Large"))
      ),
    aes(x = Size, y = foldRMR, fill = Size),
    position = position_jitter(width = 0.25), alpha = 0.5,
    pch = 21, colour = "black"
  ) +
  geom_hline(yintercept = 1, colour = "grey30", linetype = "dashed") +
  geom_errorbar(aes(ymin = LL, ymax = UL),
                colour = "grey80",
                width = 0.25
  ) +
  geom_errorbar(aes(ymin = LL, ymax = UL),
                colour = "black",
                width = 0.25
  ) +
  geom_point(aes(x = Size, y = foldRMR, fill = Size),
             size = 4, colour = "black", pch = 21) +
  theme_classic() +
  xlab("Tarsus Length") +
  scale_x_discrete(labels = c("Mean - 1 SD",
                              "Mean",
                              "Mean + 1 SD")) +
  ylim(c(0, 3)) +
  ylab("Resting Metabolism\n(Fold Change from Thermoneutrality)") +
  scale_fill_manual(values = c("#B35050", "#EABCBC", "white"),
                    guide = NULL) +
  theme(
    axis.text = element_text(family = "Noto Sans"),
    axis.title = element_text(family = "Noto Sans"),
    legend.text = element_text(family = "Noto Sans", size = 8),
    legend.title = element_text(family = "Noto Sans", size = 8),
    legend.position = "right"
  )
```

```
ggsave("../plots/tarsusPlotHot8Weeks.pdf", dpi = 800,
       width = 7, height = 7,
      tarsusPlot)

allMorphology <- (p2 + massPlot) / (p3 + tarsusPlot) +
  plot_annotation(tag_levels = "A")

allMorphology
```

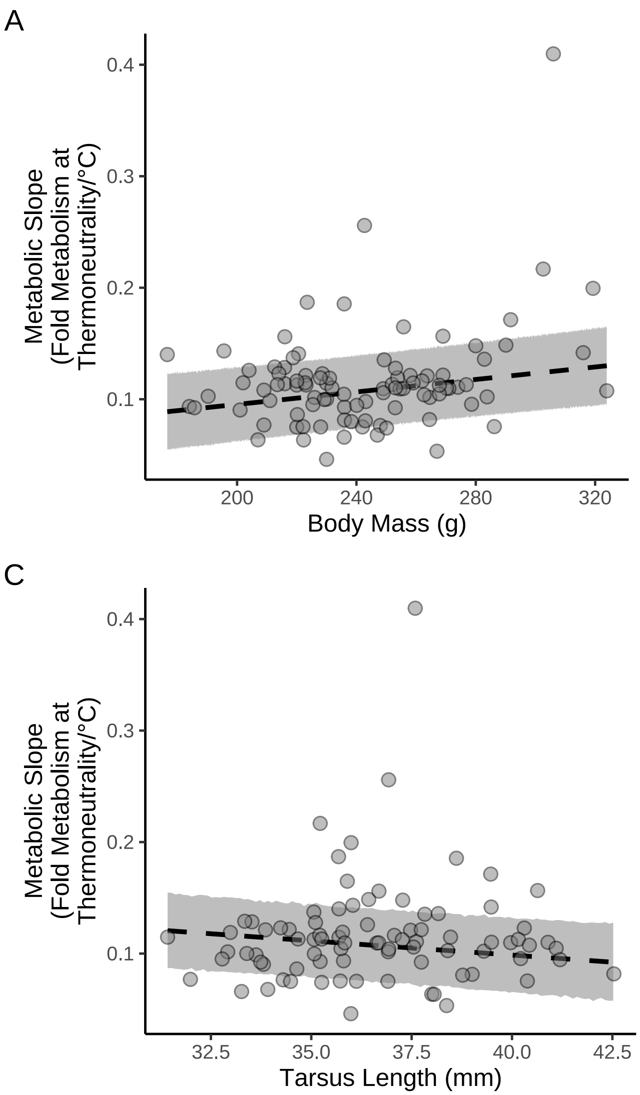


***Figure 71:*** *Conditional effects of post-hatch rearing environments (10°C, 20°C or 30°C) and morphometry (tarsus length [mm] and body mass [g]) on physiological responses to heat in eight week old Japanese quail. Dots represent raw data points per individual. Large dots in panel A and B represent estimated effects at average morphometric values and errorbars indicate +/- one standard error around means. Lines in panels C to F indicate estimate effects across given parameters and grouping (assuming mild [20°C] rearing conditions) and ribbons indicate standard errors around effects. All estimated effects are drawn from a Bayesian path analysis.*

```
ggsave("../plots/morphologyEffectsResistanceHeat8Weeks.pdf",
  allMorphology,
  dpi = 800, height = 7, width = 9
```

```
)

showtext.auto(enable = "FALSE")
```

Similar to above, we then estimate costs of mismatching morphology with expectations of Bergmann's and Allen's rules in the heat. Here, mismatches represent having a body mass two standard deviations above the mean or relative tarsus length two standard deviations below the mean.

```
data.frame(
  "Size" = c("Average", "Large (2x s.d. > mean)"),
  "pretreatment" = "B",
  "mass" = c(mean(functionModel8WeeksHot$data$mass, na.rm = T),
             mean(functionModel8WeeksHot$data$mass, na.rm = T) +
               2*sd(functionModel8WeeksHot$data$mass, na.rm = T)),
  "tarsus" = mean(functionModel8WeeksHot$data$tarsus, na.rm = T)
) %>%
  mutate("slope" = predict(functionModel8WeeksHot, newdata = .,
                              resp = "slope", re_form = NA,
                              robust = TRUE)[, "Estimate"]) %>%
  mutate(slope = slope +
         mean(slopeDataHot8$slope, na.rm = T)
  ) %>%
  select("Body Mass" = Size,
         "Metabolic Slope" = slope)
```

```
## # A tibble: 2 x 2
##   `Body Mass`            `Metabolic Slope`
##   <chr>                              <dbl>
## 1 Average                            0.108
## 2 Large (2x s.d. > mean)             0.125
```

```
data.frame(
  "Size" = c("Average", "Small (2x s.d. < mean)"),
  "pretreatment" = "B",
  "tarsus" = c(mean(functionModel8WeeksHot$data$tarsus, na.rm = T),
             mean(functionModel8WeeksHot$data$tarsus, na.rm = T) -
               2*sd(functionModel8WeeksHot$data$tarsus, na.rm = T)),
  "mass" = mean(functionModel8WeeksHot$data$mass, na.rm = T)
) %>%
  mutate("slope" = predict(functionModel8WeeksHot, newdata = .,
                              resp = "slope", re_form = NA,
                              robust = TRUE)[, "Estimate"]) %>%
  mutate(slope = slope +
         mean(slopeDataHot8$slope, na.rm = T)
  ) %>%
  select("Tarsus Length" = Size,
         "Metabolic Slope" = slope)
```

```
## # A tibble: 2 x 2
##   `Tarsus Length`        `Metabolic Slope`
##   <chr>                              <dbl>
## 1 Average                            0.108
## 2 Small (2x s.d. < mean)             0.120
```

```
# Conducting formal comparisons

mismatchDFMass <- data.frame(
  "Size" = c("Average", "Large (2x s.d. > mean)"),
  "pretreatment" = "B",
  "mass" = c(mean(functionModel8WeeksHot$data$mass, na.rm = T),
             mean(functionModel8WeeksHot$data$mass, na.rm = T) +
               2*sd(functionModel8WeeksHot$data$mass, na.rm = T)),
  "tarsus" = 0
) %>% predict(functionModel8WeeksHot, newdata = .,
              robust = TRUE, re_form = NA,
              resp = "slope", summary = FALSE) %>%
  as.data.frame() %>%
```

```
  rename("Average" = "V1", "Large" = "V2") %>%
  mutate_all(.funs = function(x)
    {x + mean(slopeDataHot8$slope, na.rm = T)}
    )

mismatchDFMassTest <- build_hdf(
  vars = list(
    mismatchDFMass$Average,
    mismatchDFMass$Large
    ),
  priors = list(
    rnorm(nrow(mismatchDFMass), 0, 0.01),
    rnorm(nrow(mismatchDFMass), 0, 0.01)
    ),
  names = c("Average", "Large")
)

mismatchTestMass <- hypothesis_df("Large > Average",
                                  mismatchDFMassTest,
                                  class = "b", alpha = 0.05
                                  )

mismatchTestMass$hypothesis$Hypothesis =
  "Large Metabolic Slope > Average Slope"

# Summarising

caption <- paste0(
  "Results of a non-linear hypothesis ",
  "tests comparing predicted metabolic slopes in the heat (30°C - 40°C) ",
  "among eight week old Japanese quail of average or atypically large ",
  "body size (mean + 2 standard deviations). Posterior probabilities are ",
  "calculated using the Savage-Dickey ",
  "density ratio method."
)

mismatchTestMassTable <- mismatchTestMass$hypothesis %>%
  mutate(Hypothesis = c(
    "Large Metabolic Slope > Average Slope"
  )) %>%
  select(-c(Evid.Ratio, Star)) %>%
  mutate(
    Estimate = round(Estimate, digits = 4),
    "Est.Error" = round(Est.Error, digits = 4),
    "CI.Lower" = round(CI.Lower, digits = 4),
    "CI.Upper" = round(CI.Upper, digits = 4)
  ) %>%
  rename(
    "Difference Between Metabolic Slopes" = Estimate,
    "Standard Error" = Est.Error,
    `2.5 % CI` = "CI.Lower",
    `97.5 % CI` = "CI.Upper",
    "Posterior Probability" = Post.Prob
  ) %>%
  kbl(.,
    longtable = T, booktabs = T, format = "latex",
    caption = caption
  ) %>%
  column_spec(column = c(1:2), width = "2.5cm") %>%
  column_spec(column = c(3:10), width = "2cm") %>%
  kable_styling(latex_options = "striped")

mismatchTestMassTable
```

***Table 35:*** *Results of a non-linear hypothesis tests comparing predicted metabolic slopes in the heat (30°C - 40°C) among eight week old Japanese quail of average or atypically large body size (mean + 2 standard deviations). Posterior probabilities are calculated using the Savage-Dickey density ratio method.*

| Hypothesis | Difference Between Metabolic Slopes | Standard Error | 2.5 % CI | 97.5 % CI | Posterior Probability |
|---|---|---|---|---|---|
| Large Metabolic Slope > Average Slope | 0.0183 | 0.0523 | -0.0646 | 0.103 | 0.657125 |

```
# Tarsus length comparison

mismatchDFTarsus <- data.frame(
  "Size" = c("Average", "Short (2x s.d. < mean)"),
  "pretreatment" = "B",
  "tarsus" = c(mean(functionModel8WeeksHot$data$tarsus, na.rm = T),
             mean(functionModel8WeeksHot$data$tarsus, na.rm = T) -
               2*sd(functionModel8WeeksHot$data$tarsus, na.rm = T)),
  "mass" = 0
) %>% predict(functionModel8WeeksHot, newdata = .,
              robust = TRUE, re_form = NA,
              resp = "slope", summary = FALSE) %>%
  as.data.frame() %>%
  rename("Average" = "V1", "Long" = "V2") %>%
  mutate_all(.funs = function(x)
    {x + mean(slopeDataHot8$slope, na.rm = T)}
    )

mismatchDFTarsusTest <- build_hdf(
  vars = list(
    mismatchDFTarsus$Average,
    mismatchDFTarsus$Long
    ),
  priors = list(
    rnorm(nrow(mismatchDFTarsus), 0, 0.01),
    rnorm(nrow(mismatchDFTarsus), 0, 0.01)
    ),
  names = c("Average", "Short")
)

mismatchTestTarsus <- hypothesis_df("Short > Average",
                                    mismatchDFTarsusTest,
                                    class = "b", alpha = 0.05
                                    )

mismatchTestTarsus$hypothesis$Hypothesis =
  "Short Tarsus Metabolic Slope > Average Slope"

caption <- paste0(
  "Results of a non-linear hypothesis ",
  "tests comparing predicted metabolic slopes in the cold (30°C - 10°C) ",
  "among eight week old Japanese quail of average or atypically long ",
  "tarsus length (mean + 2 standard deviations). Posterior probabilities are ",
  "calculated using the Savage-Dickey ",
  "density ratio method."
)

mismatchTestTarsusTable <- mismatchTestTarsus$hypothesis %>%
  mutate(Hypothesis = c(
    "Long Metabolic Slope > Average Slope"
  )) %>%
  select(-c(Evid.Ratio, Star)) %>%
  mutate(
    Estimate = round(Estimate, digits = 4),
    "Est.Error" = round(Est.Error, digits = 4),
```

```
    "CI.Lower" = round(CI.Lower, digits = 4),
    "CI.Upper" = round(CI.Upper, digits = 4)
  ) %>%
  rename(
    "Difference Between Metabolic Slopes" = Estimate,
    "Standard Error" = Est.Error,
    `2.5 % CI` = "CI.Lower",
    `97.5 % CI` = "CI.Upper",
    "Posterior Probability" = Post.Prob
  ) %>%
  kbl(.,
    longtable = T, booktabs = T, format = "latex",
    caption = caption
  ) %>%
  column_spec(column = c(1:2), width = "2.5cm") %>%
  column_spec(column = c(3:10), width = "2cm") %>%
  kable_styling(latex_options = "striped")

mismatchTestTarsusTable
```

***Table 36:*** *Results of a non-linear hypothesis tests comparing predicted metabolic slopes in the cold (30°C - 10°C) among eight week old Japanese quail of average or atypically long tarsus length (mean + 2 standard deviations). Posterior probabilities are calculated using the Savage-Dickey density ratio method.*

| Hypothesis | Difference Between Metabolic Slopes | Standard Error | 2.5 % CI | 97.5 % CI | Posterior Probability |
|---|---|---|---|---|---|
| Long Metabolic Slope > Average Slope | 0.0115 | 0.0514 | -0.0683 | 0.0936 | 0.600875 |

```
# Further, plotting estimated effects of body size enlargement, relative to
# average, on estimated rates of metabolic heat production

posteriorProbabilities <- bind_rows(
  lapply(X = seq(0, 5, by = 0.05),
       FUN = function(x){
         data <- data.frame(
          "tarsus" = 0,
          "pretreatment" = "B",
          "mass" = c(0, x * sd(
            functionModel8WeeksHot$data$mass,
            na.rm = T
          )),
          "batch" = "B"
        ) %>% predict(functionModel8WeeksHot, newdata = .,
              robust = TRUE, resp = "slope",
              summary = FALSE) %>%
           as.data.frame() %>%
           rename("Average" = "V1", "Large" = "V2") %>%
           mutate_all(.funs = function(x)
             {x + mean(slopeDataHot8$slope, na.rm = T)}
             )

         testDF <- build_hdf(
           vars = list(
             data$Average,
             data$Large
             ),
           priors = list(
             rnorm(nrow(data), 0, 0.01),
             rnorm(nrow(data), 0, 0.01)
             ),
           names = c("Average", "Large")
         )
```

```
        pp <- hypothesis_df("Large > Average", testDF,
                            class = "b", alpha = 0.5)$hypothesis$Post.Prob

        return(data.frame("sd" = x, "pp" = pp))
      }
  )
)

bind_rows(
    lapply(
      X = seq(0.1, 5, by = 0.1),
      FUN = function(x) {
        data <- data.frame(
          "tarsus" = 0,
          "pretreatment" = "B",
          "mass" = c(0, x * sd(
            functionModel8WeeksHot$data$mass,
            na.rm = T
          )),
          "batch" = "B"
        ) %>%
          predict(functionModel8WeeksHot,
            newdata = .,
            robust = TRUE, ndraws = 8000,
            resp = "slope", summary = FALSE
          ) %>%
          as.data.frame() %>%
          mutate_all(.funs = function(x){x + mean(slopeDataHot8$slope)}) %>%
          mutate("deltaRaw" = ((V2/V1)-1)*100) %>%
          summarise(
            "delta" = median(deltaRaw),
            "LL80" = quantile(deltaRaw, 0.1, type = 8),
            "UL80" = quantile(deltaRaw, 0.9, type = 8),
            "LL95" = quantile(deltaRaw, 0.025, type = 8),
            "UL95" = quantile(deltaRaw, 0.975, type = 8)
          ) %>%
          mutate("sd" = x) %>%
          select(sd, delta, LL80, UL80, LL95, UL95)
        return(data)
      }
    )
) %>%
  ggplot(aes(x = sd, y = delta)) +
  geom_ribbon(aes(ymin = LL95, ymax = UL95),
              fill = "#DECCC1", alpha = 0.5) +
  geom_ribbon(aes(ymin = LL80, ymax = UL80),
              fill = "#DECCC1", alpha = 0.8) +
  geom_smooth(method = "lm", colour = "black", linetype = "dashed",
              se = FALSE) +
  geom_line(data = posteriorProbabilities, colour = "black",
            linetype = "solid", aes(x = sd, y = pp*600)) +
  geom_hline(yintercept = 0, linetype = "dotted",
             colour = "grey50") +
  xlab("Standard Deviations\nAbove Mean Mass") +
  scale_y_continuous(name = "Increase in\nMetabolic Slopes (%)",
                     sec.axis = sec_axis(trans=~./6,
                                         name="Posterior Probability (%)")
                     ) +
  theme_classic()
```

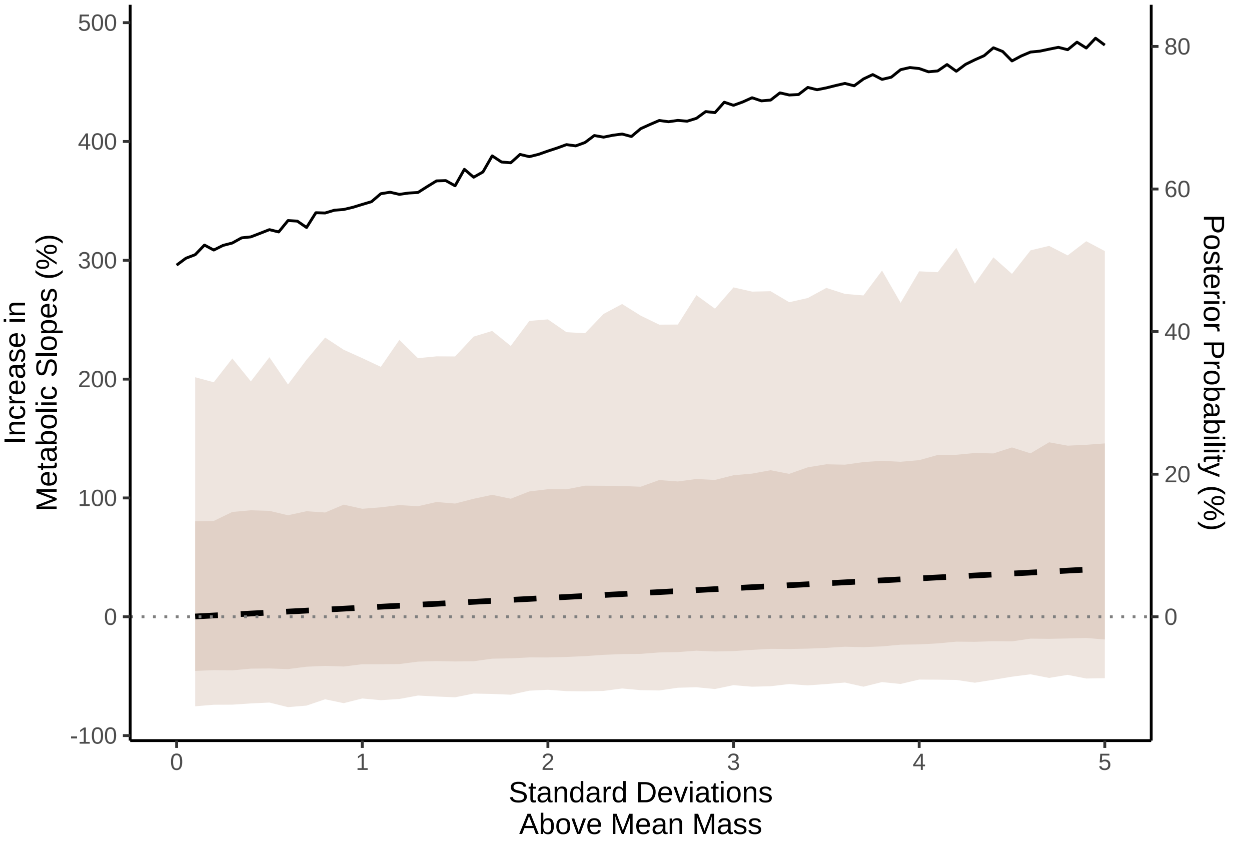


***Figure 72:*** *Effect of increasing mass (in standard deviations) above the average on the percent change of metabolic slope above 40°C in mature Japanese quail. Effects are estimated from a Bayesian path analysis. The dark ribbon represents 50% credible intervals around the estimated change in metabolic slope and the light ribbon represents 95% credible intervals around the predicted change. The dashed line represents the predicted relationship between the change in metabolic slope and relative increase in body mass above the average. The solid line indicate the posterior probability of an increase in metabolic slope among individuals of a given size above average. All predictions assume rearing in mild conditions (20°C) and average tarsus length. The horizontal dotted line indicates 0.*

```
# Last, it's important to not that having a large body size and average sized
# appendages is unlikely. Below, we calculate the odds that body mass is two
# standard deviations above the mean, but tarsus length is less than one quarter
# of a standard deviation above the mean

hyp <- paste0(
  "tarsus_mass*",
  2 * sd(functionModel8WeeksHot$data$mass),
  " < ",
  0.25 * sd(functionModel8WeeksHot$data$tarsus)
)
hypothesis(hypothesis = hyp, functionModel8WeeksHot)$hypothesis$Post.Prob
```

```
## [1] 0.007875
```

```
# Probability is low. As such, we calculate the assumed costs of mismatching
# body size when allometry is accounted for

data.frame(
  "Size" = c("Average", "Large (2x s.d. > mean)"),
  "pretreatment" = "B",
  "mass" = c(mean(functionModel8WeeksHot$data$mass, na.rm = T),
             mean(functionModel8WeeksHot$data$mass, na.rm = T) +
```

```
               2*sd(functionModel8WeeksHot$data$mass, na.rm = T))
) %>%
  mutate("tarsus" = predict(functionModel8WeeksHot, newdata = .,
                            resp = "tarsus", re_form = NA,
                            robust = TRUE)[, "Estimate"]) %>%
  mutate("slope" = predict(functionModel8WeeksHot, newdata = .,
                           resp = "slope", re_form = NA,
                           robust = TRUE)[, "Estimate"]) %>%
  mutate(slope = slope +
         mean(slopeDataHot8$slope, na.rm = T)
  ) %>%
  select("Body Mass" = Size,
         "Metabolic Slope" = slope)
```

```
## # A tibble: 2 x 2
##   `Body Mass`            `Metabolic Slope`
##   <chr>                              <dbl>
## 1 Average                            0.109
## 2 Large (2x s.d. > mean)             0.121
```

```
mismatchDFMassAllometric <- data.frame(
  "Size" = c("Average", "Large (2x s.d. > mean)"),
  "pretreatment" = "B",
  "mass" = c(
    mean(functionModel8WeeksHot$data$mass, na.rm = T),
    mean(functionModel8WeeksHot$data$mass, na.rm = T) +
      2 * sd(functionModel8WeeksHot$data$mass, na.rm = T)
  )
) %>%
  mutate("tarsus" = predict(functionModel8WeeksHot,
    newdata = .,
    robust = TRUE, re_form = NA,
    resp = "tarsus", summary = TRUE
  )[, "Estimate"]) %>%
  predict(functionModel8WeeksHot,
    newdata = .,
    robust = TRUE, re_form = NA,
    resp = "slope", summary = FALSE
  ) %>%
  as.data.frame() %>%
  rename("Average" = "V1", "Large" = "V2") %>%
  mutate_all(.funs = function(x) {
    x + mean(slopeDataHot8$slope, na.rm = T)
  })

mismatchDFMassTestAllometric <- build_hdf(
  vars = list(
    mismatchDFMassAllometric$Average,
    mismatchDFMassAllometric$Large
  ),
  priors = list(
    rnorm(nrow(mismatchDFMassAllometric), 0, 0.01),
    rnorm(nrow(mismatchDFMassAllometric), 0, 0.01)
  ),
  names = c("Average", "Large")
)

mismatchTestMassAllometric <- hypothesis_df("Large > Average",
  mismatchDFMassTestAllometric,
  class = "b", alpha = 0.05
)

mismatchTestMassAllometric$hypothesis$Hypothesis <-
  "Large Metabolic Slope > Average Slope"

# And summarising

caption <- paste0(
```

```
  "Results of a non-linear hypothesis ",
  "tests comparing predicted metabolic slopes in the heat (30°C - 40°C) ",
  "among eight week old Japanese quail of average or atypically large ",
  "body size (mean + 2 standard deviations), while assuming ",
  "that tarsus length scales allometrically . Posterior probabilities are ",
  "calculated using the Savage-Dickey ",
  "density ratio method."
)

mismatchTestMassTable <- mismatchTestMassAllometric$hypothesis %>%
  mutate(Hypothesis = c(
    "Large Metabolic Slope > Average Slope"
  )) %>%
  select(-c(Evid.Ratio, Star)) %>%
  mutate(
    Estimate = round(Estimate, digits = 4),
    "Est.Error" = round(Est.Error, digits = 4),
    "CI.Lower" = round(CI.Lower, digits = 4),
    "CI.Upper" = round(CI.Upper, digits = 4)
  ) %>%
  rename(
    "Difference Between Metabolic Slopes" = Estimate,
    "Standard Error" = Est.Error,
    `2.5 % CI` = "CI.Lower",
    `97.5 % CI` = "CI.Upper",
    "Posterior Probability" = Post.Prob
  ) %>%
  kbl(.,
    longtable = T, booktabs = T, format = "latex",
    caption = caption
  ) %>%
  column_spec(column = c(1:2), width = "2.5cm") %>%
  column_spec(column = c(3:10), width = "2cm") %>%
  kable_styling(latex_options = "striped")

mismatchTestMassTable
```

***Table 37:*** *Results of a non-linear hypothesis tests comparing predicted metabolic slopes in the heat (30°C - 40°C) among eight week old Japanese quail of average or atypically large body size (mean + 2 standard deviations), while assuming that tarsus length scales allometrically . Posterior probabilities are calculated using the Savage-Dickey density ratio method.*

| Hypothesis | Difference Between Metabolic Slopes | Standard Error | 2.5 % CI | 97.5 % CI | Posterior Probability |
|---|---|---|---|---|---|
| Large Metabolic Slope > Average Slope | 0.0139 | 0.0525 | -0.0709 | 0.0964 | 0.6235 |

We now scale model coefficients to represent the effects of changing a predictor by one standard deviation (or categorical level) on standard deviations in the response variable. After, we use these transformed coefficients to visualise direct, indirect, and total effect of individual predictors on metabolic slopes in the heat among mature quail.

```
scaledBetasHot3 <- as.data.frame(functionModel3WeeksHot) %>%
  mutate(b_mass_pretreatmentA = b_mass_pretreatmentA/
           sd(functionModel3WeeksHot$data$mass),
         b_mass_pretreatmentC = b_mass_pretreatmentC/
           sd(functionModel3WeeksHot$data$mass),
         b_tarsus_pretreatmentA = b_tarsus_pretreatmentA/
           sd(functionModel3WeeksHot$data$tarsus),
         b_tarsus_pretreatmentC = b_tarsus_pretreatmentC/
           sd(functionModel3WeeksHot$data$tarsus),
         b_tarsus_mass =
```

```
           (b_tarsus_mass*sd(functionModel3WeeksHot$data$mass))/
           sd(functionModel3WeeksHot$data$tarsus),
         b_slope_mass =
           (b_slope_mass*sd(functionModel3WeeksHot$data$mass))/
           sd(functionModel3WeeksHot$data$slope),
         b_slope_tarsus =
           (b_slope_tarsus*sd(functionModel3WeeksHot$data$tarsus))/
           sd(functionModel3WeeksHot$data$slope),
         b_slope_pretreatmentA = b_slope_pretreatmentA/
           sd(functionModel3WeeksHot$data$slope),
         b_slope_pretreatmentC = b_slope_pretreatmentC/
           sd(functionModel3WeeksHot$data$slope),
  )

fullEffectThreeWeeksBetasHot <- scaledBetasHot3 %>%
  mutate("Effects" = "Direct Effects") %>%
  mutate(
    "Body Mass" = b_slope_mass,
    "Tarsus Length" = b_slope_tarsus,
    "Cold Rearing\n(10°C)" = b_slope_pretreatmentA,
    "Warm Rearing\n(30°C)" = b_slope_pretreatmentC
  ) %>%
  select(Effects, `Body Mass`, `Tarsus Length`,
         `Cold Rearing\n(10°C)`,
         `Warm Rearing\n(30°C)`) %>%
  rbind(
    .,
    scaledBetasHot3 %>%
      mutate("Effects" = "Indirect Effects") %>%
      mutate(
        "Body Mass" = b_tarsus_mass *
          b_slope_tarsus,
        "Tarsus Length" = NA,
        "Cold Rearing\n(10°C)" =
          b_tarsus_pretreatmentA * b_slope_tarsus +
          b_mass_pretreatmentA * b_slope_mass +
          b_mass_pretreatmentA * b_tarsus_mass *
          b_slope_tarsus,
        "Warm Rearing\n(30°C)" =
          b_tarsus_pretreatmentC * b_slope_tarsus +
          b_mass_pretreatmentC * b_slope_mass +
          b_mass_pretreatmentC * b_tarsus_mass *
          b_slope_tarsus,
      ) %>%
      select(Effects, `Body Mass`, `Tarsus Length`,
             `Cold Rearing\n(10°C)`, `Warm Rearing\n(30°C)`)
  ) %>%
  rbind(., scaledBetasHot3 %>%
    mutate("Effects" = "Total Effects") %>%
    mutate(
      "Body Mass" =
        b_slope_mass +
          b_tarsus_mass * b_slope_tarsus,
      "Tarsus Length" =
        b_slope_tarsus,
      "Cold Rearing\n(10°C)" =
        b_slope_pretreatmentA +
          b_mass_pretreatmentA *
        b_slope_mass +
          b_tarsus_pretreatmentA *
        b_slope_tarsus +
          b_mass_pretreatmentA *
        b_tarsus_mass * b_slope_tarsus,
      "Warm Rearing\n(30°C)" =
         b_slope_pretreatmentC +
          b_mass_pretreatmentC *
        b_slope_mass +
```

```
          b_tarsus_pretreatmentC *
        b_slope_tarsus +
          b_mass_pretreatmentC *
        b_tarsus_mass * b_slope_tarsus,
    ) %>%
    select(Effects, `Body Mass`, `Tarsus Length`,
             `Cold Rearing\n(10°C)`, `Warm Rearing\n(30°C)`)
    ) %>%
  pivot_longer(c(-Effects), names_to = "var", values_to = "values") %>%
  mutate(var = factor(var,
                      levels = c("Tarsus Length",
                                 "Body Mass",
                                 "Warm Rearing\n(30°C)",
                                 "Cold Rearing\n(10°C)")))

fullEffectThreeWeeksHot <- fullEffectThreeWeeksBetasHot %>%
  ggplot(aes(x = values, y = var, fill = var)) +
  facet_wrap(~Effects) +
  stat_halfeye(normalize = "xy", colour = "black", alpha = 0.7) +
  geom_vline(xintercept = 0, linetype = "dashed",
             colour = "black") +
  xlab(".                   Effect on Metabolic Slope (standard deviations)") +
  scale_fill_manual(values = c("black", "grey50", "#CD5C5C", "#7BB4E3")) +
  theme_classic() +
  theme(
    legend.position = "none", axis.title.y = element_blank(),
    axis.text.y = element_text(size = 11, colour = "black",
                               family = "Noto Sans"),
    axis.title.x = element_text(family = "Noto Sans", hjust = -0.005),
    axis.text.x = element_text(family = "Noto Sans")
  )

showtext_auto(enable = TRUE)

ggsave("../plots/indirectEffectsOnResistance3WeeksHot_3ScaledFold.pdf",
       fullEffectThreeWeeksHot,
  dpi = 800, width = 8, height = 6
)
showtext_auto(enable = FALSE)

# Summarising results

caption <- paste0(
  "Direct, indirect, and total effects of morphology and ",
  "rearing temperature on metabolic slope ",
  "(fold metabolism at thermoneutrality/°C) ",
  "of three week old Japanese quail. Metabolic slopes ",
  "represent those observed above thermoneutrality ",
  "(30°C - 40°C). Effects are derived from ",
  "a Bayesian path analysis and represent those predicted for a ",
  "change in one standard deviation (or categorical level) of ",
  "a given predictor on the standard deviation of metabolic ",
  "slopes. Estimates indicate posterior medians and credible ",
  "intervels (CIs) indicate quantile intervals."
)

week3ResultsScaledHot <-
  fullEffectThreeWeeksBetasHot %>%
  filter(!is.na(values) & !is.nan(values)) %>%
  group_by(Effects, var) %>%
  summarise(
    "Estimate" = median(values),
    "50\\% CIs" = paste0(
      "[",
      round(
        quantile(values, probs = 0.1, type = 8),
        digits = 4
```

```
    ),
    ", ",
    round(
      quantile(values, probs = 0.9, type = 8),
      digits = 4
    ),
    "]"
  ),
  "95\\% CIs" = paste0(
    "[",
    round(
      quantile(values, probs = 0.025, type = 8),
      digits = 4
    ),
    ", ",
    round(
      quantile(values, probs = 0.975, type = 8),
      digits = 4
    ),
    "]"
  )
) %>%
mutate("Effects" = gsub("[[:space:]].*", "", Effects)) %>%
select(
  "Predictor" = "var", "Effect Level" = "Effects",
  Estimate, `50\\% CIs`, `95\\% CIs`
) %>%
arrange(Predictor, `Effect Level`) %>%
kbl(.,
  longtable = T, booktabs = T, format = "latex", escape = FALSE,
  caption = caption
) %>%
column_spec(column = c(1:2), width = "2.2cm") %>%
column_spec(column = c(3:10), width = "1.9cm") %>%
kable_styling(latex_options = "striped")

week3ResultsScaledHot
```

***Table 38:*** *Direct, indirect, and total effects of morphology and rearing temperature on metabolic slope (fold metabolism at thermoneutrality/°C) of three week old Japanese quail. Metabolic slopes represent those observed above thermoneutrality (30°C - 40°C). Effects are derived from a Bayesian path analysis and represent those predicted for a change in one standard deviation (or categorical level) of a given predictor on the standard deviation of metabolic slopes. Estimates indicate posterior medians and credible intervels (CIs) indicate quantile intervals.*

| Predictor | Effect Level | Estimate | 50% CIs | 95% CIs |
|---|---|---|---|---|
| Tarsus Length | Direct | -0.0906719 | [-0.1946, 0.0225] | [-0.2492, 0.086] |
| Tarsus Length | Total | -0.0906719 | [-0.1946, 0.0225] | [-0.2492, 0.086] |
| Body Mass | Direct | 0.0959500 | [-0.0183, 0.2023] | [-0.0821, 0.2565] |
| Body Mass | Indirect | -0.0272623 | [-0.0669, 0.0066] | [-0.0937, 0.0274] |
| Body Mass | Total | 0.0652326 | [-0.0295, 0.1572] | [-0.0797, 0.21] |
| Warm Rearing (30°C) | Direct | -0.3700967 | [-0.5898, -0.1261] | [-0.7118, 0.0306] |
| Warm Rearing (30°C) | Indirect | -0.0331177 | [-0.1215, 0.026] | [-0.1888, 0.0648] |
| Warm Rearing (30°C) | Total | -0.4105403 | [-0.6369, -0.1663] | [-0.761, -0.0047] |
| Cold Rearing (10°C) | Direct | -0.1310159 | [-0.3611, 0.0839] | [-0.4876, 0.1963] |

| | | | | |
|---|---|---|---|---|
| Cold Rearing (10°C) | Indirect | -0.0432073 | [-0.1321, 0.019] | [-0.1961, 0.0572] |
| Cold Rearing (10°C) | Total | -0.1828865 | [-0.406, 0.0305] | [-0.5257, 0.147] |

```
# Continuing for data from mature quail

scaledBetasHot8 <- as.data.frame(functionModel8WeeksHot) %>%
  mutate(b_mass_pretreatmentA = b_mass_pretreatmentA/
           sd(functionModel8WeeksHot$data$mass),
         b_mass_pretreatmentC = b_mass_pretreatmentC/
           sd(functionModel8WeeksHot$data$mass),
         b_tarsus_pretreatmentA = b_tarsus_pretreatmentA/
           sd(functionModel8WeeksHot$data$tarsus),
         b_tarsus_pretreatmentC = b_tarsus_pretreatmentC/
           sd(functionModel8WeeksHot$data$tarsus),
         b_tarsus_mass =
           (b_tarsus_mass*sd(functionModel8WeeksHot$data$mass))/
           sd(functionModel8WeeksHot$data$tarsus),
         b_slope_mass =
           (b_slope_mass*sd(functionModel8WeeksHot$data$mass))/
           sd(functionModel8WeeksHot$data$slope),
         b_slope_tarsus =
           (b_slope_tarsus*sd(functionModel8WeeksHot$data$tarsus))/
           sd(functionModel8WeeksHot$data$slope),
         b_slope_pretreatmentA = b_slope_pretreatmentA/
           sd(functionModel8WeeksHot$data$slope),
         b_slope_pretreatmentC = b_slope_pretreatmentC/
           sd(functionModel8WeeksHot$data$slope),
  )

fullEffectEightWeeksBetasHot <- scaledBetasHot8 %>%
  mutate("Effects" = "Direct Effects") %>%
  mutate(
    "Body Mass" = b_slope_mass,
    "Tarsus Length" = b_slope_tarsus,
    "Cold Rearing\n(10°C)" = b_slope_pretreatmentA,
    "Warm Rearing\n(30°C)" = b_slope_pretreatmentC
  ) %>%
  select(Effects, `Body Mass`, `Tarsus Length`,
         `Cold Rearing\n(10°C)`,
         `Warm Rearing\n(30°C)`) %>%
  rbind(
    .,
    scaledBetasHot8 %>%
      mutate("Effects" = "Indirect Effects") %>%
      mutate(
        "Body Mass" = b_tarsus_mass *
          b_slope_tarsus,
        "Tarsus Length" = NA,
        "Cold Rearing\n(10°C)" =
          b_tarsus_pretreatmentA * b_slope_tarsus +
          b_mass_pretreatmentA * b_slope_mass +
          b_mass_pretreatmentA * b_tarsus_mass *
          b_slope_tarsus,
        "Warm Rearing\n(30°C)" =
          b_tarsus_pretreatmentC * b_slope_tarsus +
          b_mass_pretreatmentC * b_slope_mass +
          b_mass_pretreatmentC * b_tarsus_mass *
          b_slope_tarsus,
      ) %>%
      select(Effects, `Body Mass`, `Tarsus Length`,
             `Cold Rearing\n(10°C)`, `Warm Rearing\n(30°C)`)
  ) %>%
  rbind(., scaledBetasHot8 %>%
    mutate("Effects" = "Total Effects") %>%
```

```
    mutate(
      "Body Mass" =
        b_slope_mass +
          b_tarsus_mass * b_slope_tarsus,
      "Tarsus Length" =
        b_slope_tarsus,
      "Cold Rearing\n(10°C)" =
        b_slope_pretreatmentA +
          b_mass_pretreatmentA *
        b_slope_mass +
          b_tarsus_pretreatmentA *
        b_slope_tarsus +
          b_mass_pretreatmentA *
        b_tarsus_mass * b_slope_tarsus,
      "Warm Rearing\n(30°C)" =
         b_slope_pretreatmentC +
          b_mass_pretreatmentC *
        b_slope_mass +
          b_tarsus_pretreatmentC *
        b_slope_tarsus +
          b_mass_pretreatmentC *
        b_tarsus_mass * b_slope_tarsus,
    ) %>%
    select(Effects, `Body Mass`, `Tarsus Length`,
             `Cold Rearing\n(10°C)`, `Warm Rearing\n(30°C)`)
    ) %>%
  pivot_longer(c(-Effects), names_to = "var", values_to = "values") %>%
  mutate(var = factor(var,
                      levels = c("Tarsus Length",
                                 "Body Mass",
                                 "Warm Rearing\n(30°C)",
                                 "Cold Rearing\n(10°C)")))

fullEffectEightWeeksHot <- fullEffectEightWeeksBetasHot %>%
  ggplot(aes(x = values, y = var, fill = var)) +
  facet_wrap(~Effects) +
  stat_halfeye(normalize = "xy", colour = "black", alpha = 0.7) +
  geom_vline(xintercept = 0, linetype = "dashed",
             colour = "black") +
  xlab(".                   Effect on Metabolic Slope (standard deviations)") +
  scale_fill_manual(values = c("black", "grey50", "#CD5C5C", "#7BB4E3")) +
  theme_classic() +
  theme(
    legend.position = "none", axis.title.y = element_blank(),
    axis.text.y = element_text(size = 11, colour = "black",
                           family = "Noto Sans"),
    axis.title.x = element_text(family = "Noto Sans", hjust = -0.005),
    axis.text.x = element_text(family = "Noto Sans")
  )

showtext_auto(enable = TRUE)

fullEffectThreeWeeksHot/fullEffectEightWeeksHot +
  plot_annotation(tag_levels = "A")
```

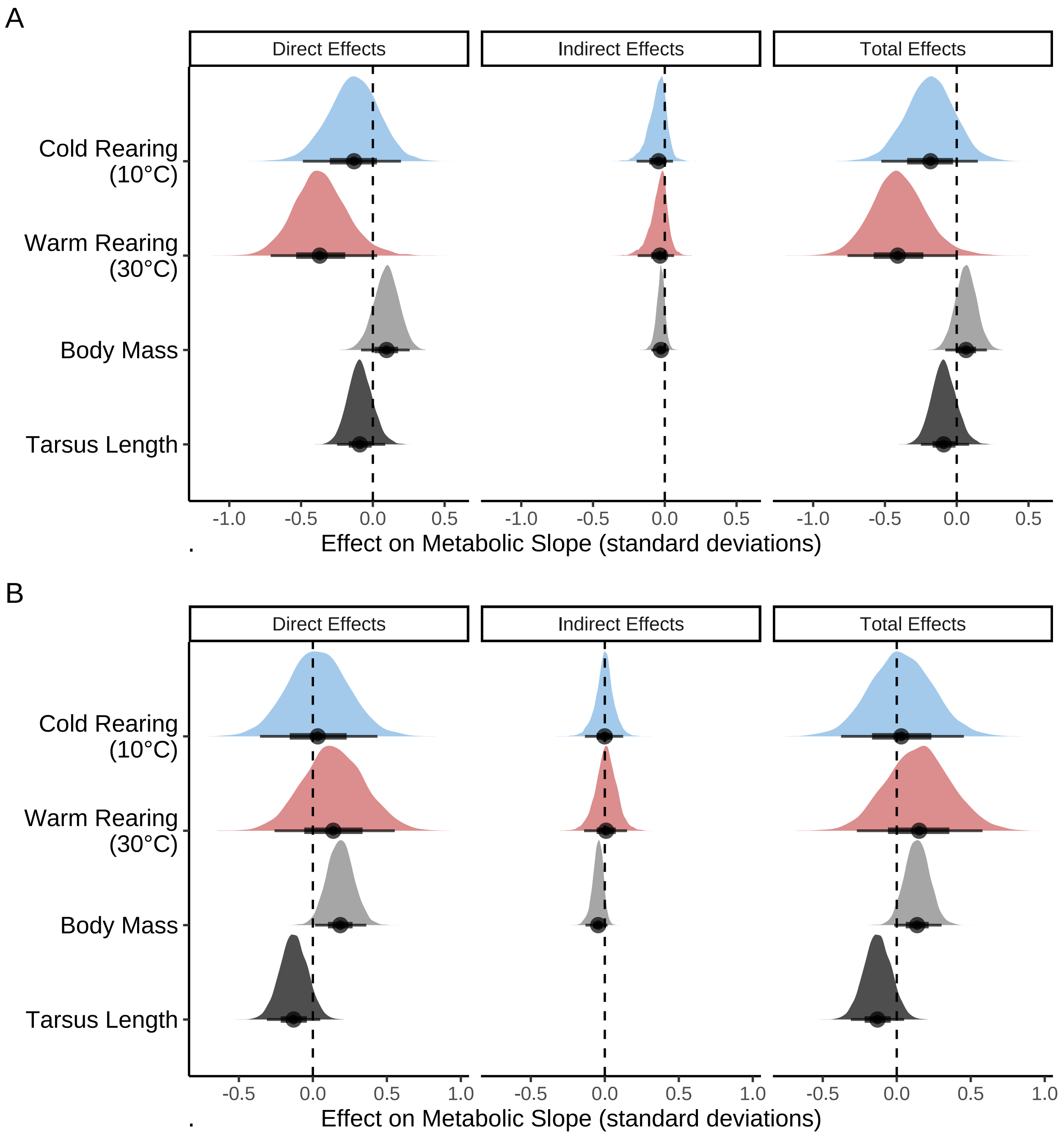


***Figure 73:*** *Effects of morphology (body mass, g, and tarsus length, mm) on metabolic slopes (in standard deviations) at ambient temperatures above thermoneutrality (30°C - 40°C) in both three week old (panel A) and eight week old (panel B) Japanese quail. Distributions represent those derived from posteriors of Bayesian path analyses.*

```
ggsave("../plots/indirectEffectsOnResistance8WeeksHot_3ScaledFold.pdf",
       fullEffectEightWeeksHot,
  dpi = 800, width = 8, height = 6
)
showtext_auto(enable = FALSE)

# Summarising results
```

```
caption <- paste0("Direct, indirect, and total effects of morphology and ",
                  "rearing temperature on metabolic slope ",
                  "(fold metabolism at thermoneutrality/ ",
                  "°C) ",
                  "of eight week old Japanese quail. Metabolic slopes represent ",
                  "those observed above thermoneutrality (30°C - 40°C). ",
                  "Effects are derived from ",
                  "a Bayesian path analysis and represent those predicted for a ",
                  "change in one standard deviation (or categorical level) of ",
                  "a given predictor on the standard deviation of metabolic ",
                  "slopes. Estimates indicate posterior medians and credible ",
                  "intervels (CIs) indicate quantile intervals."
)

week8ResultsScaledHot <-
  fullEffectEightWeeksBetasHot %>%
  filter(!is.na(values) & !is.nan(values)) %>%
  group_by(Effects, var) %>%
  summarise(
    "Estimate" = median(values),
    "50\\% CIs" = paste0(
      "[",
      round(
        quantile(values, probs = 0.15, type = 8),
        digits = 4
      ),
      ", ",
      round(
        quantile(values, probs = 0.95, type = 8),
        digits = 4
      ),
      "]"
    ),
    "95\\% CIs" = paste0(
      "[",
      round(
        quantile(values, probs = 0.025, type = 8),
        digits = 4
      ),
      ", ",
      round(
        quantile(values, probs = 0.975, type = 8),
        digits = 4
      ),
      "]"
    )
  ) %>%
  mutate("Effects" = gsub("[[:space:]].*", "", Effects)) %>%
  select(
    "Predictor" = "var", "Effect Level" = "Effects",
    Estimate, `50\\% CIs`, `95\\% CIs`
  ) %>%
  arrange(Predictor, `Effect Level`) %>%
  kbl(.,
    longtable = T, booktabs = T, format = "latex", escape = FALSE,
    caption = caption
  ) %>%
  column_spec(column = c(1:2), width = "2.2cm") %>%
  column_spec(column = c(3:10), width = "1.9cm") %>%
  kable_styling(latex_options = "striped")

week8ResultsScaledHot
```

***Table 39:*** *Direct, indirect, and total effects of morphology and rearing temperature on metabolic slope (fold metabolism at thermoneutrality/ °C) of eight week old Japanese quail. Metabolic slopes represent those observed above thermoneutrality (30°C - 40°C). Effects are derived from a Bayesian path analysis and represent those predicted for a change in one standard deviation (or categorical level) of a given predictor on the standard deviation of metabolic slopes. Estimates indicate posterior medians and credible intervels (CIs) indicate quantile intervals.*

| Predictor | Effect Level | Estimate | 50% CIs | 95% CIs |
|---|---|---|---|---|
| Tarsus Length | Direct | -0.1301246 | [-0.2236, 0.0223] | [-0.3105, 0.0496] |
| Tarsus Length | Total | -0.1301246 | [-0.2236, 0.0223] | [-0.3105, 0.0496] |
| Body Mass | Direct | 0.1853977 | [0.0951, 0.3335] | [0.015, 0.3611] |
| Body Mass | Indirect | -0.0444896 | [-0.0838, 0.0077] | [-0.1306, 0.0186] |
| Body Mass | Total | 0.1380549 | [0.054, 0.2728] | [-0.017, 0.303] |
| Warm Rearing (30°C) | Direct | 0.1377110 | [-0.0744, 0.4885] | [-0.2593, 0.5549] |
| Warm Rearing (30°C) | Indirect | 0.0085513 | [-0.0618, 0.1228] | [-0.14, 0.1505] |
| Warm Rearing (30°C) | Total | 0.1515528 | [-0.0782, 0.5094] | [-0.2701, 0.5794] |
| Cold Rearing (10°C) | Direct | 0.0330645 | [-0.1732, 0.3733] | [-0.3569, 0.4366] |
| Cold Rearing (10°C) | Indirect | -0.0006204 | [-0.0608, 0.1006] | [-0.1344, 0.1241] |
| Cold Rearing (10°C) | Total | 0.0306080 | [-0.1838, 0.3801] | [-0.3766, 0.4539] |

Again, the above path analyses are repeated but while relativising tarsus length measurements by body mass. This is achieved by calculating tarsus length residuals from simple regressions of tarsus length by body mass (priors for models with three week data: intercept = $\mathcal{N}[30, 2.5]$, body mass = $S\mathcal{N}[0, 0.25, 5]$; priors for models with eight week data: intercept = $\mathcal{N}[35, 2.5]$, body mass = $S\mathcal{N}[0, 0.25, 5]$), then loading these residuals into the full analysis. Priors for these path analyses remain the same as those used previously with response variables unscaled.

Given that these adjusted path analyses are conducted merely for comparative purposes (i.e. for comparison with models analyses where tarsus length is not relativised), residuals are not rigorously scrutinised, nor are model coefficients plotted in detail. Instead, final model outcomes are printed after checking for prior suitability and chain convergence.

```
functionModel3WeeksHotResidualPPCheck <-
  brm(
    data = slopeDataHot3 %>%
      mutate(
        mass = mass - mean(mass, na.rm = T),
        pretreatment = ifelse(pretreatment == "cold", "A",
          ifelse(pretreatment == "neutral", "B", "C")
        ),
        slope = slope -
          mean(slope, na.rm = T)
      ) %>%
      mutate(pretreatment = factor(pretreatment,
        levels = c("B", "A", "C")
      )) %>%
      drop_na() %>%
      merge(., data %>%
        select(ring, "batch" = exp) %>%
        distinct(),
```

```
      by = "ring", all.x = TRUE
      ) %>%
      mutate(
        "residualTarsus" =
          residuals(
            brm(tarsusLengthMean ~ mass,
              data = .,
              prior = c(
                set_prior("skew_normal(0, 0.25, 5)",
                          class = "b"),
                set_prior("normal(35, 2.5)",
                          class = "Intercept")
              ),
              family = "gaussian",
              iter = 50000, warmup = 10000, cores = 1,
              chains = 4, thin = 20,
              silent = TRUE, refresh = 0,
              file = "./models/residualTarsusForPathAnalysisPPCheck.Rds"
            ), robust = TRUE
          )[, "Estimate"]
      ),
    family = "gaussian",
    bf(mass ~ pretreatment + (1 | batch)) +
      bf(residualTarsus ~ pretreatment + (1 | batch)) +
      bf(slope ~ mass + residualTarsus + pretreatment + (1 | batch)) +
      set_rescor(FALSE),
    prior = c(
      set_prior("normal(0, 10)",
        class = "Intercept",
        resp = "mass"
      ),
      set_prior("normal(0, 15)",
        class = "b",
        coef = "pretreatmentA",
        resp = "mass"
      ),
      set_prior("normal(0, 15)",
        class = "b",
        coef = "pretreatmentC",
        resp = "mass"
      ),
      set_prior("exponential(2.5)",
        class = "sd",
        group = "batch",
        resp = "mass"
      ),
      set_prior("exponential(0.05)",
        class = "sigma",
        resp = "mass"
      ),
      set_prior("normal(0, 2.5)",
        class = "Intercept",
        resp = "residualTarsus"
      ),
      set_prior("normal(0, 2.5",
        class = "b",
        coef = "pretreatmentA",
        resp = "residualTarsus"
      ),
      set_prior("normal(0, 2.5)",
        class = "b",
        coef = "pretreatmentC",
        resp = "residualTarsus"
      ),
      set_prior("exponential(2)",
        class = "sd",
        group = "batch",
```

```
        resp = "residualTarsus"
      ),
      set_prior("exponential(1)",
        class = "sigma",
        resp = "residualTarsus"
      ),
      set_prior("normal(0, 0.125)",
        class = "Intercept",
        resp = "slope"
      ),
      set_prior("normal(0, 0.125)",
        class = "b",
        coef = "pretreatmentA",
        resp = "slope"
      ),
      set_prior("normal(0, 0.125)",
        class = "b",
        coef = "pretreatmentC",
        resp = "slope"
      ),
      set_prior("normal(0, 0.004)",
        class = "b",
        coef = "mass",
        resp = "slope"
      ),
      set_prior("normal(0, 0.015)",
        class = "b",
        coef = "residualTarsus",
        resp = "slope"
      ),
      set_prior("exponential(50)",
        class = "sd",
        group = "batch",
        resp = "slope"
      ),
      set_prior("exponential(10)",
        class = "sigma",
        resp = "slope"
      )
    ),
    iter = 50000, warmup = 10000, cores = 4, chains = 4, thin = 20,
    control = list(adapt_delta = .97, max_treedepth = 14),
    silent = TRUE, refresh = 0,
    sample_prior = "only",
    file = "./models/threeWeekFunctionModelHotResidualPPCheck.Rds"
  )

functionModel8WeeksHotResidualPPCheck <-
  brm(
    data = slopeDataHot8 %>%
      mutate(
        mass = mass - mean(mass, na.rm = T),
        pretreatment = ifelse(pretreatment == "cold", "A",
          ifelse(pretreatment == "neutral", "B", "C")
        ),
        slope = slope -
          mean(slope, na.rm = T)
      ) %>%
      mutate(pretreatment = factor(pretreatment,
        levels = c("B", "A", "C")
      )) %>%
      drop_na() %>%
      merge(., data %>%
        select(ring, "batch" = exp) %>%
        distinct(),
      by = "ring", all.x = TRUE
      ) %>%
```

```
    mutate(
      "residualTarsus" =
        residuals(
          brm(tarsusLengthMean ~ mass,
            data = .,
            prior = c(
              set_prior("skew_normal(0, 0.25, 5)",
                        class = "b"),
              set_prior("normal(35, 2.5)",
                        class = "Intercept")
            ),
            family = "gaussian",
            iter = 50000, warmup = 10000, cores = 1,
            chains = 4, thin = 20,
            silent = TRUE, refresh = 0
          ), robust = TRUE
        )[, "Estimate"]
    ),
  family = "gaussian",
  bf(mass ~ pretreatment + (1 | batch)) +
    bf(residualTarsus ~ pretreatment + (1 | batch)) +
    bf(slope ~ mass + residualTarsus + pretreatment + (1 | batch)) +
    set_rescor(FALSE),
  prior = c(
    set_prior("normal(0, 10)",
      class = "Intercept",
      resp = "mass"
    ),
    set_prior("normal(0, 25)",
      class = "b",
      coef = "pretreatmentA",
      resp = "mass"
    ),
    set_prior("normal(0, 25)",
      class = "b",
      coef = "pretreatmentC",
      resp = "mass"
    ),
    set_prior("exponential(2.5)",
      class = "sd",
      group = "batch",
      resp = "mass"
    ),
    set_prior("exponential(0.05)",
      class = "sigma",
      resp = "mass"
    ),
    set_prior("normal(0, 3)",
      class = "Intercept",
      resp = "residualTarsus"
    ),
    set_prior("normal(0, 3)",
      class = "b",
      coef = "pretreatmentA",
      resp = "residualTarsus"
    ),
    set_prior("normal(0, 3)",
      class = "b",
      coef = "pretreatmentC",
      resp = "residualTarsus"
    ),
    set_prior("exponential(2)",
      class = "sd",
      group = "batch",
      resp = "residualTarsus"
    ),
    set_prior("exponential(0.75)",
```

```
        class = "sigma",
        resp = "residualTarsus"
      ),
      set_prior("normal(0, 0.1)",
        class = "Intercept",
        resp = "slope"
      ),
      set_prior("normal(0, 0.1)",
        class = "b",
        coef = "pretreatmentA",
        resp = "slope"
      ),
      set_prior("normal(0, 0.1)",
        class = "b",
        coef = "pretreatmentC",
        resp = "slope"
      ),
      set_prior("normal(0, 0.0025)",
        class = "b",
        coef = "mass",
        resp = "slope"
      ),
      set_prior("normal(0, 0.03)",
        class = "b",
        coef = "residualTarsus",
        resp = "slope"
      ),
      set_prior("exponential(50)",
        class = "sd",
        group = "batch",
        resp = "slope"
      ),
      set_prior("exponential(10)",
        class = "sigma",
        resp = "slope"
      )
    ),
    iter = 50000, warmup = 10000, cores = 4, chains = 4, thin = 20,
    control = list(adapt_delta = .97, max_treedepth = 14),
    silent = TRUE, refresh = 0,
    sample_prior = "only",
    file = "./models/eightWeekFunctionModelHotResidualPPCheckNew.Rds"
  )

p1 <- pp_check2(functionModel3WeeksHotResidualPPCheck,
  resp = "slope",
  xlab = paste0("Metabolic Slope (Fold Metabolism\n",
                "at Thermoneutrality/°C)")
)

p2 <- pp_check2(functionModel8WeeksHotResidualPPCheck,
  resp = "slope",
  xlab = paste0("Metabolic Slope (Fold Metabolism\n",
                "at Thermoneutrality/°C)")
  )

(p1 + p2) + plot_annotation(tag_levels = "A")
```

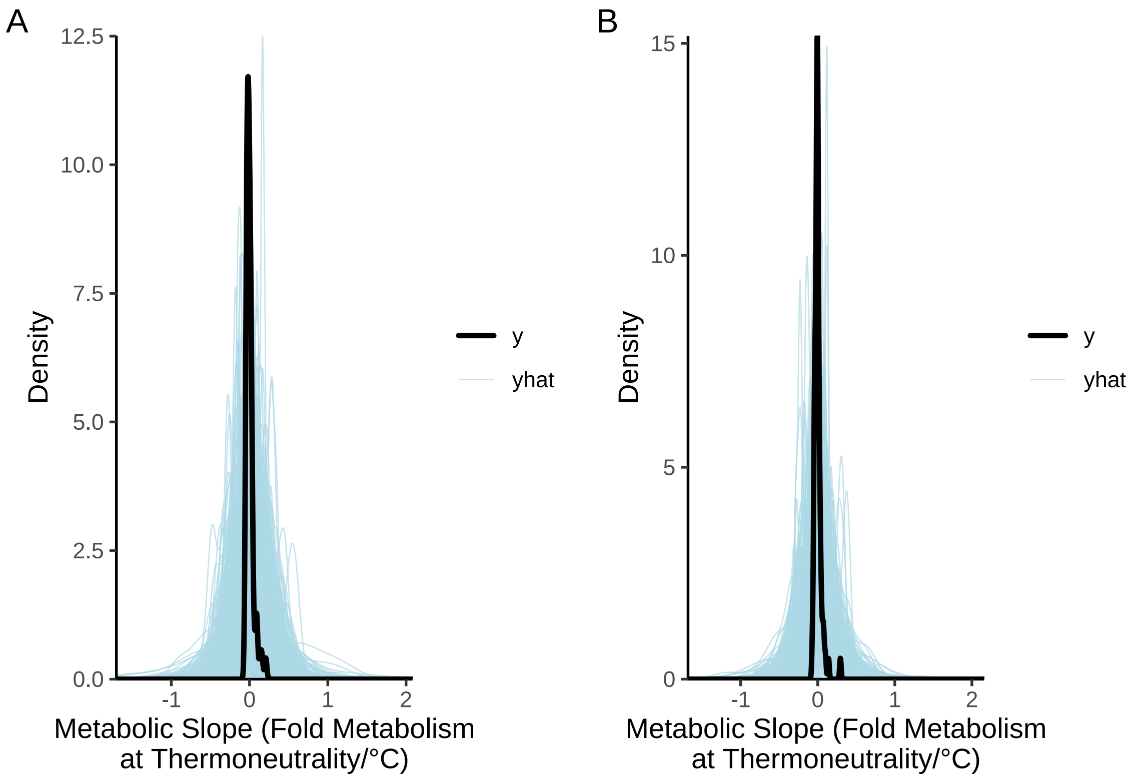


***Figure 74:*** *Prior predictive checks for two Bayesian path analyses predicting physiological responses to heat in three and eight week old Japanese quail. Tarsus length in these models is relativised by body mass (i.e. as tarsus length residuals). Black lines represent true densities of metabolic slopes (fold metabolism at thermoneutrality/°C) while blue lines represent densities estimated from model priors alone.*

```
# Priors very wide, but suitable.
```

```
functionModel3WeeksHotResidual <-
  brm(
    data = slopeDataHot3 %>%
      mutate(
        mass = mass - mean(mass, na.rm = T),
        pretreatment = ifelse(pretreatment == "cold", "A",
          ifelse(pretreatment == "neutral", "B", "C")
        ),
        slope = slope -
          mean(slope, na.rm = T)
      ) %>%
      mutate(pretreatment = factor(pretreatment,
        levels = c("B", "A", "C")
      )) %>%
      drop_na() %>%
      merge(., data %>%
        select(ring, "batch" = exp) %>%
        distinct(),
      by = "ring", all.x = TRUE
      ) %>%
      mutate(
        "residualTarsus" =
          residuals(
            brm(tarsusLengthMean ~ mass,
              data = .,
              prior = c(
                set_prior("skew_normal(0, 0.25, 5)",
                          class = "b"),
```

```
                set_prior("normal(35, 2.5)",
                          class = "Intercept")
              ),
              family = "gaussian",
              iter = 50000, warmup = 10000, cores = 1,
              chains = 4, thin = 20,
              silent = TRUE, refresh = 0
            ), robust = TRUE
          )[, "Estimate"]
      ),
    family = "gaussian",
    bf(mass ~ pretreatment + (1 | batch)) +
      bf(residualTarsus ~ pretreatment + (1 | batch)) +
      bf(slope ~ mass + residualTarsus + pretreatment + (1 | batch)) +
      set_rescor(FALSE),
    prior = c(
      set_prior("normal(0, 10)",
        class = "Intercept",
        resp = "mass"
      ),
      set_prior("normal(0, 15)",
        class = "b",
        coef = "pretreatmentA",
        resp = "mass"
      ),
      set_prior("normal(0, 15)",
        class = "b",
        coef = "pretreatmentC",
        resp = "mass"
      ),
      set_prior("exponential(2.5)",
        class = "sd",
        group = "batch",
        resp = "mass"
      ),
      set_prior("exponential(0.05)",
        class = "sigma",
        resp = "mass"
      ),
      set_prior("normal(0, 2.5)",
        class = "Intercept",
        resp = "residualTarsus"
      ),
      set_prior("normal(0, 2.5",
        class = "b",
        coef = "pretreatmentA",
        resp = "residualTarsus"
      ),
      set_prior("normal(0, 2.5)",
        class = "b",
        coef = "pretreatmentC",
        resp = "residualTarsus"
      ),
      set_prior("exponential(2)",
        class = "sd",
        group = "batch",
        resp = "residualTarsus"
      ),
      set_prior("exponential(1)",
        class = "sigma",
        resp = "residualTarsus"
      ),
      set_prior("normal(0, 0.125)",
        class = "Intercept",
        resp = "slope"
      ),
      set_prior("normal(0, 0.125)",
```

```
        class = "b",
        coef = "pretreatmentA",
        resp = "slope"
      ),
      set_prior("normal(0, 0.125)",
        class = "b",
        coef = "pretreatmentC",
        resp = "slope"
      ),
      set_prior("normal(0, 0.004)",
        class = "b",
        coef = "mass",
        resp = "slope"
      ),
      set_prior("normal(0, 0.015)",
        class = "b",
        coef = "residualTarsus",
        resp = "slope"
      ),
      set_prior("exponential(50)",
        class = "sd",
        group = "batch",
        resp = "slope"
      ),
      set_prior("exponential(10)",
        class = "sigma",
        resp = "slope"
      )
    ),
    iter = 50000, warmup = 10000, cores = 4, chains = 4, thin = 20,
    control = list(adapt_delta = .97, max_treedepth = 14),
    silent = TRUE, refresh = 0,
    file = "./models/threeWeekFunctionModelHotResidual.Rds"
  )

functionModel8WeeksHotResidual <-
  brm(
    data = slopeDataHot8 %>%
      mutate(
        mass = mass - mean(mass, na.rm = T),
        pretreatment = ifelse(pretreatment == "cold", "A",
          ifelse(pretreatment == "neutral", "B", "C")
        ),
        slope = slope -
          mean(slope, na.rm = T)
      ) %>%
      mutate(pretreatment = factor(pretreatment,
        levels = c("B", "A", "C")
      )) %>%
      drop_na() %>%
      merge(., data %>%
        select(ring, "batch" = exp) %>%
        distinct(),
      by = "ring", all.x = TRUE
      ) %>%
      mutate(
        "residualTarsus" =
          residuals(
            brm(tarsusLengthMean ~ mass,
              data = .,
              prior = c(
                set_prior("skew_normal(0, 0.25, 5)",
                          class = "b"),
                set_prior("normal(35, 2.5)",
                          class = "Intercept")
              ),
              family = "gaussian",
```

```
              iter = 50000, warmup = 10000, cores = 1,
              chains = 4, thin = 20,
              silent = TRUE, refresh = 0
            ), robust = TRUE
          )[, "Estimate"]
      ),
    family = "gaussian",
    bf(mass ~ pretreatment + (1 | batch)) +
      bf(residualTarsus ~ pretreatment + (1 | batch)) +
      bf(slope ~ mass + residualTarsus + pretreatment + (1 | batch)) +
      set_rescor(FALSE),
    prior = c(
      set_prior("normal(0, 10)",
        class = "Intercept",
        resp = "mass"
      ),
      set_prior("normal(0, 25)",
        class = "b",
        coef = "pretreatmentA",
        resp = "mass"
      ),
      set_prior("normal(0, 25)",
        class = "b",
        coef = "pretreatmentC",
        resp = "mass"
      ),
      set_prior("exponential(2.5)",
        class = "sd",
        group = "batch",
        resp = "mass"
      ),
      set_prior("exponential(0.05)",
        class = "sigma",
        resp = "mass"
      ),
      set_prior("normal(0, 3)",
        class = "Intercept",
        resp = "residualTarsus"
      ),
      set_prior("normal(0, 3)",
        class = "b",
        coef = "pretreatmentA",
        resp = "residualTarsus"
      ),
      set_prior("normal(0, 3)",
        class = "b",
        coef = "pretreatmentC",
        resp = "residualTarsus"
      ),
      set_prior("exponential(2)",
        class = "sd",
        group = "batch",
        resp = "residualTarsus"
      ),
      set_prior("exponential(0.75)",
        class = "sigma",
        resp = "residualTarsus"
      ),
      set_prior("normal(0, 0.1)",
        class = "Intercept",
        resp = "slope"
      ),
      set_prior("normal(0, 0.1)",
        class = "b",
        coef = "pretreatmentA",
        resp = "slope"
      ),
```

```
      set_prior("normal(0, 0.1)",
        class = "b",
        coef = "pretreatmentC",
        resp = "slope"
      ),
      set_prior("normal(0, 0.0025)",
        class = "b",
        coef = "mass",
        resp = "slope"
      ),
      set_prior("normal(0, 0.03)",
        class = "b",
        coef = "residualTarsus",
        resp = "slope"
      ),
      set_prior("exponential(50)",
        class = "sd",
        group = "batch",
        resp = "slope"
      ),
      set_prior("exponential(10)",
        class = "sigma",
        resp = "slope"
      )
    ),
    iter = 50000, warmup = 10000, cores = 4, chains = 4, thin = 20,
    control = list(adapt_delta = .97, max_treedepth = 14),
    silent = TRUE, refresh = 0,
    file = "./models/eightWeekFunctionModelHotResidualNew.Rds"
  )

# Minimal divergent transitions in both models.

p1 <- mcmc_neff(neff_ratio(functionModel3WeeksHotResidual)) +
  xlab(
    TeX('$Three~Week~N_{eff}/N$')
    ) +
  theme_classic() +
  theme(axis.text.y = element_blank(),
        axis.ticks.y = element_blank(),
        legend.position = "none")

p2 <- mcmc_rhat(rhat(functionModel3WeeksHotResidual)) +
  xlab(
    TeX('$\\hat{R}~(Three~Weeks)$')
    ) +
  theme_classic() +
  theme(axis.text.y = element_blank(),
        axis.ticks.y = element_blank(),
        legend.position = "none")

p3 <- functionModel3WeeksHotResidual$data %>%
  mutate(
    "Residuals" =
      residuals(functionModel3WeeksHotResidual,
        type = "ordinary",
        resp = "slope",
        robust = TRUE
      )[, "Estimate"]
  ) %>%
  ggplot(aes(x = Residuals)) +
  geom_density() +
  xlab("Three Week Metabolic\nSlope Residuals") +
  ylab("Density") +
  theme_classic()

p4 <- mcmc_neff(neff_ratio(functionModel8WeeksHotResidual)) +
```

```
  xlab(
    TeX('$Eight~Week~N_{eff}/N$')
    ) +
  theme_classic() +
  theme(axis.text.y = element_blank(),
        axis.ticks.y = element_blank(),
        legend.position = "none")

p5 <- mcmc_rhat(rhat(functionModel8WeeksHotResidual)) +
  xlab(
    TeX('$\\hat{R}~(Eight~Weeks)$')
  ) +
  theme_classic() +
  theme(axis.text.y = element_blank(),
        axis.ticks.y = element_blank(),
        legend.position = "none")

p6 <- functionModel8WeeksHotResidual$data %>%
  mutate(
    "Residuals" =
      residuals(functionModel8WeeksHotResidual,
        type = "ordinary",
        resp = "slope",
        robust = TRUE
      )[, "Estimate"]
  ) %>%
  ggplot(aes(x = Residuals)) +
  geom_density() +
  xlab("Eight Week Metabolic\nSlope Residuals") +
  ylab("Density") +
  theme_classic()

(p1 + p2 + p3) / (p4 + p5 + p6) +
  plot_annotation(tag_levels = "A")
```

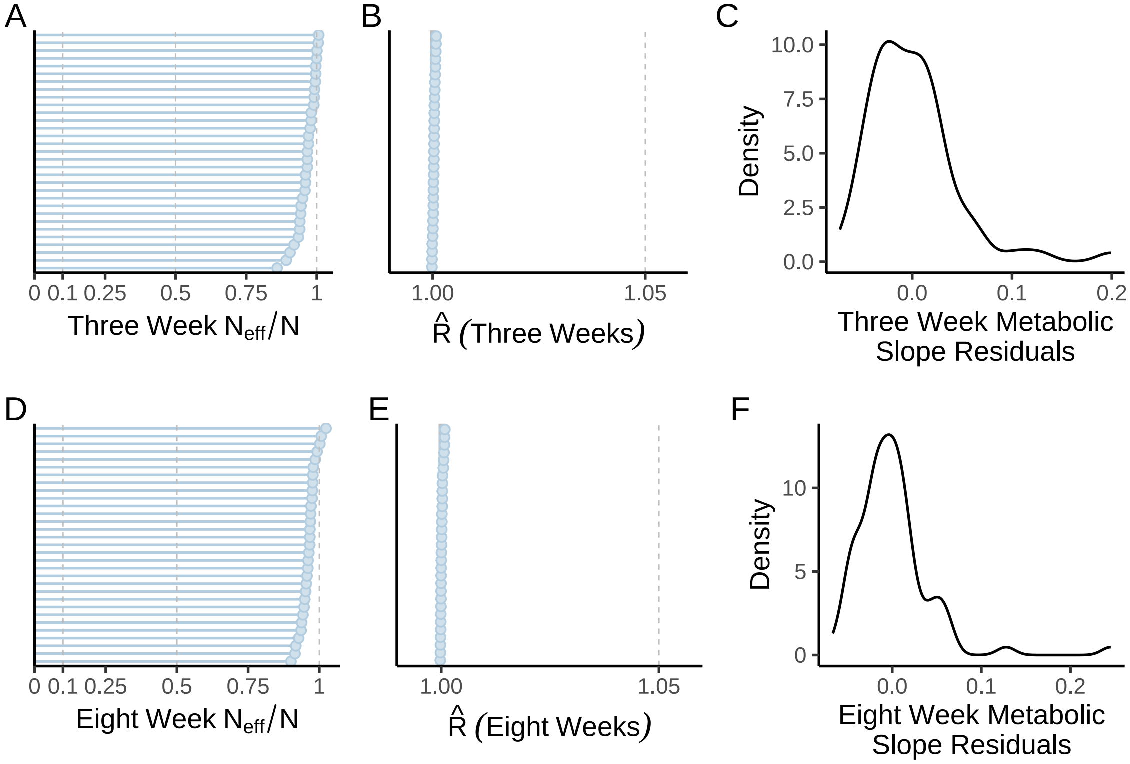


***Figure 75:*** *Gelman-Rubin ($\hat{R}$) statistics, ratio of effective sample sizes to sample sizes ($N_{eff}/N$), and residual densities from Bayesian path analyses predicting physiological responses to heat in three and eight week old Japanese quail. Tarsus length in these models is relativised by body mass (i.e. as tarsus length residuals). In panels A, B, D, and E, blue dots indicate Rhat or Neff/N values for a model parameter.*

```
# Chains evidently converged, autocorrelation between
# draws appears limited, and residuals reasonably normal.
# Extracting model outcomes for comparison.

caption <- paste0("Results from a Bayesian path analysis ",
                  "predicting metabolic responses to heat (metabolic slopes ",
                  "[fold metabolism at thermoneutrality/",
                  "°C] as a ",
                  "function of body mass (g), relative tarsus length ",
                  "(residuals; mm) and rearing conditions in three ",
                  "week old Japanese quail. Physiological measurements ",
                  "are made at ambient temperatures between ",
                  "30°C and ",
                  "40°C. Cold rearing indicates ",
                  "post-hatch rearing at 10°C, ",
                  "relative to 20°C (intercept), ",
                  "or 30°C ('warm rearing'). ",
                  "CI indicates quantile ",
                  "intervals and BF indicates Bayes Factors."
)

week3ResultsHeatResiduals <-
  as.data.frame(functionModel3WeeksHotResidual) %>%
  summarise_all(., .funs = median) %>%
  pivot_longer(everything(),
    names_to = "Parameter",
    values_to = "Estimate"
  ) %>%
  merge(., quantileCIs(functionModel3WeeksHotResidual, cis = c(50, 95)),
    by = "Parameter", all.x = TRUE
```

```
) %>%
filter(grepl("b_|sd_", Parameter)) %>%
rowwise() %>%
mutate("BF" = ifelse(Estimate < 0,
  (2 * mean(as.data.frame(
    functionModel3WeeksHotResidual
  )[, Parameter] <= 0)) /
    (2 * mean(as.data.frame(
      functionModel3WeeksHotResidual
    )[, Parameter] >= 0)),
  (2 * mean(as.data.frame(
    functionModel3WeeksHotResidual
  )[, Parameter] >= 0)) /
    (2 * mean(as.data.frame(
      functionModel3WeeksHotResidual
    )[, Parameter] <= 0))
)) %>%
ungroup() %>%
mutate(
  "Estimate" = round(Estimate, digits = 4),
  "BF" = round(BF, digits = 4),
  "N" = nrow(functionModel3WeeksHotResidual$data)
) %>%
mutate("Parameter" = ifelse(grepl("b_", Parameter),
  gsub("b_", "", Parameter),
  gsub(
    "Intercept", "batch",
    gsub(".*__", "", Parameter)
  )
)) %>%
mutate(
  "Response" = gsub("_.*", "", Parameter),
  "Parameter" = gsub(".*_", "", Parameter)
) %>%
merge(., tribble(
  ~Response, ~response, ~level,
  "mass", "Body Mass (g)", "A",
  "slope",
  "Metabolic Slope", "D",
  "residualTarsus", "Residual Tarsus Length (mm)", "B"
),
by = "Response"
) %>%
merge(., tribble(
  ~Parameter, ~parameter, ~number,
  "Intercept", "Intercept", "1",
  "mass", "Body Mass (g)", "4",
  "residualTarsus", "Residual Tarsus Length (mm)", "5",
  "pretreatmentA", "Cold Rearing", "2",
  "pretreatmentC", "Warm Rearing", "3",
  "batch", "Egg Batch [mu]", "6"
),
by = "Parameter"
) %>%
mutate(
  `50\\% HDI` = paste0("(", paste(
    round(Low_CI_50, digits = 4),
    round(High_CI_50, digits = 4),
    sep = ", "
  ), ")"),
  `95\\% HDI` = paste0("(", paste(
    round(Low_CI_95, digits = 4),
    round(High_CI_95, digits = 4),
    sep = ", "
  ), ")")
) %>%
select(-c(Low_CI_50, High_CI_50, Low_CI_95, High_CI_95)) %>%
```

```
  select(
    "Response" = "response", "Parameter" = "parameter", N,
    Estimate, `50\\% HDI`, `95\\% HDI`, BF, level, number
  ) %>%
  arrange(level, number) %>%
  select(-c(level, number)) %>%
  kbl(.,
    longtable = T, booktabs = T, format = "latex", escape = FALSE,
    caption = caption
  ) %>%
  column_spec(column = c(1:2), width = "2.2cm") %>%
  column_spec(column = c(3:10), width = "1.9cm") %>%
  kable_styling(latex_options = "striped")

week3ResultsHeatResiduals
```

***Table 40:*** *Results from a Bayesian path analysis predicting metabolic responses to heat (metabolic slopes [fold metabolism at thermoneutrality/°C] as a function of body mass (g), relative tarsus length (residuals; mm) and rearing conditions in three week old Japanese quail. Physiological measurements are made at ambient temperatures between 30°C and 40°C. Cold rearing indicates post-hatch rearing at 10°C, relative to 20°C (intercept), or 30°C ('warm rearing'). CI indicates quantile intervals and BF indicates Bayes Factors.*

| Response | Parameter | N | Estimate | 50% HDI | 95% HDI | BF |
|---|---|---|---|---|---|---|
| Body Mass (g) | Intercept | 69 | 3.7888 | (1.4092, 6.1479) | (-3.1511, 10.6327) | 6.1749 |
| Body Mass (g) | Cold Rearing | 69 | -8.5927 | (-11.5325, -5.7089) | (-16.8472, -0.0264) | 39.2010 |
| Body Mass (g) | Warm Rearing | 69 | 0.5114 | (-2.5549, 3.6148) | (-8.2087, 9.2989) | 1.1840 |
| Body Mass (g) | Egg Batch [mu] | 69 | 0.2670 | (0.1108, 0.5459) | (0.0108, 1.4297) | Inf |
| Residual Tarsus Length (mm) | Intercept | 69 | -0.7626 | (-1.318, -0.2091) | (-2.4045, 1.067) | 4.5402 |
| Residual Tarsus Length (mm) | Cold Rearing | 69 | 0.5778 | (0.0712, 1.08) | (-0.9312, 2.0621) | 3.5300 |
| Residual Tarsus Length (mm) | Warm Rearing | 69 | 1.4723 | (0.8897, 2.0681) | (-0.2768, 3.2153) | 19.5656 |
| Residual Tarsus Length (mm) | Egg Batch [mu] | 69 | 0.9404 | (0.6569, 1.2789) | (0.1586, 2.188) | Inf |
| Metabolic Slope | Intercept | 69 | 0.0115 | (2e-04, 0.0226) | (-0.0227, 0.0499) | 3.0241 |
| Metabolic Slope | Cold Rearing | 69 | -0.0184 | (-0.03, -0.0072) | (-0.0526, 0.0149) | 6.6118 |
| Metabolic Slope | Warm Rearing | 69 | -0.0080 | (-0.0197, 0.0039) | (-0.0428, 0.0267) | 2.0936 |
| Metabolic Slope | Body Mass (g) | 69 | -0.0005 | (-7e-04, -2e-04) | (-0.0013, 3e-04) | 6.8355 |
| Metabolic Slope | Residual Tarsus Length (mm) | 69 | 0.0042 | (0.0026, 0.0058) | (-5e-04, 0.0089) | 24.7235 |
| Metabolic Slope | Egg Batch [mu] | 69 | 0.0153 | (0.0083, 0.0237) | (9e-04, 0.0527) | Inf |

```
# save_kable(week3ResultsHeatResiduals,
# "../tables/pathAnalysisInHeat3WeeksResiduals.html")

caption <- paste0(
  "Results from a Bayesian path analysis ",
  "predicting metabolic responses to heat (metabolic slopes ",
  "[fold metabolism at thermoneutrality/",
  "°C] as a ",
  "function of body mass (g), relative tarsus length ",
  "(residuals; mm) and rearing conditions in eight ",
  "week old Japanese quail. Physiological measurements ",
```

```
  "are made at ambient temperatures between ",
  "30°C and ",
  "40°C. Cold rearing indicates ",
  "post-hatch rearing at 10°C, ",
  "relative to 20°C (intercept), ",
  "or 30°C ('warm rearing'). ",
  "CI indicates quantile ",
  "intervals and BF indicates Bayes Factors."
)

week8ResultsHeatResiduals <-
  as.data.frame(functionModel8WeeksHotResidual) %>%
  summarise_all(., .funs = median) %>%
  pivot_longer(everything(),
    names_to = "Parameter",
    values_to = "Estimate"
  ) %>%
  merge(., quantileCIs(functionModel8WeeksHotResidual, cis = c(50, 95)),
    by = "Parameter", all.x = TRUE
  ) %>%
  filter(grepl("b_|sd_", Parameter)) %>%
  rowwise() %>%
  mutate("BF" = ifelse(Estimate < 0,
    (2 * mean(as.data.frame(
      functionModel8WeeksHotResidual
    )[, Parameter] <= 0)) /
      (2 * mean(as.data.frame(
        functionModel8WeeksHotResidual
      )[, Parameter] >= 0)),
    (2 * mean(as.data.frame(
      functionModel8WeeksHotResidual
    )[, Parameter] >= 0)) /
      (2 * mean(as.data.frame(
        functionModel8WeeksHotResidual
      )[, Parameter] <= 0))
  )) %>%
  ungroup() %>%
  mutate(
    "Estimate" = round(Estimate, digits = 4),
    "BF" = round(BF, digits = 4),
    "N" = nrow(functionModel8WeeksHotResidual$data)
  ) %>%
  mutate("Parameter" = ifelse(grepl("b_", Parameter),
    gsub("b_", "", Parameter),
    gsub(
      "Intercept", "batch",
      gsub(".*__", "", Parameter)
    )
  )) %>%
  mutate(
    "Response" = gsub("_.*", "", Parameter),
    "Parameter" = gsub(".*_", "", Parameter)
  ) %>%
  merge(., tribble(
    ~Response, ~response, ~level,
    "mass", "Body Mass (g)", "A",
    "slope",
    "Metabolic Slope", "D",
    "residualTarsus", "Residual Tarsus Length (mm)", "B"
  ),
  by = "Response"
  ) %>%
  merge(., tribble(
    ~Parameter, ~parameter, ~number,
    "Intercept", "Intercept", "1",
    "mass", "Body Mass (g)", "4",
    "residualTarsus", "Residual Tarsus Length (mm)", "5",
```

```
    "pretreatmentA", "Cold Rearing", "2",
    "pretreatmentC", "Warm Rearing", "3",
    "batch", "Egg Batch [mu]", "6"
  ),
  by = "Parameter"
  ) %>%
  mutate(
    `50\\% HDI` = paste0("(", paste(
      round(Low_CI_50, digits = 4),
      round(High_CI_50, digits = 4),
      sep = ", "
    ), ")"),
    `95\\% HDI` = paste0("(", paste(
      round(Low_CI_95, digits = 4),
      round(High_CI_95, digits = 4),
      sep = ", "
    ), ")")
  ) %>%
  select(-c(Low_CI_50, High_CI_50, Low_CI_95, High_CI_95)) %>%
  select(
    "Response" = "response", "Parameter" = "parameter", N,
    Estimate, `50\\% HDI`, `95\\% HDI`, BF, level, number
  ) %>%
  arrange(level, number) %>%
  select(-c(level, number)) %>%
  kbl(.,
    longtable = T, booktabs = T, format = "latex", escape = FALSE,
    caption = caption
  ) %>%
  column_spec(column = c(1:2), width = "2.2cm") %>%
  column_spec(column = c(3:10), width = "1.9cm") %>%
  kable_styling(latex_options = "striped")

week8ResultsHeatResiduals
```

***Table 41:*** *Results from a Bayesian path analysis predicting metabolic responses to heat (metabolic slopes [fold metabolism at thermoneutrality/°C] as a function of body mass (g), relative tarsus length (residuals; mm) and rearing conditions in eight week old Japanese quail. Physiological measurements are made at ambient temperatures between 30°C and 40°C. Cold rearing indicates post-hatch rearing at 10°C, relative to 20°C (intercept), or 30°C ('warm rearing'). CI indicates quantile intervals and BF indicates Bayes Factors.*

| Response | Parameter | N | Estimate | 50% HDI | 95% HDI | BF |
|---|---|---|---|---|---|---|
| Body Mass (g) | Intercept | 84 | -5.6989 | (-9.4298, -2.0578) | (-16.9765, 5.332) | 5.7227 |
| Body Mass (g) | Cold Rearing | 84 | 6.0008 | (0.715, 11.0206) | (-9.7552, 21.3628) | 3.4370 |
| Body Mass (g) | Warm Rearing | 84 | 12.7077 | (7.3425, 17.9642) | (-2.8686, 27.9488) | 18.3237 |
| Body Mass (g) | Egg Batch [mu] | 84 | 0.2842 | (0.1172, 0.572) | (0.0105, 1.4781) | Inf |
| Residual Tarsus Length (mm) | Intercept | 84 | -0.5064 | (-0.8693, -0.1187) | (-1.6063, 0.8082) | 4.1613 |
| Residual Tarsus Length (mm) | Cold Rearing | 84 | 0.5440 | (0.0995, 0.9776) | (-0.7726, 1.7808) | 3.9020 |
| Residual Tarsus Length (mm) | Warm Rearing | 84 | 0.8544 | (0.4475, 1.264) | (-0.3495, 2.0594) | 11.4224 |
| Residual Tarsus Length (mm) | Egg Batch [mu] | 84 | 0.5603 | (0.3264, 0.8497) | (0.0342, 1.6656) | Inf |
| Metabolic Slope | Intercept | 84 | -0.0068 | (-0.016, 0.0029) | (-0.0368, 0.0283) | 2.1571 |
| Metabolic Slope | Cold Rearing | 84 | 0.0052 | (-0.0041, 0.0145) | (-0.0221, 0.0327) | 1.8269 |
| Metabolic Slope | Warm Rearing | 84 | 0.0122 | (0.0026, 0.0217) | (-0.0161, 0.0392) | 4.2049 |
| Metabolic Slope | Body Mass (g) | 84 | 0.0004 | (3e-04, 5e-04) | (0, 7e-04) | 77.4314 |

| | | | | | | |
|---|---|---|---|---|---|---|
| Metabolic Slope | Residual Tarsus Length (mm) | 84 | -0.0041 | (-0.0058, -0.0025) | (-0.0088, 8e-04) | 19.0000 |
| Metabolic Slope | Egg Batch [mu] | 84 | 0.0160 | (0.0098, 0.0243) | (0.0012, 0.0529) | Inf |

```
#save_kable(week8ResultsHeatResiduals,
# "../tables/pathAnalysisInHeat8WeeksResiduals.html")
```
