## Supplementary material for "No evidence that shrinking and shapeshifting meaningfully affect how birds respond to warming and cooling": Supp. 4

### Effect of Morphology on Evaporative Cooling in the Heat

#### Overview

In desert bird species, body mass has been shown to influence the rate at which individuals lose water to evaporation in the heat (McKechnie et al, 2021). Similarly, in male great tits (*Parus major*), tarsus surface area has been shown to correlated with evaporative cooling efficiency at temperatures above thermoneutrality (Playà-Montmany et al, 2021). In light of these findings, we sought to test whether alignment with Bergmann's and Allen's rules may benefit individuals by increasing their capacity to contend with metabolic heat production through evaporative cooling. Below, we collate measurements of evaporative water loss in developing (three weeks of age) and adult (eight weeks of age) Japanese quail, visualise them for evidence of oddities/errors, convert them to estimates of evaporative heat loss (described in the main text of Tabh et al, 2024), then combine them with resting metabolic measurements to estimate evaporative cooling efficiency per individual (here, represented as evaporative heat loss/resting metabolism, each in Watts). Once complete, we test whether: (1) increases in evaporative heat loss between thermoneutrality (30°C) and hot environments (40°C), and (2) evaporative cooling efficiency, vary by body mass and tarsus length in our quail.

We begin this document by loading in R packages and functions necessary for data collation, visualisation, and analysis. We then proceed to importing our data, inspecting it for oddities and errors, and using it to visualise raw trends. Last, models testing effects of morphology on evaporative heat loss and cooling efficiency are produced for evaluation.

```
def.chunk.hook <- knitr::knit_hooks$get("chunk")
knitr::knit_hooks$set(chunk = function(x, options) {
  x <- def.chunk.hook(x, options)
  ifelse(options$size != "normalsize",
    paste0("\n \\", options$size, "\n\n", x, "\n\n \\normalsize"), x
  )
})

knitr::opts_chunk$set(fig.pos = "H", out.extra = "")
knitr::opts_chunk$set(size = "footnotesize")

# First loading in packages

library("tidyverse")
library("easypackages")

packageList <- c("bayesplot", "brms", "brmsMethods",
                 "doParallel", "foreach", "ggpubr",
                 "kableExtra", "latex2exp", "patchwork",
                 "priorsense", "showtext", "tidybayes",
                 "wesanderson")
```

```
libraries(packageList)

caption <- paste0("R packages and their respective versions used for",
                  " data organisation and analysis in this study."
)

sapply(packageList, function(x) {
  y <- as.character(packageVersion(x))
  return(y)
}, simplify = FALSE) %>%
  enframe(., name = "Package", value = "Version") %>%
  as.data.frame(.) %>%
  kbl(.,
    longtable = T, booktabs = T,
    caption = caption
  ) %>%
  kable_styling(latex_options = "striped")
```

***Table 1:*** *R packages and their respective versions used for data organisation and analysis in this study.*

| Package | Version |
|---|---|
| bayesplot | 1.11.1 |
| brms | 2.20.4 |
| brmsMethods | 0.0.0.9000 |
| doParallel | 1.0.17 |
| foreach | 1.5.2 |
| ggpubr | 0.6.0 |
| kableExtra | 1.3.4 |
| latex2exp | 0.9.6 |
| patchwork | 1.2.0 |
| priorsense | 0.0.0.9000 |
| showtext | 0.9.6 |
| tidybayes | 3.0.4 |
| wesanderson | 0.3.6.9000 |

```
# Loading additional functions

pp_check2 <- function(model, resp = NA, ndraws = 500,
                      xlab = "label", colour = "lightblue") {
  require(brms)
  require(ggplot2)
  stopifnot("Model must be a brmsfit object" = is.brmsfit(model))

  if (is.na(resp)) {
    resp <- model$formula$resp
  }

  p1 <- brms::pp_check(model, ndraws = ndraws, resp = resp) +
    scale_colour_manual(
      values = c("black", colour),
      labels = c("y", "yhat"),
```

```
      name = NULL
    ) +
    xlab(xlab) +
    ylab("Density") +
    theme_classic()
  return(p1)
}

chainCheck <- function(model, rDig = 3) {
  require(brms)
  stopifnot("Model must be a brmsfit object" = is.brmsfit(model))

  Rhat <- paste0(
    "Rhat range: ",
    round(min(rhat(model)), digits = rDig),
    " - ",
    round(max(rhat(model)), digits = rDig)
  )
  Neff <- paste0(
    "Neff/N range: ",
    round(min(neff_ratio(model)), digits = rDig),
    " - ",
    round(max(neff_ratio(model)), digits = rDig)
  )
  cat(paste0(Rhat, "\n", Neff))
}

quantileCIs <- function(x, rnd = 3, cis = c(50, 95), sci_note = FALSE) {
  require(tidyverse)

  if (class(x)[1] != "brmsfit") {
    return("x must be a brmsfit object.")
  }
  if (length(cis) != 2) {
    return("cis must be a vector of integers with length 2")
  }

  prbs = c()
  nColNames = c()
  for (i in 1:length(cis)){
    prbs = c(prbs, c(0.5 - (cis[i]/100)/2, 0.5 + (cis[i]/100)/2))
    nColNames = c(nColNames,
                  paste0("Low_CI_", cis[i]),
                  paste0("High_CI_", cis[i])
    )
  }

  modelFrame = as.data.frame(x)

  Results <- apply(modelFrame, MARGIN = 2, FUN = quantile,
        probs = prbs, type = 8) %>%
    t() %>%
    as.data.frame() %>%
```

```
    rownames_to_column(var = "par") %>%
    `colnames<-`(c("Parameter", nColNames))

  if (sci_note == FALSE) {
    Results <- apply(modelFrame, MARGIN = 2, FUN = quantile,
        probs = prbs, type = 8) %>%
    t() %>%
    as.data.frame() %>%
    rownames_to_column(var = "par") %>%
    `colnames<-`(c("Parameter", nColNames))

  } else if (sci_note == TRUE) {
    Results <- apply(modelFrame, MARGIN = 2, FUN = quantile,
        probs = prbs, type = 8) %>%
    t() %>%
    as.data.frame() %>%
    rownames_to_column(var = "par") %>%
    `colnames<-`(c("Parameter", nColNames)) %>%
    mutate_at(.vars = vars(-Parameter),
              .funs = function(x){
                return(format(x, scientific = TRUE))
              }
    )
  }

  return(Results)
}

# Installing font

font_add_google(name = "Noto Sans", family = "Noto Sans")

# Setting our working directory

setwd(
  paste0("/Users/joshuatabh/Documents/",
           "researchProjects/lund/functionOfAllen/functionData"
         )
)
```

Here, our evaporative water loss, morphometry, and resting metabolism data are imported and bound. Collation and filtration of morphometric and metabolism data are described in our previous supplemental documents entitled "Data Organisation and Analysis of Growth" and "Metabolic Slope and Repeatability Analyses" respectively. Evaporative heat loss and evaporative cooling efficiency data are then visualised for potential errors using Cleveland dotplots.

```
# Morphometric data are loaded.

data <- read.csv("compiledData.csv")

# Next, metabolic data are loaded and filtered as previously described.

{
  vo2Data <- bind_rows(
    read.csv("exp1VO2.csv"),
```

```
    read.csv("exp2VO2.csv"),
    read.csv("exp3VO2.csv")
  ) %>%
    select(-c(startTime, endTime, initialTb, fileName)) %>%
    mutate(pretreatment = ifelse(pretreatment == "control",
      "neutral", pretreatment
    ))

  all <- merge(
    data %>%
      select(-Ta),
    vo2Data %>%
      distinct() %>%
      select(-c(pretreatment, posttreatment, birdID)),
    by = c("ring", "week", "exp"),
    all.x = TRUE
  ) %>%
    distinct()

  all <- all %>%
    select(
      ring, birdID, sex, exp, week, pretreatment,
      posttreatment, treatment, Ta, mass, wingLength,
      meanTb, tarsusLengthMean, tarsusLengthSD,
      tarsusHeightMean, tarsusHeightSD,
      tarsusCalibration, VO2, RMR
    )

  all <- all %>%
    mutate(Ta = ifelse(exp == "C" & Ta > 28 & Ta < 31, 30, Ta)) %>%
    mutate(Ta = ifelse(exp == "C" & Ta > 38 & Ta < 41, 40, Ta)) %>%
    filter(Ta %in% c(10, 20, 30, 40))

  all <- rbind(
    all %>%
      filter(ring == "B14" & week == "3") %>%
      group_by(Ta) %>%
      mutate(
        "tarsusLengthMean" = mean(tarsusLengthMean, na.rm = T),
        "tarsusHeightMean" = mean(tarsusHeightMean, na.rm = T)
      ) %>%
      mutate("n" = 1:2) %>%
      ungroup() %>%
      arrange(n, Ta) %>%
      slice(1:4) %>%
      select(-n),
    all %>%
      filter(!(ring == "B14" & week == "3"))
  ) %>%
    arrange(exp, week, ring, Ta)
}

# Eevaporative water lass data are now imported and bound with all other data.

vh2o = all %>%
  merge(.,
        bind_rows(
          read.csv("exp1VH2O.csv"),
          read.csv("exp2VH2O.csv"),
          read.csv("exp3VH2O.csv") %>%
            mutate(Ta = ifelse(Ta > 28 & Ta < 31, 30, Ta)) %>%
            mutate(Ta = ifelse(Ta > 38 & Ta < 41, 40, Ta)) %>%
            mutate(week = ifelse(week == 4, 3, 8))
        ), by =
          c("ring", "birdID", "exp", "week",
            "pretreatment", "posttreatment", "Ta"),
        all.x = TRUE
```

```
  ) %>% filter(Ta %in% c(10, 20, 30, 40)) %>%
  mutate(ehl = (VH2O*2.406)/60) %>%
  mutate(ecc = ehl/RMR)

# Will data loaded, we check the spread of evaporative heat loss measures by
# ambient temperature

ggplot(
  vh2o %>%
    filter(week %in% c(3, 8)) %>%
    group_by(week) %>%
    mutate("ID" = 1:n()) %>%
    ungroup() %>%
    mutate(
      "week" = paste0("Age = ", week, " weeks"),
      "Ta" = paste0(Ta, "°C")
    ),
  aes(x = ID, y = ehl)
) +
  facet_grid(week ~ Ta, scales = "free") +
  geom_point(size = 2, colour = "black",
             fill = "slateblue", alpha = 0.7) +
  geom_rect(
    data = vh2o %>%
      filter(week %in% c(3, 8)) %>%
      group_by(week, Ta) %>%
      summarise(
        "Mean" = mean(ehl, na.rm = T),
        "LCL" = Mean - 4 * sd(ehl, na.rm = T),
        "UCL" = Mean + 4 * sd(ehl, na.rm = T),
        "ID" = 1, "ehl" = 1
      ) %>%
      ungroup() %>%
      mutate(
        "week" = paste0("Age = ", week, " weeks"),
        "Ta" = paste0(Ta, "°C")
      ) %>%
      as.data.frame(),
    aes(xmin = -Inf, xmax = Inf, ymin = LCL, ymax = UCL),
    colour = "black", fill = "grey70", alpha = 0.5
  ) +
  theme_classic() +
  xlab("Sample Number") +
  ylab("Evaporative Heat Loss (W)")
```

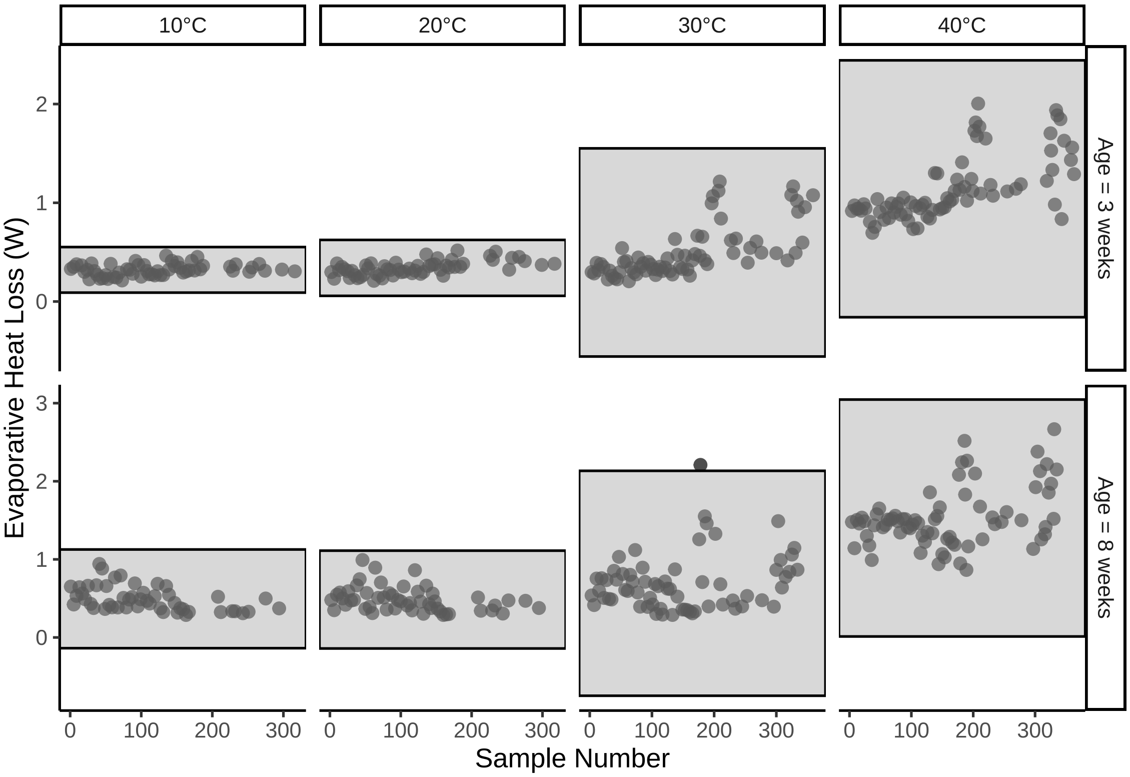


***Figure 1:*** *Cleveland dotplot of evaporative heat loss (W) by ambient temperature of collection, each drawn from adult Japanese quail. Dots represent raw data points. Grey rectangles indicate areas captured by means +/- 4 standard deviations.*

```
ggplot(
  vh2o %>%
    filter(week %in% c(3, 8)) %>%
    group_by(week) %>%
    mutate("ID" = 1:n()) %>%
    ungroup() %>%
    mutate(
      "week" = paste0("Age = ", week, " weeks"),
      "Ta" = paste0(Ta, "°C")
    ),
  aes(x = ID, y = ecc)
) +
  facet_grid(week ~ Ta, scales = "free") +
  geom_point(size = 2, colour = "black",
             fill = "slateblue", alpha = 0.7) +
  geom_rect(
    data = vh2o %>%
      filter(week %in% c(3, 8)) %>%
      group_by(week, Ta) %>%
      summarise(
        "Mean" = mean(ecc, na.rm = T),
        "LCL" = Mean - 4 * sd(ecc, na.rm = T),
        "UCL" = Mean + 4 * sd(ecc, na.rm = T),
        "ID" = 1, "ecc" = 1
      ) %>%
      ungroup() %>%
      mutate(
        "week" = paste0("Age = ", week, " weeks"),
        "Ta" = paste0(Ta, "°C")
      ) %>%
      as.data.frame(),
```

```
    aes(xmin = -Inf, xmax = Inf, ymin = LCL, ymax = UCL),
    colour = "black", fill = "grey70", alpha = 0.5
  ) +
  theme_classic() +
  xlab("Sample Number") +
  ylab("Evaporative Cooling Efficiency (EHL/RMR)")
```

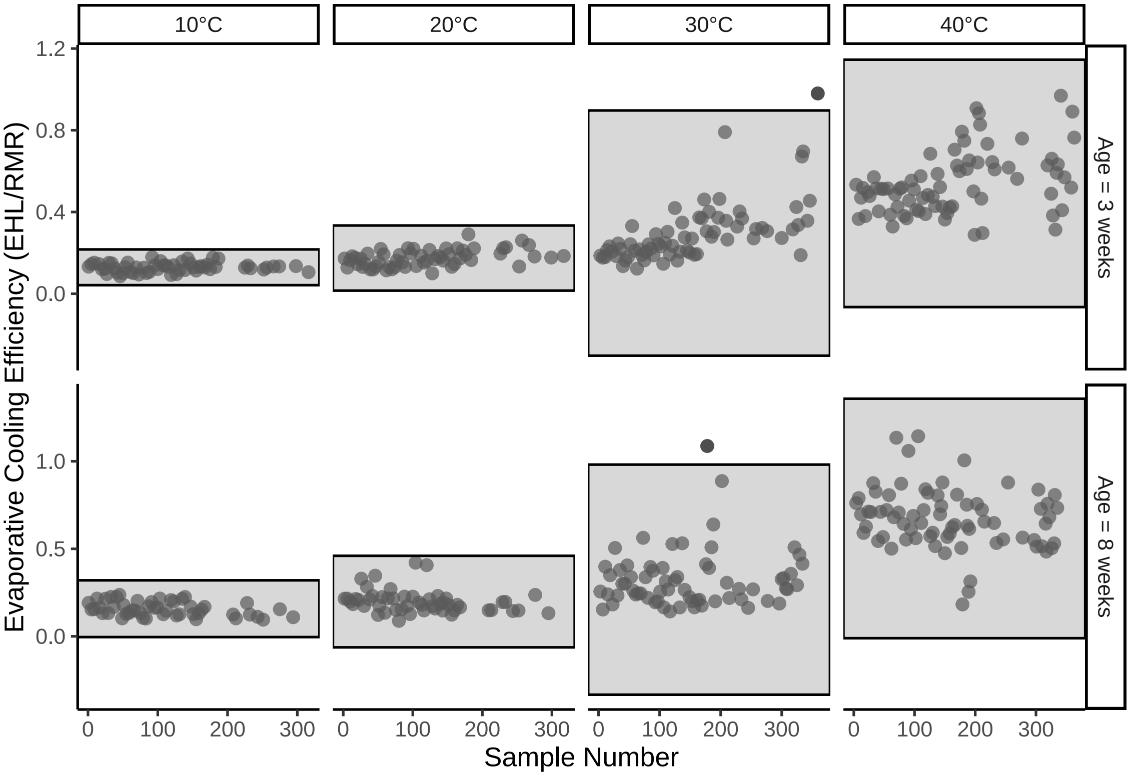


***Figure 2:*** *Cleveland dotplot of evaporative cooling efficiency (evaporative heat loss/resting metabolism, each in Watts) by ambient temperature of collection, each drawn from adult Japanese quail. Dots represent raw data points. Grey rectangles indicate areas captured by means +/- 4 standard deviations.*

Some measurements of evaporative heat loss and evaporative cooling efficiency are notably high at 30°C, however, these values still fit within the distribution of biologically realistic measurements (i.e. by comparison with measurements drawn at 40°C). For this reason, these measurements are retained and we proceed to visualing raw trends in our data.

```
p1 <- vh2o %>%
  filter(week %in% c(3,8)) %>%
  mutate(week = paste0(week, " Weeks")) %>%
  ggplot(aes(x = Ta, y = ehl)) +
  facet_wrap(~week) +
  geom_line(aes(group = ring), colour = "grey50", alpha = 0.3) +
  geom_point(pch = 21, size = 1.5, colour = "black", fill = "grey50",
             alpha = 0.3) +
  stat_summary(geom = "line", fun = "mean", colour = "black") +
  stat_summary(geom = "errorbar", fun.data = "mean_cl_boot", width = 2,
               colour = "black") +
  stat_summary(geom = "point", fun = "mean", pch = 21, size = 4,
               colour = "black", fill = "lightblue3") +
  xlab("Ambient Temperature (°C)") +
  ylab("Evaporative Heat\nLoss (W)") +
  theme_classic()
```

```
p2 <- vh2o %>%
  filter(week %in% c(3,8)) %>%
  mutate(week = paste0(week, " Weeks")) %>%
  ggplot(aes(x = Ta, y = ecc)) +
  facet_wrap(~week) +
  geom_line(aes(group = ring), colour = "grey50", alpha = 0.3) +
  geom_point(pch = 21, size = 1.5, colour = "black", fill = "grey50",
             alpha = 0.3) +
  stat_summary(geom = "line", fun = "mean", colour = "black") +
  stat_summary(geom = "errorbar", fun.data = "mean_cl_boot", width = 2,
               colour = "black") +
  stat_summary(geom = "point", fun = "mean", pch = 21, size = 4,
               colour = "black", fill = "lightblue3") +
  xlab("Ambient Temperature (°C)") +
  ylab("Evaporative Cooling\nEfficiency (EHL/RMR)") +
  theme_classic()

p1/p2 + plot_annotation(tag_levels = "A")
```

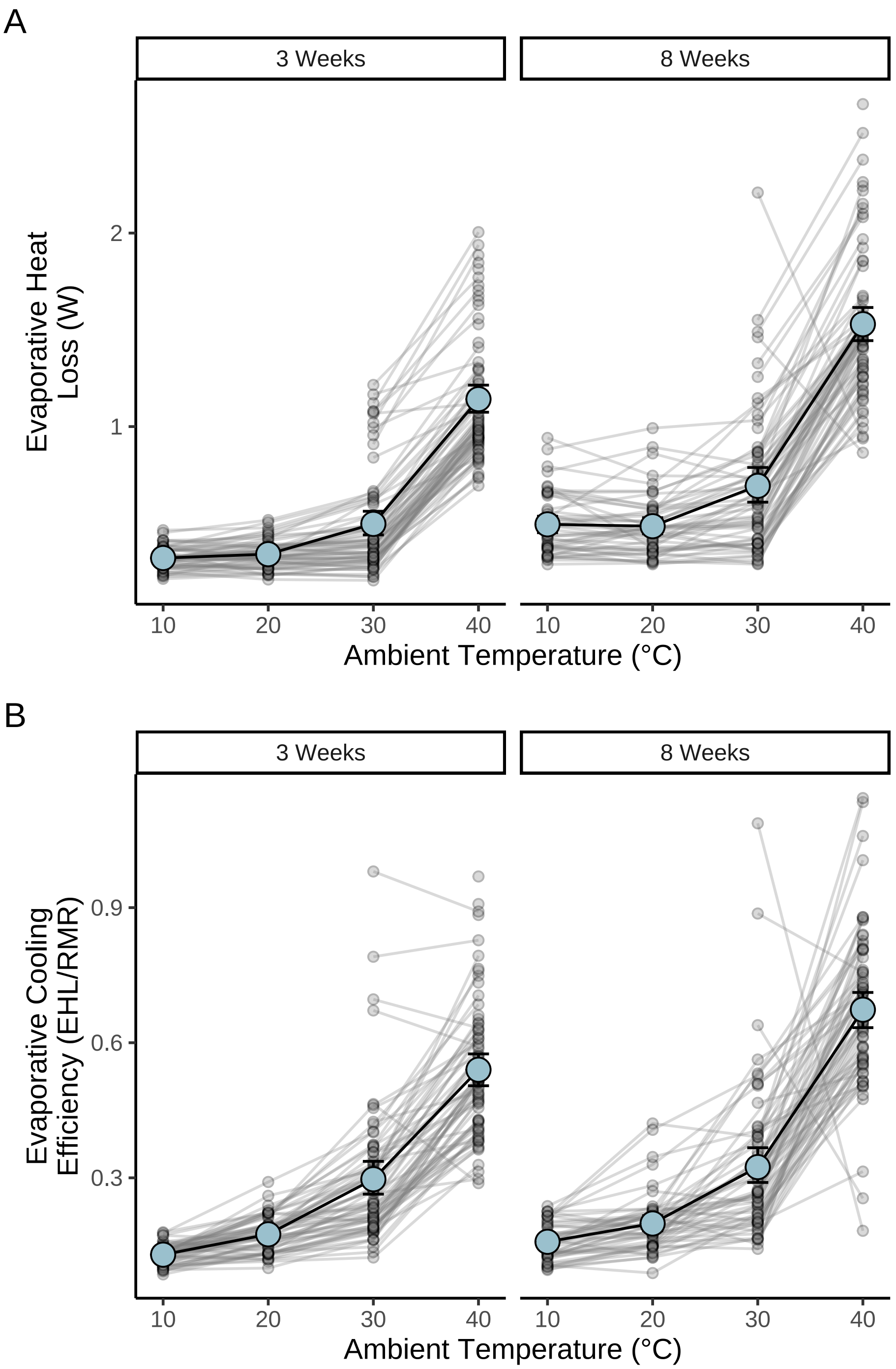


***Figure 3:*** *Effect of ambient temperature (°C) on evaporative heat production and cooling efficiency in developing (3 weeks) and adult (8 weeks) Japanese quail. Evaporative cooling efficiency represents the ratio of evaporative heat loss (W) to metabolic heat production (W) at a given temperature. Large dots represent means and errorbars indicate +/- one standard error. Small dots represent raw values and lines connect measurements drawn from the same individual.*

Next, we explore raw correlations between morphology (i.e. body mass and tarsus length) and both evaporative heat loss and evaporative cooling efficiency at 40°C. Raw effects of rearing condition on evaporative heat loss and cooling efficiency are also visualised.

```r
p1 <- vh2o %>%
  filter(week %in% c(3,8) & Ta == 40) %>%
  mutate(week = paste0(week, " Weeks")) %>%
  ggplot(aes(x = mass, y = ehl)) +
  facet_wrap(~week, scales = "free") +
  geom_point(pch = 21, size = 2, colour = "black", fill = "grey50",
             alpha = 0.3) +
  xlab("Body Mass (g)") +
  ylab("Evaporative Heat\nLoss (W)") +
  theme_classic()

p2 <- vh2o %>%
  filter(week %in% c(3,8) & Ta == 40) %>%
  mutate(week = paste0(week, " Weeks")) %>%
  ggplot(aes(x = mass, y = ecc)) +
  facet_wrap(~week, scales = "free") +
  geom_point(pch = 21, size = 2, colour = "black", fill = "grey50",
             alpha = 0.3) +
  xlab("Body Mass (g)") +
  ylab("Evaporative Cooling\nEfficiency (EHL/RMR)") +
  theme_classic()

p1/p2
```

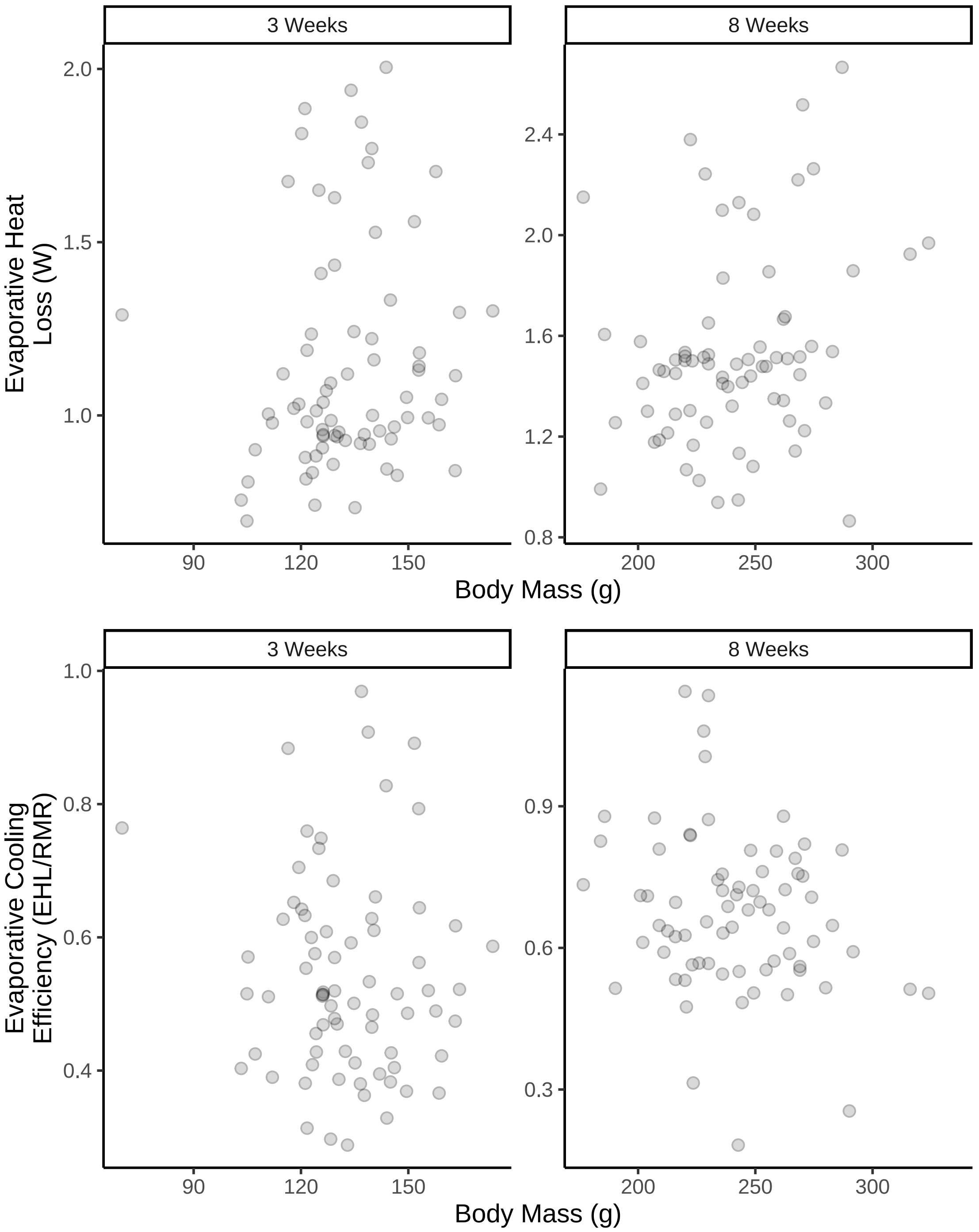


***Figure 4:*** *Evaporative heat loss and cooling efficiency as a function of body mass in developing (3 weeks) and adult (8 weeks) Japanese quail (n = 82). Small dots represent raw values. Quail were reared at either 10°C, 20°C, or 30°C until at least three weeks of age.*

```
p1 <- vh2o %>%
  filter(week %in% c(3,8) & Ta == 40) %>%
  mutate(week = paste0(week, " Weeks")) %>%
  ggplot(aes(x = tarsusLengthMean, y = ehl)) +
  facet_wrap(~week, scales = "free") +
  geom_point(pch = 21, size = 2, colour = "black", fill = "grey50",
             alpha = 0.3) +
```

```
  xlab("Tarsus Length (mm)") +
  ylab("Evaporative Heat Loss (W)") +
  theme_classic()

p2 <- vh2o %>%
  filter(week %in% c(3,8) & Ta == 40) %>%
  mutate(week = paste0(week, " Weeks")) %>%
  ggplot(aes(x = tarsusLengthMean, y = ecc)) +
  facet_wrap(~week, scales = "free") +
  geom_point(pch = 21, size = 2, colour = "black", fill = "grey50",
             alpha = 0.3) +
  xlab("Tarsus Length (mm)") +
  ylab("Evaporative Cooling\nEfficiency (EHL/RMR)") +
  theme_classic()

p1/p2
```

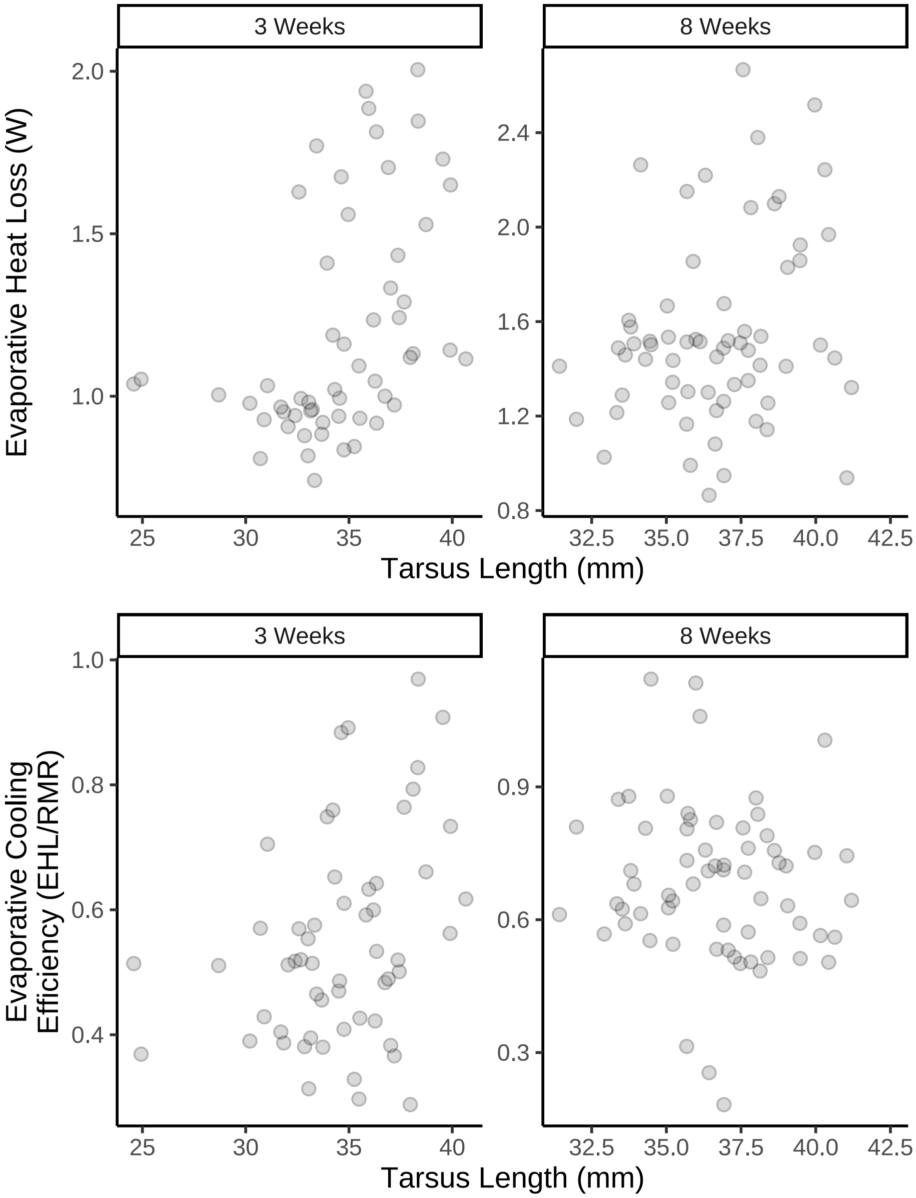


***Figure 5:*** *Evaporative heat loss and cooling efficiency as a function of tarsus length (mm) in developing (3 weeks) and adult (8 weeks) Japanese quail (n = 82). Small dots represent raw values. Quail were reared at either 10°C, 20°C, or 30°C until at least three weeks of age.*

```
p1 <- vh2o %>%
  filter(week %in% c(3, 8) & Ta == 40) %>%
  mutate(week = paste0(week, " Weeks")) %>%
  ggplot(aes(x = ehl, fill = pretreatment)) +
  facet_wrap(~week, scales = "free") +
  geom_density(alpha = 0.3) +
  xlab("Evaporative Heat Loss (W)") +
  ylab("Density") +
  scale_fill_manual(
    values = c("#7BB4E3", "black", "#CD5C5C"),
    name = "Rearing\nTreatment",
    labels = c(
      "Cold (10°C)",
      "Mild (20°C)",
```

```
      "Warm (30°C)"
    )
  ) +
  theme_classic() +
  theme(legend.position = "bottom")

p2 <- vh2o %>%
  filter(week %in% c(3, 8) & Ta == 40) %>%
  mutate(week = paste0(week, " Weeks")) %>%
  ggplot(aes(x = ecc, fill = pretreatment)) +
  facet_wrap(~week, scales = "free") +
  geom_density(alpha = 0.3) +
  xlab("Evaporative Cooling Efficiency (EHL/RMR)") +
  ylab("Density") +
  scale_fill_manual(
    values = c("#7BB4E3", "black", "#CD5C5C"),
    name = "Rearing\nTreatment",
    labels = c(
      "Cold (10°C)",
      "Mild (20°C)",
      "Warm (30°C)"
    )
  ) +
  theme_classic() +
  theme(legend.position = "bottom")

p1 / p2
```

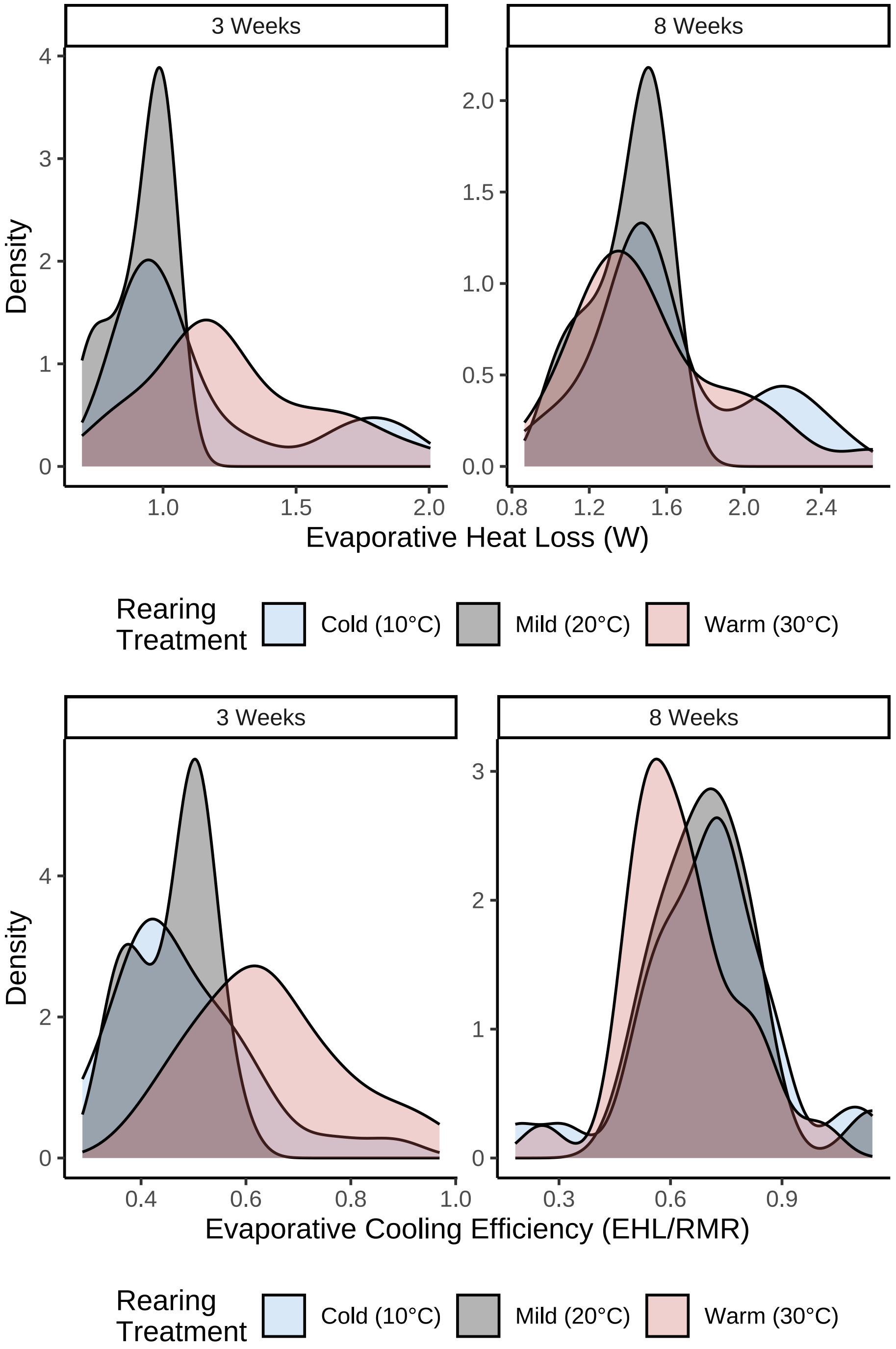


***Figure 6:*** *Density of evaporative heat loss and cooling efficiency measurements among Japanese quail reared at different temperatures (10°C, 20°C, or 30°C) until at least 3 weeks of age (n = 82). Measurements were taken at 40°C.*

Variance in body evaporative heat loss (W) and evaporative cooling efficiency may differ subtly among rearing treatments. Model residuals are therefore scrutinised below to assess whether remaining variance is indeed treatment-dependent. Last, we check whether variance in each metric differs between experimental batch 3 and experimental batches 1 and 2 owing to increased flow rates at this time.

```r
p1 <- vh2o %>%
  rename("batch" = exp) %>%
  filter(week %in% c(3, 8) & Ta == 40) %>%
  mutate(week = paste0(week, " Weeks")) %>%
  ggplot(aes(x = ehl, fill = pretreatment)) +
  facet_wrap(~week, scales = "free") +
  geom_density(alpha = 0.3) +
  xlab("Evaporative Heat Loss (W)") +
  ylab("Density") +
  scale_fill_manual(
    values = c("#7BB4E3", "black", "#CD5C5C"),
    name = "Experimental\nBatch",
    labels = c(
      "1",
      "2",
      "3"
    )
  ) +
  theme_classic() +
  theme(legend.position = "bottom")

p2 <- vh2o %>%
  rename("batch" = exp) %>%
  filter(week %in% c(3, 8) & Ta == 40) %>%
  mutate(week = paste0(week, " Weeks")) %>%
  ggplot(aes(x = ecc, fill = pretreatment)) +
  facet_wrap(~week, scales = "free") +
  geom_density(alpha = 0.3) +
  xlab("Evaporative Cooling Efficiency (EHL/RMR)") +
  ylab("Density") +
  scale_fill_manual(
    values = c("#7BB4E3", "black", "#CD5C5C"),
    name = "Experimental\nBatch",
    labels = c(
      "1",
      "2",
      "3"
    )
  ) +
  theme_classic() +
  theme(legend.position = "bottom")

p1 / p2
```

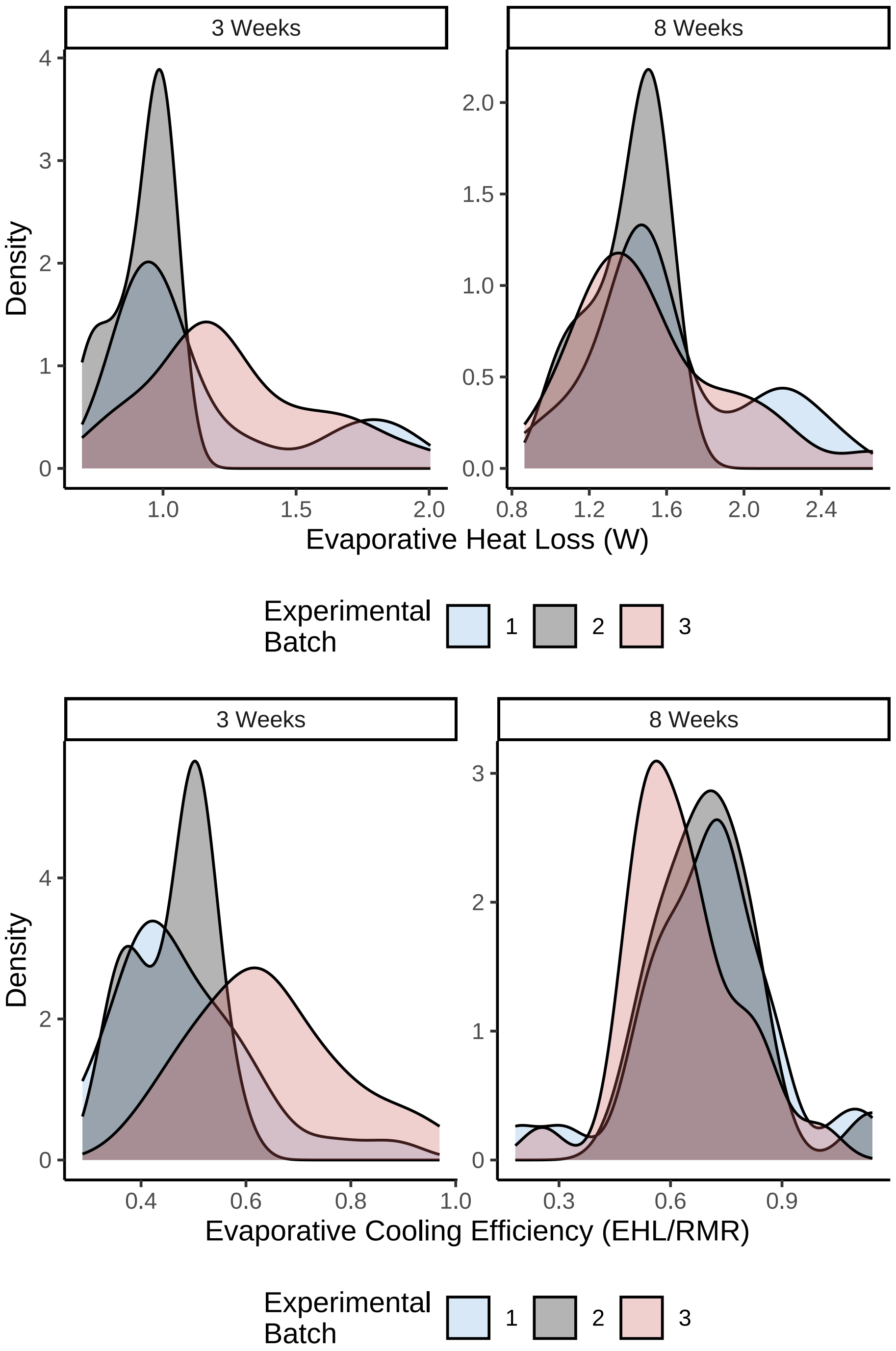


***Figure 7:*** *Density of evaporative heat loss and cooling efficiency measurements among Japanese quail derived from three different egg (or experimental) batches. Measurements were taken at 40°C.*

Variance appears particular large among experimental batch 3 for measures of evaporative cooling efficiency at three weeks of age. Given this, and that evaporative cooling efficiency was calculated using resting metabolism values which also appeared to vary considerably in experimental batch 3 at this age (refer to supplemental file 2), our error term for our model predicting evaporative cooling efficiency during development (below) is permitted to vary by experimental batch.

### Testing effects of morphology and thermal history on evaporative cooling

#### Effects in developing individuals

In our study (Tabh et al, 2024) we show that rearing temperature influences developmental trajectories and mature morphology of Japanese quail. However, in a previous study, we have also shown that rearing temperatures can directly effect evaporative heat loss and cooling capacities in this species (Persson et al, 2024). As such, to evaluate direct effects of morphology on evaporative heat loss responses and cooling efficiency while controlling for known effects of rearing conditions on each, we constructed two Bayesian path analyses depicted in the figure below.

```
data.frame(x = c(-5:5), y = c(-5:5)) %>%
  ggplot(aes(x = x, y = y)) +
  annotate("text", x = 0, y = 5, label = paste0("Developmental, ",
           "Thermal\nEnvironment"), colour = "black") +
  annotate("text", x = -2.5, y = 2.5, label = "Body Mass (g)",
           colour = "black") +
  annotate("text", x = 2.5, y = 2.5, label = "Appendage Length\n(mm)",
           colour = "black") +
  annotate("text", x = 0, y = -0.4,
           label = paste0("[1] Evaporative Heat Loss\n",
           "Response OR\n",
           "[2] Evaporative Cooling\n",
           "Efficiency"),
           colour = "black") +
  geom_segment(
    lineend = "round", linejoin = "round",
    size = 0.3, linetype = "solid", colour = "grey10",
    aes(x = -0.25, y = 4.6, xend = -2.5, yend = 2.75),
    arrow = arrow(length = unit(0.2, "cm"))
  ) +
  geom_segment(
    lineend = "round", linejoin = "round",
    size = 0.3, linetype = "solid", colour = "grey10",
    aes(x = 0.25, y = 4.6, xend = 2.5, yend = 2.75),
    arrow = arrow(length = unit(0.2, "cm"))
  ) +
  geom_segment(
    lineend = "round", linejoin = "round",
    size = 0.3, linetype = "solid", colour = "grey10",
    aes(x = -1.25, y = 2.5, xend = 1, yend = 2.5),
    arrow = arrow(length = unit(0.2, "cm"))
  ) +
  geom_segment(
    lineend = "round", linejoin = "round",
    size = 0.3, linetype = "dotted", colour = "grey10",
    aes(x = -2.5, y = 2.25, xend = -0.25, yend = 0.25),
    arrow = arrow(length = unit(0.2, "cm"))
  ) +
  geom_segment(
    lineend = "round", linejoin = "round",
    size = 0.3, linetype = "longdash", colour = "grey10",
    aes(x = 2.5, y = 2.25, xend = 0.25, yend = 0.25),
    arrow = arrow(length = unit(0.2, "cm"))
  ) +
  geom_curve(
    lineend = "round",
    size = 0.3, linetype = "solid", colour = "grey10",
    aes(x = -1.75, y = 4.75, xend = -1.75, yend = 0.25),
    arrow = arrow(length = unit(0.2, "cm"))
  ) +
  xlim(c(-6, 6)) +
  ylim(c(-0.7, 6)) +
  theme_void()
```

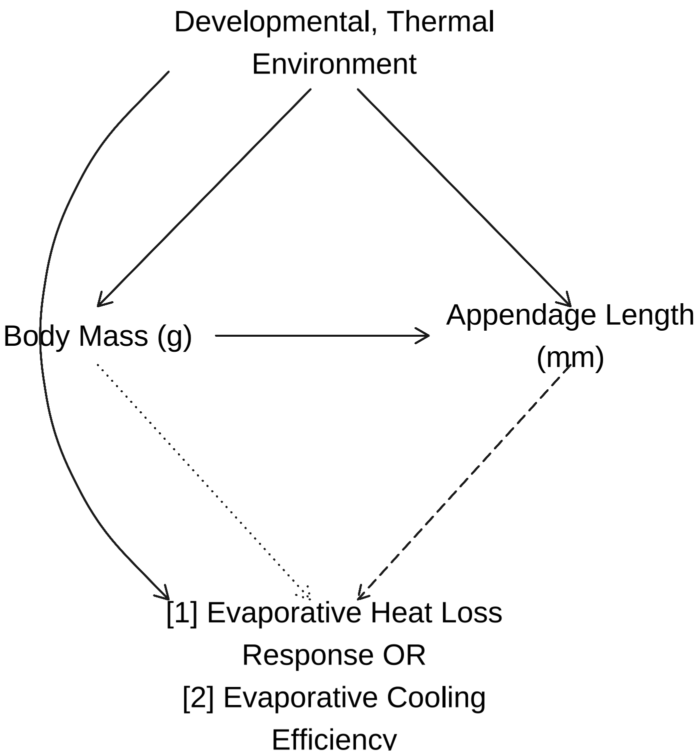


***Figure 8:*** *Flow-chart describing expected effects of developmental, thermal environment on morphology, and subsequently, thermal physiology of Japanese quail. Dashed and dotted lines represent effects predicted under a adaptive hypotheses of Allen's rule and Bergmann's rule respectively.*

Similar to our previous analyses (described in our supplement entitled: "Metabolic Slope and Repeatability Analyses"), equations within these path analyses were as follows:

$$Body\ Mass_j \sim \beta_{a0} + \beta_{a1} \cdot Cold\ Rearing_j + \beta_{a2} \cdot Warm\ Rearing_j + \mu_{0a} + \epsilon_a$$

$$Tarsus\ Length_j \sim \beta_{b0} + \beta_{b1} \cdot Cold\ Rearing_j + \beta_{b2} \cdot Warm\ Rearing_j + \beta_{b3} \cdot Body\ Mass_j + \mu_{0b} + \epsilon_b$$

and either:

$$Fold\ Evaporative\ Heat\ Loss_j \sim \beta_{c0} + \beta_{c1} \cdot Cold\ Rearing_j + \beta_{c2} \cdot Warm\ Rearing_j + \beta_{c3} \cdot Body\ Mass_j + \beta_{c4} \cdot Tarsus\ Length_j + \mu_{0c} + \epsilon_c$$

where:

$$Fold\ Evaporative\ Heat\ Loss_j = \frac{Evaporative\ Heat\ Loss_{40°\mathrm{C}j}}{Evaporative\ Heat\ Loss_{30°\mathrm{C}j}}$$

or:

$$Evaporative\ Cooling\ Efficiency_j \sim \beta_{c0} + \beta_{c1} \cdot Cold\ Rearing_j + \beta_{c2} \cdot Warm\ Rearing_j + \beta_{c3} \cdot Body\ Mass_j + \beta_{c4} \cdot Tarsus\ Length_j + \mu_{0c} + \epsilon_{cj}$$

where evaporative cooling efficiency is measured as:

$$Evaporative\ Cooling\ Efficiency_j = \frac{Evaporative\ Heat\ Loss_{40°\mathrm{C}j}}{Metabolic\ Heat\ Production_{40°\mathrm{C}j}}$$

and:

$$\epsilon_{cj} \sim e^{(\tau_0 + \tau_1 \cdot Batch_{2j} + \tau_2 \cdot Batch_{3j})}$$

In all cases, $j$ represents individual (and thus, observation) identity, "warm rearing" and "cold rearing" represent binomial parameters with "0" corresponding to "false" and "1" corresponding to "true", $\mu_{0a}$ - $\mu_{0c}$ represent group-level intercepts of egg batch on each response variable, and body mass and tarsus length are mean-centred to improve interpretability of model intercepts($\beta_0$ terms). For models predicting body mass and tarsus length, $\epsilon$ is assumed to be normally distributed and centred at 0. For model 3 (predicting evaproative cooling efficiency) epsilon is allowed to vary by experimental batch. There, $\tau_0$ indicates the natural log-transformed error term for experimental batch 1, $\tau_1$ indicates the change in error associated with measures derived from experimental batch 2, and $\tau_2$ indicates the change in error associated with measures derived from experimental batch 3; $Batch_{2j}$ and $Batch_{3j}$ indicate binomial variables pertaining to whether a slope from individual $j$ was derived from experimental batch 2 or 3 respectively (0 = no; 1 = yes). Last, residuals are assumed to be uncorrelated across models. All measurement values refer to those collected during development (3 weeks of age).

For our first two equations (i.e. those predicting mean-centred body mass and tarsus length), priors were as follows (regardless of which path analyses they were encompassed within):

$$\beta_{a0} \sim \mathcal{N}(0, 5)$$
$$\beta_{a1} \sim \mathcal{N}(0, 15)$$
$$\beta_{a2} \sim \mathcal{N}(0, 15)$$
$$\mu_{0a} \sim \exp(2.5)$$
$$\epsilon_a \sim \exp(0.15)$$

$$\beta_{b0} \sim \mathcal{N}(0, 2.5)$$
$$\beta_{b1} \sim \mathcal{N}(0, 2.5)$$
$$\beta_{b2} \sim \mathcal{N}(0, 2.5)$$
$$\beta_{b3} \sim S\mathcal{N}(0, 0.25, 5)$$
$$\mu_{0b} \sim \exp(2)$$
$$\epsilon_b \sim \exp(1)$$

Development and justification of these priors is described in our previous supplement ( Metabolic Slope and Repeatability Analyses"). For our equation predicting evaporative heat loss responses at 40°C, priors were informed from Persson et al (2024) and were as follows:

$$\beta_{c0} \sim \mathcal{N}(2.5, 1)$$
$$\beta_{c1} \sim \mathcal{N}(0, 0.5)$$
$$\beta_{c2} \sim \mathcal{N}(0, 0.5)$$

$$\beta_{c3} \sim \mathcal{N}(0, 0.025)$$
$$\beta_{c4} \sim \mathcal{N}(0, 0.05)$$
$$\mu_{0c} \sim \exp(10)$$
$$\epsilon_c \sim \exp(5)$$

Those for our equation predicting evaporative cooling efficiency were also informed by Persson et al (2024) and were as follows:

$$\beta_{c0} \sim \mathcal{N}(0.5, 0.2)$$
$$\beta_{c1} \sim \mathcal{N}(0, 0.25)$$
$$\beta_{c2} \sim \mathcal{N}(0, 0.25)$$
$$\beta_{c3} \sim \mathcal{N}(0, 0.01)$$
$$\beta_{c4} \sim \mathcal{N}(0, 0.1)$$
$$\mu_{0c} \sim \exp(15)$$
$$\tau_0 \sim \mathcal{N}(-2, 1)$$
$$\tau_1 \sim \mathcal{N}(0, 0.5)$$
$$\tau_2 \sim \mathcal{N}(1, 1)$$

To ensure suitability of our priors, we first draw posterior predictions from prior distributions alone. These predictions are then compared against true data distributions (i.e. as a "prior predictive check").

```
ehlModel3WeeksPPCheck <- brm(
  data = vh2o %>%
    filter(week == "3" & Ta %in% c(30, 40)) %>%
    select(Ta, ring, pretreatment,
           mass, tarsusLengthMean, ehl,
           "batch" = exp) %>%
    pivot_wider(
      id_cols = c("ring", "batch",
                  "pretreatment", "mass",
                  "tarsusLengthMean"),
      values_from = "ehl",
      names_from = "Ta"
    ) %>%
    mutate("foldEhl" = `40` / `30`) %>%
    mutate(pretreatment = ifelse(pretreatment == "neutral", "B",
      ifelse(pretreatment == "cold", "A", "C")
    )) %>%
    mutate(pretreatment = factor(pretreatment,
      levels = c("B", "A", "C")
    )) %>%
    distinct() %>%
    mutate(
      mass = mass - mean(mass, na.rm = T),
      tarsus = tarsusLengthMean - mean(tarsusLengthMean, na.rm = T)
    ),
  family = "gaussian",
  bf(mass ~ pretreatment + (1 | batch)) +
    bf(tarsus ~ mass + pretreatment + (1 | batch)) +
    bf(foldEhl ~ tarsus + mass + pretreatment + (1 | batch)) +
```

```
    set_rescor(FALSE),
  prior = c(
    set_prior("normal(0, 5)",
      class = "Intercept",
      resp = "mass"
    ),
    set_prior("normal(0, 15)",
      class = "b",
      coef = "pretreatmentA",
      resp = "mass"
    ),
    set_prior("normal(0, 15)",
      class = "b",
      coef = "pretreatmentC",
      resp = "mass"
    ),
    set_prior("exponential(2.5)",
      class = "sd",
      group = "batch",
      resp = "mass"
    ),
    set_prior("exponential(0.15)",
      class = "sigma",
      resp = "mass"
    ),
    set_prior("normal(0, 2.5)",
      class = "Intercept",
      resp = "tarsus"
    ),
    set_prior("normal(0, 2.5)",
      class = "b",
      coef = "pretreatmentA",
      resp = "tarsus"
    ),
    set_prior("normal(0, 2.5)",
      class = "b",
      coef = "pretreatmentC",
      resp = "tarsus"
    ),
    set_prior("skew_normal(0, 0.25, 5)",
      class = "b",
      coef = "mass",
      resp = "tarsus"
    ),
    set_prior("exponential(2)",
      class = "sd",
      group = "batch",
      resp = "tarsus"
    ),
    set_prior("exponential(1)",
      class = "sigma",
      resp = "tarsus"
    ),
    set_prior("normal(2.5, 1)",
      class = "Intercept",
      resp = "foldEhl"
    ),
    set_prior("normal(0, 0.5)",
      class = "b",
      coef = "pretreatmentA",
      resp = "foldEhl"
    ),
    set_prior("normal(0, 0.5)",
      class = "b",
      coef = "pretreatmentC",
      resp = "foldEhl"
    ),
```

```
    set_prior("normal(0, 0.025)",
      class = "b",
      coef = "mass",
      resp = "foldEhl"
    ),
    set_prior("normal(0, 0.5)",
      class = "b",
      coef = "tarsus",
      resp = "foldEhl"
    ),
    set_prior("exponential(10)",
      class = "sd",
      group = "batch",
      resp = "foldEhl"
    ),
    set_prior("exponential(5)",
      class = "sigma",
      resp = "foldEhl"
    )
  ),
  iter = 50000, warmup = 10000, cores = 4, chains = 4, thin = 20,
  control = list(adapt_delta = .97, max_treedepth = 14),
  sample_prior = "only",
  silent = TRUE, refresh = 0,
  file = "./models/heatLossModel3WeeksPPCheck.Rds",
)

pp1 <- pp_check2(ehlModel3WeeksPPCheck, resp = "foldEhl",
                 xlab = "Evaporative Heat\nLoss Response (Fold From 30°C)") +
  theme(legend.position = "none")

efficiencyModel3WeeksPPCheck <- brm(
  data = vh2o %>%
    filter(week == "3" & Ta == 40) %>%
    select(Ta, ring, pretreatment,
      mass, tarsusLengthMean, ecc,
      "batch" = exp
    ) %>%
    mutate(
      pretreatment =
        ifelse(pretreatment == "neutral", "B",
          ifelse(pretreatment == "cold", "A", "C")
        )
    ) %>%
    mutate(pretreatment = factor(pretreatment,
      levels = c("B", "A", "C")
    )) %>%
    distinct() %>%
    mutate(
      mass = mass - mean(mass, na.rm = T),
      tarsus = tarsusLengthMean -
        mean(tarsusLengthMean, na.rm = T)
    ),
  family = "gaussian",
  bf(mass ~ pretreatment + (1 | batch)) +
    bf(tarsus ~ mass + pretreatment + (1 | batch)) +
    bf(
      ecc ~ tarsus + mass + pretreatment + (1 | batch),
      sigma ~ batch
    ) +
    set_rescor(FALSE),
  prior = c(
    set_prior("normal(0, 5)",
      class = "Intercept",
      resp = "mass"
    ),
    set_prior("normal(0, 15)",
```

```
      class = "b",
      coef = "pretreatmentA",
      resp = "mass"
    ),
    set_prior("normal(0, 15)",
      class = "b",
      coef = "pretreatmentC",
      resp = "mass"
    ),
    set_prior("exponential(2.5)",
      class = "sd",
      group = "batch",
      resp = "mass"
    ),
    set_prior("exponential(0.15)",
      class = "sigma",
      resp = "mass"
    ),
    set_prior("normal(0, 2.5)",
      class = "Intercept",
      resp = "tarsus"
    ),
    set_prior("normal(0, 2.5)",
      class = "b",
      coef = "pretreatmentA",
      resp = "tarsus"
    ),
    set_prior("normal(0, 2.5)",
      class = "b",
      coef = "pretreatmentC",
      resp = "tarsus"
    ),
    set_prior("skew_normal(0, 0.25, 5)",
      class = "b",
      coef = "mass",
      resp = "tarsus"
    ),
    set_prior("exponential(2)",
      class = "sd",
      group = "batch",
      resp = "tarsus"
    ),
    set_prior("exponential(1)",
      class = "sigma",
      resp = "tarsus"
    ),
    set_prior("normal(0.5, 0.2)",
      class = "Intercept",
      resp = "ecc"
    ),
    set_prior("normal(0, 0.25)",
      class = "b",
      coef = "pretreatmentA",
      resp = "ecc"
    ),
    set_prior("normal(0, 0.25)",
      class = "b",
      coef = "pretreatmentC",
      resp = "ecc"
    ),
    set_prior("normal(0, 0.01)",
      class = "b",
      coef = "mass",
      resp = "ecc"
    ),
    set_prior("normal(0, 0.1)",
      class = "b",
```

```r
      coef = "tarsus",
      resp = "ecc"
    ),
    set_prior("exponential(15)",
      class = "sd",
      group = "batch",
      resp = "ecc"
    ),
    set_prior("normal(-2, 1)",
      dpar = "sigma",
      class = "Intercept",
      resp = "ecc"
    ),
    set_prior("normal(0, 0.5)",
      dpar = "sigma",
      class = "b",
      coef = "batchB",
      resp = "ecc"
    ),
    set_prior("normal(1, 1)",
      dpar = "sigma",
      class = "b",
      coef = "batchC",
      resp = "ecc"
    )
  ),
  iter = 50000, warmup = 10000, cores = 4, chains = 4, thin = 20,
  control = list(adapt_delta = .97, max_treedepth = 14),
  silent = TRUE, refresh = 0,
  sample_prior = "only",
  file = "./models/efficiencyModel3WeeksPPCheck.Rds"
)

pp2 <- pp_check2(efficiencyModel3WeeksPPCheck, resp = "ecc",
                 xlab = "Evaporative Cooling\nEfficiency (EHL/RMR)")

pp1 + pp2 + plot_annotation(tag_levels = "A")
```

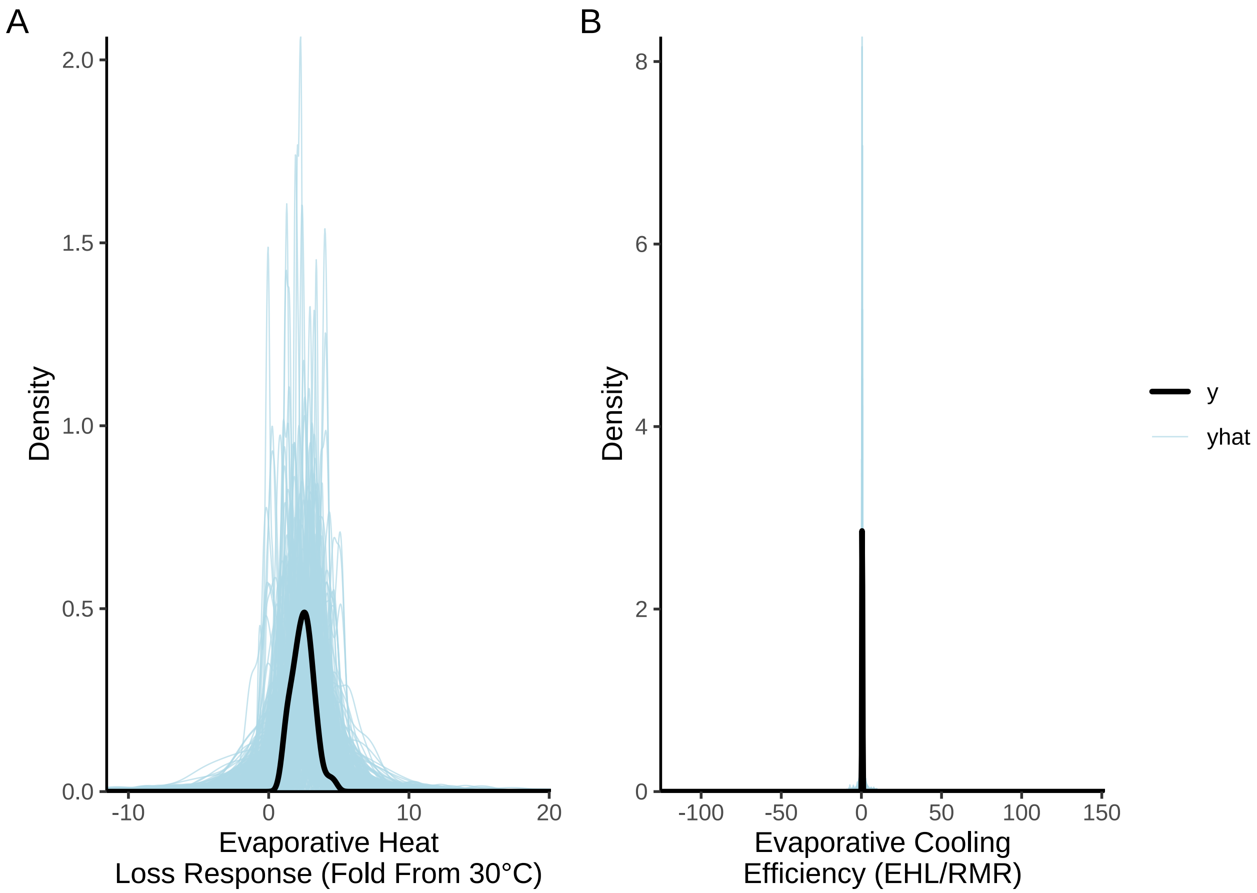


***Figure 9:*** *Prior predictive checks for two Bayesian path analyses ultimately predicting evaporative heat loss responses ('EHL'; fold from that observed at 30°C in W) and evaporative cooling efficiency ('ECE'; the ratio of evaporative heat-loss to metabolic heat production, each in W) in three week old Japanese quail. Black lines represent true EHL and ECE densities while blue lines represent densities estimated from model priors alone.*

For both analyses, priors are evidently suitable and unlikely to be constraining. We therefore proceed with constructing our complete path analyses below. After construction, we: (1) check for evidence of adequate chain mixing (i.e. a Gelman-Rubin statistic near 1), (2) check for evidence of uncorrelated chain sampling by parameters (effective sample size to sample size ratio near 1), and (3) visually compare our posterior predictions against true data distributions.

```
ehlModel3Weeks <- brm(
  data = vh2o %>%
    filter(week == "3" & Ta %in% c(30, 40)) %>%
    select(Ta, ring, pretreatment,
      mass, tarsusLengthMean, ehl,
      "batch" = exp
    ) %>%
    pivot_wider(
      id_cols = c(
        "ring", "batch",
        "pretreatment", "mass",
        "tarsusLengthMean"
      ),
      values_from = "ehl",
      names_from = "Ta"
    ) %>%
```

```
    mutate("foldEhl" = `40` / `30`) %>%
    mutate(pretreatment = ifelse(pretreatment == "neutral", "B",
      ifelse(pretreatment == "cold", "A", "C")
    )) %>%
    mutate(pretreatment = factor(pretreatment,
      levels = c("B", "A", "C")
    )) %>%
    distinct() %>%
    mutate(
      mass = mass - mean(mass, na.rm = T),
      tarsus = tarsusLengthMean - mean(tarsusLengthMean, na.rm = T)
    ),
  family = "gaussian",
  bf(mass ~ pretreatment + (1 | batch)) +
    bf(tarsus ~ mass + pretreatment + (1 | batch)) +
    bf(foldEhl ~ tarsus + mass + pretreatment + (1 | batch)) +
    set_rescor(FALSE),
  prior = c(
    set_prior("normal(0, 5)",
      class = "Intercept",
      resp = "mass"
    ),
    set_prior("normal(0, 15)",
      class = "b",
      coef = "pretreatmentA",
      resp = "mass"
    ),
    set_prior("normal(0, 15)",
      class = "b",
      coef = "pretreatmentC",
      resp = "mass"
    ),
    set_prior("exponential(2.5)",
      class = "sd",
      group = "batch",
      resp = "mass"
    ),
    set_prior("exponential(0.15)",
      class = "sigma",
      resp = "mass"
    ),
    set_prior("normal(0, 2.5)",
      class = "Intercept",
      resp = "tarsus"
    ),
    set_prior("normal(0, 2.5)",
      class = "b",
      coef = "pretreatmentA",
      resp = "tarsus"
    ),
    set_prior("normal(0, 2.5)",
      class = "b",
      coef = "pretreatmentC",
      resp = "tarsus"
    ),
    set_prior("skew_normal(0, 0.25, 5)",
      class = "b",
      coef = "mass",
      resp = "tarsus"
    ),
    set_prior("exponential(2)",
      class = "sd",
      group = "batch",
      resp = "tarsus"
    ),
    set_prior("exponential(1)",
      class = "sigma",
```

```
      resp = "tarsus"
    ),
    set_prior("normal(2.5, 1)",
      class = "Intercept",
      resp = "foldEhl"
    ),
    set_prior("normal(0, 0.5)",
      class = "b",
      coef = "pretreatmentA",
      resp = "foldEhl"
    ),
    set_prior("normal(0, 0.5)",
      class = "b",
      coef = "pretreatmentC",
      resp = "foldEhl"
    ),
    set_prior("normal(0, 0.025)",
      class = "b",
      coef = "mass",
      resp = "foldEhl"
    ),
    set_prior("normal(0, 0.5)",
      class = "b",
      coef = "tarsus",
      resp = "foldEhl"
    ),
    set_prior("exponential(10)",
      class = "sd",
      group = "batch",
      resp = "foldEhl"
    ),
    set_prior("exponential(5)",
      class = "sigma",
      resp = "foldEhl"
    )
  ),
  iter = 50000, warmup = 10000, cores = 4, chains = 4, thin = 20,
  control = list(adapt_delta = .97, max_treedepth = 14),
  silent = TRUE, refresh = 0,
  file = "./models/heatLossModel3Weeks.Rds",
)

pp1a <- mcmc_neff(neff_ratio(ehlModel3Weeks)) +
  xlab(
    TeX("$\\overset{Evaporative~Heat~Loss}{Response~(N_{eff}/N)}$")
  ) +
  theme_classic() +
  theme(
    axis.text.y = element_blank(),
    axis.ticks.y = element_blank(),
    legend.position = "none"
  )

pp1b <- mcmc_rhat(rhat(ehlModel3Weeks)) +
  xlab(
    TeX("$\\overset{Evaporative~Heat~Loss}{Response~(\\hat{R}})$")
  ) +
  theme_classic() +
  theme(
    axis.text.y = element_blank(),
    axis.ticks.y = element_blank(),
    legend.position = "none"
  )

pp1c <- pp_check2(ehlModel3Weeks,
  resp = "foldEhl",
  xlab = "Evaporative Heat\nLoss Response (Fold From 30°C)"
```

```
) +
  theme(legend.position = "none")

pp1d <- ehlModel3Weeks$data %>%
  mutate(
    "Fit" = fitted(ehlModel3Weeks, resp = "foldEhl")[, "Estimate"],
    "FitSE" = fitted(ehlModel3Weeks, resp = "foldEhl")[, "Est.Error"]
  ) %>%
  ggplot(aes(x = Fit, y = foldEhl)) +
  geom_errorbarh(aes(xmin = Fit - FitSE, xmax = Fit + FitSE),
    height = 0.25, colour = "black", alpha = 0.8
  ) +
  geom_point(
    size = 2, pch = 21, colour = "black", fill = "lightblue2",
    alpha = 0.8
  ) +
  geom_smooth(
    method = "lm", colour = "black", formula = y ~ 0 + x,
    linetype = "dashed", se = FALSE
  ) +
  xlab("Fitted Evaporative Heat\nLoss (Fold from 30°C)") +
  ylab("Evaporative Heat Loss\n(Fold from 30°C)") +
  theme_classic()

efficiencyModel3Weeks <- brm(
  data = vh2o %>%
    filter(week == "3" & Ta == 40) %>%
    select(Ta, ring, pretreatment,
      mass, tarsusLengthMean, ecc,
      "batch" = exp
    ) %>%
    mutate(
      pretreatment =
        ifelse(pretreatment == "neutral", "B",
          ifelse(pretreatment == "cold", "A", "C")
        )
    ) %>%
    mutate(pretreatment = factor(pretreatment,
      levels = c("B", "A", "C")
    )) %>%
    distinct() %>%
    mutate(
      mass = mass - mean(mass, na.rm = T),
      tarsus = tarsusLengthMean -
        mean(tarsusLengthMean, na.rm = T)
    ),
  family = "gaussian",
  bf(mass ~ pretreatment + (1 | batch)) +
    bf(tarsus ~ mass + pretreatment + (1 | batch)) +
    bf(ecc ~ tarsus + mass + pretreatment + (1 | batch),
     sigma ~ batch) +
    set_rescor(FALSE),
  prior = c(
    set_prior("normal(0, 5)",
      class = "Intercept",
      resp = "mass"
    ),
    set_prior("normal(0, 15)",
      class = "b",
      coef = "pretreatmentA",
      resp = "mass"
    ),
    set_prior("normal(0, 15)",
      class = "b",
      coef = "pretreatmentC",
      resp = "mass"
    ),
```

```
    set_prior("exponential(2.5)",
      class = "sd",
      group = "batch",
      resp = "mass"
    ),
    set_prior("exponential(0.15)",
      class = "sigma",
      resp = "mass"
    ),
    set_prior("normal(0, 2.5)",
      class = "Intercept",
      resp = "tarsus"
    ),
    set_prior("normal(0, 2.5)",
      class = "b",
      coef = "pretreatmentA",
      resp = "tarsus"
    ),
    set_prior("normal(0, 2.5)",
      class = "b",
      coef = "pretreatmentC",
      resp = "tarsus"
    ),
    set_prior("skew_normal(0, 0.25, 5)",
      class = "b",
      coef = "mass",
      resp = "tarsus"
    ),
    set_prior("exponential(2)",
      class = "sd",
      group = "batch",
      resp = "tarsus"
    ),
    set_prior("exponential(1)",
      class = "sigma",
      resp = "tarsus"
    ),
    set_prior("normal(0.5, 0.2)",
      class = "Intercept",
      resp = "ecc"
    ),
    set_prior("normal(0, 0.25)",
      class = "b",
      coef = "pretreatmentA",
      resp = "ecc"
    ),
    set_prior("normal(0, 0.25)",
      class = "b",
      coef = "pretreatmentC",
      resp = "ecc"
    ),
    set_prior("normal(0, 0.01)",
      class = "b",
      coef = "mass",
      resp = "ecc"
    ),
    set_prior("normal(0, 0.1)",
      class = "b",
      coef = "tarsus",
      resp = "ecc"
    ),
    set_prior("exponential(15)",
      class = "sd",
      group = "batch",
      resp = "ecc"
    ),
    set_prior("normal(-2, 1)",
```

```
      dpar = "sigma",
      class = "Intercept",
      resp = "ecc"
    ),
    set_prior("normal(0, 0.5)",
      dpar = "sigma",
      class = "b",
      coef = "batchB",
      resp = "ecc"
    ),
    set_prior("normal(1, 1)",
      dpar = "sigma",
      class = "b",
      coef = "batchC",
      resp = "ecc"
    )
  ),
  iter = 50000, warmup = 10000, cores = 4, chains = 4, thin = 20,
  control = list(adapt_delta = .97, max_treedepth = 14),
  silent = TRUE, refresh = 0,
  file = "./models/efficiencyModel3Weeks.Rds"
)

pp2a <- mcmc_neff(neff_ratio(efficiencyModel3Weeks)) +
  xlab(
    TeX("$\\overset{Evaporative~Cooling}{Efficiency~N_{eff}/N}$")
  ) +
  theme_classic() +
  theme(
    axis.text.y = element_blank(),
    axis.ticks.y = element_blank(),
    legend.position = "none"
  )

pp2b <- mcmc_rhat(rhat(efficiencyModel3Weeks)) +
  xlab(
    TeX("$\\overset{Evaporative~Cooling~Efficiency}{\\hat{R}}$")
  ) +
  theme_classic() +
  theme(
    axis.text.y = element_blank(),
    axis.ticks.y = element_blank(),
    legend.position = "none"
  )

pp2c <- pp_check2(efficiencyModel3Weeks,
  resp = "ecc",
  xlab = "Evaporative Cooling\nEfficiency (EHL/RMR)"
) + scale_x_continuous(n.breaks = 3)

pp2d <- efficiencyModel3Weeks$data %>%
  mutate(
    "Fit" = fitted(efficiencyModel3Weeks,
      resp = "ecc"
    )[, "Estimate"],
    "FitSE" = fitted(efficiencyModel3Weeks,
      resp = "ecc"
    )[, "Est.Error"]
  ) %>%
  ggplot(aes(x = Fit, y = ecc)) +
  geom_errorbarh(aes(xmin = Fit - FitSE, xmax = Fit + FitSE),
    height = 0.05, colour = "black", alpha = 0.8
  ) +
  geom_point(
    size = 2, pch = 21, colour = "black", fill = "lightblue2",
    alpha = 0.8
  ) +
```

```
  geom_smooth(
    method = "lm", colour = "black", formula = y ~ 0 + x,
    linetype = "dashed", se = FALSE
  ) +
  xlab("Fitted Evaporative\nCooling Efficiency (EHL/RMR)") +
  ylab("Evaporative Cooling\nEfficiency (EHL/RMR)") +
  theme_classic()

((pp1a + pp1b) /
  (pp1c + pp1d) /
  (pp2a + pp2b) /
  (pp2c + pp2d)) +
  plot_annotation(tag_levels = "A")
```

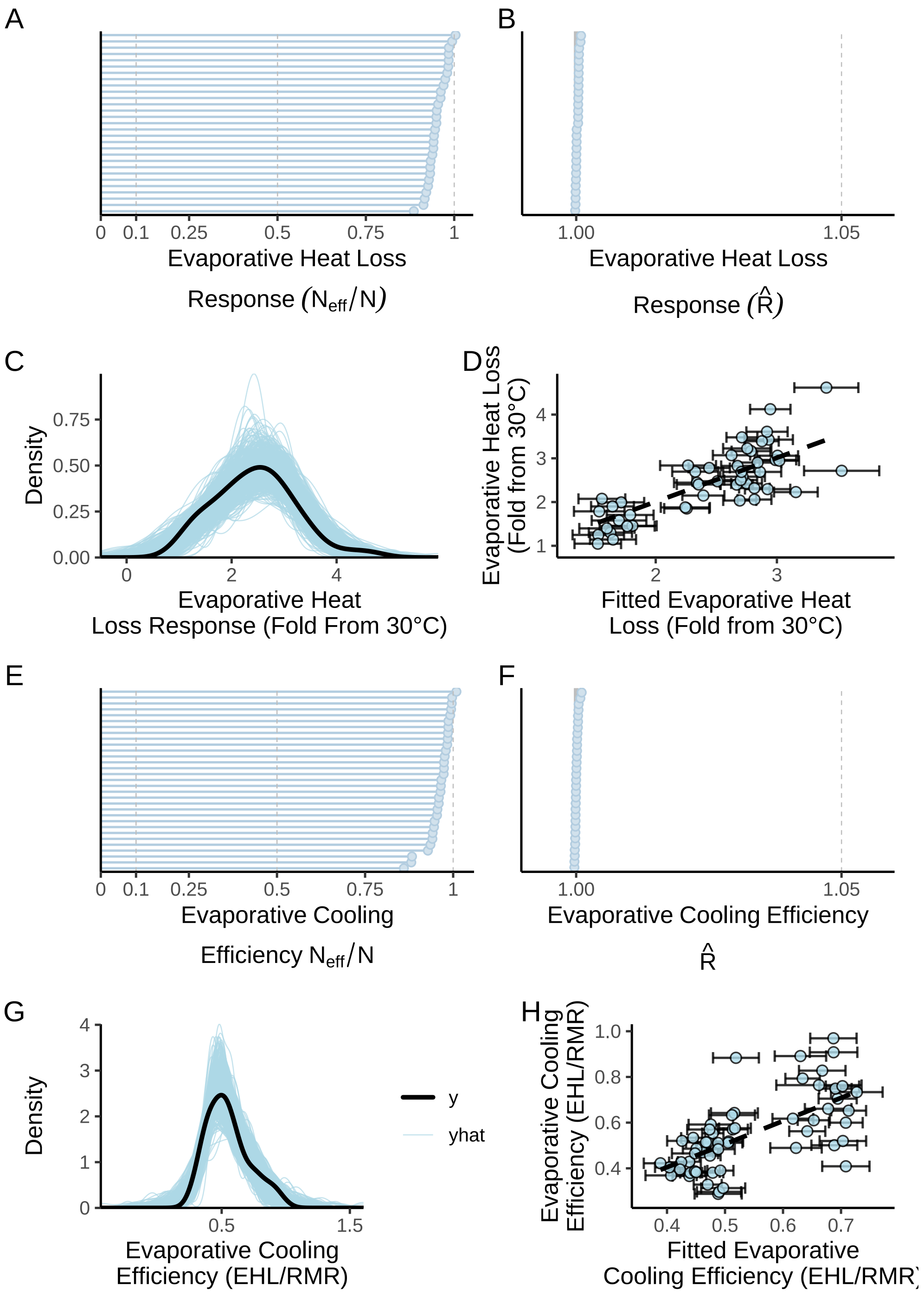


***Figure 10:*** *Model and posterior predictive checks for two Bayesian path analyses predicting, ultimately, evaporative heat loss ('EHL'; fold from that observed at 30°C in W) and evaporative cooling efficiency ('ECE'; the ratio of evaporative heat-loss to metabolic heat production, each in W) in three week old Japanese quail. Panels A and E display parameter-specific effective sample size to sample size ratios ($N_{eff}/N$) for each model, while panels B and F display parameter specific Gelman-Rubin statistics ($\hat{R}$). Panels C and G display posterior predictive checks; black lines represent true EHL or ECE densities while blue lines represent densities estimated from model posteriors. In panels D and H, dots represent individual predictions plotted against their true values. Errorbars indicate +/- one standard error around predictions and dashed black lines indicate lines of best fit estimated by ggplot2 (Wickham, 2011) while assuming a y-intercept at 0.*

Next, we visualise the spread of model residuals (here, as medians) against model predictors and expectations. After, we check posterior densities per model predictor for evidence of skewing or kurtosis.

```
rp1 <- ehlModel3Weeks$data %>%
  mutate(
    "Res" =
      residuals(ehlModel3Weeks,
        resp = "foldEhl",
        robust = TRUE
      )[, "Estimate"]
  ) %>%
  ggplot(aes(sample = Res)) +
  stat_qq(colour = "grey50") +
  stat_qq_line() +
  xlab("Theoretical EHL\nResidual Quantiles") +
  ylab("Sample ELH\nResidual Quantiles") +
  theme_classic()

rp2 <- ehlModel3Weeks$data %>%
  mutate(
    "Res" =
      residuals(ehlModel3Weeks,
        resp = "foldEhl",
        robust = TRUE
      )[, "Estimate"]
  ) %>%
  mutate(mass = mass + mean(subset(vh2o, week == "3")$mass, na.rm = T)) %>%
  ggplot(aes(x = mass, y = Res)) +
  geom_point(pch = 21, colour = "black", fill = "grey20", alpha = 0.5) +
  xlab("Body Mass (g)") +
  ylab("Evaporative Heat\nLoss Residuals") +
  theme_classic()

rp3 <- ehlModel3Weeks$data %>%
  mutate(
    "Res" =
      residuals(ehlModel3Weeks,
        resp = "foldEhl",
        robust = TRUE
      )[, "Estimate"]
  ) %>%
  mutate(tarsus = tarsus +
    mean(subset(vh2o, week == "3")$tarsusLengthMean, na.rm = T)) %>%
  ggplot(aes(x = tarsus, y = Res)) +
  geom_point(pch = 21, colour = "black", fill = "grey20", alpha = 0.5) +
  xlab("Tarsus Length (mm)") +
  ylab("Evaporative Heat\nLoss Residuals") +
  theme_classic()

rp4 <- ehlModel3Weeks$data %>%
  mutate(
    "Res" =
      residuals(ehlModel3Weeks,
        resp = "foldEhl",
        robust = TRUE
      )[, "Estimate"]
  ) %>%
  mutate(pretreatment = factor(pretreatment, levels = c("A", "B", "C"))) %>%
  ggplot(aes(x = pretreatment, y = Res)) +
  geom_boxplot(fill = "lightblue2") +
  geom_point(size = 2, position = position_jitter(width = 0.25)) +
  scale_x_discrete(
    name = "Rearing Treatment",
    labels = c(
      "Cold\n(10°C)",
      "Mild\n(20°C)",
      "Warm\n(30°C)"
```

```
    )
  ) +
  xlab("Rearing Treatment") +
  ylab("Evaporative Heat\nLoss Residuals") +
  theme_classic()

rp5 <- efficiencyModel3Weeks$data %>%
  mutate(
    "Res" =
      residuals(efficiencyModel3Weeks,
        resp = "ecc",
        robust = TRUE
      )[, "Estimate"]
  ) %>%
  ggplot(aes(sample = Res)) +
  stat_qq(colour = "grey50") +
  stat_qq_line() +
  xlab("Theoretical ECE\nResidual Quantiles") +
  ylab("Sample ECE\nResidual Quantiles") +
  theme_classic()

rp6 <- efficiencyModel3Weeks$data %>%
  mutate(
    "Res" =
      residuals(efficiencyModel3Weeks,
        resp = "ecc",
        robust = TRUE
      )[, "Estimate"]
  ) %>%
  mutate(mass = mass + mean(subset(vh2o, week == "3")$mass, na.rm = T)) %>%
  ggplot(aes(x = mass, y = Res)) +
  geom_point(pch = 21, colour = "black", fill = "grey20", alpha = 0.5) +
  xlab("Body Mass (g)") +
  ylab("Evaporative Cooling\nEfficiency Residuals") +
  theme_classic()

rp7 <- efficiencyModel3Weeks$data %>%
  mutate(
    "Res" =
      residuals(efficiencyModel3Weeks,
        resp = "ecc",
        robust = TRUE
      )[, "Estimate"]
  ) %>%
  mutate(tarsus = tarsus +
    mean(subset(vh2o, week == "3")$tarsusLengthMean, na.rm = T)) %>%
  ggplot(aes(x = tarsus, y = Res)) +
  geom_point(pch = 21, colour = "black", fill = "grey20", alpha = 0.5) +
  xlab("Tarsus Length (mm)") +
  ylab("Evaporative Cooling\nEfficiency Residuals") +
  theme_classic()

rp8 <- efficiencyModel3Weeks$data %>%
  mutate(
    "Res" =
      residuals(efficiencyModel3Weeks,
        resp = "ecc",
        robust = TRUE
      )[, "Estimate"]
  ) %>%
  mutate(pretreatment = factor(pretreatment, levels = c("A", "B", "C"))) %>%
  ggplot(aes(x = pretreatment, y = Res)) +
  geom_boxplot(fill = "lightblue2") +
  geom_point(size = 2, position = position_jitter(width = 0.25)) +
  scale_x_discrete(
    name = "Rearing Treatment",
    labels = c(
```

```
      "Cold\n(10°C)",
      "Mild\n(20°C)",
      "Warm\n(30°C)"
    )
  ) +
  xlab("Rearing Treatment") +
  ylab("Evaporative Cooling\nEfficiency Residuals") +
  theme_classic()

((rp1 + rp2)/
  (rp3 + rp4)/
  (rp5 + rp6)/
  (rp7 + rp8)
) + plot_annotation(tag_levels = "A")
```

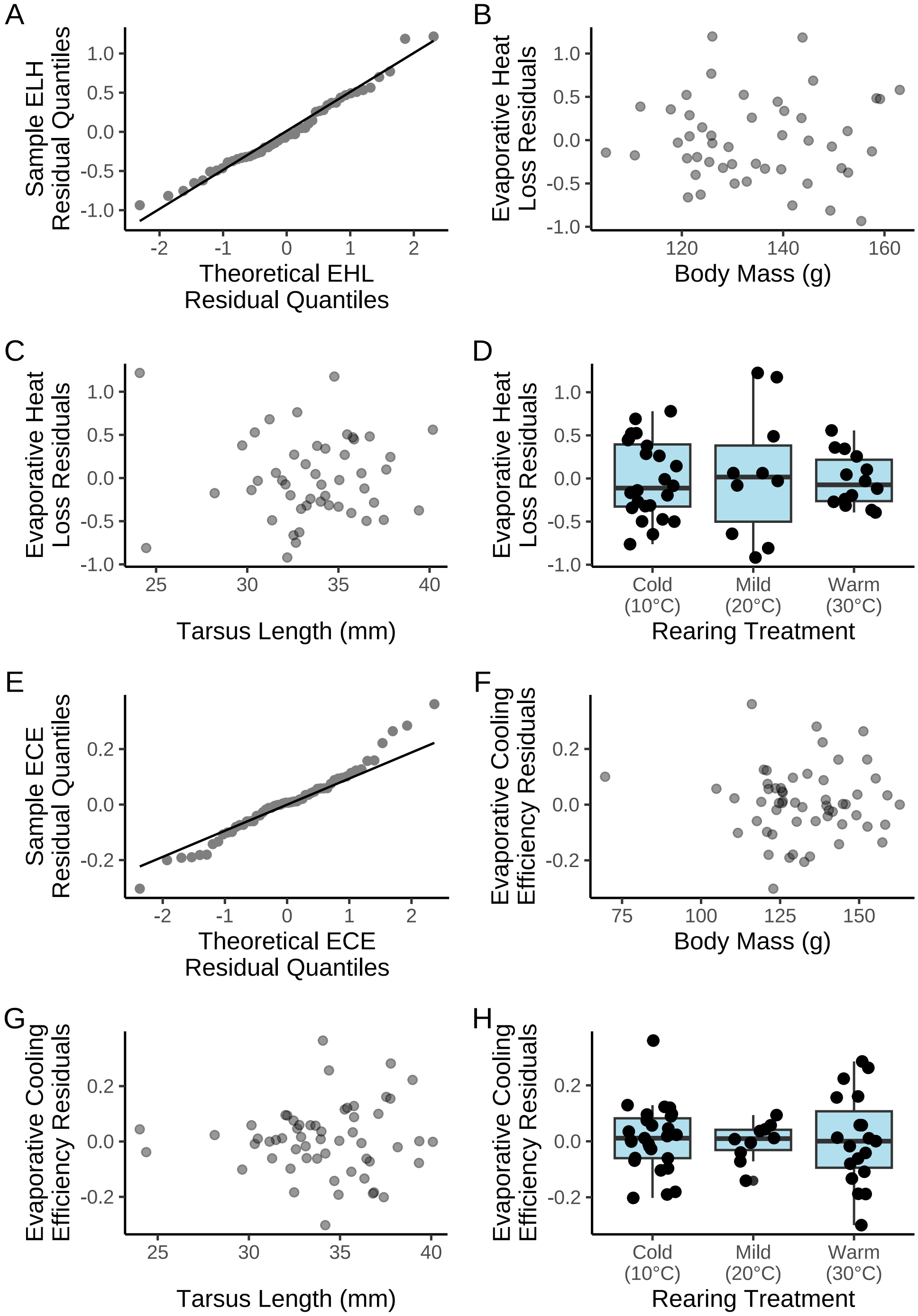


***Figure 11:*** *Residual diagnostics from two Bayesian path analyses predicting, ultimately, evaporative heat loss ('EHL'; fold from that observed at 30°C in W) and evaporative cooling efficiency ('ECE'; the ratio of evaporative heat-loss to metabolic heat production, each in W) in three week old Japanese quail. Panels A and B display traditional 'qq-plots', with theoretic and sample residual quantiles regressed against each other. Dots represent individual samples. Remaining panels display median residual, per raw data point (small dots) by model predictors (here, body mass [g], tarsus length [mm] and rearing condition (10°C, 20°C, or 30°C until the time of measurement). Boxplots in panels D and H display medians (centre horizontal bar), first and third quantiles (lower and upper limits of boxes respectively) and ranges excluding outliers (whiskers). 'EHL' indicates evaporative heat loss in watts, and ECE indicates evaporative cooling efficiency (the ratio of evaporative heat loss by metabolic heat production, each in watts).*

Residual spreads are largely homogenous and normal. However, one individual with an extremely low mass is evident in our efficiency analysis but not our relative heat loss analysis. This individual is therefore removed from our efficiency analysis and our analysis re-run.

```
efficiencyModel3Weeks <- brm(
  data = vh2o %>%
    filter(week == "3" & Ta == 40) %>%
    select(Ta, ring, pretreatment,
      mass, tarsusLengthMean, ecc,
      "batch" = exp
    ) %>%
    mutate(
      pretreatment =
        ifelse(pretreatment == "neutral", "B",
          ifelse(pretreatment == "cold", "A", "C")
        )
    ) %>%
    mutate(pretreatment = factor(pretreatment,
      levels = c("B", "A", "C")
    )) %>%
    distinct() %>%
    filter(mass > 75) %>%
    mutate(
      mass = mass - mean(mass, na.rm = T),
      tarsus = tarsusLengthMean -
        mean(tarsusLengthMean, na.rm = T)
    ),
  family = "gaussian",
  bf(mass ~ pretreatment + (1 | batch)) +
    bf(tarsus ~ mass + pretreatment + (1 | batch)) +
    bf(ecc ~ tarsus + mass + pretreatment + (1 | batch),
       sigma ~ batch) +
    set_rescor(FALSE),
  prior = c(
    set_prior("normal(0, 5)",
      class = "Intercept",
      resp = "mass"
    ),
    set_prior("normal(0, 15)",
      class = "b",
      coef = "pretreatmentA",
      resp = "mass"
    ),
    set_prior("normal(0, 15)",
      class = "b",
      coef = "pretreatmentC",
      resp = "mass"
    ),
    set_prior("exponential(2.5)",
      class = "sd",
      group = "batch",
      resp = "mass"
    ),
    set_prior("exponential(0.15)",
      class = "sigma",
      resp = "mass"
    ),
    set_prior("normal(0, 2.5)",
      class = "Intercept",
      resp = "tarsus"
    ),
    set_prior("normal(0, 2.5)",
      class = "b",
      coef = "pretreatmentA",
      resp = "tarsus"
    ),
    set_prior("normal(0, 2.5)",
      class = "b",
```

```
    coef = "pretreatmentC",
    resp = "tarsus"
  ),
  set_prior("skew_normal(0, 0.25, 5)",
    class = "b",
    coef = "mass",
    resp = "tarsus"
  ),
  set_prior("exponential(2)",
    class = "sd",
    group = "batch",
    resp = "tarsus"
  ),
  set_prior("exponential(1)",
    class = "sigma",
    resp = "tarsus"
  ),
  set_prior("normal(0.5, 0.2)",
    class = "Intercept",
    resp = "ecc"
  ),
  set_prior("normal(0, 0.25)",
    class = "b",
    coef = "pretreatmentA",
    resp = "ecc"
  ),
  set_prior("normal(0, 0.25)",
    class = "b",
    coef = "pretreatmentC",
    resp = "ecc"
  ),
  set_prior("normal(0, 0.01)",
    class = "b",
    coef = "mass",
    resp = "ecc"
  ),
  set_prior("normal(0, 0.1)",
    class = "b",
    coef = "tarsus",
    resp = "ecc"
  ),
  set_prior("exponential(15)",
    class = "sd",
    group = "batch",
    resp = "ecc"
  ),
  set_prior("normal(-2, 1)",
    dpar = "sigma",
    class = "Intercept",
    resp = "ecc"
  ),
  set_prior("normal(0, 0.5)",
    dpar = "sigma",
    class = "b",
    coef = "batchB",
    resp = "ecc"
  ),
  set_prior("normal(1, 1)",
    dpar = "sigma",
    class = "b",
    coef = "batchC",
    resp = "ecc"
  )
),
iter = 50000, warmup = 10000, cores = 4, chains = 4, thin = 20,
control = list(adapt_delta = .97, max_treedepth = 14),
silent = TRUE, refresh = 0,
```

```
  file = "./models/efficiencyModel3WeeksRevised.Rds"
)

ppa <- mcmc_neff(neff_ratio(efficiencyModel3Weeks)) +
  xlab(
    TeX("$\\overset{Evaporative~Cooling}{Efficiency~N_{eff}/N}$")
  ) +
  theme_classic() +
  theme(
    axis.text.y = element_blank(),
    axis.ticks.y = element_blank(),
    legend.position = "none"
  )

ppb <- mcmc_rhat(rhat(efficiencyModel3Weeks)) +
  xlab(
    TeX("$\\overset{Evaporative~Cooling~Efficiency}{\\hat{R}}$")
  ) +
  theme_classic() +
  theme(
    axis.text.y = element_blank(),
    axis.ticks.y = element_blank(),
    legend.position = "none"
  )

ppc <- pp_check2(efficiencyModel3Weeks,
  resp = "ecc",
  xlab = "Evaporative Cooling\nEfficiency (EHL/RMR)"
) + scale_x_continuous(n.breaks = 3)

(ppa + ppb)/ppc + plot_annotation(tag_levels = "A")
```

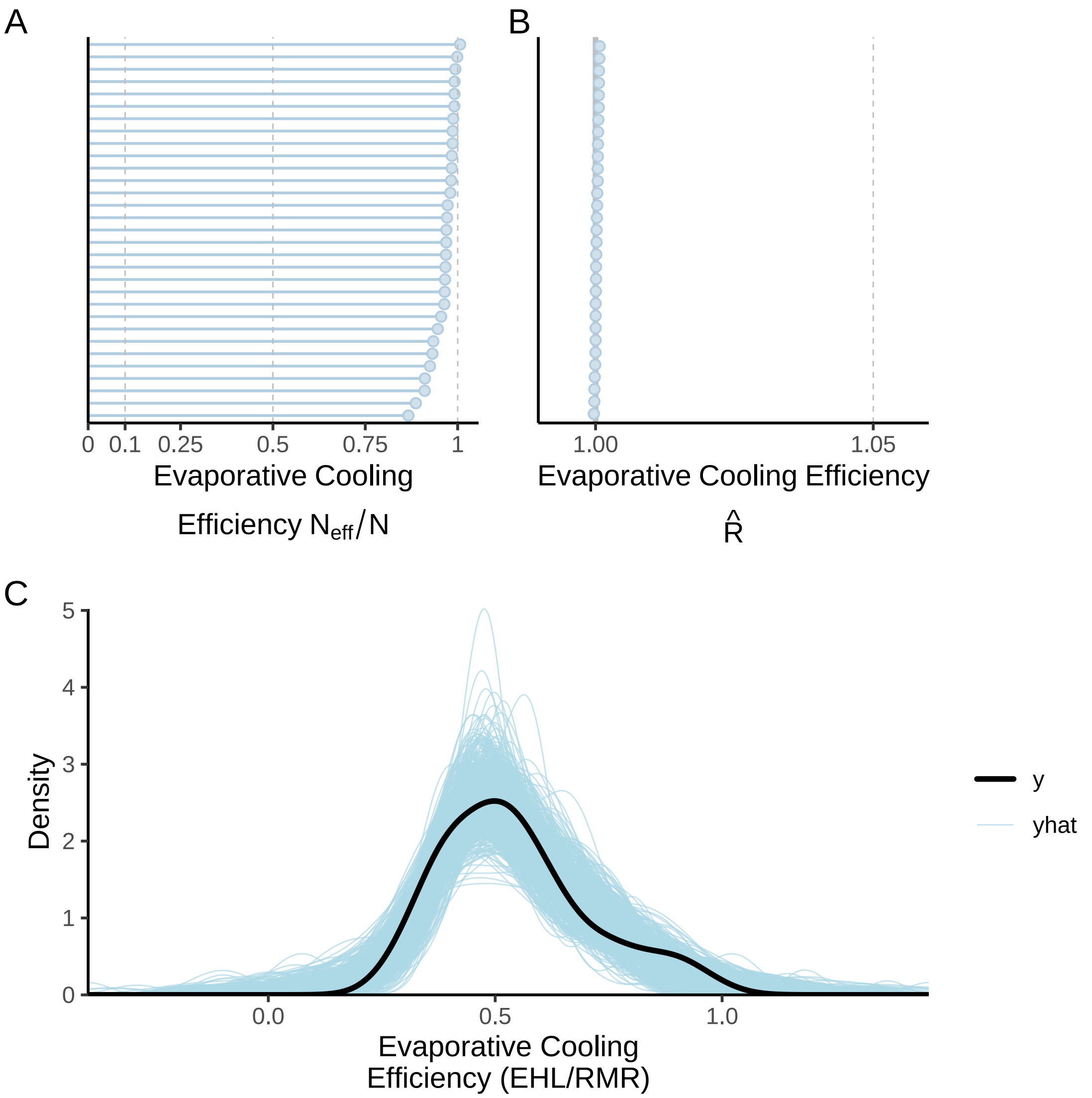


***Figure 12:*** *Model and posterior predictive checks for a Bayesian path analysis predicting evaporative cooling efficiency ('ECE'; the ratio of evaporative heat-loss to metabolic heat production, each in W) in three week old Japanese quail. Panel A displays parameter-specific effective sample size to sample size ratios ($N_{eff}/N$), panel B displays parameter specific Gelman-Rubin statistics ($\hat{R}$), and panel panels C displays a posterior predictive check; black lines represent true ECE densities while blue lines represent densities estimated from model posteriors.*

Now we proceed to plotting model coefficients.

```
as.data.frame(ehlModel3Weeks) %>%
  pivot_longer(everything(), names_to = "par", values_to = "values") %>%
  filter(grepl("b_|sd_", par)) %>%
  merge(., tribble(~par, ~Par,
                   "b_mass_Intercept", "Mass\nIntercept",
                   "b_mass_pretreatmentA", "Mass ~\nCold Rearing",
                   "b_mass_pretreatmentC", "Mass ~\nWarm Rearing",
                   "sd_batch__mass_Intercept", "Mass ~\nBatch",
                   "b_tarsus_Intercept", "Tarsus\nIntercept",
                   "b_tarsus_pretreatmentA", "Tarsus ~\nCold Rearing",
                   "b_tarsus_pretreatmentC", "Tarsus ~\nWarm Rearing",
```

```
                    "b_tarsus_mass", "Tarsus ~\nMass",
                    "sd_batch__tarsus_Intercept", "Tarsus ~\nBatch",
                    "b_foldEhl_Intercept", "EHL\nIntercept",
                    "b_foldEhl_pretreatmentA", "EHL ~\nCold Rearing",
                    "b_foldEhl_pretreatmentC", "EHL ~\nWarm Rearing",
                    "b_foldEhl_mass", "EHL ~ Mass",
                    "b_foldEhl_tarsus", "EHL ~\nTarsus",
                    "sd_batch__foldEhl_Intercept", "EHL ~\nBatch",
                    ),
        by = "par", all.x = TRUE) %>%
  ggplot(aes(x = values)) +
  facet_wrap(~Par, scales = "free") +
  geom_density(colour = "black", fill = "white") +
  geom_vline(xintercept = 0, colour = "darkred", linetype = "solid") +
  scale_x_continuous(n.breaks = 3) +
  scale_y_continuous(n.breaks = 3) +
  xlab("Values") +
  ylab("Density") +
  theme_classic()
```

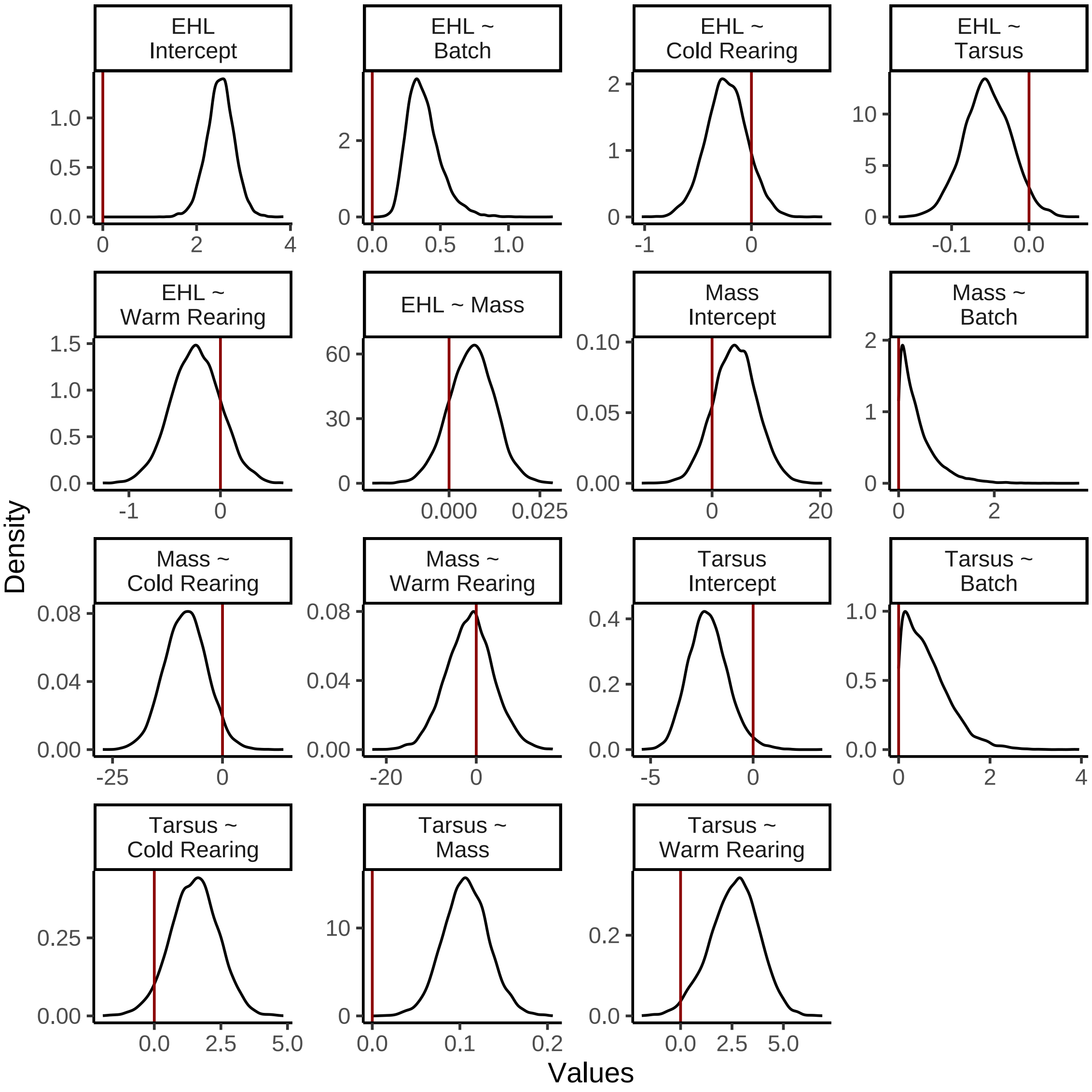


***Figure 13:*** *Posterior densities for coefficients from a Bayesian path analysis predicting body mass (g), tarsus length (mm) and evaporative heat loss responses ('EHL'; fold from that observed at 30°C in W) in three week old Japanese quail. Red vertical lines label 0. Response and predictor variables for which densities refer are indicated on the left side and right side of tildes respectively. 'Cold Rearing' indicates rearing at 10°C, 'Warm Rearing' indicates rearing at 30°C, and 'Batch' indicates the batch of eggs from which an individual was derived. All coefficients except batch are population level.*

```
as.data.frame(efficiencyModel3Weeks) %>%
  pivot_longer(everything(), names_to = "par", values_to = "values") %>%
  filter(grepl("b_|sd_", par)) %>%
  merge(., tribble(~par, ~Par,
                   "b_mass_Intercept", "Mass\nIntercept",
                   "b_mass_pretreatmentA", "Mass ~\nCold Rearing",
                   "b_mass_pretreatmentC", "Mass ~\nWarm Rearing",
                   "sd_batch__mass_Intercept", "Mass ~\nBatch",
                   "b_tarsus_Intercept", "Tarsus\nIntercept",
                   "b_tarsus_pretreatmentA", "Tarsus ~\nCold Rearing",
                   "b_tarsus_pretreatmentC", "Tarsus ~\nWarm Rearing",
                   "b_tarsus_mass", "Tarsus ~\nMass",
                   "sd_batch__tarsus_Intercept", "Tarsus ~\nBatch",
```

```
                    "b_ecc_Intercept", "ECC\nIntercept",
                    "b_ecc_pretreatmentA", "ECC ~\nCold Rearing",
                    "b_ecc_pretreatmentC", "ECC ~\nWarm Rearing",
                    "b_ecc_mass", "ECC ~ Mass",
                    "b_ecc_tarsus", "ECC ~\nTarsus",
                    "sd_batch__ecc_Intercept", "ECC ~\nBatch",
                    ),
        by = "par", all.x = TRUE) %>%
  ggplot(aes(x = values)) +
  facet_wrap(~Par, scales = "free") +
  geom_density(colour = "black", fill = "white") +
  geom_vline(xintercept = 0, colour = "darkred", linetype = "solid") +
  scale_x_continuous(n.breaks = 3) +
  scale_y_continuous(n.breaks = 3) +
  xlab("Values") +
  ylab("Density") +
  theme_classic()
```

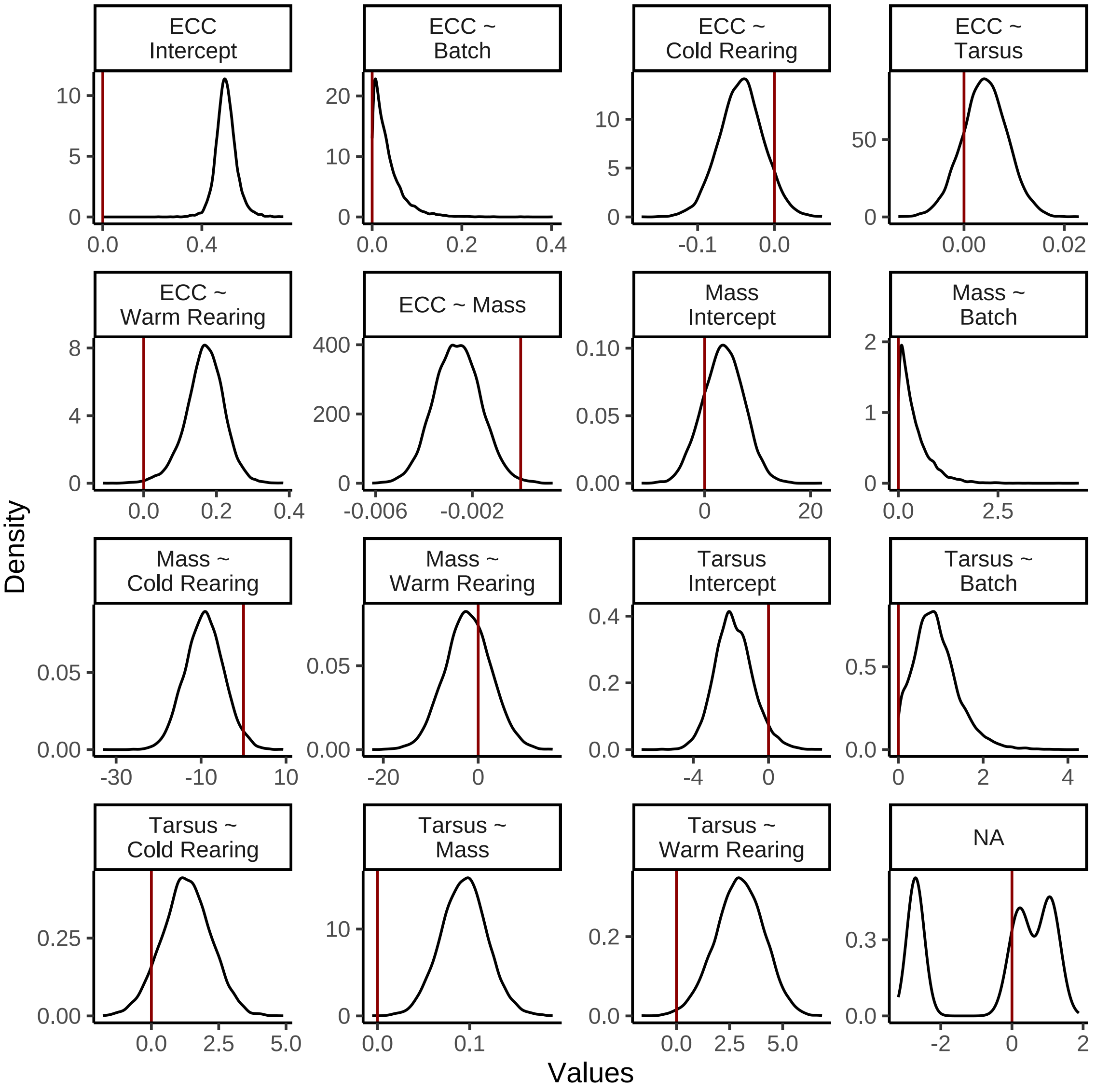


***Figure 14:*** *Posterior densities for coefficients from a Bayesian path analysis predicting body mass (g), tarsus length (mm) and evaporative cooling efficiency ('ECC', the ratio of evaporative heat loss in watts to evaporative heat production in watts) in three week old Japanese quail. Red vertical lines label 0. Response and predictor variables for which densities refer are indicated on the left side and right side of tildes respectively. 'Cold Rearing' indicates rearing at 10°C, 'Warm Rearing' indicates rearing at 30°C, and 'Batch' indicates the batch of eggs from which an individual was derived. All coefficients except batch are population level.*

Given that some densities are slightly skewed or leptokurtic, we chose to summarise posteriors as medians and estimate error around these summaries using quantile-based credible intervals. Following this summarisation, we estimate variance explained by our total models (using Bayesian $R^2$ values) and by morphometric predictors (using partial Bayesian $R^2$ values).

```
caption <- paste0(
  "Results of Bayesian path analysis testing the ",
  "effects of morphology and rearing temperature on relative evaporative ",
  "heat loss at 40°C in three week old Japanese quail. Relative evapoative ",
  "heat loss represents the Evaporative cooling efficiency ",
  "fold change from 30°C, per individual. Estimates indicate posterior ",
```

```
  "medians and credible intervels (CIs) indicate quantile intervals."
)

heatLoss3WeeksResults <-
  as.data.frame(ehlModel3Weeks) %>%
  summarise_all(., .funs = median) %>%
  pivot_longer(everything(),
    names_to = "Parameter",
    values_to = "Estimate"
  ) %>%
  merge(., quantileCIs(ehlModel3Weeks, cis = c(50, 95)),
    by = "Parameter", all.x = TRUE
  ) %>%
  filter(grepl("b_|sd_", Parameter)) %>%
  rowwise() %>%
  mutate("BF" = ifelse(Estimate < 0,
    (2 * mean(as.data.frame(
      ehlModel3Weeks
    )[, Parameter] <= 0)) /
      (2 * mean(as.data.frame(
        ehlModel3Weeks
      )[, Parameter] >= 0)),
    (2 * mean(as.data.frame(
      ehlModel3Weeks
    )[, Parameter] >= 0)) /
      (2 * mean(as.data.frame(
        ehlModel3Weeks
      )[, Parameter] <= 0))
  )) %>%
  ungroup() %>%
  mutate(
    "Estimate" = round(Estimate, digits = 4),
    "BF" = round(BF, digits = 4),
    "N" = nrow(ehlModel3Weeks$data)
  ) %>%
  mutate("Parameter" = ifelse(grepl("b_", Parameter),
    gsub("b_", "", Parameter),
    gsub(
      "Intercept", "batch",
      gsub(".*__", "", Parameter)
    )
  )) %>%
  mutate(
    "Response" = gsub("_.*", "", Parameter),
    "Parameter" = gsub(".*_", "", Parameter)
  ) %>%
  merge(., tribble(
    ~Response, ~response, ~level,
    "mass", "Body Mass (g)", "A",
    "foldEhl", "Evaporative Heat Loss (fold from 30°C)", "D",
    "tarsus", "Tarsus Length (mm)", "B"
  ),
  by = "Response"
  ) %>%
  merge(., tribble(
    ~Parameter, ~parameter, ~number,
    "Intercept", "Intercept", "1",
    "mass", "Body Mass (g)", "4",
    "tarsus", "Tarsus Length (mm)", "5",
    "pretreatmentA", "Cold Rearing", "2",
    "pretreatmentC", "Warm Rearing", "3",
    "batch", "Egg Batch [mu]", "6"
  ),
  by = "Parameter"
  ) %>%
  mutate(
    `50\\% CI` = paste0("(", paste(
```

```
      round(Low_CI_50, digits = 4),
      round(High_CI_50, digits = 4),
      sep = ", "
    ), ")"),
    `95\\% CI` = paste0("(", paste(
      round(Low_CI_95, digits = 4),
      round(High_CI_95, digits = 4),
      sep = ", "
    ), ")")
  ) %>%
  select(-c(Low_CI_50, High_CI_50, Low_CI_95, High_CI_95)) %>%
  select(
    "Response" = "response", "Parameter" = "parameter", N,
    Estimate, `50\\% CI`, `95\\% CI`, BF, level, number
  ) %>%
  arrange(level, number) %>%
  select(-c(level, number)) %>%
  kbl(.,
    longtable = T, booktabs = T, format = "latex", escape = FALSE,
    caption = caption
  ) %>%
  column_spec(column = c(1:2), width = "2.1cm") %>%
  column_spec(column = c(3:10), width = "1.8cm") %>%
  kable_styling(latex_options = "striped")

heatLoss3WeeksResults
```

***Table 2:*** *Results of Bayesian path analysis testing the effects of morphology and rearing temperature on relative evaporative heat loss at 40°C in three week old Japanese quail. Relative evapoative heat loss represents the Evaporative cooling efficiency fold change from 30°C, per individual. Estimates indicate posterior medians and credible intervels (CIs) indicate quantile intervals.*

| Response | Parameter | N | Estimate | 50% CI | 95% CI | BF |
|---|---|---|---|---|---|---|
| Body Mass (g) | Intercept | 48 | 4.2748 | (1.5227, 6.914) | (-3.6383, 12.1768) | 5.9025 |
| Body Mass (g) | Cold Rearing | 48 | -8.3386 | (-11.6305, -5.1198) | (-17.759, 0.9783) | 24.6410 |
| Body Mass (g) | Warm Rearing | 48 | -1.3179 | (-4.9282, 2.0196) | (-11.8457, 9.079) | 1.5633 |
| Body Mass (g) | Egg Batch [mu] | 48 | 0.2802 | (0.1135, 0.5461) | (0.0087, 1.466) | Inf |
| Tarsus Length (mm) | Intercept | 48 | -2.2295 | (-2.8441, -1.579) | (-3.9262, -0.1067) | 46.6190 |
| Tarsus Length (mm) | Cold Rearing | 48 | 1.5287 | (0.9192, 2.1187) | (-0.2886, 3.2543) | 20.2766 |
| Tarsus Length (mm) | Warm Rearing | 48 | 2.6689 | (1.8442, 3.4288) | (0.1685, 4.8282) | 55.7376 |
| Tarsus Length (mm) | Body Mass (g) | 48 | 0.1067 | (0.0897, 0.1241) | (0.0564, 0.1591) | Inf |
| Tarsus Length (mm) | Egg Batch [mu] | 48 | 0.5265 | (0.2314, 0.9006) | (0.0203, 1.8637) | Inf |
| Evaporative Heat Loss (fold from 30°C) | Intercept | 48 | 2.5094 | (2.3213, 2.6949) | (1.9145, 3.074) | Inf |
| Evaporative Heat Loss (fold from 30°C) | Cold Rearing | 48 | -0.2339 | (-0.3593, -0.1059) | (-0.6129, 0.1546) | 7.8692 |
| Evaporative Heat Loss (fold from 30°C) | Warm Rearing | 48 | -0.2682 | (-0.4523, -0.0858) | (-0.8018, 0.2734) | 5.2257 |
| Evaporative Heat Loss (fold from 30°C) | Body Mass (g) | 48 | 0.0063 | (0.0021, 0.0104) | (-0.0062, 0.0186) | 5.2745 |

| | | | | | | |
|---|---|---|---|---|---|---|
| Evaporative Heat Loss (fold from 30°C) | Tarsus Length (mm) | 48 | -0.0543 | (-0.0743, -0.0328) | (-0.1133, 0.0062) | 23.9221 |
| Evaporative Heat Loss (fold from 30°C) | Egg Batch [mu] | 48 | 0.3620 | (0.2915, 0.4471) | (0.1908, 0.6772) | Inf |

```
caption <- paste0(
  "Results of Bayesian path analysis testing the ",
  "effects of morphology and rearing temperature on evaporative ",
  "cooling efficiency at 40°C in three week old Japanese quail. ",
  "Evaporative cooling efficiency represents the ratio of ",
  "evaporative heat loss (in W) to metabolic heat production ",
  "(again, in W). Estimates indicate posterior medians and credible ",
  "intervels (CIs) indicate quantile intervals."
)

efficiency3WeeksResults <-
  as.data.frame(efficiencyModel3Weeks) %>%
  summarise_all(., .funs = median) %>%
  pivot_longer(everything(),
    names_to = "Parameter",
    values_to = "Estimate"
  ) %>%
  merge(., quantileCIs(efficiencyModel3Weeks,
                       cis = c(50, 95)),
    by = "Parameter", all.x = TRUE
  ) %>%
  filter(grepl("b_|sd_", Parameter)) %>%
  rowwise() %>%
  mutate("BF" = ifelse(Estimate < 0,
    (2 * mean(as.data.frame(
      efficiencyModel3Weeks
    )[, Parameter] <= 0)) /
      (2 * mean(as.data.frame(
        efficiencyModel3Weeks
      )[, Parameter] >= 0)),
    (2 * mean(as.data.frame(
      efficiencyModel3Weeks
    )[, Parameter] >= 0)) /
      (2 * mean(as.data.frame(
        efficiencyModel3Weeks
      )[, Parameter] <= 0))
  )) %>%
  ungroup() %>%
  mutate(
    "Estimate" = round(Estimate, digits = 4),
    "BF" = round(BF, digits = 4),
    "N" = nrow(efficiencyModel3Weeks$data)
  ) %>%
  mutate("Parameter" = ifelse(grepl("b_", Parameter),
    gsub("b_", "", Parameter),
    gsub(
      "Intercept", "batch",
      gsub(".*__", "", Parameter)
    )
  )) %>%
  mutate(
    "Response" = gsub("_.*", "", Parameter),
    "Parameter" = gsub(".*_", "", Parameter)
  ) %>%
  merge(., tribble(
    ~Response, ~response, ~level,
    "mass", "Body Mass (g)", "A",
    "ecc", "Evaporative Cooling Efficiency", "D",
    "tarsus", "Tarsus Length (mm)", "B"
  ),
```

```
  by = "Response"
  ) %>%
  merge(., tribble(
    ~Parameter, ~parameter, ~number,
    "Intercept", "Intercept", "1",
    "mass", "Body Mass (g)", "4",
    "tarsus", "Tarsus Length (mm)", "5",
    "pretreatmentA", "Cold Rearing", "2",
    "pretreatmentC", "Warm Rearing", "3",
    "batch", "Egg Batch [mu]", "6"
  ),
  by = "Parameter"
  ) %>%
  mutate(
    `50\\% CI` = paste0("(", paste(
      round(Low_CI_50, digits = 4),
      round(High_CI_50, digits = 4),
      sep = ", "
    ), ")"),
    `95\\% CI` = paste0("(", paste(
      round(Low_CI_95, digits = 4),
      round(High_CI_95, digits = 4),
      sep = ", "
    ), ")")
  ) %>%
  select(-c(Low_CI_50, High_CI_50, Low_CI_95, High_CI_95)) %>%
  select(
    "Response" = "response", "Parameter" = "parameter", N,
    Estimate, `50\\% CI`, `95\\% CI`, BF, level, number
  ) %>%
  arrange(level, number) %>%
  select(-c(level, number)) %>%
  kbl(.,
    longtable = T, booktabs = T, format = "latex", escape = FALSE,
    caption = caption
  ) %>%
  column_spec(column = c(1:2), width = "2.1cm") %>%
  column_spec(column = c(3:10), width = "1.8cm") %>%
  kable_styling(latex_options = "striped")

efficiency3WeeksResults
```

***Table 3:*** *Results of Bayesian path analysis testing the effects of morphology and rearing temperature on evaporative cooling efficiency at 40°C in three week old Japanese quail. Evaporative cooling efficiency represents the ratio of evaporative heat loss (in W) to metabolic heat production (again, in W). Estimates indicate posterior medians and credible intervels (CIs) indicate quantile intervals.*

| Response | Parameter | N | Estimate | 50% CI | 95% CI | BF |
|---|---|---|---|---|---|---|
| Body Mass (g) | Intercept | 55 | 3.5530 | (0.8995, 6.1473) | (-4.0248, 10.9654) | 4.3476 |
| Body Mass (g) | Cold Rearing | 55 | -9.2038 | (-12.3038, -6.2072) | (-17.9892, -0.0403) | 40.2371 |
| Body Mass (g) | Warm Rearing | 55 | -2.1775 | (-5.3675, 1.1206) | (-11.5769, 7.4524) | 2.0315 |
| Body Mass (g) | Egg Batch [mu] | 55 | 0.2724 | (0.1131, 0.538) | (0.0087, 1.401) | Inf |
| Tarsus Length (mm) | Intercept | 55 | -1.9040 | (-2.5327, -1.1989) | (-3.7442, 0.3711) | 21.4090 |
| Tarsus Length (mm) | Cold Rearing | 55 | 1.3001 | (0.7015, 1.9087) | (-0.5165, 3.0769) | 11.8205 |
| Tarsus Length (mm) | Warm Rearing | 55 | 2.9828 | (2.2164, 3.758) | (0.6538, 5.1493) | 162.2653 |
| Tarsus Length (mm) | Body Mass (g) | 55 | 0.0942 | (0.077, 0.1105) | (0.0441, 0.1443) | 3999.0000 |
| Tarsus Length (mm) | Egg Batch [mu] | 55 | 0.8320 | (0.5241, 1.1777) | (0.0643, 2.1173) | Inf |

| | | | | | | |
|---|---|---|---|---|---|---|
| Evaporative Cooling Efficiency | Intercept | 55 | 0.4960 | (0.4728, 0.5207) | (0.4183, 0.5887) | Inf |
| Evaporative Cooling Efficiency | Cold Rearing | 55 | -0.0428 | (-0.0621, -0.024) | (-0.0996, 0.0147) | 13.4665 |
| Evaporative Cooling Efficiency | Warm Rearing | 55 | 0.1699 | (0.1366, 0.2025) | (0.0536, 0.268) | 234.2941 |
| Evaporative Cooling Efficiency | Body Mass (g) | 55 | -0.0026 | (-0.0033, -0.002) | (-0.0045, -7e-04) | 194.1220 |
| Evaporative Cooling Efficiency | Tarsus Length (mm) | 55 | 0.0042 | (0.0012, 0.0071) | (-0.0048, 0.0131) | 4.6140 |
| Evaporative Cooling Efficiency | Egg Batch [mu] | 55 | 0.0238 | (0.01, 0.0453) | (9e-04, 0.1237) | Inf |

Partial $R^2$ values are hear calculated by removing the focal predictor (i.e. either body mass or tarsus length) from models within path analyses predicting evaporative heat loss or cooling measurements. Models are then re-run and total $R^2$ values compared between original and parameter-excluding models. The extent to which $R^2$ values are improved by inclusion of the target predictor is assummed as its partial $R^2$.

```
## Calculating total R2 values

caption = paste0("Estimates of fit for each element of a Bayesian path ",
                 "analysis predicting evaporative heat loss responses ",
                 "(fold from 30°C; Model 'A') and evaporative cooling ",
                 "efficiency (Model 'B') in developing Japanese quail ",
                 "(3 weeks of age). Response variables refer to those ",
                 "measured at 40°C. Credible intervals are quantile intervals.")

brms::bayes_R2(ehlModel3Weeks, ndraws = 1000,
               robust = TRUE) %>%
  as.data.frame() %>%
  rownames_to_column("var") %>%
  merge(., tribble(
    ~var, ~Var,
    "R2mass", "Body Mass (g)",
    "R2tarsus", "Tarsus Length (mm)",
    "R2foldEhl",
      "Evaporative Heat Loss Response"
  ), by = c("var")) %>%
  mutate(Estimate = round(Estimate, digits = 4),
         Est.Error = round(Est.Error, digits = 4),
         "95\\% CI" = paste0("[", round(Q2.5, digits = 4),
                             ", ", round(Q97.5, digits = 4),
                             "]"
         ),
         "Model" = "A"
         ) %>%
  rbind(., brms::bayes_R2(efficiencyModel3Weeks, ndraws = 1000,
               robust = TRUE) %>%
  as.data.frame() %>%
  rownames_to_column("var") %>%
  merge(., tribble(
    ~var, ~Var,
    "R2mass", "Body Mass (g)",
    "R2tarsus", "Tarsus Length (mm)",
    "R2ecc",
      "Evaporative Cooling Efficiency"
  ), by = c("var")) %>%
  mutate(Estimate = round(Estimate, digits = 4),
```

```
        Est.Error = round(Est.Error, digits = 4),
        "95\\% CI" = paste0("[", round(Q2.5, digits = 4),
                            ", ", round(Q97.5, digits = 4),
                            "]"
        ),
        "Model" = "B"
        )) %>%
  select(
    Model, "Response" = Var, "R\\textsuperscript{2}" = Estimate,
    "Standard Error" = Est.Error,
    `95\\% CI`
  ) %>%
  kbl(.,
    longtable = T, booktabs = T, format = "latex",
    caption = caption, escape = FALSE
  ) %>%
  column_spec(column = c(1:10), width = "2.5cm")  %>%
  kable_styling(latex_options = "striped")
```

***Table 4:*** *Estimates of fit for each element of a Bayesian path analysis predicting evaporative heat loss responses (fold from 30°C; Model 'A') and evaporative cooling efficiency (Model 'B') in developing Japanese quail (3 weeks of age). Response variables refer to those measured at 40°C. Credible intervals are quantile intervals.*

| Model | Response | $R^2$ | Standard Error | 95% CI |
|---|---|---|---|---|
| A | Evaporative Heat Loss Response | 0.5679 | 0.0687 | [0.3799, 0.6688] |
| A | Body Mass (g) | 0.0850 | 0.0650 | [0.0072, 0.2271] |
| A | Tarsus Length (mm) | 0.4000 | 0.0825 | [0.2167, 0.5337] |
| B | Evaporative Cooling Efficiency | 0.4166 | 0.0651 | [0.281, 0.5246] |
| B | Body Mass (g) | 0.0957 | 0.0634 | [0.0115, 0.2311] |
| B | Tarsus Length (mm) | 0.4385 | 0.0775 | [0.2515, 0.5598] |

```
# Calculating Partial R2 values.

ehlModel3WeeksMassR2 <- brm(
  data = vh2o %>%
    filter(week == "3" & Ta %in% c(30, 40)) %>%
    select(Ta, ring, pretreatment,
      mass, tarsusLengthMean, ehl,
      "batch" = exp
    ) %>%
    pivot_wider(
      id_cols = c(
        "ring", "batch",
        "pretreatment", "mass",
        "tarsusLengthMean"
      ),
      values_from = "ehl",
      names_from = "Ta"
    ) %>%
    mutate("foldEhl" = `40` / `30`) %>%
    mutate(pretreatment = ifelse(pretreatment == "neutral", "B",
      ifelse(pretreatment == "cold", "A", "C")
    )) %>%
    mutate(pretreatment = factor(pretreatment,
      levels = c("B", "A", "C")
    )) %>%
    distinct() %>%
    mutate(
      mass = mass - mean(mass, na.rm = T),
```

```
    tarsus = tarsusLengthMean - mean(tarsusLengthMean, na.rm = T)
  ),
  family = "gaussian",
  bf(mass ~ pretreatment + (1 | batch)) +
    bf(tarsus ~ mass + pretreatment + (1 | batch)) +
    bf(foldEhl ~ tarsus + pretreatment + (1 | batch)) +
    set_rescor(FALSE),
  prior = c(
    set_prior("normal(0, 5)",
      class = "Intercept",
      resp = "mass"
    ),
    set_prior("normal(0, 15)",
      class = "b",
      coef = "pretreatmentA",
      resp = "mass"
    ),
    set_prior("normal(0, 15)",
      class = "b",
      coef = "pretreatmentC",
      resp = "mass"
    ),
    set_prior("exponential(2.5)",
      class = "sd",
      group = "batch",
      resp = "mass"
    ),
    set_prior("exponential(0.15)",
      class = "sigma",
      resp = "mass"
    ),
    set_prior("normal(0, 2.5)",
      class = "Intercept",
      resp = "tarsus"
    ),
    set_prior("normal(0, 2.5)",
      class = "b",
      coef = "pretreatmentA",
      resp = "tarsus"
    ),
    set_prior("normal(0, 2.5)",
      class = "b",
      coef = "pretreatmentC",
      resp = "tarsus"
    ),
    set_prior("skew_normal(0, 0.25, 5)",
      class = "b",
      coef = "mass",
      resp = "tarsus"
    ),
    set_prior("exponential(2)",
      class = "sd",
      group = "batch",
      resp = "tarsus"
    ),
    set_prior("exponential(1)",
      class = "sigma",
      resp = "tarsus"
    ),
    set_prior("normal(2.5, 1)",
      class = "Intercept",
      resp = "foldEhl"
    ),
    set_prior("normal(0, 0.5)",
      class = "b",
      coef = "pretreatmentA",
      resp = "foldEhl"
```

```
    ),
    set_prior("normal(0, 0.5)",
      class = "b",
      coef = "pretreatmentC",
      resp = "foldEhl"
    ),
    set_prior("normal(0, 0.5)",
      class = "b",
      coef = "tarsus",
      resp = "foldEhl"
    ),
    set_prior("exponential(10)",
      class = "sd",
      group = "batch",
      resp = "foldEhl"
    ),
    set_prior("exponential(5)",
      class = "sigma",
      resp = "foldEhl"
    )
  ),
  iter = 50000, warmup = 10000, cores = 4, chains = 4, thin = 20,
  control = list(adapt_delta = .97, max_treedepth = 14),
  silent = TRUE, refresh = 0,
  file = "./models/heatLossModel3WeeksMassR2.Rds",
)

efficiencyModel3WeeksMassR2 <- brm(
  data = vh2o %>%
    filter(week == "3" & Ta == 40) %>%
    select(Ta, ring, pretreatment,
      mass, tarsusLengthMean, ecc,
      "batch" = exp
    ) %>%
    mutate(
      pretreatment =
        ifelse(pretreatment == "neutral", "B",
          ifelse(pretreatment == "cold", "A", "C")
        )
    ) %>%
    mutate(pretreatment = factor(pretreatment,
      levels = c("B", "A", "C")
    )) %>%
    distinct() %>%
    mutate(
      mass = mass - mean(mass, na.rm = T),
      tarsus = tarsusLengthMean -
        mean(tarsusLengthMean, na.rm = T)
    ),
  family = "gaussian",
  bf(mass ~ pretreatment + (1 | batch)) +
    bf(tarsus ~ mass + pretreatment + (1 | batch)) +
    bf(ecc ~ tarsus + pretreatment + (1 | batch),
     sigma ~ batch) +
    set_rescor(FALSE),
  prior = c(
    set_prior("normal(0, 5)",
      class = "Intercept",
      resp = "mass"
    ),
    set_prior("normal(0, 15)",
      class = "b",
      coef = "pretreatmentA",
      resp = "mass"
    ),
    set_prior("normal(0, 15)",
      class = "b",
```

```
      coef = "pretreatmentC",
      resp = "mass"
    ),
    set_prior("exponential(2.5)",
      class = "sd",
      group = "batch",
      resp = "mass"
    ),
    set_prior("exponential(0.15)",
      class = "sigma",
      resp = "mass"
    ),
    set_prior("normal(0, 2.5)",
      class = "Intercept",
      resp = "tarsus"
    ),
    set_prior("normal(0, 2.5)",
      class = "b",
      coef = "pretreatmentA",
      resp = "tarsus"
    ),
    set_prior("normal(0, 2.5)",
      class = "b",
      coef = "pretreatmentC",
      resp = "tarsus"
    ),
    set_prior("skew_normal(0, 0.25, 5)",
      class = "b",
      coef = "mass",
      resp = "tarsus"
    ),
    set_prior("exponential(2)",
      class = "sd",
      group = "batch",
      resp = "tarsus"
    ),
    set_prior("exponential(1)",
      class = "sigma",
      resp = "tarsus"
    ),
    set_prior("normal(0.5, 0.2)",
      class = "Intercept",
      resp = "ecc"
    ),
    set_prior("normal(0, 0.25)",
      class = "b",
      coef = "pretreatmentA",
      resp = "ecc"
    ),
    set_prior("normal(0, 0.25)",
      class = "b",
      coef = "pretreatmentC",
      resp = "ecc"
    ),
    set_prior("normal(0, 0.1)",
      class = "b",
      coef = "tarsus",
      resp = "ecc"
    ),
    set_prior("exponential(15)",
      class = "sd",
      group = "batch",
      resp = "ecc"
    ),
    set_prior("normal(-2, 1)",
      dpar = "sigma",
      class = "Intercept",
```

```
      resp = "ecc"
    ),
    set_prior("normal(0, 0.5)",
      dpar = "sigma",
      class = "b",
      coef = "batchB",
      resp = "ecc"
    ),
    set_prior("normal(1, 1)",
      dpar = "sigma",
      class = "b",
      coef = "batchC",
      resp = "ecc"
    )
  ),
  iter = 50000, warmup = 10000, cores = 4, chains = 4, thin = 20,
  control = list(adapt_delta = .97, max_treedepth = 14),
  silent = TRUE, refresh = 0,
  file = "./models/efficiencyModel3WeeksMassR2.Rds"
)

ehlModel3WeeksTarsusR2 <- brm(
  data = vh2o %>%
    filter(week == "3" & Ta %in% c(30, 40)) %>%
    select(Ta, ring, pretreatment,
      mass, tarsusLengthMean, ehl,
      "batch" = exp
    ) %>%
    pivot_wider(
      id_cols = c(
        "ring", "batch",
        "pretreatment", "mass",
        "tarsusLengthMean"
      ),
      values_from = "ehl",
      names_from = "Ta"
    ) %>%
    mutate("foldEhl" = `40` / `30`) %>%
    mutate(pretreatment = ifelse(pretreatment == "neutral", "B",
      ifelse(pretreatment == "cold", "A", "C")
    )) %>%
    mutate(pretreatment = factor(pretreatment,
      levels = c("B", "A", "C")
    )) %>%
    distinct() %>%
    mutate(
      mass = mass - mean(mass, na.rm = T),
      tarsus = tarsusLengthMean - mean(tarsusLengthMean, na.rm = T)
    ),
  family = "gaussian",
  bf(mass ~ pretreatment + (1 | batch)) +
    bf(tarsus ~ mass + pretreatment + (1 | batch)) +
    bf(foldEhl ~ mass + pretreatment + (1 | batch)) +
    set_rescor(FALSE),
  prior = c(
    set_prior("normal(0, 5)",
      class = "Intercept",
      resp = "mass"
    ),
    set_prior("normal(0, 15)",
      class = "b",
      coef = "pretreatmentA",
      resp = "mass"
    ),
    set_prior("normal(0, 15)",
      class = "b",
      coef = "pretreatmentC",
```

```
    resp = "mass"
  ),
  set_prior("exponential(2.5)",
    class = "sd",
    group = "batch",
    resp = "mass"
  ),
  set_prior("exponential(0.15)",
    class = "sigma",
    resp = "mass"
  ),
  set_prior("normal(0, 2.5)",
    class = "Intercept",
    resp = "tarsus"
  ),
  set_prior("normal(0, 2.5)",
    class = "b",
    coef = "pretreatmentA",
    resp = "tarsus"
  ),
  set_prior("normal(0, 2.5)",
    class = "b",
    coef = "pretreatmentC",
    resp = "tarsus"
  ),
  set_prior("skew_normal(0, 0.25, 5)",
    class = "b",
    coef = "mass",
    resp = "tarsus"
  ),
  set_prior("exponential(2)",
    class = "sd",
    group = "batch",
    resp = "tarsus"
  ),
  set_prior("exponential(1)",
    class = "sigma",
    resp = "tarsus"
  ),
  set_prior("normal(2.5, 1)",
    class = "Intercept",
    resp = "foldEhl"
  ),
  set_prior("normal(0, 0.5)",
    class = "b",
    coef = "pretreatmentA",
    resp = "foldEhl"
  ),
  set_prior("normal(0, 0.5)",
    class = "b",
    coef = "pretreatmentC",
    resp = "foldEhl"
  ),
  set_prior("normal(0, 0.025)",
    class = "b",
    coef = "mass",
    resp = "foldEhl"
  ),
  set_prior("exponential(10)",
    class = "sd",
    group = "batch",
    resp = "foldEhl"
  ),
  set_prior("exponential(5)",
    class = "sigma",
    resp = "foldEhl"
  )
```

```
  ),
  iter = 50000, warmup = 10000, cores = 4, chains = 4, thin = 20,
  control = list(adapt_delta = .97, max_treedepth = 14),
  silent = TRUE, refresh = 0,
  file = "./models/heatLossModel3WeeksTarsusR2.Rds",
)

efficiencyModel3WeeksTarsusR2 <- brm(
  data = vh2o %>%
    filter(week == "3" & Ta == 40) %>%
    select(Ta, ring, pretreatment,
      mass, tarsusLengthMean, ecc,
      "batch" = exp
    ) %>%
    mutate(
      pretreatment =
        ifelse(pretreatment == "neutral", "B",
          ifelse(pretreatment == "cold", "A", "C")
        )
    ) %>%
    mutate(pretreatment = factor(pretreatment,
      levels = c("B", "A", "C")
    )) %>%
    distinct() %>%
    mutate(
      mass = mass - mean(mass, na.rm = T),
      tarsus = tarsusLengthMean -
        mean(tarsusLengthMean, na.rm = T)
    ),
  family = "gaussian",
  bf(mass ~ pretreatment + (1 | batch)) +
    bf(tarsus ~ mass + pretreatment + (1 | batch)) +
    bf(ecc ~ tarsus + mass + pretreatment + (1 | batch),
     sigma ~ batch) +
    set_rescor(FALSE),
  prior = c(
    set_prior("normal(0, 5)",
      class = "Intercept",
      resp = "mass"
    ),
    set_prior("normal(0, 15)",
      class = "b",
      coef = "pretreatmentA",
      resp = "mass"
    ),
    set_prior("normal(0, 15)",
      class = "b",
      coef = "pretreatmentC",
      resp = "mass"
    ),
    set_prior("exponential(2.5)",
      class = "sd",
      group = "batch",
      resp = "mass"
    ),
    set_prior("exponential(0.15)",
      class = "sigma",
      resp = "mass"
    ),
    set_prior("normal(0, 2.5)",
      class = "Intercept",
      resp = "tarsus"
    ),
    set_prior("normal(0, 2.5)",
      class = "b",
      coef = "pretreatmentA",
      resp = "tarsus"
```

```
    ),
    set_prior("normal(0, 2.5)",
      class = "b",
      coef = "pretreatmentC",
      resp = "tarsus"
    ),
    set_prior("skew_normal(0, 0.25, 5)",
      class = "b",
      coef = "mass",
      resp = "tarsus"
    ),
    set_prior("exponential(2)",
      class = "sd",
      group = "batch",
      resp = "tarsus"
    ),
    set_prior("exponential(1)",
      class = "sigma",
      resp = "tarsus"
    ),
    set_prior("normal(0.5, 0.2)",
      class = "Intercept",
      resp = "ecc"
    ),
    set_prior("normal(0, 0.25)",
      class = "b",
      coef = "pretreatmentA",
      resp = "ecc"
    ),
    set_prior("normal(0, 0.25)",
      class = "b",
      coef = "pretreatmentC",
      resp = "ecc"
    ),
    set_prior("normal(0, 0.01)",
      class = "b",
      coef = "mass",
      resp = "ecc"
    ),
    set_prior("normal(0, 0.1)",
      class = "b",
      coef = "tarsus",
      resp = "ecc"
    ),
    set_prior("exponential(15)",
      class = "sd",
      group = "batch",
      resp = "ecc"
    ),
    set_prior("normal(-2, 1)",
      dpar = "sigma",
      class = "Intercept",
      resp = "ecc"
    ),
    set_prior("normal(0, 0.5)",
      dpar = "sigma",
      class = "b",
      coef = "batchB",
      resp = "ecc"
    ),
    set_prior("normal(1, 1)",
      dpar = "sigma",
      class = "b",
      coef = "batchC",
      resp = "ecc"
    )
  ),
```

```
  iter = 50000, warmup = 10000, cores = 4, chains = 4, thin = 20,
  control = list(adapt_delta = .97, max_treedepth = 14),
  silent = TRUE, refresh = 0,
  file = "./models/efficiencyModel3WeeksTarsusR2.Rds"
)

heatLossMassPR2 <- brms::bayes_R2(ehlModel3Weeks,
                          ndraws = 1000,
                          robust = TRUE, resp = "foldEhl",
                          summary = FALSE
) %>%
  as.data.frame() %>%
  select("fullR2" = R2foldEhl) %>%
  cbind(
    .,
    brms::bayes_R2(ehlModel3WeeksMassR2,
                   ndraws = 1000,
                   robust = TRUE, resp = "foldEhl",
                   summary = FALSE
    )
  ) %>%
  mutate("pR2" = round(fullR2 - R2foldEhl, digits = 3)) %>%
  pull(pR2)

efficiencyMassPR2 <- brms::bayes_R2(efficiencyModel3Weeks,
                          ndraws = 1000,
                          robust = TRUE, resp = "ecc",
                          summary = FALSE
) %>%
  as.data.frame() %>%
  select("fullR2" = R2ecc) %>%
  cbind(
    .,
    brms::bayes_R2(efficiencyModel3WeeksMassR2,
                   ndraws = 1000,
                   robust = TRUE, resp = "ecc",
                   summary = FALSE
    )
  ) %>%
  mutate("pR2" = round(fullR2 - R2ecc, digits = 3)) %>%
  pull(pR2)

heatLossTarsusPR2 <- brms::bayes_R2(ehlModel3Weeks,
                          ndraws = 1000,
                          robust = TRUE, resp = "foldEhl",
                          summary = FALSE
) %>%
  as.data.frame() %>%
  select("fullR2" = R2foldEhl) %>%
  cbind(
    .,
    brms::bayes_R2(ehlModel3WeeksTarsusR2,
                   ndraws = 1000,
                   robust = TRUE, resp = "foldEhl",
                   summary = FALSE
    )
  ) %>%
  mutate("pR2" = round(fullR2 - R2foldEhl, digits = 3)) %>%
  pull(pR2)

efficiencyTarsusPR2 <- brms::bayes_R2(efficiencyModel3Weeks,
                          ndraws = 1000,
                          robust = TRUE, resp = "ecc",
                          summary = FALSE
) %>%
  as.data.frame() %>%
  select("fullR2" = R2ecc) %>%
```

```
  cbind(
    .,
    brms::bayes_R2(efficiencyModel3WeeksTarsusR2,
                   ndraws = 1000,
                   robust = TRUE, resp = "ecc",
                   summary = FALSE
    )
  ) %>%
  mutate("pR2" = round(fullR2 - R2ecc, digits = 3)) %>%
  pull(pR2)

caption = paste0('Variance in evaporative heat loss (fold from that at 30°C) ',
                 'and evaporative cooling efficiency ',
                 '(EHL/MHP) at 40°C explained by morphometry in three week ',
                 'old Japanese quail. Variance explained is represented as ',
                 'partial R\\textsuperscript{2} values.')

partialR2ThreeWeeks <- data.frame(
  "Response" = rep(
    c(
      "Fold Evaporative Heat Loss",
      "Evaporative Cooling Efficiency"
    ),
    each = 2
  ),
  "Predictor" = rep(
    c(
      "Body Mass (g)",
      "Tarsus Length (mm)"
    ),
    2
  )
) %>%
  mutate(
    "Mass Partial R\\textsuperscript{2}" =
      lapply(
        X = list(
          heatLossMassPR2,
          efficiencyMassPR2,
          heatLossTarsusPR2,
          efficiencyTarsusPR2
        ),
        FUN = function(x) {
          round(median(x), digits = 4)
        }
      ) %>%
        unlist(),
    "95\\% CI" = lapply(
      X = list(
        heatLossMassPR2,
        efficiencyMassPR2,
        heatLossTarsusPR2,
        efficiencyTarsusPR2
      ),
      FUN = function(x) {
        paste0(
          "[",
          paste0(
            round(
              quantile(x, probs = c(0.025, 0.975), type = 8),
              digits = 4
            ),
            collapse = ", "
          ),
          "]"
        )
      }
```

```
    )
  ) %>%
  kbl(.,
      longtable = T, booktabs = T,
      format = "latex", escape = FALSE,
      caption = caption
  ) %>%
  column_spec(column = c(1:2), width = "2.0cm")  %>%
  column_spec(column = c(3:10), width = "1.9cm") %>%
  kable_styling(latex_options = "striped")

partialR2ThreeWeeks
```

***Table 5:*** *Variance in evaporative heat loss (fold from that at 30°C) and evaporative cooling efficiency (EHL/MHP) at 40°C explained by morphometry in three week old Japanese quail. Variance explained is represented as partial $R^2$ values.*

| Response | Predictor | Mass Partial $R^2$ | 95% CI |
|---|---|---|---|
| Fold Evaporative Heat Loss | Body Mass (g) | 0.010 | [-0.2056, 0.2193] |
| Fold Evaporative Heat Loss | Tarsus Length (mm) | 0.052 | [-0.132, 0.2517] |
| Evaporative Cooling Efficiency | Body Mass (g) | 0.019 | [-0.1743, 0.2327] |
| Evaporative Cooling Efficiency | Tarsus Length (mm) | -0.010 | [-0.1777, 0.163] |

Next, we visualise conditional effects of morphology and rearing conditions on evaporative heat loss responses and evaporative cooling efficiency in our developing quail. Conditional effects are estimated from our path analysis posteriors while assuming that all other parameters lay at their average.

```
## Producing morphology plots

heatLoss3WeeksMassPlot <-
  expand.grid(
    "mass" = with(
      ehlModel3Weeks$data,
      seq(min(mass), max(mass), by = 1)
    ),
    "tarsus" = 0,
    "pretreatment" = "B"
  ) %>%
  mutate(
    "foldEhl" = predict(ehlModel3Weeks,
      newdata = .,
      re_form = NA, robust = TRUE,
      resp = "foldEhl"
    )[, "Estimate"],
    "SE" = predict(ehlModel3Weeks,
      newdata = .,
      re_form = NA, robust = TRUE,
      resp = "foldEhl"
    )[, "Est.Error"]
  ) %>%
  mutate(mass = mass + mean(subset(vh2o, week == 3)$mass, na.rm = T)) %>%
  ggplot(aes(x = mass, y = foldEhl)) +
  geom_ribbon(aes(ymin = foldEhl - SE, ymax = foldEhl + SE),
    alpha = 0.5, fill = "#DECCC1"
```

```
  ) +
  geom_point(
    data = ehlModel3Weeks$data %>%
      mutate(mass = mass +
        mean(subset(vh2o, week == 3)$mass, na.rm = T)),
    aes(x = mass, y = foldEhl),
    alpha = 0.5
  ) +
  geom_smooth(
    method = "lm", colour = "black",
    linetype = "dashed", se = FALSE
  ) +
  xlab("Body Mass (g)") +
  ylab("Evaporative Heat Loss\n(Fold From 30°C)") +
  theme_classic() +
  theme(
    axis.title = element_text(family = "Noto Sans"),
    axis.text = element_text(family = "Noto Sans")
  )

heatLoss3WeeksTarsusPlot <-
  expand.grid(
    "tarsus" = with(
      ehlModel3Weeks$data,
      seq(min(tarsus), max(tarsus), by = 1)
    ),
    "mass" = 0,
    "pretreatment" = "B"
  ) %>%
  mutate(
    "foldEhl" = predict(ehlModel3Weeks,
      newdata = .,
      re_form = NA, robust = TRUE,
      resp = "foldEhl"
    )[, "Estimate"],
    "SE" = predict(ehlModel3Weeks,
      newdata = .,
      re_form = NA, robust = TRUE,
      resp = "foldEhl"
    )[, "Est.Error"]
  ) %>%
  mutate(tarsus = tarsus +
           mean(subset(vh2o, week == 3)$tarsusLengthMean, na.rm = T)) %>%
  ggplot(aes(x = tarsus, y = foldEhl)) +
  geom_ribbon(aes(ymin = foldEhl - SE, ymax = foldEhl + SE),
    alpha = 0.5, fill = "#DECCC1"
  ) +
  geom_point(
    data = ehlModel3Weeks$data %>%
      mutate(tarsus = tarsus +
        mean(subset(vh2o, week == 3)$tarsusLengthMean, na.rm = T)),
    aes(x = tarsus, y = foldEhl),
    alpha = 0.5
  ) +
  geom_smooth(
    method = "lm", colour = "black",
    linetype = "dashed", se = FALSE
  ) +
  xlab("Tarsus Length (mm)") +
  ylab("Evaporative Heat Loss\n(Fold From 30°C)") +
  theme_classic() +
  theme(
    axis.title = element_text(family = "Noto Sans"),
    axis.text = element_text(family = "Noto Sans")
  )

efficiency3WeeksMassPlot <-
```

```
  expand.grid(
    "mass" = with(
      efficiencyModel3Weeks$data,
      seq(min(mass), max(mass), by = 1)
    ),
    "tarsus" = 0,
    "pretreatment" = "B",
    "batch" = "A"
  ) %>%
  mutate(
    "ecc" = predict(efficiencyModel3Weeks,
      newdata = .,
      re_form = NA, robust = TRUE,
      resp = "ecc"
    )[, "Estimate"],
    "SE" = predict(efficiencyModel3Weeks,
      newdata = .,
      re_form = NA, robust = TRUE,
      resp = "ecc"
    )[, "Est.Error"]
  ) %>%
  mutate(mass = mass + mean(subset(vh2o, week == 3)$mass, na.rm = T)) %>%
  ggplot(aes(x = mass, y = ecc)) +
  geom_ribbon(aes(ymin = ecc - SE, ymax = ecc + SE),
    alpha = 0.5, fill = "#DECCC1"
  ) +
  geom_point(
    data = efficiencyModel3Weeks$data %>%
      mutate(mass = mass +
        mean(subset(vh2o, week == 3)$mass, na.rm = T)),
    aes(x = mass, y = ecc),
    alpha = 0.5
  ) +
  geom_smooth(
    method = "lm", colour = "black",
    linetype = "dashed", se = FALSE
  ) +
  xlab("Body Mass (g)") +
  ylab("Evaporative Cooling\nEfficiency (EHL/RMR)") +
  theme_classic() +
  theme(
    axis.title = element_text(family = "Noto Sans"),
    axis.text = element_text(family = "Noto Sans")
  )

efficiency3WeeksTarsusPlot <-
  expand.grid(
    "tarsus" = with(
      efficiencyModel3Weeks$data,
      seq(min(tarsus), max(tarsus), by = 1)
    ),
    "mass" = 0,
    "pretreatment" = "B",
    "batch" = "A"
  ) %>%
  mutate(
    "ecc" = predict(efficiencyModel3Weeks,
      newdata = .,
      re_form = NA, robust = TRUE,
      resp = "ecc"
    )[, "Estimate"],
    "SE" = predict(efficiencyModel3Weeks,
      newdata = .,
      re_form = NA, robust = TRUE,
      resp = "ecc"
    )[, "Est.Error"]
  ) %>%
```

```
  mutate(tarsus = tarsus +
           mean(subset(vh2o, week == 3)$tarsusLengthMean, na.rm = T)) %>%
  ggplot(aes(x = tarsus, y = ecc)) +
  geom_ribbon(aes(ymin = ecc - SE, ymax = ecc + SE),
    alpha = 0.5, fill = "#DECCC1"
  ) +
  geom_point(
    data = efficiencyModel3Weeks$data %>%
      mutate(tarsus = tarsus +
        mean(subset(vh2o, week == 3)$tarsusLengthMean, na.rm = T)),
    aes(x = tarsus, y = ecc),
    alpha = 0.5
  ) +
  geom_smooth(
    method = "lm", colour = "black",
    linetype = "dashed", se = FALSE
  ) +
  xlab("Tarsus Length (mm)") +
  ylab("Evaporative Cooling\nEfficiency (EHL/RMR)") +
  theme_classic() +
  theme(
    axis.title = element_text(family = "Noto Sans"),
    axis.text = element_text(family = "Noto Sans")
  )

(heatLoss3WeeksMassPlot + heatLoss3WeeksTarsusPlot) /
  (efficiency3WeeksMassPlot + efficiency3WeeksTarsusPlot) +
  plot_annotation(tag_level = "A")
```

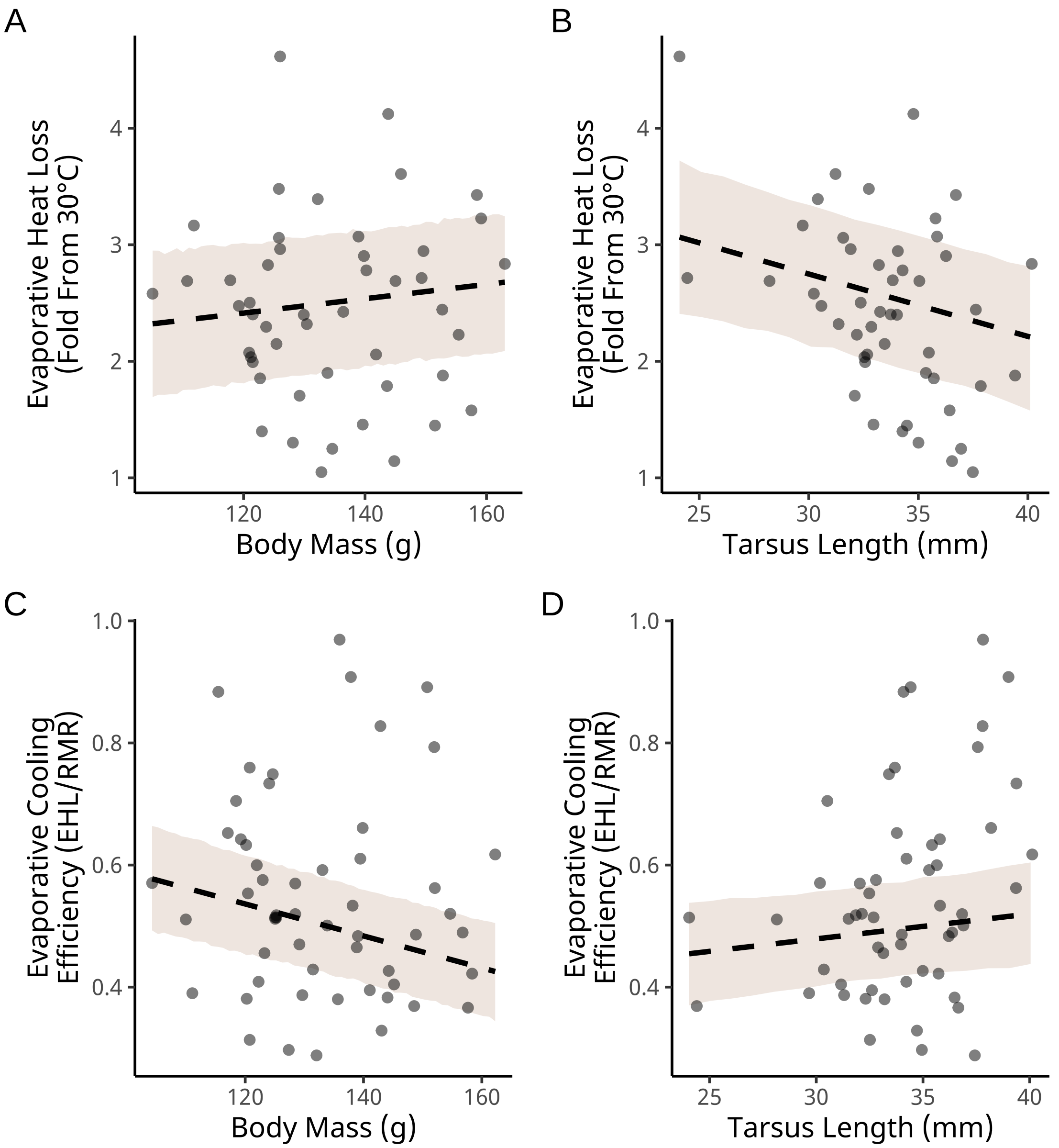


***Figure 15:*** *Effects of morphology on evaporative heat loss responses (panels A and B) and evaporative cooling efficiency (panels C and D) in developing Japanese quail (three weeks of age; n = 48). Evaporative heat loss responses represent fold increases in evaporative heat loss, in Watts, between 30°C and 40°C. Evaporative cooling efficiency represents the ratio between evaporative heat loss (in Watts) and metabolic heat production (in Watts). Dashed lines indicate predicted relationships, as estimated from Bayesian path analyses. Ribbons represents +/- one standard error around predicted lines of best fit. Small dots indicate raw data points.*

```
heatLoss3WeeksTreatmentPlot <-
  expand.grid(
    "mass" = 0,
    "tarsus" = 0,
    "pretreatment" = c("A", "B", "C")
  ) %>%
  mutate(
    "foldEhl" = predict(ehlModel3Weeks,
      newdata = .,
```

```
      re_form = NA, robust = TRUE,
      resp = "foldEhl"
    )[, "Estimate"],
    "SE" = predict(ehlModel3Weeks,
      newdata = .,
      re_form = NA, robust = TRUE,
      resp = "foldEhl"
    )[, "Est.Error"]
  ) %>%
  mutate(pretreatment = factor(pretreatment, levels = c("A", "B", "C"))) %>%
  ggplot(aes(x = pretreatment, y = foldEhl, fill = pretreatment)) +
  geom_point(
    data = ehlModel3Weeks$data %>%
      mutate(pretreatment = factor(pretreatment, levels = c("A", "B", "C"))),
    aes(x = pretreatment, y = foldEhl),
    alpha = 0.5, position = position_jitter(width = 0.25)
  ) +
  geom_errorbar(aes(ymin = foldEhl - SE, ymax = foldEhl + SE),
    colour = "black", width = 0.25
  ) +
  geom_point(
    pch = 21, colour = "black", size = 4
  ) +
  scale_x_discrete(name = "Rearing Treatment",
                   labels = c("Cold (10°C)",
                              "Mild (20°C)",
                              "Warm (30°C)")
                   ) +
  ylab("Evaporative Heat Loss\nResponse (Fold From 30°C)") +
  scale_fill_manual(values = c("#7BB4E3", "black", "#CD5C5C")) +
  theme_classic() +
  theme(
    axis.title = element_text(family = "Noto Sans"),
    axis.text = element_text(family = "Noto Sans"),
    legend.position = "none"
  )

efficiency3WeeksTreatmentPlot <-
  expand.grid(
    "mass" = 0,
    "tarsus" = 0,
    "pretreatment" = c("A", "B", "C"),
    "batch" = "A"
  ) %>%
  mutate(
    "ecc" = predict(efficiencyModel3Weeks,
      newdata = .,
      re_form = NA, robust = TRUE,
      resp = "ecc"
    )[, "Estimate"],
    "SE" = predict(efficiencyModel3Weeks,
      newdata = .,
      re_form = NA, robust = TRUE,
      resp = "ecc"
    )[, "Est.Error"]
  ) %>%
  mutate(pretreatment = factor(pretreatment, levels = c("A", "B", "C"))) %>%
  ggplot(aes(x = pretreatment, y = ecc, fill = pretreatment)) +
  geom_point(
    data = efficiencyModel3Weeks$data %>%
      mutate(pretreatment = factor(pretreatment, levels = c("A", "B", "C"))),
    aes(x = pretreatment, y = ecc),
    alpha = 0.5, position = position_jitter(width = 0.25)
  ) +
  geom_errorbar(aes(ymin = ecc - SE, ymax = ecc + SE),
    colour = "black", width = 0.25
  ) +
```

```
  geom_point(
    pch = 21, colour = "black", size = 4
  ) +
  scale_x_discrete(name = "Rearing Treatment",
                   labels = c("Cold (10°C)",
                              "Mild (20°C)",
                              "Warm (30°C)")
                   ) +
  ylab("Evaporative Cooling\nEfficiency (EHL/RMR)") +
  scale_fill_manual(values = c("#7BB4E3", "black", "#CD5C5C")) +
  theme_classic() +
  theme(
    axis.title = element_text(family = "Noto Sans"),
    axis.text = element_text(family = "Noto Sans"),
    legend.position = "none"
  )

(heatLoss3WeeksTreatmentPlot / efficiency3WeeksTreatmentPlot) +
  plot_annotation(tag_levels = "A")
```

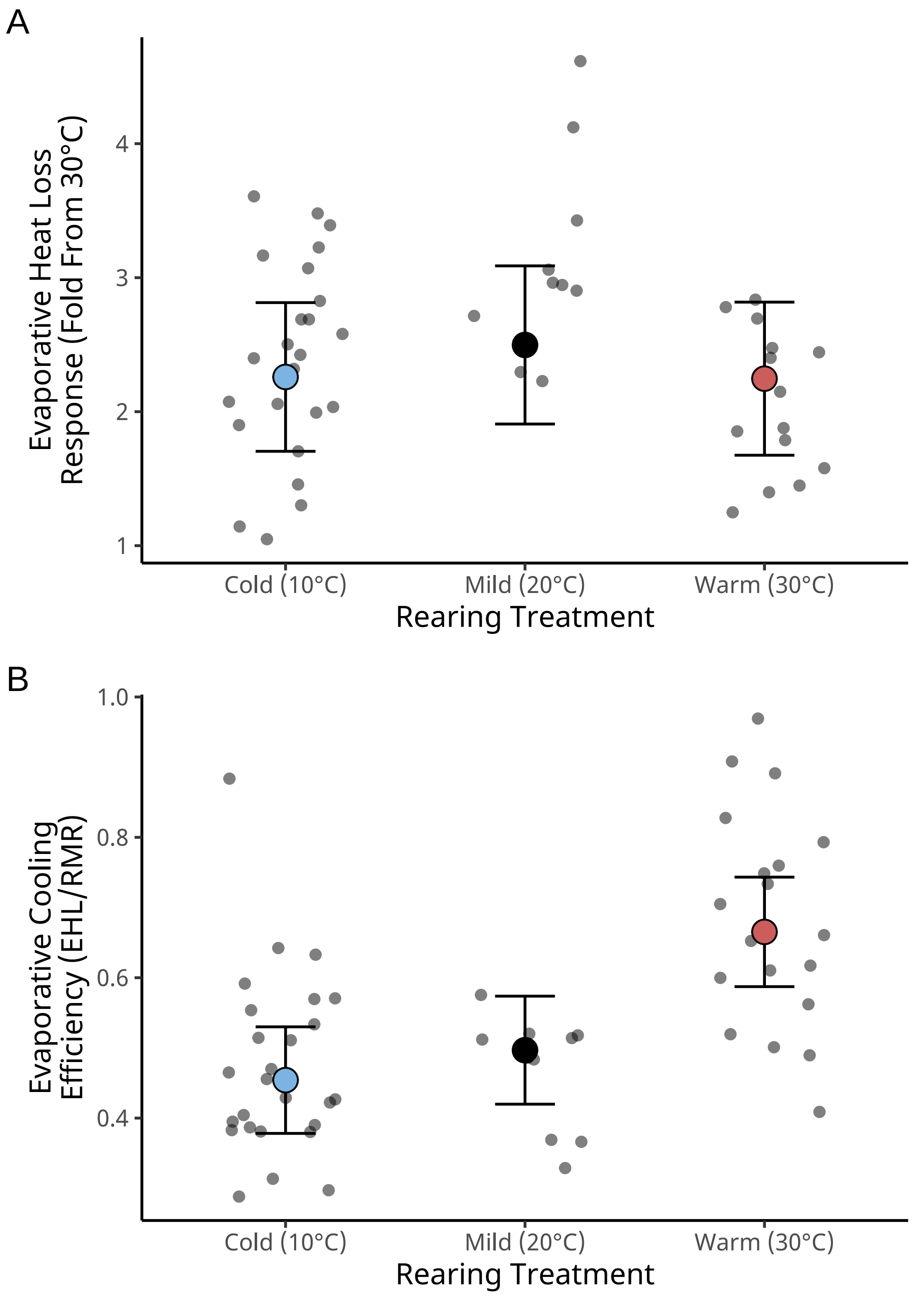


***Figure 16:*** *Effects of rearing temperature on evaporative heat loss responses (panel A) and evaporative cooling efficiency (panel B) in developing Japanese quail (three weeks of age; n = 48). Evaporative heat loss responses represent fold increases in evaporative heat loss, in Watts, between 30°C and 40°C. Evaporative cooling efficiency represents the ratio between evaporative heat loss (in Watts) and metabolic heat production (in Watts). Dashed lines indicate predicted relationships, as estimated from Bayesian path analyses. Ribbons represents +/- one standard error around predicted lines of best fit. Small dots indicate raw data points.*

To quantify direct and indirect effects of morphology or rearing treatment on final evaporative cooling phenotypes, we combine posterior estimates from each model within our path analyses. To ease interpretation and cross-comparability of effects, however, we first scale individual coefficients to represent the effect of changing a given predictor variable by one standard deviation (or categorical level) on a change, in standard deviations, of the response variable. Direct and indirect effects are both visualised as densities and summarised at medians (± quantile intervals).

```
# Scaling coefficients

scaledBetasHeatLoss3 <- as.data.frame(ehlModel3Weeks) %>%
  mutate(
    b_mass_pretreatmentA = b_mass_pretreatmentA /
      sd(ehlModel3Weeks$data$mass),
    b_mass_pretreatmentC = b_mass_pretreatmentC /
      sd(ehlModel3Weeks$data$mass),
    b_tarsus_pretreatmentA = b_tarsus_pretreatmentA /
      sd(ehlModel3Weeks$data$tarsus),
    b_tarsus_pretreatmentC = b_tarsus_pretreatmentC /
      sd(ehlModel3Weeks$data$tarsus),
    b_tarsus_mass =
      (b_tarsus_mass * sd(ehlModel3Weeks$data$mass)) /
        sd(ehlModel3Weeks$data$tarsus),
    b_foldEhl_mass =
      (b_foldEhl_mass * sd(ehlModel3Weeks$data$mass)) /
        sd(ehlModel3Weeks$data$foldEhl),
    b_foldEhl_tarsus =
      (b_foldEhl_tarsus * sd(ehlModel3Weeks$data$tarsus)) /
        sd(ehlModel3Weeks$data$foldEhl),
    b_foldEhl_pretreatmentA = b_foldEhl_pretreatmentA /
      sd(ehlModel3Weeks$data$foldEhl),
    b_foldEhl_pretreatmentC = b_foldEhl_pretreatmentC /
      sd(ehlModel3Weeks$data$foldEhl),
  )

scaledBetasEfficiency3 <- as.data.frame(efficiencyModel3Weeks) %>%
  mutate(
    b_mass_pretreatmentA = b_mass_pretreatmentA /
      sd(efficiencyModel3Weeks$data$mass),
    b_mass_pretreatmentC = b_mass_pretreatmentC /
      sd(efficiencyModel3Weeks$data$mass),
    b_tarsus_pretreatmentA = b_tarsus_pretreatmentA /
      sd(efficiencyModel3Weeks$data$tarsus),
    b_tarsus_pretreatmentC = b_tarsus_pretreatmentC /
      sd(efficiencyModel3Weeks$data$tarsus),
    b_tarsus_mass =
      (b_tarsus_mass * sd(efficiencyModel3Weeks$data$mass)) /
        sd(efficiencyModel3Weeks$data$tarsus),
    b_ecc_mass =
      (b_ecc_mass * sd(efficiencyModel3Weeks$data$mass)) /
        sd(efficiencyModel3Weeks$data$ecc),
    b_ecc_tarsus =
      (b_ecc_tarsus * sd(efficiencyModel3Weeks$data$tarsus)) /
        sd(efficiencyModel3Weeks$data$ecc),
    b_ecc_pretreatmentA = b_ecc_pretreatmentA /
      sd(efficiencyModel3Weeks$data$ecc),
    b_ecc_pretreatmentC = b_ecc_pretreatmentC /
      sd(efficiencyModel3Weeks$data$ecc),
  )

# Calculating effects

fullEffectHeatLoss3 <- scaledBetasHeatLoss3 %>%
  mutate("Effects" = "Direct Effects") %>%
  mutate(
    "Body Mass" = b_foldEhl_mass,
    "Tarsus Length" = b_foldEhl_tarsus,
    "Cold Rearing\n(10°C)" = b_foldEhl_pretreatmentA,
    "Warm Rearing\n(30°C)" = b_foldEhl_pretreatmentC
  ) %>%
  select(
    Effects, `Body Mass`, `Tarsus Length`,
    `Cold Rearing\n(10°C)`,
```

```
    `Warm Rearing\n(30°C)`
  ) %>%
  rbind(
    .,
    scaledBetasHeatLoss3 %>%
      mutate("Effects" = "Indirect Effects") %>%
      mutate(
        "Body Mass" = b_tarsus_mass *
          b_foldEhl_tarsus,
        "Tarsus Length" = NA,
        "Cold Rearing\n(10°C)" =
          b_tarsus_pretreatmentA * b_foldEhl_tarsus +
            b_mass_pretreatmentA * b_foldEhl_mass +
            b_mass_pretreatmentA * b_tarsus_mass *
              b_foldEhl_tarsus,
        "Warm Rearing\n(30°C)" =
          b_tarsus_pretreatmentC * b_foldEhl_tarsus +
            b_mass_pretreatmentC * b_foldEhl_mass +
            b_mass_pretreatmentC * b_tarsus_mass *
              b_foldEhl_tarsus,
      ) %>%
      select(
        Effects, `Body Mass`, `Tarsus Length`,
        `Cold Rearing\n(10°C)`, `Warm Rearing\n(30°C)`
      )
  ) %>%
  rbind(., scaledBetasHeatLoss3 %>%
    mutate("Effects" = "Total Effects") %>%
    mutate(
      "Body Mass" =
        b_foldEhl_mass +
          b_tarsus_mass * b_foldEhl_tarsus,
      "Tarsus Length" =
        b_foldEhl_tarsus,
      "Cold Rearing\n(10°C)" =
        b_foldEhl_pretreatmentA +
          b_mass_pretreatmentA *
            b_foldEhl_mass +
          b_tarsus_pretreatmentA *
            b_foldEhl_tarsus +
          b_mass_pretreatmentA *
            b_tarsus_mass * b_foldEhl_tarsus,
      "Warm Rearing\n(30°C)" =
        b_foldEhl_pretreatmentC +
          b_mass_pretreatmentC *
            b_foldEhl_mass +
          b_tarsus_pretreatmentC *
            b_foldEhl_tarsus +
          b_mass_pretreatmentC *
            b_tarsus_mass * b_foldEhl_tarsus,
    ) %>%
    select(
      Effects, `Body Mass`, `Tarsus Length`,
      `Cold Rearing\n(10°C)`, `Warm Rearing\n(30°C)`
    )) %>%
  pivot_longer(c(-Effects), names_to = "var", values_to = "values") %>%
  mutate(var = factor(var,
    levels = c(
      "Tarsus Length",
      "Body Mass",
      "Warm Rearing\n(30°C)",
      "Cold Rearing\n(10°C)"
    )
  ))

fullEffectEfficiency3 <- scaledBetasEfficiency3 %>%
  mutate("Effects" = "Direct Effects") %>%
```

```
mutate(
  "Body Mass" = b_ecc_mass,
  "Tarsus Length" = b_ecc_tarsus,
  "Cold Rearing\n(10°C)" = b_ecc_pretreatmentA,
  "Warm Rearing\n(30°C)" = b_ecc_pretreatmentC
) %>%
select(
  Effects, `Body Mass`, `Tarsus Length`,
  `Cold Rearing\n(10°C)`,
  `Warm Rearing\n(30°C)`
) %>%
rbind(
  .,
  scaledBetasEfficiency3 %>%
    mutate("Effects" = "Indirect Effects") %>%
    mutate(
      "Body Mass" = b_tarsus_mass *
        b_ecc_tarsus,
      "Tarsus Length" = NA,
      "Cold Rearing\n(10°C)" =
        b_tarsus_pretreatmentA * b_ecc_tarsus +
          b_mass_pretreatmentA * b_ecc_mass +
          b_mass_pretreatmentA * b_tarsus_mass *
            b_ecc_tarsus,
      "Warm Rearing\n(30°C)" =
        b_tarsus_pretreatmentC * b_ecc_tarsus +
          b_mass_pretreatmentC * b_ecc_mass +
          b_mass_pretreatmentC * b_tarsus_mass *
            b_ecc_tarsus,
    ) %>%
    select(
      Effects, `Body Mass`, `Tarsus Length`,
      `Cold Rearing\n(10°C)`, `Warm Rearing\n(30°C)`
    )
) %>%
rbind(., scaledBetasEfficiency3 %>%
  mutate("Effects" = "Total Effects") %>%
  mutate(
    "Body Mass" =
      b_ecc_mass +
        b_tarsus_mass * b_ecc_tarsus,
    "Tarsus Length" =
      b_ecc_tarsus,
    "Cold Rearing\n(10°C)" =
      b_ecc_pretreatmentA +
        b_mass_pretreatmentA *
          b_ecc_mass +
        b_tarsus_pretreatmentA *
          b_ecc_tarsus +
        b_mass_pretreatmentA *
          b_tarsus_mass * b_ecc_tarsus,
    "Warm Rearing\n(30°C)" =
      b_ecc_pretreatmentC +
        b_mass_pretreatmentC *
          b_ecc_mass +
        b_tarsus_pretreatmentC *
          b_ecc_tarsus +
        b_mass_pretreatmentC *
          b_tarsus_mass * b_ecc_tarsus,
  ) %>%
  select(
    Effects, `Body Mass`, `Tarsus Length`,
    `Cold Rearing\n(10°C)`, `Warm Rearing\n(30°C)`
  )) %>%
pivot_longer(c(-Effects), names_to = "var", values_to = "values") %>%
mutate(var = factor(var,
  levels = c(
```

```
      "Tarsus Length",
      "Body Mass",
      "Warm Rearing\n(30°C)",
      "Cold Rearing\n(10°C)"
    )
  ))

# Plotting effects

fullEffectHeatLoss3Plot <- fullEffectHeatLoss3 %>%
  ggplot(aes(x = values, y = var, fill = var)) +
  facet_wrap(~Effects) +
  stat_halfeye(normalize = "xy", colour = "black", alpha = 0.7) +
  geom_vline(
    xintercept = 0, linetype = "dashed",
    colour = "black"
  ) +
  xlab("Effect on Fold Evaporative\nHeat Loss (standard deviations)") +
  scale_fill_manual(values = c("black", "grey50", "#CD5C5C", "#7BB4E3")) +
  theme_classic() +
  theme(
    legend.position = "none", axis.title.y = element_blank(),
    axis.text.y = element_text(
      size = 11, colour = "black",
      family = "Noto Sans"
    ),
    axis.title.x = element_text(family = "Noto Sans", hjust = -0.005),
    axis.text.x = element_text(family = "Noto Sans")
  )

fullEffectEfficiency3Plot <- fullEffectEfficiency3 %>%
  ggplot(aes(x = values, y = var, fill = var)) +
  facet_wrap(~Effects) +
  stat_halfeye(normalize = "xy", colour = "black", alpha = 0.7) +
  geom_vline(
    xintercept = 0, linetype = "dashed",
    colour = "black"
  ) +
  xlab("Effect on Evaporative Cooling\nEfficiency (standard deviations)") +
  scale_fill_manual(values = c("black", "grey50", "#CD5C5C", "#7BB4E3")) +
  theme_classic() +
  theme(
    legend.position = "none", axis.title.y = element_blank(),
    axis.text.y = element_text(
      size = 11, colour = "black",
      family = "Noto Sans"
    ),
    axis.title.x = element_text(family = "Noto Sans", hjust = -0.005),
    axis.text.x = element_text(family = "Noto Sans")
  )

(fullEffectHeatLoss3Plot /
  fullEffectEfficiency3Plot) +
  plot_annotation(tag_level = "A")
```

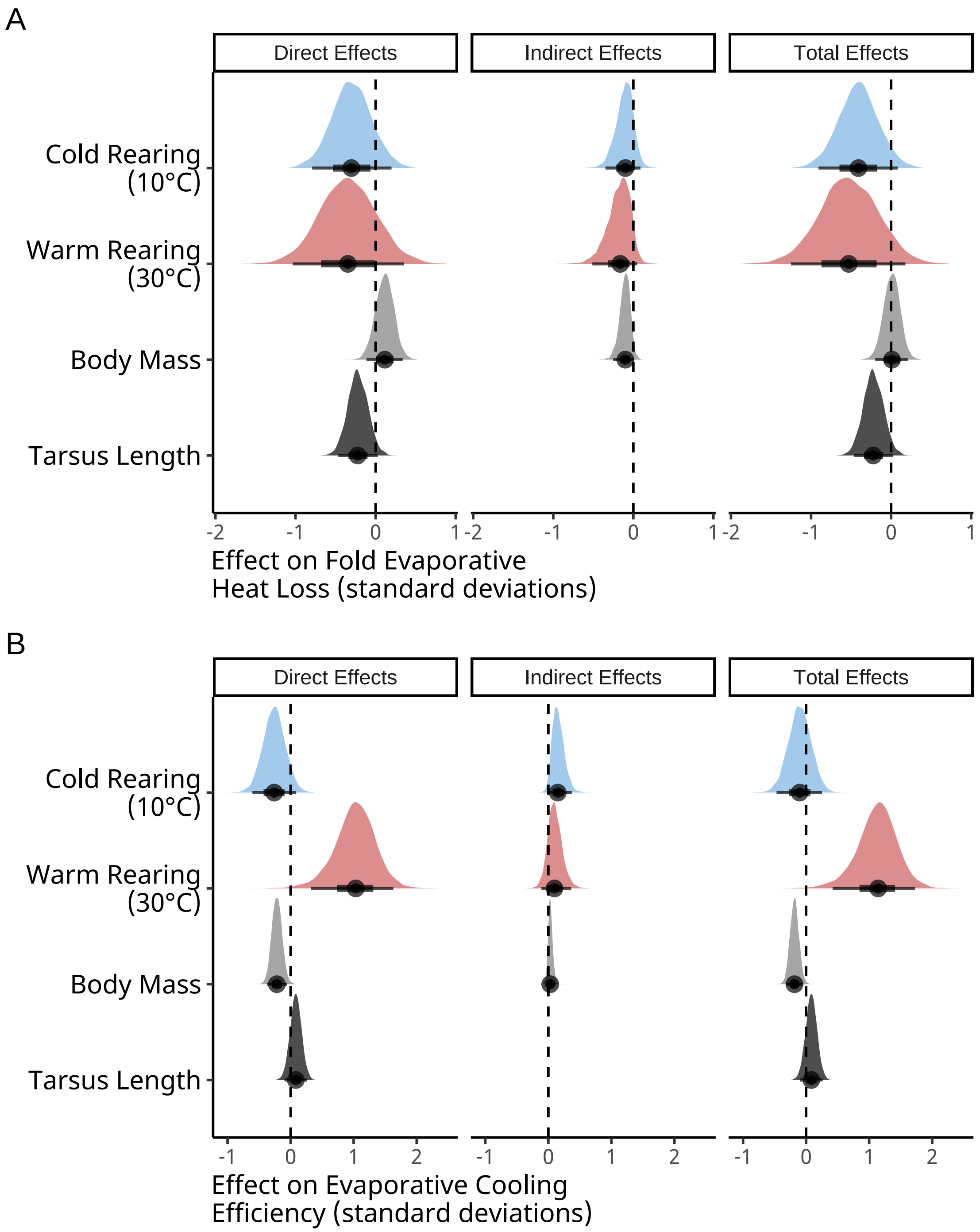


***Figure 17:*** *Direct and indirect effects of rearing temperature and morphology (here, body mass and tarsus length) on evaporative heat loss responses (panel A) and evaporative cooling efficiency (panel B) in developing Japanese quail (three weeks of age; n = 48). Evaporative heat loss responses represent fold increases in evaporative heat loss, in Watts, between 30°C and 40°C. Evaporative cooling efficiency represents the ratio between evaporative heat loss (in Watts) and metabolic heat production (in Watts). Effects indicate how much, in standard deviations, a response variables would be altered by changing a predictor variable by one standard deviation. Densities are derived from posteriors of Bayesian path analyses. Dashed lines indicate 0.*

```
# And printing outcomes

caption <- paste0("Direct, indirect, and total effects of morphology and ",
                  "rearing temperature on fold evaporative heat loss at 40°C",
                  "(relative to 30°C) of three week old Japanese quail. ",
                  "Effects are derived from ",
                  "a Bayesian path analysis and represent those predicted for a ",
                  "change in one standard deviation (or categorical level) of ",
                  "a given predictor on the standard deviation of metabolic ",
                  "slopes. Estimates indicate posterior medians and credible ",
                  "intervels (CIs) indicate quantile intervals."
)

heatLossResultsScaled3 <-
  fullEffectHeatLoss3 %>%
  filter(!is.na(values) & !is.nan(values)) %>%
  group_by(Effects, var) %>%
  summarise(
    "Estimate" = median(values),
    "50\\% CIs" = paste0(
      "[",
      round(
        quantile(values, probs = 0.1, type = 8),
        digits = 4
      ),
      ", ",
      round(
        quantile(values, probs = 0.9, type = 8),
        digits = 4
      ),
      "]"
    ),
    "95\\% CIs" = paste0(
      "[",
      round(
        quantile(values, probs = 0.025, type = 8),
        digits = 4
      ),
      ", ",
      round(
        quantile(values, probs = 0.975, type = 8),
        digits = 4
      ),
      "]"
    )
  ) %>%
  mutate("Effects" = gsub("[[:space:]].*", "", Effects)) %>%
  select(
    "Predictor" = "var", "Effect Level" = "Effects",
    Estimate, `50\\% CIs`, `95\\% CIs`
  ) %>%
  arrange(Predictor, `Effect Level`) %>%
  kbl(.,
    longtable = T, booktabs = T, format = "latex", escape = FALSE,
    caption = caption
  ) %>%
  column_spec(column = c(1:2), width = "2.2cm") %>%
  column_spec(column = c(3:10), width = "1.9cm") %>%
  kable_styling(latex_options = "striped")

heatLossResultsScaled3
```

***Table 6:*** *Direct, indirect, and total effects of morphology and rearing temperature on fold evaporative heat loss at 40°C(relative to 30°C) of three week old Japanese quail. Effects are derived from a Bayesian path analysis and represent those predicted for a change in one standard deviation (or categorical level) of a given predictor on the standard deviation of metabolic slopes. Estimates indicate posterior medians and credible intervels (CIs) indicate quantile intervals.*

| Predictor | Effect Level | Estimate | 50% CIs | 95% CIs |
|---|---|---|---|---|
| Tarsus Length | Direct | -0.2236154 | [-0.382, -0.059] | [-0.4664, 0.0257] |
| Tarsus Length | Total | -0.2236154 | [-0.382, -0.059] | [-0.4664, 0.0257] |
| Body Mass | Direct | 0.1159303 | [-0.0326, 0.2572] | [-0.1142, 0.3388] |
| Body Mass | Indirect | -0.1003261 | [-0.1919, -0.0254] | [-0.2526, 0.0111] |
| Body Mass | Total | 0.0109036 | [-0.1231, 0.1381] | [-0.1987, 0.2078] |
| Warm Rearing (30°C) | Direct | -0.3469091 | [-0.7881, 0.1064] | [-1.037, 0.3536] |
| Warm Rearing (30°C) | Indirect | -0.1641438 | [-0.3737, -0.0173] | [-0.5121, 0.0479] |
| Warm Rearing (30°C) | Total | -0.5293582 | [-0.9917, -0.0599] | [-1.2497, 0.1787] |
| Cold Rearing (10°C) | Direct | -0.3024596 | [-0.6152, 0.0196] | [-0.7926, 0.2] |
| Cold Rearing (10°C) | Indirect | -0.0996831 | [-0.2577, 0.021] | [-0.3484, 0.0893] |
| Cold Rearing (10°C) | Total | -0.4085589 | [-0.7255, -0.0894] | [-0.9079, 0.0818] |

```r
caption <- paste0("Direct, indirect, and total effects of morphology and ",
                  "rearing temperature on evaporative cooling efficiency ",
                  "at 40°C  of three week old Japanese quail. ",
                  "Evaporative cooling efficiency represents the ratio of ",
                  "evaporative heat loss (in W) to metabolic heat production ",
                  "(again, in W). Effects are derived from ",
                  "a Bayesian path analysis and represent those predicted for a ",
                  "change in one standard deviation (or categorical level) of ",
                  "a given predictor on the standard deviation of metabolic ",
                  "slopes. Estimates indicate posterior medians and credible ",
                  "intervels (CIs) indicate quantile intervals."
)

efficiencyResultsScaled3 <-
  fullEffectEfficiency3 %>%
  filter(!is.na(values) & !is.nan(values)) %>%
  group_by(Effects, var) %>%
  summarise(
    "Estimate" = median(values),
    "50\\% CIs" = paste0(
      "[",
      round(
        quantile(values, probs = 0.1, type = 8),
        digits = 4
      ),
      ", ",
      round(
        quantile(values, probs = 0.9, type = 8),
        digits = 4
      ),
      "]"
    ),
    "95\\% CIs" = paste0(
      "[",
```

```
      round(
        quantile(values, probs = 0.025, type = 8),
        digits = 4
      ),
      ", ",
      round(
        quantile(values, probs = 0.975, type = 8),
        digits = 4
      ),
      "]"
    )
  ) %>%
  mutate("Effects" = gsub("[[:space:]].*", "", Effects)) %>%
  select(
    "Predictor" = "var", "Effect Level" = "Effects",
    Estimate, `50\\% CIs`, `95\\% CIs`
  ) %>%
  arrange(Predictor, `Effect Level`) %>%
  kbl(.,
    longtable = T, booktabs = T, format = "latex", escape = FALSE,
    caption = caption
  ) %>%
  column_spec(column = c(1:2), width = "2.2cm") %>%
  column_spec(column = c(3:10), width = "1.9cm") %>%
  kable_styling(latex_options = "striped")

efficiencyResultsScaled3
```

***Table 7:*** *Direct, indirect, and total effects of morphology and rearing temperature on evaporative cooling efficiency at 40°C of three week old Japanese quail. Evaporative cooling efficiency represents the ratio of evaporative heat loss (in W) to metabolic heat production (again, in W). Effects are derived from a Bayesian path analysis and represent those predicted for a change in one standard deviation (or categorical level) of a given predictor on the standard deviation of metabolic slopes. Estimates indicate posterior medians and credible intervels (CIs) indicate quantile intervals.*

| Predictor | Effect Level | Estimate | 50% CIs | 95% CIs |
|---|---|---|---|---|
| Tarsus Length | Direct | 0.0830262 | [-0.0338, 0.1968] | [-0.0967, 0.2624] |
| Tarsus Length | Total | 0.0830262 | [-0.0338, 0.1968] | [-0.0967, 0.2624] |
| Body Mass | Direct | -0.2177851 | [-0.3198, -0.1141] | [-0.3721, -0.0589] |
| Body Mass | Indirect | 0.0300385 | [-0.0122, 0.08] | [-0.0382, 0.1135] |
| Body Mass | Total | -0.1850131 | [-0.2738, -0.0953] | [-0.3201, -0.0492] |
| Warm Rearing (30°C) | Direct | 1.0336979 | [0.6131, 1.4099] | [0.3263, 1.631] |
| Warm Rearing (30°C) | Indirect | 0.0971794 | [-0.0348, 0.2538] | [-0.1107, 0.3623] |
| Warm Rearing (30°C) | Total | 1.1419407 | [0.726, 1.5037] | [0.4201, 1.7256] |
| Cold Rearing (10°C) | Direct | -0.2606734 | [-0.4863, -0.0345] | [-0.606, 0.0892] |
| Cold Rearing (10°C) | Indirect | 0.1485293 | [0.0448, 0.2849] | [-0.0011, 0.3708] |
| Cold Rearing (10°C) | Total | -0.1024252 | [-0.3353, 0.1291] | [-0.4683, 0.2512] |

Finally, we estimate the tentative consequences of misalignment with Bergmann's and Allen's rule in the heat (i.e. by having an atypically large body or atypically short tarsi). Consequences here represent changes in evaporative heat loss responses and evaporative cooling efficiency at 40°C relative to average.

```
data.frame(
  "Size" = c("Average", "Large (2x s.d. > mean)"),
  "pretreatment" = "B",
  "mass" = c(
    mean(ehlModel3Weeks$data$mass, na.rm = T),
    mean(ehlModel3Weeks$data$mass, na.rm = T) +
      2 * sd(ehlModel3Weeks$data$mass, na.rm = T)
  ),
  "tarsus" = 0,
  "batch" = "A"
) %>%
  mutate(
    "ehl" = predict(ehlModel3Weeks,
      newdata = .,
      resp = "foldEhl",
      re_form = NA,
      robust = TRUE
    )[, "Estimate"],
    "se1" = predict(ehlModel3Weeks,
      newdata = .,
      resp = "foldEhl",
      re_form = NA,
      robust = TRUE
    )[, "Est.Error"],
    "ecc" = predict(efficiencyModel3Weeks,
      newdata = .,
      resp = "ecc",
      re_form = NA,
      robust = TRUE
    )[, "Estimate"],
    "se2" = predict(efficiencyModel3Weeks,
      newdata = .,
      resp = "ecc",
      re_form = NA,
      robust = TRUE
    )[, "Est.Error"]
  ) %>%
  mutate_if(is.numeric, round, digits = 4) %>%
  mutate(
    "Body Size" = Size,
    "Evaporative Heat Loss" = paste0(ehl, " [",
                                     se1, "]"
    ),
    "Evaporative Cooling Efficiency" = paste0(ecc, " [",
                                     se2, "]"
    )
  ) %>%
  select(`Body Size`, `Evaporative Heat Loss`, `Evaporative Cooling Efficiency`)
```

```
## # A tibble: 2 x 3
##   `Body Size`            `Evaporative Heat Loss` Evaporative Cooling Efficienc~1
##   <chr>                  <chr>                   <chr>
## 1 Average                2.5116 [0.58]           0.4968 [0.0762]
## 2 Large (2x s.d. > mean) 2.6679 [0.5957]         0.425 [0.0769]
## # i abbreviated name: 1: `Evaporative Cooling Efficiency`
```

```
data.frame(
  "Size" = c("Average", "Short (2x s.d. < mean)"),
  "pretreatment" = "B",
  "tarsus" = c(
    mean(ehlModel3Weeks$data$tarsus, na.rm = T),
    mean(ehlModel3Weeks$data$tarsus, na.rm = T) -
      2 * sd(ehlModel3Weeks$data$tarsus, na.rm = T)
  ),
  "mass" = 0,
  "batch" = "A"
) %>%
  mutate(
```

```
    "ehl" = predict(ehlModel3Weeks,
      newdata = .,
      resp = "foldEhl",
      re_form = NA,
      robust = TRUE
    )[, "Estimate"],
    "se1" = predict(ehlModel3Weeks,
      newdata = .,
      resp = "foldEhl",
      re_form = NA,
      robust = TRUE
    )[, "Est.Error"],
    "ecc" = predict(efficiencyModel3Weeks,
      newdata = .,
      resp = "ecc",
      re_form = NA,
      robust = TRUE
    )[, "Estimate"],
    "se2" = predict(efficiencyModel3Weeks,
      newdata = .,
      resp = "ecc",
      re_form = NA,
      robust = TRUE
    )[, "Est.Error"]
  ) %>%
  mutate_if(is.numeric, round, digits = 4) %>%
  mutate(
    "Tarsus Length" = Size,
    "Evaporative Heat Loss" = paste0(ehl, " [",
                                     se1, "]"
    ),
    "Evaporative Cooling Efficiency" = paste0(ecc, " [",
                                     se2, "]"
    )
  ) %>%
  select(`Tarsus Length`, `Evaporative Heat Loss`,
         `Evaporative Cooling Efficiency`)
```

```
## # A tibble: 2 x 3
##   `Tarsus Length`        `Evaporative Heat Loss` Evaporative Cooling Efficienc~1
##   <chr>                  <chr>                   <chr>
## 1 Average                2.5654 [0.5825]         0.4937 [0.0775]
## 2 Short (2x s.d. < mean) 2.8992 [0.6095]         0.4653 [0.0804]
## # i abbreviated name: 1: `Evaporative Cooling Efficiency`
```

##### Effects in mature individuals

In this subsection, we evaluate effects of rearing temperature and morphology on evaporative heat loss responses and evaporative cooling efficiency in mature Japanese quail (here, eight weeks of age). To do so, we repeat our above described path analyses but while: (1) only using data obtained from our mature quail, and (2) broadening and shifting our model priors to account for known changes in morphology and physiology observed between early development and maturity (see Persson et al, 2024). Model equations are repeated here for convenience.

$$Body\ Mass_j \sim \beta_{a0} + \beta_{a1} \cdot Cold\ Rearing_j + \beta_{a2} \cdot Warm\ Rearing_j + \\ \mu_{0a} + \epsilon_a$$

$$Tarsus\ Length_j \sim \beta_{b0} + \beta_{b1} \cdot Cold\ Rearing_j + \beta_{b2} \cdot Warm\ Rearing_j + \beta_{b3} \cdot Body\ Mass_j + \\ \mu_{0b} + \epsilon_b$$

and either:

$$Fold\ Evaporative\ Heat\ Loss_j \sim \beta_{c0} + \beta_{c1} \cdot Cold\ Rearing_j + \beta_{c2} \cdot Warm\ Rearing_j + \beta_{c3} \cdot Body\ Mass_j + \beta_{c4} \cdot Tarsus\ Length_j + \mu_{0c} + \epsilon_c$$

where:

$$Fold\ Evaporative\ Heat\ Loss_j = \frac{Evaporative\ Heat\ Loss_{40°\mathrm{C}j}}{Evaporative\ Heat\ Loss_{30°\mathrm{C}j}}$$

or:

$$Evaporative\ Cooling\ Efficiency_j \sim \beta_{c0} + \beta_{c1} \cdot Cold\ Rearing_j + \beta_{c2} \cdot Warm\ Rearing_j + \beta_{c3} \cdot Body\ Mass_j + \beta_{c4} \cdot Tarsus\ Length_j + \mu_{0c} + \epsilon_c$$

where evaporative cooling efficiency is measured as:

$$Evaporative\ Cooling\ Efficiency_j = \frac{Evaporative\ Heat\ Loss_{40°\mathrm{C}j}}{Metabolic\ Heat\ Production_{40°\mathrm{C}j}}$$

Model terms remain as previously described.

Priors for our new path analyses are therefore as follows:

$$\beta_{a0} \sim \mathcal{N}(0, 10)$$
$$\beta_{a1} \sim \mathcal{N}(0, 25)$$
$$\beta_{a2} \sim \mathcal{N}(0, 25)$$
$$\mu_{0a} \sim exp(2.5)$$
$$\epsilon_a \sim exp(0.05)$$

$$\beta_{b0} \sim \mathcal{N}(0, 3)$$
$$\beta_{b1} \sim \mathcal{N}(0, 3)$$
$$\beta_{b2} \sim \mathcal{N}(0, 3)$$
$$\beta_{b3} \sim S\mathcal{N}(0, 0.25, 5)$$
$$\mu_{0b} \sim exp(2)$$
$$\epsilon_b \sim exp(0.75)$$

and either:

$$\beta_{c0} \sim \mathcal{N}(2.5, 1)$$
$$\beta_{c1} \sim \mathcal{N}(0, 0.5)$$
$$\beta_{c2} \sim \mathcal{N}(0, 0.5)$$
$$\beta_{c3} \sim \mathcal{N}(0, 0.025)$$
$$\beta_{c4} \sim \mathcal{N}(0, 0.05)$$
$$\mu_{0c} \sim \exp(5)$$
$$\epsilon_c \sim \exp(5)$$

for parameters predicting evaporative heat loss responses, and:

$$\beta_{c0} \sim \mathcal{N}(0.75, 0.2)$$
$$\beta_{c1} \sim \mathcal{N}(0, 0.25)$$
$$\beta_{c2} \sim \mathcal{N}(0, 0.25)$$
$$\beta_{c3} \sim \mathcal{N}(0, 0.01)$$
$$\beta_{c4} \sim \mathcal{N}(0, 0.1)$$
$$\mu_{0c} \sim \exp(15)$$
$$\epsilon_c \sim \exp(5)$$

for parameters predicting evaporative cooling efficiency.

Similar to our above analyses, suitability of priors is first assessed using prior predictive checks.

```
ehlModel8WeeksPPCheck <- brm(
  data = vh2o %>%
    filter(week == "8" & Ta %in% c(30, 40)) %>%
    select(Ta, ring, pretreatment,
           mass, tarsusLengthMean, ehl,
           "batch" = exp) %>%
    pivot_wider(
      id_cols = c("ring", "batch",
                  "pretreatment", "mass",
                  "tarsusLengthMean"),
      values_from = "ehl",
      names_from = "Ta"
    ) %>%
    mutate("foldEhl" = `40` / `30`) %>%
    mutate(pretreatment = ifelse(pretreatment == "neutral", "B",
      ifelse(pretreatment == "cold", "A", "C")
    )) %>%
    mutate(pretreatment = factor(pretreatment,
      levels = c("B", "A", "C")
    )) %>%
    distinct() %>%
    mutate(
      mass = mass - mean(mass, na.rm = T),
      tarsus = tarsusLengthMean - mean(tarsusLengthMean, na.rm = T)
    ),
  family = "gaussian",
  bf(mass ~ pretreatment + (1 | batch)) +
    bf(tarsus ~ mass + pretreatment + (1 | batch)) +
    bf(foldEhl ~ tarsus + mass + pretreatment + (1 | batch)) +
    set_rescor(FALSE),
  prior = c(
    set_prior("normal(0, 10)",
        class = "Intercept",
        resp = "mass"
      ),
      set_prior("normal(0, 25)",
        class = "b",
        coef = "pretreatmentA",
        resp = "mass"
      ),
      set_prior("normal(0, 25)",
        class = "b",
        coef = "pretreatmentC",
        resp = "mass"
      ),
```

```
      set_prior("exponential(2.5)",
        class = "sd",
        group = "batch",
        resp = "mass"
      ),
      set_prior("exponential(0.05)",
        class = "sigma",
        resp = "mass"
      ),
      set_prior("normal(0, 3)",
        class = "Intercept",
        resp = "tarsus"
      ),
      set_prior("normal(0, 3)",
        class = "b",
        coef = "pretreatmentA",
        resp = "tarsus"
      ),
      set_prior("normal(0, 3)",
        class = "b",
        coef = "pretreatmentC",
        resp = "tarsus"
      ),
      set_prior("skew_normal(0, 0.25, 5)",
        class = "b",
        coef = "mass",
        resp = "tarsus"
      ),
      set_prior("exponential(2)",
        class = "sd",
        group = "batch",
        resp = "tarsus"
      ),
      set_prior("exponential(0.75)",
        class = "sigma",
        resp = "tarsus"
      ),
    set_prior("normal(2.5, 1)",
      class = "Intercept",
      resp = "foldEhl"
    ),
    set_prior("normal(0, 0.5)",
      class = "b",
      coef = "pretreatmentA",
      resp = "foldEhl"
    ),
    set_prior("normal(0, 0.5)",
      class = "b",
      coef = "pretreatmentC",
      resp = "foldEhl"
    ),
    set_prior("normal(0, 0.025)",
      class = "b",
      coef = "mass",
      resp = "foldEhl"
    ),
    set_prior("normal(0, 0.5)",
      class = "b",
      coef = "tarsus",
      resp = "foldEhl"
    ),
    set_prior("exponential(5)",
      class = "sd",
      group = "batch",
      resp = "foldEhl"
    ),
    set_prior("exponential(5)",
```

```
      class = "sigma",
      resp = "foldEhl"
    )
  ),
  iter = 50000, warmup = 10000, cores = 4, chains = 4, thin = 20,
  control = list(adapt_delta = .97, max_treedepth = 14),
  sample_prior = "only",
  silent = TRUE, refresh = 0,
  file = "./models/heatLossModel8WeeksPPCheck.Rds",
)

pp1 <- pp_check2(ehlModel8WeeksPPCheck, resp = "foldEhl",
                 xlab = "Evaporative Heat\nLoss Responses (Fold From 30°C)") +
  theme(legend.position = "none")

efficiencyModel8WeeksPPCheck <- brm(
  data = vh2o %>%
    filter(week == "8" & Ta == 40) %>%
    select(Ta, ring, pretreatment,
      mass, tarsusLengthMean, ecc,
      "batch" = exp
    ) %>%
    mutate(
      pretreatment =
        ifelse(pretreatment == "neutral", "B",
          ifelse(pretreatment == "cold", "A", "C")
        )
    ) %>%
    mutate(pretreatment = factor(pretreatment,
      levels = c("B", "A", "C")
    )) %>%
    distinct() %>%
    mutate(
      mass = mass - mean(mass, na.rm = T),
      tarsus = tarsusLengthMean -
        mean(tarsusLengthMean, na.rm = T)
    ),
  family = "gaussian",
  bf(mass ~ pretreatment + (1 | batch)) +
    bf(tarsus ~ mass + pretreatment + (1 | batch)) +
    bf(ecc ~ tarsus + mass + pretreatment + (1 | batch)) +
    set_rescor(FALSE),
  prior = c(
    set_prior("normal(0, 10)",
        class = "Intercept",
        resp = "mass"
      ),
      set_prior("normal(0, 25)",
        class = "b",
        coef = "pretreatmentA",
        resp = "mass"
      ),
      set_prior("normal(0, 25)",
        class = "b",
        coef = "pretreatmentC",
        resp = "mass"
      ),
      set_prior("exponential(2.5)",
        class = "sd",
        group = "batch",
        resp = "mass"
      ),
      set_prior("exponential(0.05)",
        class = "sigma",
        resp = "mass"
      ),
      set_prior("normal(0, 3)",
```

```
        class = "Intercept",
        resp = "tarsus"
      ),
      set_prior("normal(0, 3)",
        class = "b",
        coef = "pretreatmentA",
        resp = "tarsus"
      ),
      set_prior("normal(0, 3)",
        class = "b",
        coef = "pretreatmentC",
        resp = "tarsus"
      ),
      set_prior("skew_normal(0, 0.25, 5)",
        class = "b",
        coef = "mass",
        resp = "tarsus"
      ),
      set_prior("exponential(2)",
        class = "sd",
        group = "batch",
        resp = "tarsus"
      ),
      set_prior("exponential(0.75)",
        class = "sigma",
        resp = "tarsus"
      ),
    set_prior("normal(0.75, 0.2)",
      class = "Intercept",
      resp = "ecc"
    ),
    set_prior("normal(0, 0.25)",
      class = "b",
      coef = "pretreatmentA",
      resp = "ecc"
    ),
    set_prior("normal(0, 0.25)",
      class = "b",
      coef = "pretreatmentC",
      resp = "ecc"
    ),
    set_prior("normal(0, 0.01)",
      class = "b",
      coef = "mass",
      resp = "ecc"
    ),
    set_prior("normal(0, 0.1)",
      class = "b",
      coef = "tarsus",
      resp = "ecc"
    ),
    set_prior("exponential(15)",
      class = "sd",
      group = "batch",
      resp = "ecc"
    ),
    set_prior("exponential(5)",
      class = "sigma",
      resp = "ecc"
    )
  ),
  iter = 50000, warmup = 10000, cores = 4, chains = 4, thin = 20,
  control = list(adapt_delta = .97, max_treedepth = 14),
  silent = TRUE, refresh = 0,
  sample_prior = "only",
  file = "./models/efficiencyModel8WeeksPPCheck.Rds"
)
```

```
pp2 <- pp_check2(efficiencyModel8WeeksPPCheck, resp = "ecc",
                 xlab = "Evaporative Cooling\nEfficiency (EHL/RMR)")

pp1 + pp2 + plot_annotation(tag_levels = "A")
```

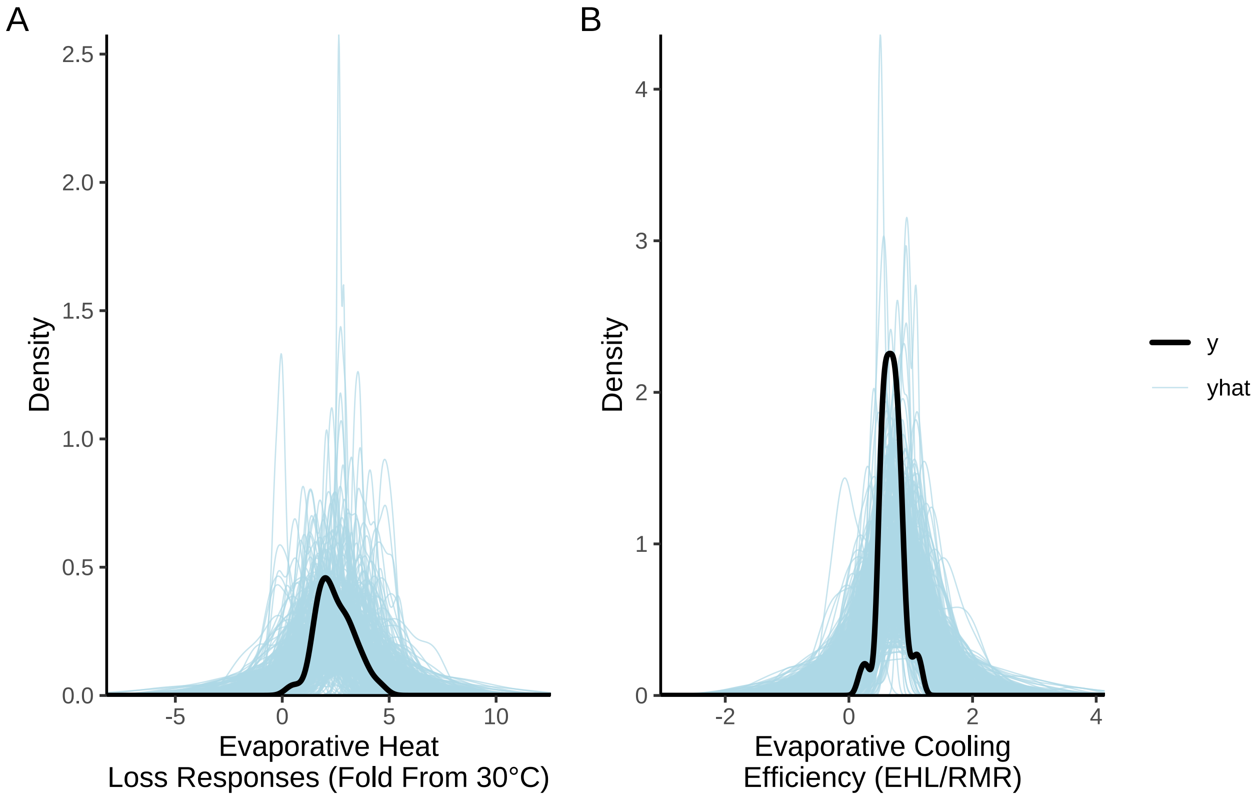


***Figure 18:*** *Prior predictive checks for two Bayesian path analyses ultimately predicting evaporative heat loss ('EHL'; fold from that observed at 30°C in W) and evaporative cooling efficiency ('ECE'; the ratio of evaporative heat-loss to metabolic heat production, each in W) in eight week old Japanese quail. Black lines represent true EHL and ECE densities while blue lines represent densities estimated from model priors alone.*

Predictions from priors generally capture true data densities well. We thus proceed with their use in the construction of full models below.

```
ehlModel8Weeks <- brm(
  data = vh2o %>%
    filter(week == "8" & Ta %in% c(30, 40)) %>%
    select(Ta, ring, pretreatment,
      mass, tarsusLengthMean, ehl,
      "batch" = exp
    ) %>%
    pivot_wider(
      id_cols = c(
        "ring", "batch",
        "pretreatment", "mass",
        "tarsusLengthMean"
      ),
      values_from = "ehl",
      names_from = "Ta"
    ) %>%
    mutate("foldEhl" = `40` / `30`) %>%
    mutate(pretreatment = ifelse(pretreatment == "neutral", "B",
```

```
      ifelse(pretreatment == "cold", "A", "C")
    )) %>%
    mutate(pretreatment = factor(pretreatment,
      levels = c("B", "A", "C")
    )) %>%
    distinct() %>%
    mutate(
      mass = mass - mean(mass, na.rm = T),
      tarsus = tarsusLengthMean - mean(tarsusLengthMean, na.rm = T)
    ),
  family = "gaussian",
  bf(mass ~ pretreatment + (1 | batch)) +
    bf(tarsus ~ mass + pretreatment + (1 | batch)) +
    bf(foldEhl ~ tarsus + mass + pretreatment + (1 | batch)) +
    set_rescor(FALSE),
  prior = c(
    set_prior("normal(0, 10)",
      class = "Intercept",
      resp = "mass"
    ),
    set_prior("normal(0, 25)",
      class = "b",
      coef = "pretreatmentA",
      resp = "mass"
    ),
    set_prior("normal(0, 25)",
      class = "b",
      coef = "pretreatmentC",
      resp = "mass"
    ),
    set_prior("exponential(2.5)",
      class = "sd",
      group = "batch",
      resp = "mass"
    ),
    set_prior("exponential(0.05)",
      class = "sigma",
      resp = "mass"
    ),
    set_prior("normal(0, 3)",
      class = "Intercept",
      resp = "tarsus"
    ),
    set_prior("normal(0, 3)",
      class = "b",
      coef = "pretreatmentA",
      resp = "tarsus"
    ),
    set_prior("normal(0, 3)",
      class = "b",
      coef = "pretreatmentC",
      resp = "tarsus"
    ),
    set_prior("skew_normal(0, 0.25, 5)",
      class = "b",
      coef = "mass",
      resp = "tarsus"
    ),
    set_prior("exponential(2)",
      class = "sd",
      group = "batch",
      resp = "tarsus"
    ),
    set_prior("exponential(0.75)",
      class = "sigma",
      resp = "tarsus"
    ),
```

```
    set_prior("normal(2.5, 1)",
      class = "Intercept",
      resp = "foldEhl"
    ),
    set_prior("normal(0, 0.5)",
      class = "b",
      coef = "pretreatmentA",
      resp = "foldEhl"
    ),
    set_prior("normal(0, 0.5)",
      class = "b",
      coef = "pretreatmentC",
      resp = "foldEhl"
    ),
    set_prior("normal(0, 0.025)",
      class = "b",
      coef = "mass",
      resp = "foldEhl"
    ),
    set_prior("normal(0, 0.5)",
      class = "b",
      coef = "tarsus",
      resp = "foldEhl"
    ),
    set_prior("exponential(5)",
      class = "sd",
      group = "batch",
      resp = "foldEhl"
    ),
    set_prior("exponential(5)",
      class = "sigma",
      resp = "foldEhl"
    )
  ),
  iter = 50000, warmup = 10000, cores = 4, chains = 4, thin = 20,
  control = list(adapt_delta = .97, max_treedepth = 14),
  silent = TRUE, refresh = 0,
  file = "./models/heatLossModel8Weeks.Rds",
)

pp1a <- mcmc_neff(neff_ratio(ehlModel8Weeks)) +
  xlab(
    TeX("$\\overset{Evaporative~Heat~Loss}{Response~(N_{eff}/N})$")
  ) +
  theme_classic() +
  theme(
    axis.text.y = element_blank(),
    axis.ticks.y = element_blank(),
    legend.position = "none"
  )

pp1b <- mcmc_rhat(rhat(ehlModel8Weeks)) +
  xlab(
    TeX("$\\overset{Evaporative~Heat~Loss}{Response~(\\hat{R}})$")
  ) +
  theme_classic() +
  theme(
    axis.text.y = element_blank(),
    axis.ticks.y = element_blank(),
    legend.position = "none"
  )

pp1c <- pp_check2(ehlModel8Weeks,
  resp = "foldEhl",
  xlab = "Evaporative Heat\nLoss Response (Fold From 30°C)"
) +
  theme(legend.position = "none")
```

```
pp1d <- ehlModel8Weeks$data %>%
  mutate(
    "Fit" = fitted(ehlModel8Weeks, resp = "foldEhl")[, "Estimate"],
    "FitSE" = fitted(ehlModel8Weeks, resp = "foldEhl")[, "Est.Error"]
  ) %>%
  ggplot(aes(x = Fit, y = foldEhl)) +
  geom_errorbarh(aes(xmin = Fit - FitSE, xmax = Fit + FitSE),
    height = 0.25, colour = "black", alpha = 0.8
  ) +
  geom_point(
    size = 2, pch = 21, colour = "black", fill = "lightblue2",
    alpha = 0.8
  ) +
  geom_smooth(
    method = "lm", colour = "black", formula = y ~ 0 + x,
    linetype = "dashed", se = FALSE
  ) +
  xlab("Fitted Evaporative Heat\nLoss Response (Fold from 30°C)") +
  ylab("Evaporative Heat Loss\nResponse(Fold from 30°C)") +
  theme_classic()

efficiencyModel8Weeks <- brm(
  data = vh2o %>%
    filter(week == "8" & Ta == 40) %>%
    select(Ta, ring, pretreatment,
      mass, tarsusLengthMean, ecc,
      "batch" = exp
    ) %>%
    mutate(
      pretreatment =
        ifelse(pretreatment == "neutral", "B",
          ifelse(pretreatment == "cold", "A", "C")
        )
    ) %>%
    mutate(pretreatment = factor(pretreatment,
      levels = c("B", "A", "C")
    )) %>%
    distinct() %>%
    mutate(
      mass = mass - mean(mass, na.rm = T),
      tarsus = tarsusLengthMean -
        mean(tarsusLengthMean, na.rm = T)
    ),
  family = "gaussian",
  bf(mass ~ pretreatment + (1 | batch)) +
    bf(tarsus ~ mass + pretreatment + (1 | batch)) +
    bf(ecc ~ tarsus + mass + pretreatment + (1 | batch)) +
    set_rescor(FALSE),
  prior = c(
    set_prior("normal(0, 10)",
      class = "Intercept",
      resp = "mass"
    ),
    set_prior("normal(0, 25)",
      class = "b",
      coef = "pretreatmentA",
      resp = "mass"
    ),
    set_prior("normal(0, 25)",
      class = "b",
      coef = "pretreatmentC",
      resp = "mass"
    ),
    set_prior("exponential(2.5)",
      class = "sd",
      group = "batch",
      resp = "mass"
```

```
    ),
    set_prior("exponential(0.05)",
      class = "sigma",
      resp = "mass"
    ),
    set_prior("normal(0, 3)",
      class = "Intercept",
      resp = "tarsus"
    ),
    set_prior("normal(0, 3)",
      class = "b",
      coef = "pretreatmentA",
      resp = "tarsus"
    ),
    set_prior("normal(0, 3)",
      class = "b",
      coef = "pretreatmentC",
      resp = "tarsus"
    ),
    set_prior("skew_normal(0, 0.25, 5)",
      class = "b",
      coef = "mass",
      resp = "tarsus"
    ),
    set_prior("exponential(2)",
      class = "sd",
      group = "batch",
      resp = "tarsus"
    ),
    set_prior("exponential(0.75)",
      class = "sigma",
      resp = "tarsus"
    ),
    set_prior("normal(0.75, 0.2)",
      class = "Intercept",
      resp = "ecc"
    ),
    set_prior("normal(0, 0.25)",
      class = "b",
      coef = "pretreatmentA",
      resp = "ecc"
    ),
    set_prior("normal(0, 0.25)",
      class = "b",
      coef = "pretreatmentC",
      resp = "ecc"
    ),
    set_prior("normal(0, 0.01)",
      class = "b",
      coef = "mass",
      resp = "ecc"
    ),
    set_prior("normal(0, 0.1)",
      class = "b",
      coef = "tarsus",
      resp = "ecc"
    ),
    set_prior("exponential(15)",
      class = "sd",
      group = "batch",
      resp = "ecc"
    ),
    set_prior("exponential(5)",
      class = "sigma",
      resp = "ecc"
    )
  ),
```

```
  iter = 50000, warmup = 10000, cores = 4, chains = 4, thin = 20,
  control = list(adapt_delta = .97, max_treedepth = 14),
  silent = TRUE, refresh = 0,
  file = "./models/efficiencyModel8Weeks.Rds"
)

pp2a <- mcmc_neff(neff_ratio(efficiencyModel8Weeks)) +
  xlab(
    TeX("$\\overset{Evaporative~Cooling}{Efficiency~N_{eff}/N}$")
  ) +
  theme_classic() +
  theme(
    axis.text.y = element_blank(),
    axis.ticks.y = element_blank(),
    legend.position = "none"
  )

pp2b <- mcmc_rhat(rhat(efficiencyModel8Weeks)) +
  xlab(
    TeX("$\\overset{Evaporative~Cooling~Efficiency}{\\hat{R}}$")
  ) +
  theme_classic() +
  theme(
    axis.text.y = element_blank(),
    axis.ticks.y = element_blank(),
    legend.position = "none"
  )

pp2c <- pp_check2(efficiencyModel8Weeks,
  resp = "ecc",
  xlab = "Evaporative Cooling\nEfficiency (EHL/RMR)"
)

pp2d <- efficiencyModel8Weeks$data %>%
  mutate(
    "Fit" = fitted(efficiencyModel8Weeks,
      resp = "ecc"
    )[, "Estimate"],
    "FitSE" = fitted(efficiencyModel8Weeks,
      resp = "ecc"
    )[, "Est.Error"]
  ) %>%
  ggplot(aes(x = Fit, y = ecc)) +
  geom_errorbarh(aes(xmin = Fit - FitSE, xmax = Fit + FitSE),
    height = 0.05, colour = "black", alpha = 0.8
  ) +
  geom_point(
    size = 2, pch = 21, colour = "black", fill = "lightblue2",
    alpha = 0.8
  ) +
  geom_smooth(
    method = "lm", colour = "black", formula = y ~ 0 + x,
    linetype = "dashed", se = FALSE
  ) +
  xlab("Fitted Evaporative\nCooling Efficiency (EHL/RMR)") +
  ylab("Evaporative Cooling\nEfficiency (EHL/RMR)") +
  theme_classic()

((pp1a + pp1b) /
  (pp1c + pp1d) /
  (pp2a + pp2b) /
  (pp2c + pp2d)) +
  plot_annotation(tag_levels = "A")
```

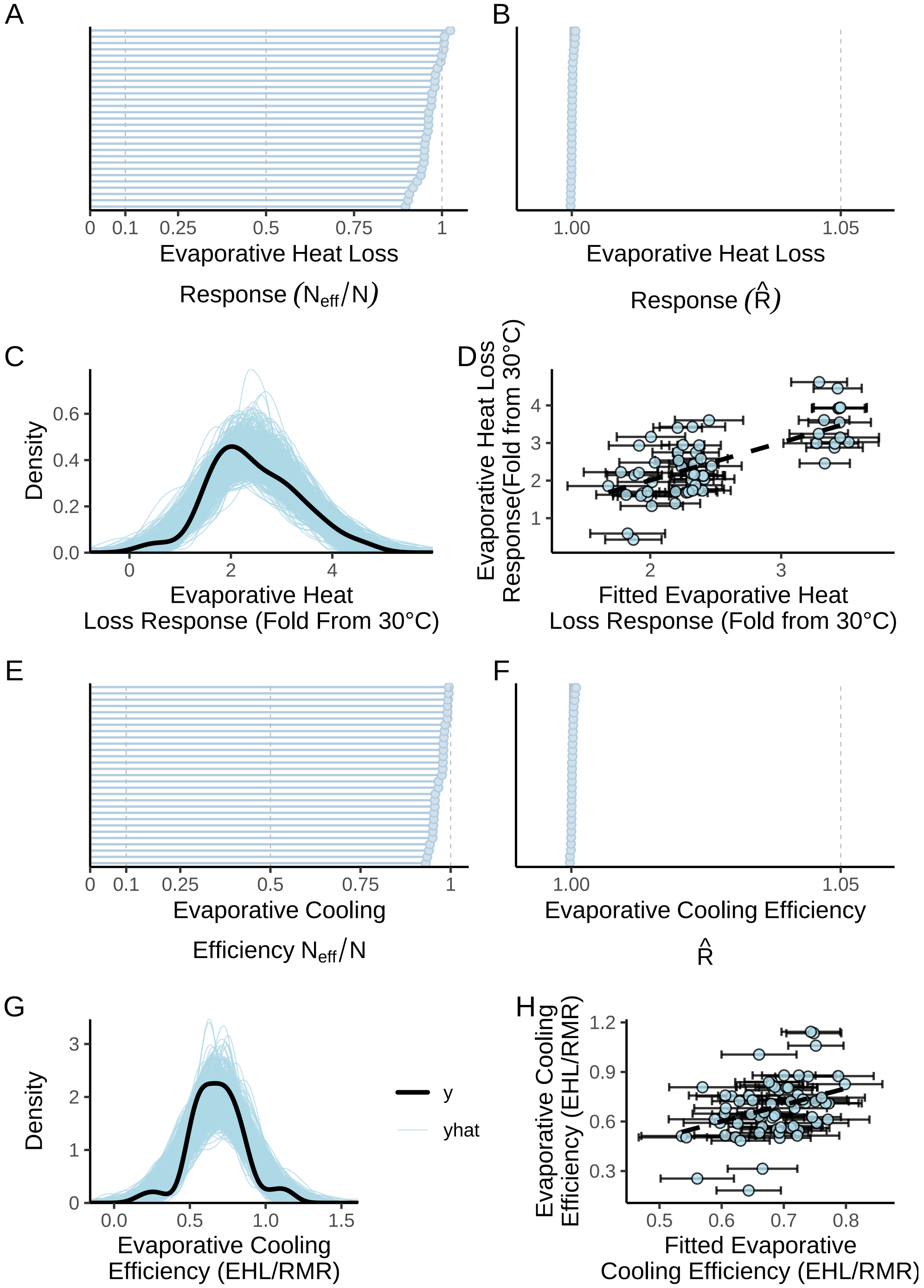


***Figure 19:*** *Model and posterior predictive checks for two Bayesian path analyses predicting, ultimately, evaporative heat loss ('EHL'; fold from that observed at 30°C in W) and evaporative cooling efficiency ('ECE'; the ratio of evaporative heat-loss to metabolic heat production, each in W) in eight week old Japanese quail. Panels A and E display parameter-specific effective sample size to sample size ratios ($N_{eff}/N$) for each model, while panels B and F display parameter specific Gelman-Rubin statistics ($\hat{R}$). Panels C and G display posterior predictive checks; black lines represent true EHL or ECE densities while blue lines represent densities estimated from model posteriors. In panels D and H, dots represent individual predictions plotted against their true values. Errorbars indicate +/- one standard error around predictions and dashed black lines indicate lines of best fit estimated by ggplot2 (Wickham, 2011) while assuming a y-intercept at 0.*

Chains are clearly well mixed and posterior estimates are unlikely to be biased by autocorrelation between chain draws. As such, we next plot model residuals (here, as medians) against model predictors and expectations to check for evidence of outlaying samples or overt heteroskedasticity.

```
rp1 <- ehlModel8Weeks$data %>%
  mutate(
    "Res" =
      residuals(ehlModel8Weeks,
        resp = "foldEhl",
        robust = TRUE
      )[, "Estimate"]
  ) %>%
  ggplot(aes(sample = Res)) +
  stat_qq(colour = "grey50") +
  stat_qq_line() +
  xlab("Theoretical EHL\nResidual Quantiles") +
  ylab("Sample ELH\nResidual Quantiles") +
  theme_classic()

rp2 <- ehlModel8Weeks$data %>%
  mutate(
    "Res" =
      residuals(ehlModel8Weeks,
        resp = "foldEhl",
        robust = TRUE
      )[, "Estimate"]
  ) %>%
  mutate(mass = mass + mean(subset(vh2o, week == "8")$mass, na.rm = T)) %>%
  ggplot(aes(x = mass, y = Res)) +
  geom_point(pch = 21, colour = "black", fill = "grey20", alpha = 0.5) +
  xlab("Body Mass (g)") +
  ylab("Evaporative Heat\nLoss Rate Residuals") +
  theme_classic()

rp3 <- ehlModel8Weeks$data %>%
  mutate(
    "Res" =
      residuals(ehlModel8Weeks,
        resp = "foldEhl",
        robust = TRUE
      )[, "Estimate"]
  ) %>%
  mutate(tarsus = tarsus +
    mean(subset(vh2o, week == "8")$tarsusLengthMean, na.rm = T)) %>%
  ggplot(aes(x = tarsus, y = Res)) +
  geom_point(pch = 21, colour = "black", fill = "grey20", alpha = 0.5) +
  xlab("Tarsus Length (mm)") +
  ylab("Evaporative Heat\nLoss Rate Residuals") +
  theme_classic()

rp4 <- ehlModel8Weeks$data %>%
  mutate(
    "Res" =
      residuals(ehlModel8Weeks,
        resp = "foldEhl",
        robust = TRUE
      )[, "Estimate"]
  ) %>%
  mutate(pretreatment = factor(pretreatment, levels = c("A", "B", "C"))) %>%
  ggplot(aes(x = pretreatment, y = Res)) +
  geom_boxplot(fill = "lightblue2") +
  geom_point(size = 2, position = position_jitter(width = 0.25)) +
  scale_x_discrete(
    name = "Rearing Treatment",
    labels = c(
      "Cold\n(10°C)",
      "Mild\n(20°C)",
```

```
      "Warm\n(30°C)"
    )
  ) +
  xlab("Rearing Treatment") +
  ylab("Evaporative Heat\nLoss Rate Residuals") +
  theme_classic()

rp5 <- efficiencyModel8Weeks$data %>%
  mutate(
    "Res" =
      residuals(efficiencyModel8Weeks,
        resp = "ecc",
        robust = TRUE
      )[, "Estimate"]
  ) %>%
  ggplot(aes(sample = Res)) +
  stat_qq(colour = "grey50") +
  stat_qq_line() +
  xlab("Theoretical ECE\nResidual Quantiles") +
  ylab("Sample ECE\nResidual Quantiles") +
  theme_classic()

rp6 <- efficiencyModel8Weeks$data %>%
  mutate(
    "Res" =
      residuals(efficiencyModel8Weeks,
        resp = "ecc",
        robust = TRUE
      )[, "Estimate"]
  ) %>%
  mutate(mass = mass + mean(subset(vh2o, week == "8")$mass, na.rm = T)) %>%
  ggplot(aes(x = mass, y = Res)) +
  geom_point(pch = 21, colour = "black", fill = "grey20", alpha = 0.5) +
  xlab("Body Mass (g)") +
  ylab("Evaporative Cooling\nEfficiency Residuals") +
  theme_classic()

rp7 <- efficiencyModel8Weeks$data %>%
  mutate(
    "Res" =
      residuals(efficiencyModel8Weeks,
        resp = "ecc",
        robust = TRUE
      )[, "Estimate"]
  ) %>%
  mutate(tarsus = tarsus +
    mean(subset(vh2o, week == "8")$tarsusLengthMean, na.rm = T)) %>%
  ggplot(aes(x = tarsus, y = Res)) +
  geom_point(pch = 21, colour = "black", fill = "grey20", alpha = 0.5) +
  xlab("Tarsus Length (mm)") +
  ylab("Evaporative Cooling\nEfficiency Residuals") +
  theme_classic()

rp8 <- efficiencyModel8Weeks$data %>%
  mutate(
    "Res" =
      residuals(efficiencyModel8Weeks,
        resp = "ecc",
        robust = TRUE
      )[, "Estimate"]
  ) %>%
  mutate(pretreatment = factor(pretreatment, levels = c("A", "B", "C"))) %>%
  ggplot(aes(x = pretreatment, y = Res)) +
  geom_boxplot(fill = "lightblue2") +
  geom_point(size = 2, position = position_jitter(width = 0.25)) +
  scale_x_discrete(
    name = "Rearing Treatment",
```

```
    labels = c(
      "Cold\n(10°C)",
      "Mild\n(20°C)",
      "Warm\n(30°C)"
    )
  ) +
  xlab("Rearing Treatment") +
  ylab("Evaporative Cooling\nEfficiency Residuals") +
  theme_classic()

((rp1 + rp2)/
  (rp3 + rp4)/
  (rp5 + rp6)/
  (rp7 + rp8)
) + plot_annotation(tag_levels = "A")
```

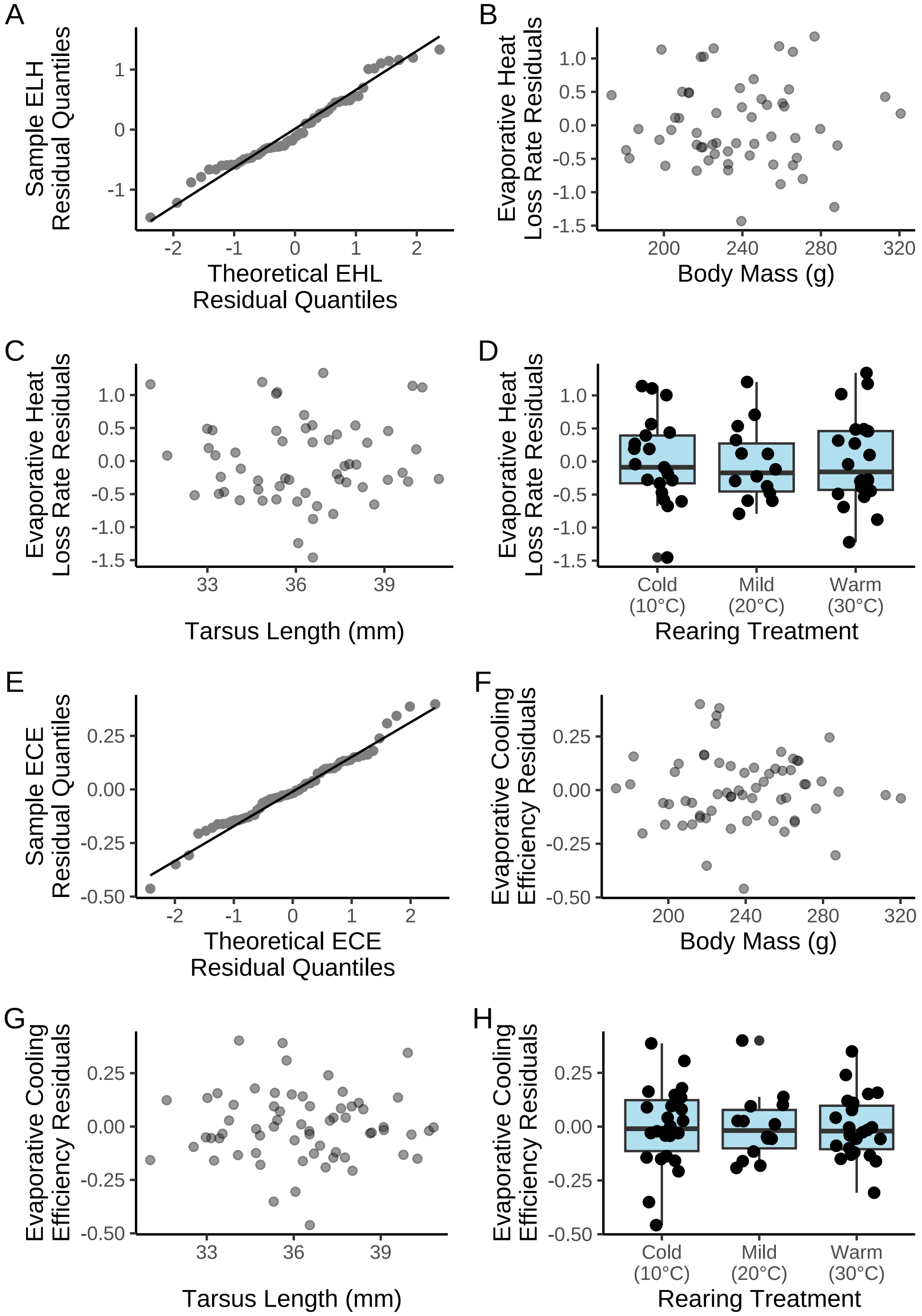


***Figure 20:*** *Residual diagnostics from two Bayesian path analyses predicting evaporative heat loss ('EHL'; fold from that observed at 30°C in W) and evaporative cooling efficiency ('ECE'; the ratio of evaporative heat-loss to metabolic heat production, each in W) in eight week old Japanese quail. Panels A and B display traditional 'qq-plots', with theoretic and sample residual quantiles regressed against each other. Dots represent individual samples. Remaining panels display median residual, per raw data point (small dots) by model predictors (here, body mass [g], tarsus length [mm] and rearing condition (10°C, 20°C, or 30°C until the time of measurement). Boxplots in panels D and H display medians (centre horizontal bar), first and third quantiles (lower and upper limits of boxes respectively) and ranges excluding outliers (whiskers). 'EHL' indicates evaporative heat loss in watts, and ECE indicates evaporative cooling efficiency (the ratio of evaporative heat loss by metabolic heat production, each in watts).*

Residuals appear both normally- and evenly-distributed (by predictor variables). Next, we plot model coefficient density to evaluate possible skewing, multimodality etc.

```
as.data.frame(ehlModel8Weeks) %>%
  pivot_longer(everything(), names_to = "par", values_to = "values") %>%
  filter(grepl("b_|sd_", par)) %>%
  merge(., tribble(~par, ~Par,
                   "b_mass_Intercept", "Mass\nIntercept",
                   "b_mass_pretreatmentA", "Mass ~\nCold Rearing",
                   "b_mass_pretreatmentC", "Mass ~\nWarm Rearing",
                   "sd_batch__mass_Intercept", "Mass ~\nBatch",
                   "b_tarsus_Intercept", "Tarsus\nIntercept",
                   "b_tarsus_pretreatmentA", "Tarsus ~\nCold Rearing",
                   "b_tarsus_pretreatmentC", "Tarsus ~\nWarm Rearing",
                   "b_tarsus_mass", "Tarsus ~\nMass",
                   "sd_batch__tarsus_Intercept", "Tarsus ~\nBatch",
                   "b_foldEhl_Intercept", "EHL\nIntercept",
                   "b_foldEhl_pretreatmentA", "EHL ~\nCold Rearing",
                   "b_foldEhl_pretreatmentC", "EHL ~\nWarm Rearing",
                   "b_foldEhl_mass", "EHL ~ Mass",
                   "b_foldEhl_tarsus", "EHL ~\nTarsus",
                   "sd_batch__foldEhl_Intercept", "EHL ~\nBatch",
                   ),
        by = "par", all.x = TRUE) %>%
  ggplot(aes(x = values)) +
  facet_wrap(~Par, scales = "free") +
  geom_density(colour = "black", fill = "white") +
  geom_vline(xintercept = 0, colour = "darkred", linetype = "solid") +
  scale_x_continuous(n.breaks = 3) +
  scale_y_continuous(n.breaks = 3) +
  xlab("Values") +
  ylab("Density") +
  theme_classic()
```

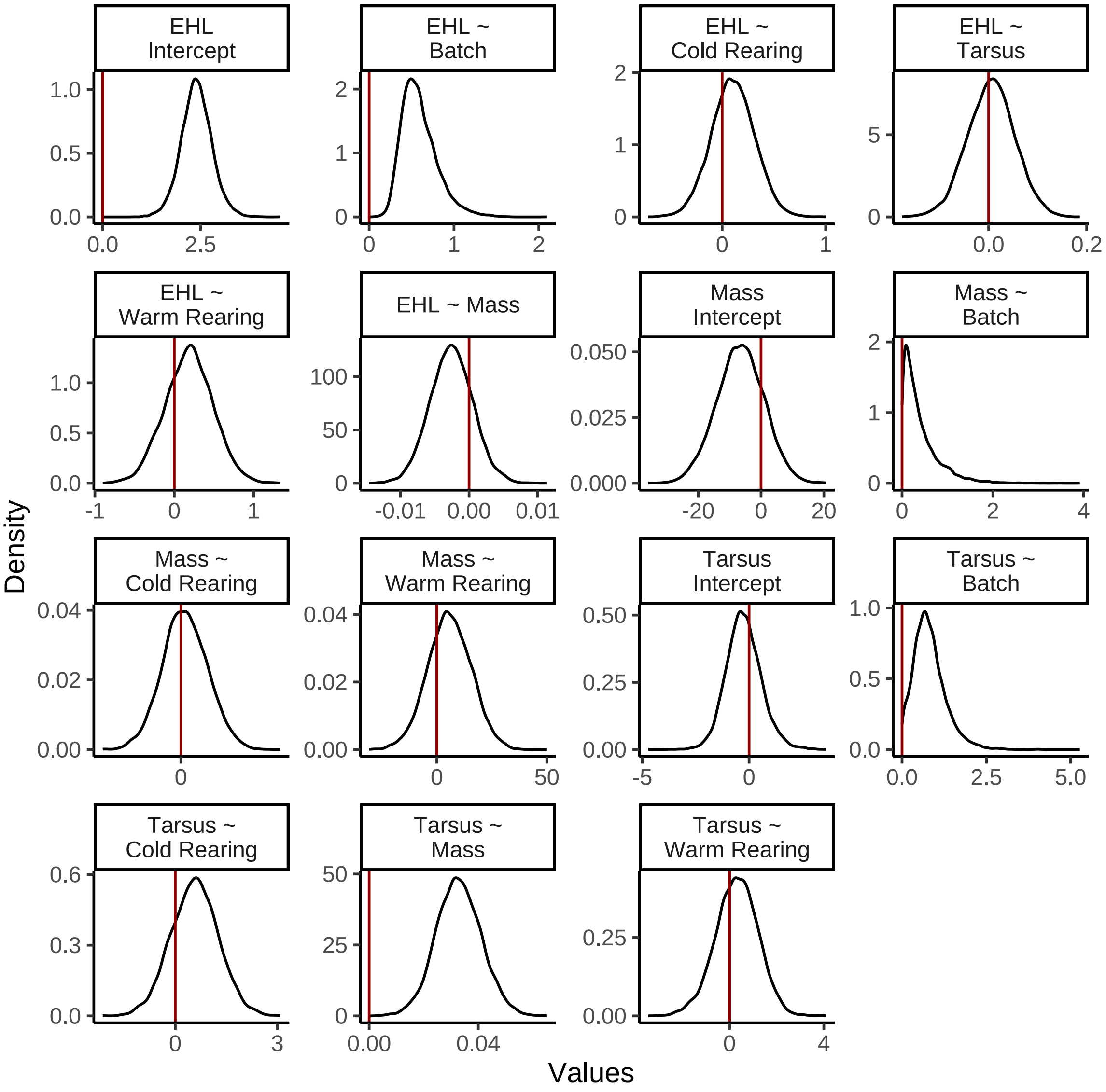


***Figure 21:*** *Posterior densities for coefficients from a Bayesian path analysis predicting body mass (g), tarsus length (mm) and evaporative heat loss responses ('EHL'; fold from that observed at 30°C in W) in eight week old Japanese quail. Red vertical lines label 0. Response and predictor variables for which densities refer are indicated on the left side and right side of tildes respectively. 'Cold Rearing' indicates rearing at 10°C, 'Warm Rearing' indicates rearing at 30°C, and 'Batch' indicates the batch of eggs from which an individual was derived. All coefficients except batch are population level.*

```
as.data.frame(efficiencyModel8Weeks) %>%
  pivot_longer(everything(), names_to = "par", values_to = "values") %>%
  filter(grepl("b_|sd_", par)) %>%
  merge(., tribble(~par, ~Par,
                   "b_mass_Intercept", "Mass\nIntercept",
                   "b_mass_pretreatmentA", "Mass ~\nCold Rearing",
                   "b_mass_pretreatmentC", "Mass ~\nWarm Rearing",
                   "sd_batch__mass_Intercept", "Mass ~\nBatch",
                   "b_tarsus_Intercept", "Tarsus\nIntercept",
                   "b_tarsus_pretreatmentA", "Tarsus ~\nCold Rearing",
                   "b_tarsus_pretreatmentC", "Tarsus ~\nWarm Rearing",
                   "b_tarsus_mass", "Tarsus ~\nMass",
                   "sd_batch__tarsus_Intercept", "Tarsus ~\nBatch",
```

```
                    "b_ecc_Intercept", "ECC\nIntercept",
                    "b_ecc_pretreatmentA", "ECC ~\nCold Rearing",
                    "b_ecc_pretreatmentC", "ECC ~\nWarm Rearing",
                    "b_ecc_mass", "ECC ~ Mass",
                    "b_ecc_tarsus", "ECC ~\nTarsus",
                    "sd_batch__ecc_Intercept", "ECC ~\nBatch",
                    ),
        by = "par", all.x = TRUE) %>%
  ggplot(aes(x = values)) +
  facet_wrap(~Par, scales = "free") +
  geom_density(colour = "black", fill = "white") +
  geom_vline(xintercept = 0, colour = "darkred", linetype = "solid") +
  scale_x_continuous(n.breaks = 3) +
  scale_y_continuous(n.breaks = 3) +
  xlab("Values") +
  ylab("Density") +
  theme_classic()
```

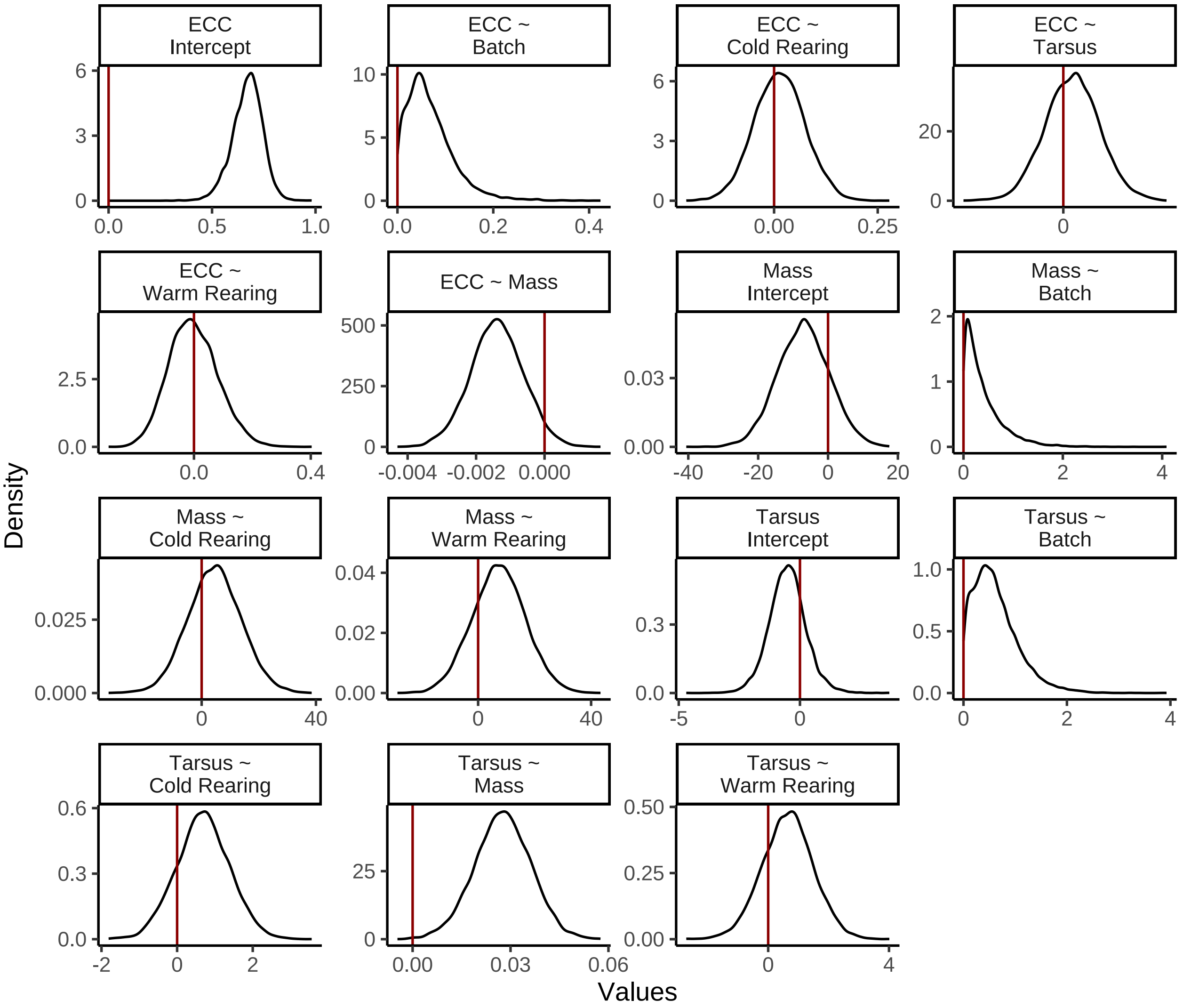


***Figure 22:*** *Posterior densities for coefficients from a Bayesian path analysis predicting body mass (g), tarsus length (mm) and evaporative cooling efficiency ('ECC', the ratio of evaporative heat loss in watts to evaporative heat production in watts) in eight week old Japanese quail. Red vertical lines label 0. Response and predictor variables for which densities refer are indicated on the left side and right side of tildes respectively. 'Cold Rearing' indicates rearing at 10°C, 'Warm Rearing' indicates rearing at 30°C, and 'Batch' indicates the batch of eggs from which an individual was derived. All coefficients except batch are population level.*

As before, we proceed by summarising centrality of posteriors with medians. Credible intervals around medians are estimated using quantiles.

```
caption <- paste0(
  "Results of Bayesian path analysis testing the ",
  "effects of morphology and rearing temperature on relative evaporative ",
  "heat loss at 40°C in eight week old Japanese quail. Relative evapoative ",
  "heat loss represents the Evaporative cooling efficiency ",
  "fold change from 30°C, per individual. Estimates indicate posterior ",
  "medians and credible intervals (CIs) indicate quantile intervals."
)

heatLoss8WeeksResults <-
  as.data.frame(ehlModel8Weeks) %>%
```

```
summarise_all(., .funs = median) %>%
pivot_longer(everything(),
  names_to = "Parameter",
  values_to = "Estimate"
) %>%
merge(., quantileCIs(ehlModel8Weeks, cis = c(50, 95)),
  by = "Parameter", all.x = TRUE
) %>%
filter(grepl("b_|sd_", Parameter)) %>%
rowwise() %>%
mutate("BF" = ifelse(Estimate < 0,
  (2 * mean(as.data.frame(
    ehlModel8Weeks
  )[, Parameter] <= 0)) /
    (2 * mean(as.data.frame(
      ehlModel8Weeks
    )[, Parameter] >= 0)),
  (2 * mean(as.data.frame(
    ehlModel8Weeks
  )[, Parameter] >= 0)) /
    (2 * mean(as.data.frame(
      ehlModel8Weeks
    )[, Parameter] <= 0))
)) %>%
ungroup() %>%
mutate(
  "Estimate" = round(Estimate, digits = 4),
  "BF" = round(BF, digits = 4),
  "N" = nrow(ehlModel3Weeks$data)
) %>%
mutate("Parameter" = ifelse(grepl("b_", Parameter),
  gsub("b_", "", Parameter),
  gsub(
    "Intercept", "batch",
    gsub(".*__", "", Parameter)
  )
)) %>%
mutate(
  "Response" = gsub("_.*", "", Parameter),
  "Parameter" = gsub(".*_", "", Parameter)
) %>%
merge(., tribble(
  ~Response, ~response, ~level,
  "mass", "Body Mass (g)", "A",
  "foldEhl", "Evaporative Heat Loss (fold from 30°C)", "D",
  "tarsus", "Tarsus Length (mm)", "B"
),
by = "Response"
) %>%
merge(., tribble(
  ~Parameter, ~parameter, ~number,
  "Intercept", "Intercept", "1",
  "mass", "Body Mass (g)", "4",
  "tarsus", "Tarsus Length (mm)", "5",
  "pretreatmentA", "Cold Rearing", "2",
  "pretreatmentC", "Warm Rearing", "3",
  "batch", "Egg Batch [mu]", "6"
),
by = "Parameter"
) %>%
mutate(
  `50\\% CI` = paste0("(", paste(
    round(Low_CI_50, digits = 4),
    round(High_CI_50, digits = 4),
    sep = ", "
  ), ")"),
  `95\\% CI` = paste0("(", paste(
```

```
      round(Low_CI_95, digits = 4),
      round(High_CI_95, digits = 4),
      sep = ", "
    ), ")")
  ) %>%
  select(-c(Low_CI_50, High_CI_50, Low_CI_95, High_CI_95)) %>%
  select(
    "Response" = "response", "Parameter" = "parameter", N,
    Estimate, `50\\% CI`, `95\\% CI`, BF, level, number
  ) %>%
  arrange(level, number) %>%
  select(-c(level, number)) %>%
  kbl(.,
    longtable = T, booktabs = T, format = "latex", escape = FALSE,
    caption = caption
  ) %>%
  column_spec(column = c(1:2), width = "2.1cm") %>%
  column_spec(column = c(3:10), width = "1.8cm") %>%
  kable_styling(latex_options = "striped")

heatLoss8WeeksResults
```

***Table 8:*** *Results of Bayesian path analysis testing the effects of morphology and rearing temperature on relative evaporative heat loss at 40°C in eight week old Japanese quail. Relative evapoative heat loss represents the Evaporative cooling efficiency fold change from 30°C, per individual. Estimates indicate posterior medians and credible intervels (CIs) indicate quantile intervals.*

| Response | Parameter | N | Estimate | 50% CI | 95% CI | BF |
|---|---|---|---|---|---|---|
| Body Mass (g) | Intercept | 48 | -6.5190 | (-11.5978, -1.5163) | (-21.5355, 7.9502) | 4.2016 |
| Body Mass (g) | Cold Rearing | 48 | 1.7124 | (-4.833, 8.5139) | (-17.5112, 21.6191) | 1.2936 |
| Body Mass (g) | Warm Rearing | 48 | 5.6182 | (-0.9137, 12.4524) | (-13.7622, 24.9382) | 2.5556 |
| Body Mass (g) | Egg Batch [mu] | 48 | 0.2702 | (0.1168, 0.54) | (0.0104, 1.4658) | Inf |
| Tarsus Length (mm) | Intercept | 48 | -0.2848 | (-0.7924, 0.2659) | (-1.8037, 1.506) | 1.7884 |
| Tarsus Length (mm) | Cold Rearing | 48 | 0.5756 | (0.0994, 1.0315) | (-0.7892, 1.9047) | 3.8164 |
| Tarsus Length (mm) | Warm Rearing | 48 | 0.3237 | (-0.2802, 0.9088) | (-1.5568, 2.0195) | 1.7787 |
| Tarsus Length (mm) | Body Mass (g) | 48 | 0.0327 | (0.0272, 0.0384) | (0.0166, 0.0494) | Inf |
| Tarsus Length (mm) | Egg Batch [mu] | 48 | 0.7489 | (0.4846, 1.062) | (0.0656, 1.9595) | Inf |
| Evaporative Heat Loss (fold from 30°C) | Intercept | 48 | 2.3953 | (2.1488, 2.6514) | (1.6216, 3.2044) | Inf |
| Evaporative Heat Loss (fold from 30°C) | Cold Rearing | 48 | 0.1034 | (-0.0353, 0.2428) | (-0.307, 0.511) | 2.2680 |
| Evaporative Heat Loss (fold from 30°C) | Warm Rearing | 48 | 0.1887 | (-0.0164, 0.3879) | (-0.4086, 0.7784) | 2.7471 |
| Evaporative Heat Loss (fold from 30°C) | Body Mass (g) | 48 | -0.0026 | (-0.0046, -5e-04) | (-0.0087, 0.0036) | 3.9659 |
| Evaporative Heat Loss (fold from 30°C) | Tarsus Length (mm) | 48 | 0.0058 | (-0.0272, 0.0372) | (-0.0907, 0.0989) | 1.2106 |
| Evaporative Heat Loss (fold from 30°C) | Egg Batch [mu] | 48 | 0.5600 | (0.4435, 0.709) | (0.2801, 1.1062) | Inf |

```
caption <- paste0(
  "Results of Bayesian path analysis testing the ",
  "effects of morphology and rearing temperature on evaporative ",
  "cooling efficiency at 40°C in eight week old Japanese quail. ",
  "Evaporative cooling efficiency represents the ratio of ",
  "evaporative heat loss (in W) to metabolic heat production ",
  "(again, in W). Estimates indicate posterior medians and credible ",
  "intervels (CIs) indicate quantile intervals."
)

efficiency8WeeksResults <-
  as.data.frame(efficiencyModel8Weeks) %>%
  summarise_all(., .funs = median) %>%
  pivot_longer(everything(),
    names_to = "Parameter",
    values_to = "Estimate"
  ) %>%
  merge(., quantileCIs(efficiencyModel8Weeks,
                       cis = c(50, 95)),
    by = "Parameter", all.x = TRUE
  ) %>%
  filter(grepl("b_|sd_", Parameter)) %>%
  rowwise() %>%
  mutate("BF" = ifelse(Estimate < 0,
    (2 * mean(as.data.frame(
      efficiencyModel8Weeks
    )[, Parameter] <= 0)) /
      (2 * mean(as.data.frame(
        efficiencyModel8Weeks
      )[, Parameter] >= 0)),
    (2 * mean(as.data.frame(
      efficiencyModel8Weeks
    )[, Parameter] >= 0)) /
      (2 * mean(as.data.frame(
        efficiencyModel8Weeks
      )[, Parameter] <= 0))
  )) %>%
  ungroup() %>%
  mutate(
    "Estimate" = round(Estimate, digits = 4),
    "BF" = round(BF, digits = 4),
    "N" = nrow(efficiencyModel8Weeks$data)
  ) %>%
  mutate("Parameter" = ifelse(grepl("b_", Parameter),
    gsub("b_", "", Parameter),
    gsub(
      "Intercept", "batch",
      gsub(".*__", "", Parameter)
    )
  )) %>%
  mutate(
    "Response" = gsub("_.*", "", Parameter),
    "Parameter" = gsub(".*_", "", Parameter)
  ) %>%
  merge(., tribble(
    ~Response, ~response, ~level,
    "mass", "Body Mass (g)", "A",
    "ecc", "Evaporative Cooling Efficiency", "D",
    "tarsus", "Tarsus Length (mm)", "B"
  ),
  by = "Response"
  ) %>%
  merge(., tribble(
    ~Parameter, ~parameter, ~number,
    "Intercept", "Intercept", "1",
    "mass", "Body Mass (g)", "4",
    "tarsus", "Tarsus Length (mm)", "5",
```

```
    "pretreatmentA", "Cold Rearing", "2",
    "pretreatmentC", "Warm Rearing", "3",
    "batch", "Egg Batch [mu]", "6"
  ),
  by = "Parameter"
  ) %>%
  mutate(
    `50\\% CI` = paste0("(", paste(
      round(Low_CI_50, digits = 4),
      round(High_CI_50, digits = 4),
      sep = ", "
    ), ")"),
    `95\\% CI` = paste0("(", paste(
      round(Low_CI_95, digits = 4),
      round(High_CI_95, digits = 4),
      sep = ", "
    ), ")")
  ) %>%
  select(-c(Low_CI_50, High_CI_50, Low_CI_95, High_CI_95)) %>%
  select(
    "Response" = "response", "Parameter" = "parameter", N,
    Estimate, `50\\% CI`, `95\\% CI`, BF, level, number
  ) %>%
  arrange(level, number) %>%
  select(-c(level, number)) %>%
  kbl(.,
    longtable = T, booktabs = T, format = "latex", escape = FALSE,
    caption = caption
  ) %>%
  column_spec(column = c(1:2), width = "2.1cm") %>%
  column_spec(column = c(3:10), width = "1.8cm") %>%
  kable_styling(latex_options = "striped")

efficiency8WeeksResults
```

***Table 9:*** *Results of Bayesian path analysis testing the effects of morphology and rearing temperature on evaporative cooling efficiency at 40°C in eight week old Japanese quail. Evaporative cooling efficiency represents the ratio of evaporative heat loss (in W) to metabolic heat production (again, in W). Estimates indicate posterior medians and credible intervels (CIs) indicate quantile intervals.*

| Response | Parameter | N | Estimate | 50% CI | 95% CI | BF |
|---|---|---|---|---|---|---|
| Body Mass (g) | Intercept | 64 | -7.0209 | (-12.0914, -2.0795) | (-21.5148, 7.7252) | 4.8097 |
| Body Mass (g) | Cold Rearing | 64 | 4.5796 | (-1.7198, 10.7346) | (-13.7908, 23.0482) | 2.2116 |
| Body Mass (g) | Warm Rearing | 64 | 7.3107 | (1.0485, 13.5708) | (-11.1297, 25.9208) | 3.5951 |
| Body Mass (g) | Egg Batch [mu] | 64 | 0.2708 | (0.1132, 0.5467) | (0.009, 1.4372) | Inf |
| Tarsus Length (mm) | Intercept | 64 | -0.5196 | (-0.9998, -0.0512) | (-2.0378, 1.0594) | 3.3692 |
| Tarsus Length (mm) | Cold Rearing | 64 | 0.6863 | (0.2204, 1.1517) | (-0.7044, 2.0642) | 4.9657 |
| Tarsus Length (mm) | Warm Rearing | 64 | 0.6582 | (0.0875, 1.205) | (-1.0304, 2.2949) | 3.4944 |
| Tarsus Length (mm) | Body Mass (g) | 64 | 0.0277 | (0.0219, 0.0333) | (0.0101, 0.0441) | 999.0000 |
| Tarsus Length (mm) | Egg Batch [mu] | 64 | 0.5417 | (0.2962, 0.8422) | (0.0329, 1.7005) | Inf |
| Evaporative Cooling Efficiency | Intercept | 64 | 0.6752 | (0.6247, 0.719) | (0.5145, 0.8025) | Inf |
| Evaporative Cooling Efficiency | Cold Rearing | 64 | 0.0130 | (-0.0283, 0.054) | (-0.106, 0.1348) | 1.4067 |

| | | | | | | |
|---|---|---|---|---|---|---|
| Evaporative Cooling Efficiency | Warm Rearing | 64 | -0.0061 | (-0.0609, 0.0537) | (-0.1566, 0.1715) | 1.1215 |
| Evaporative Cooling Efficiency | Body Mass (g) | 64 | -0.0014 | (-0.0019, -9e-04) | (-0.0029, 1e-04) | 27.3688 |
| Evaporative Cooling Efficiency | Tarsus Length (mm) | 64 | 0.0037 | (-0.0038, 0.0109) | (-0.0178, 0.0253) | 1.6792 |
| Evaporative Cooling Efficiency | Egg Batch [mu] | 64 | 0.0576 | (0.033, 0.0918) | (0.0033, 0.1916) | Inf |

Our results suggest that morphology (in particular, body mass) may influence: (1) the rate at which individuals increase their evaporative heat loss above 40°C, and (2) the efficiency by which evaporative cooling is achieved at maturity. To evaluate the extent to which variance in these two response variables are explained by morphology, we below calculate partial $R^2$ values for body mass and tarsus length from our above path analyses. Calculation of partial $R^2$ values is described above (subsection "Effects in developing individuals").

```
## Calculating total R2 values

caption = paste0("Estimates of fit for each element of a Bayesian path ",
                 "analysis predicting evaporative heat loss responses ",
                 "(fold from 30°C; Model 'A') and evaporative cooling ",
                 "efficiency (Model 'B') in mature Japanese quail ",
                 "(8 weeks of age). Response variables refer to those ",
                 "measured at 40°C. Credible intervals are quantile intervals.")

brms::bayes_R2(ehlModel8Weeks, ndraws = 1000,
               robust = TRUE) %>%
  as.data.frame() %>%
  rownames_to_column("var") %>%
  merge(., tribble(
    ~var, ~Var,
    "R2mass", "Body Mass (g)",
    "R2tarsus", "Tarsus Length (mm)",
    "R2foldEhl",
      "Evaporative Heat Loss Response"
  ), by = c("var")) %>%
  mutate(Estimate = round(Estimate, digits = 4),
         Est.Error = round(Est.Error, digits = 4),
         "95\\% CI" = paste0("[", round(Q2.5, digits = 4),
                             ", ", round(Q97.5, digits = 4),
                             "]"
         ),
         "Model" = "A"
         ) %>%
  rbind(., brms::bayes_R2(efficiencyModel8Weeks, ndraws = 1000,
               robust = TRUE) %>%
  as.data.frame() %>%
  rownames_to_column("var") %>%
  merge(., tribble(
    ~var, ~Var,
    "R2mass", "Body Mass (g)",
    "R2tarsus", "Tarsus Length (mm)",
    "R2ecc",
      "Evaporative Cooling Efficiency"
  ), by = c("var")) %>%
  mutate(Estimate = round(Estimate, digits = 4),
         Est.Error = round(Est.Error, digits = 4),
         "95\\% CI" = paste0("[", round(Q2.5, digits = 4),
                             ", ", round(Q97.5, digits = 4),
                             "]"
         ),
         "Model" = "B"
```

```
        )) %>%
  select(
    Model, "Response" = Var, "R\\textsuperscript{2}" = Estimate,
    "Standard Error" = Est.Error,
    `95\\% CI`
  ) %>%
  kbl(.,
    longtable = T, booktabs = T, format = "latex",
    caption = caption, escape = FALSE
  ) %>%
  column_spec(column = c(1:10), width = "2.5cm")  %>%
  kable_styling(latex_options = "striped")
```

***Table 10:*** *Estimates of fit for each element of a Bayesian path analysis predicting evaporative heat loss responses (fold from 30°C; Model 'A') and evaporative cooling efficiency (Model 'B') in mature Japanese quail (8 weeks of age). Response variables refer to those measured at 40°C. Credible intervals are quantile intervals.*

| Model | Response | $R^2$ | Standard Error | 95% CI |
|---|---|---|---|---|
| A | Evaporative Heat Loss Response | 0.4552 | 0.0758 | [0.2758, 0.573] |
| A | Body Mass (g) | 0.0257 | 0.0253 | [0.0012, 0.1118] |
| A | Tarsus Length (mm) | 0.3457 | 0.0868 | [0.1589, 0.478] |
| B | Evaporative Cooling Efficiency | 0.1683 | 0.0744 | [0.0461, 0.311] |
| B | Body Mass (g) | 0.0244 | 0.0253 | [0.001, 0.1077] |
| B | Tarsus Length (mm) | 0.2457 | 0.0775 | [0.1046, 0.3918] |

```
# Partial R2 calculations

ehlModel8WeeksMassR2 <- brm(
  data = vh2o %>%
    filter(week == "8" & Ta %in% c(30, 40)) %>%
    select(Ta, ring, pretreatment,
      mass, tarsusLengthMean, ehl,
      "batch" = exp
    ) %>%
    pivot_wider(
      id_cols = c(
        "ring", "batch",
        "pretreatment", "mass",
        "tarsusLengthMean"
      ),
      values_from = "ehl",
      names_from = "Ta"
    ) %>%
    mutate("foldEhl" = `40` / `30`) %>%
    mutate(pretreatment = ifelse(pretreatment == "neutral", "B",
      ifelse(pretreatment == "cold", "A", "C")
    )) %>%
    mutate(pretreatment = factor(pretreatment,
      levels = c("B", "A", "C")
    )) %>%
    distinct() %>%
    mutate(
      mass = mass - mean(mass, na.rm = T),
      tarsus = tarsusLengthMean - mean(tarsusLengthMean, na.rm = T)
    ),
  family = "gaussian",
  bf(mass ~ pretreatment + (1 | batch)) +
    bf(tarsus ~ mass + pretreatment + (1 | batch)) +
    bf(foldEhl ~ tarsus + pretreatment + (1 | batch)) +
```

```
    set_rescor(FALSE),
  prior = c(
        set_prior("normal(0, 10)",
      class = "Intercept",
      resp = "mass"
    ),
    set_prior("normal(0, 25)",
      class = "b",
      coef = "pretreatmentA",
      resp = "mass"
    ),
    set_prior("normal(0, 25)",
      class = "b",
      coef = "pretreatmentC",
      resp = "mass"
    ),
    set_prior("exponential(2.5)",
      class = "sd",
      group = "batch",
      resp = "mass"
    ),
    set_prior("exponential(0.05)",
      class = "sigma",
      resp = "mass"
    ),
    set_prior("normal(0, 3)",
      class = "Intercept",
      resp = "tarsus"
    ),
    set_prior("normal(0, 3)",
      class = "b",
      coef = "pretreatmentA",
      resp = "tarsus"
    ),
    set_prior("normal(0, 3)",
      class = "b",
      coef = "pretreatmentC",
      resp = "tarsus"
    ),
    set_prior("skew_normal(0, 0.25, 5)",
      class = "b",
      coef = "mass",
      resp = "tarsus"
    ),
    set_prior("exponential(2)",
      class = "sd",
      group = "batch",
      resp = "tarsus"
    ),
    set_prior("exponential(0.75)",
      class = "sigma",
      resp = "tarsus"
    ),
    set_prior("normal(2.5, 1)",
      class = "Intercept",
      resp = "foldEhl"
    ),
    set_prior("normal(0, 0.5)",
      class = "b",
      coef = "pretreatmentA",
      resp = "foldEhl"
    ),
    set_prior("normal(0, 0.5)",
      class = "b",
      coef = "pretreatmentC",
      resp = "foldEhl"
    ),
```

```
    set_prior("normal(0, 0.5)",
      class = "b",
      coef = "tarsus",
      resp = "foldEhl"
    ),
    set_prior("exponential(5)",
      class = "sd",
      group = "batch",
      resp = "foldEhl"
    ),
    set_prior("exponential(5)",
      class = "sigma",
      resp = "foldEhl"
    )
  ),
  iter = 50000, warmup = 10000, cores = 4, chains = 4, thin = 20,
  control = list(adapt_delta = .97, max_treedepth = 14),
  silent = TRUE, refresh = 0,
  file = "./models/heatLossModel8WeeksMassR2.Rds",
)

efficiencyModel8WeeksMassR2 <- brm(
  data = vh2o %>%
    filter(week == "8" & Ta == 40) %>%
    select(Ta, ring, pretreatment,
      mass, tarsusLengthMean, ecc,
      "batch" = exp
    ) %>%
    mutate(
      pretreatment =
        ifelse(pretreatment == "neutral", "B",
          ifelse(pretreatment == "cold", "A", "C")
        )
    ) %>%
    mutate(pretreatment = factor(pretreatment,
      levels = c("B", "A", "C")
    )) %>%
    distinct() %>%
    mutate(
      mass = mass - mean(mass, na.rm = T),
      tarsus = tarsusLengthMean -
        mean(tarsusLengthMean, na.rm = T)
    ),
  family = "gaussian",
  bf(mass ~ pretreatment + (1 | batch)) +
    bf(tarsus ~ mass + pretreatment + (1 | batch)) +
    bf(ecc ~ tarsus + pretreatment + (1 | batch)) +
    set_rescor(FALSE),
  prior = c(
 set_prior("normal(0, 10)",
      class = "Intercept",
      resp = "mass"
    ),
    set_prior("normal(0, 25)",
      class = "b",
      coef = "pretreatmentA",
      resp = "mass"
    ),
    set_prior("normal(0, 25)",
      class = "b",
      coef = "pretreatmentC",
      resp = "mass"
    ),
    set_prior("exponential(2.5)",
      class = "sd",
      group = "batch",
      resp = "mass"
```

```
    ),
    set_prior("exponential(0.05)",
      class = "sigma",
      resp = "mass"
    ),
    set_prior("normal(0, 3)",
      class = "Intercept",
      resp = "tarsus"
    ),
    set_prior("normal(0, 3)",
      class = "b",
      coef = "pretreatmentA",
      resp = "tarsus"
    ),
    set_prior("normal(0, 3)",
      class = "b",
      coef = "pretreatmentC",
      resp = "tarsus"
    ),
    set_prior("skew_normal(0, 0.25, 5)",
      class = "b",
      coef = "mass",
      resp = "tarsus"
    ),
    set_prior("exponential(2)",
      class = "sd",
      group = "batch",
      resp = "tarsus"
    ),
    set_prior("exponential(0.75)",
      class = "sigma",
      resp = "tarsus"
    ),
    set_prior("normal(0.75, 0.2)",
      class = "Intercept",
      resp = "ecc"
    ),
    set_prior("normal(0, 0.25)",
      class = "b",
      coef = "pretreatmentA",
      resp = "ecc"
    ),
    set_prior("normal(0, 0.25)",
      class = "b",
      coef = "pretreatmentC",
      resp = "ecc"
    ),
    set_prior("normal(0, 0.1)",
      class = "b",
      coef = "tarsus",
      resp = "ecc"
    ),
    set_prior("exponential(15)",
      class = "sd",
      group = "batch",
      resp = "ecc"
    ),
    set_prior("exponential(5)",
      class = "sigma",
      resp = "ecc"
    )
  ),
  iter = 50000, warmup = 10000, cores = 4, chains = 4, thin = 20,
  control = list(adapt_delta = .97, max_treedepth = 14),
  silent = TRUE, refresh = 0,
  file = "./models/efficiencyModelWeeksMassR2.Rds"
)
```

```
ehlModel8WeeksTarsusR2 <- brm(
  data = vh2o %>%
    filter(week == "8" & Ta %in% c(30, 40)) %>%
    select(Ta, ring, pretreatment,
      mass, tarsusLengthMean, ehl,
      "batch" = exp
    ) %>%
    pivot_wider(
      id_cols = c(
        "ring", "batch",
        "pretreatment", "mass",
        "tarsusLengthMean"
      ),
      values_from = "ehl",
      names_from = "Ta"
    ) %>%
    mutate("foldEhl" = `40` / `30`) %>%
    mutate(pretreatment = ifelse(pretreatment == "neutral", "B",
      ifelse(pretreatment == "cold", "A", "C")
    )) %>%
    mutate(pretreatment = factor(pretreatment,
      levels = c("B", "A", "C")
    )) %>%
    distinct() %>%
    mutate(
      mass = mass - mean(mass, na.rm = T),
      tarsus = tarsusLengthMean - mean(tarsusLengthMean, na.rm = T)
    ),
  family = "gaussian",
  bf(mass ~ pretreatment + (1 | batch)) +
    bf(tarsus ~ mass + pretreatment + (1 | batch)) +
    bf(foldEhl ~ mass + pretreatment + (1 | batch)) +
    set_rescor(FALSE),
  prior = c(
    set_prior("normal(0, 10)",
      class = "Intercept",
      resp = "mass"
    ),
    set_prior("normal(0, 25)",
      class = "b",
      coef = "pretreatmentA",
      resp = "mass"
    ),
    set_prior("normal(0, 25)",
      class = "b",
      coef = "pretreatmentC",
      resp = "mass"
    ),
    set_prior("exponential(2.5)",
      class = "sd",
      group = "batch",
      resp = "mass"
    ),
    set_prior("exponential(0.05)",
      class = "sigma",
      resp = "mass"
    ),
    set_prior("normal(0, 3)",
      class = "Intercept",
      resp = "tarsus"
    ),
    set_prior("normal(0, 3)",
      class = "b",
      coef = "pretreatmentA",
      resp = "tarsus"
    ),
    set_prior("normal(0, 3)",
```

```
      class = "b",
      coef = "pretreatmentC",
      resp = "tarsus"
    ),
    set_prior("skew_normal(0, 0.25, 5)",
      class = "b",
      coef = "mass",
      resp = "tarsus"
    ),
    set_prior("exponential(2)",
      class = "sd",
      group = "batch",
      resp = "tarsus"
    ),
    set_prior("exponential(0.75)",
      class = "sigma",
      resp = "tarsus"
    ),
    set_prior("normal(2.5, 1)",
      class = "Intercept",
      resp = "foldEhl"
    ),
    set_prior("normal(0, 0.5)",
      class = "b",
      coef = "pretreatmentA",
      resp = "foldEhl"
    ),
    set_prior("normal(0, 0.5)",
      class = "b",
      coef = "pretreatmentC",
      resp = "foldEhl"
    ),
    set_prior("normal(0, 0.025)",
      class = "b",
      coef = "mass",
      resp = "foldEhl"
    ),
    set_prior("exponential(5)",
      class = "sd",
      group = "batch",
      resp = "foldEhl"
    ),
    set_prior("exponential(5)",
      class = "sigma",
      resp = "foldEhl"
    )
  ),
  iter = 50000, warmup = 10000, cores = 4, chains = 4, thin = 20,
  control = list(adapt_delta = .97, max_treedepth = 14),
  silent = TRUE, refresh = 0,
  file = "./models/heatLossModel8WeeksTarsusR2.Rds",
)

efficiencyModel8WeeksTarsusR2 <- brm(
  data = vh2o %>%
    filter(week == "8" & Ta == 40) %>%
    select(Ta, ring, pretreatment,
      mass, tarsusLengthMean, ecc,
      "batch" = exp
    ) %>%
    mutate(
      pretreatment =
        ifelse(pretreatment == "neutral", "B",
          ifelse(pretreatment == "cold", "A", "C")
        )
    ) %>%
    mutate(pretreatment = factor(pretreatment,
```

```
      levels = c("B", "A", "C")
    )) %>%
    distinct() %>%
    mutate(
      mass = mass - mean(mass, na.rm = T),
      tarsus = tarsusLengthMean -
        mean(tarsusLengthMean, na.rm = T)
    ),
  family = "gaussian",
  bf(mass ~ pretreatment + (1 | batch)) +
    bf(tarsus ~ mass + pretreatment + (1 | batch)) +
    bf(ecc ~ tarsus + mass + pretreatment + (1 | batch)) +
    set_rescor(FALSE),
  prior = c(
set_prior("normal(0, 10)",
      class = "Intercept",
      resp = "mass"
    ),
    set_prior("normal(0, 25)",
      class = "b",
      coef = "pretreatmentA",
      resp = "mass"
    ),
    set_prior("normal(0, 25)",
      class = "b",
      coef = "pretreatmentC",
      resp = "mass"
    ),
    set_prior("exponential(2.5)",
      class = "sd",
      group = "batch",
      resp = "mass"
    ),
    set_prior("exponential(0.05)",
      class = "sigma",
      resp = "mass"
    ),
    set_prior("normal(0, 3)",
      class = "Intercept",
      resp = "tarsus"
    ),
    set_prior("normal(0, 3)",
      class = "b",
      coef = "pretreatmentA",
      resp = "tarsus"
    ),
    set_prior("normal(0, 3)",
      class = "b",
      coef = "pretreatmentC",
      resp = "tarsus"
    ),
    set_prior("skew_normal(0, 0.25, 5)",
      class = "b",
      coef = "mass",
      resp = "tarsus"
    ),
    set_prior("exponential(2)",
      class = "sd",
      group = "batch",
      resp = "tarsus"
    ),
    set_prior("exponential(0.75)",
      class = "sigma",
      resp = "tarsus"
    ),
    set_prior("normal(0.75, 0.2)",
      class = "Intercept",
```

```
      resp = "ecc"
    ),
    set_prior("normal(0, 0.25)",
      class = "b",
      coef = "pretreatmentA",
      resp = "ecc"
    ),
    set_prior("normal(0, 0.25)",
      class = "b",
      coef = "pretreatmentC",
      resp = "ecc"
    ),
    set_prior("normal(0, 0.01)",
      class = "b",
      coef = "mass",
      resp = "ecc"
    ),
    set_prior("exponential(15)",
      class = "sd",
      group = "batch",
      resp = "ecc"
    ),
    set_prior("exponential(5)",
      class = "sigma",
      resp = "ecc"
    )
  ),
  iter = 50000, warmup = 10000, cores = 4, chains = 4, thin = 20,
  control = list(adapt_delta = .97, max_treedepth = 14),
  silent = TRUE, refresh = 0,
  file = "./models/efficiencyModel8WeeksTarsusR2.Rds"
)

heatLossMassPR2 <- brms::bayes_R2(ehlModel8Weeks,
                          ndraws = 1000,
                          robust = TRUE, resp = "foldEhl",
                          summary = FALSE
) %>%
  as.data.frame() %>%
  select("fullR2" = R2foldEhl) %>%
  cbind(
    .,
    brms::bayes_R2(ehlModel8WeeksMassR2,
                   ndraws = 1000,
                   robust = TRUE, resp = "foldEhl",
                   summary = FALSE
    )
  ) %>%
  mutate("pR2" = round(fullR2 - R2foldEhl, digits = 3)) %>%
  pull(pR2)

efficiencyMassPR2 <- brms::bayes_R2(efficiencyModel8Weeks,
                          ndraws = 1000,
                          robust = TRUE, resp = "ecc",
                          summary = FALSE
) %>%
  as.data.frame() %>%
  select("fullR2" = R2ecc) %>%
  cbind(
    .,
    brms::bayes_R2(efficiencyModel8WeeksMassR2,
                   ndraws = 1000,
                   robust = TRUE, resp = "ecc",
                   summary = FALSE
    )
  ) %>%
  mutate("pR2" = round(fullR2 - R2ecc, digits = 3)) %>%
```

```
  pull(pR2)

heatLossTarsusPR2 <- brms::bayes_R2(ehlModel8Weeks,
                          ndraws = 1000,
                          robust = TRUE, resp = "foldEhl",
                          summary = FALSE
) %>%
  as.data.frame() %>%
  select("fullR2" = R2foldEhl) %>%
  cbind(
    .,
    brms::bayes_R2(ehlModel8WeeksTarsusR2,
                   ndraws = 1000,
                   robust = TRUE, resp = "foldEhl",
                   summary = FALSE
    )
  ) %>%
  mutate("pR2" = round(fullR2 - R2foldEhl, digits = 3)) %>%
  pull(pR2)

efficiencyTarsusPR2 <- brms::bayes_R2(efficiencyModel8Weeks,
                          ndraws = 1000,
                          robust = TRUE, resp = "ecc",
                          summary = FALSE
) %>%
  as.data.frame() %>%
  select("fullR2" = R2ecc) %>%
  cbind(
    .,
    brms::bayes_R2(efficiencyModel8WeeksTarsusR2,
                   ndraws = 1000,
                   robust = TRUE, resp = "ecc",
                   summary = FALSE
    )
  ) %>%
  mutate("pR2" = round(fullR2 - R2ecc, digits = 3)) %>%
  pull(pR2)

caption = paste0('Variance in evaporative heat loss (fold from that at 30°C) ',
                 'and evaporative cooling efficiency ',
                 '(EHL/MHP) at 40°C explained by morphometry in eight week ',
                 'old Japanese quail. Variance explained is represented as ',
                 'partial R\\textsuperscript{2} values.')

partialR2EightWeeks <- data.frame(
  "Response" = rep(
    c(
      "Fold Evaporative Heat Loss",
      "Evaporative Cooling Efficiency"
    ),
    each = 2
  ),
  "Predictor" = rep(
    c(
      "Body Mass (g)",
      "Tarsus Length (mm)"
    ),
    2
  )
) %>%
  mutate(
    "Mass Partial R\\textsuperscript{2}" =
      lapply(
        X = list(
          heatLossMassPR2,
          efficiencyMassPR2,
          heatLossTarsusPR2,
```

```
        efficiencyTarsusPR2
      ),
      FUN = function(x) {
        round(median(x), digits = 4)
      }
    ) %>%
      unlist(),
  "95\\% CI" = lapply(
    X = list(
      heatLossMassPR2,
      efficiencyMassPR2,
      heatLossTarsusPR2,
      efficiencyTarsusPR2
    ),
    FUN = function(x) {
      paste0(
        "[",
        paste0(
          round(
            quantile(x, probs = c(0.025, 0.975), type = 8),
            digits = 4
          ),
          collapse = ", "
        ),
        "]"
      )
    }
  )
) %>%
kbl(.,
    longtable = T, booktabs = T,
    format = "latex", escape = FALSE,
    caption = caption
) %>%
column_spec(column = c(1:2), width = "2.0cm")  %>%
column_spec(column = c(3:10), width = "1.9cm") %>%
kable_styling(latex_options = "striped")

partialR2EightWeeks
```

***Table 11:*** *Variance in evaporative heat loss (fold from that at 30°C) and evaporative cooling efficiency (EHL/MHP) at 40°C explained by morphometry in eight week old Japanese quail. Variance explained is represented as partial $R^2$ values.*

| Response | Predictor | Mass Partial $R^2$ | 95% CI |
|---|---|---|---|
| Fold Evaporative Heat Loss | Body Mass (g) | 0.0115 | [-0.214, 0.236] |
| Fold Evaporative Heat Loss | Tarsus Length (mm) | 0.0520 | [-0.1283, 0.2433] |
| Evaporative Cooling Efficiency | Body Mass (g) | -0.0015 | [-0.2247, 0.2107] |
| Evaporative Cooling Efficiency | Tarsus Length (mm) | 0.0030 | [-0.191, 0.189] |

Below, conditional effects of body mass, tarsus length, and rearing treatment on both evaporative heat loss responses and evaporative cooling efficiency (both at 40°C) are plotted. All plots assume otherwise average morphology or mild rearing (20°C).

```
heatLoss8WeeksMassPlot <-
  expand.grid(
    "mass" = with(
      ehlModel8Weeks$data,
      seq(min(mass), max(mass), by = 1)
    ),
    "tarsus" = 0,
    "pretreatment" = "B"
  ) %>%
  mutate(
    "foldEhl" = predict(ehlModel8Weeks,
      newdata = .,
      re_form = NA, robust = TRUE,
      resp = "foldEhl"
    )[, "Estimate"],
    "SE" = predict(ehlModel8Weeks,
      newdata = .,
      re_form = NA, robust = TRUE,
      resp = "foldEhl"
    )[, "Est.Error"]
  ) %>%
  mutate(mass = mass + mean(subset(vh2o, week == 8)$mass, na.rm = T)) %>%
  ggplot(aes(x = mass, y = foldEhl)) +
  geom_ribbon(aes(ymin = foldEhl - SE, ymax = foldEhl + SE),
    alpha = 0.5, fill = "#B35050"
  ) +
  geom_point(
    data = ehlModel8Weeks$data %>%
      mutate(mass = mass +
        mean(subset(vh2o, week == 8)$mass, na.rm = T)),
    aes(x = mass, y = foldEhl),
    alpha = 0.9
  ) +
  geom_smooth(
    method = "lm", colour = "black",
    linetype = "dashed", se = FALSE
  ) +
  xlab("Body Mass (g)") +
  ylab("Evaporative Heat Loss\n(Fold From 30°C)") +
  theme_classic() +
  theme(
    axis.title = element_text(family = "Noto Sans"),
    axis.text = element_text(family = "Noto Sans")
  )

heatLoss8WeeksTarsusPlot <-
  expand.grid(
    "tarsus" = with(
      ehlModel8Weeks$data,
      seq(min(tarsus), max(tarsus), by = 1)
    ),
    "mass" = 0,
    "pretreatment" = "B"
  ) %>%
  mutate(
    "foldEhl" = predict(ehlModel8Weeks,
      newdata = .,
      re_form = NA, robust = TRUE,
      resp = "foldEhl"
    )[, "Estimate"],
    "SE" = predict(ehlModel8Weeks,
      newdata = .,
      re_form = NA, robust = TRUE,
      resp = "foldEhl"
    )[, "Est.Error"]
  ) %>%
  mutate(tarsus = tarsus +
```

```
          mean(subset(vh2o, week == 8)$tarsusLengthMean, na.rm = T)) %>%
  ggplot(aes(x = tarsus, y = foldEhl)) +
  geom_ribbon(aes(ymin = foldEhl - SE, ymax = foldEhl + SE),
    alpha = 0.5, fill = "#B35050"
  ) +
  geom_point(
    data = ehlModel8Weeks$data %>%
      mutate(tarsus = tarsus +
        mean(subset(vh2o, week == 8)$tarsusLengthMean, na.rm = T)),
    aes(x = tarsus, y = foldEhl),
    alpha = 0.5
  ) +
  geom_smooth(
    method = "lm", colour = "black",
    linetype = "dashed", se = FALSE
  ) +
  xlab("Tarsus Length (mm)") +
  ylab("Evaporative Heat Loss\n(Fold From 30°C)") +
  theme_classic() +
  theme(
    axis.title = element_text(family = "Noto Sans"),
    axis.text = element_text(family = "Noto Sans")
  )

efficiency8WeeksMassPlot <-
  expand.grid(
    "mass" = with(
      efficiencyModel8Weeks$data,
      seq(min(mass), max(mass), by = 1)
    ),
    "tarsus" = 0,
    "pretreatment" = "B"
  ) %>%
  mutate(
    "ecc" = predict(efficiencyModel8Weeks,
      newdata = .,
      re_form = NA, robust = TRUE,
      resp = "ecc"
    )[, "Estimate"],
    "SE" = predict(efficiencyModel8Weeks,
      newdata = .,
      re_form = NA, robust = TRUE,
      resp = "ecc"
    )[, "Est.Error"]
  ) %>%
  mutate(mass = mass + mean(subset(vh2o, week == 8)$mass, na.rm = T)) %>%
  ggplot(aes(x = mass, y = ecc)) +
  geom_ribbon(aes(ymin = ecc - SE, ymax = ecc + SE),
    alpha = 0.5, fill = "#B35050"
  ) +
  geom_point(
    data = efficiencyModel8Weeks$data %>%
      mutate(mass = mass +
        mean(subset(vh2o, week == 8)$mass, na.rm = T)),
    aes(x = mass, y = ecc)
  ) +
  geom_smooth(
    method = "lm", colour = "black",
    linetype = "dashed", se = FALSE
  ) +
  xlab("Body Mass (g)") +
  ylab("Evaporative Cooling\nEfficiency (EHL/RMR)") +
  theme_classic() +
  theme(
    axis.title = element_text(family = "Noto Sans"),
    axis.text = element_text(family = "Noto Sans")
  )
```

```r
efficiency8WeeksTarsusPlot <-
  expand.grid(
    "tarsus" = with(
      efficiencyModel8Weeks$data,
      seq(min(tarsus), max(tarsus), by = 1)
    ),
    "mass" = 0,
    "pretreatment" = "B"
  ) %>%
  mutate(
    "ecc" = predict(efficiencyModel8Weeks,
      newdata = .,
      re_form = NA, robust = TRUE,
      resp = "ecc"
    )[, "Estimate"],
    "SE" = predict(efficiencyModel8Weeks,
      newdata = .,
      re_form = NA, robust = TRUE,
      resp = "ecc"
    )[, "Est.Error"]
  ) %>%
  mutate(tarsus = tarsus +
           mean(subset(vh2o, week == 8)$tarsusLengthMean, na.rm = T)) %>%
  ggplot(aes(x = tarsus, y = ecc)) +
  geom_ribbon(aes(ymin = ecc - SE, ymax = ecc + SE),
    alpha = 0.5, fill = "#DECCC1"
  ) +
  geom_point(
    data = efficiencyModel8Weeks$data %>%
      mutate(tarsus = tarsus +
        mean(subset(vh2o, week == 8)$tarsusLengthMean, na.rm = T)),
    aes(x = tarsus, y = ecc),
    alpha = 0.5
  ) +
  geom_smooth(
    method = "lm", colour = "black",
    linetype = "dashed", se = FALSE
  ) +
  xlab("Tarsus Length (mm)") +
  ylab("Evaporative Cooling\nEfficiency (EHL/RMR)") +
  theme_classic() +
  theme(
    axis.title = element_text(family = "Noto Sans"),
    axis.text = element_text(family = "Noto Sans")
  )

(heatLoss8WeeksMassPlot + heatLoss8WeeksTarsusPlot) /
  (efficiency8WeeksMassPlot + efficiency8WeeksTarsusPlot) +
  plot_annotation(tag_level = "A")
```

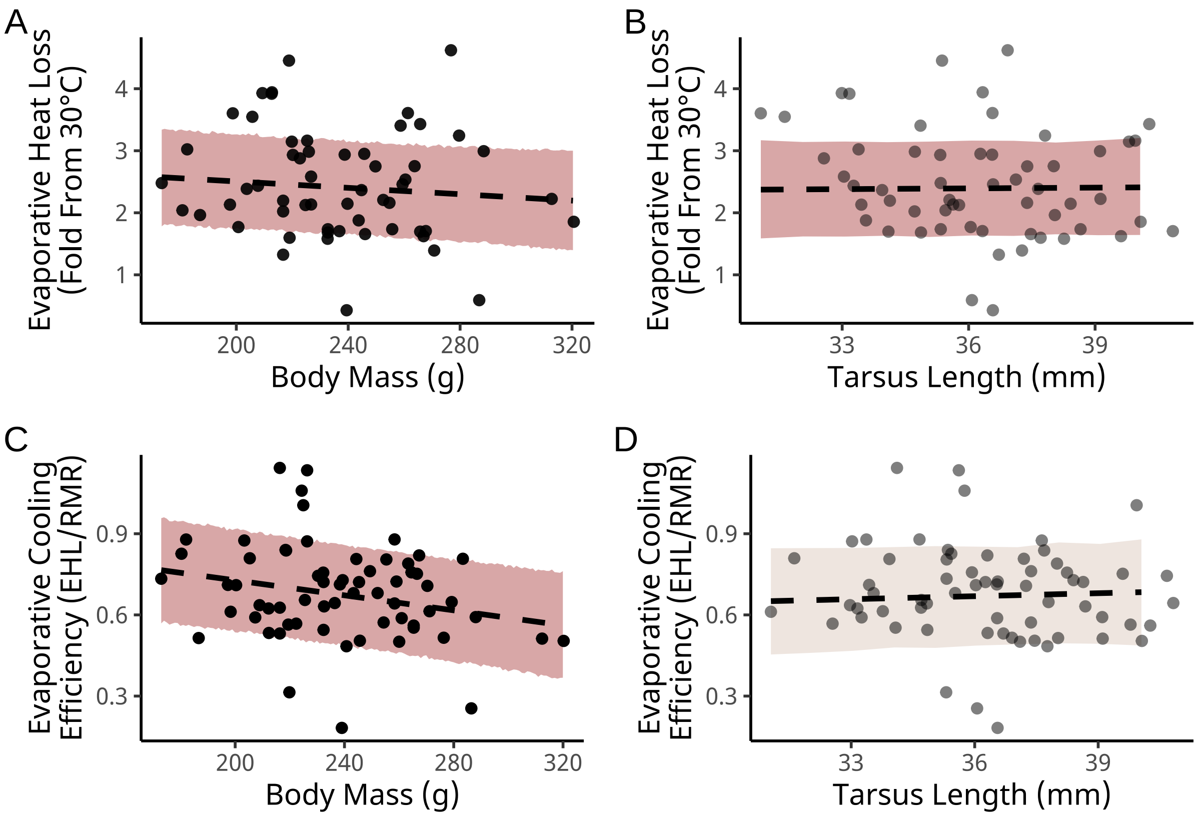


***Figure 23:*** *Effects of morphology on evaporative heat loss responses (panels A and B) and evaporative cooling efficiency (panels C and D) in mature Japanese quail (eight weeks of age; n = 57). Evaporative heat loss responses represent fold increases in evaporative heat loss, in Watts, between 30°C and 40°C. Evaporative cooling efficiency represents the ratio between evaporative heat loss (in Watts) and metabolic heat production (in Watts). Dashed lines indicate predicted relationships, as estimated from Bayesian path analyses. Ribbons represents +/- one standard error around predicted lines of best fit. Small dots indicate raw data points.*

```
heatLoss8WeeksTreatmentPlot <-
  expand.grid(
    "mass" = 0,
    "tarsus" = 0,
    "pretreatment" = c("A", "B", "C")
  ) %>%
  mutate(
    "foldEhl" = predict(ehlModel8Weeks,
      newdata = .,
      re_form = NA, robust = TRUE,
      resp = "foldEhl"
    )[, "Estimate"],
    "SE" = predict(ehlModel8Weeks,
      newdata = .,
      re_form = NA, robust = TRUE,
      resp = "foldEhl"
    )[, "Est.Error"]
  ) %>%
  mutate(pretreatment = factor(pretreatment, levels = c("A", "B", "C"))) %>%
  ggplot(aes(x = pretreatment, y = foldEhl, fill = pretreatment)) +
  geom_point(
    data = ehlModel8Weeks$data %>%
      mutate(pretreatment = factor(pretreatment, levels = c("A", "B", "C"))),
    aes(x = pretreatment, y = foldEhl),
    alpha = 0.5, position = position_jitter(width = 0.25)
```

```
  ) +
  geom_errorbar(aes(ymin = foldEhl - SE, ymax = foldEhl + SE),
    colour = "black", width = 0.25
  ) +
  geom_point(
    pch = 21, colour = "black", size = 4
  ) +
  scale_x_discrete(name = "Rearing Treatment",
                   labels = c("Cold (10°C)",
                              "Mild (20°C)",
                              "Warm (30°C)")
                   ) +
  ylab("Evaporative Heat Loss\nResponse (Fold From 30°C)") +
  scale_fill_manual(values = c("#7BB4E3", "black", "#CD5C5C")) +
  theme_classic() +
  theme(
    axis.title = element_text(family = "Noto Sans"),
    axis.text = element_text(family = "Noto Sans"),
    legend.position = "none"
  )

efficiency8WeeksTreatmentPlot <-
  expand.grid(
    "mass" = 0,
    "tarsus" = 0,
    "pretreatment" = c("A", "B", "C")
  ) %>%
  mutate(
    "ecc" = predict(efficiencyModel8Weeks,
      newdata = .,
      re_form = NA, robust = TRUE,
      resp = "ecc"
    )[, "Estimate"],
    "SE" = predict(efficiencyModel8Weeks,
      newdata = .,
      re_form = NA, robust = TRUE,
      resp = "ecc"
    )[, "Est.Error"]
  ) %>%
  mutate(pretreatment = factor(pretreatment, levels = c("A", "B", "C"))) %>%
  ggplot(aes(x = pretreatment, y = ecc, fill = pretreatment)) +
  geom_point(
    data = efficiencyModel8Weeks$data %>%
      mutate(pretreatment = factor(pretreatment, levels = c("A", "B", "C"))),
    aes(x = pretreatment, y = ecc),
    alpha = 0.5, position = position_jitter(width = 0.25)
  ) +
  geom_errorbar(aes(ymin = ecc - SE, ymax = ecc + SE),
    colour = "black", width = 0.25
  ) +
  geom_point(
    pch = 21, colour = "black", size = 4
  ) +
  scale_x_discrete(name = "Rearing Treatment",
                   labels = c("Cold (10°C)",
                              "Mild (20°C)",
                              "Warm (30°C)")
                   ) +
  ylab("Evaporative Cooling\nEfficiency (EHL/RMR)") +
  scale_fill_manual(values = c("#7BB4E3", "black", "#CD5C5C")) +
  theme_classic() +
  theme(
    axis.title = element_text(family = "Noto Sans"),
    axis.text = element_text(family = "Noto Sans"),
    legend.position = "none"
  )
```

```
(heatLoss8WeeksTreatmentPlot / efficiency8WeeksTreatmentPlot) +
  plot_annotation(tag_levels = "A")
```

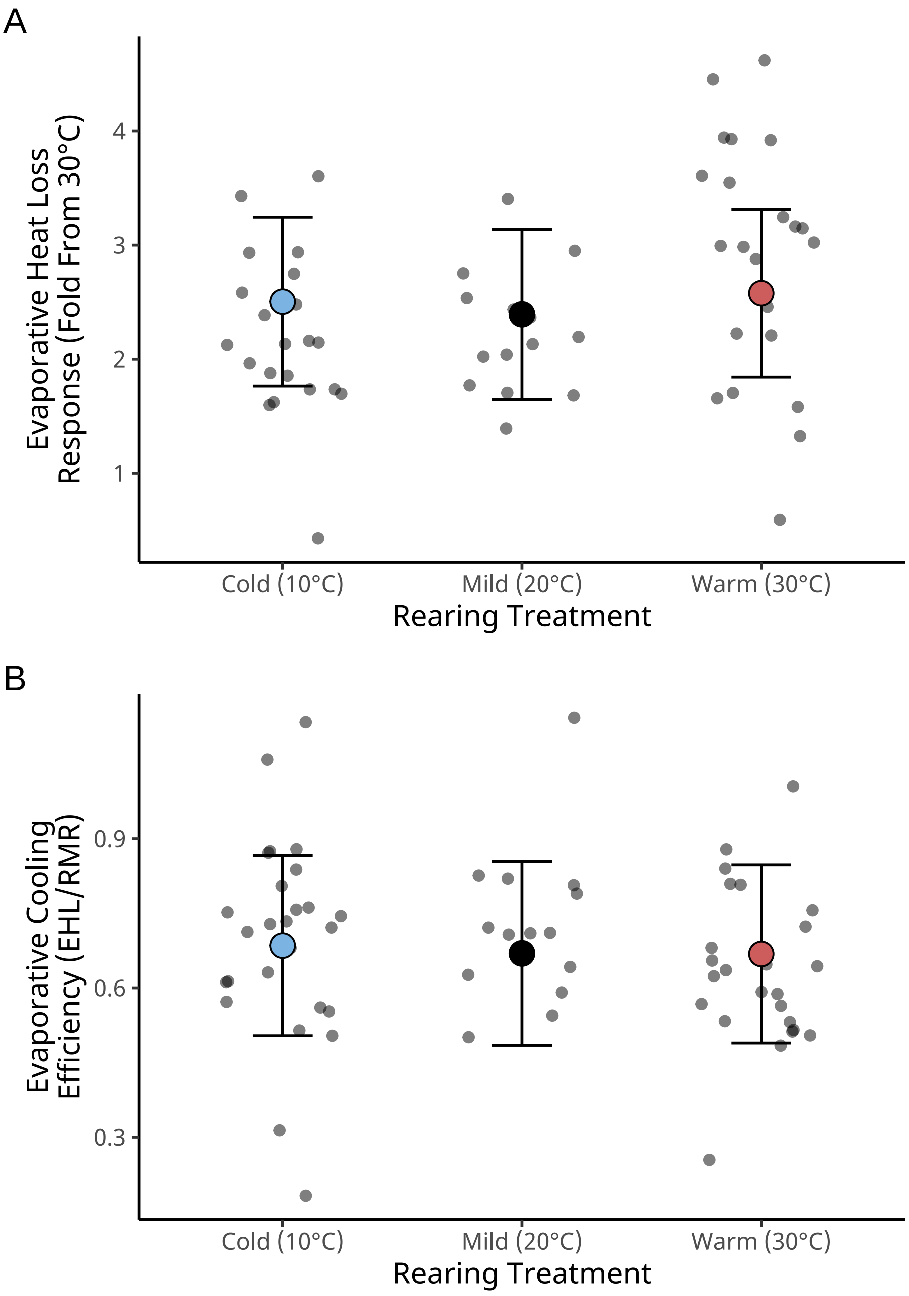


***Figure 24:*** *Effects of rearing temperature on evaporative heat loss responses (panel A) and evaporative cooling efficiency (panel B) in mature Japanese quail (eight weeks of age; n = 57). Evaporative heat loss responses represent fold increases in evaporative heat loss, in Watts, between 30°C and 40°C. Evaporative cooling efficiency represents the ratio between evaporative heat loss (in Watts) and metabolic heat production (in Watts). Dashed lines indicate predicted relationships, as estimated from Bayesian path analyses. Ribbons represents +/- one standard error around predicted lines of best fit. Small dots indicate raw data points.*

Last, we again estimate and visualise the direct and indirect effects of rearing conditions (and body mass) on evaporative heat loss responses and evaporative cooling efficiency (both at 40°C). Similar to our estimations for developing quail, model coefficients are first scaled to represent the effect of changing a given predictor variable by one standard deviation (or categorical level) on a change, in standard deviations, of the response variable.

```
# Coefficients are first scaled

scaledBetasHeatLoss8 <- as.data.frame(ehlModel8Weeks) %>%
  mutate(
    b_mass_pretreatmentA = b_mass_pretreatmentA /
      sd(ehlModel8Weeks$data$mass),
    b_mass_pretreatmentC = b_mass_pretreatmentC /
      sd(ehlModel8Weeks$data$mass),
    b_tarsus_pretreatmentA = b_tarsus_pretreatmentA /
      sd(ehlModel8Weeks$data$tarsus),
    b_tarsus_pretreatmentC = b_tarsus_pretreatmentC /
      sd(ehlModel8Weeks$data$tarsus),
    b_tarsus_mass =
      (b_tarsus_mass * sd(ehlModel8Weeks$data$mass)) /
        sd(ehlModel8Weeks$data$tarsus),
    b_foldEhl_mass =
      (b_foldEhl_mass * sd(ehlModel8Weeks$data$mass)) /
        sd(ehlModel8Weeks$data$foldEhl),
    b_foldEhl_tarsus =
      (b_foldEhl_tarsus * sd(ehlModel8Weeks$data$tarsus)) /
        sd(ehlModel8Weeks$data$foldEhl),
    b_foldEhl_pretreatmentA = b_foldEhl_pretreatmentA /
      sd(ehlModel8Weeks$data$foldEhl),
    b_foldEhl_pretreatmentC = b_foldEhl_pretreatmentC /
      sd(ehlModel8Weeks$data$foldEhl),
  )

scaledBetasEfficiency8 <- as.data.frame(efficiencyModel8Weeks) %>%
  mutate(
    b_mass_pretreatmentA = b_mass_pretreatmentA /
      sd(efficiencyModel8Weeks$data$mass),
    b_mass_pretreatmentC = b_mass_pretreatmentC /
      sd(efficiencyModel8Weeks$data$mass),
    b_tarsus_pretreatmentA = b_tarsus_pretreatmentA /
      sd(efficiencyModel8Weeks$data$tarsus),
    b_tarsus_pretreatmentC = b_tarsus_pretreatmentC /
      sd(efficiencyModel8Weeks$data$tarsus),
    b_tarsus_mass =
      (b_tarsus_mass * sd(efficiencyModel8Weeks$data$mass)) /
        sd(efficiencyModel8Weeks$data$tarsus),
    b_ecc_mass =
      (b_ecc_mass * sd(efficiencyModel8Weeks$data$mass)) /
        sd(efficiencyModel8Weeks$data$ecc),
    b_ecc_tarsus =
      (b_ecc_tarsus * sd(efficiencyModel8Weeks$data$tarsus)) /
        sd(efficiencyModel8Weeks$data$ecc),
    b_ecc_pretreatmentA = b_ecc_pretreatmentA /
      sd(efficiencyModel8Weeks$data$ecc),
    b_ecc_pretreatmentC = b_ecc_pretreatmentC /
      sd(efficiencyModel8Weeks$data$ecc),
  )

# Effects are now calculated

fullEffectHeatLoss8 <- scaledBetasHeatLoss8 %>%
  mutate("Effects" = "Direct Effects") %>%
  mutate(
    "Body Mass" = b_foldEhl_mass,
    "Tarsus Length" = b_foldEhl_tarsus,
    "Cold Rearing\n(10°C)" = b_foldEhl_pretreatmentA,
    "Warm Rearing\n(30°C)" = b_foldEhl_pretreatmentC
  ) %>%
  select(
```

```
  Effects, `Body Mass`, `Tarsus Length`,
  `Cold Rearing\n(10°C)`,
  `Warm Rearing\n(30°C)`
) %>%
rbind(
  .,
  scaledBetasHeatLoss8 %>%
    mutate("Effects" = "Indirect Effects") %>%
    mutate(
      "Body Mass" = b_tarsus_mass *
        b_foldEhl_tarsus,
      "Tarsus Length" = NA,
      "Cold Rearing\n(10°C)" =
        b_tarsus_pretreatmentA * b_foldEhl_tarsus +
          b_mass_pretreatmentA * b_foldEhl_mass +
          b_mass_pretreatmentA * b_tarsus_mass *
            b_foldEhl_tarsus,
      "Warm Rearing\n(30°C)" =
        b_tarsus_pretreatmentC * b_foldEhl_tarsus +
          b_mass_pretreatmentC * b_foldEhl_mass +
          b_mass_pretreatmentC * b_tarsus_mass *
            b_foldEhl_tarsus,
    ) %>%
    select(
      Effects, `Body Mass`, `Tarsus Length`,
      `Cold Rearing\n(10°C)`, `Warm Rearing\n(30°C)`
    )
) %>%
rbind(., scaledBetasHeatLoss8 %>%
  mutate("Effects" = "Total Effects") %>%
  mutate(
    "Body Mass" =
      b_foldEhl_mass +
        b_tarsus_mass * b_foldEhl_tarsus,
    "Tarsus Length" =
      b_foldEhl_tarsus,
    "Cold Rearing\n(10°C)" =
      b_foldEhl_pretreatmentA +
        b_mass_pretreatmentA *
          b_foldEhl_mass +
        b_tarsus_pretreatmentA *
          b_foldEhl_tarsus +
        b_mass_pretreatmentA *
          b_tarsus_mass * b_foldEhl_tarsus,
    "Warm Rearing\n(30°C)" =
      b_foldEhl_pretreatmentC +
        b_mass_pretreatmentC *
          b_foldEhl_mass +
        b_tarsus_pretreatmentC *
          b_foldEhl_tarsus +
        b_mass_pretreatmentC *
          b_tarsus_mass * b_foldEhl_tarsus,
  ) %>%
  select(
    Effects, `Body Mass`, `Tarsus Length`,
    `Cold Rearing\n(10°C)`, `Warm Rearing\n(30°C)`
  )) %>%
pivot_longer(c(-Effects), names_to = "var", values_to = "values") %>%
mutate(var = factor(var,
  levels = c(
    "Tarsus Length",
    "Body Mass",
    "Warm Rearing\n(30°C)",
    "Cold Rearing\n(10°C)"
  )
))
```

```
fullEffectEfficiency8 <- scaledBetasEfficiency8 %>%
  mutate("Effects" = "Direct Effects") %>%
  mutate(
    "Body Mass" = b_ecc_mass,
    "Tarsus Length" = b_ecc_tarsus,
    "Cold Rearing\n(10°C)" = b_ecc_pretreatmentA,
    "Warm Rearing\n(30°C)" = b_ecc_pretreatmentC
  ) %>%
  select(
    Effects, `Body Mass`, `Tarsus Length`,
    `Cold Rearing\n(10°C)`,
    `Warm Rearing\n(30°C)`
  ) %>%
  rbind(
    .,
    scaledBetasEfficiency8 %>%
      mutate("Effects" = "Indirect Effects") %>%
      mutate(
        "Body Mass" = b_tarsus_mass *
          b_ecc_tarsus,
        "Tarsus Length" = NA,
        "Cold Rearing\n(10°C)" =
          b_tarsus_pretreatmentA * b_ecc_tarsus +
            b_mass_pretreatmentA * b_ecc_mass +
            b_mass_pretreatmentA * b_tarsus_mass *
              b_ecc_tarsus,
        "Warm Rearing\n(30°C)" =
          b_tarsus_pretreatmentC * b_ecc_tarsus +
            b_mass_pretreatmentC * b_ecc_mass +
            b_mass_pretreatmentC * b_tarsus_mass *
              b_ecc_tarsus,
      ) %>%
      select(
        Effects, `Body Mass`, `Tarsus Length`,
        `Cold Rearing\n(10°C)`, `Warm Rearing\n(30°C)`
      )
  ) %>%
  rbind(., scaledBetasEfficiency8 %>%
    mutate("Effects" = "Total Effects") %>%
    mutate(
      "Body Mass" =
        b_ecc_mass +
          b_tarsus_mass * b_ecc_tarsus,
      "Tarsus Length" =
        b_ecc_tarsus,
      "Cold Rearing\n(10°C)" =
        b_ecc_pretreatmentA +
          b_mass_pretreatmentA *
            b_ecc_mass +
          b_tarsus_pretreatmentA *
            b_ecc_tarsus +
          b_mass_pretreatmentA *
            b_tarsus_mass * b_ecc_tarsus,
      "Warm Rearing\n(30°C)" =
        b_ecc_pretreatmentC +
          b_mass_pretreatmentC *
            b_ecc_mass +
          b_tarsus_pretreatmentC *
            b_ecc_tarsus +
          b_mass_pretreatmentC *
            b_tarsus_mass * b_ecc_tarsus,
    ) %>%
    select(
      Effects, `Body Mass`, `Tarsus Length`,
      `Cold Rearing\n(10°C)`, `Warm Rearing\n(30°C)`
    )) %>%
  pivot_longer(c(-Effects), names_to = "var", values_to = "values") %>%
```

```
  mutate(var = factor(var,
    levels = c(
      "Tarsus Length",
      "Body Mass",
      "Warm Rearing\n(30°C)",
      "Cold Rearing\n(10°C)"
    )
  ))

# And effects are plotted

fullEffectHeatLoss8Plot <- fullEffectHeatLoss8 %>%
  ggplot(aes(x = values, y = var, fill = var)) +
  facet_wrap(~Effects) +
  stat_halfeye(normalize = "xy", colour = "black", alpha = 0.7) +
  geom_vline(
    xintercept = 0, linetype = "dashed",
    colour = "black"
  ) +
  xlab("Effect on Fold Evaporative\nHeat Loss (standard deviations)") +
  scale_fill_manual(values = c("black", "grey50", "#CD5C5C", "#7BB4E3")) +
  theme_classic() +
  theme(
    legend.position = "none", axis.title.y = element_blank(),
    axis.text.y = element_text(
      size = 11, colour = "black",
      family = "Noto Sans"
    ),
    axis.title.x = element_text(family = "Noto Sans", hjust = -0.005),
    axis.text.x = element_text(family = "Noto Sans")
  )

fullEffectEfficiency8Plot <- fullEffectEfficiency8 %>%
  ggplot(aes(x = values, y = var, fill = var)) +
  facet_wrap(~Effects) +
  stat_halfeye(normalize = "xy", colour = "black", alpha = 0.7) +
  geom_vline(
    xintercept = 0, linetype = "dashed",
    colour = "black"
  ) +
  xlab("Effect on Evaporative Cooling\nEfficiency (standard deviations)") +
  scale_fill_manual(values = c("black", "grey50", "#CD5C5C", "#7BB4E3")) +
  theme_classic() +
  theme(
    legend.position = "none", axis.title.y = element_blank(),
    axis.text.y = element_text(
      size = 11, colour = "black",
      family = "Noto Sans"
    ),
    axis.title.x = element_text(family = "Noto Sans", hjust = -0.005),
    axis.text.x = element_text(family = "Noto Sans")
  )

(fullEffectHeatLoss8Plot /
  fullEffectEfficiency8Plot) +
  plot_annotation(tag_level = "A")
```

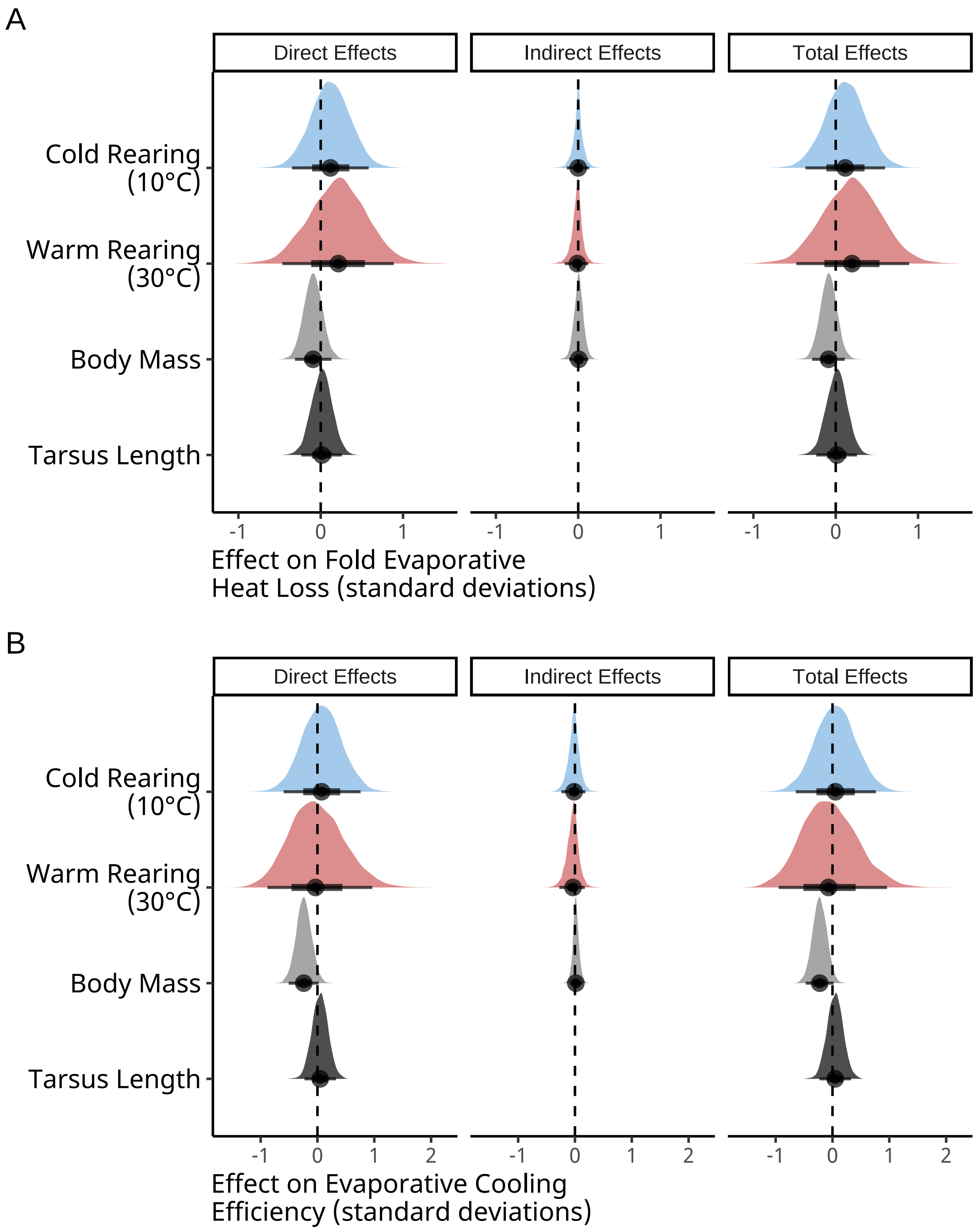


***Figure 25:*** *Direct and indirect effects of rearing temperature and morphology (here, body mass and tarsus length) on evaporative heat loss responses (panel A) and evaporative cooling efficiency (panel B) in developing Japanese quail (eight weeks of age; n = 57). Evaporative heat loss responses represent fold increases in evaporative heat loss, in Watts, between 30°C and 40°C. Evaporative cooling efficiency represents the ratio between evaporative heat loss (in Watts) and metabolic heat production (in Watts). Effects indicate how much, in standard deviations, a response variables would be altered by changing a predictor variable by one standard deviation. Densities are derived from posteriors of Bayesian path analyses. Dashed lines indicate 0.*

```
# EFfects are summarised below

caption <- paste0("Direct, indirect, and total effects of morphology and ",
                  "rearing temperature on fold evaporative heat loss at 40°C",
                  "(relative to 30°C) of eight week old Japanese quail. ",
                  "Effects are derived from ",
                  "a Bayesian path analysis and represent those predicted for a ",
                  "change in one standard deviation (or categorical level) of ",
                  "a given predictor on the standard deviation of metabolic ",
                  "slopes. Estimates indicate posterior medians and credible ",
                  "intervels (CIs) indicate quantile intervals."
)

heatLossResultsScaled8 <-
  fullEffectHeatLoss8 %>%
  filter(!is.na(values) & !is.nan(values)) %>%
  group_by(Effects, var) %>%
  summarise(
    "Estimate" = median(values),
    "50\\% CIs" = paste0(
      "[",
      round(
        quantile(values, probs = 0.1, type = 8),
        digits = 4
      ),
      ", ",
      round(
        quantile(values, probs = 0.9, type = 8),
        digits = 4
      ),
      "]"
    ),
    "95\\% CIs" = paste0(
      "[",
      round(
        quantile(values, probs = 0.025, type = 8),
        digits = 4
      ),
      ", ",
      round(
        quantile(values, probs = 0.975, type = 8),
        digits = 4
      ),
      "]"
    )
  ) %>%
  mutate("Effects" = gsub("[[:space:]].*", "", Effects)) %>%
  select(
    "Predictor" = "var", "Effect Level" = "Effects",
    Estimate, `50\\% CIs`, `95\\% CIs`
  ) %>%
  arrange(Predictor, `Effect Level`) %>%
  kbl(.,
    longtable = T, booktabs = T, format = "latex", escape = FALSE,
    caption = caption
  ) %>%
  column_spec(column = c(1:2), width = "2.2cm") %>%
  column_spec(column = c(3:10), width = "1.9cm") %>%
  kable_styling(latex_options = "striped")

heatLossResultsScaled8
```

***Table 12:*** *Direct, indirect, and total effects of morphology and rearing temperature on fold evaporative heat loss at 40°C(relative to 30°C) of eight week old Japanese quail. Effects are derived from a Bayesian path analysis and represent those predicted for a change in one standard deviation (or categorical level) of a given predictor on the standard deviation of metabolic slopes. Estimates indicate posterior medians and credible intervels (CIs) indicate quantile intervals.*

| Predictor | Effect Level | Estimate | 50% CIs | 95% CIs |
|---|---|---|---|---|
| Tarsus Length | Direct | 0.0152228 | [-0.1487, 0.1736] | [-0.2366, 0.2578] |
| Tarsus Length | Total | 0.0152228 | [-0.1487, 0.1736] | [-0.2366, 0.2578] |
| Body Mass | Direct | -0.0925756 | [-0.2344, 0.0477] | [-0.3122, 0.1303] |
| Body Mass | Indirect | 0.0061148 | [-0.0679, 0.0788] | [-0.1111, 0.1236] |
| Body Mass | Total | -0.0863202 | [-0.2156, 0.0407] | [-0.2858, 0.1106] |
| Warm Rearing (30°C) | Direct | 0.2151993 | [-0.2393, 0.6526] | [-0.466, 0.8877] |
| Warm Rearing (30°C) | Indirect | -0.0101400 | [-0.0961, 0.0609] | [-0.1634, 0.1235] |
| Warm Rearing (30°C) | Total | 0.1981495 | [-0.2524, 0.643] | [-0.479, 0.8918] |
| Cold Rearing (10°C) | Direct | 0.1179637 | [-0.1852, 0.4228] | [-0.3501, 0.5828] |
| Cold Rearing (10°C) | Indirect | -0.0000902 | [-0.0781, 0.0731] | [-0.1386, 0.1329] |
| Cold Rearing (10°C) | Total | 0.1164526 | [-0.1953, 0.434] | [-0.3684, 0.6002] |

```
caption <- paste0("Direct, indirect, and total effects of morphology and ",
                  "rearing temperature on evaporative cooling efficiency ",
                  "at 40°C  of eight week old Japanese quail. ",
                  "Evaporative cooling efficiency represents the ratio of ",
                  "evaporative heat loss (in W) to metabolic heat production ",
                  "(again, in W). Effects are derived from ",
                  "a Bayesian path analysis and represent those predicted for a ",
                  "change in one standard deviation (or categorical level) of ",
                  "a given predictor on the standard deviation of metabolic ",
                  "slopes. Estimates indicate posterior medians and credible ",
                  "intervels (CIs) indicate quantile intervals."
)

efficiencyResultsScaled8 <-
  fullEffectEfficiency8 %>%
  filter(!is.na(values) & !is.nan(values)) %>%
  group_by(Effects, var) %>%
  summarise(
    "Estimate" = median(values),
    "50\\% CIs" = paste0(
      "[",
      round(
        quantile(values, probs = 0.1, type = 8),
        digits = 4
      ),
      ", ",
      round(
        quantile(values, probs = 0.9, type = 8),
        digits = 4
      ),
      "]"
    ),
    "95\\% CIs" = paste0(
      "[",
```

```
    round(
      quantile(values, probs = 0.025, type = 8),
      digits = 4
    ),
    ", ",
    round(
      quantile(values, probs = 0.975, type = 8),
      digits = 4
    ),
    "]"
  )
) %>%
mutate("Effects" = gsub("[[:space:]].*", "", Effects)) %>%
select(
  "Predictor" = "var", "Effect Level" = "Effects",
  Estimate, `50\\% CIs`, `95\\% CIs`
) %>%
arrange(Predictor, `Effect Level`) %>%
kbl(.,
  longtable = T, booktabs = T, format = "latex", escape = FALSE,
  caption = caption
) %>%
column_spec(column = c(1:2), width = "2.2cm") %>%
column_spec(column = c(3:10), width = "1.9cm") %>%
kable_styling(latex_options = "striped")

efficiencyResultsScaled8
```

***Table 13:*** *Direct, indirect, and total effects of morphology and rearing temperature on evaporative cooling efficiency at 40°C of eight week old Japanese quail. Evaporative cooling efficiency represents the ratio of evaporative heat loss (in W) to metabolic heat production (again, in W). Effects are derived from a Bayesian path analysis and represent those predicted for a change in one standard deviation (or categorical level) of a given predictor on the standard deviation of metabolic slopes. Estimates indicate posterior medians and credible intervels (CIs) indicate quantile intervals.*

| Predictor | Effect Level | Estimate | 50% CIs | 95% CIs |
|---|---|---|---|---|
| Tarsus Length | Direct | 0.0478407 | [-0.1367, 0.2263] | [-0.2293, 0.3259] |
| Tarsus Length | Total | 0.0478407 | [-0.1367, 0.2263] | [-0.2293, 0.3259] |
| Body Mass | Direct | -0.2413953 | [-0.4121, -0.068] | [-0.507, 0.0217] |
| Body Mass | Indirect | 0.0151252 | [-0.0487, 0.0858] | [-0.0912, 0.1323] |
| Body Mass | Total | -0.2248253 | [-0.3838, -0.0638] | [-0.471, 0.0188] |
| Warm Rearing (30°C) | Direct | -0.0343423 | [-0.6007, 0.6072] | [-0.8809, 0.9651] |
| Warm Rearing (30°C) | Indirect | -0.0336755 | [-0.175, 0.0844] | [-0.2744, 0.1724] |
| Warm Rearing (30°C) | Total | -0.0694908 | [-0.6463, 0.5756] | [-0.9451, 0.9611] |
| Cold Rearing (10°C) | Direct | 0.0730438 | [-0.361, 0.5172] | [-0.5965, 0.7586] |
| Cold Rearing (10°C) | Indirect | -0.0156016 | [-0.1453, 0.0998] | [-0.2378, 0.1844] |
| Cold Rearing (10°C) | Total | 0.0525784 | [-0.3974, 0.5142] | [-0.6452, 0.7635] |

Again, we estimate the consequences of misalignment with Allen's and Bergmann's rule on evaporative heat loss responses and evaporative cooling efficiency at 40°C. Misalignment here represents have a relative tarsus length two standard deviations below average (Allen's Rule) or a body mass two standard deviations above average (Bergmann's rule).

```
data.frame(
  "Size" = c("Average", "Large (2x s.d. > mean)"),
  "pretreatment" = "B",
  "mass" = c(
    mean(efficiencyModel8Weeks$data$mass, na.rm = T),
    mean(efficiencyModel8Weeks$data$mass, na.rm = T) +
      2 * sd(efficiencyModel8Weeks$data$mass, na.rm = T)
  ),
  "tarsus" = 0
) %>%
  mutate(
    "ehl" = predict(ehlModel8Weeks,
      newdata = .,
      resp = "foldEhl",
      re_form = NA,
      robust = TRUE
    )[, "Estimate"],
    "se1" = predict(ehlModel8Weeks,
      newdata = .,
      resp = "foldEhl",
      re_form = NA,
      robust = TRUE
    )[, "Est.Error"],
    "ecc" = predict(efficiencyModel8Weeks,
      newdata = .,
      resp = "ecc",
      re_form = NA,
      robust = TRUE
    )[, "Estimate"],
    "se2" = predict(efficiencyModel8Weeks,
      newdata = .,
      resp = "ecc",
      re_form = NA,
      robust = TRUE
    )[, "Est.Error"]
  ) %>%
  mutate_if(is.numeric, round, digits = 4) %>%
  mutate(
    "Body Size" = Size,
    "Evaporative Heat Loss" = paste0(ehl, " [",
                                     se1, "]"
    ),
    "Evaporative Cooling Efficiency" = paste0(ecc, " [",
                                     se2, "]"
    )
  ) %>%
  select(`Body Size`, `Evaporative Heat Loss`, `Evaporative Cooling Efficiency`)
```

```
## # A tibble: 2 x 3
##   `Body Size`            `Evaporative Heat Loss` Evaporative Cooling Efficienc~1
##   <chr>                  <chr>                   <chr>
## 1 Average                2.414  [0.7532]         0.6752  [0.1894]
## 2 Large (2x s.d. > mean) 2.2437  [0.788]         0.5885  [0.1935]
## # i abbreviated name: 1: `Evaporative Cooling Efficiency`
```

```
data.frame(
  "Size" = c("Average", "Short (2x s.d. < mean)"),
  "pretreatment" = "B",
  "tarsus" = c(
    mean(efficiencyModel8Weeks$data$tarsus, na.rm = T),
    mean(efficiencyModel8Weeks$data$tarsus, na.rm = T) -
      2 * sd(efficiencyModel8Weeks$data$tarsus, na.rm = T)
  ),
  "mass" = 0
) %>%
  mutate(
```

```
    "ehl" = predict(ehlModel8Weeks,
      newdata = .,
      resp = "foldEhl",
      re_form = NA,
      robust = TRUE
    )[, "Estimate"],
    "se1" = predict(ehlModel8Weeks,
      newdata = .,
      resp = "foldEhl",
      re_form = NA,
      robust = TRUE
    )[, "Est.Error"],
    "ecc" = predict(efficiencyModel8Weeks,
      newdata = .,
      resp = "ecc",
      re_form = NA,
      robust = TRUE
    )[, "Estimate"],
    "se2" = predict(efficiencyModel8Weeks,
      newdata = .,
      resp = "ecc",
      re_form = NA,
      robust = TRUE
    )[, "Est.Error"]
  ) %>%
  mutate_if(is.numeric, round, digits = 4) %>%
  mutate(
    "Tarsus Length" = Size,
    "Evaporative Heat Loss" = paste0(ehl, " [",
                                     se1, "]"
    ),
    "Evaporative Cooling Efficiency" = paste0(ecc, " [",
                                     se2, "]"
    )
  ) %>%
  select(`Tarsus Length`, `Evaporative Heat Loss`,
         `Evaporative Cooling Efficiency`)
```

```
## # A tibble: 2 x 3
##   `Tarsus Length`        `Evaporative Heat Loss` Evaporative Cooling Efficienc~1
##   <chr>                  <chr>                   <chr>
## 1 Average                2.4043 [0.7547]         0.6671 [0.1849]
## 2 Short (2x s.d. < mean) 2.3634 [0.7769]         0.6494 [0.1886]
## # i abbreviated name: 1: `Evaporative Cooling Efficiency`
```
