## Supplementary material for "No evidence that shrinking and shapeshifting meaningfully affect how birds respond to warming and cooling": Supp. 5

### Estimating Lower Critical Temperature

#### Overview

Previous studies have argued that alignment with Allen's and Bergmann's rules provide thermal advantages in the cold by decreasing an individual's lower critical temperature (e.g. Kendeigh, 1969). In this document, we test whether: (1) lower critical temperature evidently varies among adult quail from our sample population, and (2) whether lower critical temperature does indeed co-vary with adult body mass and/or limb length (here, tarsus length; mm), as might be predicted by Allen's and Bergmann's rules.

##### Methods and analysis

To estimate mean lower critical temperature in our quail, we first measured resting metabolism (mL $O_2$/min) in a subsample of adult birds (n = 40; age 20 weeks) at 5°C intervals ranging from 0°C to 35°C. Measurements were obtained by flow-through respirometry following methods described for adult quail in the main text of Tabh et al (2024). Body mass of individuals was measured using a digital scale immediately before respirometry; tarsus length was measured digitally (described in Tabh et al 2024) at 12 weeks of age.

To obtain lower critical temperature estimates from resting metabolism data, we constructed a Bayesian piece-wise regression with resting metabolism as the Gaussian-distributed response variable and ambient temperature (continuous) as the sole population-level predictor. In this model, we assumed that resting metabolism was constant at ambient temperatures above a critical limit (the lower critical temperature, here defined as a break-point) and decreased at a constant linear rate below the critical limit (defined as "thermal resistance"). To evaluate the degree to which lower critical temperatures might differ by individual, our break-point, thermal resistance, and metabolism intercepts were allowed to vary by individual around a population mean (i.e. by inclusion of population; and group-level intercepts for each parameter). Further, to simplify model interpretation, we defined 35°C, an ambient temperature known to fall within thermoneutrality for quail (Ben-Hamo et al, 2010), as our regression intercept and relativised remaining temperature values as absolute deviations from 35°C (e.g. 35°C = 0°C, 0°C = 35°C). By doing so, we defined metabolism at thermoneutrality as our model "reference point".

Priors for our model break-point and and resting metabolism at thermoneutrality (model intercept) were skew-normal with $\xi$, $\omega$, and $\alpha$ values of 6.5 and 4.5, 5 and 2, and 2.5 and 4 respectively (setting means at approximately 5°C and 6 mL $O_2$/min; see Ben-Hamo et al, 2010 and Persson et al, 2024), while that for thermal resistance was also skew-normal ($\xi = 0.07$, $\omega = 0.1$, $\alpha = 9$; informed by Saarela and Heldmaier, 1978; Persson et al, 2024). Priors for individual variation around break-points, resting metabolism at thermoneutrality, and thermal resistance were exponential with lambda ($\lambda$) values of 0.5, 0.5, and 10 respectively.

To account for any skewness in posterior densities, model coefficients are summarised as posterior medians to and confidence around coefficients was calculated at 50% and 95% (via quantile intervals). All analyses were conducted using R Statistical Software (2023; version 4.2.3) and executed first on a linux platform (kernel: 5.15.0-46-generic).

Below, we begin by loading in R-packages required for analysis, then importing and checking data for anomalies.

```
def.chunk.hook <- knitr::knit_hooks$get("chunk")
knitr::knit_hooks$set(chunk = function(x, options) {
  x <- def.chunk.hook(x, options)
  ifelse(options$size != "normalsize",
    paste0("\n \\", options$size, "\n\n", x, "\n\n \\normalsize"), x
  )
})

knitr::opts_chunk$set(fig.pos = "H", out.extra = "")
knitr::opts_chunk$set(size = "footnotesize")

# First loading in packages

library("tidyverse")
library("easypackages")

packageList <- c("bayesplot", "brms", "brmsMethods",
                 "doParallel", "foreach", "ggpubr",
                 "kableExtra", "latex2exp", "patchwork",
                 "priorsense", "showtext", "tidybayes",
                 "wesanderson")

libraries(packageList)

caption <- paste0("R packages and their respective versions used for",
                  " data organisation and analysis in this study."
)

sapply(packageList, function(x) {
  y <- as.character(packageVersion(x))
  return(y)
}, simplify = FALSE) %>%
  enframe(., name = "Package", value = "Version") %>%
  as.data.frame(.) %>%
  kbl(.,
    longtable = T, booktabs = T,
    caption = caption
  ) %>%
  kable_styling(latex_options = "striped")
```

***Table 1:*** *R packages and their respective versions used for data organisation and analysis in this study.*

| Package | Version |
|---|---|
| bayesplot | 1.11.1 |
| brms | 2.20.4 |
| brmsMethods | 0.0.0.9000 |
| doParallel | 1.0.17 |
| foreach | 1.5.2 |
| ggpubr | 0.6.0 |
| kableExtra | 1.3.4 |
| latex2exp | 0.9.6 |
| patchwork | 1.2.0 |
| priorsense | 0.0.0.9000 |

| | |
|---|---|
| showtext | 0.9.6 |
| tidybayes | 3.0.4 |
| wesanderson | 0.3.6.9000 |

```r
# Loading additional functions

pp_check2 <- function(model, resp = NA, ndraws = 500,
                      xlab = "label", colour = "lightblue") {
  require(brms)
  require(ggplot2)
  stopifnot("Model must be a brmsfit object" = is.brmsfit(model))

  if (is.na(resp)) {
    resp <- model$formula$resp
  }

  p1 <- brms::pp_check(model, ndraws = ndraws, resp = resp) +
    scale_colour_manual(
      values = c("black", colour),
      labels = c("y", "yhat"),
      name = NULL
    ) +
    xlab(xlab) +
    ylab("Density") +
    theme_classic()
  return(p1)
}

chainCheck <- function(model, rDig = 3) {
  require(brms)
  stopifnot("Model must be a brmsfit object" = is.brmsfit(model))

  Rhat <- paste0(
    "Rhat range: ",
    round(min(rhat(model)), digits = rDig),
    " - ",
    round(max(rhat(model)), digits = rDig)
  )
  Neff <- paste0(
    "Neff/N range: ",
    round(min(neff_ratio(model)), digits = rDig),
    " - ",
    round(max(neff_ratio(model)), digits = rDig)
  )
  cat(paste0(Rhat, "\n", Neff))
}

quantileCIs <- function(x, rnd = 3, cis = c(50, 95), sci_note = FALSE) {
  require(tidyverse)

  if (class(x)[1] != "brmsfit") {
    return("x must be a brmsfit object.")
  }
```

```
  if (length(cis) != 2) {
    return("cis must be a vector of integers with length 2")
  }

  prbs = c()
  nColNames = c()
  for (i in 1:length(cis)){
    prbs = c(prbs, c(0.5 - (cis[i]/100)/2, 0.5 + (cis[i]/100)/2))
    nColNames = c(nColNames,
                  paste0("Low_CI_", cis[i]),
                  paste0("High_CI_", cis[i])
    )
  }

  modelFrame = as.data.frame(x)

  Results <- apply(modelFrame, MARGIN = 2, FUN = quantile,
        probs = prbs, type = 8) %>%
    t() %>%
    as.data.frame() %>%
    rownames_to_column(var = "par") %>%
    `colnames<-`(c("Parameter", nColNames))

  if (sci_note == FALSE) {
    Results <- apply(modelFrame, MARGIN = 2, FUN = quantile,
        probs = prbs, type = 8) %>%
    t() %>%
    as.data.frame() %>%
    rownames_to_column(var = "par") %>%
    `colnames<-`(c("Parameter", nColNames))

  } else if (sci_note == TRUE) {
    Results <- apply(modelFrame, MARGIN = 2, FUN = quantile,
        probs = prbs, type = 8) %>%
    t() %>%
    as.data.frame() %>%
    rownames_to_column(var = "par") %>%
    `colnames<-`(c("Parameter", nColNames)) %>%
    mutate_at(.vars = vars(-Parameter),
              .funs = function(x){
                return(format(x, scientific = TRUE))
              }
    )
  }

  return(Results)
}

# Installing font

font_add_google(name = "Noto Sans", family = "Noto Sans")

# Setting working directory
```

```
setwd(paste0("/Users/joshuatabh/Documents/",
  "researchProjects/lund/functionOfAllen/functionData")
)

# Loading in and restructuring metabolism data for use

rmrData <- read.csv("lctRMRData.csv") %>%
  rename("fileName" = FileName_truncated) %>%
  merge(., read.csv("lctRMRBirdIDs.csv") %>%
    select(
      "ring" = Ring, "sex" = Sex, Chamber,
      fileName = "respirometry_file"
    ),
  by = c("fileName", "Chamber"), all.x = TRUE
  ) %>%
  rowwise() %>%
  mutate("vo2" = ifelse(Chamber == 1,
    VO2_1,
    ifelse(Chamber == "2",
      VO2_2,
      ifelse(Chamber == 3,
        VO2_3,
        VO2_4
      )
    )
  )) %>%
  ungroup() %>%
  filter(FileNumber >= 2) %>%
  merge(., data.frame(
    "FileNumber" = seq(2, 9, by = 1),
    "Ta" = seq(35, 0, by = -5)
  ),
  by = "FileNumber", all.x = TRUE
  ) %>%
  select(ring, sex,
    "chamber" = Chamber,
    Ta, "VO2" = vo2, fileName
  ) %>%
  filter(Ta <= 35)

# Relativising ambient temperature measures with respect to known
# thermoneutrality

rmrData$relativeTa = abs(rmrData$Ta - 35)

# Producing dotplot to inspect data visually

rmrData %>%
  group_by(Ta) %>%
  mutate("Row" = 1:n()) %>%
  mutate(Ta = paste0(Ta, "°C")) %>%
  ungroup() %>%
  ggplot(
```

```
  aes(x = Row, y = VO2)) +
  facet_wrap(~Ta, scales = "free") +
  geom_point() +
  xlab("Sample Number") +
  ylab(bquote(Resting ~ Metabolism ~ (mL ~ O[2] / min))) +
  theme_classic() +
  theme(legend.position = "none")
```

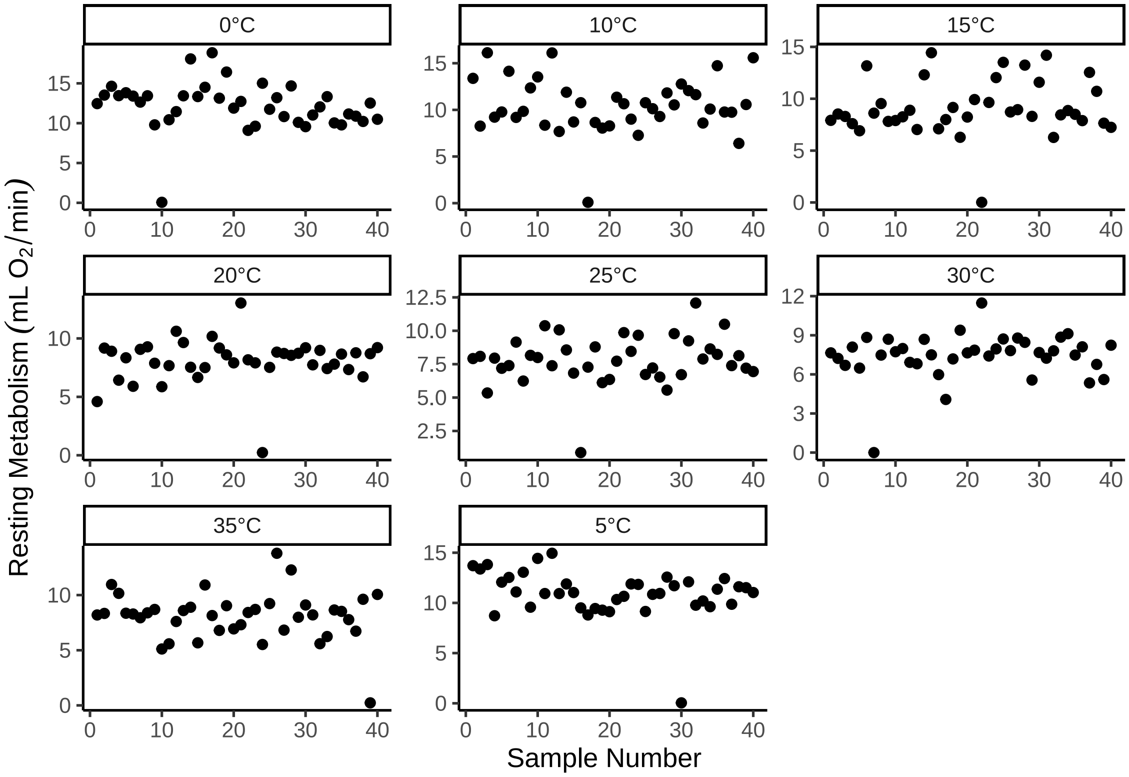


***Figure 1:*** *Cleveland dotplot of resting metabolism values by ambient temperature of collection, each drawn from adult Japanese quail. Dots represent raw data points.*

```
# Several points indicating an unrealistic resting metabolism of 0.
# Filtering these points out and proceeding.

rmrData = rmrData %>%
  filter(VO2 > 3)
```

Next, we check the suitability of our piece-wise regression priors. This is achieved by estimating possible resting metabolism values from our priors alone, then comparing these values against true resting metabolism measurements.

```
# First constructing model formula

bform <- bf(
  VO2 ~ Intercept +
    (resistance * (relativeTa - breakPoint)) * step(relativeTa - breakPoint),
  Intercept + resistance + breakPoint ~ 1 + (1|ring),
  nl = TRUE
)
```

```r
# Constructing model priors

breakPriors <- c(
  set_prior("skew_normal(4.5, 2, 4)", nlpar = "Intercept"),
  set_prior("skew_normal(6.5, 5, 2.5)", nlpar = "breakPoint"),
  set_prior("skew_normal(0.07, 0.01, 9)", nlpar = "resistance"),
  set_prior("exponential(0.5)", nlpar = "Intercept", class = "sd"),
  set_prior("exponential(0.5)", nlpar = "breakPoint", class = "sd"),
  set_prior("exponential(10)", nlpar = "resistance", class = "sd")
)

# Fitting model but only sampling from priors

priorCheck <- brm(bform, data = rmrData,
            prior = breakPriors,
            iter = 50000, warmup = 10000, thin = 10,
            cores = 4, chains = 4,
            silent = TRUE, refresh = 0,
            sample_prior = "only",
            file = "./models/lowerCriticalTemperatureAnalysisPPCheck.Rds"
            )

pp_check2(priorCheck,
  xlab = TeX('$Resting~Metabolism~(mL~O_{2}/min)$')) +
  xlim(c(0, 50))
```

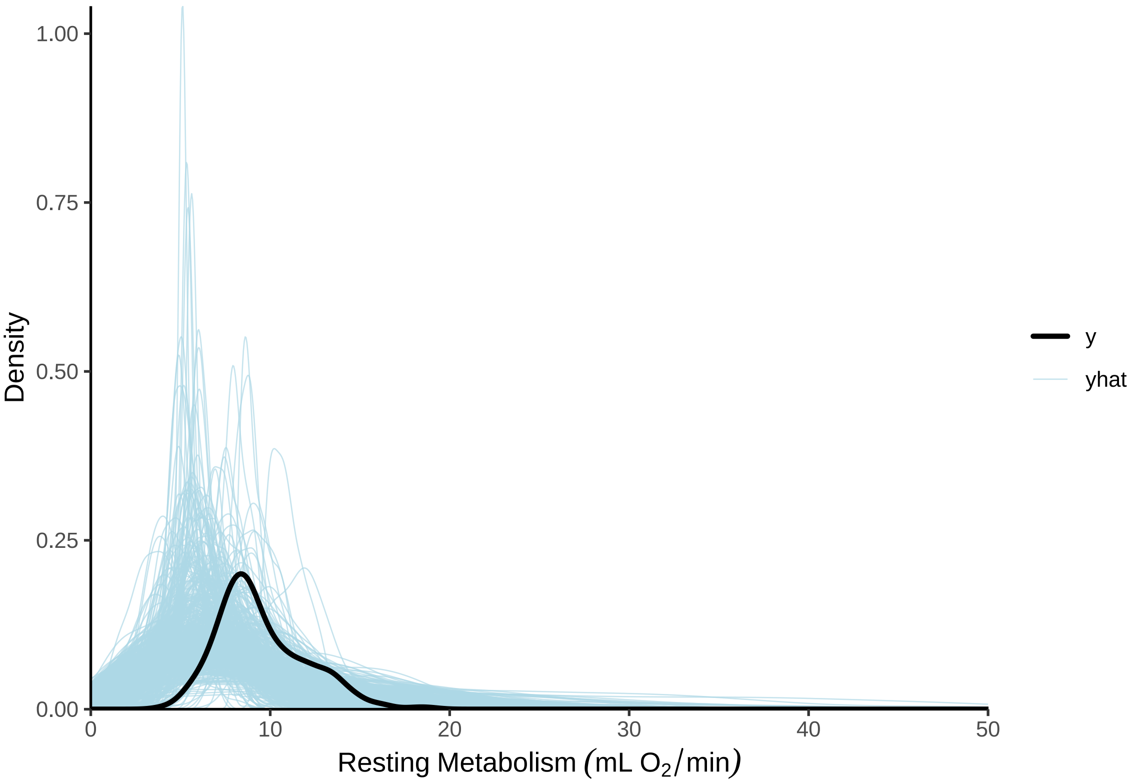


***Figure 2:*** *Prior predictive check for a Bayesian piece-wise regression predicting resting metabolism (mL $O_2$/min) of adult Japanese quail by ambient temperature (raw ambient temperatures ranging from 0°C - 30°C). Resting metabolism is assumed to remain constant at ambient temperatures above an estimated breakpoint (the lower critical temperature) and increase linearly below that ambient temperature. Light blue lines represent densities of resting metabolism values as predicted by model priors alone. The dark blue line represent the true density of resting metabolism values. Clear overlap between the dark blue and light blue lines indicates that priors are suitable.*

Given clear overlap between the density of true resting metabolism values and those derived from model priors, we proceed with executing our full model as parameterised.

```
model <- brm(bform, data = rmrData,
            prior = breakPriors,
            iter = 100000, warmup = 10000, thin = 10,
            cores = 4, chains = 4,
            silent = TRUE, refresh = 0,
            file = "./models/lowerCriticalTemperatureAnalysis.Rds"
            )

# Few divergent transitions in HMC chains; little concern.
# Checking Gelman-Rubin statistics (Rhat values) and ratio of effective
# sample sizes to sample sizes.

chainCheck(model)
```

```
## Rhat range: 1 - 1
## Neff/N range: 0.895 - 1.003
```

```
# Rhat values all close to 1 indicating clear chain mixing.
# Also, ratio of effective sample sizes to sample sizes all near 1,
# suggesting little autocorrelation within sample draws.

# Viewing posterior densities and fitted trends

pp_check2(model,
  xlab = TeX('$Resting~Metabolism~(mL~O_{2}/min)$')) +
  xlim(c(0, 50))
```

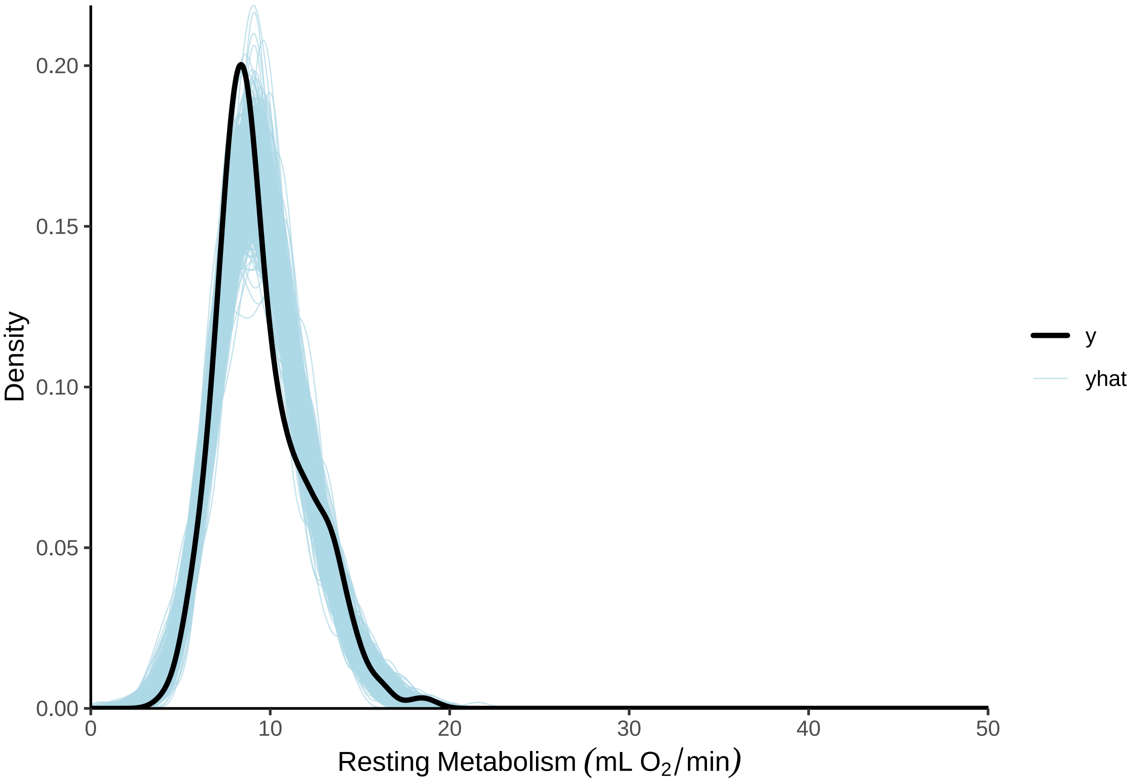


***Figure 3:*** *Posterior predictive check for a Bayesian piece-wise regression predicting resting metabolism (mL $O_2$/min) of adult Japanese quail by ambient temperature (raw ambient temperatures ranging from 0°C - 30°C).Light blue lines represent densities of resting metabolism values as predicted by model posteriors. The dark blue line represent the true density of resting metabolism values.*

```
model$data %>%
  mutate("fit" = fitted(model, robust = TRUE)[, "Estimate"]) %>%
  mutate(Ta = 35 - relativeTa) %>%
  ggplot(aes(x = Ta, y = fit, colour = ring)) +
  geom_point(data = rmrData, aes(x = Ta, y = VO2)) +
  geom_line() +
  facet_wrap(~ring) +
  scale_color_grey() +
  xlab("Ambient Temperature (°C)") +
  ylab(TeX('$Resting~Metabolism~(mL~O_{2}/min)$')) +
  theme_classic() +
  theme(legend.position = "none")
```

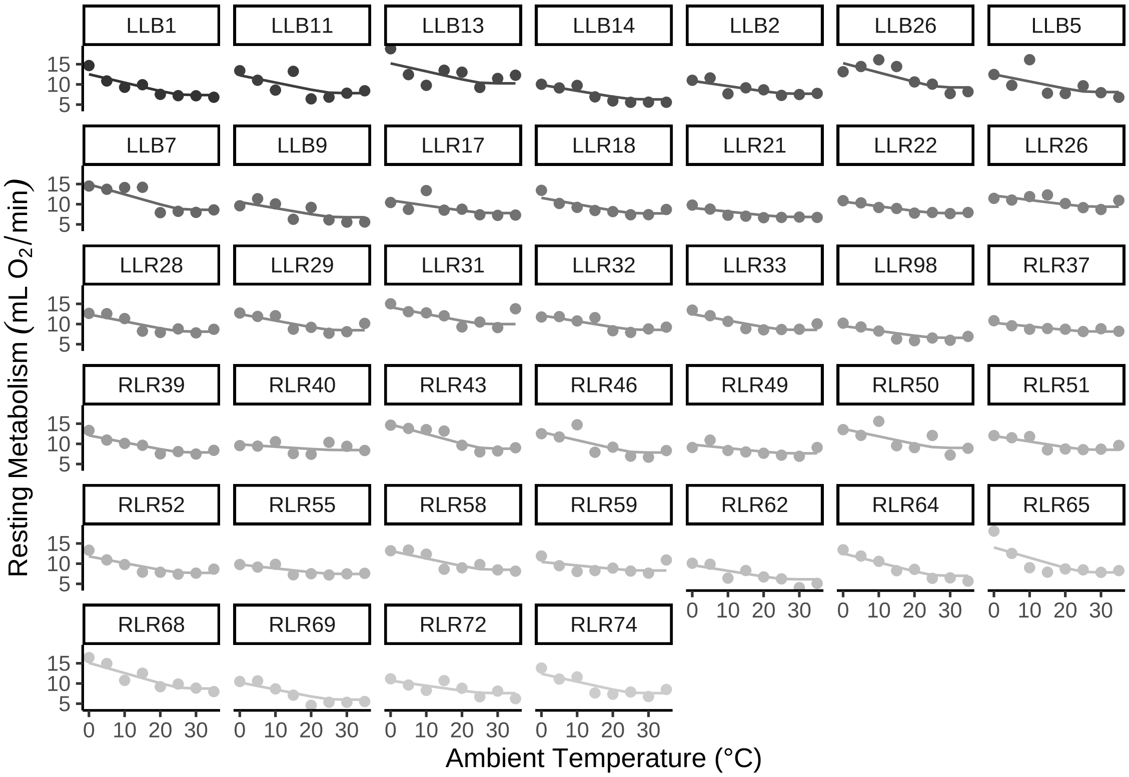


***Figure 4:*** *Predictions from a Bayesian piece-wise regression correlating resting metabolism (mL $O_2$/min) with ambient temperature in adult Japanese quail. Dots represent raw values per individual and lines represent lines of best fit derived from regression.*

Next, we check model residuals for abnormalities.

```
p1 = model$data %>%
  mutate("Residuals" = residuals(model, robust = TRUE)[,"Estimate"]) %>%
  ggplot(aes(x = Residuals)) +
  geom_density() +
  xlab("Ordinary Residuals") +
  ylab("Density") +
  theme_classic()

p2 = model$data %>%
  mutate("Residuals" = residuals(model, robust = TRUE)[,"Estimate"]) %>%
  mutate("Ta" = 30 - relativeTa) %>%
  ggplot(aes(x = Ta, y = Residuals)) +
  geom_point() +
  geom_smooth(colour = "black", linetype = "dashed", se = FALSE) +
```

```
  xlab("Ambient Temperature (°C)") +
  ylab("Ordinary Residuals") +
  theme_classic()

p3 = model$data %>%
  mutate("Residuals" = residuals(model, robust = TRUE)[,"Estimate"],
         "Fitted" = fitted(model, robust = TRUE)[,"Estimate"]) %>%
  ggplot(aes(x = Fitted, y = Residuals)) +
  geom_point() +
  xlab(TeX('$Fitted~Values~(mL~O_{2}/min)$')) +
  ylab("Ordinary Residuals") +
  theme_classic()

(p1 + p2)/p3 + plot_annotation(tag_levels = "A")
```

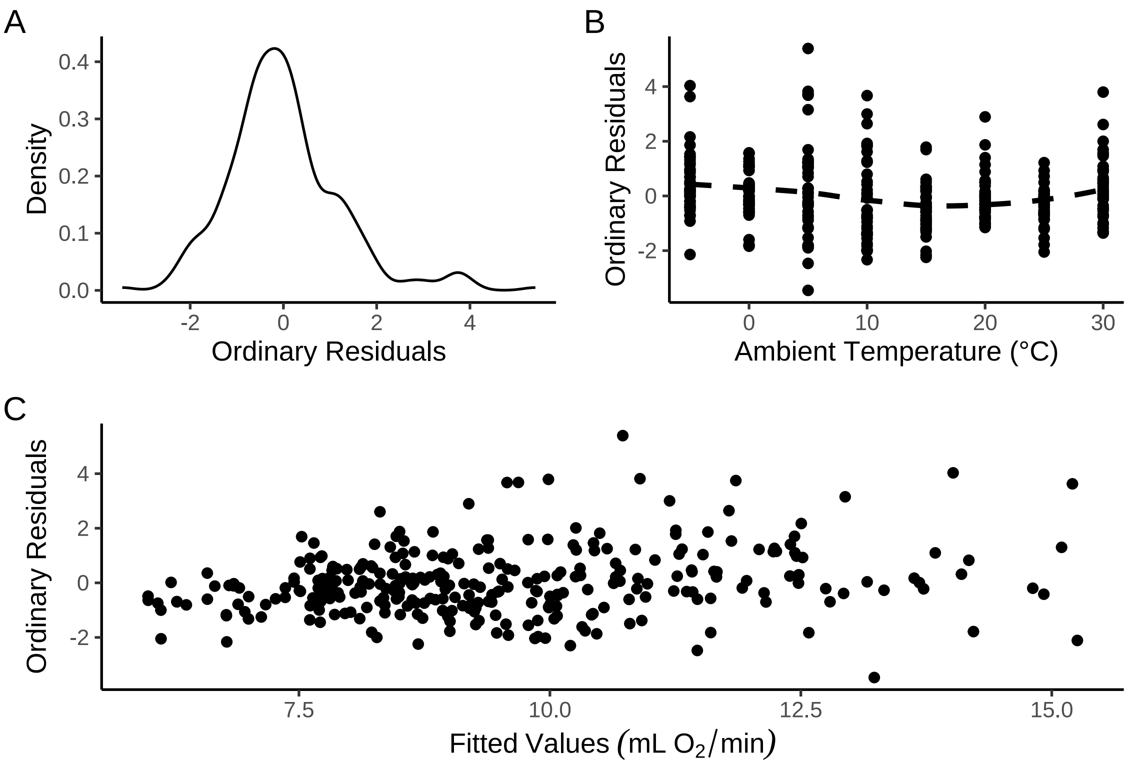


***Figure 5:*** *Spread of ordinary residuals (here, medians) from a Bayesian piece-wise regression predicting resting metabolism (mL $O_2$/min) of adult Japanese quail across ambient temperature (°C). Dots represent individual data points (medians). In panel B, the dotted line represents a loess line of best fit as estimated by the R package ggplot2 (Wickham, 2011). Fitted values in panel C represent posterior medians.*

```
# Reasonable spread
```

Densities of our model coefficients are below visualised and summarised at their medians ± 50% and 95% quantile intervals.

```
as.data.frame(model) %>%
  pivot_longer(everything(), names_to = "par", values_to = "values") %>%
  filter(grepl("b_|sd_", par)) %>%
  mutate(values = ifelse(par == "b_breakPoint_Intercept",
                         35 - values, values),
         values = ifelse(par == "b_resistance_Intercept",
                         values*-1, values)) %>%
  merge(.,
```

```
        tribble(~par, ~Parameter,
                "b_Intercept_Intercept", "Metabolism at 35°C\n(mL O2/min)",
                "b_resistance_Intercept", "Thermal Resistance\n(mL O2/min/°C)",
                "b_breakPoint_Intercept", "Lower Critical Temperature\n(°C)",
                "sd_ring__Intercept_Intercept",
                "Group-level\nMetabolism Intercept",
                "sd_ring__resistance_Intercept",
                "Group-level\nResistance Intercept",
                "sd_ring__breakPoint_Intercept",
                "Group-level Lower\nCritical Temperature\nIntercept"),
        by = 'par', all.x = TRUE
  ) %>%
  ggplot(aes(x = values)) +
    facet_wrap(~Parameter, scales = "free", strip.position = "bottom") +
    geom_density() +
    ylab("Density") +
    theme_classic() +
    theme(axis.title.x = element_blank(),
          strip.background = element_blank(),
          strip.placement = "outside",
          strip.text.x = element_text(size = 10))
```

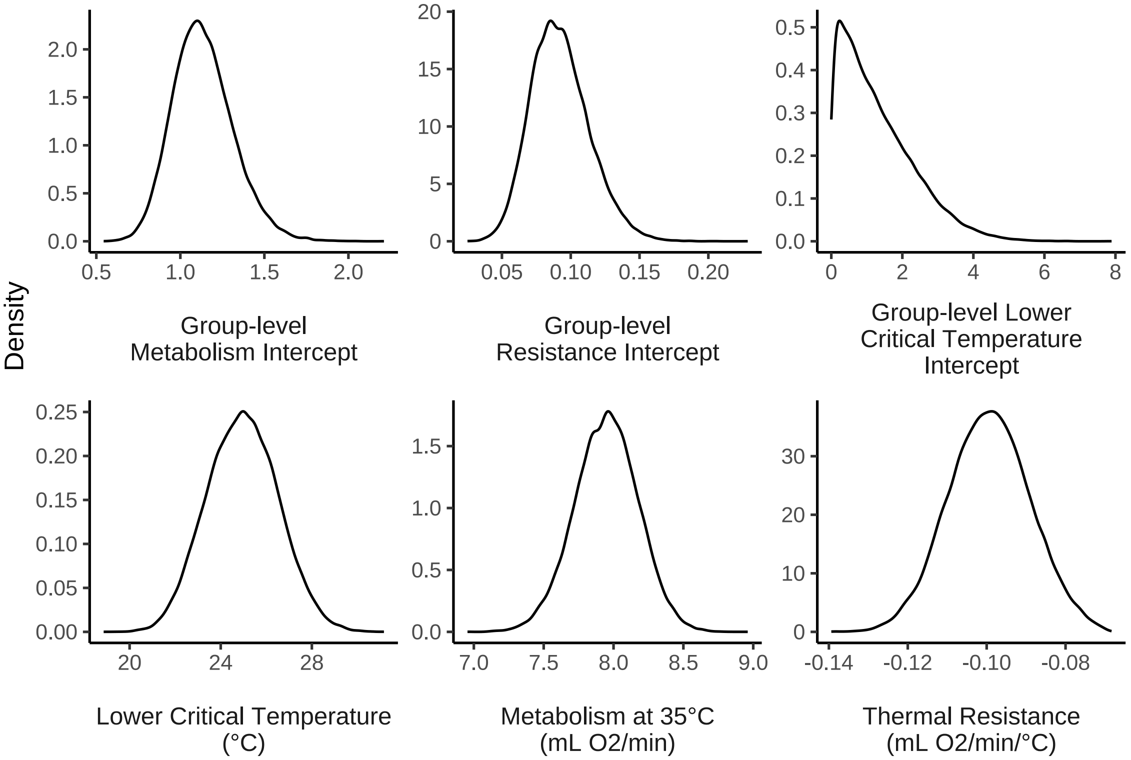


***Figure 6:*** *Posterior densities of coefficients from a Bayesian piece-wise regression predicting resting metabolism (mL $O_2$/min) of adult Japanese quail across ambient temperature (°C).*

```
caption = paste0("Results of a Bayesian piece-wise regression ",
                 "predicting resting metabolism (mL O\\textsubscript{2}/min) ",
                 "of adult Japanese quail across ambient temperature ",
                 "(°C). Break-points (here, lower critical ",
                 "temperature), thermal resistance, and metabolism at ",
                 "thermoneutrality was allowed to vary by individual. ",
                 "Estimates represent posterior medians and credible ",
                 "intervals represent quantile intervals.")

as.data.frame(model) %>%
```

```
  mutate("b_breakPoint_Intercept" = 35 - b_breakPoint_Intercept,
         "b_resistance_Intercept" = b_resistance_Intercept*-1) %>%
  summarise_all(.funs = function(x) round(median(x), digits = 3)) %>%
  pivot_longer(everything(), names_to = "par", values_to = "median") %>%
  merge(.,
  apply(as.data.frame(model),
        FUN = quantile, probs = c(0.025, 0.1, 0.9, 0.975),
        MARGIN = 2) %>%
  t() %>%
  as.data.frame() %>%
  mutate_all(.funs = round, digits = 3) %>%
  rownames_to_column(var = "par") %>%
  filter(grepl("b_|sd_|sigma", par)) %>%
  mutate(`50\\% CIs` = paste0("[", `10%`, ", ", `90%`, "]"),
         `95\\% CIs` = paste0("[", `2.5%`, ", ", `97.5%`, "]")
         ), by = "par",
  all.y = TRUE) %>%
  merge(.,
        tribble(~par, ~Parameter,
                "b_Intercept_Intercept",
                "Metabolism at 35°C (mL O\\textsubscript{2}/min)",
                "b_resistance_Intercept",
                "Thermal Resistance (mL O\\textsubscript{2}/min/°C)",
                "b_breakPoint_Intercept",
                "Lower Critical Temperature (°C)",
                "sd_ring__Intercept_Intercept",
                "Bird ID (Metabolism Intercept)",
                "sd_ring__resistance_Intercept",
                "Bird ID (Thermal Resistance)",
                "sd_ring__breakPoint_Intercept",
                "Bird ID (Lower Critical Temperature)",
                "sigma", "Sigma"),
        by = 'par', all.x = TRUE
  ) %>%
  select(Parameter, "Estimate" = median, `50\\% CIs`, `95\\% CIs`) %>%
  kbl(.,
    longtable = T, booktabs = T, format = "latex",
    caption = caption, escape = FALSE
  ) %>%
  kable_styling(latex_options = "striped")
```

***Table 2:*** *Results of a Bayesian piece-wise regression predicting resting metabolism (mL $O_2$/min) of adult Japanese quail across ambient temperature (°C). Break-points (here, lower critical temperature), thermal resistance, and metabolism at thermoneutrality was allowed to vary by individual. Estimates represent posterior medians and credible intervals represent quantile intervals.*

| Parameter | Estimate | 50% CIs | 95% CIs |
|---|---|---|---|
| Lower Critical Temperature (°C) | 24.962 | [8.026, 12.051] | [6.968, 13.004] |
| Metabolism at 35°C (mL $O_2$/min) | 7.951 | [7.658, 8.237] | [7.49, 8.388] |
| Thermal Resistance (mL $O_2$/min/°C) | -0.099 | [0.086, 0.113] | [0.079, 0.12] |
| Bird ID (Lower Critical Temperature) | 1.063 | [0.175, 2.75] | [0.043, 3.794] |
| Bird ID (Metabolism Intercept) | 1.123 | [0.916, 1.371] | [0.818, 1.529] |
| Bird ID (Thermal Resistance) | 0.091 | [0.067, 0.121] | [0.055, 0.138] |
| Sigma | 1.354 | [1.275, 1.441] | [1.235, 1.491] |

Finally, we test whether individual body mass and tarsus length predict lower critical temperature in quail.

```
base = 35 - as.data.frame(model)$b_breakPoint_Intercept
grab = as.data.frame(model) %>%
  select(starts_with("r_ring__breakPoint")) %>%
  mutate_all(.funs = function(x){return(base - x)}) %>%
  summarise_all(.funs = mean) %>%
  pivot_longer(everything(), names_to = "ring", values_to = "LCT") %>%
```

```
   mutate(ring = gsub("\\,.*", "",
    gsub(".*\\[", "", ring)
    )
  )

base <- 35 - as.data.frame(model)$b_breakPoint_Intercept

modData <- as.data.frame(model) %>%
  select(starts_with("r_ring__breakPoint")) %>%
  mutate_all(.funs = function(x){return(base - x)}) %>%
  summarise_all(.funs = mean) %>%
  pivot_longer(everything(), names_to = "ring", values_to = "LCT") %>%
   mutate(ring = gsub("\\,.*", "",
    gsub(".*\\[", "", ring)
    )
  ) %>%
  merge(., read.csv("lctRMRBirdIDs.csv") %>%
  select("ring" = Ring, "mass" = mass_in),
  by = "ring", all.x = TRUE) %>%
  merge(., read.csv("springWeek12Tarsus.csv"),
        by = "ring", all.x = TRUE)

lctModel <- brm(
  data = modData %>%
    select(ring, LCT, mass, tarsus) %>%
    drop_na(),
  bf(LCT ~ mass + tarsus),
  family = "gaussian",
  prior = c(
    set_prior("normal(25, 2.5)", class = "Intercept"),
    set_prior("normal(0, 0.1)", class = "b", coef = "mass"),
    set_prior("normal(0, 0.5)", class = "b", coef = "tarsus"),
    set_prior("exponential(0.5)", class = "sigma")
  ),
  iter = 100000, warmup = 50000, thin = 10,
  chains = 4, cores = 4,
  control = list(adapt_delta = .98, max_treedepth = 14),
  silent = TRUE, refresh = 0,
  file = "./models/lowerCriticalTemperatureMorphometryModel.Rds"
)

as.data.frame(lctModel) %>%
  summarise_all(., .funs = median) %>%
  pivot_longer(everything(), names_to = "Parameter",
               values_to = "Estimate") %>%
  merge(., quantileCIs(lctModel, cis = c(50, 95)),
        by = "Parameter", all.x = TRUE) %>%
  filter(grepl("b_|sd_", Parameter)) %>%
  rowwise() %>%
  mutate("BF" = ifelse(Estimate < 0,
                       (2 * mean(as.data.frame(
                         lctModel)[, Parameter] <= 0)) /
    (2 * mean(as.data.frame(
      lctModel)[, Parameter] >= 0)),
  (2 * mean(as.data.frame(
    lctModel)[, Parameter] >= 0)) /
    (2 * mean(as.data.frame(
      lctModel)[, Parameter] <= 0))
  )) %>%
  ungroup() %>%
  mutate(
    "Estimate" = round(Estimate, digits = 4),
    "BF" = round(BF, digits = 4),
    "N" = nrow(lctModel$data)
  ) %>%
  mutate("Parameter" = gsub("b_|sd_", "",
                            Parameter)) %>%
```

```
  mutate(
    "Response" = "Lower Critical Temperature",
    "Parameter" = gsub(".*_", "", Parameter)
  ) %>%
  merge(., tribble(
    ~Parameter, ~parameter, ~number,
    "Intercept", "Intercept", "1",
    "mass", "Body Mass (g)", "2",
    "tarsus", "Tarsus Length (mm)", "3"
  ),
  by = "Parameter"
  ) %>%
  mutate(
    `50\\% CI` = paste0("(", paste(
      round(Low_CI_50, digits = 4),
      round(High_CI_50, digits = 4),
      sep = ", "
    ), ")"),
    `95\\% CI` = paste0("(", paste(
      round(Low_CI_95, digits = 4),
      round(High_CI_95, digits = 4),
      sep = ", "
    ), ")")
  ) %>%
  select(-c(Low_CI_50, High_CI_50, Low_CI_95, High_CI_95)) %>%
  select(
    Response, "Parameter" = "parameter", N,
    Estimate, `50\\% CI`, `95\\% CI`, BF, number
  ) %>%
  arrange(number) %>%
  select(-c(number)) %>%
  kbl(.,
    longtable = T, booktabs = T, format = "latex",
    caption = caption, escape = FALSE
  ) %>%
  column_spec(column = c(1:10), width = "2cm")  %>%
  kable_styling(latex_options = "striped")
```

***Table 3:*** *Results of a Bayesian piece-wise regression predicting resting metabolism (mL $O_2$/min) of adult Japanese quail across ambient temperature (°C). Break-points (here, lower critical temperature), thermal resistance, and metabolism at thermoneutrality was allowed to vary by individual. Estimates represent posterior medians and credible intervals represent quantile intervals.*

| Response | Parameter | N | Estimate | 50% CI | 95% CI | BF |
|---|---|---|---|---|---|---|
| Lower Critical Temperature | Intercept | 34 | 24.6642 | (24.0768, 25.2474) | (22.8899, 26.4075) | Inf |
| Lower Critical Temperature | Body Mass (g) | 34 | -0.0010 | (-0.0018, -3e-04) | (-0.0033, 0.0012) | 4.6148 |
| Lower Critical Temperature | Tarsus Length (mm) | 34 | 0.0137 | (-0.0036, 0.0313) | (-0.038, 0.0665) | 2.3490 |

```
showtext_auto()

lctPlot <- ggplot(lctModel$data, aes(x = mass, y = LCT)) +
  geom_ribbon(data = expand.grid("mass" = seq(min(lctModel$data$mass, na.rm = T),
                                            max(lctModel$data$mass, na.rm = T),
                                            by = 1
                                            ),
                                "tarsus" = mean(lctModel$data$tarsus)) %>%
              mutate("LCT" = predict(lctModel, newdata = ., robust = TRUE)[, "Estimate"],
                     "SE" = predict(lctModel, newdata = ., robust = TRUE)[, "Est.Error"]) %>%
              mutate("LCL" = LCT - SE,
                     "UCL" = LCT + SE),
            aes(x = mass, ymin = LCL, ymax = UCL),
```

```
            fill = "#DECCC1", alpha = 0.4) +
  geom_point(pch = 21, size = 2, colour = "black", fill = "#DECCC1", alpha = 0.8) +
  geom_line(data = expand.grid("mass" = seq(min(lctModel$data$mass, na.rm = T),
                                             max(lctModel$data$mass, na.rm = T),
                                             by = 1
                                             ),
                                "tarsus" = mean(lctModel$data$tarsus)) %>%
              mutate("LCT" = predict(lctModel, newdata = ., robust = TRUE)[, "Estimate"],
                     "LCL" = predict(lctModel, newdata = ., robust = TRUE)[, "Q2.5"],
                     "UCL" = predict(lctModel, newdata = ., robust = TRUE)[, "Q97.5"]),
            aes(x = mass, y = LCT),
            colour = "black", linetype = "dashed") +
  xlab("Body Mass (g)") +
  ylab("Lower Critical Temperature (°C)") +
  ylim(c(24, 26)) +
  theme_classic() +
  theme(
    # axis.text = element_text(family = "Noto Sans", size = 16),
    # axis.title = element_text(family = "Noto Sans", size = 16)
    axis.text = element_text(family = "Noto Sans"),
    axis.title = element_text(family = "Noto Sans")
  )

lctPlot
```

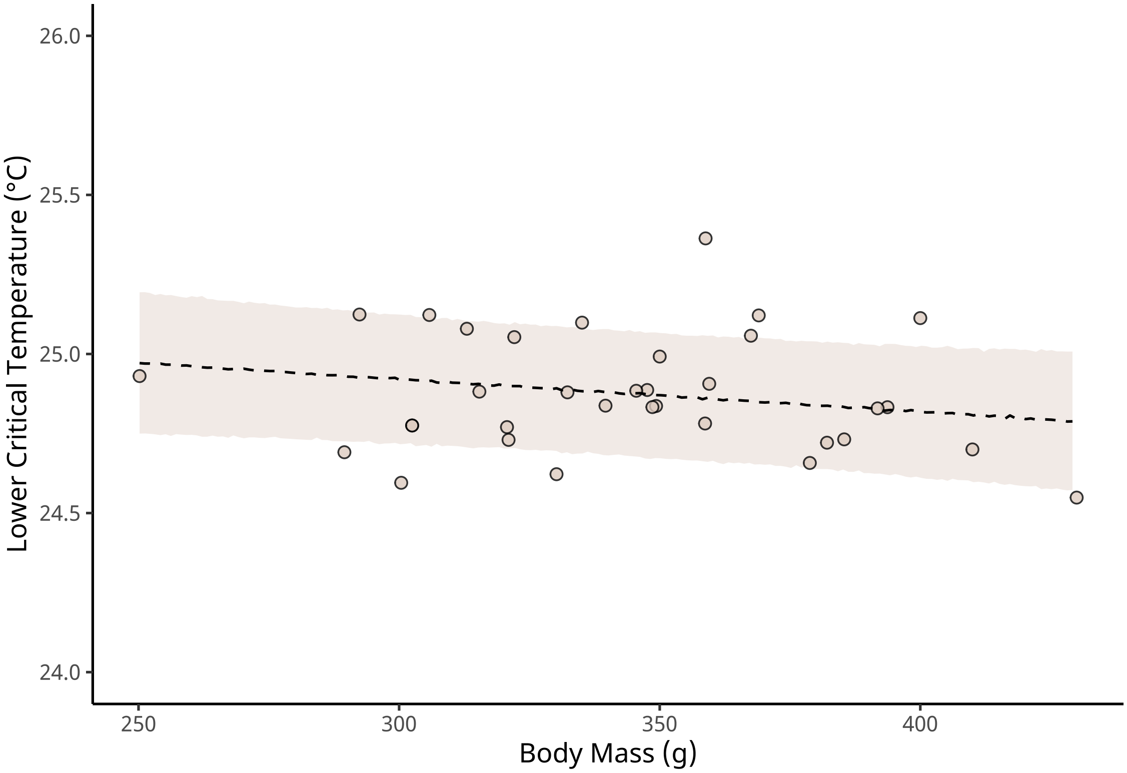


***Figure 7:*** *Lower critical temperature (°C) of adult Japanese quail as a function of body mass (g). Lower critical temperature is estimate from a Bayesian mixed effects break-point regression with resting metabolism (ml $O_2$/min) as the response variable and ambient temperature (°C) as the sole population-level predictor. Dots represent individual values and the dashed line represents the predicted line of best fit, as estimated from a Bayesian linear model. The ribbon indicates +/- one standard error around the line of best fit.*

```
ggsave("../plots/lctPlot.pdf", dpi = 800,
       width = 7, height = 6.5,
       lctPlot)
```

```r
showtext_auto(enable = FALSE)
```
