## Supplementary material for "No evidence that shrinking and shapeshifting meaningfully affect how birds respond to warming and cooling": Supp. 6

### Body Temperature Analyses

#### Body Temperature Responses to Cold Exposures

Contrasting long-standing assumptions, we found that neither body size (proxied by body mass) nor tarsus length influenced the rate at which adult Japanese quail increased there metabolism across a cold challenge (from 30°C to 10°C; Tabh et al 2024). This finding suggests that neither variable delectably affected the rates at which body heat was lost to the environment, and thus, the rates at which heat compensatory heat production was demanded. However, it is also possible that body size and tarsus length did indeed influence rates of body heat loss, and those with highest rates of heat loss suffered comparative declines in body temperature owing to insufficient rates of compensatory heat production. To evaluate this possibility, we tested whether body mass or tarsus length influenced the degree to which core body temperature of our quail changed between thermoneutral conditions (30°C) and our lowest temperature exposure (10°C).

To measure core body temperatures of adult quail throughout cold exposures, we used remotely-monitored temperature sensitive passive integrated transponder (PIT) tags (LifeChip BioTherm, Destron Fearing, South St. Paul, MN, USA; 2.1 × 12 mm; < 0.5% of body mass) implanted in the peritoneum. Tags were calibrated and implanted between 13 and 16 days of age following methods described in Persson et al (2024). During cold exposures, temperatures from tags were read passively but at a frequency of approximately 5 reads/min. For our analyses, we quantified cold-induced changes in body temperature as the difference in mean body temperature (°C) displayed at 10°C and 30°C, per individual.

To analyses whether and how body mass and tarsus length influenced body temperature responses to the cold, we used a Bayesian path analysis similar to those described in our main text (Tabh et al, 2024). Using a path analysis allowed us to control for, and measures, direct and indirect effects of rearing conditions (here, 10°C, 20°C, or 30°C) among quail on morphology and body temperature responses. Here, our path analysis was composed of three models with uncorrelated residuals as follows:

$$Body\ Mass_j \sim \beta_{a0} + \beta_{a1} \cdot Cold\ Rearing_j + \beta_{a2} \cdot Warm\ Rearing_j + \mu_{0a} + \epsilon_a$$

$$Tarsus\ Length_j \sim \beta_{b0} + \beta_{b1} \cdot Cold\ Rearing_j + \beta_{b2} \cdot Warm\ Rearing_j + \beta_{b3} \cdot Body\ Mass_j + \mu_{0b} + \epsilon_b$$

and:

$$Delta\ Tb_j \sim \beta_{c0} + \beta_{c1} \cdot Cold\ Rearing_j + \beta_{c2} \cdot Warm\ Rearing_j + \beta_{c3} \cdot Body\ Mass_j + \beta_{c4} \cdot Tarsus\ Length_j + \mu_{0c} + \epsilon_c$$

where $j$ represents individual (or, observation) identity, "warm rearing" and "cold rearing" represent binomial terms with "0" implying "false" and "1" implying"true", $\mu_{0a}$ -$\mu_{0c}$ representing group-level intercepts, per response variable, of egg batch, *Delta Tb* indicating the change in mean body temperature observed at 10°C and 30°C, and body mass and tarsus length measures being mean-centred to simplify model intercepts ($\beta_0$ terms). Error terms ($\epsilon$) were normally distributed with a mean of zero.

Priors for the first two models in our path analysis, including their derivations, are described in the main text and in supplemental files 2 and 3. Briefly, these priors were as follows:

$$\beta_{a0} \sim \mathcal{N}(0, 10)$$
$$\beta_{a1} \sim \mathcal{N}(0, 25)$$
$$\beta_{a2} \sim \mathcal{N}(0, 25)$$
$$\mu0a \sim \exp(2.5)$$
$$\epsilon_a \sim \exp(0.15)$$

$$\beta_{b0} \sim \mathcal{N}(0, 2.5)$$
$$\beta_{b1} \sim \mathcal{N}(0, 2.5)$$
$$\beta_{b2} \sim \mathcal{N}(0, 2.5)$$
$$\beta_{b3} \sim S\mathcal{N}(0, 0.25, 5)$$
$$\mu0b \sim \exp(2)$$
$$\epsilon_b \sim \exp(1)$$

For the third model in our path analysis, priors were weak and moderately informed by Persson et al (2024). For the effects of body mass and tarsus length on body temperature responses, we assumed that effects larger than the range of body temperature responses divided by the range of the given morphological variable were unlikely (here, standard deviation of a normally-distributed prior $= 0.5 \cdot \frac{Range[Delta\ Tb]}{Range[Mass\ or\ Tarsus]}$). Priors for this model were therefore as follows:

$$\beta_{c0} \sim \mathcal{N}(1, 1)$$
$$\beta_{c1} \sim \mathcal{N}(0, 1)$$
$$\beta_{c2} \sim \mathcal{N}(0, 1)$$
$$\beta_{c3} \sim \mathcal{N}(0, 0.01)$$
$$\beta_{c4} \sim \mathcal{N}(0, 0.1)$$
$$\mu0c \sim \exp(2.5)$$
$$\epsilon_c \sim \exp(2.5)$$

Below, we begin by importing all packages and functions required for this analysis. We then set our working

directly to simplify import and collation of our data.

```
def.chunk.hook <- knitr::knit_hooks$get("chunk")
knitr::knit_hooks$set(chunk = function(x, options) {
  x <- def.chunk.hook(x, options)
  ifelse(options$size != "normalsize",
         paste0("\n \\", options$size, "\n\n", x, "\n\n \\normalsize"), x)
})

knitr::opts_chunk$set(fig.pos = "H", out.extra = "")
```

```r
knitr::opts_chunk$set(size = "footnotesize")

library("tidyverse")
library("easypackages")

packageList <- c("bayesplot", "brms", "doParallel",
                 "foreach", "ggpubr", "kableExtra",
                 "latex2exp", "patchwork", "priorsense",
                 "showtext", "tidybayes", "wesanderson")

libraries(packageList)
library("brmsMethods")

# Printing package version numbers

caption <- paste0("R packages and their respective versions used for",
                  " data organisation and analysis in this study."
)

sapply(packageList, function(x) {
  y <- as.character(packageVersion(x))
  return(y)
}, simplify = FALSE) %>%
  enframe(., name = "Package", value = "Version") %>%
  as.data.frame(.) %>%
  kbl(.,
    longtable = T, booktabs = T,
    caption = caption
  ) %>%
  kable_styling(latex_options = "striped")
```

***Table 1:*** *R packages and their respective versions used for data organisation and analysis in this study.*

| Package | Version |
|---|---|
| bayesplot | 1.11.1 |
| brms | 2.20.4 |
| doParallel | 1.0.17 |
| foreach | 1.5.2 |
| ggpubr | 0.6.0 |
| kableExtra | 1.3.4 |
| latex2exp | 0.9.6 |
| patchwork | 1.2.0 |
| priorsense | 0.0.0.9000 |
| showtext | 0.9.6 |
| tidybayes | 3.0.4 |
| wesanderson | 0.3.6.9000 |

```r
# Adding custom functions

## A function to calculate the mode of a vector
```

```
md <- function(x) {
  all_values <- unique(x)
  all_values[which.max(tabulate(match(x, all_values)))]
}

# A function to cleanly view autocorrelation between posterior
# draws of specified coefficients/variables

clean_ac <- function(x, prs = NA, names = NA) {
  require("rstan")

  if (class(x)[1] != "brmsfit") {
    return("x must be a brmsfit object.")
  }

  if (is.na(prs[1])) {
    return(stan_ac(x$fit))
  }

  if (!is.na(prs[1]) & is.na(names[1])) {
    return(stan_ac(x$fit, pars = prs))
  }

  if (length(prs) != length(names)) {
    return("Length of pars and names must be equal.")
  }

  Base_plot <- stan_ac(x$fit, pars = prs, fill = nice_pink)
  Base_plot$data$parameters <- as.character(Base_plot$data$parameters)

  for (i in 1:length(prs)) {
    Base_plot$data$parameters[c(which(Base_plot$data$parameters == prs[i]))] <-
      names[i]
  }
  Base_plot$data$parameters <- as.factor(Base_plot$data$parameters)

  return(Base_plot)
}

# A function to simplify the output of bayestestR's hdi function.

simple_hdi <- function(x, rnd = 3, cis = c(50, 95), sci_note = FALSE) {
  if (class(x)[1] != "brmsfit") {
    return("x must be a brmsfit object.")
  }
  if (length(cis) != 2) {
    return("cis must be a vector of integers with length 2")
  }

  HDI_low <- bayestestR::hdi(x, effects = "all", ci = min(cis) / 100)
  HDI_high <- bayestestR::hdi(x, effects = "all", ci = max(cis) / 100)

  if (sci_note == FALSE) {
```

```
    Results <- data.frame(
      "Parameter" = HDI_low$Parameter,
      "1" = round(HDI_low$CI_low, rnd),
      "2" = round(HDI_low$CI_high, rnd),
      "3" = round(HDI_high$CI_low, rnd),
      "4" = round(HDI_high$CI_high, rnd)
    )
    colnames(Results)[c(2:5)] <-
      c(paste0("Low_HDI_", min(cis)), paste0("High_HDI_", min(cis)),
        paste0("Low_HDI_", max(cis)), paste0("High_HDI_", max(cis)))
  } else if (sci_note == TRUE) {
    Results <- data.frame(
      "Parameter" = HDI_low$Parameter,
      "1" = format(round(HDI_low$CI_low, rnd), scientific = TRUE),
      "2" = format(round(HDI_low$CI_high, rnd), scientific = TRUE),
      "3" = format(round(HDI_high$CI_low, rnd), scientific = TRUE),
      "4" = format(round(HDI_high$CI_high, rnd), scientific = TRUE)
    )
    colnames(Results)[c(2:5)] <- c(paste0("Low_HDI_", min(cis)),
                                   paste0("High_HDI_", min(cis)),
                                   paste0("Low_HDI_", max(cis)),
                                   paste0("High_HDI_", max(cis)))
  }

  return(Results)
}

modeHDI <- function(x, cis = c(50, 95), rnd = 4, collapse = FALSE){
  stopifnot("x must be a 'brmsfit' object" = class(x) == "brmsfit",
            "collapse must be logical TRUE/FALSE" = is.logical(collapse))

  out <- lapply(X = as.data.frame(x), MARGIN = 2, FUN = ggdist::mode_hdi,
                .width = c(cis/100))
  hold <- names(out)
  out <- out %>%
    map2(hold, ~mutate(.x, name = .y)) %>%
    bind_rows() %>%
    mutate(y = round(y, digits = rnd),
           ymin = round(ymin, digits = rnd),
           ymax = round(ymax, digits = rnd)) %>%
    select("par" = name, "mode" = y, "lcl" = ymin,
           "ucl"= ymax, "confidenceLevel" = .width)

  if (collapse == FALSE){
    return(out)
  } else if (collapse == TRUE){
    out <- out %>%
      mutate("cis" = paste0("[", lcl, ", ", ucl, "]")) %>%
      select(-c(lcl, ucl))

    return(out)
  }
}
```

```r
# A function to calculate quantile intervals from a brmsfit object

quantileCIs <- function(x, rnd = 3, cis = c(50, 95), sci_note = FALSE) {
  require(tidyverse)

  if (class(x)[1] != "brmsfit") {
    return("x must be a brmsfit object.")
  }
  if (length(cis) != 2) {
    return("cis must be a vector of integers with length 2")
  }

  prbs = c()
  nColNames = c()
  for (i in 1:length(cis)){
    prbs = c(prbs, c(0.5 - (cis[i]/100)/2, 0.5 + (cis[i]/100)/2))
    nColNames = c(nColNames,
                  paste0("Low_CI_", cis[i]),
                  paste0("High_CI_", cis[i])
    )
  }

  modelFrame = as.data.frame(x)

  Results <- apply(modelFrame, MARGIN = 2, FUN = quantile,
        probs = prbs, type = 8) %>%
    t() %>%
    as.data.frame() %>%
    rownames_to_column(var = "par") %>%
    `colnames<-`(c("Parameter", nColNames))

  if (sci_note == FALSE) {
    Results <- apply(modelFrame, MARGIN = 2, FUN = quantile,
        probs = prbs, type = 8) %>%
    t() %>%
    as.data.frame() %>%
    rownames_to_column(var = "par") %>%
    `colnames<-`(c("Parameter", nColNames))

  } else if (sci_note == TRUE) {
    Results <- apply(modelFrame, MARGIN = 2, FUN = quantile,
        probs = prbs, type = 8) %>%
    t() %>%
    as.data.frame() %>%
    rownames_to_column(var = "par") %>%
    `colnames<-`(c("Parameter", nColNames)) %>%
    mutate_at(.vars = vars(-Parameter),
              .funs = function(x){
                return(format(x, scientific = TRUE))
              }
    )
  }
```

```r
  return(Results)
}

# A function that allows users to assign multiple objects
# to different variable names at once. The below is reported by "ellbur" at
# https://strugglingthroughproblems.wordpress.com/author/ellbur/page/3/.

{
  "%=%" <- function(l, r, ...) UseMethod("%=%")

  "%=%.lbunch" <- function(l, r, ..., List = NA) {
    Envir <- as.environment(-1)

    if (!is.na(List)) {
      l <- List[[1]]
      r <- List[[2]]
    }

    if (length(r) > length(l)) {
      warning("RHS has more args than LHS. Only first",
              length(l), "used.")
    }

    if (length(l) > length(r)) {
      warning("LHS has more args than RHS. RHS will be repeated.")
      r <- extendToMatch(r, l)
    }

    for (II in 1:length(l)) {
      do.call("<-", list(l[[II]], r[[II]]), envir = Envir)
    }
  }

  extendToMatch <- function(source, destin) {
    s <- length(source)
    d <- length(destin)

    if (d == 1 && s > 1 && !is.null(as.numeric(destin))) {
      d <- destin
    }

    dif <- d - s
    if (dif > 0) {
      source <- rep(source, ceiling(d / s))[1:d]
    }
    return(source)
  }

  g <- function(...) {
    List <- as.list(substitute(list(...)))[-1L]
    class(List) <- "lbunch"
    return(List)
  }
```

```r
}

# A function to simplify output of bayes_R2 from brms

simpleR2 <- function(x, ndraws = 1000, roundDigits = 5,
                     robust = TRUE) {
  stopifnot("x must be a 'brmsfit' object" = class(x) == "brmsfit",
            "robust must be logical (TRUE/FALSE)" = is.logical(robust))
  grab <- brms::bayes_R2(x, ndraws = ndraws, robust = robust)
  toPrint <- paste0(
    "R2 = ", round(grab[, "Estimate"], digits = roundDigits),
    " [",
    round(grab[, "Q2.5"], digits = roundDigits),
    ",",
    round(grab[, "Q97.5"], digits = roundDigits),
    "]"
  )
  cat(toPrint)
}

# A function to calculate the position of a skew-normal
# distribution given its mean, omega, and alpha values

skewxi <- function(mean, omega, alpha) {
  delta <- alpha / (sqrt(1 + alpha^2))
  xi <- mean - omega * delta * sqrt(2 / pi)
  return(xi)
}

# A function to produce clean posterior or prior
# predictive checks, based upon "pp_check" from the R-package 'brms'.

pp_check2 <- function(model, resp = NA, ndraws = 500,
                      xlab = "label", colour = "lightblue") {
  require(brms)
  require(ggplot2)
  stopifnot("Model must be a brmsfit object" = is.brmsfit(model))

  if (is.na(resp)) {
    resp <- model$formula$resp
  }

  p1 <- brms::pp_check(model, ndraws = ndraws, resp = resp) +
    scale_colour_manual(
      values = c("black", colour),
      labels = c("y", "yhat"),
      name = NULL
    ) +
    xlab(xlab) +
    ylab("Density") +
    theme_classic()
  return(p1)
}
```

```r
# A function to summarise and print Gelman-Rubin statistics and effective
# sample sizes to sample sizes for brmsfit objects.

neffBase <- function(x){
  stopifnot("Model must be a brmsfit object" = is.brmsfit(x))
  out <- as.data.frame(brms::neff_ratio(x)) %>%
    rownames_to_column(var = "var") %>%
    filter(!(var %in% c("lprior", "lp__"))) %>%
    pull(.)
  return(out)
}

chainCheck <- function(model, rDig = 3) {
  require(brms)
  stopifnot("Model must be a brmsfit object" = is.brmsfit(model))

  Rhat <- paste0(
    "Rhat range: ",
    round(min(rhat(model)), digits = rDig),
    " - ",
    round(max(rhat(model)), digits = rDig)
  )
  Neff <- paste0(
    "Neff/N range: ",
    round(min(neffBase(model)), digits = rDig),
    " - ",
    round(max(neffBase(model)), digits = rDig)
  )
  cat(paste0(Rhat, "\n", Neff))
}

# Setting working directory.

setwd(
  paste0("/Users/joshuatabh/Documents/researchProjects/",
  "lund/functionOfAllen/functionData"
  )
)
```

Next, we import and organise our data pertaining to body temperature measurements. Raw body temperature means are then plotted to check for erroneous measurements usine a Cleveland dot plot.

```r
# Loading and binding data as described in Supplement 3
# "Metabolic Slope and Repeatability Analyses"

{
  all <- merge(
    read.csv("compiledData.csv") %>%
      select(-Ta),
    bind_rows(
      read.csv("exp1VO2.csv"),
      read.csv("exp2VO2.csv"),
      read.csv("exp3VO2.csv")
    ) %>%
      select(-c(startTime, endTime, initialTb, fileName)) %>%
      mutate(pretreatment = ifelse(pretreatment == "control",
```

```
        "neutral", pretreatment
      )) %>%
      distinct() %>%
      select(-c(pretreatment, posttreatment, birdID)),
    by = c("ring", "week", "exp"),
    all.x = TRUE
  ) %>%
    distinct() %>%
    select(
      ring, birdID, sex, exp, week, pretreatment,
      posttreatment, treatment, Ta, mass, wingLength,
      meanTb, tarsusLengthMean, tarsusLengthSD,
      tarsusHeightMean, tarsusHeightSD,
      tarsusCalibration, VO2, RMR
    ) %>%
    mutate(Ta = ifelse(exp == "C" & Ta > 28 & Ta < 31, 30, Ta)) %>%
    mutate(Ta = ifelse(exp == "C" & Ta > 38 & Ta < 41, 40, Ta)) %>%
    filter(Ta %in% c(10, 20, 30, 40))

  all <- rbind(
    all %>%
      filter(ring == "B14" & week == "3") %>%
      group_by(Ta) %>%
      mutate(
        "tarsusLengthMean" = mean(tarsusLengthMean, na.rm = T),
        "tarsusHeightMean" = mean(tarsusHeightMean, na.rm = T)
      ) %>%
      mutate("n" = 1:2) %>%
      ungroup() %>%
      arrange(n, Ta) %>%
      slice(1:4) %>%
      select(-n),
    all %>%
      filter(!(ring == "B14" & week == "3"))
  ) %>%
    arrange(exp, week, ring, Ta)

  # Splitting out experiment 3 data and adding in body temperature data

  all <- all %>%
    filter(exp == "C") %>%
    select(-meanTb) %>%
    merge(.,
      read.csv("exp3Tb.csv") %>%
        select(-X) %>%
        mutate(Ta = ifelse(Ta > 28 & Ta < 31, 30, Ta)) %>%
        mutate(
          Ta = ifelse(Ta > 38 & Ta < 41, 40, Ta),
          week = ifelse(week == 4, "3",
            ifelse(week == 9, "8",
              "12"
            )
          )
        ) %>%
        filter(Ta %in% c(30, 40) & week %in% c(3, 8)) %>%
        select(ring, week, Ta, meanTb),
      by = c("ring", "week", "Ta"),
      all.x = TRUE
    ) %>%
    bind_rows(
      .,
      all %>%
        filter(exp != "C")
    ) %>%
    arrange(exp, week, ring) %>%
    filter(week == 8)
}
```

```r
# Plotting body temperature with Cleveland dotplot to check for erroneous
# measurements

lims <- all %>%
  filter(!is.na(meanTb)) %>%
  mutate(
    "Ta" = paste0(Ta, "°C")
  ) %>%
  group_by(Ta) %>%
  summarise(
    "LCL" = mean(meanTb, na.rm = T) -
      3.5 * sd(meanTb, na.rm = T),
    "UCL" = mean(meanTb, na.rm = T) +
      3.5 * sd(meanTb, na.rm = T),
    meanTb = mean(meanTb, na.rm = T)
  )

all %>%
  filter(!is.na(meanTb)) %>%
  mutate(
    "Ta" = paste0(Ta, "°C")
  ) %>%
  ggplot(aes(x = 1:nrow(.), y = meanTb)) +
  facet_wrap(~ Ta) +
  geom_rect(
    data = lims,
    colour = "black", fill = "grey80", alpha = 0.5,
    aes(
      group = Ta,
      xmin = -Inf, xmax = Inf,
      ymin = LCL, ymax = UCL),
      inherit.aes = FALSE
  ) +
  geom_point(size = 2, colour = "black", alpha = 0.7) +
  theme_classic() +
  xlab("Sample Number") +
  ylab("Mean Body Temperature (°C)")
```

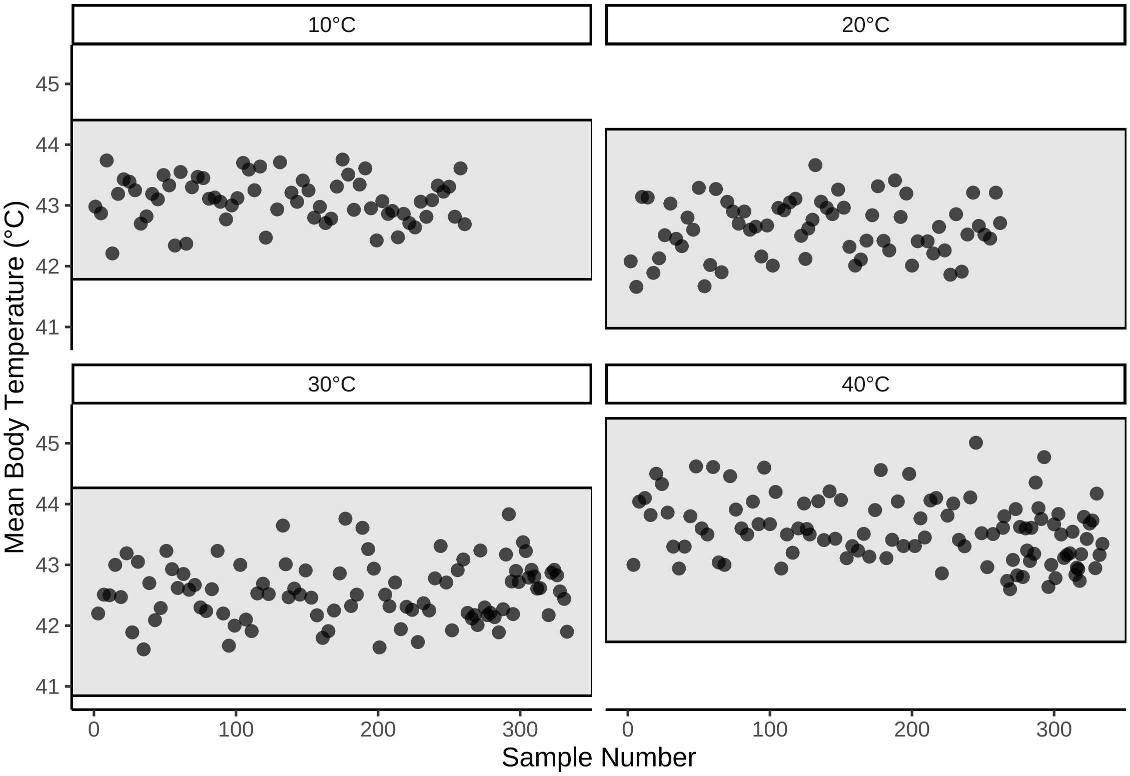


***Figure 1:*** *Cleveland dotplot of mean body temperature values (°C) by ambient temperature of collection, each drawn from adult Japanese quail. Dots represent raw data points. Rectangles indicate mean body temperatures +/- 3.5 times the standard deviation at a given ambient temperature.*

No extreme values are evident from our dotplot. Next, we calculate body temperature responses from 30°C to 10°C and visualise these values for oddities.

```
tbResponse <- rbind(
  all %>%
    filter(Ta %in% c(10, 30)) %>%
    rename(
      "batch" = exp,
      "tarsus" = tarsusLengthMean
    ) %>%
    pivot_wider(
      id_cols = c(
        "ring", "batch", "week", "pretreatment",
        "mass", "tarsus"
      ), names_from = "Ta",
      values_from = "meanTb"
    ) %>%
    mutate(deltaTb = `10` - `30`) %>%
    select(-c(`10`, `30`)) %>%
    mutate("challenge" = "cold"),

  all %>%
    filter(Ta %in% c(30, 40)) %>%
    rename(
      "batch" = exp,
      "tarsus" = tarsusLengthMean
    ) %>%
    pivot_wider(
      id_cols = c(
        "ring", "batch", "week", "pretreatment",
```

```
        "mass", "tarsus"
      ), names_from = "Ta",
      values_from = "meanTb"
    ) %>%
    mutate(deltaTb = `40` - `30`) %>%
    select(-c(`30`, `40`)) %>%
    mutate("challenge" = "warm")
)

tbResponse %>%
  filter(!is.na(deltaTb) & challenge == "cold") %>%
  ggplot(aes(x = 1:nrow(.), y = deltaTb)) +
  geom_rect(aes(xmin = -Inf, xmax = Inf,
                ymin = mean(deltaTb) - 3.5*sd(deltaTb),
                ymax = mean(deltaTb) + 3.5*sd(deltaTb)
                ),
            fill = "grey80", alpha = 0.5, colour = "black") +
  geom_hline(yintercept = 0, linetype = "dashed", colour = "black") +
  geom_point(size = 2, colour = "black", alpha = 0.7) +
  scale_y_continuous(sec.axis = sec_axis( trans=~., name="Second Axis",
                                          breaks = c(1.5, -0.5),
                                          labels = c("Increasing Tb",
                                                     "Decreasing Tb"))
                     ) +
  theme_classic() +
  theme(axis.text.y.right = element_text(angle = 270),
        axis.ticks.y.right = element_blank(),
        axis.title.y.right = element_blank()) +
  xlab("Sample Number") +
  ylab("Change in\nBody Temperature (°C)")
```

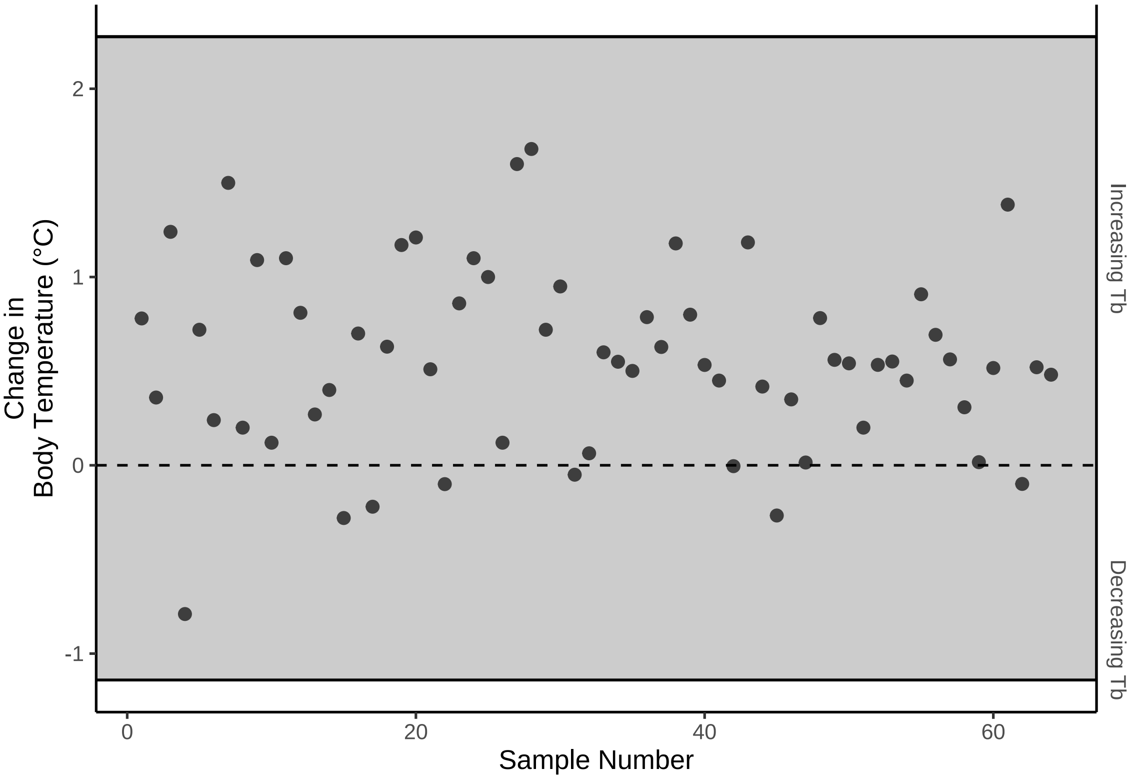


***Figure 2:*** *Cleveland dotplot of mean changes in body temperature (°C) between 10°C and 30°C, each drawn from adult Japanese quail. Again, dots represent raw data points and rectangles indicate means +/- 3.5 times the standard deviation at a given ambient temperature. The dashed line indicates no change in body temperature.*

Raw effects of morphology and rearing conditions on body temperature responses to cold are visualised to build information for model adjustments, if needed.

```
p1 <- tbResponse %>%
  filter(!is.na(deltaTb) & challenge == "cold") %>%
  ggplot(aes(x = mass, y = deltaTb)) +
  geom_point(size = 2, colour = "black", alpha = 0.7) +
  geom_smooth(
    method = "lm", se = FALSE, linetype = "dashed",
    colour = "black"
  ) +
  geom_hline(yintercept = 0, linetype = "dashed", colour = "firebrick4") +
  theme_classic() +
  xlab("Body Mass (g)") +
  ylab("Change in\nBody Temperature (°C)")

p2 <- tbResponse %>%
  filter(!is.na(deltaTb) & challenge == "cold") %>%
  ggplot(aes(x = tarsus, y = deltaTb)) +
  geom_point(size = 2, colour = "black", alpha = 0.7) +
  geom_smooth(
    method = "lm", se = FALSE, linetype = "dashed",
    colour = "black"
  ) +
  geom_hline(yintercept = 0, linetype = "dashed", colour = "firebrick4") +
  theme_classic() +
  xlab("Tarsus Length (mm)") +
  ylab("Change in\nBody Temperature (°C)")
```

```r
p3 <- tbResponse %>%
  filter(!is.na(deltaTb) & challenge == "cold") %>%
  mutate(pretreatment = ifelse(pretreatment == "cold",
    "Cold (10°C)",
    ifelse(pretreatment == "neutral",
      "Mild (20°C)", "Warm (30°C)"
    )
  )) %>%
  ggplot(aes(x = pretreatment, y = deltaTb, fill = pretreatment)) +
  geom_point(
    size = 1, colour = "black", alpha = 0.5,
    position = position_jitter(width = 0.25),
    pch = 21
  ) +
  stat_summary(
    geom = "errorbar", fun.data = "mean_se",
    colour = "black", width = 0.25,
    position = position_dodge(width = 0.35)
  ) +
  stat_summary(
    geom = "point", fun = "mean", pch = 21,
    colour = "black", size = 3,
    position = position_dodge(width = 0.35)
  ) +
  geom_hline(yintercept = 0, linetype = "dashed", colour = "firebrick4") +
  scale_fill_manual(values = c("#7BB4E3", "black", "#CD5C5C")) +
  theme_classic() +
  theme(
    axis.title.x = element_blank(),
    legend.position = "none"
  ) +
  ylab("Change in\nBody Temperature (°C)")

(p1 + p2) / p3 + plot_annotation(tag_levels = "A")
```

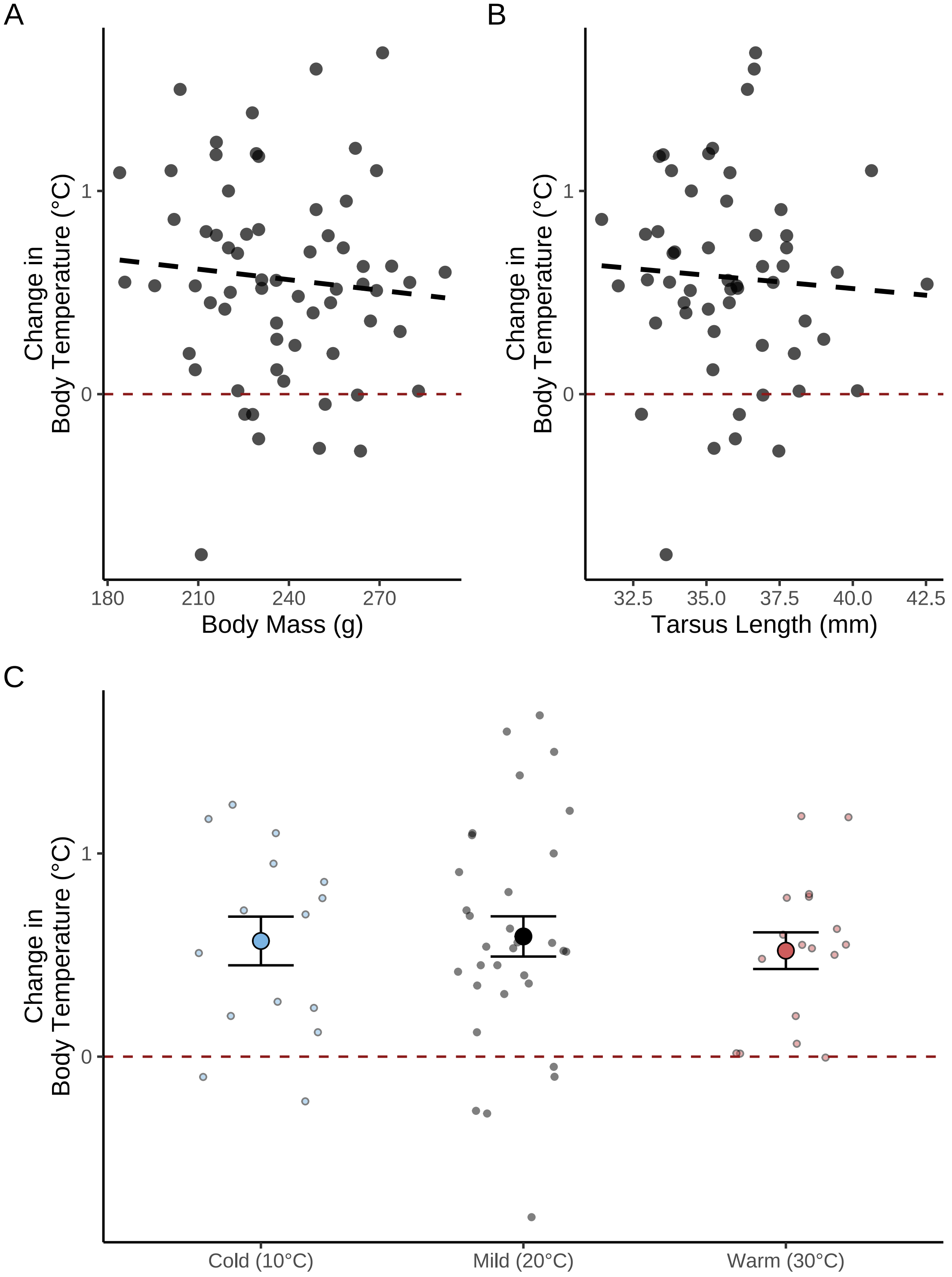


***Figure 3:*** *Effects of body mass (g), tarsus length (mm), and rearing treatment on changes in body temperature (°C) in responses to a cold exposure (10°C) in adult Japanese quail. Baseline body temperatures represent those measured at thermoneutrality (30°C). Small dots represent raw values per individual. Horizontal and dashed red lines indicate no change in body temperature. Black dashed lines represent raw lines of best fit estimated from the R package ggplot2 (Wickham, 2011). In panel C, large dots indicate means per rearing treatment and errorbars indicate standard errors.*

Last, we visually check whether variance in body temperature responses differ between rearing treatments or sources of individuals (here, egg batches).

```
# Checking whether variance in responses by treatment temperature

p1 <- tbResponse %>%
  filter(!is.na(deltaTb) & challenge == "cold") %>%
  ggplot(aes(x = deltaTb, fill = pretreatment)) +
  geom_density(colour = "black", alpha = 0.5) +
  scale_fill_manual(name = "Rearing Treatment",
                    labels = c("Cold (10°C)",
                               "Mild (20°C)",
                               "Warm (30°C)"),
                    values = c("#7BB4E3", "black", "#CD5C5C")
                    ) +
  theme_classic() +
  theme(axis.title.x = element_blank(),
        legend.position = "bottom") +
  ylab("Change in\nBody Temperature (°C)")

p2 <- tbResponse %>%
  filter(!is.na(deltaTb) & challenge == "cold") %>%
  ggplot(aes(x = deltaTb, fill = batch)) +
  geom_density(colour = "black", alpha = 0.5) +
  scale_fill_manual(name = "Egg\nBatch",
                    values = c("#7BB4E3", "black")
                    ) +
  theme_classic() +
  theme(axis.title.x = element_blank(),
        legend.position = "bottom") +
  ylab("Change in\nBody Temperature (°C)")

(p1 + p2) + plot_annotation(tag_levels = "A")
```

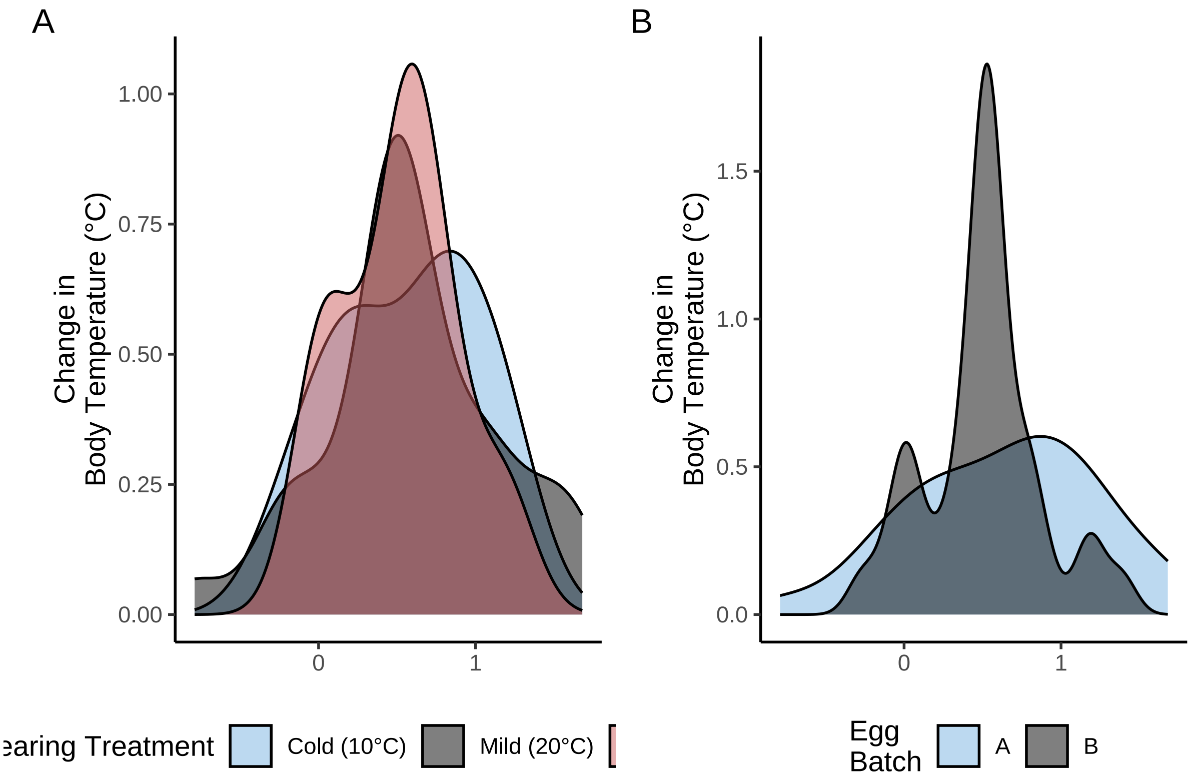


***Figure 4:*** *Density of body temperature responses (°C) to cold exposure (10°C) in adult Japanese quail from different rearing conditions or egg batches. Baseline body temperatures represent those measured at thermoneutrality (30°C).*

```
# No evidence of heteroskedasticity.
```

We now proceed to constructing our path analysis as described previously. To test the suitability of our model priors, we first check how reasonably predictions from our priors alone overlay with the true density of our body temperature response values.

```
tbResponseCold8WeeksPPCheck <-
  brm(
    data = tbResponse %>%
      filter(challenge == "cold") %>%
      mutate(
        mass = mass - mean(mass, na.rm = T),
        tarsus = tarsus - mean(tarsus, na.rm = T),
        pretreatment = ifelse(pretreatment == "cold", "A",
          ifelse(pretreatment == "neutral", "B", "C")
        )
      ) %>%
      mutate(pretreatment = factor(pretreatment,
        levels = c("B", "A", "C")
      )) %>%
      drop_na(),
    family = "gaussian",
    bf(mass ~ pretreatment + (1 | batch)) +
      bf(tarsus ~ mass + pretreatment + (1 | batch)) +
      bf(deltaTb ~ tarsus + mass + pretreatment + (1 | batch)) +
      set_rescor(FALSE),
    prior = c(
      set_prior("normal(0, 10)",
        class = "Intercept",
        resp = "mass"
```

```
    ),
    set_prior("normal(0, 25)",
      class = "b",
      coef = "pretreatmentA", resp = "mass"
    ),
    set_prior("normal(0, 25)",
      class = "b",
      coef = "pretreatmentC", resp = "mass"
    ),
    set_prior("exponential(2.5)",
      class = "sd",
      group = "batch", resp = "mass"
    ),
    set_prior("exponential(0.15)",
      class = "sigma",
      resp = "mass"
    ),
    set_prior("normal(0, 2.5)",
      class = "Intercept",
      resp = "tarsus"
    ),
    set_prior("normal(0, 2.5)",
      class = "b",
      coef = "pretreatmentA", resp = "tarsus"
    ),
    set_prior("normal(0, 2.5)",
      class = "b",
      coef = "pretreatmentC", resp = "tarsus"
    ),
    set_prior("skew_normal(0, 0.25, 5)",
      class = "b",
      coef = "mass",
      resp = "tarsus"
    ),
    set_prior("exponential(2)",
      class = "sd",
      group = "batch",
      resp = "tarsus"
    ),
    set_prior("exponential(1)",
      class = "sigma",
      resp = "tarsus"
    ),
    set_prior("normal(1, 1)",
      class = "Intercept",
      resp = "deltaTb"
    ),
    set_prior("normal(0, 1)",
      class = "b",
      coef = "pretreatmentA",
      resp = "deltaTb"
    ),
    set_prior("normal(0, 1)",
      class = "b",
      coef = "pretreatmentC",
      resp = "deltaTb"
    ),
    set_prior("normal(0, 0.01)",
      class = "b",
      coef = "mass",
      resp = "deltaTb"
    ),
    set_prior("normal(0, 0.1)",
      class = "b",
      coef = "tarsus",
      resp = "deltaTb"
    ),
```

```
      set_prior("exponential(2.5)",
        class = "sd",
        group = "batch",
        resp = "deltaTb"
      ),
      set_prior("exponential(2.5)",
        class = "sigma",
        resp = "deltaTb"
      )
    ),
    iter = 50000, warmup = 10000, cores = 4, chains = 4, thin = 20,
    control = list(adapt_delta = .97, max_treedepth = 14),
    silent = TRUE, refresh = 0,
    sample_prior = "only",
    file = "./models/tbResponseToColdPPCheck.Rds"
  )

pp_check2(tbResponseCold8WeeksPPCheck,
          xlab = "Change in\nBody Temperature (°C)",
          resp = "deltaTb"
          )
```

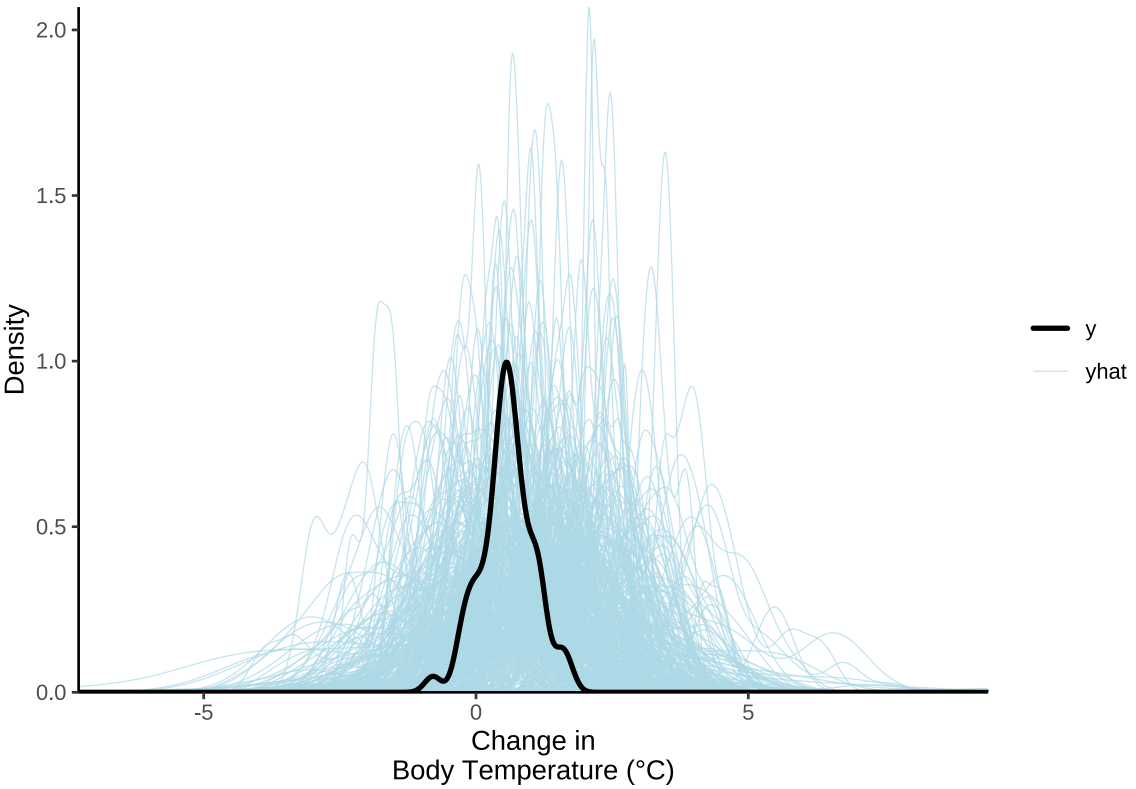


***Figure 5:*** *Overlay of predicted (blue) and true (black) body temperature response (°C) densities, where predicted densities are derived from priors in a Bayesian path analyses. Clear overlap betwen the black and blue lines suggests that model priors are reasonable with respect to the data.*

Our priors clearly capture the density of true body temperature responses well without obvious constraints. We therefore proceed to constructing and evaluating our full path analysis.

```
tbResponseCold8Weeks <-
  brm(
    data = tbResponse %>%
```

```
    filter(challenge == "cold") %>%
    mutate(
      mass = mass - mean(mass, na.rm = T),
      tarsus = tarsus - mean(tarsus, na.rm = T),
      pretreatment = ifelse(pretreatment == "cold", "A",
        ifelse(pretreatment == "neutral", "B", "C")
      )
    ) %>%
    mutate(pretreatment = factor(pretreatment,
      levels = c("B", "A", "C")
    )) %>%
    drop_na(),
  family = "gaussian",
  bf(mass ~ pretreatment + (1 | batch)) +
    bf(tarsus ~ mass + pretreatment + (1 | batch)) +
    bf(deltaTb ~ tarsus + mass + pretreatment + (1 | batch)) +
    set_rescor(FALSE),
  prior = c(
    set_prior("normal(0, 10)",
      class = "Intercept",
      resp = "mass"
    ),
    set_prior("normal(0, 25)",
      class = "b",
      coef = "pretreatmentA", resp = "mass"
    ),
    set_prior("normal(0, 25)",
      class = "b",
      coef = "pretreatmentC", resp = "mass"
    ),
    set_prior("exponential(2.5)",
      class = "sd",
      group = "batch", resp = "mass"
    ),
    set_prior("exponential(0.15)",
      class = "sigma",
      resp = "mass"
    ),
    set_prior("normal(0, 2.5)",
      class = "Intercept",
      resp = "tarsus"
    ),
    set_prior("normal(0, 2.5)",
      class = "b",
      coef = "pretreatmentA", resp = "tarsus"
    ),
    set_prior("normal(0, 2.5)",
      class = "b",
      coef = "pretreatmentC", resp = "tarsus"
    ),
    set_prior("skew_normal(0, 0.25, 5)",
      class = "b",
      coef = "mass",
      resp = "tarsus"
    ),
    set_prior("exponential(2)",
      class = "sd",
      group = "batch",
      resp = "tarsus"
    ),
    set_prior("exponential(1)",
      class = "sigma",
      resp = "tarsus"
    ),
    set_prior("normal(1, 1)",
      class = "Intercept",
      resp = "deltaTb"
```

```
      ),
      set_prior("normal(0, 1)",
        class = "b",
        coef = "pretreatmentA",
        resp = "deltaTb"
      ),
      set_prior("normal(0, 1)",
        class = "b",
        coef = "pretreatmentC",
        resp = "deltaTb"
      ),
      set_prior("normal(0, 0.01)",
        class = "b",
        coef = "mass",
        resp = "deltaTb"
      ),
      set_prior("normal(0, 0.1)",
        class = "b",
        coef = "tarsus",
        resp = "deltaTb"
      ),
      set_prior("exponential(2.5)",
        class = "sd",
        group = "batch",
        resp = "deltaTb"
      ),
      set_prior("exponential(2.5)",
        class = "sigma",
        resp = "deltaTb"
      )
    ),
    iter = 50000, warmup = 10000, cores = 4, chains = 4, thin = 20,
    control = list(adapt_delta = .97, max_treedepth = 14),
    silent = TRUE, refresh = 0,
    file = "./models/tbResponseToCold.Rds"
  )
```

A few (n = 17) divergent transitions were detected during our Hamiltonian Monte Carlo (HMC) chain sampling. These divergences are identified and visualised with respect to all samples to determine their cause.

```
grab <- paste0(grep("b_|sd_|sigma_",
                    get_variables(tbResponseCold8Weeks), value = TRUE),
  collapse = "|"
)

posteriorDraws <- as.array(tbResponseCold8Weeks)
divergences <- nuts_params(tbResponseCold8Weeks) %>%
  filter(Parameter == "divergent__" & Value == "1") %>%
  select(".chain" = Chain, ".iteration" = Iteration) %>%
  mutate("divergent" = "TRUE")

spread_draws(tbResponseCold8Weeks, !!sym(grab), regex = TRUE) %>%
  select(-c(.draw)) %>%
  pivot_longer(-c(".chain", ".iteration"),
               names_to = "par", values_to = "values") %>%
  mutate("chain" = paste0("Chain ", .chain)) %>%
  merge(.,
    tribble(
      ~par, ~Par,
      "b_mass_Intercept", " a0",
      "b_mass_pretreatmentA", " a1",
      "b_mass_pretreatmentC", " a2",
      "sd_batch__mass_Intercept", " 0a",
      "sigma_mass", " a",
      "b_tarsus_Intercept", " b0",
```

```
    "b_tarsus_pretreatmentA", " b1",
    "b_tarsus_pretreatmentC", " b2",
    "b_tarsus_mass", " b3",
    "sd_batch__tarsus_Intercept", " 0b",
    "sigma_tarsus", " b",
    "b_deltaTb_Intercept", " c0",
    "b_deltaTb_pretreatmentA", " c1",
    "b_deltaTb_pretreatmentC", " c2",
    "b_deltaTb_mass", " c3",
    "b_deltaTb_tarsus", " c4",
    "sd_batch__deltaTb_Intercept", " 0c",
    "sigma_deltaTb", " c"
  ),
  by = "par", all.x = TRUE
) %>%
merge(., divergences, by = c(".chain", ".iteration"), all.x = TRUE) %>%
mutate(divergent = ifelse(is.na(divergent), "FALSE", divergent)) %>%
ggplot(aes(x = .iteration, y = values,
           fill = divergent, alpha = divergent)) +
facet_grid(Par ~ chain, scales = "free") +
geom_point(pch = 21, colour = "black") +
scale_fill_manual(values = c("grey90", "darkred"),
                  name = "Divergent") +
scale_alpha_manual(values = c(0.2, 1), name = "Divergent") +
xlab("Iteration") +
ylab("Values") +
theme_classic() +
theme(
  axis.text.y = element_blank(),
  axis.ticks.y = element_blank()
)
```

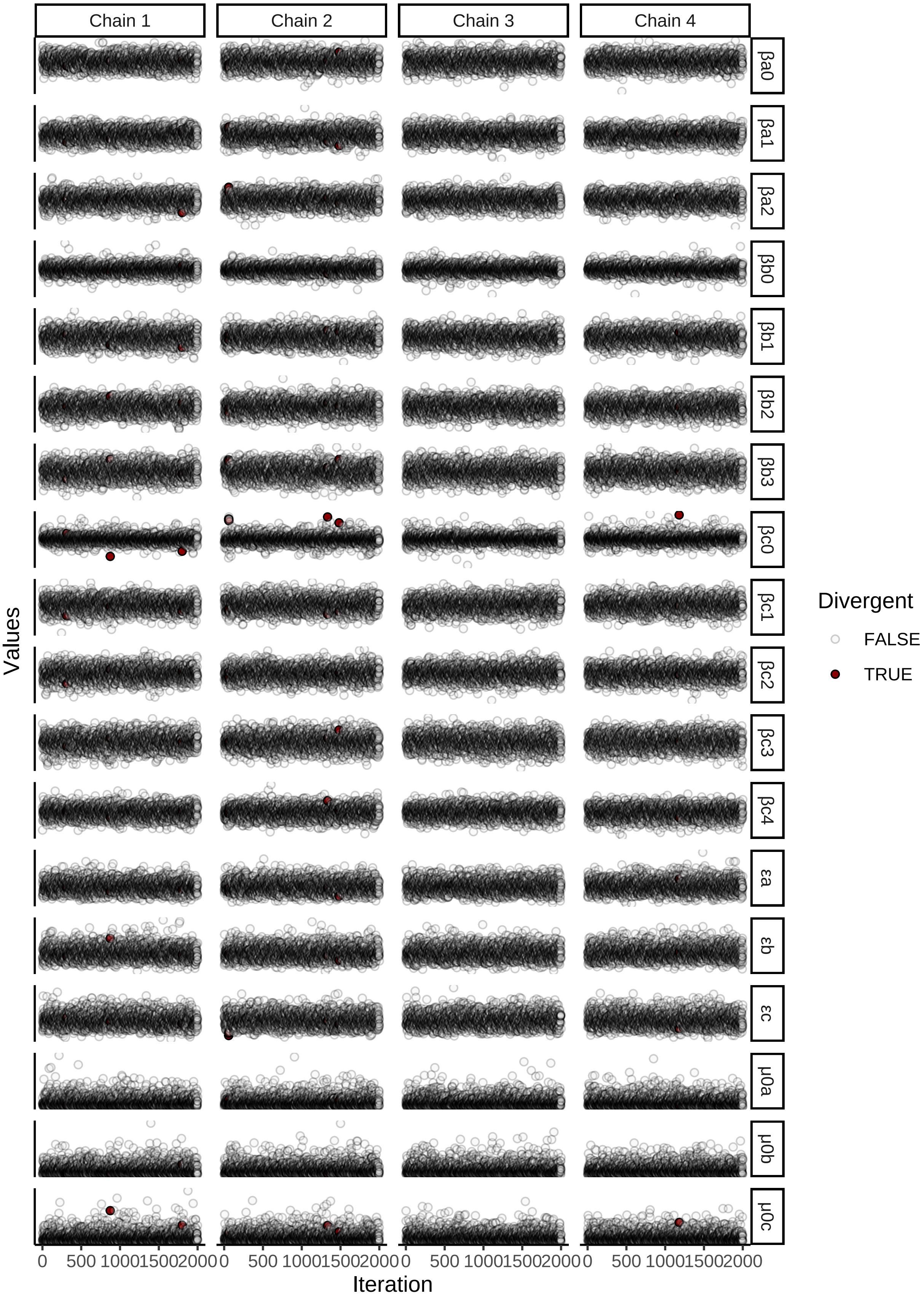


***Figure 6:*** *Posterior draws from Bayesian path analysis predicting body temperature responses (°C) to the cold (10°C) as a direct and indirect function of body mass (g) and tarsus length (mm) in adult old Japanese quail. Draws are seperated by model parameter and Hamiltonian Monte Carlo (HMC) chains.*

Divergent transitions appear to occur when the value for our intercept on body temperature responses is either large or small. We check whether these divergences lie on a numeric scale.

```
spread_draws(tbResponseCold8Weeks, !!sym(grab), regex = TRUE) %>%
  select(-c(.draw)) %>%
  pivot_longer(-c(".chain", ".iteration"),
               names_to = "par", values_to = "values") %>%
  mutate("chain" = paste0("Chain ", .chain)) %>%
  filter(par == "b_deltaTb_Intercept") %>%
  merge(., divergences, by = c(".chain", ".iteration"), all.x = TRUE) %>%
  mutate(divergent = ifelse(is.na(divergent), "FALSE", divergent)) %>%
  ggplot(aes(x = .iteration, y = values,
             fill = divergent, alpha = divergent)) +
  geom_point(pch = 21, colour = "black") +
  geom_hline(yintercept = -1, linetype = "dashed", colour = "firebrick4") +
  geom_hline(yintercept = 3, linetype = "dashed", colour = "firebrick4") +
  scale_fill_manual(values = c("grey90", "darkred"),
                    name = "Divergent") +
  scale_alpha_manual(values = c(0.2, 1), name = "Divergent") +
  xlab("Iteration") +
  ylab("Body Temperature Response (°C)\nIntercept") +
  theme_classic()
```

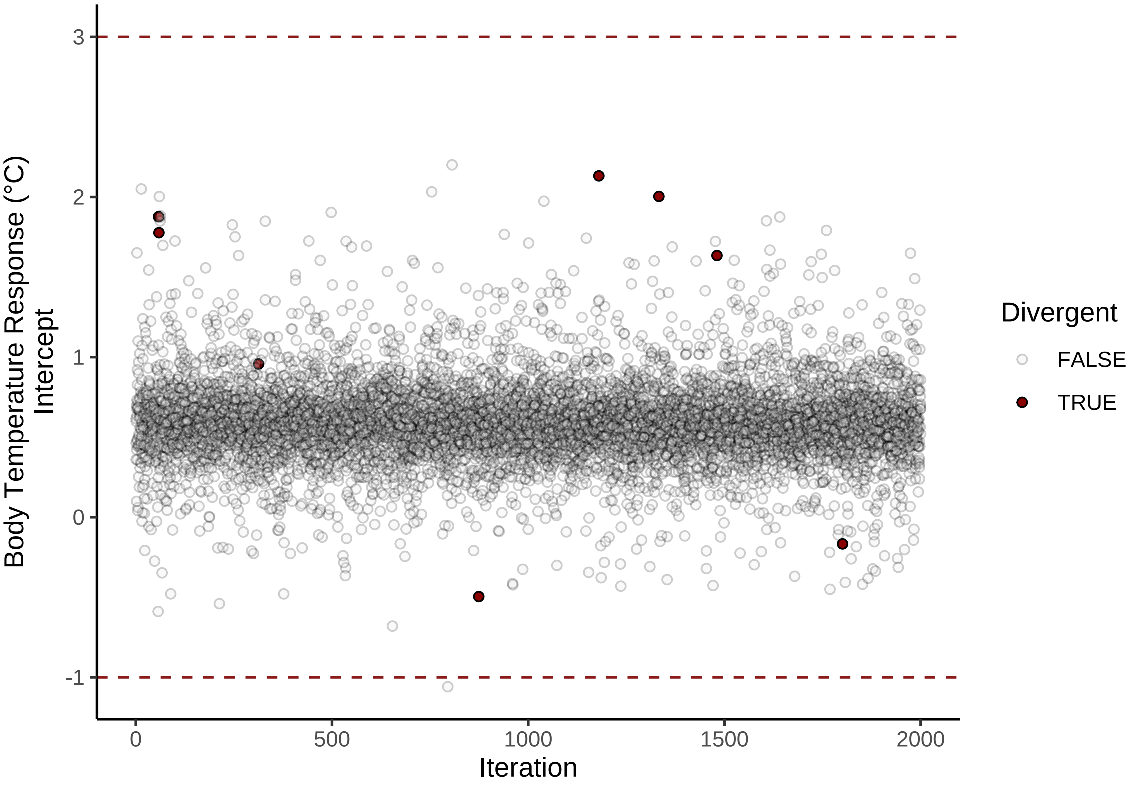


***Figure 7:*** *Posterior draws from Bayesian path analysis predicting body temperature responses (°C) to the cold (10°C) as a direct and indirect function of body mass (g) and tarsus length (mm) in adult old Japanese quail. Estimate body temperature response intercepts from draws are plotted against sample iteration. Dashed red lines indicate 95% limits of our prior on the body temperature response intercept.*

Our prior on the intercept of body temperature responses to cold appears slightly too generous and mismatched with our data. We therefore restrict the standard deviation of our prior to 0.5, rather than 1, and reconstruct our model.

```
tbResponseCold8Weeks <-
  brm(
    data = tbResponse %>%
      filter(challenge == "cold") %>%
      mutate(
        mass = mass - mean(mass, na.rm = T),
        tarsus = tarsus - mean(tarsus, na.rm = T),
        pretreatment = ifelse(pretreatment == "cold", "A",
          ifelse(pretreatment == "neutral", "B", "C")
        )
      ) %>%
      mutate(pretreatment = factor(pretreatment,
        levels = c("B", "A", "C")
      )) %>%
      drop_na(),
    family = "gaussian",
    bf(mass ~ pretreatment + (1 | batch)) +
      bf(tarsus ~ mass + pretreatment + (1 | batch)) +
      bf(deltaTb ~ tarsus + mass + pretreatment + (1 | batch)) +
      set_rescor(FALSE),
    prior = c(
      set_prior("normal(0, 10)",
        class = "Intercept",
        resp = "mass"
      ),
      set_prior("normal(0, 25)",
        class = "b",
        coef = "pretreatmentA", resp = "mass"
      ),
      set_prior("normal(0, 25)",
        class = "b",
        coef = "pretreatmentC", resp = "mass"
      ),
      set_prior("exponential(2.5)",
        class = "sd",
        group = "batch", resp = "mass"
      ),
      set_prior("exponential(0.15)",
        class = "sigma",
        resp = "mass"
      ),
      set_prior("normal(0, 2.5)",
        class = "Intercept",
        resp = "tarsus"
      ),
      set_prior("normal(0, 2.5)",
        class = "b",
        coef = "pretreatmentA", resp = "tarsus"
      ),
      set_prior("normal(0, 2.5)",
        class = "b",
        coef = "pretreatmentC", resp = "tarsus"
      ),
      set_prior("skew_normal(0, 0.25, 5)",
        class = "b",
        coef = "mass",
        resp = "tarsus"
      ),
      set_prior("exponential(2)",
        class = "sd",
        group = "batch",
        resp = "tarsus"
      ),
      set_prior("exponential(1)",
        class = "sigma",
        resp = "tarsus"
      ),
```

```
      set_prior("normal(1, 0.5)",
        class = "Intercept",
        resp = "deltaTb"
      ),
      set_prior("normal(0, 1)",
        class = "b",
        coef = "pretreatmentA",
        resp = "deltaTb"
      ),
      set_prior("normal(0, 1)",
        class = "b",
        coef = "pretreatmentC",
        resp = "deltaTb"
      ),
      set_prior("normal(0, 0.01)",
        class = "b",
        coef = "mass",
        resp = "deltaTb"
      ),
      set_prior("normal(0, 0.1)",
        class = "b",
        coef = "tarsus",
        resp = "deltaTb"
      ),
      set_prior("exponential(2.5)",
        class = "sd",
        group = "batch",
        resp = "deltaTb"
      ),
      set_prior("exponential(2.5)",
        class = "sigma",
        resp = "deltaTb"
      )
    ),
    iter = 50000, warmup = 10000, cores = 4, chains = 4, thin = 20,
    control = list(adapt_delta = .97, max_treedepth = 14),
    silent = TRUE, refresh = 0,
    file = "./models/tbResponseToColdRevised.Rds"
  )
```

Divergent transitions are almost entirely resolved. We therefore proceed to assessing: (1) convergence of our HMC chains (here, using Gelman-Rubin statistics), (2) evidence of within chain autocorrelation (here, using the ratio of effective sample sizes to sample sizes), and (3) the capacity of our model to predict true densities, and individual measures, of body temperature responses to cold.

```
p1 <- mcmc_rhat(rhat(tbResponseCold8Weeks)) +
  theme(legend.position = "none") +
  xlab(
    TeX('$\\hat{R}$')
  )

p2 <- mcmc_neff(neff_ratio(tbResponseCold8Weeks), size = 2) +
  theme(legend.position = "none") +
  xlab(
    TeX('$N_{eff}/N~Ratio$')
  )

p3 <- pp_check2(tbResponseCold8Weeks,
          xlab = "Change in\nBody Temperature (°C)",
          resp = "deltaTb"
          )
p4 <- tbResponseCold8Weeks$data %>%
  mutate("fit" = fitted(tbResponseCold8Weeks, resp = "deltaTb",
                        robust = TRUE)[,"Estimate"]) %>%
  ggplot(aes(x = fit, y = deltaTb)) +
  geom_point(size = 2, colour = "black", pch = 21, fill = "grey70") +
```

```
  geom_smooth(method = "lm", linetype = "dashed",
              colour = "black", se = FALSE) +
  xlab("Fitted Body Temperature\nResponse (°C)") +
  ylab("True Body Temperature\nResponse (°C)") +
  theme_classic()

(p1 + p2)/(p3 + p4) + plot_annotation(tag_levels = "A")
```

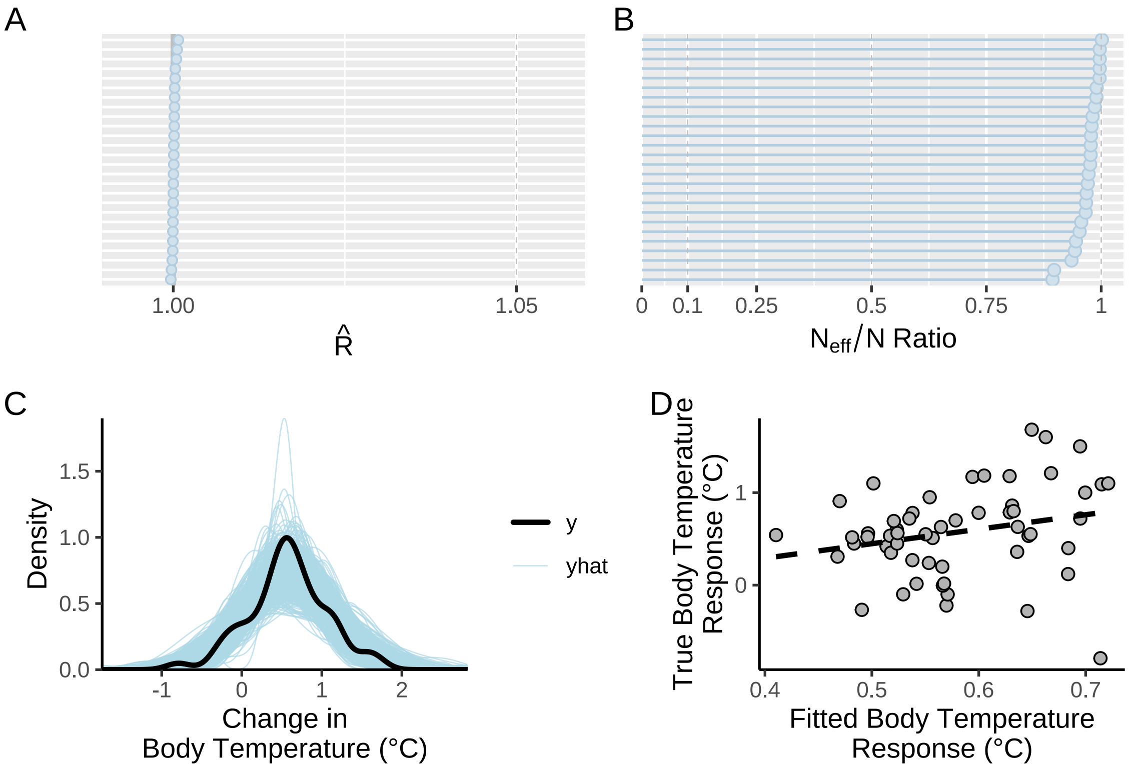


***Figure 8:*** *Validations for a Bayesian path analysis ultimately predicting body temperature responses (°C) to cold (10°C) in eight week old Japanese quail. Baseline body temperatures represent those measured at 30°C. Panel A displays displays Gelman-Rubin statistics for all model parameters. Panel B displays the ratio of effective sample sizes to sample sizes, again, for each model parameter. Panel C displays densities of true (y) and predicted (yhat) values of body temperature responses to cold. Panel D displays true and fitted body temperature responses. The dashed line in panel D represents that line of best fit as estimated by the R package ggplot2 (Wickham 2011).*

Chains have evidently converged and we see no evidence of problematic within-chain autocorrelation. Predictions from our model also reasonably predict true body temperature response values. Next, we visualise model posteriors to check for evidence of heteroskedasticity or high-influence measures.

```
g(p1, p2, p3, p4, p5, p6) %=% list(
  ggplot(
    tbResponseCold8Weeks$data %>%
      mutate(
        "residuals" =
          residuals(tbResponseCold8Weeks,
            type = "ordinary",
            robust = TRUE,
            resp = "deltaTb"
          )[, "Estimate"]
      ),
    aes(x = residuals)
```

```
) +
  geom_density(colour = "black") +
  theme_classic() +
  xlab("Ordinary Residuals") +
  ylab("Denstiy"),

ggplot(
  tbResponseCold8Weeks$data %>%
    mutate(
      "residuals" =
        residuals(tbResponseCold8Weeks,
          type = "ordinary",
          robust = TRUE,
          resp = "deltaTb"
        )[, "Estimate"]
    ), aes(sample = residuals)
) +
  stat_qq(colour = "grey50") +
  stat_qq_line() +
  xlab("Theoretical") +
  ylab("Sample") +
  theme_classic(),

ggplot(
  tbResponseCold8Weeks$data %>%
    mutate(
      "residuals" =
        residuals(tbResponseCold8Weeks,
          type = "ordinary",
          robust = TRUE,
          resp = "deltaTb"
        )[, "Estimate"],
      "resSE" = residuals(tbResponseCold8Weeks,
        type = "ordinary",
        robust = TRUE,
        resp = "deltaTb"
      )[, "Est.Error"]
    ),
  aes(x = mass, y = residuals)
) +
  geom_errorbar(
    aes(
      x = mass, ymin = residuals - resSE,
      ymax = residuals + resSE
    ),
    colour = "black", width = 2
  ) +
  geom_point(
    size = 2, pch = 21, colour = "black",
    fill = "grey70"
  ) +
  theme_classic() +
  xlab("Body Mass\n(g; Mean-Centred)") +
  ylab("Ordinary Residuals"),

ggplot(
  tbResponseCold8Weeks$data %>%
    mutate(
      "residuals" =
        residuals(tbResponseCold8Weeks,
          type = "ordinary",
          robust = TRUE,
          resp = "deltaTb"
        )[, "Estimate"],
      "resSE" = residuals(tbResponseCold8Weeks,
        type = "ordinary",
        robust = TRUE,
```

```
        resp = "deltaTb"
      )[, "Est.Error"]
    ),
  aes(x = tarsus, y = residuals)
) +
  geom_errorbar(
    aes(
      x = tarsus, ymin = residuals - resSE,
      ymax = residuals + resSE
    ),
    colour = "black", width = 2
  ) +
  geom_point(
    size = 2, pch = 21, colour = "black",
    fill = "grey70"
  ) +
  theme_classic() +
  xlab("Tarsus Length\n(mm; Mean-Centred)") +
  ylab("Ordinary Residuals"),

ggplot(
  tbResponseCold8Weeks$data %>%
    mutate(
      "residuals" =
        residuals(tbResponseCold8Weeks,
          type = "ordinary",
          robust = TRUE,
          resp = "deltaTb"
        )[, "Estimate"],
      pretreatment = ifelse(pretreatment == "B",
        "Mild (20°C)",
        ifelse(pretreatment == "A",
          "Cold (10°C)", "Warm (30°C)"
        )
      )
    ),
  aes(x = pretreatment, y = residuals)
) +
  geom_boxplot(colour = "black", alpha = 0.5, fill = "grey70") +
  geom_point(
    size = 1.5, colour = "black",
    position = position_jitter(width = 0.25)
  ) +
  theme_classic() +
  xlab("Rearing Treatment") +
  ylab("Ordinary Residuals"),

ggplot(
  tbResponseCold8Weeks$data %>%
    mutate(
      "residuals" =
        residuals(tbResponseCold8Weeks,
          type = "ordinary",
          robust = TRUE,
          resp = "deltaTb"
        )[, "Estimate"]
    ),
  aes(x = residuals, fill = batch)
) +
  geom_density(colour = "black", alpha = 0.5) +
  scale_fill_manual(
    values = c("black", "grey70"),
    name = "Egg Batch"
  ) +
  theme_classic() +
  xlab("Ordinary Residuals") +
  ylab("Density")
```

```
)

(p1 + p2) / (p3 + p4) / (p5 + p6) +
  plot_annotation(tag_levels = "A")
```

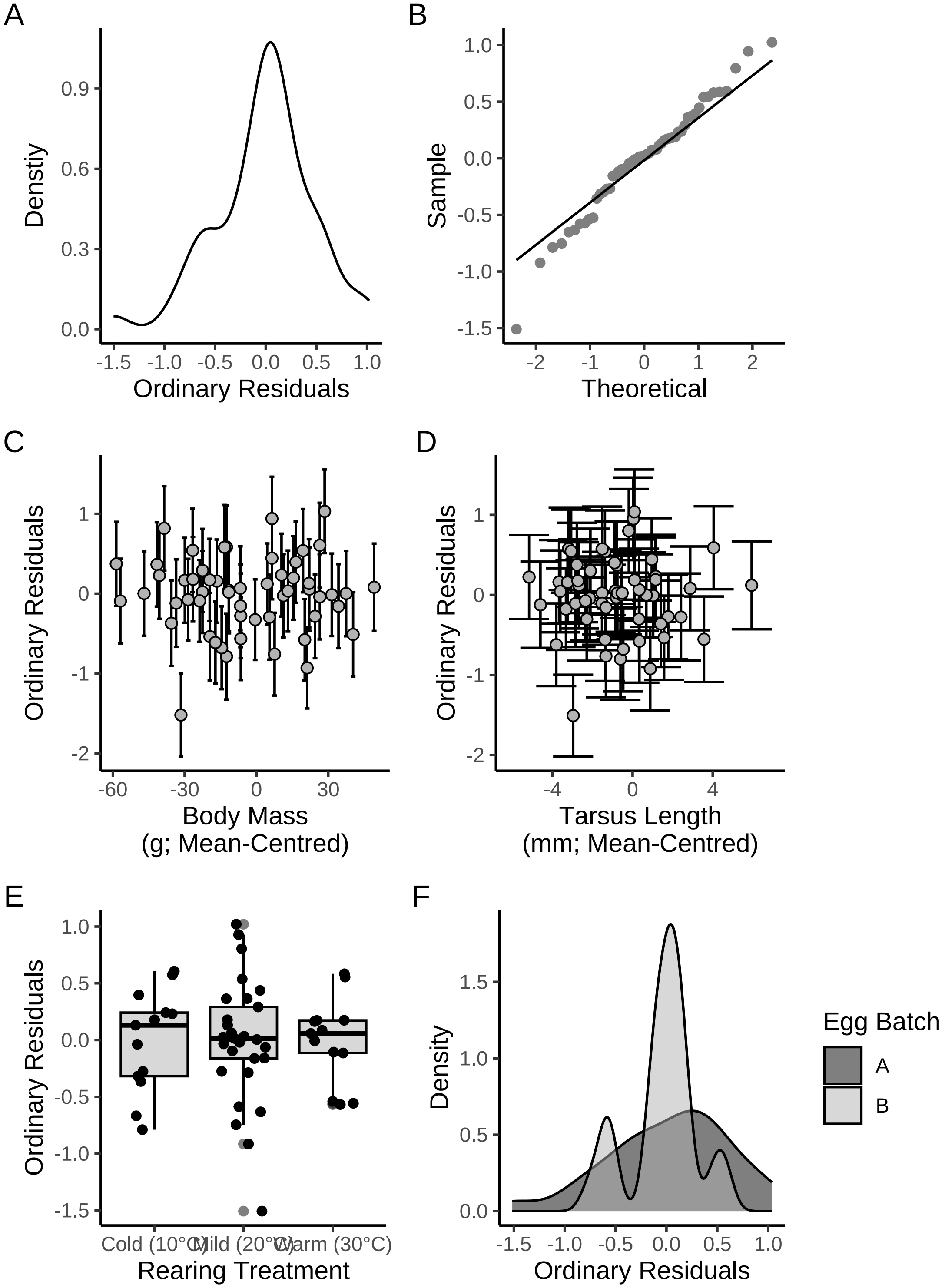


***Figure 9:*** *Ordinary residuals from a Bayesian path analysis ultimately predicting body temperature responses (°C) to cold (10°C) in eight week old Japanese quail. Panel A displays the density of median body temperature response residuals, panel B displays theoretical against sample residuals (qq-plot), panel C displays median residuals against mean-centred body mass (g), panel D displays median residuals against mean-centred tarsus length (mm), panel E displays median residuals against rearing treatment, and panel F displays median residuals against egg source number (or batch). Errorbars in panels B and C indicate one median absolute deviation around median residuals.*

No obvious concerns. We proceed by calculating Bayesian $R^2$ for models within our path analysis, then plotting posterior coefficients to check for evidence of skewing.

```
# Checking R2 values

caption <- paste0("R\\textsuperscript{2} for Bayesian path analysis ",
                  "ultimate predicting body temperature responses (°C) to cold ",
                  "(10°C) in eight week old Japanese quail. Baseline body ",
                  "temperatures represent those measured at thermoneutrality ",
                  "(30°C)"
)

brms::bayes_R2(tbResponseCold8Weeks,
  robust = TRUE, ndraws = 1000
) %>%
  as.data.frame() %>%
  rownames_to_column(var = "Response") %>%
  mutate(Response = ifelse(Response == "R2mass", "Body Mass (g)",
    ifelse(Response == "R2tarsus", "Tarsus Length (mm)",
      "Change in Tb (°C)"
    )
  )) %>%
  mutate(
    Estimate = round(Estimate, digits = 4),
    Est.Error = round(Est.Error, digits = 4),
    Q2.5 = round(Q2.5, digits = 4),
    Q97.5 = round(Q97.5, digits = 4)
  ) %>%
  rename(
    "R\\textsuperscript{2}" = Estimate,
    "Standard Error" = Est.Error,
    `2.5\\%CI` = "Q2.5", `97.5\\% CI` = "Q97.5"
  ) %>%
  kbl(.,
    longtable = T, booktabs = T, format = "latex",
    caption = caption, escape = FALSE
  ) %>%
  kable_styling(latex_options = "striped")
```

***Table 2:*** *$R^2$ for Bayesian path analysis ultimate predicting body temperature responses (°C) to cold (10°C) in eight week old Japanese quail. Baseline body temperatures represent those measured at thermoneutrality (30°C)*

| Response | $R^2$ | Standard Error | 2.5%CI | 97.5% CI |
|---|---|---|---|---|
| Body Mass (g) | 0.0229 | 0.0239 | 0.0008 | 0.1041 |
| Tarsus Length (mm) | 0.2947 | 0.0853 | 0.1193 | 0.4352 |
| Change in Tb (°C) | 0.0868 | 0.0529 | 0.0187 | 0.2128 |

```
as.data.frame(tbResponseCold8Weeks) %>%
    select(
      "Intercept" = b_deltaTb_Intercept,
      "Body Mass (g)" = b_deltaTb_mass,
      "Tarsus Length\n(mm)" = b_deltaTb_tarsus,
      "Cold Rearing\n(10°C)" = b_deltaTb_pretreatmentA,
      "Warm Rearing\n(30°C)" = b_deltaTb_pretreatmentC,
      "Egg Batch (mu)" = sd_batch__deltaTb_Intercept,
      "Sigma" = sigma_deltaTb
    ) %>%
    pivot_longer(everything(), names_to = "Par",
                 values_to = "Coefs") %>%
    ggplot(aes(x = Coefs)) +
    facet_wrap(~Par, scales = "free", ncol = 2) +
    geom_density(colour = "black", alpha = 0.5, fill = "grey70") +
    geom_vline(xintercept = 0, linetype = "dashed", colour = "black") +
```

```
    ylab("Density") +
    theme_classic() +
    theme(axis.title.x = element_blank())
```

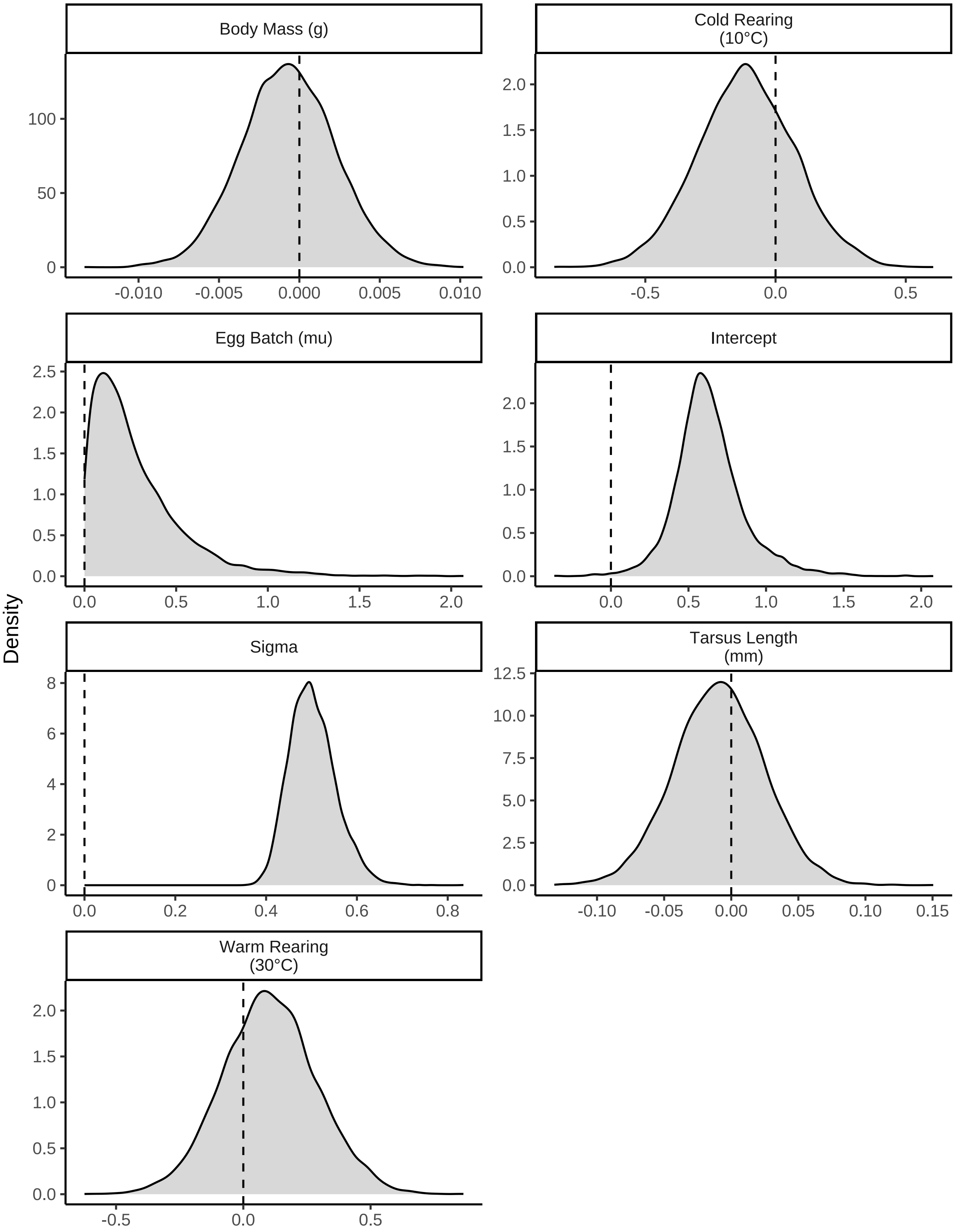


***Figure 10:*** *Posterior densities for model coefficients derived from a Bayesian path analysis ultimately predicting body temperature responses (°C) to cold (10°C) in eight week old Japanese quail. Densities are split by their respective response values (indicated with titles). Vertical dashed lines indicate 0.*

```
# Subtle skewing
```

We now summarise estimates from our path analysis, with central estimates of our coefficients being calculated as their medians and credible intervals as quantile intervals around medians.

```
caption <- paste0("Results from a Bayesian path analysis ",
                  "ultimately predicting body temperature responses to a ",
                  "cold exposure (10°C) relative to thermoneutrality (30°C)",
                  "in adult Japanese quail. Body temperature responses are ",
                  "predicted as a function of body mass (g), tarsus length ",
                  "(mm) and rearing treatment. ",
                  "Cold rearing indicates post-hatch rearing at ",
                  "10°C, relative to ",
                  "20°C (intercept), or ",
                  "30°C ('warm rearing'). ",
                  "Coefficients represent medians and credible intervals ",
                  "(CIs) represent quantile intervals. ",
                  "BF indicates Bayes Factors."
)

as.data.frame(tbResponseCold8Weeks) %>%
  summarise_all(., .funs = median) %>%
  pivot_longer(everything(),
    names_to = "Parameter",
    values_to = "Estimate"
  ) %>%
  merge(., quantileCIs(tbResponseCold8Weeks, cis = c(50, 95)),
    by = "Parameter", all.x = TRUE
  ) %>%
  filter(grepl("b_|sd_", Parameter)) %>%
  rowwise() %>%
  mutate("BF" = ifelse(Estimate < 0,
    (2 * mean(as.data.frame(
      tbResponseCold8Weeks
    )[, Parameter] <= 0)) /
      (2 * mean(as.data.frame(
        tbResponseCold8Weeks
      )[, Parameter] >= 0)),
    (2 * mean(as.data.frame(
      tbResponseCold8Weeks
    )[, Parameter] >= 0)) /
      (2 * mean(as.data.frame(
        tbResponseCold8Weeks
      )[, Parameter] <= 0))
  )) %>%
  ungroup() %>%
  mutate(
    "Estimate" = round(Estimate, digits = 4),
    "BF" = round(BF, digits = 4),
    "N" = nrow(tbResponseCold8Weeks$data)
  ) %>%
  rowwise() %>%
  mutate(Parameter = gsub("deltaTb", "deltaT", Parameter)) %>%
  ungroup() %>%
  mutate("Parameter" = ifelse(grepl("b_", Parameter),
                              gsub("b_", "", Parameter),
                              gsub("Intercept", "batch",
                                   gsub(".*__", "", Parameter)
                                   )
                              )
         ) %>%
  mutate(
    "Response" = gsub("_.*", "", Parameter),
    "Parameter" = gsub(".*_", "", Parameter)
  ) %>%
  merge(., tribble(
```

```
    ~Response, ~response, ~level,
    "mass", "Body Mass (g)", "A",
    "deltaT",
    "Delta Body Temperature (°C)", "D",
    "tarsus", "Tarsus Length (mm)",
    "B"
  ),
  by = "Response"
  ) %>%
  merge(., tribble(
    ~Parameter, ~parameter, ~number,
    "Intercept", "Intercept", "1",
    "mass", "Body Mass (g)", "4",
    "tarsus", "Tarsus Length (mm)", "5",
    "pretreatmentA", "Cold Rearing", "2",
    "pretreatmentC", "Warm Rearing", "3",
    "batch", "Egg Batch [mu]", "6"
  ),
  by = "Parameter"
  ) %>%
  mutate(
    `50\\% CI` = paste0("(", paste(
      round(Low_CI_50, digits = 4),
      round(High_CI_50, digits = 4),
      sep = ", "
    ), ")"),
    `95\\% CI` = paste0("(", paste(
      round(Low_CI_95, digits = 4),
      round(High_CI_95, digits = 4),
      sep = ", "
    ), ")")
  ) %>%
  select(-c(Low_CI_50, High_CI_50, Low_CI_95, High_CI_95)) %>%
  select(
    "Response" = "response", "Parameter" = "parameter", N,
    Estimate, `50\\% CI`, `95\\% CI`, BF, level, number
  ) %>%
  arrange(level, number) %>%
  select(-c(level, number)) %>%
  kbl(.,
    longtable = T, booktabs = T, format = "latex", escape = FALSE,
    caption = caption
  ) %>%
  column_spec(column = c(1:2), width = "2.2cm") %>%
  column_spec(column = c(3:10), width = "1.9cm") %>%
  kable_styling(latex_options = "striped")
```

***Table 3:*** *Results from a Bayesian path analysis ultimately predicting body temperature responses to a cold exposure (10°C) relative to thermoneutrality (30°C)in adult Japanese quail. Body temperature responses are predicted as a function of body mass (g), tarsus length (mm) and rearing treatment. Cold rearing indicates post-hatch rearing at 10°C, relative to 20°C (intercept), or 30°C ('warm rearing'). Coefficients represent medians and credible intervals (CIs) represent quantile intervals. BF indicates Bayes Factors.*

| Response | Parameter | N | Estimate | 50% CI | 95% CI | BF |
|---|---|---|---|---|---|---|
| Body Mass (g) | Intercept | 55 | -4.9297 | (-8.0227, -1.9287) | (-14.0289, 4.2278) | 6.1878 |
| Body Mass (g) | Cold Rearing | 55 | 3.4914 | (-2.1106, 8.8435) | (-13.353, 19.4883) | 1.9597 |
| Body Mass (g) | Warm Rearing | 55 | 0.9890 | (-4.2715, 6.7385) | (-14.9981, 16.8916) | 1.2290 |
| Body Mass (g) | Egg Batch [mu] | 55 | 0.2701 | (0.1144, 0.5448) | (0.01, 1.4313) | Inf |
| Tarsus Length (mm) | Intercept | 55 | -0.6777 | (-0.982, -0.3828) | (-1.6749, 0.3089) | 12.2013 |

| | | | | | | |
|---|---|---|---|---|---|---|
| Tarsus Length (mm) | Cold Rearing | 55 | 0.3764 | (-0.0789, 0.8243) | (-0.9394, 1.6457) | 2.4261 |
| Tarsus Length (mm) | Warm Rearing | 55 | 0.1908 | (-0.2542, 0.6285) | (-1.1094, 1.5409) | 1.5898 |
| Tarsus Length (mm) | Body Mass (g) | 55 | 0.0440 | (0.037, 0.0507) | (0.0235, 0.0644) | Inf |
| Tarsus Length (mm) | Egg Batch [mu] | 55 | 0.2593 | (0.1081, 0.5019) | (0.01, 1.3701) | Inf |
| Delta Body Temperature (°C) | Intercept | 55 | 0.6135 | (0.5033, 0.7432) | (0.233, 1.1632) | 241.4242 |
| Delta Body Temperature (°C) | Cold Rearing | 55 | -0.1150 | (-0.2381, 0.0111) | (-0.4854, 0.258) | 2.6714 |
| Delta Body Temperature (°C) | Warm Rearing | 55 | 0.1011 | (-0.0218, 0.2228) | (-0.2566, 0.4744) | 2.5119 |
| Delta Body Temperature (°C) | Body Mass (g) | 55 | -0.0007 | (-0.0026, 0.0013) | (-0.0063, 0.0052) | 1.4737 |
| Delta Body Temperature (°C) | Tarsus Length (mm) | 55 | -0.0091 | (-0.0315, 0.0135) | (-0.075, 0.0569) | 1.5592 |
| Delta Body Temperature (°C) | Egg Batch [mu] | 55 | 0.2089 | (0.1037, 0.3797) | (0.0106, 0.9472) | Inf |

### Body Temperature Responses to Heat Exposures

Unlike metabolic responses to cold exposure, we found that metabolic responses to heat exposure (40°C) were indeed influenced by body size and tarsus length among adult Japanese quail (Tabh et al 2024). Specifically, adult quail with relatively large body sizes and short tarsi displayed larger increases in metabolism during a heat exposure than those with relatively small body sizes and long tarsi. However, the magnitude of these effects were small, limiting their biological implications. Nevertheless, it is still possible that body size and tarsus length also influenced how body temperature responded to heat exposure, potentially increasing the biological implications of being large with atypically short appendages in the heat. To test this, we again evaluated whether body temperature responses to a heat exposure varied by body mass and tarsus length, while controlling for prior temperature exposure. This was achieved by quantified the mean change in body temperature displayed between that measured in the heat (40°C) and that measured at thermoneutrality (30°C) then modelling this change as a function of body mass, tarsus length, and rearing condition. Given that rearing condition also influences body mass and tarsus length in our quail (Tabh et al 2024), however, this model was nested with a path analysis identical to that described for body temperature responses to the cold above, but with body temperature responses to heat (°C) as the ultimate response variable.

For this analysis, priors for our models predicting body mass and tarsus length on their own remained the same as those described previously (i.e. for our path analysis predicting body temperature responses to the cold). Priors for our model predicting body temperature responses to heat, however, differed slightly, but were equally informed by Persson et al (2024) and the observed ranges of body temperature responses and morphometric measurements. For the effects of body mass and tarsus length on body temperature responses, we again assumed that effects larger than the range of body temperature responses divided by the range of the given morphological variable were unlikely (standard deviation of a normally-distributed prior = $0.5 \cdot \frac{Range[Delta\ Tb]}{Range[Mass\ or\ Tarsus]}$). Priors for our model predicting body temperature responses to heat were therefore as follows:

$$\beta_{c0} \sim \mathcal{N}(1, 1)$$

$$\beta_{c1} \sim \mathcal{N}(0, 1)$$

$$\beta_{c2} \sim \mathcal{N}(0, 1)$$
$$\beta_{c3} \sim \mathcal{N}(0, 0.01)$$
$$\beta_{c4} \sim \mathcal{N}(0, 0.15)$$
$$\mu 0c \sim \exp(2.5)$$
$$\epsilon_c \sim \exp(2.5)$$

Before constructing our path analysis, we check raw body temperature response values for oddities, then assess suitability of our priors using a prior predictive check (as described above).

```
tbResponse %>%
  filter(!is.na(deltaTb) & challenge == "warm") %>%
  ggplot(aes(x = 1:nrow(.), y = deltaTb)) +
  geom_rect(aes(xmin = -Inf, xmax = Inf,
                ymin = mean(deltaTb) - 3.5*sd(deltaTb),
                ymax = mean(deltaTb) + 3.5*sd(deltaTb)
                ),
            fill = "grey80", alpha = 0.5, colour = "black") +
  geom_hline(yintercept = 0, linetype = "dashed", colour = "black") +
  geom_point(size = 2, colour = "black", alpha = 0.7) +
  scale_y_continuous(sec.axis = sec_axis( trans=~., name="Second Axis",
                                          breaks = c(1.5, -0.2),
                                          labels = c("Increasing Tb",
                                                     "Decreasing Tb"))
                     ) +
  theme_classic() +
  theme(axis.text.y.right = element_text(angle = 270),
        axis.ticks.y.right = element_blank(),
        axis.title.y.right = element_blank()) +
  xlab("Sample Number") +
  ylab("Change in\nBody Temperature (°C)")
```

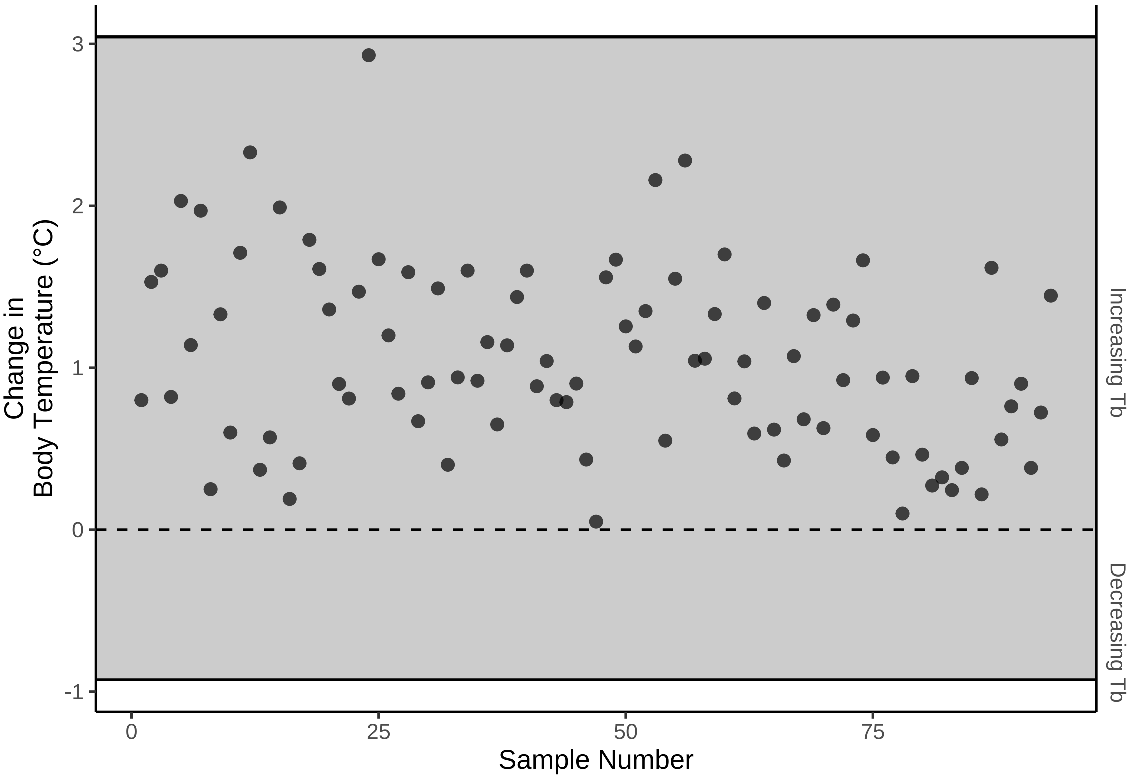


***Figure 11:*** *Cleveland dotplot of mean changes in body temperature (°C) between 40°C and 30°C, each drawn from adult Japanese quail. Again, dots represent raw data points and rectangles indicate means 3.5 times the standard deviation at a given ambient temperature. The dashed line indicates no change in body temperature.*

```
tbResponseHot8WeeksPPCheck <-
  brm(
    data = tbResponse %>%
      filter(week == "8" & challenge == "warm") %>%
      mutate(
        mass = mass - mean(mass, na.rm = T),
        tarsus = tarsus - mean(tarsus, na.rm = T),
        pretreatment = ifelse(pretreatment == "cold", "A",
          ifelse(pretreatment == "neutral", "B", "C")
        )
      ) %>%
      mutate(pretreatment = factor(pretreatment,
                                   levels = c("B", "A", "C"))
             ) %>%
      drop_na(),
    family = "gaussian",
    bf(mass ~ pretreatment + (1 | batch)) +
      bf(tarsus ~ mass + pretreatment + (1 | batch)) +
      bf(deltaTb ~ tarsus + mass + pretreatment + (1 | batch)) +
      set_rescor(FALSE),
    prior = c(
      set_prior("normal(0, 10)",
        class = "Intercept",
        resp = "mass"
      ),
      set_prior("normal(0, 25)",
        class = "b",
        coef = "pretreatmentA",
        resp = "mass"
      ),
      set_prior("normal(0, 25)",
```

```
      class = "b",
      coef = "pretreatmentC",
      resp = "mass"
    ),
    set_prior("exponential(2.5)",
      class = "sd",
      group = "batch",
      resp = "mass"
    ),
    set_prior("exponential(0.15)",
      class = "sigma",
      resp = "mass"
    ),
    set_prior("normal(0, 2.5)",
      class = "Intercept",
      resp = "tarsus"
    ),
    set_prior("normal(0, 2.5)",
      class = "b",
      coef = "pretreatmentA",
      resp = "tarsus"
    ),
    set_prior("normal(0, 2.5)",
      class = "b",
      coef = "pretreatmentC",
      resp = "tarsus"
    ),
    set_prior("skew_normal(0, 0.25, 5)",
      class = "b",
      coef = "mass",
      resp = "tarsus"
    ),
    set_prior("exponential(2)",
      class = "sd",
      group = "batch",
      resp = "tarsus"
    ),
    set_prior("exponential(1)",
      class = "sigma",
      resp = "tarsus"
    ),
    set_prior("normal(1, 1)",
      class = "Intercept",
      resp = "deltaTb"
    ),
    set_prior("normal(0, 1)",
      class = "b",
      coef = "pretreatmentA",
      resp = "deltaTb"
    ),
    set_prior("normal(0, 1)",
      class = "b",
      coef = "pretreatmentC",
      resp = "deltaTb"
    ),
    set_prior("normal(0, 0.01)",
      class = "b",
      coef = "mass",
      resp = "deltaTb"
    ),
    set_prior("normal(0, 0.15)",
      class = "b",
      coef = "tarsus",
      resp = "deltaTb"
    ),
    set_prior("exponential(2.5)",
      class = "sd",
```

```
        group = "batch",
        resp = "deltaTb"
      ),
      set_prior("exponential(2.5)",
        class = "sigma",
        resp = "deltaTb"
      )
    ),
    iter = 50000, warmup = 10000, cores = 4, chains = 4, thin = 20,
    control = list(adapt_delta = .97, max_treedepth = 14),
    silent = TRUE, refresh = 0,
    sample_prior = "only",
    file = "./models/bodyTemperatureResponsesToHeatPPCheck.Rds"
  )

pp_check2(tbResponseHot8WeeksPPCheck,
          xlab = "Change in\nBody Temperature (°C)",
          resp = "deltaTb"
          )
```

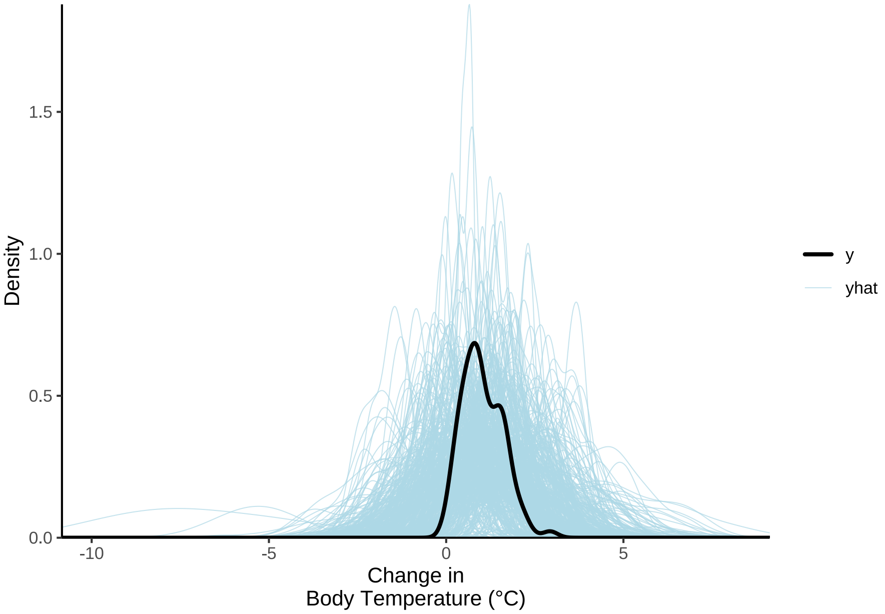


***Figure 12:*** *Overlay of predicted (blue) and true (black) body temperature response (°C) densities, where predicted densities are derived from priors in a Bayesian path analyses. Clear overlap betwen the black and blue lines suggests that model priors are reasonable with respect to the data.*

Priors appear suitable. Our full analysis is therefore constructed and chains checked for convergence and evidence of autocorrelation. Further, the capacity of analysis to predict true body temperature responses to heat is visually assessed.

```
tbResponseHot8Weeks <-
  brm(
    data = tbResponse %>%
      filter(week == "8" & challenge == "warm") %>%
      mutate(
        mass = mass - mean(mass, na.rm = T),
        tarsus = tarsus - mean(tarsus, na.rm = T),
        pretreatment = ifelse(pretreatment == "cold", "A",
          ifelse(pretreatment == "neutral", "B", "C")
        )
      ) %>%
```

```
    mutate(pretreatment = factor(pretreatment,
                                 levels = c("B", "A", "C"))
           ) %>%
    drop_na(),
  family = "gaussian",
  bf(mass ~ pretreatment + (1 | batch)) +
    bf(tarsus ~ mass + pretreatment + (1 | batch)) +
    bf(deltaTb ~ tarsus + mass + pretreatment + (1 | batch)) +
    set_rescor(FALSE),
  prior = c(
    set_prior("normal(0, 10)",
      class = "Intercept",
      resp = "mass"
    ),
    set_prior("normal(0, 25)",
      class = "b",
      coef = "pretreatmentA",
      resp = "mass"
    ),
    set_prior("normal(0, 25)",
      class = "b",
      coef = "pretreatmentC",
      resp = "mass"
    ),
    set_prior("exponential(2.5)",
      class = "sd",
      group = "batch",
      resp = "mass"
    ),
    set_prior("exponential(0.15)",
      class = "sigma",
      resp = "mass"
    ),
    set_prior("normal(0, 2.5)",
      class = "Intercept",
      resp = "tarsus"
    ),
    set_prior("normal(0, 2.5)",
      class = "b",
      coef = "pretreatmentA",
      resp = "tarsus"
    ),
    set_prior("normal(0, 2.5)",
      class = "b",
      coef = "pretreatmentC",
      resp = "tarsus"
    ),
    set_prior("skew_normal(0, 0.25, 5)",
      class = "b",
      coef = "mass",
      resp = "tarsus"
    ),
    set_prior("exponential(2)",
      class = "sd",
      group = "batch",
      resp = "tarsus"
    ),
    set_prior("exponential(1)",
      class = "sigma",
      resp = "tarsus"
    ),
    set_prior("normal(1, 1)",
      class = "Intercept",
      resp = "deltaTb"
    ),
    set_prior("normal(0, 1)",
      class = "b",
```

```
        coef = "pretreatmentA",
        resp = "deltaTb"
      ),
      set_prior("normal(0, 1)",
        class = "b",
        coef = "pretreatmentC",
        resp = "deltaTb"
      ),
      set_prior("normal(0, 0.01)",
        class = "b",
        coef = "mass",
        resp = "deltaTb"
      ),
      set_prior("normal(0, 0.15)",
        class = "b",
        coef = "tarsus",
        resp = "deltaTb"
      ),
      set_prior("exponential(2.5)",
        class = "sd",
        group = "batch",
        resp = "deltaTb"
      ),
      set_prior("exponential(2.5)",
        class = "sigma",
        resp = "deltaTb"
      )
    ),
    iter = 50000, warmup = 10000, cores = 4, chains = 4, thin = 20,
    control = list(adapt_delta = .97, max_treedepth = 14),
    silent = TRUE, refresh = 0,
    file = "./models/bodyTemperatureResponsesToHeat.Rds"
  )

p1 <- mcmc_rhat(rhat(tbResponseHot8Weeks)) +
  theme(legend.position = "none") +
  xlab(
    TeX('$\\hat{R}$')
  )

p2 <- mcmc_neff(neff_ratio(tbResponseHot8Weeks), size = 2) +
  theme(legend.position = "none") +
  xlab(
    TeX('$N_{eff}/N~Ratio$')
  )

p3 <- pp_check2(tbResponseHot8Weeks,
          xlab = "Change in\nBody Temperature (°C)",
          resp = "deltaTb"
          )
p4 <- tbResponseHot8Weeks$data %>%
  mutate("fit" = fitted(tbResponseHot8Weeks, resp = "deltaTb",
                         robust = TRUE)[,"Estimate"]) %>%
  ggplot(aes(x = fit, y = deltaTb)) +
  geom_point(size = 2, colour = "black", pch = 21, fill = "grey70") +
  geom_smooth(method = "lm", linetype = "dashed",
              colour = "black", se = FALSE) +
  xlab("Fitted Body Temperature\nResponse (°C)") +
  ylab("True Body Temperature\nResponse (°C)") +
  theme_classic()

(p1 + p2)/(p3 + p4) + plot_annotation(tag_levels = "A")
```

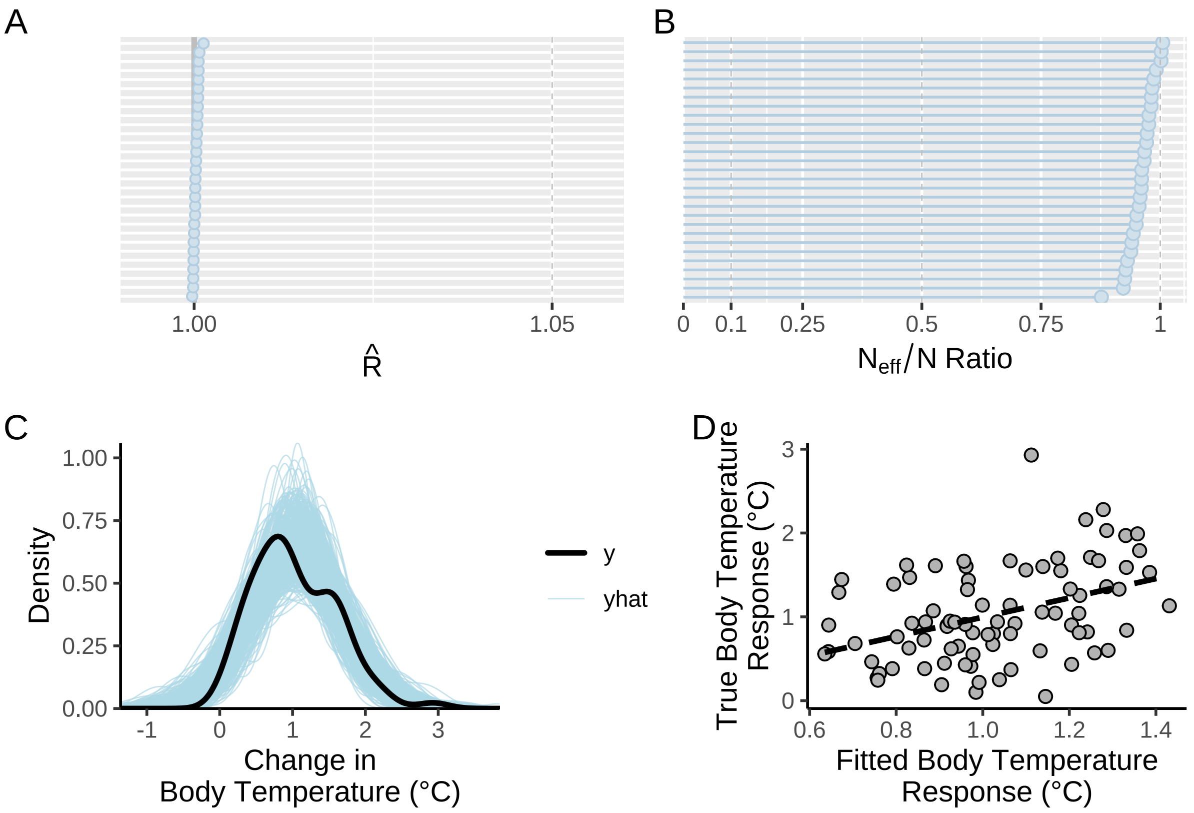


***Figure 13:*** *Validations for a Bayesian path analysis ultimately predicting body temperature responses (°C) to heat (40°C) in eight week old Japanese quail. Baseline body temperatures represent those measured at 30°C. Panel A displays displays Gelman-Rubin statistics for all model parameters. Panel B displays the ratio of effective sample sizes to sample sizes, again, for each model parameter. Panel C displays densities of true (y) and predicted (yhat) values of body temperature responses to cold. Panel D displays true and fitted body temperature responses. The dashed line in panel D represents that line of best fit as estimated by the R package ggplot2 (Wickham 2011).*

Again, our HMC chains appear well converged with little evidence of autocorrelation, and predictions from our analysis reasonably overlay with true body temperature responses. Next, distributions of residuals from our analysis are visually diagnosed.

```
g(p1, p2, p3, p4, p5, p6) %=% list(
  ggplot(
    tbResponseHot8Weeks$data %>%
      mutate(
        "residuals" =
          residuals(tbResponseHot8Weeks,
            type = "ordinary",
            robust = TRUE,
            resp = "deltaTb"
          )[, "Estimate"]
      ),
    aes(x = residuals)
  ) +
    geom_density(colour = "black") +
    theme_classic() +
    xlab("Ordinary Residuals") +
    ylab("Denstiy"),

  ggplot(
    tbResponseHot8Weeks$data %>%
```

```
    mutate(
      "residuals" =
        residuals(tbResponseHot8Weeks,
          type = "ordinary",
          robust = TRUE,
          resp = "deltaTb"
        )[, "Estimate"]
    ), aes(sample = residuals)
) +
  stat_qq(colour = "grey50") +
  stat_qq_line() +
  xlab("Theoretical") +
  ylab("Sample") +
  theme_classic(),

ggplot(
  tbResponseHot8Weeks$data %>%
    mutate(
      "residuals" =
        residuals(tbResponseHot8Weeks,
          type = "ordinary",
          robust = TRUE,
          resp = "deltaTb"
        )[, "Estimate"],
      "resSE" = residuals(tbResponseHot8Weeks,
        type = "ordinary",
        robust = TRUE,
        resp = "deltaTb"
      )[, "Est.Error"]
    ),
  aes(x = mass, y = residuals)
) +
  geom_errorbar(
    aes(
      x = mass, ymin = residuals - resSE,
      ymax = residuals + resSE
    ),
    colour = "black", width = 2
  ) +
  geom_point(
    size = 2, pch = 21, colour = "black",
    fill = "grey70"
  ) +
  theme_classic() +
  xlab("Body Mass\n(g; Mean-Centred)") +
  ylab("Ordinary Residuals"),

ggplot(
  tbResponseHot8Weeks$data %>%
    mutate(
      "residuals" =
        residuals(tbResponseHot8Weeks,
          type = "ordinary",
          robust = TRUE,
          resp = "deltaTb"
        )[, "Estimate"],
      "resSE" = residuals(tbResponseHot8Weeks,
        type = "ordinary",
        robust = TRUE,
        resp = "deltaTb"
      )[, "Est.Error"]
    ),
  aes(x = tarsus, y = residuals)
) +
  geom_errorbar(
    aes(
      x = tarsus, ymin = residuals - resSE,
```

```
        ymax = residuals + resSE
      ),
      colour = "black", width = 2
    ) +
    geom_point(
      size = 2, pch = 21, colour = "black",
      fill = "grey70"
    ) +
    theme_classic() +
    xlab("Tarsus Length\n(mm; Mean-Centred)") +
    ylab("Ordinary Residuals"),

  ggplot(
    tbResponseHot8Weeks$data %>%
      mutate(
        "residuals" =
          residuals(tbResponseHot8Weeks,
            type = "ordinary",
            robust = TRUE,
            resp = "deltaTb"
          )[, "Estimate"],
        pretreatment = ifelse(pretreatment == "B",
          "Mild (20°C)",
          ifelse(pretreatment == "A",
            "Cold (10°C)", "Warm (30°C)"
          )
        )
      ),
    aes(x = pretreatment, y = residuals)
  ) +
    geom_boxplot(colour = "black", alpha = 0.5, fill = "grey70") +
    geom_point(
      size = 1.5, colour = "black",
      position = position_jitter(width = 0.25)
    ) +
    theme_classic() +
    xlab("Rearing Treatment") +
    ylab("Ordinary Residuals"),

  ggplot(
    tbResponseHot8Weeks$data %>%
      mutate(
        "residuals" =
          residuals(tbResponseHot8Weeks,
            type = "ordinary",
            robust = TRUE,
            resp = "deltaTb"
          )[, "Estimate"]
      ),
    aes(x = residuals, fill = batch)
  ) +
    geom_density(colour = "black", alpha = 0.5) +
    scale_fill_manual(
      values = c("black", "grey40", "grey90"),
      name = "Egg Batch"
    ) +
    theme_classic() +
    xlab("Ordinary Residuals") +
    ylab("Density")
)

(p1 + p2) / (p3 + p4) / (p5 + p6) +
  plot_annotation(tag_levels = "A")
```

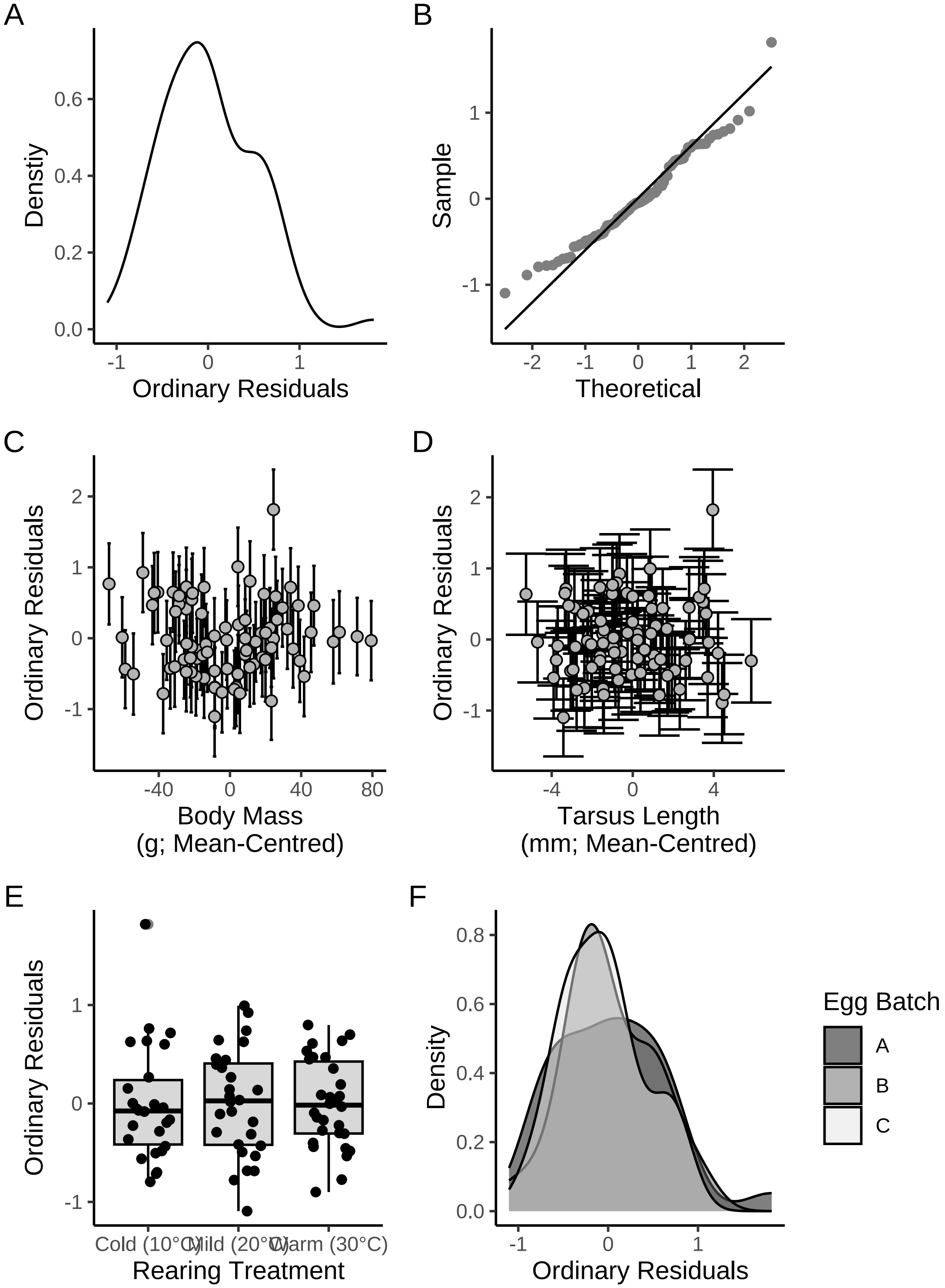


***Figure 14:*** *Ordinary residuals from a Bayesian path analysis ultimately predicting body temperature responses (°C) to heat (40°C) in eight week old Japanese quail. Panel A displays the density of median body temperature response residuals, panel B displays theoretical against sample residuals (qq-plot), panel C displays median residuals against mean-centred body mass (g), panel D displays median residuals against mean-centred tarsus length (mm), panel E displays median residuals against rearing treatment, and panel F displays median residuals against egg source number (or batch). Errorbars in panels B and C indicate one median absolute deviation around median residuals.*

One measurement appears to deviate strongly from others and from expectations of error normality. We therefore check whether this value has undue influence on model outcomes by using Pareto-smoothed, leave-one-out ("LOO") cross-validation (Vehtari et al, 2017). Influence values per data point (here, Pareto K) are then visualised for extremes.

```
loo(tbResponseHot8Weeks)$diagnostics$pareto_k %>%
  as_tibble() %>%
  ggplot(aes(x = 1:nrow(.), y = value)) +
  geom_point(pch = 21, colour = "black", fill = "grey50", size = 2) +
  geom_hline(yintercept = 0.5, colour = "black", linetype = "dashed") +
  geom_hline(yintercept = 0.7, colour = "red4", linetype = "dashed") +
  xlab("Sample") +
  ylab("Pareto K") +
  theme_classic()
```

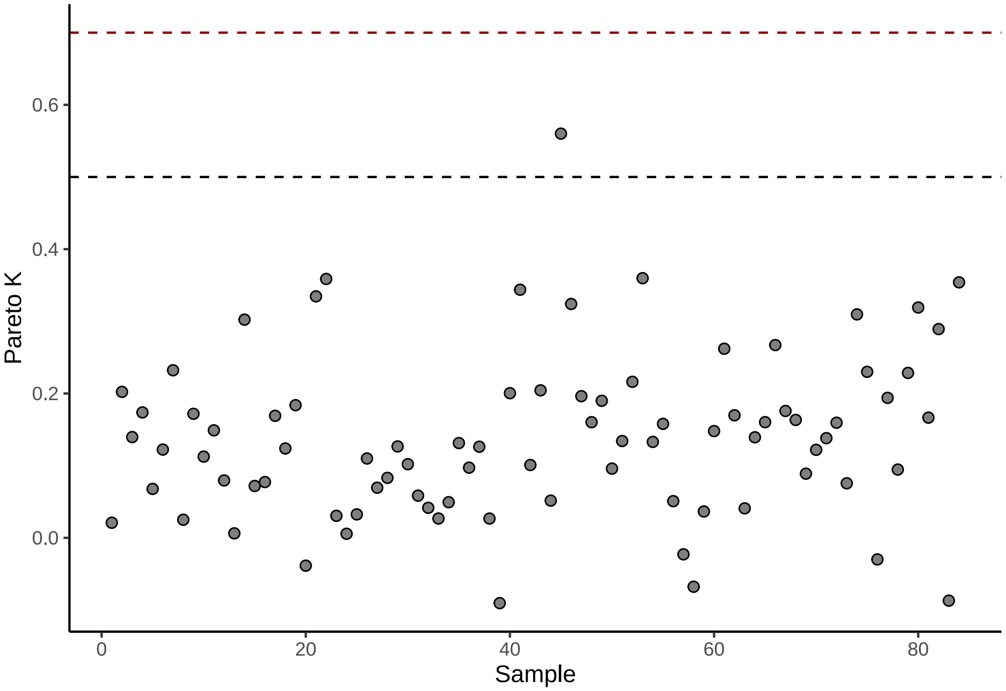


***Figure 15:*** *Relative importance of individual body temperature response values (dots; derived from mature quail) on outcomes of our Bayesian path analysis, as estimated using leave-one-out cross validations. Importance is measured here as Pareto K (Vehtari et al, 2017). The black, dashed horizontal line indicated a Pareto K of 0.5 and the red horizontal dashed line indicates as Pareto K of 0.7, above which, samples hold significant importance on analysis outcomes.*

Importance of one data point is high and as such, this point is inspected.

```
caption <- paste0("Potential body temperature response outlier ",
                  "among adult week old Japanese quail with ",
                  "body temperature measured at 30°C and 40°C."
)

tbResponseHot8Weeks$data %>%
  mutate("paretoK" = loo(tbResponseHot8Weeks)$diagnostics$pareto_k) %>%
  filter(paretoK > 0.5) %>%
  select(-c(pretreatment, mass, tarsus, batch)) %>%
  merge(., tbResponse %>%
          filter(challenge == "warm"),
        by = c("deltaTb"),
```

```
        all.x = TRUE) %>%
  merge(., all %>%
          filter(Ta == 30) %>%
          select(ring, week, meanTb) %>%
          rename("Tb at 30°C" = meanTb),
        by = c("ring", "week"),
        all.x = TRUE) %>%
    merge(., all %>%
          filter(Ta == 40) %>%
          select(ring, week, meanTb) %>%
          rename("Tb at 40°C" = meanTb),
        by = c("ring", "week"),
        all.x = TRUE) %>%
  mutate(pretreatment = ifelse(pretreatment == "cold",
                               "Cold (10°C)",
                               ifelse(pretreatment == "neutral",
                                      "Mild (20°C)", "Warm (30°C)")
                               )
         ) %>%
  select("Bird ID" = ring,
         "Rearing Treatment" = pretreatment,
         "Egg Batch" = batch,
         "Mass (g)" = mass,
         "Tarsus Length (mm)" = tarsus,
         `Tb at 30°C`,
         `Tb at 40°C`,
         "Delta Tb (°C)" = deltaTb,
         "Pareto K" = paretoK) %>%
  kbl(.,
    longtable = T, booktabs = T, format = "latex", escape = FALSE,
    caption = caption
  ) %>%
  column_spec(column = c(1:10), width = "1.4cm") %>%
  kable_styling(latex_options = "striped")
```

***Table 4:*** *Potential body temperature response outlier among adult week old Japanese quail with body temperature measured at 30°C and 40°C.*

| Bird ID | Rearing Treatment | Egg Batch | Mass (g) | Tarsus Length (mm) | Tb at 30°C | Tb at 40°C | Delta Tb (°C) | Pareto K |
|---|---|---|---|---|---|---|---|---|
| R20 | Mild (20°C) | B | 264.5 | 42.537 | 42.319 | 43.45058 | 1.131581 | 0.5600915 |

All measurements are within the range of expectations for adult Japanese quail. For this reason, the value is retained and we proceed to estimating variance in our data explained by our path analysis and visualing posterior densities.

```
# Checking R2 values

caption <- paste0("R\\textsuperscript{2} for Bayesian path analysis ",
                  "ultimate predicting body temperature responses (°C) to heat ",
                  "(40°C) in eight week old Japanese quail. Baseline body ",
                  "temperatures represent those measured at thermoneutrality ",
                  "(30°C)"
)

brms::bayes_R2(tbResponseHot8Weeks,
  robust = TRUE, ndraws = 1000
) %>%
  as.data.frame() %>%
  rownames_to_column(var = "Response") %>%
  mutate(Response = ifelse(Response == "R2mass", "Body Mass (g)",
```

```
    ifelse(Response == "R2tarsus", "Tarsus Length (mm)",
      "Change in Tb (°C)"
    )
  )) %>%
  mutate(
    Estimate = round(Estimate, digits = 4),
    Est.Error = round(Est.Error, digits = 4),
    Q2.5 = round(Q2.5, digits = 4),
    Q97.5 = round(Q97.5, digits = 4)
  ) %>%
  rename(
    "R\\textsuperscript{2}" = Estimate,
    "Standard Error" = Est.Error,
    `2.5\\%CI` = "Q2.5", `97.5\\% CI` = "Q97.5"
  ) %>%
  kbl(.,
    longtable = T, booktabs = T, format = "latex",
    caption = caption, escape = FALSE
  ) %>%
  kable_styling(latex_options = "striped")
```

***Table 5:*** *$R^2$ for Bayesian path analysis ultimate predicting body temperature responses (°C) to heat (40°C) in eight week old Japanese quail. Baseline body temperatures represent those measured at thermoneutrality (30°C)*

| Response | $R^2$ | Standard Error | 2.5%CI | 97.5% CI |
|---|---|---|---|---|
| Body Mass (g) | 0.0412 | 0.0351 | 0.0029 | 0.1294 |
| Tarsus Length (mm) | 0.3249 | 0.0697 | 0.1673 | 0.4436 |
| Change in Tb (°C) | 0.1733 | 0.0646 | 0.0600 | 0.2880 |

```
# R2 of temperature responses slightly improved relative to that estimated
# in the cold.

as.data.frame(tbResponseHot8Weeks) %>%
    select(
      "Intercept" = b_deltaTb_Intercept,
      "Body Mass (g)" = b_deltaTb_mass,
      "Tarsus Length\n(mm)" = b_deltaTb_tarsus,
      "Cold Rearing\n(10°C)" = b_deltaTb_pretreatmentA,
      "Warm Rearing\n(30°C)" = b_deltaTb_pretreatmentC,
      "Egg Batch (mu)" = sd_batch__deltaTb_Intercept,
      "Sigma" = sigma_deltaTb
    ) %>%
    pivot_longer(everything(), names_to = "Par",
                 values_to = "Coefs") %>%
    ggplot(aes(x = Coefs)) +
    facet_wrap(~Par, scales = "free", ncol = 2) +
    geom_density(colour = "black", alpha = 0.5, fill = "grey70") +
    geom_vline(xintercept = 0, linetype = "dashed", colour = "black") +
    ylab("Density") +
    theme_classic() +
    theme(axis.title.x = element_blank())
```

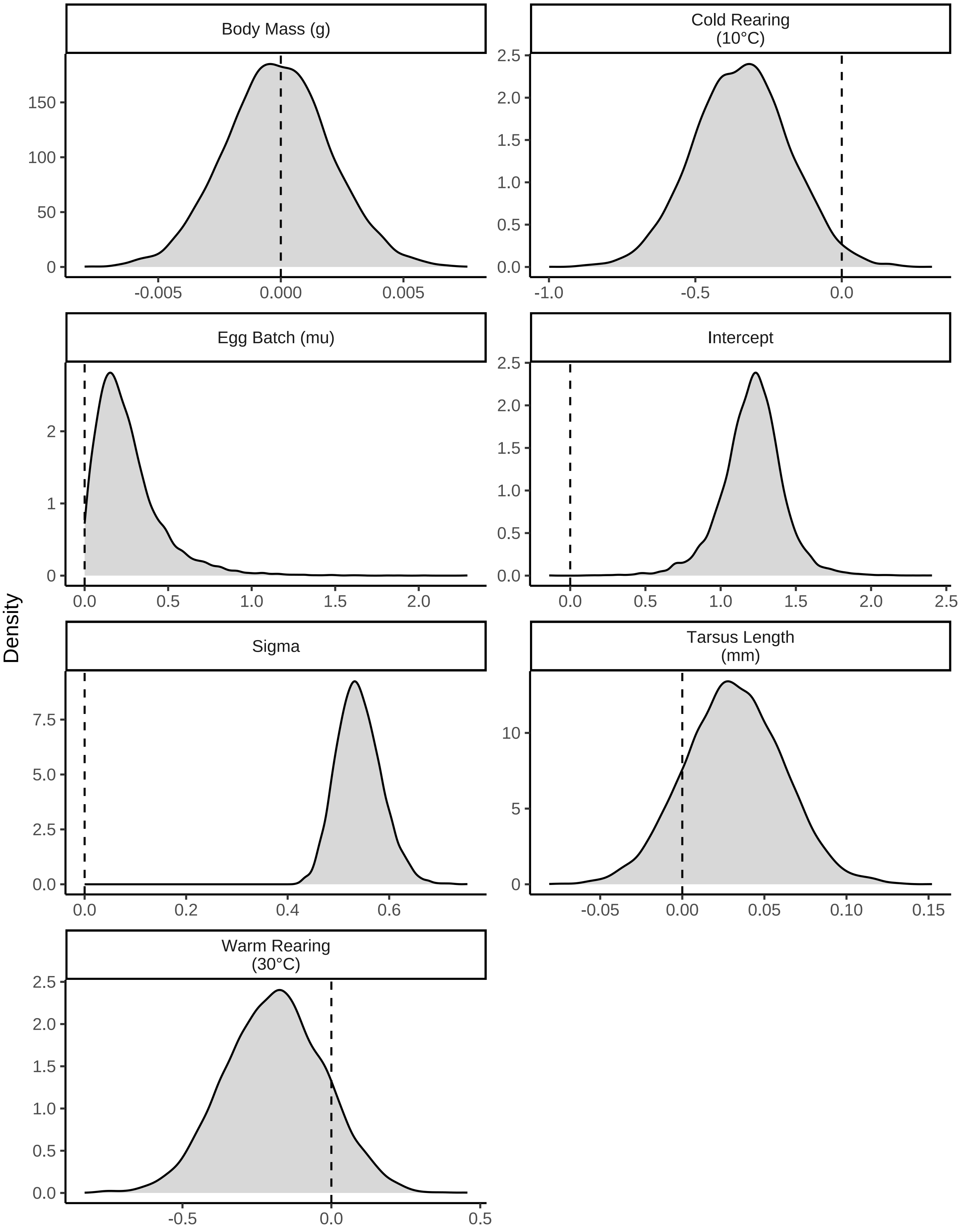


***Figure 16:*** *Posterior densities for model coefficients derived from a Bayesian path analysis ultimately predicting body temperature responses (°C) to heat (40°C) in eight week old Japanese quail. Densities are split by their respective response values (indicated with titles). Vertical dashed lines indicate 0.*

Last, we summarise estimates from our path analysis while again using posterior medians to represent model coefficients and quantiles to estimate credible intervals around medians.

```
caption <- paste0("Results from a Bayesian path analysis ",
                  "ultimately predicting body temperature responses to a ",
                  "heat exposure (40°C) relative to thermoneutrality (30°C)",
                  "in adult Japanese quail. Body temperature responses are ",
                  "predicted as a function of body mass (g), tarsus length ",
                  "(mm) and rearing treatment. ",
                  "Cold rearing indicates post-hatch rearing at ",
                  "10°C, relative to ",
                  "20°C (intercept), or ",
                  "30°C ('warm rearing'). ",
                  "Coefficients represent medians and credible intervals ",
                  "(CIs) represent quantile intervals. ",
                  "BF indicates Bayes Factors."
)

as.data.frame(tbResponseHot8Weeks) %>%
  summarise_all(., .funs = median) %>%
  pivot_longer(everything(),
    names_to = "Parameter",
    values_to = "Estimate"
  ) %>%
  merge(., quantileCIs(tbResponseHot8Weeks, cis = c(50, 95)),
    by = "Parameter", all.x = TRUE
  ) %>%
  filter(grepl("b_|sd_", Parameter)) %>%
  rowwise() %>%
  mutate("BF" = ifelse(Estimate < 0,
    (2 * mean(as.data.frame(
      tbResponseHot8Weeks
    )[, Parameter] <= 0)) /
      (2 * mean(as.data.frame(
        tbResponseHot8Weeks
      )[, Parameter] >= 0)),
    (2 * mean(as.data.frame(
      tbResponseHot8Weeks
    )[, Parameter] >= 0)) /
      (2 * mean(as.data.frame(
        tbResponseHot8Weeks
      )[, Parameter] <= 0))
  )) %>%
  ungroup() %>%
  mutate(
    "Estimate" = round(Estimate, digits = 4),
    "BF" = round(BF, digits = 4),
    "N" = nrow(tbResponseHot8Weeks$data)
  ) %>%
  rowwise() %>%
  mutate(Parameter = gsub("deltaTb", "deltaT", Parameter)) %>%
  ungroup() %>%
  mutate("Parameter" = ifelse(grepl("b_", Parameter),
                              gsub("b_", "", Parameter),
                              gsub("Intercept", "batch",
                                   gsub(".*__", "", Parameter)
                                   )
                              )
         ) %>%
  mutate(
    "Response" = gsub("_.*", "", Parameter),
    "Parameter" = gsub(".*_", "", Parameter)
  ) %>%
  merge(., tribble(
    ~Response, ~response, ~level,
    "mass", "Body Mass (g)", "A",
    "deltaT",
```

```
    "Delta Body Temperature (°C)", "D",
    "tarsus", "Tarsus Length (mm)",
    "B"
  ),
  by = "Response"
  ) %>%
  merge(., tribble(
    ~Parameter, ~parameter, ~number,
    "Intercept", "Intercept", "1",
    "mass", "Body Mass (g)", "4",
    "tarsus", "Tarsus Length (mm)", "5",
    "pretreatmentA", "Cold Rearing", "2",
    "pretreatmentC", "Warm Rearing", "3",
    "batch", "Egg Batch [mu]", "6"
  ),
  by = "Parameter"
  ) %>%
  mutate(
    `50\\% CI` = paste0("(", paste(
      round(Low_CI_50, digits = 4),
      round(High_CI_50, digits = 4),
      sep = ", "
    ), ")"),
    `95\\% CI` = paste0("(", paste(
      round(Low_CI_95, digits = 4),
      round(High_CI_95, digits = 4),
      sep = ", "
    ), ")")
  ) %>%
  select(-c(Low_CI_50, High_CI_50, Low_CI_95, High_CI_95)) %>%
  select(
    "Response" = "response", "Parameter" = "parameter", N,
    Estimate, `50\\% CI`, `95\\% CI`, BF, level, number
  ) %>%
  arrange(level, number) %>%
  select(-c(level, number)) %>%
  kbl(.,
    longtable = T, booktabs = T, format = "latex", escape = FALSE,
    caption = caption
  ) %>%
  column_spec(column = c(1:2), width = "2.2cm") %>%
  column_spec(column = c(3:10), width = "1.9cm") %>%
  kable_styling(latex_options = "striped")
```

***Table 6:*** *Results from a Bayesian path analysis ultimately predicting body temperature responses to a heat exposure (40°C) relative to thermoneutrality (30°C)in adult Japanese quail. Body temperature responses are predicted as a function of body mass (g), tarsus length (mm) and rearing treatment. Cold rearing indicates post-hatch rearing at 10°C, relative to 20°C (intercept), or 30°C ('warm rearing'). Coefficients represent medians and credible intervals (CIs) represent quantile intervals. BF indicates Bayes Factors.*

| Response | Parameter | N | Estimate | 50% CI | 95% CI | BF |
|---|---|---|---|---|---|---|
| Body Mass (g) | Intercept | 84 | -6.5551 | (-10.1365, -3.0202) | (-16.9546, 3.9759) | 8.4899 |
| Body Mass (g) | Cold Rearing | 84 | 2.4098 | (-2.7582, 7.5298) | (-12.4358, 17.161) | 1.6429 |
| Body Mass (g) | Warm Rearing | 84 | 12.0673 | (7.1913, 17.1146) | (-2.613, 26.436) | 17.1406 |
| Body Mass (g) | Egg Batch [mu] | 84 | 0.2807 | (0.1156, 0.5617) | (0.0102, 1.4918) | Inf |
| Tarsus Length (mm) | Intercept | 84 | -0.4942 | (-0.8773, -0.0981) | (-1.644, 0.8971) | 3.9170 |
| Tarsus Length (mm) | Cold Rearing | 84 | 0.5397 | (0.1245, 0.9692) | (-0.6746, 1.7766) | 4.2840 |
| Tarsus Length (mm) | Warm Rearing | 84 | 0.7450 | (0.3287, 1.1552) | (-0.4758, 1.9066) | 7.7146 |

| | | | | | | |
|---|---|---|---|---|---|---|
| Tarsus Length (mm) | Body Mass (g) | 84 | 0.0315 | (0.0262, 0.0366) | (0.0165, 0.0467) | Inf |
| Tarsus Length (mm) | Egg Batch [mu] | 84 | 0.6614 | (0.422, 0.9537) | (0.0582, 1.7962) | Inf |
| Delta Body Temperature (°C) | Intercept | 84 | 1.2174 | (1.0973, 1.3277) | (0.7639, 1.6044) | 7999.0000 |
| Delta Body Temperature (°C) | Cold Rearing | 84 | -0.3410 | (-0.4535, -0.2332) | (-0.6619, -0.0222) | 52.3333 |
| Delta Body Temperature (°C) | Warm Rearing | 84 | -0.1882 | (-0.3031, -0.0774) | (-0.5169, 0.1322) | 6.8663 |
| Delta Body Temperature (°C) | Body Mass (g) | 84 | -0.0001 | (-0.0015, 0.0013) | (-0.0043, 0.0041) | 1.1069 |
| Delta Body Temperature (°C) | Tarsus Length (mm) | 84 | 0.0312 | (0.0111, 0.0517) | (-0.0276, 0.0898) | 5.7739 |
| Delta Body Temperature (°C) | Egg Batch [mu] | 84 | 0.2152 | (0.1243, 0.3391) | (0.0186, 0.7917) | Inf |
